# In-host evolution of *Yersinia enterocolitica* during a chronic human infection

**DOI:** 10.1101/2024.06.12.598599

**Authors:** Savin Cyril, Lê-Bury Pierre, Guglielmini Julien, Douché Thibaut, Buzelé Rodolphe, Le Brun Cécile, Bastides Frédéric, François Maud, Birmelé Béatrice, Guichard Laura, Cabanel Nicolas, Dortet Laurent, Matondo Mariette, Dussurget Olivier, Carniel Elisabeth, Lanotte Philippe, Pizarro-Cerdá Javier

## Abstract

Following a pacemaker implantation, a 75-years-old patient suffered from five successive bacteremia episodes between in 1999 and 2013 despite long-term antibiotic treatment, with intermittent vegetation apparition on the device atrial lead. Four blood isolates, identified as *Yersinia enterocolitica* bioserotype 4/O:3, were further genetically and phenotypically characterized. Phylogenetic reconstruction showed that the patient was chronically infected by the same strain, which evolved within the host for 14 years. Single-nucleotide polymorphism (SNP) analysis indicates that the last two isolates evolved in parallel and formed two independent lineages within the host. Pan-genome analysis and genome comparison showed that their common evolution was characterized by 41 small insertion/deletion events, loss of three large DNA fragments and mutations in 140 genes. A phylogenetic analysis by maximum likelihood identified two genes presenting a positive selection signal, suggesting that these mutations provided a survival advantage to bacteria during chronic infection. Quinolone resistance in the last two isolates was acquired through a so far undescribed deletion in the *gyrA* gene.

Mass-spectrometry analysis revealed a strong proteome remodeling in the last two isolates which was correlated with a truncation in the stringent response regulator DksA. A reduced carbon, energy and purine metabolism supports their severe growth defects *in vitro*. 3^rd^-generation cephalosporin resistance of the last isolate was correlated with a truncation of OmpF, the main porin translocating antibiotics through the outer-membrane, as well as an increased production of BlaA and AmpC β-lactamases.

This is the first report of genetic and phenotypic changes associated to within-host adaptation of a pathogenic *Yersinia* species under antibiotic pressure.

## Introduction

Bacterial exposure to a novel environment in a host creates conditions where evolution might take place. As observed for cancer cells, a bacterial clone can experience selection for an accumulation of genetic variants that promote long-term survival and clonal expansion^1^. This scenario has been observed in cystic fibrosis patients who can be chronically infected with pathogenic bacteria. In a study from a single patient infected for 20 years with *Burkholderia multivorans*, bacterial adaptation to respiratory airways was associated with mutations affecting metabolism, cell envelope, and biofilm formation^2^. In a different study concerning 474 *P. aeruginosa* isolates from 34 cystic fibrosis patients, convergent evolution of 52 genes involved in antibiotic resistance, motility, and biofilm formation was observed^3^. Host-adaptation through rapid evolution of a *Salmonella enterica* serotype Enteritidis causing a chronic systemic infection was documented in an immunocompromised patient^4^. Genetic diversification of *Staphylococcus epidermidis* has also been described during a 16-week pacemaker-associated endocarditis, leading to increased biofilm formation, reduced growth rate, and antibiotic tolerance^5^.

In-host evolution has not been reported for infections associated with *Yersinia* spp. Yersinioses include fulminant infections such as plague caused by *Yersinia pestis*, as well as mild or severe enteritis caused by *Yersinia pseudotuberculosis* or *Yersinia enterocolitica* (*Ye)*^6,7^. The latter represents the third most common cause of diarrhea from bacterial origin in temperate and cold countries^8^. Systemic *Ye* infections occur mostly in elderly patients with underlying disorders such as diabetes, iron overload, or cirrhosis^9^.

Persistent *Ye* infections in patients causing relapses for several years have been reported^10^, but the potential genetic relationships between the isolated strains was not determined. This study is the first report of long-term, in-host evolution of a pathogenic *Yersinia* species in a patient presenting iterative episodes of bacteremia over 14 years.

## Results

### Case description

A 75-year-old woman presented with an atrioventricular block that led to a pacemaker implantation in September 1998. In December 1999, the patient had a first septicemic episode resulting in the isolation of a *Ye* strain (Ye.1) (Fig 1). She received ceftriaxone and netilmicin treatment for 4 weeks and recovered. In January 2000, the patient experienced a second episode of bacteremia with isolation of another *Ye* strain (Ye.2). She was treated with the same antibiotics for four weeks, followed by 18 months of ciprofloxacin therapy (until July 2001), during which monthly blood cultures were negative. A third bacteremia occurred in August 2001 with isolation of a third *Ye* (Ye.X, not kept in collection) that was resistant to nalidixic acid but susceptible to ciprofloxacin, which had been used for treatment. The patient received a long-term ceftriaxone therapy until July 2005, and monthly blood cultures were negative. In June 2006, the patient was hospitalized for a sepsis but had a negative blood culture. For the first time, echocardiography evidenced vegetations on the pacemaker atrial lead, suggesting endocarditis due to bacterial growth on the cardiac device (Fig S1). A 6-year ceftriaxone treatment was administered, leading to vegetation disappearance. In December 2012, the patient suffered from pneumonia, and ceftriaxone treatment was replaced by spiramycin. In January 2013, she presented with another bacteremia with isolation of a fourth *Ye* strain (Ye.3) and vegetations reappeared on the pacemaker atrial lead. Strain Ye.3, like Ye.X, was resistant to nalidixic acid. The patient was treated with piperacillin/tazobactam/amikacin for one week, followed by a 9-month ceftriaxone therapy. The patient had a last bacteremia in October 2013, during which a fifth *Ye* strain (Ye.4) was isolated. Minimum inhibitory concentration (MIC) of ceftriaxone increased from 0.19 mg/L (Ye.3) to 2 mg/L (Ye.4) leading to antimicrobial treatment modification with cotrimoxazole. Three months later, the patient died of heart failure.

**Figure 1.**
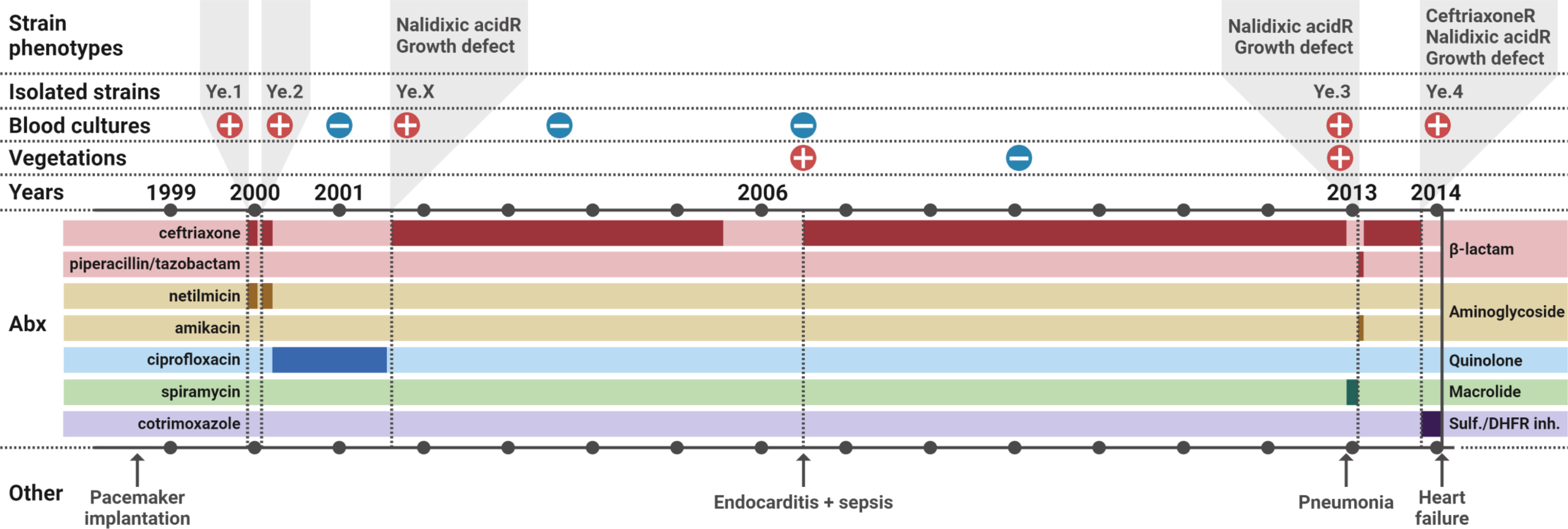
Clinical case description of *Ye* chronic infection. Positive and negative symbols reflect the blood culture results, or the observation of vegetations on the pacemaker atrial lead. Periods of antibiotics treatment are shown in a darker color on the chronograph, with the associated treatment names on the left and antibiotic classes on the right. Abx: antibiotics. CeftriaxoneR: resistant to ceftriaxone. Nalidixic acidR: resistant to nalidixic acid. Sulf: sulfonamide. DHFR inh: dihydrofolate reductase inhibitor.

### Phenotypic characterization of *Ye* isolates shows in-host evolution of antibiotic susceptibility during long term treatment

The five strains isolated during the iterative bacteremia were initially identified as *Ye* using a VITEK2 GN card. Except for Ye.X, which was not kept in collection, strains were subjected to phenotypical characterization: Ye.1 and Ye.2 (also referred as “early isolates”) were identified as *Ye* bioserotype 4/O:3; due to a severe growth defect, Ye.3 and Ye.4 (also referred as “late isolates”) could not be characterized by phenotypic methods. A substantial increase in mid-exponential phase doubling time at 37°C was measured, from 49 and 44 minutes for Ye.1 and Ye.2 to 5h38 and 4h46 for Ye.3 and Ye.4, respectively. Doubling times at 28°C were slightly longer, with 57 minutes for Ye.1 and Ye.2, 6h49 for Ye.3 and 5h28 for Ye.4 (Fig S2).

After isolation of the four strains, we parallelly assessed the susceptibility of Ye.1 to Ye.4 to 33 different antibiotics or antibiotics combinations (Table S1). All strains were resistant to the penicillins amoxicillin and ticarcillin. Ye.1 and Ye.2 were resistant to first-generation cephalosporin (C1G) (data not shown) and they were susceptible to all other tested antibiotics, as usually observed for *Ye* bioserotype 4/O:3, although Ye.2 showed a small resistance increase to several β-lactams and netilmicin, the aminoglycoside (AG) which was administered in combination with the 3^rd^ generation cephalosporin (C3G) ceftriaxone between Ye.1 and Ye.2 isolation. Ye.3 and Ye.4 acquired a resistance to the quinolone nalidixic acid, as observed with Ye.X, and Ye.4 became resistant to the second-generation cephalosporin (C2G) cefoxitin.

To further characterize the antibiotic profile of the isolated strains, the MIC of 10 antibiotics and antibiotics combinations was measured (Table 1). While the MIC of different antibiotics classes such as cephalosporin, AG and fluoroquinolone (FQ) slightly increased between Ye.1 and Ye.2, an unexpected AG and FQ MIC decrease was observed for the late isolates Ye.3 and Ye.4 compared to the early isolates Ye.1 and Ye.2. Furthermore, Ye.4 was fully resistant to the C3G ceftriaxone as well as the C2G cefoxitin, as observed before, and was slightly more resistant than Ye.3 to piperacillin-tazobactam, the combination of a penicillin and a β-lactamase inhibitor administered before ceftriaxone between Ye.3 and Ye.4 isolation.

**Table 1.** Minimum inhibitory concentration (mg/L) of the 4 Ye strains on 10 tibiotics and antibiotics combinations assessed by Etest.

| Antibiotic class | Antibiotic | Ye.1 | Ye.2 | Ye.3 | Ye.4 |
| --- | --- | --- | --- | --- | --- |
| β-lactam | ceftriaxone<sup>123</sup> | 0,047 | 0,094 | 0,094 | >256 |
|  | cefoxitin | 4 | 12 | 12 | >256 |
|  | piperacillin-tazobactam<sup>3</sup> | 0,094 | 0,25 | <0,016 | 0,094 |
| aminoglycosides and<br>aminoglycosides-like | netilmicin<sup>12</sup> | 0,75 | 3 | 0,06 | 0,06 |
|  | amikacin<sup>3</sup> | 3 | 3 | 2 | 1 |
|  | tobramycin | 1 | 1 | 0,09 | 0,09 |
|  | gentamicin | 0,5 | 1 | 0,09 | 0,09 |
|  | spectinomycin | 12 | 24 | 0,09 | 0,09 |
| Quinolone | ciprofloxacin<sup>2</sup> | 0,008 | 0,012 | 0,002 | 0,032 |
|  | levofloxacin | 0,032 | 0,032 | 0,008 | 0,032 |

<sup>1</sup>Administred treatment between Ye.1 and Ye.2 isolation
<sup>2</sup>Administred treatment between Ye.2 and Ye.3 isolation
<sup>3</sup>Administred treatment between Ye.3 and Ye.4 isolation

### Genetic analysis supports a chronic *Ye* infection, unravels a new quinolone resistance mutation in *gyrA* and reveals a OmpF-mediated cephalosporin resistance

Taxonomic assignment of the 4 isolated strains was obtained based on a core-genome multilocus sequence typing (cgMLST) with 500 core genes using whole-genome *de novo* sequence, confirming that strains Ye.1 to Ye.4 belong to the *Ye* genotype 4, corresponding to bioserotype 4/O:3.

To investigate whether the multiple infection episodes were independent or due to a chronic colonization by a unique evolving bacterial strain, we studied the genetic relatedness of strains Ye.1 to Ye.4 by a core-genome single-nucleotide polymorphism analysis, with inclusion of 259 additional *Ye* biotype 4 strains isolated between 1963 and 2016 (Table S2). The phylogenetic tree reconstruction based on analysis of 4,738 SNPs showed that Ye.1 to Ye.4 were closely related and formed a unique clade separated from other *Ye* strains (Fig 2A). This result strongly suggests that the patient suffered from a chronic infection due to a unique evolving *Ye* 4/O:3 strain.

**Figure 2.**
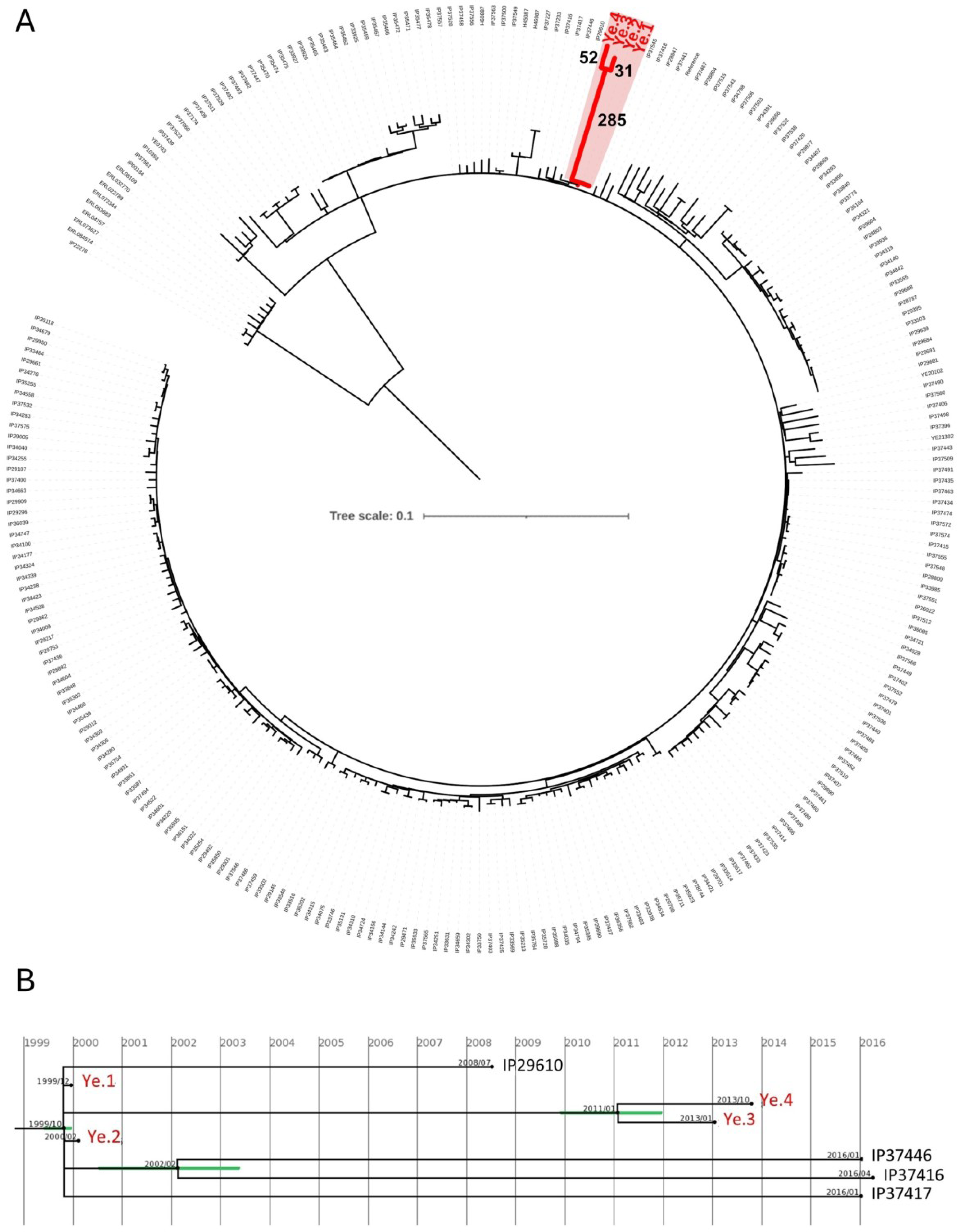
Phylogenetic reconstruction. (A) Maximum likelihood phylogeny reconstructed with RAxML under GTR model with 100 bootstraps based on 4,738 SNPs identified in 262 strains. The genome of the Y11 strain (NC_017564) was used as reference for variant calling. Numbers close to the branches indicate SNPs numbers. (B) Estimation of the date of internal nodes using least-squares dating (https://lsdating.pasteur.fr) among the 4 strains isolated from the patient. Green bars represent the standard deviation of the estimated date.

We next analyzed the molecular events that characterized the in-host evolution of Ye.1 to Ye.4. Genomic comparison and pan-genome phylogenetic analysis did not evidence any gene acquisition or small insertion/deletion events that were common and specific to the four *Ye* isolates. Ye.1 (isolated in December 1999) and Ye.2 (isolated in January 2000) were identical except for the loss in Ye.1 of a 1.5 kb region (between nucleotides 4,282,424 and 4,283,952 from the Y11 reference genome), indicating that evolution had already started within the first months of infection. The use of the least-squares dating tool (https://lsdating.pasteur.fr) estimated the closest internal node for Ye.1 and Ye.2 in October 1999, suggesting that the patient was contaminated at least two months before the first bacteremic episode (Fig 2B). The closest internal node for Ye.3 (isolated in January 2013) and Ye.4 (isolated in October 2013) was January 2011 (Fig 2B). However, a truncation in the mutation frequency decline *mfd* gene and the slow growth phenotype of the late isolates should be considered, thus the divergence between Ye.3 and Ye.4 could have happened earlier. Their common ancestor had therefore evolved in the patient for at least 11 years and accumulated 285 SNPs before diverging into Ye.3 and Ye.4, which since accumulated additional 31 and 52 SNPs, respectively (Table S3). 140 genes containing mutations vertically acquired from their common ancestor (synapomorphies) were identified in the Ye.3 and Ye.4 genomes, including the *gyrA* gene (see below and Table S4). Ye.3 and Ye.4 shared three identical large deletions: a 2-gene deletion of 2.5 kb, a 9-gene deletion of 9.9 kb, and a 28-gene deletion of 32.7 kb (Table 2). These strains also shared 41 small insertions/deletions (Table 2). Whereas 6 genes did not exhibit any frameshift, 35 genes displayed a mutation leading to a truncated form of the product, and thus to loss-of-function. Truncated genes and deleted regions were found to be involved in essential physiological functions such as transcription, translation, replication, respiration, division, and carbohydrate and iron metabolism (Table 2).

**Table 2.** Small insertions/deletions and list of CDS absent in Ye.3 and Ye.4 strains’ genome, along log2(foldchange) of protein abundance in all pairwise comparisons. Loci are expressed on Y11 reference genome. P-value and adjusted p-value can be found in Table S6. 0 corresponds to proteins not detected in any conditions of the comparison. 15 corresponds to proteins detected only in the first condition of the comparison.

| Genbank id | Locus | Old locus | Status | Product | Ye.1<br>vs<br>Ye.2 | Ye.1<br>vs<br>Ye.3 | Ye.1<br>vs<br>Ye.4 | Ye.2<br>vs<br>Ye.3 | Ye.2<br>vs<br>Ye.4 | Ye.3<br>vs<br>Ye.4 |
| --- | --- | --- | --- | --- | --- | --- | --- | --- | --- | --- |
| Small insertions/deletions |  |  |  |  |  |  |  |  |  |  |
| WP_005165697.1 | Y11_RS00200 | Y11_00411 | complete | NADP-dependent oxaloacetate-decarboxylating malate dehydrogenase MaeB | -0,26 | 0,35 | 1,72 | 0,61 | 1,97 | 1,37 |
| WP_005164830.1 | Y11_RS00450 | Y11_00921 | truncated | Glucokinase Glk | 0,38 | 4,86 | 4,31 | 4,43 | 3,95 | 0 |
| WP_013650157.1 | Y11_RS01245 | Y11_02641 | complete | DNA topoisomerase (ATP-hydrolyzing) subunit A GyrA | 0,06 | -0,75 | -0,25 | -0,72 | -0,96 | -0,24 |
| WP_005162896.1 | Y11_RS01655 | Y11_03551 | complete | HAAAP family serine/threonine permease | -0,93 | -0,81 | 0,7 | 0,12 | 1,63 | 1,51 |
| WP_005158003.1 | Y11_RS02570 | Y11_05521 | truncated | transcription-repair coupling factor Mfd | -0,02 | 3,35 | 3,19 | 3,38 | 3,21 | -0,17 |
| WP_112999146.1 | Y11_RS02795 | Y11_06001 | truncated | sensor domain-containing diguanylate cyclase | 0,44 | 1,32 | 15 | 0,82 | 15 | 0 |
| WP_013649985.1 | Y11_RS03165 | Y11_06821 | truncated | MFS transporter | 1,2 | 15 | 15 | 15 | 15 | 0 |
| WP_076706727.1 | Y11_RS03165 | Y11_06821 | truncated | MFS transporter | 1,2 | 15 | 15 | 15 | 15 | 0 |
| WP_050344735.1 | Y11_RS03400 | Y11_07381 | truncated | RNA chaperone ProQ | 0,36 | 2,64 | 1,47 | 2,28 | 1,11 | -1,17 |
| WP_005165635.1 | Y11_RS03825 | Y11_08261 | truncated | MATE family efflux transporter | 0 | 0 | 0 | 0 | 0 | 0 |
| WP_001534187.1 | Y11_RS04455 | Y11_09631 | truncated | hypothetical protein | 0 | 0 | 0 | 0 | 0 | 0 |
| WP_145535939.1 | Y11_RS04925 | Y11_10641 | truncated | type I DNA topoisomerase TopA | 0,57 | 1,09 | -0,12 | 0,52 | -0,69 | -1,21 |
| WP_005157750.1 | Y11_RS07710 | Y11_16461 | truncated | APC family permease | 0 | 0 | 0 | 0 | 0 | 0 |
| WP_005158842.1 | Y11_RS08345 | Y11_17791 | truncated | urea ABC transporter ATP-binding subunit Urte | 0 | 0 | 0 | 0 | 0 | 0 |
| WP_145531626.1 | Y11_RS08850 | Y11_18831 | truncated | two-component system sensor histidine kinase KdpD | -0,57 | 0,59 | 0,27 | 1,16 | 0,84 | -0,32 |
| WP_050879682.1 | Y11_RS09025 | Y11_19131 | truncated | amino acid ABC transporter permease | 3,16 | 15 | 15 | 15 | 15 | 0 |
| WP_005167417.1 | Y11_RS10470 | Y11_22201 | truncated | MFS transporter | 0 | 0 | 0 | 0 | 0 | 0 |
| WP_085910232.1 | Y11_RS10470 | Y11_22201 | truncated | MFS transporter | 0 | 0 | 0 | 0 | 0 | 0 |
| WP_005163236.1 | Y11_RS10625 | Y11_22531 | truncated | beta-ketoacyl-[acyl-carrier-protein] synthase family protein | 0 | 0 | 0 | 0 | 0 | 0 |
| WP_005163313.1 | Y11_RS10785 | Y11_22871 | truncated | Aminopeptidase | 0,67 | 2,46 | 1,95 | 1,78 | 1,27 | -0,52 |
| WP_005160711.1 | Y11_RS11590 | Y11_24591 | truncated | hypothetical protein | 0 | 0 | 0 | 0 | 0 | 0 |
| WP_005180337.1 | Y11_RS11725 | Y11_24871 | truncated | 3',5'-cyclic-AMP phosphodiesterase - CpdA | 1,49 | 2,66 | 2,09 | 1,17 | 0,63 | 0 |
| WP_016266722.1 | Y11_RS12715 | Y11_26981 | truncated | toxin subunit | 0 | 0 | 0 | 0 | 0 | 0 |
| WP_011815289.1 | Y11_RS13360 | Y11_28281 | truncated | class II fructose-bisphosphatase GlpX | -0,55 | 3,5 | 2,93 | 4,05 | 3,47 | -0,58 |
| WP_005161086.1 | Y11_RS14235 | Y11_30151 | truncated | DUF3748 domain-containing protein | 0 | 0 | 0 | 0 | 0 | 0 |
| WP_049679210.1 | Y11_RS14685 | Y11_31111 | truncated | dipeptide transporter permease DppB | 0 | 0 | 0 | 0 | 0 | 0 |
| WP_019250468.1 | Y11_RS14860 | Y11_31471 | truncated | inorganic phosphate transporter PitA | -0,31 | 15 | 0,61 | 15 | 1,02 | 0 |
| WP_005157438.1 | Y11_RS14960 | Y11_31681 | truncated | MurR/RpiR family transcriptional regulator | -0,2 | 1,81 | 2,63 | 2,01 | 2,83 | 0,82 |
| WP_014609301.1 | Y11_RS15690 | Y11_33251 | truncated | DUF494 domain-containing protein | 0 | 0 | 0 | 0 | 0 | 0 |
| WP_005181075.1 | Y11_RS15890 | Y11_33631 | truncated | guanosine pentaphosphate phosphohydrolase/exopolyposphatase GppA | 0,23 | 2,92 | 15 | 2,63 | 15 | 0 |
| WP_005166113.1 | Y11_RS16045 | Y11_33921 | truncated | DUF484 domain-containing protein | 1,61 | 15 | 15 | 15 | 15 | 0 |
| WP_005165794.1 | Y11_RS16565 | Y11_34951 | complete | DNA-directed RNA polymerase subunit beta' RpoC | -0,1 | -0,06 | -0,41 | 0,05 | -0,3 | -0,35 |
| WP_005156737.1 | Y11_RS16685 | Y11_35131 | truncated | cation/acetate symporter ActP | -0,51 | 15 | 15 | 15 | 15 | 0 |
| WP_005156732.1 | Y11_RS16695 | Y11_35151 | truncated | acetate--CoA ligase Acs | -0,29 | 3,98 | 15 | 4,27 | 15 | 15 |
| WP_011815418.1 | Y11_RS17110 | Y11_36011 | truncated | miniconductance mechanosensitive channel MscM | -0,24 | 2,59 | 2,75 | 2,83 | 2,99 | 0,16 |
| WP_005156285.1 | Y11_RS17420 | Y11_36641 | truncated | 23S rRNA (uridine(2552)-2'-O)-methyltransferase RlmE | 0,36 | 15 | 2,4 | 15 | 2,09 | 0 |
| WP_023160195.1 | Y11_RS17940 | Y11_37741 | truncated | dihydroxyacetone kinase subunit DhaM | -0,65 | 1,25 | 0,66 | 1,91 | 1,31 | -0,6 |
| WP_004706052.1 | Y11_RS18210 | Y11_38281 | complete | two-component system response regulator ArcA | 0,65 | 0,37 | 0,17 | -0,28 | -0,48 | -0,2 |
| WP_049530647.1 | Y11_RS18720 | Y11_39351 | truncated | TonB-dependent siderophore receptor | 0 | 0 | 0 | 0 | 0 | 0 |
| WP_005156764.1 | Y11_RS18820 | Y11_39571 | truncated | RNA polymerase-binding protein DksA | -0,03 | 4,47 | 4,42 | 4,51 | 4,46 | -0,05 |
| KGA74145.1 |  |  | complete | hypothetical protein DJ62_2388 |  |  |  |  |  |  |
| Large deletion of 9,919 bp (positions 887742 to 897661) - 9 genes |  |  |  |  |  |  |  |  |  |  |
| WP_072076553.1 | Y11_RS03970 | Y11_08571 | deleted | dipeptide/tripeptide permease DtpA | 0,45 | 15 | 15 | 15 | 15 | 0 |
| WP_005166379.1 | Y11_RS03975 | Y11_08591 | deleted | LysR family transcriptional regulator | -0,47 | 15 | 15 | 15 | 15 | 0 |
| WP_005166377.1 | Y11_RS03980 | Y11_08601 | deleted | oxidoreductase | 0 | 0 | 0 | 0 | 0 | 0 |
| WP_005178982.1 |  |  | deleted | hypothetical protein |  |  |  |  |  |  |
| WP_005166373.1 | Y11_RS03990 | Y11_08621 | deleted | peptide ABC transporter ATP-binding protein SapF | -0,01 | 15 | 15 | 15 | 15 | 0 |
| WP_005166371.1 | Y11_RS03995 | Y11_08631 | deleted | peptide ABC transporter ATP-binding protein SapD | 0,31 | 15 | 4,23 | 15 | 3,72 | 0 |
| WP_005166369.1 | Y11_RS04000 | Y11_08641 | deleted | peptide ABC transporter permease SapC | 0 | 0 | 0 | 0 | 0 | 0 |
| WP_005169521.1 | Y11_RS04005 | Y11_08651 | deleted | peptide ABC transporter permease SapB | 0 | 0 | 0 | 0 | 0 | 0 |
| WP_005166365.1 | Y11_RS04010 | Y11_08661 | deleted | peptide ABC transporter substrate-binding protein SapA | -0,14 | 15 | 15 | 15 | 15 | 0 |
| Large deletion of 32,673 bp (positions 1039738 to 1072411) - 28 genes |  |  |  |  |  |  |  |  |  |  |
| WP_005164789.1 | Y11_RS04705 | Y11_10131 | deleted | DUF2813 domain-containing protein | 0 | 0 | 0 | 0 | 0 | 0 |
| YP_001006252.1 | Y11_RS04710 |  | deleted | type I secretion system permease/ATPase | 0 | 0 | 0 | 0 | 0 | 0 |
| WP_005164791.1 | Y11_RS04720 |  | deleted | HlyD family type I secretion periplasmic adaptor subunit | 0 | 0 | 0 | 0 | 0 | 0 |
| WP_005164793.1 | Y11_RS04725 | Y11_10171 | deleted | peptidase C39 | 0 | 0 | 0 | 0 | 0 | 0 |
| WP_023161026.1 | Y11_RS04730 | Y11_10191 | deleted | formate--tetrahydrofolate ligase Fhs | -0,72 | 2,88 | 2,59 | 3,6 | 3,29 | -0,3 |
| WP_005164796.1 | Y11_RS04735 | Y11_10201 | deleted | DUF2569 domain-containing protein | 0 | 0 | 0 | 0 | 0 | 0 |
| WP_005164797.1 | Y11_RS04740 | Y11_10211 | deleted | electron transport complex subunit RsxA | 0 | 0 | 0 | 0 | 0 | 0 |
| WP_005164800.1 | Y11_RS04745 | Y11_10221 | deleted | electron transport complex subunit RsxB | 0 | 0 | 0 | 0 | 0 | 0 |
| WP_014608949.1 | Y11_RS04750 | Y11_10231 | deleted | electron transport complex subunit RsxC | 0 | 0 | 0 | 0 | 0 | 0 |
| WP_005164805.1 | Y11_RS04755 |  | deleted | hypothetical protein | 0 | 0 | 0 | 0 | 0 | 0 |
| WP_005164807.1 | Y11_RS04760 | Y11_10261 | deleted | right oriC-binding transcriptional activator | 0 | 0 | 0 | 0 | 0 | 0 |
| WP_020283189.1 | Y11_RS04765 | Y11_10281 | deleted | electron transport complex subunit RsxD | 0 | 0 | 0 | 0 | 0 | 0 |
| WP_005164811.1 | Y11_RS04770 | Y11_10291 | deleted | electron transport complex subunit RsxG | -0,06 | 15 | 15 | 15 | 15 | 0 |
| WP_005164813.1 | Y11_RS04775 | Y11_10301 | deleted | electron transport complex subunit RsxE | -1,57 | 0 | 0 | 15 | 15 | 0 |
| WP_005164815.1 | Y11_RS04780 | Y11_10311 | deleted | endonuclease III Nth | 0,39 | 15 | 4,92 | 15 | 4,54 | 0 |
| WP_016266185.1 | Y11_RS04785 | Y11_10321 | deleted | pore-forming cytotoxin subunit YaxB | -2,57 | 2,34 | 1,78 | 4,89 | 4,35 | 0 |
| WP_005164820.1 | Y11_RS04790 | Y11_10331 | deleted | alpha-xenorhabdolysin family binary toxin YaxA | -2,49 | 2,11 | 2,38 | 4,6 | 4,87 | 0 |
| WP_005166280.1 | Y11_RS04805 | Y11_10361 | deleted | oxidoreductase | 0 | 0 | 0 | 0 | 0 | 0 |
| WP_005178976.1 |  |  | deleted | hypothetical protein |  |  |  |  |  |  |
| WP_005166281.1 | Y11_RS04810 | Y11_10381 | deleted | exoribonuclease II Rnb | 0,07 | 15 | 2,6 | 15 | 2,52 | 0 |
| WP_005166282.1 | Y11_RS04815 | Y11_10401 | deleted | carbon starvation protein A | 2,39 | 2,34 | 15 | -0,05 | 15 | 0 |
| WP_005166289.1 | Y11_RS04820 |  | deleted | DUF466 domain-containing protein | 0 | 0 | 0 | 0 | 0 | 0 |
| WP_005166294.1 | Y11_RS04825 | Y11_10421 | deleted | fructose-6-phosphate aldolase Fsa | -0,6 | 4,69 | 4,86 | 5,3 | 5,46 | 0,18 |
| WP_005166295.1 | Y11_RS04830 | Y11_10431 | deleted | DeoR/GlpR transcriptional regulator YciT | -0,65 | 15 | 15 | 15 | 15 | 0 |
| WP_005166298.1 | Y11_RS04835 | Y11_10441 | deleted | DUF2164 domain-containing protein | -0,77 | 15 | 15 | 15 | 15 | 0 |
| WP_005166300.1 | Y11_RS04840 | Y11_10451 | deleted | L-ribulose-5-phosphate 4-epimerase AraD | 0 | 0 | 0 | 0 | 0 | 0 |
| WP_005166303.1 | Y11_RS04845 | Y11_10461 | deleted | glycine zipper 2TM domain-containing protein OsmB | 0,87 | 15 | 15 | 15 | 15 | 0 |
| WP_002224869.1 |  |  | deleted | gluconate 5-dehydrogenase |  |  |  |  |  |  |
| Large deletion of 2,533 bp (positions 4134104 to 4136637) - 2 genes |  |  |  |  |  |  |  |  |  |  |
| WP_002228219.1 | Y11_RS18810 | Y11_39551 | deleted | polynucleotide adenyllyltransferase PcnB | -0,08 | 4,28 | 4,13 | 4,37 | 4,21 | 0 |
| WP_005156770.1 | Y11_RS18815 | Y11_39561 | deleted | tRNA glutamyl-Q(34) synthetase GluQRS | -0,58 | 15 | 15 | 15 | 15 | 0 |

A positive selection signal was detected in all isolated strains, in the acyl-CoA thioesterase II, which regulates levels of acyl-CoAs, free fatty acids, and coenzyme A. In position 280 of this enzyme, a glycine is normally conserved in diverse eukaryotic and bacterial species^11^ while in Ye.1 to Ye.4 a serine was present. Position 280 is located next to important residues involved in two catalytic triads (Q278 together with N204 and T228, as well as E279 together with H58 and S107) and a dimer interface (V277 and V281) and could possibly affect the catalytic and functional activities of the acyl-Coa thioesterase. Another positive selection signal was present in a phosphate transporter, for which the C-terminal region displayed a deletion observed only in Ye.3 and Ye.4 that affects two cytoplasmic and two transmembrane domains. While we did not investigate the specific contribution of this deletion to the functionality of the phosphate transporter, the positive selection signal suggests that the promoted changes provided a survival advantage (Table S5).

The genetic basis of antibiotic resistance was investigated upon bacterial genome sequencing. Comparison of *gyrA* sequences of Ye.1 to Ye.4 with those of the reference strain Y11 and strain IP38477, which are susceptible to nalidixic acid, showed that Ye.3 and Ye.4 displayed the same 3-nucleotide deletion at positions 245 to 247 in the quinolone resistance-determining region (QRDR) of *gyrA* (Fig S3A). This so far undescribed deletion leads to replacement of aspartic acid 82 and serine 83 with a unique glycine without any frameshift, explaining the genetic basis of the resistance to quinolones in Ye.3 and Ye.4 (Fig S3B). Additionally, a nucleotide substitution introduced a stop codon in position 78 of the Ye.4 OmpF porin, truncating the protein at one fifth of its length. Porin mutations are usually associated with a decreased antibiotics uptake through the outer-membrane and could explain cefoxitin and ceftriaxone resistance.

### Comparative proteomics of *Ye* isolates reveals an extensive metabolic rewiring, reduced virulence determinant expression and possibly OmpF-mediated antibiotics resistance

To further characterize *Ye* evolution during the course of infection, we analyzed proteomes of the four isolates by mass spectrometry. We measured between 2,086 and 2,287 proteins in the 20 samples (5 replicates of the 4 strains), for a total of 2,408 different proteins identified among all samples against a total database containing 4,424 unique proteins of the 4 isolates. All pairwise comparisons for protein differential abundance between strains were performed. 277 proteins were differentially abundant between the two early isolates Ye.1 and Ye.2 and 323 proteins between the two late isolates Ye.3 and Ye.4. In contrast, 845 to 1,070 proteins were differentially abundant in the 4 pairwise comparisons of early versus late isolates, which account for almost half of the identified proteome (Dataset S1). These proteome differences correlate with the genetic distances of early and late isolates.

The annotated proteins from Ye.1 to Ye.4 were mapped to the *Ye* genotype 4 reference strain Y11 (Table S6). Gene set enrichment analysis (GSEA) on biological process gene ontology (GO) terms revealed isolate-specific variations as well as common trends in early versus lates isolate comparisons (Fig 3). The loss of the pYV virulence plasmid (probably a laboratory artifact) in all strains but Ye.1 could be observed through the enrichment of the type III secretion system term in Ye.1. Ye.2 was specifically characterized by an enriched galactose metabolism. Translation and tRNA processing terms were strongly upregulated in late isolates and even more in Ye.4. Amino acid biosynthetic process was the most specific enriched term in late isolates versus early isolates. Mapping of the protein abundance comparisons on the global metabolic map of Y11 (Fig S4, S5) and on specific KEGG pathways (Fig S6) shows a strong metabolic rewiring from early to late isolates, such as a decreased purine, pyrimidine or riboflavin metabolism, one-carbon pool by folate, carbohydrate metabolism and oxidative phosphorylation, and a perturbated amino acid metabolism such as increased histidine, serine, tryptophan or leucine biosynthesis, or reduced arginine or methionine biosynthesis and glycine cleavage (Fig S6).

**Figure 3.**
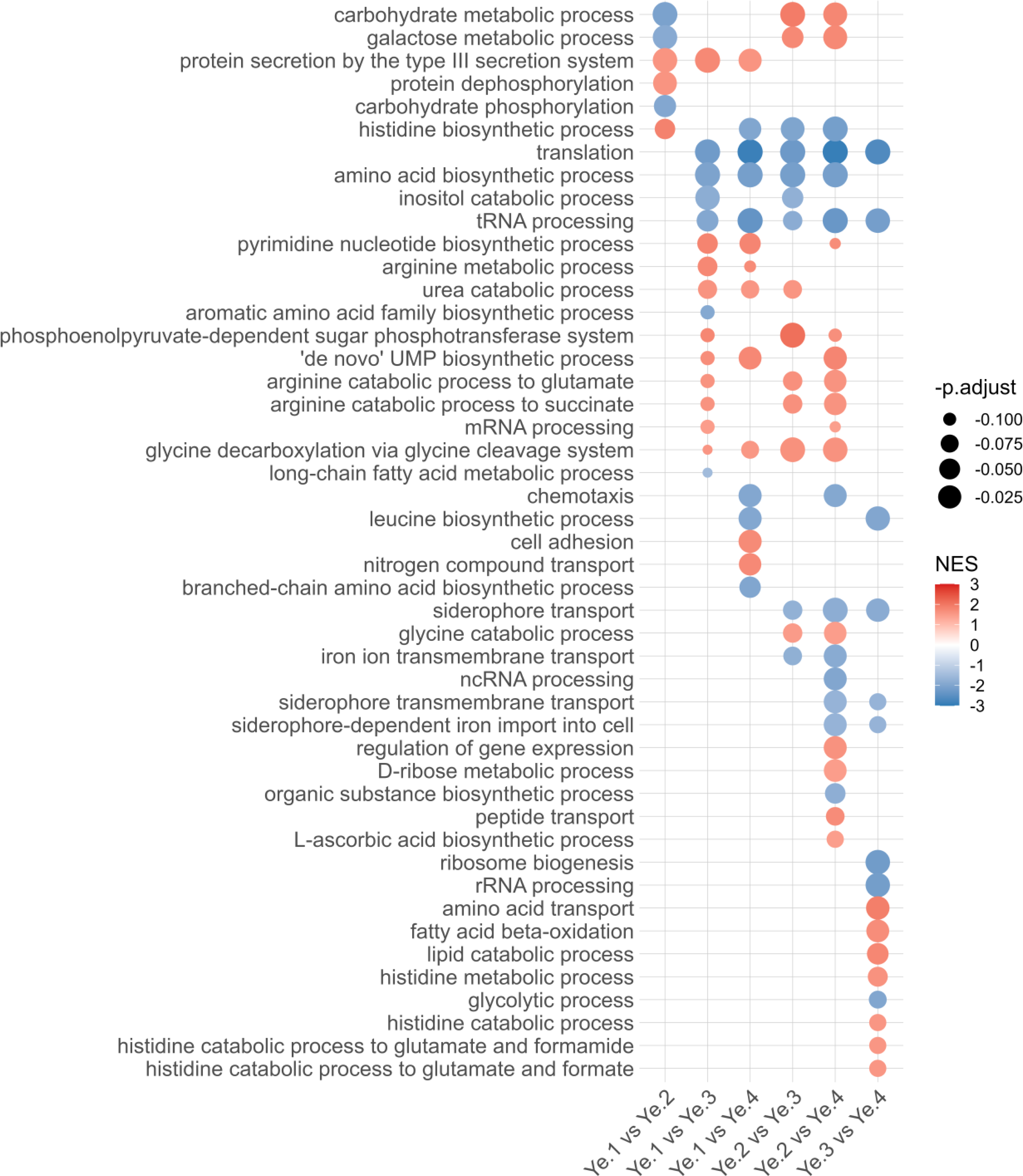
Gene set enrichment analysis (GSEA) of biological process gene ontology terms across pairwise comparisons of *Ye* proteomes. Positive normalized enrichment scores (NES) are shown in red and correspond to enriched terms in the first strain in the comparison, while negative NES are shown in blue and correspond to enriched term in the second strain in the comparison. Multiple testing is adjusted by the Benjamini-Hochberg procedure.

Virulence-associated proteins such as the attachment locus Ail, the fimbriae MyfA, the serine protease ecotin, superoxide dismutases or the positive regulator RovA were less abundant in late isolates, correlating with the increased abundance of the negative virulence regulators RovM or YmoA (Table 3). Additionally, a mutation in the initiation codon of the invasin Inv completely abrogated its production in Ye.4. On the other hand, the chemotaxic operon *cheAWYZ* was more expressed in late isolates, along an increased expression of quorum sensing autoinducer-1 *yenI* and a decreased expression of autoinducer-2 operon *lsrGFBDCARK*. The invasin Inv was only slightly downregulated in Ye.3 but strongly in Ye.4 compared to early isolates.

**Table 3.** Log2(foldchange) of the proteome pairwise comparisons for selected proteins. Proteins were mapped on the Y11 reference genome. P-value and adjusted p-value can be found in Table S6. 0 corresponds to proteins not detected in any conditions of the comparison. +15 and -15 corresponds to proteins detected only in the first or second condition of the comparison, respectively.

|  |  |  |  | Log2(foldchange) |  |  |  |  |  |
| --- | --- | --- | --- | --- | --- | --- | --- | --- | --- |
| Locus | Old Locus | Gene | Product | Ye.1 | Ye.1 | Ye.1 | Ye.2 | Ye.2 | Ye.3 |
|  |  |  |  | vs | vs | vs | vs | vs | vs |
|  |  |  |  | Ye.2 | Ye.3 | Ye.4 | Ye.3 | Ye.4 | Ye.4 |
| Virulence-associated proteins |  |  |  |  |  |  |  |  |  |
| Y11_RS00110 | Y11_00221 | ail | attachment invasion locus protein Ail | 0 | 2,62 | 3,25 | 2,63 | 3,25 | 0,62 |
| Y11_RS01220 | Y11_02591 | eco | serine protease inhibitor ecotin | -0,05 | 1,35 | 1,66 | 1,39 | 1,71 | 0,32 |
| Y11_RS06755 | Y11_14481 | inv | autotransporter invasin Inv | 0,53 | 1,28 | 4,99 | 0,75 | 4,47 | 3,71 |
| Y11_RS01540 | Y11_03301 | myfA | fimbrial polyadhesin major subunit MyfA | -0,27 | 0,62 | 1,18 | 0,89 | 1,47 | 0,57 |
| Y11_RS01020 | Y11_02141 | rovM | virulence transcriptional regulator RovM | 1,02 | -1,57 | -2,01 | -2,59 | -3,03 | -0,44 |
| Y11_RS14390 | Y11_30461 | sodA | superoxide dismutase [Mn] | 0,33 | 1,09 | 1,13 | 0,75 | 0,79 | 0,04 |
| Y11_RS03855 | Y11_08321 | sodB | superoxide dismutase [Fe] | 0,06 | 1,73 | 1,96 | 1,67 | 1,91 | 0,23 |
| Y11_RS10840 | Y11_22981 | sodC | superoxide dismutase family protein [Cu-Zn] | 0,7 | 3,08 | 3,5 | 2,38 | 2,8 | 0,42 |
| Y11_RS09565 | Y11_20331 | ymoA | expression modulating protein YmoA | -0,19 | -2,41 | -3 | -2,22 | -2,81 | -0,59 |
| Chemotaxis proteins |  |  |  |  |  |  |  |  |  |
| Y11_RS06745 | Y11_14461 | flgM | anti-sigma-28 factor FlgM | -0,56 | -1,01 | -1,56 | -0,44 | -1 | -0,55 |
| Y11_RS06785 | Y11_14551 | cheZ | protein phosphatase CheZ | -0,6 | -2,08 | -2,86 | -1,48 | -2,26 | -0,78 |
| Y11_RS06790 | Y11_14561 | cheY | chemotaxis response regulator CheY | -0,06 | -1,74 | -1,96 | -1,68 | -1,9 | -0,22 |
| Y11_RS06795 | Y11_14571 |  | chemotaxis response regulator protein-glutamate methylesterase | 0 | -0,34 | -0,14 | -0,85 | -0,76 | 0,15 |
| Y11_RS06805 | Y11_14591 |  | methyl-accepting chemotaxis protein | 0 | -15 | -15 | -2,19 | -2,48 | -0,31 |
| Y11_RS06815 | Y11_14621 | cheW | chemotaxis protein CheW | 0 | -2,6 | -3,26 | -2,59 | -3,26 | -0,67 |
| Y11_RS06820 | Y11_14641 | cheA | chemotaxis protein CheA | 0 | -0,75 | -2,55 | -0,94 | -2,71 | -1,8 |
| Y11_RS11490 | Y11_24361 | flgM | flagellar biosynthesis anti-sigma factor FlgM | 0,08 | 1,54 | 1,43 | 1,45 | 1,35 | -0,11 |
| Quorum sensing proteins |  |  |  |  |  |  |  |  |  |
| Y11_RS02270 | Y11_04881 | yenR | LuxR family transcriptional regulator YenR | -0,31 | 0,58 | 1,3 | 0 | 0 | 0 |
| Y11_RS02275 | Y11_04891 | yenI | acyl-homoserine-lactone synthase | -0,36 | -2,42 | -2,17 | -2,06 | -1,79 | 0,26 |
| Y11_RS10410 | Y11_22111 | luxS | S-ribosylhomocysteine lyase | -0,04 | 0,16 | 0,38 | 0,2 | 0,42 | 0,22 |
| Y11_RS17855 | Y11_37571 | lsrG | (4S)-4-hydroxy-5-phosphonooxypentane-2,3-dione isomerase | -0,3 | 1,2 | 1,94 | 1,49 | 2,23 | 0,74 |
| Y11_RS17860 | Y11_37581 | lsrF | 3-hydroxy-5-phosphonooxypentane-2,4-dione thiolase | -0,51 | 2,05 | 2,68 | 2,56 | 3,18 | 0,62 |
| Y11_RS17865 | Y11_37591 | lsrB | autoinducer 2 ABC transporter substrate-binding protein LsrB | 0,03 | 2,81 | 3,84 | 2,78 | 3,81 | 1,03 |
| Y11_RS17870 | Y11_37601 | lsrD | autoinducer 2 ABC transporter permease LsrD | 0 | 0 | 0 | 0 | 0 | 0 |
| Y11_RS17875 | Y11_37611 | lsrC | autoinducer 2 ABC transporter permease LsrC | -0,91 | 0,08 | 15 | 0,98 | 15 | 0 |
| Y11_RS17880 | Y11_37621 | lsrA | autoinducer 2 ABC transporter ATP-binding protein LsrA | -0,45 | 15 | 15 | 15 | 15 | 0 |
| Y11_RS17885 | Y11_37631 | lsrR | transcriptional regulator LsrR | 0,15 | 1,25 | 1,43 | 1,1 | 1,28 | 0,18 |
| Y11_RS17890 | Y11_37641 | lsrK | autoinducer-2 kinase | -0,15 | 0,81 | 1,3 | 0,96 | 1,45 | 0,49 |
| Iron import and storage proteins |  |  |  |  |  |  |  |  |  |
| Y11_RS02915 | Y11_06251 | fcuA | Ferrichrome receptor protein FcuA | 1,51 | -0,54 | -1,43 | -2,04 | -2,94 | -0,9 |
| Y11_RS03015 | Y11_06491 | ftnA | non-heme ferritin | -0,27 | 1,46 | 2,04 | 1,73 | 2,31 | 0,58 |
| Y11_RS05035 | Y11_10881 | tonB | TonB system transport protein TonB | 0,98 | -1,07 | -1,96 | -2,05 | -2,94 | -0,89 |
| Y11_RS05710 | Y11_12361 |  | siderophore-interacting protein | 0,76 | -0,55 | -1,22 | -1,31 | -1,98 | -0,67 |
| Y11_RS08900 | Y11_18931 | fur | ferric iron uptake transcriptional regulator | -0,18 | 0,43 | 1,37 | 0,61 | 1,55 | 0,94 |
| Y11_RS11645 | Y11_24711 | exbD | TonB system transport protein ExbD | 0,37 | -1,41 | -2,68 | -1,78 | -3,06 | -1,28 |
| Y11_RS11650 | Y11_24721 | exbB | tol-pal system-associated acyl-CoA thioesterase | 1,79 | 0,31 | -0,86 | -1,49 | -2,65 | -1,16 |
| Y11_RS15495 | Y11_32861 | bfr | bacterioferritin | 0,68 | 3,87 | 4,66 | 3,19 | 3,98 | 0,79 |
| Y11_RS16005 | Y11_33831 | hemY | protoheme IX biogenesis protein HemY | 0,13 | 1,29 | 1,56 | 1,17 | 1,44 | 0,27 |
| Y11_RS16010 | Y11_33841 | hemX | uroporphyrinogen-III C-methyltransferase | 0,51 | 1,09 | 1,57 | 0,57 | 1,06 | 0,48 |
| Y11_RS16920 | Y11_35641 |  | TonB-dependent siderophore receptor | 0,49 | -2,35 | -3,5 | -2,84 | -3,99 | -1,15 |
| Y11_RS16930 | Y11_35671 | hmuV | Hemin import ATP-binding protein HmuV | 0 | 0 | 0 | 0 | 0 | 0 |
| Y11_RS16935 | Y11_35681 | hmuU | Hemin transport system permease protein HmuU | 0 | 0 | 0 | 0 | 0 | 0 |
| Y11_RS16940 | Y11_35691 | hmuT | hemin ABC transporter substrate-binding protein | 0,88 | -1,2 | -2,82 | -2,08 | -3,7 | -1,62 |
| Y11_RS16945 | Y11_35701 | hmuS | hemin-degrading factor | 0,29 | -2,71 | -4,85 | -2,99 | -5,13 | -2,14 |
| Y11_RS16950 | Y11_35711 | hmuR | TonB-dependent hemoglobin/transferrin/lactoferrin family receptor | 1,85 | -1,42 | -2,95 | -3,27 | -4,79 | -1,52 |
| Y11_RS18015 | Y11_37891 | fecA | Iron(III) dicitrate transport protein FecA | 2 | 0,26 | -0,74 | -1,73 | -2,75 | -1,01 |
| Beta-lactamase associated proteins |  |  |  |  |  |  |  |  |  |
| Y11_RS04605 | Y11_09911 | blaA | class A beta-lactamase BlaA | 0,45 | -1,37 | -3,01 | -1,83 | -3,46 | -1,63 |
| Y11_RS05870 | Y11_12651 | blaB | class C beta-lactamase BlaB (AmpC) | 1,03 | -0,03 | -0,97 | -1,05 | -1,99 | -0,94 |
| Y11_RS05875 | Y11_12661 | ampR | LysR family transcriptional regulator AmpR | 0,46 | -2,15 | -0,65 | -2,6 | -1,11 | 1,49 |
| Y11_RS18675 | Y11_39241 | ampD | 1,6-anhydro-N-acetylmuramyl-L-alanine amidase AmpD | 0 | -0,06 | 0,19 | -0,15 | 0,05 | 0,24 |
| Y11_RS18680 | Y11_39251 | ampE | beta-lactamase regulator AmpE | -0,49 | 0,82 | 2,58 | 1,31 | 3,06 | 1,75 |
| Porins |  |  |  |  |  |  |  |  |  |
| Y11_RS01275 | Y11_02741 | ompC | porin OmpC | -1,48 | 0,9 | -0,15 | 2,38 | 1,34 | -1,04 |
| Y11_RS02065 | Y11_04441 | ompF | porin OmpF | 1,4 | -1,37 | 3,58 | -2,77 | 2,18 | 4,95 |
| Y11_RS02150 | Y11_04631 | ompA | porin OmpA | -0,24 | 0,53 | 0,26 | 0,77 | 0,5 | -0,27 |
| Y11_RS05005 | Y11_10821 | ompW | outer membrane protein OmpW | 0,92 | -2,55 | -1,52 | -3,47 | -2,44 | 1,03 |
| Y11_RS05640 | Y11_12201 |  | porin | 1,22 | -2,84 | -2,99 | -4,07 | -4,19 | -0,15 |
| Y11_RS08265 | Y11_17621 | aqpZ | aquaporin Z | -0,79 | 2,64 | 4,48 | 3,43 | 5,27 | 1,84 |
| Y11_RS08275 | Y11_17651 | ompF2 | porin OmpF2 | 0,19 | -0,96 | -1,92 | -1,15 | -2,12 | -0,96 |
| Y11_RS15180 | Y11_32151 | ompR | two-component system response regulator OmpR | -1,92 | 0,83 | 0,58 | 2,75 | 2,5 | -0,25 |
| Y11_RS15185 | Y11_32161 | envZ | two-component system sensor histidine kinase EnvZ | -1,26 | 0,69 | 0 | 1,95 | 1,26 | -0,69 |
| Efflux pumps |  |  |  |  |  |  |  |  |  |
| Y11_RS00155 | Y11_00321 | acrD | multidrug efflux RND transporter permease AcrD | -0,54 | -2,03 | -0,85 | -1,49 | -0,31 | 1,18 |
| Y11_RS00315 | Y11_00681 | macA | macrolide-specific efflux protein MacA | 0 | -15 | 0 | -15 | 0 | 1,4 |
| Y11_RS09485 | Y11_20131 | acrR | multidrug efflux transporter transcriptional repressor AcrR | 0 | 0 | 0 | 0 | 0 | 0 |
| Y11_RS09490 | Y11_20141 | acrA | efflux RND transporter adaptor subunit AcrA | 0,22 | 0,82 | 0,87 | 0,6 | 0,65 | 0,05 |
| Y11_RS09495 | Y11_20151 | acrB | efflux RND transporter permease AcrB | 0,14 | 1,2 | 1,08 | 1,06 | 0,94 | -0,11 |
| Y11_RS10240 | Y11_21811 | emrA | multidrug efflux MFS transporter periplasmic adaptor subunit EmrA | 0 | -0,33 | -2,77 | -0,39 | -2,86 | -2,45 |
| Y11_RS10245 | Y11_21821 | emrB | multidrug efflux MFS transporter permease subunit EmrB | 0 | 0 | 0 | 0 | 0 | 0 |
| Y11_RS11740 | Y11_24901 | tolC | outer membrane channel protein TolC | 0,05 | -0,18 | -0,18 | -0,23 | -0,23 | 0 |
| Y11_RS18200 | Y11_38261 | robA | MDR efflux pump AcrAB transcriptional activator RobA | -0,86 | 0,76 | 1,2 | 0,13 | 2,06 | 0,44 |
| Y11_RS20600 | Y11_43311 | mdtC | multidrug efflux RND transporter permease subunit MdtC | -0,16 | -0,51 | -0,37 | -0,34 | -0,21 | 0,14 |
| Y11_RS20605 | Y11_43321 | mdtB | MdtB/MuxB family multidrug efflux RND transporter permease subunit | -0,06 | 0,11 | 0,28 | 0,17 | 0,34 | 0,17 |
| Y11_RS20610 | Y11_43331 | mdtA | MdtA/MuxA family multidrug efflux RND transporter periplasmic adaptor subunit | -0,53 | -0,64 | -0,04 | -0,12 | 0,49 | 0,61 |

Iron import systems such as the hemin import system encoded by the *hmuRSTUV* operon or the TonB/ExbB/ExbD transport system were more abundant in the late isolates, while iron storage abundance such as the non-heme ferritin FtnA or the bacterioferritin Bfr was decreased. All these iron metabolism protein changes correlated with a decreased abundance of the iron metabolism repressor Fur, showing a probable adaptation to the iron-scarce environment during *Ye* evolution in the bloodstream.

We looked at the proteins level of resistance and susceptibility factors such as β-lactamase, porin or efflux pump. An upregulation of the category A β-lactamase BlaA was observed for the late strains, negatively correlating with the regulator AmpE. More importantly, the category C BlaB (or AmpC) β-lactamase was more abundant in the ceftriaxone-resistant isolate Ye.4 compared to all other isolates. In addition to its truncation in Ye.4, the porin OmpF was less abundant in Ye.2 compared to Ye.1, probably causing a decreased permeability of the outer-membrane to antibiotics (Table 3). The OmpC porin showed variable abundance levels, correlating with OmpR/EnZ two components system abundance, while other porins such à OmpW and two other porins (Y11_17651 and Y11_12201) were strongly upregulated in the late isolates.

### Truncation of the stringent response regulator DksA is one of the main drivers of proteome changes

To better understand which genetic difference could be the cause of the overall changing proteome, the correlation of the 6 pairwise comparisons was assessed against 57 published *Ye* transcriptome comparisons (Table S7) that we previously gathered on the Yersiniomics database^12^. Interestingly, early vs late isolate comparisons had a correlation coefficient ranging from -0.29 to - 0.43 with the comparisons of a *Ye* WT vs Δ*dksA* mutant and was one of the most correlated experiments in our dataset (Table S8, Fig S7). These transcriptomes originated from a recent RNA-Sequencing study exploring the *Ye* stringent response on *Ye* bioserotype 1B/O:8 in WT, Δ*dksA*, Δ*relA*Δ*spoT and* Δ*dksA*Δ*relA*Δ*spoT* strains^13^. We considered the correlations with our proteomes as high considering the variation in the *Ye* genotype and culture conditions used, as well as the analytic differences between RNA-sequencing based transcriptomics and mass spectrometry-based proteomics. Strikingly, a truncation in *dksA* gene is also observed in our late isolates (Table 2), where only the 72 first amino acids (AA) out of 153 AA remains. This truncation could explain the differential abundance of ribosome-associated proteins such as the ribosome maturation factor RimM, ribosome hibernation promoting factor Hpf or ribosome-associated translation inhibitor RaiA, ultimately leading to ribosomal protein upregulation. tRNA-associated proteins were also more abundant in late isolates in a *dksA*-dependent way (Table S9). Other protein abundance changes correlating with *dksA* inactivation include the more abundant previously mentioned hemin transporter subunits HmuR, HmuS, HmuT, magnesium/cobalt transporter CorA, sulfurtransferase TusA, the shikimate and proline pathway enzyme AroB, AroK, ProA and ProB, DNA-binding transcriptional regulator Fis, two of the porins (Y11_12201, Y11_17651) or the acetate uptake transporter SatP among other. On the other hand, *dksA* inactivation was associated to the downregulation of the previously mentioned *lsr* operon as well as the urease *ureABCDEFGD* and glycogen utilization *glgBXCAP* operons, of the ferritins FtnA and Bfr, stress response protein ElaB, superoxide dismutase SodC, entericidin A/B, glycine cleavage system enzyme GcvT or fatty acid degradation enzymes FadA and FadB (Table S9). Interestingly, *dksA* deletion also correlated with a strong decrease in *acs* and *actP* genes, which were described as truncated in late isolates. Most importantly, the stationary phase sigma factor RpoS (σ^S^) expression was *dksA*-dependent, and probably explain most of the regulated proteins such as ribosomal and metabolic enzymes.

## Discussion

This is the first report of a chronic *Ye* infection case due to a strain evolving in the same patient. Between December 1999 and October 2013, antibiotics were apparently successful in the treatment of five independent *Ye* bacteremia episodes; however, discontinuation of antibiotics was always followed by a new bacteremia. The appearance of vegetations observed by echocardiography in 2006, and their disappearance during therapy, argue for *Ye* growth on the atrial lead of the pacemaker and release into the bloodstream during the antibiotic discontinuation periods. Genomic and proteomic analyses of four *Ye* strains (Ye.1 to Ye.4) isolated from the patient allowed us to identify different mechanisms to resist to various antibiotic classes. Resistance to nalidixic acid was already observed from August 2001 after long-term ciprofloxacin treatment. Resistance to quinolones usually involves mutations in the QRDR of the *gyrA* gene, leading either to (i) a change of glycine 81 into cysteine, (ii) a change of serine 83 into arginine, isoleucine, or cysteine, or (ii) a change of aspartic acid 87 into a tyrosine, asparagine, or glycine^14–17^. In Ye.3 and Ye.4, we observed a previously undescribed deletion of 3 nucleotides in the *gyrA* QRDR, leading to substitution of aspartic acid 82 and serine 83 by a unique glycine. Our work thus identifies a novel mutation in *gyrA* likely leading to quinolone resistance.

After the long-term ciprofloxacin treatment was changed for the C3G ceftriaxone, Ye.4 became resistant to cefoxitin and ceftriaxone by abrogating OmpF porin production, a widely shared strategy to reduce membrane permeability to antibiotics^18^ and shown to be induced in *Escherichia coli* upon exposure to ceftriaxone^19^. A previous work showing OmpF mutations (the highest molecular weight called YOMP-C in this previous work) selected by cefoxitin treatment in *Ye* and leading to increased β-lactam resistance supports the antibiotic import via this porin^20^. β-lactamase induction has long been studied in *Ye*^21^ and a recent *in silico* study suggested an increased BlaB inducibility in *Ye* biotype 4^22^. The class A β-lactamase BlaA and class C β-lactamase BlaB (or AmpC) were more abundant in the Ye.4 isolate, probably playing a synergistic role together with porin inactivation, as both β-lactamases have been shown to be involved in ceftriaxone resistance in *Ye* biotype 1B^23^. Moreover, the non-specific MIC increased for various antibiotics classes in Ye.2 could also be explained by a slightly reduced OmpF expression compared to Ye.1. Interestingly, the macrolide efflux pump MacA was more expressed in Ye.3, which was isolated during the pneumonia treatment with the macrolide spiramycin. However, no specific genetic event could be linked to all these proteome variations.

While no ciprofloxacin resistance was observed, Ye.X (which was isolated after ciprofloxacin treatment), Ye.3 and Ye.4 displayed a severe growth defect *in vitro*, which could account for increased antibiotics tolerance. Two major biological processes contribute indeed to antibiotic tolerance: biofilm formation^24,25^ and slow growth and/or metabolic dormancy^5,26,27^. In the late isolates, we identified mutations in genes involved in metabolism or growth in several gene clusters. The cumulative mutations in genes encoding the RNA polymerase subunits RpoC^28^ and RpoD, the pDNA topoisomerase ParE^29^, the 50S ribosomal protein RplF^30,31^, the rRNA methyltransferase RlmE/RrmJ/FtsJ^32,33^ or the stringent response regulator DksA^34,35^ could account for slow growth. Proteomics analysis of the isolates revealed a strong metabolic rewiring with reduced carbohydrate utilization and reduced purine, pyrimidine and riboflavin folate metabolism, together with decreased respiration, which overall corroborate the slow growth phenotype.

Vegetations detected in the pacemaker argue for the formation of a biofilm by the *Ye* strains. *Ye* biofilm production has been reported on medical devices such as catheters or feeding tubes^25,36^. We identified several mutations in Ye.3 and Ye.4 which can be associated with loss of motility and planktonic growth, favoring biofilm formation: for example, mutation of the flagellar genes *flgD* in both Ye.3 and Ye.4, and *flgG* and *flgA* in Ye.4. Of note, loss-of-function mutations associated with decreased motility have been identified in *Burkholderia spp.* and *P. aeruginosa* isolates from cystic fibrosis patients^2,3^. In addition, flagellar assembly is repressed in adverse and nutrient-poor environments owing to its energetic cost^37^. In this direction, no flagellar proteins were detected for any isolate in our culture conditions, in agreement with the reported low flagellar expression in *Yersinia* species at 37°C^38^. Additionally, a decreased expression or inactivation of adherence and invasion factors to host cells such as Ail, MyfA or Inv could favor adhesion to an abiotic surface by other surface factors. However, proteomics observations regarding higher chemotaxis and quorum sensing AI-1 as well as lower quorum sensing AI-2 protein levels in late isolates prevent us to draw definitive conclusions on biofilm formation capacities of these isolates. A recent study characterizing the *Ye* stringent response also showed reduced motility and biofilm formation in a Δ*dksA* mutant, while a (p)ppGpp mutant Δ*relA*Δ*spoT* had an increased biofilm capability^34^, suggesting that our late isolates, truncated in DksA due to the third large deletion described in these isolates, had reduced abilities to form biofilms. However, these experiment from the literature were performed at 26°C and are not representative of bacterial *in vivo* behavior at 37°C.

Strikingly, late isolates seem more sensitive to various aminoglycosides and fluoroquinolones. DksA truncation could explain this phenotype, as a *dksA* mutation increased susceptibility to ciprofloxacin in *Y. pseudotuberculosis*^35^ and to chloramphenicol and ampicillin in *Ye*^34^. However, this effect is species and condition specific as, in *Acinetobacter baumannii,* conflicting results have been reported concerning ciprofloxacin susceptibility^39,40^. Interestingly, a *dksA* mutation in *Pseudomonas aeruginosa* increased on the other hand survival when exposed to ciprofloxacin, strengthening the interest in studying bacterial tolerance in addition to resistance^41^. Finally, a *rlmE* (or *rrmJ*/*ftsJ*) mutation, found in the late isolates, also led to an increased susceptibility to chloramphenicol, gentamicin, spectinomycin and other antibiotics in *E. coli*^32^.

In addition to the evolutive pressure of antibiotics, *Ye* faced the host immune system, nutritional immunity as well as a specific environment requiring a metabolic switch to adapt to a new lifestyle in the blood^42^. Amino acid requirements such as tryptophan for survival in the blood could explain the perturbated amino acid biosynthesis pathways in the late isolates. Recent studies also highlighted the role of glycine and other amino acid in complement susceptibility, and metabolism adaptation could be a way to better survive the complement cascade^43–45^. Iron scarcity in the bloodstream favored the upregulation of iron transport and reduced abundance of iron storage proteins. Interestingly, these regulations were also observed in the transcriptome of the *dksA* mutant^13^.

DksA regulation of ribosome production, modulation of regulators such as RpoS or Fis or amino acid promoters is well documented in model bacteria^46–51^ and was recently validated in *Ye* by transcriptomics^13^. We could observe these effects in the proteome of the late isolates lacking a functional DksA. DksA regulates the stringent response that the bacteria can experience during nutrient starvation, but is also linked to antibiotic exposure or immune stress^52,53^. The pppGpp pyrophosphatase GppA (or Ppx) was also mutated in the late isolates, probably reinforcing unbalanced (p)ppGpp levels and a dysregulated stringent response. As a substantial number of protein changes are associated with DksA truncation and that a strong phenotypic change is induced by *dksA* inactivation^13,34,54^, this mutation could have been specifically selected during in-host evolution, for example for the previously described iron transport. However, as the deletion has a pleiotropic impact on the global transcriptome, it does not allow to distinguish a specific selection process from side-effects of the deletion. Moreover, it remains to be deciphered if the main selection pressure was the antibiotic treatment or the adaptation to the host environment, to temper a detrimental stringent response on the long run for example. Additionally, the Ye.4 isolate showed an even closer proteome to a Δ*dksA* or Δ*relA*Δ*spoT* mutants than Ye.3, perceptible in the higher ribosomal content or decreased fatty acid degradation pathway for example, suggesting that ceftriaxone-resistant Ye.4 possess a more dysregulated stringent response which could be due toOmpF inactivation, reducing nutrient import. Alternatively, stringent response being an adaptation mechanism to a varying environment, its inactivation could have been simply selected by the evolution of the bacteria in a controlled and non-changing environment, the blood. This mechanism could synergistically happen along genome size reduction, a hallmark of environment specialization.

Interestingly, loss of the pore-forming toxin YaxAB (*Y11_10331/Y11_10321*) as well as reduced virulence factor abundance such as Ail, MyfA, superoxide dismutases, ecotin, invasin or the regulator RovA suggest that bacteria defective for pathogenic functions that carry a fitness cost can be selected to favor host adaptation, as argued for *Staphylococcus aureus*^31,55,56^. Finally, *Ye* late isolates displayed a mutation in *mfd*, previously showed to be required for diversification and antibiotics resistance^57,58^. This suggest that the late slow-growing isolates accumulated less mutations and evolved even slower than expected, while most of the in-host evolving strains previously described showed increased mutation rate^59^.

In conclusion, we report the unusual clinical case of a patient with iterative bacteremia and endocarditis due to a *Ye* bioserotype 4/O:3 strain evolving over a period of 14 years. The last two isolates, which were simultaneously evolving in the patient, accumulated mutations that allowed antibiotic resistance (a previously unknown substitution conferring resistance to quinolones) and possibly antibiotic tolerance. The latter phenotype was associated with a reduction in genome size and loss of essential functions for growth *in vitro*, while retaining the ability to survive in the host. The last isolated strains caused bacteremia despite the continuity of the antibiotic treatment, highlighting the transition occurring between early onset tolerance and resistance, a critical problem in antibiotic treatment that should urgently be addressed^60–62^. Our findings have implications in case management, as they reinforce the need for early pacemaker removal in addition to antibiotherapy^63^. They may also guide future design of more effective treatments for chronic and biofilm-associated bacterial infections. If the infected material cannot be removed as it was the case in our study, the choice of the right antibiotic is crucial. It must be well tolerated by the patient, be active on the biofilm and should not promote the emergence of resistance nor tolerance. It appears here that long-term ceftriaxone has been well tolerated, however it induced tolerance and resistance to this antibiotic. Investigation of the mechanisms modulating antibiotic tolerance is therefore a critical issue for the treatment of long-term infections.

## Materials and Methods

### Phenotypic characterization

Bacterial strains were identified using the VITEK2 GN card (bioMérieux) and were sent to the *Yersinia* National Reference Laboratory (Institut Pasteur) for species confirmation/characterization as described^64^. For the growth curves, bacteria were precultured in 5 mL lysogeny broth (LB) at 37°C under agitation at 200 rotation per minute (rpm) for 16 to 96 hours. Precultures were diluted to OD_600nm_=0.1, and 2.5 mL were inoculated in 250 mL LB. Incubations were performed either at 28°C or 37°C under agitation at 200 rpm. Optical density at 600 nm was recorded at different time points to monitor growth of the isolates. Mid-exponential phase growth rate was calculated between 2 and 4 hours for Ye.1 and Ye.2, and between 7 and 31 hours for Ye.3 and Ye.4.

### Antibiotic resistance profiling

The antibiotic resistance profiles of the bacterial strains were first determined using VITEK2 (bioMérieux) and by disk diffusion method on Mueller-Hinton agar supplemented with 5% horse blood (bioMérieux) for growth deficient isolates during the course of the clinical case. A more comprehensive antibiotic resistance profiling and minimal inhibitory concentration (MIC) measurement was respectively performed by disk diffusion method and Etest on Mueller-Hinton agar supplemented with 5% horse blood (bioMérieux) for the four strains kept in collection at the end of the clinical case. Interpretation was performed according to the recommendations of the Antibiogram Committee of the French Society for Microbiology.

### Whole genome sequencing and analysis

Bacterial genomes were sequenced and whole genome-based taxonomic assignment was obtained using a 500-gene cgMLST-*Yersinia* as previously described^65^. 229 epidemiologically unrelated *Ye* 4/O:3 strains isolated between 1982 and 2016 were also sequenced and public genomes of 34 additional strains were used for subsequent analysis (Table S2). Paired-end FASTQ files were used for variant calling using the Y11 reference strain (accession number: NC_017564) with Snippy version 4.1.0^66^. Putative recombinogenic regions were detected and masked with Gubbins version 2.3.4^67^. A maximum-likelihood phylogenetic tree was built from an alignment of 4,738 chromosomal SNPs with RAxML version 8.2.8^68^. Tree visualization was created with iTOL^69^.

### Pan-genome phylogenetic analysis

A total of 96 additional *Ye* strains were randomly selected together with the Ye.1-4 for Phylogenetic Analysis by Maximum Likelihood (PAML)^70^. A pan-genome was constructed using PPanGGOLiN^71^. Each gene of the pan-genome (3,824 genes) was translated to proteins and aligned using MAFFT v7.407^72^, allowing to search for genes specific to *Ye* isolates. Back translation to nucleotide was performed to obtain codon alignment, and a phylogenetic tree was built for each gene using IQ-TREE v1.6.7.2^73^. IQ-TREE’s ModelFinder^74^ was used to estimate the best variant of the General Time Reversible (GTR) model. The 3,612 genes for which the strains Ye.3 and Ye.4 are not sisters in the tree were removed. Synapomorphies were analyzed using a dedicated script (https://gitlab.pasteur.fr/GIPhy/findSynapomorphies/). PAML tests detected genes under positive selection pressure.

### Proteomics sample preparation

Ye.1 and Ye.2 strains were grown overnight in LB at 37°C under agitation at 200 rpm. 200 µL of the culture were plated on 5 tryptic soy agar (TSA) plates and incubated for 24 hours at 37°C. Ye.3 and Ye.4 were grown for 3 days in LB at 37°C under agitation at 200 rpm, then 200 µL of the culture were plated on 5 TSA plates and incubated for 48 hours at 37°C. For all Ye strains, the bacteria were resuspended with an inoculation loop from the TSA plates in 1 mL phosphate buffered saline (PBS) and washed two times by centrifugation and resuspension in PBS. Bacterial proteins were extracted as explained previously ^75^. Briefly, bacteria were lysed with 20 μL trifluoroacetic acid (TFA) for 10 minutes and transferred to a sterile 1.5 mL protein low-binding tube. 200 μL of 2M Tris were added, then 24 μL of 100mM tris(2-carboxyethyl)phosphine (TCEP), 400 mM chloroacetamide (CAA) for a final concentration of 10 mM TCEP and 40 mM CAA. Samples were incubated at 95°C for 5 minutes and proteins were quantified using tryptophan fluorescence in a Synergy H1M microplate reader (BioTek) with an excitation wavelength of 280 nm and an emission wavelength of 360 nm. 100 µg of proteins were aliquoted for each sample, and 5 volumes of water were added.

Proteins were digested for 11 hours at 37°C under agitation at 600 rpm on a ThermoMix (Thermo) using Sequencing Grade Modified Trypsin (Promega - V5111) with a 1:50 ratio (enzyme:protein) before stopping the digestion by addition of TFA to reach a final pH lower than 2. 20 µg of digested peptides were desalted using the Stage-Tips method^76^ using C18 Empore disc, eluted with acetonitrile (ACN) 80%, formic acid (FA) 0.1%. Finally, the peptide solutions were speed-vac dried and resuspended in ACN 2%, FA 0.1% buffer. For each sample, absorbance at 280 nm was measured with a Nanodrop^TM^ 2000 spectrophotometer (Thermo Scientific) to inject an equivalent of DO = 1.

For the spectral library preparation for data-independent analysis (DIA), 2 µg of each digested sample were pooled before proceeding to a peptide fractionation. The fractionation was performed with a poly(styrene-divinylbenzene) reverse phase sulfonate (SDB-RPS) Stage-Tips method as previously described ^76,77^. 20µg of the pooled sample was loaded into 3 SDB-RPS (Empore^TM^, 66886-U) discs stacked on a P200 tip and 8 serial elutions were applied as following: elution buffer 1 (Ammonium formate (AmF) 60 mM, ACN 20%, FA 0.5%), elution buffer 2 (AmF 80 mM, ACN 30%, FA 0.5%), elution buffer 3 (AmF 95 mM, ACN 40%, FA 0.5%), elution buffer 4 (AmF 110 mM, ACN 50%, FA 0.5%), elution buffer 5 (AmF 130 mM, ACN 60%, FA 0.5%), elution buffer 6 (AmF 150 mM, ACN 70%, FA 0.5%) and elution buffer 7 (ACN 80%, ammonium hydroxide 5%). All fractions were speed-vac dried and resuspended with ACN 2%, FA 0.1% before injection. For all fractions and individual samples, iRT peptides were spiked as recommended by Biognosys (Biognosys - Ki-3002-1).

### LC-MS/MS proteomics

For the creation of the spectral library, a nanochromatographic system (Proxeon EASY-nLC 1200 - Thermo Scientific) was coupled online with a Q Exactive^TM^ HF mass spectrometer (Thermo Scientific). Peptides from each fraction and from the pooled sample were injected into a capillary column picotip silica emitter tip (home-made column, 39cm x 75 µm ID, 1.9 µm particles, 100 Å pore size, ReproSil-Pur Basic C18 - Dr. Maisch GmbH, Ammerbuch-Entringen, Germany) after an equilibration step in 100 % mobile phase A (H_2_O, FA 0.1%). Peptides were eluted with a multi-step gradient from 2 to 7% mobile phase B (ACN 80% / FA 0.1%) in 5 min, 7 to 23% B in 70 min, 23 to 45% B in 30 min and 45 to 95% B in 5 min at a flow rate of 250 nL/min for up to 132 min. Column temperature was set to 60°C. Mass spectra were acquired using Xcalibur software using a data-dependent Top 10 method with a survey scans (300-1700 m/z) at a resolution of 60,000 and MS/MS scans (fixed first mass 100 m/z) at a resolution of 15,000. The AGC target and maximum injection time for the survey scans and the MS/MS scans were set to 3.0×10^6^, 100 ms and 1.0×10^5^, 45 ms respectively. The isolation window was set to 1.6 m/z and normalized collision energy fixed to 28 for HCD fragmentation. We used a minimum AGC target of 2.0×10^3^ for an intensity threshold of 4.4×10^4^. Unassigned precursor ion charge states as well as 1, 7, 8 and >8 charged states were rejected and peptide match was disable. Exclude isotopes was enabled and selected ions were dynamically excluded for 30 seconds. For the DIA analysis, mass spectra were acquired in DIA mode with the same nanochromatographic system coupled on-line to a Q Exactive^TM^ HF mass spectrometer. For each sample, 1 μg of peptides was injected into the same capillary column picotip silica emitter tip after an equilibration step in 100 % mobile phase A (H_2_O, FA 0.1%). Peptides were eluted using the same multi-step gradient and temperature for up to 125min. MS data was acquired using the Xcalibur software with a scan range from 295 to 1170 m/z. The DIA method consisted in a succession of one MS scan at a resolution of 60,000 and 36 consecutive MS/MS scans with 1 Da overlapping windows (isolation window = 25 m/z) at 30,000 resolution. The AGC (Automatic Gain Control) target and maximum injection time for MS and MS/MS scans were set to 3.0×10^6^, 60 ms and 2.0×10^5^, auto respectively. The normalized collision energy was set to 28 for HCD fragmentation.

### Proteome bioinformatic analysis

MaxQuant analyses: DDA raw files were processes using MaxQuant software version 1.6.6.0^78^ with Andromeda search engine^79^. The MS/MS spectra were searched against a personal *Yersinia enterocolitica* database (4,424 entries). The database was created by combining the four different *Yersinia enterocolitica* strains (Ye.1, Ye.2, Ye.3 and Ye.4). Sequences with at least one different amino acid were considered as unique and specific of the corresponding strain. Sequences with 100% similarity were reported once. FASTA headers contains the traceability of each strain.

Variable modifications (methionine oxidation and N-terminal acetylation) and fixed modification (cysteine carbamidomethylation) were set for the search and trypsin with a maximum of two missed cleavages was chosen. The minimum peptide length was set to 7 amino acids and the false discovery rate (FDR) for peptide and protein identification was set to 0.01. The main search peptide tolerance was set to 4.5 ppm and to 20 ppm for the MS/MS match tolerance. Second peptides were enabled to identify co-fragmentation events.

Spectronaut analyses: Spectronaut v. 16.0.220606 (Biognosys AG)^80^ was used for DIA-MS data analyses. Data extraction was performed using the default BGS Factory Settings. Briefly, for identification, both precursor and protein FDR were controlled at 1%. For quantification, Qvalue was used for precursor filtering and global imputation strategy was used; peptides were grouped based on stripped sequences. Cross Run Normalization was enabled.

### Proteome statistical analysis

To find the proteins more abundant in one condition than in another, the intensities quantified using Spectronaut were compared. Only proteins identified with at least one peptide and with at least four intensity values in one of the two compared conditions were kept for further statistics. First, proteins absent in a condition and present in another are put aside. These proteins can directly be assumed differentially abundant between the conditions. Then, intensities of the remaining proteins were first log-transformed (log2). Intensity values were normalized by median centering within conditions (section 3.5 in this reference^81^). Missing values were imputed using the impute.slsa function of the R package imp4p^82^. Statistical testing was conducted using a limma t-test thanks to the R package limma^83^. An adaptive Benjamini-Hochberg procedure was applied on the resulting p-values thanks to the function adjust.p of the cp4p R package^84^ using the robust method described in this reference^85^ to estimate the proportion of true null hypotheses among the set of statistical tests. The proteins associated to an adjusted p-value inferior to a FDR level of 1% and an absolute log_2_(fold-change) superior to 1 have been considered as significant and differentially abundant proteins. Finally, the proteins of interest are therefore those which emerge from this statistical analysis supplemented by those which are present from one condition and absent in another (“ID table” in the S1 Dataset). Annotated proteins of Ye.1 to Ye.4 strains were blasted on the Y11 reference proteome (RefSeq annotation from December 2022, accession GCF_000253175.1), and the best hit for each Y11 protein was kept. For each pairwise comparison based on the Ye.1 to Ye.4 loci database describe above, the duplicated Y11 loci were removed by keeping the less differentially expressed value, assuming the duplicated loci came from an annotation of a protein carrying a SNP. The R package clusterProfiler v4.6.2 was used for enrichment and gene set enrichment analysis^86^. The GSEA function was used based on the Y11 loci by mapping to an in-house annotation file extracted from the Y11 Gene Ontology terms downloaded from the QuickGo website accessed on May 25^th^ 2023^87^. The gseKEGG function was used with the online KEGG dataset for the “yey” organism accessed on May 25^th^ 2023. Analysis was performed with the detected proteins as statistical background. Foldchange data were transformed to be uploaded on the iPath3 website^88^. Foldchange data were also mapped to specific KEGG pathways using the R package pathview^89^. Published transcriptomes were downloaded from the Yersiniomics website ^12^. *Ye* 8081, Y1 and Y11 reference genomes were blasted to find homologous loci, and proteomes and transcriptomes were fused in a unique dataframe. The rquery.cormat function was used to compute the Pearson correlations of the foldchanges and display the correlation matrix.

## Data Availability Section

The datasets and computer code produced in this study are available in the following databases:

- Genome sequences: ENA PRJEB19854 (sample ID starting with ERS and accession numbers starting with FWC) *for Y. enterocolitica* 1 (Ye.1) (ERS1580350, FWCF02000001-FWCF02000181), Ye.2 (ERS1580351, FWCE02000001-FWCE02000227), Ye.3 (ERS1580352, FWCB02000001-FWCB02000195), and Ye.4 (ERS1580353, FWCD02000001-FWCD02000173)
- The mass spectrometry proteomics data have been deposited to the ProteomeXchange Consortium via the PRIDE partner repository^90^ with the dataset identifier PXD043567

## Acknowledgements

This work was funded by Santé Publique France (SpF, Saint-Maurice, France), Institut Pasteur, Direction Générale de l’Armement, Agence Innovation Défense, Fondation pour la Recherche Médicale (FDT20220401-711 5222), the Inception program (Investissement d’Avenir grant ANR-16-CONV-0005), ANRS Emerging Infectious Disease (ANRS0349b) and Université Paris Cité. The *Yersinia* Research Unit is a member of the LabEX IBEID (ANR-10LBX-62-IBEID). We thank Vincent Enouf and Andreea Alexandru (P2M platform, Institut Pasteur, Paris, France) for the sequencing of the strains, Daniel Boury (SID’COM, University of Tours) for transesophageal echocardiography image processing, and Pr. Frédéric Patat (University Hospital of Tours) for transesophageal echocardiography image interpretation.

Some figures were created with BioRender.

## AUTHOR CONTRIBUTIONS

C.S. and P.L.-B. planned, performed and analyzed experiments with the help of L.M., N.C. and L.G. J.G. set up and performed pan-genome together with PAML analysis. R.B., C.L.-B., F.B., M.F. and B.M. are the physicians who took care of the patient and shared all the clinico-epidemiological data. L.D. performed antibiograms and MIC measurement. T.D. performed mass spectrometry-based proteomics. P.L.-B. and O.D. were involved in proteome analysis and identification of antibiotic tolerance mechanism. C.S., P.L.-B. and J.P.-C. interpreted data and wrote the manuscript. E.C., O.D., M. M., P.L. and J.P.-C. supervised the work. C.S. and P.L.-B. are shared first authors and P.L. and J.P.-C. are shared last authors.

