## Supplemental Files for "In-host evolution of *Yersinia enterocolitica* during a chronic human infection"

**Figure S1.** Vegetation on pacemaker lead. Transesophageal echocardiography performed July 27th, 2006, showing large vegetations on the pacemaker lead in the right atrium. Vegetations formed a sheath around the electrode, measuring 1.6cm by 0.8cm

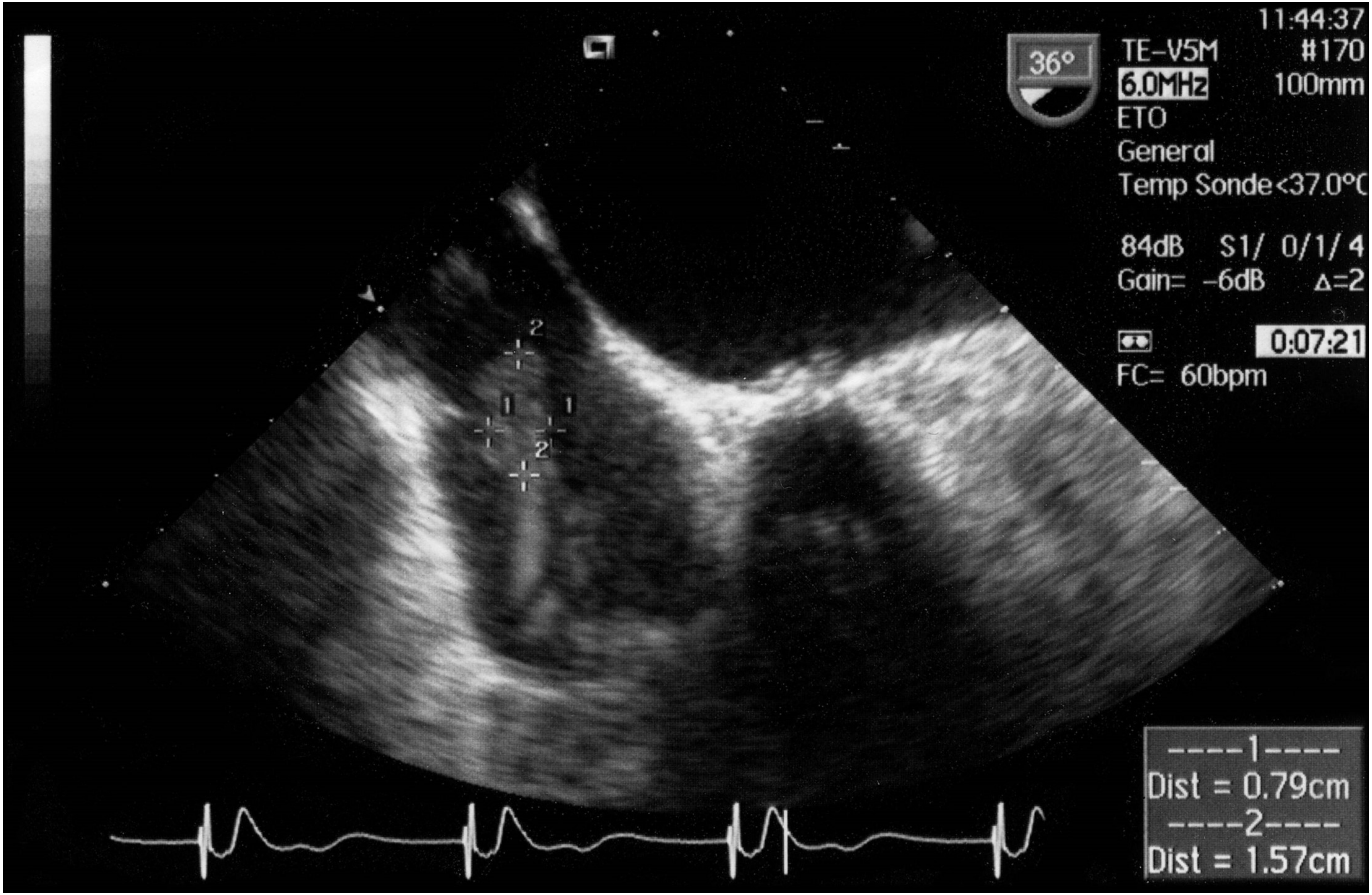

**Figure S2.** Growth curves of the 4 isolates in lysogeny broth (LB) at 28°C and 37°C, plotted (A) until 72 hours and (B) until 216 hours.

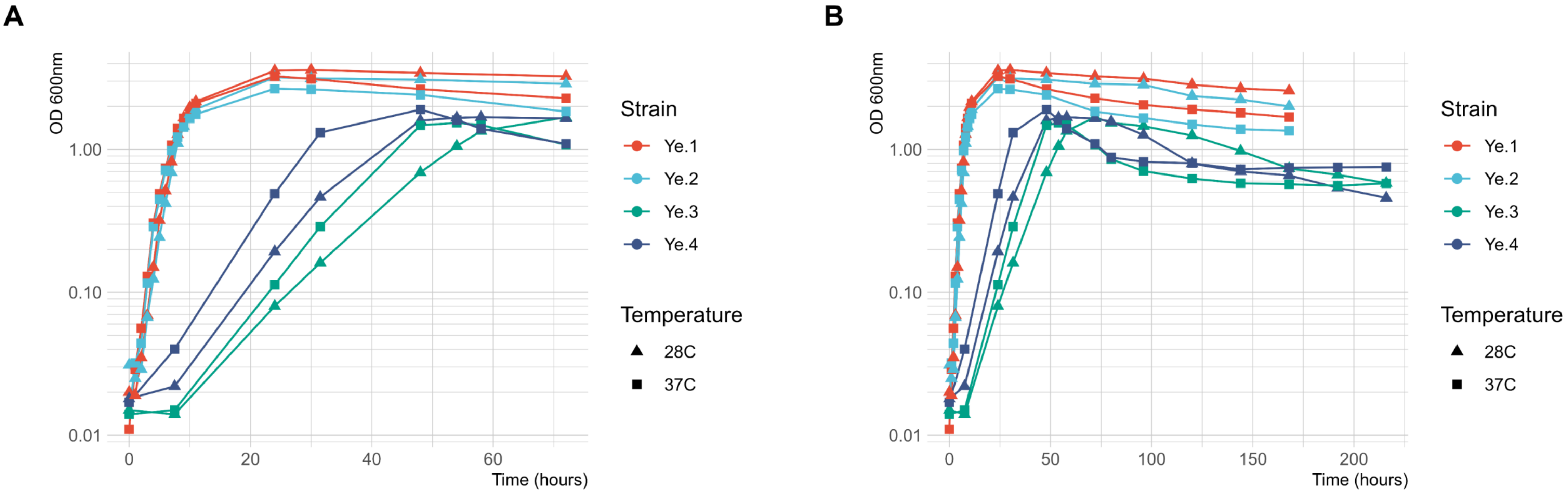

**Figure S3.** Alignments for (A) the *gyrA* gene nucleotide sequences and (B) the predicted *gyrA* product amino acid sequences, from the 4 patient strains, the Y11 reference strain and IP38477 (sensitive to nalidixic acid).

**A.**

```
Y11         ATGAGCGACCTTGCCAGAGAAATAACACCGGTCAACATCGAGGAAGAGCTGAAAAGCTCC   60
IP38477     ATGAGCGACCTTGCCAGAGAAATAACACCGGTCAACATCGAGGAAGAGCTGAAAAGCTCC   60
Ye.1        ATGAGCGACCTTGCCAGAGAAATAACACCGGTCAACATCGAGGAAGAGCTGAAAAGCTCC   60
Ye.2        ATGAGCGACCTTGCCAGAGAAATAACACCGGTCAACATCGAGGAAGAGCTGAAAAGCTCC   60
Ye.3        ATGAGCGACCTTGCCAGAGAAATAACACCGGTCAACATCGAGGAAGAGCTGAAAAGCTCC   60
Ye.4        ATGAGCGACCTTGCCAGAGAAATAACACCGGTCAACATCGAGGAAGAGCTGAAAAGCTCC   60
            ************************************************************

Y11         TATCTGGATTATGCGATGTCCGTTATTGTCGGACGTGCTTTGCCAGATGTCCGGGATGGA   120
IP38477     TATCTGGATTATGCGATGTCCGTTATTGTCGGACGTGCTTTGCCAGATGTCCGGGATGGA   120
Ye.1        TATCTGGATTATGCGATGTCCGTTATTGTCGGACGTGCTTTGCCAGATGTCCGGGATGGA   120
Ye.2        TATCTGGATTATGCGATGTCCGTTATTGTCGGACGTGCTTTGCCAGATGTCCGGGATGGA   120
Ye.3        TATCTGGATTATGCGATGTCCGTTATTGTCGGACGTGCTTTGCCAGATGTCCGGGATGGA   120
Ye.4        TATCTGGATTATGCGATGTCCGTTATTGTCGGACGTGCTTTGCCAGATGTCCGGGATGGA   120
            ************************************************************

Y11         CTGAAACCGGTGCACCGTCGCGTACTGTATGCGATGAATGTACTGGGTAATGACTGGAAT   180
IP38477     CTGAAACCGGTGCACCGTCGCGTACTGTATGCGATGAATGTACTGGGTAATGACTGGAAT   180
Ye.1        CTGAAACCGGTGCACCGTCGCGTACTGTATGCGATGAATGTACTGGGTAATGACTGGAAT   180
Ye.2        CTGAAACCGGTGCACCGTCGCGTACTGTATGCGATGAATGTACTGGGTAATGACTGGAAT   180
Ye.3        CTGAAACCGGTGCACCGTCGCGTACTGTATGCGATGAATGTACTGGGTAATGACTGGAAT   180
Ye.4        CTGAAACCGGTGCACCGTCGCGTACTGTATGCGATGAATGTACTGGGTAATGACTGGAAT   180
            ************************************************************

Y11         AAACCATACAAAAAAATCGGCCCGTGTAGTCGGGGACGTTATCGGTAAATATCACCCGCAT   240
IP38477     AAACCATACAAAAAAATCGGCCCGTGTAGTCGGGGACGTTATCGGTAAATATCACCCGCAT   240
Ye.1        AAACCATACAAAAAAATCGGCCCGTGTAGTCGGGGACGTTATCGGTAAATATCACCCGCAT   240
Ye.2        AAACCATACAAAAAAATCGGCCCGTGTAGTCGGGGACGTTATCGGTAAATATCACCCGCAT   240
Ye.3        AAACCATACAAAAAAATCGGCCCGTGTAGTCGGGGACGTTATCGGTAAATATCACCCGCAT   240
Ye.4        AAACCATACAAAAAAATCGGCCCGTGTAGTCGGGGACGTTATCGGTAAATATCACCCGCAT   240
            ************************************************************

Y11         GGTGACAGCGCGGTCTACGACACTATAGTGCGTATGGCCCAGCCGTTCTCACTGCGCTAT   300
IP38477     GGTGACAGCGCGGTCTACGACACTATAGTGCGTATGGCCCAGCCGTTCTCACTGCGCTAT   300
Ye.1        GGTGACAGCGCGGTCTACGACACTATAGTGCGTATGGCCCAGCCGTTCTCACTGCGCTAT   300
Ye.2        GGTGACAGCGCGGTCTACGACACTATAGTGCGTATGGCCCAGCCGTTCTCACTGCGCTAT   300
Ye.3        GGTG---GCGCGGTCTACGACACTATAGTGCGTATGGCCCAGCCGTTCTCACTGCGCTAT   297
Ye.4        GGTG---GCGCGGTCTACAACACTATAGTGCGTATGGCCCAGCCGTTCTCACTGCGCTAT   297
            ****   *********** *****************************************

Y11         ATGCTGGTGGATGGGCAGGGTAACTTCGGTTCCGTTGATGGCGACTCCGCCGCAGCGATG   360
IP38477     ATGCTGGTGGATGGGCAGGGTAACTTCGGTTCCGTTGATGGCGACTCCGCCGCAGCGATG   360
Ye.1        ATGCTGGTGGATGGGCAGGGTAACTTCGGTTCCGTTGATGGCGACTCCGCCGCAGCGATG   360
Ye.2        ATGCTGGTGGATGGGCAGGGTAACTTCGGTTCCGTTGATGGCGACTCCGCCGCAGCGATG   360
Ye.3        ATGCTGGTGGATGGGCAGGGTAACTTCGGTTCCGTTGATGGCGACTCCGCCGCAGCGATG   357
Ye.4        ATGCTGGTGGATGGGCAGGGTAACTTCGGTTCCGTTGATGGCGACTCCGCCGCAGCGATG   357
            ************************************************************

Y11         CGTTATACCGAAATCCGTATGTCTAAAATTGCTCACGAACTGTTGGCGGACTTAGAAAAA   420
IP38477     CGTTATACCGAAATCCGTATGTCTAAAATTGCTCACGAACTGTTGGCGGACTTAGAAAAA   420
Ye.1        CGTTATACCGAAATCCGTATGTCTAAAATTGCTCACGAACTGTTGGCGGACTTAGAAAAA   420
Ye.2        CGTTATACCGAAATCCGTATGTCTAAAATTGCTCACGAACTGTTGGCGGACTTAGAAAAA   420
```

```
Ye.3       CGTTATACCGAAATCCGTATGTCTAAAATTGCTCACGAACTGTTGGCGGACTTAGAAAAA   417
Ye.4       CGTTATACCGAAATCCGTATGTCTAAAATTGCTCACGAACTGTTGGCGGACTTAGAAAAA   417
           ************************************************************

Y11        GATACCGTTGACTTCGTGCCGAACTATGACGGTACGGAGCAAATTCCTGCCGTAATGCCG   480
IP38477    GATACCGTTGACTTCGTGCCGAACTATGACGGTACGGAGCAAATTCCTGCCGTAATGCCG   480
Ye.1       GATACCGTTGACTTCGTGCCGAACTATGACGGTACGGAGCAAATTCCTGCCGTAATGCCG   480
Ye.2       GATACCGTTGACTTCGTGCCGAACTATGACGGTACGGAGCAAATTCCTGCCGTAATGCCG   480
Ye.3       GATACCGTTGACTTCGTGCCGAACTATGACGGTACGGAGCAAATTCCTGCCGTAATGCCG   477
Ye.4       GATACCGTTGACTTCGTGCCGAACTATGACGGTACGGAGCAAATTCCTGCCGTAATGCCG   477
           ************************************************************

Y11        ACCAAAATCCCTAACTTGCTGGTGAACGGCTCGTCAGGTATTGCCGTCGGGATGGCAACC   540
IP38477    ACCAAAATCCCTAACTTGCTGGTGAACGGCTCGTCAGGTATTGCCGTCGGGATGGCAACC   540
Ye.1       ACCAAAATCCCTAACTTGCTGGTGAACGGCTCGTCAGGTATTGCCGTCGGGATGGCAACC   540
Ye.2       ACCAAAATCCCTAACTTGCTGGTGAACGGCTCGTCAGGTATTGCCGTCGGGATGGCAACC   540
Ye.3       ACCAAAATCCCTAACTTGCTGGTGAACGGCTCGTCAGGTATTGCCGTCGGGATGGCAACC   537
Ye.4       ACCAAAATCCCTAACTTGCTGGTGAACGGCTCGTCAGGTATTGCCGTCGGGATGGCAACC   537
           ************************************************************

Y11        AATATTCCGCCGCATAACCTTTCTGAGGTTATTGATGGCTGTCTGGCCTATATCGAAGAT   600
IP38477    AATATTCCGCCGCATAACCTTTCTGAGGTTATTGATGGCTGTCTGGCCTATATCGAAGAT   600
Ye.1       AATATTCCGCCGCATAACCTTTCTGAGGTTATTGATGGCTGTCTGGCCTATATCGAAGAT   600
Ye.2       AATATTCCGCCGCATAACCTTTCTGAGGTTATTGATGGCTGTCTGGCCTATATCGAAGAT   600
Ye.3       AATATTCCGCCGCATAACCTTTCTGAGGTTATTGATGGCTGTCTGGCCTATATCGAAGAT   597
Ye.4       AATATTCCGCCGCATAACCTTTCTGAGGTTATTGATGGCTGTCTGGCCTATATCGAAGAT   597
           ************************************************************

Y11        GAAAACATCACCATTGAAGGGTTGATGGAGTACATCCCGGGGCCAGATTTCCCCACTGCT   660
IP38477    GAAAACATCACCATTGAAGGGTTGATGGAGTACATCCCGGGGCCAGATTTCCCCACTGCT   660
Ye.1       GAAAACATCACCATTGAAGGGTTGATGGAGTACATCCCGGGGCCAGATTTCCCCACTGCT   660
Ye.2       GAAAACATCACCATTGAAGGGTTGATGGAGTACATCCCGGGGCCAGATTTCCCCACTGCT   660
Ye.3       GAAAACATCACCATTGAAGGGTTGATGGAGTACATCCCGGGGCCAGATTTCCCCACTGCT   657
Ye.4       GAAAACATCACCATTGAAGGGTTGATGGAGTACATCCCGGGGCCAGATTTCCCCACTGCT   657
           ************************************************************

Y11        GCGATTATCAATGGTCGCCGTGGTATTGAAGAAGCTTATCGTACTGGCCGTGGCAAGGTG   720
IP38477    GCGATTATCAATGGTCGCCGTGGTATTGAAGAAGCTTATCGTACTGGCCGTGGCAAGGTG   720
Ye.1       GCGATTATCAATGGTCGCCGTGGTATTGAAGAAGCTTATCGTACTGGCCGTGGCAAGGTG   720
Ye.2       GCGATTATCAATGGTCGCCGTGGTATTGAAGAAGCTTATCGTACTGGCCGTGGCAAGGTG   720
Ye.3       GCGATTATCAATGGTCGCCGTGGTATTGAAGAAGCTTATCGTACTGGCCGTGGCAAGGTG   717
Ye.4       GCGATTATCAATGGTCGCCGTGGTATTGAAGAAGCTTATCGTACTGGCCGTGGCAAGGTG   717
           ************************************************************

Y11        TATATCCGTGCCCGTGCTGAAGTTGAGGCTGACGCTAAAACCGGTCGCGAAACCATTATT   780
IP38477    TATATCCGTGCCCGTGCTGAAGTTGAGGCTGACGCTAAAACCGGTCGCGAAACCATTATT   780
Ye.1       TATATCCGTGCCCGTGCTGAAGTTGAGGCTGACGCTAAAACCGGTCGCGAAACCATTATT   780
Ye.2       TATATCCGTGCCCGTGCTGAAGTTGAGGCTGACGCTAAAACCGGTCGCGAAACCATTATT   780
Ye.3       TATATCCGTGCCCGTGCTGAAGTTGAGGCTGACGCTAAAACCGGTCGCGAAACCATTATT   777
Ye.4       TATATCCGTGCCCGTGCTGAAGTTGAGGCTGACGCTAAAACCGGTCGCGAAACCATTATT   777
           ************************************************************

Y11        GTTCACGAGATCCCGTATCAGGTGAACAAGGCGCGGTTGATTGAAAAAATCGCCGAGCTG   840
IP38477    GTTCACGAGATCCCGTATCAGGTGAACAAGGCGCGGTTGATTGAAAAAATCGCCGAGCTG   840
Ye.1       GTTCACGAGATCCCGTATCAGGTGAACAAGGCGCGGTTGATTGAAAAAATCGCCGAGCTG   840
Ye.2       GTTCACGAGATCCCGTATCAGGTGAACAAGGCGCGGTTGATTGAAAAAATCGCCGAGCTG   840
Ye.3       GTTCACGAGATCCCGTATCAGGTGAACAAGGCGCGGTTGATTGAAAAAATCGCCGAGCTG   837
Ye.4       GTTCACGAGATCCCGTATCAGGTGAACAAGGCGCGGTTGATTGAAAAAATCGCCGAGCTG   837
           ************************************************************

Y11        GTTAAAGAAAAAACGCGTAGAAGGTATCAGTGCGTTGCGTGATGAGTCTGATAAAGACGGC   900
IP38477    GTTAAAGAAAAAACGCGTAGAAGGTATCAGTGCGTTGCGTGATGAGTCTGATAAAGACGGC   900
Ye.1       GTTAAAGAAAAAACGCGTAGAAGGTATCAGTGCGTTGCGTGATGAGTCTGATAAAGACGGC   900
```

```
Ye.2      GTTAAAGAAAAACGCGTAGAAGGTATCAGTGCGTTGCGTGATGAGTCTGATAAAGACGGC   900
Ye.3      GTTAAAGAAAAACGCGTAGAAGGTATCAGTGCGTTGCGTGATGAGTCTGATAAAGACGGC   897
Ye.4      GTTAAAGAAAAACGCGTAGAAGGTATCAGTGCGTTGCGTGATGAGTCTGATAAAGACGGC   897
          ************************************************************

Y11       ATGCGTATCGTGATTGAAATCAAACGTGATGCTGTCGGGGAAGTGGTGCTGAACAACCTC   960
IP38477   ATGCGTATCGTGATTGAAATCAAACGTGATGCTGTCGGGGAAGTGGTGCTGAACAACCTC   960
Ye.1      ATGCGTATCGTGATTGAAATCAAACGTGATGCTGTCGGGGAAGTGGTGCTGAACAACCTC   960
Ye.2      ATGCGTATCGTGATTGAAATCAAACGTGATGCTGTCGGGGAAGTGGTGCTGAACAACCTC   960
Ye.3      ATGCGTATCGTGATTGAAATCAAACGTGATGCTGTCGGGGAAGTGGTGCTGAACAACCTC   957
Ye.4      ATGCGTATCGTGATTGAAATCAAACGTGATGCTGTCGGGGAAGTGGTGCTGAACAACCTC   957
          ************************************************************

Y11       TACTCTCTGACGCAATTGCAGGTGACTTTCGGTATCAATATGGTGGGTCTGTCTCAAGGG   1020
IP38477   TACTCTCTGACGCAATTGCAGGTGACTTTCGGTATCAATATGGTGGGTCTGTCTCAAGGG   1020
Ye.1      TACTCTCTGACGCAATTGCAGGTGACTTTCGGTATCAATATGGTGGGTCTGTCTCAAGGG   1020
Ye.2      TACTCTCTGACGCAATTGCAGGTGACTTTCGGTATCAATATGGTGGGTCTGTCTCAAGGG   1020
Ye.3      TACTCTCTGACGCAATTGCAGGTGACTTTCGGTATCAATATGGTGGGTCTGTCTCAAGGG   1017
Ye.4      TACTCTCTGACGCAATTGCAGGTGACTTTCGGTATCAATATGGTGGGTCTGTCTCAAGGG   1017
          ************************************************************

Y11       CAGCCTAAGTTGCTTAACCTGAAAGACATTTTGGTTGCTTTCGTGCGCCACCGCCGTGAA   1080
IP38477   CAGCCTAAGTTGCTTAACCTGAAAGACATTTTGGTTGCTTTCGTGCGCCACCGCCGTGAA   1080
Ye.1      CAGCCTAAGTTGCTTAACCTGAAAGACATTTTGGTTGCTTTCGTGCGCCACCGCCGTGAA   1080
Ye.2      CAGCCTAAGTTGCTTAACCTGAAAGACATTTTGGTTGCTTTCGTGCGCCACCGCCGTGAA   1080
Ye.3      CAGCCTAAGTTGCTTAACCTGAAAGACATTTTGGTTGCTTTCGTGCGCCACCGCCGTGAA   1077
Ye.4      CAGCCTAAGTTGCTTAACCTGAAAGACATTTTGGTTGCTTTCGTGCGCCACCGCCGTGAA   1077
          ************************************************************

Y11       GTGGTGACTCGCCGTACCATTTTTGAACTGCGTAAAGCACGTGACCGCGCACATATCCTT   1140
IP38477   GTGGTGACTCGCCGTACCATTTTTGAACTGCGTAAAGCACGTGACCGCGCACATATCCTT   1140
Ye.1      GTGGTGACTCGCCGTACCATTTTTGAACTGCGTAAAGCACGTGACCGCGCACATATCCTT   1140
Ye.2      GTGGTGACTCGCCGTACCATTTTTGAACTGCGTAAAGCACGTGACCGCGCACATATCCTT   1140
Ye.3      GTGGTGACTCGCCGTACCATTTTTGAACTGCGTAAAGCACGTGACCGCGCACATATCCTT   1137
Ye.4      GTGGTGACTCGCCGTACCATTTTTGAACTGCGTAAAGCACGTGACCGCGCACATATCCTT   1137
          ************************************************************

Y11       GAAGCGCTGGCTATTGCACTGGCTAACATCGATCCGATTATCGAGTTGATCCGCCGTGCA   1200
IP38477   GAAGCGCTGGCTATTGCACTGGCTAACATCGATCCGATTATCGAGTTGATCCGCCGTGCA   1200
Ye.1      GAAGCGCTGGCTATTGCACTGGCTAACATCGATCCGATTATCGAGTTGATCCGCCGTGCA   1200
Ye.2      GAAGCGCTGGCTATTGCACTGGCTAACATCGATCCGATTATCGAGTTGATCCGCCGTGCA   1200
Ye.3      GAAGCGCTGGCTATTGCACTGGCTAACATCGATCCGATTATCGAGTTGATCCGCCGTGCA   1197
Ye.4      GAAGCGCTGGCTATTGCACTGGCTAACATCGATCCGATTATCGAGTTGATCCGCCGTGCA   1197
          ************************************************************

Y11       GCAACACCTGCCGAAGCGAAAGCCGGCTTGATTGCCAGCCCATGGGAGTTAGGTAACGTT   1260
IP38477   GCAACACCTGCCGAAGCGAAAGCCGGCTTGATTGCCAGCCCATGGGAGTTAGGTAACGTT   1260
Ye.1      GCAACACCTGCCGAAGCGAAAGCCGGCTTGATTGCCAGCCCATGGGAGTTAGGTAACGTT   1260
Ye.2      GCAACACCTGCCGAAGCGAAAGCCGGCTTGATTGCCAGCCCATGGGAGTTAGGTAACGTT   1260
Ye.3      GCAACACCTGCCGAAGCGAAAGCCGGCTTGATTGCCAGCCCATGGGAGTTAGGTAACGTT   1257
Ye.4      GCAACACCTGCCGAAGCGAAAGCCGGCTTGATTGCCAGCCCATGGGAGTTAGGTAACGTT   1257
          ************************************************************

Y11       GCAGCCATGTTGGAACGTGCCGGTGGTGATGCTGCCCGCCCTGAATGGCTGGAAGCTGAG   1320
IP38477   GCAGCCATGTTGGAACGTGCCGGTGGTGATGCTGCCCGCCCTGAATGGCTGGAAGCTGAG   1320
Ye.1      GCAGCCATGTTGGAACGTGCCGGTGGTGATGCTGCCCGCCCTGAATGGCTGGAAGCTGAG   1320
Ye.2      GCAGCCATGTTGGAACGTGCCGGTGGTGATGCTGCCCGCCCTGAATGGCTGGAAGCTGAG   1320
Ye.3      GCAGCCATGTTGGAACGTGCCGGTGGTGATGCTGCCCGCCCTGAATGGCTGGAAGCTGAG   1317
Ye.4      GCAGCCATGTTGGAACGTGCCGGTGGTGATGCTGCCCGCCCTGAATGGCTGGAAGCTGAG   1317
          ************************************************************

Y11       TTCGGTATCCGTGACGGCAAATATTATCTCACCGAGCAGCAAGCTCAGGCGATTTTGGAT   1380
IP38477   TTCGGTATCCGTGACGGCAAATATTATCTCACCGAGCAGCAAGCTCAGGCGATTTTGGAT   1380
Ye.1      TTCGGTATCCGTGACGGCAAATATTATCTCACCGAGCAGCAAGCTCAGGCGATTTTGGAT   1380
```

```
Ye.2        TTCGGTATCCGTGACGGCAAATATTATCTCACCGAGCAGCAAGCTCAGGCGATTTTGGAT   1380
Ye.3        TTCGGTATCCGTGACGGCAAATATTATCTCACCGAGCAGCAAGCTCAGGCGATTTTGGAT   1377
Ye.4        TTCGGTATCCGTGACGGCAAATATTATCTCACCGAGCAGCAAGCTCAGGCGATTTTGGAT   1377
            ************************************************************

Y11         CTGCGTTTGCAGAAACTGACCGGCCTGGAGCATGAAAAACTGCTGGATGAGTATAAAGAG   1440
IP38477     CTGCGTTTGCAGAAACTGACCGGCCTGGAGCATGAAAAACTGCTGGATGAGTATAAAGAG   1440
Ye.1        CTGCGTTTGCAGAAACTGACCGGCCTGGAGCATGAAAAACTGCTGGATGAGTATAAAGAG   1440
Ye.2        CTGCGTTTGCAGAAACTGACCGGCCTGGAGCATGAAAAACTGCTGGATGAGTATAAAGAG   1440
Ye.3        CTGCGTTTGCAGAAACTGACCGGCCTGGAGCATGAAAAACTGCTGGATGAGTATAAAGAG   1437
Ye.4        CTGCGTTTGCAGAAACTGACCGGCCTGGAGCATGAAAAACTGCTGGATGAGTATAAAGAG   1437
            ************************************************************

Y11         CTGCTGACTGTCATTGCCGAGCTGATCTTTATTCTGGAAAATCCAGAGCGCTTGATGGAA   1500
IP38477     CTGCTGACTGTCATTGCCGAGCTGATCTTTATTCTGGAAAATCCAGAGCGCTTGATGGAA   1500
Ye.1        CTGCTGACTGTCATTGCCGAGCTGATCTTTATTCTGGAAAATCCAGAGCGCTTGATGGAA   1500
Ye.2        CTGCTGACTGTCATTGCCGAGCTGATCTTTATTCTGGAAAATCCAGAGCGCTTGATGGAA   1500
Ye.3        CTGCTGACTGTCATTGCCGAGCTGATCTTTATTCTGGAAAATCCAGAGCGCTTGATGGAA   1497
Ye.4        CTGCTGACTGTCATTGCCGAGCTGATCTTTATTCTGGAAAATCCAGAGCGCTTGATGGAA   1497
            ************************************************************

Y11         GTTATCCGCGAAGAGTTAGTGGCGATTAAAGAGCTATATAATGATGCGCGCCGTACCGAA   1560
IP38477     GTTATCCGCGAAGAGTTAGTGGCGATTAAAGAGCTATATAATGATGCGCGCCGTACCGAA   1560
Ye.1        GTTATCCGCGAAGAGTTAGTGGCGATTAAAGAGCTATATAATGATGCGCGCCGTACCGAA   1560
Ye.2        GTTATCCGCGAAGAGTTAGTGGCGATTAAAGAGCTATATAATGATGCGCGCCGTACCGAA   1560
Ye.3        GTTATCCGCGAAGAGTTAGTGGCGATTAAAGAGCTATATAATGATGCGCGCCGTACCGAA   1557
Ye.4        GTTATCCGCGAAGAGTTAGTGGCGATTAAAGAGCTATATAATGATGCGCGCCGTACCGAA   1557
            ************************************************************

Y11         ATCACCGCGAATACCTCTGACATCAATATTGAAGACCTGATTAATCAGGAAGATGTGGTT   1620
IP38477     ATCACCGCGAATACCTCTGACATCAATATTGAAGACCTGATTAATCAGGAAGATGTGGTT   1620
Ye.1        ATCACCGCGAATACCTCTGACATCAATATTGAAGACCTGATTAATCAGGAAGATGTGGTT   1620
Ye.2        ATCACCGCGAATACCTCTGACATCAATATTGAAGACCTGATTAATCAGGAAGATGTGGTT   1620
Ye.3        ATCACCGCGAATACCTCTGACATCAATATTGAAGACCTGATTAATCAGGAAGATGTGGTT   1617
Ye.4        ATCACCGCGAATACCTCTGACATCAATATTGAAGACCTGATTAATCAGGAAGATGTGGTT   1617
            ************************************************************

Y11         GTGACATTGTCTCATCAGGGCTATGTCAAATACCAACCTCTGTCTGATTACGAAGCTCAG   1680
IP38477     GTGACATTGTCTCATCAGGGCTATGTCAAATACCAACCTCTGTCTGATTACGAAGCTCAG   1680
Ye.1        GTGACATTGTCTCATCAGGGCTATGTCAAATACCAACCTCTGTCTGATTACGAAGCTCAG   1680
Ye.2        GTGACATTGTCTCATCAGGGCTATGTCAAATACCAACCTCTGTCTGATTACGAAGCTCAG   1680
Ye.3        GTGATATTGTCTCATCAGGGCTATGTCAAATACCAACCTCTGTCTGATTACGAAGCTCAG   1677
Ye.4        GTGATATTGTCTCATCAGGGCTATGTCAAATACCAACCTCTGTCTGATTACGAAGCTCAG   1677
            **** *******************************************************

Y11         CGTCGTGGTGGTAAAGGTAAATCAGCTGCGCGTATTAAAGAAGAAGACTTCATTGATCGC   1740
IP38477     CGTCGTGGTGGTAAAGGTAAATCAGCTGCGCGTATTAAAGAAGAAGACTTCATTGATCGC   1740
Ye.1        CGTCGTGGTGGTAAAGGTAAATCAGCTGCGCGTATTAAAGAAGAAGACTTCATTGATCGC   1740
Ye.2        CGTCGTGGTGGTAAAGGTAAATCAGCTGCGCGTATTAAAGAAGAAGACTTCATTGATCGC   1740
Ye.3        CGTCGTGGTGGTAAAGGTAAATCAGCTGCGCGTATTAAAGAAGAAGACTTCATTGATCGC   1737
Ye.4        CGTCGTGGTGGTAAAGGTAAATCAGCTGCGCGTATTAAAGAAGAAGACTTCATTGATCGC   1737
            ************************************************************

Y11         CTGCTGGTCGCCAACACCCACGATACTATTTTGTGCTTCTCCAGCCGTGGCCGTCTCTAT   1800
IP38477     CTGCTGGTCGCCAACACCCACGATACTATTTTGTGCTTCTCCAGCCGTGGCCGTCTCTAT   1800
Ye.1        CTGCTGGTCGCCAACACCCACGATACTATTTTGTGCTTCTCCAGCCGTGGCCGTCTCTAT   1800
Ye.2        CTGCTGGTCGCCAACACCCACGATACTATTTTGTGCTTCTCCAGCCGTGGCCGTCTCTAT   1800
Ye.3        CTGCTGGTCGCCAACACCCACGATACTATTTTGTGCTTCTCCAGCCGTGGCCGTCTCTAT   1797
Ye.4        CTGCTGGTCGCCAACACCCACGATACTATTTTGTGCTTCTCCAGCCGTGGCCGTCTCTAT   1797
            ************************************************************

Y11         TGGATGAAGGTCTATCAGTTGCCGGAAGCCAGTCGTGGCGCACGTGGTCGTCCGATCGTC   1860
IP38477     TGGATGAAGGTCTATCAGTTGCCGGAAGCCAGTCGTGGCGCACGTGGTCGTCCGATCGTC   1860
Ye.1        TGGATGAAGGTCTATCAGTTGCCGGAAGCCAGTCGTGGCGCACGTGGTCGTCCGATCGTC   1860
```

```
Ye.2        TGGATGAAGGTCTATCAGTTGCCGGAAGCCAGTCGTGGCGCACGTGGTCGTCCGATCGTC   1860
Ye.3        TGGATGAAGGTCTATCAGTTGCCGGAAGCCAGTCGTGGCGCACGTGGTCGTCCGATCGTC   1857
Ye.4        TGGATGAAGGTCTATCAGTTGCCGGAAGCCAGTCGTGGCGCACGTGGTCGTCCGATCGTC   1857
            ************************************************************

Y11         AACTTGTTGCCGCTGGAGCCAAATGAGCGTATCACCGCCATTCTGCCGGTGCGCGAATAC   1920
IP38477     AACTTGTTGCCGCTGGAGCCAAATGAGCGTATCACCGCCATTCTGCCGGTGCGCGAATAC   1920
Ye.1        AACTTGTTGCCGCTGGAGCCAAATGAGCGTATCACCGCCATTCTGCCGGTGCGCGAATAC   1920
Ye.2        AACTTGTTGCCGCTGGAGCCAAATGAGCGTATCACCGCCATTCTGCCGGTGCGCGAATAC   1920
Ye.3        AACTTGTTGCCGCTGGAGCCAAATGAGCGTATCACCGCCATTCTGCCGGTGCGCGAATAC   1917
Ye.4        AACTTGTTGCCGCTGGAGCCAAATGAGCGTATCACCGCCATTCTGCCGGTGCGCGAATAC   1917
            ************************************************************

Y11         GAAGAAGGTCGTCACATCTTTATGGCTACCGCCAGCGGTACCGTGAAGAAAACCGCACTG   1980
IP38477     GAAGAAGGTCGTCACATCTTTATGGCTACCGCCAGCGGTACCGTGAAGAAAACCGCACTG   1980
Ye.1        GAAGAAGGTCGTCACATCTTTATGGCTACCGCCAGCGGTACCGTGAAGAAAACCGCACTG   1980
Ye.2        GAAGAAGGTCGTCACATCTTTATGGCTACCGCCAGCGGTACCGTGAAGAAAACCGCACTG   1980
Ye.3        GAAGAAGGTCGTCACATCTTTATGGCTACCGCCAGCGGTACCGTGAAGAAAACCGCACTG   1977
Ye.4        GAAGAAGGTCGTCACATCTTTATGGCTACCGCCAGCGGTACCGTGAAGAAAACCGCACTG   1977
            ************************************************************

Y11         ACCGAGTTTAGCCGTCCACGCAGTGCCGGTATTATTGCCGTCAATCTGAATGAAGGCGAT   2040
IP38477     ACCGAGTTTAGCCGTCCACGCAGTGCCGGTATTATTGCCGTCAATCTGAATGAAGGCGAT   2040
Ye.1        ACCGAGTTTAGCCGTCCACGCAGTGCCGGTATTATTGCCGTCAATCTGAATGAAGGCGAT   2040
Ye.2        ACCGAGTTTAGCCGTCCACGCAGTGCCGGTATTATTGCCGTCAATCTGAATGAAGGCGAT   2040
Ye.3        ACCGAGTTTAGCCGTCCACGCAGTGCCGGTATTATTGCCGTCAATCTGAATGAAGGCGAT   2037
Ye.4        ACCGAGTTTAGCCGTCCACGCAGTGCCGGTATTATTGCCGTCAATCTGAATGAAGGCGAT   2037
            ************************************************************

Y11         GAACTGATTGGTGTCGATCTGACCGATGGCACTAACGAAGTCATGCTGTTCTCTGCATTG   2100
IP38477     GAACTGATTGGTGTCGATCTGACCGATGGCACTAACGAAGTCATGCTGTTCTCTGCATTG   2100
Ye.1        GAACTGATTGGTGTCGATCTGACCGATGGCACTAACGAAGTCATGCTGTTCTCTGCATTG   2100
Ye.2        GAACTGATTGGTGTCGATCTGACCGATGGCACTAACGAAGTCATGCTGTTCTCTGCATTG   2100
Ye.3        GAACTGATTGGTGTCGATCTGACCGATGGCACTAACGAAGTCATGCTGTTCTCTGCATTG   2097
Ye.4        GAACTGATTGGTGTCGATCTGACCGATGGCACTAACGAAGTCATGCTGTTCTCTGCATTG   2097
            ************************************************************

Y11         GGTAAAGTGGTTCGCTTCCCTGAATCGCAGGTCCGTTCGATGGGCCGTACCGCGACCGGT   2160
IP38477     GGTAAAGTGGTTCGCTTCCCTGAATCGCAGGTCCGTTCGATGGGCCGTACCGCGACCGGT   2160
Ye.1        GGTAAAGTGGTTCGCTTCCCTGAATCGCAGGTCCGTTCGATGGGCCGTACCGCGACCGGT   2160
Ye.2        GGTAAAGTGGTTCGCTTCCCTGAATCGCAGGTCCGTTCGATGGGCCGTACCGCGACCGGT   2160
Ye.3        GGTAAAGTGGTTCGCTTCCCTGAATCGCAGGTCCGTTCGATGGGCCGTACCGCGACCGGT   2157
Ye.4        GGTAAAGTGGTTCGCTTCCCTGAATCGCAGGTCCGTTCGATGGGCCGTACCGCGACCGGT   2157
            ************************************************************

Y11         GTACGCGGTATCAACCTCAATGGCGACGATCGGGTTATTTCTCTTATCATCCCTCGTGGC   2220
IP38477     GTACGCGGTATCAACCTCAATGGCGACGATCGGGTTATTTCTCTTATCATCCCTCGTGGC   2220
Ye.1        GTACGCGGTATCAACCTCAATGGCGACGATCGGGTTATTTCTCTTATCATCCCTCGTGGC   2220
Ye.2        GTACGCGGTATCAACCTCAATGGCGACGATCGGGTTATTTCTCTTATCATCCCTCGTGGC   2220
Ye.3        GTACGCGGTATCAACCTCAATGGCGACGATCGGGTTATTTCTCTTATCATCCCTCGTGGC   2217
Ye.4        GTACGCGGTATCAACCTCAATGGCGACGATCGGGTTATTTCTCTTATCATCCCTCGTGGC   2217
            ************************************************************

Y11         GATGGTGAAATCCTGACTGTGACTGAAAACGGTTACGGTAAACGTACCGCAGTGGAAGAA   2280
IP38477     GATGGTGAAATCCTGACTGTGACTGAAAACGGTTACGGTAAACGTACCGCAGTGGAAGAA   2280
Ye.1        GATGGTGAAATCCTGACTGTGACTGAAAACGGTTACGGTAAACGTACCGCAGTGGAAGAA   2280
Ye.2        GATGGTGAAATCCTGACTGTGACTGAAAACGGTTACGGTAAACGTACCGCAGTGGAAGAA   2280
Ye.3        GATGGTGAAATCCTGACTGTGACTGAAAACGGTTACGGTAAACGTACCGCAGTGGAAGAA   2277
Ye.4        GATGGTGAAATCCTGACTGTGACTGAAAACGGTTACGGTAAACGTACCGCAGTGGAAGAA   2277
            ************************************************************

Y11         TATCCAACCAAGTCCCGTGCGACTCAGGGGGTTATCTCCATTAAAGTCAGTGAGCGTAAT   2340
IP38477     TATCCAACCAAGTCCCGTGCGACTCAGGGGGTTATCTCCATTAAAGTCAGTGAGCGTAAT   2340
Ye.1        TATCCAACCAAGTCCCGTGCGACTCAGGGGGTTATCTCCATTAAAGTCAGTGAGCGTAAT   2340
```

```
Ye.2        TATCCAACCAAGTCCCGTGCGACTCAGGGGGTTATCTCCATTAAAGTCAGTGAGCGTAAT   2340
Ye.3        TATCCAACCAAGTCCCGTGCGACTCAGGGGGTTATCTCCATTAAAGTCAGTGAGCGTAAT   2337
Ye.4        TATCCAACCAAGTCCCGTGCGACTCAGGGGGTTATCTCCATTAAAGTCAGTGAGCGTAAT   2337
            ************************************************************

Y11         GGTAAGGTTGTTGGGGCGGTACAAGTCGCGCCGACTGACCAAATCATGATGATCACCGAT   2400
IP38477     GGTAAGGTTGTTGGGGCGGTACAAGTCGCGCCGACTGACCAAATCATGATGATCACCGAT   2400
Ye.1        GGTAAGGTTGTTGGGGCGGTACAAGTCGCGCCGACTGACCAAATCATGATGATCACCGAT   2400
Ye.2        GGTAAGGTTGTTGGGGCGGTACAAGTCGCGCCGACTGACCAAATCATGATGATCACCGAT   2400
Ye.3        GGTAAGGTTGTTGGGGCGGTACAAGTCGCGCCGACTGACCAAATCATGATGATCACCGAT   2397
Ye.4        GGTAAGGTTGTTGGGGCGGTACAAGTCGCGCCGACTGACCAAATCATGATGATCACCGAT   2397
            ************************************************************

Y11         GCCGGTACACTGGTACGTACCCGCGTATCAGAGGTGAGTGTTGTGGGGCGTAATACCCAG   2460
IP38477     GCCGGTACACTGGTACGTACCCGCGTATCAGAGGTGAGTGTTGTGGGGCGTAATACCCAG   2460
Ye.1        GCCGGTACACTGGTACGTACCCGCGTATCAGAGGTGAGTGTTGTGGGGCGTAATACCCAG   2460
Ye.2        GCCGGTACACTGGTACGTACCCGCGTATCAGAGGTGAGTGTTGTGGGGCGTAATACCCAG   2460
Ye.3        GCCGGTACACTGGTACATACCCGCGTATCAGAGGTGAGTGTTGTGGGGCGTAATACCCAG   2457
Ye.4        GCCGGTACACTGGTACATACCCGCGTATCAGAGGTGAGTGTTGTGGGGCGTAATACCCAG   2457
            **************** *******************************************

Y11         GGTGTGACACTTATCCGTACCACTGAAGACGAGCATGTCGTTGGCCTGCAACGTGTAGCG   2520
IP38477     GGTGTGACACTTATCCGTACCACTGAAGACGAGCATGTCGTTGGCCTGCAACGTGTAGCG   2520
Ye.1        GGTGTGACACTTATCCGTACCACTGAAGACGAGCATGTCGTTGGCCTGCAACGTGTAGCG   2520
Ye.2        GGTGTGACACTTATCCGTACCACTGAAGACGAGCATGTCGTTGGCCTGCAACGTGTAGCG   2520
Ye.3        GGTGTGACACTTATCCGTACCACTGAAGACGAGCATGTCGTTGGCCTGCAACGTGTAGCG   2517
Ye.4        GGTGTGACACTTATCCGTACCACTGAAGACGAGCATGTCGTTGGCCTGCAACGTGTAGCG   2517
            ************************************************************

Y11         GAGCCAGAAGAGGATGATAACATCCTTGAGGGTGAATCATTAGAAGGTGAAGAGGGCAGC   2580
IP38477     GAGCCAGAAGAGGATGATAACATCCTTGAGGGTGAATCATTAGAAGGTGAAGAGGGCAGC   2580
Ye.1        GAGCCAGAAGAGGATGATAACATCCTTGAGGGTGAATCATTAGAAGGTGAAGAGGGCAGC   2580
Ye.2        GAGCCAGAAGAGGATGATAACATCCTTGAGGGTGAATCATTAGAAGGTGAAGAGGGCAGC   2580
Ye.3        GAGCCAGAAGAGGATGATAACATCCTTGAGGGTGAATCATTAGAAGGTGAAGAGGGCAGC   2577
Ye.4        GAGCCAGAAGAGGATGATAACATCCTTGAGGGTGAATCATTAGAAGGTGAAGAGGGCAGC   2577
            ************************************************************

Y11         GAGGAAAATACTGCACTGAATGCACCAGAAGATGAAGATGCTGCTGACGAAGCAGAAGAC   2640
IP38477     GAGGAAAATACTGCACTGAATGCACCAGAAGATGAAGATGCTGCTGACGAAGCAGAAGAC   2640
Ye.1        GAGGAAAATACTGCACTGAATGCACCAGAAGATGAAGATGCTGCTGACGAAGCAGAAGAC   2640
Ye.2        GAGGAAAATACTGCACTGAATGCACCAGAAGATGAAGATGCTGCTGACGAAGCAGAAGAC   2640
Ye.3        GAGGAAAATACTGCACTGAATGCACCAGAAGATGAAGATGCTGCTGACGAAGCAGAAGAC   2637
Ye.4        GAGGAAAATACTGCACTGAATGCACCAGAAGATGAAGATGCTGCTGACGAAGCAGAAGAC   2637
            ************************************************************

Y11         GATGACAATAATGTGTAA   2658
IP38477     GATGACAATAATGTGTAA   2658
Ye.1        GATGACAATAATGTGTAA   2658
Ye.2        GATGACAATAATGTGTAA   2658
Ye.3        GATGACAATAATGTGTAA   2655
Ye.4        GATGACAATAATGTGTAA   2655
            ******************
```

**B.**

```
Y11         MSDLAREITPVNIEEELKSSYLDYAMSVIVGRALPDVRDGLKPVHRRVLYAMNVLGNDWN   60
IP38477     MSDLAREITPVNIEEELKSSYLDYAMSVIVGRALPDVRDGLKPVHRRVLYAMNVLGNDWN   60
Ye.1        MSDLAREITPVNIEEELKSSYLDYAMSVIVGRALPDVRDGLKPVHRRVLYAMNVLGNDWN   60
Ye.2        MSDLAREITPVNIEEELKSSYLDYAMSVIVGRALPDVRDGLKPVHRRVLYAMNVLGNDWN   60
Ye.3        MSDLAREITPVNIEEELKSSYLDYAMSVIVGRALPDVRDGLKPVHRRVLYAMNVLGNDWN   60
Ye.4        MSDLAREITPVNIEEELKSSYLDYAMSVIVGRALPDVRDGLKPVHRRVLYAMNVLGNDWN   60
            ************************************************************

Y11         KPYKKSARVVGDVIGKYHPHGDSAVYDTIVRMAQPFSLRYMLVDGQGNFGSVDGDSAAAM   120
IP38477     KPYKKSARVVGDVIGKYHPHGDSAVYDTIVRMAQPFSLRYMLVDGQGNFGSVDGDSAAAM   120
Ye.1        KPYKKSARVVGDVIGKYHPHGDSAVYDTIVRMAQPFSLRYMLVDGQGNFGSVDGDSAAAM   120
Ye.2        KPYKKSARVVGDVIGKYHPHGDSAVYDTIVRMAQPFSLRYMLVDGQGNFGSVDGDSAAAM   120
Ye.3        KPYKKSARVVGDVIGKYHPHGG-AVYDTIVRMAQPFSLRYMLVDGQGNFGSVDGDSAAAM   119
Ye.4        KPYKKSARVVGDVIGKYHPHGG-AVYNTIVRMAQPFSLRYMLVDGQGNFGSVDGDSAAAM   119
            *********************. ***:*********************************

Y11         RYTEIRMSKIAHELLADLEKDTVDFVPNYDGTEQIPAVMPTKIPNLLVNGSSGIAVGMAT   180
IP38477     RYTEIRMSKIAHELLADLEKDTVDFVPNYDGTEQIPAVMPTKIPNLLVNGSSGIAVGMAT   180
Ye.1        RYTEIRMSKIAHELLADLEKDTVDFVPNYDGTEQIPAVMPTKIPNLLVNGSSGIAVGMAT   180
Ye.2        RYTEIRMSKIAHELLADLEKDTVDFVPNYDGTEQIPAVMPTKIPNLLVNGSSGIAVGMAT   180
Ye.3        RYTEIRMSKIAHELLADLEKDTVDFVPNYDGTEQIPAVMPTKIPNLLVNGSSGIAVGMAT   179
Ye.4        RYTEIRMSKIAHELLADLEKDTVDFVPNYDGTEQIPAVMPTKIPNLLVNGSSGIAVGMAT   179
            ************************************************************

Y11         NIPPHNLSEVIDGCLAYIEDENITIEGLMEYIPGPDFPTAAIINGRRGIEEAYRTGRGKV   240
IP38477     NIPPHNLSEVIDGCLAYIEDENITIEGLMEYIPGPDFPTAAIINGRRGIEEAYRTGRGKV   240
Ye.1        NIPPHNLSEVIDGCLAYIEDENITIEGLMEYIPGPDFPTAAIINGRRGIEEAYRTGRGKV   240
Ye.2        NIPPHNLSEVIDGCLAYIEDENITIEGLMEYIPGPDFPTAAIINGRRGIEEAYRTGRGKV   240
Ye.3        NIPPHNLSEVIDGCLAYIEDENITIEGLMEYIPGPDFPTAAIINGRRGIEEAYRTGRGKV   239
Ye.4        NIPPHNLSEVIDGCLAYIEDENITIEGLMEYIPGPDFPTAAIINGRRGIEEAYRTGRGKV   239
            ************************************************************

Y11         YIRARAEVEADAKTGRETIIVHEIPYQVNKARLIEKIAELVKEKRVEGISALRDESDKDG   300
IP38477     YIRARAEVEADAKTGRETIIVHEIPYQVNKARLIEKIAELVKEKRVEGISALRDESDKDG   300
Ye.1        YIRARAEVEADAKTGRETIIVHEIPYQVNKARLIEKIAELVKEKRVEGISALRDESDKDG   300
Ye.2        YIRARAEVEADAKTGRETIIVHEIPYQVNKARLIEKIAELVKEKRVEGISALRDESDKDG   300
Ye.3        YIRARAEVEADAKTGRETIIVHEIPYQVNKARLIEKIAELVKEKRVEGISALRDESDKDG   299
Ye.4        YIRARAEVEADAKTGRETIIVHEIPYQVNKARLIEKIAELVKEKRVEGISALRDESDKDG   299
            ************************************************************

Y11         MRIVIEIKRDAVGEVVLNNLYSLTQLQVTFGINMVGLSQGQPKLLNLKDILVAFVRHRRE   360
IP38477     MRIVIEIKRDAVGEVVLNNLYSLTQLQVTFGINMVGLSQGQPKLLNLKDILVAFVRHRRE   360
Ye.1        MRIVIEIKRDAVGEVVLNNLYSLTQLQVTFGINMVGLSQGQPKLLNLKDILVAFVRHRRE   360
Ye.2        MRIVIEIKRDAVGEVVLNNLYSLTQLQVTFGINMVGLSQGQPKLLNLKDILVAFVRHRRE   360
Ye.3        MRIVIEIKRDAVGEVVLNNLYSLTQLQVTFGINMVGLSQGQPKLLNLKDILVAFVRHRRE   359
Ye.4        MRIVIEIKRDAVGEVVLNNLYSLTQLQVTFGINMVGLSQGQPKLLNLKDILVAFVRHRRE   359
            ************************************************************

Y11         VVTRRTIFELRKARDRAHILEALAIALANIDPIIELIRRAATPAEAKAGLIASPWELGNV   420
IP38477     VVTRRTIFELRKARDRAHILEALAIALANIDPIIELIRRAATPAEAKAGLIASPWELGNV   420
Ye.1        VVTRRTIFELRKARDRAHILEALAIALANIDPIIELIRRAATPAEAKAGLIASPWELGNV   420
Ye.2        VVTRRTIFELRKARDRAHILEALAIALANIDPIIELIRRAATPAEAKAGLIASPWELGNV   420
Ye.3        VVTRRTIFELRKARDRAHILEALAIALANIDPIIELIRRAATPAEAKAGLIASPWELGNV   419
Ye.4        VVTRRTIFELRKARDRAHILEALAIALANIDPIIELIRRAATPAEAKAGLIASPWELGNV   419
            ************************************************************

Y11         AAMLERAGGDAARPEWLEAEFGIRDGKYYLTEQQAQAILDLRLQKLTGLEHEKLLDEYKE   480
IP38477     AAMLERAGGDAARPEWLEAEFGIRDGKYYLTEQQAQAILDLRLQKLTGLEHEKLLDEYKE   480
```

```
Ye.1       AAMLERAGGDAARPEWLEAEFGIRDGKYYLTEQQAQAILDLRLQKLTGLEHEKLLDEYKE   480
Ye.2       AAMLERAGGDAARPEWLEAEFGIRDGKYYLTEQQAQAILDLRLQKLTGLEHEKLLDEYKE   480
Ye.3       AAMLERAGGDAARPEWLEAEFGIRDGKYYLTEQQAQAILDLRLQKLTGLEHEKLLDEYKE   479
Ye.4       AAMLERAGGDAARPEWLEAEFGIRDGKYYLTEQQAQAILDLRLQKLTGLEHEKLLDEYKE   479
           ************************************************************

Y11        LLTVIAELIFILENPERLMEVIREELVAIKELYNDARRTEITANTSDINIEDLINQEDVV   540
IP38477    LLTVIAELIFILENPERLMEVIREELVAIKELYNDARRTEITANTSDINIEDLINQEDVV   540
Ye.1       LLTVIAELIFILENPERLMEVIREELVAIKELYNDARRTEITANTSDINIEDLINQEDVV   540
Ye.2       LLTVIAELIFILENPERLMEVIREELVAIKELYNDARRTEITANTSDINIEDLINQEDVV   540
Ye.3       LLTVIAELIFILENPERLMEVIREELVAIKELYNDARRTEITANTSDINIEDLINQEDVV   539
Ye.4       LLTVIAELIFILENPERLMEVIREELVAIKELYNDARRTEITANTSDINIEDLINQEDVV   539
           ************************************************************

Y11        VTLSHQGYVKYQPLSDYEAQRRGGKGKSAARIKEEDFIDRLLVANTHDTILCFSSRGRLY   600
IP38477    VTLSHQGYVKYQPLSDYEAQRRGGKGKSAARIKEEDFIDRLLVANTHDTILCFSSRGRLY   600
Ye.1       VTLSHQGYVKYQPLSDYEAQRRGGKGKSAARIKEEDFIDRLLVANTHDTILCFSSRGRLY   600
Ye.2       VTLSHQGYVKYQPLSDYEAQRRGGKGKSAARIKEEDFIDRLLVANTHDTILCFSSRGRLY   600
Ye.3       VILSHQGYVKYQPLSDYEAQRRGGKGKSAARIKEEDFIDRLLVANTHDTILCFSSRGRLY   599
Ye.4       VILSHQGYVKYQPLSDYEAQRRGGKGKSAARIKEEDFIDRLLVANTHDTILCFSSRGRLY   599
           * **********************************************************

Y11        WMKVYQLPEASRGARGRPIVNLLPLEPNERITAILPVREYEEGRHIFMATASGTVKKTAL   660
IP38477    WMKVYQLPEASRGARGRPIVNLLPLEPNERITAILPVREYEEGRHIFMATASGTVKKTAL   660
Ye.1       WMKVYQLPEASRGARGRPIVNLLPLEPNERITAILPVREYEEGRHIFMATASGTVKKTAL   660
Ye.2       WMKVYQLPEASRGARGRPIVNLLPLEPNERITAILPVREYEEGRHIFMATASGTVKKTAL   660
Ye.3       WMKVYQLPEASRGARGRPIVNLLPLEPNERITAILPVREYEEGRHIFMATASGTVKKTAL   659
Ye.4       WMKVYQLPEASRGARGRPIVNLLPLEPNERITAILPVREYEEGRHIFMATASGTVKKTAL   659
           ************************************************************

Y11        TEFSRPRSAGIIAVNLNEGDELIGVDLTDGTNEVMLFSALGKVVRFPESQVRSMGRTATG   720
IP38477    TEFSRPRSAGIIAVNLNEGDELIGVDLTDGTNEVMLFSALGKVVRFPESQVRSMGRTATG   720
Ye.1       TEFSRPRSAGIIAVNLNEGDELIGVDLTDGTNEVMLFSALGKVVRFPESQVRSMGRTATG   720
Ye.2       TEFSRPRSAGIIAVNLNEGDELIGVDLTDGTNEVMLFSALGKVVRFPESQVRSMGRTATG   720
Ye.3       TEFSRPRSAGIIAVNLNEGDELIGVDLTDGTNEVMLFSALGKVVRFPESQVRSMGRTATG   719
Ye.4       TEFSRPRSAGIIAVNLNEGDELIGVDLTDGTNEVMLFSALGKVVRFPESQVRSMGRTATG   719
           ************************************************************

Y11        VRGINLNGDDRVISLIIPRGDGEILTVTENGYGKRTAVEEYPTKSRATQGVISIKVSERN   780
IP38477    VRGINLNGDDRVISLIIPRGDGEILTVTENGYGKRTAVEEYPTKSRATQGVISIKVSERN   780
Ye.1       VRGINLNGDDRVISLIIPRGDGEILTVTENGYGKRTAVEEYPTKSRATQGVISIKVSERN   780
Ye.2       VRGINLNGDDRVISLIIPRGDGEILTVTENGYGKRTAVEEYPTKSRATQGVISIKVSERN   780
Ye.3       VRGINLNGDDRVISLIIPRGDGEILTVTENGYGKRTAVEEYPTKSRATQGVISIKVSERN   779
Ye.4       VRGINLNGDDRVISLIIPRGDGEILTVTENGYGKRTAVEEYPTKSRATQGVISIKVSERN   779
           ************************************************************

Y11        GKVVGAVQVAPTDQIMMITDAGTLVRTRVSEVSVVGRNTQGVTLIRTTEDEHVVGLQRVA   840
IP38477    GKVVGAVQVAPTDQIMMITDAGTLVRTRVSEVSVVGRNTQGVTLIRTTEDEHVVGLQRVA   840
Ye.1       GKVVGAVQVAPTDQIMMITDAGTLVRTRVSEVSVVGRNTQGVTLIRTTEDEHVVGLQRVA   840
Ye.2       GKVVGAVQVAPTDQIMMITDAGTLVRTRVSEVSVVGRNTQGVTLIRTTEDEHVVGLQRVA   840
Ye.3       GKVVGAVQVAPTDQIMMITDAGTLVHTRVSEVSVVGRNTQGVTLIRTTEDEHVVGLQRVA   839
Ye.4       GKVVGAVQVAPTDQIMMITDAGTLVHTRVSEVSVVGRNTQGVTLIRTTEDEHVVGLQRVA   839
           *************************:**********************************

Y11        EPEEDDNILEGESLEGEEGSEENTALNAPEDEDAADEAEDDDNNV* 885
IP38477    EPEEDDNILEGESLEGEEGSEENTALNAPEDEDAADEAEDDDNNV* 885
Ye.1       EPEEDDNILEGESLEGEEGSEENTALNAPEDEDAADEAEDDDNNV* 885
Ye.2       EPEEDDNILEGESLEGEEGSEENTALNAPEDEDAADEAEDDDNNV* 885
Ye.3       EPEEDDNILEGESLEGEEGSEENTALNAPEDEDAADEAEDDDNNV* 884
Ye.4       EPEEDDNILEGESLEGEEGSEENTALNAPEDEDAADEAEDDDNNV* 884
           *********************************************
```

**Figure S4.** Metabolic map of *Ye* Y11 strain displayed on iPath 3 (https://pathways.embl.de/)

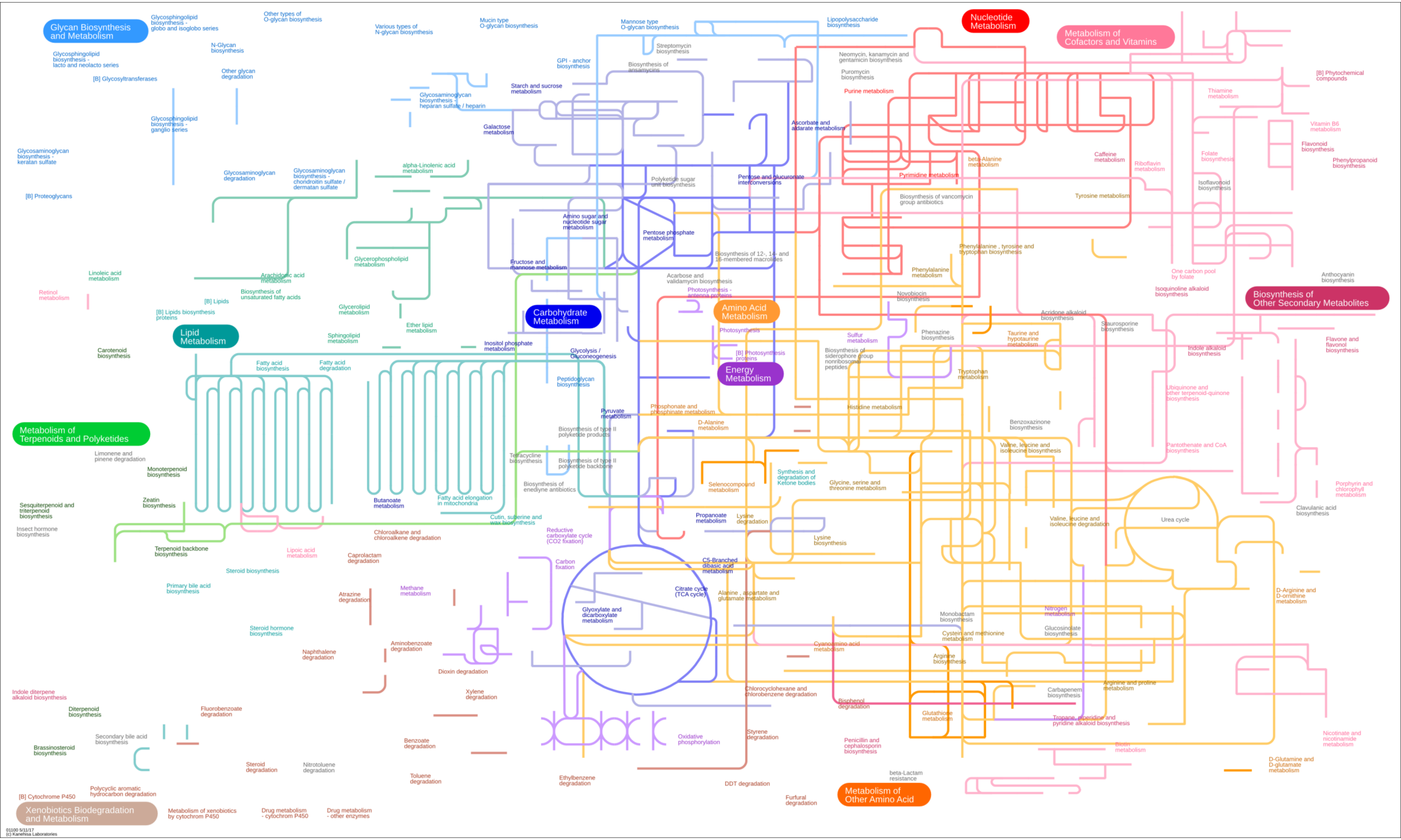

**Figure S5.** Protein abundance foldchanges mapped on the Y11 metabolic map visualized with iPath3 for (A) Ye.1 vs Ye.2, (B) Ye.1 vs Ye.3, (C) Ye.1 vs Ye.4, (D) Ye.2 vs Ye.3, (E) Ye.2 vs Ye.4 and (F) Ye.3 vs Ye.4. Red lines show upregulated paths in the first term of the comparison, while blue lines stand for upregulated paths in the second term of the comparison. Black lines represent pathways with |log2(foldchange)|<0.5, and grey lines stand for pathways with no detected proteins in our data. Line width is proportional to log2(foldchange) and is capped at |log2(foldchange)|=3. All greater foldchange are represented with the same width for better clarity.

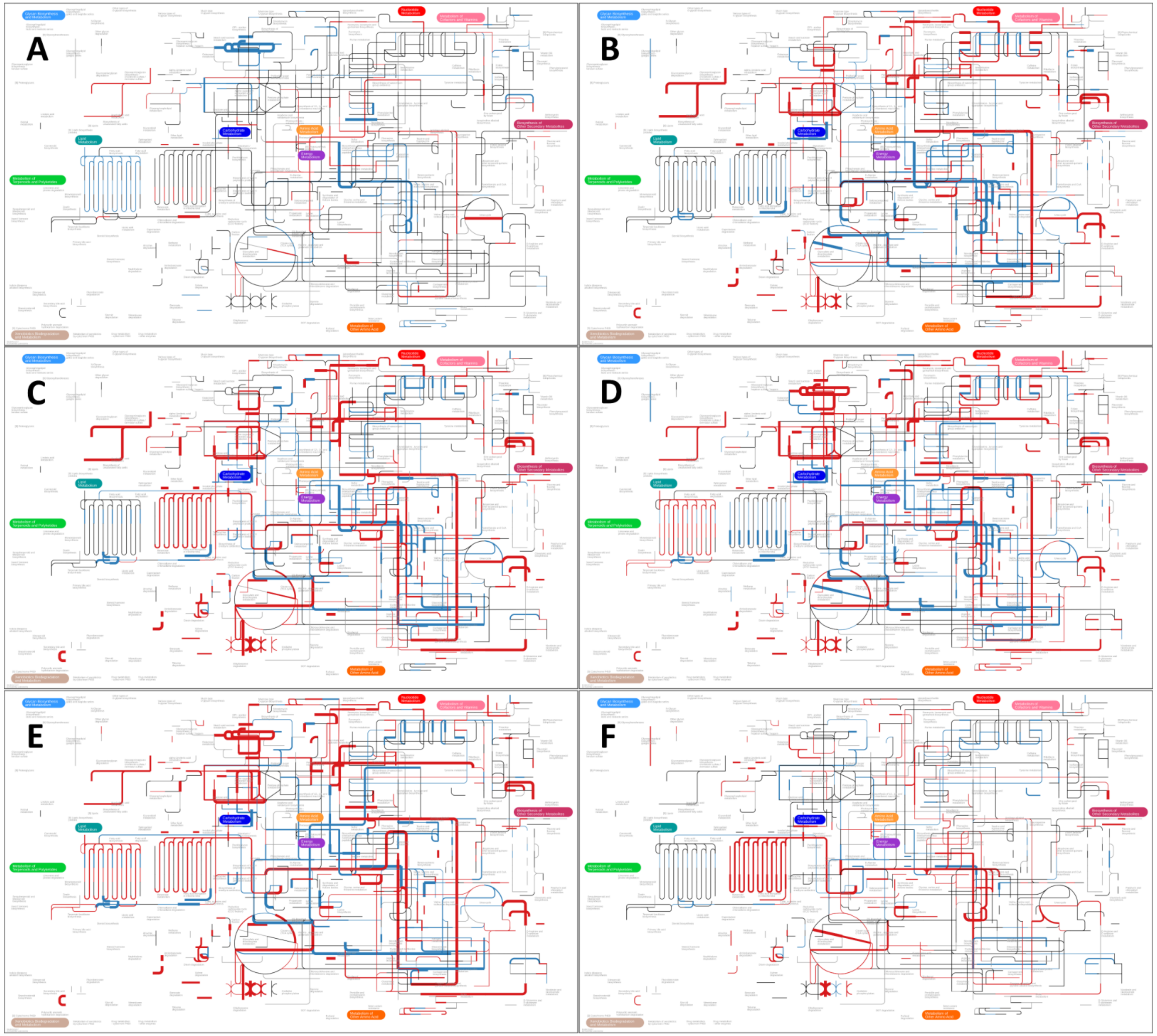

**Figure S6.** Protein abundance foldchanges mapped on the Y11 KEGG pathways.

- more abundant
- equally abundant
- less abundant
- not detected
- not existing in *Y. enterocolitica* Y11

#### Comparisons inside a gene

| | | | |
|---|---|---|---|
| Ye.1 vs Ye.2 | Ye.1 vs Ye.3 | Ye.1 vs Ye.4 | Ye.3 vs Ye.4 |

#### Foldchange scale

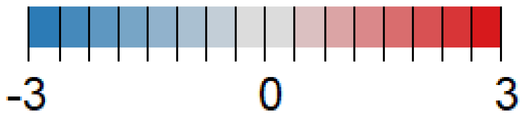

### Fig S6 table of content

| KEGG pathway link | page |
|---|---|
| Carbohydrate metabolism | |
| 00010 Glycolysis / Gluconeogenesis | 3 |
| 00020 Citrate cycle (TCA cycle) | 4 |
| 00030 Pentose phosphate pathway | 5 |
| 00040 Pentose and glucuronate interconversions | 6 |
| 00051 Fructose and mannose metabolism | 7 |
| 00052 Galactose metabolism | 8 |
| 00053 Ascorbate and aldarate metabolism | 9 |
| 00500 Starch and sucrose metabolism | 10 |
| 00520 Amino sugar and nucleotide sugar metabolism | 11 |
| 00620 Pyruvate metabolism | 12 |
| 00630 Glyoxylate and dicarboxylate metabolism | 13 |
| 00640 Propanoate metabolism | 14 |
| 00650 Butanoate metabolism | 15 |
| 00660 C5-Branched dibasic acid metabolism | 16 |
| 00562 Inositol phosphate metabolism | 17 |
| Energy metabolism | |
| 00190 Oxidative phosphorylation | 18 |
| 00680 Methane metabolism | 19 |
| 00910 Nitrogen metabolism | 20 |
| 00920 Sulfur metabolism | 21 |
| Lipid metabolism | |
| 00061 Fatty acid biosynthesis | 22 |
| 00071 Fatty acid degradation | 23 |
| 00561 Glycerolipid metabolism | 24 |
| 00564 Glycerophospholipid metabolism | 25 |
| 00565 Ether lipid metabolism | 26 |
| 00600 Sphingolipid metabolism | 27 |
| 00592 alpha-Linolenic acid metabolism | 28 |
| 01040 Biosynthesis of unsaturated fatty acids | 29 |
| Nucleotide metabolism | |
| 00230 Purine metabolism | 30 |
| 00240 Pyrimidine metabolism | 31 |
| Amino acid metabolism | |
| 00250 Alanine, aspartate and glutamate metabolism | 32 |
| 00260 Glycine, serine and threonine metabolism | 33 |
| 00270 Cysteine and methionine metabolism | 34 |
| 00280 Valine, leucine and isoleucine degradation | 35 |
| 00290 Valine, leucine and isoleucine biosynthesis | 36 |
| 00300 Lysine biosynthesis | 37 |
| 00310 Lysine degradation | 38 |
| 00220 Arginine biosynthesis | 39 |
| 00330 Arginine and proline metabolism | 40 |
| 00340 Histidine metabolism | 41 |
| 00350 Tyrosine metabolism | 42 |
| 00360 Phenylalanine metabolism | 43 |
| 00380 Tryptophan metabolism | 44 |
| 00400 Phenylalanine, tyrosine and tryptophan biosynthesis | 45 |
| Metabolism of other amino acids | |
| 00410 beta-Alanine metabolism | 46 |
| 00430 Taurine and hypotaurine metabolism | 47 |
| 00440 Phosphonate and phosphinate metabolism | 48 |
| 00450 Selenocompound metabolism | 49 |
| 00460 Cyanoamino acid metabolism | 50 |
| 00470 D-Amino acid metabolism | 51 |
| 00480 Glutathione metabolism | 52 |
| Glycan biosynthesis and metabolism | |
| 00541 O-Antigen nucleotide sugar biosynthesis | 53 |
| 00550 Peptidoglycan biosynthesis | 54 |

| KEGG pathway link | page |
|---|---|
| Metabolism of cofactors and vitamins | |
| 00730 Thiamine metabolism | 55 |
| 00740 Riboflavin metabolism | 56 |
| 00750 Vitamin B6 metabolism | 57 |
| 00760 Nicotinate and nicotinamide metabolism | 58 |
| 00770 Pantothenate and CoA biosynthesis | 59 |
| 00780 Biotin metabolism | 60 |
| 00785 Lipoic acid metabolism | 61 |
| 00790 Folate biosynthesis | 62 |
| 00670 One carbon pool by folate | 63 |
| 00860 Porphyrin metabolism | 64 |
| 00130 Ubiquinone and other terpenoid-quinone biosynthesis | 65 |
| Metabolism of terpenoids and polyketides | |
| 00900 Terpenoid backbone biosynthesis | 66 |
| 00907 Pinene, camphor and geraniol degradation | 67 |
| 00523 Polyketide sugar unit biosynthesis | 68 |
| 01053 Biosynthesis of siderophore group nonribosomal peptides | 69 |
| Genetic Information Processing | |
| Transcription | |
| 03020 RNA polymerase | 70 |
| Translation | |
| 03010 Ribosome | 71 |
| 00970 Aminoacyl-tRNA biosynthesis | 72 |
| Folding, sorting and degradation | |
| 03060 Protein export | 73 |
| 04122 Sulfur relay system | 74 |
| 03018 RNA degradation | 75 |
| Replication and repair | |
| 03030 DNA replication | 76 |
| 03410 Base excision repair | 77 |
| 03420 Nucleotide excision repair | 78 |
| 03430 Mismatch repair | 79 |
| 03440 Homologous recombination | 80 |
| Environmental Information Processing | |
| Membrane transport | |
| 02010 ABC transporters | 81 |
| 02060 Phosphotransferase system (PTS) | 82 |
| 03070 Bacterial secretion system | 83 |
| Signal transduction | |
| 02020 Two-component system | 84-85 |
| Cellular Processes | |
| Cellular community - prokaryotes | |
| 02024 Quorum sensing | 86 |
| Cell motility | |
| 02030 Bacterial chemotaxis | 87 |
| 02040 Flagellar assembly | 88 |
| Human Diseases | |
| Drug resistance: antimicrobial | |
| 01501 beta-Lactam resistance | 89 |
| 01502 Vancomycin resistance | 90 |
| 01503 Cationic antimicrobial peptide (CAMP) resistance | 91 |

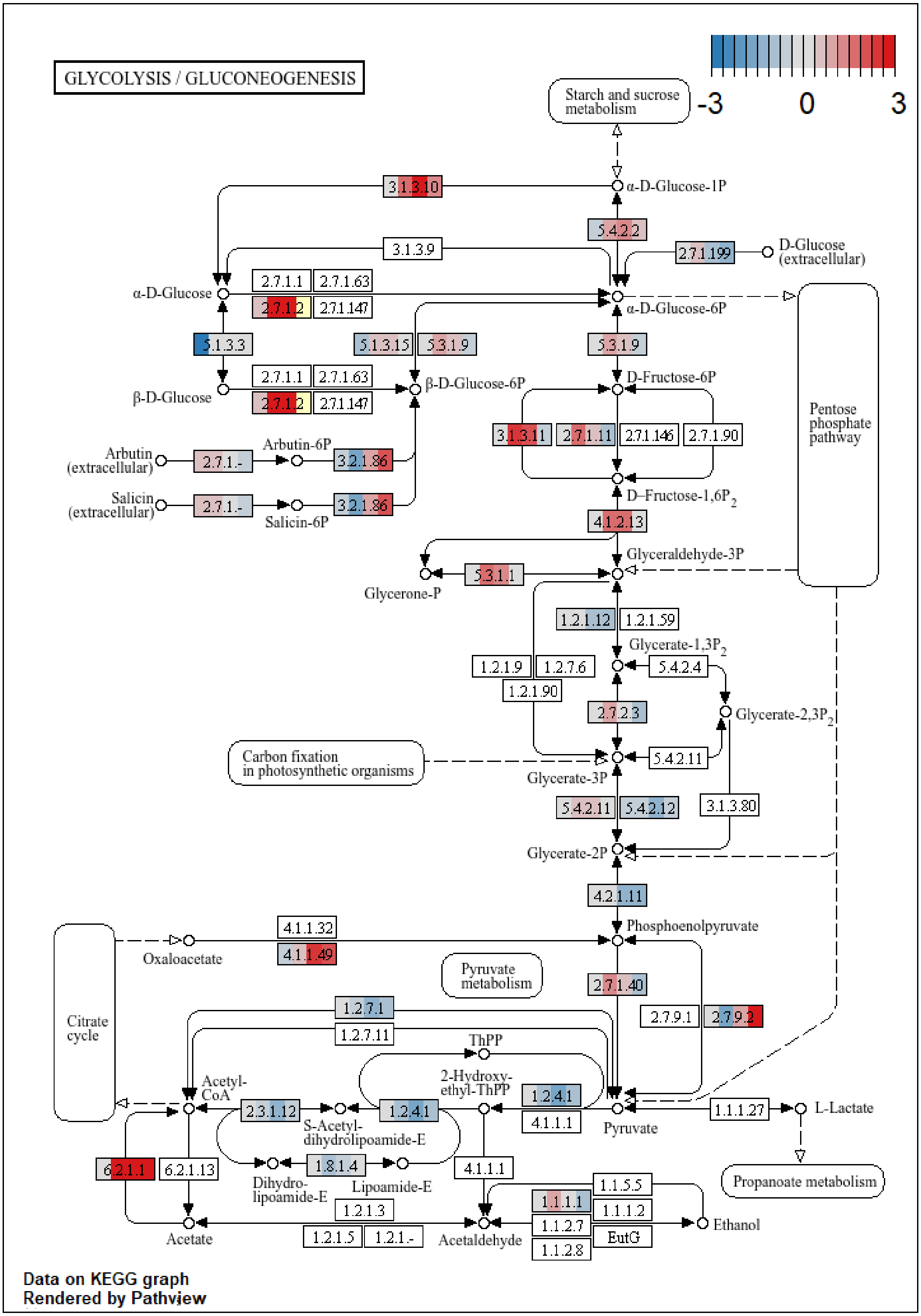
GLYCOLYSIS / GLUCONEOGENESIS
-3
0
3
Starch and sucrose metabolism
α-D-Glucose-1P
3.1.3.10
5.4.2.2
3.1.3.9
2.7.1.199
D-Glucose (extracellular)
α-D-Glucose
2.7.1.1
2.7.1.63
2.7.1.2
2.7.1.147
α-D-Glucose-6P
5.1.3.3
5.1.3.15
5.3.1.9
β-D-Glucose
β-D-Glucose-6P
D-Fructose-6P
Pentose phosphate pathway
3.1.3.11
2.7.1.11
2.7.1.146
2.7.1.90
Arbutin (extracellular)
2.7.1.-
Arbutin-6P
3.2.1.86
Salicin (extracellular)
Salicin-6P
D-Fructose-1,6P2
4.1.2.13
Glyceraldehyde-3P
Glycerone-P
5.3.1.1
1.2.1.12
1.2.1.59
Glycerate-1,3P2
1.2.1.9
1.2.7.6
5.4.2.4
1.2.1.90
2.7.2.3
Glycerate-2,3P2
Carbon fixation in photosynthetic organisms
5.4.2.11
Glycerate-3P
5.4.2.12
3.1.3.80
Glycerate-2P
4.2.1.11
Phosphoenolpyruvate
4.1.1.32
4.1.1.49
Citrate cycle
Oxaloacetate
Pyruvate metabolism
2.7.1.40
1.2.7.1
1.2.7.11
2.7.9.1
2.7.9.2
ThPP
2-Hydroxy-ethyl-ThPP
1.2.4.1
Acetyl-CoA
2.3.1.12
S-Acetyl-dihydrolipoamide-E
4.1.1.1
Pyruvate
1.1.1.27
L-Lactate
6.2.1.1
6.2.1.13
1.8.1.4
Dihydro-lipoamide-E
Lipoamide-E
Propanoate metabolism
1.1.5.5
1.1.1.1
1.1.1.2
1.2.1.3
1.1.2.7
Ethanol
Acetate
1.2.1.5
1.2.1.-
Acetaldehyde
EutG
1.1.2.8
Data on KEGG graph
Rendered by Pathview

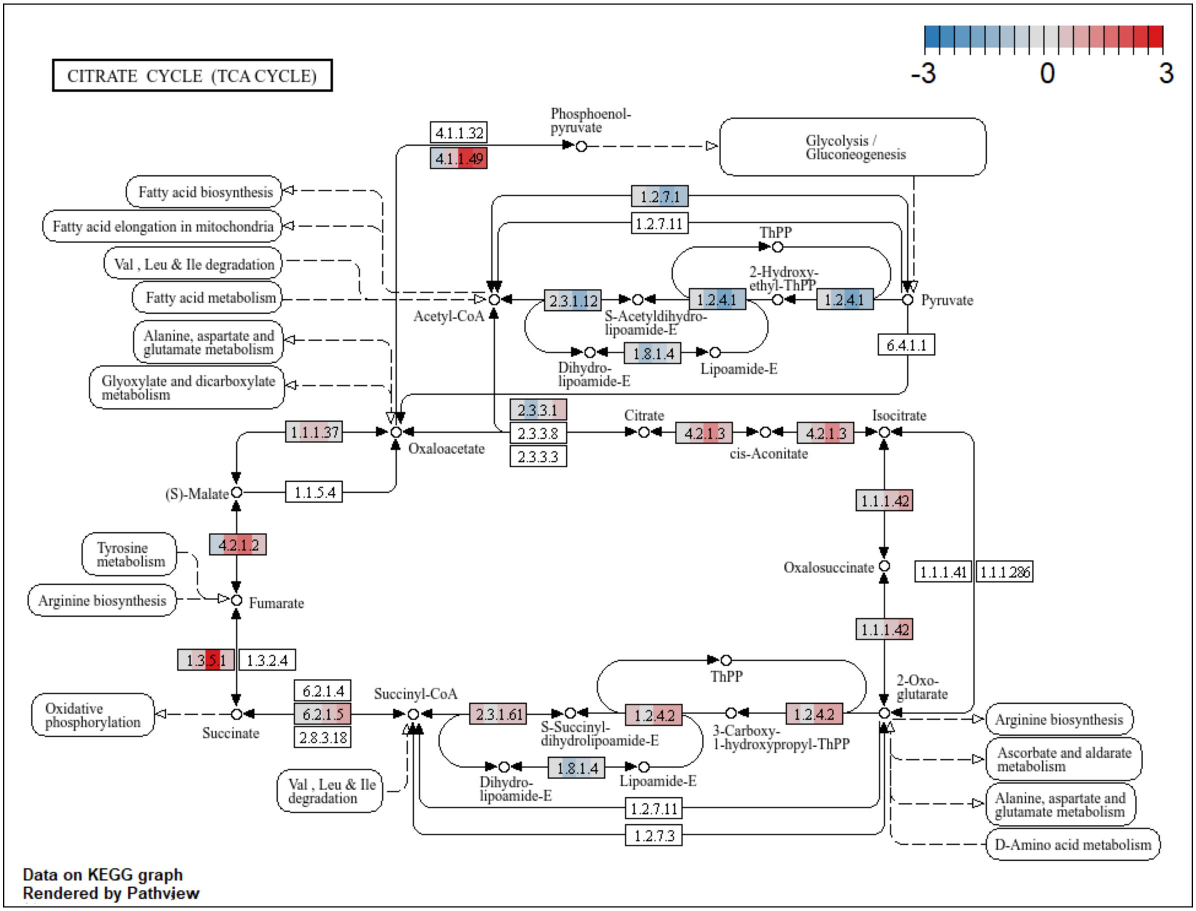
CITRATE CYCLE (TCA CYCLE)
-3
0
3
Phosphoenol-pyruvate
Glycolysis / Gluconeogenesis
4.1.1.32
4.1.1.49
Fatty acid biosynthesis
Fatty acid elongation in mitochondria
Val , Leu & Ile degradation
Fatty acid metabolism
Alanine, aspartate and glutamate metabolism
Glyoxylate and dicarboxylate metabolism
1.2.7.1
1.2.7.11
ThPP
2-Hydroxy-ethyl-ThPP
2.3.1.12
1.2.4.1
1.2.4.1
Pyruvate
Acetyl-CoA
S-Acetyldihydro-lipoamide-E
1.8.1.4
Dihydro-lipoamide-E
Lipoamide-E
6.4.1.1
2.3.3.1
2.3.3.8
2.3.3.3
Citrate
4.2.1.3
4.2.1.3
cis-Aconitate
Isocitrate
1.1.1.37
Oxaloacetate
(S)-Malate
1.1.5.4
1.1.1.42
4.2.1.2
Tyrosine metabolism
Arginine biosynthesis
Fumarate
Oxalosuccinate
1.1.1.41
1.1.1.286
1.1.1.42
1.3.5.1
1.3.2.4
6.2.1.4
6.2.1.5
2.8.3.18
Succinyl-CoA
2.3.1.61
S-Succinyl-dihydrolipoamide-E
1.2.4.2
1.2.4.2
3-Carboxy-1-hydroxypropyl-ThPP
2-Oxo-glutarate
Oxidative phosphorylation
Succinate
Val , Leu & Ile degradation
Dihydro-lipoamide-E
1.8.1.4
Lipoamide-E
1.2.7.11
1.2.7.3
Arginine biosynthesis
Ascorbate and aldarate metabolism
Alanine, aspartate and glutamate metabolism
D-Amino acid metabolism
Data on KEGG graph
Rendered by Pathview

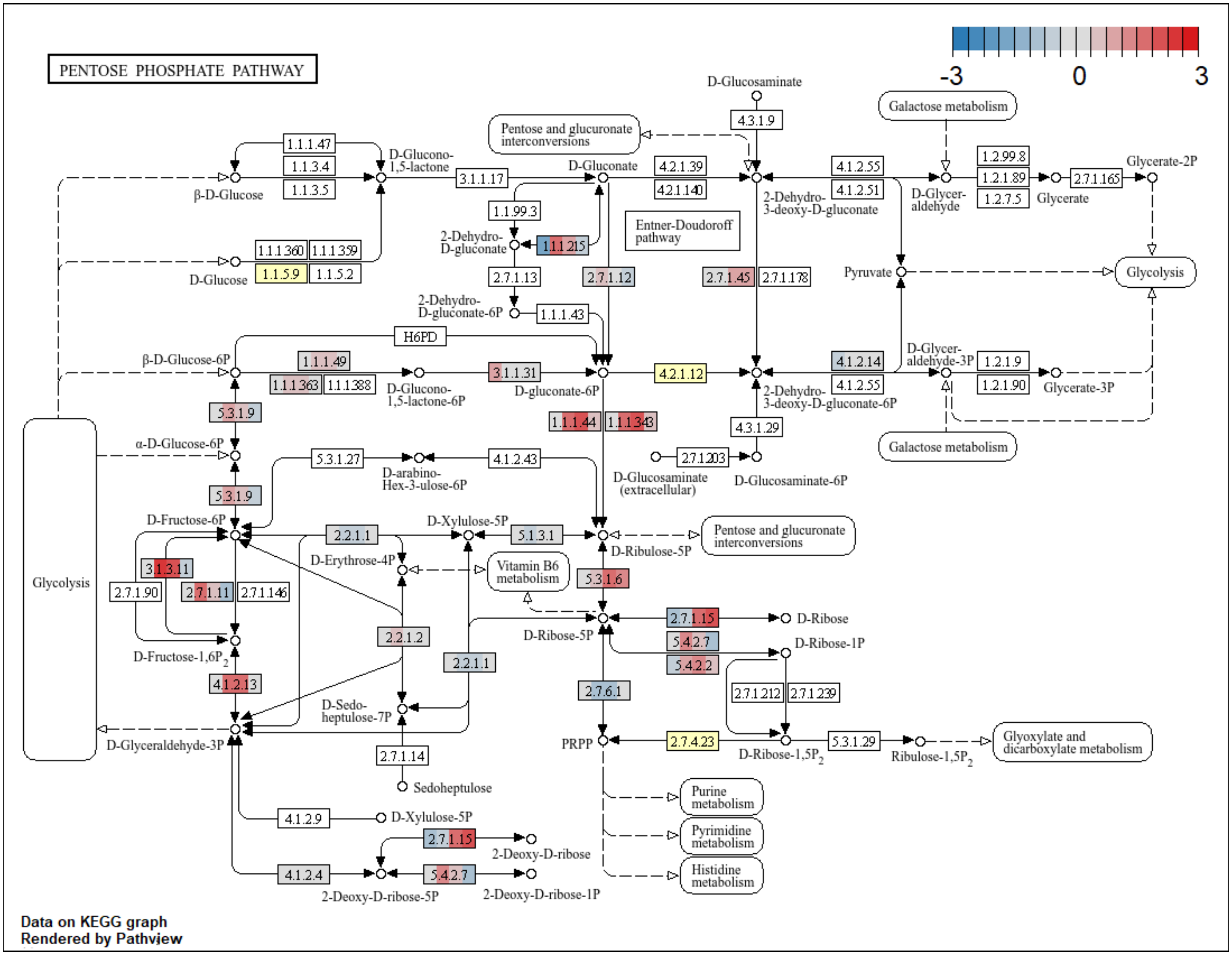
PENTOSE PHOSPHATE PATHWAY
-3
0
3
D-Glucosaminate
Galactose metabolism
Pentose and glucuronate interconversions
D-Glucono-1,5-lactone
β-D-Glucose
D-Gluconate
2-Dehydro-3-deoxy-D-gluconate
D-Glyceraldehyde
Glycerate
Glycerate-2P
2-Dehydro-D-gluconate
Entner-Doudoroff pathway
D-Glucose
Pyruvate
Glycolysis
2-Dehydro-D-gluconate-6P
H6PD
β-D-Glucose-6P
D-Glucono-1,5-lactone-6P
D-gluconate-6P
2-Dehydro-3-deoxy-D-gluconate-6P
D-Glyceraldehyde-3P
Glycerate-3P
α-D-Glucose-6P
D-arabino-Hex-3-ulose-6P
D-Glucosamine (extracellular)
D-Glucosamine-6P
D-Fructose-6P
D-Xylulose-5P
D-Ribulose-5P
D-Erythrose-4P
Vitamin B6 metabolism
D-Fructose-1,6P2
D-Ribose-5P
D-Ribose
D-Ribose-1P
D-Sedo-heptulose-7P
PRPP
D-Ribose-1,5P2
Ribulose-1,5P2
Glyoxylate and dicarboxylate metabolism
D-Glyceraldehyde-3P
Sedoheptulose
Purine metabolism
Pyrimidine metabolism
Histidine metabolism
2-Deoxy-D-ribose
2-Deoxy-D-ribose-5P
2-Deoxy-D-ribose-1P
Data on KEGG graph
Rendered by Pathview

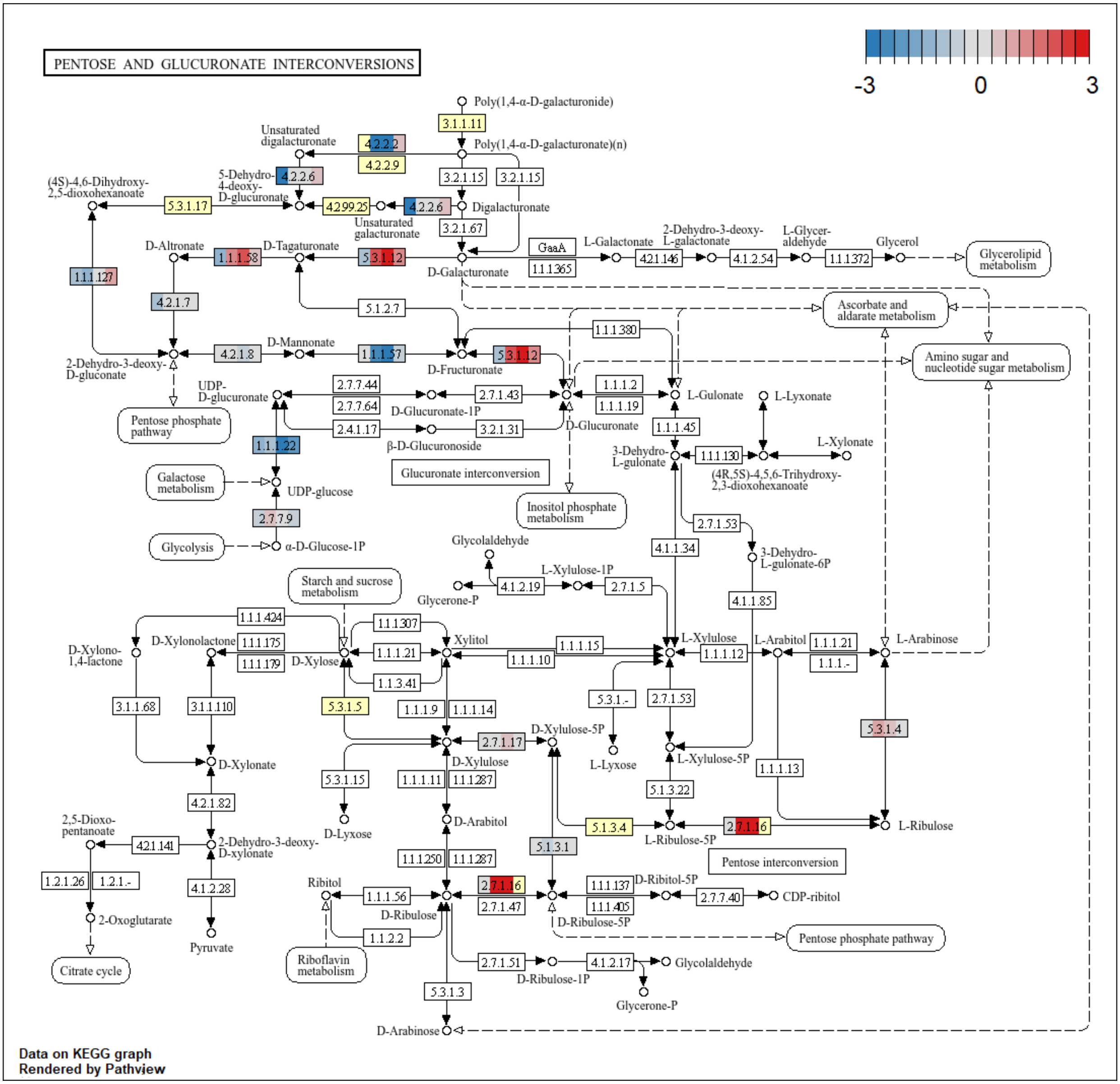
PENTOSE AND GLUCURONATE INTERCONVERSIONS
-3
0
3
Poly(1,4-α-D-galacturonide)
3.1.1.11
Poly(1,4-α-D-galacturonate)(n)
Unsaturated digalacturonate
4.2.2.2
4.2.2.9
3.2.1.15
3.2.1.15
5-Dehydro-4-deoxy-D-glucuronate
4.2.2.6
(4S)-4,6-Dihydroxy-2,5-dioxohexanoate
5.3.1.17
4.2.99.25
4.2.2.6
Digalacturonate
Unsaturated galacturonate
3.2.1.67
D-Altronate
1.1.1.58
D-Tagaturonate
5.3.1.12
D-Galacturonate
GaaA
1.1.1.365
L-Galactonate
4.2.1.146
2-Dehydro-3-deoxy-L-galactonate
4.1.2.54
L-Glyceraldehyde
1.1.1.372
Glycerol
Glycerolipid metabolism
1.1.1.127
4.2.1.7
5.1.2.7
Ascorbate and aldarate metabolism
1.1.1.380
2-Dehydro-3-deoxy-D-gluconate
4.2.1.8
D-Mannonate
1.1.1.57
D-Fructuronate
5.3.1.12
Amino sugar and nucleotide sugar metabolism
UDP-D-glucuronate
2.7.7.44
2.7.7.64
D-Glucuronate-1P
2.7.1.43
1.1.1.2
1.1.1.19
L-Gulonate
L-Lyxonate
Pentose phosphate pathway
2.4.1.17
3.2.1.31
β-D-Glucuronoside
D-Glucuronate
1.1.1.45
1.1.1.22
3-Dehydro-L-gulonate
1.1.1.130
L-Xylonate
Galactose metabolism
UDP-glucose
Glucuronate interconversion
(4R,5S)-4,5,6-Trihydroxy-2,3-dioxohexanoate
Inositol phosphate metabolism
2.7.7.9
2.7.1.53
Glycolysis
α-D-Glucose-1P
Glycolaldehyde
4.1.1.34
3-Dehydro-L-gulonate-6P
L-Xylulose-1P
Starch and sucrose metabolism
4.1.2.19
2.7.1.5
Glycerone-P
4.1.1.85
1.1.1.424
1.1.1.307
D-Xylono-1,4-lactone
D-Xylonolactone
1.1.1.175
1.1.1.21
Xylitol
1.1.1.15
L-Xylulose
L-Arabitol
1.1.1.21
L-Arabinose
1.1.1.179
D-Xylose
1.1.1.10
1.1.1.12
1.1.1.-
1.1.3.41
3.1.1.68
3.1.1.110
5.3.1.5
1.1.1.9
1.1.1.14
5.3.1.-
2.7.1.53
5.3.1.4
D-Xylulose-5P
2.7.1.17
D-Xylonate
D-Xylulose
L-Lyxose
L-Xylulose-5P
5.3.1.15
1.1.1.11
1.1.1.287
1.1.1.13
4.2.1.82
5.1.3.22
2,5-Dioxopentanoate
D-Lyxose
D-Arabitol
5.1.3.4
2.7.1.16
L-Ribulose
4.2.1.141
2-Dehydro-3-deoxy-D-xylonate
5.1.3.1
L-Ribulose-5P
Pentose interconversion
1.1.1.250
1.1.1.287
1.2.1.26
1.2.1.-
4.1.2.28
Ribitol
2.7.1.16
1.1.1.137
D-Ribitol-5P
1.1.1.56
2.7.1.47
1.1.1.405
2.7.7.40
CDP-ribitol
2-Oxoglutarate
D-Ribulose
D-Ribulose-5P
Pyruvate
1.1.2.2
Pentose phosphate pathway
Riboflavin metabolism
2.7.1.51
4.1.2.17
Glycolaldehyde
Citrate cycle
D-Ribulose-1P
5.3.1.3
Glycerone-P
D-Arabinose
Data on KEGG graph
Rendered by Pathview

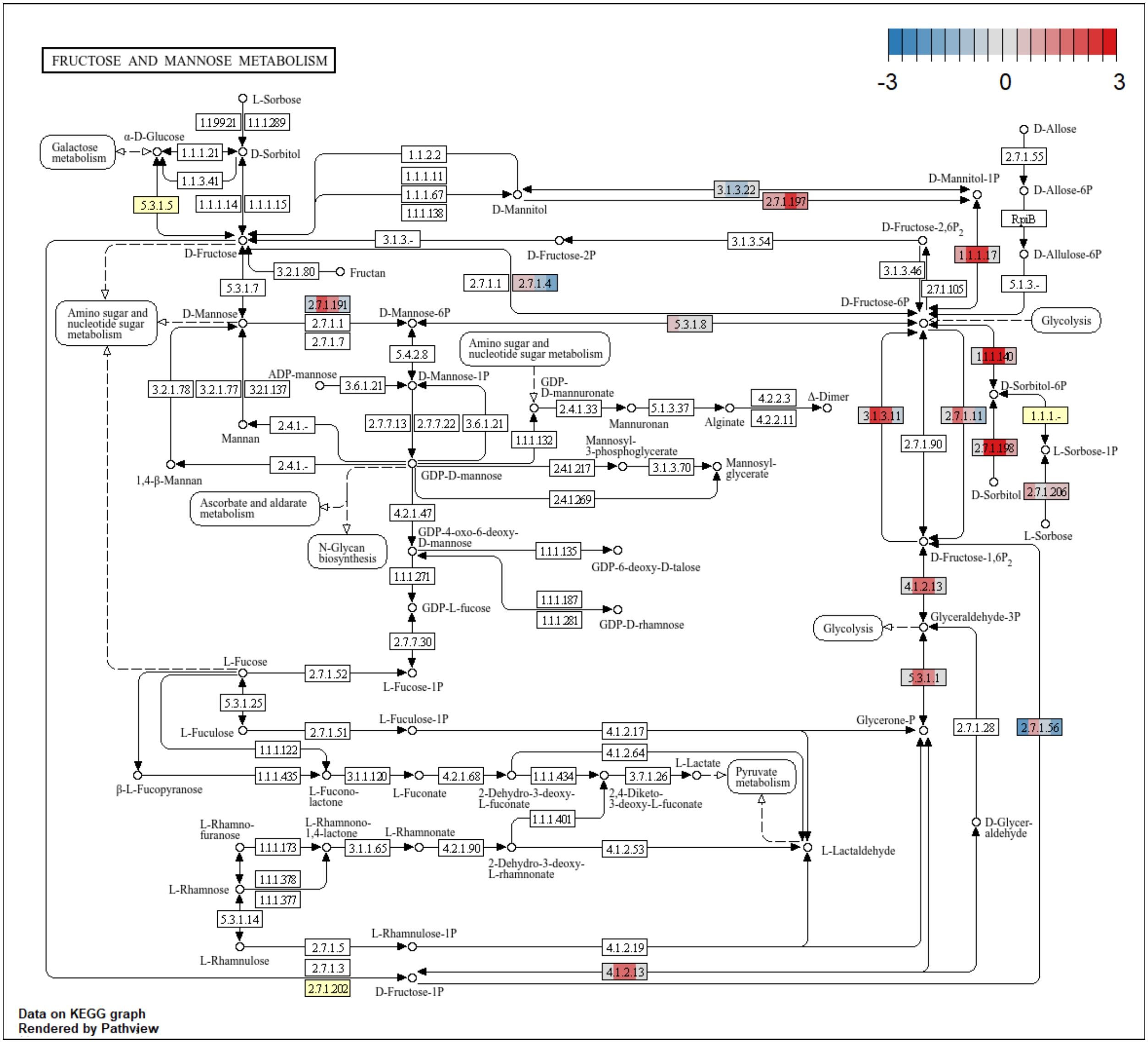

FRUCTOSE AND MANNOSE METABOLISM
-3
0
3
L-Sorbose
1.1.99.21
1.1.1.289
Galactose metabolism
α-D-Glucose
1.1.1.21
D-Sorbitol
1.1.3.41
5.3.1.5
1.1.1.14
1.1.1.15
1.1.2.2
1.1.1.11
1.1.1.67
1.1.1.138
D-Mannitol
3.1.3.22
2.7.1.197
D-Mannitol-1P
D-Allose
2.7.1.55
D-Allose-6P
RpiB
D-Allulose-6P
5.1.3.-
3.1.3.-
D-Fructose-2P
3.1.3.54
D-Fructose-2,6P2
1.1.1.17
D-Fructose
3.2.1.80
Fructan
2.7.1.1
2.7.1.4
3.1.3.46
2.7.1.105
D-Fructose-6P
Glycolysis
5.3.1.7
Amino sugar and nucleotide sugar metabolism
D-Mannose
2.7.1.191
2.7.1.1
2.7.1.7
D-Mannose-6P
5.3.1.8
5.4.2.8
D-Mannose-1P
Amino sugar and nucleotide sugar metabolism
1.1.1.140
D-Sorbitol-6P
ADP-mannose
3.6.1.21
3.2.1.78
3.2.1.77
3.2.1.137
GDP-D-mannuronate
2.4.1.33
Mannuronan
5.1.3.37
Alginate
4.2.2.3
4.2.2.11
Δ-Dimer
3.1.3.11
2.7.1.11
1.1.1.-
2.7.7.13
2.7.7.22
3.6.1.21
1.1.1.132
2.4.1.-
Mannan
2.7.1.90
2.7.1.198
L-Sorbose-1P
Mannosyl-3-phosphoglycerate
2.4.1.-
1,4-β-Mannan
GDP-D-mannose
2.4.1.217
3.1.3.70
Mannosyl-glycerate
D-Sorbitol
2.7.1.206
2.4.1.269
Ascorbate and aldarate metabolism
4.2.1.47
L-Sorbose
GDP-4-oxo-6-deoxy-D-mannose
N-Glycan biosynthesis
1.1.1.135
GDP-6-deoxy-D-talose
D-Fructose-1,6P2
1.1.1.271
4.1.2.13
1.1.1.187
1.1.1.281
GDP-L-fucose
GDP-D-rhamnose
Glyceraldehyde-3P
Glycolysis
2.7.7.30
L-Fucose
2.7.1.52
L-Fucose-1P
5.3.1.1
5.3.1.25
L-Fuculose-1P
Glycerone-P
2.7.1.28
2.7.1.56
L-Fuculose
2.7.1.51
4.1.2.17
1.1.1.122
4.1.2.64
L-Lactate
β-L-Fucopyranose
1.1.1.435
3.1.1.120
4.2.1.68
1.1.1.434
3.7.1.26
Pyruvate metabolism
L-Fucono-lactone
L-Fuconate
2-Dehydro-3-deoxy-L-fuconate
2,4-Diketo-3-deoxy-L-fuconate
1.1.1.401
L-Rhamno-furanose
L-Rhamnono-1,4-lactone
L-Rhamnonate
D-Glycer-aldehyde
1.1.1.173
3.1.1.65
4.2.1.90
4.1.2.53
L-Lactaldehyde
2-Dehydro-3-deoxy-L-rhamnonate
1.1.1.378
L-Rhamnose
1.1.1.377
5.3.1.14
L-Rhamnulose-1P
L-Rhamnulose
2.7.1.5
4.1.2.19
2.7.1.3
4.1.2.13
2.7.1.202
D-Fructose-1P
Data on KEGG graph
Rendered by Pathview

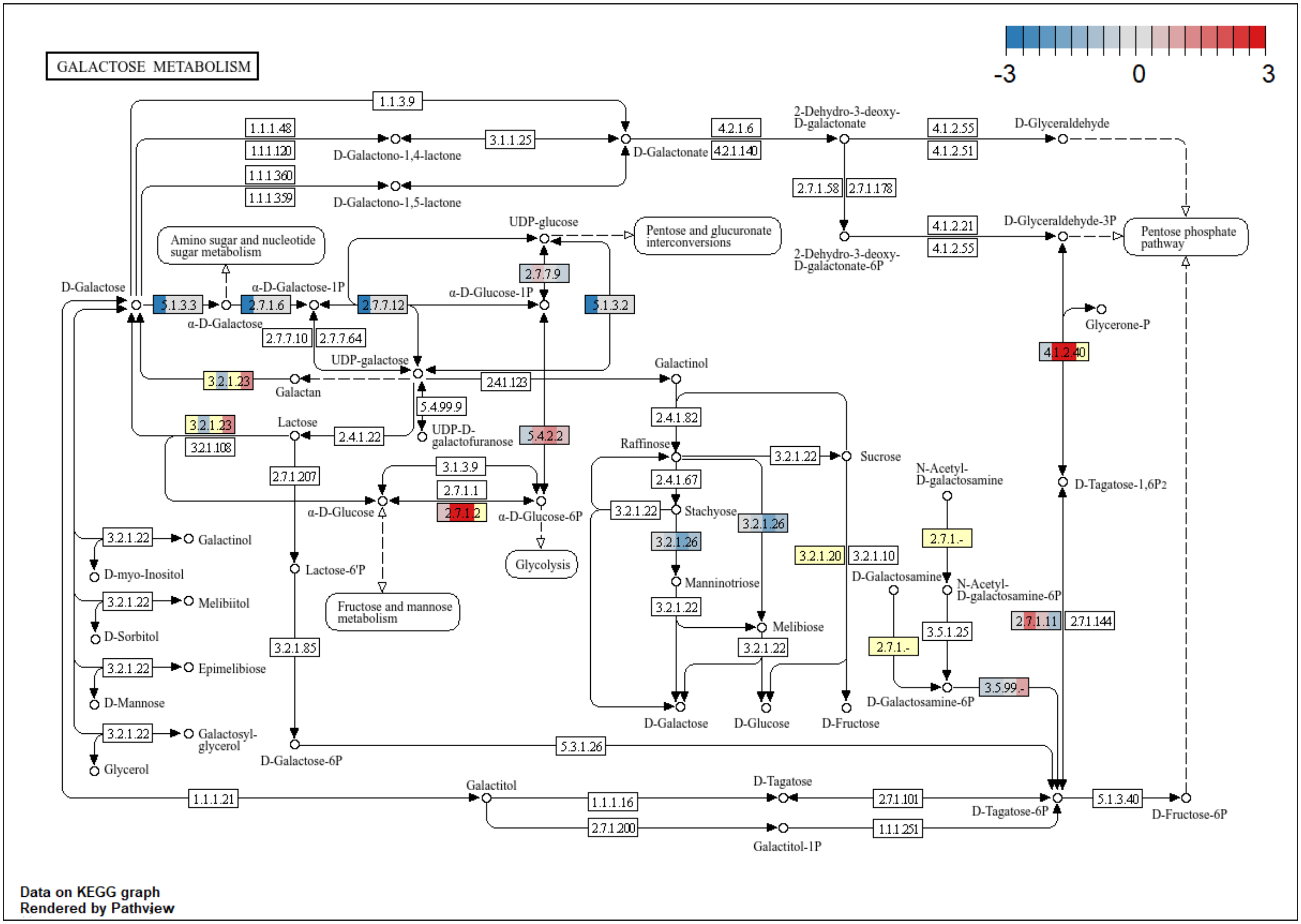
GALACTOSE METABOLISM
-3
0
3
D-Galactose
α-D-Galactose-1P
α-D-Galactose
UDP-glucose
α-D-Glucose-1P
UDP-galactose
Galactan
Lactose
UDP-D-galactofuranose
α-D-Glucose
α-D-Glucose-6P
Glycolysis
Lactose-6'P
Fructose and mannose metabolism
D-Galactose-6P
Galactinol
D-myo-Inositol
Melibiitol
D-Sorbitol
Epimelibiose
D-Mannose
Galactosyl-glycerol
Glycerol
Galactitol
Galactitol-1P
D-Tagatose
D-Tagatose-6P
D-Fructose-6P
D-Galactono-1,4-lactone
D-Galactono-1,5-lactone
D-Galactonate
2-Dehydro-3-deoxy-D-galactonate
2-Dehydro-3-deoxy-D-galactonate-6P
D-Glyceraldehyde
D-Glyceraldehyde-3P
Pentose phosphate pathway
Pentose and glucuronate interconversions
Amino sugar and nucleotide sugar metabolism
Glycerone-P
D-Tagatose-1,6P2
Raffinose
Sucrose
Stachyose
Manninotriose
Melibiose
D-Galactose
D-Glucose
D-Fructose
N-Acetyl-D-galactosamine
N-Acetyl-D-galactosamine-6P
D-Galactosamine
D-Galactosamine-6P
1.1.3.9
1.1.1.48
1.1.1.120
1.1.1.360
1.1.1.359
3.1.1.25
4.2.1.6
4.2.1.140
4.1.2.55
4.1.2.51
2.7.1.58
2.7.1.178
4.1.2.21
4.1.2.55
5.1.3.3
2.7.1.6
2.7.7.12
2.7.7.10
2.7.7.64
2.7.7.9
5.1.3.2
3.2.1.23
2.4.1.123
5.4.99.9
3.2.1.23
3.2.1.108
2.4.1.22
5.4.2.2
2.7.1.207
3.1.3.9
2.7.1.1
2.7.1.2
3.2.1.85
3.2.1.22
3.2.1.22
3.2.1.22
3.2.1.22
5.3.1.26
1.1.1.21
1.1.1.16
2.7.1.200
2.7.1.101
1.1.1.251
5.1.3.40
2.4.1.82
3.2.1.22
2.4.1.67
3.2.1.22
3.2.1.26
3.2.1.26
3.2.1.22
3.2.1.22
3.2.1.20
3.2.1.10
4.1.2.40
2.7.1.-
3.5.1.25
2.7.1.11
2.7.1.144
2.7.1.-
3.5.99.-
Data on KEGG graph
Rendered by Pathview

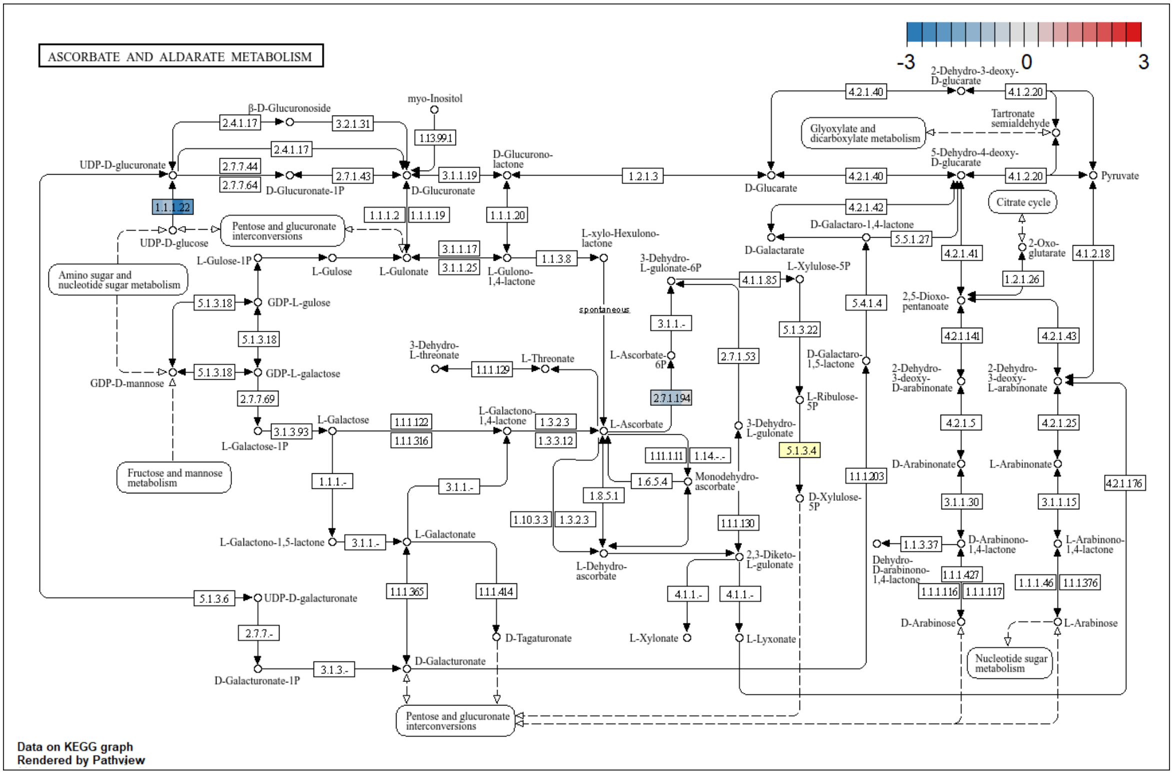
ASCORBATE AND ALDARATE METABOLISM
-3
0
3
UDP-D-glucuronate
β-D-Glucuronoside
myo-Inositol
D-Glucuronate
D-Glucurono-lactone
D-Glucurate
D-Glucuronate-1P
UDP-D-glucose
Pentose and glucuronate interconversions
Amino sugar and nucleotide sugar metabolism
L-Gulose-1P
L-Gulose
L-Gulonate
L-Gulono-1,4-lactone
L-xylo-Hexulono-lactone
spontaneous
GDP-L-gulose
GDP-L-galactose
GDP-D-mannose
L-Galactose-1P
L-Galactose
L-Galactono-1,4-lactone
L-Ascorbate
L-Ascorbate-6P
3-Dehydro-L-gulonate-6P
3-Dehydro-L-threonate
L-Threonate
Monodehydro-ascorbate
L-Dehydro-ascorbate
2,3-Diketo-L-gulonate
3-Dehydro-L-gulonate
L-Xylulose-5P
L-Ribulose-5P
D-Xylulose-5P
L-Xylonate
L-Lyxonate
Fructose and mannose metabolism
L-Galactono-1,5-lactone
L-Galactonate
D-Tagaturonate
UDP-D-galacturonate
D-Galacturonate-1P
D-Galacturonate
D-Galactarate
D-Galactaro-1,4-lactone
D-Galactaro-1,5-lactone
2-Dehydro-3-deoxy-D-glucarate
Tartronate semialdehyde
Glyoxylate and dicarboxylate metabolism
5-Dehydro-4-deoxy-D-glucarate
Pyruvate
Citrate cycle
2-Oxo-glutarate
2,5-Dioxo-pentanoate
2-Dehydro-3-deoxy-D-arabinonate
2-Dehydro-3-deoxy-L-arabinonate
D-Arabinonate
L-Arabinonate
D-Arabinono-1,4-lactone
L-Arabinono-1,4-lactone
Dehydro-D-arabinono-1,4-lactone
D-Arabinose
L-Arabinose
Nucleotide sugar metabolism
2.4.1.17
3.2.1.31
1.13.99.1
2.7.7.44
2.7.7.64
2.7.1.43
3.1.1.19
1.2.1.3
4.2.1.40
4.1.2.20
1.1.1.22
1.1.1.2
1.1.1.19
1.1.1.20
3.1.1.17
3.1.1.25
1.1.3.8
4.1.1.85
5.1.3.18
3.1.1.-
2.7.1.194
2.7.1.53
5.1.3.22
5.4.1.4
5.5.1.27
4.2.1.42
4.2.1.41
4.1.2.18
1.2.1.26
4.2.1.43
1.1.1.129
2.7.7.69
3.1.3.93
1.1.1.122
1.1.1.316
1.3.2.3
1.3.3.12
1.11.1.11
1.14.-.-
1.6.5.4
1.8.5.1
1.10.3.3
5.1.3.4
1.1.1.203
1.1.1.130
4.1.1.-
4.2.1.5
4.2.1.25
4.2.1.176
3.1.1.30
3.1.1.15
1.1.3.37
1.1.1.427
1.1.1.116
1.1.1.117
1.1.1.46
1.1.1.376
1.1.1.-
1.1.1.365
1.1.1.414
5.1.3.6
2.7.7.-
3.1.3.-
Data on KEGG graph
Rendered by Pathview

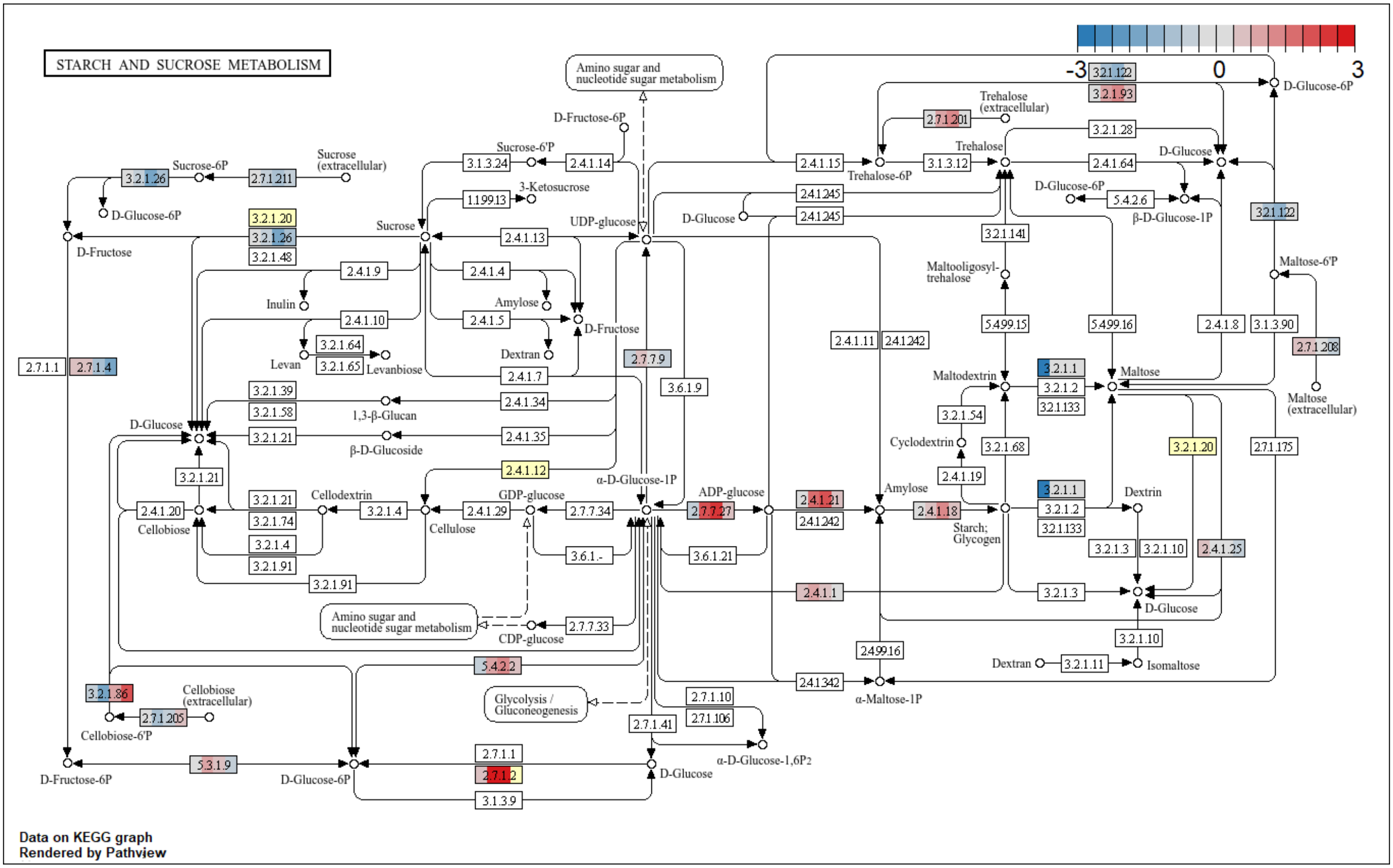
STARCH AND SUCROSE METABOLISM
-3
0
3
Amino sugar and nucleotide sugar metabolism
Glycolysis / Gluconeogenesis
D-Glucose-6P
D-Fructose-6P
Sucrose-6'P
Sucrose-6P
Sucrose (extracellular)
3-Ketosucrose
UDP-glucose
Sucrose
D-Fructose
Inulin
Levan
Levanbiose
Amylose
Dextran
1,3-β-Glucan
β-D-Glucoside
D-Glucose
Cellodextrin
Cellobiose
Cellulose
GDP-glucose
CDP-glucose
α-D-Glucose-1P
ADP-glucose
Amylose
Starch; Glycogen
Trehalose (extracellular)
Trehalose-6P
Trehalose
β-D-Glucose-1P
Maltooligosyl-trehalose
Maltodextrin
Maltose
Maltose-6'P
Maltose (extracellular)
Cyclodextrin
Dextrin
Isomaltose
α-Maltose-1P
α-D-Glucose-1,6P2
Cellobiose-6'P
Cellobiose (extracellular)
3.2.1.26
2.7.1.211
3.2.1.20
3.2.1.48
2.4.1.9
2.4.1.10
3.2.1.64
3.2.1.65
2.7.1.1
2.7.1.4
3.2.1.39
3.2.1.58
3.2.1.21
3.2.1.74
3.2.1.4
3.2.1.91
2.4.1.20
3.1.3.24
2.4.1.14
1.1.99.13
2.4.1.13
2.4.1.4
2.4.1.5
2.4.1.7
2.4.1.34
2.4.1.35
2.4.1.12
2.4.1.29
2.7.7.34
3.6.1.-
2.7.7.33
2.7.7.9
3.6.1.9
2.7.7.27
3.6.1.21
5.4.2.2
2.7.1.10
2.7.1.106
2.7.1.41
2.7.1.2
3.1.3.9
5.3.1.9
3.2.1.86
2.7.1.205
2.4.1.15
3.1.3.12
2.4.1.245
2.7.1.201
3.2.1.122
3.2.1.93
3.2.1.28
2.4.1.64
5.4.2.6
3.2.1.141
5.4.99.15
5.4.99.16
2.4.1.8
3.1.3.90
2.7.1.208
2.4.1.11
2.4.1.242
2.4.1.21
2.4.1.18
3.2.1.1
3.2.1.2
3.2.1.133
3.2.1.54
3.2.1.68
2.4.1.19
3.2.1.3
3.2.1.10
2.4.1.25
2.7.1.175
2.4.1.1
2.4.99.16
2.4.1.342
3.2.1.11
Data on KEGG graph
Rendered by Pathview

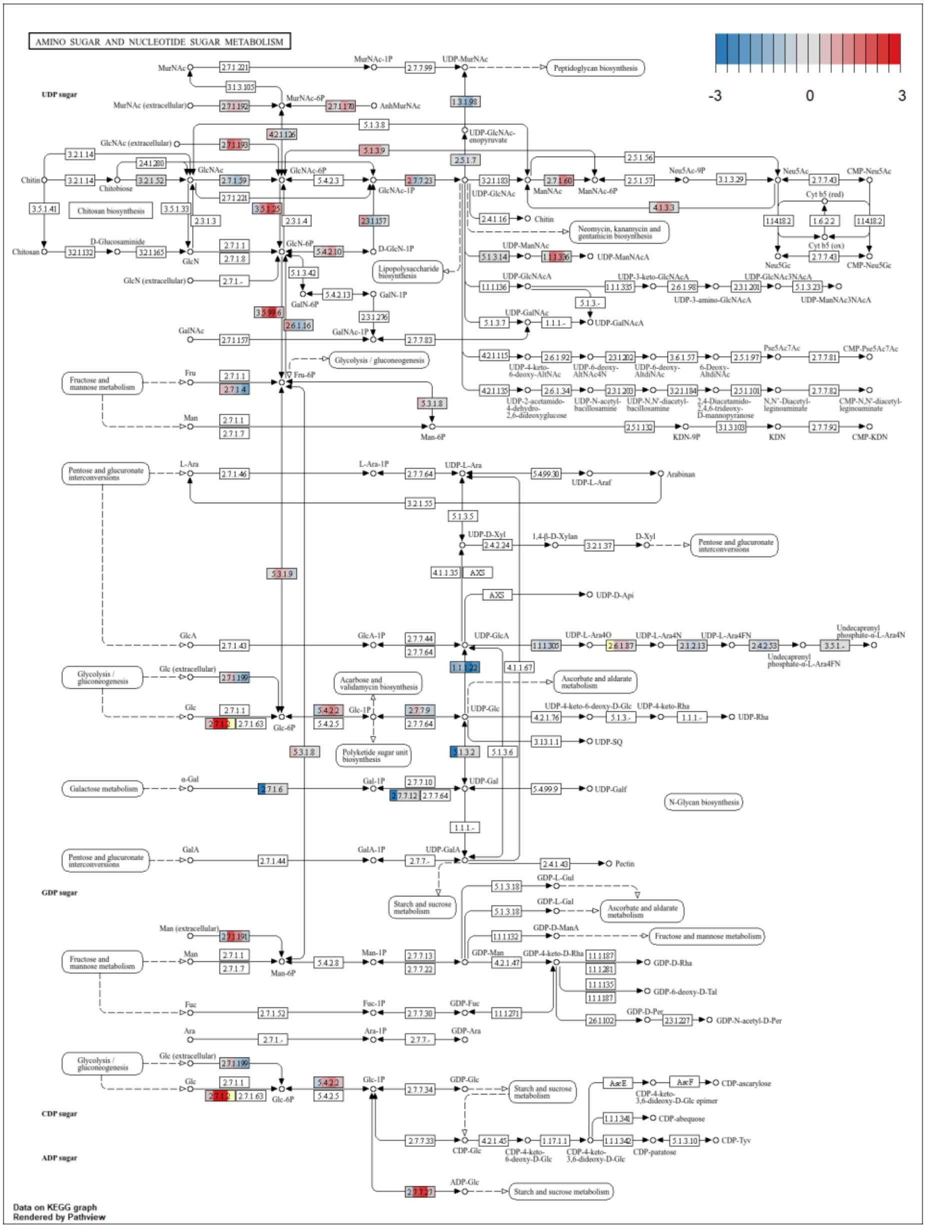
AMINO SUGAR AND NUCLEOTIDE SUGAR METABOLISM
-3
0
3
UDP sugar
GDP sugar
CDP sugar
ADP sugar
Peptidoglycan biosynthesis
Chitosan biosynthesis
Fructose and mannose metabolism
Pentose and glucuronate interconversions
Glycolysis / gluconeogenesis
Galactose metabolism
Starch and sucrose metabolism
Ascorbate and aldarate metabolism
N-Glycan biosynthesis
Data on KEGG graph
Rendered by Pathview

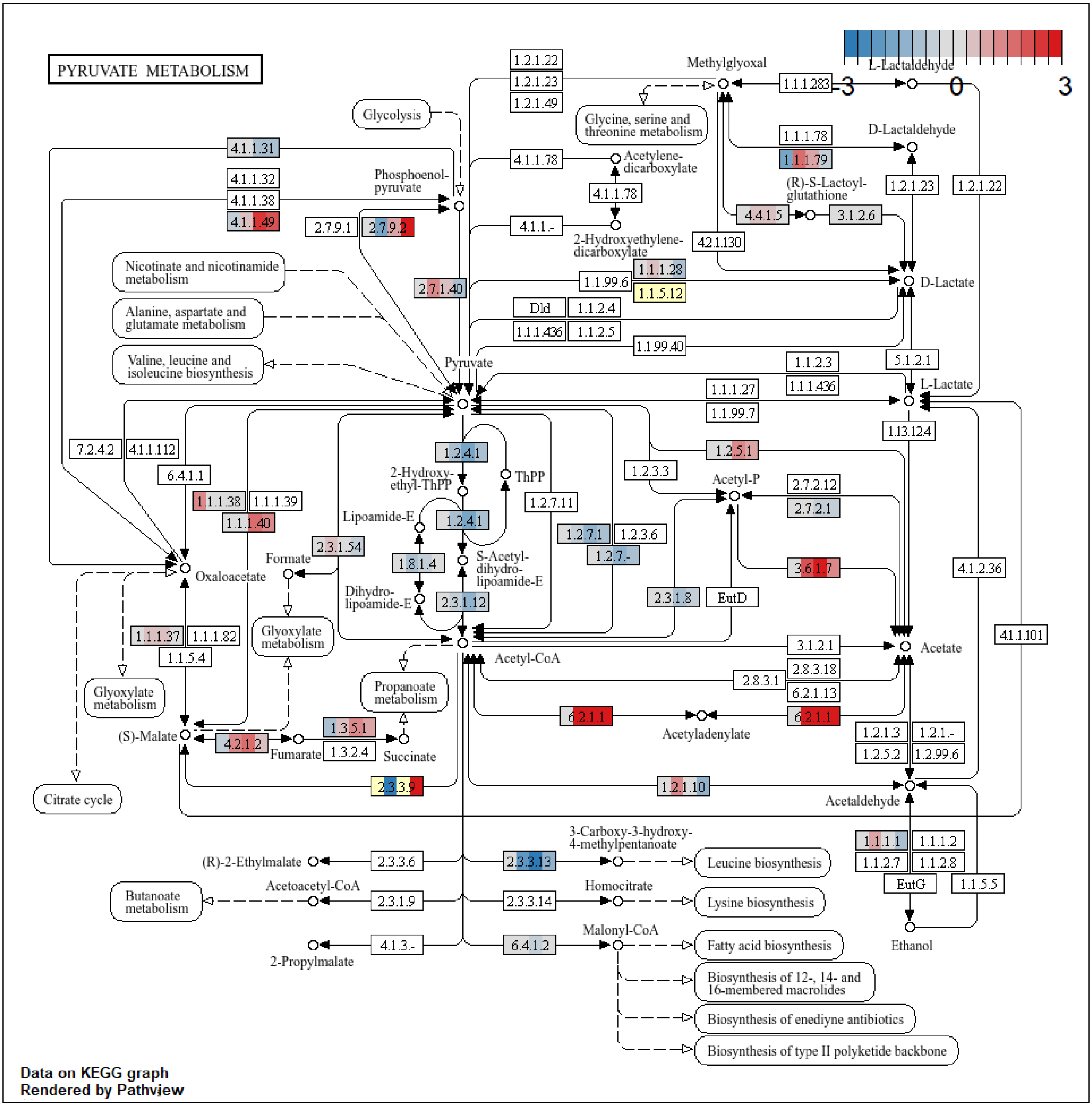

PYRUVATE METABOLISM
-3
0
3
Methylglyoxal
L-Lactaldehyde
D-Lactaldehyde
Glycolysis
Glycine, serine and threonine metabolism
Phosphoenol-pyruvate
Acetylene-dicarboxylate
2-Hydroxyethylene-dicarboxylate
(R)-S-Lactoyl-glutathione
D-Lactate
L-Lactate
Nicotinate and nicotinamide metabolism
Alanine, aspartate and glutamate metabolism
Valine, leucine and isoleucine biosynthesis
Pyruvate
2-Hydroxy-ethyl-ThPP
ThPP
Lipoamide-E
S-Acetyl-dihydro-lipoamide-E
Dihydro-lipoamide-E
Acetyl-P
Oxaloacetate
Formate
Glyoxylate metabolism
Acetyl-CoA
Acetate
Propanoate metabolism
(S)-Malate
Fumarate
Succinate
Acetyladenylate
Acetaldehyde
Citrate cycle
Ethanol
3-Carboxy-3-hydroxy-4-methylpentanoate
(R)-2-Ethylmalate
Leucine biosynthesis
Butanoate metabolism
Acetoacetyl-CoA
Homocitrate
Lysine biosynthesis
2-Propylmalate
Malonyl-CoA
Fatty acid biosynthesis
Biosynthesis of 12-, 14- and 16-membered macrolides
Biosynthesis of enediyne antibiotics
Biosynthesis of type II polyketide backbone
1.2.1.22
1.2.1.23
1.2.1.49
1.1.1.283
1.1.1.78
1.1.1.79
4.1.1.31
4.1.1.32
4.1.1.38
4.1.1.49
2.7.9.1
2.7.9.2
4.1.1.78
4.1.1.-
4.4.1.5
3.1.2.6
4.2.1.130
1.1.1.28
1.1.99.6
1.1.5.12
2.7.1.40
D-ld
1.1.2.4
1.1.1.436
1.1.2.5
1.1.99.40
1.1.2.3
5.1.2.1
1.1.1.27
1.1.99.7
1.13.12.4
7.2.4.2
4.1.1.112
6.4.1.1
1.2.4.1
1.2.5.1
1.2.3.3
2.7.2.12
2.7.2.1
1.1.1.38
1.1.1.39
1.1.1.40
1.2.7.11
2.3.1.54
1.2.7.1
1.2.3.6
1.2.7.-
1.8.1.4
3.6.1.7
4.1.2.36
2.3.1.12
2.3.1.8
EutD
1.1.1.37
1.1.1.82
1.1.5.4
3.1.2.1
4.1.1.101
2.8.3.18
2.8.3.1
6.2.1.13
6.2.1.1
1.3.5.1
4.2.1.2
1.3.2.4
1.2.1.3
1.2.1.-
1.2.5.2
1.2.99.6
2.3.3.9
1.2.1.10
1.1.1.1
1.1.1.2
1.1.2.7
1.1.2.8
EutG
1.1.5.5
2.3.3.6
2.3.3.13
2.3.1.9
2.3.3.14
4.1.3.-
6.4.1.2
Data on KEGG graph
Rendered by Pathview

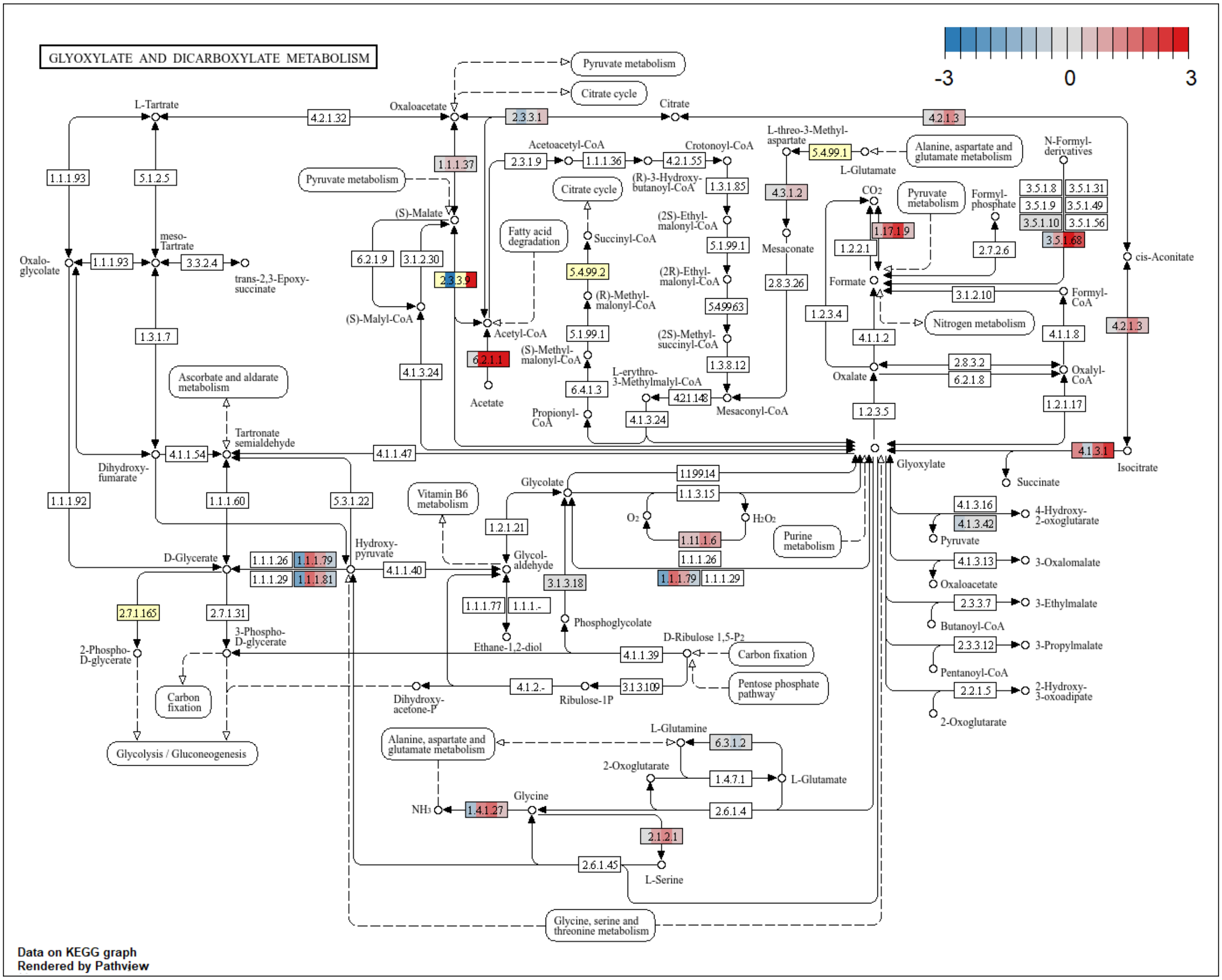
GLYOXYLATE AND DICARBOXYLATE METABOLISM
-3
0
3
Pyruvate metabolism
Citrate cycle
L-Tartrate
Oxaloacetate
Citrate
Acetoacetyl-CoA
Crotonoyl-CoA
L-threo-3-Methyl-aspartate
Alanine, aspartate and glutamate metabolism
N-Formyl-derivatives
L-Glutamate
Mesaconate
CO2
Formate
Formyl-phosphate
Formyl-CoA
cis-Aconitate
Oxalo-glycolate
meso-Tartrate
trans-2,3-Epoxy-succinate
(S)-Malate
(S)-Malyl-CoA
Fatty acid degradation
Succinyl-CoA
Acetyl-CoA
Acetate
(R)-Methyl-malonyl-CoA
(S)-Methyl-malonyl-CoA
Propionyl-CoA
(2S)-Ethyl-malonyl-CoA
(2R)-Ethyl-malonyl-CoA
(2S)-Methyl-succinyl-CoA
L-erythro-3-Methylmalyl-CoA
Mesaconyl-CoA
Nitrogen metabolism
Oxalate
Oxalyl-CoA
Ascorbate and aldarate metabolism
Tartronate semialdehyde
Dihydroxy-fumarate
Glyoxylate
Isocitrate
Succinate
Glycolate
Vitamin B6 metabolism
O2
H2O2
Purine metabolism
D-Glycerate
Hydroxy-pyruvate
Glycol-aldehyde
4-Hydroxy-2-oxoglutarate
Pyruvate
3-Oxalomalate
Oxaloacetate
3-Ethylmalate
Butanoyl-CoA
3-Propylmalate
Pentanoyl-CoA
2-Hydroxy-3-oxoadipate
2-Oxoglutarate
2-Phospho-D-glycerate
3-Phospho-D-glycerate
Phosphoglycolate
Ethane-1,2-diol
D-Ribulose 1,5-P2
Carbon fixation
Pentose phosphate pathway
Dihydroxy-acetone-P
Ribulose-1P
Glycolysis / Gluconeogenesis
L-Glutamine
2-Oxoglutarate
L-Glutamate
NH3
Glycine
L-Serine
Glycine, serine and threonine metabolism
Data on KEGG graph
Rendered by Pathview

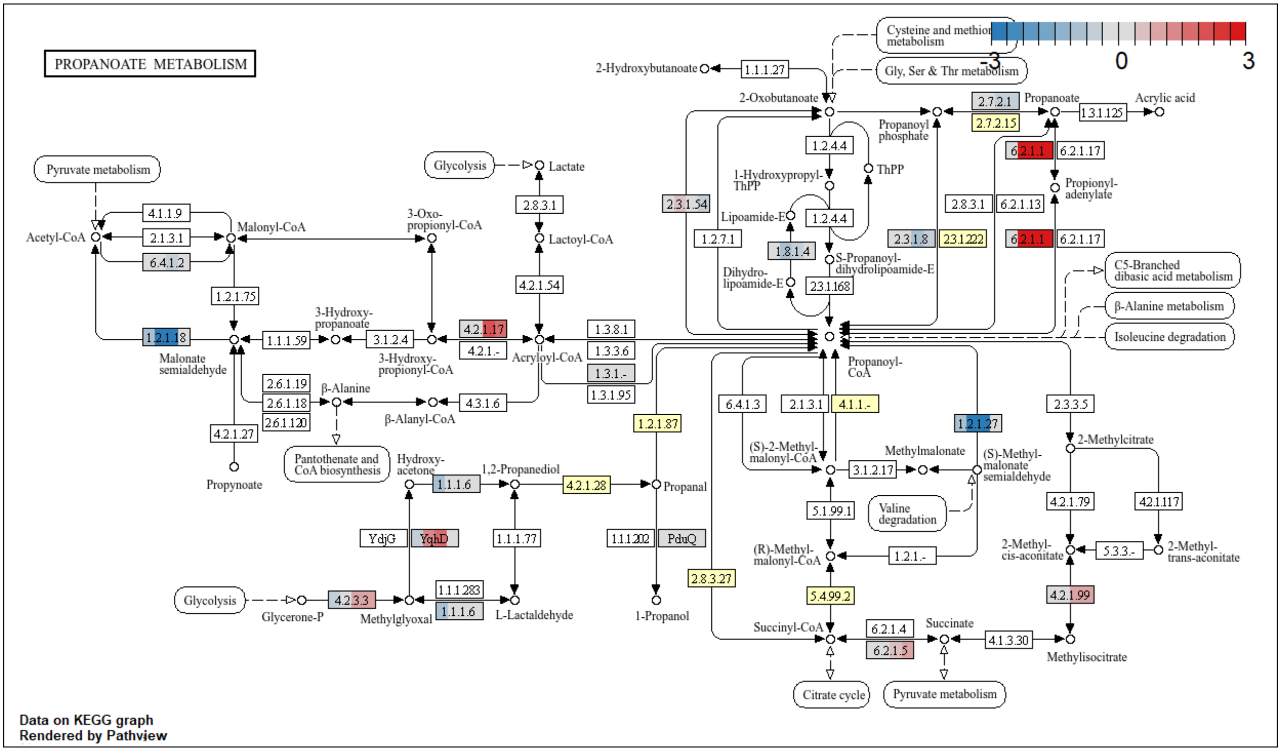
PROPANOATE METABOLISM
-3
0
3
Cysteine and methionine metabolism
Gly, Ser & Thr metabolism
2-Hydroxybutanoate
2-Oxobutanoate
Propanoate
Acrylic acid
Propanoyl phosphate
ThPP
1-Hydroxypropyl-ThPP
Lipoamide-E
Dihydro-lipoamide-E
S-Propanoyl-dihydrolipoamide-E
Propionyl-adenylate
C5-Branched dibasic acid metabolism
β-Alanine metabolism
Isoleucine degradation
Propanoyl-CoA
Pyruvate metabolism
Acetyl-CoA
Malonyl-CoA
3-Oxo-propionyl-CoA
Glycolysis
Lactate
Lactoyl-CoA
3-Hydroxy-propanoate
3-Hydroxy-propionyl-CoA
Acryloyl-CoA
Malonate semialdehyde
β-Alanine
β-Alanyl-CoA
Pantothenate and CoA biosynthesis
Propynoate
Hydroxy-acetone
1,2-Propanediol
Propanal
1-Propanol
Glycerone-P
Methylglyoxal
L-Lactaldehyde
(S)-2-Methyl-malonyl-CoA
Methylmalonate
(S)-Methyl-malonate semialdehyde
Valine degradation
(R)-Methyl-malonyl-CoA
2-Methylcitrate
2-Methyl-cis-aconitate
2-Methyl-trans-aconitate
Methylisocitrate
Succinyl-CoA
Succinate
Citrate cycle
Data on KEGG graph
Rendered by Pathview

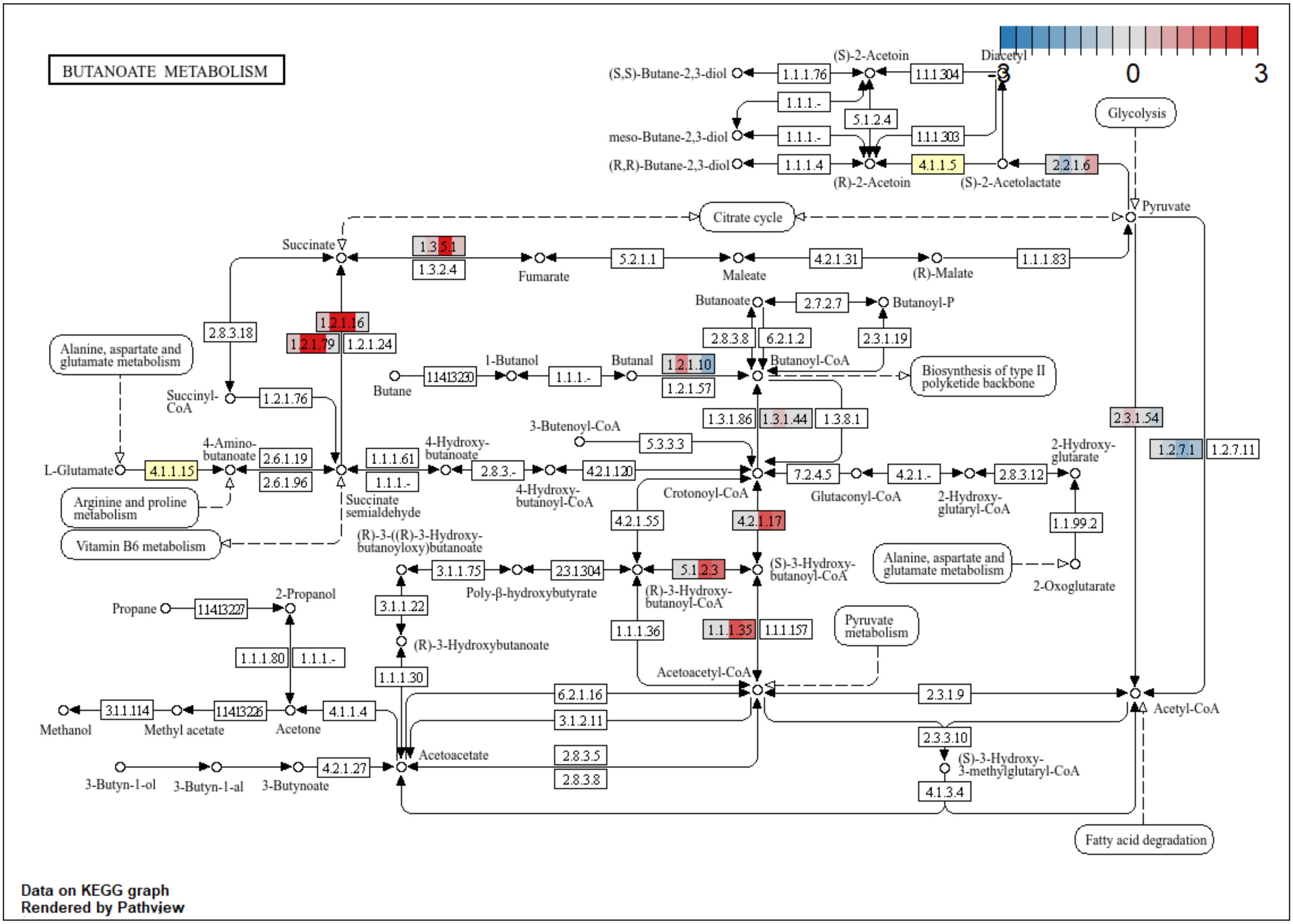
BUTANOATE METABOLISM
-3
0
3
(S,S)-Butane-2,3-diol
(S)-2-Acetoin
Diacetyl
meso-Butane-2,3-diol
(R,R)-Butane-2,3-diol
(R)-2-Acetoin
(S)-2-Acetolactate
Glycolysis
Pyruvate
Citrate cycle
Succinate
Fumarate
Maleate
(R)-Malate
Butanoate
Butanoyl-P
Butanoyl-CoA
Biosynthesis of type II polyketide backbone
Alanine, aspartate and glutamate metabolism
Succinyl-CoA
1-Butanol
Butanal
Butane
3-Butenoyl-CoA
4-Amino-butanoate
4-Hydroxy-butanoate
L-Glutamate
Succinate semialdehyde
4-Hydroxy-butanoyl-CoA
Crotonoyl-CoA
Glutaconyl-CoA
2-Hydroxy-glutaryl-CoA
2-Hydroxy-glutarate
Arginine and proline metabolism
Vitamin B6 metabolism
(R)-3-((R)-3-Hydroxy-butanoyloxy)butanoate
Poly-β-hydroxybutyrate
(R)-3-Hydroxy-butanoyl-CoA
(S)-3-Hydroxy-butanoyl-CoA
Alanine, aspartate and glutamate metabolism
2-Oxoglutarate
Propane
2-Propanol
(R)-3-Hydroxybutanoate
Pyruvate metabolism
Acetoacetyl-CoA
Acetyl-CoA
Methanol
Methyl acetate
Acetone
Acetoacetate
(S)-3-Hydroxy-3-methylglutaryl-CoA
3-Butyn-1-ol
3-Butyn-1-al
3-Butynoate
Fatty acid degradation
Data on KEGG graph
Rendered by Pathview

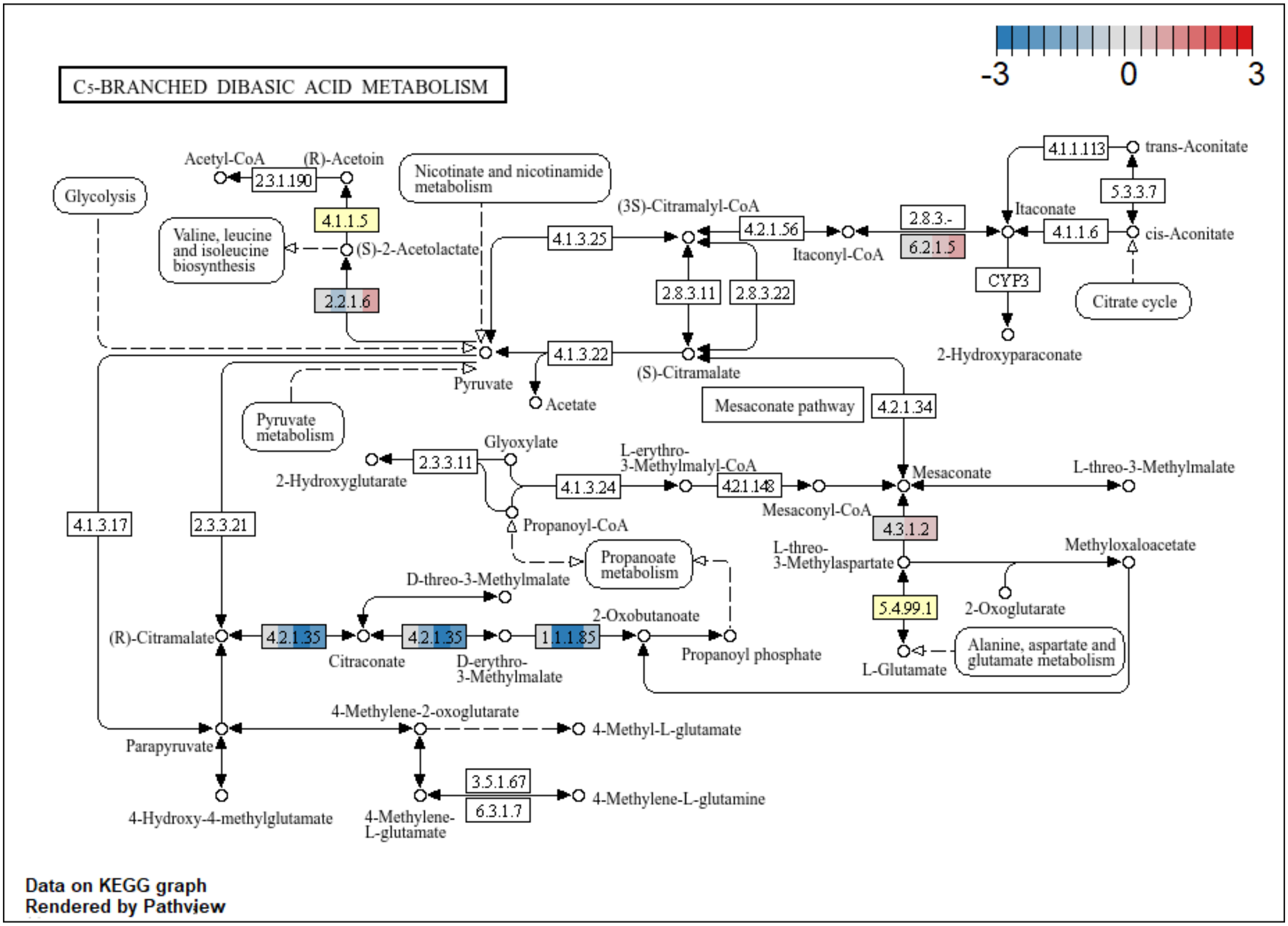
C5-BRANCHED DIBASIC ACID METABOLISM
-3
0
3
Acetyl-CoA
(R)-Acetoin
2.3.1.190
Glycolysis
Nicotinate and nicotinamide metabolism
4.1.1.5
Valine, leucine and isoleucine biosynthesis
(S)-2-Acetolactate
2.2.1.6
(3S)-Citramalyl-CoA
4.1.3.25
4.2.1.56
Itaconyl-CoA
2.8.3.-
6.2.1.5
4.1.1.113
trans-Aconitate
5.3.3.7
Itaconate
4.1.1.6
cis-Aconitate
CYP3
Citrate cycle
2-Hydroxyparaconate
2.8.3.11
2.8.3.22
4.1.3.22
Pyruvate
Acetate
(S)-Citramalate
Mesaconate pathway
4.2.1.34
Pyruvate metabolism
Glyoxylate
2.3.3.11
2-Hydroxyglutarate
L-erythro-3-Methylmalyl-CoA
4.1.3.24
4.2.1.148
Mesaconate
L-threo-3-Methylmalate
Mesaconyl-CoA
Propanoyl-CoA
4.3.1.2
L-threo-3-Methylaspartate
Methyloxaloacetate
4.1.3.17
2.3.3.21
Propanoate metabolism
D-threo-3-Methylmalate
5.4.99.1
2-Oxoglutarate
2-Oxobutanoate
(R)-Citramalate
4.2.1.35
4.2.1.35
1.1.1.85
Citraconate
D-erythro-3-Methylmalate
Propanoyl phosphate
L-Glutamate
Alanine, aspartate and glutamate metabolism
4-Methylene-2-oxoglutarate
4-Methyl-L-glutamate
Parapyruvate
3.5.1.67
6.3.1.7
4-Methylene-L-glutamine
4-Hydroxy-4-methylglutamate
4-Methylene-L-glutamate
Data on KEGG graph
Rendered by Pathview

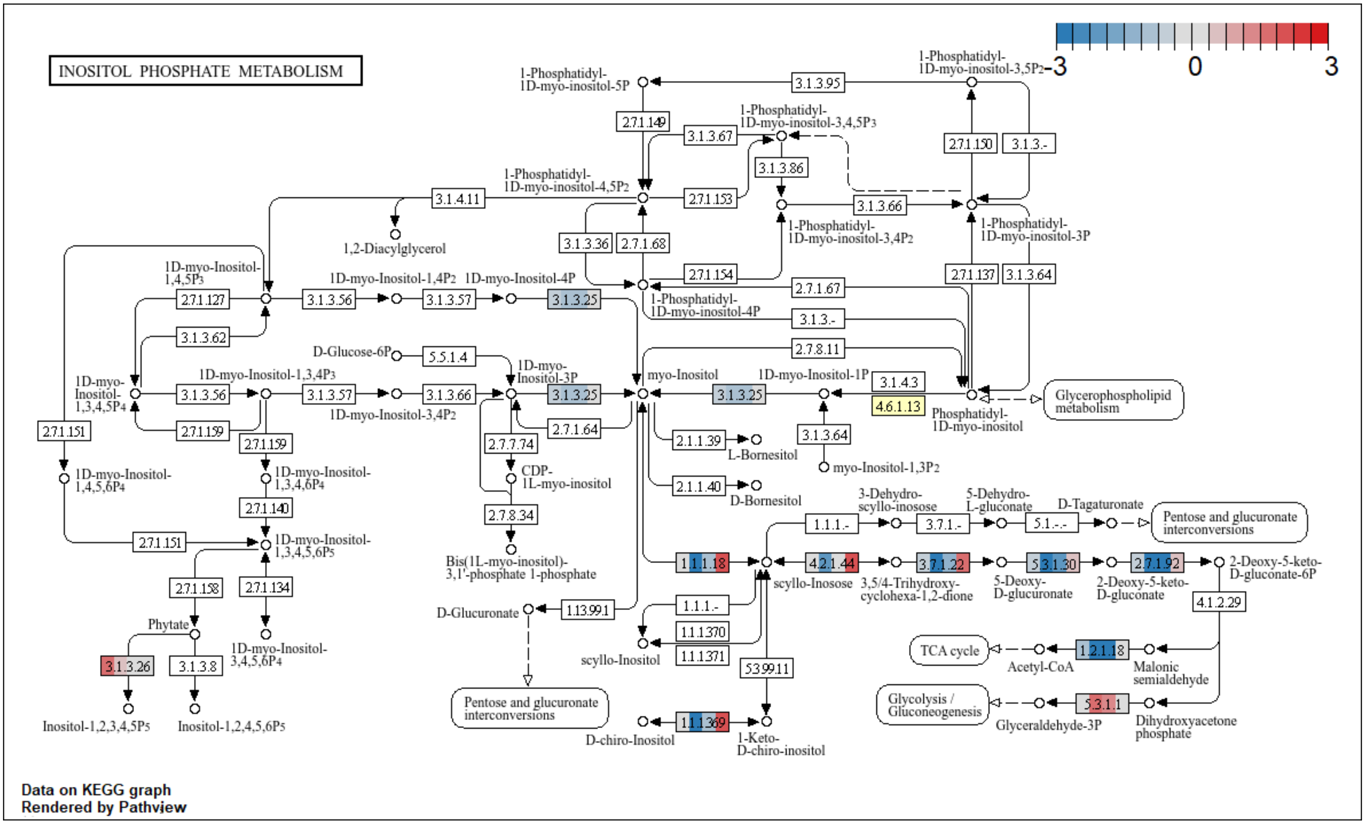
INOSITOL PHOSPHATE METABOLISM
-3
0
3
Data on KEGG graph
Rendered by Pathview

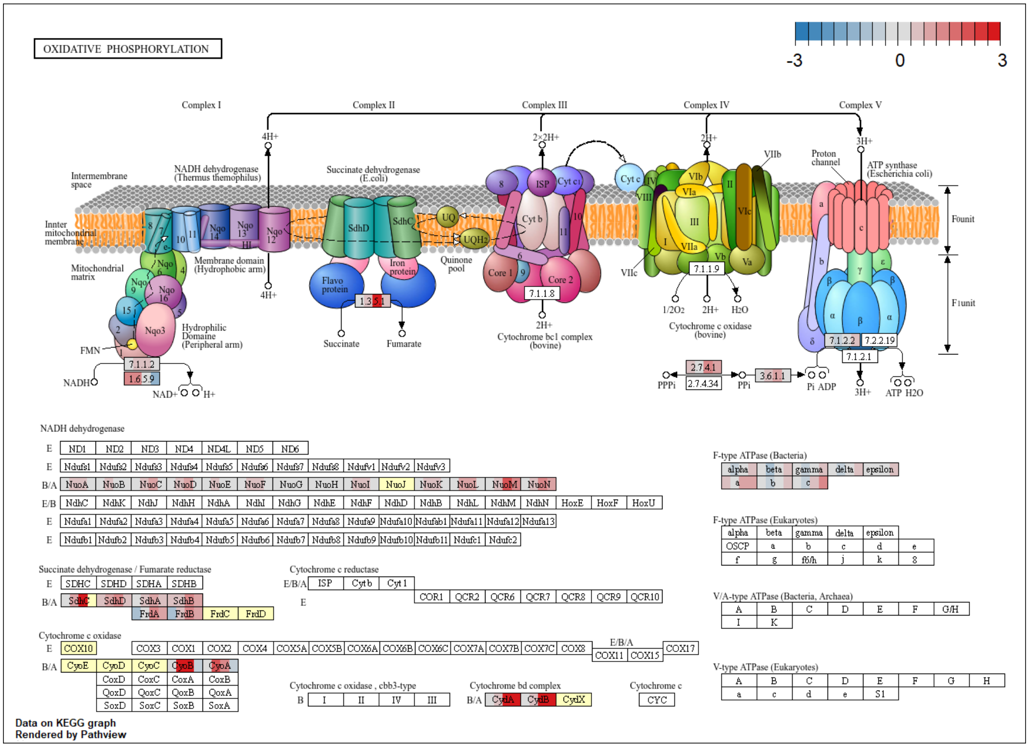
OXIDATIVE PHOSPHORYLATION
-3
0
3
Complex I
Complex II
Complex III
Complex IV
Complex V
NADH dehydrogenase (Thermus themophilus)
Succinate dehydrogenase (E.coli)
Intermembrane space
Innter mitochondrial membrane
Mitochondrial matrix
Membrane domain (Hydrophobic arm)
Hydrophilic Domaine (Peripheral arm)
Flavo protein
Iron protein
Quinone pool
UQ
UQH2
SdhC
SdhD
Cytochrome bc1 complex (bovine)
Cytochrome c oxidase (bovine)
Proton channel
ATP synthase (Escherichia coli)
Founit
F1unit
FMN
NADH
NAD+
H+
Succinate
Fumarate
1/2O2
2H+
H2O
PPPi
PPi
Pi ADP
3H+
ATP H2O
7.1.1.2
1.6.5.9
1.3.5.1
7.1.1.8
7.1.1.9
2.7.4.1
2.7.4.34
3.6.1.1
7.1.2.2
7.2.2.19
7.1.2.1
NADH dehydrogenase
E ND1 ND2 ND3 ND4 ND4L ND5 ND6
E Ndufs1 Ndufs2 Ndufs3 Ndufs4 Ndufs5 Ndufs6 Ndufs7 Ndufs8 Ndufv1 Ndufv2 Ndufv3
B/A NuoA NuoB NuoC NuoD NuoE NuoF NuoG NuoH NuoI NuoJ NuoK NuoL NuoM NuoN
E/B NdhC NdhK NdhJ NdhH NdhA NdhI NdhG NdhE NdhF NdhD NdhB NdhL NdhM NdhN HoxE HoxF HoxU
E Ndufa1 Ndufa2 Ndufa3 Ndufa4 Ndufa5 Ndufa6 Ndufa7 Ndufa8 Ndufa9 Ndufa10 Ndufab1 Ndufa11 Ndufa12 Ndufa13
E Ndufb1 Ndufb2 Ndufb3 Ndufb4 Ndufb5 Ndufb6 Ndufb7 Ndufb8 Ndufb9 Ndufb10 Ndufb11 Ndufc1 Ndufc2
Succinate dehydrogenase / Fumarate reductase
E SDHC SDHD SDHA SDHB
B/A SdhC SdhD SdhA SdhB
FrdA FrdB FrdC FrdD
Cytochrome c reductase
E/B/A ISP Cyt b Cyt 1
E COR1 QCR2 QCR6 QCR7 QCR8 QCR9 QCR10
Cytochrome c oxidase
E COX10 COX3 COX1 COX2 COX4 COX5A COX5B COX6A COX6B COX6C COX7A COX7B COX7C COX8
E/B/A COX11 COX15 COX17
B/A CyoE CyoD CyoC CyoB CyoA
CoxD CoxC CoxA CoxB
QoxD QoxC QoxB QoxA
SoxD SoxC SoxB SoxA
Cytochrome c oxidase , cbb3-type
B I II IV III
Cytochrome bd complex
B/A CydA CydB CydX
Cytochrome c
CYC
F-type ATPase (Bacteria)
alpha beta gamma delta epsilon
a b c
F-type ATPase (Eukaryotes)
alpha beta gamma delta epsilon
OSCP a b c d e
f g f6/h j k 8
V/A-type ATPase (Bacteria, Archaea)
A B C D E F G/H
I K
V-type ATPase (Eukaryotes)
A B C D E F G H
a c d e S1
Data on KEGG graph
Rendered by Pathview

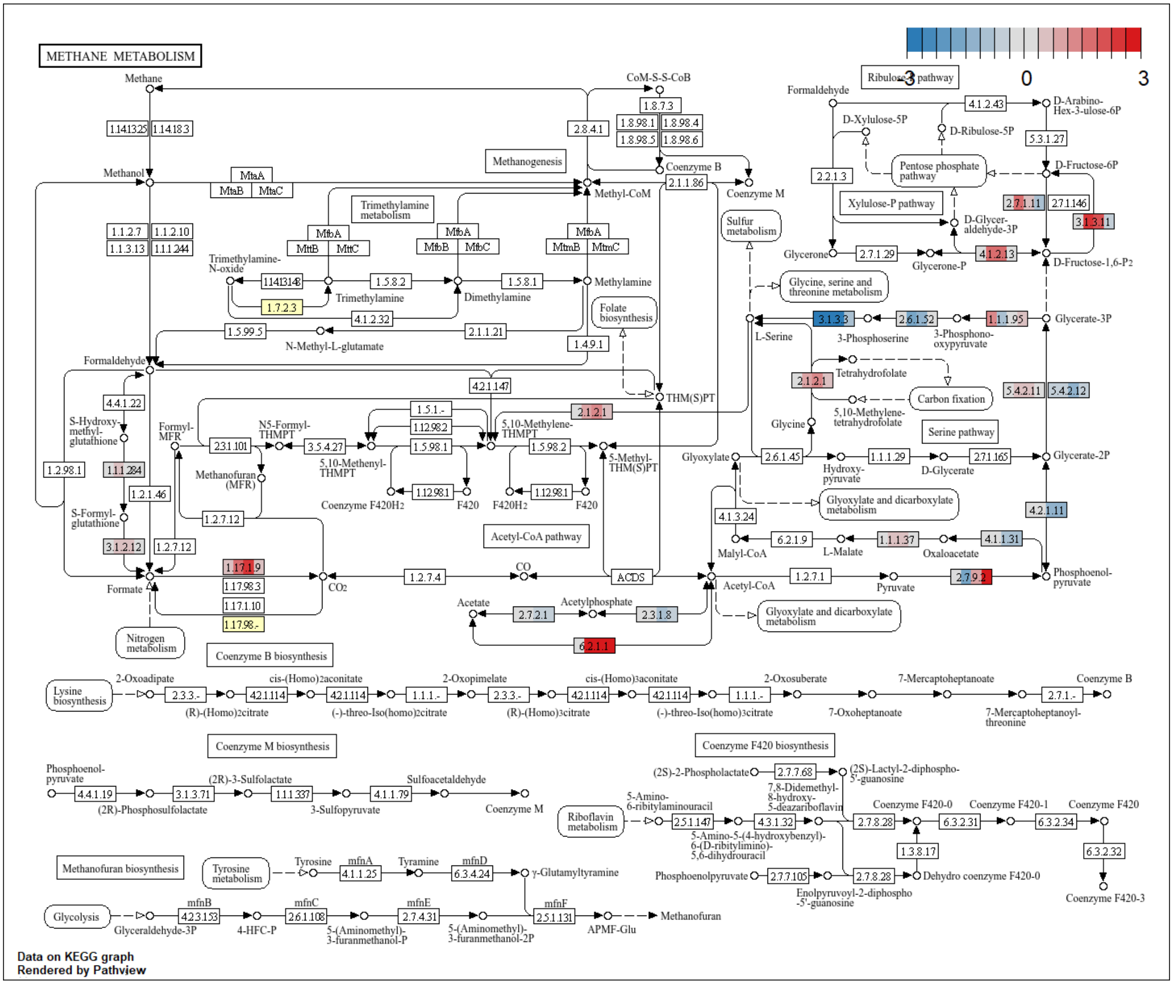
METHANE METABOLISM
-3
0
3
Methane
CoM-S-S-CoB
Formaldehyde
Ribulose-P pathway
D-Arabino-Hex-3-ulose-6P
D-Xylulose-5P
D-Ribulose-5P
Pentose phosphate pathway
D-Fructose-6P
Xylulose-P pathway
D-Glycer-aldehyde-3P
Glycerone
Glycerone-P
D-Fructose-1,6-P2
Methanogenesis
Methanol
Coenzyme B
Coenzyme M
Methyl-CoM
Trimethylamine metabolism
Sulfur metabolism
Trimethylamine-N-oxide
Trimethylamine
Dimethylamine
Methylamine
Folate biosynthesis
Glycine, serine and threonine metabolism
N-Methyl-L-glutamate
L-Serine
3-Phosphoserine
3-Phospho-oxypyruvate
Glycerate-3P
Tetrahydrofolate
Carbon fixation
5,10-Methylene-tetrahydrofolate
Glycine
Serine pathway
S-Hydroxy-methyl-glutathione
Formyl-MFR
N5-Formyl-THMPT
5,10-Methylene-THMPT
THM(S)PT
5,10-Methenyl-THMPT
Methanofuran (MFR)
Coenzyme F420H2
F420
F420H2
F420
5-Methyl-THM(S)PT
Glyoxylate
Hydroxy-pyruvate
D-Glycerate
Glycerate-2P
Glyoxylate and dicarboxylate metabolism
S-Formyl-glutathione
Acetyl-CoA pathway
Malyl-CoA
L-Malate
Oxaloacetate
Phosphoenol-pyruvate
Formate
CO2
CO
Acetyl-CoA
Pyruvate
Acetate
Acetylphosphate
Glyoxylate and dicarboxylate metabolism
Nitrogen metabolism
Coenzyme B biosynthesis
Lysine biosynthesis
2-Oxoadipate
(R)-(Homo)2citrate
cis-(Homo)2aconitate
(-)-threo-Iso(homo)2citrate
2-Oxopimelate
(R)-(Homo)3citrate
cis-(Homo)3aconitate
(-)-threo-Iso(homo)3citrate
2-Oxosuberate
7-Oxoheptanoate
7-Mercaptoheptanoate
7-Mercaptoheptanoyl-threonine
Coenzyme B
Coenzyme M biosynthesis
Phosphoenol-pyruvate
(2R)-Phosphosulfolactate
(2R)-3-Sulfolactate
3-Sulfopyruvate
Sulfoacetaldehyde
Coenzyme M
Coenzyme F420 biosynthesis
(2S)-2-Phospholactate
(2S)-Lactyl-2-diphospho-5'-guanosine
Riboflavin metabolism
5-Amino-6-ribitylaminouracil
7,8-Didemethyl-8-hydroxy-5-deazariboflavin
5-Amino-5-(4-hydroxybenzyl)-6-(D-ribitylimino)-5,6-dihydrouracil
Coenzyme F420-0
Coenzyme F420-1
Coenzyme F420
Phosphoenolpyruvate
Enolpyruvoyl-2-diphospho-5'-guanosine
Dehydro coenzyme F420-0
Coenzyme F420-3
Methanofuran biosynthesis
Tyrosine metabolism
Tyrosine
Tyramine
γ-Glutamyltyramine
Glycolysis
Glyceraldehyde-3P
4-HFC-P
5-(Aminomethyl)-3-furanmethanol-P
5-(Aminomethyl)-3-furanmethanol-2P
APMF-Glu
Methanofuran
Data on KEGG graph
Rendered by Pathview

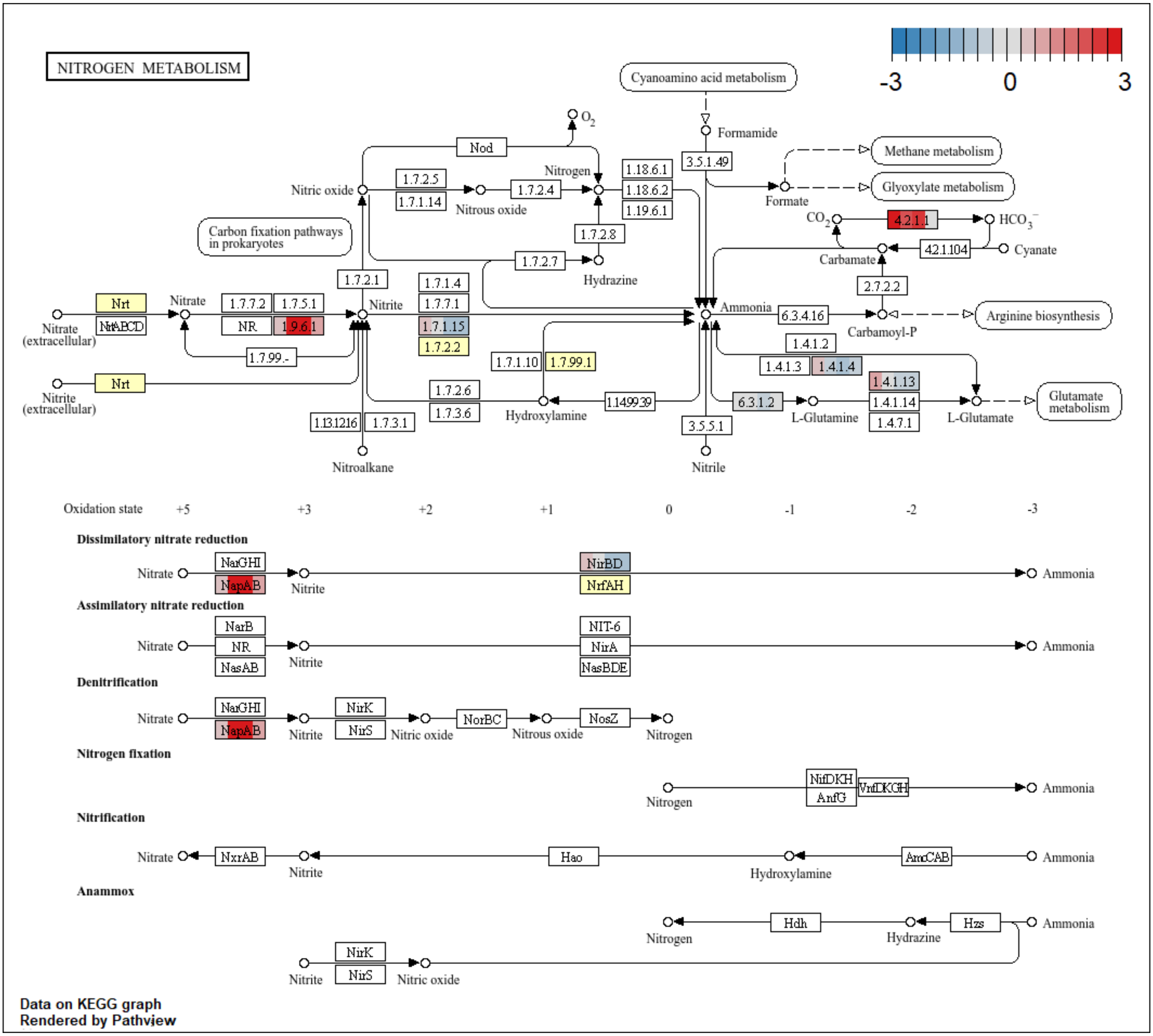

NITROGEN METABOLISM
-3
0
3
Cyanoamino acid metabolism
Formamide
Methane metabolism
Glyoxylate metabolism
Formate
Nod
Nitrogen
Nitric oxide
Nitrous oxide
Carbon fixation pathways in prokaryotes
Hydrazine
Carbamate
Cyanate
Nitrate
Nitrite
Ammonia
Carbamoyl-P
Arginine biosynthesis
Nitrate (extracellular)
Nitrite (extracellular)
Nrt
NrtABCD
NR
Hydroxylamine
L-Glutamine
L-Glutamate
Glutamate metabolism
Nitroalkane
Nitrile
Oxidation state +5 +3 +2 +1 0 -1 -2 -3
Dissimilatory nitrate reduction
NarGHI
NapAB
NirBD
NrfAH
Assimilatory nitrate reduction
NarB
NasAB
NIT-6
NirA
NasBDE
Denitrification
NirK
NirS
NorBC
NosZ
Nitrogen fixation
NifDKH
AnfG
VnfDKGH
Nitrification
NxrAB
Hao
AmoCAB
Anammox
Hdh
Hzs
Data on KEGG graph
Rendered by Pathview

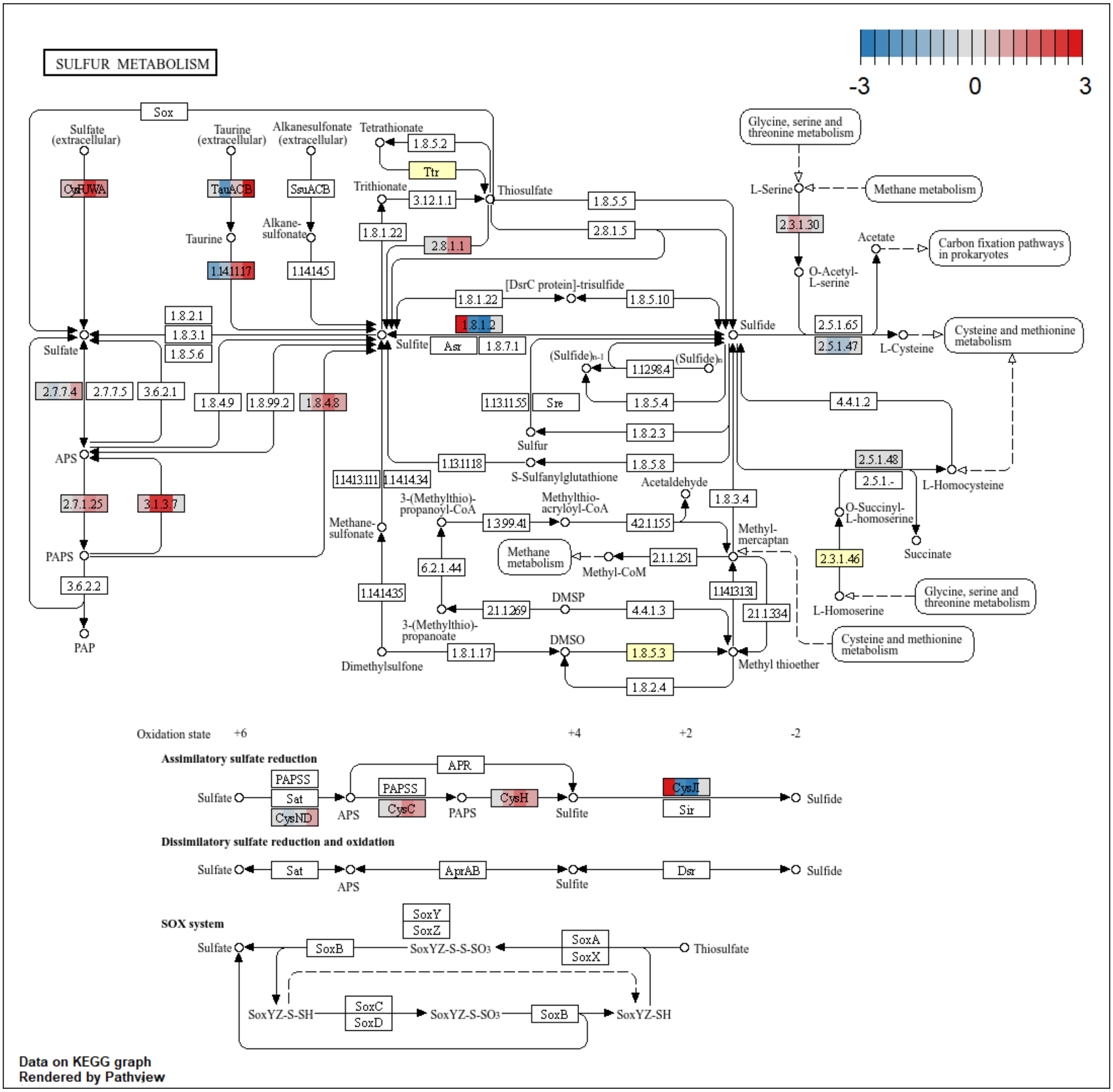
SULFUR METABOLISM
-3
0
3
Sox
Sulfate (extracellular)
Taurine (extracellular)
Alkanesulfonate (extracellular)
Tetrathionate
1.8.5.2
Ttr
Thiosulfate
Trithionate
3.12.1.1
1.8.5.5
2.8.1.5
CysPUWA
TauACB
SsuACB
Taurine
Alkane-sulfonate
1.8.1.22
2.8.1.1
1.14.11.17
1.14.14.5
[DsrC protein]-trisulfide
1.8.1.22
1.8.5.10
1.8.1.2
Sulfide
1.8.2.1
1.8.3.1
1.8.5.6
Sulfate
Sulfite
Asr
1.8.7.1
(Sulfide)n-1
(Sulfide)n
1.12.98.4
2.7.7.4
2.7.7.5
3.6.2.1
1.8.4.9
1.8.99.2
1.8.4.8
1.13.11.55
Sre
1.8.5.4
Sulfur
1.8.2.3
APS
1.13.11.18
1.8.5.8
S-Sulfanylglutathione
1.14.13.111
1.14.14.34
2.7.1.25
3.1.3.7
3-(Methylthio)-propanoyl-CoA
Methylthio-acryloyl-CoA
Acetaldehyde
1.8.3.4
Methane-sulfonate
1.3.99.41
4.2.1.155
Methyl-mercaptan
PAPS
Methane metabolism
2.1.1.251
Methyl-CoM
6.2.1.44
1.14.14.35
1.14.13.131
3.6.2.2
DMSP
2.1.1.269
4.4.1.3
2.1.1.334
3-(Methylthio)-propanoate
DMSO
1.8.1.17
1.8.5.3
Dimethylsulfone
Methyl thioether
1.8.2.4
PAP
Glycine, serine and threonine metabolism
L-Serine
Methane metabolism
2.3.1.30
Acetate
Carbon fixation pathways in prokaryotes
O-Acetyl-L-serine
2.5.1.65
2.5.1.47
L-Cysteine
Cysteine and methionine metabolism
4.4.1.2
2.5.1.48
2.5.1.-
L-Homocysteine
O-Succinyl-L-homoserine
2.3.1.46
Succinate
L-Homoserine
Glycine, serine and threonine metabolism
Cysteine and methionine metabolism
Oxidation state
+6
+4
+2
-2
Assimilatory sulfate reduction
APR
PAPSS
Sat
CysND
APS
PAPSS
CysC
PAPS
CysH
Sulfite
CysJI
Sir
Sulfide
Dissimilatory sulfate reduction and oxidation
Sulfate
Sat
APS
AprAB
Sulfite
Dsr
Sulfide
SOX system
SoxY
SoxZ
SoxA
SoxX
Sulfate
SoxB
SoxYZ-S-S-SO3
Thiosulfate
SoxYZ-S-SH
SoxC
SoxD
SoxYZ-S-SO3
SoxB
SoxYZ-SH
Data on KEGG graph
Rendered by Pathview

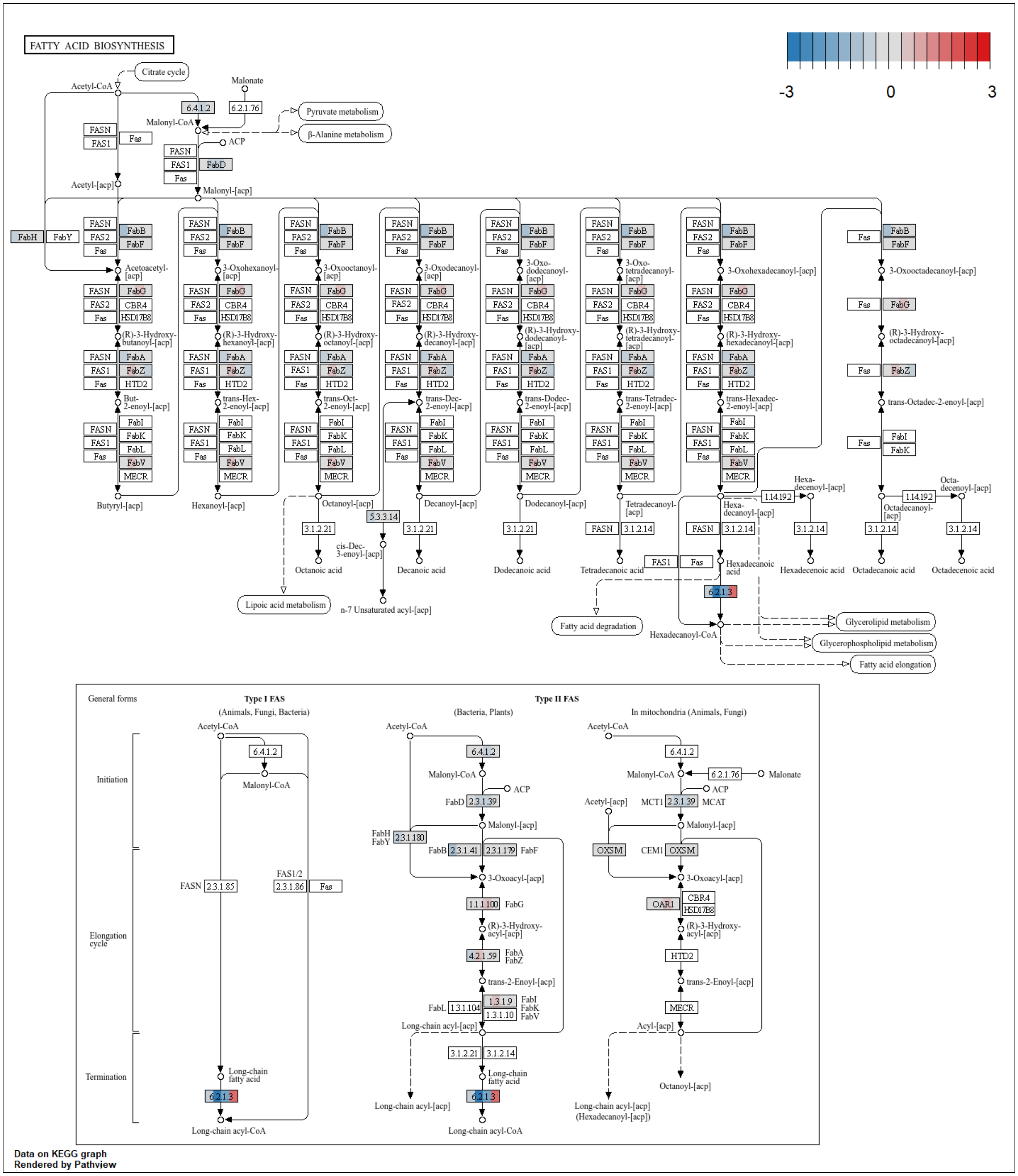
FATTY ACID BIOSYNTHESIS
-3
0
3
Citrate cycle
Acetyl-CoA
Malonate
Malonyl-CoA
Pyruvate metabolism
β-Alanine metabolism
Acetyl-[acp]
Malonyl-[acp]
Butyryl-[acp]
Hexanoyl-[acp]
Octanoyl-[acp]
Decanoyl-[acp]
Dodecanoyl-[acp]
Tetradecanoyl-[acp]
Octanoic acid
Decanoic acid
Dodecanoic acid
Tetradecanoic acid
Hexadecanoic acid
Hexadecenoic acid
Octadecanoic acid
Octadecenoic acid
Lipoic acid metabolism
n-7 Unsaturated acyl-[acp]
Fatty acid degradation
Hexadecanoyl-CoA
Glycerolipid metabolism
Glycerophospholipid metabolism
Fatty acid elongation
General forms
Type I FAS
Type II FAS
(Animals, Fungi, Bacteria)
(Bacteria, Plants)
In mitochondria (Animals, Fungi)
Initiation
Elongation cycle
Termination
Long-chain fatty acid
Long-chain acyl-CoA
Long-chain acyl-[acp]
Data on KEGG graph
Rendered by Pathview

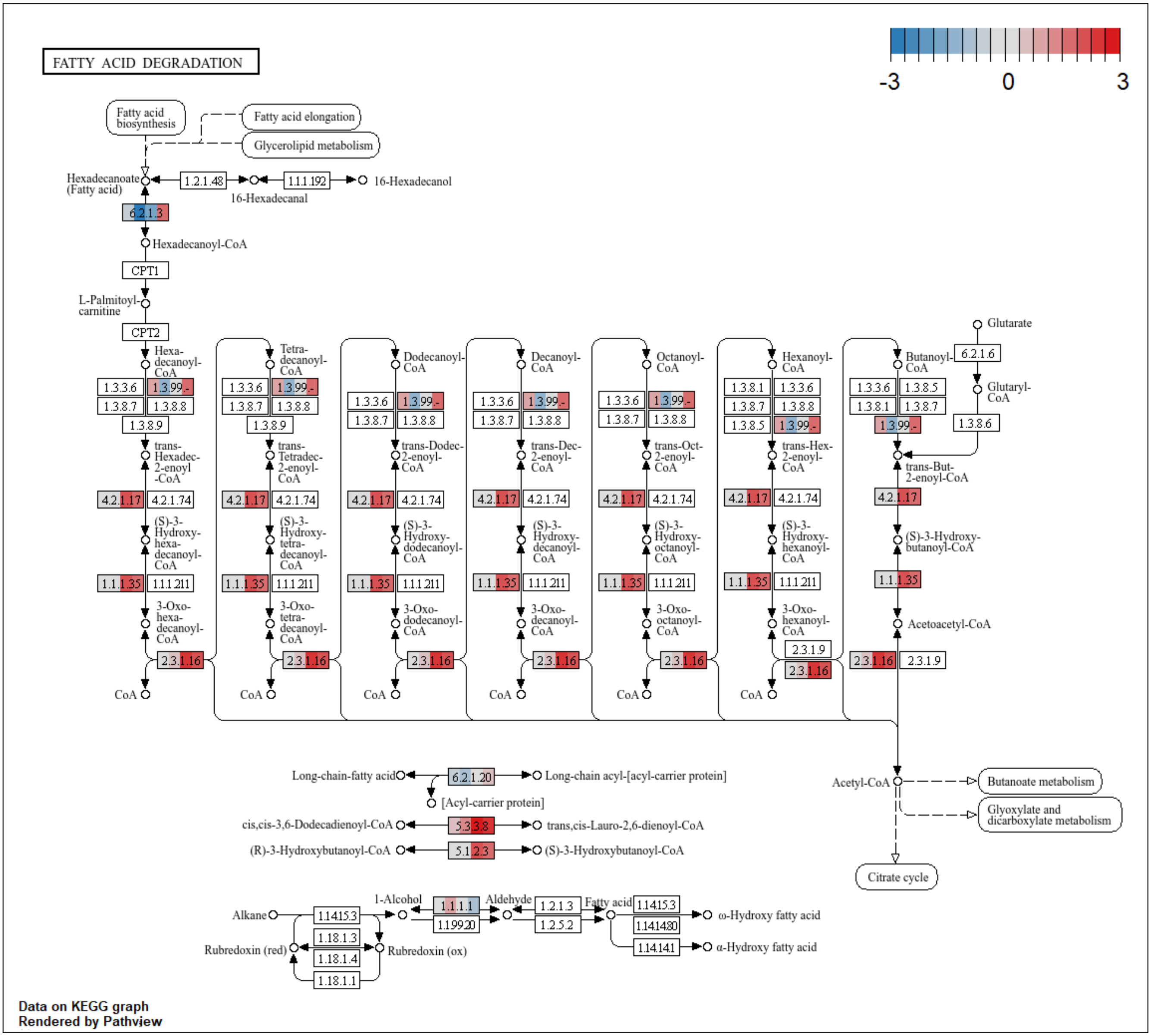
FATTY ACID DEGRADATION
-3
0
3
Fatty acid biosynthesis
Fatty acid elongation
Glycerolipid metabolism
Hexadecanoate (Fatty acid)
16-Hexadecanal
16-Hexadecanol
Hexadecanoyl-CoA
CPT1
L-Palmitoyl-carnitine
CPT2
Glutarate
Glutaryl-CoA
Acetoacetyl-CoA
Long-chain-fatty acid
Long-chain acyl-[acyl-carrier protein]
[Acyl-carrier protein]
cis,cis-3,6-Dodecadienoyl-CoA
trans,cis-Lauro-2,6-dienoyl-CoA
(R)-3-Hydroxybutanoyl-CoA
(S)-3-Hydroxybutanoyl-CoA
Acetyl-CoA
Butanoate metabolism
Glyoxylate and dicarboxylate metabolism
Citrate cycle
Alkane
1-Alcohol
Aldehyde
Fatty acid
ω-Hydroxy fatty acid
α-Hydroxy fatty acid
Rubredoxin (red)
Rubredoxin (ox)
Data on KEGG graph
Rendered by Pathview

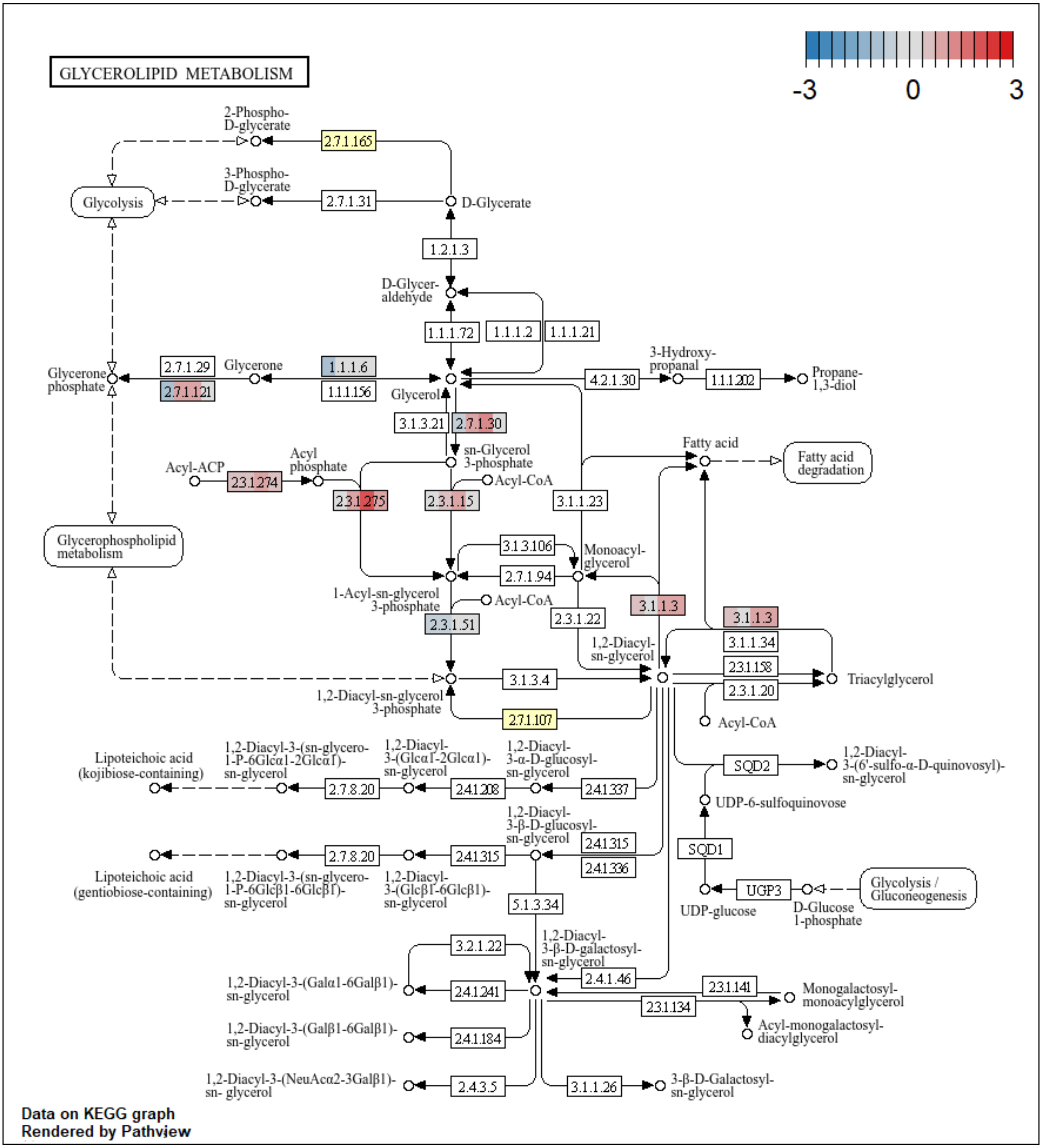
GLYCEROLIPID METABOLISM
-3
0
3
2-Phospho-D-glycerate
2.7.1.165
3-Phospho-D-glycerate
2.7.1.31
Glycolysis
D-Glycerate
1.2.1.3
D-Glycer-aldehyde
1.1.1.72
1.1.1.2
1.1.1.21
Glycerone phosphate
2.7.1.29
2.7.1.121
Glycerone
1.1.1.6
1.1.1.156
Glycerol
4.2.1.30
3-Hydroxy-propanal
1.1.1.202
Propane-1,3-diol
3.1.3.21
2.7.1.30
sn-Glycerol 3-phosphate
Acyl-ACP
2.3.1.274
Acyl phosphate
2.3.1.275
2.3.1.15
Acyl-CoA
3.1.1.23
Fatty acid
Fatty acid degradation
Glycerophospholipid metabolism
3.1.3.106
Monoacyl-glycerol
2.7.1.94
1-Acyl-sn-glycerol 3-phosphate
Acyl-CoA
2.3.1.51
2.3.1.22
3.1.1.3
3.1.1.3
3.1.1.34
1,2-Diacyl-sn-glycerol
2.3.1.158
2.3.1.20
Triacylglycerol
3.1.3.4
1,2-Diacyl-sn-glycerol 3-phosphate
2.7.1.107
Acyl-CoA
Lipoteichoic acid (kojibiose-containing)
1,2-Diacyl-3-(sn-glycero-1-P-6Glcα1-2Glcα1)-sn-glycerol
2.7.8.20
1,2-Diacyl-3-(Glcα1-2Glcα1)-sn-glycerol
2.4.1.208
1,2-Diacyl-3-α-D-glucosyl-sn-glycerol
2.4.1.337
SQD2
1,2-Diacyl-3-(6'-sulfo-α-D-quinovosyl)-sn-glycerol
UDP-6-sulfoquinovose
1,2-Diacyl-3-β-D-glucosyl-sn-glycerol
2.4.1.315
2.4.1.336
SQD1
2.7.8.20
2.4.1.315
Lipoteichoic acid (gentiobiose-containing)
1,2-Diacyl-3-(sn-glycero-1-P-6Glcβ1-6Glcβ1)-sn-glycerol
1,2-Diacyl-3-(Glcβ1-6Glcβ1)-sn-glycerol
5.1.3.34
UGP3
UDP-glucose
D-Glucose 1-phosphate
Glycolysis / Gluconeogenesis
3.2.1.22
1,2-Diacyl-3-β-D-galactosyl-sn-glycerol
2.4.1.46
1,2-Diacyl-3-(Galα1-6Galβ1)-sn-glycerol
2.4.1.241
2.3.1.141
2.3.1.134
Monogalactosyl-monoacylglycerol
Acyl-monogalactosyl-diacylglycerol
1,2-Diacyl-3-(Galβ1-6Galβ1)-sn-glycerol
2.4.1.184
1,2-Diacyl-3-(NeuAcα2-3Galβ1)-sn- glycerol
2.4.3.5
3.1.1.26
3-β-D-Galactosyl-sn-glycerol
Data on KEGG graph
Rendered by Pathview

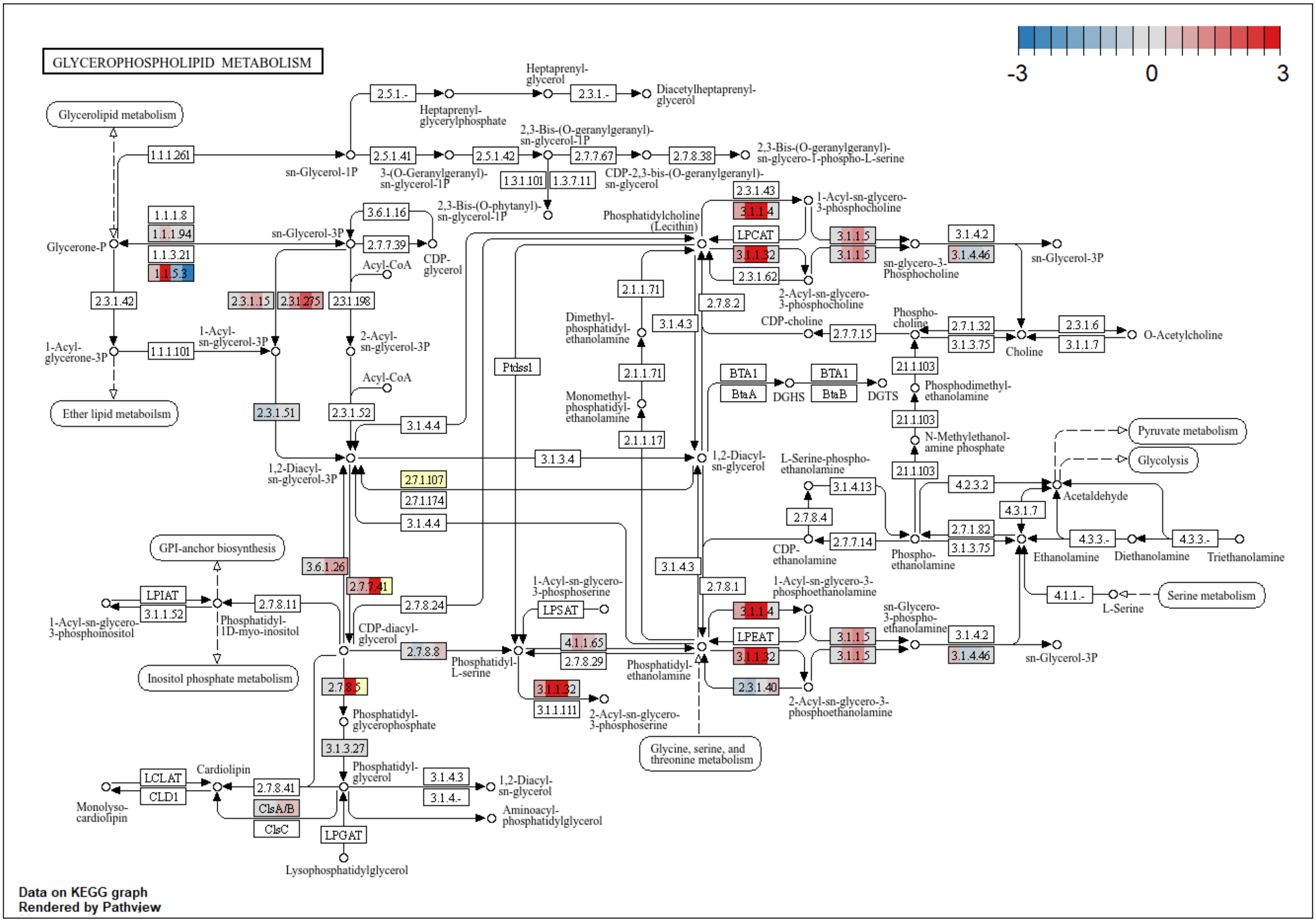
GLYCEROPHOSPHOLIPID METABOLISM
-3
0
3
Glycerolipid metabolism
Glycerone-P
sn-Glycerol-1P
sn-Glycerol-3P
1-Acyl-glycerone-3P
1-Acyl-sn-glycerol-3P
2-Acyl-sn-glycerol-3P
1,2-Diacyl-sn-glycerol-3P
Ether lipid metabolism
Phosphatidylcholine (Lecithin)
Phosphatidylethanolamine
Phosphatidyl-L-serine
CDP-diacylglycerol
Cardiolipin
Choline
Ethanolamine
Acetaldehyde
GPI-anchor biosynthesis
Inositol phosphate metabolism
Pyruvate metabolism
Glycolysis
Serine metabolism
Glycine, serine, and threonine metabolism
Lysophosphatidylglycerol
Data on KEGG graph
Rendered by Pathview

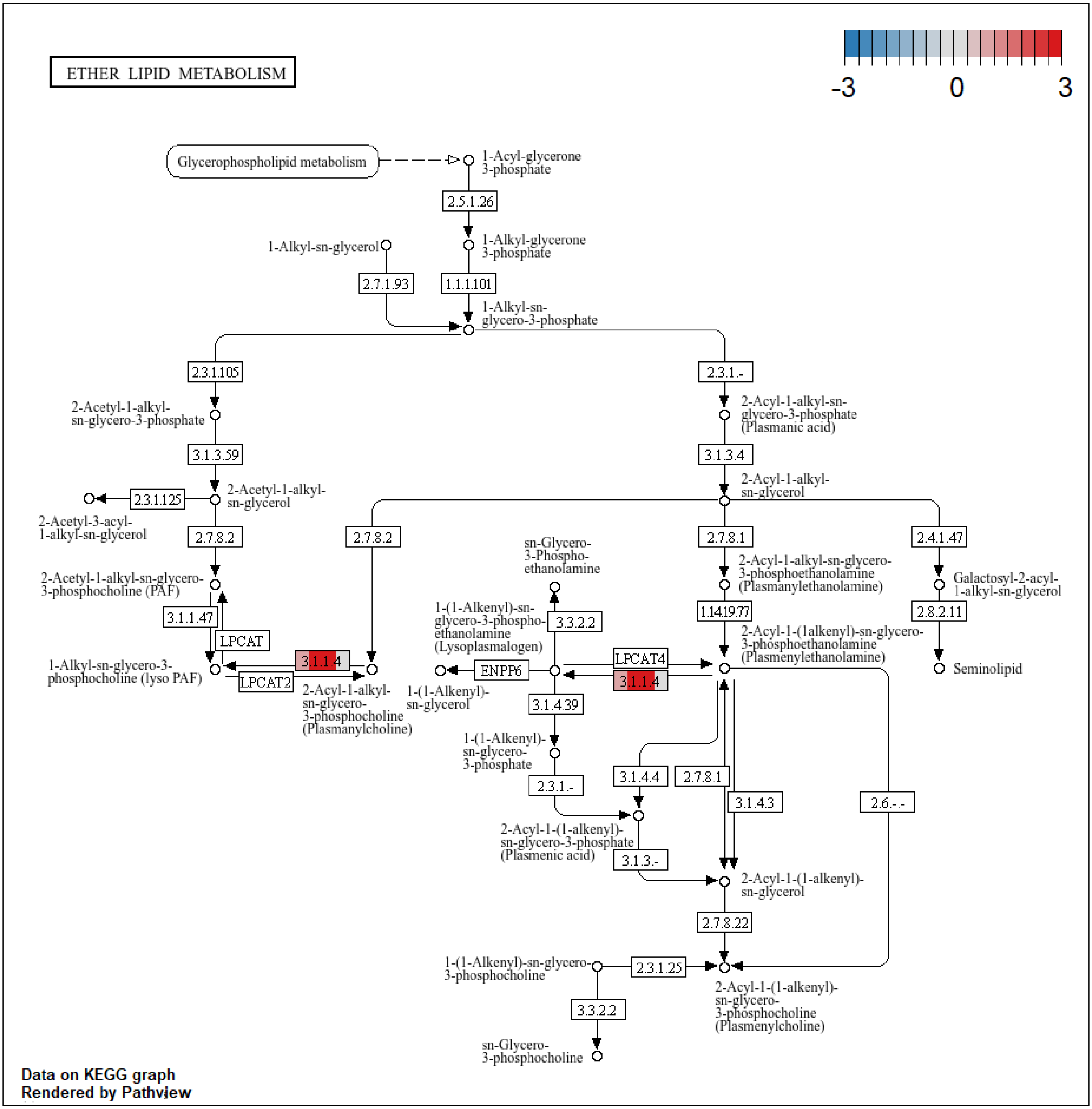
ETHER LIPID METABOLISM
-3
0
3
Glycerophospholipid metabolism
1-Acyl-glycerone 3-phosphate
2.5.1.26
1-Alkyl-sn-glycerol
1-Alkyl-glycerone 3-phosphate
2.7.1.93
1.1.1.101
1-Alkyl-sn-glycero-3-phosphate
2.3.1.105
2.3.1.-
2-Acetyl-1-alkyl-sn-glycero-3-phosphate
2-Acyl-1-alkyl-sn-glycero-3-phosphate (Plasmanic acid)
3.1.3.59
3.1.3.4
2.3.1.125
2-Acetyl-1-alkyl-sn-glycerol
2-Acyl-1-alkyl-sn-glycerol
2-Acetyl-3-acyl-1-alkyl-sn-glycerol
2.7.8.2
2.7.8.2
2.7.8.1
2.4.1.47
sn-Glycero-3-Phospho-ethanolamine
2-Acyl-1-alkyl-sn-glycero-3-phosphoethanolamine (Plasmanylethanolamine)
Galactosyl-2-acyl-1-alkyl-sn-glycerol
2-Acetyl-1-alkyl-sn-glycero-3-phosphocholine (PAF)
1-(1-Alkenyl)-sn-glycero-3-phospho-ethanolamine (Lysoplasmalogen)
3.3.2.2
1.14.19.77
2.8.2.11
3.1.1.47
LPCAT
2-Acyl-1-(1alkenyl)-sn-glycero-3-phosphoethanolamine (Plasmenylethanolamine)
1-Alkyl-sn-glycero-3-phosphocholine (lyso PAF)
3.1.1.4
LPCAT4
ENPP6
3.1.1.4
Seminolipid
LPCAT2
2-Acyl-1-alkyl-sn-glycero-3-phosphocholine (Plasmanylcholine)
1-(1-Alkenyl)-sn-glycerol
3.1.4.39
1-(1-Alkenyl)-sn-glycero-3-phosphate
3.1.4.4
2.7.8.1
2.3.1.-
3.1.4.3
2.6.-.-
2-Acyl-1-(1-alkenyl)-sn-glycero-3-phosphate (Plasmenic acid)
3.1.3.-
2-Acyl-1-(1-alkenyl)-sn-glycerol
2.7.8.22
1-(1-Alkenyl)-sn-glycero-3-phosphocholine
2.3.1.25
2-Acyl-1-(1-alkenyl)-sn-glycero-3-phosphocholine (Plasmenylcholine)
3.3.2.2
sn-Glycero-3-phosphocholine
Data on KEGG graph
Rendered by Pathview

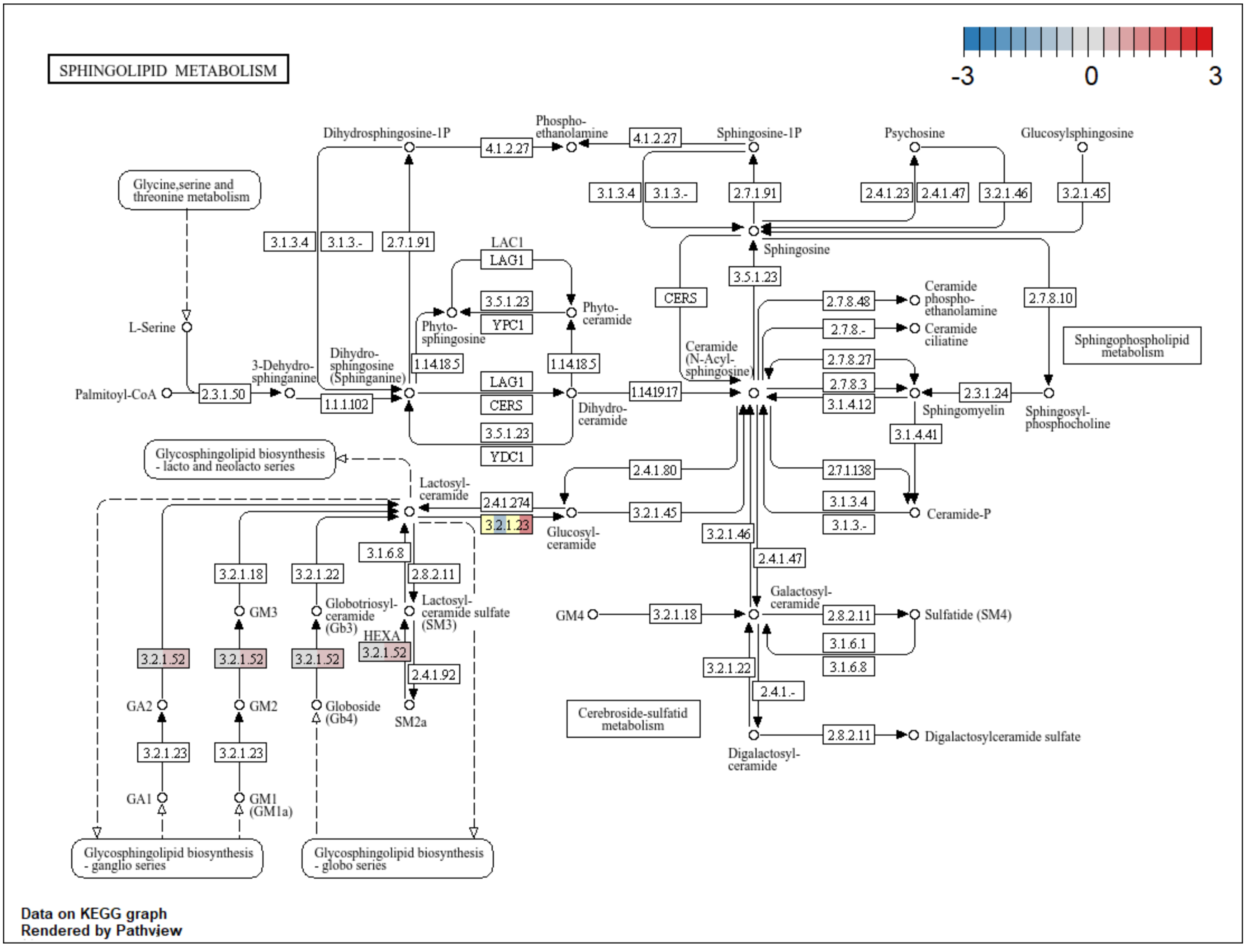
SPHINGOLIPID METABOLISM
-3
0
3
Dihydrosphingosine-1P
Phospho-ethanolamine
Sphingosine-1P
Psychosine
Glucosylsphingosine
Glycine,serine and threonine metabolism
L-Serine
Palmitoyl-CoA
3-Dehydro-sphinganine
Dihydro-sphingosine (Sphinganine)
Phyto-sphingosine
Phyto-ceramide
Dihydro-ceramide
Sphingosine
Ceramide (N-Acyl-sphingosine)
Ceramide phospho-ethanolamine
Ceramide ciliatine
Sphingomyelin
Sphingosyl-phosphocholine
Sphingophospholipid metabolism
Ceramide-P
Glycosphingolipid biosynthesis - lacto and neolacto series
Lactosyl-ceramide
Glucosyl-ceramide
Galactosyl-ceramide
GM4
Sulfatide (SM4)
Digalactosyl-ceramide
Digalactosylceramide sulfate
Cerebroside-sulfatid metabolism
GM3
GM2
GM1 (GM1a)
GA2
GA1
Globotriosyl-ceramide (Gb3)
Globoside (Gb4)
Lactosyl-ceramide sulfate (SM3)
SM2a
HEXA
LAC1
LAG1
YPC1
CERS
YDC1
Glycosphingolipid biosynthesis - ganglio series
Glycosphingolipid biosynthesis - globo series
Data on KEGG graph
Rendered by Pathview

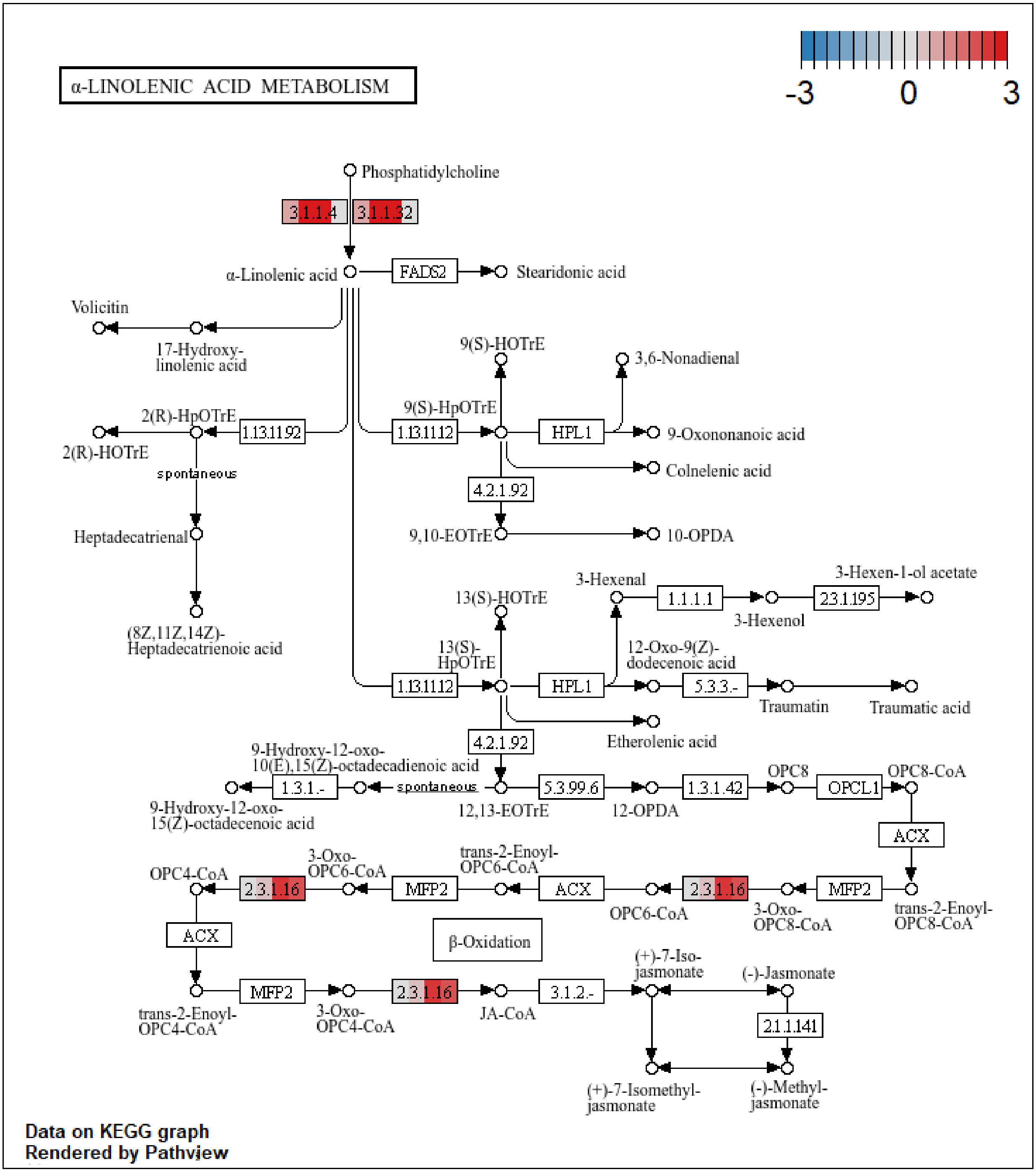

α-LINOLENIC ACID METABOLISM
-3
0
3
Phosphatidylcholine
3.1.1.4
3.1.1.32
α-Linolenic acid
FADS2
Stearidonic acid
Volicitin
17-Hydroxy-linolenic acid
9(S)-HOTrE
3,6-Nonadienal
2(R)-HpOTrE
1.13.11.92
9(S)-HpOTrE
1.13.11.12
HPL1
9-Oxononanoic acid
2(R)-HOTrE
spontaneous
Colnelenic acid
4.2.1.92
Heptadecatrienal
9,10-EOTrE
10-OPDA
3-Hexenal
1.1.1.1
3-Hexen-1-ol acetate
2.3.1.195
13(S)-HOTrE
3-Hexenol
(8Z,11Z,14Z)-Heptadecatrienoic acid
12-Oxo-9(Z)-dodecenoic acid
13(S)-HpOTrE
1.13.11.12
HPL1
5.3.3.-
Traumatin
Traumatic acid
Etherolenic acid
4.2.1.92
9-Hydroxy-12-oxo-10(E),15(Z)-octadecadienoic acid
1.3.1.-
spontaneous
5.3.99.6
1.3.1.42
OPC8
OPCL1
OPC8-CoA
9-Hydroxy-12-oxo-15(Z)-octadecenoic acid
12,13-EOTrE
12-OPDA
ACX
OPC4-CoA
3-Oxo-OPC6-CoA
trans-2-Enoyl-OPC6-CoA
2.3.1.16
MFP2
ACX
2.3.1.16
MFP2
OPC6-CoA
3-Oxo-OPC8-CoA
trans-2-Enoyl-OPC8-CoA
ACX
β-Oxidation
(+)-7-Iso-jasmonate
(-)-Jasmonate
MFP2
2.3.1.16
3.1.2.-
trans-2-Enoyl-OPC4-CoA
3-Oxo-OPC4-CoA
JA-CoA
2.1.1.141
(+)-7-Isomethyl-jasmonate
(-)-Methyl-jasmonate
Data on KEGG graph
Rendered by Pathview

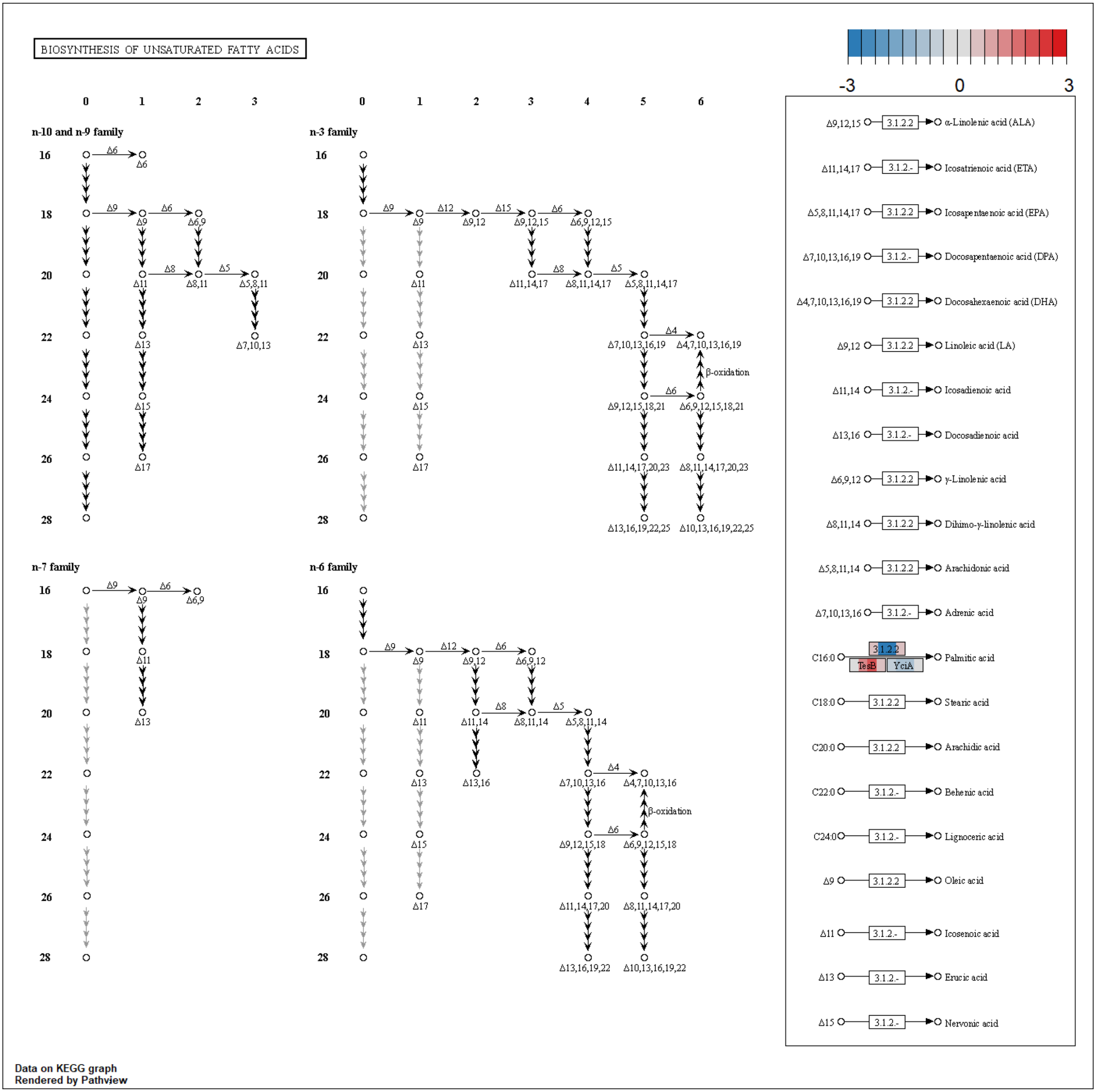
BIOSYNTHESIS OF UNSATURATED FATTY ACIDS
-3
0
3
n-10 and n-9 family
n-3 family
n-7 family
n-6 family
β-oxidation
Δ9,12,15 3.1.2.2 α-Linolenic acid (ALA)
Δ11,14,17 3.1.2.- Icosatrienoic acid (ETA)
Δ5,8,11,14,17 3.1.2.2 Icosapentaenoic acid (EPA)
Δ7,10,13,16,19 3.1.2.- Docosapentaenoic acid (DPA)
Δ4,7,10,13,16,19 3.1.2.2 Docosahexaenoic acid (DHA)
Δ9,12 3.1.2.2 Linoleic acid (LA)
Δ11,14 3.1.2.- Icosadienoic acid
Δ13,16 3.1.2.- Docosadienoic acid
Δ6,9,12 3.1.2.2 γ-Linolenic acid
Δ8,11,14 3.1.2.2 Dihimo-γ-linolenic acid
Δ5,8,11,14 3.1.2.2 Arachidonic acid
Δ7,10,13,16 3.1.2.- Adrenic acid
C16:0 3.1.2.2 TesB YciA Palmitic acid
C18:0 3.1.2.2 Stearic acid
C20:0 3.1.2.2 Arachidic acid
C22:0 3.1.2.- Behenic acid
C24:0 3.1.2.- Lignoceric acid
Δ9 3.1.2.2 Oleic acid
Δ11 3.1.2.- Icosenoic acid
Δ13 3.1.2.- Erucic acid
Δ15 3.1.2.- Nervonic acid
Data on KEGG graph
Rendered by Pathview

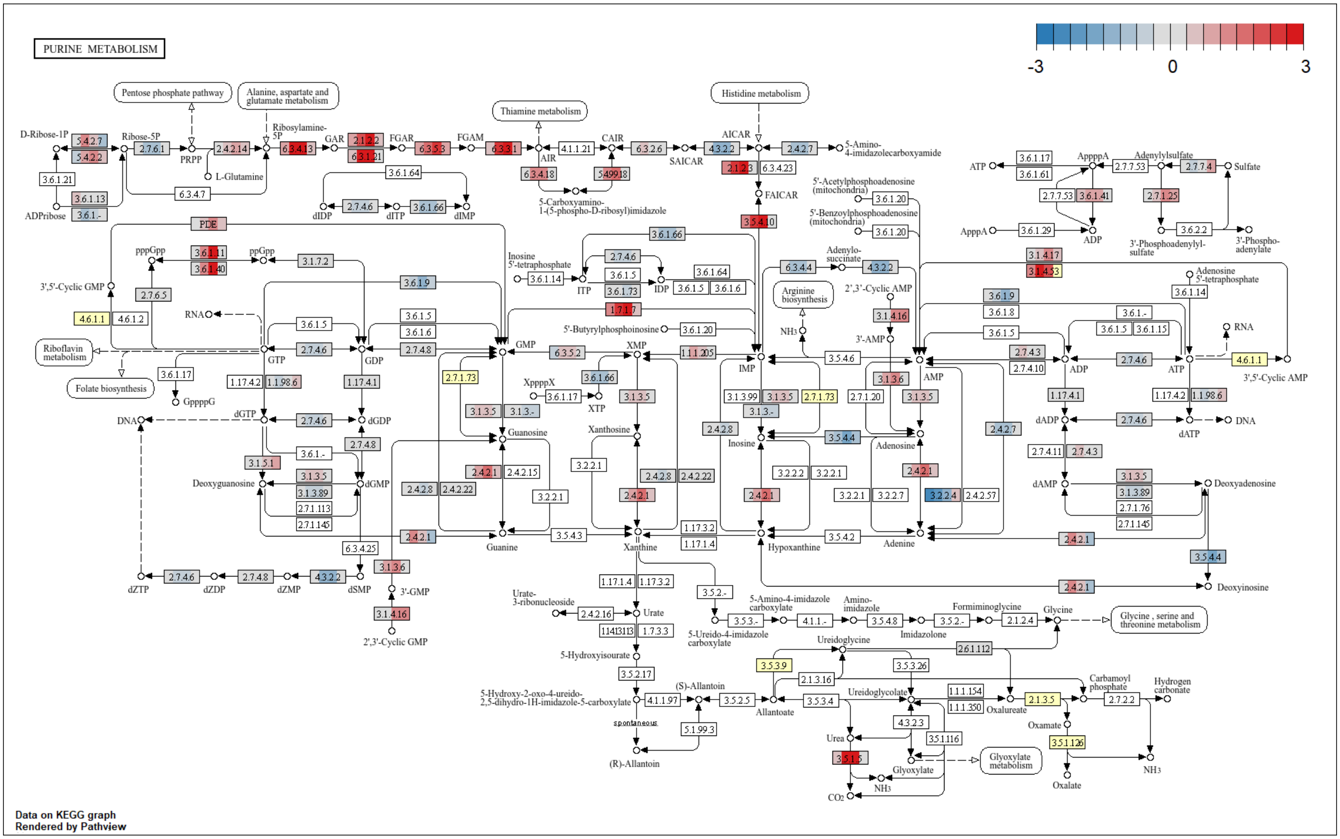
PURINE METABOLISM
-3
0
3
Pentose phosphate pathway
Alanine, aspartate and glutamate metabolism
Thiamine metabolism
Histidine metabolism
D-Ribose-1P
Ribose-5P
PRPP
Ribosylamine-5P
L-Glutamine
GAR
FGAR
FGAM
AIR
CAIR
SAICAR
AICAR
5-Amino-4-imidazolecarboxyamide
FAICAR
ADPRibose
5-Carboxyamino-1-(5-phospho-D-ribosyl)imidazole
dIDP
dITP
dIMP
PDE
pppGpp
ppGpp
3',5'-Cyclic GMP
RNA
Riboflavin metabolism
Folate biosynthesis
GTP
GDP
GMP
GppppG
DNA
dGTP
dGDP
dGMP
Deoxyguanosine
Guanosine
Guanine
Inosine 5'-tetraphosphate
ITP
IDP
IMP
XppppX
XTP
XMP
Xanthosine
Xanthine
5'-Butyrylphosphoinosine
Inosine
Hypoxanthine
Arginine biosynthesis
NH3
Adenylo-succinate
2',3'-Cyclic AMP
3'-AMP
AMP
Adenosine
Adenine
5'-Acetylphosphoadenosine (mitochondria)
5'-Benzoylphosphoadenosine (mitochondria)
ATP
ApppA
Adenylylsulfate
Sulfate
AppppA
ADP
3'-Phosphoadenylyl-sulfate
3'-Phospho-adenylate
Adenosine 5'-tetraphosphate
3',5'-Cyclic AMP
dADP
dATP
dAMP
Deoxyadenosine
Deoxyinosine
dZTP
dZDP
dZMP
dSMP
3'-GMP
2',3'-Cyclic GMP
Urate-3-ribonucleoside
Urate
5-Hydroxyisourate
5-Hydroxy-2-oxo-4-ureido-2,5-dihydro-1H-imidazole-5-carboxylate
(S)-Allantoin
(R)-Allantoin
spontaneous
5-Ureido-4-imidazole carboxylate
5-Amino-4-imidazole carboxylate
Amino-imidazole
Imidazolone
Formiminoglycine
Glycine
Glycine, serine and threonine metabolism
Ureidoglycine
Allantoate
Ureidoglycolate
Oxalureate
Carbamoyl phosphate
Hydrogen carbonate
Oxamate
Oxalate
Urea
Glyoxylate
Glyoxylate metabolism
CO2
Data on KEGG graph
Rendered by Pathview

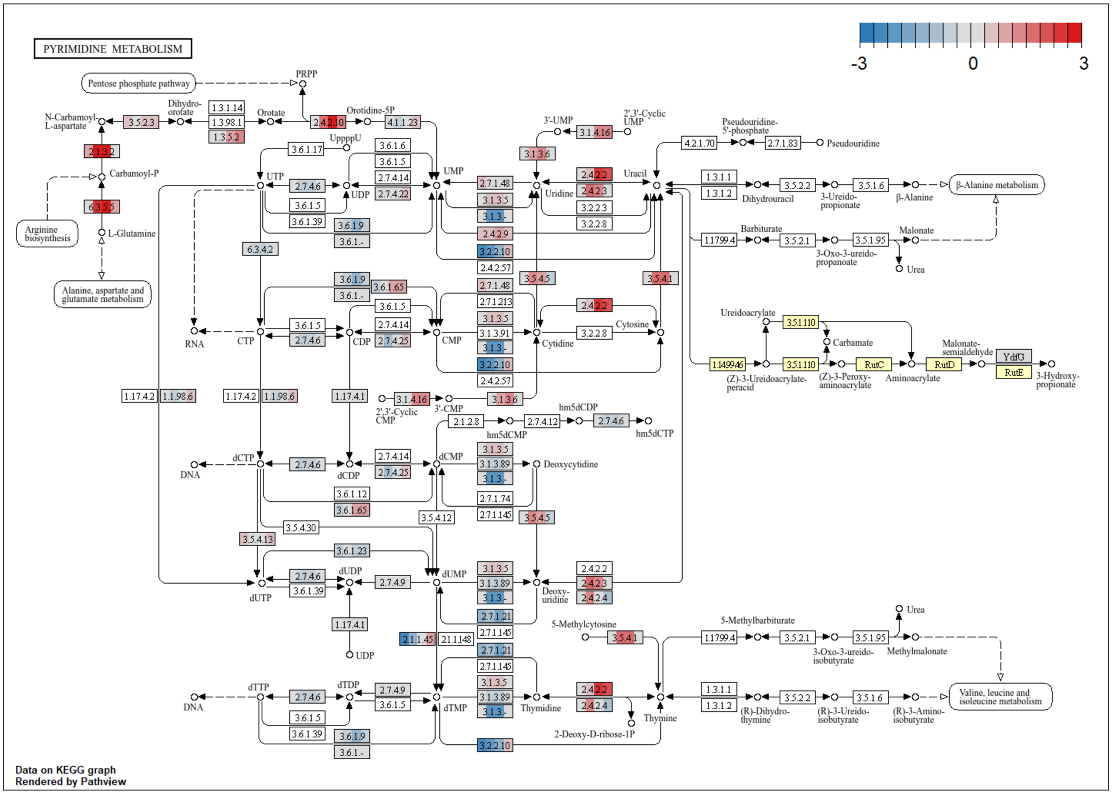

PYRIMIDINE METABOLISM
-3
0
3
Pentose phosphate pathway
PRPP
N-Carbamoyl-L-aspartate
Dihydro-orotate
Orotate
Orotidine-5P
UMP
UppppU
UTP
UDP
Carbamoyl-P
L-Glutamine
Arginine biosynthesis
Alanine, aspartate and glutamate metabolism
3'-UMP
2',3'-Cyclic UMP
Uridine
Uracil
Pseudouridine-5'-phosphate
Pseudouridine
Dihydrouracil
3-Ureido-propionate
β-Alanine
β-Alanine metabolism
Barbiturate
3-Oxo-3-ureido-propanoate
Malonate
Urea
RNA
CTP
CDP
CMP
Cytidine
Cytosine
Ureidoacrylate
Carbamate
Malonate-semialdehyde
(Z)-3-Ureidoacrylate-peracid
(Z)-3-Peroxy-aminoacrylate
Aminoacrylate
3-Hydroxy-propionate
RutC
RutD
RutE
YdfG
2',3'-Cyclic CMP
3'-CMP
hm5dCMP
hm5dCDP
hm5dCTP
DNA
dCTP
dCDP
dCMP
Deoxycytidine
dUTP
dUDP
dUMP
Deoxy-uridine
5-Methylcytosine
5-Methylbarbiturate
3-Oxo-3-ureido-isobutyrate
Methylmalonate
Valine, leucine and isoleucine metabolism
dTTP
dTDP
dTMP
Thymidine
Thymine
2-Deoxy-D-ribose-1P
(R)-Dihydro-thymine
(R)-3-Ureido-isobutyrate
(R)-3-Amino-isobutyrate
Data on KEGG graph
Rendered by Pathview

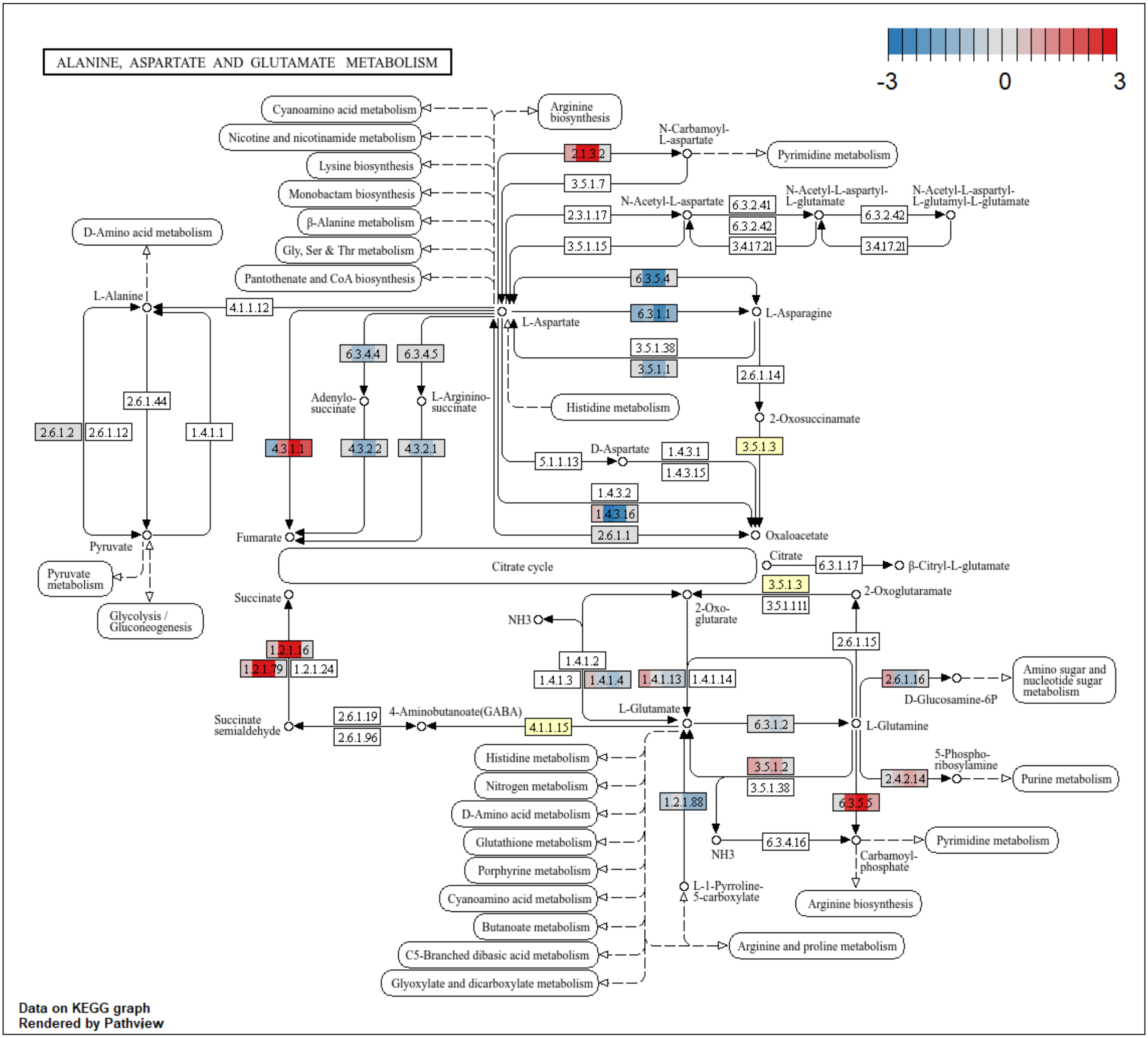

ALANINE, ASPARTATE AND GLUTAMATE METABOLISM
-3
0
3
Cyanoamino acid metabolism
Nicotine and nicotinamide metabolism
Lysine biosynthesis
Monobactam biosynthesis
β-Alanine metabolism
Gly, Ser & Thr metabolism
Pantothenate and CoA biosynthesis
Arginine biosynthesis
N-Carbamoyl-L-aspartate
Pyrimidine metabolism
2.1.3.2
3.5.1.7
2.3.1.17
3.5.1.15
N-Acetyl-L-aspartate
6.3.2.41
6.3.2.42
3.4.17.21
N-Acetyl-L-aspartyl-L-glutamate
N-Acetyl-L-aspartyl-L-glutamyl-L-glutamate
D-Amino acid metabolism
L-Alanine
4.1.1.12
6.3.5.4
6.3.1.1
L-Aspartate
L-Asparagine
3.5.1.38
3.5.1.1
2.6.1.14
6.3.4.4
6.3.4.5
Adenylo-succinate
L-Arginino-succinate
Histidine metabolism
2-Oxosuccinamate
2.6.1.44
2.6.1.2
2.6.1.12
1.4.1.1
4.3.1.1
4.3.2.2
4.3.2.1
3.5.1.3
D-Aspartate
5.1.1.13
1.4.3.1
1.4.3.15
1.4.3.2
1.4.3.16
2.6.1.1
Oxaloacetate
Pyruvate
Fumarate
Pyruvate metabolism
Glycolysis / Gluconeogenesis
Citrate cycle
Citrate
6.3.1.17
β-Citryl-L-glutamate
Succinate
3.5.1.3
3.5.1.111
2-Oxoglutaramate
NH3
2-Oxo-glutarate
2.6.1.15
1.2.1.16
1.2.1.79
1.2.1.24
1.4.1.2
1.4.1.3
1.4.1.4
1.4.1.13
1.4.1.14
2.6.1.16
Amino sugar and nucleotide sugar metabolism
D-Glucosamine-6P
Succinate semialdehyde
2.6.1.19
2.6.1.96
4-Aminobutanoate(GABA)
4.1.1.15
L-Glutamate
6.3.1.2
L-Glutamine
5-Phospho-ribosylamine
Histidine metabolism
Nitrogen metabolism
D-Amino acid metabolism
Glutathione metabolism
Porphyrine metabolism
Cyanoamino acid metabolism
Butanoate metabolism
C5-Branched dibasic acid metabolism
Glyoxylate and dicarboxylate metabolism
3.5.1.2
3.5.1.38
2.4.2.14
Purine metabolism
1.2.1.88
6.3.5.5
NH3
6.3.4.16
Pyrimidine metabolism
Carbamoyl-phosphate
L-1-Pyrroline-5-carboxylate
Arginine biosynthesis
Arginine and proline metabolism
Data on KEGG graph
Rendered by Pathview

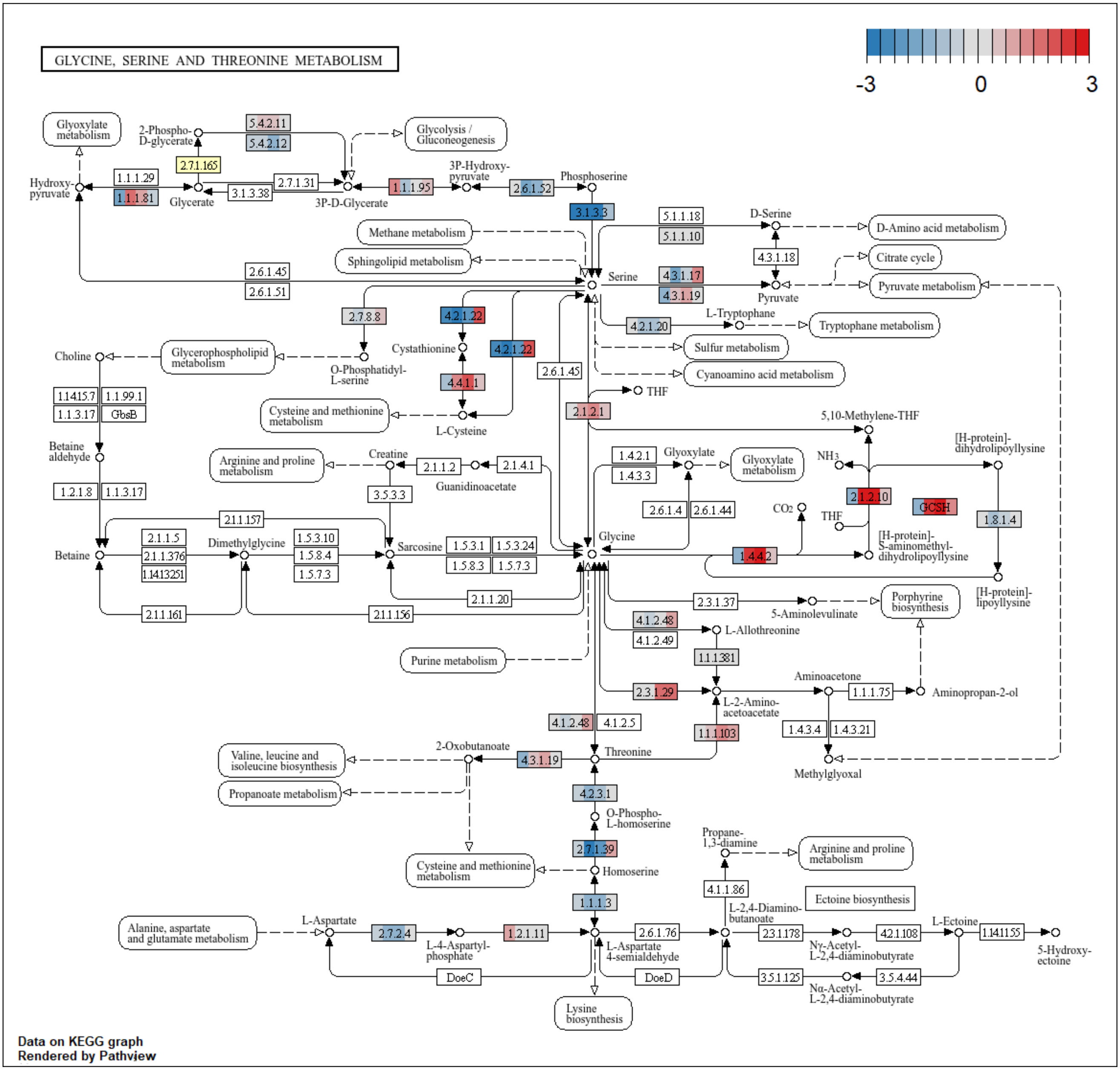
GLYCINE, SERINE AND THREONINE METABOLISM
-3
0
3
Glyoxylate metabolism
2-Phospho-D-glycerate
5.4.2.11
5.4.2.12
Glycolysis / Gluconeogenesis
2.7.1.165
Hydroxy-pyruvate
1.1.1.29
1.1.1.81
Glycerate
3.1.3.38
2.7.1.31
3P-D-Glycerate
1.1.1.95
3P-Hydroxy-pyruvate
2.6.1.52
Phosphoserine
3.1.3.3
5.1.1.18
5.1.1.10
D-Serine
D-Amino acid metabolism
Methane metabolism
Sphingolipid metabolism
4.3.1.18
Citrate cycle
2.6.1.45
2.6.1.51
Serine
4.3.1.17
4.3.1.19
Pyruvate
Pyruvate metabolism
2.7.8.8
4.2.1.22
4.2.1.20
L-Tryptophane
Tryptophane metabolism
Choline
Glycerophospholipid metabolism
O-Phosphatidyl-L-serine
Cystathionine
4.2.1.22
Sulfur metabolism
Cyanoamino acid metabolism
4.4.1.1
2.6.1.45
THF
1.1.41.5.7
1.1.99.1
1.1.3.17
GbsB
Cysteine and methionine metabolism
L-Cysteine
2.1.2.1
5,10-Methylene-THF
Betaine aldehyde
Arginine and proline metabolism
Creatine
2.1.1.2
2.1.4.1
Guanidinoacetate
1.4.2.1
1.4.3.3
Glyoxylate
Glyoxylate metabolism
NH3
[H-protein]-dihydrolipoyllysine
3.5.3.3
2.6.1.4
2.6.1.44
CO2
2.1.2.10
GCSH
1.2.1.8
1.1.3.17
2.1.1.157
THF
1.8.1.4
2.1.1.5
2.1.1.376
1.14.13.251
Dimethylglycine
1.5.3.10
1.5.8.4
1.5.7.3
Sarcosine
1.5.3.1
1.5.3.24
1.5.8.3
1.5.7.3
Glycine
[H-protein]-S-aminomethyl-dihydrolipoyllysine
1.4.4.2
Betaine
[H-protein]-lipoyllysine
2.1.1.20
2.3.1.37
5-Aminolevulinate
Porphyrine biosynthesis
2.1.1.161
2.1.1.156
4.1.2.48
4.1.2.49
L-Allothreonine
Purine metabolism
1.1.1.381
Aminoacetone
2.3.1.29
L-2-Amino-acetoacetate
1.1.1.75
Aminopropan-2-ol
4.1.2.48
4.1.2.5
1.1.1.103
1.4.3.4
1.4.3.21
2-Oxobutanoate
Valine, leucine and isoleucine biosynthesis
4.3.1.19
Threonine
Methylglyoxal
Propanoate metabolism
4.2.3.1
O-Phospho-L-homoserine
Propane-1,3-diamine
2.7.1.39
Arginine and proline metabolism
Cysteine and methionine metabolism
Homoserine
4.1.1.86
1.1.1.3
L-2,4-Diamino-butanoate
Ectoine biosynthesis
Alanine, aspartate and glutamate metabolism
L-Aspartate
2.7.2.4
1.2.1.11
2.6.1.76
2.3.1.178
4.2.1.108
L-Ectoine
1.14.11.55
L-4-Aspartyl-phosphate
L-Aspartate 4-semialdehyde
Nγ-Acetyl-L-2,4-diaminobutyrate
5-Hydroxy-ectoine
DoeC
DoeD
3.5.1.125
3.5.4.44
Nα-Acetyl-L-2,4-diaminobutyrate
Lysine biosynthesis
Data on KEGG graph
Rendered by Pathview

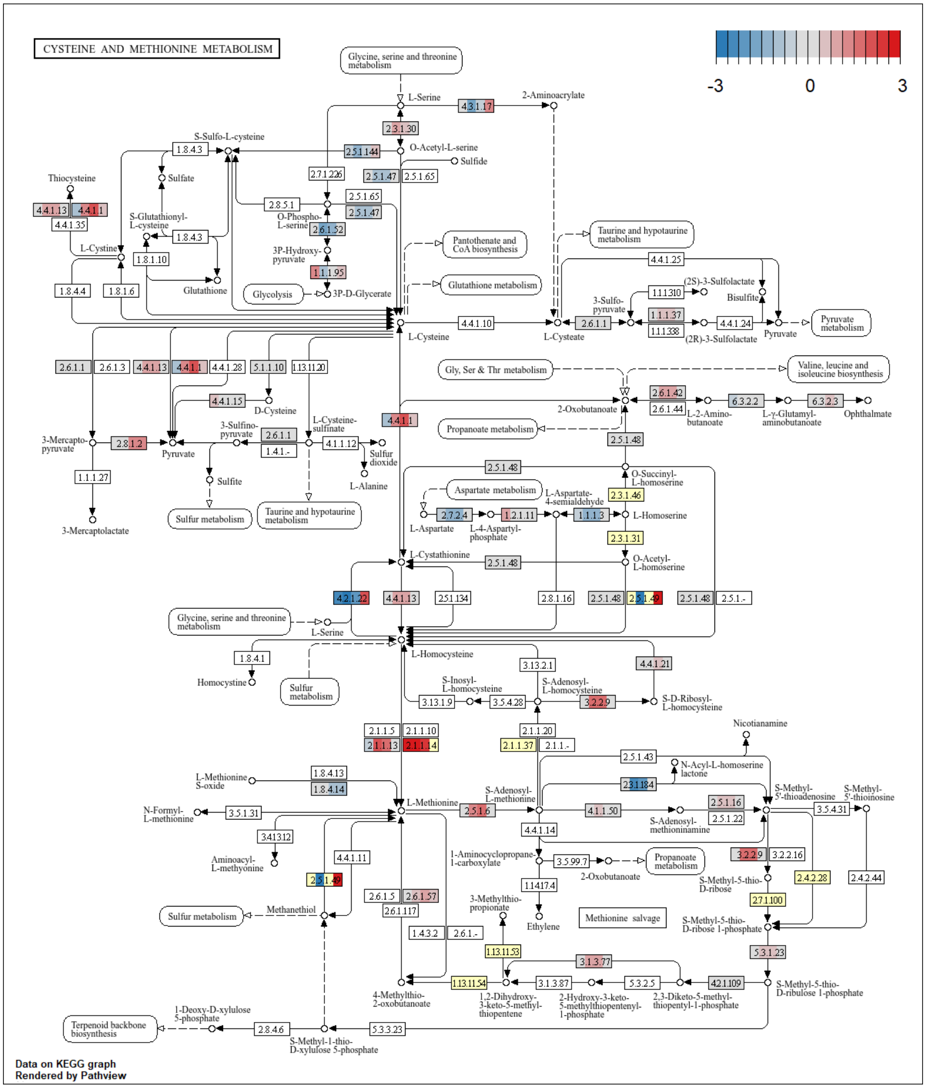
CYSTEINE AND METHIONINE METABOLISM
-3
0
3
Glycine, serine and threonine metabolism
L-Serine
4.3.1.17
2-Aminoacrylate
2.3.1.30
S-Sulfo-L-cysteine
1.8.4.3
2.5.1.144
O-Acetyl-L-serine
Sulfide
2.7.1.226
2.5.1.47
2.5.1.65
Thiocysteine
Sulfate
2.5.1.65
4.4.1.13
4.4.1.1
2.8.5.1
2.5.1.47
4.4.1.35
S-Glutathionyl-L-cysteine
O-Phospho-L-serine
2.6.1.52
Taurine and hypotaurine metabolism
1.8.4.3
Pantothenate and CoA biosynthesis
L-Cystine
3P-Hydroxy-pyruvate
1.8.1.10
4.4.1.25
1.1.1.95
Glutathione metabolism
(2S)-3-Sulfolactate
1.8.4.4
1.8.1.6
Glutathione
Glycolysis
3P-D-Glycerate
1.1.1.310
Bisulfite
3-Sulfo-pyruvate
1.1.1.37
4.4.1.10
2.6.1.1
4.4.1.24
Pyruvate metabolism
L-Cysteine
L-Cysteate
1.1.1.338
(2R)-3-Sulfolactate
Pyruvate
2.6.1.1
2.6.1.3
4.4.1.13
4.4.1.1
4.4.1.28
5.1.1.10
1.13.11.20
Gly, Ser & Thr metabolism
Valine, leucine and isoleucine biosynthesis
4.4.1.15
D-Cysteine
2.6.1.42
2.6.1.44
6.3.2.2
6.3.2.3
4.4.1.1
2-Oxobutanoate
L-2-Amino-butanoate
L-γ-Glutamyl-aminobutanoate
Ophthalmate
3-Sulfino-pyruvate
L-Cysteine-sulfinate
Propanoate metabolism
3-Mercapto-pyruvate
2.8.1.2
Pyruvate
2.6.1.1
1.4.1.-
4.1.1.12
Sulfur dioxide
2.5.1.48
2.5.1.48
1.1.1.27
Sulfite
L-Alanine
O-Succinyl-L-homoserine
Aspartate metabolism
L-Aspartate-4-semialdehyde
2.3.1.46
Sulfur metabolism
Taurine and hypotaurine metabolism
2.7.2.4
1.2.1.11
1.1.1.3
L-Homoserine
3-Mercaptolactate
L-Aspartate
L-4-Aspartyl-phosphate
2.3.1.31
L-Cystathionine
2.5.1.48
O-Acetyl-L-homoserine
4.2.1.22
4.4.1.13
2.5.1.134
2.8.1.16
2.5.1.48
2.5.1.49
2.5.1.48
2.5.1.-
Glycine, serine and threonine metabolism
L-Serine
L-Homocysteine
1.8.4.1
3.13.2.1
4.4.1.21
Homocystine
Sulfur metabolism
S-Inosyl-L-homocysteine
S-Adenosyl-L-homocysteine
3.13.1.9
3.5.4.28
3.2.2.9
S-D-Ribosyl-L-homocysteine
2.1.1.5
2.1.1.10
2.1.1.20
Nicotianamine
2.1.1.13
2.1.1.14
2.1.1.37
2.1.1.-
2.5.1.43
N-Acyl-L-homoserine lactone
L-Methionine S-oxide
1.8.4.13
1.8.4.14
2.3.1.184
S-Methyl-5'-thioadenosine
S-Methyl-5'-thioinosine
L-Methionine
S-Adenosyl-L-methionine
2.5.1.6
4.1.1.50
2.5.1.16
N-Formyl-L-methionine
3.5.1.31
S-Adenosyl-methioninamine
2.5.1.22
3.5.4.31
3.4.13.12
4.4.1.14
1-Aminocyclopropane-1-carboxylate
3.5.99.7
Propanoate metabolism
3.2.2.9
3.2.2.16
Aminoacyl-L-methyonine
4.4.1.11
2-Oxobutanoate
S-Methyl-5-thio-D-ribose
2.4.2.28
2.4.2.44
2.5.1.49
1.14.17.4
2.6.1.5
2.6.1.57
3-Methylthio-propionate
2.7.1.100
Sulfur metabolism
Methanethiol
2.6.1.117
Ethylene
Methionine salvage
S-Methyl-5-thio-D-ribose 1-phosphate
1.4.3.2
2.6.1.-
1.13.11.53
5.3.1.23
3.1.3.77
1.13.11.54
3.1.3.87
5.3.2.5
4.2.1.109
4-Methylthio-2-oxobutanoate
1,2-Dihydroxy-3-keto-5-methyl-thiopentene
2-Hydroxy-3-keto-5-methylthiopentenyl-1-phosphate
2,3-Diketo-5-methyl-thiopentyl-1-phosphate
S-Methyl-5-thio-D-ribulose 1-phosphate
1-Deoxy-D-xylulose 5-phosphate
Terpenoid backbone biosynthesis
2.8.4.6
5.3.3.23
S-Methyl-1-thio-D-xylulose 5-phosphate
Data on KEGG graph
Rendered by Pathview

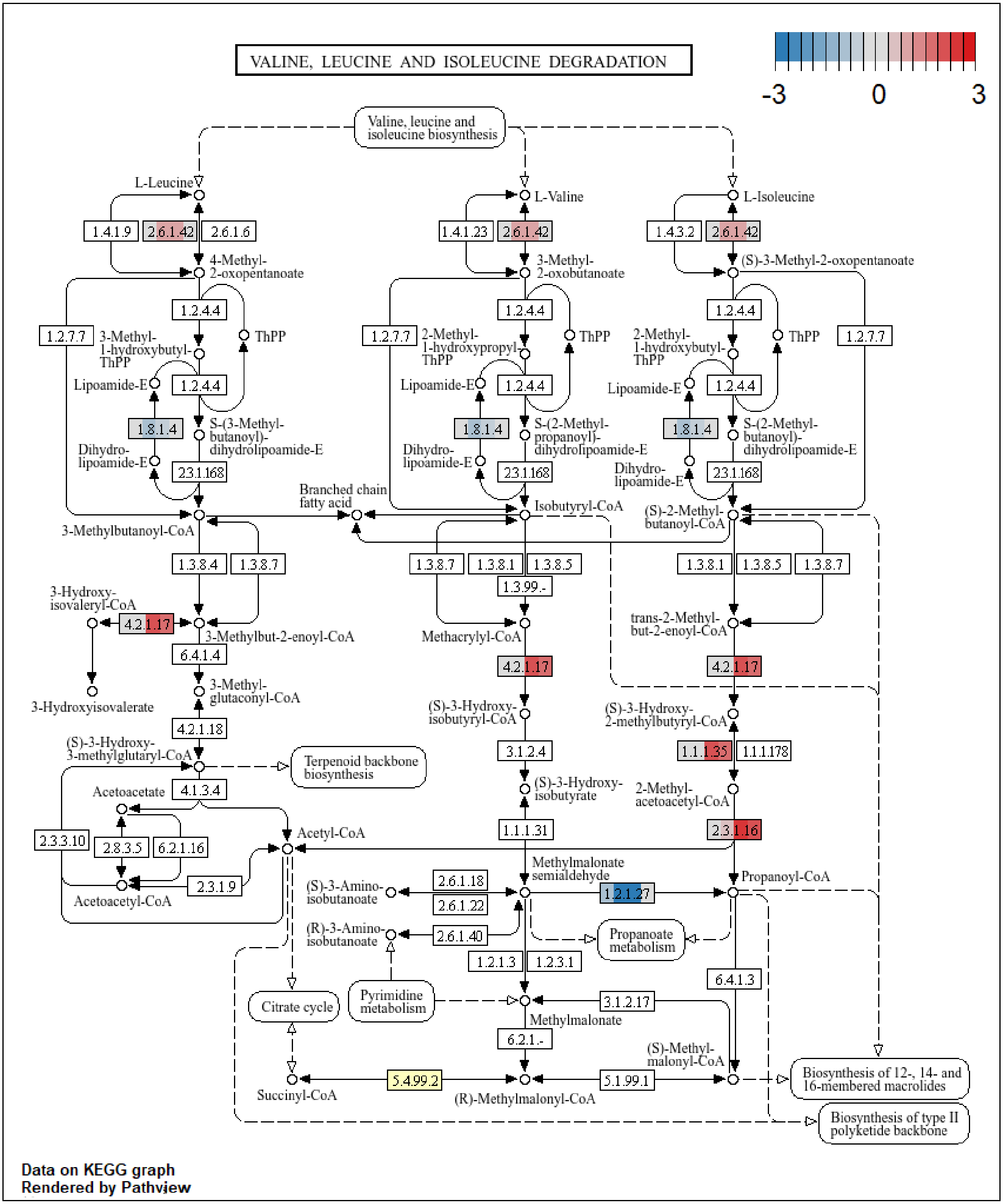

VALINE, LEUCINE AND ISOLEUCINE DEGRADATION
-3
0
3
Valine, leucine and isoleucine biosynthesis
L-Leucine
L-Valine
L-Isoleucine
1.4.1.9
2.6.1.42
2.6.1.6
1.4.1.23
1.4.3.2
4-Methyl-2-oxopentanoate
3-Methyl-2-oxobutanoate
(S)-3-Methyl-2-oxopentanoate
1.2.4.4
1.2.7.7
ThPP
3-Methyl-1-hydroxybutyl-ThPP
2-Methyl-1-hydroxypropyl-ThPP
2-Methyl-1-hydroxybutyl-ThPP
Lipoamide-E
1.8.1.4
S-(3-Methylbutanoyl)-dihydrolipoamide-E
S-(2-Methylpropanoyl)-dihydrolipoamide-E
S-(2-Methylbutanoyl)-dihydrolipoamide-E
Dihydrolipoamide-E
2.3.1.168
Branched chain fatty acid
3-Methylbutanoyl-CoA
Isobutyryl-CoA
(S)-2-Methylbutanoyl-CoA
1.3.8.4
1.3.8.7
1.3.8.1
1.3.8.5
1.3.99.-
3-Hydroxy-isovaleryl-CoA
4.2.1.17
3-Methylbut-2-enoyl-CoA
Methacrylyl-CoA
trans-2-Methylbut-2-enoyl-CoA
6.4.1.4
3-Methylglutaconyl-CoA
3-Hydroxyisovalerate
4.2.1.18
(S)-3-Hydroxyisobutyryl-CoA
(S)-3-Hydroxy-2-methylbutyryl-CoA
(S)-3-Hydroxy-3-methylglutaryl-CoA
Terpenoid backbone biosynthesis
3.1.2.4
1.1.1.35
1.1.1.178
Acetoacetate
4.1.3.4
(S)-3-Hydroxyisobutyrate
2-Methylacetoacetyl-CoA
2.3.3.10
2.8.3.5
6.2.1.16
Acetyl-CoA
1.1.1.31
2.3.1.16
Methylmalonate semialdehyde
Propanoyl-CoA
2.3.1.9
Acetoacetyl-CoA
(S)-3-Amino-isobutanoate
2.6.1.18
2.6.1.22
1.2.1.27
(R)-3-Amino-isobutanoate
2.6.1.40
Propanoate metabolism
1.2.1.3
1.2.3.1
6.4.1.3
Citrate cycle
Pyrimidine metabolism
3.1.2.17
Methylmalonate
6.2.1.-
(S)-Methylmalonyl-CoA
Succinyl-CoA
5.4.99.2
(R)-Methylmalonyl-CoA
5.1.99.1
Biosynthesis of 12-, 14- and 16-membered macrolides
Biosynthesis of type II polyketide backbone
Data on KEGG graph
Rendered by Pathview

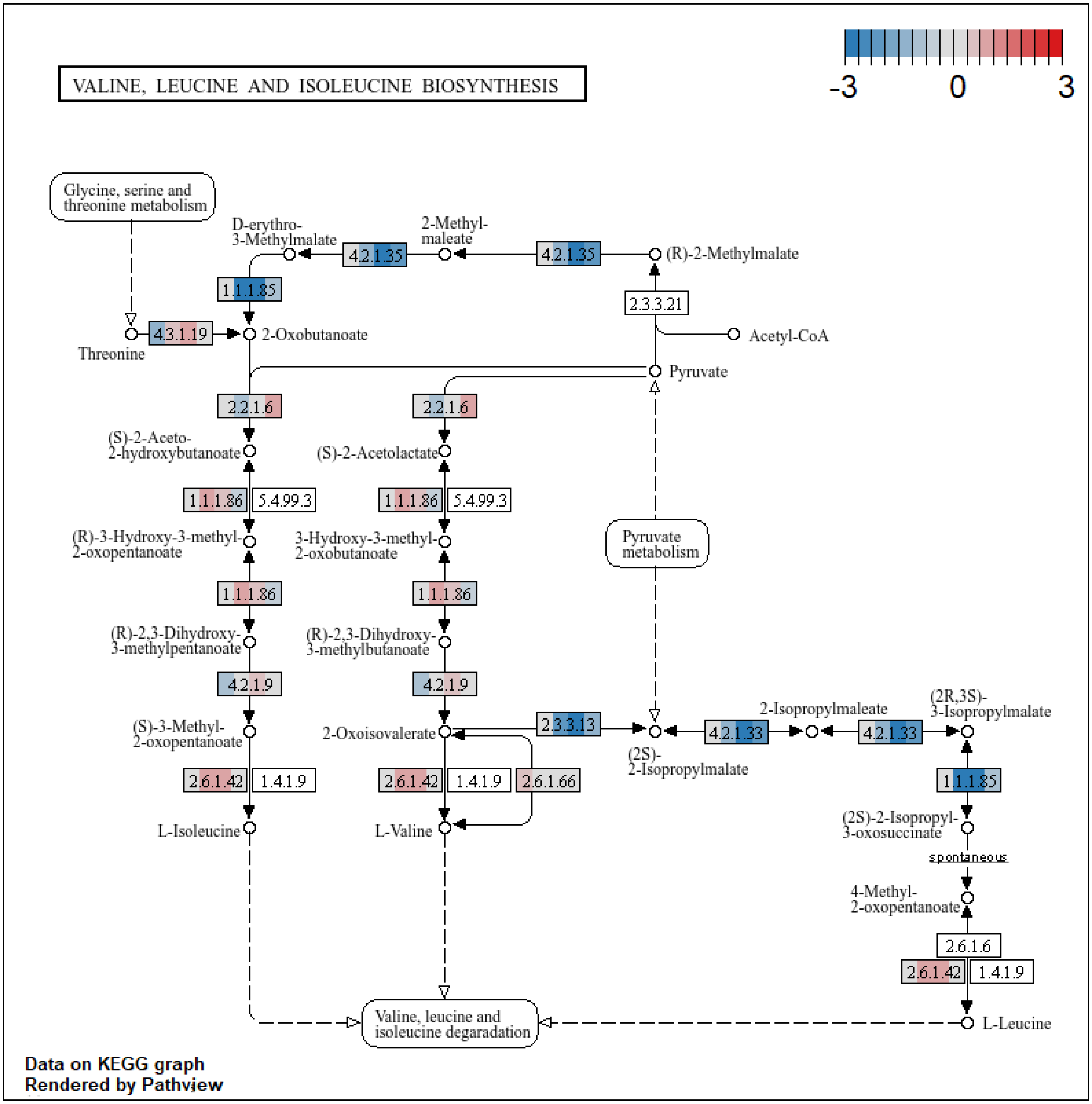

VALINE, LEUCINE AND ISOLEUCINE BIOSYNTHESIS
-3
0
3
Glycine, serine and threonine metabolism
D-erythro-3-Methylmalate
4.2.1.35
2-Methyl-maleate
4.2.1.35
(R)-2-Methylmalate
1.1.1.85
2.3.3.21
Acetyl-CoA
4.3.1.19
2-Oxobutanoate
Threonine
Pyruvate
2.2.1.6
2.2.1.6
(S)-2-Aceto-2-hydroxybutanoate
(S)-2-Acetolactate
1.1.1.86
5.4.99.3
1.1.1.86
5.4.99.3
(R)-3-Hydroxy-3-methyl-2-oxopentanoate
3-Hydroxy-3-methyl-2-oxobutanoate
Pyruvate metabolism
1.1.1.86
1.1.1.86
(R)-2,3-Dihydroxy-3-methylpentanoate
(R)-2,3-Dihydroxy-3-methylbutanoate
4.2.1.9
4.2.1.9
2.3.3.13
2-Isopropylmaleate
(2R,3S)-3-Isopropylmalate
(S)-3-Methyl-2-oxopentanoate
2-Oxoisovalerate
4.2.1.33
4.2.1.33
(2S)-2-Isopropylmalate
2.6.1.42
1.4.1.9
2.6.1.42
1.4.1.9
2.6.1.66
1.1.1.85
(2S)-2-Isopropyl-3-oxosuccinate
L-Isoleucine
L-Valine
spontaneous
4-Methyl-2-oxopentanoate
2.6.1.6
2.6.1.42
1.4.1.9
Valine, leucine and isoleucine degaradation
L-Leucine
Data on KEGG graph
Rendered by Pathview

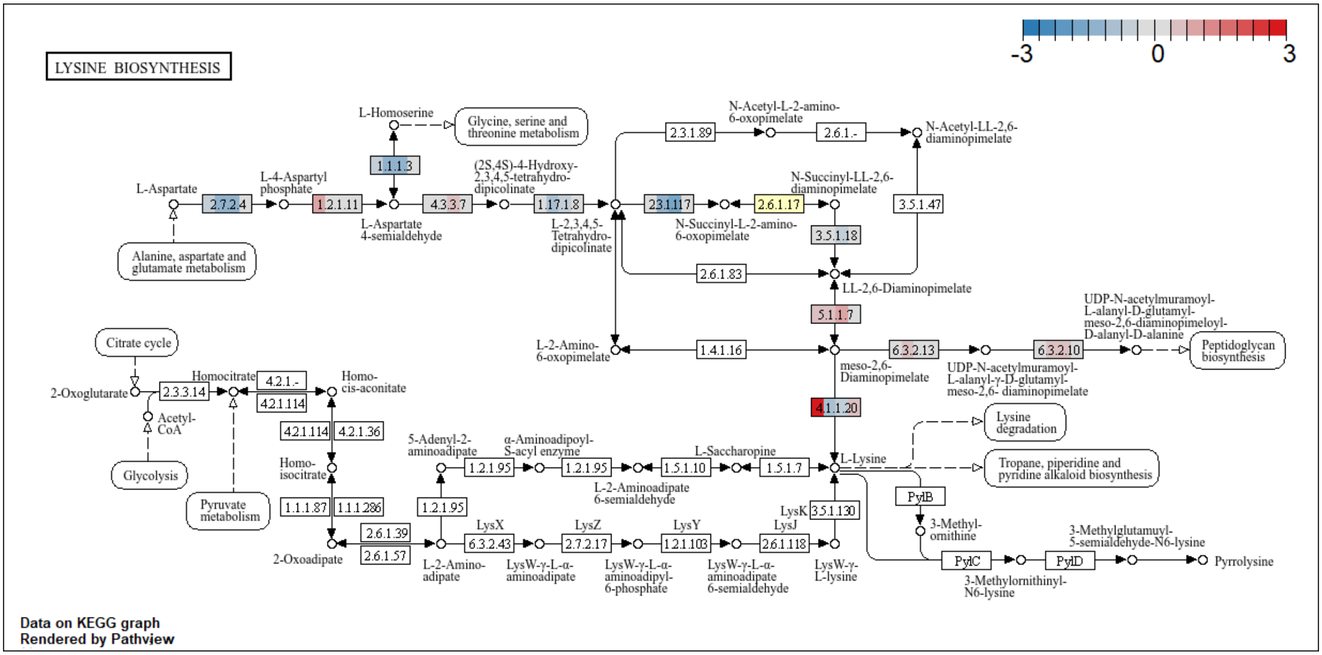

LYSINE BIOSYNTHESIS
-3
0
3
L-Homoserine
Glycine, serine and threonine metabolism
N-Acetyl-L-2-amino-6-oxopimelate
N-Acetyl-LL-2,6-diaminopimelate
L-Aspartate
L-4-Aspartyl phosphate
(2S,4S)-4-Hydroxy-2,3,4,5-tetrahydro-dipicolinate
N-Succinyl-LL-2,6-diaminopimelate
L-Aspartate 4-semialdehyde
L-2,3,4,5-Tetrahydro-dipicolinate
N-Succinyl-L-2-amino-6-oxopimelate
Alanine, aspartate and glutamate metabolism
LL-2,6-Diaminopimelate
UDP-N-acetylmuramoyl-L-alanyl-D-glutamyl-meso-2,6-diaminopimeloyl-D-alanyl-D-alanine
Citrate cycle
L-2-Amino-6-oxopimelate
Peptidoglycan biosynthesis
meso-2,6-Diaminopimelate
UDP-N-acetylmuramoyl-L-alanyl-γ-D-glutamyl-meso-2,6-diaminopimelate
Homocitrate
Homo-cis-aconitate
2-Oxoglutarate
Acetyl-CoA
Lysine degradation
5-Adenyl-2-aminoadipate
α-Aminoadipoyl-S-acyl enzyme
L-Saccharopine
Homo-isocitrate
L-Lysine
Glycolysis
Tropane, piperidine and pyridine alkaloid biosynthesis
L-2-Aminoadipate 6-semialdehyde
Pyruvate metabolism
3-Methyl-ornithine
3-Methylglutamuyl-5-semialdehyde-N6-lysine
2-Oxoadipate
L-2-Amino-adipate
LysW-γ-L-α-aminoadipate
LysW-γ-L-α-aminoadipyl-6-phosphate
LysW-γ-L-α-aminoadipate 6-semialdehyde
LysW-γ-L-lysine
Pyrrolysine
3-Methylornithinyl-N6-lysine
2.7.2.4
1.2.1.11
1.1.1.3
4.3.3.7
1.17.1.8
2.3.1.89
2.6.1.-
2.3.1.117
2.6.1.17
3.5.1.47
3.5.1.18
2.6.1.83
5.1.1.7
1.4.1.16
6.3.2.13
6.3.2.10
4.1.1.20
2.3.3.14
4.2.1.-
4.2.1.114
4.2.1.36
1.1.1.87
1.1.1.286
1.2.1.95
1.5.1.10
1.5.1.7
2.6.1.39
2.6.1.57
6.3.2.43
2.7.2.17
1.2.1.103
2.6.1.118
3.5.1.130
LysX
LysZ
LysY
LysK
LysJ
PylB
PylC
PylD
Data on KEGG graph
Rendered by Pathview

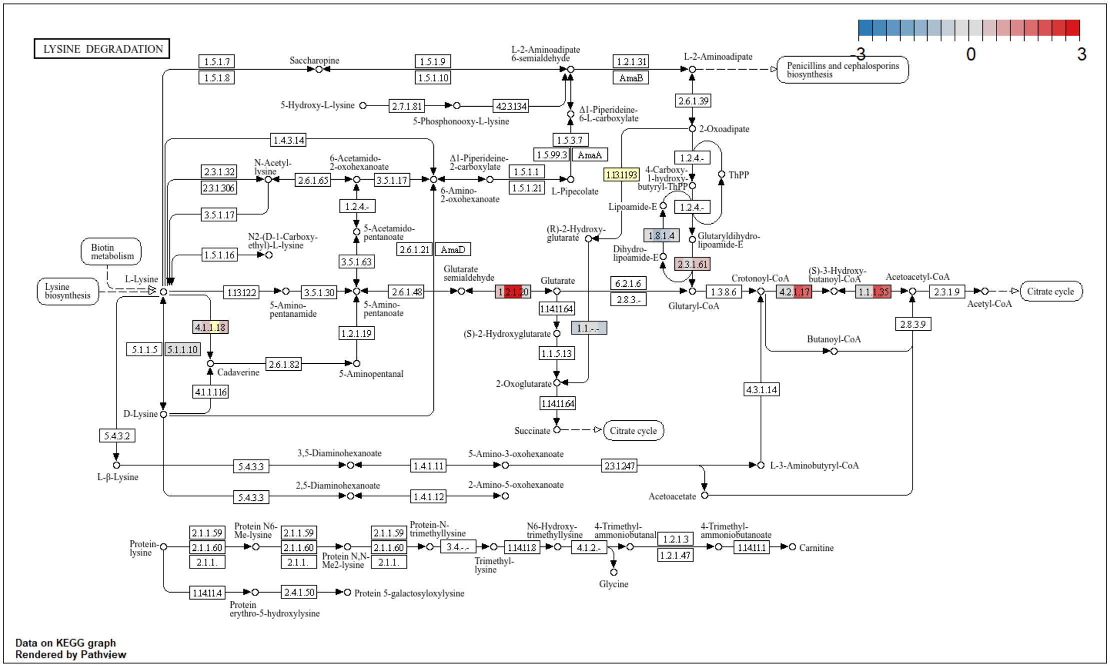

LYSINE DEGRADATION
-3
0
3
Saccharopine
L-2-Aminoadipate 6-semialdehyde
L-2-Aminoadipate
Penicillins and cephalosporins biosynthesis
5-Hydroxy-L-lysine
5-Phosphonooxy-L-lysine
Δ1-Piperideine-6-L-carboxylate
2-Oxoadipate
N-Acetyl-lysine
6-Acetamido-2-oxohexanoate
Δ1-Piperideine-2-carboxylate
6-Amino-2-oxohexanoate
L-Pipecolate
ThPP
4-Carboxy-1-hydroxy-butyryl-ThPP
Lipoamide-E
5-Acetamido-pentanoate
N2-(D-1-Carboxy-ethyl)-L-lysine
(R)-2-Hydroxy-glutarate
Glutaryldihydro-lipoamide-E
Dihydro-lipoamide-E
Biotin metabolism
Lysine biosynthesis
L-Lysine
Glutarate semialdehyde
Glutarate
Crotonoyl-CoA
(S)-3-Hydroxy-butanoyl-CoA
Acetoacetyl-CoA
Citrate cycle
Acetyl-CoA
5-Amino-pentanamide
5-Amino-pentanoate
Glutaryl-CoA
(S)-2-Hydroxyglutarate
Butanoyl-CoA
Cadaverine
5-Aminopentanal
2-Oxoglutarate
D-Lysine
Succinate
Citrate cycle
L-β-Lysine
3,5-Diaminohexanoate
5-Amino-3-oxohexanoate
L-3-Aminobutyryl-CoA
2,5-Diaminohexanoate
2-Amino-5-oxohexanoate
Acetoacetate
Protein-lysine
Protein N6-Me-lysine
Protein N,N-Me2-lysine
Protein-N-trimethyllysine
Trimethyl-lysine
N6-Hydroxy-trimethyllysine
4-Trimethyl-ammoniobutanal
Glycine
4-Trimethyl-ammoniobutanoate
Carnitine
Protein erythro-5-hydroxylysine
Protein 5-galactosyloxylysine
1.5.1.7
1.5.1.8
1.5.1.9
1.5.1.10
1.2.1.31
AmaB
2.7.1.81
4.2.3.134
2.6.1.39
1.4.3.14
1.5.3.7
1.5.99.3
AmaA
2.3.1.32
2.3.1.306
2.6.1.65
3.5.1.17
1.5.1.1
1.5.1.21
1.13.11.93
1.2.4.-
3.5.1.17
1.2.4.-
1.5.1.16
2.6.1.21
AmaD
3.5.1.63
1.8.1.4
2.3.1.61
1.13.12
3.5.1.30
2.6.1.48
1.2.1.20
6.2.1.6
2.8.3.-
1.3.8.6
4.2.1.17
1.1.1.35
2.3.1.9
4.1.1.18
1.14.11.64
1.1.-.-
2.8.3.9
5.1.1.5
5.1.1.10
2.6.1.82
1.2.1.19
1.1.5.13
4.1.1.116
4.3.1.14
1.14.11.64
5.4.3.2
5.4.3.3
1.4.1.11
2.3.1.247
5.4.3.3
1.4.1.12
2.1.1.59
2.1.1.60
2.1.1.
3.4.-.-
1.14.11.8
4.1.2.-
1.2.1.3
1.2.1.47
1.14.11.1
1.14.11.4
2.4.1.50
Data on KEGG graph
Rendered by Pathview

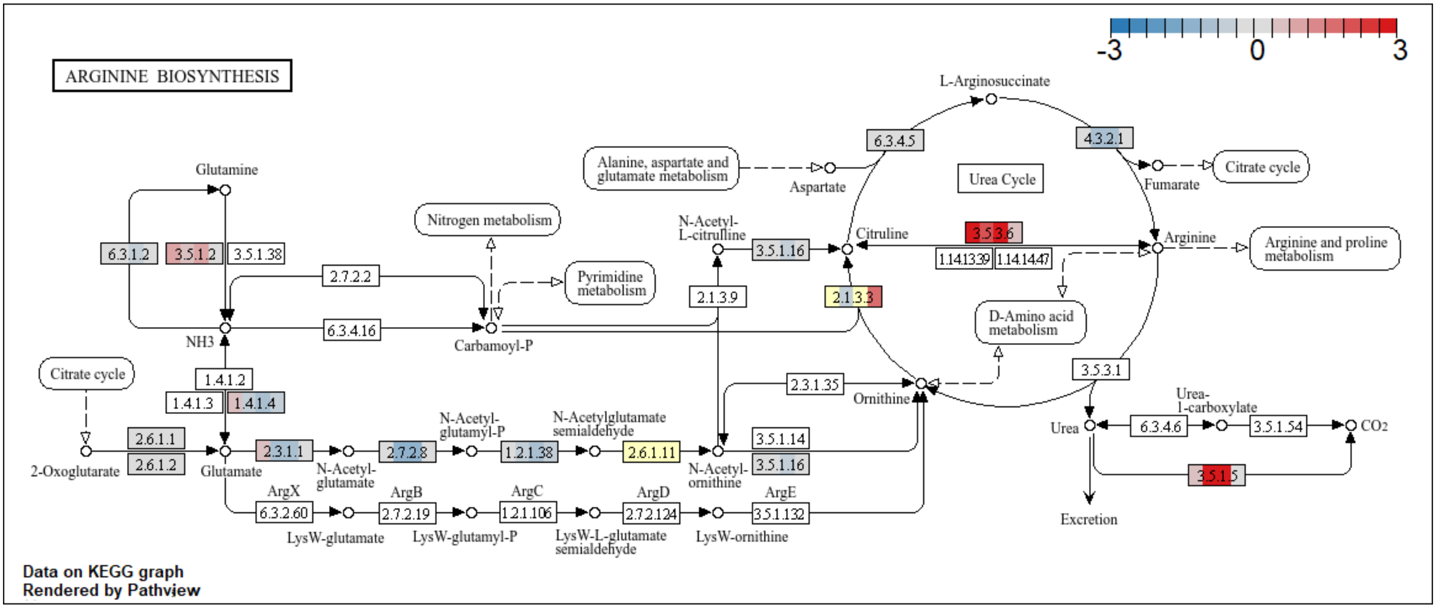
ARGININE BIOSYNTHESIS
-3
0
3
L-Arginosuccinate
6.3.4.5
4.3.2.1
Alanine, aspartate and glutamate metabolism
Aspartate
Urea Cycle
Citrate cycle
Fumarate
Glutamine
Nitrogen metabolism
N-Acetyl-L-citrulline
Citruline
3.5.3.6
Arginine
Arginine and proline metabolism
6.3.1.2
3.5.1.2
3.5.1.38
2.7.2.2
Pyrimidine metabolism
3.5.1.16
1.14.13.39
1.14.14.47
2.1.3.9
2.1.3.3
D-Amino acid metabolism
NH3
6.3.4.16
Carbamoyl-P
Citrate cycle
1.4.1.2
1.4.1.3
1.4.1.4
2.3.1.35
Ornithine
3.5.3.1
Urea-1-carboxylate
Urea
6.3.4.6
3.5.1.54
CO2
2.6.1.1
2.6.1.2
2-Oxoglutarate
Glutamate
2.3.1.1
N-Acetyl-glutamate
N-Acetyl-glutamyl-P
2.7.2.8
1.2.1.38
N-Acetylglutamate semialdehyde
2.6.1.11
N-Acetyl-ornithine
3.5.1.14
3.5.1.16
3.5.1.5
Excretion
ArgX
6.3.2.60
ArgB
2.7.2.19
ArgC
1.2.1.106
ArgD
2.7.2.124
ArgE
3.5.1.132
LysW-glutamate
LysW-glutamyl-P
LysW-L-glutamate semialdehyde
LysW-ornithine
Data on KEGG graph
Rendered by Pathview

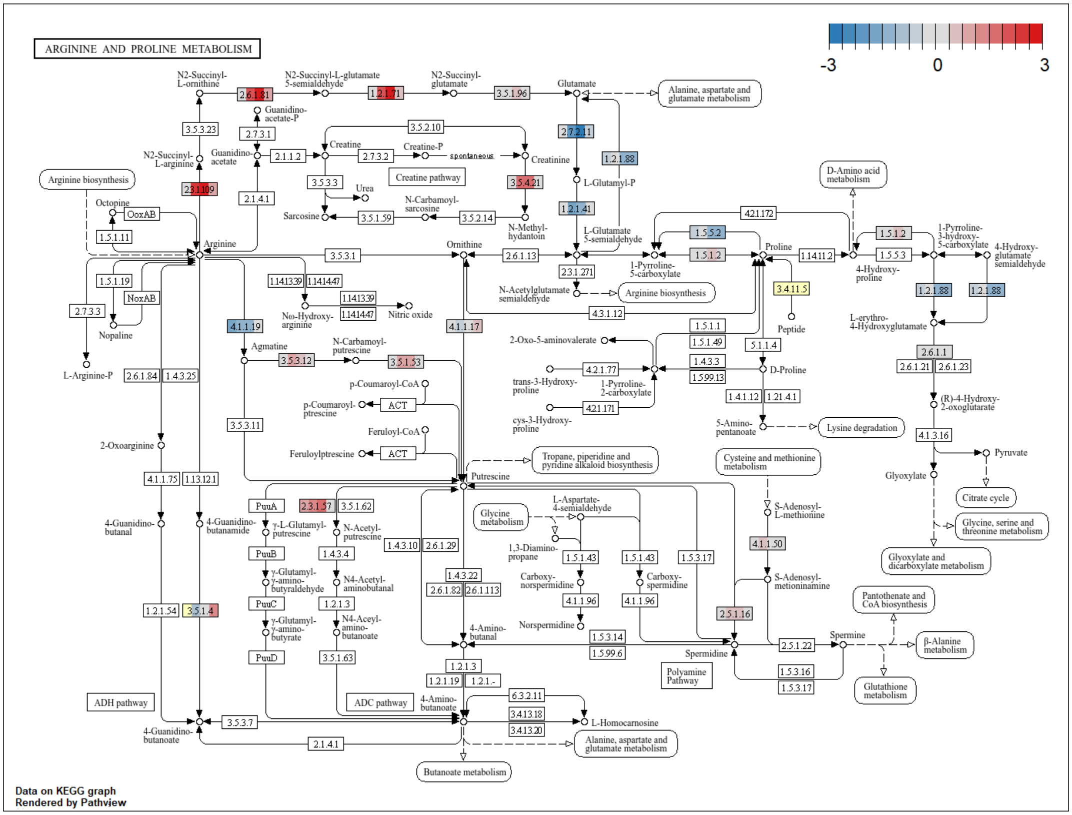

ARGININE AND PROLINE METABOLISM
-3
0
3
N2-Succinyl-L-ornithine
N2-Succinyl-L-glutamate 5-semialdehyde
N2-Succinyl-glutamate
Glutamate
Alanine, aspartate and glutamate metabolism
Guanidino-acetate-P
N2-Succinyl-L-arginine
Guanidino-acetate
Creatine
Creatine-P
spontaneous
Creatinine
Creatine pathway
Arginine biosynthesis
Urea
Octopine
OoxAB
Sarcosine
N-Carbamoyl-sarcosine
N-Methyl-hydantoin
L-Glutamyl-P
L-Glutamate 5-semialdehyde
D-Amino acid metabolism
Arginine
Ornithine
Proline
1-Pyrroline-5-carboxylate
1-Pyrroline-3-hydroxy-5-carboxylate
4-Hydroxy-proline
4-Hydroxy-glutamate semialdehyde
NoxAB
Nopaline
L-Arginine-P
Nω-Hydroxy-arginine
Nitric oxide
N-Acetylglutamate semialdehyde
Peptide
L-erythro-4-Hydroxyglutamate
Agmatine
N-Carbamoyl-putrescine
2-Oxo-5-aminovalerate
trans-3-Hydroxy-proline
D-Proline
cys-3-Hydroxy-proline
p-Coumaroyl-CoA
p-Coumaroyl-ptrescine
ACT
Feruloyl-CoA
Feruloylptrescine
5-Amino-pentanoate
Lysine degradation
(R)-4-Hydroxy-2-oxoglutarate
Pyruvate
2-Oxoarginine
Tropane, piperidine and pyridine alkaloid biosynthesis
Cysteine and methionine metabolism
Glyoxylate
Citrate cycle
Putrescine
PuuA
4-Guanidino-butanal
4-Guanidino-butanamide
γ-L-Glutamyl-putrescine
N-Acetyl-putrescine
Glycine metabolism
L-Aspartate-4-semialdehyde
S-Adenosyl-L-methionine
Glycine, serine and threonine metabolism
PuuB
1,3-Diamino-propane
Glyoxylate and dicarboxylate metabolism
γ-Glutamyl-γ-amino-butyraldehyde
N4-Acetyl-aminobutanal
Carboxy-norspermidine
Carboxy-spermidine
S-Adenosyl-methioninamine
Pantothenate and CoA biosynthesis
PuuC
γ-Glutamyl-γ-amino-butyrate
N4-Acetyl-amino-butanoate
Norspermidine
4-Amino-butanal
Spermidine
Spermine
β-Alanine metabolism
PuuD
Polyamine Pathway
Glutathione metabolism
ADH pathway
ADC pathway
4-Amino-butanoate
L-Homocarnosine
4-Guanidino-butanoate
Alanine, aspartate and glutamate metabolism
Butanoate metabolism
Data on KEGG graph
Rendered by Pathview

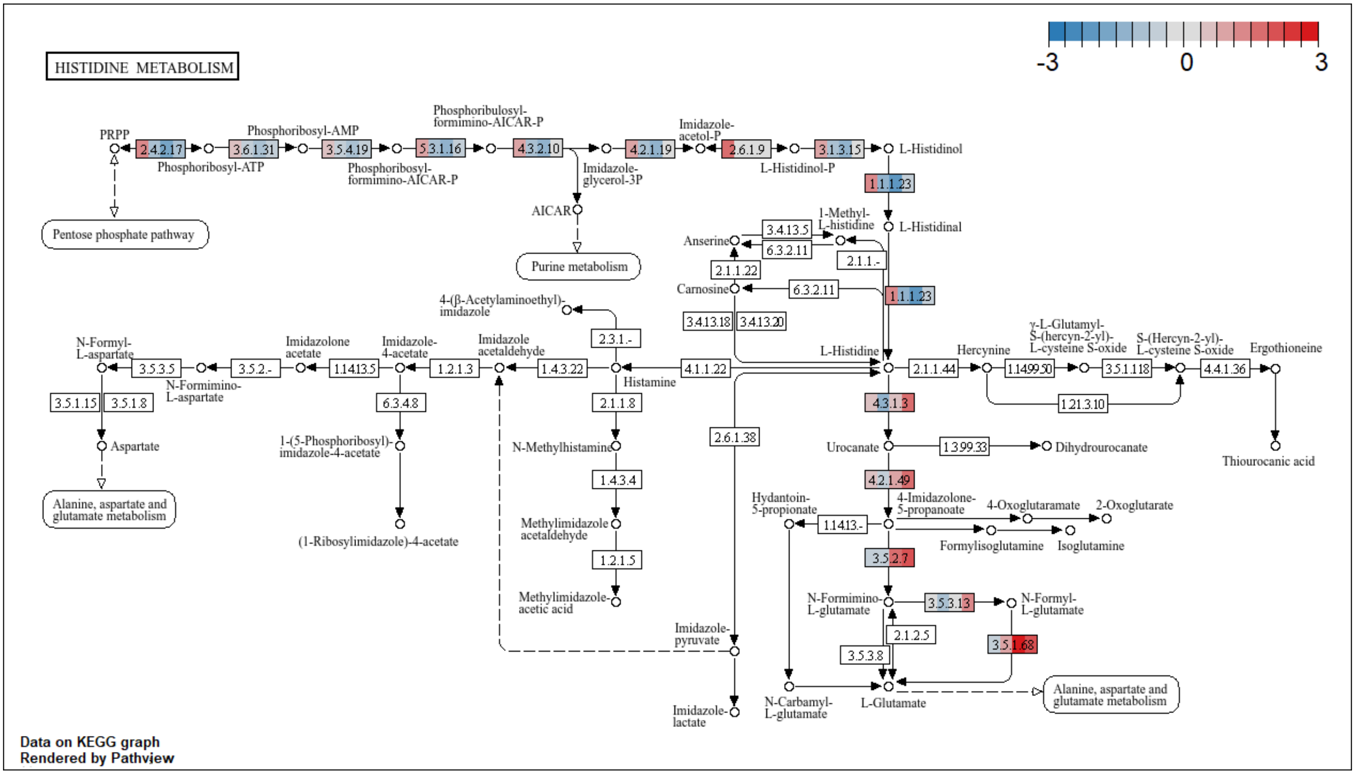

HISTIDINE METABOLISM
-3
0
3
PRPP
Phosphoribosyl-AMP
Phosphoribosyl-ATP
Phosphoribulosyl-formimino-AICAR-P
Phosphoribosyl-formimino-AICAR-P
Imidazole-acetol-P
Imidazole-glycerol-3P
L-Histidinol-P
L-Histidinol
L-Histidinal
AICAR
Pentose phosphate pathway
Purine metabolism
Anserine
Carnosine
1-Methyl-L-histidine
4-(β-Acetylaminoethyl)-imidazole
L-Histidine
Histamine
Hercynine
Ergothioneine
Thiourocanic acid
Urocanate
Dihydrourocanate
N-Formyl-L-aspartate
N-Formimino-L-aspartate
Aspartate
Imidazolone acetate
Imidazole-4-acetate
Imidazole acetaldehyde
1-(5-Phosphoribosyl)-imidazole-4-acetate
(1-Ribosylimidazole)-4-acetate
N-Methylhistamine
Methylimidazole acetaldehyde
Methylimidazole-acetic acid
Imidazole-pyruvate
Imidazole-lactate
Hydantoin-5-propionate
4-Imidazolone-5-propanoate
4-Oxoglutaramate
2-Oxoglutarate
Formylisoglutamine
Isoglutamine
N-Formimino-L-glutamate
N-Formyl-L-glutamate
N-Carbamyl-L-glutamate
L-Glutamate
Alanine, aspartate and glutamate metabolism
Data on KEGG graph
Rendered by Pathview

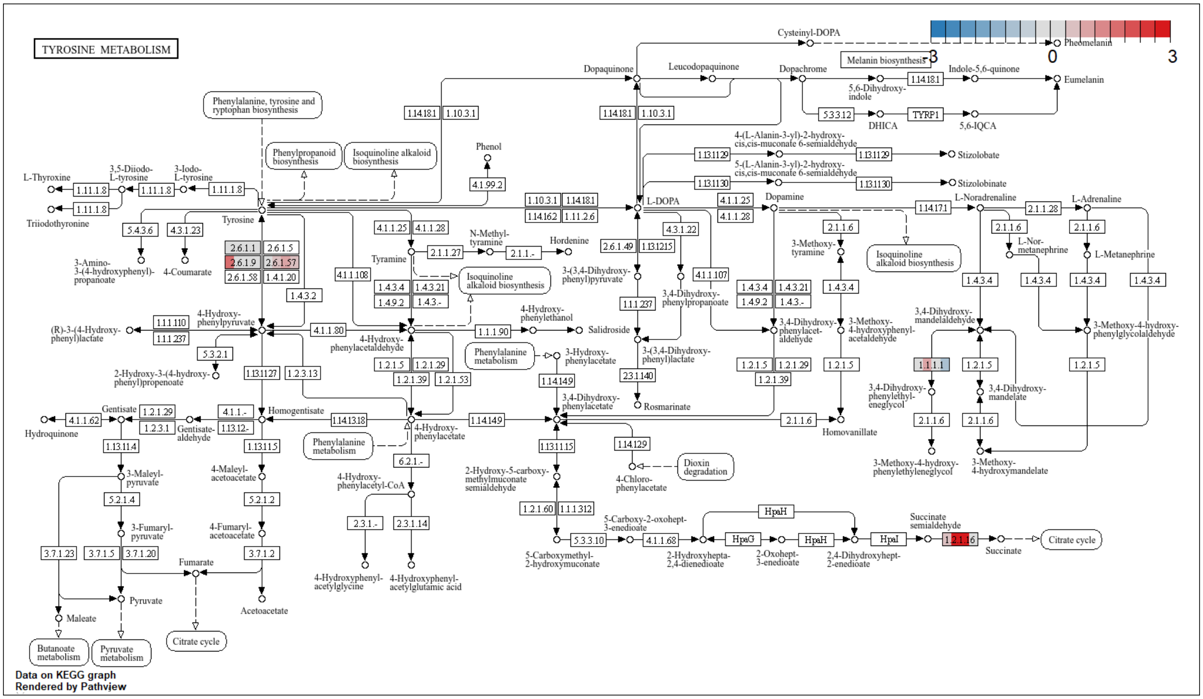
TYROSINE METABOLISM
-3
0
3
Cysteinyl-DOPA
Pheomelanin
Melanin biosynthesis
Dopaquinone
Leucodopaquinone
Dopachrome
Indole-5,6-quinone
Eumelanin
5,6-Dihydroxy-indole
DHICA
5,6-IQCA
TYRP1
Phenylalanine, tyrosine and tryptophan biosynthesis
Phenylpropanoid biosynthesis
Isoquinoline alkaloid biosynthesis
Phenol
L-Thyroxine
3,5-Diiodo-L-tyrosine
3-Iodo-L-tyrosine
Triiodothyronine
Tyrosine
L-DOPA
Dopamine
L-Noradrenaline
L-Adrenaline
Stizolobate
Stizolobinate
N-Methyl-tyramine
Hordenine
Tyramine
3-Methoxy-tyramine
L-Nor-metanephrine
L-Metanephrine
3-Amino-3-(4-hydroxyphenyl)-propanoate
4-Coumarate
4-Hydroxy-phenylpyruvate
(R)-3-(4-Hydroxy-phenyl)lactate
2-Hydroxy-3-(4-hydroxy-phenyl)propenoate
4-Hydroxy-phenylacetaldehyde
4-Hydroxy-phenylethanol
Salidroside
3-Hydroxy-phenylacetate
Phenylalanine metabolism
3-(3,4-Dihydroxy-phenyl)pyruvate
3,4-Dihydroxy-phenylpropanoate
3-(3,4-Dihydroxy-phenyl)lactate
Rosmarinate
3,4-Dihydroxy-phenylacetaldehyde
3-Methoxy-4-hydroxyphenyl-acetaldehyde
3,4-Dihydroxy-mandelaldehyde
3-Methoxy-4-hydroxy-phenylglycolaldehyde
3,4-Dihydroxy-phenylethyleneglycol
3,4-Dihydroxy-mandelate
Homovanillate
3-Methoxy-4-hydroxy-phenylethyleneglycol
3-Methoxy-4-hydroxymandelate
Gentisate
Gentisate-aldehyde
Hydroquinone
Homogentisate
4-Hydroxy-phenylacetate
3,4-Dihydroxy-phenylacetate
4-Chloro-phenylacetate
Dioxin degradation
3-Maleyl-pyruvate
4-Maleyl-acetoacetate
4-Hydroxy-phenylacetyl-CoA
2-Hydroxy-5-carboxy-methylmuconate semialdehyde
5-Carboxy-2-oxohept-3-enedioate
Succinate semialdehyde
Succinate
Citrate cycle
3-Fumaryl-pyruvate
4-Fumaryl-acetoacetate
Fumarate
Pyruvate
Maleate
Acetoacetate
4-Hydroxyphenyl-acetylglycine
4-Hydroxyphenyl-acetylglutamic acid
5-Carboxymethyl-2-hydroxymuconate
2-Hydroxyhepta-2,4-dienedioate
2-Oxohept-3-enedioate
2,4-Dihydroxyhept-2-enedioate
HpaH
HpaG
HpaI
Butanoate metabolism
Pyruvate metabolism
Citrate cycle
Data on KEGG graph
Rendered by Pathview

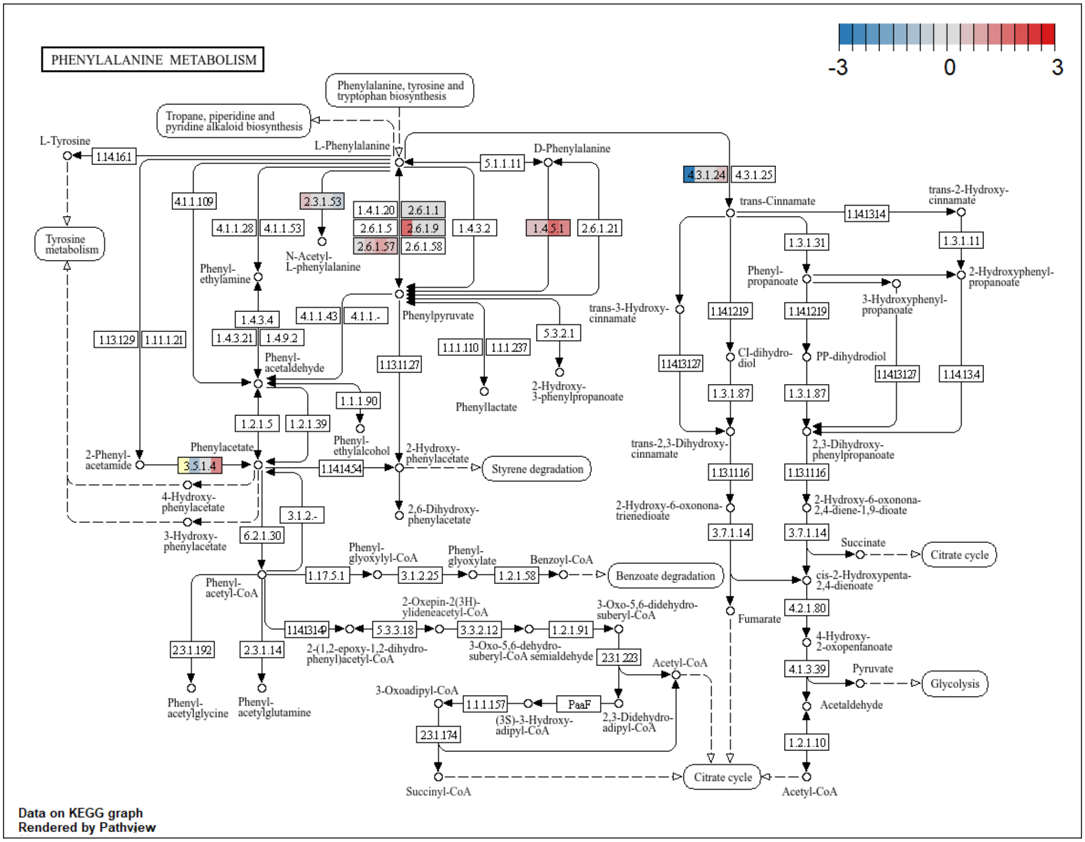
PHENYLALANINE METABOLISM
-3
0
3
Phenylalanine, tyrosine and tryptophan biosynthesis
Tropane, piperidine and pyridine alkaloid biosynthesis
L-Tyrosine
L-Phenylalanine
D-Phenylalanine
Tyrosine metabolism
1.14.16.1
5.1.1.11
4.3.1.24
4.3.1.25
trans-Cinnamate
trans-2-Hydroxy-cinnamate
1.14.13.14
4.1.1.109
2.3.1.53
1.4.1.20
2.6.1.1
2.6.1.5
2.6.1.9
2.6.1.57
2.6.1.58
1.4.3.2
1.4.5.1
2.6.1.21
4.1.1.28
4.1.1.53
N-Acetyl-L-phenylalanine
Phenyl-ethylamine
1.3.1.31
1.3.1.11
Phenyl-propanoate
2-Hydroxyphenyl-propanoate
3-Hydroxyphenyl-propanoate
1.4.3.4
4.1.1.43
4.1.1.-
Phenylpyruvate
trans-3-Hydroxy-cinnamate
1.14.12.19
1.13.12.9
1.11.1.21
1.4.3.21
1.4.9.2
5.3.2.1
Phenyl-acetaldehyde
1.1.1.110
1.1.1.237
1.13.11.27
1.14.13.127
Cl-dihydro-diol
PP-dihydrodiol
1.14.13.4
2-Hydroxy-3-phenylpropanoate
Phenyllactate
1.1.1.90
1.3.1.87
1.2.1.5
1.2.1.39
Phenyl-ethylalcohol
trans-2,3-Dihydroxy-cinnamate
2,3-Dihydroxy-phenylpropanoate
2-Phenyl-acetamide
Phenylacetate
3.5.1.4
2-Hydroxy-phenylacetate
1.14.14.54
Styrene degradation
1.13.11.16
4-Hydroxy-phenylacetate
2,6-Dihydroxy-phenylacetate
3.1.2.-
2-Hydroxy-6-oxonona-trienedioate
2-Hydroxy-6-oxonona-2,4-diene-1,9-dioate
3-Hydroxy-phenylacetate
6.2.1.30
3.7.1.14
Succinate
Citrate cycle
Phenyl-glyoxylyl-CoA
Phenyl-glyoxylate
Benzoyl-CoA
Phenyl-acetyl-CoA
1.17.5.1
3.1.2.25
1.2.1.58
Benzoate degradation
cis-2-Hydroxypenta-2,4-dienoate
2-Oxepin-2(3H)-ylideneacetyl-CoA
3-Oxo-5,6-didehydro-suberyl-CoA
4.2.1.80
Fumarate
1.14.13.149
5.3.3.18
3.3.2.12
1.2.1.91
4-Hydroxy-2-oxopentanoate
2.3.1.192
2.3.1.14
2-(1,2-epoxy-1,2-dihydro-phenyl)acetyl-CoA
3-Oxo-5,6-dehydro-suberyl-CoA semialdehyde
2.3.1.223
Acetyl-CoA
4.1.3.39
Pyruvate
Glycolysis
3-Oxoadipyl-CoA
1.1.1.157
PaaF
Acetaldehyde
Phenyl-acetylglycine
Phenyl-acetylglutamine
(3S)-3-Hydroxy-adipyl-CoA
2,3-Didehydro-adipyl-CoA
2.3.1.174
1.2.1.10
Succinyl-CoA
Citrate cycle
Acetyl-CoA
Data on KEGG graph
Rendered by Pathview

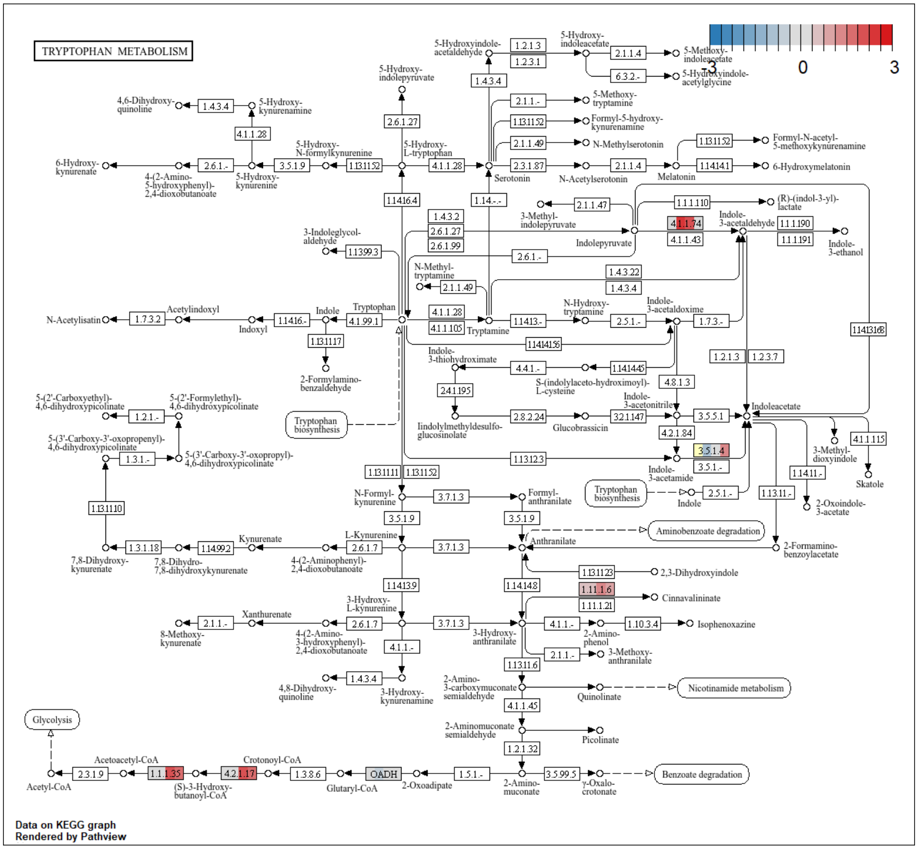
TRYPTOPHAN METABOLISM
-3
0
3
5-Hydroxyindole-acetaldehyde
5-Hydroxy-indoleacetate
5-Methoxy-indoleacetate
5-Hydroxyindole-acetylglycine
5-Hydroxy-indolepyruvate
4,6-Dihydroxy-quinoline
5-Hydroxy-kynurenamine
5-Methoxy-tryptamine
Formyl-5-hydroxy-kynurenamine
N-Methylserotonin
Formyl-N-acetyl-5-methoxykynurenamine
6-Hydroxy-kynurenate
5-Hydroxy-N-formylkynurenine
5-Hydroxy-L-tryptophan
Serotonin
N-Acetylserotonin
Melatonin
6-Hydroxymelatonin
4-(2-Amino-5-hydroxyphenyl)-2,4-dioxobutanoate
5-Hydroxy-kynurenine
(R)-(indol-3-yl)-lactate
3-Methyl-indolepyruvate
Indole-3-acetaldehyde
Indole-3-ethanol
3-Indoleglycol-aldehyde
Indolepyruvate
N-Methyl-tryptamine
N-Acetylisatin
Acetylindoxyl
Indoxyl
Indole
Tryptophan
Tryptamine
N-Hydroxy-tryptamine
Indole-3-acetaldoxime
2-Formylamino-benzaldehyde
Indole-3-thiohydroximate
S-(indolylaceto-hydroximoyl)-L-cysteine
Indole-3-acetonitrile
Indoleacetate
5-(2'-Carboxyethyl)-4,6-dihydroxypicolinate
5-(2'-Formylethyl)-4,6-dihydroxypicolinate
Tryptophan biosynthesis
Iindolylmethyldesulfo-glucosinolate
Glucobrassicin
5-(3'-Carboxy-3'-oxopropenyl)-4,6-dihydroxypicolinate
5-(3'-Carboxy-3'-oxopropyl)-4,6-dihydroxypicolinate
3-Methyl-dioxyindole
Skatole
Indole-3-acetamide
N-Formyl-kynurenine
Formyl-anthranilate
2-Oxoindole-3-acetate
Aminobenzoate degradation
Anthranilate
2-Formamino-benzoylacetate
7,8-Dihydroxy-kynurenate
7,8-Dihydro-7,8-dihydroxykynurenate
Kynurenate
4-(2-Aminophenyl)-2,4-dioxobutanoate
L-Kynurenine
2,3-Dihydroxyindole
Cinnavalininate
Xanthurenate
8-Methoxy-kynurenate
4-(2-Amino-3-hydroxyphenyl)-2,4-dioxobutanoate
3-Hydroxy-L-kynurenine
3-Hydroxy-anthranilate
2-Amino-phenol
Isophenoxazine
3-Methoxy-anthranilate
4,8-Dihydroxy-quinoline
3-Hydroxy-kynurenamine
2-Amino-3-carboxymuconate semialdehyde
Quinolinate
Nicotinamide metabolism
Glycolysis
2-Aminomuconate semialdehyde
Picolinate
Acetoacetyl-CoA
Crotonoyl-CoA
Acetyl-CoA
(S)-3-Hydroxy-butanoyl-CoA
Glutaryl-CoA
2-Oxoadipate
2-Amino-muconate
γ-Oxalo-crotonate
Benzoate degradation
OADH
Data on KEGG graph
Rendered by Pathview

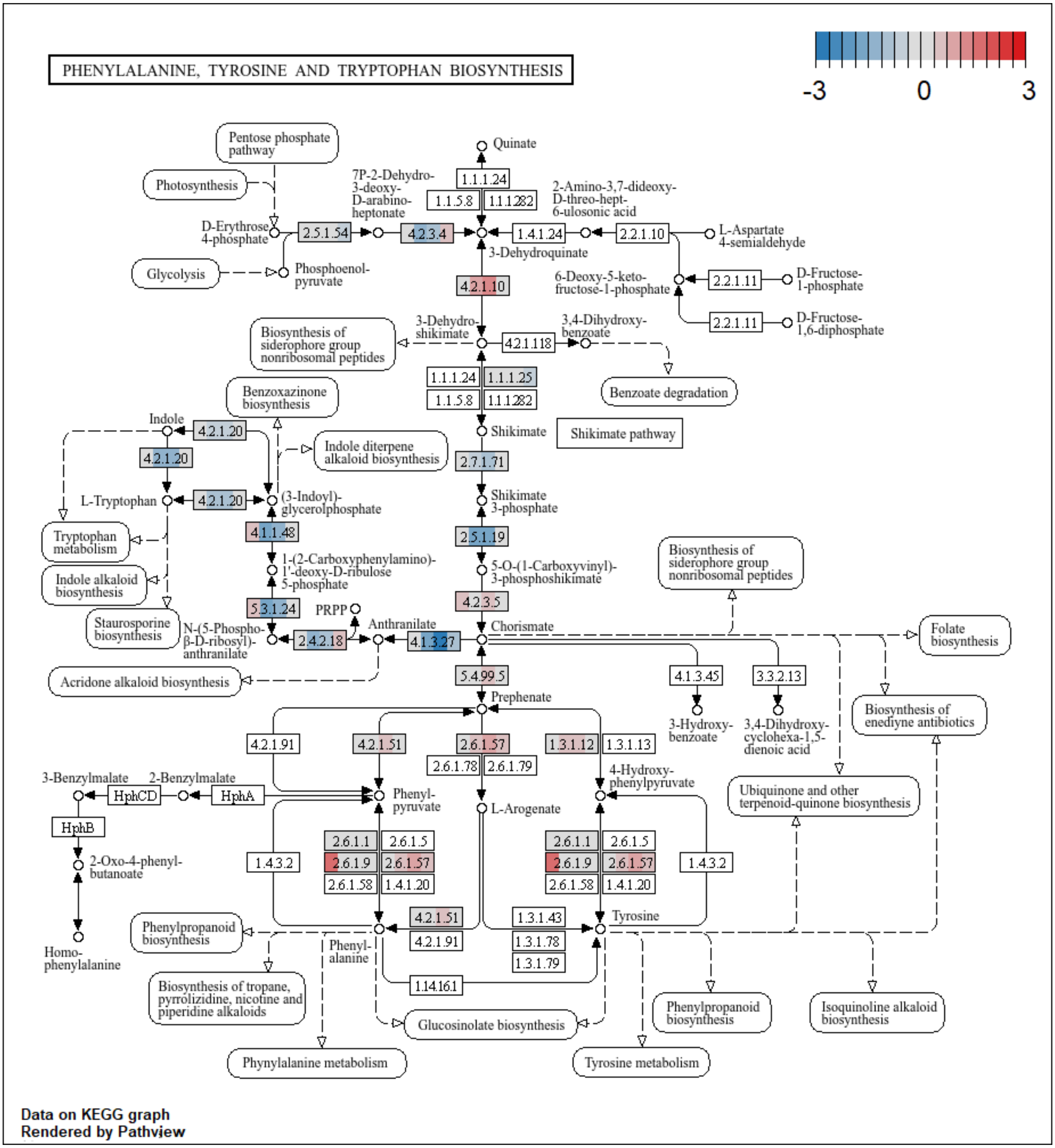
PHENYLALANINE, TYROSINE AND TRYPTOPHAN BIOSYNTHESIS
-3
0
3
Pentose phosphate pathway
Photosynthesis
D-Erythrose 4-phosphate
Glycolysis
Phosphoenol-pyruvate
2.5.1.54
7P-2-Dehydro-3-deoxy-D-arabino-heptonate
4.2.3.4
Quinate
1.1.1.24
1.1.5.8
1.1.1.282
2-Amino-3,7-dideoxy-D-threo-hept-6-ulosonic acid
1.4.1.24
2.2.1.10
L-Aspartate 4-semialdehyde
3-Dehydroquinate
4.2.1.10
6-Deoxy-5-keto-fructose-1-phosphate
2.2.1.11
D-Fructose-1-phosphate
2.2.1.11
D-Fructose-1,6-diphosphate
Biosynthesis of siderophore group nonribosomal peptides
3-Dehydro-shikimate
4.2.1.118
3,4-Dihydroxy-benzoate
Benzoate degradation
1.1.1.24
1.1.1.25
1.1.5.8
1.1.1.282
Benzoxazinone biosynthesis
Indole
4.2.1.20
4.2.1.20
Shikimate
Shikimate pathway
Indole diterpene alkaloid biosynthesis
2.7.1.71
L-Tryptophan
4.2.1.20
(3-Indoyl)-glycerolphosphate
Shikimate 3-phosphate
Tryptophan metabolism
4.1.1.48
2.5.1.19
1-(2-Carboxyphenylamino)-1'-deoxy-D-ribulose 5-phosphate
5-O-(1-Carboxyvinyl)-3-phosphoshikimate
Biosynthesis of siderophore group nonribosomal peptides
Indole alkaloid biosynthesis
5.3.1.24
PRPP
4.2.3.5
Staurosporine biosynthesis
N-(5-Phospho-β-D-ribosyl)-anthranilate
2.4.2.18
Anthranilate
4.1.3.27
Chorismate
Folate biosynthesis
Acridone alkaloid biosynthesis
5.4.99.5
4.1.3.45
3.3.2.13
Prephenate
3-Hydroxy-benzoate
3,4-Dihydroxy-cyclohexa-1,5-dienoic acid
Biosynthesis of enediyne antibiotics
4.2.1.91
4.2.1.51
2.6.1.57
1.3.1.12
1.3.1.13
2.6.1.78
2.6.1.79
3-Benzylmalate
2-Benzylmalate
HphCD
HphA
Phenyl-pyruvate
L-Arogenate
4-Hydroxy-phenylpyruvate
Ubiquinone and other terpenoid-quinone biosynthesis
HphB
2-Oxo-4-phenyl-butanoate
1.4.3.2
2.6.1.1
2.6.1.5
2.6.1.9
2.6.1.57
2.6.1.58
1.4.1.20
2.6.1.1
2.6.1.5
2.6.1.9
2.6.1.57
2.6.1.58
1.4.1.20
1.4.3.2
Homo-phenylalanine
Phenylpropanoid biosynthesis
4.2.1.51
4.2.1.91
Phenyl-alanine
1.3.1.43
1.3.1.78
1.3.1.79
Tyrosine
1.14.16.1
Biosynthesis of tropane, pyrrolizidine, nicotine and piperidine alkaloids
Glucosinolate biosynthesis
Phenylpropanoid biosynthesis
Isoquinoline alkaloid biosynthesis
Phynylalanine metabolism
Tyrosine metabolism
Data on KEGG graph
Rendered by Pathview

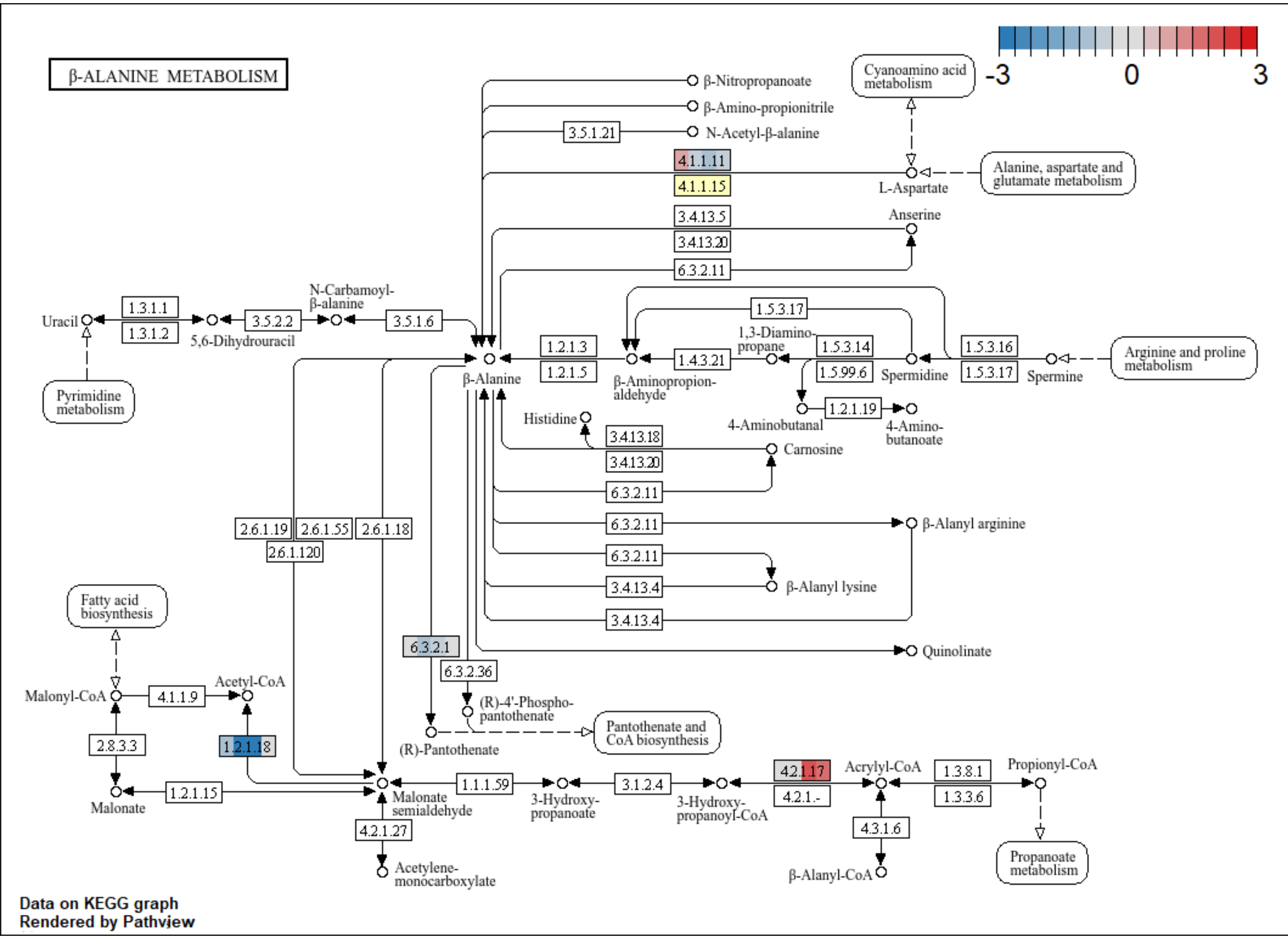

β-ALANINE METABOLISM
-3
0
3
β-Nitropropanoate
β-Amino-propionitrile
N-Acetyl-β-alanine
3.5.1.21
Cyanoamino acid metabolism
4.1.1.11
4.1.1.15
L-Aspartate
Alanine, aspartate and glutamate metabolism
Anserine
3.4.13.5
3.4.13.20
6.3.2.11
Uracil
1.3.1.1
1.3.1.2
5,6-Dihydrouracil
3.5.2.2
N-Carbamoyl-β-alanine
3.5.1.6
Pyrimidine metabolism
β-Alanine
1.2.1.3
1.2.1.5
β-Aminopropion-aldehyde
1.4.3.21
1,3-Diamino-propane
1.5.3.17
1.5.3.14
1.5.99.6
Spermidine
1.5.3.16
1.5.3.17
Spermine
Arginine and proline metabolism
4-Aminobutanal
1.2.1.19
4-Amino-butanoate
Histidine
3.4.13.18
3.4.13.20
Carnosine
6.3.2.11
6.3.2.11
β-Alanyl arginine
6.3.2.11
β-Alanyl lysine
3.4.13.4
3.4.13.4
Quinolinate
2.6.1.19
2.6.1.55
2.6.1.18
2.6.1.120
Fatty acid biosynthesis
6.3.2.1
6.3.2.36
(R)-4'-Phospho-pantothenate
Pantothenate and CoA biosynthesis
(R)-Pantothenate
Malonyl-CoA
4.1.1.9
Acetyl-CoA
2.8.3.3
1.2.1.18
Malonate
1.2.1.15
Malonate semialdehyde
1.1.1.59
3-Hydroxy-propanoate
3.1.2.4
3-Hydroxy-propanoyl-CoA
4.2.1.17
4.2.1.-
Acrylyl-CoA
1.3.8.1
1.3.3.6
Propionyl-CoA
4.3.1.6
β-Alanyl-CoA
Propanoate metabolism
4.2.1.27
Acetylene-monocarboxylate
Data on KEGG graph
Rendered by Pathview

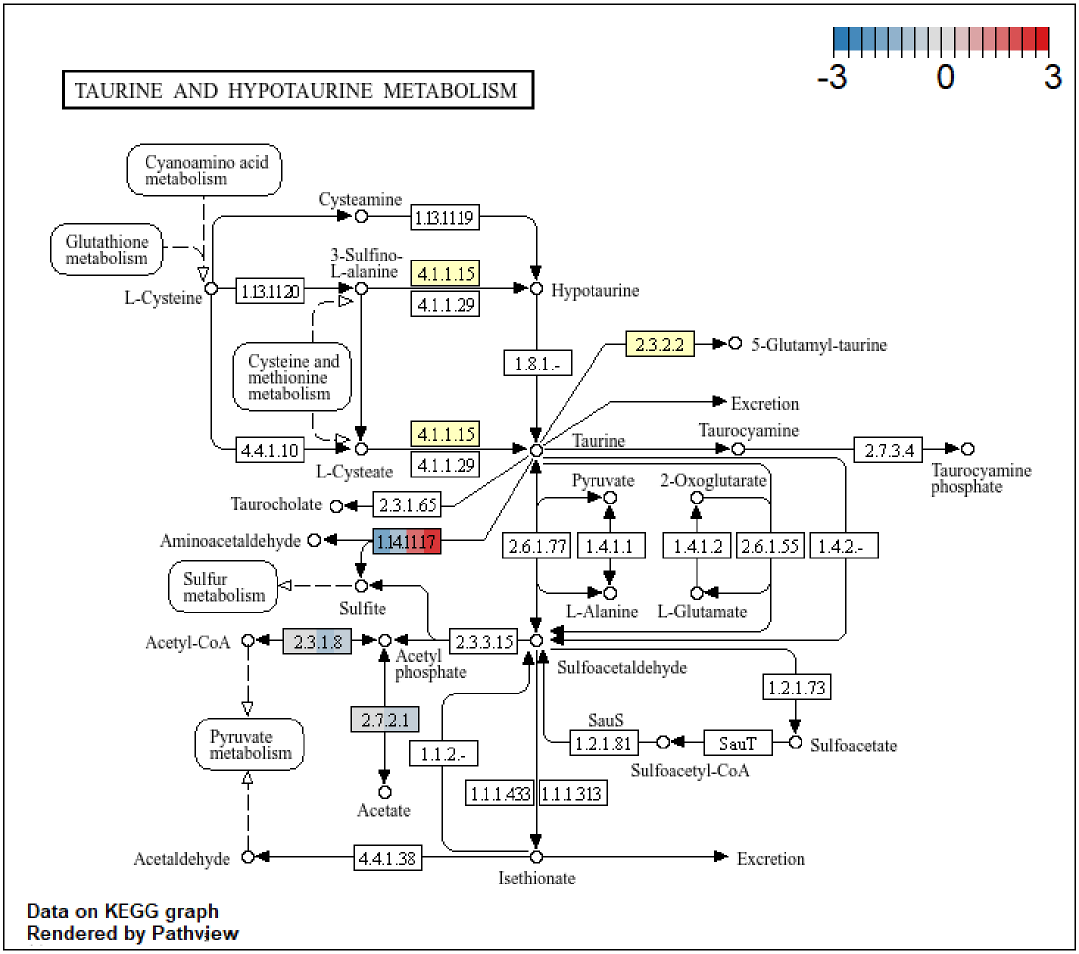

-3
0
3
TAURINE AND HYPOTAURINE METABOLISM
Cyanoamino acid metabolism
Glutathione metabolism
Cysteamine
1.13.11.19
3-Sulfino-L-alanine
4.1.1.15
4.1.1.29
Hypotaurine
L-Cysteine
1.13.11.20
Cysteine and methionine metabolism
1.8.1.-
2.3.2.2
5-Glutamyl-taurine
Excretion
Taurine
Taurocyamine
4.4.1.10
L-Cysteate
2.7.3.4
Taurocyamine phosphate
Taurocholate
2.3.1.65
Pyruvate
2-Oxoglutarate
Aminoacetaldehyde
1.14.11.17
2.6.1.77
1.4.1.1
1.4.1.2
2.6.1.55
1.4.2.-
Sulfur metabolism
Sulfite
L-Alanine
L-Glutamate
Acetyl-CoA
2.3.1.8
Acetyl phosphate
2.3.3.15
Sulfoacetaldehyde
1.2.1.73
Pyruvate metabolism
2.7.2.1
SauS
1.2.1.81
SauT
Sulfoacetate
1.1.2.-
Sulfoacetyl-CoA
Acetate
1.1.1.433
1.1.1.313
Acetaldehyde
4.4.1.38
Excretion
Isethionate
Data on KEGG graph
Rendered by Pathview

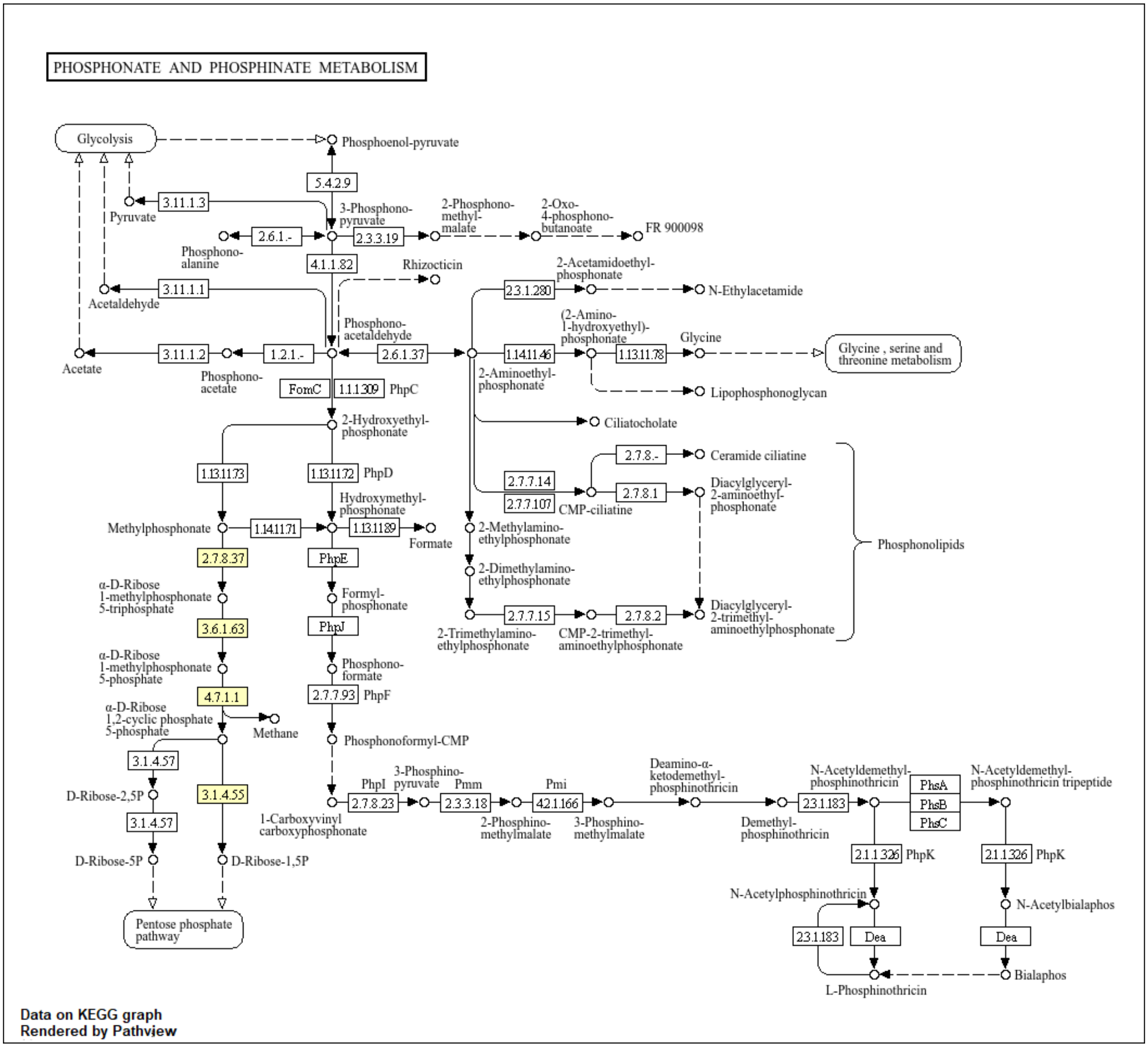

PHOSPHONATE AND PHOSPHINATE METABOLISM
Glycolysis
Phosphoenol-pyruvate
5.4.2.9
3.11.1.3
Pyruvate
3-Phosphono-pyruvate
2-Phosphono-methyl-malate
2-Oxo-4-phosphono-butanoate
FR 900098
2.6.1.-
2.3.3.19
Phosphono-alanine
4.1.1.82
Rhizocticin
3.11.1.1
Acetaldehyde
2-Acetamidoethyl-phosphonate
2.3.1.280
N-Ethylacetamide
Phosphono-acetaldehyde
(2-Amino-1-hydroxyethyl)-phosphonate
Glycine
Glycine , serine and threonine metabolism
3.11.1.2
1.2.1.-
2.6.1.37
1.14.11.46
1.13.11.78
Acetate
Phosphono-acetate
2-Aminoethyl-phosphonate
FomC
1.1.1.309
PhpC
Lipophosphonoglycan
2-Hydroxyethyl-phosphonate
Ciliatocholate
2.7.8.-
Ceramide ciliatine
1.13.11.73
1.13.11.72
PhpD
2.7.7.14
2.7.8.1
Diacylglyceryl-2-aminoethyl-phosphonate
Hydroxymethyl-phosphonate
2.7.7.107
CMP-ciliatine
Methylphosphonate
1.14.11.71
1.13.11.89
Formate
2-Methylamino-ethylphosphonate
Phosphonolipids
2.7.8.37
PhpE
2-Dimethylamino-ethylphosphonate
α-D-Ribose 1-methylphosphonate 5-triphosphate
Formyl-phosphonate
Diacylglyceryl-2-trimethyl-aminoethylphosphonate
3.6.1.63
PhpJ
2.7.7.15
2.7.8.2
2-Trimethylamino-ethylphosphonate
CMP-2-trimethyl-aminoethylphosphonate
α-D-Ribose 1-methylphosphonate 5-phosphate
Phosphono-formate
4.7.1.1
2.7.7.93
PhpF
α-D-Ribose 1,2-cyclic phosphate 5-phosphate
Methane
Phosphonoformyl-CMP
3.1.4.57
3-Phosphino-pyruvate
Deamino-α-ketodemethyl-phosphinothricin
N-Acetyldemethyl-phosphinothricin
N-Acetyldemethyl-phosphinothricin tripeptide
PhpI
Pmm
Pmi
PhsA
D-Ribose-2,5P
3.1.4.55
2.7.8.23
2.3.3.18
4.2.1.166
2.3.1.183
PhsB
PhsC
3.1.4.57
1-Carboxyvinyl carboxyphosphonate
2-Phosphino-methylmalate
3-Phosphino-methylmalate
Demethyl-phosphinothricin
2.1.1.326
PhpK
2.1.1.326
PhpK
D-Ribose-5P
D-Ribose-1,5P
N-Acetylphosphinothricin
N-Acetylbialaphos
Pentose phosphate pathway
2.3.1.183
Dea
Dea
Bialaphos
L-Phosphinothricin
Data on KEGG graph
Rendered by Pathview

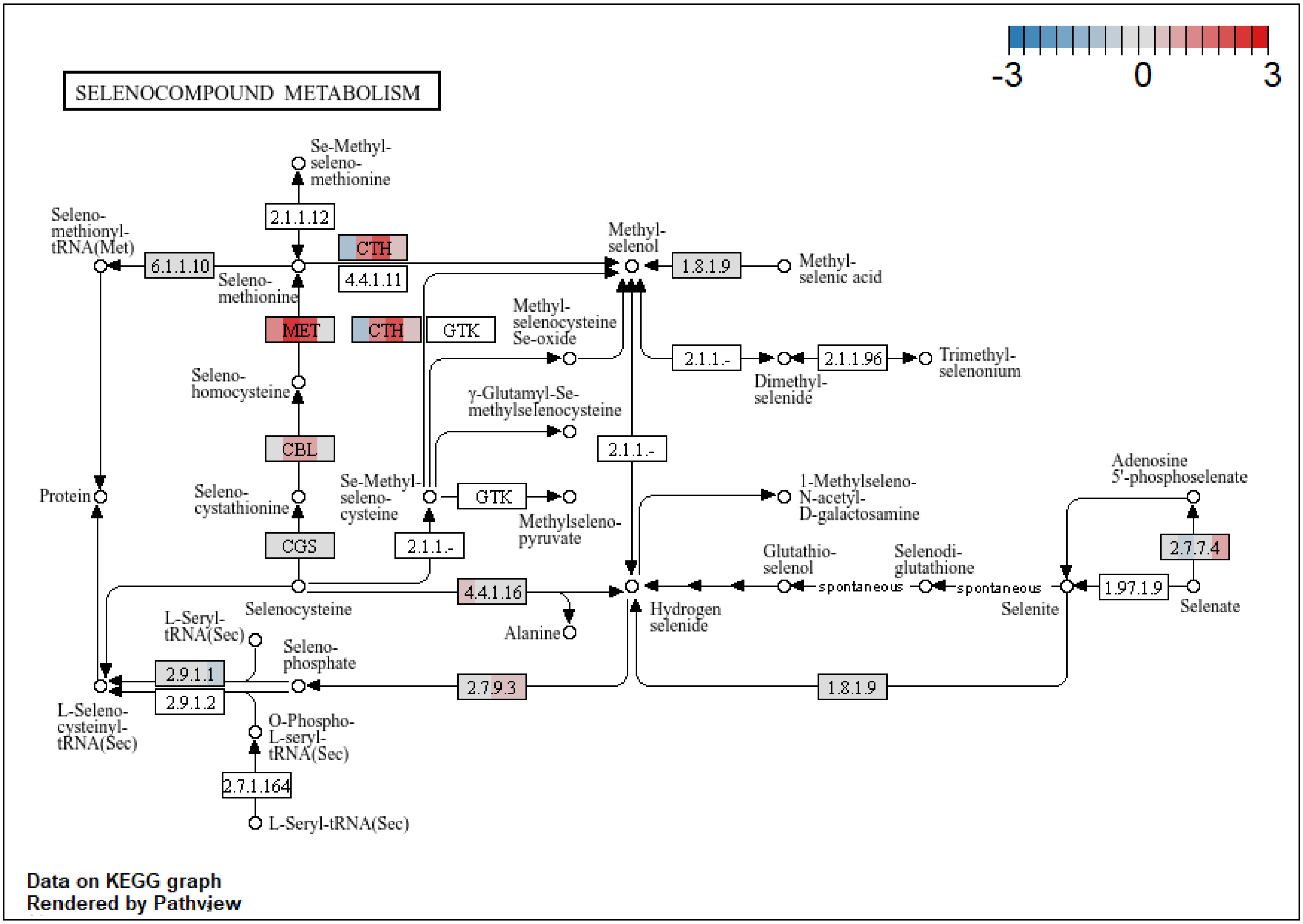
SELENOCOMPOUND METABOLISM
-3
0
3
Se-Methyl-seleno-methionine
2.1.1.12
Seleno-methionyl-tRNA(Met)
6.1.1.10
Seleno-methionine
CTH
4.4.1.11
Methyl-selenol
1.8.1.9
Methyl-selenic acid
MET
CTH
GTK
Methyl-selenocysteine Se-oxide
2.1.1.-
Dimethyl-selenide
2.1.1.96
Trimethyl-selenonium
Seleno-homocysteine
γ-Glutamyl-Se-methylselenocysteine
CBL
2.1.1.-
Protein
Seleno-cystathionine
Se-Methyl-seleno-cysteine
GTK
Methylseleno-pyruvate
1-Methylseleno-N-acetyl-D-galactosamine
Adenosine 5'-phosphoselenate
CGS
2.1.1.-
2.7.7.4
Glutathio-selenol
Selenodi-glutathione
spontaneous
spontaneous
1.97.1.9
Selenocysteine
4.4.1.16
Hydrogen selenide
Selenite
Selenate
Alanine
L-Seryl-tRNA(Sec)
Seleno-phosphate
2.9.1.1
2.9.1.2
2.7.9.3
1.8.1.9
L-Seleno-cysteinyl-tRNA(Sec)
O-Phospho-L-seryl-tRNA(Sec)
2.7.1.164
L-Seryl-tRNA(Sec)
Data on KEGG graph
Rendered by Pathview

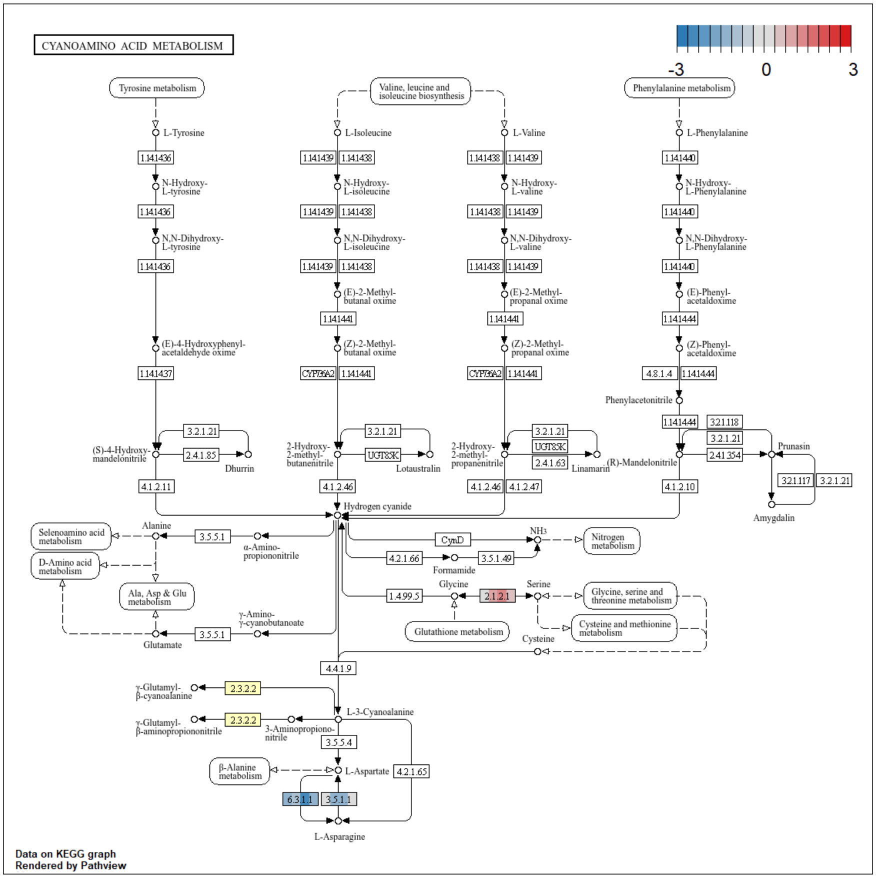

CYANOAMINO ACID METABOLISM
-3
0
3
Tyrosine metabolism
Valine, leucine and isoleucine biosynthesis
Phenylalanine metabolism
L-Tyrosine
L-Isoleucine
L-Valine
L-Phenylalanine
N-Hydroxy-L-tyrosine
N-Hydroxy-L-isoleucine
N-Hydroxy-L-valine
N-Hydroxy-L-Phenylalanine
N,N-Dihydroxy-L-tyrosine
N,N-Dihydroxy-L-isoleucine
N,N-Dihydroxy-L-valine
N,N-Dihydroxy-L-Phenylalanine
(E)-2-Methyl-butanal oxime
(E)-2-Methyl-propanal oxime
(E)-Phenyl-acetaldoxime
(E)-4-Hydroxyphenyl-acetaldehyde oxime
(Z)-2-Methyl-butanal oxime
(Z)-2-Methyl-propanal oxime
(Z)-Phenyl-acetaldoxime
Phenylacetonitrile
(S)-4-Hydroxy-mandelonitrile
Dhurrin
2-Hydroxy-2-methyl-butanenitrile
Lotaustralin
2-Hydroxy-2-methyl-propanenitrile
Linamarin
(R)-Mandelonitrile
Prunasin
Amygdalin
Hydrogen cyanide
Selenoamino acid metabolism
D-Amino acid metabolism
Alanine
α-Amino-propiononitrile
Ala, Asp & Glu metabolism
γ-Amino-γ-cyanobutanoate
Glutamate
CynD
NH3
Nitrogen metabolism
Formamide
Glycine
Serine
Glycine, serine and threonine metabolism
Glutathione metabolism
Cysteine and methionine metabolism
Cysteine
γ-Glutamyl-β-cyanoalanine
γ-Glutamyl-β-aminopropiononitrile
3-Aminopropiono-nitrile
L-3-Cyanoalanine
β-Alanine metabolism
L-Aspartate
L-Asparagine
Data on KEGG graph
Rendered by Pathview

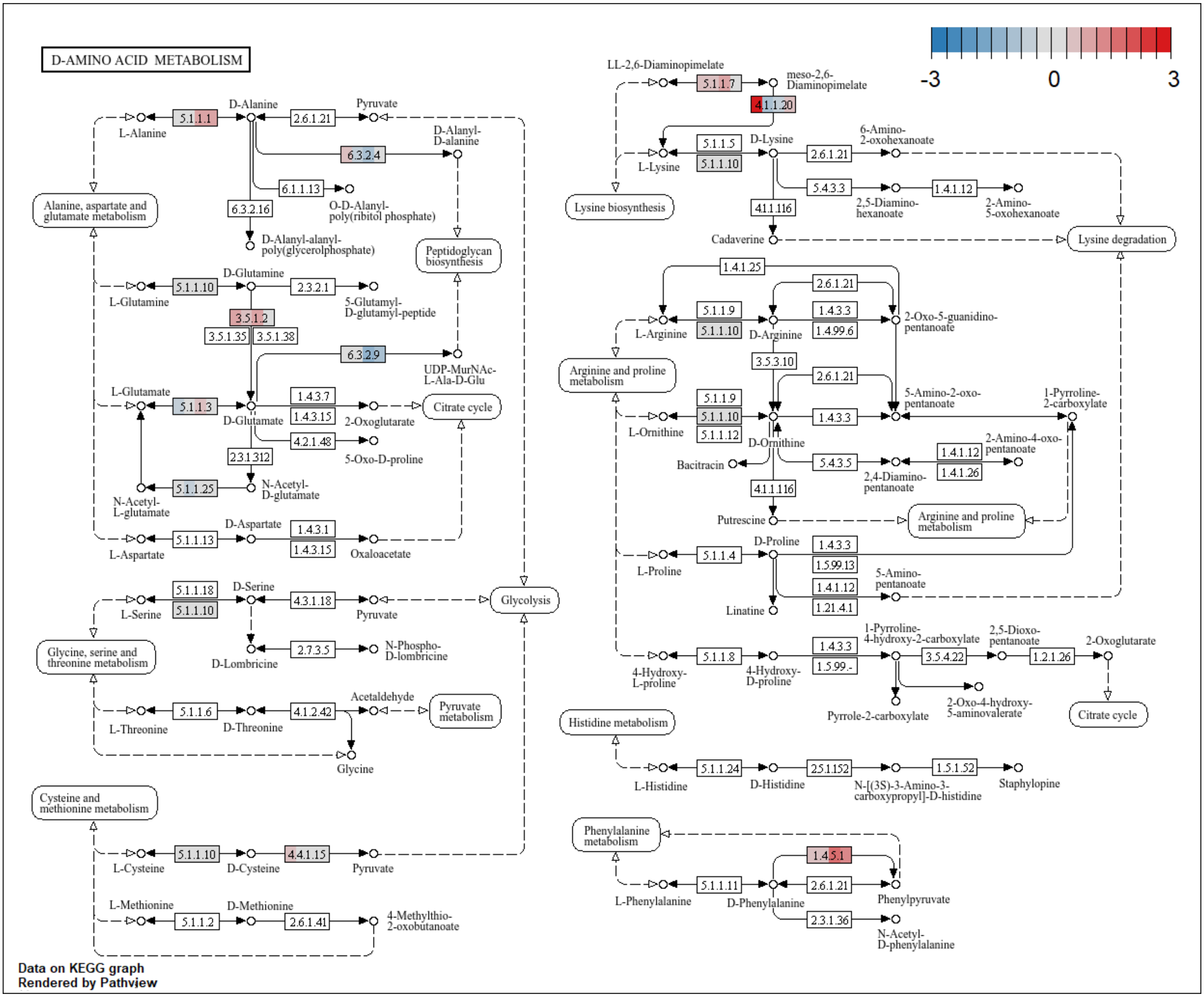
D-AMINO ACID METABOLISM
-3
0
3
L-Alanine
5.1.1.1
D-Alanine
2.6.1.21
Pyruvate
D-Alanyl-D-alanine
6.3.2.4
6.1.1.13
O-D-Alanyl-poly(ribitol phosphate)
6.3.2.16
D-Alanyl-alanyl-poly(glycerolphosphate)
Alanine, aspartate and glutamate metabolism
Peptidoglycan biosynthesis
L-Glutamine
5.1.1.10
D-Glutamine
2.3.2.1
5-Glutamyl-D-glutamyl-peptide
3.5.1.2
3.5.1.35
3.5.1.38
6.3.2.9
UDP-MurNAc-L-Ala-D-Glu
L-Glutamate
5.1.1.3
D-Glutamate
1.4.3.7
1.4.3.15
2-Oxoglutarate
Citrate cycle
4.2.1.48
5-Oxo-D-proline
2.3.1.312
N-Acetyl-L-glutamate
5.1.1.25
N-Acetyl-D-glutamate
L-Aspartate
5.1.1.13
D-Aspartate
1.4.3.1
1.4.3.15
Oxaloacetate
L-Serine
5.1.1.18
5.1.1.10
D-Serine
4.3.1.18
Pyruvate
Glycolysis
Glycine, serine and threonine metabolism
D-Lombricine
2.7.3.5
N-Phospho-D-lombricine
L-Threonine
5.1.1.6
D-Threonine
4.1.2.42
Acetaldehyde
Pyruvate metabolism
Glycine
Cysteine and methionine metabolism
L-Cysteine
5.1.1.10
D-Cysteine
4.4.1.15
Pyruvate
L-Methionine
5.1.1.2
D-Methionine
2.6.1.41
4-Methylthio-2-oxobutanoate
LL-2,6-Diaminopimelate
5.1.1.7
meso-2,6-Diaminopimelate
4.1.1.20
L-Lysine
5.1.1.5
5.1.1.10
D-Lysine
2.6.1.21
6-Amino-2-oxohexanoate
5.4.3.3
2,5-Diamino-hexanoate
1.4.1.12
2-Amino-5-oxohexanoate
Lysine biosynthesis
4.1.1.116
Cadaverine
Lysine degradation
1.4.1.25
2.6.1.21
L-Arginine
5.1.1.9
5.1.1.10
D-Arginine
1.4.3.3
1.4.99.6
2-Oxo-5-guanidino-pentanoate
Arginine and proline metabolism
3.5.3.10
2.6.1.21
L-Ornithine
5.1.1.9
5.1.1.10
5.1.1.12
D-Ornithine
1.4.3.3
5-Amino-2-oxo-pentanoate
1-Pyrroline-2-carboxylate
Bacitracin
5.4.3.5
2,4-Diamino-pentanoate
1.4.1.12
1.4.1.26
2-Amino-4-oxo-pentanoate
4.1.1.116
Putrescine
Arginine and proline metabolism
L-Proline
5.1.1.4
D-Proline
1.4.3.3
1.5.99.13
1.4.1.12
1.21.4.1
5-Amino-pentanoate
Linatine
4-Hydroxy-L-proline
5.1.1.8
4-Hydroxy-D-proline
1.4.3.3
1.5.99.-
1-Pyrroline-4-hydroxy-2-carboxylate
3.5.4.22
2,5-Dioxo-pentanoate
1.2.1.26
2-Oxoglutarate
Pyrrole-2-carboxylate
2-Oxo-4-hydroxy-5-aminovalerate
Citrate cycle
Histidine metabolism
L-Histidine
5.1.1.24
D-Histidine
2.5.1.152
N-[(3S)-3-Amino-3-carboxypropyl]-D-histidine
1.5.1.52
Staphylopine
Phenylalanine metabolism
1.4.5.1
L-Phenylalanine
5.1.1.11
D-Phenylalanine
2.6.1.21
Phenylpyruvate
2.3.1.36
N-Acetyl-D-phenylalanine
Data on KEGG graph
Rendered by Pathview

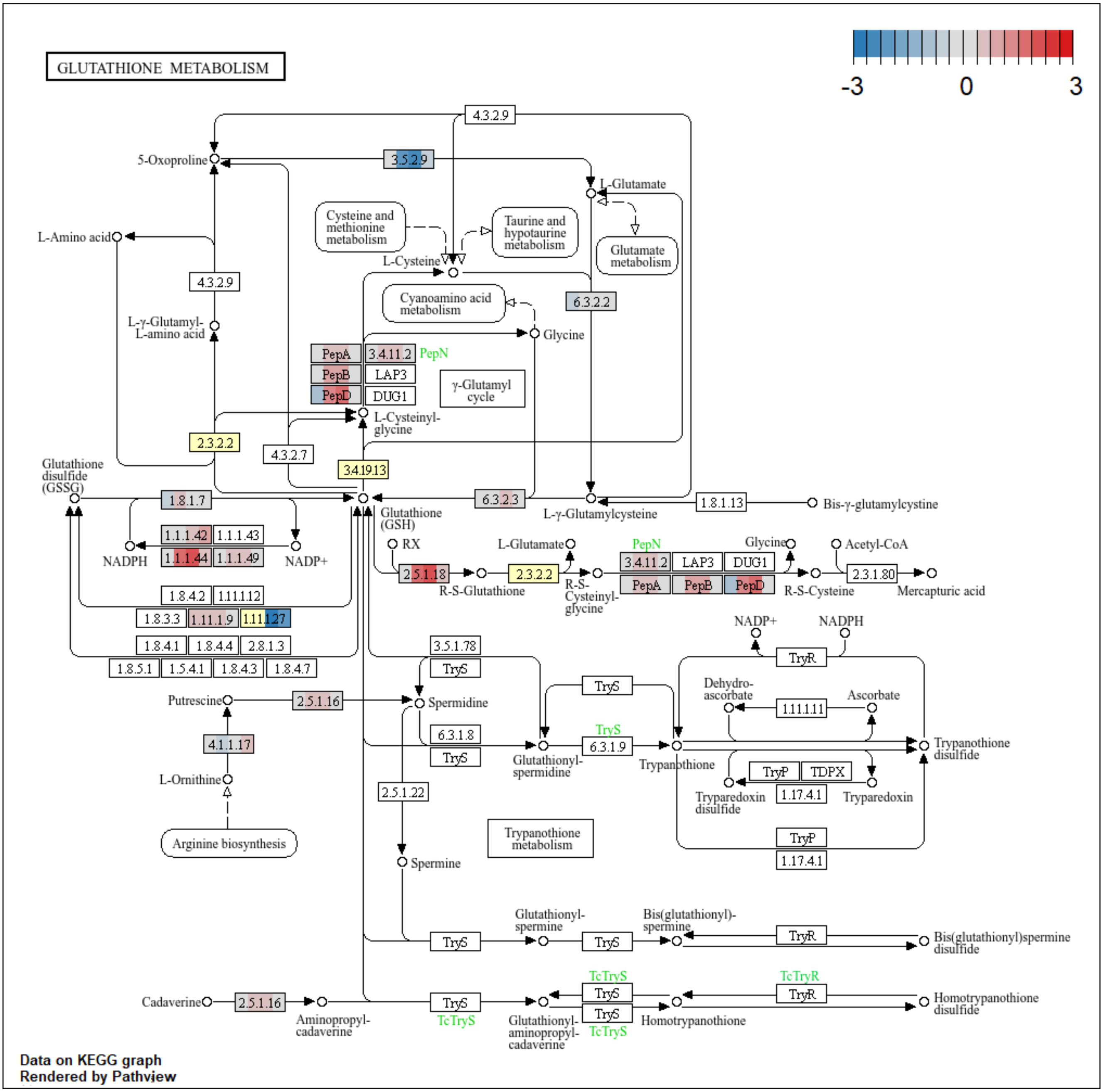
GLUTATHIONE METABOLISM
-3
0
3
4.3.2.9
5-Oxoproline
3.5.2.9
L-Glutamate
Cysteine and methionine metabolism
Taurine and hypotaurine metabolism
Glutamate metabolism
L-Amino acid
L-Cysteine
4.3.2.9
Cyanoamino acid metabolism
6.3.2.2
L-γ-Glutamyl-L-amino acid
Glycine
PepA
3.4.11.2
PepN
PepB
LAP3
PepD
DUG1
γ-Glutamyl cycle
L-Cysteinyl-glycine
2.3.2.2
4.3.2.7
3.4.19.13
Glutathione disulfide (GSSG)
1.8.1.7
6.3.2.3
L-γ-Glutamylcysteine
1.8.1.13
Bis-γ-glutamylcystine
Glutathione (GSH)
1.1.1.42
1.1.1.43
NADPH
1.1.1.44
1.1.1.49
NADP+
RX
L-Glutamate
PepN
Glycine
Acetyl-CoA
2.5.1.18
2.3.2.2
3.4.11.2
LAP3
DUG1
2.3.1.80
R-S-Glutathione
R-S-Cysteinyl-glycine
PepA
PepB
PepD
R-S-Cysteine
Mercapturic acid
1.8.4.2
1.11.1.12
1.8.3.3
1.11.1.9
1.11.1.27
NADP+
NADPH
1.8.4.1
1.8.4.4
2.8.1.3
3.5.1.78
TryR
1.8.5.1
1.5.4.1
1.8.4.3
1.8.4.7
TryS
TryS
Dehydro-ascorbate
Putrescine
2.5.1.16
Spermidine
Ascorbate
1.11.1.11
TryS
4.1.1.17
6.3.1.8
6.3.1.9
Trypanothione disulfide
TryS
Glutathionyl-spermidine
Trypanothione
L-Ornithine
TryP
TDPX
2.5.1.22
Trypanoredoxin disulfide
1.17.4.1
Trypanoredoxin
Arginine biosynthesis
Trypanothione metabolism
TryP
1.17.4.1
Spermine
Glutathionyl-spermine
Bis(glutathionyl)-spermine
TryS
TryS
TryR
Bis(glutathionyl)spermine disulfide
TcTryS
TcTryR
TryS
TryR
Cadaverine
2.5.1.16
TryS
TryS
Homotrypanothione disulfide
Aminopropyl-cadaverine
TcTryS
Glutathionyl-aminopropyl-cadaverine
TcTryS
Homotrypanothione
Data on KEGG graph
Rendered by Pathview

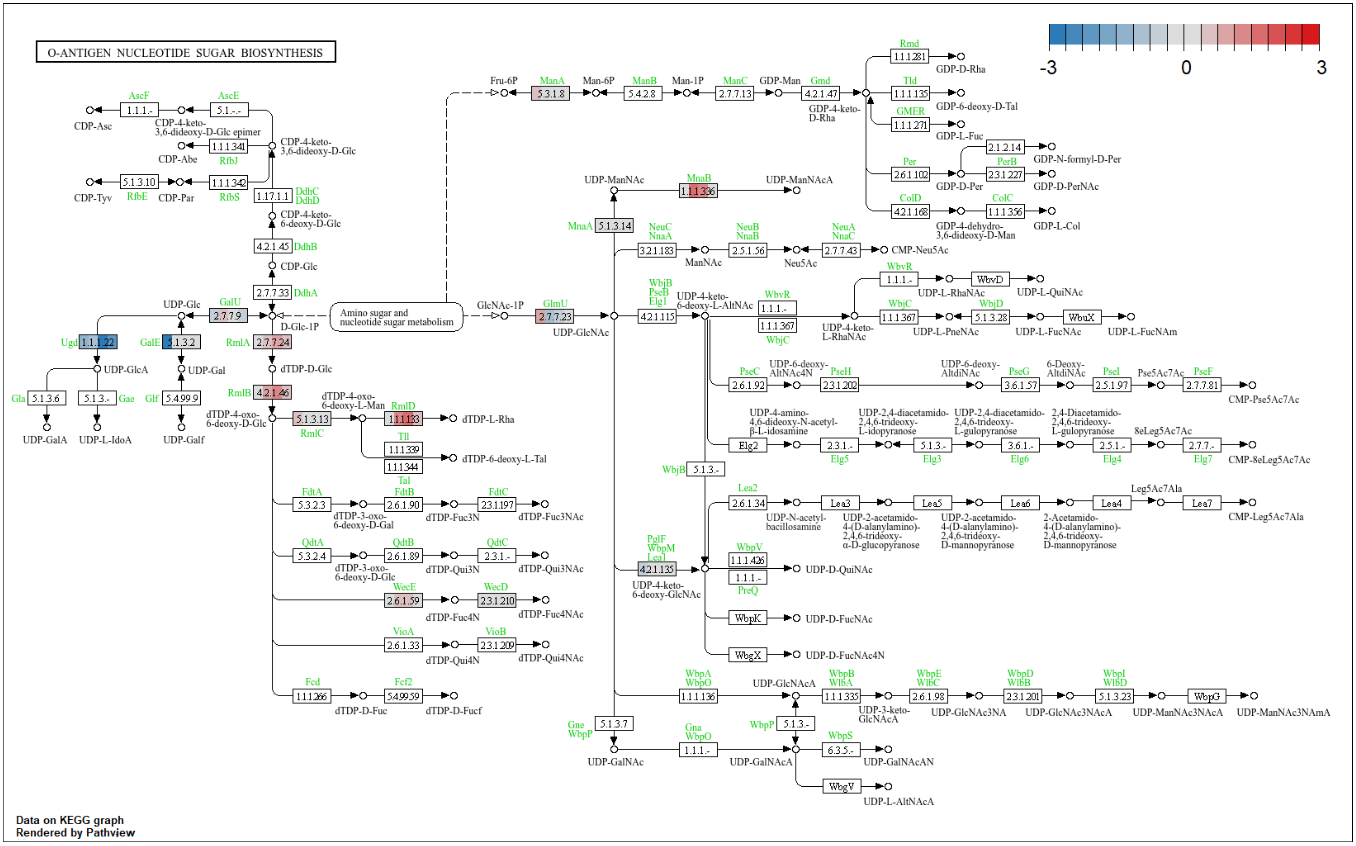
O-ANTIGEN NUCLEOTIDE SUGAR BIOSYNTHESIS
-3
0
3
Amino sugar and nucleotide sugar metabolism
Fru-6P
Man-6P
Man-1P
GDP-Man
GDP-4-keto-D-Rha
GDP-D-Rha
GDP-6-deoxy-D-Tal
GDP-L-Fuc
GDP-D-Per
GDP-N-formyl-D-Per
GDP-D-PerNAc
GDP-4-dehydro-3,6-dideoxy-D-Man
GDP-L-Col
CDP-Asc
CDP-4-keto-3,6-dideoxy-D-Glc epimer
CDP-Abe
CDP-Tyv
CDP-Par
CDP-4-keto-3,6-dideoxy-D-Glc
CDP-4-keto-6-deoxy-D-Glc
CDP-Glc
UDP-Glc
D-Glc-1P
UDP-GlcA
UDP-Gal
UDP-GalA
UDP-L-IdoA
UDP-Galf
dTDP-D-Glc
dTDP-4-oxo-6-deoxy-D-Glc
dTDP-4-oxo-6-deoxy-L-Man
dTDP-L-Rha
dTDP-6-deoxy-L-Tal
dTDP-3-oxo-6-deoxy-D-Gal
dTDP-Fuc3N
dTDP-Fuc3NAc
dTDP-3-oxo-6-deoxy-D-Glc
dTDP-Qui3N
dTDP-Qui3NAc
dTDP-Fuc4N
dTDP-Fuc4NAc
dTDP-Qui4N
dTDP-Qui4NAc
dTDP-D-Fuc
dTDP-D-Fucf
GlcNAc-1P
UDP-GlcNAc
UDP-ManNAc
UDP-ManNAcA
ManNAc
Neu5Ac
CMP-Neu5Ac
UDP-4-keto-6-deoxy-L-AltNAc
UDP-4-keto-L-RhaNAc
UDP-L-RhaNAc
UDP-L-QuiNAc
UDP-L-PneNAc
UDP-L-FucNAc
UDP-L-FucNAm
UDP-6-deoxy-AltNAc4N
UDP-6-deoxy-AltdiNAc
6-Deoxy-AltdiNAc
Pse5Ac7Ac
CMP-Pse5Ac7Ac
UDP-4-amino-4,6-dideoxy-N-acetyl-β-L-idosamine
UDP-2,4-diacetamido-2,4,6-trideoxy-L-idopyranose
UDP-2,4-diacetamido-2,4,6-trideoxy-L-gulopyranose
2,4-Diacetamido-2,4,6-trideoxy-L-gulopyranose
8eLeg5Ac7Ac
CMP-8eLeg5Ac7Ac
UDP-N-acetyl-bacillosamine
UDP-2-acetamido-4-(D-alanylamino)-2,4,6-trideoxy-α-D-glucopyranose
UDP-2-acetamido-4-(D-alanylamino)-2,4,6-trideoxy-D-mannopyranose
2-Acetamido-4-(D-alanylamino)-2,4,6-trideoxy-D-mannopyranose
Leg5Ac7Ala
CMP-Leg5Ac7Ala
UDP-4-keto-6-deoxy-GlcNAc
UDP-D-QuiNAc
UDP-D-FucNAc
UDP-D-FucNAc4N
UDP-GlcNAcA
UDP-3-keto-GlcNAcA
UDP-GlcNAc3NA
UDP-GlcNAc3NAcA
UDP-ManNAc3NAcA
UDP-ManNAc3NAmA
UDP-GalNAc
UDP-GalNAcA
UDP-GalNAcAN
UDP-L-AltNAcA
Data on KEGG graph
Rendered by Pathview

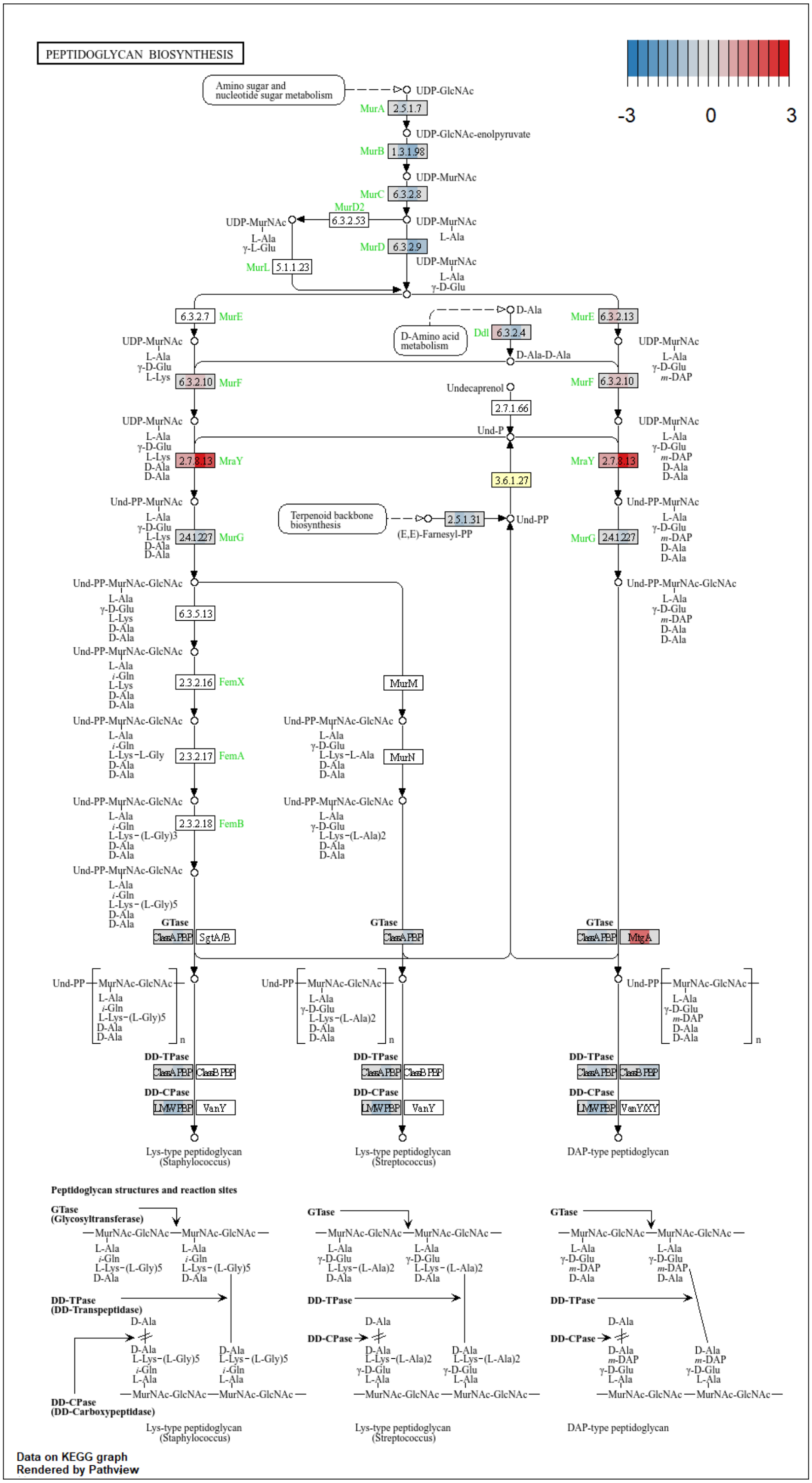
PEPTIDOGLYCAN BIOSYNTHESIS
-3
0
3
Amino sugar and nucleotide sugar metabolism
UDP-GlcNAc
MurA 2.5.1.7
UDP-GlcNAc-enolpyruvate
MurB 1.3.1.98
UDP-MurNAc
MurC 6.3.2.8
MurD2 6.3.2.53
MurD 6.3.2.9
MurL 5.1.1.23
MurE 6.3.2.7
MurE 6.3.2.13
D-Amino acid metabolism
D-Ala
Ddl 6.3.2.4
D-Ala-D-Ala
MurF 6.3.2.10
Undecaprenol
2.7.1.66
Und-P
MraY 2.7.8.13
3.6.1.27
Terpenoid backbone biosynthesis
2.5.1.31
(E,E)-Farnesyl-PP
Und-PP
MurG 2.4.1.227
Und-PP-MurNAc-GlcNAc
6.3.5.13
2.3.2.16 FemX
MurM
2.3.2.17 FemA
MurN
2.3.2.18 FemB
GTase
ClassA PBP
SgtA/B
MtgA
DD-TPase
ClassB PBP
DD-CPase
LMW PBP
VanY
VanY/X/Y
Lys-type peptidoglycan (Staphylococcus)
Lys-type peptidoglycan (Streptococcus)
DAP-type peptidoglycan
Peptidoglycan structures and reaction sites
GTase (Glycosyltransferase)
DD-TPase (DD-Transpeptidase)
DD-CPase (DD-Carboxypeptidase)
Data on KEGG graph
Rendered by Pathview

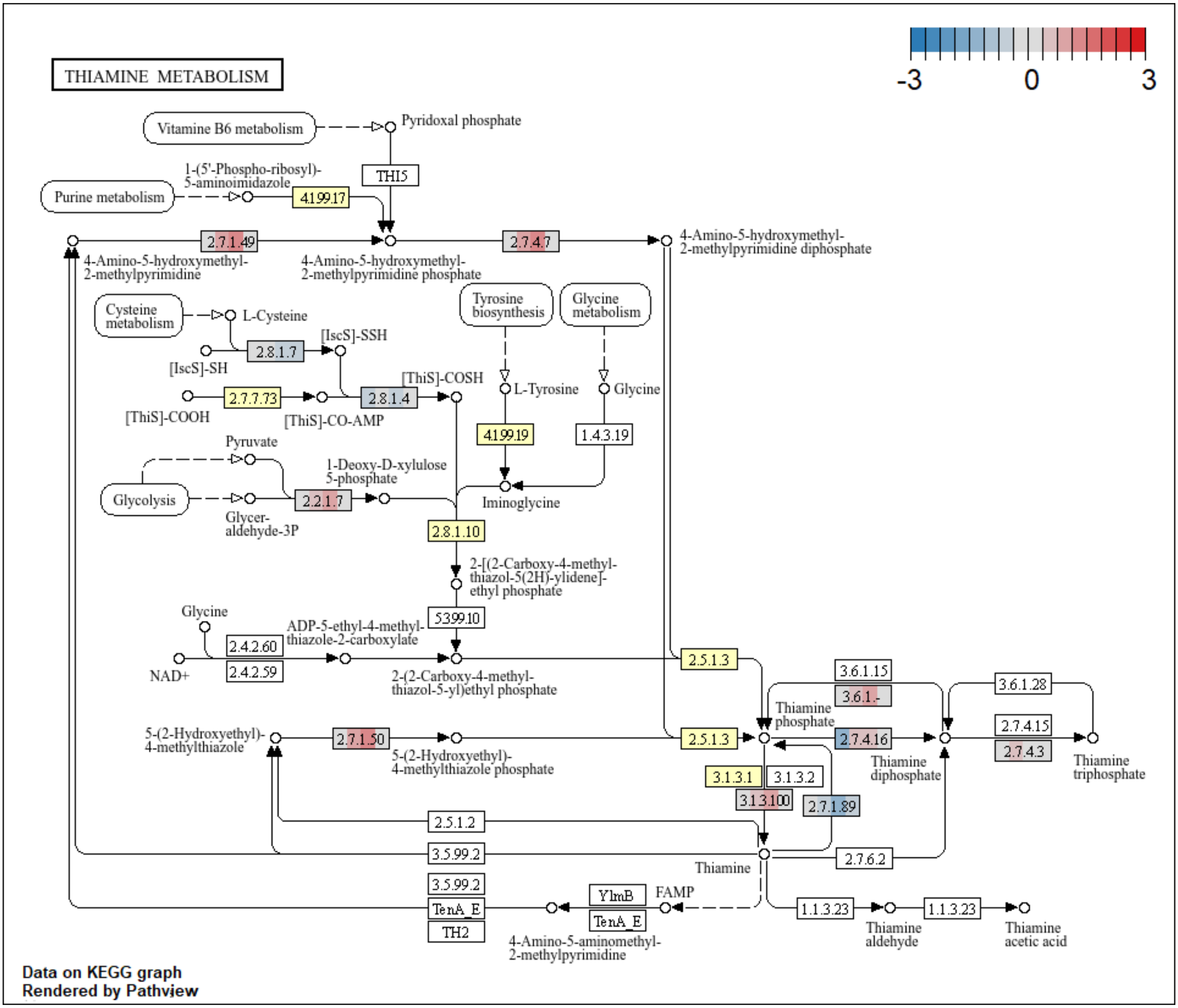
THIAMINE METABOLISM
-3
0
3
Vitamine B6 metabolism
Pyridoxal phosphate
THI5
Purine metabolism
1-(5'-Phospho-ribosyl)-5-aminoimidazole
4.1.99.17
2.7.1.49
2.7.4.7
4-Amino-5-hydroxymethyl-2-methylpyrimidine
4-Amino-5-hydroxymethyl-2-methylpyrimidine phosphate
4-Amino-5-hydroxymethyl-2-methylpyrimidine diphosphate
Cysteine metabolism
L-Cysteine
[IscS]-SSH
2.8.1.7
[IscS]-SH
Tyrosine biosynthesis
Glycine metabolism
[ThiS]-COSH
2.7.7.73
2.8.1.4
[ThiS]-COOH
[ThiS]-CO-AMP
L-Tyrosine
Glycine
4.1.99.19
1.4.3.19
Pyruvate
1-Deoxy-D-xylulose 5-phosphate
Glycolysis
2.2.1.7
Glyceraldehyde-3P
Iminoglycine
2.8.1.10
2-[(2-Carboxy-4-methyl-thiazol-5(2H)-ylidene]-ethyl phosphate
5.3.99.10
Glycine
2.4.2.60
ADP-5-ethyl-4-methyl-thiazole-2-carboxylate
NAD+
2.4.2.59
2-(2-Carboxy-4-methyl-thiazol-5-yl)ethyl phosphate
2.5.1.3
3.6.1.15
3.6.1.-
3.6.1.28
Thiamine phosphate
5-(2-Hydroxyethyl)-4-methylthiazole
2.7.1.50
2.5.1.3
2.7.4.16
2.7.4.15
2.7.4.3
5-(2-Hydroxyethyl)-4-methylthiazole phosphate
Thiamine diphosphate
Thiamine triphosphate
3.1.3.1
3.1.3.2
3.1.3.100
2.7.1.89
2.5.1.2
3.5.99.2
2.7.6.2
Thiamine
3.5.99.2
YlmB
FAMP
TenA_E
TenA_E
TH2
1.1.3.23
1.1.3.23
4-Amino-5-aminomethyl-2-methylpyrimidine
Thiamine aldehyde
Thiamine acetic acid
Data on KEGG graph
Rendered by Pathview

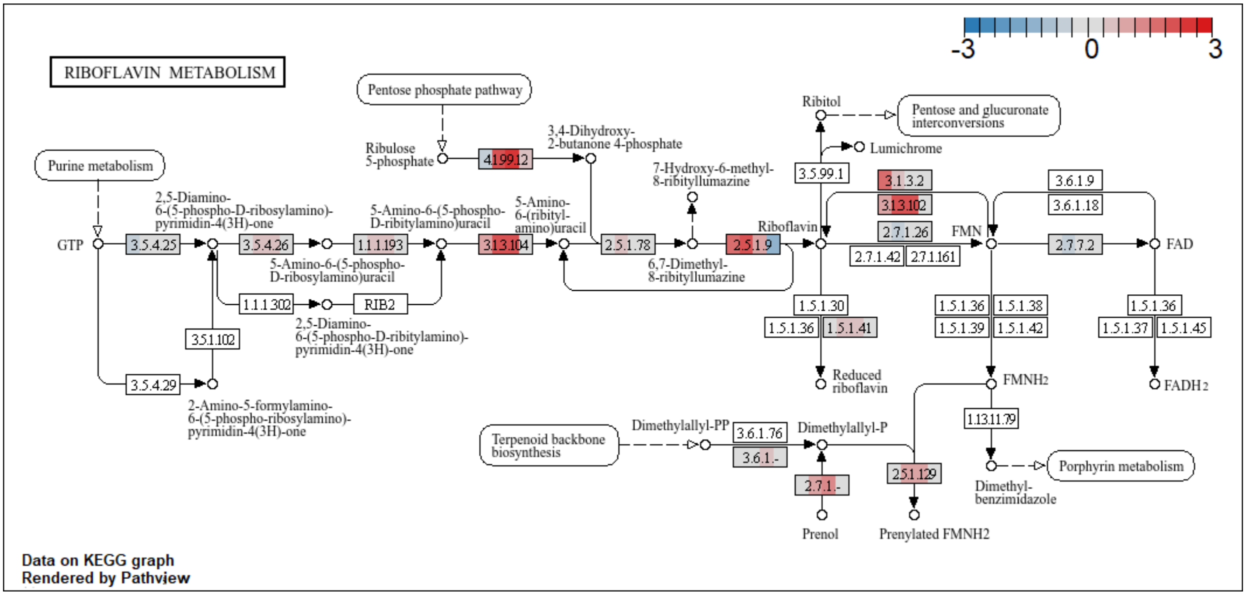
-3
0
3
RIBOFLAVIN METABOLISM
Pentose phosphate pathway
Ribulose 5-phosphate
4.1.99.12
3,4-Dihydroxy-2-butanone 4-phosphate
Purine metabolism
GTP
3.5.4.25
2,5-Diamino-6-(5-phospho-D-ribosylamino)-pyrimidin-4(3H)-one
3.5.4.26
5-Amino-6-(5-phospho-D-ribosylamino)uracil
1.1.1.193
5-Amino-6-(5-phospho-D-ribitylamino)uracil
3.1.3.104
5-Amino-6-(ribitylamino)uracil
2.5.1.78
7-Hydroxy-6-methyl-8-ribityllumazine
6,7-Dimethyl-8-ribityllumazine
2.5.1.9
Riboflavin
Ribitol
Pentose and glucuronate interconversions
Lumichrome
3.5.99.1
3.1.3.2
3.1.3.102
2.7.1.26
FMN
2.7.1.42
2.7.1.161
3.6.1.9
3.6.1.18
2.7.7.2
FAD
1.1.1.302
RIB2
2,5-Diamino-6-(5-phospho-D-ribitylamino)-pyrimidin-4(3H)-one
3.5.1.102
3.5.4.29
2-Amino-5-formylamino-6-(5-phospho-ribosylamino)-pyrimidin-4(3H)-one
1.5.1.30
1.5.1.36
1.5.1.41
Reduced riboflavin
1.5.1.36
1.5.1.38
1.5.1.39
1.5.1.42
FMNH2
1.5.1.36
1.5.1.37
1.5.1.45
FADH2
1.13.11.79
Dimethyl-benzimidazole
Porphyrin metabolism
Terpenoid backbone biosynthesis
Dimethylallyl-PP
3.6.1.76
3.6.1.-
Dimethylallyl-P
2.7.1.-
Prenol
2.5.1.129
Prenylated FMNH2
Data on KEGG graph
Rendered by Pathview

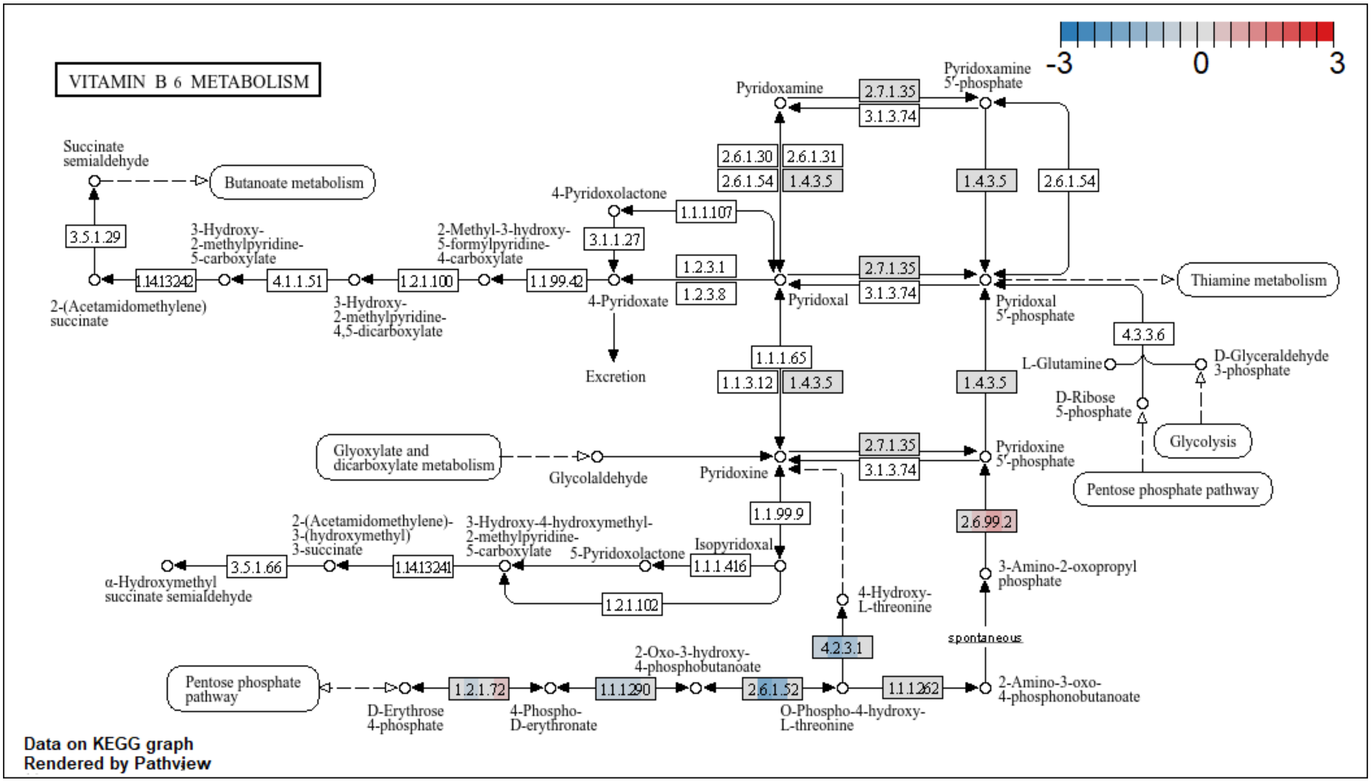
VITAMIN B 6 METABOLISM
-3
0
3
Pyridoxamine
Pyridoxamine 5'-phosphate
2.7.1.35
3.1.3.74
2.6.1.30
2.6.1.31
2.6.1.54
1.4.3.5
1.4.3.5
2.6.1.54
Succinate semialdehyde
Butanoate metabolism
4-Pyridoxolactone
1.1.1.107
3.5.1.29
3-Hydroxy-2-methylpyridine-5-carboxylate
2-Methyl-3-hydroxy-5-formylpyridine-4-carboxylate
3.1.1.27
1.2.3.1
1.2.3.8
2.7.1.35
3.1.3.74
Thiamine metabolism
1.14.13.242
4.1.1.51
1.2.1.100
1.1.99.42
4-Pyridoxate
Pyridoxal
Pyridoxal 5'-phosphate
2-(Acetamidomethylene) succinate
3-Hydroxy-2-methylpyridine-4,5-dicarboxylate
4.3.3.6
1.1.1.65
L-Glutamine
D-Glyceraldehyde 3-phosphate
Excretion
1.1.3.12
1.4.3.5
1.4.3.5
D-Ribose 5-phosphate
Glyoxylate and dicarboxylate metabolism
2.7.1.35
3.1.3.74
Glycolysis
Pyridoxine 5'-phosphate
Glycolaldehyde
Pyridoxine
Pentose phosphate pathway
1.1.99.9
2.6.99.2
2-(Acetamidomethylene)-3-(hydroxymethyl)-3-succinate
3-Hydroxy-4-hydroxymethyl-2-methylpyridine-5-carboxylate
5-Pyridoxolactone
Isopyridoxal
3.5.1.66
1.14.13.241
1.1.1.416
3-Amino-2-oxopropyl phosphate
α-Hydroxymethyl succinate semialdehyde
1.2.1.102
4-Hydroxy-L-threonine
spontaneous
4.2.3.1
2-Oxo-3-hydroxy-4-phosphobutanoate
Pentose phosphate pathway
1.2.1.72
1.1.1.290
2.6.1.52
1.1.1.262
2-Amino-3-oxo-4-phosphonobutanoate
D-Erythrose 4-phosphate
4-Phospho-D-erythronate
O-Phospho-4-hydroxy-L-threonine
Data on KEGG graph
Rendered by Pathview

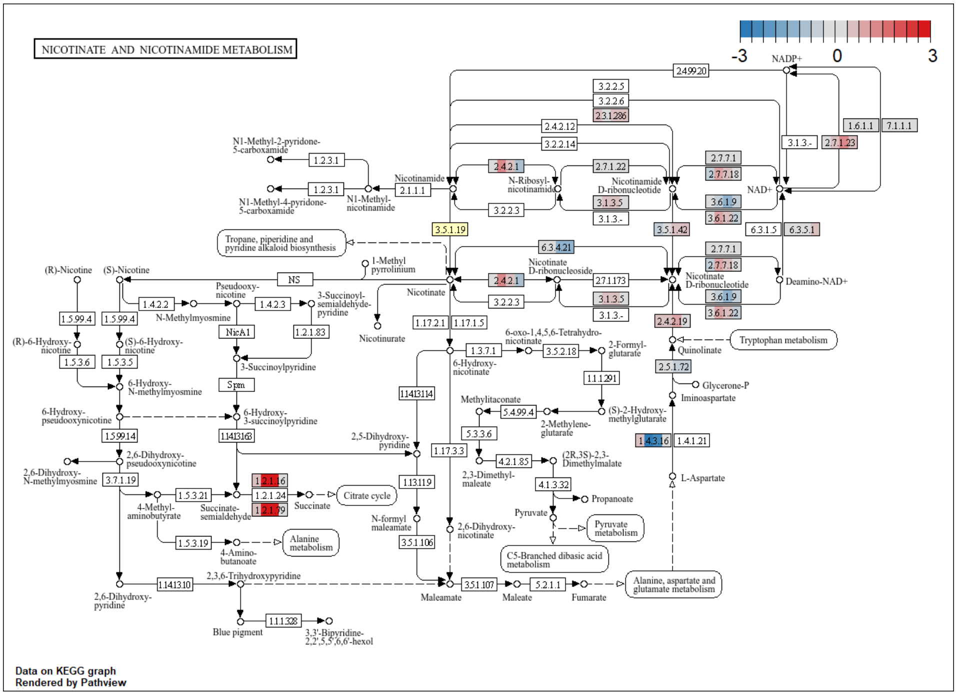
NICOTINATE AND NICOTINAMIDE METABOLISM
-3
0
3
Data on KEGG graph
Rendered by Pathview

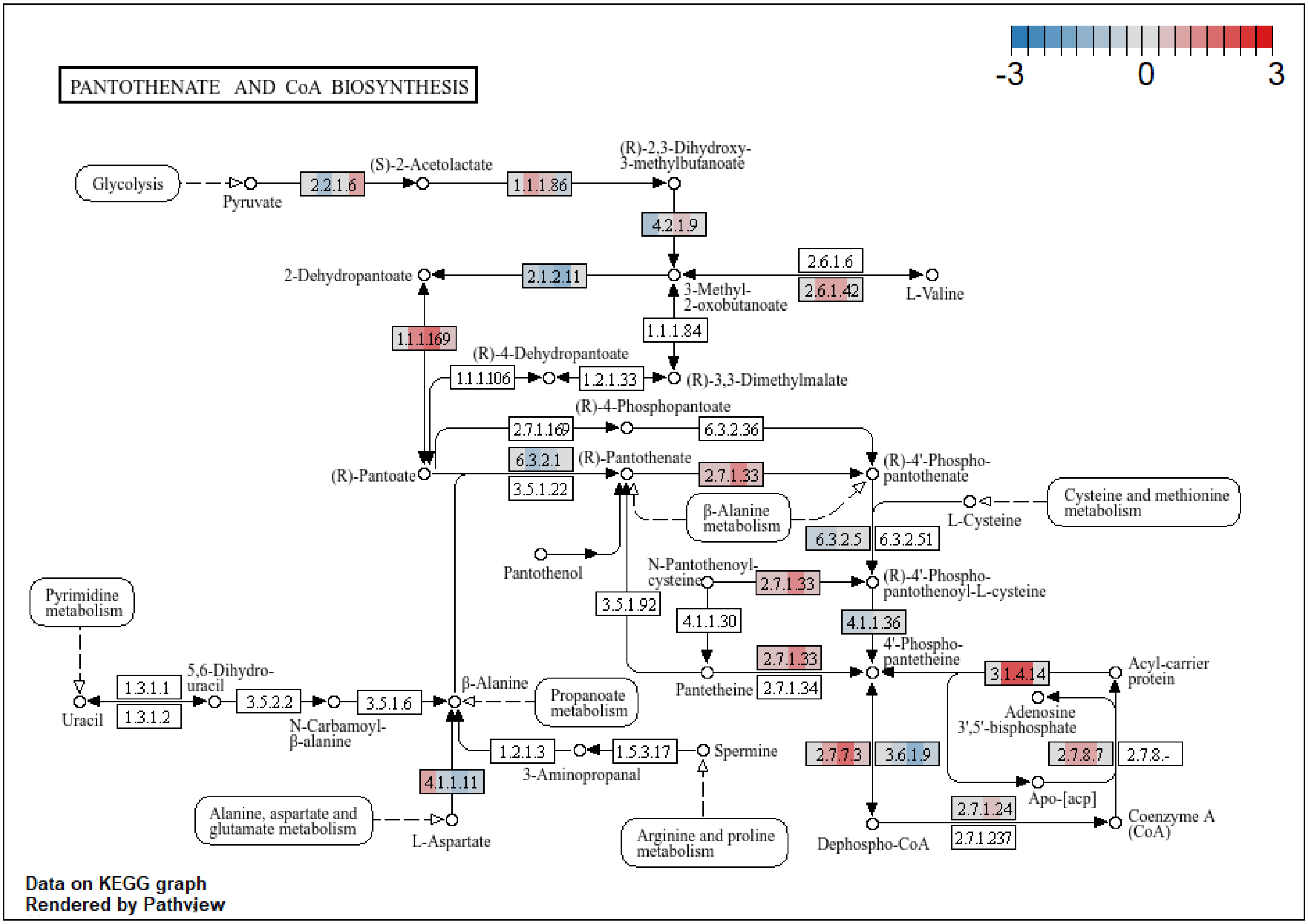
PANTOTHENATE AND CoA BIOSYNTHESIS
-3
0
3
Glycolysis
Pyruvate
2.2.1.6
(S)-2-Acetolactate
1.1.1.86
(R)-2,3-Dihydroxy-3-methylbutanoate
4.2.1.9
2-Dehydropantoate
2.1.2.11
2.6.1.6
2.6.1.42
L-Valine
3-Methyl-2-oxobutanoate
1.1.1.169
1.1.1.84
(R)-4-Dehydropantoate
1.1.1.106
1.2.1.33
(R)-3,3-Dimethylmalate
(R)-4-Phosphopantoate
2.7.1.169
6.3.2.36
6.3.2.1
(R)-Pantothenate
(R)-Pantoate
3.5.1.22
2.7.1.33
(R)-4'-Phosphopantothenate
β-Alanine metabolism
L-Cysteine
Cysteine and methionine metabolism
6.3.2.5
6.3.2.51
Pantothenol
N-Pantothenoyl-cysteine
2.7.1.33
(R)-4'-Phospho-pantothenoyl-L-cysteine
Pyrimidine metabolism
3.5.1.92
4.1.1.30
4.1.1.36
4'-Phospho-pantetheine
2.7.1.33
5,6-Dihydro-uracil
1.3.1.1
1.3.1.2
Uracil
3.5.2.2
N-Carbamoyl-β-alanine
3.5.1.6
β-Alanine
Propanoate metabolism
Pantetheine
2.7.1.34
3.1.4.14
Acyl-carrier protein
Adenosine 3',5'-bisphosphate
1.2.1.3
1.5.3.17
Spermine
2.7.7.3
3.6.1.9
2.7.8.7
2.7.8.-
3-Aminopropanal
4.1.1.11
Apo-[acp]
Alanine, aspartate and glutamate metabolism
L-Aspartate
Arginine and proline metabolism
2.7.1.24
2.7.1.237
Dephospho-CoA
Coenzyme A (CoA)
Data on KEGG graph
Rendered by Pathview

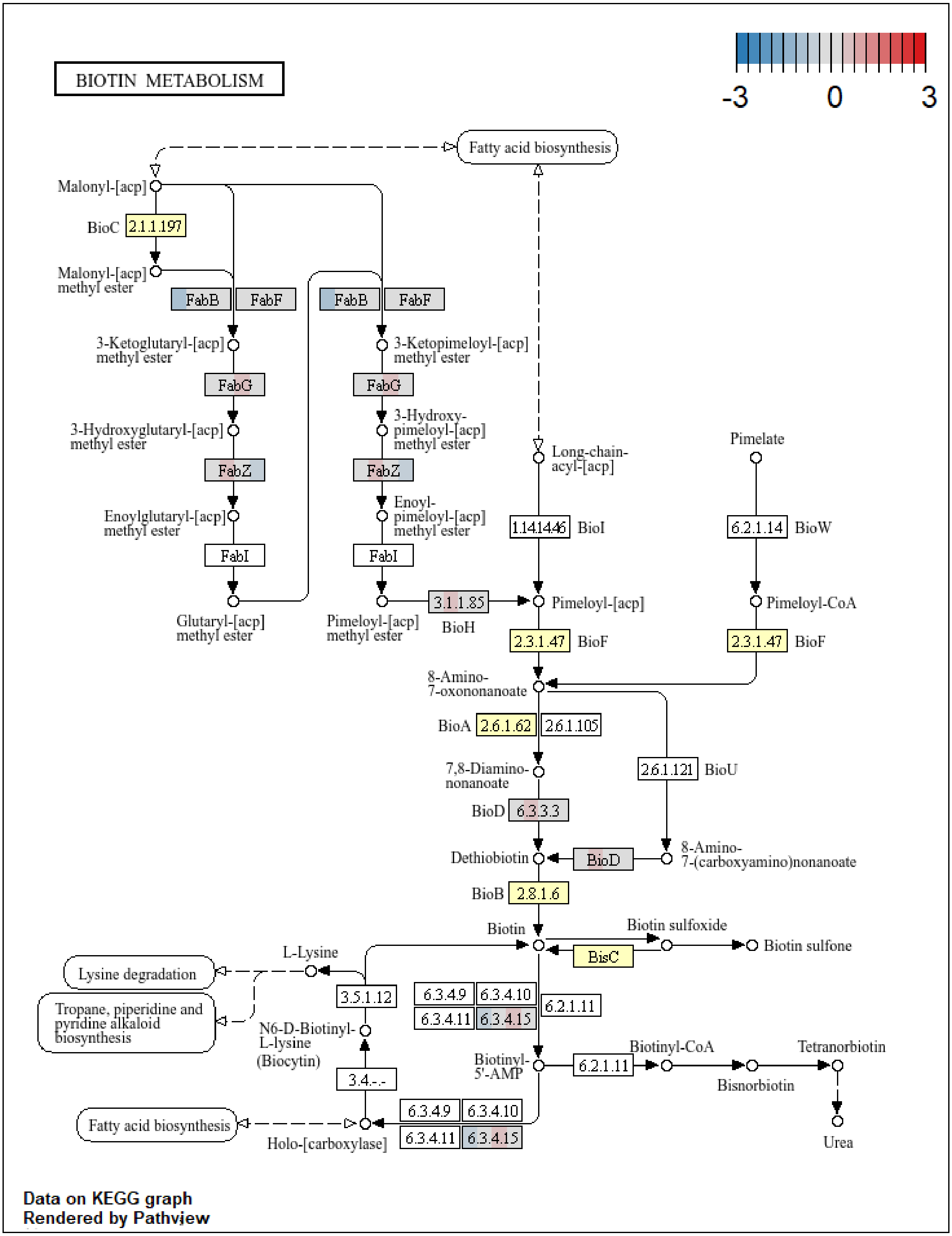
BIOTIN METABOLISM
-3
0
3
Fatty acid biosynthesis
Malonyl-[acp]
BioC
2.1.1.197
Malonyl-[acp] methyl ester
FabB
FabF
FabB
FabF
3-Ketoglutaryl-[acp] methyl ester
3-Ketopimeloyl-[acp] methyl ester
FabG
FabG
3-Hydroxyglutaryl-[acp] methyl ester
3-Hydroxy-pimeloyl-[acp] methyl ester
FabZ
FabZ
Enoylglutaryl-[acp] methyl ester
Enoyl-pimeloyl-[acp] methyl ester
FabI
FabI
Glutaryl-[acp] methyl ester
Pimeloyl-[acp] methyl ester
3.1.1.85
BioH
Long-chain-acyl-[acp]
1.14.14.46
BioI
Pimelate
6.2.1.14
BioW
Pimeloyl-[acp]
Pimeloyl-CoA
2.3.1.47
BioF
2.3.1.47
BioF
8-Amino-7-oxononanoate
BioA
2.6.1.62
2.6.1.105
2.6.1.121
BioU
7,8-Diamino-nonanoate
BioD
6.3.3.3
Dethiobiotin
BioD
8-Amino-7-(carboxyamino)nonanoate
BioB
2.8.1.6
Biotin
Biotin sulfoxide
Biotin sulfone
BisC
L-Lysine
Lysine degradation
Tropane, piperidine and pyridine alkaloid biosynthesis
3.5.1.12
6.3.4.9
6.3.4.10
6.3.4.11
6.3.4.15
6.2.1.11
N6-D-Biotinyl-L-lysine (Biocytin)
3.4.-.-
Biotinyl-5'-AMP
6.2.1.11
Biotinyl-CoA
Bisnorbiotin
Tetranorbiotin
Urea
6.3.4.9
6.3.4.10
6.3.4.11
6.3.4.15
Fatty acid biosynthesis
Holo-[carboxylase]
Data on KEGG graph
Rendered by Pathview

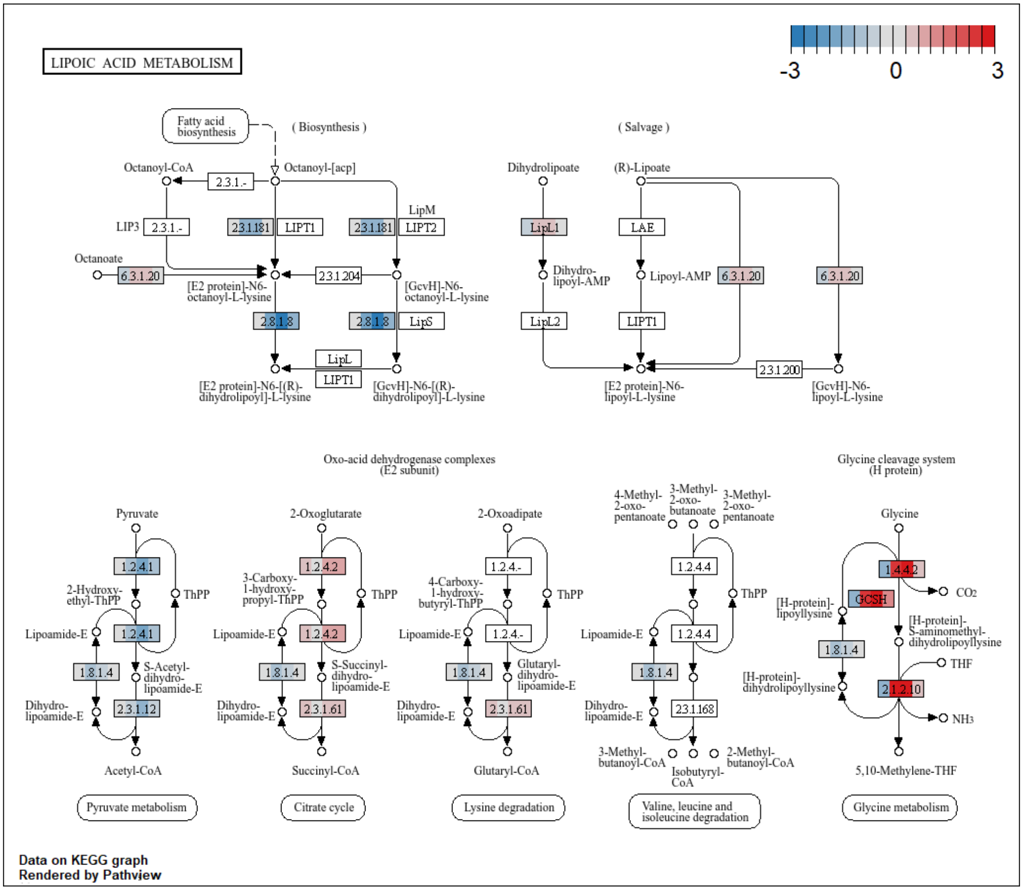

LIPOIC ACID METABOLISM
-3
0
3
Fatty acid biosynthesis
( Biosynthesis )
( Salvage )
Octanoyl-CoA
Octanoyl-[acp]
Dihydrolipoate
(R)-Lipoate
LipM
LIP3
LIPT1
LIPT2
LipL1
LAE
Octanoate
[E2 protein]-N6-octanoyl-L-lysine
[GcvH]-N6-octanoyl-L-lysine
Dihydro-lipoyl-AMP
Lipoyl-AMP
LipS
LipL2
LipL
[E2 protein]-N6-[(R)-dihydrolipoyl]-L-lysine
[GcvH]-N6-[(R)-dihydrolipoyl]-L-lysine
[E2 protein]-N6-lipoyl-L-lysine
[GcvH]-N6-lipoyl-L-lysine
Oxo-acid dehydrogenase complexes (E2 subunit)
Glycine cleavage system (H protein)
Pyruvate
2-Oxoglutarate
2-Oxoadipate
Glycine
ThPP
Lipoamide-E
Dihydro-lipoamide-E
Acetyl-CoA
Succinyl-CoA
Glutaryl-CoA
5,10-Methylene-THF
GCSH
CO2
THF
NH3
Pyruvate metabolism
Citrate cycle
Lysine degradation
Valine, leucine and isoleucine degradation
Glycine metabolism
Data on KEGG graph
Rendered by Pathview

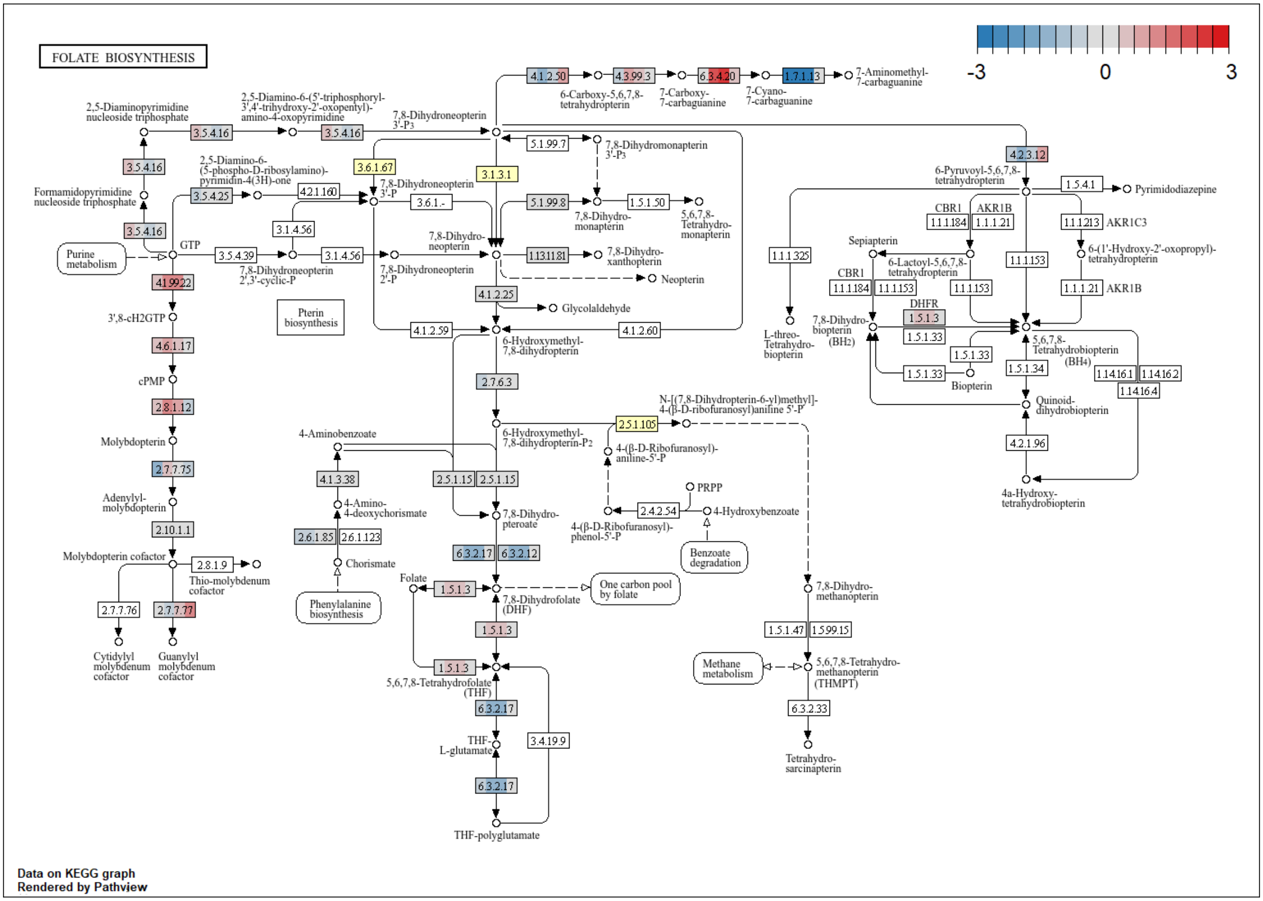
FOLATE BIOSYNTHESIS
-3
0
3
2,5-Diaminopyrimidine nucleoside triphosphate
2,5-Diamino-6-(5'-triphosphoryl-3',4'-trihydroxy-2'-oxopentyl)-amino-4-oxopyrimidine
7,8-Dihydroneopterin 3'-P3
Formamidopyrimidine nucleoside triphosphate
2,5-Diamino-6-(5-phospho-D-ribosylamino)-pyrimidin-4(3H)-one
7,8-Dihydroneopterin 3'-P
Purine metabolism
GTP
7,8-Dihydroneopterin 2',3'-cyclic-P
7,8-Dihydroneopterin 2'-P
7,8-Dihydro-neopterin
3',8-cH2GTP
Pterin biosynthesis
cPMP
Molybdopterin
Adenylyl-molybdopterin
Molybdopterin cofactor
Thio-molybdenum cofactor
Cytidylyl molybdenum cofactor
Guanylyl molybdenum cofactor
6-Carboxy-5,6,7,8-tetrahydropterin
7-Carboxy-7-carbaguanine
7-Cyano-7-carbaguanine
7-Aminomethyl-7-carbaguanine
7,8-Dihydromonapterin 3'-P3
7,8-Dihydro-monapterin
5,6,7,8-Tetrahydro-monapterin
7,8-Dihydro-xanthopterin
Neopterin
Glycolaldehyde
6-Hydroxymethyl-7,8-dihydropterin
6-Hydroxymethyl-7,8-dihydropterin-P2
N-[(7,8-Dihydropterin-6-yl)methyl]-4-(β-D-ribofuranosyl)aniline 5'-P
4-(β-D-Ribofuranosyl)-aniline-5'-P
PRPP
4-Hydroxybenzoate
4-(β-D-Ribofuranosyl)-phenol-5'-P
Benzoate degradation
4-Aminobenzoate
4-Amino-4-deoxychorismate
Chorismate
Phenylalanine biosynthesis
7,8-Dihydro-pteroate
Folate
7,8-Dihydrofolate (DHF)
One carbon pool by folate
5,6,7,8-Tetrahydrofolate (THF)
THF-L-glutamate
THF-polyglutamate
7,8-Dihydro-methanopterin
5,6,7,8-Tetrahydro-methanopterin (THMPT)
Methane metabolism
Tetrahydro-sarcinapterin
6-Pyruvoyl-5,6,7,8-tetrahydropterin
Pyrimidodiazepine
CBR1
AKR1B
AKR1C3
Sepiapterin
6-Lactoyl-5,6,7,8-tetrahydropterin
6-(1'-Hydroxy-2'-oxopropyl)-tetrahydropterin
L-threo-Tetrahydro-biopterin
7,8-Dihydro-biopterin (BH2)
DHFR
5,6,7,8-Tetrahydrobiopterin (BH4)
Biopterin
Quinoid-dihydrobiopterin
4a-Hydroxy-tetrahydrobiopterin
3.5.4.16
3.5.4.25
4.2.1.160
3.6.1.67
3.1.3.1
3.6.1.-
3.1.4.56
3.5.4.39
4.1.99.22
4.6.1.17
2.8.1.12
2.7.7.75
2.10.1.1
2.8.1.9
2.7.7.76
2.7.7.77
4.1.2.50
4.3.99.3
6.3.4.20
1.7.1.13
5.1.99.7
5.1.99.8
1.5.1.50
1.13.11.81
4.1.2.25
4.1.2.59
4.1.2.60
2.7.6.3
2.5.1.105
2.4.2.54
4.1.3.38
2.5.1.15
2.6.1.85
2.6.1.123
6.3.2.17
6.3.2.12
1.5.1.3
3.4.19.9
1.5.1.47
1.5.99.15
6.3.2.33
4.2.3.12
1.5.4.1
1.1.1.184
1.1.1.21
1.1.1.213
1.1.1.325
1.1.1.153
1.5.1.33
1.5.1.34
1.14.16.1
1.14.16.2
1.14.16.4
4.2.1.96
Data on KEGG graph
Rendered by Pathview

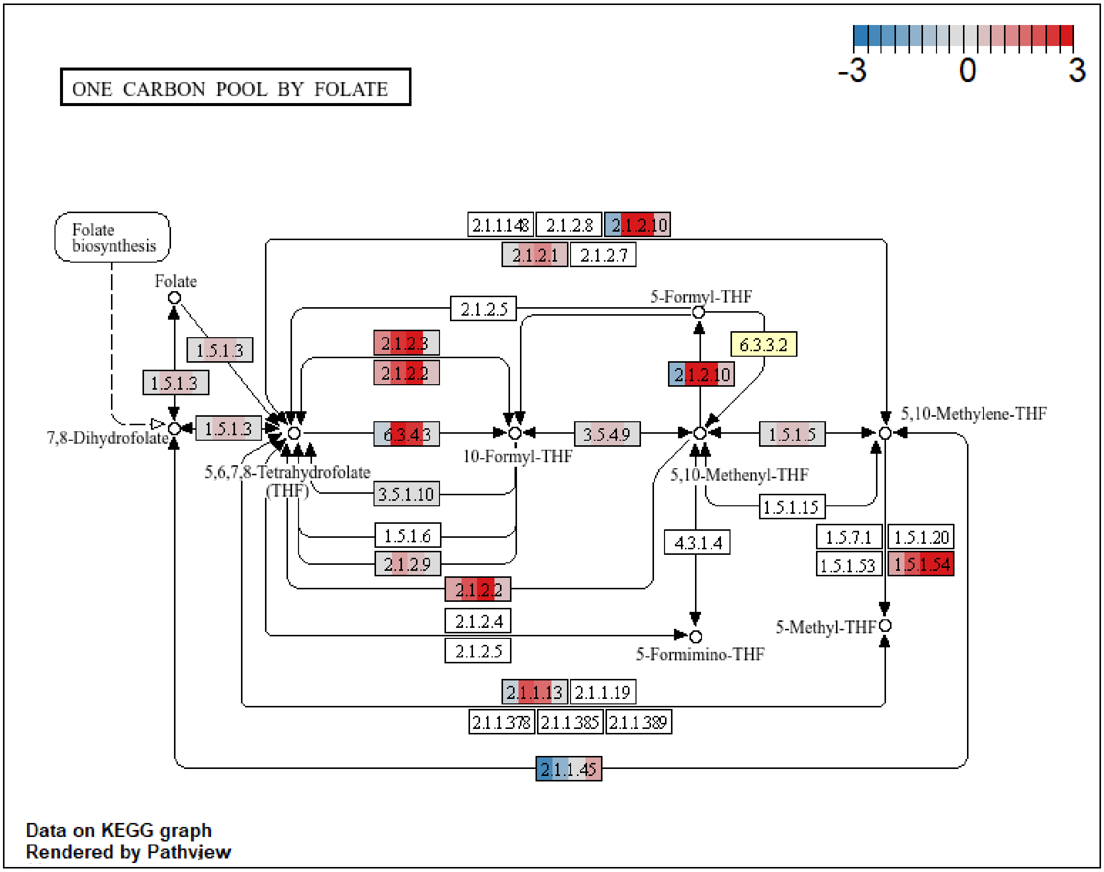
ONE CARBON POOL BY FOLATE
-3
0
3
Folate biosynthesis
Folate
7,8-Dihydrofolate
5,6,7,8-Tetrahydrofolate (THF)
10-Formyl-THF
5-Formyl-THF
5,10-Methenyl-THF
5,10-Methylene-THF
5-Methyl-THF
5-Formimino-THF
2.1.1.148
2.1.2.8
2.1.2.10
2.1.2.1
2.1.2.7
2.1.2.5
2.1.2.3
2.1.2.2
6.3.4.3
3.5.4.9
6.3.3.2
1.5.1.3
1.5.1.5
1.5.1.15
3.5.1.10
1.5.1.6
2.1.2.9
4.3.1.4
1.5.7.1
1.5.1.20
1.5.1.53
1.5.1.54
2.1.2.4
2.1.1.13
2.1.1.19
2.1.1.378
2.1.1.385
2.1.1.389
2.1.1.45
Data on KEGG graph
Rendered by Pathview

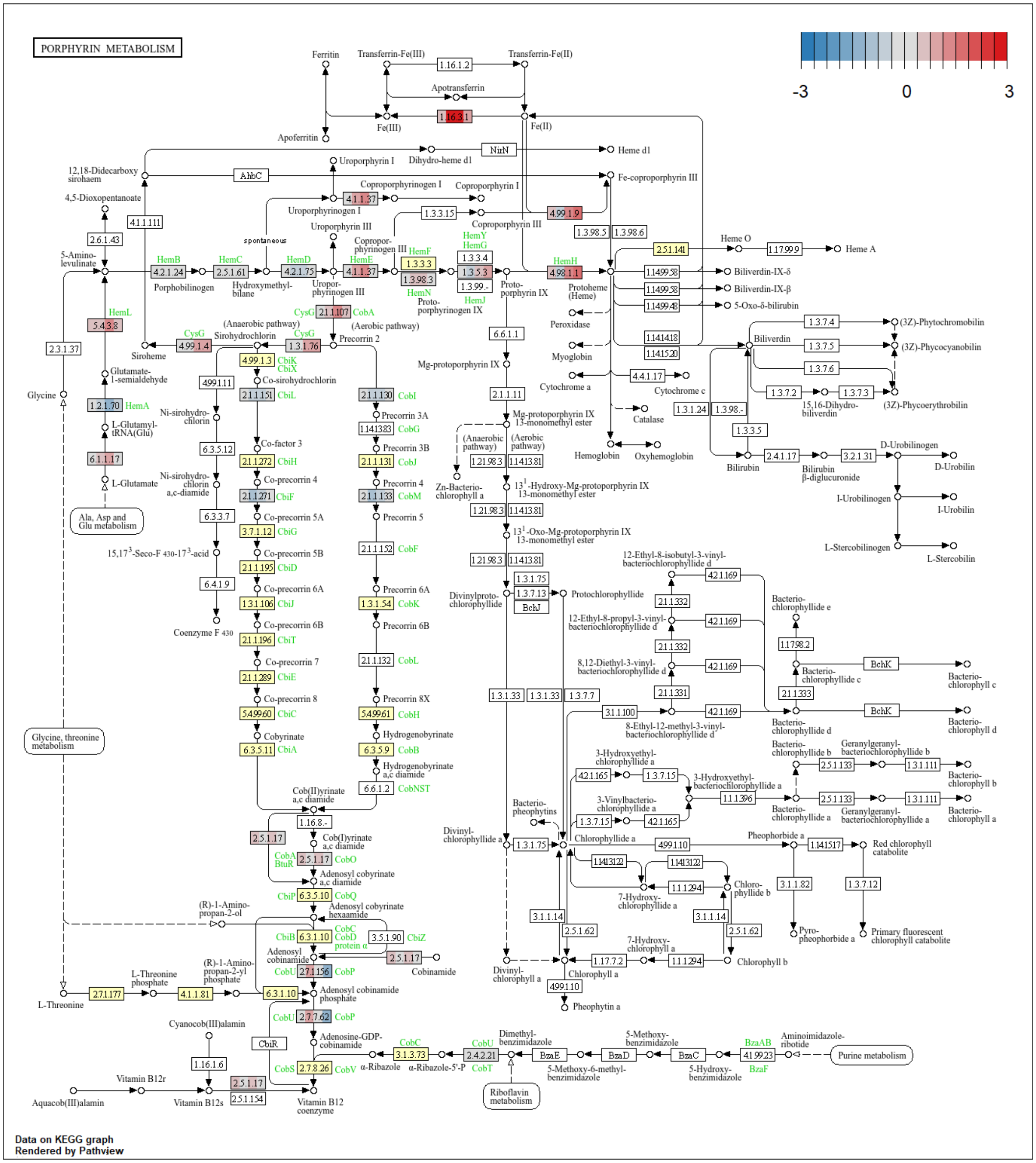
PORPHYRIN METABOLISM
-3
0
3
Ferritin
Transferrin-Fe(III)
1.16.1.2
Transferrin-Fe(II)
Apotransferrin
1.16.3.1
Fe(III)
Fe(II)
Apoferritin
NirN
Heme d1
Dihydro-heme d1
Uroporphyrin I
12,18-Didecarboxy siroheam
AhbC
Fe-coproporphyrin III
4,5-Dioxopentanoate
Coproporphyrinogen I
Coproporphyrin I
4.1.1.37
Uroporphyrinogen I
1.3.3.15
4.99.1.9
Coproporphyrin III
4.1.1.111
2.6.1.43
spontaneous
Uroporphyrin III
1.3.98.5
1.3.98.6
Heme O
2.5.1.141
1.17.99.9
Heme A
5-Amino-levulinate
HemB
HemC
HemD
HemE
HemF
HemY
HemG
HemH
4.2.1.24
2.5.1.61
4.2.1.75
1.3.3.3
1.3.3.4
1.3.98.3
1.3.5.3
1.3.99.-
4.98.1.1
Porphobilinogen
Hydroxymethyl-bilane
Uroporphyrinogen III
Coproporphyrinogen III
HemN
HemJ
Protoporphyrinogen IX
Protoporphyrin IX
Protoheme (Heme)
1.14.99.58
1.14.99.48
Biliverdin-IX-δ
Biliverdin-IX-β
5-Oxo-δ-bilirubin
HemL
CysG
2.1.1.107
CobA
5.4.3.8
4.99.1.4
1.3.1.76
(Anaerobic pathway)
(Aerobic pathway)
Sirohydrochlorin
Precorrin 2
Peroxidase
1.3.7.4
(3Z)-Phytochromobilin
Siroheme
4.99.1.3
CbiK
CbiX
6.6.1.1
Myoglobin
1.14.14.18
1.14.15.20
Biliverdin
1.3.7.5
(3Z)-Phycocyanobilin
Glutamate-1-semialdehyde
Co-sirohydrochlorin
Mg-protoporphyrin IX
1.3.7.6
Glycine
1.2.1.70
HemA
4.99.1.11
2.1.1.151
CbiL
2.1.1.130
CobI
Cytochrome a
4.4.1.17
Cytochrome c
L-Glutamyl-tRNA(Glu)
Ni-sirohydro-chlorin
Precorrin 3A
2.1.1.11
Mg-protoporphyrin IX 13-monomethyl ester
1.3.1.24
1.3.98.-
1.3.7.2
1.3.7.3
15,16-Dihydro-biliverdin
(3Z)-Phycoerythrobilin
Catalase
1.14.13.83
CobG
6.3.5.12
Co-factor 3
Precorrin 3B
1.3.3.5
(Anaerobic pathway)
(Aerobic pathway)
Hemoglobin
Oxyhemoglobin
6.1.1.17
2.1.1.272
CbiH
2.1.1.131
CobJ
1.21.98.3
1.14.13.81
Bilirubin
2.4.1.17
3.2.1.31
D-Urobilinogen
D-Urobilin
Bilirubin β-diglucuronide
L-Glutamate
Ni-sirohydro-chlorin a,c-diamide
Co-precorrin 4
Precorrin 4
2.1.1.271
CbiF
2.1.1.133
CobM
Zn-Bacterio-chlorophyll a
13[1]-Hydroxy-Mg-protoporphyrin IX 13-monomethyl ester
I-Urobilinogen
I-Urobilin
Ala, Asp and Glu metabolism
6.3.3.7
Co-precorrin 5A
Precorrin 5
3.7.1.12
CbiG
15,17[3]-Seco-F 430-17[3]-acid
Co-precorrin 5B
2.1.1.152
CobF
13[1]-Oxo-Mg-protoporphyrin IX 13-monomethyl ester
L-Stercobilinogen
L-Stercobilin
2.1.1.195
CbiD
12-Ethyl-8-isobutyl-3-vinyl-bacteriochlorophyllide d
4.2.1.169
6.4.1.9
Co-precorrin 6A
Precorrin 6A
1.3.1.75
1.3.7.13
BchJ
Divinylproto-chlorophyllide
Protochlorophyllide
2.1.1.332
Bacterio-chlorophyllide e
1.3.1.106
CbiJ
1.3.1.54
CobK
Coenzyme F 430
Co-precorrin 6B
Precorrin 6B
12-Ethyl-8-propyl-3-vinyl-bacteriochlorophyllide d
2.1.1.196
CbiT
1.17.98.2
Co-precorrin 7
2.1.1.132
CobL
8,12-Diethyl-3-vinyl-bacteriochlorophyllide d
BchK
Bacterio-chlorophyllide c
Bacterio-chlorophyll c
2.1.1.289
CbiE
2.1.1.331
2.1.1.333
Co-precorrin 8
Precorrin 8X
1.3.1.33
1.3.7.7
3.1.1.100
8-Ethyl-12-methyl-3-vinyl-bacteriochlorophyllide d
Bacterio-chlorophyllide d
Bacterio-chlorophyll d
5.4.99.60
CbiC
5.4.99.61
CobH
Glycine, threonine metabolism
Cobyrinate
Hydrogenobyrinate
6.3.5.11
CbiA
6.3.5.9
CobB
3-Hydroxyethyl-chlorophyllide a
Bacterio-chlorophyllide b
Geranylgeranyl-bacteriochlorophyllide b
2.5.1.133
1.3.1.111
Bacterio-chlorophyll b
Hydrogenobyrinate a,c diamide
4.2.1.165
1.3.7.15
3-Hydroxyethyl-bacteriochlorophyllide a
Cob(II)yrinate a,c diamide
6.6.1.2
CobNST
Bacterio-pheophytins
3-Vinylbacterio-chlorophyllide a
1.1.1.396
Bacterio-chlorophyllide a
Geranylgeranyl-bacteriochlorophyllide a
Bacterio-chlorophyll a
1.16.8.-
Divinyl-chlorophyllide a
Chlorophyllide a
Pheophorbide a
2.5.1.17
Cob(I)yrinate a,c diamide
4.99.1.10
1.14.15.17
Red chlorophyll catabolite
CobA
BtuR
CobO
1.14.13.122
Adenosyl cobyrinate a,c diamide
1.1.1.294
Chloro-phyll b
3.1.1.82
1.3.7.12
7-Hydroxy-chlorophyllide a
CbiP
6.3.5.10
CobQ
(R)-1-Amino-propan-2-ol
Adenosyl cobyrinate hexaamide
3.1.1.14
CbiB
6.3.1.10
CobC
CobD
3.5.1.90
CbiZ
protein α
2.5.1.62
Pyro-pheophorbide a
Primary fluorescent chlorophyll catabolite
Adenosyl cobinamide
Cobinamide
7-Hydroxy-chlorophyll a
L-Threonine phosphate
(R)-1-Amino-propan-2-yl phosphate
CobU
2.7.1.156
CobP
1.17.7.2
Chlorophyll b
Divinyl-chlorophyll a
Chlorophyll a
L-Threonine
2.7.1.177
4.1.1.81
Adenosyl cobinamide phosphate
Pheophytin a
2.7.7.62
Cyanocob(III)alamin
CbiR
Adenosine-GDP-cobinamide
Dimethyl-benzimidazole
5-Methoxy-benzimidazole
Aminoimidazole-ribotide
3.1.3.73
2.4.2.21
BzaE
BzaD
BzaC
BzaAB
4.1.99.23
Purine metabolism
1.16.1.6
α-Ribazole
α-Ribazole-5'-P
CobT
5-Methoxy-6-methyl-benzimidazole
5-Hydroxy-benzimidazole
BzaF
CobS
2.7.8.26
CobV
Vitamin B12r
2.5.1.154
Riboflavin metabolism
Aquacob(III)alamin
Vitamin B12s
Vitamin B12 coenzyme
Data on KEGG graph
Rendered by Pathview

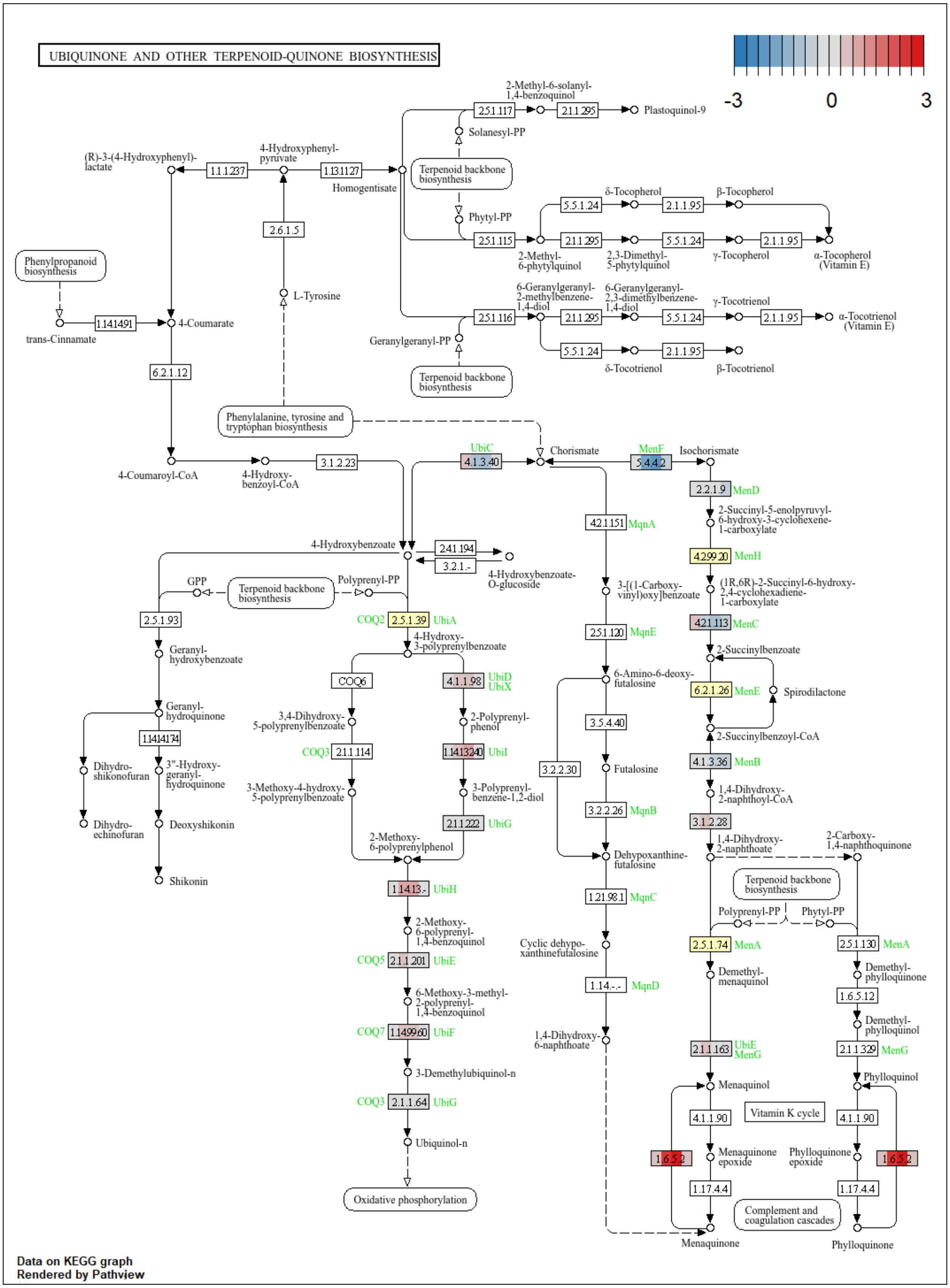
UBIQUINONE AND OTHER TERPENOID-QUINONE BIOSYNTHESIS
-3
0
3
2-Methyl-6-solanyl-1,4-benzoquinol
2.5.1.117
2.1.1.295
Plastoquinol-9
Solanesyl-PP
(R)-3-(4-Hydroxyphenyl)-lactate
4-Hydroxyphenyl-pyruvate
1.1.1.237
1.13.11.27
Homogentisate
Terpenoid backbone biosynthesis
2.6.1.5
Phytyl-PP
δ-Tocopherol
β-Tocopherol
5.5.1.24
2.1.1.95
2.5.1.115
2.1.1.295
2-Methyl-6-phytylquinol
2,3-Dimethyl-5-phytylquinol
γ-Tocopherol
α-Tocopherol (Vitamin E)
Phenylpropanoid biosynthesis
L-Tyrosine
trans-Cinnamate
1.14.14.91
4-Coumarate
6-Geranylgeranyl-2-methylbenzene-1,4-diol
6-Geranylgeranyl-2,3-dimethylbenzene-1,4-diol
γ-Tocotrienol
2.5.1.116
α-Tocotrienol (Vitamin E)
Geranylgeranyl-PP
δ-Tocotrienol
β-Tocotrienol
6.2.1.12
Phenylalanine, tyrosine and tryptophan biosynthesis
UbiC
4.1.3.40
Chorismate
MenF
5.4.4.2
Isochorismate
4-Coumaroyl-CoA
4-Hydroxy-benzoyl-CoA
3.1.2.23
2.2.1.9
MenD
2-Succinyl-5-enolpyruvyl-6-hydroxy-3-cyclohexene-1-carboxylate
4.2.1.151
MqnA
4-Hydroxybenzoate
2.4.1.194
3.2.1.-
4-Hydroxybenzoate-O-glucoside
4.2.99.20
MenH
GPP
Polyprenyl-PP
3-[(1-Carboxy-vinyl)oxy]benzoate
(1R,6R)-2-Succinyl-6-hydroxy-2,4-cyclohexadiene-1-carboxylate
2.5.1.93
COQ2
2.5.1.39
UbiA
2.5.1.120
MqnE
4.2.1.113
MenC
4-Hydroxy-3-polyprenylbenzoate
Geranyl-hydroxybenzoate
2-Succinylbenzoate
COQ6
4.1.1.98
UbiD
UbiX
6-Amino-6-deoxy-futalosine
6.2.1.26
MenE
Spirodilactone
Geranyl-hydroquinone
3,4-Dihydroxy-5-polyprenylbenzoate
2-Polyprenyl-phenol
3.5.4.40
2-Succinylbenzoyl-CoA
1.14.14.174
COQ3
2.1.1.114
1.14.13.240
UbiI
Dihydro-shikonofuran
3''-Hydroxy-geranyl-hydroquinone
3.2.2.30
Futalosine
4.1.3.36
MenB
3-Methoxy-4-hydroxy-5-polyprenylbenzoate
3-Polyprenyl-benzene-1,2-diol
1,4-Dihydroxy-2-naphthoyl-CoA
Dihydro-echinofuran
Deoxyshikonin
2.1.1.222
UbiG
3.2.2.26
MqnB
3.1.2.28
2-Methoxy-6-polyprenylphenol
Dehypoxanthine-futalosine
1,4-Dihydroxy-2-naphthoate
2-Carboxy-1,4-naphthoquinone
Shikonin
1.14.13.-
UbiH
1.21.98.1
MqnC
Polyprenyl-PP
Phytyl-PP
2-Methoxy-6-polyprenyl-1,4-benzoquinol
Cyclic dehypo-xanthinefutalosine
2.5.1.74
MenA
2.5.1.130
MenA
COQ5
2.1.1.201
UbiE
Demethyl-menaquinol
Demethyl-phylloquinone
1.14.-.-
MqnD
1.6.5.12
6-Methoxy-3-methyl-2-polyprenyl-1,4-benzoquinol
1,4-Dihydroxy-6-naphthoate
Demethyl-phylloquinol
COQ7
1.14.99.60
UbiF
2.1.1.163
UbiE
MenG
2.1.1.329
MenG
3-Demethylubiquinol-n
Menaquinol
Phylloquinol
Vitamin K cycle
COQ3
2.1.1.64
UbiG
4.1.1.90
4.1.1.90
Ubiquinol-n
1.6.5.2
Menaquinone epoxide
Phylloquinone epoxide
1.6.5.2
1.17.4.4
1.17.4.4
Oxidative phosphorylation
Complement and coagulation cascades
Menaquinone
Phylloquinone
Data on KEGG graph
Rendered by Pathview

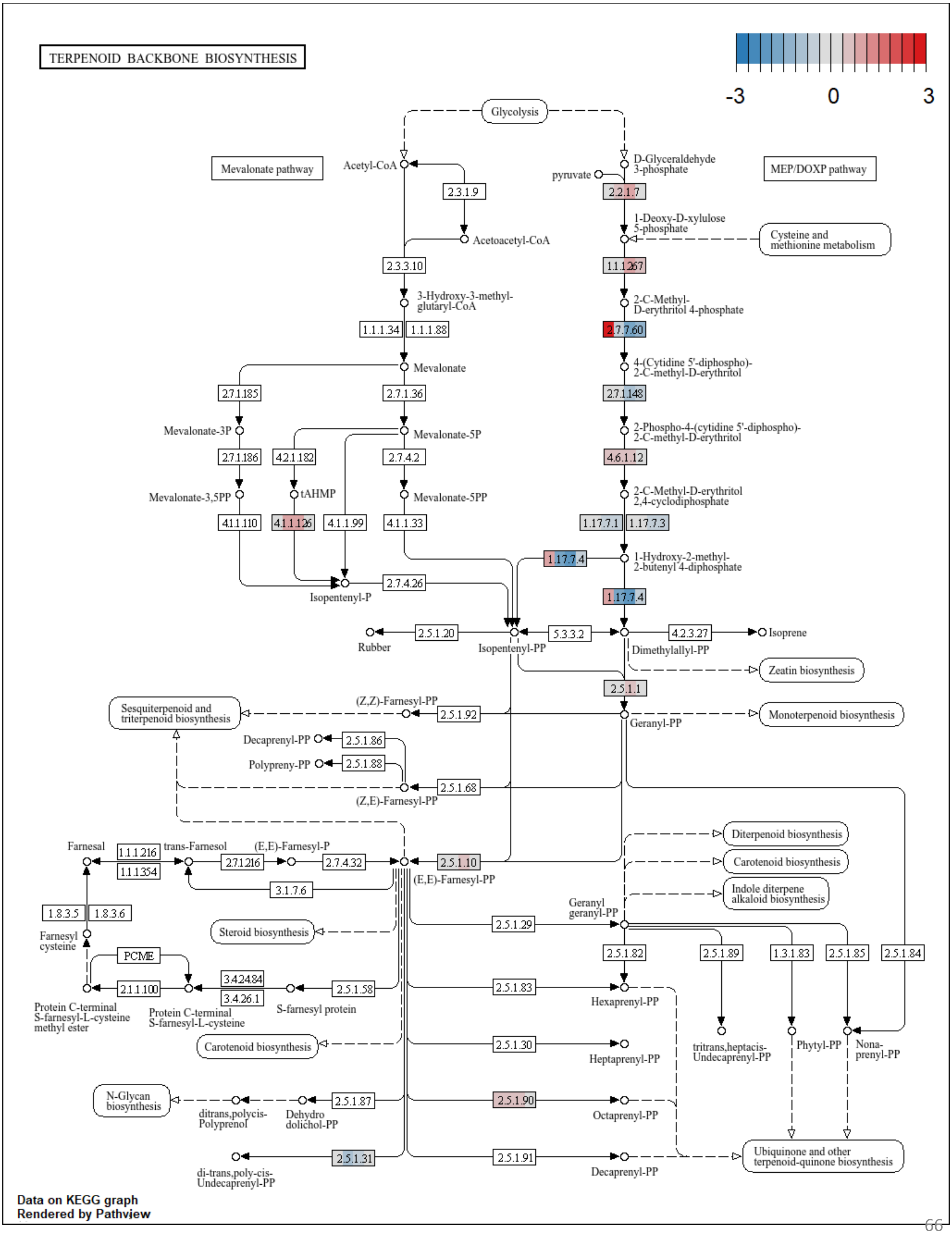
TERPENOID BACKBONE BIOSYNTHESIS
-3
0
3
Glycolysis
Mevalonate pathway
Acetyl-CoA
2.3.1.9
Acetoacetyl-CoA
2.3.3.10
3-Hydroxy-3-methyl-glutaryl-CoA
1.1.1.34
1.1.1.88
Mevalonate
2.7.1.185
2.7.1.36
Mevalonate-3P
Mevalonate-5P
2.7.1.186
4.2.1.182
2.7.4.2
Mevalonate-3,5PP
tAHMP
Mevalonate-5PP
4.1.1.110
4.1.1.126
4.1.1.99
4.1.1.33
Isopentenyl-P
2.7.4.26
D-Glyceraldehyde 3-phosphate
pyruvate
MEP/DOXP pathway
2.2.1.7
1-Deoxy-D-xylulose 5-phosphate
Cysteine and methionine metabolism
1.1.1.267
2-C-Methyl-D-erythritol 4-phosphate
2.7.7.60
4-(Cytidine 5'-diphospho)-2-C-methyl-D-erythritol
2.7.1.148
2-Phospho-4-(cytidine 5'-diphospho)-2-C-methyl-D-erythritol
4.6.1.12
2-C-Methyl-D-erythritol 2,4-cyclodiphosphate
1.17.7.1
1.17.7.3
1.17.7.4
1-Hydroxy-2-methyl-2-butenyl 4-diphosphate
1.17.7.4
Rubber
2.5.1.20
Isopentenyl-PP
5.3.3.2
Dimethylallyl-PP
4.2.3.27
Isoprene
Zeatin biosynthesis
2.5.1.1
Sesquiterpenoid and triterpenoid biosynthesis
(Z,Z)-Farnesyl-PP
2.5.1.92
Geranyl-PP
Monoterpenoid biosynthesis
Decaprenyl-PP
2.5.1.86
Polypreny-PP
2.5.1.88
2.5.1.68
(Z,E)-Farnesyl-PP
Farnesal
1.1.1.216
1.1.1.354
trans-Farnesol
2.7.1.216
(E,E)-Farnesyl-P
2.7.4.32
2.5.1.10
(E,E)-Farnesyl-PP
3.1.7.6
Diterpenoid biosynthesis
Carotenoid biosynthesis
Indole diterpene alkaloid biosynthesis
1.8.3.5
1.8.3.6
Farnesyl cysteine
Steroid biosynthesis
2.5.1.29
Geranyl geranyl-PP
PCME
3.4.24.84
3.4.26.1
2.1.1.100
2.5.1.58
S-farnesyl protein
Protein C-terminal S-farnesyl-L-cysteine methyl ester
Protein C-terminal S-farnesyl-L-cysteine
2.5.1.83
Hexaprenyl-PP
2.5.1.82
2.5.1.89
1.3.1.83
2.5.1.85
2.5.1.84
Carotenoid biosynthesis
2.5.1.30
Heptaprenyl-PP
tritrans,heptacis-Undecaprenyl-PP
Phytyl-PP
Nona-prenyl-PP
N-Glycan biosynthesis
ditrans,polycis-Polyprenol
Dehydro dolichol-PP
2.5.1.87
2.5.1.90
Octaprenyl-PP
Ubiquinone and other terpenoid-quinone biosynthesis
2.5.1.31
di-trans,poly-cis-Undecaprenyl-PP
2.5.1.91
Decaprenyl-PP
Data on KEGG graph
Rendered by Pathview

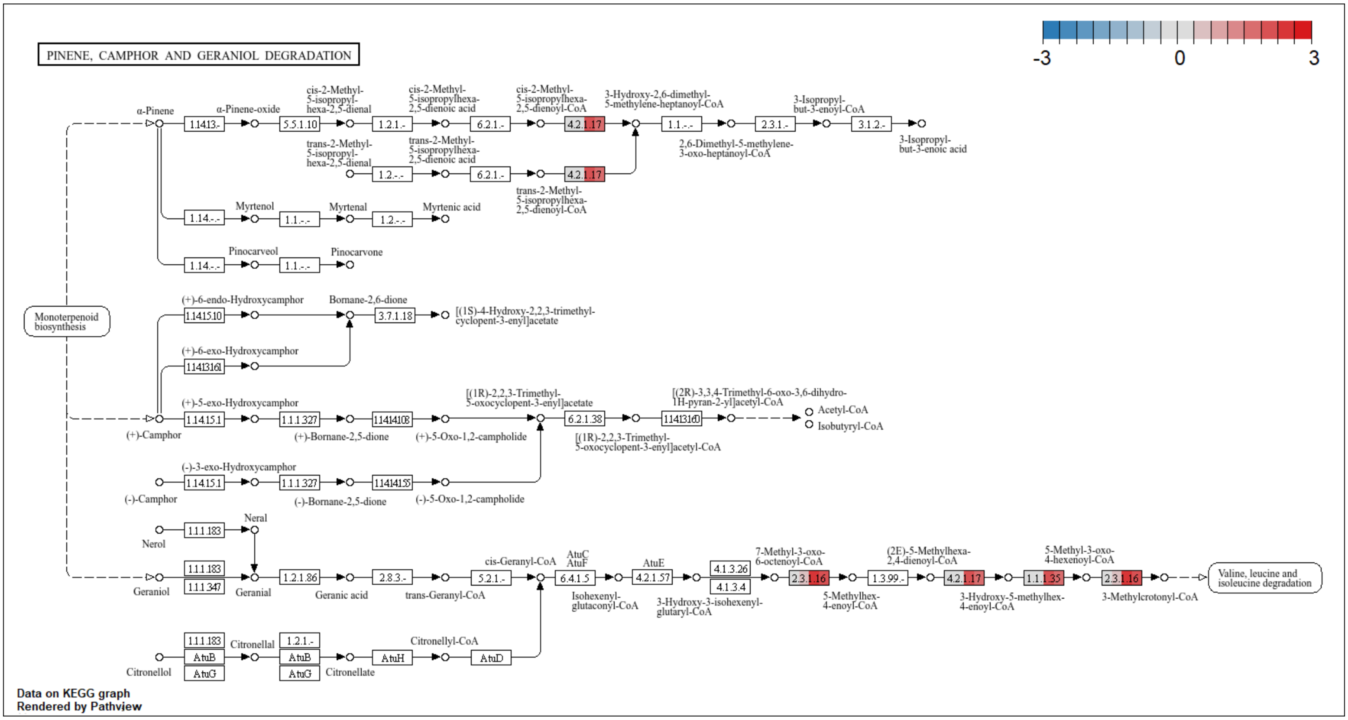
PINENE, CAMPHOR AND GERANIOL DEGRADATION
-3
0
3
α-Pinene
α-Pinene-oxide
cis-2-Methyl-5-isopropyl-hexa-2,5-dienal
cis-2-Methyl-5-isopropylhexa-2,5-dienoic acid
cis-2-Methyl-5-isopropylhexa-2,5-dienoyl-CoA
3-Hydroxy-2,6-dimethyl-5-methylene-heptanoyl-CoA
3-Isopropyl-but-3-enoyl-CoA
3-Isopropyl-but-3-enoic acid
2,6-Dimethyl-5-methylene-3-oxo-heptanoyl-CoA
trans-2-Methyl-5-isopropyl-hexa-2,5-dienal
trans-2-Methyl-5-isopropylhexa-2,5-dienoic acid
trans-2-Methyl-5-isopropylhexa-2,5-dienoyl-CoA
Myrtenol
Myrtenal
Myrtenic acid
Pinocarveol
Pinocarvone
Monoterpenoid biosynthesis
(+)-6-endo-Hydroxycamphor
Bornane-2,6-dione
[(1S)-4-Hydroxy-2,2,3-trimethyl-cyclopent-3-enyl]acetate
(+)-6-exo-Hydroxycamphor
(+)-5-exo-Hydroxycamphor
[(1R)-2,2,3-Trimethyl-5-oxocyclopent-3-enyl]acetate
[(2R)-3,3,4-Trimethyl-6-oxo-3,6-dihydro-1H-pyran-2-yl]acetyl-CoA
Acetyl-CoA
Isobutyryl-CoA
(+)-Camphor
(+)-Bornane-2,5-dione
(+)-5-Oxo-1,2-campholide
[(1R)-2,2,3-Trimethyl-5-oxocyclopent-3-enyl]acetyl-CoA
(-)-3-exo-Hydroxycamphor
(-)-Camphor
(-)-Bornane-2,5-dione
(-)-5-Oxo-1,2-campholide
Nerol
Neral
Geraniol
Geranial
Geranic acid
trans-Geranyl-CoA
cis-Geranyl-CoA
Isohexenyl-glutaconyl-CoA
3-Hydroxy-3-isohexenyl-glutaryl-CoA
7-Methyl-3-oxo-6-octenoyl-CoA
5-Methylhex-4-enoyl-CoA
(2E)-5-Methylhexa-2,4-dienoyl-CoA
3-Hydroxy-5-methylhex-4-enoyl-CoA
5-Methyl-3-oxo-4-hexenoyl-CoA
3-Methylcrotonyl-CoA
Valine, leucine and isoleucine degradation
Citronellol
Citronellal
Citronellate
Citronellyl-CoA
1.14.13.-
5.5.1.10
1.2.1.-
6.2.1.-
4.2.1.17
1.1.-.-
2.3.1.-
3.1.2.-
1.2.-.-
1.14.-.-
1.14.15.10
3.7.1.18
1.14.15.1
1.1.1.327
6.2.1.38
1.1.1.183
1.1.1.347
1.2.1.86
2.8.3.-
5.2.1.-
6.4.1.5
4.2.1.57
4.1.3.26
4.1.3.4
2.3.1.16
1.3.99.-
1.1.1.35
AtuC
AtuF
AtuE
AtuB
AtuG
AtuH
AtuD
Data on KEGG graph
Rendered by Pathview

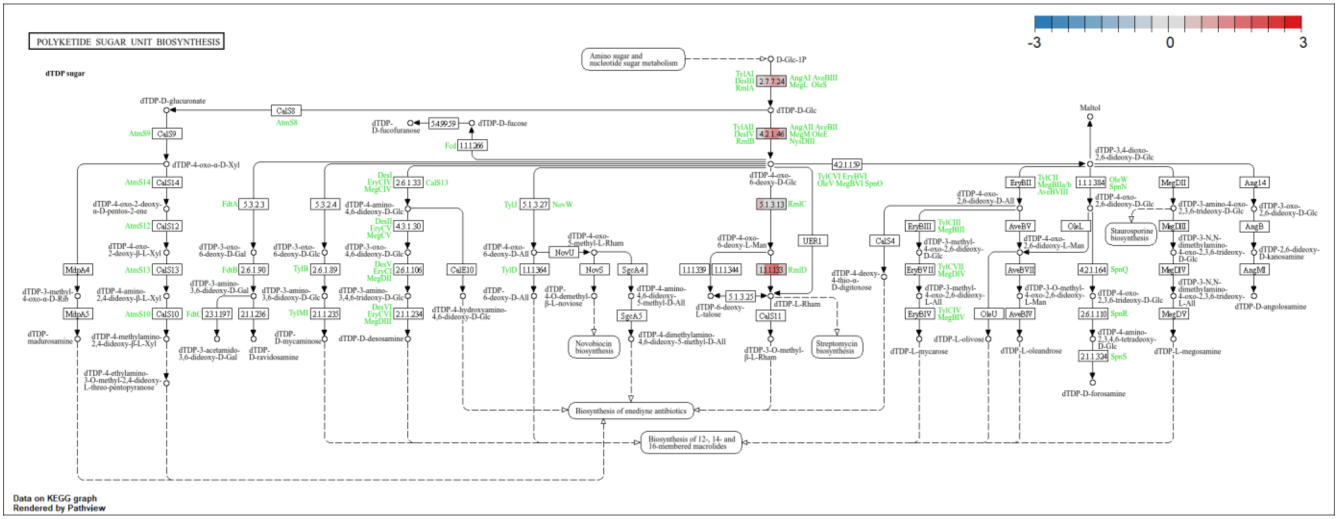
POLYKETIDE SUGAR UNIT BIOSYNTHESIS
dTDP sugar
-3
0
3
Amino sugar and nucleotide sugar metabolism
D-Glc-1P
dTDP-D-Glc
dTDP-D-glucuronate
Maltol
Biosynthesis of enediyne antibiotics
Biosynthesis of 12-, 14- and 16-membered macrolides
Novobiocin biosynthesis
Streptomycin biosynthesis
Staurosporine biosynthesis
Data on KEGG graph
Rendered by Pathview

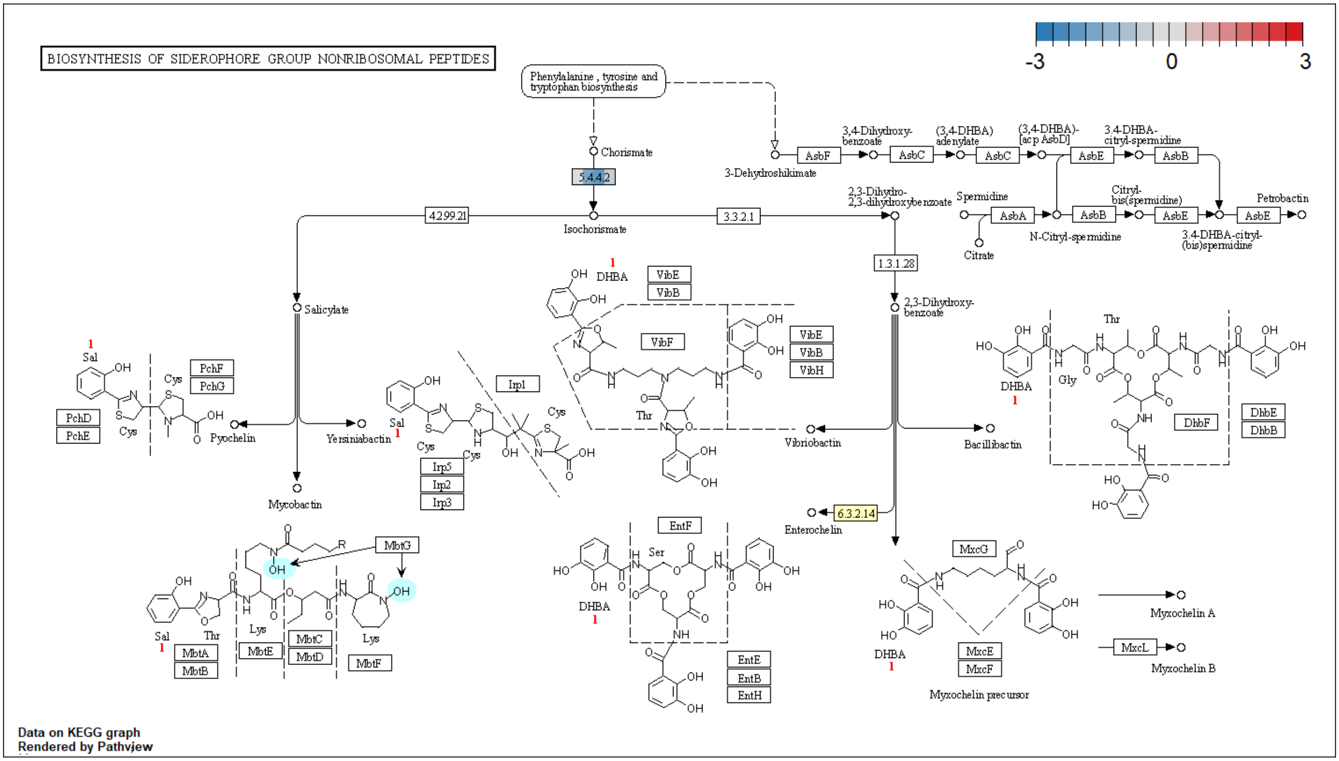

BIOSYNTHESIS OF SIDEROPHORE GROUP NONRIBOSOMAL PEPTIDES
-3
0
3
Phenylalanine, tyrosine and tryptophan biosynthesis
Chorismate
5.4.4.2
Isochorismate
4.2.99.21
3.3.2.1
3-Dehydroshikimate
AsbF
3,4-Dihydroxy-benzoate
AsbC
(3,4-DHBA) adenylate
(3,4-DHBA)-[acp AsbD]
AsbE
3,4-DHBA-citryl-spermidine
AsbB
2,3-Dihydro-2,3-dihydroxybenzoate
Spermidine
AsbA
N-Citryl-spermidine
Citrate
Citryl-bis(spermidine)
3,4-DHBA-citryl-(bis)spermidine
Petrobactin
1.3.1.28
2,3-Dihydroxy-benzoate
Salicylate
PchF
PchG
PchD
PchE
Pyochelin
Yersiniabactin
Irp1
Irp5
Irp2
Irp3
Mycobactin
DHBA
VibE
VibB
VibF
VibH
Vibriobactin
Bacillibactin
DhbF
DhbE
DhbB
6.3.2.14
Enterochelin
EntF
EntE
EntB
EntH
MbtG
MbtA
MbtB
MbtE
MbtC
MbtD
MbtF
MxcG
MxcE
MxcF
MxcL
Myxochelin A
Myxochelin B
Myxochelin precursor
Sal
Cys
Thr
Lys
Ser
Gly
Data on KEGG graph
Rendered by Pathview

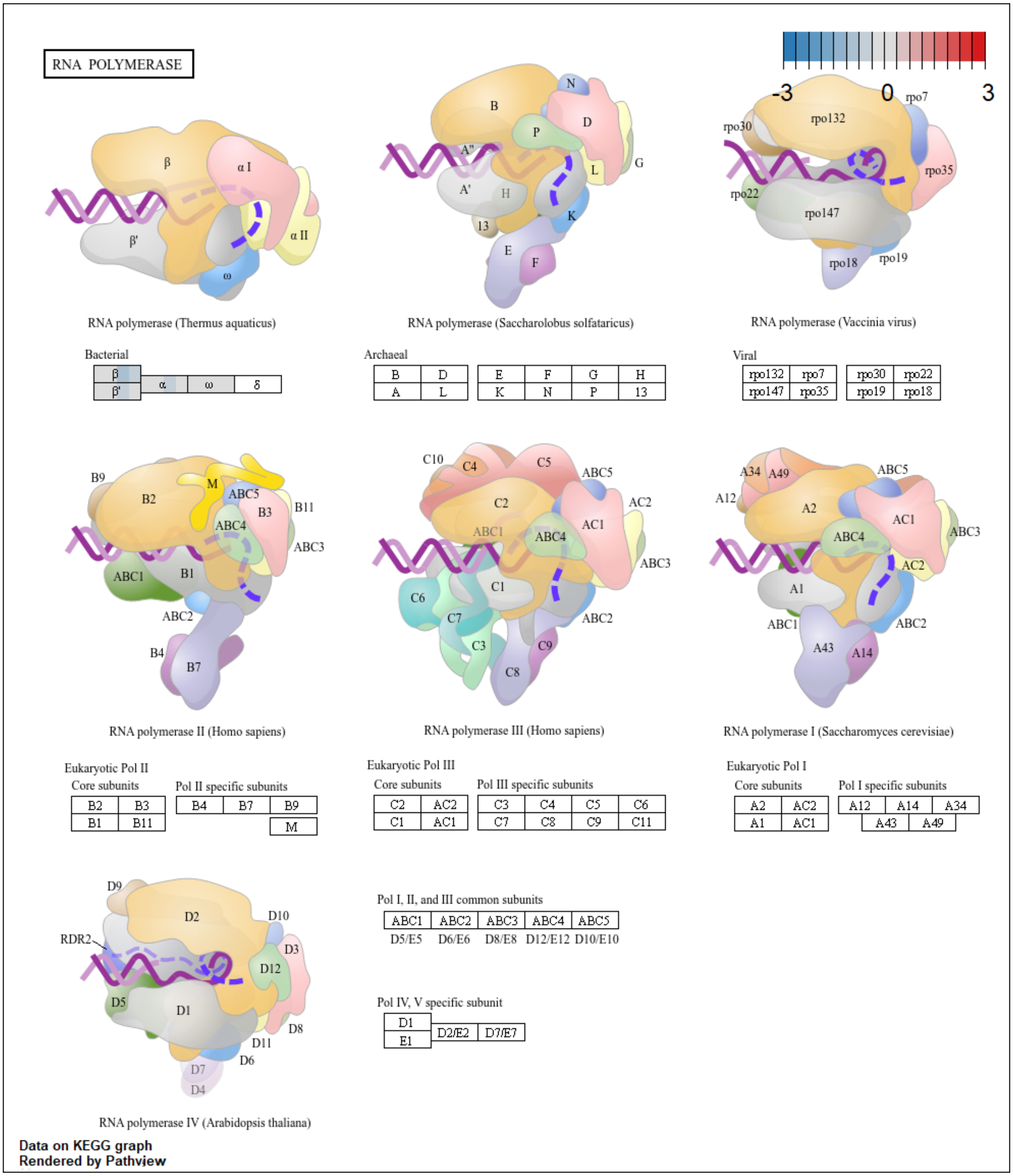

RNA POLYMERASE
-3
0
3
RNA polymerase (Thermus aquaticus)
β
α I
β'
α II
ω
Bacterial
β
β'
α
ω
δ
RNA polymerase (Saccharolobus solfataricus)
B
N
D
P
A''
L
G
A'
H
K
13
E
F
Archaeal
B
D
A
L
E
F
G
H
K
N
P
13
RNA polymerase (Vaccinia virus)
rpo7
rpo30
rpo132
rpo35
rpo22
rpo147
rpo18
rpo19
Viral
rpo132
rpo7
rpo147
rpo35
rpo30
rpo22
rpo19
rpo18
RNA polymerase II (Homo sapiens)
B9
B2
M
ABC5
B3
B11
ABC4
ABC3
ABC1
B1
ABC2
B4
B7
Eukaryotic Pol II
Core subunits
B2
B3
B1
B11
Pol II specific subunits
B4
B7
B9
M
RNA polymerase III (Homo sapiens)
C10
C4
C5
ABC5
C2
AC2
ABC1
AC1
ABC4
ABC3
C1
C6
C7
ABC2
C3
C9
C8
Eukaryotic Pol III
Core subunits
C2
AC2
C1
AC1
Pol III specific subunits
C3
C4
C5
C6
C7
C8
C9
C11
RNA polymerase I (Saccharomyces cerevisiae)
A34
A49
ABC5
A12
A2
AC1
ABC3
ABC4
AC2
A1
ABC1
ABC2
A43
A14
Eukaryotic Pol I
Core subunits
A2
AC2
A1
AC1
Pol I specific subunits
A12
A14
A34
A43
A49
RNA polymerase IV (Arabidopsis thaliana)
D9
D2
D10
RDR2
D3
D12
D5
D1
D8
D11
D6
D7
D4
Pol I, II, and III common subunits
ABC1
ABC2
ABC3
ABC4
ABC5
D5/E5
D6/E6
D8/E8
D12/E12
D10/E10
Pol IV, V specific subunit
D1
E1
D2/E2
D7/E7
Data on KEGG graph
Rendered by Pathview

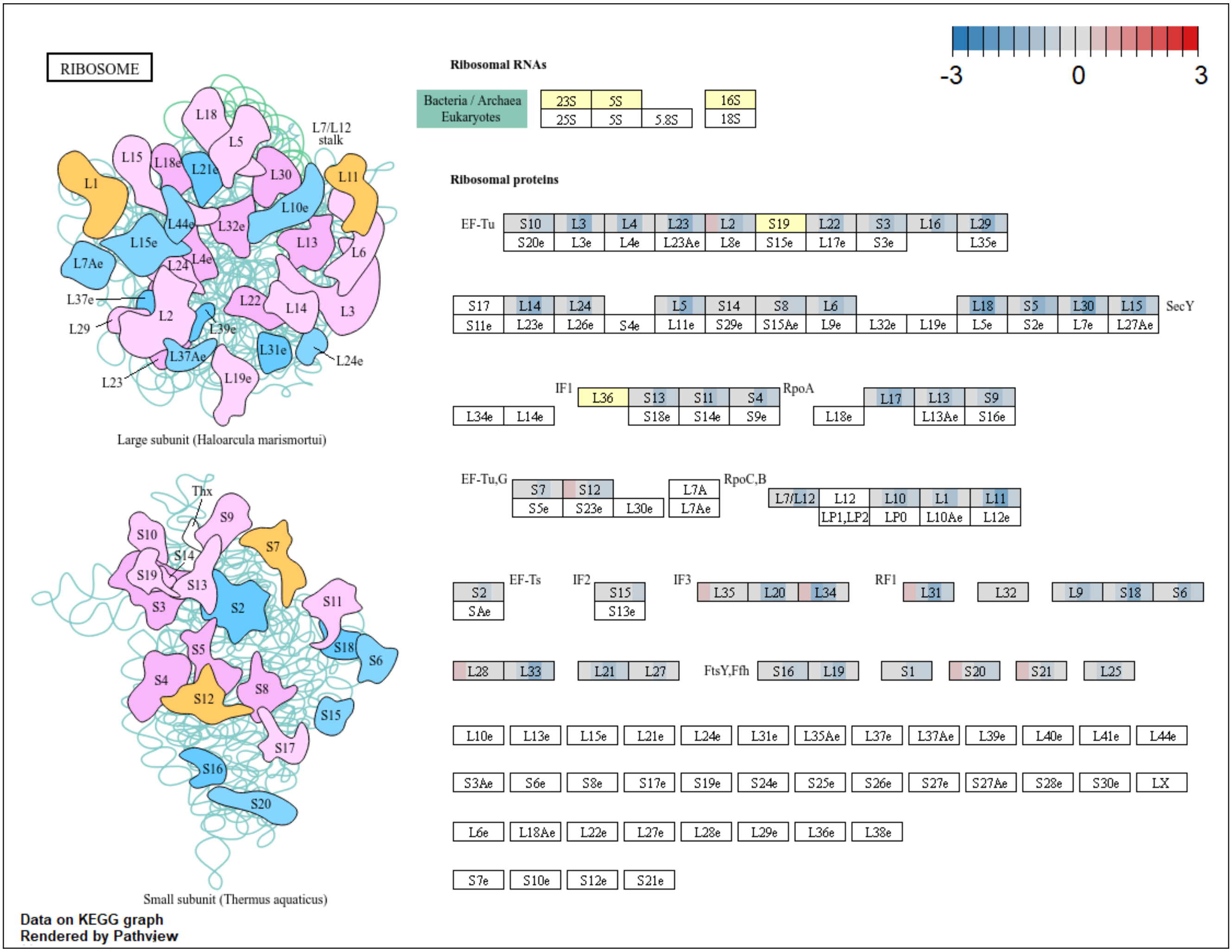
RIBOSOME
Ribosomal RNAs
Bacteria / Archaea
Eukaryotes
23S
5S
16S
25S
5S
5.8S
18S
Ribosomal proteins
-3
0
3
EF-Tu
SecY
IF1
RpoA
EF-Tu,G
RpoC,B
EF-Ts
IF2
IF3
RF1
FtsY,Ffh
Large subunit (Haloarcula marismortui)
Small subunit (Thermus aquaticus)
Data on KEGG graph
Rendered by Pathview

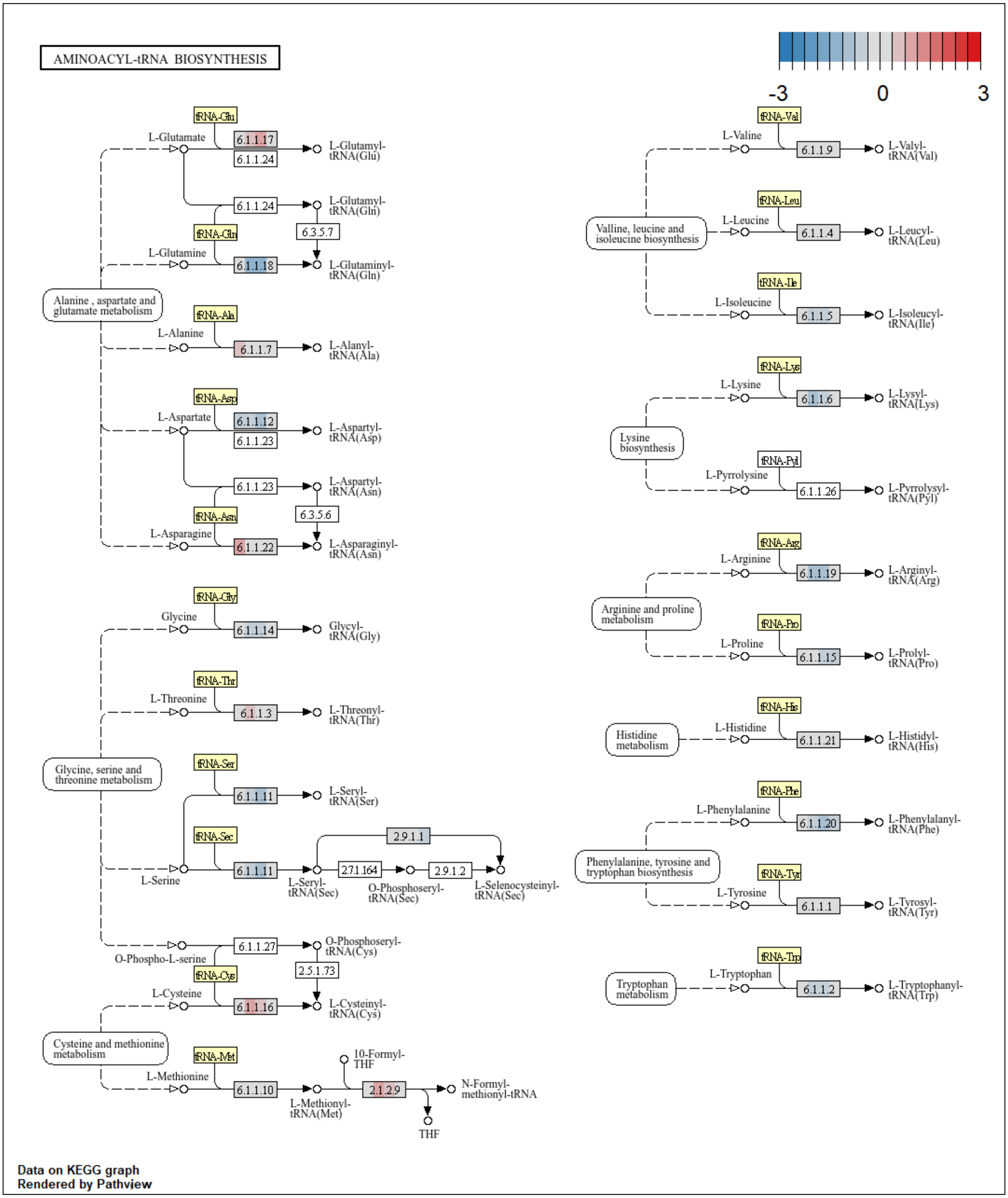
AMINOACYL-tRNA BIOSYNTHESIS
-3
0
3
Alanine , aspartate and glutamate metabolism
Glycine, serine and threonine metabolism
Cysteine and methionine metabolism
Valline, leucine and isoleucine biosynthesis
Lysine biosynthesis
Arginine and proline metabolism
Histidine metabolism
Phenylalanine, tyrosine and tryptophan biosynthesis
Tryptophan metabolism
Data on KEGG graph
Rendered by Pathview

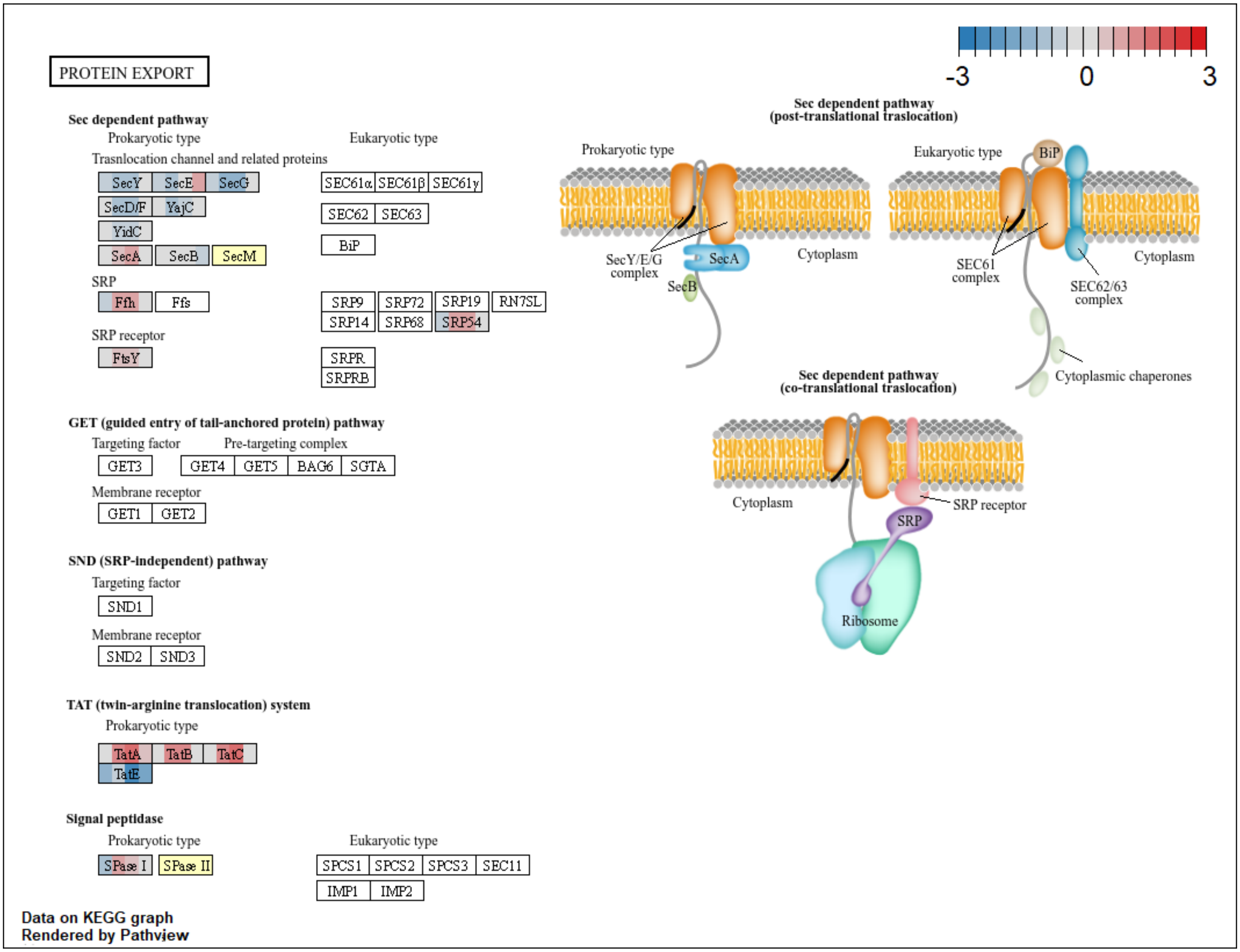

-3
0
3
PROTEIN EXPORT
Sec dependent pathway
Prokaryotic type
Eukaryotic type
Traslocation channel and related proteins
SecY
SecE
SecG
SecD/F
YajC
YidC
SecA
SecB
SecM
SEC61α
SEC61β
SEC61γ
SEC62
SEC63
BiP
SRP
Ffh
Ffs
SRP9
SRP72
SRP19
RN7SL
SRP14
SRP68
SRP54
SRP receptor
FtsY
SRPR
SRPRB
GET (guided entry of tail-anchored protein) pathway
Targeting factor
Pre-targeting complex
GET3
GET4
GET5
BAG6
SGTA
Membrane receptor
GET1
GET2
SND (SRP-independent) pathway
Targeting factor
SND1
Membrane receptor
SND2
SND3
TAT (twin-arginine translocation) system
Prokaryotic type
TatA
TatB
TatC
TatE
Signal peptidase
Prokaryotic type
Eukaryotic type
SPase I
SPase II
SPCS1
SPCS2
SPCS3
SEC11
IMP1
IMP2
Sec dependent pathway
(post-translational traslocation)
Prokaryotic type
Eukaryotic type
BiP
SecY/E/G
complex
SecA
SecB
Cytoplasm
SEC61
complex
SEC62/63
complex
Cytoplasmic chaperones
Sec dependent pathway
(co-translational traslocation)
Cytoplasm
SRP receptor
SRP
Ribosome
Data on KEGG graph
Rendered by Pathview

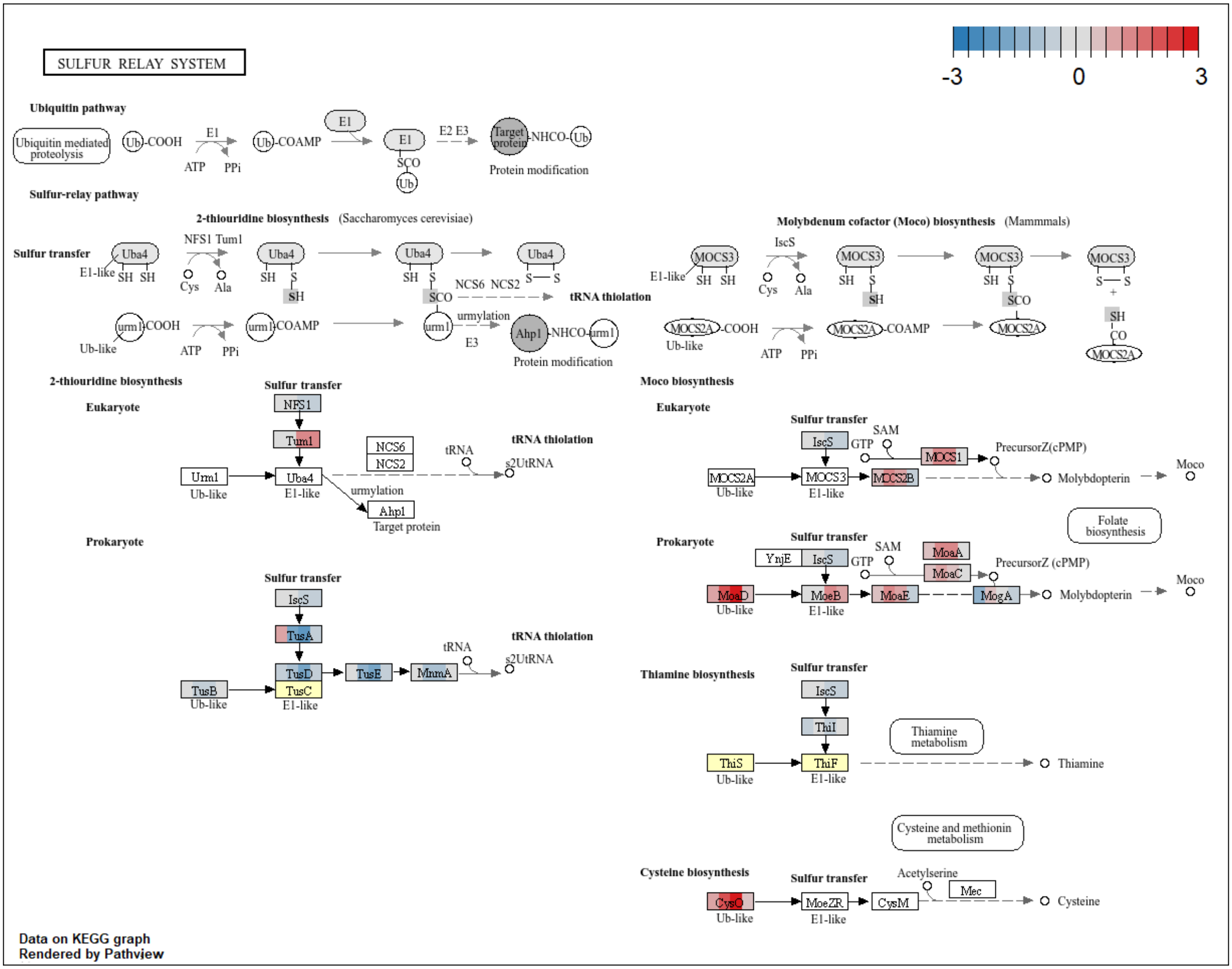
SULFUR RELAY SYSTEM
-3
0
3
Ubiquitin pathway
Ubiquitin mediated proteolysis
Protein modification
Sulfur-relay pathway
2-thiouridine biosynthesis (Saccharomyces cerevisiae)
Molybdenum cofactor (Moco) biosynthesis (Mammmals)
Sulfur transfer
tRNA thiolation
2-thiouridine biosynthesis
Eukaryote
Prokaryote
Moco biosynthesis
Thiamine biosynthesis
Cysteine biosynthesis
Folate biosynthesis
Thiamine metabolism
Cysteine and methionin metabolism
Data on KEGG graph
Rendered by Pathview

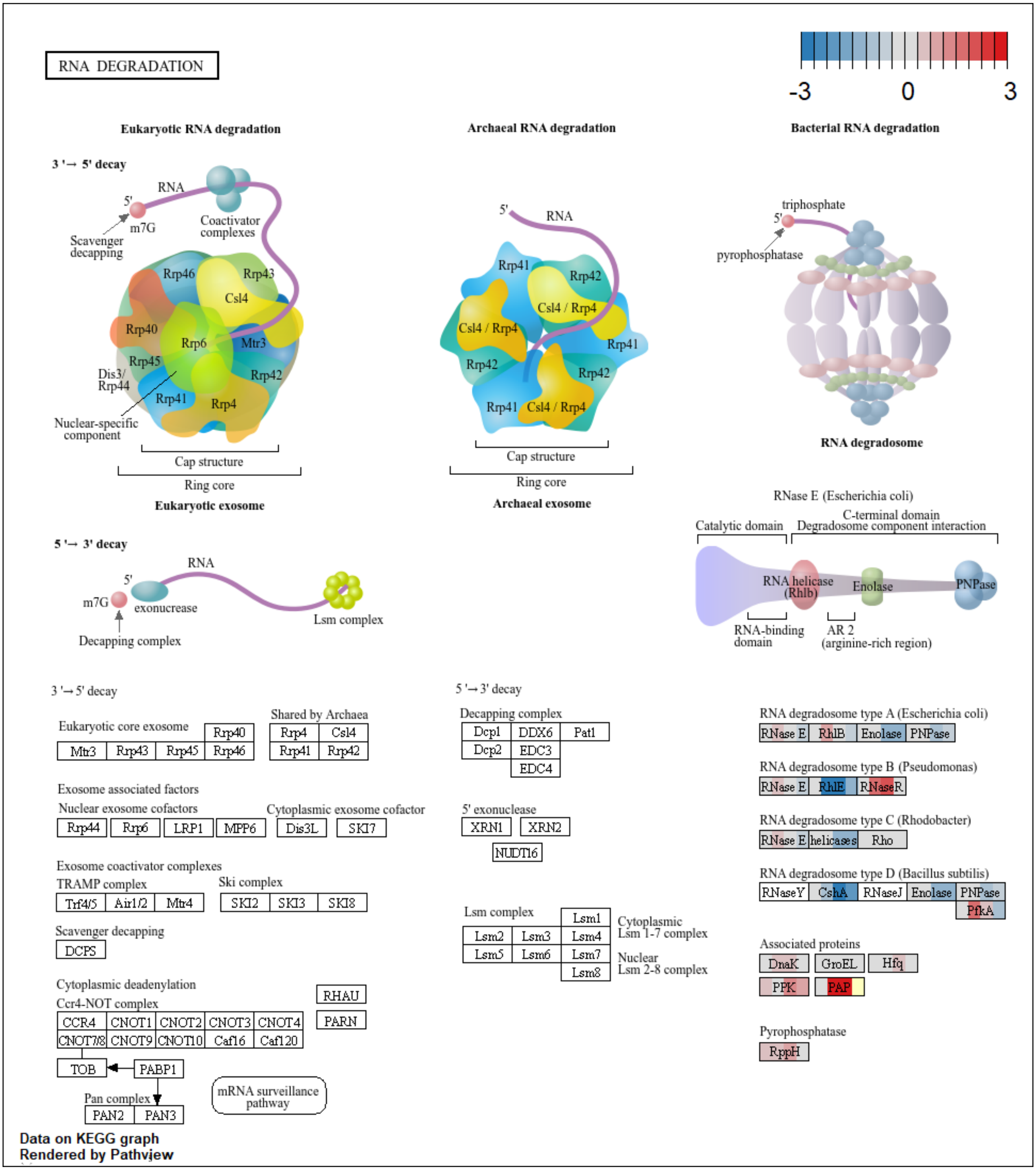
RNA DEGRADATION
-3
0
3
Eukaryotic RNA degradation
Archaeal RNA degradation
Bacterial RNA degradation
3 '→ 5' decay
RNA
5'
m7G
Scavenger decapping
Coactivator complexes
Rrp46
Rrp43
Csl4
Rrp40
Rrp6
Mtr3
Rrp45
Dis3/ Rrp44
Rrp42
Rrp41
Rrp4
Nuclear-specific component
Cap structure
Ring core
Eukaryotic exosome
Csl4 / Rrp4
Archaeal exosome
triphosphate
pyrophosphatase
RNA degradosome
RNase E (Escherichia coli)
Catalytic domain
C-terminal domain
Degradosome component interaction
RNA helicase (Rhlb)
Enolase
PNPase
RNA-binding domain
AR 2 (arginine-rich region)
5 '→ 3' decay
exonucrease
Lsm complex
Decapping complex
3 '→ 5' decay
Eukaryotic core exosome
Shared by Archaea
Rrp40
Rrp4
Csl4
Mtr3
Rrp43
Rrp45
Rrp46
Rrp41
Rrp42
Exosome associated factors
Nuclear exosome cofactors
Cytoplasmic exosome cofactor
Rrp44
Rrp6
LRP1
MPP6
Dis3L
SKI7
Exosome coactivator complexes
TRAMP complex
Ski complex
Trf4/5
Air1/2
Mtr4
SKI2
SKI3
SKI8
Scavenger decapping
DCPS
Cytoplasmic deadenylation
Ccr4-NOT complex
CCR4
CNOT1
CNOT2
CNOT3
CNOT4
CNOT7/8
CNOT9
CNOT10
Caf16
Caf120
RHAU
PARN
TOB
PABP1
Pan complex
PAN2
PAN3
mRNA surveillance pathway
5 '→ 3' decay
Decapping complex
Dcp1
DDX6
Pat1
Dcp2
EDC3
EDC4
5' exonuclease
XRN1
XRN2
NUDT16
Lsm complex
Lsm1
Lsm2
Lsm3
Lsm4
Lsm5
Lsm6
Lsm7
Lsm8
Cytoplasmic Lsm 1-7 complex
Nuclear Lsm 2-8 complex
RNA degradosome type A (Escherichia coli)
RNase E
RhlB
Enolase
PNPase
RNA degradosome type B (Pseudomonas)
RNase E
RhlE
RNaseR
RNA degradosome type C (Rhodobacter)
RNase E
helicases
Rho
RNA degradosome type D (Bacillus subtilis)
RNaseY
CshA
RNaseJ
Enolase
PNPase
PfkA
Associated proteins
DnaK
GroEL
Hfq
PPK
PAP
Pyrophosphatase
RppH
Data on KEGG graph
Rendered by Pathview

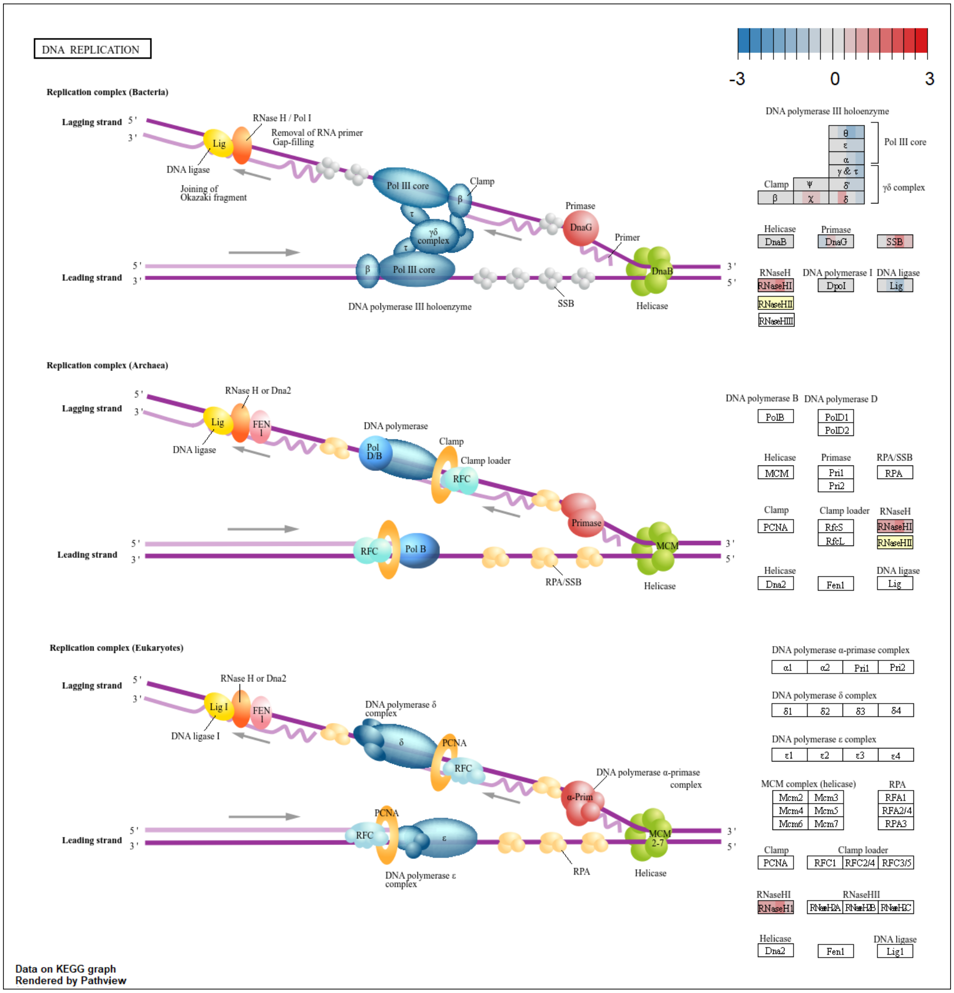
DNA REPLICATION
-3
0
3
Replication complex (Bacteria)
Lagging strand
Leading strand
RNase H / Pol I
Removal of RNA primer
Gap-filling
DNA ligase
Joining of
Okazaki fragment
Pol III core
Clamp
Primase
DnaG
Primer
DnaB
DNA polymerase III holoenzyme
SSB
Helicase
Pol III core
γδ complex
Replication complex (Archaea)
RNase H or Dna2
DNA ligase
DNA polymerase
Clamp
Clamp loader
RFC
Primase
MCM
RPA/SSB
Pol B
DNA polymerase B
DNA polymerase D
Replication complex (Eukaryotes)
RNase H or Dna2
DNA ligase I
DNA polymerase δ complex
PCNA
DNA polymerase α-primase complex
α-Prim
DNA polymerase ε complex
RPA
MCM complex (helicase)
RNaseH
RNaseHII
RNaseHIII
DNA polymerase I
DNA ligase
Data on KEGG graph
Rendered by Pathview

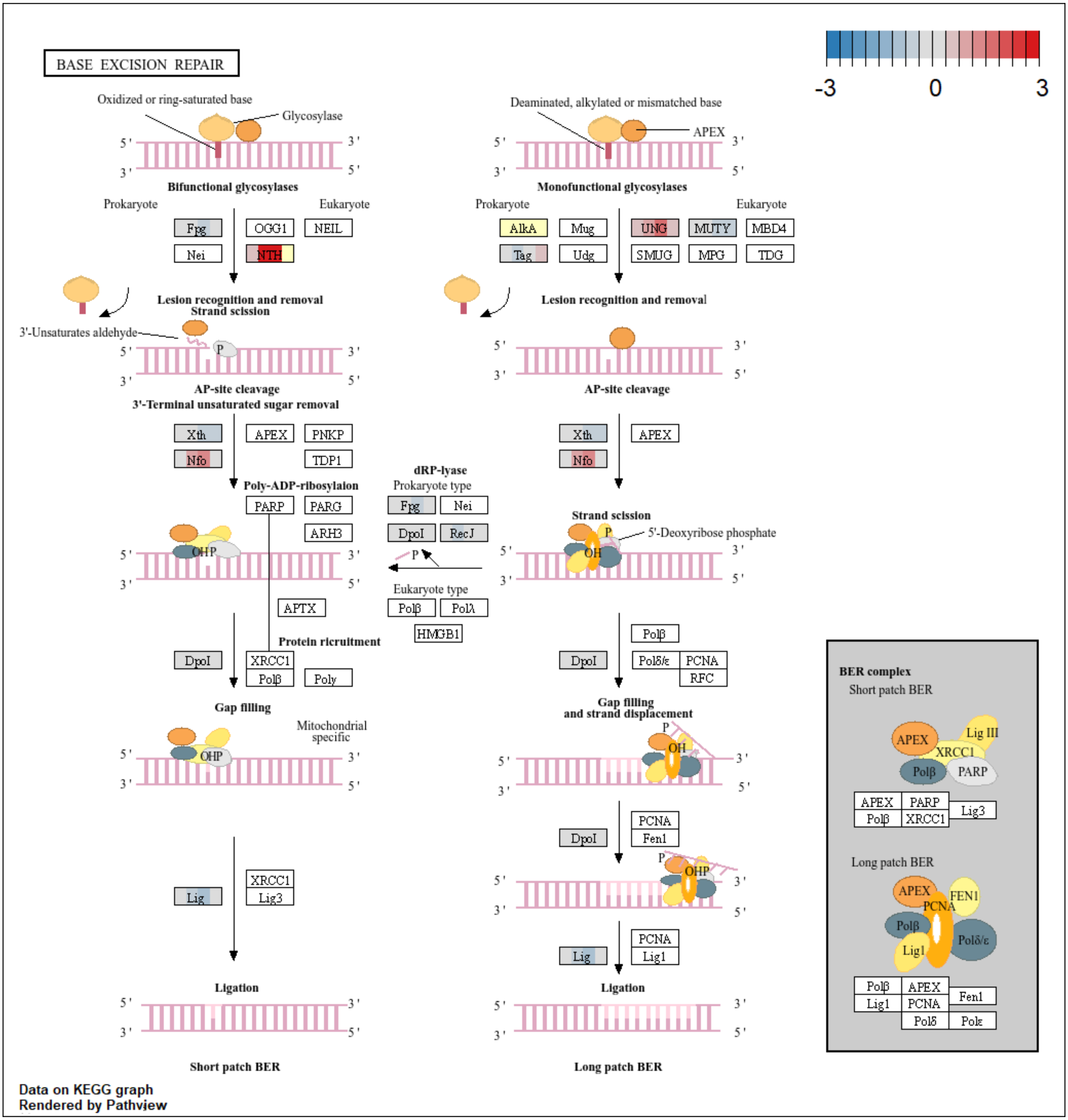
BASE EXCISION REPAIR
-3
0
3
Oxidized or ring-saturated base
Glycosylase
Bifunctional glycosylases
Prokaryote
Eukaryote
Fpg
Nei
OGG1
NEIL
NTH
Deaminated, alkylated or mismatched base
APEX
Monofunctional glycosylases
AlkA
Mug
Tag
Udg
UNG
MUTY
MBD4
SMUG
MPG
TDG
Lesion recognition and removal
Strand scission
3'-Unsaturates aldehyde
AP-site cleavage
3'-Terminal unsaturated sugar removal
Xth
Nfo
APEX
PNKP
TDP1
Poly-ADP-ribosylaion
PARP
PARG
ARH3
dRP-lyase
Prokaryote type
Fpg
Nei
DpoI
RecJ
Eukaryote type
Polβ
Polλ
HMGB1
Strand scission
5'-Deoxyribose phosphate
APTX
Protein ricruitment
DpoI
XRCC1
Polβ
Polγ
Gap filling
Mitochondrial specific
Polβ
Polδ/ε
PCNA
RFC
Gap filling and strand displacement
PCNA
Fen1
Lig
XRCC1
Lig3
Lig1
Ligation
Short patch BER
Long patch BER
BER complex
Short patch BER
APEX
Lig III
XRCC1
Polβ
PARP
Lig3
Long patch BER
FEN1
PCNA
Polδ/ε
Lig1
Fen1
Polδ
Polε
Data on KEGG graph
Rendered by Pathview

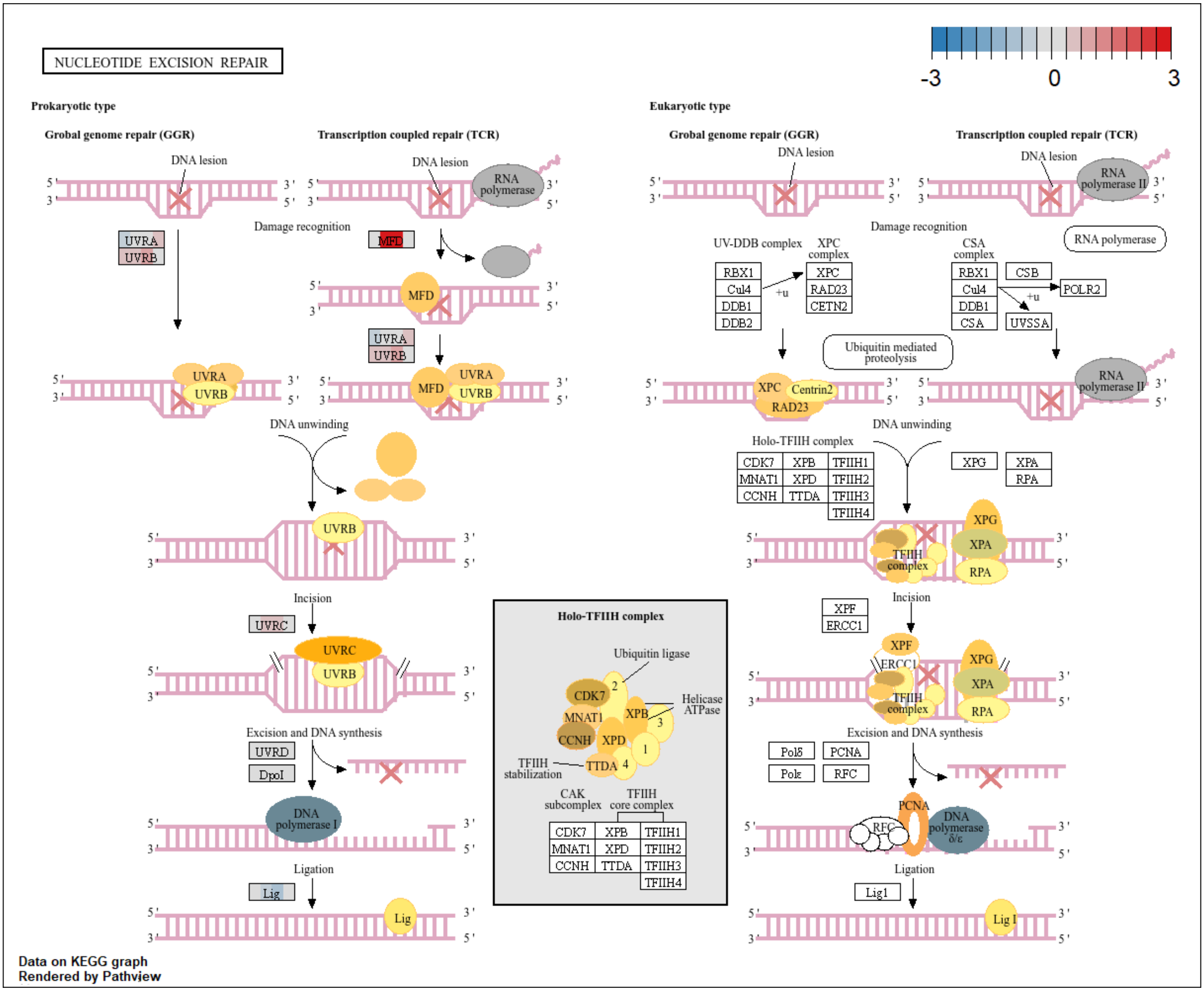
NUCLEOTIDE EXCISION REPAIR
-3
0
3
Prokaryotic type
Grobal genome repair (GGR)
Transcription coupled repair (TCR)
Eukaryotic type
DNA lesion
RNA polymerase
RNA polymerase II
Damage recognition
UVRA
UVRB
MFD
UVRC
UVRD
DpoI
DNA polymerase I
Lig
DNA unwinding
Incision
Excision and DNA synthesis
Ligation
UV-DDB complex
XPC complex
CSA complex
RBX1
Cul4
DDB1
DDB2
XPC
RAD23
CETN2
CSB
POLR2
CSA
UVSSA
+u
Ubiquitin mediated proteolysis
Centrin2
Holo-TFIIH complex
CDK7
XPB
TFIIH1
MNAT1
XPD
TFIIH2
CCNH
TTDA
TFIIH3
TFIIH4
XPG
XPA
RPA
TFIIH complex
XPF
ERCC1
Polδ
PCNA
Polε
RFC
DNA polymerase δ/ε
Lig1
Lig I
Ubiquitin ligase
Helicase ATPase
TFIIH stabilization
CAK subcomplex
TFIIH core complex
Data on KEGG graph
Rendered by Pathview

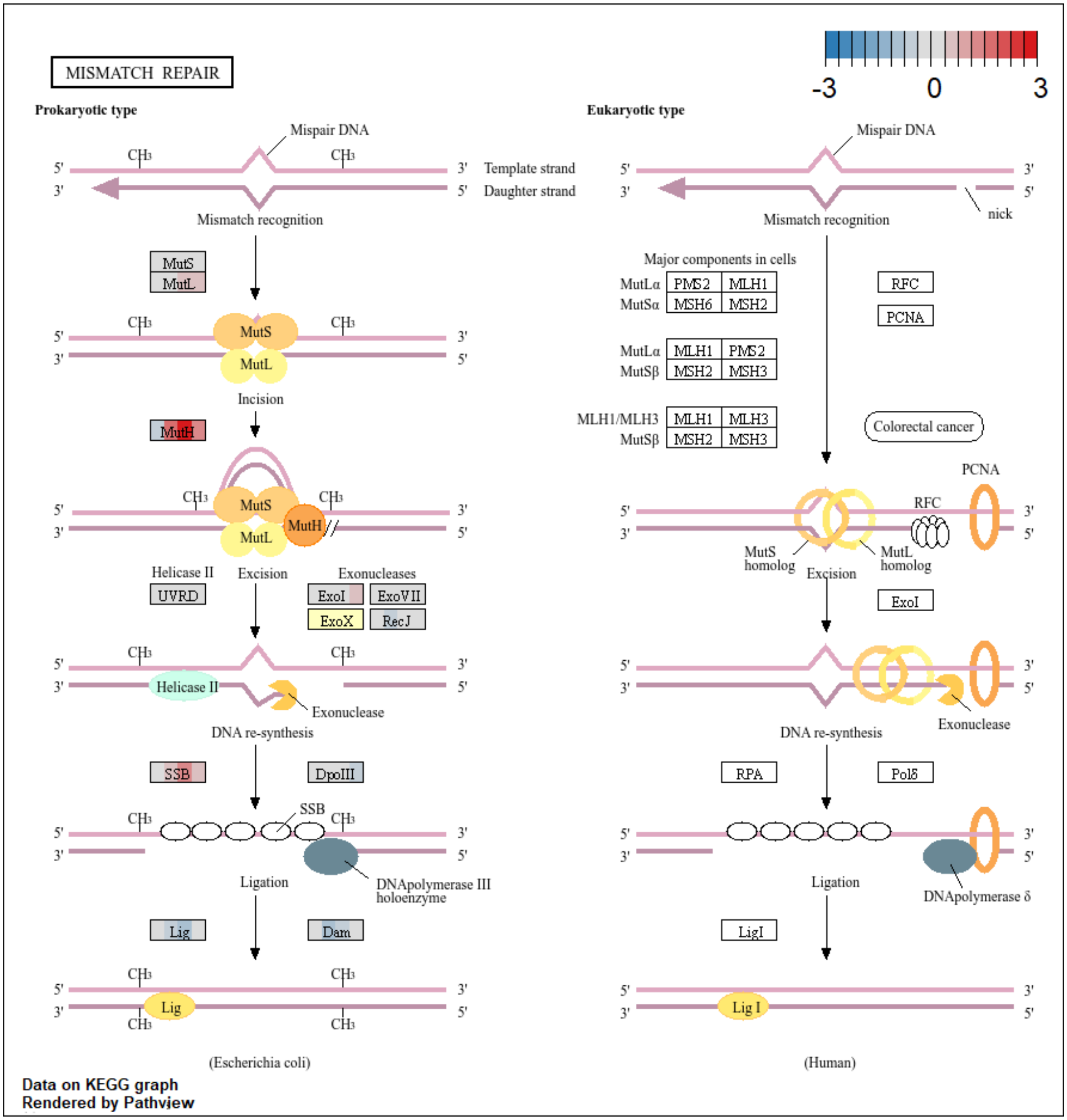
MISMATCH REPAIR
-3
0
3
Prokaryotic type
Eukaryotic type
Mispair DNA
Template strand
Daughter strand
Mismatch recognition
nick
MutS
MutL
Major components in cells
MutLα PMS2 MLH1
MutSα MSH6 MSH2
MutLα MLH1 PMS2
MutSβ MSH2 MSH3
MLH1/MLH3 MLH1 MLH3
MutSβ MSH2 MSH3
RFC
PCNA
Colorectal cancer
Incision
MutH
MutS homolog
MutL homolog
Helicase II
UVRD
Excision
Exonucleases
ExoI
ExoVII
ExoX
RecJ
Exonuclease
DNA re-synthesis
SSB
DpoIII
RPA
Polδ
DNApolymerase III holoenzyme
DNApolymerase δ
Ligation
Lig
Dam
LigI
Lig I
(Escherichia coli)
(Human)
Data on KEGG graph
Rendered by Pathview

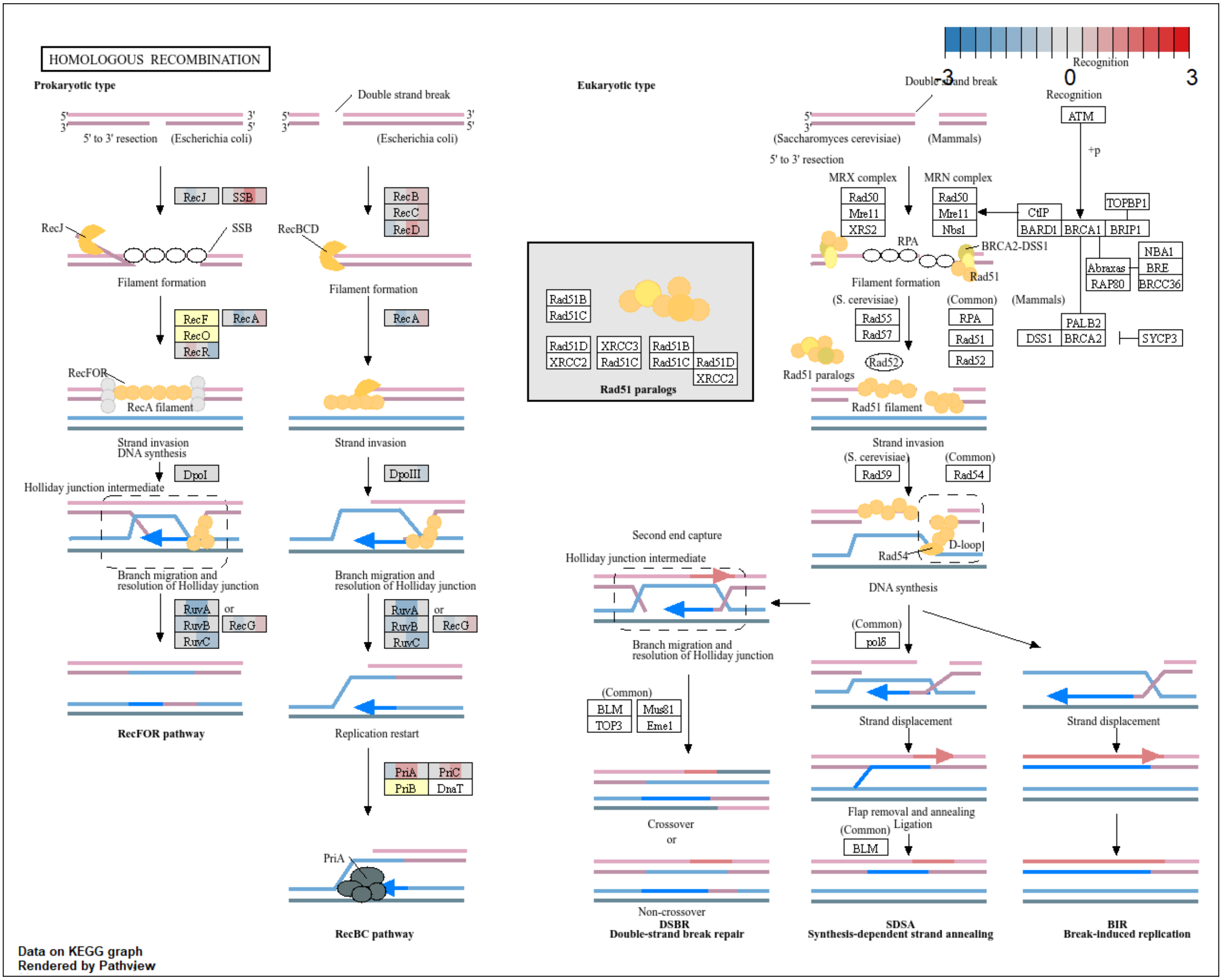
HOMOLOGOUS RECOMBINATION
-3
0
3
Recognition
Prokaryotic type
Eukaryotic type
Double strand break
5' to 3' resection
(Escherichia coli)
(Saccharomyces cerevisiae)
(Mammals)
Recognition
ATM
+p
MRX complex
MRN complex
Rad50
Mre11
XRS2
Nbs1
CtIP
BARD1
BRCA1
BRIP1
TOPBP1
NBA1
Abraxas
BRE
RAP80
BRCC36
RPA
BRCA2-DSS1
Rad51
RecJ
SSB
RecB
RecC
RecD
RecBCD
Filament formation
(S. cerevisiae)
(Common)
Rad55
Rad57
Rad52
PALB2
DSS1
BRCA2
SYCP3
RecF
RecO
RecR
RecA
RecFOR
RecA filament
Rad51B
Rad51C
Rad51D
XRCC2
XRCC3
Rad51 paralogs
Rad51 filament
Strand invasion
DNA synthesis
Rad59
Rad54
DpoI
DpoIII
Holliday junction intermediate
D-loop
Second end capture
Branch migration and resolution of Holliday junction
RuvA
RuvB
RuvC
or
RecG
pol δ
BLM
TOP3
Mus81
Eme1
Strand displacement
Replication restart
PriA
PriC
PriB
DnaT
Crossover
Non-crossover
Flap removal and annealing
Ligation
RecFOR pathway
RecBC pathway
DSBR
Double-strand break repair
SDSA
Synthesis-dependent strand annealing
BIR
Break-induced replication
Data on KEGG graph
Rendered by Pathview

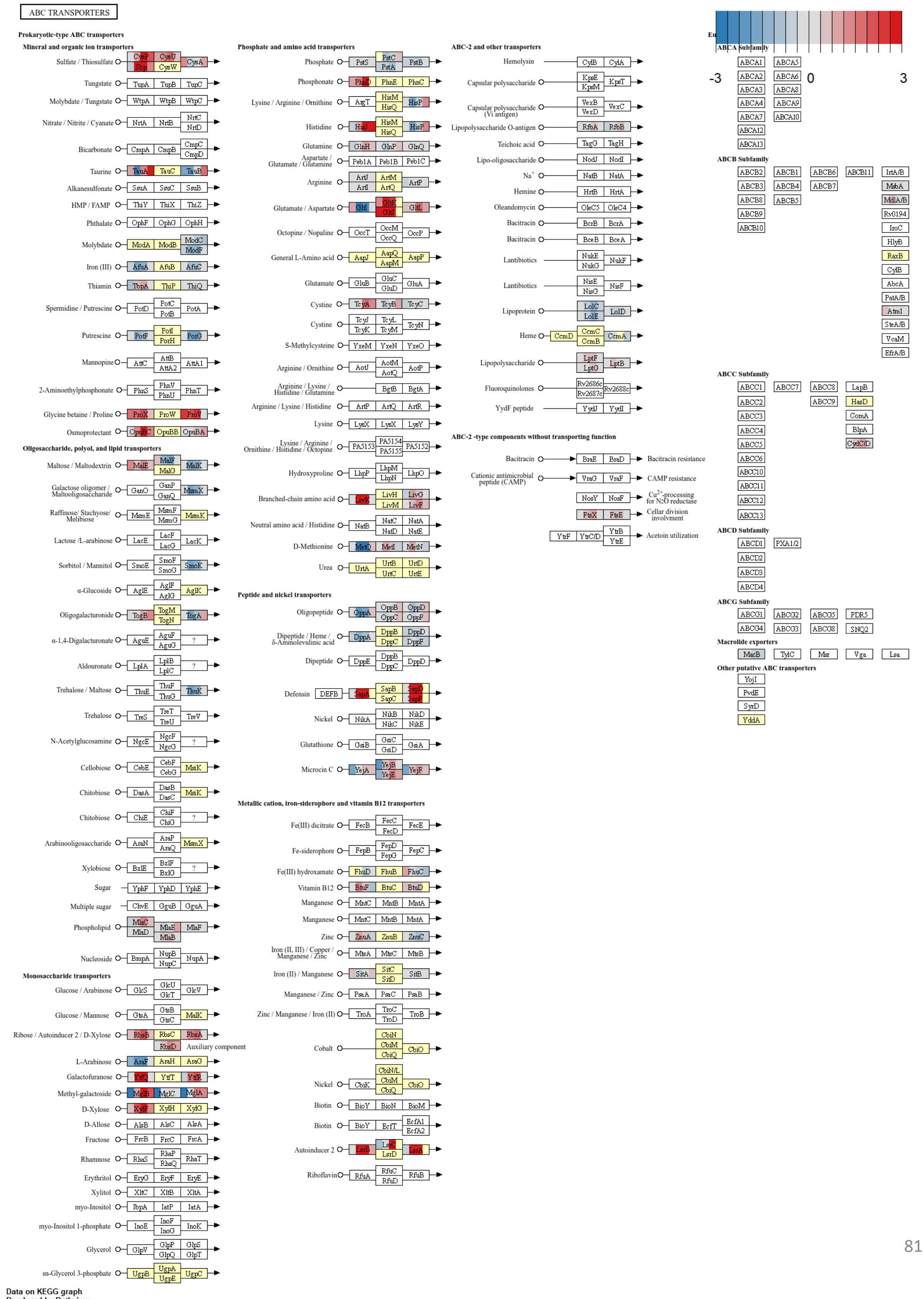

ABC TRANSPORTERS
-3
0
3
Prokaryotic-type ABC transporters
Mineral and organic ion transporters
Oligosaccharide, polyol, and lipid transporters
Monosaccharide transporters
Phosphate and amino acid transporters
Peptide and nickel transporters
Metallic cation, iron-siderophore and vitamin B12 transporters
ABC-2 and other transporters
ABC-2 -type components without transporting function
ABCA Subfamily
ABCB Subfamily
ABCC Subfamily
ABCD Subfamily
ABCG Subfamily
Macrolide exporters
Other putative ABC transporters
Data on KEGG graph
Rendered by Pathview

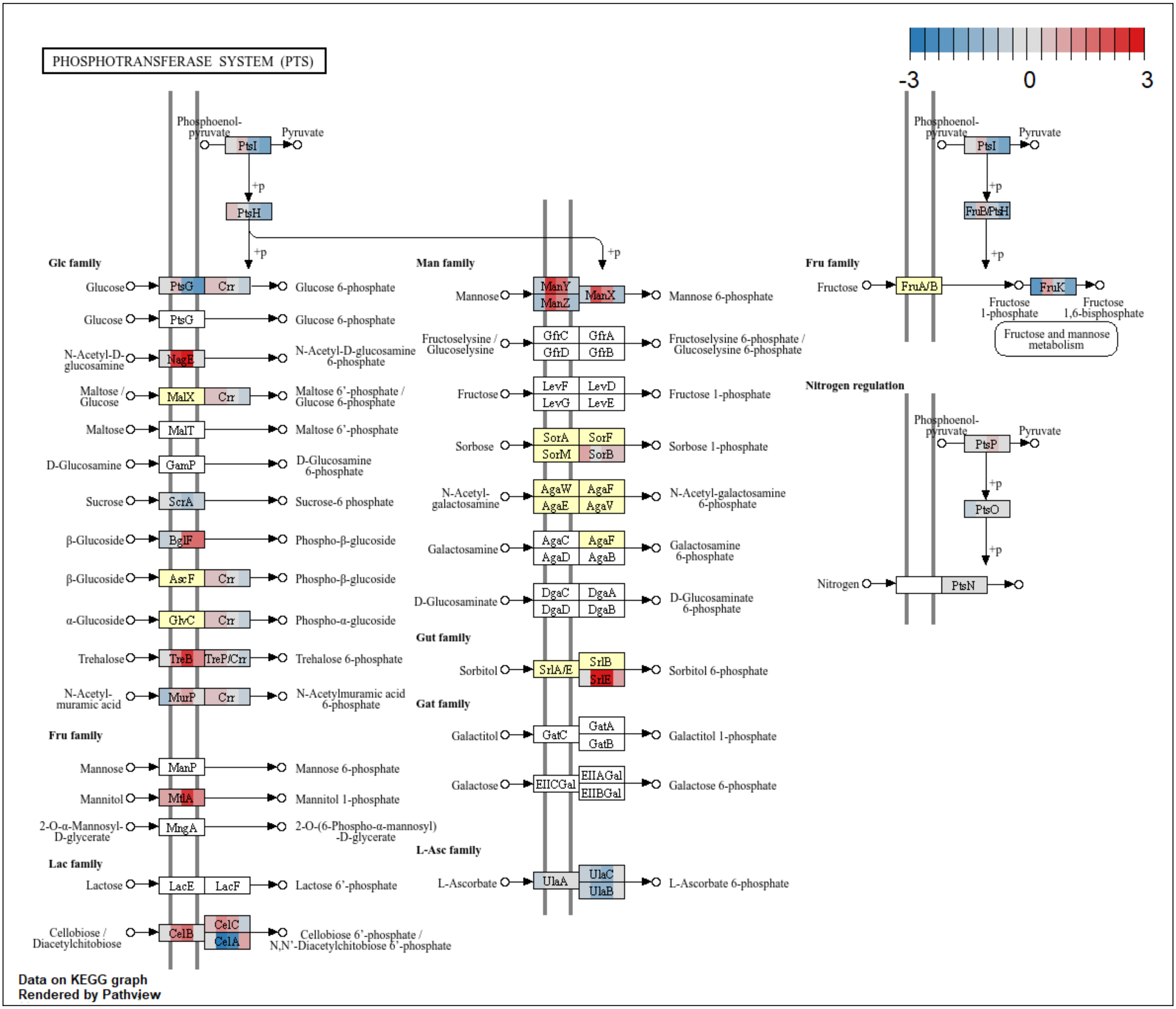
PHOSPHOTRANSFERASE SYSTEM (PTS)
-3
0
3
Phosphoenol-pyruvate
Pyruvate
PtsI
+p
PtsH
Glc family
Glucose
PtsG
Crr
Glucose 6-phosphate
NagE
N-Acetyl-D-glucosamine
N-Acetyl-D-glucosamine 6-phosphate
Maltose / Glucose
MalX
Maltose 6'-phosphate / Glucose 6-phosphate
Maltose
MalT
Maltose 6'-phosphate
D-Glucosamine
GamP
D-Glucosamine 6-phosphate
Sucrose
ScrA
Sucrose-6 phosphate
β-Glucoside
BglF
Phospho-β-glucoside
AscF
α-Glucoside
GlvC
Phospho-α-glucoside
Trehalose
TreB
TreP/Crr
Trehalose 6-phosphate
N-Acetyl-muramic acid
MurP
N-Acetylmuramic acid 6-phosphate
Fru family
Mannose
ManP
Mannose 6-phosphate
Mannitol
MtlA
Mannitol 1-phosphate
2-O-α-Mannosyl-D-glycerate
MngA
2-O-(6-Phospho-α-mannosyl)-D-glycerate
Lac family
Lactose
LacE
LacF
Lactose 6'-phosphate
Cellobiose / Diacetylchitobiose
CelB
CelC
CelA
Cellobiose 6'-phosphate / N,N'-Diacetylchitobiose 6'-phosphate
Man family
ManY
ManZ
ManX
Fructoselysine / Glucoselysine
GfrC
GfrD
GfrA
GfrB
Fructoselysine 6-phosphate / Glucoselysine 6-phosphate
Fructose
LevF
LevG
LevD
LevE
Fructose 1-phosphate
Sorbose
SorA
SorM
SorF
SorB
Sorbose 1-phosphate
N-Acetyl-galactosamine
AgaW
AgaE
AgaF
AgaV
N-Acetyl-galactosamine 6-phosphate
Galactosamine
AgaC
AgaD
AgaB
Galactosamine 6-phosphate
D-Glucosaminate
DgaC
DgaD
DgaA
DgaB
D-Glucosaminate 6-phosphate
Gut family
Sorbitol
SrlA/E
SrlB
SrlE
Sorbitol 6-phosphate
Gat family
Galactitol
GatC
GatA
GatB
Galactitol 1-phosphate
Galactose
EIICGal
EIIAGal
EIIBGal
Galactose 6-phosphate
L-Asc family
L-Ascorbate
UlaA
UlaC
UlaB
L-Ascorbate 6-phosphate
FruB/PtsH
FruA/B
FruK
Fructose 1-phosphate
Fructose 1,6-bisphosphate
Fructose and mannose metabolism
Nitrogen regulation
PtsP
PtsO
Nitrogen
PtsN
Data on KEGG graph
Rendered by Pathview

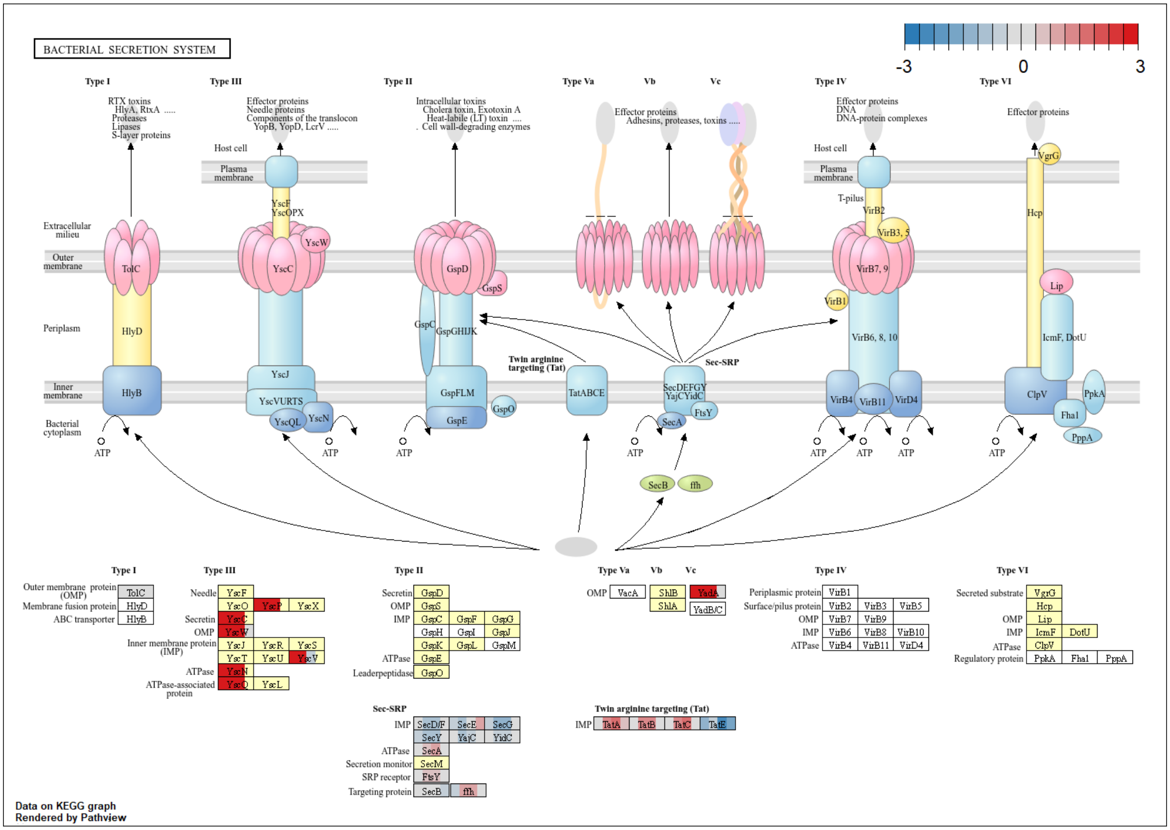

BACTERIAL SECRETION SYSTEM
-3
0
3
Type I
Type III
Type II
Type Va
Vb
Vc
Type IV
Type VI
Twin arginine targeting (Tat)
Sec-SRP
Data on KEGG graph
Rendered by Pathview

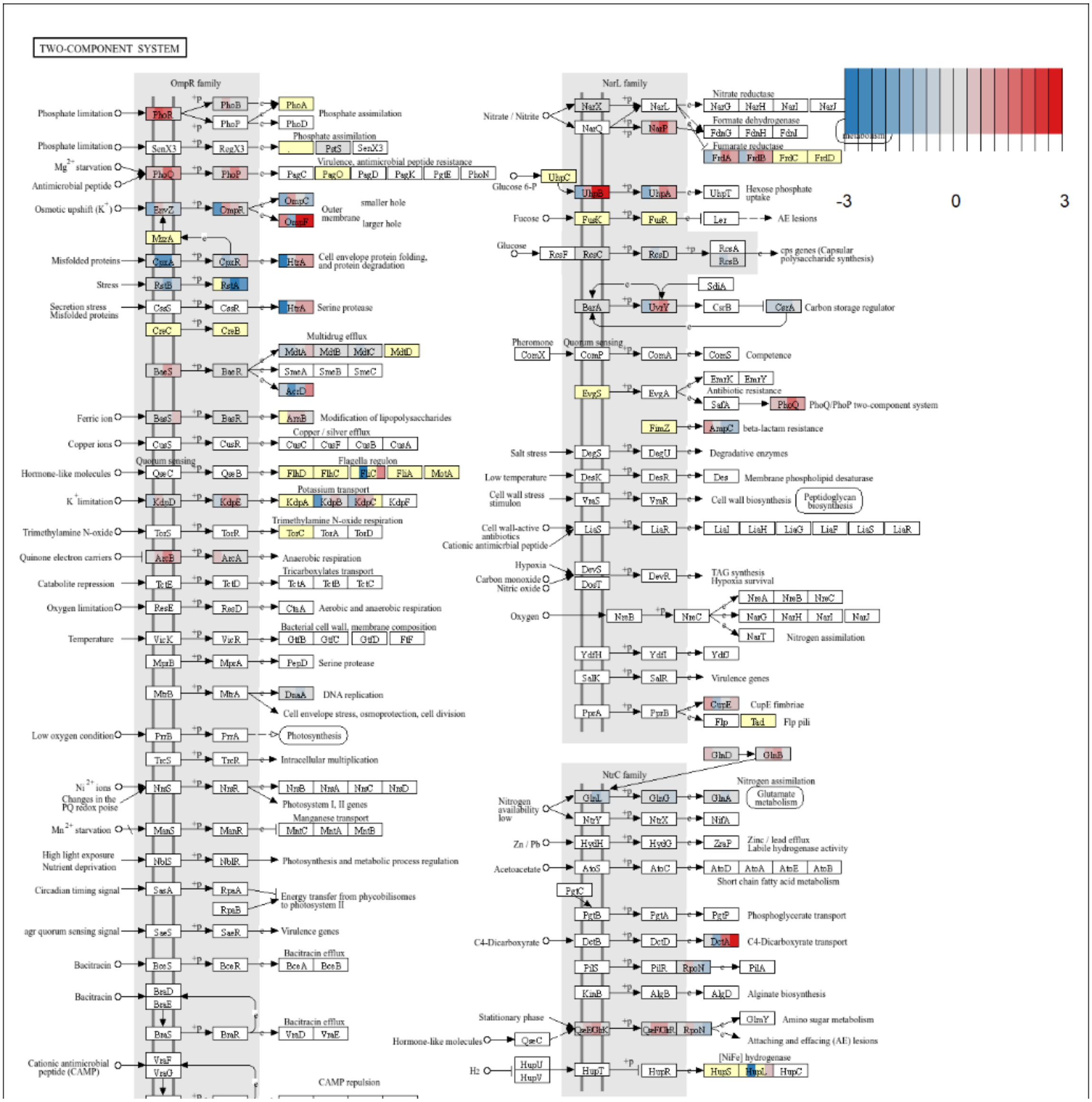
TWO-COMPONENT SYSTEM
OmpR family
NarL family
NtrC family
-3
0
3
Phosphate limitation
PhoR
PhoB
PhoP
PhoA
PhoD
Phosphate assimilation
SenX3
RegX3
PstS
Mg2+ starvation
Antimicrobial peptide
PhoQ
Virulence, antimicrobial peptide resistance
PagC
PagO
PagD
PagK
PgtE
PhoN
Osmotic upshift (K+)
EnvZ
OmpR
OmpC
OmpF
Outer membrane
smaller hole
larger hole
MzrA
Misfolded proteins
CpxA
CpxR
HtrA
Cell envelope protein folding, and protein degradation
Stress
RstB
RstA
Secretion stress
Misfolded proteins
CssS
CssR
Serine protease
CreC
CreB
Multidrug efflux
MdtA
MdtB
MdtC
MdtD
BaeS
BaeR
SmeA
SmeB
SmeC
AcrD
Ferric ion
BasS
BasR
ArnB
Modification of lipopolysaccharides
Copper ions
CusS
CusR
Copper / silver efflux
CusC
CusF
CusB
CusA
Quorum sensing
Hormone-like molecules
QseC
QseB
Flagella regulon
FlhD
FlhC
FliC
FliA
MotA
K+ limitation
KdpD
KdpE
Potassium transport
KdpA
KdpB
KdpC
KdpF
Trimethylamine N-oxide
TorS
TorR
Trimethylamine N-oxide respiration
TorC
TorA
TorD
Quinone electron carriers
ArcB
ArcA
Anaerobic respiration
Catabolite repression
TctE
TctD
Tricarboxylates transport
TctA
TctB
TctC
Oxygen limitation
ResE
ResD
CtaA
Aerobic and anaerobic respiration
Temperature
VicK
VicR
Bacterial cell wall, membrane composition
MprB
MprA
PepD
Serine protease
MtrB
MtrA
DnaA
DNA replication
Cell envelope stress, osmoprotection, cell division
Low oxygen condition
PrrB
PrrA
Photosynthesis
TrcS
TrcR
Intracellular multiplication
Ni2+ ions
Changes in the PQ redox poise
NrsS
NrsR
NrsB
NrsA
NrsC
NrsD
Photosystem I, II genes
Mn2+ starvation
ManS
ManR
Manganese transport
MntC
MntA
MntB
High light exposure
Nutrient deprivation
NblS
NblR
Photosynthesis and metabolic process regulation
Circadian timing signal
SasA
RpaA
RpaB
Energy transfer from phycobilisomes to photosystem II
agr quorum sensing signal
SaeS
SaeR
Virulence genes
Bacitracin
BceS
BceR
Bacitracin efflux
BceA
BceB
BraD
BraE
BraS
BraR
VraD
VraE
Cationic antimicrobial peptide (CAMP)
VraF
VraG
CAMP repulsion
Nitrate / Nitrite
NarX
NarQ
NarL
NarP
Nitrate reductase
NarG
NarH
NarI
NarJ
Formate dehydrogenase
FdnG
FdnH
FdnI
Fumarate reductase
FrdA
FrdB
FrdC
FrdD
UhpC
Glucose 6-P
UhpB
UhpA
UhpT
Hexose phosphate uptake
Fucose
FusK
FusR
Ler
AE lesions
Glucose
RcsF
RcsC
RcsD
RcsA
RcsB
cps genes (Capsular polysaccharide synthesis)
SdiA
BarA
UvrY
CsrB
CsrA
Carbon storage regulator
Pheromone
ComX
ComP
ComA
ComS
Competence
EvgS
EvgA
EmrK
EmrY
Antibiotic resistance
SafA
PhoQ/PhoP two-component system
FimZ
AmpC
beta-lactam resistance
Salt stress
DegS
DegU
Degradative enzymes
Low temperature
DesK
DesR
Des
Membrane phospholipid desaturase
Cell wall stress stimulon
VraS
VraR
Cell wall biosynthesis
Peptidoglycan biosynthesis
Cell wall-active antibiotics
Cationic antimicrobial peptide
LiaS
LiaR
LiaI
LiaH
LiaG
LiaF
Hypoxia
Carbon monoxide
Nitric oxide
DevS
DosT
DevR
TAG synthesis
Hypoxia survival
Oxygen
NreB
NreC
NreA
Nitrogen assimilation
YdfH
YdfI
YdfJ
SalK
SalR
PprA
PprB
CupE
CupE fimbriae
Flp
Tad
Flp pili
GlnD
GlnB
Nitrogen assimilation
Nitrogen availability low
GlnL
GlnG
GlnA
Glutamate metabolism
NtrY
NtrX
NifA
Zn / Pb
HydH
HydG
ZraP
Zinc / lead efflux
Labile hydrogenase activity
Acetoacetate
AtoS
AtoC
AtoD
AtoA
AtoE
AtoB
Short chain fatty acid metabolism
PgtC
PgtB
PgtA
PgtP
Phosphoglycerate transport
C4-Dicarboxylate
DctB
DctD
DctA
C4-Dicarboxylate transport
PilS
PilR
RpoN
PilA
KinB
AlgB
AlgD
Alginate biosynthesis
Stationary phase
Hormone-like molecules
QseC
GlmY
Amino sugar metabolism
Attaching and effacing (AE) lesions
H2
HupU
HupV
HupT
HupR
[NiFe] hydrogenase
HupS
HupL
HupC

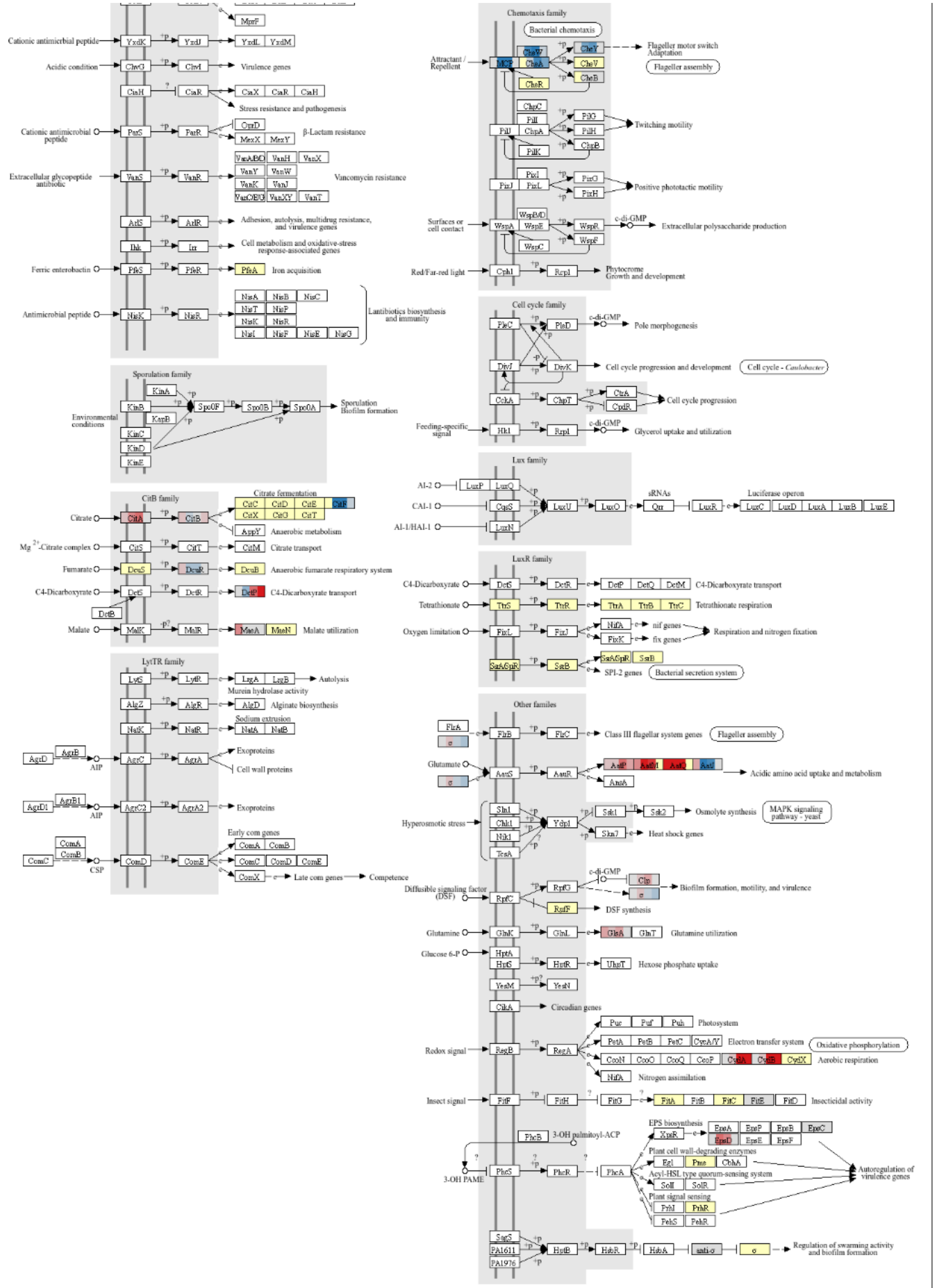

Data on KEGG graph
Rendered by Pathview

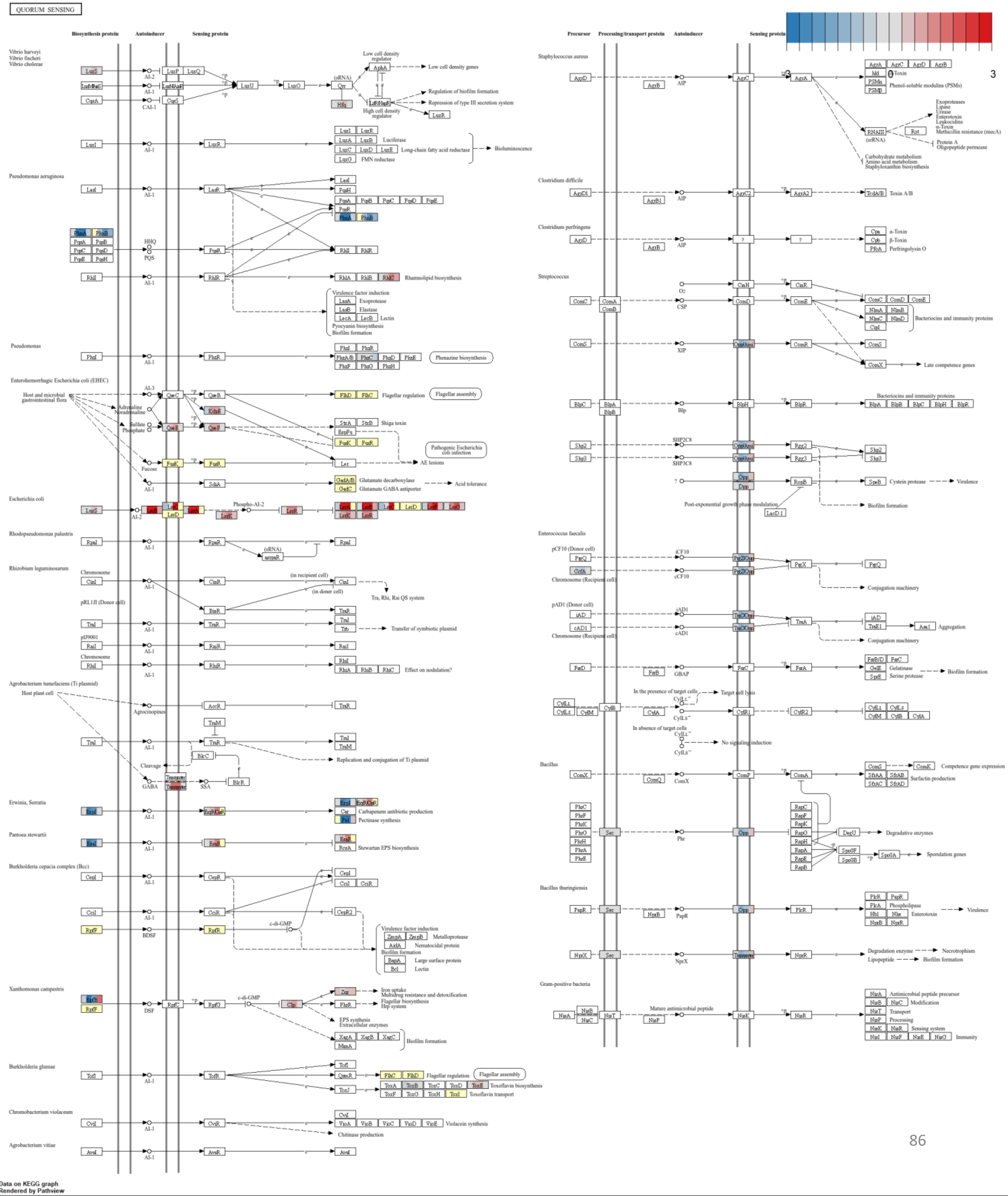

QUORUM SENSING
Biosynthesis protein
Autoinducer
Sensing protein
Vibrio harveyi
Vibrio fischeri
Vibrio cholerae
Low cell density regulator
Low cell density genes
Regulation of biofilm formation
Repression of type III secretion system
High cell density regulator
Luciferase
Long-chain fatty acid reductase
FMN reductase
Bioluminescence
Pseudomonas aeruginosa
Rhamnolipid biosynthesis
Virulence factor induction
Exoprotease
Elastase
Lectin
Pyocyanin biosynthesis
Biofilm formation
Pseudomonas
Phenazine biosynthesis
Enterohemorrhagic Escherichia coli (EHEC)
Host and microbial gastrointestinal flora
Adrenaline
Noradrenaline
Sulfate
Phosphate
Fucose
Flagellar regulation
Flagellar assembly
Shiga toxin
Pathogenic Escherichia coli infection
AE lesions
Glutamate decarboxylase
Glutamate GABA antiporter
Acid tolerance
Escherichia coli
Phospho-AI-2
Rhodopseudomonas palustris
Rhizobium leguminosarum
Chromosome
(in recipient cell)
(in donor cell)
Tra, Rhi, Rai QS system
pRL1JI (Donor cell)
Transfer of symbiotic plasmid
pIJ9001
Effect on nodulation?
Agrobacterium tumefaciens (Ti plasmid)
Host plant cell
Agrocinopines
Replication and conjugation of Ti plasmid
Cleavage
GABA
SSA
Erwinia, Serratia
Carbapenem antibiotic production
Pectinase synthesis
Pantoea stewartii
Stewartan EPS biosynthesis
Burkholderia cepacia complex (Bcc)
c-di-GMP
BDSF
Virulence factor induction
Metalloprotease
Nematocidal protein
Biofilm formation
Large surface protein
Lectin
Xanthomonas campestris
DSF
Iron uptake
Multidrug resistance and detoxification
Flagellar biosynthesis
Hrp system
EPS synthesis
Extracellular enzymes
Biofilm formation
Burkholderia glumae
Flagellar regulation
Flagellar assembly
Toxoflavin biosynthesis
Toxoflavin transport
Chromobacterium violaceum
Violacein synthesis
Chitinase production
Agrobacterium vitiae
Precursor
Processing/transport protein
Autoinducer
Sensing protein
Staphylococcus aureus
AIP
Phenol-soluble modulins (PSMs)
Exoproteases
Lipase
Urease
Enterotoxin
Leukocidins
α-Toxin
Methicillin resistance (mecA)
Protein A
Oligopeptide permease
Carbohydrate metabolism
Amino acid metabolism
Staphyloxanthin biosynthesis
Clostridium difficile
Toxin A/B
Clostridium perfringens
α-Toxin
β-Toxin
Perfringolysin O
Streptococcus
CSP
XIP
Bacteriocins and immunity proteins
Late competence genes
Cysteine protease
Virulence
Post-exponential growth phase modulation
Biofilm formation
Enterococcus faecalis
pCF10 (Donor cell)
Chromosome (Recipient cell)
Conjugation machinery
pAD1 (Donor cell)
Aggregation
GBAP
Gelatinase
Serine protease
Biofilm formation
In the presence of target cells
Target cell lysis
In absence of target cells
No signaling induction
Bacillus
Competence gene expression
Surfactin production
Degradative enzymes
Sporulation genes
Bacillus thuringiensis
Phospholipase
Enterotoxin
Virulence
Degradation enzyme
Necrotrophism
Lipopeptide
Gram-positive bacteria
Mature antimicrobial peptide
Antimicrobial peptide precursor
Modification
Transport
Processing
Sensing system
Immunity
Data on KEGG graph
Rendered by Pathview

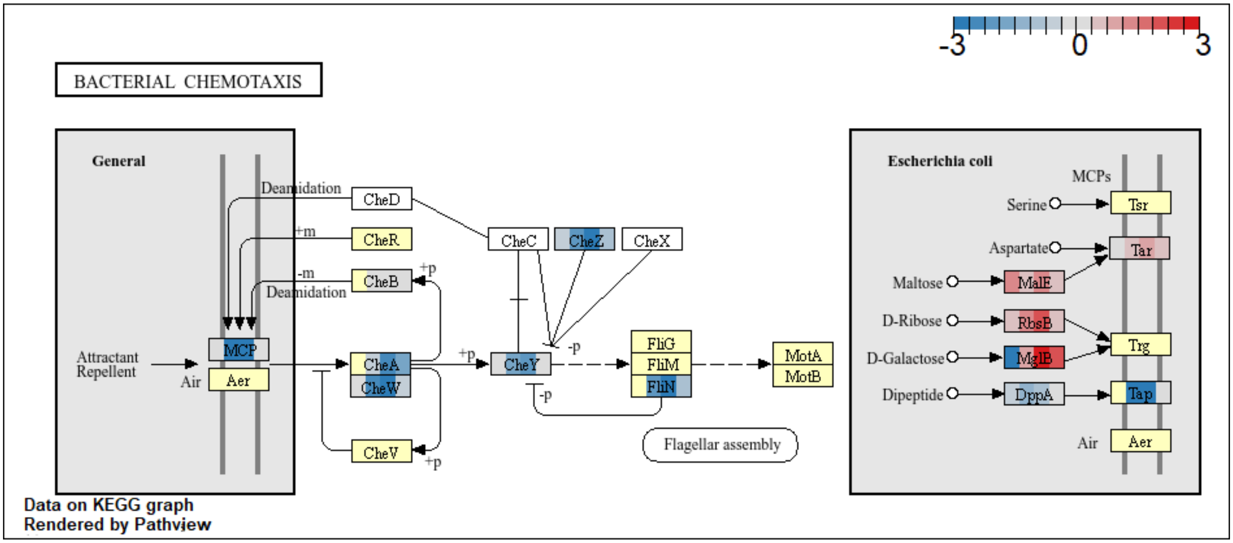
-3
0
3
BACTERIAL CHEMOTAXIS
General
Deamidation
CheD
+m
CheR
-m
CheB
+p
Deamidation
CheC
CheZ
CheX
-p
Attractant
Repellent
MCP
Air
Aer
CheA
CheW
+p
CheY
-p
FliG
FliM
FliN
MotA
MotB
CheV
+p
Flagellar assembly
Escherichia coli
MCPs
Serine
Tsr
Aspartate
Tar
Maltose
MalE
D-Ribose
RbsB
Trg
D-Galactose
MglB
Dipeptide
DppA
Tap
Air
Aer
Data on KEGG graph
Rendered by Pathview

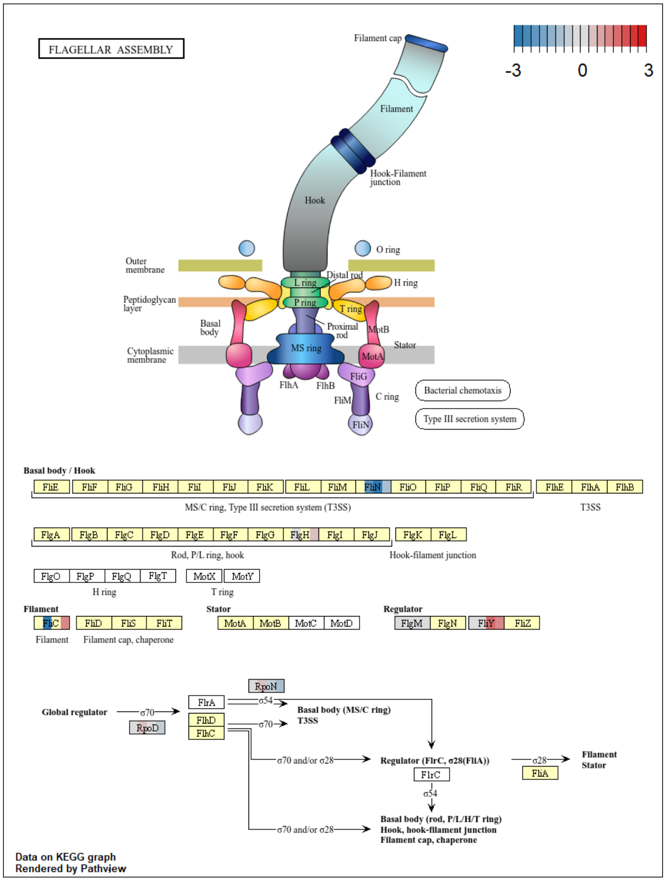

FLAGELLAR ASSEMBLY
-3
0
3
Filament cap
Filament
Hook-Filament junction
Hook
O ring
Outer membrane
Distal rod
L ring
H ring
Peptidoglycan layer
P ring
T ring
Basal body
Proximal rod
MotB
Stator
Cytoplasmic membrane
MS ring
MotA
FliG
FlhA
FlhB
FliM
C ring
FliN
Bacterial chemotaxis
Type III secretion system
Basal body / Hook
FliE
FliF
FliG
FliH
FliI
FliJ
FliK
FliL
FliM
FliN
FliO
FliP
FliQ
FliR
FlhE
FlhA
FlhB
MS/C ring, Type III secretion system (T3SS)
T3SS
FlgA
FlgB
FlgC
FlgD
FlgE
FlgF
FlgG
FlgH
FlgI
FlgJ
FlgK
FlgL
Rod, P/L ring, hook
Hook-filament junction
FlgO
FlgP
FlgQ
FlgT
MotX
MotY
H ring
T ring
Filament
FliC
FliD
FliS
FliT
Filament
Filament cap, chaperone
Stator
MotA
MotB
MotC
MotD
Regulator
FlgM
FlgN
FliY
FliZ
RpoN
FlrA
σ54
Global regulator
σ70
RpoD
FlhD
FlhC
σ70
Basal body (MS/C ring)
T3SS
σ70 and/or σ28
Regulator (FlrC, σ28(FliA))
FlrC
σ54
σ28
FliA
Filament
Stator
σ70 and/or σ28
Basal body (rod, P/L/H/T ring)
Hook, hook-filament junction
Filament cap, chaperone
Data on KEGG graph
Rendered by Pathview

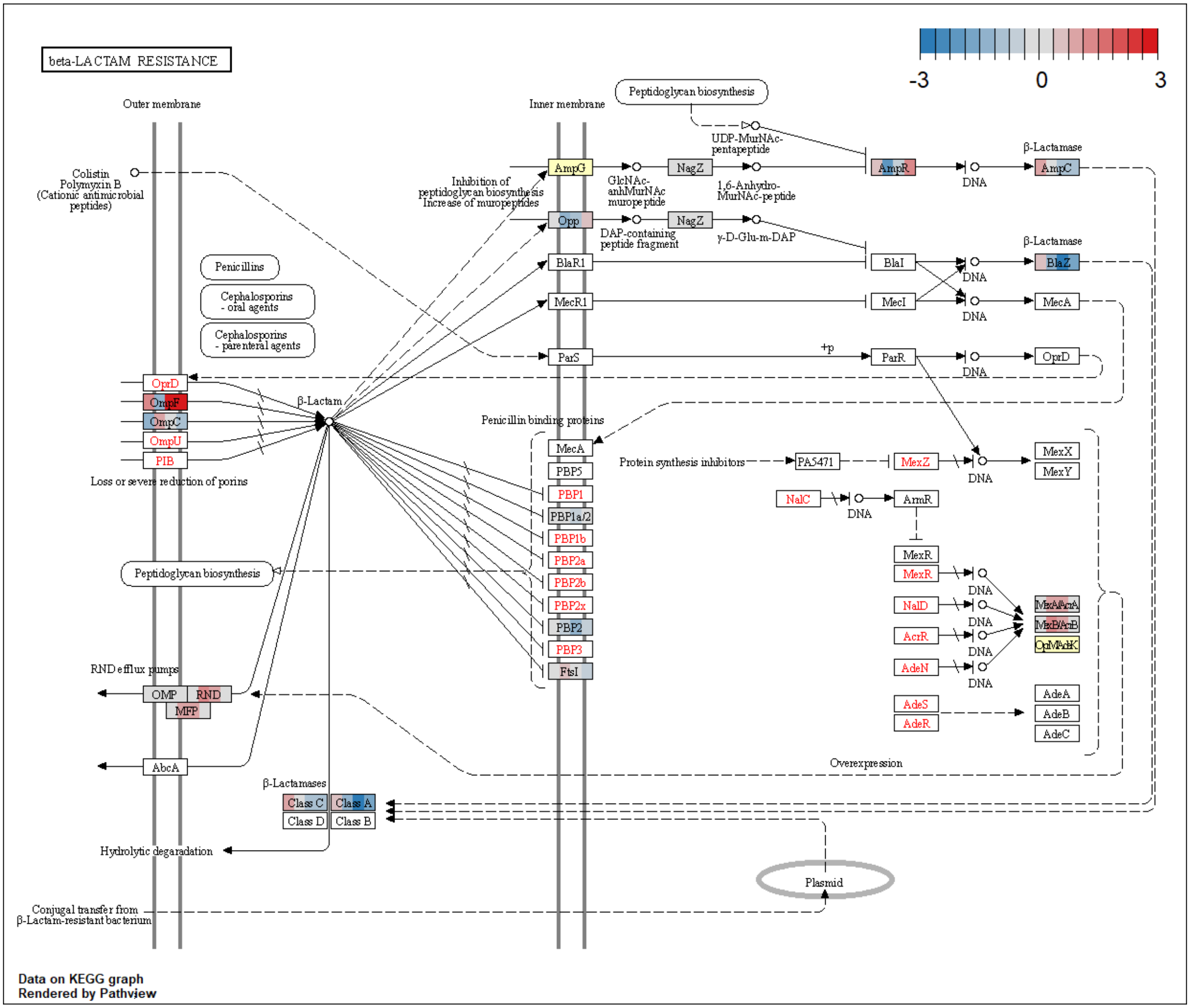
beta-LACTAM RESISTANCE
-3
0
3
Outer membrane
Inner membrane
Peptidoglycan biosynthesis
UDP-MurNAc-pentapeptide
β-Lactamase
AmpG
NagZ
AmpR
AmpC
DNA
Colistin
Polymyxin B
(Cationic antimicrobial peptides)
Inhibition of peptidoglycan biosynthesis
Increase of muropeptides
GlcNAc-anhMurNAc muropeptide
1,6-Anhydro-MurNAc-peptide
Opp
DAP-containing peptide fragment
γ-D-Glu-m-DAP
BlaR1
BlaI
BlaZ
MecR1
MecI
MecA
Penicillins
Cephalosporins - oral agents
Cephalosporins - parenteral agents
ParS
+p
ParR
OprD
OmpF
OmpC
OmpU
PIB
β-Lactam
Penicillin binding proteins
Loss or severe reduction of porins
Protein synthesis inhibitors
PA5471
MexZ
MexX
MexY
PBP5
PBP1
PBP1a/2
PBP1b
PBP2a
PBP2b
PBP2x
PBP2
PBP3
FtsI
NalC
ArmR
MexR
NalD
AcrR
AdeN
AdeS
AdeR
AdeA
AdeB
AdeC
Overexpression
RND efflux pumps
OMP
RND
MFP
AbcA
β-Lactamases
Class C
Class A
Class D
Class B
Hydrolytic degaradation
Plasmid
Conjugal transfer from β-Lactam-resistant bacterium
Data on KEGG graph
Rendered by Pathview

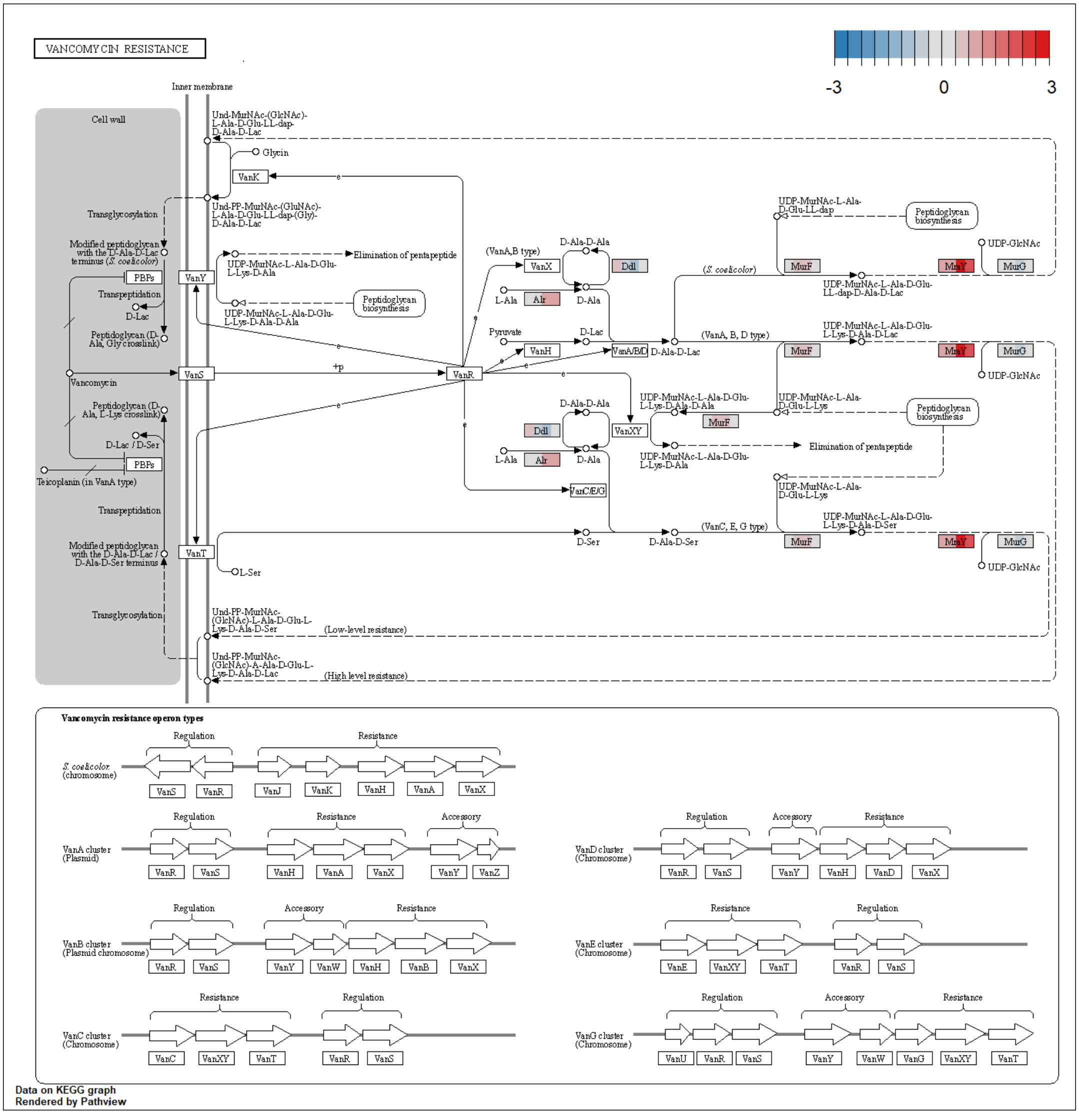
VANCOMYCIN RESISTANCE
-3
0
3
Inner membrane
Cell wall
Transglycosylation
Transpeptidation
Vancomycin
Teicoplanin (in VanA type)
PBPs
VanY
VanS
VanR
VanT
VanK
VanX
VanH
VanA/B/D
VanXY
VanC/E/G
Ddl
Alr
MurF
MraY
MurG
Glycin
Pyruvate
D-Ala-D-Ala
D-Ala-D-Lac
D-Ala-D-Ser
L-Ala
D-Ala
D-Lac
D-Ser
L-Ser
UDP-GlcNAc
Elimination of pentapeptide
Peptidoglycan biosynthesis
(VanA,B type)
(VanA, B, D type)
(VanC, E, G type)
(S. coelicolor)
(Low-level resistance)
(High level resistance)
Vancomycin resistance operon types
Regulation
Resistance
Accessory
S. coelicolor (chromosome)
VanA cluster (Plasmid)
VanB cluster (Plasmid chromosome)
VanC cluster (Chromosome)
VanD cluster (Chromosome)
VanE cluster (Chromosome)
VanG cluster (Chromosome)
Data on KEGG graph
Rendered by Pathview

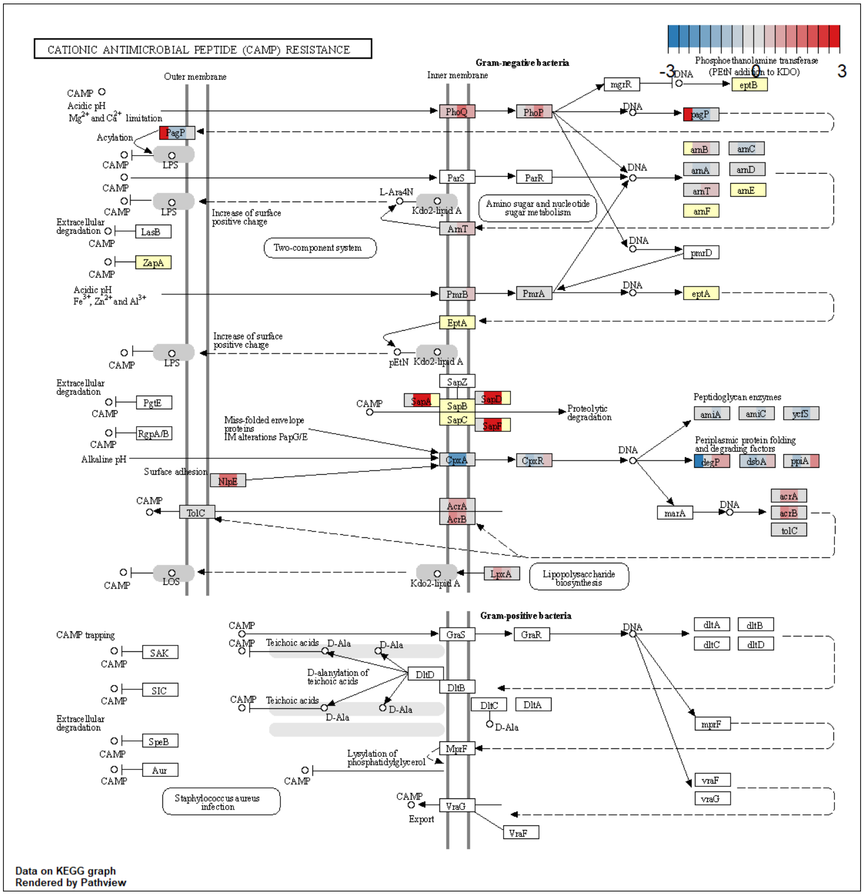
CATIONIC ANTIMICROBIAL PEPTIDE (CAMP) RESISTANCE
-3
0
3
Gram-negative bacteria
Outer membrane
Inner membrane
Phosphoethanolamine transferase (PEtN addition to KDO)
Two-component system
Amino sugar and nucleotide sugar metabolism
Lipopolysaccharide biosynthesis
Peptidoglycan enzymes
Periplasmic protein folding and degrading factors
Gram-positive bacteria
CAMP trapping
Staphylococcus aureus infection
Data on KEGG graph
Rendered by Pathview

**Figure S7.** Correlation map of the 6 proteome comparisons of the 4 clinical isolates and 57 additional Ye transcriptome comparisons. Dot sizes are proportional to Pearson coefficient absolute value. Biological conditions associated to each comparison are described in Table S7.

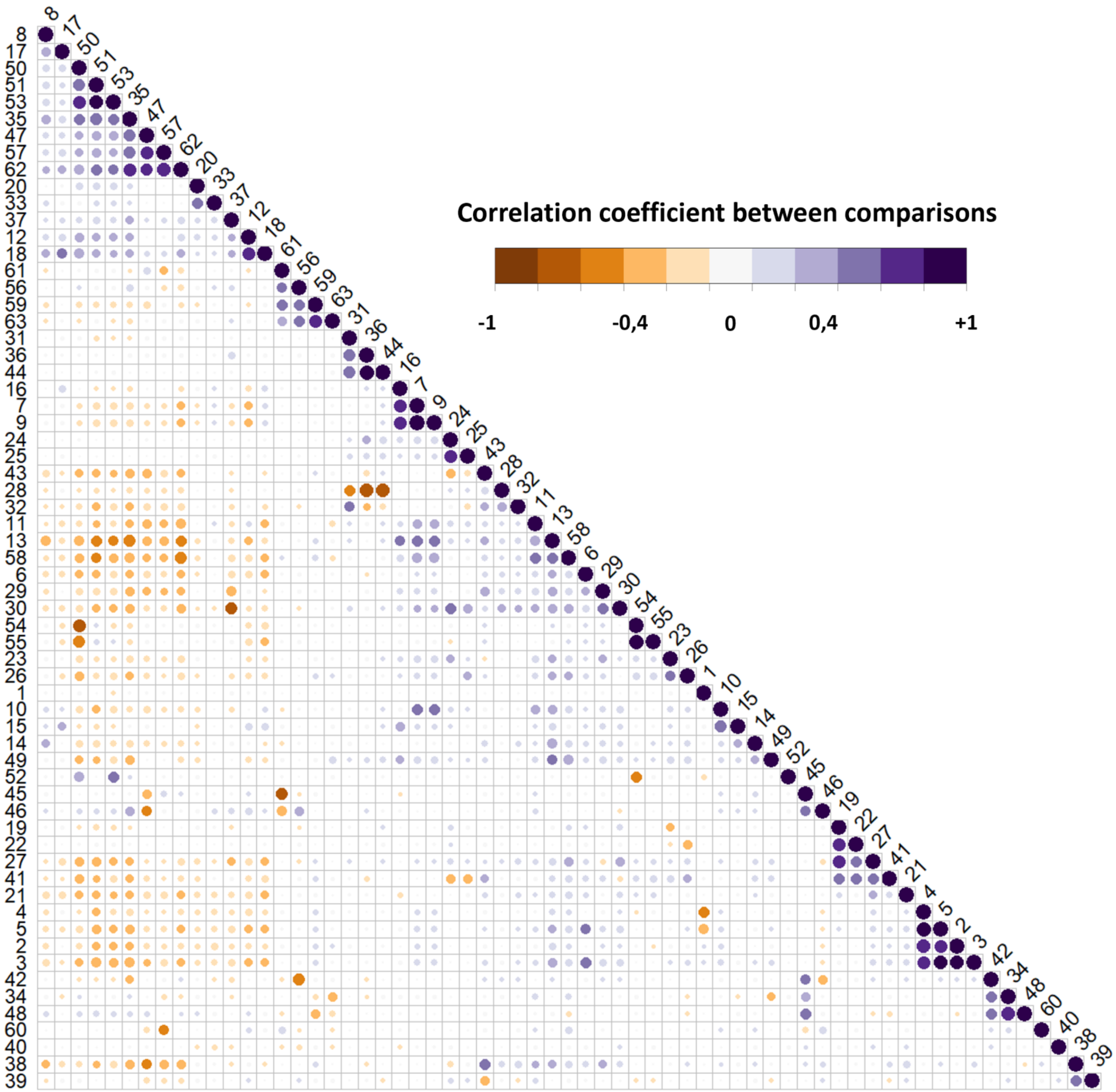

**Table S1.** Antibiotic susceptibility profiling of the 4 Ye strains on 33 antibiotics and antibiotics combinations, assessed by disk diffusion assay. 3G: 3rd generation. 4G: 4th generation. 5G: 5th generation. -: susceptible. R: resistance. r: increased resistance compared to previous strain.

| Class | Subclass | Antibiotic | Ye.1 | Ye.2 | Ye.3 | Ye.4 |
|---|---|---|---|---|---|---|
| β-lactam | penicillin (4G) | piperacillin-tazobactam | - | - | - | - |
| | penicillin (4G) | piperacillin | - | r | - | - |
| | penicillin (4G) | ticarcillin | R | R | R | R |
| | penicillin (3G) | amoxicillin | R | R | R | R |
| | carbapenem | ertapenem | - | - | - | - |
| | penicillin | ticarcillin-clavulanate | - | - | - | - |
| | cephem (3G) | ceftazidime | - | r | - | - |
| | cephem (2G) | cefoxitin | - | r | - | R |
| | penicillin (4G) | temocillin | - | - | - | - |
| | carbapenem | imipenem | - | - | - | - |
| | penicillin (3G) | amoxicillin-clavulanate | - | - | - | - |
| | cephem (3G) | cefotaxime | - | - | - | - |
| | cephem (3G) | ceftazidime-avibactam | - | - | - | - |
| | carbapenem | meropenem | - | - | - | - |
| | monobactam | aztreonam | - | - | - | - |
| | cephem (4G) | cefepime | - | - | - | - |
| | cephem (3G) | moxalactam | - | - | - | - |
| | penicillin | mecillinam | - | - | - | - |
| | cephem (5G) | ceftolozan-tazobactam | - | r | - | - |
| | cephem | cefiderocol | - | - | - | - |
| quinolone | quinolone | nalidixic acid | - | - | R | R |
| | fluoroquinolone | levofloxacine | - | - | - | - |
| | fluoroquinolone | ciprofloxacine | - | - | - | - |
| tetracycline | | tigecycline | - | - | - | - |
| fosfomycin | | fosfomycin | - | - | - | - |
| nitrofuran | | nitrofurantoin | - | - | - | - |
| sulfonamide-trimethoprime | | sulfamethoxazole-trimethoprime | - | - | - | - |
| amphenicol | | chloramphenicol | - | - | - | - |
| polymyxin | | colistin | - | - | - | - |
| aminoglycosides | | gentamicin | - | - | - | - |
| | | amikacin | - | - | - | - |
| | | tobramycin | - | - | - | - |
| | | netilmycin | - | r | - | - |

**Table S2.** Genomes of 263 *Ye* biotype 4 used for core-genome single-nucleotide polymorphism analysis

| Strain | Species | Biotype | Serotype | Category | Isolation date | Origin | Country | Genomic data |
|---|---|---|---|---|---|---|---|---|
| Ye.1 | Y.enterocolitica | 4 | O:3 | Clinical | 1999-12-15 | blood | France | this study |
| Ye.2 | Y.enterocolitica | 4 | O:3 | Clinical | 2000-02-10 | blood | France | this study |
| Ye.3 | Y.enterocolitica | 4 | O:3 | Clinical | 2013-01-12 | blood | France | this study |
| Ye.4 | Y.enterocolitica | 4 | O:3 | Clinical | 2013-10-03 | blood | France | this study |
| IP00134 | Y.enterocolitica | 4 | O:3 | Clinical | 1963 | Ganglion | Sweden | Saraka et al., PNTD, 2017 |
| IP10393 | Y.enterocolitica | 4 | O:3 | Clinical | 1982 | stool | France | Reuter et al., PNAS, 2014 |
| IP28847 | Y.enterocolitica | 4 | O:3 | Clinical | 1988-05-03 | stool | Russia | This study |
| IP22276 | Y.enterocolitica | 4 | O:3 | Clinical | 1991-05-15 | stool | Australia | Reuter et al., PNAS, 2014 |
| IP26656 | Y.enterocolitica | 4 | O:3 | Clinical | 1999-10-04 | stool | France | Reuter et al., PNAS, 2014 |
| IP29063 | Y.enterocolitica | 4 | O:3 | Clinical | 2005-12-08 | stool | France | This study |
| IP28744 | Y.enterocolitica | 4 | O:3 | Clinical | 2006-05-29 | stool | France | This study |
| IP28787 | Y.enterocolitica | 4 | O:3 | Clinical | 2006-07-18 | stool | France | This study |
| IP28804 | Y.enterocolitica | 4 | O:3 | Clinical | 2006-07-31 | stool | France | This study |
| IP28800 | Y.enterocolitica | 4 | O:3 | Clinical | 2006-08-03 | blood | France | This study |
| IP28803 | Y.enterocolitica | 4 | O:3 | Clinical | 2006-08-07 | stool | France | This study |
| IP28892 | Y.enterocolitica | 4 | O:3 | Clinical | 2006-09-27 | stool | France | This study |
| IP29005 | Y.enterocolitica | 4 | O:3 | Clinical | 2007-02-22 | stool | France | This study |
| IP29012 | Y.enterocolitica | 4 | O:3 | Clinical | 2007-03-06 | stool | France | This study |
| IP29069 | Y.enterocolitica | 4 | O:3 | Clinical | 2007-03-30 | stool | France | This study |
| IP29107 | Y.enterocolitica | 4 | O:3 | Clinical | 2007-05-14 | stool | France | This study |
| IP29145 | Y.enterocolitica | 4 | O:3 | Clinical | 2007-07-05 | blood | France | This study |
| IP29217 | Y.enterocolitica | 4 | O:3 | Clinical | 2007-09-05 | stool | France | This study |
| IP29296 | Y.enterocolitica | 4 | O:3 | Clinical | 2007-10-08 | blood | France | This study |
| IP29301 | Y.enterocolitica | 4 | O:3 | Clinical | 2007-10-11 | stool | France | This study |
| IP29395 | Y.enterocolitica | 4 | O:3 | Veterinary | 2008-01-07 | stool | France | This study |
| IP29402 | Y.enterocolitica | 4 | O:3 | Clinical | 2008-01-17 | stool | France | This study |
| IP29471 | Y.enterocolitica | 4 | O:3 | Clinical | 2008-04-01 | stool | France | This study |
| IP29604 | Y.enterocolitica | 4 | O:3 | Clinical | 2008-07-01 | stool | France | This study |
| IP29610 | Y.enterocolitica | 4 | O:3 | Clinical | 2008-07-07 | stool | France | Reuter et al., PNAS, 2014 |
| IP29681 | Y.enterocolitica | 4 | O:3 | Clinical | 2008-08-01 | stool | France | This study |
| IP29639 | Y.enterocolitica | 4 | O:3 | Clinical | 2008-08-04 | stool | France | This study |
| IP29661 | Y.enterocolitica | 4 | O:3 | Clinical | 2008-08-12 | stool | France | This study |
| IP29690 | Y.enterocolitica | 4 | O:3 | Clinical | 2008-09-02 | stool | France | This study |
| IP29684 | Y.enterocolitica | 4 | O:3 | Clinical | 2008-09-06 | stool | France | This study |
| IP29688 | Y.enterocolitica | 4 | O:3 | Clinical | 2008-09-07 | stool | France | This study |
| IP29691 | Y.enterocolitica | 4 | O:3 | Clinical | 2008-09-16 | stool | France | This study |
| IP29701 | Y.enterocolitica | 4 | O:3 | Clinical | 2008-09-20 | stool | France | This study |
| IP29708 | Y.enterocolitica | 4 | O:3 | Clinical | 2008-09-23 | stool | France | This study |
| IP29753 | Y.enterocolitica | 4 | O:3 | Clinical | 2008-11-19 | stool | France | This study |
| IP29877 | Y.enterocolitica | 4 | O:3 | Clinical | 2009-04 | stool | France | This study |
| IP29890 | Y.enterocolitica | 4 | O:3 | Clinical | 2009-04-30 | stool | France | This study |
| IP29909 | Y.enterocolitica | 4 | O:3 | Clinical | 2009-05 | stool | France | This study |
| IP29950 | Y.enterocolitica | 4 | O:3 | Clinical | 2009-06-07 | stool | France | This study |
| IP29962 | Y.enterocolitica | 4 | O:3 | Clinical | 2009-06-30 | stool | France | This study |
| IP33484 | Y.enterocolitica | 4 | O:3 | Clinical | 2009-08-14 | stool | France | This study |
| IP33483 | Y.enterocolitica | 4 | O:3 | Clinical | 2009-08-17 | stool | France | This study |
| IP33502 | Y.enterocolitica | 4 | O:3 | Clinical | 2009-09-05 | stool | France | This study |
| IP33503 | Y.enterocolitica | 4 | O:3 | Clinical | 2009-09-07 | stool | France | This study |
| IP33514 | Y.enterocolitica | 4 | O:3 | Clinical | 2009-09-21 | stool | France | This study |
| IP33517 | Y.enterocolitica | 4 | O:3 | Clinical | 2009-09-24 | stool | France | This study |
| IP33540 | Y.enterocolitica | 4 | O:3 | Clinical | 2009-10-28 | stool | France | This study |
| IP33555 | Y.enterocolitica | 4 | O:3 | Clinical | 2009-11-18 | stool | France | This study |
| IP33925 | Y.enterocolitica | 4 | O:3 | Veterinary | 2009-11-23 | Pig (tongue) | Ivory Coast | This study |
| IP33926 | Y.enterocolitica | 4 | O:3 | Veterinary | 2009-11-23 | Pig (meat) | Ivory Coast | This study |
| IP33569 | Y.enterocolitica | 4 | O:3 | Clinical | 2009-12-10 | stool | France | This study |
| IP33927 | Y.enterocolitica | 4 | O:3 | Veterinary | 2009-12-11 | Pig (stool) | Ivory Coast | This study |
| IP33587 | Y.enterocolitica | 4 | O:3 | Clinical | 2010-01-20 | stool | France | This study |
| IP33631 | Y.enterocolitica | 4 | O:3 | Clinical | 2010-03-08 | stool | France | This study |
| IP33746 | Y.enterocolitica | 4 | O:3 | Clinical | 2010-04-26 | stool | France | This study |
| IP33750 | Y.enterocolitica | 4 | O:3 | Clinical | 2010-05-25 | stool | France | This study |
| IP33773 | Y.enterocolitica | 4 | O:3 | Clinical | 2010-07-15 | stool | France | This study |
| IP33840 | Y.enterocolitica | 4 | O:3 | Clinical | 2010-08-27 | stool | France | This study |
| IP33851 | Y.enterocolitica | 4 | O:3 | Clinical | 2010-09-03 | stool | France | This study |

| | | | | | | | | |
|---|---|---|---|---|---|---|---|---|
| IP33848 | Y.enterocolitica | 4 | O:3 | Clinical | 2010-09-04 | stool | France | This study |
| IP33895 | Y.enterocolitica | 4 | O:3 | Clinical | 2010-10-09 | stool | France | This study |
| IP33916 | Y.enterocolitica | 4 | O:3 | Clinical | 2010-11-09 | blood | France | This study |
| IP33936 | Y.enterocolitica | 4 | O:3 | Clinical | 2010-11-13 | Ascites | France | This study |
| IP33938 | Y.enterocolitica | 4 | O:3 | Clinical | 2010-11-20 | stool | France | This study |
| IP33985 | Y.enterocolitica | 4 | O:3 | Clinical | 2011-01-06 | stool | France | This study |
| IP34009 | Y.enterocolitica | 4 | O:3 | Clinical | 2011-01-22 | stool | France | This study |
| IP34022 | Y.enterocolitica | 4 | O:3 | Clinical | 2011-02-08 | stool | France | This study |
| IP34028 | Y.enterocolitica | 4 | O:3 | Clinical | 2011-02-14 | stool | France | This study |
| IP34035 | Y.enterocolitica | 4 | O:3 | Clinical | 2011-02-22 | stool | France | This study |
| IP34040 | Y.enterocolitica | 4 | O:3 | Clinical | 2011-03-05 | stool | France | This study |
| IP34075 | Y.enterocolitica | 4 | O:3 | Clinical | 2011-03-30 | stool | France | This study |
| IP34100 | Y.enterocolitica | 4 | O:3 | Clinical | 2011-04-13 | stool | France | This study |
| IP34140 | Y.enterocolitica | 4 | O:3 | Clinical | 2011-05-10 | stool | France | This study |
| IP34144 | Y.enterocolitica | 4 | O:3 | Clinical | 2011-05-11 | stool | France | This study |
| IP34166 | Y.enterocolitica | 4 | O:3 | Clinical | 2011-05-25 | stool | France | This study |
| IP34177 | Y.enterocolitica | 4 | O:3 | Clinical | 2011-06-16 | stool | France | This study |
| IP34242 | Y.enterocolitica | 4 | O:3 | Clinical | 2011-06-17 | blood | France | This study |
| IP34220 | Y.enterocolitica | 4 | O:3 | Clinical | 2011-07-08 | blood | France | This study |
| IP34238 | Y.enterocolitica | 4 | O:3 | Clinical | 2011-07-18 | stool | France | This study |
| IP34255 | Y.enterocolitica | 4 | O:3 | Clinical | 2011-07-21 | stool | France | This study |
| IP34251 | Y.enterocolitica | 4 | O:3 | Clinical | 2011-07-25 | stool | France | This study |
| IP34283 | Y.enterocolitica | 4 | O:3 | Clinical | 2011-08-03 | stool | France | This study |
| IP34276 | Y.enterocolitica | 4 | O:3 | Clinical | 2011-08-13 | stool | France | This study |
| IP34280 | Y.enterocolitica | 4 | O:3 | Clinical | 2011-08-17 | stool | France | This study |
| IP34293 | Y.enterocolitica | 4 | O:3 | Clinical | 2011-08-22 | stool | France | This study |
| IP34305 | Y.enterocolitica | 4 | O:3 | Clinical | 2011-08-28 | stool | France | This study |
| IP34302 | Y.enterocolitica | 4 | O:3 | Clinical | 2011-08-29 | stool | France | This study |
| IP34303 | Y.enterocolitica | 4 | O:3 | Clinical | 2011-08-29 | stool | France | This study |
| IP34315 | Y.enterocolitica | 4 | O:3 | Clinical | 2011-08-30 | stool | France | This study |
| IP34310 | Y.enterocolitica | 4 | O:3 | Clinical | 2011-09-01 | stool | France | This study |
| IP34324 | Y.enterocolitica | 4 | O:3 | Clinical | 2011-09-01 | stool | France | This study |
| IP34319 | Y.enterocolitica | 4 | O:3 | Clinical | 2011-09-03 | stool | France | This study |
| IP34321 | Y.enterocolitica | 4 | O:3 | Clinical | 2011-09-12 | stool | France | This study |
| IP34339 | Y.enterocolitica | 4 | O:3 | Clinical | 2011-09-16 | stool | France | This study |
| IP34391 | Y.enterocolitica | 4 | O:3 | Clinical | 2011-11-25 | stool | France | This study |
| IP34407 | Y.enterocolitica | 4 | O:3 | Clinical | 2011-12-07 | stool | France | This study |
| IP34421 | Y.enterocolitica | 4 | O:3 | Clinical | 2011-12-19 | stool | France | This study |
| IP34423 | Y.enterocolitica | 4 | O:3 | Clinical | 2011-12-21 | blood | France | This study |
| IP34460 | Y.enterocolitica | 4 | O:3 | Clinical | 2012-01-20 | stool | France | This study |
| IP34508 | Y.enterocolitica | 4 | O:3 | Clinical | 2012-03-06 | stool | France | This study |
| IP34522 | Y.enterocolitica | 4 | O:3 | Clinical | 2012-03-19 | stool | France | This study |
| IP34534 | Y.enterocolitica | 4 | O:3 | Clinical | 2012-03-23 | stool | France | This study |
| IP34558 | Y.enterocolitica | 4 | O:3 | Clinical | 2012-04-03 | stool | France | This study |
| IP34604 | Y.enterocolitica | 4 | O:3 | Clinical | 2012-05-14 | stool | France | This study |
| IP34601 | Y.enterocolitica | 4 | O:3 | Clinical | 2012-05-15 | stool | France | This study |
| IP34663 | Y.enterocolitica | 4 | O:3 | Clinical | 2012-06-18 | stool | France | This study |
| IP34659 | Y.enterocolitica | 4 | O:3 | Clinical | 2012-06-22 | stool | France | This study |
| IP34679 | Y.enterocolitica | 4 | O:3 | Clinical | 2012-07-06 | stool | France | This study |
| IP34721 | Y.enterocolitica | 4 | O:3 | Clinical | 2012-08-03 | stool | France | This study |
| IP34724 | Y.enterocolitica | 4 | O:3 | Clinical | 2012-08-07 | stool | France | This study |
| IP34747 | Y.enterocolitica | 4 | O:3 | Clinical | 2012-08-20 | stool | France | This study |
| IP34794 | Y.enterocolitica | 4 | O:3 | Clinical | 2012-09-13 | stool | France | This study |
| IP34798 | Y.enterocolitica | 4 | O:3 | Clinical | 2012-09-19 | stool | France | This study |
| IP35459 | Y.enterocolitica | 4 | O:3 | Veterinary | 2012-09-24 | stool | Ivory Coast | Saraka et al., PNTD, 2017 |
| IP34842 | Y.enterocolitica | 4 | O:3 | Clinical | 2012-10-26 | stool | France | This study |
| IP34931 | Y.enterocolitica | 4 | O:3 | Clinical | 2013-01-02 | stool | France | This study |
| IP35462 | Y.enterocolitica | 4 | O:3 | Veterinary | 2013-03-01 | stool | Ivory Coast | Saraka et al., PNTD, 2017 |
| IP35463 | Y.enterocolitica | 4 | O:3 | Veterinary | 2013-03-01 | stool | Ivory Coast | Saraka et al., PNTD, 2017 |
| IP35464 | Y.enterocolitica | 4 | O:3 | Veterinary | 2013-03-01 | stool | Ivory Coast | Saraka et al., PNTD, 2017 |
| IP35465 | Y.enterocolitica | 4 | O:3 | Veterinary | 2013-03-07 | stool | Ivory Coast | Saraka et al., PNTD, 2017 |
| IP35466 | Y.enterocolitica | 4 | O:3 | Veterinary | 2013-03-07 | stool | Ivory Coast | Saraka et al., PNTD, 2017 |
| IP35467 | Y.enterocolitica | 4 | O:3 | Veterinary | 2013-04-08 | stool | Ivory Coast | Saraka et al., PNTD, 2017 |
| IP35088 | Y.enterocolitica | 4 | O:3 | Clinical | 2013-04-26 | stool | France | This study |
| IP35470 | Y.enterocolitica | 4 | O:3 | Veterinary | 2013-05-03 | stool | Ivory Coast | Saraka et al., PNTD, 2017 |
| IP35104 | Y.enterocolitica | 4 | O:3 | Clinical | 2013-05-15 | stool | France | This study |

| | | | | | | | | |
|---|---|---|---|---|---|---|---|---|
| IP35118 | Y.enterocolitica | 4 | O:3 | Clinical | 2013-05-15 | stool | France | This study |
| IP35477 | Y.enterocolitica | 4 | O:3 | Clinical | 2013-05-23 | stool | Ivory Coast | Saraka et al., PNTD, 2017 |
| IP35131 | Y.enterocolitica | 4 | O:3 | Clinical | 2013-05-29 | stool | France | This study |
| IP35471 | Y.enterocolitica | 4 | O:3 | Veterinary | 2013-07-11 | stool | Ivory Coast | Saraka et al., PNTD, 2017 |
| IP35472 | Y.enterocolitica | 4 | O:3 | Veterinary | 2013-07-11 | stool | Ivory Coast | Saraka et al., PNTD, 2017 |
| IP35213 | Y.enterocolitica | 4 | O:3 | Clinical | 2013-07-12 | stool | France | This study |
| IP35255 | Y.enterocolitica | 4 | O:3 | Clinical | 2013-07-20 | stool | France | This study |
| IP35254 | Y.enterocolitica | 4 | O:3 | Clinical | 2013-07-23 | stool | France | This study |
| IP35474 | Y.enterocolitica | 4 | O:3 | Veterinary | 2013-08-13 | stool | Ivory Coast | Saraka et al., PNTD, 2017 |
| IP35475 | Y.enterocolitica | 4 | O:3 | Veterinary | 2013-08-13 | stool | Ivory Coast | Saraka et al., PNTD, 2017 |
| IP35478 | Y.enterocolitica | 4 | O:3 | Clinical | 2013-08-13 | stool | Ivory Coast | Saraka et al., PNTD, 2017 |
| IP35382 | Y.enterocolitica | 4 | O:3 | Clinical | 2013-09-09 | stool | France | This study |
| IP35395 | Y.enterocolitica | 4 | O:3 | Clinical | 2013-09-22 | stool | France | This study |
| IP35439 | Y.enterocolitica | 4 | O:3 | Clinical | 2013-10-16 | stool | France | This study |
| IP35711 | Y.enterocolitica | 4 | O:3 | Clinical | 2014-03-18 | stool | France | This study |
| IP35728 | Y.enterocolitica | 4 | O:3 | Clinical | 2014-03-21 | stool | France | This study |
| IP35754 | Y.enterocolitica | 4 | O:3 | Clinical | 2014-04-02 | stool | France | This study |
| IP35764 | Y.enterocolitica | 4 | O:3 | Clinical | 2014-04-02 | stool | France | This study |
| IP35850 | Y.enterocolitica | 4 | O:3 | Clinical | 2014-05-17 | stool | France | This study |
| IP35933 | Y.enterocolitica | 4 | O:3 | Clinical | 2014-06-16 | stool | France | This study |
| IP35923 | Y.enterocolitica | 4 | O:3 | Clinical | 2014-06-17 | stool | France | This study |
| IP35935 | Y.enterocolitica | 4 | O:3 | Clinical | 2014-06-19 | stool | France | This study |
| IP36022 | Y.enterocolitica | 4 | O:3 | Clinical | 2014-07-19 | blood | France | This study |
| IP36039 | Y.enterocolitica | 4 | O:3 | Clinical | 2014-07-24 | stool | France | This study |
| IP36085 | Y.enterocolitica | 4 | O:3 | Clinical | 2014-08-17 | stool | France | This study |
| IP36202 | Y.enterocolitica | 4 | O:3 | Clinical | 2014-09 | stool | France | This study |
| IP36151 | Y.enterocolitica | 4 | O:3 | Clinical | 2014-09-08 | stool | France | This study |
| IP36356 | Y.enterocolitica | 4 | O:3 | Clinical | 2014-12-15 | stool | France | This study |
| IP37447 | Y.enterocolitica | 4 | O:3 | Clinical | 2015-01-25 | stool | France | This study |
| IP37060 | Y.enterocolitica | 4 | O:3 | Clinical | 2015-08-26 | stool | France | This study |
| IP37174 | Y.enterocolitica | 4 | O:3 | Clinical | 2015-10-03 | stool | France | This study |
| IP37227 | Y.enterocolitica | 4 | O:3 | Clinical | 2015-10-20 | stool | France | This study |
| IP37233 | Y.enterocolitica | 4 | O:3 | Clinical | 2015-10-27 | stool | France | This study |
| IP37402 | Y.enterocolitica | 4 | O:3 | Clinical | 2015-12-25 | stool | France | This study |
| IP37401 | Y.enterocolitica | 4 | O:3 | Clinical | 2015-12-29 | stool | France | This study |
| IP37403 | Y.enterocolitica | 4 | O:3 | Clinical | 2015-12-29 | stool | France | This study |
| IP37406 | Y.enterocolitica | 4 | O:3 | Clinical | 2015-12-30 | stool | France | This study |
| IP37423 | Y.enterocolitica | 4 | O:3 | Clinical | 2015-12-30 | stool | France | This study |
| IP37405 | Y.enterocolitica | 4 | O:3 | Clinical | 2015-12-31 | stool | France | This study |
| IP37407 | Y.enterocolitica | 4 | O:3 | Clinical | 2015-12-31 | stool | France | This study |
| IP37415 | Y.enterocolitica | 4 | O:3 | Clinical | 2015-12-31 | stool | France | This study |
| IP37400 | Y.enterocolitica | 4 | O:3 | Clinical | 2016-01-02 | stool | France | This study |
| IP37434 | Y.enterocolitica | 4 | O:3 | Clinical | 2016-01-04 | stool | France | This study |
| IP37409 | Y.enterocolitica | 4 | O:3 | Clinical | 2016-01-05 | stool | France | This study |
| IP37418 | Y.enterocolitica | 4 | O:3 | Clinical | 2016-01-06 | stool | France | This study |
| IP37414 | Y.enterocolitica | 4 | O:3 | Clinical | 2016-01-07 | stool | France | This study |
| IP37420 | Y.enterocolitica | 4 | O:3 | Clinical | 2016-01-07 | stool | France | This study |
| IP37416 | Y.enterocolitica | 4 | O:3 | Clinical | 2016-01-08 | stool | France | This study |
| IP37417 | Y.enterocolitica | 4 | O:3 | Clinical | 2016-01-09 | stool | France | This study |
| IP37443 | Y.enterocolitica | 4 | O:3 | Clinical | 2016-01-10 | stool | France | This study |
| IP37425 | Y.enterocolitica | 4 | O:3 | Clinical | 2016-01-11 | stool | France | This study |
| IP37436 | Y.enterocolitica | 4 | O:3 | Clinical | 2016-01-11 | stool | France | This study |
| IP37446 | Y.enterocolitica | 4 | O:3 | Clinical | 2016-01-11 | stool | France | This study |
| IP37449 | Y.enterocolitica | 4 | O:3 | Clinical | 2016-01-11 | stool | France | This study |
| IP37452 | Y.enterocolitica | 4 | O:3 | Clinical | 2016-01-11 | stool | France | This study |
| IP37456 | Y.enterocolitica | 4 | O:3 | Clinical | 2016-01-11 | stool | France | This study |
| IP37492 | Y.enterocolitica | 4 | O:3 | Clinical | 2016-01-11 | stool | France | This study |
| IP37433 | Y.enterocolitica | 4 | O:3 | Clinical | 2016-01-12 | stool | France | This study |
| IP37435 | Y.enterocolitica | 4 | O:3 | Clinical | 2016-01-12 | stool | France | This study |
| IP37437 | Y.enterocolitica | 4 | O:3 | Clinical | 2016-01-12 | stool | France | This study |
| IP37461 | Y.enterocolitica | 4 | O:3 | Clinical | 2016-01-12 | stool | France | This study |
| IP37440 | Y.enterocolitica | 4 | O:3 | Clinical | 2016-01-13 | stool | France | This study |
| IP37460 | Y.enterocolitica | 4 | O:3 | Clinical | 2016-01-13 | stool | France | This study |
| IP37462 | Y.enterocolitica | 4 | O:3 | Clinical | 2016-01-13 | stool | France | This study |
| IP37441 | Y.enterocolitica | 4 | O:3 | Clinical | 2016-01-14 | stool | France | This study |
| IP37459 | Y.enterocolitica | 4 | O:3 | Clinical | 2016-01-14 | stool | France | This study |

| | | | | | | | | |
|---|---|---|---|---|---|---|---|---|
| IP37463 | Y.enterocolitica | 4 | O:3 | Clinical | 2016-01-15 | stool | France | This study |
| IP37396 | Y.enterocolitica | 4 | O:3 | Clinical | 2016-01-16 | stool | France | This study |
| IP37458 | Y.enterocolitica | 4 | O:3 | Clinical | 2016-01-17 | stool | France | This study |
| IP37439 | Y.enterocolitica | 4 | O:3 | Clinical | 2016-01-18 | stool | France | This study |
| IP37466 | Y.enterocolitica | 4 | O:3 | Clinical | 2016-01-18 | stool | France | This study |
| IP37467 | Y.enterocolitica | 4 | O:3 | Clinical | 2016-01-20 | stool | France | This study |
| IP37480 | Y.enterocolitica | 4 | O:3 | Clinical | 2016-01-20 | stool | France | This study |
| IP37483 | Y.enterocolitica | 4 | O:3 | Clinical | 2016-01-22 | stool | France | This study |
| IP37493 | Y.enterocolitica | 4 | O:3 | Clinical | 2016-01-22 | stool | France | This study |
| IP37474 | Y.enterocolitica | 4 | O:3 | Clinical | 2016-01-23 | stool | France | This study |
| IP37482 | Y.enterocolitica | 4 | O:3 | Clinical | 2016-01-23 | stool | France | This study |
| IP37491 | Y.enterocolitica | 4 | O:3 | Clinical | 2016-01-23 | stool | France | This study |
| IP37478 | Y.enterocolitica | 4 | O:3 | Clinical | 2016-01-25 | stool | France | This study |
| IP37486 | Y.enterocolitica | 4 | O:3 | Clinical | 2016-01-25 | stool | France | This study |
| IP37500 | Y.enterocolitica | 4 | O:3 | Clinical | 2016-01-26 | stool | France | This study |
| IP37490 | Y.enterocolitica | 4 | O:3 | Clinical | 2016-01-27 | stool | France | This study |
| IP37499 | Y.enterocolitica | 4 | O:3 | Clinical | 2016-01-27 | stool | France | This study |
| IP37494 | Y.enterocolitica | 4 | O:3 | Clinical | 2016-01-28 | stool | France | This study |
| IP37498 | Y.enterocolitica | 4 | O:3 | Clinical | 2016-01-28 | stool | France | This study |
| IP37503 | Y.enterocolitica | 4 | O:3 | Clinical | 2016-01-29 | stool | France | This study |
| IP37506 | Y.enterocolitica | 4 | O:3 | Clinical | 2016-01-30 | stool | France | This study |
| IP37510 | Y.enterocolitica | 4 | O:3 | Clinical | 2016-01-31 | stool | France | This study |
| IP37511 | Y.enterocolitica | 4 | O:3 | Clinical | 2016-02-01 | blood | France | This study |
| IP37523 | Y.enterocolitica | 4 | O:3 | Clinical | 2016-02-01 | stool | France | This study |
| IP37509 | Y.enterocolitica | 4 | O:3 | Clinical | 2016-02-03 | stool | France | This study |
| IP37512 | Y.enterocolitica | 4 | O:3 | Clinical | 2016-02-04 | stool | France | This study |
| IP37515 | Y.enterocolitica | 4 | O:3 | Clinical | 2016-02-04 | stool | France | This study |
| IP37522 | Y.enterocolitica | 4 | O:3 | Clinical | 2016-02-08 | stool | France | This study |
| IP37528 | Y.enterocolitica | 4 | O:3 | Clinical | 2016-02-08 | stool | France | This study |
| IP37529 | Y.enterocolitica | 4 | O:3 | Clinical | 2016-02-09 | stool | France | This study |
| IP37551 | Y.enterocolitica | 4 | O:3 | Clinical | 2016-02-09 | stool | France | This study |
| IP37532 | Y.enterocolitica | 4 | O:3 | Clinical | 2016-02-10 | stool | France | This study |
| IP37536 | Y.enterocolitica | 4 | O:3 | Clinical | 2016-02-10 | stool | France | This study |
| IP37538 | Y.enterocolitica | 4 | O:3 | Clinical | 2016-02-10 | stool | France | This study |
| IP37543 | Y.enterocolitica | 4 | O:3 | Clinical | 2016-02-10 | stool | France | This study |
| IP37552 | Y.enterocolitica | 4 | O:3 | Clinical | 2016-02-11 | stool | France | This study |
| IP37555 | Y.enterocolitica | 4 | O:3 | Clinical | 2016-02-11 | stool | France | This study |
| IP37535 | Y.enterocolitica | 4 | O:3 | Clinical | 2016-02-12 | stool | France | This study |
| IP37545 | Y.enterocolitica | 4 | O:3 | Clinical | 2016-02-12 | stool | France | This study |
| IP37565 | Y.enterocolitica | 4 | O:3 | Clinical | 2016-02-12 | stool | France | This study |
| IP37549 | Y.enterocolitica | 4 | O:3 | Clinical | 2016-02-13 | stool | France | This study |
| IP37546 | Y.enterocolitica | 4 | O:3 | Clinical | 2016-02-15 | stool | France | This study |
| IP37548 | Y.enterocolitica | 4 | O:3 | Clinical | 2016-02-15 | stool | France | This study |
| IP37556 | Y.enterocolitica | 4 | O:3 | Clinical | 2016-02-15 | stool | France | This study |
| IP37557 | Y.enterocolitica | 4 | O:3 | Clinical | 2016-02-16 | stool | France | This study |
| IP37563 | Y.enterocolitica | 4 | O:3 | Clinical | 2016-02-16 | stool | France | This study |
| IP37566 | Y.enterocolitica | 4 | O:3 | Clinical | 2016-02-17 | stool | France | This study |
| IP37562 | Y.enterocolitica | 4 | O:3 | Clinical | 2016-02-18 | stool | France | This study |
| IP37560 | Y.enterocolitica | 4 | O:3 | Clinical | 2016-02-19 | stool | France | This study |
| IP37561 | Y.enterocolitica | 4 | O:3 | Clinical | 2016-02-19 | stool | France | This study |
| IP37572 | Y.enterocolitica | 4 | O:3 | Clinical | 2016-02-19 | stool | France | This study |
| IP37575 | Y.enterocolitica | 4 | O:3 | Clinical | 2016-02-19 | Hematoma | France | This study |
| IP37574 | Y.enterocolitica | 4 | O:3 | Clinical | 2016-02-25 | blood | France | This study |
| ERL022789 | Y.enterocolitica | 4 | O:3 | Veterinary | NK* | Dog | New Zealand | Reuter et al., PNAS, 2014 |
| ERL032770 | Y.enterocolitica | 4 | O:3 | Clinical | NK* | blood | New Zealand | Reuter et al., PNAS, 2014 |
| ERL04757 | Y.enterocolitica | 4 | O:3 | Clinical | NK* | blood | New Zealand | Reuter et al., PNAS, 2014 |
| ERL063683 | Y.enterocolitica | 4 | nd | Clinical | NK* | blood | New Zealand | Reuter et al., PNAS, 2014 |
| ERL072344 | Y.enterocolitica | 4 | O:3 | Veterinary | NK* | NK* | New Zealand | Reuter et al., PNAS, 2014 |
| ERL073627 | Y.enterocolitica | 4 | O:3 | Clinical | NK* | blood | New Zealand | Reuter et al., PNAS, 2014 |
| ERL08109 | Y.enterocolitica | 4 | O:3 | Clinical | NK* | blood | New Zealand | Reuter et al., PNAS, 2014 |
| ERL084574 | Y.enterocolitica | 4 | O:3 | Clinical | NK* | blood | New Zealand | Reuter et al., PNAS, 2014 |
| H450/87 | Y.enterocolitica | 4 | O:3 | Clinical | NK* | NK* | Unknown | Reuter et al., PNAS, 2014 |
| H469/87 | Y.enterocolitica | 4 | O:3 | Veterinary | NK* | Pig | Unknown | Reuter et al., PNAS, 2014 |
| H608/87 | Y.enterocolitica | 4 | O:3 | Clinical | NK* | NK* | Unknown | Reuter et al., PNAS, 2014 |
| Y11 | Y.enterocolitica | 4 | O:3 | Clinical | NK* | stool | Germany | Batzilla et al., J Bacteriol, 2011 |
| YE07/03 | Y.enterocolitica | 4 | O:3 | Clinical | NK* | stool | United Kingdom | Reuter et al., PNAS, 2014 |

| | | | | | | | | |
|---|---|---|---|---|---|---|---|---|
| YE201/02 | Y.enterocolitica | 4 | O:3 | Veterinary | NK* | Pig | United Kingdom | Reuter et al., PNAS, 2014 |
| YE213/02 | Y.enterocolitica | 4 | O:3 | Veterinary | NK* | Pig | United Kingdom | Reuter et al., PNAS, 2014 |

*NK: not known

**Table S3 - SNP specific**

| SNP specific to Ye.3 | | | | | | | | | | | | |
|---|---|---|---|---|---|---|---|---|---|---|---|---|
| Chromosome_ref | Position | Reference | Ye.3 | Gene | locus_tag | protein | protein id | location | type | impact | nucleotide mutation | AA mutation |
| NC_017564 | 49601 | G | A | | Y11_RS00275 | MurR/RpiR family transcriptional regulator | WP_005164882.1] | 49589..50467 | missense_variant | MODERATE | c.13G>A | p.Ala5Thr |
| NC_017564 | 97308 | G | A | | Y11_RS00505 | LysE family transporter | WP_005158710.1] | complement(96732..97370) | synonymous_variant | LOW | c.63C>T | p.Pro21Pro |
| NC_017564 | 174798 | G | A | folC | Y11_RS00875 | bifunctional tetrahydrofolate / synthase/dihydrofolate synthase | WP_012303924.1 | 174454..175722 | synonymous_variant | LOW | c.345G>A | p.Gly115Gly |
| NC_017564 | 309481 | C | T | yejF | Y11_RS01415 | microcin C ABC transporter ATP-binding protein YejF | WP_012105426.1 | complement(309214..310806) | synonymous_variant | LOW | c.1326G>A | p.Leu442Leu |
| NC_017564 | 363013 | G | A | tRNA-Ala | Y11_RS00380 | tRNA-Ala | | | intragenic_variant | MODIFIER | n.363013G>A | |
| NC_017564 | 370589 | G | A | deoR | Y11_RS01660 | DNA-binding transcriptional repressor DeoR | WP_005162894.1] | complement(370092..370856) | missense_variant | MODERATE | c.268C>T | p.Arg90Cys |
| NC_017564 | 595386 | G | A | lolE | Y11_RS02585 | lipoprotein-releasing ABC transporter permease subunit LolE | WP_005158010.1] | 594559..595806 | synonymous_variant | LOW | c.828G>A | p.Ala276Ala |
| NC_017564 | 815340 | C | T | pheT | Y11_RS03630 | phenylalanine--tRNA ligase subunit bet | WP_005161568.1 | 814531..816918 | synonymous_variant | LOW | c.810C>T | p.Asp270Asp |
| NC_017564 | 1026845 | C | T | | Y11_RS04655 | MalY/PatB family protein | WP_005164772.1 | 1026228..1027406 | synonymous_variant | LOW | c.636C>T | p.Asp212Asp |
| NC_017564 | 1251521 | C | T | | Y11_RS21225 | DUF2474 domain-containing protein | WP_005178681.1 | 1251379..1251540 | missense_variant | MODERATE | c.143C>T | p.Ala48Val |
| NC_017564 | 1277973 | C | T | | Y11_RS05805 | tetratricopeptide repeat-containing protein | WP_005161315.1] | complement(1277240..1278049) | missense_variant | MODERATE | c.77G>A | p.Arg26Gln |
| NC_017564 | 1488668 | G | A | mtfA | Y11_RS06985 | DgsA anti-repressor MtfA | WP_005166192.1 | 1488361..1489164 | stop_gained | HIGH | c.308G>A | p.Trp103* |
| NC_017564 | 1662165 | C | T | galU | Y11_RS07795 | UTP--glucose-1-phosphate uridylyltransferase GalU | WP_005157804.1 | complement(1661582..1662472) | missense_variant | MODERATE | c.299G>A | p.Arg100His |
| NC_017564 | 1699363 | G | A | | Y11_RS00380 | | | | intragenic_variant | MODIFIER | n.1699363G>A | |
| NC_017564 | 1811756 | G | A | | Y11_RS08470 | ABC transporter permease | WP_005158920.1 | 1811069..1812175 | missense_variant | MODERATE | c.688G>A | p.Ala230Thr |
| NC_017564 | 2071463 | C | T | lon | Y11_RS09650 | endopeptidase La | WP_005163502.1 | complement(2070791..2073145) | missense_variant | MODERATE | c.1683G>A | p.Met561Ile |
| NC_017564 | 2115898 | C | T | | Y11_RS00380 | | | | intragenic_variant | MODIFIER | n.2115898C>T | |
| NC_017564 | 2222543 | C | T | tRNA-Ala | Y11_RS00380 | tRNA-Ala | | | intragenic_variant | MODIFIER | n.2222543C>T | |
| NC_017564 | 2375049 | C | T | | Y11_RS10980 | YqgE/AlgH family protein | WP_020282839.1 | 2374791..2375354 | missense_variant | MODERATE | c.259C>T | p.His87Tyr |
| NC_017564 | 2906451 | G | A | | Y11_RS13380 | type IV secretion protein Rhs | WP_005165348.1] | 2904276..2909213 | missense_variant | MODERATE | c.2176G>A | p.Gly726Ser |
| NC_017564 | 3035734 | C | T | | Y11_RS00380 | | | | intragenic_variant | MODIFIER | n.3035734C>T | |
| NC_017564 | 3346896 | C | T | hslR | Y11_RS15200 | ribosome-associated heat shock protein Hsp15 | WP_005159667.1] | complement(3346780..3347187) | missense_variant | MODERATE | c.292G>A | p.Glu98Lys |
| NC_017564 | 3579020 | C | T | | Y11_RS16350 | tyrosine-protein phosphatase | WP_005178104.1 | 3578535..3579332 | synonymous_variant | LOW | c.486C>T | p.Asp162Asp |
| NC_017564 | 3674155 | G | A | | Y11_RS16815 | EscV/YscV/HrcV family type III secretion system export apparatus protein | WP_005156632.1 | 3672706..3674745 | missense_variant | MODERATE | c.1450G>A | p.Gly484Ser |
| NC_017564 | 3698014 | G | A | | Y11_RS16925 | PLP-dependent aminotransferase family protein | WP_005156555.1 | 3697930..3699381 | missense_variant | MODERATE | c.85G>A | p.Val29Ile |
| NC_017564 | 3728878 | GC | AA | sugE | Y11_RS17075 | quaternary ammonium compound efflux SMR transporter SugE | WP_014609340.1 | 3728624..3728938 | missense_variant | MODERATE | c.255_256delGCinsAA | p.Arg86Ser |
| NC_017564 | 3728878 | GC | AA | sugE | Y11_RS17075 | quaternary ammonium compound efflux SMR transporter SugE | WP_014609340.1 | 3728624..3728938 | missense_variant | MODERATE | c.255_256delGCinsAA | p.Arg86Ser |
| NC_017564 | 3842142 | C | T | | Y11_RS17590 | alginate lyase family protein | WP_005156235.1] | 3841651..3842817 | synonymous_variant | LOW | c.492C>T | p.Pro164Pro |
| NC_017564 | 3842630 | C | T | | Y11_RS17590 | alginate lyase family protein | WP_005156235.1] | 3841651..3842817 | missense_variant | MODERATE | c.980C>T | p.Pro327Leu |
| NC_017564 | 3864728 | G | A | | Y11_RS17710 | ABC transporter substrate-binding protein | WP_005156162.1 | complement(3863502..3865070) | missense_variant | MODERATE | c.343C>T | p.Arg115Cys |
| NC_017564 | 4057978 | G | A | leuB | Y11_RS18490 | 3-isopropylmalate dehydrogenase | WP_014609375.1] | complement(4057492..4058583) | synonymous_variant | LOW | c.606C>T | p.Ser202Ser |

| SNP specific to Ye.4 | | | | | | | | | | | | |
|---|---|---|---|---|---|---|---|---|---|---|---|---|
| Chromosome_ref | Position | Reference | Ye.4 | Gene | locus_tag | protein | protein id | location | type | impact | nucleotide mutation | AA mutation |
| NC_017564 | 81749 | G | A | | Y11_RS00435 | indolepyruvate decarboxylase | WP_005164836.1 | 80299..81960 | stop_gained | HIGH | c.1454G>A | p.Trp485* |
| NC_017564 | 102936 | C | T | | Y11_RS00530 | APC family permease | WP_005158698.1 | complement(102823..104166) | missense_variant | MODERATE | c.1231G>A | p.Val411Met |
| NC_017564 | 237245 | C | T | | Y11_RS00380 | | | | intragenic_variant | MODIFIER | n.237245C>T | |
| NC_017564 | 268161 | G | A | gyrA | Y11_RS01245 | DNA topoisomerase (ATP-hydrolyzing) subunit A | WP_005159304.1 | 267903..270560 | missense_variant | MODERATE | c.259G>A | p.Asp87Asn |
| NC_017564 | 276445 | G | A | rcsD | Y11_RS01260 | phosphotransferase RcsD | WP_005177600.1 | complement(274457..277150) | missense_variant | MODERATE | c.706C>T | p.Pro236Ser |
| NC_017564 | 407337 | G | A | clpA | Y11_RS01830 | ATP-dependent Clp protease ATP-binding subunit ClpA | WP_005162786.1 | 407121..409397 | missense_variant | MODERATE | c.217G>A | p.Glu73Lys |
| NC_017564 | 471261 | C | T | ompF | Y11_RS02065 | porin OmpF | WP_005159982.1 | complement(470378..471490) | stop_gained | HIGH | c.233G>A | p.Trp78* |
| NC_017564 | 931523 | G | A | hrpA | Y11_RS04160 | ATP-dependent RNA helicase HrpA | WP_005160948.1 | 928586..932473 | missense_variant | MODERATE | c.2938G>A | p.Glu980Lys |
| NC_017564 | 950132 | C | T | sbmC | Y11_RS04255 | DNA gyrase inhibitor SbmC | WP_005160901.1 | 949861..950334 | missense_variant | MODERATE | c.272C>T | p.Pro91Leu |
| NC_017564 | 989672 | C | T | | Y11_RS04430 | esterase-like activity of phytase family protein | WP_005165537.1 | 988486..989832 | missense_variant | MODERATE | c.1187C>T | p.Thr396Ile |
| NC_017564 | 1107108 | C | T | | Y11_RS05015 | YciC family protein | WP_014608963.1 | 1106795..1107562 | missense_variant | MODERATE | c.314C>T | p.Thr105Ile |
| NC_017564 | 1250175 | G | A | | Y11_RS05680 | cytochrome ubiquinol oxidase subunit I | WP_005161246.1 | 1248920..1250368 | missense_variant | MODERATE | c.1256G>A | p.Gly419Asp |
| NC_017564 | 1266669 | G | A | mdtH | Y11_RS05750 | multidrug efflux MFS transporter MdtH | WP_005161289.1 | 1266421..1267626 | missense_variant | MODERATE | c.249G>A | p.Met83Ile |
| NC_017564 | 1295959 | C | T | | Y11_RS00380 | | | | intragenic_variant | MODIFIER | n.1295959C>T | |
| NC_017564 | 1301150 | C | T | | Y11_RS05955 | M15 family metallopeptidase | WP_005161419.1 | 1300955..1301350 | missense_variant | MODERATE | c.196C>T | p.His66Tyr |
| NC_017564 | 1378582 | G | A | putP | Y11_RS06485 | sodium/proline symporter PutP | WP_005177983.1 | complement(1378390..1379874) | synonymous_variant | LOW | c.1293C>T | p.Leu431Leu |
| NC_017564 | 1381121 | G | A | putA | Y11_RS06490 | trifunctional transcriptional regulator/proline dehydrogenase/L-glutamate gamma-semialdehyde dehydrogenase | WP_005160276.1 | 1380401..1384390 | missense_variant | MODERATE | c.721G>A | p.Gly241Ser |
| NC_017564 | 1424592 | G | A | flgG | Y11_RS06710 | flagellar basal-body rod protein FlgG | WP_005160438.1 | complement(1424408..1425190) | missense_variant | MODERATE | c.599C>T | p.Ala200Val |
| NC_017564 | 1429318 | C | T | flgA | Y11_RS06740 | flagellar basal body P-ring formation protein FlgA | WP_005160464.1 | 1429252..1429926 | stop_gained | HIGH | c.67C>T | p.Gln23* |
| NC_017564 | 1433374 | C | T | inv | Y11_RS06755 | inverse autotransporter invasin Inv | WP_014609051.1 | complement(1430869..1433376) | initiator_codon_variant | LOW | c.3G>A | p.Met1? |
| NC_017564 | 1540420 | G | A | | Y11_RS07220 | ABC transporter substrate-binding protein | WP_005165439.1 | 1539751..1540782 | missense_variant | MODERATE | c.670G>A | p.Gly224Ser |
| NC_017564 | 1638801 | G | A | | Y11_RS07685 | ABC transporter permease | WP_005157737.1 | 1638706..1639689 | synonymous_variant | LOW | c.96G>A | p.Val32Val |
| NC_017564 | 1787309 | G | A | urtB | Y11_RS08360 | urea ABC transporter permease subunit UrtB | WP_014609102.1 | complement(1786712..1788313) | synonymous_variant | LOW | c.1005C>T | p.Thr335Thr |
| NC_017564 | 1935973 | C | T | corC | Y11_RS09005 | CNNM family magnesium/cobalt transport protein CorC | WP_005158416.1 | 1935157..1936035 | missense_variant | MODERATE | c.817C>T | p.Arg273Cys |
| NC_017564 | 1963052 | C | T | | Y11_RS09145 | metallophosphoesterase | WP_005158354.1 | complement(1962861..1963505) | missense_variant | MODERATE | c.454G>A | p.Glu152Lys |
| NC_017564 | 1975952 | G | A | | Y11_RS00380 | | | | intragenic_variant | MODIFIER | n.1975952G>A | |
| NC_017564 | 2022198 | G | A | hemH | Y11_RS09425 | ferrochelatase | WP_005158187.1 | complement(2021995..2022978) | missense_variant | MODERATE | c.781C>T | p.His261Tyr |

| | | | | | | | | | | | | |
|---|---|---|---|---|---|---|---|---|---|---|---|---|
| NC_017564 | 2038641 | CGTTC | TGTTA | acrB | Y11_RS09495 | multidrug efflux RND transporter permease subunit | WP_005158148.1 | 2037650..2040802 | missense_variant | MODERATE | c.992_996delCGTTCinsTGTTA | p.ProPhe331LeuLeu |
| NC_017564 | 2039040 | G | A | acrB | Y11_RS09495 | multidrug efflux RND transporter permease subunit | WP_005158148.1 | 2037650..2040802 | missense_variant | MODERATE | c.1391G>A | p.Gly464Glu |
| NC_017564 | 2142508 | G | A | | Y11_RS00380 | | | | intragenic_variant | MODIFIER | n.2142508G>A | |
| NC_017564 | 2195957 | C | T | mtnA | Y11_RS10130 | S-methyl-5-thioribose-1-phosphate isomerase | WP_005164971.1 | 2195063..2196103 | missense_variant | MODERATE | c.895C>T | p.His299Tyr |
| NC_017564 | 2335296 | G | A | cobA | Y11_RS10780 | uroporphyrinogen-III C-methyltransferase | WP_005163311.1 | complement(2334405..2335823) | synonymous_variant | LOW | c.528C>T | p.Arg176Arg |
| NC_017564 | 2350148 | G | A | serA | Y11_RS10865 | phosphoglycerate dehydrogenase | WP_005163339.1 | complement(2349556..2350797) | missense_variant | MODERATE | c.650C>T | p.Thr217Ile |
| NC_017564 | 2374897 | G | A | | Y11_RS10980 | YqgE/AlgH family protein | WP_020282839.1 | 2374791..2375354 | missense_variant | MODERATE | c.107G>A | p.Gly36Asp |
| NC_017564 | 2599791 | C | T | | Y11_RS12105 | NADPH-dependent 2,4-dienoyl-CoA reductase | WP_014609190.1 | 2598223..2600244 | synonymous_variant | LOW | c.1569C>T | p.Ser523Ser |
| NC_017564 | 2655984 | C | T | arcB | Y11_RS12360 | aerobic respiration two-component sensor histidine kinase ArcB | WP_005162394.1 | complement(2654296..2656632) | missense_variant | MODERATE | c.649G>A | p.Glu217Lys |
| NC_017564 | 2656579 | C | T | arcB | Y11_RS12360 | aerobic respiration two-component sensor histidine kinase ArcB | WP_005162394.1 | complement(2654296..2656632) | synonymous_variant | LOW | c.54G>A | p.Leu18Leu |
| NC_017564 | 2667005 | C | T | rpsI | Y11_RS12400 | 30S ribosomal protein S9 | WP_005162424.1 | complement(2666950..2667342) | missense_variant | MODERATE | c.338G>A | p.Arg113His |
| NC_017564 | 2881034 | G | A | metF | Y11_RS13280 | methylenetetrahydrofolate reductase | WP_005163851.1 | complement(2880966..2881850) | missense_variant | MODERATE | c.817C>T | p.His273Tyr |
| NC_017564 | 2941323 | C | T | | Y11_RS13555 | Spy/CpxP family protein refolding chaperone | WP_005164704.1 | complement(2940913..2941410) | missense_variant | MODERATE | c.88G>A | p.Gly30Arg |
| NC_017564 | 2976987 | G | A | pdeR | Y11_RS13730 | cyclic di-GMP phosphodiesterase | WP_005165873.1 | 2976363..2978357 | missense_variant | MODERATE | c.625G>A | p.Gly209Arg |
| NC_017564 | 3090987 | G | A | gyrB | Y11_RS14215 | DNA topoisomerase (ATP-hydrolyzing) subunit B | WP_005161097.1 | 3088702..3091116 | missense_variant | MODERATE | c.2286G>A | p.Met762Ile |
| NC_017564 | 3245271 | G | A | yapE | Y11_RS14815 | autotransporter adhesin YapE | WP_005157376.1 | 3243036..3246278 | missense_variant | MODERATE | c.2236G>A | p.Val746Ile |
| NC_017564 | 3338974 | G | A | | Y11_RS15165 | hypothetical protein | WP_005178422.1 | complement(3338206..3339036) | synonymous_variant | LOW | c.63C>T | p.Ala21Ala |
| NC_017564 | 3491775 | G | A | rffA | Y11_RS15945 | dTDP-4-amino-4,6-dideoxygalactose transaminase | WP_013649042.1 | 3491154..3492284 | missense_variant | MODERATE | c.622G>A | p.Glu208Lys |
| NC_017564 | 3556799 | G | A | livF | Y11_RS16255 | high-affinity branched-chain amino acid ABC transporter ATP-binding protein LivF | WP_005163929.1 | 3556671..3557372 | synonymous_variant | LOW | c.129G>A | p.Leu43Leu |
| NC_017564 | 3852561 | G | A | phnF | Y11_RS22520 | phosphonate metabolism transcriptional regulator PhnF | WP_005156203.1 | complement(3851851..3852576) | missense_variant | MODERATE | c.16C>T | p.Pro6Ser |
| NC_017564 | 3863881 | G | A | | Y11_RS17710 | ABC transporter substrate-binding protein | WP_005156162.1 | complement(3863502..3865070) | missense_variant | MODERATE | c.1190C>T | p.Pro397Leu |
| NC_017564 | 4013902 | C | T | carA | Y11_RS18330 | glutamine-hydrolyzing carbamoyl-phosphate synthase small subunit | WP_005156985.1 | 4012938..4014101 | missense_variant | MODERATE | c.965C>T | p.Pro322Leu |
| NC_017564 | 4026646 | C | T | lptD | Y11_RS18385 | LPS assembly protein LptD | WP_005156954.1 | complement(4024986..4027352) | missense_variant | MODERATE | c.707G>A | p.Ser236Asn |
| NC_017564 | 4200200 | G | A | | Y11_RS00380 | | | | intragenic_variant | MODIFIER | n.4200200G>A | |
| NC_017564 | 4457465 | G | A | suhB | Y11_RS20330 | inositol-1-monophosphatase | WP_005159504.1 | complement(4456847..4457653) | synonymous_variant | LOW | c.189C>T | p.Thr63Thr |

**Table S4.** Synapomorphic genes

| Ppanggolin ID | Genbank ID | Description |
|---|---|---|
| H60887_CDS_0252 | WP_005166068 | phosphoribosylamine--glycine ligase |
| H60887_CDS_1258 | YP_001005538 | hypothetical protein |
| H60887_CDS_2071 | WP_005165195 | PhoPQ-regulated protein |
| H60887_CDS_3051 | WP_005166190 | LysR family transcriptional regulator |
| IP37498_CDS_0391 | WP_016266285 | RNA polymerase sigma factor RpoD |
| H60887_CDS_3187 | WP_005160260 | deferrochelatase/peroxidase EfeB |
| H60887_CDS_0385 | YP_001008098 | phosphoribulokinase 1 |
| H60887_CDS_1374 | WP_072079261 | helix-turn-helix transcriptional regulator |
| H60887_CDS_2167 | WP_005159304 | DNA topoisomerase (ATP-hydrolyzing) subunit A (*gyrA* ) |
| H60887_CDS_3205 | WP_005163567 | PAS and helix-turn-helix domain-containing protein |
| H60887_CDS_2097 | WP_005159073 | thymidine phosphorylase |
| H60887_CDS_3181 | WP_005177983 | sodium/proline symporter PutP |
| H60887_CDS_0365 | WP_005159706 | SPOR domain-containing protein |
| H60887_CDS_1362 | WP_005157647 | cobalt-factor II C(20)-methyltransferase |
| H60887_CDS_2147 | WP_005158596 | endonuclease SmrB |
| H60887_CDS_2219 | WP_005165599 | ubiquinone biosynthesis regulatory protein kinase UbiB |
| H60887_CDS_3357 | YP_001008380 | hypothetical protein |
| H60887_CDS_0562 | WP_005161207 | low affinity potassium transporter Kup |
| H60887_CDS_1459 | WP_005166443 | formimidoylglutamate deiminase |
| H60887_CDS_2222 | WP_005165593 | Sec-independent protein translocase subunit TatC |
| H60887_CDS_0257 | WP_005166266 | GNAT family N-acetyltransferase |
| H60887_CDS_1293 | WP_005163311 | uroporphyrinogen-III C-methyltransferase |
| H60887_CDS_2073 | YP_001004950 | two-component response regulator |
| H60887_CDS_3110 | WP_162197869 | methionine synthase |
| H60887_CDS_0273 | WP_005160457 | flagellar hook assembly protein FlgD |
| H60887_CDS_1762 | YP_001008190 | phosphate transport protein |
| H60887_CDS_2362 | WP_005179944 | TonB-dependent siderophore receptor |
| H60887_CDS_3672 | WP_005165565 | inositol monophosphatase |
| H60887_CDS_0812 | WP_016266346 | L-threonylcarbamoyladenylate synthase type 1 TsaC |
| H60887_CDS_1782 | WP_005157438 | MurR/RpiR family transcriptional regulator |
| H60887_CDS_0211 | WP_005166512 | ATP-dependent chaperone ClpB |
| H60887_CDS_1243 | WP_005164872 | sulfate/thiosulfate ABC transporter ATP-binding protein CysA |
| H60887_CDS_2033 | WP_005157912 | N-methyl-L-tryptophan oxidase |
| H60887_CDS_3048 | WP_005162867 | putrescine ABC transporter permease PotH |
| IP37401_CDS_2750 | WP_005163368 | family 20 glycosylhydrolase |
| H60887_CDS_2314 | WP_016266165 | zinc transporter ZntB |
| H60887_CDS_3536 | WP_011816794 | excinuclease ABC subunit B |
| H60887_CDS_0644 | WP_005166111 | diaminopimelate epimerase |
| H60887_CDS_1574 | WP_013650082 | L,D-transpeptidase |
| H60887_CDS_2326 | WP_005163605 | prolyl aminopeptidase |
| H60887_CDS_0158 | WP_057621728 | phage major capsid protein, P2 family |
| H60887_CDS_1190 | WP_005157168 | DNA-binding domain-containing protein |
| H60887_CDS_1989 | WP_005164939 | LysR family transcriptional regulator |
| H60887_CDS_2872 | WP_005165887 | bifunctional GTP diphosphokinase/guanosine-3',5'-bis pyrophosphate 3'-pyrophosphohydrolase |
| H60887_CDS_3993 | WP_005166177 | ShlB/FhaC/HecB family hemolysin secretion/activation protein |
| H60887_CDS_1438 | WP_005165324 | phage tail tape measure protein |
| H60887_CDS_2184 | WP_005180990 | envelope stress sensor histidine kinase CpxA |
| H60887_CDS_3323 | YP_001005932 | PTS system glucose-specific transporter subunit IIBC |
| H60887_CDS_0555 | WP_005161186 | tRNA uridine-5-carboxymethylaminomethyl(34) synthesis enzyme MnmG |
| H60887_CDS_1440 | WP_005165328 | phage late control D family protein |
| H60887_CDS_3687 | WP_005159375 | 6-phospho-beta-glucosidase |
| H60887_CDS_0919 | WP_005157341 | cellulose synthase operon protein YhjQ |
| H60887_CDS_1832 | WP_005159455 | two-component system response regulator GlrR |
| H60887_CDS_2448 | WP_005160774 | DNA topoisomerase IV subunit B |
| H60887_CDS_3709 | WP_005165857 | type I DNA topoisomerase |
| 191430_CDS_2033 | WP_071841733 | glycine cleavage system transcriptional repressor |
| H60887_CDS_1004 | WP_005156784 | carbonate dehydratase |
| H60887_CDS_1899 | WP_014609259 | 6-N-hydroxylaminopurine resistance protein |
| H60887_CDS_2611 | WP_005163370 | ferric iron uptake transcriptional regulator |
| H60887_CDS_3739 | WP_005165738 | phage-like membrane protein |
| H60887_CDS_2370 | WP_005160583 | phosphate regulon sensor histidine kinase PhoR |
| H60887_CDS_3677 | WP_005159346 | methyl-accepting chemotaxis protein |
| H60887_CDS_0826 | WP_011816964 | DNA polymerase III subunit alpha |
| H60887_CDS_1794 | WP_005157471 | acetolactate synthase AlsS |
| H60887_CDS_2441 | WP_005160755 | YgiQ family radical SAM protein |
| H60887_CDS_0142 | YP_001008053 | 50S ribosomal protein L6 |
| H60887_CDS_1145 | WP_005161116 | methyl-accepting chemotaxis protein |

| | | |
|---|---|---|
| H60887_CDS_1984 | WP_005164944 | carboxy terminal-processing peptidase |
| H60887_CDS_2821 | WP_016266231 | fimbrial biogenesis outer membrane usher protein |
| H60887_CDS_3928 | WP_005160713 | type II and III secretion system protein family protein |
| H60887_CDS_0111 | WP_005165794 | DNA-directed RNA polymerase subunit beta' |
| H60887_CDS_1082 | WP_005162233 | bifunctional hydroxymethylpyrimidine kinase/phosphomethylpyrimidine kinase |
| H60887_CDS_1966 | WP_016266222 | chemotaxis protein CheA |
| H60887_CDS_2761 | WP_005161635 | succinate CoA transferase |
| H60887_CDS_3872 | WP_005157047 | transaldolase |
| 191430_CDS_0267 | WP_005178371 | EAL domain-containing protein |
| H60887_CDS_0988 | WP_016266118 | phosphate acetyltransferase |
| H60887_CDS_1836 | WP_005159441 | phosphoribosylformylglycinamidine synthase |
| H60887_CDS_2545 | WP_005158394 | endolytic peptidoglycan transglycosylase RlpA |
| H60887_CDS_3720 | WP_005165542 | biotin/lipoyl-binding protein |
| 191430_CDS_3879 | WP_005158334 | aromatic amino acid lyase |
| H60887_CDS_1017 | WP_005158160 | mechanosensitive channel MscK |
| H60887_CDS_1955 | WP_005164506 | HAMP domain-containing protein |
| H60887_CDS_2720 | WP_005165719 | maltose/maltodextrin ABC transporter ATP-binding protein MalK |
| H60887_CDS_3799 | WP_005166203 | DUF1120 domain-containing protein |
| H60887_CDS_1537 | WP_005156284 | ATP-dependent zinc metalloprotease FtsH |
| H60887_CDS_2296 | WP_005164625 | serine/threonine transporter SstT |
| H60887_CDS_3487 | WP_005158806 | alginate lyase family protein |
| H60887_CDS_0641 | WP_005166106 | class I adenylate cyclase |
| H60887_CDS_1566 | WP_005159990 | aminopeptidase N |
| H60887_CDS_3358 | WP_005164159 | bifunctional acetaldehyde-CoA/alcohol dehydrogenase |
| H60887_CDS_0580 | WP_005165945 | peptidoglycan DD-metalloendopeptidase family protein |
| H60887_CDS_1483 | WP_005162319 | hydrogenase 4 subunit B |
| H60887_CDS_2242 | WP_005160498 | glycerophosphoryl diester phosphodiesterase |
| H60887_CDS_3374 | WP_050326492 | S8 family serine peptidase |
| H60887_CDS_0165 | WP_005162408 | ClpXP protease specificity-enhancing factor |
| H60887_CDS_1215 | WP_005157106 | glutathione synthase |
| H60887_CDS_1990 | WP_005164938 | MFS transporter |
| H60887_CDS_3037 | WP_005162896 | HAAAP family serine/threonine permease |
| IP35131_CDS_0782 | WP_005165352 | multidrug efflux MFS transporter EmrD |
| H60887_CDS_1307 | WP_005163282 | DNA mismatch repair protein MutS |
| H60887_CDS_2075 | WP_005159031 | MDR efflux pump AcrAB transcriptional activator RobA |
| H60887_CDS_3127 | WP_005163006 | long-chain-fatty-acid--CoA ligase FadD |
| H60887_CDS_0333 | YP_001007135 | succinate dehydrogenase cytochrome b556 small membrane subunit |
| H60887_CDS_1331 | WP_005157709 | BMC domain-containing protein |
| H60887_CDS_3575 | WP_005158665 | 6-phospho-beta-glucosidase |
| H60887_CDS_0729 | WP_005165050 | nitrogen regulation protein NR(II) |
| H60887_CDS_1591 | WP_004391091 | 30S ribosomal protein S1 |
| H60887_CDS_2330 | WP_005163607 | hypothetical protein |
| H60887_CDS_3582 | WP_005159102 | ABC transporter substrate-binding protein |
| H60887_CDS_0174 | YP_001005015 | penicillin-binding protein 3 |
| H60887_CDS_1220 | WP_005157092 | YeeE/YedE family protein |
| H60887_CDS_1995 | WP_016266154 | fimbrial biogenesis outer membrane usher protein |
| H60887_CDS_3040 | WP_005162886 | undecaprenyl-diphosphate phosphatase |
| IP35463_CDS_3158 | WP_005164219 | PHP domain-containing protein |
| H60887_CDS_0465 | WP_005164089 | glutamate-1-semialdehyde 2,1-aminomutase |
| H60887_CDS_1403 | WP_023161097 | adhesin/invasin protein |
| H60887_CDS_2175 | WP_014609236 | murein hydrolase activator EnvC |
| H60887_CDS_3243 | WP_005165647 | superoxide dismutase [Fe] |
| H60887_CDS_0475 | WP_005164102 | GTP diphosphokinase |
| 191430_CDS_3430 | WP_005163484 | PLP-dependent cysteine synthase family protein |
| H60887_CDS_1013 | WP_050326420 | efflux RND transporter permease subunit |
| H60887_CDS_1924 | WP_014609156 | Na/Pi cotransporter family protein |
| H60887_CDS_2715 | WP_005165714 | chorismate lyase |
| H60887_CDS_3756 | WP_005165775 | MFS transporter |
| H60887_CDS_0589 | YP_001005186 | anaerobic C4-dicarboxylate transporter |
| H60887_CDS_1533 | WP_005166341 | N-acetylmuramoyl-L-alanine amidase |
| H60887_CDS_2289 | WP_020282746 | hypothetical protein |
| H60887_CDS_3406 | WP_005162911 | D-hexose-6-phosphate mutarotase |
| H60887_CDS_0624 | WP_014609354 | phni protein |
| H60887_CDS_0760 | WP_005163155 | AsmA2 domain-containing protein |
| H60887_CDS_1715 | WP_005161239 | succinylglutamate-semialdehyde dehydrogenase |
| H60887_CDS_2355 | WP_014609131 | glycosyl transferase |
| H60887_CDS_3657 | WP_005163404 | biotin-dependent carboxyltransferase family protein |
| H60887_CDS_0763 | WP_005163164 | type II toxin-antitoxin system HipA family toxin |
| 191430_CDS_1101 | WP_014609197 | TIGR01212 family radical SAM protein |
| H60887_CDS_0995 | WP_005156446 | phosphatidylserine decarboxylase |

| | | |
|---|---|---|
| H60887_CDS_1837 | WP_005159436 | membrane-bound lytic murein transglycosylase MltF |
| H60887_CDS_2599 | WP_005156333 | HlyC/CorC family transporter |
| H60887_CDS_3731 | WP_001386054 | phage/plasmid replication protein, II/X family |

**Table S5.** PAML results

| Ppanggolin ID | Genbank ID | Description | Positive selection signal | Strains involved |
|---|---|---|---|---|
| H60887_CDS_1751 | WP_005163474 | acyl-CoA thioesterase II [Yersinia enterocolitica]. | shorter C-terminal part | Ye.3 and Ye.4 |
| H60887_CDS_1762 | YP_001008190 | phosphate transport protein [Yersinia enterocolitica subsp. enterocolitica 8081]. | G280S | Ye.1, Ye.2, Ye.3 and Ye.4 |

**Tables S6.** Ye.1 to Ye.4 proteome pairwise comparisons. Proteins were mapped on the Y11 reference genome. 0 corresponds to proteins not detected in any conditions of the comparison. +15 and -15 corresponds to proteins detected only in the first or second condition of the comparison, respectively.

| | | | | Log2(foldchange) | | | | | | p-value | | | | | | adjusted p-value | | | | | | | | | |
|---|---|---|---|---|---|---|---|---|---|---|---|---|---|---|---|---|---|---|---|---|---|---|---|---|---|
| Locus | Old Locus | Gene | Product | Ye.1 vs Ye.2 | Ye.1 vs Ye.3 | Ye.1 vs Ye.4 | Ye.2 vs Ye.3 | Ye.2 vs Ye.4 | Ye.3 vs Ye.4 | Ye.1 vs Ye.2 | Ye.1 vs Ye.3 | Ye.1 vs Ye.4 | Ye.2 vs Ye.3 | Ye.2 vs Ye.4 | Ye.3 vs Ye.4 | Ye.1 vs Ye.2 | Ye.1 vs Ye.3 | Ye.1 vs Ye.4 | Ye.2 vs Ye.3 | Ye.2 vs Ye.4 | Ye.3 vs Ye.4 | Position start | Position end | Length | Strand |
| Y11_RS00005 | Y11_00001 | hda | DnaA inactivator Hda | -0,14 | 0,47 | 0,32 | 0,61 | 0,46 | -0,15 | 5,6E-02 | 1,3E-03 | 5,1E-02 | 2,4E-04 | 1,1E-02 | 3,8E-01 | 0,0E+00 | 0,0E+00 | 0,0E+00 | 0,0E+00 | 0,0E+00 | 0,0E+00 | 13 | 720 | 708 | + |
| Y11_RS00010 | Y11_00011 | arsC | arsenate reductase (glutaredoxin) | 0,52 | 1,16 | 1,69 | 0,63 | 1,17 | 0,53 | 2,7E-05 | 1,2E-07 | 2,5E-06 | 1,2E-05 | 3,9E-05 | 6,5E-03 | 0,0E+00 | 4,6E-09 | 5,0E-08 | 0,0E+00 | 1,1E-06 | 0,0E+00 | 768 | 1121 | 354 | - |
| Y11_RS00015 | Y11_00021 | | M48 family metallopeptidase | 0,2 | 0,57 | 1 | 0,37 | 0,8 | 0,43 | 6,3E-02 | 7,4E-04 | 2,9E-05 | 7,1E-04 | 1,4E-05 | 3,0E-03 | 0,0E+00 | 0,0E+00 | 0,0E+00 | 0,0E+00 | 0,0E+00 | 0,0E+00 | 1133 | 2659 | 1527 | - |
| Y11_RS00025 | Y11_00051 | bcp | thioredoxin-dependent thiol peroxidase | 0,28 | 0,41 | 0,79 | 0,13 | 0,51 | 0,38 | 5,3E-04 | 1,0E-04 | 1,1E-05 | 9,6E-02 | 4,9E-04 | 4,2E-03 | 0,0E+00 | 0,0E+00 | 0,0E+00 | 0,0E+00 | 0,0E+00 | 0,0E+00 | 4441 | 4911 | 471 | - |
| Y11_RS00030 | Y11_00061 | | glycine cleavage system transcriptional repressor | -0,14 | 0,45 | 0,23 | 0,59 | 0,37 | -0,22 | 3,1E-01 | 1,8E-03 | 2,4E-02 | 1,7E-03 | 1,3E-02 | 2,4E-02 | 0,0E+00 | 0,0E+00 | 0,0E+00 | 0,0E+00 | 0,0E+00 | 0,0E+00 | 4914 | 5483 | 570 | - |
| Y11_RS00035 | Y11_00071 | dapA | 4-hydroxy-tetrahydrodipicolinate synthase | 0,01 | 0,25 | 0,44 | 0,24 | 0,43 | 0,19 | 9,0E-01 | 3,4E-03 | 1,4E-04 | 4,2E-03 | 1,6E-04 | 2,3E-03 | 0,0E+00 | 0,0E+00 | 0,0E+00 | 0,0E+00 | 0,0E+00 | 0,0E+00 | 5648 | 6529 | 882 | + |
| Y11_RS00040 | Y11_00081 | bamC | outer membrane protein assembly factor BamC | -0,04 | 0,09 | 0,26 | 0,13 | 0,3 | 0,17 | 2,3E-01 | 1,1E-02 | 2,2E-03 | 1,0E-03 | 8,4E-04 | 1,6E-02 | 0,0E+00 | 0,0E+00 | 0,0E+00 | 0,0E+00 | 0,0E+00 | 0,0E+00 | 6546 | 7604 | 1059 | + |
| Y11_RS00045 | Y11_00091 | | phosphoribosylaminoimidazolesuccinocarboxamide synthase | 0,21 | 0,38 | 0,16 | 0,16 | -0,05 | -0,21 | 5,2E-02 | 1,3E-02 | 1,7E-01 | 2,7E-01 | 7,0E-01 | 1,9E-01 | 0,0E+00 | 0,0E+00 | 0,0E+00 | 0,0E+00 | 0,0E+00 | 0,0E+00 | 7817 | 8530 | 714 | + |
| Y11_RS00050 | Y11_00101 | | zinc metallopeptidase | 0,29 | 1,72 | 1,43 | 1,43 | 1,14 | -0,29 | 3,6E-03 | 5,6E-09 | 2,6E-05 | 1,0E-07 | 1,7E-04 | 1,3E-01 | 0,0E+00 | 4,0E-10 | 4,0E-07 | 3,4E-09 | 4,2E-06 | 0,0E+00 | 8732 | 9601 | 870 | + |
| Y11_RS00110 | Y11_00221 | ail | attachment invasion locus protein Ail | 0 | 2,62 | 3,25 | 2,63 | 3,25 | 0,62 | 9,7E-01 | 4,5E-11 | 1,5E-09 | 6,8E-10 | 3,8E-09 | 4,1E-04 | 0,0E+00 | 1,4E-11 | 1,3E-10 | 6,5E-11 | 4,5E-10 | 0,0E+00 | 17349 | 17885 | 537 | + |
| Y11_RS00130 | Y11_00261 | | M15 family metallopeptidase | 0,2 | 0 | -0,07 | -0,2 | -0,26 | -0,06 | 1,4E-01 | 9,8E-01 | 5,8E-01 | 9,1E-02 | 4,9E-02 | 5,3E-01 | 0,0E+00 | 0,0E+00 | 0,0E+00 | 0,0E+00 | 0,0E+00 | 0,0E+00 | 19548 | 20252 | 705 | - |
| Y11_RS00135 | Y11_00271 | dapE | succinyl-diaminopimelate desuccinylase | 0,12 | -0,33 | -0,4 | -0,44 | -0,51 | -0,07 | 6,1E-02 | 3,1E-03 | 9,4E-04 | 9,1E-05 | 2,9E-05 | 4,4E-01 | 0,0E+00 | 0,0E+00 | 0,0E+00 | 0,0E+00 | 0,0E+00 | 0,0E+00 | 20255 | 21382 | 1128 | - |
| Y11_RS00140 | Y11_00281 | | ArsC family reductase | 0,74 | -0,25 | -1,02 | -0,99 | -1,76 | -0,76 | 1,4E-06 | 2,8E-02 | 8,0E-07 | 2,4E-06 | 2,0E-09 | 4,5E-05 | 0,0E+00 | 0,0E+00 | 1,9E-08 | 0,0E+00 | 2,8E-10 | 0,0E+00 | 21384 | 21806 | 423 | - |
| Y11_RS00155 | Y11_00321 | acrD | multidrug efflux RND transporter permease AcrD | -0,54 | -2,03 | -0,85 | -1,49 | -0,31 | 1,18 | 1,5E-05 | 2,8E-07 | 9,6E-06 | 1,8E-06 | 3,0E-03 | 2,5E-05 | 0,0E+00 | 9,8E-09 | 0,0E+00 | 4,3E-08 | 0,0E+00 | 2,6E-06 | 24554 | 27685 | 3132 | - |
| Y11_RS00160 | Y11_00331 | | helix-turn-helix domain-containing protein | 0,42 | 1,33 | 2,78 | 0,91 | 2,36 | 1,45 | 3,3E-01 | 2,2E-04 | 2,8E-06 | 4,4E-02 | 3,5E-04 | 1,1E-04 | 0,0E+00 | 4,7E-06 | 5,5E-08 | 0,0E+00 | 8,4E-06 | 9,1E-06 | 27895 | 28173 | 279 | + |
| Y11_RS00165 | Y11_00341 | | response regulator | 0,28 | 1,29 | 1,91 | 1,01 | 1,63 | 0,63 | 1,2E-02 | 6,9E-08 | 2,2E-10 | 4,9E-06 | 3,7E-08 | 7,7E-06 | 0,0E+00 | 3,0E-09 | 3,7E-11 | 1,1E-07 | 2,7E-09 | 0,0E+00 | 28212 | 28841 | 630 | - |
| Y11_RS00175 | Y11_00361 | napD | chaperone NapD | -0,32 | 0,96 | 1,6 | 1,28 | 1,92 | 0,64 | 1,4E-01 | 9,9E-03 | 4,3E-04 | 4,0E-03 | 2,9E-04 | 1,3E-01 | 0,0E+00 | 0,0E+00 | 5,3E-06 | 6,6E-05 | 7,1E-06 | 0,0E+00 | 29839 | 30105 | 267 | + |
| Y11_RS00180 | Y11_00371 | napA | nitrate reductase catalytic subunit NapA | 1,57 | 3,2 | 3,76 | 1,63 | 2,19 | 0,56 | 2,8E-05 | 1,2E-10 | 6,9E-10 | 1,4E-05 | 3,2E-06 | 3,5E-04 | 1,3E-06 | 2,4E-11 | 8,1E-11 | 2,8E-07 | 1,2E-07 | 0,0E+00 | 30102 | 32597 | 2496 | + |
| Y11_RS00185 | Y11_00381 | napB | nitrate reductase cytochrome c-type subunit | -0,7 | 2,46 | 3,93 | 3,16 | 4,58 | 1,48 | 4,1E-04 | 2,7E-07 | 2,1E-09 | 1,3E-07 | 3,0E-09 | 5,9E-05 | 0,0E+00 | 9,5E-09 | 1,7E-10 | 4,1E-09 | 3,7E-10 | 5,1E-06 | 32668 | 33135 | 468 | + |
| Y11_RS00190 | Y11_00391 | napC | cytochrome c-type protein NapC | -0,6 | -0,08 | 1,04 | 0,53 | 1,65 | 1,12 | 1,2E-01 | 8,0E-01 | 5,5E-02 | 1,9E-01 | 1,2E-02 | 4,8E-02 | 0,0E+00 | 0,0E+00 | 6,2E-04 | 0,0E+00 | 2,6E-04 | 2,6E-03 | 33163 | 33762 | 600 | + |
| Y11_RS00195 | Y11_00401 | nudK | GDP-mannose pyrophosphatase NudK | -0,09 | 0,71 | 0,71 | 0,8 | 0,8 | 0 | 6,9E-01 | 6,9E-03 | 4,2E-03 | 4,1E-03 | 2,5E-03 | 1,0E+00 | 0,0E+00 | 0,0E+00 | 0,0E+00 | 0,0E+00 | 0,0E+00 | 0,0E+00 | 33944 | 34519 | 576 | + |
| Y11_RS00200 | Y11_00411 | maeB | NADP-dependent oxaloacetate-decarboxylating malate dehydrogenase | -0,26 | 0,35 | 1,72 | 0,61 | 1,97 | 1,37 | 7,1E-05 | 4,5E-04 | 4,3E-09 | 1,5E-05 | 2,4E-09 | 2,3E-07 | 0,0E+00 | 0,0E+00 | 2,8E-10 | 0,0E+00 | 3,2E-10 | 6,2E-08 | 34740 | 37019 | 2280 | + |
| Y11_RS00220 | Y11_00451 | | YaiI/YqxD family protein | 0,05 | 0,33 | 0,09 | 0,29 | 0,05 | -0,24 | 7,2E-01 | 1,1E-03 | 5,2E-01 | 3,5E-02 | 7,8E-01 | 1,0E-01 | 0,0E+00 | 0,0E+00 | 0,0E+00 | 0,0E+00 | 0,0E+00 | 0,0E+00 | 39743 | 40201 | 459 | - |
| Y11_RS00230 | Y11_00471 | | GNAT family acetyltransferase | 0,2 | 1,17 | 1,24 | 0,97 | 1,04 | 0,07 | 2,5E-01 | 3,7E-05 | 4,3E-05 | 6,7E-05 | 8,0E-05 | 5,7E-01 | 0,0E+00 | 8,4E-07 | 6,2E-07 | 0,0E+00 | 2,1E-06 | 0,0E+00 | 41373 | 41798 | 426 | + |
| Y11_RS00235 | Y11_00481 | | YgiW/YdeI family stress tolerance OB fold protein | -0,12 | 0,93 | 1,37 | 1,05 | 1,49 | 0,44 | 8,2E-02 | 5,4E-07 | 3,7E-05 | 5,0E-08 | 1,6E-05 | 2,7E-02 | 0,0E+00 | 0,0E+00 | 5,5E-07 | 1,9E-09 | 5,1E-07 | 0,0E+00 | 41952 | 42365 | 414 | - |
| Y11_RS00240 | Y11_00491 | | RpoE-regulated lipoprotein | -0,02 | 0,46 | 1,31 | 0,47 | 1,32 | 0,85 | 9,3E-01 | 4,3E-02 | 2,1E-03 | 1,4E-02 | 1,2E-03 | 1,4E-02 | 0,0E+00 | 0,0E+00 | 2,5E-05 | 0,0E+00 | 2,7E-05 | 0,0E+00 | 42749 | 43360 | 612 | + |
| Y11_RS00245 | Y11_00501 | | Dyp-type peroxidase | -0,66 | 0,52 | 0,97 | 1,18 | 1,63 | 0,45 | 2,3E-06 | 9,3E-04 | 6,2E-07 | 3,6E-06 | 1,9E-08 | 3,9E-03 | 0,0E+00 | 0,0E+00 | 0,0E+00 | 7,9E-08 | 1,6E-09 | 0,0E+00 | 43536 | 44441 | 906 | + |
| Y11_RS00255 | Y11_00531 | | dihydrodipicolinate synthase family protein | 0,19 | 0,79 | 1,09 | 0,61 | 0,9 | 0,3 | 1,4E-02 | 7,8E-07 | 8,5E-08 | 1,0E-05 | 6,3E-07 | 1,5E-03 | 0,0E+00 | 0,0E+00 | 2,8E-09 | 0,0E+00 | 0,0E+00 | 0,0E+00 | 46069 | 46959 | 891 | - |
| Y11_RS00260 | Y11_00541 | | putative N-acetylmannosamine-6-phosphate 2-epimerase | 0,96 | 1,07 | 0,81 | 0,11 | -0,14 | -0,26 | 6,2E-03 | 2,7E-03 | 2,1E-02 | 7,0E-01 | 6,6E-01 | 4,2E-01 | 0,0E+00 | 5,4E-05 | 0,0E+00 | 0,0E+00 | 0,0E+00 | 0,0E+00 | 47042 | 47710 | 669 | - |
| Y11_RS00265 | Y11_00561 | | N-acetylmannosamine kinase | 0,31 | 0,27 | 1,26 | -0,03 | 0,95 | 0,98 | 1,1E-02 | 6,0E-03 | 2,4E-07 | 7,0E-01 | 6,3E-06 | 6,1E-07 | 0,0E+00 | 0,0E+00 | 6,7E-09 | 0,0E+00 | 0,0E+00 | 0,0E+00 | 47987 | 48895 | 909 | + |
| Y11_RS00270 | Y11_00571 | | YhcH/YjgK/YiaL family protein | 0,19 | -1,14 | 0,25 | -1,32 | 0,07 | 1,39 | 2,7E-01 | 8,6E-05 | 9,5E-02 | 1,2E-04 | 7,1E-01 | 4,1E-05 | 0,0E+00 | 1,9E-06 | 0,0E+00 | 2,2E-06 | 0,0E+00 | 3,7E-06 | 48929 | 49393 | 465 | + |
| Y11_RS00275 | Y11_00581 | | MurR/RpiR family transcriptional regulator | -0,53 | -1,13 | -0,13 | -0,6 | 0,4 | 1 | 1,3E-03 | 6,0E-06 | 4,4E-01 | 2,5E-04 | 2,8E-02 | 1,5E-04 | 0,0E+00 | 1,5E-07 | 0,0E+00 | 0,0E+00 | 0,0E+00 | 0,0E+00 | 49589 | 50467 | 879 | + |
| Y11_RS00285 | Y11_00611 | | sulfate ABC transporter substrate-binding protein | 0,13 | 1,01 | 3,21 | 0,88 | 3,09 | 2,2 | 9,3E-02 | 4,8E-07 | 1,2E-09 | 1,8E-07 | 7,6E-10 | 1,2E-08 | 0,0E+00 | 1,5E-08 | 1,2E-10 | 0,0E+00 | 1,5E-10 | 8,3E-09 | 52866 | 53894 | 1029 | + |
| Y11_RS00290 | Y11_00621 | cysT | sulfate/thiosulfate ABC transporter permease CysT | 0,81 | 1,17 | 1,83 | 0,35 | 1,01 | 0,66 | 9,0E-02 | 2,0E-02 | 2,1E-03 | 3,7E-01 | 2,7E-02 | 9,7E-02 | 0,0E+00 | 3,8E-04 | 2,4E-05 | 0,0E+00 | 5,6E-04 | 0,0E+00 | 53894 | 54730 | 837 | + |
| Y11_RS00300 | Y11_00641 | cysA | sulfate/thiosulfate ABC transporter ATP-binding protein CysA | 1,37 | -0,7 | 0,26 | -2,07 | -1,11 | 0,96 | 3,1E-06 | 5,1E-03 | 1,1E-01 | 8,9E-07 | 8,7E-06 | 6,2E-04 | 1,8E-07 | 0,0E+00 | 0,0E+00 | 2,2E-08 | 2,9E-07 | 0,0E+00 | 55595 | 56686 | 1092 | + |
| Y11_RS00305 | Y11_00651 | cysM | cysteine synthase CysM | -0,71 | -1,23 | -0,71 | -0,52 | 0 | 0,52 | 2,0E-05 | 1,6E-08 | 6,1E-05 | 5,7E-05 | 1,0E+00 | 2,4E-04 | 0,0E+00 | 8,8E-10 | 0,0E+00 | 0,0E+00 | 0,0E+00 | 0,0E+00 | 56906 | 57787 | 882 | + |
| Y11_RS00315 | Y11_00681 | | efflux RND transporter periplasmic adaptor subunit | 0 | -15 | 0 | -15 | 0 | 1,4 | 0,0E+00 | 0,0E+00 | 0,0E+00 | 0,0E+00 | 0,0E+00 | 5,7E-03 | 0,0E+00 | 0,0E+00 | 0,0E+00 | 0,0E+00 | 0,0E+00 | 3,4E-04 | 59840 | 61048 | 1209 | - |
| Y11_RS00325 | Y11_00711 | | ATP-binding protein | -0,04 | -2,26 | -0,96 | -2,22 | -0,92 | 1,3 | 7,3E-01 | 7,0E-07 | 8,5E-07 | 1,7E-06 | 2,0E-05 | 4,2E-05 | 0,0E+00 | 2,1E-08 | 0,0E+00 | 4,0E-08 | 0,0E+00 | 3,7E-06 | 61945 | 63312 | 1368 | + |
| Y11_RS00330 | Y11_00721 | crr | PTS glucose transporter subunit IIA | 0,39 | 0,69 | 0,1 | 0,3 | -0,29 | -0,59 | 1,4E-03 | 9,7E-07 | 5,5E-01 | 2,3E-03 | 1,1E-01 | 4,2E-03 | 0,0E+00 | 0,0E+00 | 0,0E+00 | 0,0E+00 | 0,0E+00 | 0,0E+00 | 63364 | 63873 | 510 | - |
| Y11_RS00335 | Y11_00731 | ptsI | phosphoenolpyruvate-protein phosphotransferase PtsI | 0,28 | 0,51 | -1,24 | 0,23 | -1,52 | -1,75 | 1,7E-03 | 1,4E-04 | 5,6E-05 | 2,3E-02 | 1,5E-05 | 6,7E-06 | 0,0E+00 | 0,0E+00 | 8,0E-07 | 0,0E+00 | 4,6E-07 | 8,2E-07 | 63922 | 65649 | 1728 | - |
| Y11_RS00340 | Y11_00741 | ptsH | phosphocarrier protein Hpr | 0,39 | 0,36 | -0,88 | -0,03 | -1,27 | -1,24 | 4,3E-03 | 1,7E-03 | 6,6E-02 | 7,9E-01 | 1,7E-02 | 1,8E-02 | 0,0E+00 | 0,0E+00 | 0,0E+00 | 0,0E+00 | 3,5E-04 | 1,0E-03 | 65697 | 65954 | 258 | - |
| Y11_RS00345 | Y11_00751 | cysK | cysteine synthase A | -0,19 | 0,05 | 0,66 | 0,24 | 0,85 | 0,61 | 5,7E-02 | 6,6E-01 | 3,0E-04 | 5,7E-02 | 4,5E-05 | 1,4E-03 | 0,0E+00 | 0,0E+00 | 0,0E+00 | 0,0E+00 | 0,0E+00 | 0,0E+00 | 66406 | 67374 | 969 | - |
| Y11_RS00350 | Y11_00761 | cysZ | sulfate transporter CysZ | -0,11 | 1,1 | 1,45 | 1,27 | 1,64 | 0,35 | 8,7E-01 | 1,3E-01 | 5,9E-02 | 8,5E-02 | 3,3E-02 | 6,1E-01 | 0,0E+00 | 2,5E-03 | 6,5E-04 | 1,3E-03 | 6,9E-04 | 0,0E+00 | 67564 | 68331 | 768 | - |
| Y11_RS00355 | Y11_00771 | zipA | cell division protein ZipA | 0,27 | 0,63 | 0,92 | 0,36 | 0,65 | 0,29 | 1,7E-03 | 1,0E-05 | 2,3E-06 | 7,0E-04 | 3,8E-05 | 8,8E-03 | 0,0E+00 | 0,0E+00 | 0,0E+00 | 0,0E+00 | 0,0E+00 | 0,0E+00 | 68551 | 69534 | 984 | + |
| Y11_RS00360 | Y11_00781 | ligA | NAD-dependent DNA ligase LigA | -0,08 | -0,46 | -0,81 | -0,38 | -0,72 | -0,34 | 5,7E-02 | 2,4E-06 | 2,5E-07 | 4,9E-06 | 3,6E-07 | 1,1E-04 | 0,0E+00 | 0,0E+00 | 0,0E+00 | 0,0E+00 | 0,0E+00 | 0,0E+00 | 69610 | 71628 | 2019 | + |
| Y11_RS00365 | Y11_00791 | | DUF3820 family protein | -0,63 | -1,97 | -2,33 | -1,34 | -1,69 | -0,36 | 1,9E-01 | 1,0E-03 | 3,8E-04 | 7,1E-04 | 1,9E-04 | 3,8E-02 | 0,0E+00 | 2,0E-05 | 4,7E-06 | 1,2E-05 | 4,8E-06 | 0,0E+00 | 71643 | 71858 | 216 | + |
| Y11_RS00390 | Y11_00831 | gltX | glutamate--tRNA ligase | 0,27 | 0,52 | 0,84 | 0,26 | 0,58 | 0,32 | 3,0E-05 | 1,7E-07 | 3,0E-07 | 9,2E-05 | 7,6E-06 | 4,8E-04 | 0,0E+00 | 0,0E+00 | 0,0E+00 | 0,0E+00 | 0,0E+00 | 0,0E+00 | 73347 | 74762 | 1416 | - |
| Y11_RS00415 | Y11_00841 | | NupC/NupG family nucleoside CNT transporter | 0,02 | -0,19 | -0,87 | -0,22 | -0,9 | -0,68 | 9,1E-01 | 4,6E-01 | 2,1E-02 | 2,9E-01 | 8,5E-03 | 5,3E-02 | 0,0E+00 | 0,0E+00 | 0,0E+00 | 0,0E+00 | 0,0E+00 | 0,0E+00 | 75633 | 76817 | 1185 | - |
| Y11_RS00430 | Y11_00871 | | aldo/keto reductase | -2,9 | 0,6 | 0,72 | 3,5 | 3,61 | 0,11 | 6,9E-11 | 2,8E-05 | 4,2E-06 | 1,6E-11 | 5,4E-12 | 1,2E-01 | 2,7E-11 | 0,0E+00 | 0,0E+00 | 6,2E-12 | 8,8E-12 | 0,0E+00 | 79086 | 80075 | 990 | - |
| Y11_RS00435 | Y11_00891 | | alpha-keto acid decarboxylase family protein | -0,19 | 2,35 | 2,2 | 2,54 | 2,39 | -0,14 | 4,1E-01 | 5,4E-06 | 6,3E-06 | 2,8E-09 | 3,6E-10 | 1,3E-01 | 0,0E+00 | 1,4E-07 | 1,1E-07 | 1,8E-10 | 9,7E-11 | 0,0E+00 | 80299 | 81960 | 1662 | + |
| Y11_RS00450 | Y11_00921 | glk | glucokinase | 0,38 | 4,86 | 4,31 | 4,43 | 3,95 | 0 | 9,0E-03 | 2,7E-11 | 9,1E-11 | 5,4E-11 | 1,0E-09 | 0,0E+00 | 0,0E+00 | 9,7E-12 | 2,3E-11 | 1,2E-11 | 1,7E-10 | 0,0E+00 | 83583 | 84554 | 972 | + |
| Y11_RS00470 | Y11_00961 | | sensor histidine kinase | -1,66 | 0 | 0 | 15 | 15 | 0 | 4,4E-06 | 0,0E+00 | 0,0E+00 | 0,0E+00 | 0,0E+00 | 0,0E+00 | 2,5E-07 | 0,0E+00 | 0,0E+00 | 0,0E+00 | 0,0E+00 | 0,0E+00 | 88398 | 90098 | 1701 | - |
| Y11_RS00475 | Y11_00971 | | DUF1479 domain-containing protein | 0,51 | 0,91 | 1,83 | 0,4 | 1,33 | 0,93 | 6,8E-05 | 5,0E-08 | 1,9E-08 | 2,9E-04 | 6,8E-07 | 3,2E-06 | 0,0E+00 | 0,0E+00 | 8,6E-10 | 0,0E+00 | 3,0E-08 | 0,0E+00 | 90269 | 91516 | 1248 | - |
| Y11_RS00485 | Y11_00991 | alaC | alanine transaminase | 0,12 | -0,39 | -0,3 | -0,51 | -0,42 | 0,09 | 1,4E-01 | 1,2E-03 | 1,5E-02 | 1,9E-05 | 7,4E-04 | 3,2E-01 | 0,0E+00 | 0,0E+00 | 0,0E+00 | 0,0E+00 | 0,0E+00 | 0,0E+00 | 92422 | 93657 | 1236 | + |
| Y11_RS00495 | Y11_01011 | | hypothetical protein | 0,29 | -0,97 | -0,61 | -1,26 | -0,88 | 0,38 | 5,3E-02 | 1,6E-03 | 8,2E-04 | 7,2E-04 | 6,3E-04 | 1,4E-01 | 0,0E+00 | 0,0E+00 | 0,0E+00 | 1,2E-05 | 0,0E+00 | 0,0E+00 | 94766 | 95197 | 432 | - |
| Y11_RS00510 | Y11_01051 | | ABC transporter ATP-binding protein | -0,42 | -0,24 | -0,31 | 0,18 | 0,12 | -0,07 | 6,7E-02 | 2,4E-01 | 1,6E-01 | 3,5E-01 | 5,7E-01 | 7,2E-01 | 0,0E+00 | 0,0E+00 | 0,0E+00 | 0,0E+00 | 0,0E+00 | 0,0E+00 | 97479 | 98528 | 1050 | - |
| Y11_RS00520 | Y11_01071 | | iron ABC transporter substrate-binding protein | -1,2 | 0,46 | -0,39 | 1,66 | 0,81 | -0,85 | 6,7E-07 | 7,6E-04 | 5,4E-04 | 1,6E-07 | 1,3E-05 | 9,2E-06 | 4,5E-08 | 0,0E+00 | 0,0E+00 | 5,1E-09 | 0,0E+00 | 0,0E+00 | 100198 | 101214 | 1017 | - |
| Y11_RS00525 | Y11_01081 | mltB | lytic murein transglycosylase B | 0 | -15 | -15 | -15 | -15 | -0,73 | 0,0E+00 | 0,0E+00 | 0,0E+00 | 0,0E+00 | 0,0E+00 | 8,3E-03 | 0,0E+00 | 0,0E+00 | 0,0E+00 | 0,0E+00 | 0,0E+00 | 0,0E+00 | 101526 | 102602 | 1077 | + |
| Y11_RS00535 | Y11_01101 | | amidohydrolase | -1,34 | -0,38 | -0,36 | 0,96 | 0,98 | 0,02 | 1,7E-08 | 3,9E-05 | 3,3E-04 | 3,6E-07 | 1,0E-06 | 7,9E-01 | 2,0E-09 | 0,0E+00 | 0,0E+00 | 0,0E+00 | 0,0E+00 | 0,0E+00 | 104482 | 106212 | 1731 | - |
| Y11_RS00540 | Y11_01111 | | TetR family transcriptional regulator | 1,17 | 0,74 | 0,78 | -0,43 | -0,39 | 0,04 | 1,8E-04 | 3,0E-03 | 2,8E-03 | 3,9E-04 | 2,4E-03 | 6,3E-01 | 7,5E-06 | 0,0E+00 | 0,0E+00 | 0,0E+00 | 0,0E+00 | 0,0E+00 | 106372 | 106962 | 591 | - |
| Y11_RS00550 | Y11_01131 | | ABC transporter permease | 0,14 | -0,7 | 0,14 | -0,84 | 0 | 0,84 | 4,7E-01 | 2,9E-02 | 7,1E-01 | 1,0E-02 | 1,0E+00 | 6,8E-02 | 0,0E+00 | 0,0E+00 | 0,0E+00 | 0,0E+00 | 0,0E+00 | 0,0E+00 | 108946 | 110091 | 1146 | + |
| Y11_RS00555 | Y11_01141 | | ABC transporter ATP-binding protein | -0,04 | -0,98 | -0,52 | -0,94 | -0,48 | 0,46 | 7,2E-01 | 5,8E-06 | 7,9E-04 | 2,2E-06 | 5,4E-04 | 4,0E-05 | 0,0E+00 | 0,0E+00 | 0,0E+00 | 0,0E+00 | 0,0E+00 | 0,0E+00 | 110085 | 110891 | 807 | + |
| Y11_RS00560 | Y11_01151 | | MCE family protein | -0,08 | -0,89 | -0,4 | -0,81 | -0,33 | 0,48 | 3,9E-01 | 1,5E-06 | 2,4E-02 | 1,1E-05 | 6,4E-02 | 1,0E-02 | 0,0E+00 | 0,0E+00 | 0,0E+00 | 0,0E+00 | 0,0E+00 | 0,0E+00 | 110892 | 111833 | 942 | + |

| | | | | | | | | | | | | | | | | | | | | | | | | |
|---|---|---|---|---|---|---|---|---|---|---|---|---|---|---|---|---|---|---|---|---|---|---|---|---|
| Y11_RS00565 | Y11_01161 | | ABC-type transport auxiliary lipoprotein family protein | 0,13 | -0,55 | -0,3 | -0,68 | -0,43 | 0,25 | 2,1E-01 | 2,1E-04 | 1,9E-01 | 6,0E-05 | 7,8E-02 | 2,8E-01 | 0,0E+00 | 0,0E+00 | 0,0E+00 | 0,0E+00 | 0,0E+00 | 0,0E+00 | 111830 | 112465 | 636 | + |
| Y11_RS00590 | Y11_01221 | | LacI family DNA-binding transcriptional regulator | -0,27 | 0,26 | 15 | 0 | 0 | 0 | 4,6E-01 | 3,1E-01 | 0,0E+00 | 0,0E+00 | 0,0E+00 | 0,0E+00 | 0,0E+00 | 0,0E+00 | 0,0E+00 | 0,0E+00 | 0,0E+00 | 0,0E+00 | 114698 | 115729 | 1032 | - |
| Y11_RS00605 | Y11_01251 | | DUF3861 domain-containing protein | -0,39 | 0,08 | 0,69 | 0,47 | 1,08 | 0,61 | 3,8E-03 | 4,7E-01 | 1,0E-03 | 4,5E-03 | 8,8E-05 | 4,2E-03 | 0,0E+00 | 0,0E+00 | 0,0E+00 | 0,0E+00 | 2,4E-06 | 0,0E+00 | 119069 | 119380 | 312 | - |
| Y11_RS00610 | Y11_01261 | | PadR family transcriptional regulator | 0,4 | 0,76 | 0,67 | 0,36 | 0,27 | -0,09 | 2,3E-02 | 2,4E-04 | 5,5E-02 | 5,3E-02 | 4,2E-01 | 7,7E-01 | 0,0E+00 | 0,0E+00 | 0,0E+00 | 0,0E+00 | 0,0E+00 | 0,0E+00 | 119380 | 120006 | 627 | - |
| Y11_RS00615 | Y11_01271 | | alkylphosphonate utilization protein | 0,56 | 0,79 | 1,5 | 0,23 | 0,94 | 0,72 | 1,7E-01 | 1,1E-01 | 1,5E-02 | 4,5E-01 | 3,1E-02 | 1,4E-01 | 0,0E+00 | 0,0E+00 | 1,7E-04 | 0,0E+00 | 0,0E+00 | 0,0E+00 | 120271 | 120609 | 339 | - |
| Y11_RS00635 | Y11_01301 | ccmA | cytochrome c biogenesis heme-transporting ATPase CcmA | -0,7 | 0,18 | 0 | 0,88 | 0,19 | -0,64 | 1,1E-02 | 3,8E-01 | 0,0E+00 | 1,6E-02 | 4,3E-01 | 1,7E-02 | 0,0E+00 | 0,0E+00 | 0,0E+00 | 0,0E+00 | 0,0E+00 | 0,0E+00 | 123077 | 123703 | 627 | + |
| Y11_RS00655 | Y11_01351 | ccmE | cytochrome c maturation protein CcmE | -0,51 | -0,83 | -0,61 | -0,31 | -0,09 | 0,22 | 5,4E-02 | 6,1E-03 | 4,7E-02 | 7,1E-03 | 5,7E-01 | 1,9E-01 | 0,0E+00 | 0,0E+00 | 0,0E+00 | 0,0E+00 | 0,0E+00 | 0,0E+00 | 125512 | 126006 | 495 | + |
| Y11_RS00660 | Y11_01361 | | heme lyase CcmF/NrfE family subunit | -0,02 | 0,54 | 0,45 | 0,56 | 0,47 | -0,09 | 8,8E-01 | 4,8E-02 | 4,3E-03 | 3,8E-02 | 1,8E-03 | 6,7E-01 | 0,0E+00 | 0,0E+00 | 0,0E+00 | 0,0E+00 | 0,0E+00 | 0,0E+00 | 126003 | 127970 | 1968 | + |
| Y11_RS00665 | Y11_01371 | | DsbE family thiol:disulfide interchange protein | -0,13 | 0,6 | 0,74 | 0,72 | 0,87 | 0,14 | 5,1E-01 | 1,1E-02 | 6,2E-04 | 7,9E-03 | 8,2E-04 | 4,1E-01 | 0,0E+00 | 0,0E+00 | 0,0E+00 | 0,0E+00 | 0,0E+00 | 0,0E+00 | 127970 | 128536 | 567 | + |
| Y11_RS00670 | Y11_01381 | | cytochrome c-type biogenesis protein CcmH | -0,5 | -1,18 | -1,38 | -0,67 | -0,88 | -0,21 | 2,4E-02 | 2,3E-05 | 2,3E-06 | 8,1E-03 | 1,2E-03 | 1,6E-01 | 0,0E+00 | 5,4E-07 | 4,7E-08 | 0,0E+00 | 0,0E+00 | 0,0E+00 | 128533 | 129039 | 507 | + |
| Y11_RS00675 | Y11_01391 | ccmI | c-type cytochrome biogenesis protein CcmI | -0,35 | -0,31 | -0,26 | 0,04 | 0,08 | 0,04 | 1,2E-02 | 1,8E-02 | 1,1E-01 | 7,1E-01 | 5,9E-01 | 7,8E-01 | 0,0E+00 | 0,0E+00 | 0,0E+00 | 0,0E+00 | 0,0E+00 | 0,0E+00 | 129039 | 130259 | 1221 | + |
| Y11_RS00680 | Y11_01401 | mlaA | phospholipid-binding lipoprotein MlaA | -0,52 | -0,88 | -0,5 | -0,36 | 0,02 | 0,38 | 1,6E-03 | 1,3E-05 | 1,1E-03 | 8,3E-03 | 8,6E-01 | 3,1E-03 | 0,0E+00 | 0,0E+00 | 0,0E+00 | 0,0E+00 | 0,0E+00 | 0,0E+00 | 130296 | 131060 | 765 | + |
| Y11_RS00685 | Y11_01411 | fadL | long-chain fatty acid transporter FadL | 2 | -1,43 | 0,55 | -3,43 | -1,45 | 1,99 | 8,5E-13 | 2,9E-08 | 2,0E-04 | 2,8E-11 | 1,5E-07 | 7,2E-08 | 1,8E-12 | 1,5E-09 | 0,0E+00 | 8,5E-12 | 8,8E-09 | 2,8E-08 | 131153 | 132424 | 1272 | - |
| Y11_RS00690 | Y11_01421 | | YfcZ/YiiS family protein | 1,14 | 0,09 | 0,52 | -1,05 | -0,62 | 0,43 | 7,2E-09 | 3,0E-01 | 4,7E-02 | 1,2E-06 | 2,3E-02 | 1,0E-01 | 1,1E-09 | 0,0E+00 | 0,0E+00 | 3,0E-08 | 0,0E+00 | 0,0E+00 | 132855 | 133166 | 312 | + |
| Y11_RS00695 | Y11_01431 | fadI | acetyl-CoA C-acyltransferase FadI | 0,15 | -0,32 | 1,44 | -0,47 | 1,29 | 1,76 | 1,6E-01 | 3,6E-03 | 1,2E-07 | 2,7E-03 | 3,0E-06 | 1,0E-07 | 0,0E+00 | 0,0E+00 | 3,7E-09 | 0,0E+00 | 1,1E-07 | 3,7E-08 | 133475 | 134785 | 1311 | + |
| Y11_RS00700 | Y11_01441 | fadJ | fatty acid oxidation complex subunit alpha FadJ | 0,11 | -0,55 | 0,47 | -0,66 | 0,36 | 1,01 | 1,8E-01 | 1,3E-04 | 3,2E-04 | 4,6E-05 | 2,1E-03 | 2,4E-06 | 0,0E+00 | 0,0E+00 | 0,0E+00 | 0,0E+00 | 0,0E+00 | 3,6E-07 | 134785 | 137022 | 2238 | + |
| Y11_RS00705 | Y11_01451 | sixA | phosphohistidine phosphatase SixA | 0,59 | 0,9 | 0,37 | 0,32 | -0,22 | -0,53 | 2,2E-04 | 1,4E-06 | 2,8E-03 | 2,3E-02 | 1,2E-01 | 1,2E-03 | 0,0E+00 | 0,0E+00 | 0,0E+00 | 0,0E+00 | 0,0E+00 | 0,0E+00 | 137405 | 137896 | 492 | + |
| Y11_RS00710 | Y11_01461 | smrB | endonuclease SmrB | 0 | 0 | 0,27 | 0 | 0,33 | -15 | 0,0E+00 | 0,0E+00 | 2,8E-01 | 0,0E+00 | 1,6E-01 | 0,0E+00 | 0,0E+00 | 0,0E+00 | 0,0E+00 | 0,0E+00 | 0,0E+00 | 0,0E+00 | 138198 | 138728 | 531 | - |
| Y11_RS00715 | Y11_01471 | prmB | 50S ribosomal protein L3 N(5)-glutamine methyltransferase | 0,22 | 0,33 | 0,8 | 0,12 | 0,58 | 0,47 | 1,8E-02 | 8,8E-05 | 9,1E-07 | 1,1E-01 | 5,2E-05 | 1,1E-05 | 0,0E+00 | 0,0E+00 | 0,0E+00 | 0,0E+00 | 0,0E+00 | 0,0E+00 | 138935 | 139867 | 933 | + |
| Y11_RS00720 | Y11_01481 | aroC | chorismate synthase | 0,43 | 0,28 | 0,63 | -0,14 | 0,2 | 0,35 | 6,6E-04 | 6,0E-03 | 2,8E-03 | 1,0E-01 | 2,1E-01 | 4,6E-02 | 0,0E+00 | 0,0E+00 | 0,0E+00 | 0,0E+00 | 0,0E+00 | 0,0E+00 | 139944 | 141029 | 1086 | + |
| Y11_RS00725 | Y11_01491 | mepA | penicillin-insensitive murein endopeptidase | 0,37 | -0,28 | 0,13 | -0,65 | -0,24 | 0,41 | 3,4E-02 | 7,8E-02 | 4,5E-01 | 1,1E-03 | 1,6E-01 | 2,6E-02 | 0,0E+00 | 0,0E+00 | 0,0E+00 | 0,0E+00 | 0,0E+00 | 0,0E+00 | 141035 | 141862 | 828 | + |
| Y11_RS00740 | Y11_01521 | | YfcL family protein | 0 | -0,03 | -1,93 | -15 | -15 | -1,9 | 0,0E+00 | 9,1E-01 | 7,3E-05 | 0,0E+00 | 0,0E+00 | 5,5E-04 | 0,0E+00 | 0,0E+00 | 1,0E-06 | 0,0E+00 | 0,0E+00 | 3,9E-05 | 143585 | 143866 | 282 | + |
| Y11_RS00745 | Y11_01531 | mnmC | bifunctional tRNA (5-methylaminomethyl-2-thiouridine)(34)-methyltransferase MnmD/FAD-dependent 5-carboxymethylaminomethyl-2-thiouridine(34) oxidoreductase MnmC | 0,24 | -1 | -1,04 | -1,24 | -1,28 | -0,04 | 1,5E-01 | 9,9E-07 | 5,1E-05 | 1,5E-05 | 7,6E-05 | 7,4E-01 | 0,0E+00 | 2,9E-08 | 7,4E-07 | 3,0E-07 | 2,1E-06 | 0,0E+00 | 144025 | 146088 | 2064 | - |
| Y11_RS00750 | Y11_01541 | fabB | beta-ketoacyl-ACP synthase I | -0,81 | 0,06 | 0,22 | 0,87 | 1,03 | 0,16 | 5,2E-04 | 6,1E-01 | 5,8E-02 | 5,1E-04 | 1,1E-04 | 2,0E-01 | 0,0E+00 | 0,0E+00 | 0,0E+00 | 0,0E+00 | 2,8E-06 | 0,0E+00 | 146260 | 147483 | 1224 | + |
| Y11_RS00760 | Y11_01571 | | aspartate/glutamate racemase family protein | 0,24 | -0,28 | -0,33 | -0,52 | -0,56 | -0,04 | 1,1E-01 | 7,8E-03 | 5,1E-03 | 1,8E-03 | 1,3E-03 | 4,1E-01 | 0,0E+00 | 0,0E+00 | 0,0E+00 | 0,0E+00 | 0,0E+00 | 0,0E+00 | 148949 | 149644 | 696 | - |
| Y11_RS00775 | Y11_01601 | | glycoside hydrolase family 1 protein | -0,43 | -0,97 | -1,48 | -0,55 | -1,06 | -0,51 | 6,1E-02 | 6,3E-04 | 3,9E-04 | 2,7E-02 | 4,6E-03 | 8,6E-02 | 0,0E+00 | 0,0E+00 | 4,9E-06 | 0,0E+00 | 1,0E-04 | 0,0E+00 | 151945 | 153327 | 1383 | - |
| Y11_RS00790 | Y11_01631 | | LacI family transcriptional regulator | 0,03 | 0,18 | 1,19 | 0,14 | 1,16 | 1,01 | 8,2E-01 | 2,3E-01 | 2,3E-02 | 1,3E-01 | 2,2E-02 | 3,8E-02 | 0,0E+00 | 0,0E+00 | 2,6E-04 | 0,0E+00 | 4,7E-04 | 2,0E-03 | 155354 | 156310 | 957 | + |
| Y11_RS00795 | Y11_01641 | agp | bifunctional glucose-1-phosphatase/inositol phosphatase | -0,24 | 2,95 | 4,48 | 3,18 | 4,72 | 1,53 | 2,3E-03 | 1,3E-10 | 1,5E-11 | 9,9E-11 | 1,3E-11 | 2,7E-07 | 0,0E+00 | 2,5E-11 | 8,0E-12 | 1,9E-11 | 1,4E-11 | 6,9E-08 | 156584 | 157885 | 1302 | + |
| Y11_RS00820 | Y11_01701 | | DsbA family protein | 0,03 | -3,35 | -2,48 | -3,37 | -2,52 | 0,87 | 8,6E-01 | 7,2E-09 | 1,2E-05 | 6,6E-10 | 6,6E-06 | 6,0E-03 | 0,0E+00 | 4,9E-10 | 1,9E-07 | 6,4E-11 | 2,3E-07 | 0,0E+00 | 163002 | 163727 | 726 | + |
| Y11_RS00845 | Y11_01751 | | helix-turn-helix transcriptional regulator | -0,22 | 2,35 | 2,47 | 2,57 | 2,69 | 0,12 | 1,2E-02 | 2,2E-10 | 2,2E-05 | 6,0E-10 | 1,3E-05 | 6,8E-01 | 0,0E+00 | 3,8E-11 | 3,5E-07 | 5,9E-11 | 4,2E-07 | 0,0E+00 | 168027 | 168812 | 786 | + |
| Y11_RS00850 | Y11_01761 | pdxB | 4-phosphoerythronate dehydrogenase PdxB | -0,46 | -0,39 | -0,38 | 0,07 | 0,08 | 0,01 | 8,3E-05 | 7,6E-05 | 1,0E-03 | 3,5E-01 | 4,0E-01 | 9,2E-01 | 0,0E+00 | 0,0E+00 | 0,0E+00 | 0,0E+00 | 0,0E+00 | 0,0E+00 | 169031 | 170158 | 1128 | + |
| Y11_RS00855 | Y11_01771 | | aspartate-semialdehyde dehydrogenase | -0,1 | 0,23 | 0,35 | 0,33 | 0,44 | 0,11 | 6,1E-02 | 2,8E-04 | 1,6E-02 | 2,6E-05 | 4,5E-03 | 3,3E-01 | 0,0E+00 | 0,0E+00 | 0,0E+00 | 0,0E+00 | 0,0E+00 | 0,0E+00 | 170430 | 171440 | 1011 | + |
| Y11_RS00860 | Y11_01781 | truA | tRNA pseudouridine(38-40) synthase TruA | -0,3 | -0,23 | -0,47 | 0,07 | -0,18 | -0,24 | 5,5E-03 | 3,2E-02 | 1,2E-03 | 3,2E-01 | 4,4E-02 | 2,0E-02 | 0,0E+00 | 0,0E+00 | 0,0E+00 | 0,0E+00 | 0,0E+00 | 0,0E+00 | 171440 | 172282 | 843 | + |
| Y11_RS00865 | Y11_01791 | | DedA family protein | -0,05 | -2,51 | -1,64 | -2,45 | -1,59 | 0,87 | 7,9E-01 | 7,2E-07 | 5,3E-04 | 7,1E-07 | 6,2E-04 | 1,6E-02 | 0,0E+00 | 2,2E-08 | 6,5E-06 | 1,8E-08 | 1,5E-05 | 0,0E+00 | 172414 | 173076 | 663 | + |
| Y11_RS00870 | Y11_01801 | accD | acetyl-CoA carboxylase, carboxyltransferase subunit beta | 0,19 | -0,21 | -0,72 | -0,4 | -0,91 | -0,51 | 1,7E-02 | 2,3E-03 | 7,8E-06 | 2,5E-04 | 4,2E-06 | 1,2E-04 | 0,0E+00 | 0,0E+00 | 0,0E+00 | 0,0E+00 | 0,0E+00 | 0,0E+00 | 173307 | 174218 | 912 | + |
| Y11_RS00875 | Y11_01821 | folC | bifunctional tetrahydrofolate synthase/dihydrofolate synthase | -0,4 | -1,45 | -1,36 | -1,05 | -0,97 | 0,09 | 1,5E-05 | 1,2E-09 | 4,8E-08 | 9,4E-09 | 5,6E-07 | 2,4E-01 | 0,0E+00 | 1,3E-10 | 1,8E-09 | 5,0E-10 | 0,0E+00 | 0,0E+00 | 174454 | 175722 | 1269 | + |
| Y11_RS00880 | Y11_01831 | dedD | cell division protein DedD | -0,19 | -0,95 | -0,88 | -0,75 | -0,69 | 0,06 | 8,6E-02 | 1,9E-05 | 8,7E-05 | 2,9E-05 | 2,0E-04 | 6,0E-01 | 0,0E+00 | 0,0E+00 | 0,0E+00 | 0,0E+00 | 0,0E+00 | 0,0E+00 | 175762 | 176481 | 720 | + |
| Y11_RS00885 | Y11_01841 | cvpA | colicin V production protein | 0,01 | -1,25 | -1,58 | -1,26 | -1,58 | -0,32 | 9,8E-01 | 2,0E-03 | 4,3E-04 | 5,8E-04 | 1,1E-04 | 6,5E-02 | 0,0E+00 | 3,9E-05 | 5,3E-06 | 9,9E-06 | 2,8E-06 | 0,0E+00 | 176787 | 177296 | 510 | + |
| Y11_RS00890 | Y11_01851 | purF | amidophosphoribosyltransferase | 0,23 | 0,54 | 1,12 | 0,31 | 0,89 | 0,58 | 1,2E-02 | 2,9E-04 | 3,4E-07 | 1,2E-03 | 1,5E-08 | 2,4E-05 | 0,0E+00 | 0,0E+00 | 9,2E-09 | 0,0E+00 | 0,0E+00 | 0,0E+00 | 177309 | 178826 | 1518 | + |
| Y11_RS00895 | Y11_01861 | | UbiX family flavin prenyltransferase | -0,11 | 0,96 | 0,97 | 1,07 | 1,08 | 0,01 | 1,7E-01 | 3,4E-06 | 2,3E-06 | 2,4E-06 | 1,7E-06 | 9,5E-01 | 0,0E+00 | 0,0E+00 | 0,0E+00 | 5,5E-08 | 6,6E-08 | 0,0E+00 | 179156 | 179728 | 573 | + |
| Y11_RS00900 | Y11_01871 | hisJ | histidine ABC transporter substrate-binding protein HisJ | -0,39 | 1,02 | 3,72 | 1,41 | 4,11 | 2,7 | 5,1E-04 | 4,9E-06 | 3,0E-11 | 1,4E-06 | 9,0E-11 | 1,1E-08 | 0,0E+00 | 1,2E-07 | 1,3E-11 | 3,4E-08 | 4,4E-11 | 7,9E-09 | 180037 | 180819 | 783 | + |
| Y11_RS00915 | Y11_01901 | hisP | histidine ABC transporter ATP-binding protein HisP | -0,25 | -1,8 | -0,5 | -1,55 | -0,25 | 1,3 | 2,5E-01 | 2,1E-04 | 6,4E-02 | 3,8E-04 | 2,8E-01 | 1,9E-03 | 0,0E+00 | 4,5E-06 | 0,0E+00 | 6,7E-06 | 0,0E+00 | 1,2E-04 | 182359 | 183156 | 798 | + |
| Y11_RS00930 | Y11_01931 | | TIGR01777 family oxidoreductase | -0,03 | 0,94 | 2,14 | 0,97 | 2,17 | 1,2 | 5,9E-01 | 3,8E-07 | 1,0E-07 | 5,9E-08 | 6,3E-08 | 7,0E-06 | 0,0E+00 | 0,0E+00 | 3,3E-09 | 0,0E+00 | 4,2E-09 | 8,4E-07 | 189749 | 190657 | 909 | - |
| Y11_RS00935 | Y11_01941 | | EAL domain-containing protein | 1,2 | -0,16 | 0,09 | -1,37 | -1,11 | 0,25 | 1,8E-04 | 3,9E-01 | 7,5E-01 | 7,9E-05 | 4,5E-03 | 3,9E-01 | 7,4E-06 | 0,0E+00 | 0,0E+00 | 1,5E-06 | 9,9E-05 | 0,0E+00 | 191166 | 191858 | 693 | + |
| Y11_RS00940 | Y11_01951 | yfcE | phosphodiesterase | 0,38 | 1,03 | 1 | 0,66 | 0,63 | -0,03 | 2,1E-02 | 4,4E-07 | 1,5E-05 | 4,9E-04 | 2,3E-03 | 7,4E-01 | 0,0E+00 | 1,4E-08 | 2,4E-07 | 0,0E+00 | 0,0E+00 | 0,0E+00 | 192095 | 192649 | 555 | + |
| Y11_RS00945 | Y11_01961 | yfcD | NUDIX hydrolase YfcD | 0,26 | 0,87 | 0,97 | 0,62 | 0,71 | 0,09 | 3,1E-02 | 1,9E-06 | 2,8E-06 | 2,6E-04 | 1,8E-04 | 3,1E-01 | 0,0E+00 | 0,0E+00 | 0,0E+00 | 0,0E+00 | 0,0E+00 | 0,0E+00 | 192721 | 193269 | 549 | + |
| Y11_RS00955 | Y11_01991 | | PTS ascorbate transporter subunit IIC | -0,43 | -0,2 | 0,02 | 0,22 | 0,45 | 0,23 | 1,1E-01 | 3,7E-01 | 9,4E-01 | 3,6E-01 | 1,9E-01 | 4,7E-01 | 0,0E+00 | 0,0E+00 | 0,0E+00 | 0,0E+00 | 0,0E+00 | 0,0E+00 | 194852 | 196108 | 1257 | - |
| Y11_RS00960 | Y11_02001 | | PTS sugar transporter subunit IIB | -0,41 | -1,22 | -1,29 | -0,81 | -0,88 | -0,06 | 3,9E-04 | 5,0E-06 | 2,4E-03 | 1,1E-04 | 1,8E-02 | 8,4E-01 | 0,0E+00 | 1,3E-07 | 2,8E-05 | 0,0E+00 | 0,0E+00 | 0,0E+00 | 196121 | 196408 | 288 | - |
| Y11_RS00965 | Y11_02011 | | PTS sugar transporter subunit IIA | -0,39 | -1,12 | -1,39 | -0,74 | -1 | -0,27 | 2,3E-01 | 1,3E-04 | 4,1E-05 | 5,3E-02 | 1,6E-02 | 2,5E-01 | 0,0E+00 | 2,7E-06 | 6,0E-07 | 0,0E+00 | 3,3E-04 | 0,0E+00 | 196451 | 196894 | 444 | - |
| Y11_RS00970 | Y11_02021 | | LacI family transcriptional regulator | -0,11 | -0,06 | -0,79 | 0,05 | -0,68 | -0,73 | 5,9E-01 | 7,0E-01 | 1,0E-02 | 8,2E-01 | 5,2E-02 | 2,4E-02 | 0,0E+00 | 0,0E+00 | 0,0E+00 | 0,0E+00 | 0,0E+00 | 0,0E+00 | 197119 | 198144 | 1026 | + |
| Y11_RS00975 | Y11_02031 | pta | phosphate acetyltransferase | 0,08 | -0,16 | -0,81 | -0,24 | -0,88 | -0,65 | 7,9E-02 | 2,9E-04 | 3,4E-07 | 2,9E-08 | 2,6E-04 | 1,8E-07 | 0,0E+00 | 0,0E+00 | 0,0E+00 | 0,0E+00 | 0,0E+00 | 0,0E+00 | 198725 | 200863 | 2139 | - |
| Y11_RS00980 | Y11_02051 | ackA | acetate kinase | 0 | -0,21 | -0,74 | -0,21 | -0,74 | -0,53 | 1,0E+00 | 6,2E-02 | 1,2E-05 | 2,3E-02 | 3,0E-07 | 7,7E-05 | 0,0E+00 | 0,0E+00 | 0,0E+00 | 0,0E+00 | 0,0E+00 | 0,0E+00 | 201159 | 202361 | 1203 | - |
| Y11_RS00990 | Y11_02071 | | YfbU family protein | -0,09 | -0,91 | -0,79 | -0,82 | -0,7 | 0,12 | 2,0E-01 | 4,4E-06 | 2,5E-05 | 1,6E-05 | 9,1E-05 | 3,1E-01 | 0,0E+00 | 0,0E+00 | 0,0E+00 | 0,0E+00 | 0,0E+00 | 0,0E+00 | 203418 | 203912 | 495 | + |
| Y11_RS00995 | Y11_02081 | | sugar phosphatase | 0,33 | -2,33 | -3,45 | -2,66 | -3,78 | -1,12 | 3,3E-02 | 1,1E-07 | 1,2E-07 | 9,2E-08 | 8,9E-08 | 7,2E-04 | 0,0E+00 | 4,3E-09 | 3,7E-09 | 3,1E-09 | 5,6E-09 | 5,0E-05 | 203959 | 204615 | 657 | + |
| Y11_RS01000 | Y11_02091 | | hypothetical protein | -0,35 | 0,45 | 15 | 0,89 | 15 | 0 | 2,7E-01 | 6,0E-02 | 0,0E+00 | 7,4E-03 | 0,0E+00 | 0,0E+00 | 0,0E+00 | 0,0E+00 | 0,0E+00 | 0,0E+00 | 0,0E+00 | 0,0E+00 | 204806 | 205603 | 798 | - |
| Y11_RS01005 | Y11_02111 | | SLC13 family permease | -0,63 | 0,4 | 1,37 | 1,02 | 2 | 0,97 | 2,7E-04 | 3,6E-03 | 7,7E-05 | 2,0E-06 | 3,7E-06 | 5,6E-04 | 0,0E+00 | 0,0E+00 | 1,1E-06 | 4,6E-08 | 1,3E-07 | 0,0E+00 | 205966 | 207801 | 1836 | + |
| Y11_RS01010 | Y11_02121 | yfbR | 5'-deoxynucleotidase | 0,21 | -0,44 | -0,5 | -0,65 | -0,71 | -0,06 | 4,3E-02 | 9,5E-04 | 8,0E-04 | 3,9E-05 | 4,9E-05 | 5,0E-01 | 0,0E+00 | 0,0E+00 | 0,0E+00 | 0,0E+00 | 0,0E+00 | 0,0E+00 | 208033 | 208626 | 594 | - |
| Y11_RS01015 | Y11_02131 | | pyridoxal phosphate-dependent aminotransferase | 0,36 | -0,33 | -0,15 | -0,69 | -0,51 | 0,17 | 1,4E-04 | 3,1E-04 | 1,4E-01 | 1,8E-07 | 3,7E-04 | 8,3E-02 | 0,0E+00 | 0,0E+00 | 0,0E+00 | 0,0E+00 | 0,0E+00 | 0,0E+00 | 209368 | 210582 | 1215 | - |
| Y11_RS01020 | Y11_02141 | rovM | virulence transcriptional regulator RovM | 1,02 | -1,57 | -2,01 | -2,59 | -3,03 | -0,44 | 7,3E-07 | 1,9E-08 | 2,0E-08 | 2,6E-09 | 2,9E-09 | 3,2E-03 | 4,9E-08 | 1,0E-09 | 8,9E-10 | 1,8E-10 | 3,7E-10 | 0,0E+00 | 211446 | 212378 | 933 | + |
| Y11_RS01025 | Y11_02151 | nuoA | NADH-quinone oxidoreductase subunit NuoA | 0,12 | 0,69 | 0,7 | 0,57 | 0,58 | 0,01 | 2,4E-01 | 2,5E-05 | 1,0E-04 | 2,8E-04 | 7,2E-04 | 9,5E-01 | 0,0E+00 | 0,0E+00 | 0,0E+00 | 0,0E+00 | 0,0E+00 | 0,0E+00 | 213055 | 213546 | 492 | + |
| Y11_RS01030 | Y11_02161 | | NADH-quinone oxidoreductase subunit B | 0,1 | 0,54 | 0,3 | 0,44 | 0,2 | -0,24 | 1,5E-01 | 3,4E-06 | 1,4E-02 | 2,4E-04 | 1,1E-01 | 4,1E-02 | 0,0E+00 | 0,0E+00 | 0,0E+00 | 0,0E+00 | 0,0E+00 | 0,0E+00 | 213668 | 214345 | 678 | + |
| Y11_RS01035 | Y11_02171 | nuoC | NADH-quinone oxidoreductase subunit C/D | 0,16 | 0,73 | 0,76 | 0,57 | 0,6 | 0,02 | 1,2E-02 | 4,0E-07 | 3,7E-06 | 8,4E-06 | 4,0E-05 | 7,5E-01 | 0,0E+00 | 0,0E+00 | 0,0E+00 | 0,0E+00 | 0,0E+00 | 0,0E+00 | 214485 | 216281 | 1797 | + |
| Y11_RS01040 | Y11_02181 | nuoE | NADH-quinone oxidoreductase subunit NuoE | 0,35 | 0,59 | 0,35 | 0,24 | 0,01 | -0,24 | 3,0E-03 | 2,3E-04 | 1,9E-03 | 5,5E-02 | 9,5E-01 | 5,3E-02 | 0,0E+00 | 0,0E+00 | 0,0E+00 | 0,0E+00 | 0,0E+00 | 0,0E+00 | 216284 | 216847 | 564 | + |
| Y11_RS01045 | Y11_02191 | nuoF | NADH-quinone oxidoreductase subunit NuoF | 0,24 | 0,59 | 0,46 | 0,35 | 0,22 | -0,13 | 1,1E-03 | 4,8E-07 | 1,4E-06 | 6,7E-05 | 9,1E-04 | 4,4E-03 | 0,0E+00 | 0,0E+00 | 0,0E+00 | 0,0E+00 | 0,0E+00 | 0,0E+00 | 216844 | 218211 | 1368 | + |
| Y11_RS01050 | Y11_02201 | nuoG | NADH-quinone oxidoreductase subunit NuoG | 0,22 | 0,11 | -0,02 | -0,1 | -0,24 | -0,14 | 2,2E-06 | 6,4E-04 | 8,1E-01 | 4,4E-03 | 2,2E-02 | 1,5E-01 | 0,0E+00 | 0,0E+00 | 0,0E+00 | 0,0E+00 | 0,0E+00 | 0,0E+00 | 218439 | 221183 | 2745 | + |
| Y11_RS01055 | Y11_02211 | nuoH | NADH-quinone oxidoreductase subunit NuoH | 0 | 0,36 | 0,36 | 0,36 | 0,36 | 0 | 9,9E-01 | 3,5E-02 | 5,1E-02 | 1,3E-01 | 1,4E-01 | 1,0E+00 | 0,0E+00 | 0,0E+00 | 0,0E+00 | 0,0E+00 | 0,0E+00 | 0,0E+00 | 221180 | 222157 | 978 | + |
| Y11_RS01060 | Y11_02221 | nuoI | NADH-quinone oxidoreductase subunit NuoI | 0,1 | 0,69 | 0,47 | 0,6 | 0,38 | -0,22 | 1,4E-01 | 1,0E-06 | 2,1E-02 | 3,8E-06 | 5,3E-02 | 2,2E-01 | 0,0E+00 | 0,0E+00 | 0,0E+00 | 0,0E+00 | 0,0E+00 | 0,0E+00 | 222172 | 222714 | 543 | + |
| Y11_RS01070 | Y11_02241 | nuoK | NADH-quinone oxidoreductase subunit NuoK | -0,37 | 0,55 | 0,49 | 0,92 | 0,86 | -0,06 | 7,0E-02 | 2,4E-02 | 2,2E-02 | 4,3E-04 | 1,3E-04 | 6,9E-01 | 0,0E+00 | 0,0E+00 | 0,0E+00 | 0,0E+00 | 0,0E+00 | 0,0E+00 | 223269 | 223571 | 303 | + |

| | | | | | | | | | | | | | | | | | | | | | | | | | |
|---|---|---|---|---|---|---|---|---|---|---|---|---|---|---|---|---|---|---|---|---|---|---|---|---|---|
| Y11_RS01075 | Y11_02251 | nuoL | NADH-quinone oxidoreductase subunit L | 0,27 | 0,77 | 1,05 | 0,5 | 0,78 | 0,28 | 2,2E-01 | 2,1E-03 | 3,0E-03 | 6,6E-02 | 3,1E-02 | 3,5E-01 | 0,0E+00 | 0,0E+00 | 3,5E-05 | 0,0E+00 | 0,0E+00 | 0,0E+00 | 223568 | 225412 | 1845 | + |
| Y11_RS01080 | Y11_02261 | nuoM | NADH-quinone oxidoreductase subunit M | 0,03 | 0,56 | 1,96 | 0,53 | 1,84 | 1,39 | 8,5E-01 | 1,3E-01 | 7,6E-03 | 1,7E-01 | 1,2E-02 | 6,1E-02 | 0,0E+00 | 0,0E+00 | 8,7E-05 | 0,0E+00 | 2,6E-04 | 3,2E-03 | 225431 | 226969 | 1539 | + |
| Y11_RS01085 | Y11_02271 | nuoN | NADH-quinone oxidoreductase subunit NuoN | -0,06 | 0,56 | 1,41 | 0,62 | 1,48 | 0,85 | 7,3E-01 | 1,3E-01 | 2,8E-02 | 6,6E-02 | 1,9E-02 | 1,8E-01 | 0,0E+00 | 0,0E+00 | 3,2E-04 | 0,0E+00 | 4,1E-04 | 0,0E+00 | 226976 | 228433 | 1458 | + |
| Y11_RS01110 | Y11_02331 | idnO | gluconate 5-dehydrogenase | -0,46 | -0,67 | 0,1 | -0,2 | 0,55 | 0,77 | 2,0E-02 | 7,2E-02 | 7,4E-01 | 5,4E-01 | 7,0E-02 | 8,2E-02 | 0,0E+00 | 0,0E+00 | 0,0E+00 | 0,0E+00 | 0,0E+00 | 0,0E+00 | 232141 | 232905 | 765 | - |
| Y11_RS01115 | Y11_02351 | | substrate-binding domain-containing protein | 0,13 | 1,01 | 0,93 | 0,88 | 0,8 | -0,08 | 4,4E-01 | 2,0E-04 | 8,9E-04 | 1,9E-04 | 1,2E-03 | 6,3E-01 | 0,0E+00 | 4,2E-06 | 0,0E+00 | 0,0E+00 | 0,0E+00 | 0,0E+00 | 233408 | 234400 | 993 | + |
| Y11_RS01140 | Y11_02411 | elaB | stress response protein ElaB | 0,58 | 4,97 | 4,77 | 4,39 | 4,19 | -0,2 | 1,4E-02 | 5,1E-08 | 9,9E-07 | 1,6E-07 | 2,9E-06 | 6,3E-01 | 0,0E+00 | 2,3E-09 | 2,3E-08 | 5,1E-09 | 1,1E-07 | 0,0E+00 | 239846 | 240154 | 309 | + |
| Y11_RS01145 | Y11_02421 | menF | isochorismate synthase MenF | -0,1 | -1,56 | -2,08 | -1,46 | -1,97 | -0,52 | 6,3E-01 | 5,7E-05 | 1,8E-03 | 4,3E-06 | 1,7E-03 | 2,6E-01 | 0,0E+00 | 1,3E-06 | 2,1E-05 | 9,4E-08 | 3,9E-05 | 0,0E+00 | 240388 | 241752 | 1365 | + |
| Y11_RS01150 | Y11_02441 | menD | 2-succinyl-5-enolpyruvyl-6-hydroxy-3-cyclohexene-1-carboxylic-acid synthase | 0,31 | 0,09 | -0,35 | -0,22 | -0,66 | -0,44 | 4,5E-03 | 2,8E-01 | 9,6E-02 | 1,4E-02 | 6,9E-03 | 4,3E-02 | 0,0E+00 | 0,0E+00 | 0,0E+00 | 0,0E+00 | 0,0E+00 | 0,0E+00 | 241981 | 243654 | 1674 | + |
| Y11_RS01160 | Y11_02461 | menB | 1,4-dihydroxy-2-naphthoyl-CoA synthase | 0,11 | 0,07 | -0,65 | -0,05 | -0,76 | -0,72 | 5,0E-02 | 1,8E-01 | 6,1E-05 | 3,8E-01 | 2,6E-05 | 3,5E-05 | 0,0E+00 | 0,0E+00 | 0,0E+00 | 0,0E+00 | 0,0E+00 | 0,0E+00 | 244489 | 245346 | 858 | + |
| Y11_RS01165 | Y11_02471 | menC | o-succinylbenzoate synthase | 0,41 | -0,42 | -1,09 | -0,83 | -1,5 | -0,67 | 1,6E-01 | 1,3E-01 | 5,6E-03 | 8,9E-04 | 1,6E-04 | 9,6E-03 | 0,0E+00 | 0,0E+00 | 6,4E-05 | 0,0E+00 | 4,0E-06 | 0,0E+00 | 245346 | 246317 | 972 | + |
| Y11_RS01175 | Y11_02501 | katA | catalase KatA | 0,38 | 0,43 | 1,27 | 0,05 | 0,9 | 0,85 | 6,8E-06 | 4,3E-06 | 7,2E-08 | 1,9E-01 | 9,0E-07 | 1,7E-06 | 0,0E+00 | 0,0E+00 | 2,5E-09 | 0,0E+00 | 0,0E+00 | 0,0E+00 | 248349 | 249788 | 1440 | - |
| Y11_RS01180 | Y11_02511 | | YfaZ family protein | -1,3 | 0,34 | 0,33 | 1,63 | 1,63 | 0 | 8,4E-06 | 2,1E-02 | 1,1E-01 | 1,1E-06 | 2,5E-05 | 9,9E-01 | 4,3E-07 | 0,0E+00 | 0,0E+00 | 2,8E-08 | 7,3E-07 | 0,0E+00 | 250146 | 250688 | 543 | + |
| Y11_RS01190 | Y11_02531 | | nicotinamide mononucleotide deamidase-related protein YfaY | 0,52 | 0,84 | 0,4 | 0,32 | -0,12 | -0,44 | 5,0E-05 | 1,7E-07 | 7,6E-03 | 2,3E-04 | 3,1E-01 | 2,8E-03 | 0,0E+00 | 0,0E+00 | 0,0E+00 | 0,0E+00 | 0,0E+00 | 0,0E+00 | 252448 | 253650 | 1203 | + |
| Y11_RS01220 | Y11_02591 | eco | serine protease inhibitor ecotin | -0,05 | 1,35 | 1,66 | 1,39 | 1,71 | 0,32 | 5,1E-01 | 2,6E-07 | 7,9E-08 | 2,9E-08 | 1,2E-08 | 7,8E-03 | 0,0E+00 | 9,2E-09 | 2,7E-09 | 1,2E-09 | 1,2E-09 | 0,0E+00 | 261691 | 262197 | 507 | - |
| Y11_RS01230 | Y11_02611 | nrdB | ribonucleotide-diphosphate reductase subunit beta | -0,52 | 0,03 | 0,19 | 0,55 | 0,71 | 0,16 | 1,9E-06 | 4,8E-01 | 4,5E-02 | 8,1E-06 | 3,6E-05 | 1,1E-01 | 0,0E+00 | 0,0E+00 | 0,0E+00 | 0,0E+00 | 0,0E+00 | 0,0E+00 | 262953 | 264083 | 1131 | - |
| Y11_RS01235 | Y11_02621 | nrdA | ribonucleoside-diphosphate reductase subunit alpha | -0,28 | 0,19 | 0,21 | 0,46 | 0,48 | 0,02 | 1,1E-05 | 2,2E-04 | 1,7E-03 | 1,2E-09 | 1,2E-06 | 6,0E-01 | 0,0E+00 | 0,0E+00 | 0,0E+00 | 0,0E+00 | 0,0E+00 | 0,0E+00 | 264169 | 266454 | 2286 | - |
| Y11_RS01240 | Y11_02631 | ubiG | bifunctional 2-polyprenyl-6-hydroxyphenol methylase/3-demethylubiquinol 3-O-methyltransferase UbiG | 0,18 | -0,06 | -0,18 | -0,24 | -0,36 | -0,12 | 1,8E-02 | 5,9E-01 | 3,0E-01 | 4,8E-02 | 5,4E-02 | 5,2E-01 | 0,0E+00 | 0,0E+00 | 0,0E+00 | 0,0E+00 | 0,0E+00 | 0,0E+00 | 266971 | 267699 | 729 | - |
| Y11_RS01245 | Y11_02641 | gyrA | DNA topoisomerase (ATP-hydrolyzing) subunit A | 0,06 | -0,75 | -0,25 | -0,72 | -0,96 | -0,24 | 8,2E-01 | 4,9E-09 | 6,3E-01 | 2,6E-09 | 4,2E-08 | 1,8E-03 | 0,0E+00 | 0,0E+00 | 0,0E+00 | 0,0E+00 | 0,0E+00 | 0,0E+00 | 267903 | 270560 | 2658 | + |
| Y11_RS01250 | Y11_02651 | rcsC | two-component system sensor histidine kinase RcsC | 0,13 | -0,25 | -0,04 | -0,37 | -0,16 | 0,21 | 7,2E-01 | 2,9E-01 | 8,1E-01 | 3,6E-01 | 6,6E-01 | 3,6E-01 | 0,0E+00 | 0,0E+00 | 0,0E+00 | 0,0E+00 | 0,0E+00 | 0,0E+00 | 270678 | 273545 | 2868 | + |
| Y11_RS01255 | Y11_02661 | rcsB | transcriptional regulator RcsB | 0,14 | -0,39 | -0,31 | -0,53 | -0,45 | 0,09 | 7,5E-02 | 2,0E-04 | 4,6E-04 | 3,2E-05 | 5,0E-05 | 1,1E-01 | 0,0E+00 | 0,0E+00 | 0,0E+00 | 0,0E+00 | 0,0E+00 | 0,0E+00 | 273807 | 274454 | 648 | - |
| Y11_RS01260 | Y11_02671 | rcsD | phosphotransferase RcsD | 0,17 | -0,47 | -0,2 | -0,64 | -0,38 | 0,27 | 1,7E-03 | 1,4E-04 | 3,4E-02 | 8,2E-06 | 1,1E-03 | 2,4E-02 | 0,0E+00 | 0,0E+00 | 0,0E+00 | 0,0E+00 | 0,0E+00 | 0,0E+00 | 274457 | 277150 | 2694 | - |
| Y11_RS01275 | Y11_02741 | | porin OmpC | -1,48 | 0,9 | -0,15 | 2,38 | 1,34 | -1,04 | 2,3E-08 | 7,8E-08 | 1,6E-01 | 1,9E-10 | 9,0E-07 | 2,1E-06 | 2,5E-09 | 0,0E+00 | 0,0E+00 | 2,7E-11 | 3,8E-08 | 3,3E-07 | 280776 | 281879 | 1104 | + |
| Y11_RS01300 | Y11_02781 | | MBL fold metallo-hydrolase | -0,04 | -0,9 | 0,38 | -0,85 | 0,42 | 1,27 | 9,3E-01 | 1,2E-02 | 3,0E-01 | 9,1E-02 | 4,1E-01 | 1,2E-03 | 0,0E+00 | 0,0E+00 | 0,0E+00 | 0,0E+00 | 0,0E+00 | 7,7E-05 | 285656 | 286645 | 990 | + |
| Y11_RS01315 | Y11_02821 | | DinI family protein | 0 | -15 | 0 | -15 | 0 | 2,27 | 0,0E+00 | 0,0E+00 | 0,0E+00 | 0,0E+00 | 0,0E+00 | 1,0E-05 | 0,0E+00 | 0,0E+00 | 0,0E+00 | 0,0E+00 | 0,0E+00 | 1,2E-06 | 289928 | 290170 | 243 | - |
| Y11_RS01335 | Y11_02871 | | MurR/RpiR family transcriptional regulator | 0,16 | 0,19 | 1,48 | 0,03 | 1,32 | 1,28 | 4,7E-01 | 3,3E-01 | 3,6E-06 | 8,9E-01 | 1,0E-04 | 3,6E-05 | 0,0E+00 | 0,0E+00 | 6,8E-08 | 0,0E+00 | 2,7E-06 | 3,4E-06 | 292565 | 293305 | 741 | + |
| Y11_RS01340 | Y11_02891 | | PTS transporter subunit EIIC | -0,42 | 0,02 | 1,71 | 0,44 | 2,13 | 1,69 | 2,0E-02 | 9,2E-01 | 4,1E-06 | 1,2E-02 | 5,7E-08 | 2,7E-06 | 0,0E+00 | 0,0E+00 | 7,6E-08 | 0,0E+00 | 3,9E-09 | 4,0E-07 | 293471 | 294997 | 1527 | + |
| Y11_RS01345 | Y11_02901 | | 6-phospho-beta-glucosidase | -0,28 | -0,14 | 0,32 | 0,14 | 0,6 | 0,46 | 3,8E-05 | 4,0E-02 | 3,3E-04 | 5,9E-02 | 9,8E-06 | 2,9E-04 | 0,0E+00 | 0,0E+00 | 0,0E+00 | 0,0E+00 | 0,0E+00 | 0,0E+00 | 295101 | 296537 | 1437 | + |
| Y11_RS01375 | Y11_02951 | yejM | LPS biosynthesis-modulating metalloenzyme YejM | -0,19 | -0,49 | -0,49 | -0,3 | -0,3 | 0 | 8,7E-03 | 5,0E-04 | 9,8E-05 | 8,1E-03 | 2,1E-03 | 9,7E-01 | 0,0E+00 | 0,0E+00 | 0,0E+00 | 0,0E+00 | 0,0E+00 | 0,0E+00 | 299936 | 301729 | 1794 | - |
| Y11_RS01380 | Y11_02961 | | YejL family protein | -0,01 | -0,51 | -0,68 | -0,5 | -0,67 | -0,16 | 9,5E-01 | 3,2E-03 | 1,2E-03 | 1,2E-02 | 4,2E-03 | 1,7E-01 | 0,0E+00 | 0,0E+00 | 0,0E+00 | 0,0E+00 | 0,0E+00 | 0,0E+00 | 301757 | 301984 | 228 | - |
| Y11_RS01385 | Y11_02971 | yejK | nucleoid-associated protein YejK | -0,32 | -0,36 | -0,55 | -0,05 | -0,24 | -0,19 | 9,1E-04 | 4,5E-03 | 1,3E-03 | 5,6E-01 | 5,2E-02 | 1,7E-01 | 0,0E+00 | 0,0E+00 | 0,0E+00 | 0,0E+00 | 0,0E+00 | 0,0E+00 | 302162 | 303163 | 1002 | + |
| Y11_RS01390 | Y11_02981 | rplY | 50S ribosomal protein L25 | 0,29 | -0,47 | -0,65 | -0,77 | -0,94 | -0,18 | 1,2E-02 | 3,1E-02 | 9,7E-02 | 3,1E-03 | 2,7E-02 | 6,6E-01 | 0,0E+00 | 0,0E+00 | 0,0E+00 | 0,0E+00 | 0,0E+00 | 0,0E+00 | 303245 | 303532 | 288 | - |
| Y11_RS01400 | Y11_03001 | rsuA | 16S rRNA pseudouridine(516) synthase RsuA | -0,03 | -0,26 | -0,62 | -0,23 | -0,58 | -0,36 | 7,4E-01 | 6,9E-03 | 6,3E-03 | 1,4E-02 | 8,3E-03 | 5,7E-02 | 0,0E+00 | 0,0E+00 | 0,0E+00 | 0,0E+00 | 0,0E+00 | 0,0E+00 | 305913 | 306620 | 708 | + |
| Y11_RS01415 | Y11_03031 | yejF | microcin C ABC transporter ATP-binding protein YejF | -1,67 | 0,6 | 1,06 | 2,27 | 2,72 | 0,45 | 7,1E-11 | 3,2E-07 | 8,4E-06 | 1,5E-11 | 5,9E-09 | 2,8E-03 | 2,7E-11 | 0,0E+00 | 1,4E-07 | 6,1E-12 | 6,3E-10 | 0,0E+00 | 309214 | 310806 | 1593 | - |
| Y11_RS01420 | Y11_03041 | | microcin C ABC transporter permease | -1,18 | 0,49 | 1,3 | 1,67 | 2,48 | 0,81 | 3,7E-02 | 8,9E-02 | 3,0E-04 | 8,4E-03 | 6,7E-04 | 8,6E-03 | 1,2E-03 | 0,0E+00 | 3,8E-06 | 1,4E-04 | 1,6E-05 | 0,0E+00 | 310808 | 311836 | 1029 | - |
| Y11_RS01425 | Y11_03051 | | microcin C ABC transporter permease YejB | -1,95 | 0,53 | 0,8 | 2,48 | 2,75 | 0,27 | 5,1E-06 | 6,7E-03 | 4,9E-04 | 2,5E-07 | 9,3E-08 | 3,5E-02 | 2,8E-07 | 0,0E+00 | 0,0E+00 | 7,4E-09 | 5,9E-09 | 0,0E+00 | 311833 | 312933 | 1101 | - |
| Y11_RS01430 | Y11_03061 | | extracellular solute-binding protein | -1,82 | 0,19 | 0,59 | 2,01 | 2,42 | 0,4 | 5,3E-11 | 1,1E-04 | 2,9E-08 | 2,3E-11 | 6,5E-12 | 5,4E-07 | 2,2E-11 | 0,0E+00 | 0,0E+00 | 7,5E-12 | 8,8E-12 | 0,0E+00 | 312943 | 314751 | 1809 | - |
| Y11_RS01435 | Y11_03071 | | cyclic di-GMP phosphodiesterase | 0,32 | 1,92 | 2,65 | 1,6 | 2,34 | 0,73 | 4,7E-02 | 5,9E-07 | 3,3E-06 | 1,9E-05 | 1,9E-05 | 2,3E-02 | 0,0E+00 | 1,8E-08 | 6,2E-08 | 3,7E-07 | 5,9E-07 | 0,0E+00 | 314850 | 316412 | 1563 | - |
| Y11_RS01450 | Y11_03091 | | phosphatase PAP2 family protein | 1,11 | 0,81 | 1,85 | -0,3 | 0,73 | 1,04 | 7,6E-02 | 3,9E-02 | 5,7E-03 | 5,2E-01 | 2,4E-01 | 2,7E-02 | 2,5E-03 | 0,0E+00 | 6,6E-05 | 0,0E+00 | 0,0E+00 | 1,4E-03 | 317899 | 318600 | 702 | - |
| Y11_RS01455 | Y11_03101 | | GTP-binding protein | 0,36 | 0,52 | 1,04 | 0,16 | 0,67 | 0,52 | 1,9E-04 | 2,7E-05 | 3,7E-05 | 3,5E-02 | 7,4E-04 | 4,1E-03 | 0,0E+00 | 0,0E+00 | 5,5E-07 | 0,0E+00 | 0,0E+00 | 0,0E+00 | 318664 | 319647 | 984 | - |
| Y11_RS01465 | Y11_03121 | | GntR family transcriptional regulator | 0,07 | 0,61 | 0,62 | 0,53 | 0,55 | 0,01 | 6,8E-01 | 1,4E-03 | 8,4E-04 | 7,2E-03 | 5,0E-03 | 8,8E-01 | 0,0E+00 | 0,0E+00 | 0,0E+00 | 0,0E+00 | 0,0E+00 | 0,0E+00 | 320512 | 321255 | 744 | + |
| Y11_RS01470 | Y11_03131 | | fructuronate reductase | -0,19 | -2,48 | -2,95 | -2,3 | -2,76 | -0,46 | 2,1E-02 | 1,3E-09 | 1,5E-08 | 6,0E-10 | 1,5E-08 | 7,5E-03 | 0,0E+00 | 1,3E-10 | 7,1E-10 | 5,9E-11 | 1,3E-09 | 0,0E+00 | 321355 | 322827 | 1473 | - |
| Y11_RS01480 | Y11_03151 | uxuA | mannonate dehydratase | -0,04 | -0,36 | -0,42 | -0,32 | -0,39 | -0,06 | 6,7E-01 | 4,0E-03 | 1,7E-03 | 3,5E-03 | 1,4E-03 | 4,7E-01 | 0,0E+00 | 0,0E+00 | 0,0E+00 | 0,0E+00 | 0,0E+00 | 0,0E+00 | 323255 | 324445 | 1191 | + |
| Y11_RS01485 | Y11_03161 | yeiP | elongation factor P-like protein YeiP | -0,26 | -2,75 | -3,17 | -2,49 | -2,91 | -0,42 | 1,3E-01 | 8,8E-10 | 3,4E-09 | 1,4E-07 | 1,2E-07 | 3,7E-03 | 0,0E+00 | 9,9E-11 | 2,4E-10 | 4,6E-09 | 6,9E-09 | 0,0E+00 | 324722 | 325294 | 573 | - |
| Y11_RS01510 | Y11_03231 | | NUDIX domain-containing protein | -0,24 | -0,57 | -1,13 | -0,31 | -0,87 | -0,56 | 2,9E-01 | 2,9E-02 | 1,7E-03 | 4,1E-02 | 9,8E-04 | 1,1E-02 | 0,0E+00 | 0,0E+00 | 2,1E-05 | 0,0E+00 | 0,0E+00 | 0,0E+00 | 331602 | 332024 | 423 | - |
| Y11_RS01515 | Y11_03241 | fruB | fused PTS fructose transporter subunit IIA/HPr protein | -1,91 | 0,86 | -0,39 | 2,77 | 1,52 | -1,25 | 5,6E-09 | 2,4E-06 | 1,1E-01 | 2,9E-11 | 1,0E-04 | 3,8E-04 | 9,5E-10 | 0,0E+00 | 0,0E+00 | 8,5E-12 | 2,7E-06 | 2,7E-05 | 332285 | 333418 | 1134 | + |
| Y11_RS01520 | Y11_03251 | fruK | 1-phosphofructokinase | -1,9 | 0,97 | -0,55 | 2,86 | 1,34 | -1,52 | 1,4E-09 | 2,0E-07 | 4,0E-02 | 1,3E-10 | 3,6E-04 | 1,5E-04 | 3,2E-10 | 0,0E+00 | 0,0E+00 | 2,1E-11 | 8,7E-06 | 1,2E-05 | 333415 | 334353 | 939 | + |
| Y11_RS01530 | Y11_03281 | | winged helix-turn-helix domain-containing protein | 0,53 | -1,96 | -1,66 | -2,48 | -2,19 | 0,3 | 1,3E-02 | 2,4E-06 | 5,0E-03 | 6,1E-08 | 7,9E-04 | 4,9E-01 | 0,0E+00 | 6,5E-08 | 5,8E-05 | 2,2E-09 | 1,8E-05 | 0,0E+00 | 336312 | 336977 | 666 | + |
| Y11_RS01535 | | | fimbrial polyadhesin regulatory protein MyfF | 0,44 | -0,27 | 0,89 | -0,71 | 0,44 | 1,15 | 4,2E-02 | 5,0E-02 | 2,2E-04 | 8,1E-03 | 8,0E-02 | 1,2E-04 | 0,0E+00 | 0,0E+00 | 0,0E+00 | 0,0E+00 | 0,0E+00 | 9,9E-06 | 336974 | 337465 | 492 | + |
| Y11_RS01540 | Y11_03301 | myfA | fimbrial polyadhesin major subunit MyfA | -0,27 | 0,62 | 1,18 | 0,89 | 1,47 | 0,57 | 7,7E-02 | 5,1E-02 | 1,5E-04 | 7,7E-03 | 6,8E-06 | 7,5E-02 | 0,0E+00 | 0,0E+00 | 2,1E-06 | 0,0E+00 | 2,3E-07 | 0,0E+00 | 337726 | 338205 | 480 | + |
| Y11_RS01555 | Y11_03341 | nfo | deoxyribonuclease IV | 0,09 | 1,06 | 1,41 | 0,97 | 1,32 | 0,34 | 5,3E-03 | 4,1E-07 | 1,9E-09 | 6,2E-07 | 1,6E-09 | 2,6E-03 | 0,0E+00 | 1,3E-08 | 1,6E-10 | 0,0E+00 | 2,4E-10 | 0,0E+00 | 341784 | 342629 | 846 | - |
| Y11_RS01575 | Y11_03381 | | LysR family transcriptional regulator | -0,15 | 0,87 | 1,17 | 1,02 | 1,32 | 0,31 | 2,7E-02 | 2,7E-07 | 4,6E-09 | 4,0E-08 | 5,9E-10 | 5,5E-05 | 0,0E+00 | 0,0E+00 | 3,0E-10 | 1,6E-09 | 1,2E-10 | 0,0E+00 | 347096 | 347968 | 873 | + |
| Y11_RS01580 | Y11_03391 | | amino acid permease | -0,45 | -0,19 | -0,04 | 0,25 | 0,4 | 0,15 | 2,0E-02 | 2,3E-01 | 8,7E-01 | 7,7E-02 | 1,2E-01 | 5,3E-01 | 0,0E+00 | 0,0E+00 | 0,0E+00 | 0,0E+00 | 0,0E+00 | 0,0E+00 | 348213 | 349739 | 1527 | + |
| Y11_RS01590 | Y11_03411 | | ABC transporter ATP-binding protein | -0,27 | -0,34 | -0,84 | -0,08 | -0,59 | -0,5 | 1,4E-01 | 2,5E-03 | 2,0E-02 | 6,8E-01 | 1,1E-01 | 1,3E-01 | 0,0E+00 | 0,0E+00 | 0,0E+00 | 0,0E+00 | 0,0E+00 | 0,0E+00 | 351093 | 351881 | 789 | + |
| Y11_RS01595 | Y11_03421 | | ligand-gated channel protein | 1,21 | -1,56 | -1,96 | -2,77 | -3,17 | -0,4 | 7,3E-07 | 1,8E-08 | 4,6E-09 | 5,8E-10 | 2,8E-10 | 3,3E-04 | 4,9E-08 | 9,8E-10 | 3,0E-10 | 5,8E-11 | 8,3E-11 | 0,0E+00 | 352105 | 354078 | 1974 | + |
| Y11_RS01600 | Y11_03431 | | glutathione S-transferase | 0,32 | 1,03 | 1,67 | 0,72 | 1,35 | 0,64 | 2,4E-04 | 2,1E-08 | 1,0E-07 | 2,7E-07 | 4,9E-07 | 1,2E-04 | 0,0E+00 | 1,1E-09 | 3,3E-09 | 0,0E+00 | 2,3E-08 | 0,0E+00 | 354626 | 355246 | 621 | - |
| Y11_RS01615 | Y11_03471 | | MetQ/NlpA family ABC transporter substrate-binding protein | 0 | -15 | -15 | -4,58 | -2,91 | 1,65 | 0,0E+00 | 0,0E+00 | 0,0E+00 | 1,5E-06 | 4,0E-06 | 4,0E-03 | 0,0E+00 | 0,0E+00 | 0,0E+00 | 3,7E-08 | 1,4E-07 | 2,5E-04 | 357514 | 358338 | 825 | + |
| Y11_RS01630 | Y11_03501 | | PLP-dependent aspartate aminotransferase family protein | -1,08 | 1,33 | 2 | 2,41 | 3,08 | 0,67 | 1,5E-05 | 1,6E-06 | 4,2E-08 | 2,1E-08 | 2,0E-09 | 9,9E-05 | 7,2E-07 | 4,5E-08 | 1,7E-09 | 9,2E-10 | 2,9E-10 | 0,0E+00 | 360106 | 361251 | 1146 | - |
| Y11_RS01635 | Y11_03511 | | cystathionine beta-synthase | -2,28 | -4,3 | -2,1 | -2,02 | 0,18 | 2,2 | 2,4E-07 | 3,7E-10 | 4,2E-07 | 2,3E-07 | 2,5E-01 | 1,1E-07 | 1,9E-08 | 5,3E-11 | 1,1E-08 | 6,8E-09 | 0,0E+00 | 3,8E-08 | 361277 | 362644 | 1368 | - |
| Y11_RS01640 | Y11_03521 | | serine hydrolase | 0,24 | -1,26 | -1,3 | -1,5 | -1,54 | -0,04 | 2,5E-03 | 3,3E-08 | 1,9E-07 | 1,5E-11 | 3,8E-09 | 5,5E-01 | 0,0E+00 | 1,6E-09 | 5,6E-09 | 6,1E-12 | 4,5E-10 | 0,0E+00 | 363061 | 364254 | 1194 | + |
| Y11_RS01650 | Y11_03541 | | L-serine ammonia-lyase | -0,21 | -1,86 | -0,03 | -1,65 | 0,18 | 1,83 | 4,8E-02 | 1,4E-09 | 7,1E-01 | 2,5E-08 | 6,0E-02 | 4,0E-10 | 0,0E+00 | 1,4E-10 | 0,0E+00 | 1,0E-09 | 0,0E+00 | 1,1E-09 | 366024 | 367391 | 1368 | - |
| Y11_RS01655 | Y11_03551 | | HAAAP family serine/threonine permease | -0,93 | -0,81 | 0,7 | 0,12 | 1,63 | 1,51 | 2,0E-03 | 5,1E-03 | 1,2E-02 | 3,1E-01 | 7,1E-07 | 2,4E-06 | 0,0E+00 | 0,0E+00 | 0,0E+00 | 0,0E+00 | 3,1E-08 | 3,7E-07 | 367537 | 368832 | 1296 | - |
| Y11_RS01665 | Y11_03571 | deoC | deoxyribose-phosphate aldolase | -0,51 | -2 | -0,6 | -1,49 | -0,09 | 1,4 | 4,9E-04 | 8,3E-09 | 9,0E-03 | 6,5E-07 | 6,5E-01 | 5,9E-05 | 0,0E+00 | 5,3E-10 | 0,0E+00 | 1,7E-08 | 0,0E+00 | 5,1E-06 | 371044 | 371715 | 672 | - |
| Y11_RS01680 | Y11_03601 | | aspartate:alanine antiporter | -0,08 | -0,66 | -0,7 | -0,58 | -0,62 | -0,04 | 5,1E-01 | 2,8E-05 | 6,9E-03 | 1,1E-03 | 2,0E-02 | 8,5E-01 | 0,0E+00 | 0,0E+00 | 0,0E+00 | 0,0E+00 | 0,0E+00 | 0,0E+00 | 373638 | 375326 | 1689 | - |
| Y11_RS01685 | Y11_03611 | | GrxA family glutaredoxin | 0,54 | 0,12 | 0,29 | -0,42 | -0,25 | 0,17 | 4,8E-05 | 7,1E-02 | 1,4E-01 | 6,1E-04 | 2,1E-01 | 3,8E-01 | 0,0E+00 | 0,0E+00 | 0,0E+00 | 0,0E+00 | 0,0E+00 | 0,0E+00 | 375735 | 375998 | 264 | - |
| Y11_RS01690 | Y11_03621 | | YbjC family protein | 15 | -1,83 | 0,89 | -15 | -15 | 2,7 | 0,0E+00 | 1,3E-04 | 2,0E-01 | 0,0E+00 | 0,0E+00 | 2,3E-03 | 0,0E+00 | 2,7E-06 | 0,0E+00 | 0,0E+00 | 0,0E+00 | 1,5E-04 | 376313 | 376621 | 309 | + |
| Y11_RS01695 | Y11_03631 | | YbjN domain-containing protein | -0,09 | -0,08 | -0,24 | 0,02 | -0,14 | -0,16 | 4,7E-01 | 5,4E-01 | 3,8E-01 | 8,9E-01 | 6,0E-01 | 5,6E-01 | 0,0E+00 | 0,0E+00 | 0,0E+00 | 0,0E+00 | 0,0E+00 | 0,0E+00 | 376740 | 377225 | 486 | + |
| Y11_RS01700 | Y11_03641 | potF | spermidine/putrescine ABC transporter substrate-binding protein PotF | 0,01 | -1,35 | -1 | -1,36 | -1,01 | 0,35 | 8,7E-01 | 7,4E-09 | 8,1E-05 | 3,6E-08 | 9,9E-05 | 3,2E-02 | 0,0E+00 | 5,0E-10 | 0,0E+00 | 1,4E-09 | 2,6E-06 | 0,0E+00 | 377671 | 378780 | 1110 | + |
| Y11_RS01705 | Y11_03661 | potG | putrescine ABC transporter ATP-binding subunit PotG | -0,07 | -1,82 | -1,75 | -1,74 | -1,68 | 0,07 | 6,9E-01 | 1,5E-05 | 3,6E-05 | 7,8E-07 | 4,2E-06 | 7,2E-01 | 0,0E+00 | 3,4E-07 | 5,4E-07 | 2,0E-08 | 1,5E-07 | 0,0E+00 | 378947 | 380080 | 1134 | + |

| | | | | | | | | | | | | | | | | | | | | | | | | | |
|---|---|---|---|---|---|---|---|---|---|---|---|---|---|---|---|---|---|---|---|---|---|---|---|---|---|
| Y11_RS01725 | Y11_03711 | rlmC | 23S rRNA (uracil(747)-C(5))-methyltransferase RlmC | -0,04 | -0,82 | -1,7 | -0,78 | -1,65 | -0,87 | 8,4E-01 | 6,8E-04 | 5,3E-04 | 5,8E-03 | 1,2E-03 | 2,4E-02 | 0,0E+00 | 0,0E+00 | 6,6E-06 | 0,0E+00 | 2,7E-05 | 0,0E+00 | 382624 | 383751 | 1128 | + |
| Y11_RS01730 | Y11_03721 | | arginine ABC transporter substrate-binding protein | 0,11 | 0,06 | -0,07 | -0,05 | -0,17 | -0,12 | 3,1E-01 | 4,1E-01 | 6,2E-01 | 5,9E-01 | 2,5E-01 | 3,2E-01 | 0,0E+00 | 0,0E+00 | 0,0E+00 | 0,0E+00 | 0,0E+00 | 0,0E+00 | 383871 | 384620 | 750 | - |
| Y11_RS01745 | Y11_03761 | artJ | arginine ABC transporter substrate-binding protein | -0,07 | 0,36 | 0,13 | 0,42 | 0,2 | -0,23 | 3,0E-01 | 5,6E-04 | 3,2E-01 | 2,9E-04 | 1,6E-01 | 1,3E-01 | 0,0E+00 | 0,0E+00 | 0,0E+00 | 0,0E+00 | 0,0E+00 | 0,0E+00 | 386701 | 387432 | 732 | - |
| Y11_RS01750 | Y11_03771 | artP | arginine ABC transporter ATP-binding protein ArtP | -0,45 | -0,11 | -0,35 | 0,33 | 0,09 | -0,24 | 1,3E-02 | 4,7E-01 | 7,2E-02 | 1,1E-01 | 6,5E-01 | 2,8E-01 | 0,0E+00 | 0,0E+00 | 0,0E+00 | 0,0E+00 | 0,0E+00 | 0,0E+00 | 387453 | 388181 | 729 | - |
| Y11_RS01755 | Y11_03781 | | lipoprotein | 0,23 | 0,75 | 0,92 | 0,53 | 0,69 | 0,16 | 1,0E-04 | 5,8E-09 | 4,9E-04 | 8,5E-09 | 2,7E-03 | 3,4E-01 | 0,0E+00 | 0,0E+00 | 0,0E+00 | 0,0E+00 | 0,0E+00 | 0,0E+00 | 388605 | 389180 | 576 | - |
| Y11_RS01760 | Y11_03791 | | NAD(P)-dependent oxidoreductase | 0,62 | -1,3 | -0,96 | -1,92 | -1,57 | 0,35 | 2,2E-03 | 2,3E-08 | 1,6E-06 | 6,3E-07 | 4,4E-06 | 1,1E-03 | 0,0E+00 | 1,2E-09 | 0,0E+00 | 1,7E-08 | 1,5E-07 | 0,0E+00 | 389249 | 390259 | 1011 | - |
| Y11_RS01765 | Y11_03801 | | DUF2867 domain-containing protein | 0,49 | 0,3 | 1,05 | -0,19 | 0,56 | 0,75 | 3,8E-03 | 1,5E-02 | 2,1E-04 | 9,6E-02 | 9,5E-03 | 1,0E-03 | 0,0E+00 | 0,0E+00 | 2,7E-06 | 0,0E+00 | 0,0E+00 | 0,0E+00 | 390391 | 391848 | 1458 | - |
| Y11_RS01770 | Y11_03811 | ltaE | low-specificity L-threonine aldolase | 0,28 | -0,5 | 0,28 | -0,78 | 0 | 0,78 | 2,6E-04 | 4,2E-05 | 1,9E-02 | 1,1E-06 | 9,8E-01 | 6,5E-05 | 0,0E+00 | 0,0E+00 | 0,0E+00 | 0,0E+00 | 0,0E+00 | 0,0E+00 | 391845 | 392864 | 1020 | - |
| Y11_RS01780 | Y11_03831 | poxB | ubiquinone-dependent pyruvate dehydrogenase | -0,3 | 0,27 | 1,36 | 0,58 | 1,67 | 1,09 | 1,3E-02 | 1,3E-02 | 1,1E-06 | 6,2E-04 | 7,8E-07 | 1,1E-05 | 0,0E+00 | 0,0E+00 | 2,5E-08 | 0,0E+00 | 3,4E-08 | 1,3E-06 | 394121 | 395842 | 1722 | - |
| Y11_RS01800 | Y11_03871 | | ATP-dependent endonuclease | 0,72 | 0,46 | 1,57 | -0,25 | 0,85 | 1,1 | 4,6E-08 | 1,9E-05 | 1,2E-07 | 2,9E-04 | 8,1E-06 | 2,2E-06 | 0,0E+00 | 0,0E+00 | 3,8E-09 | 0,0E+00 | 0,0E+00 | 3,4E-07 | 400062 | 401726 | 1665 | + |
| Y11_RS01805 | Y11_03881 | | VirK/YbjX family protein | 0 | -15 | -15 | -15 | -15 | 0,58 | 0,0E+00 | 0,0E+00 | 0,0E+00 | 0,0E+00 | 0,0E+00 | 2,4E-01 | 0,0E+00 | 0,0E+00 | 0,0E+00 | 0,0E+00 | 0,0E+00 | 0,0E+00 | 401723 | 402706 | 984 | - |
| Y11_RS01810 | Y11_03891 | macA | macrolide transporter subunit MacA | -0,15 | -0,64 | -1,08 | -0,48 | -0,92 | -0,44 | 2,5E-04 | 5,3E-08 | 1,5E-05 | 1,9E-07 | 4,4E-05 | 5,7E-03 | 0,0E+00 | 0,0E+00 | 2,5E-07 | 0,0E+00 | 0,0E+00 | 0,0E+00 | 402898 | 404010 | 1113 | + |
| Y11_RS01815 | Y11_03901 | macB | macrolide ABC transporter ATP-binding protein/permease MacB | -0,13 | -0,06 | -0,49 | 0,06 | -0,37 | -0,43 | 8,1E-02 | 4,1E-01 | 7,6E-04 | 4,9E-01 | 8,6E-03 | 4,8E-03 | 0,0E+00 | 0,0E+00 | 0,0E+00 | 0,0E+00 | 0,0E+00 | 0,0E+00 | 404010 | 405959 | 1950 | + |
| Y11_RS01820 | Y11_03911 | cspD | cold shock-like protein CspD | 15 | -3,16 | -2,23 | -15 | -15 | 0,93 | 0,0E+00 | 3,3E-06 | 1,6E-06 | 0,0E+00 | 0,0E+00 | 1,9E-02 | 0,0E+00 | 8,6E-08 | 3,5E-08 | 0,0E+00 | 0,0E+00 | 0,0E+00 | 406158 | 406415 | 258 | - |
| Y11_RS01825 | Y11_03921 | clpS | ATP-dependent Clp protease adapter ClpS | 0,69 | 0,34 | -2,01 | -0,34 | -2,71 | -2,35 | 3,8E-02 | 1,9E-01 | 5,8E-05 | 3,4E-01 | 6,2E-05 | 9,7E-05 | 0,0E+00 | 0,0E+00 | 8,3E-07 | 0,0E+00 | 1,7E-06 | 8,0E-06 | 406775 | 407095 | 321 | + |
| Y11_RS01830 | Y11_03931 | clpA | ATP-dependent Clp protease ATP-binding subunit ClpA | 0,28 | -0,18 | -0,91 | -0,46 | -1,18 | -0,72 | 1,7E-04 | 2,4E-02 | 9,2E-09 | 2,7E-04 | 1,3E-08 | 8,5E-06 | 0,0E+00 | 0,0E+00 | 0,0E+00 | 0,0E+00 | 1,2E-09 | 0,0E+00 | 407121 | 409397 | 2277 | + |
| Y11_RS01835 | Y11_03941 | infA | translation initiation factor IF-1 | 0,19 | 0,35 | -0,09 | 0,16 | -0,28 | -0,44 | 2,8E-01 | 5,9E-02 | 7,3E-01 | 1,7E-01 | 2,2E-01 | 6,7E-02 | 0,0E+00 | 0,0E+00 | 0,0E+00 | 0,0E+00 | 0,0E+00 | 0,0E+00 | 409642 | 409860 | 219 | - |
| Y11_RS01840 | Y11_03951 | aat | leucyl/phenylalanyl-tRNA--protein transferase | 0,97 | 1,58 | 2,06 | 0,61 | 1,09 | 0,48 | 3,7E-02 | 2,7E-04 | 3,4E-05 | 1,1E-01 | 1,1E-02 | 2,0E-02 | 0,0E+00 | 5,7E-06 | 5,2E-07 | 0,0E+00 | 2,4E-04 | 0,0E+00 | 410026 | 410736 | 711 | - |
| Y11_RS01845 | Y11_03961 | cydC | cysteine/glutathione ABC transporter ATP-binding protein/permease CydC | 0,02 | 0,54 | 0,94 | 0,52 | 0,91 | 0,4 | 8,9E-01 | 6,3E-03 | 1,9E-04 | 3,3E-03 | 7,1E-05 | 9,3E-03 | 0,0E+00 | 0,0E+00 | 0,0E+00 | 0,0E+00 | 0,0E+00 | 0,0E+00 | 410970 | 412694 | 1725 | - |
| Y11_RS01850 | Y11_03971 | cydD | cysteine/glutathione ABC transporter permease/ATP-binding protein CydD | 0,27 | 0,27 | 0,58 | -0,01 | 0,31 | 0,32 | 1,6E-01 | 2,5E-02 | 4,6E-03 | 9,8E-01 | 1,7E-01 | 6,2E-02 | 0,0E+00 | 0,0E+00 | 0,0E+00 | 0,0E+00 | 0,0E+00 | 0,0E+00 | 412697 | 414463 | 1767 | - |
| Y11_RS01855 | Y11_03981 | trxB | thioredoxin-disulfide reductase | 0,14 | -0,02 | 0,09 | -0,16 | -0,05 | 0,11 | 2,1E-02 | 7,6E-01 | 2,0E-01 | 4,6E-02 | 4,9E-01 | 2,1E-01 | 0,0E+00 | 0,0E+00 | 0,0E+00 | 0,0E+00 | 0,0E+00 | 0,0E+00 | 414738 | 415700 | 963 | - |
| Y11_RS01860 | Y11_03991 | lrp | leucine-responsive transcriptional regulator Lrp | 0,83 | -0,91 | -1,14 | -1,74 | -1,97 | -0,23 | 1,4E-04 | 1,1E-05 | 1,4E-06 | 1,1E-06 | 3,6E-07 | 6,3E-02 | 0,0E+00 | 0,0E+00 | 3,1E-08 | 2,7E-08 | 1,8E-08 | 0,0E+00 | 416470 | 416964 | 495 | + |
| Y11_RS01865 | Y11_04001 | | DNA translocase FtsK | 0,28 | -0,14 | -0,75 | -0,41 | -1,02 | -0,61 | 3,3E-03 | 9,1E-02 | 5,4E-07 | 1,2E-03 | 4,5E-07 | 3,2E-05 | 0,0E+00 | 0,0E+00 | 0,0E+00 | 0,0E+00 | 2,2E-08 | 0,0E+00 | 417086 | 420700 | 3615 | + |
| Y11_RS01870 | Y11_04011 | lolA | outer membrane lipoprotein chaperone LolA | 0,37 | -0,93 | -1,15 | -1,3 | -1,52 | -0,22 | 7,5E-04 | 8,1E-07 | 8,8E-06 | 2,7E-07 | 1,9E-06 | 1,2E-01 | 0,0E+00 | 0,0E+00 | 1,5E-07 | 7,9E-09 | 7,4E-08 | 0,0E+00 | 420930 | 421541 | 612 | + |
| Y11_RS01875 | Y11_04021 | | replication-associated recombination protein A | 0,17 | -0,06 | -0,26 | -0,23 | -0,43 | -0,21 | 4,8E-02 | 4,8E-01 | 3,2E-02 | 3,1E-02 | 4,2E-03 | 1,1E-01 | 0,0E+00 | 0,0E+00 | 0,0E+00 | 0,0E+00 | 0,0E+00 | 0,0E+00 | 421549 | 422892 | 1344 | + |
| Y11_RS01880 | Y11_04031 | serS | serine--tRNA ligase | 0,18 | -0,55 | -0,87 | -0,74 | -1,06 | -0,32 | 2,0E-04 | 1,5E-06 | 3,9E-07 | 1,0E-07 | 6,9E-08 | 1,2E-03 | 0,0E+00 | 0,0E+00 | 0,0E+00 | 0,0E+00 | 4,6E-09 | 0,0E+00 | 423064 | 424356 | 1293 | + |
| Y11_RS01900 | Y11_04071 | pflA | pyruvate formate lyase 1-activating protein | -0,01 | 0,38 | 0,97 | 0,39 | 0,98 | 0,59 | 8,8E-01 | 4,9E-04 | 2,0E-07 | 7,0E-04 | 4,1E-07 | 2,0E-05 | 0,0E+00 | 0,0E+00 | 0,0E+00 | 0,0E+00 | 0,0E+00 | 0,0E+00 | 428671 | 429411 | 741 | - |
| Y11_RS01905 | Y11_04081 | pflB | formate C-acetyltransferase | 0,12 | 0,65 | 0,19 | 0,52 | 0,07 | -0,45 | 2,1E-02 | 1,8E-07 | 8,1E-03 | 1,1E-08 | 1,5E-01 | 4,4E-06 | 0,0E+00 | 0,0E+00 | 0,0E+00 | 0,0E+00 | 0,0E+00 | 0,0E+00 | 429490 | 431772 | 2283 | - |
| Y11_RS01910 | Y11_04091 | focA | formate transporter FocA | -0,9 | 1,23 | 1,77 | 2,13 | 2,66 | 0,54 | 1,3E-04 | 6,8E-04 | 5,6E-05 | 1,3E-05 | 2,3E-06 | 1,0E-01 | 0,0E+00 | 1,4E-05 | 8,0E-07 | 2,6E-07 | 8,7E-08 | 0,0E+00 | 431825 | 432682 | 858 | - |
| Y11_RS01925 | Y11_04141 | | 30S ribosomal protein S12 methylthiotransferase accessory protein YcaO | -0,15 | -2,56 | -2,45 | -2,41 | -2,29 | 0,12 | 1,8E-02 | 7,1E-12 | 1,1E-06 | 3,6E-12 | 1,7E-06 | 5,4E-01 | 0,0E+00 | 5,1E-12 | 2,4E-08 | 3,9E-12 | 6,5E-08 | 0,0E+00 | 434781 | 436547 | 1767 | - |
| Y11_RS01940 | Y11_04181 | serC | 3-phosphoserine/phosphohydroxythreonine transaminase | -0,04 | -1,51 | -1,48 | -1,47 | -1,44 | 0,03 | 4,5E-01 | 1,7E-08 | 3,0E-10 | 1,5E-08 | 1,4E-10 | 5,9E-01 | 0,0E+00 | 9,4E-10 | 4,4E-11 | 7,2E-10 | 5,9E-11 | 0,0E+00 | 438778 | 439863 | 1086 | + |
| Y11_RS01945 | Y11_04191 | aroA | 3-phosphoshikimate 1-carboxyvinyltransferase | -0,15 | -1,68 | -1,86 | -1,52 | -1,71 | -0,18 | 2,5E-02 | 2,2E-09 | 5,7E-09 | 1,2E-09 | 4,9E-09 | 2,7E-02 | 0,0E+00 | 1,9E-10 | 3,5E-10 | 9,8E-11 | 5,4E-10 | 0,0E+00 | 439990 | 441276 | 1287 | + |
| Y11_RS01950 | Y11_04201 | cmk | (d)CMP kinase | -0,05 | -0,81 | -0,21 | -0,76 | -0,16 | 0,6 | 5,7E-01 | 9,1E-07 | 1,6E-02 | 1,2E-06 | 4,1E-02 | 1,5E-06 | 0,0E+00 | 0,0E+00 | 0,0E+00 | 0,0E+00 | 0,0E+00 | 0,0E+00 | 441515 | 442207 | 693 | + |
| Y11_RS01955 | Y11_04211 | rpsA | 30S ribosomal protein S1 | 0,1 | -0,15 | -0,55 | -0,25 | -0,65 | -0,4 | 3,4E-02 | 1,9E-02 | 1,5E-06 | 8,6E-04 | 2,7E-07 | 4,8E-05 | 0,0E+00 | 0,0E+00 | 0,0E+00 | 0,0E+00 | 0,0E+00 | 0,0E+00 | 442376 | 444049 | 1674 | + |
| Y11_RS01960 | Y11_04221 | ihfB | integration host factor subunit beta | -0,45 | 1,47 | 2,02 | 1,92 | 2,47 | 0,55 | 1,0E-02 | 2,8E-06 | 1,4E-04 | 2,1E-08 | 2,4E-05 | 8,7E-02 | 0,0E+00 | 7,5E-08 | 1,9E-06 | 9,2E-10 | 7,0E-07 | 0,0E+00 | 444113 | 444397 | 285 | + |
| Y11_RS01970 | Y11_04241 | msbA | lipid A ABC transporter ATP-binding protein/permease MsbA | 0,03 | 0,17 | 0,05 | 0,14 | 0,02 | -0,12 | 6,3E-01 | 6,1E-02 | 5,7E-01 | 9,4E-02 | 8,4E-01 | 1,9E-01 | 0,0E+00 | 0,0E+00 | 0,0E+00 | 0,0E+00 | 0,0E+00 | 0,0E+00 | 447055 | 448803 | 1749 | + |
| Y11_RS01975 | Y11_04251 | lpxK | tetraacyldisaccharide 4'-kinase | 0,31 | 0,21 | -0,55 | -0,09 | -0,86 | -0,77 | 1,4E-01 | 2,2E-01 | 3,8E-02 | 5,8E-01 | 5,5E-03 | 5,6E-03 | 0,0E+00 | 0,0E+00 | 0,0E+00 | 0,0E+00 | 0,0E+00 | 0,0E+00 | 448800 | 449786 | 987 | + |
| Y11_RS01985 | Y11_04271 | | cold-shock protein | 0,02 | -4,68 | -5,3 | -4,7 | -5,31 | -0,62 | 9,2E-01 | 3,7E-09 | 1,7E-04 | 1,4E-08 | 1,7E-04 | 4,7E-01 | 0,0E+00 | 2,9E-10 | 2,2E-06 | 6,7E-10 | 4,4E-06 | 0,0E+00 | 451641 | 451850 | 210 | + |
| Y11_RS01995 | Y11_04301 | | hypothetical protein | -0,03 | -0,4 | -0,38 | -0,37 | -0,35 | 0,02 | 6,4E-01 | 5,6E-04 | 1,7E-01 | 2,2E-04 | 2,0E-01 | 9,4E-01 | 0,0E+00 | 0,0E+00 | 0,0E+00 | 0,0E+00 | 0,0E+00 | 0,0E+00 | 452707 | 452889 | 183 | + |
| Y11_RS02000 | Y11_04311 | kdsB | 3-deoxy-manno-octulosonate cytidylyltransferase | -0,53 | -0,77 | -0,95 | -0,24 | -0,43 | -0,19 | 2,9E-03 | 3,3E-05 | 1,4E-04 | 7,6E-02 | 2,7E-02 | 2,0E-01 | 0,0E+00 | 0,0E+00 | 0,0E+00 | 0,0E+00 | 0,0E+00 | 0,0E+00 | 452886 | 453638 | 753 | + |
| Y11_RS02005 | Y11_04321 | | YcbJ family phosphotransferase | 0,05 | -0,18 | -0,5 | -0,23 | -0,55 | -0,31 | 8,3E-01 | 3,9E-01 | 1,9E-01 | 2,1E-01 | 1,4E-01 | 3,5E-01 | 0,0E+00 | 0,0E+00 | 0,0E+00 | 0,0E+00 | 0,0E+00 | 0,0E+00 | 453828 | 454721 | 894 | + |
| Y11_RS02010 | Y11_04331 | elyC | envelope biogenesis factor ElyC | 0,06 | 0,21 | -0,03 | 0,14 | -0,1 | -0,24 | 7,2E-01 | 3,4E-01 | 8,7E-01 | 4,2E-01 | 4,7E-01 | 1,9E-01 | 0,0E+00 | 0,0E+00 | 0,0E+00 | 0,0E+00 | 0,0E+00 | 0,0E+00 | 454724 | 455491 | 768 | - |
| Y11_RS02015 | Y11_04351 | cmoM | tRNA uridine 5-oxyacetic acid(34) methyltransferase CmoM | 0,38 | -0,97 | -0,81 | -1,34 | -1,19 | 0,16 | 8,9E-02 | 2,6E-03 | 3,2E-03 | 3,7E-05 | 1,0E-05 | 3,6E-01 | 0,0E+00 | 0,0E+00 | 0,0E+00 | 7,2E-07 | 3,4E-07 | 0,0E+00 | 455682 | 456467 | 786 | + |
| Y11_RS02020 | Y11_04361 | mukF | chromosome partition protein MukF | -0,09 | -1,12 | -1,16 | -1,02 | -1,07 | -0,04 | 1,2E-01 | 1,3E-08 | 1,8E-08 | 1,1E-08 | 1,6E-08 | 3,4E-01 | 0,0E+00 | 7,8E-10 | 8,1E-10 | 5,8E-10 | 1,4E-09 | 0,0E+00 | 456464 | 457786 | 1323 | + |
| Y11_RS02025 | Y11_04371 | mukE | chromosome partition protein MukE | -0,18 | -1,08 | -1,32 | -0,9 | -1,13 | -0,24 | 2,4E-02 | 6,2E-06 | 3,0E-06 | 1,7E-05 | 6,7E-06 | 1,2E-01 | 0,0E+00 | 1,5E-07 | 5,8E-08 | 0,0E+00 | 2,3E-07 | 0,0E+00 | 457767 | 458495 | 729 | + |
| Y11_RS02030 | Y11_04381 | mukB | chromosome partition protein MukB | -0,12 | -0,88 | -1,02 | -0,76 | -0,91 | -0,14 | 7,0E-03 | 1,1E-12 | 2,2E-07 | 1,2E-08 | 1,3E-06 | 5,6E-02 | 0,0E+00 | 0,0E+00 | 6,2E-09 | 0,0E+00 | 0,0E+00 | 0,0E+00 | 458492 | 462937 | 4446 | + |
| Y11_RS02035 | Y11_04391 | ldtD | L,D-transpeptidase | 1,92 | 1,34 | -0,13 | -0,58 | -2,04 | -1,46 | 4,1E-07 | 4,9E-06 | 4,0E-01 | 6,4E-04 | 2,6E-07 | 2,5E-06 | 3,0E-08 | 1,2E-07 | 0,0E+00 | 0,0E+00 | 1,4E-08 | 3,8E-07 | 463261 | 465132 | 1872 | + |
| Y11_RS02045 | Y11_04411 | | MBL fold metallo-hydrolase | -0,3 | -0,89 | -1,04 | -0,59 | -0,75 | -0,16 | 7,7E-05 | 4,3E-07 | 5,7E-06 | 2,5E-06 | 4,4E-05 | 1,7E-01 | 0,0E+00 | 0,0E+00 | 1,0E-07 | 0,0E+00 | 0,0E+00 | 0,0E+00 | 466198 | 466845 | 648 | + |
| Y11_RS02060 | Y11_04431 | | aspartate/tyrosine/aromatic aminotransferase | 0,04 | -0,29 | -0,19 | -0,33 | -0,23 | 0,1 | 3,6E-01 | 2,4E-03 | 3,7E-02 | 7,7E-04 | 1,3E-02 | 2,7E-01 | 0,0E+00 | 0,0E+00 | 0,0E+00 | 0,0E+00 | 0,0E+00 | 0,0E+00 | 468932 | 470122 | 1191 | - |
| Y11_RS02065 | Y11_04441 | ompF | porin OmpF | 1,4 | -1,37 | 3,58 | -2,77 | 2,18 | 4,95 | 9,2E-08 | 7,0E-07 | 3,4E-11 | 2,1E-09 | 1,0E-09 | 1,5E-11 | 8,6E-09 | 2,1E-08 | 1,3E-11 | 1,5E-10 | 1,7E-10 | 2,3E-10 | 470378 | 471490 | 1113 | - |
| Y11_RS02070 | Y11_04451 | asnS | asparagine--tRNA ligase | 0,86 | 0,13 | 0,08 | -0,72 | -0,77 | -0,05 | 6,7E-08 | 2,2E-02 | 2,7E-01 | 2,2E-07 | 3,5E-06 | 4,9E-01 | 0,0E+00 | 0,0E+00 | 0,0E+00 | 0,0E+00 | 0,0E+00 | 0,0E+00 | 471792 | 473192 | 1401 | - |
| Y11_RS02075 | Y11_04461 | pncB | nicotinate phosphoribosyltransferase | 0,36 | -0,09 | -1,36 | -0,4 | -1,71 | -1,27 | 1,7E-02 | 4,9E-01 | 3,8E-04 | 1,2E-02 | 9,9E-05 | 7,8E-04 | 0,0E+00 | 0,0E+00 | 4,8E-06 | 0,0E+00 | 2,6E-06 | 5,3E-05 | 473449 | 474654 | 1206 | - |
| Y11_RS02080 | Y11_04471 | pepN | aminopeptidase N | 0,09 | 0,5 | 0,65 | 0,42 | 0,56 | 0,14 | 3,2E-02 | 5,0E-06 | 6,6E-04 | 8,6E-06 | 1,5E-03 | 2,8E-01 | 0,0E+00 | 0,0E+00 | 0,0E+00 | 0,0E+00 | 0,0E+00 | 0,0E+00 | 475248 | 477863 | 2616 | + |
| Y11_RS02085 | Y11_04481 | pyrD | quinone-dependent dihydroorotate dehydrogenase | 0,34 | -0,23 | 0,99 | -0,57 | 0,65 | 1,22 | 1,7E-05 | 1,6E-03 | 1,9E-09 | 1,3E-05 | 1,1E-06 | 2,9E-08 | 0,0E+00 | 0,0E+00 | 0,0E+00 | 0,0E+00 | 0,0E+00 | 1,4E-08 | 478031 | 479041 | 1011 | + |
| Y11_RS02090 | Y11_04501 | | cell division protein ZapC | 15 | -0,88 | -0,59 | -15 | -15 | 0,29 | 0,0E+00 | 2,1E-03 | 2,6E-02 | 0,0E+00 | 0,0E+00 | 1,6E-01 | 0,0E+00 | 0,0E+00 | 0,0E+00 | 0,0E+00 | 0,0E+00 | 0,0E+00 | 479217 | 479774 | 558 | + |
| Y11_RS02095 | Y11_04511 | | YcbX family protein | 0,54 | -2,03 | -2,06 | -2,57 | -2,6 | -0,03 | 5,7E-04 | 2,3E-09 | 2,2E-07 | 1,6E-09 | 5,9E-08 | 8,2E-01 | 0,0E+00 | 1,9E-10 | 6,2E-09 | 1,2E-10 | 4,0E-09 | 0,0E+00 | 479790 | 480902 | 1113 | - |
| Y11_RS02100 | Y11_04521 | rlmKL | bifunctional 23S rRNA (guanine(2069)-N(7))-methyltransferase RlmK/23S rRNA (guanine(2445)-N(2))-methyltransferase RlmL | -0,22 | -0,8 | -1,29 | -0,58 | -1,07 | -0,48 | 1,1E-02 | 7,9E-10 | 7,4E-09 | 3,0E-05 | 1,3E-06 | 2,0E-05 | 0,0E+00 | 0,0E+00 | 4,3E-10 | 0,0E+00 | 5,1E-08 | 0,0E+00 | 481002 | 483122 | 2121 | + |
| Y11_RS02105 | Y11_04531 | | ABC transporter ATP-binding protein | -0,15 | -0,73 | -1,18 | -0,58 | -1,03 | -0,45 | 1,6E-02 | 6,4E-07 | 3,7E-04 | 2,7E-06 | 8,6E-04 | 5,7E-02 | 0,0E+00 | 0,0E+00 | 4,7E-06 | 0,0E+00 | 2,0E-05 | 0,0E+00 | 483128 | 485041 | 1914 | + |
| Y11_RS02115 | Y11_04551 | pqiB | intermembrane transport protein PqiB | 0,02 | 0,1 | 0,1 | 0,08 | 0,08 | 0 | 8,5E-01 | 2,9E-01 | 4,4E-01 | 5,0E-01 | 5,8E-01 | 9,9E-01 | 0,0E+00 | 0,0E+00 | 0,0E+00 | 0,0E+00 | 0,0E+00 | 0,0E+00 | 486464 | 488116 | 1653 | + |
| Y11_RS02120 | Y11_04561 | pqiC | membrane integrity-associated transporter subunit PqiC | 0,06 | 0,93 | 1,93 | 0,87 | 1,87 | 1 | 8,0E-01 | 1,0E-02 | 1,4E-03 | 4,1E-03 | 9,6E-04 | 3,8E-02 | 0,0E+00 | 0,0E+00 | 1,7E-05 | 0,0E+00 | 2,2E-05 | 0,0E+00 | 488113 | 488664 | 552 | + |
| Y11_RS02125 | Y11_04581 | fabF | beta-ketoacyl-ACP synthase II | -0,21 | 1,17 | 1,19 | 1,38 | 1,4 | 0,02 | 3,0E-02 | 2,3E-07 | 2,9E-07 | 2,1E-08 | 3,0E-08 | 7,3E-01 | 0,0E+00 | 8,4E-09 | 7,9E-09 | 9,0E-10 | 2,3E-09 | 0,0E+00 | 489168 | 490442 | 1275 | - |
| Y11_RS02130 | Y11_04591 | | helix-turn-helix transcriptional regulator | 0 | -1,85 | -2,73 | -15 | -15 | -0,76 | 0,0E+00 | 2,4E-04 | 2,0E-04 | 0,0E+00 | 0,0E+00 | 1,5E-01 | 0,0E+00 | 5,0E-06 | 2,6E-06 | 0,0E+00 | 0,0E+00 | 0,0E+00 | 490442 | 490984 | 543 | - |
| Y11_RS02135 | Y11_04601 | fabA | bifunctional 3-hydroxydecanoyl-ACP dehydratase/trans-2-decenoyl-ACP isomerase | -0,54 | 0,18 | 0,16 | 0,72 | 0,7 | -0,02 | 7,3E-05 | 8,3E-02 | 4,7E-01 | 1,2E-04 | 1,2E-02 | 9,2E-01 | 0,0E+00 | 0,0E+00 | 0,0E+00 | 0,0E+00 | 0,0E+00 | 0,0E+00 | 491335 | 491853 | 519 | - |
| Y11_RS02140 | Y11_04611 | | Lon protease family protein | -0,13 | -0,22 | -0,6 | -0,09 | -0,47 | -0,38 | 3,2E-02 | 5,6E-04 | 8,6E-04 | 7,6E-02 | 4,0E-03 | 9,9E-03 | 0,0E+00 | 0,0E+00 | 0,0E+00 | 0,0E+00 | 0,0E+00 | 0,0E+00 | 491922 | 493673 | 1752 | - |
| Y11_RS02145 | Y11_04621 | matP | macrodomain Ter protein MatP | 0,11 | 0,92 | 1,08 | 0,81 | 0,97 | 0,17 | 4,9E-01 | 3,4E-04 | 8,3E-05 | 4,9E-05 | 5,9E-06 | 1,2E-01 | 0,0E+00 | 0,0E+00 | 1,1E-06 | 0,0E+00 | 0,0E+00 | 0,0E+00 | 493884 | 494339 | 456 | + |
| Y11_RS02150 | Y11_04631 | ompA | porin OmpA | -0,24 | 0,53 | 0,26 | 0,77 | 0,5 | -0,27 | 6,0E-03 | 6,6E-05 | 1,2E-01 | 1,6E-05 | 1,3E-02 | 1,3E-01 | 0,0E+00 | 0,0E+00 | 0,0E+00 | 0,0E+00 | 0,0E+00 | 0,0E+00 | 494470 | 495543 | 1074 | - |
| Y11_RS02165 | Y11_04671 | yccS | YccS family putative transporter | -2,02 | -1,36 | -1,58 | 0,75 | 0,49 | -0,25 | 2,6E-04 | 9,6E-04 | 2,6E-04 | 1,8E-02 | 7,5E-02 | 8,6E-02 | 1,1E-05 | 1,9E-05 | 3,4E-06 | 0,0E+00 | 0,0E+00 | 0,0E+00 | 497308 | 499443 | 2136 | - |
| Y11_RS02170 | Y11_04681 | | YccF domain-containing protein | -0,43 | 0,82 | 1,57 | 1,25 | 2 | 0,75 | 4,2E-03 | 7,5E-05 | 1,2E-04 | 8,8E-06 | 2,7E-05 | 1,3E-02 | 0,0E+00 | 0,0E+00 | 1,6E-06 | 1,8E-07 | 8,0E-07 | 0,0E+00 | 499468 | 499920 | 453 | - |

| | | | | | | | | | | | | | | | | | | | | | | | | |
|---|---|---|---|---|---|---|---|---|---|---|---|---|---|---|---|---|---|---|---|---|---|---|---|---|
| Y11_RS02175 | Y11_04691 | helD | DNA helicase IV | 0,11 | -0,34 | 0,1 | -0,45 | -0,01 | 0,43 | 1,4E-01 | 5,3E-04 | 1,5E-01 | 1,2E-04 | 8,7E-01 | 9,1E-05 | 0,0E+00 | 0,0E+00 | 0,0E+00 | 0,0E+00 | 0,0E+00 | 0,0E+00 | 500115 | 502169 | 2055 | + |
| Y11_RS02180 | Y11_04701 | | methylglyoxal synthase | -0,44 | -0,07 | 0,97 | 0,37 | 1,41 | 1,05 | 1,3E-02 | 4,0E-01 | 6,4E-06 | 1,7E-02 | 4,6E-06 | 2,7E-07 | 0,0E+00 | 0,0E+00 | 0,0E+00 | 0,0E+00 | 1,6E-07 | 6,9E-08 | 502232 | 502690 | 459 | - |
| Y11_RS02190 | Y11_04721 | | CoA-binding protein | 0,46 | 0,3 | 1,54 | -0,16 | 1,08 | 1,24 | 9,2E-03 | 1,2E-01 | 6,0E-05 | 2,5E-01 | 2,2E-04 | 2,6E-04 | 0,0E+00 | 0,0E+00 | 8,5E-07 | 0,0E+00 | 5,5E-06 | 1,9E-05 | 503889 | 504305 | 417 | + |
| Y11_RS02195 | Y11_04731 | hspQ | heat shock protein HspQ | 0,6 | 2,02 | 2,43 | 1,42 | 1,83 | 0,41 | 4,8E-01 | 2,9E-02 | 8,1E-03 | 3,0E-02 | 2,9E-03 | 2,6E-01 | 0,0E+00 | 5,6E-04 | 9,2E-05 | 4,7E-04 | 6,5E-05 | 0,0E+00 | 504388 | 504705 | 318 | - |
| Y11_RS02200 | Y11_04741 | rlmI | 23S rRNA (cytosine(1962)-C(5))-methyltransferase RlmI | 0,05 | -0,46 | -0,57 | -0,51 | -0,63 | -0,12 | 5,1E-01 | 4,1E-04 | 6,8E-03 | 2,8E-05 | 3,0E-03 | 4,6E-01 | 0,0E+00 | 0,0E+00 | 0,0E+00 | 0,0E+00 | 0,0E+00 | 0,0E+00 | 504768 | 505958 | 1191 | - |
| Y11_RS02205 | Y11_04751 | | hypothetical protein | -0,21 | 1,33 | 1,65 | 1,54 | 1,86 | 0,32 | 1,7E-01 | 9,1E-06 | 5,3E-07 | 3,0E-05 | 4,1E-06 | 9,0E-02 | 0,0E+00 | 2,2E-07 | 1,3E-08 | 5,8E-07 | 1,5E-07 | 0,0E+00 | 506208 | 506417 | 210 | + |
| Y11_RS02215 | Y11_04771 | yccX | acylphosphatase | 0,09 | 2,17 | 3,46 | 2,08 | 3,37 | 1,29 | 1,7E-01 | 1,6E-09 | 3,3E-04 | 9,5E-09 | 4,0E-04 | 5,8E-02 | 0,0E+00 | 1,5E-10 | 4,2E-06 | 5,0E-10 | 9,5E-06 | 3,1E-03 | 506815 | 507093 | 279 | + |
| Y11_RS02220 | Y11_04781 | tusE | sulfurtransferase TusE | 0,03 | -1,23 | -1,64 | -1,27 | -1,67 | -0,4 | 7,2E-01 | 7,5E-07 | 5,7E-06 | 1,8E-06 | 7,1E-06 | 3,7E-02 | 0,0E+00 | 2,3E-08 | 1,0E-07 | 4,2E-08 | 2,4E-07 | 0,0E+00 | 507121 | 507450 | 330 | - |
| Y11_RS02270 | Y11_04881 | yenR | LuxR family transcriptional regulator YenR | -0,31 | 0,58 | 1,3 | 0 | 0 | 0 | 4,0E-01 | 8,6E-02 | 4,4E-03 | 0,0E+00 | 0,0E+00 | 0,0E+00 | 0,0E+00 | 0,0E+00 | 5,0E-05 | 0,0E+00 | 0,0E+00 | 0,0E+00 | 516241 | 516975 | 735 | + |
| Y11_RS02275 | Y11_04891 | | acyl-homoserine-lactone synthase | -0,36 | -2,42 | -2,17 | -2,06 | -1,79 | 0,26 | 3,8E-02 | 1,3E-06 | 3,4E-05 | 4,3E-07 | 4,8E-05 | 3,4E-01 | 0,0E+00 | 3,8E-08 | 5,1E-07 | 1,2E-08 | 1,3E-06 | 0,0E+00 | 516968 | 517612 | 645 | - |
| Y11_RS02285 | Y11_04921 | | potassium/proton antiporter | -1,42 | 0 | 0 | 1,04 | 0,73 | 0 | 1,3E-04 | 0,0E+00 | 0,0E+00 | 5,0E-04 | 2,8E-04 | 0,0E+00 | 5,4E-06 | 0,0E+00 | 0,0E+00 | 8,5E-06 | 0,0E+00 | 0,0E+00 | 518954 | 520681 | 1728 | - |
| Y11_RS02295 | Y11_04941 | | hypothetical protein | -0,18 | 1,86 | 2,69 | 2,04 | 2,87 | 0,82 | 1,9E-02 | 2,8E-07 | 4,3E-08 | 8,1E-08 | 1,7E-08 | 1,0E-03 | 0,0E+00 | 9,7E-09 | 1,7E-09 | 2,8E-09 | 1,5E-09 | 0,0E+00 | 522721 | 522966 | 246 | - |
| Y11_RS02305 | Y11_04961 | | glucan biosynthesis protein G | -0,46 | 0,01 | 0,45 | 0,47 | 0,91 | 0,43 | 3,6E-06 | 7,5E-01 | 3,3E-03 | 8,5E-07 | 2,7E-05 | 3,5E-03 | 0,0E+00 | 0,0E+00 | 0,0E+00 | 0,0E+00 | 0,0E+00 | 0,0E+00 | 524649 | 526202 | 1554 | + |
| Y11_RS02310 | Y11_04971 | mdoH | glucans biosynthesis glucosyltransferase MdoH | -0,64 | -0,02 | 0,36 | 0,61 | 1 | 0,39 | 1,1E-05 | 7,6E-01 | 1,2E-03 | 1,2E-06 | 7,2E-08 | 1,7E-04 | 0,0E+00 | 0,0E+00 | 0,0E+00 | 0,0E+00 | 0,0E+00 | 0,0E+00 | 526195 | 528777 | 2583 | + |
| Y11_RS02320 | Y11_04991 | | peptide MFS transporter | -0,56 | 1,37 | 1,16 | 1,92 | 1,72 | -0,2 | 7,3E-02 | 2,3E-05 | 2,6E-02 | 1,0E-04 | 7,3E-03 | 6,5E-01 | 0,0E+00 | 5,3E-07 | 3,0E-04 | 1,8E-06 | 1,6E-04 | 0,0E+00 | 530426 | 531961 | 1536 | - |
| Y11_RS02335 | Y11_05031 | | RHS repeat protein | -0,53 | -0,76 | -0,99 | -0,23 | -0,46 | -0,23 | 1,0E-01 | 2,1E-02 | 2,3E-02 | 3,6E-01 | 1,9E-01 | 4,7E-01 | 0,0E+00 | 0,0E+00 | 0,0E+00 | 0,0E+00 | 0,0E+00 | 0,0E+00 | 535059 | 537962 | 2904 | - |
| Y11_RS02340 | Y11_05041 | | LpxL/LpxP family Kdo(2)-lipid IV(A) lauroyl/palmitoleoyl acyltransferasee | -0,48 | -0,09 | 0,33 | 0,39 | 0,81 | 0,42 | 6,0E-03 | 3,7E-01 | 5,4E-02 | 6,9E-03 | 8,3E-04 | 1,1E-02 | 0,0E+00 | 0,0E+00 | 0,0E+00 | 0,0E+00 | 0,0E+00 | 0,0E+00 | 538113 | 539033 | 921 | - |
| Y11_RS02375 | Y11_05111 | | rhodanese-related sulfurtransferase | 0,1 | -0,39 | -1,26 | -0,51 | -1,38 | -0,87 | 6,5E-01 | 1,6E-01 | 3,0E-03 | 6,3E-02 | 1,2E-03 | 2,6E-02 | 0,0E+00 | 0,0E+00 | 3,5E-05 | 0,0E+00 | 2,8E-05 | 0,0E+00 | 547119 | 548171 | 1053 | + |
| Y11_RS02380 | Y11_05121 | | AI-2E family transporter | 0,34 | 0,11 | 0,69 | -0,23 | 0,36 | 0,58 | 3,2E-01 | 7,6E-01 | 5,0E-02 | 5,3E-01 | 2,5E-01 | 1,2E-01 | 0,0E+00 | 0,0E+00 | 0,0E+00 | 0,0E+00 | 0,0E+00 | 0,0E+00 | 548272 | 549333 | 1062 | + |
| Y11_RS02390 | Y11_05151 | solA | N-methyl-L-tryptophan oxidase | 0,66 | -0,67 | -0,5 | -1,33 | -1,16 | 0,17 | 2,5E-05 | 2,6E-05 | 5,8E-04 | 5,8E-08 | 7,0E-07 | 7,5E-02 | 0,0E+00 | 0,0E+00 | 0,0E+00 | 2,1E-09 | 3,1E-08 | 0,0E+00 | 550687 | 551808 | 1122 | - |
| Y11_RS02400 | Y11_05171 | dinI | DNA damage-inducible protein I | 0,33 | -4,82 | -3,12 | -5,14 | -3,44 | 1,7 | 4,2E-02 | 1,1E-07 | 4,4E-05 | 4,2E-08 | 1,7E-05 | 5,2E-03 | 0,0E+00 | 4,4E-09 | 6,4E-07 | 1,6E-09 | 5,3E-07 | 3,1E-04 | 552689 | 552934 | 246 | - |
| Y11_RS02405 | Y11_05181 | pyrC | dihydroorotase | 0,59 | -0,24 | 0,27 | -0,83 | -0,32 | 0,51 | 1,8E-07 | 8,2E-04 | 2,8E-04 | 2,3E-07 | 2,4E-04 | 2,2E-05 | 0,0E+00 | 0,0E+00 | 0,0E+00 | 0,0E+00 | 0,0E+00 | 0,0E+00 | 553051 | 554097 | 1047 | - |
| Y11_RS02410 | Y11_05191 | | antibiotic biosynthesis monooxygenase | 0,08 | -0,5 | -0,31 | -0,58 | -0,39 | 0,19 | 1,6E-01 | 2,5E-06 | 1,9E-02 | 6,7E-06 | 8,5E-03 | 1,2E-01 | 0,0E+00 | 0,0E+00 | 0,0E+00 | 0,0E+00 | 0,0E+00 | 0,0E+00 | 554324 | 554611 | 288 | + |
| Y11_RS02415 | Y11_05201 | rne | ribonuclease E | 0,24 | 0,66 | 0,25 | 0,43 | 0,01 | -0,41 | 6,0E-05 | 2,8E-07 | 2,1E-03 | 1,4E-05 | 8,3E-01 | 2,2E-04 | 0,0E+00 | 0,0E+00 | 0,0E+00 | 0,0E+00 | 0,0E+00 | 0,0E+00 | 554681 | 558148 | 3468 | - |
| Y11_RS02420 | Y11_05211 | rluC | 23S rRNA pseudouridine(955/2504/2580) synthase RluC | 0,17 | -0,67 | -0,43 | -0,84 | -0,6 | 0,24 | 3,7E-03 | 3,4E-07 | 4,2E-03 | 1,5E-07 | 6,6E-04 | 6,2E-02 | 0,0E+00 | 0,0E+00 | 0,0E+00 | 0,0E+00 | 0,0E+00 | 0,0E+00 | 558726 | 559685 | 960 | + |
| Y11_RS02425 | Y11_05221 | | Maf-like protein | -0,29 | -0,35 | -0,02 | -0,06 | 0,27 | 0,33 | 7,1E-03 | 4,9E-03 | 9,2E-01 | 5,3E-01 | 2,3E-01 | 1,6E-01 | 0,0E+00 | 0,0E+00 | 0,0E+00 | 0,0E+00 | 0,0E+00 | 0,0E+00 | 559726 | 560307 | 582 | - |
| Y11_RS02430 | Y11_05231 | yceD | 23S rRNA accumulation protein YceD | 0,08 | 0,24 | 0,46 | 0,17 | 0,39 | 0,22 | 2,9E-01 | 1,4E-02 | 2,7E-04 | 8,1E-02 | 1,4E-03 | 3,8E-02 | 0,0E+00 | 0,0E+00 | 0,0E+00 | 0,0E+00 | 0,0E+00 | 0,0E+00 | 560449 | 560979 | 531 | + |
| Y11_RS02435 | Y11_05241 | rpmF | 50S ribosomal protein L32 | 0,1 | -0,09 | 0,06 | -0,2 | -0,04 | 0,16 | 2,9E-01 | 3,8E-01 | 7,0E-01 | 1,2E-02 | 7,7E-01 | 3,0E-01 | 0,0E+00 | 0,0E+00 | 0,0E+00 | 0,0E+00 | 0,0E+00 | 0,0E+00 | 560985 | 561152 | 168 | + |
| Y11_RS02440 | Y11_05251 | plsX | phosphate acyltransferase PlsX | 0,62 | 0,38 | 0,77 | -0,24 | 0,15 | 0,38 | 2,3E-03 | 4,5E-02 | 2,9E-03 | 1,3E-01 | 3,9E-01 | 6,4E-02 | 0,0E+00 | 0,0E+00 | 0,0E+00 | 0,0E+00 | 0,0E+00 | 0,0E+00 | 561189 | 562223 | 1035 | + |
| Y11_RS02445 | Y11_05261 | | ketoacyl-ACP synthase III | -0,71 | -0,37 | -0,1 | 0,34 | 0,62 | 0,28 | 1,0E-06 | 1,9E-05 | 1,5E-02 | 7,4E-04 | 5,2E-06 | 3,3E-04 | 0,0E+00 | 0,0E+00 | 0,0E+00 | 0,0E+00 | 0,0E+00 | 0,0E+00 | 562230 | 563183 | 954 | + |
| Y11_RS02450 | Y11_05271 | fabD | ACP S-malonyltransferase | -0,32 | -0,69 | -0,56 | -0,37 | -0,24 | 0,13 | 1,7E-02 | 2,6E-05 | 2,2E-03 | 1,3E-02 | 1,5E-01 | 3,6E-01 | 0,0E+00 | 0,0E+00 | 0,0E+00 | 0,0E+00 | 0,0E+00 | 0,0E+00 | 563238 | 564167 | 930 | + |
| Y11_RS02455 | Y11_05281 | fabG | 3-oxoacyl-ACP reductase FabG | 0,02 | -0,61 | -1,12 | -0,63 | -1,14 | -0,51 | 6,3E-01 | 9,9E-08 | 1,4E-05 | 1,6E-06 | 1,7E-05 | 3,3E-03 | 0,0E+00 | 0,0E+00 | 2,2E-07 | 0,0E+00 | 5,3E-07 | 0,0E+00 | 564181 | 564915 | 735 | + |
| Y11_RS02460 | Y11_05291 | acpP | acyl carrier protein | -0,15 | -0,19 | 0,39 | -0,04 | 0,54 | 0,57 | 1,5E-02 | 1,5E-02 | 1,8E-01 | 5,7E-01 | 7,3E-02 | 6,1E-02 | 0,0E+00 | 0,0E+00 | 0,0E+00 | 0,0E+00 | 0,0E+00 | 0,0E+00 | 565069 | 565305 | 237 | + |
| Y11_RS02465 | Y11_05301 | fabF | beta-ketoacyl-ACP synthase II | -0,12 | -1,02 | -1,2 | -0,9 | -1,09 | -0,19 | 6,6E-02 | 3,0E-06 | 2,0E-06 | 1,0E-05 | 5,4E-06 | 1,7E-01 | 0,0E+00 | 8,0E-08 | 4,1E-08 | 0,0E+00 | 1,9E-07 | 0,0E+00 | 565400 | 566638 | 1239 | + |
| Y11_RS02470 | Y11_05311 | pabC | aminodeoxychorismate lyase | -0,35 | 0,3 | 0,21 | 0,65 | 0,56 | -0,09 | 1,6E-02 | 8,1E-04 | 4,1E-02 | 6,9E-04 | 3,5E-03 | 3,9E-01 | 0,0E+00 | 0,0E+00 | 0,0E+00 | 0,0E+00 | 0,0E+00 | 0,0E+00 | 567053 | 567877 | 825 | + |
| Y11_RS02475 | Y11_05321 | mltG | endolytic transglycosylase MltG | -0,54 | -2,78 | -3,26 | -2,24 | -2,72 | -0,49 | 1,4E-02 | 9,6E-08 | 4,8E-08 | 2,3E-08 | 1,6E-08 | 9,2E-04 | 0,0E+00 | 4,0E-09 | 1,8E-09 | 9,8E-10 | 1,4E-09 | 0,0E+00 | 567939 | 568964 | 1026 | + |
| Y11_RS02480 | Y11_05331 | tmk | dTMP kinase | 0,18 | 0 | -0,1 | -0,18 | -0,28 | -0,1 | 5,4E-02 | 9,9E-01 | 7,5E-01 | 5,3E-02 | 3,6E-01 | 7,5E-01 | 0,0E+00 | 0,0E+00 | 0,0E+00 | 0,0E+00 | 0,0E+00 | 0,0E+00 | 568954 | 569592 | 639 | + |
| Y11_RS02485 | Y11_05341 | holB | DNA polymerase III subunit delta' | 0,02 | 0,23 | -0,49 | 0,21 | -0,51 | -0,72 | 9,1E-01 | 3,0E-01 | 1,6E-01 | 4,2E-01 | 1,8E-01 | 8,5E-02 | 0,0E+00 | 0,0E+00 | 0,0E+00 | 0,0E+00 | 0,0E+00 | 0,0E+00 | 569592 | 570599 | 1008 | + |
| Y11_RS02490 | Y11_05351 | | metal-dependent hydrolase | 0,28 | -0,82 | -1,58 | -1,1 | -1,86 | -0,76 | 4,6E-02 | 2,1E-05 | 6,2E-07 | 1,3E-05 | 6,5E-07 | 1,4E-04 | 0,0E+00 | 0,0E+00 | 1,5E-08 | 2,7E-07 | 2,9E-08 | 0,0E+00 | 570614 | 571423 | 810 | + |
| Y11_RS02495 | Y11_05361 | ptsG | PTS glucose transporter subunit IIBC | -0,16 | 0,57 | -1,64 | 0,73 | -1,48 | -2,21 | 1,8E-03 | 2,5E-05 | 4,0E-05 | 3,2E-06 | 8,0E-05 | 5,6E-06 | 0,0E+00 | 0,0E+00 | 5,8E-07 | 0,0E+00 | 2,1E-06 | 7,1E-07 | 571731 | 573164 | 1434 | + |
| Y11_RS02520 | Y11_05421 | | SgrR family transcriptional regulator | -0,3 | 0 | 0 | -0,25 | 15 | 0 | 7,8E-01 | 0,0E+00 | 0,0E+00 | 8,2E-01 | 0,0E+00 | 0,0E+00 | 0,0E+00 | 0,0E+00 | 0,0E+00 | 0,0E+00 | 0,0E+00 | 0,0E+00 | 579438 | 581228 | 1791 | - |
| Y11_RS02525 | Y11_05431 | | shikimate 5-dehydrogenase | 0,84 | 1,36 | 1,42 | 0,51 | 0,58 | 0,07 | 7,1E-04 | 2,7E-06 | 2,8E-03 | 2,5E-02 | 1,5E-01 | 8,5E-01 | 0,0E+00 | 7,2E-08 | 3,2E-05 | 0,0E+00 | 0,0E+00 | 0,0E+00 | 581376 | 582197 | 822 | + |
| Y11_RS02530 | Y11_05441 | hinT | purine nucleoside phosphoramidase | 0,1 | 1,27 | 1,34 | 1,17 | 1,24 | 0,08 | 2,8E-01 | 1,5E-07 | 1,8E-05 | 1,2E-06 | 4,6E-05 | 6,3E-01 | 0,0E+00 | 5,9E-09 | 2,8E-07 | 3,0E-08 | 1,3E-06 | 0,0E+00 | 582273 | 582626 | 354 | + |
| Y11_RS02540 | Y11_05461 | lpoB | penicillin-binding protein activator LpoB | 0,57 | 0,9 | 1,36 | 0,33 | 0,78 | 0,46 | 3,7E-04 | 1,7E-05 | 3,9E-06 | 1,1E-02 | 2,1E-04 | 5,9E-03 | 0,0E+00 | 0,0E+00 | 7,2E-08 | 0,0E+00 | 0,0E+00 | 0,0E+00 | 583123 | 583704 | 582 | + |
| Y11_RS02545 | Y11_05471 | thiK | thiamine kinase | 0,02 | -0,45 | -1,19 | -0,46 | -1,21 | -0,75 | 9,4E-01 | 6,0E-02 | 2,4E-03 | 4,7E-04 | 3,0E-04 | 6,0E-03 | 0,0E+00 | 0,0E+00 | 2,7E-05 | 0,0E+00 | 7,2E-06 | 0,0E+00 | 583685 | 584605 | 921 | + |
| Y11_RS02550 | Y11_05481 | nagZ | beta-N-acetylhexosaminidase | 0,24 | 0,29 | 0,06 | 0,05 | -0,17 | -0,23 | 5,3E-03 | 9,8E-03 | 5,0E-01 | 5,9E-01 | 1,1E-01 | 8,1E-02 | 0,0E+00 | 0,0E+00 | 0,0E+00 | 0,0E+00 | 0,0E+00 | 0,0E+00 | 584628 | 585653 | 1026 | + |
| Y11_RS02555 | Y11_05491 | ycfP | alpha/beta hydrolase YcfP | 0,68 | 1,21 | 0,63 | 0,53 | -0,05 | -0,59 | 5,2E-07 | 9,4E-08 | 4,3E-04 | 8,8E-05 | 6,4E-01 | 1,4E-03 | 0,0E+00 | 3,9E-09 | 0,0E+00 | 0,0E+00 | 0,0E+00 | 0,0E+00 | 585656 | 586198 | 543 | + |
| Y11_RS02560 | Y11_05501 | | NAD(P)/FAD-dependent oxidoreductase | 1,17 | 1,3 | 0 | 0,13 | -1,17 | -1,3 | 1,1E-06 | 5,3E-07 | 9,9E-01 | 2,9E-01 | 3,0E-05 | 1,4E-05 | 7,1E-08 | 1,7E-08 | 0,0E+00 | 0,0E+00 | 8,8E-07 | 1,6E-06 | 586577 | 587881 | 1305 | + |
| Y11_RS02565 | Y11_05511 | | glycine zipper 2TM domain-containing protein | 0 | -1,48 | -2,17 | -15 | -15 | -0,44 | 0,0E+00 | 2,3E-05 | 1,9E-06 | 0,0E+00 | 0,0E+00 | 7,1E-02 | 0,0E+00 | 5,2E-07 | 4,0E-08 | 0,0E+00 | 0,0E+00 | 0,0E+00 | 588262 | 588801 | 540 | + |
| Y11_RS02570 | Y11_05521 | mfd | transcription-repair coupling factor | -0,02 | 3,35 | 3,19 | 3,38 | 3,21 | -0,17 | 6,2E-01 | 6,8E-10 | 1,9E-09 | 8,8E-10 | 2,3E-09 | 2,7E-01 | 0,0E+00 | 8,2E-11 | 1,6E-10 | 7,9E-11 | 3,2E-10 | 0,0E+00 | 588940 | 592386 | 3447 | - |
| Y11_RS02575 | Y11_05531 | lolC | lipoprotein-releasing ABC transporter permease subunit LolC | -0,04 | -0,89 | -1,42 | -0,85 | -1,38 | -0,53 | 7,7E-01 | 4,0E-04 | 7,1E-06 | 2,1E-05 | 5,1E-08 | 1,0E-03 | 0,0E+00 | 0,0E+00 | 1,2E-07 | 0,0E+00 | 3,6E-09 | 0,0E+00 | 592648 | 593850 | 1203 | + |
| Y11_RS02580 | Y11_05541 | lolD | lipoprotein-releasing ABC transporter ATP-binding protein LolD | -0,01 | -0,55 | -0,55 | -0,53 | -0,54 | -0,01 | 7,3E-01 | 4,2E-05 | 8,4E-05 | 3,4E-05 | 7,4E-05 | 9,6E-01 | 0,0E+00 | 0,0E+00 | 0,0E+00 | 0,0E+00 | 0,0E+00 | 0,0E+00 | 593843 | 594547 | 705 | + |
| Y11_RS02585 | Y11_05551 | lolE | lipoprotein-releasing ABC transporter permease subunit LolE | -0,04 | -0,56 | -0,83 | -0,52 | -0,8 | -0,27 | 7,6E-01 | 1,9E-03 | 1,3E-04 | 1,6E-04 | 5,8E-06 | 1,6E-02 | 0,0E+00 | 0,0E+00 | 0,0E+00 | 0,0E+00 | 0,0E+00 | 0,0E+00 | 594559 | 595806 | 1248 | + |
| Y11_RS02590 | Y11_05561 | nagK | N-acetylglucosamine kinase | 0,09 | -0,65 | -0,81 | -0,74 | -0,9 | -0,16 | 4,9E-01 | 1,8E-04 | 7,5E-05 | 3,2E-05 | 1,8E-05 | 2,6E-02 | 0,0E+00 | 0,0E+00 | 0,0E+00 | 0,0E+00 | 0,0E+00 | 0,0E+00 | 596114 | 597028 | 915 | + |
| Y11_RS02595 | Y11_05571 | cobB | NAD-dependent protein deacylase | 0,46 | 0,17 | 0,57 | -0,29 | 0,11 | 0,4 | 3,6E-02 | 3,7E-01 | 3,5E-02 | 5,7E-02 | 5,7E-01 | 6,2E-02 | 0,0E+00 | 0,0E+00 | 0,0E+00 | 0,0E+00 | 0,0E+00 | 0,0E+00 | 597089 | 597925 | 837 | + |
| Y11_RS02600 | Y11_05581 | pepT | peptidase T | -0,06 | 0,04 | 0,38 | 0,1 | 0,43 | 0,34 | 7,3E-01 | 6,6E-01 | 1,2E-01 | 5,4E-01 | 1,2E-01 | 1,4E-01 | 0,0E+00 | 0,0E+00 | 0,0E+00 | 0,0E+00 | 0,0E+00 | 0,0E+00 | 598095 | 599330 | 1236 | + |
| Y11_RS02605 | Y11_05591 | | cupin domain-containing protein | -0,54 | -1,63 | -1,21 | -1,08 | -0,67 | 0,41 | 1,5E-04 | 6,2E-08 | 1,5E-06 | 1,0E-06 | 9,7E-05 | 2,9E-03 | 0,0E+00 | 2,7E-09 | 3,3E-08 | 2,6E-08 | 0,0E+00 | 0,0E+00 | 599543 | 600664 | 1122 | - |
| Y11_RS02610 | Y11_05601 | phoQ | two-component system sensor histidine kinase PhoQ | 0,39 | 0,69 | 1,55 | 0,3 | 1,15 | 0,85 | 1,3E-01 | 4,5E-03 | 2,2E-07 | 3,2E-01 | 1,5E-03 | 2,1E-03 | 0,0E+00 | 0,0E+00 | 6,4E-09 | 0,0E+00 | 3,4E-05 | 0,0E+00 | 600746 | 602203 | 1458 | - |
| Y11_RS02615 | Y11_05611 | phoP | two-component system response regulator PhoP | 0,14 | 0,5 | 1,16 | 0,35 | 1,02 | 0,67 | 9,2E-02 | 2,9E-04 | 5,8E-07 | 8,1E-03 | 8,1E-06 | 2,5E-04 | 0,0E+00 | 0,0E+00 | 1,4E-08 | 0,0E+00 | 2,7E-07 | 0,0E+00 | 602209 | 602880 | 672 | - |
| Y11_RS02620 | Y11_05621 | | outer membrane beta-barrel protein | 0,84 | -0,8 | 0,53 | -1,63 | -0,31 | 1,33 | 7,0E-06 | 4,4E-04 | 4,7E-03 | 7,6E-07 | 2,9E-02 | 3,6E-05 | 0,0E+00 | 0,0E+00 | 0,0E+00 | 1,9E-08 | 0,0E+00 | 3,4E-06 | 602952 | 603575 | 624 | - |
| Y11_RS02625 | Y11_05631 | purB | adenylosuccinate lyase | -0,28 | -1,13 | -1,17 | -0,85 | -0,89 | -0,04 | 9,1E-04 | 5,9E-11 | 3,1E-06 | 2,0E-07 | 4,6E-05 | 7,1E-01 | 0,0E+00 | 1,6E-11 | 6,0E-08 | 0,0E+00 | 0,0E+00 | 0,0E+00 | 603730 | 605100 | 1371 | - |
| Y11_RS02630 | Y11_05641 | hflD | high frequency lysogenization protein HflD | -0,37 | -1,17 | -0,74 | -0,8 | -0,38 | 0,43 | 5,4E-03 | 3,0E-07 | 5,1E-03 | 7,4E-06 | 9,1E-02 | 5,2E-02 | 0,0E+00 | 1,0E-08 | 0,0E+00 | 0,0E+00 | 0,0E+00 | 0,0E+00 | 605126 | 605752 | 627 | - |
| Y11_RS02635 | Y11_05651 | mnmA | tRNA 2-thiouridine(34) synthase MnmA | -0,31 | -0,78 | -0,81 | -0,48 | -0,5 | -0,03 | 3,9E-04 | 1,3E-05 | 3,6E-07 | 7,4E-04 | 5,1E-05 | 7,6E-01 | 0,0E+00 | 0,0E+00 | 0,0E+00 | 0,0E+00 | 0,0E+00 | 0,0E+00 | 605883 | 607004 | 1122 | - |
| Y11_RS02640 | Y11_05661 | | NUDIX hydrolase | 0,06 | 0,4 | 0,58 | 0,33 | 0,52 | 0,19 | 5,2E-01 | 5,4E-03 | 2,1E-04 | 5,9E-03 | 9,9E-05 | 6,3E-02 | 0,0E+00 | 0,0E+00 | 0,0E+00 | 0,0E+00 | 0,0E+00 | 0,0E+00 | 607190 | 607636 | 447 | - |
| Y11_RS02645 | Y11_05671 | rluE | 23S rRNA pseudouridine(2457) synthase RluE | -0,26 | -0,03 | -0,16 | 0,22 | 0,1 | -0,12 | 3,6E-02 | 7,9E-01 | 4,0E-01 | 2,7E-02 | 5,3E-01 | 4,8E-01 | 0,0E+00 | 0,0E+00 | 0,0E+00 | 0,0E+00 | 0,0E+00 | 0,0E+00 | 607629 | 608255 | 627 | - |
| Y11_RS02650 | Y11_05681 | icd | NADP-dependent isocitrate dehydrogenase | 0,17 | -0,04 | 0,73 | -0,21 | 0,57 | 0,78 | 1,8E-02 | 4,0E-01 | 2,6E-05 | 5,3E-03 | 2,6E-04 | 1,6E-05 | 0,0E+00 | 0,0E+00 | 0,0E+00 | 0,0E+00 | 0,0E+00 | 0,0E+00 | 608386 | 609639 | 1254 | + |
| Y11_RS02685 | Y11_05741 | | DUF2219 family protein | -1,11 | 1,95 | 2,46 | 3,06 | 3,57 | 0,51 | 1,2E-06 | 3,9E-10 | 4,8E-11 | 5,0E-10 | 1,3E-10 | 1,5E-05 | 7,6E-08 | 5,4E-11 | 1,5E-11 | 5,2E-11 | 5,8E-11 | 0,0E+00 | 612760 | 613719 | 960 | - |
| Y11_RS02695 | Y11_05761 | | glycosyltransferase family 2 protein | 0,83 | 0,97 | 1,13 | 0,14 | 0,3 | 0,16 | 1,0E-05 | 2,8E-06 | 8,7E-05 | 1,6E-01 | 9,5E-02 | 3,4E-01 | 0,0E+00 | 0,0E+00 | 1,2E-06 | 0,0E+00 | 0,0E+00 | 0,0E+00 | 614936 | 615796 | 861 | + |
| Y11_RS02700 | Y11_05771 | waaL-os | outer core oligosaccharide ligase WaaL-os | 0,09 | -0,04 | 0,13 | -0,13 | 0,04 | 0,17 | 4,2E-01 | 6,8E-01 | 1,6E-01 | 2,3E-01 | 6,5E-01 | 5,9E-02 | 0,0E+00 | 0,0E+00 | 0,0E+00 | 0,0E+00 | 0,0E+00 | 0,0E+00 | 615948 | 617168 | 1221 | + |

| | | | | | | | | | | | | | | | | | | | | | | | | | |
|---|---|---|---|---|---|---|---|---|---|---|---|---|---|---|---|---|---|---|---|---|---|---|---|---|---|
| Y11_RS02715 | Y11_05811 | | LacI family DNA-binding transcriptional regulator | 0,77 | 1,87 | 1,69 | 1,1 | 0,92 | -0,18 | 8,2E-06 | 7,4E-09 | 5,2E-06 | 3,9E-07 | 3,8E-04 | 2,8E-01 | 0,0E+00 | 5,0E-10 | 9,3E-08 | 1,1E-08 | 0,0E+00 | 0,0E+00 | 620394 | 621389 | 996 | - |
| Y11_RS02720 | Y11_05821 | | hypothetical protein | -0,37 | -0,06 | 0,17 | 0,31 | 0,54 | 0,23 | 1,8E-02 | 2,9E-01 | 4,7E-01 | 3,8E-02 | 6,1E-02 | 3,3E-01 | 0,0E+00 | 0,0E+00 | 0,0E+00 | 0,0E+00 | 0,0E+00 | 0,0E+00 | 621520 | 621780 | 261 | - |
| Y11_RS02745 | Y11_05871 | | bifunctional O-acetylhomoserine aminocarboxypropyltransferase/cysteine synthase | 0 | -15 | 0 | -15 | 0 | 15 | 0,0E+00 | 0,0E+00 | 0,0E+00 | 0,0E+00 | 0,0E+00 | 0,0E+00 | 0,0E+00 | 0,0E+00 | 0,0E+00 | 0,0E+00 | 0,0E+00 | 0,0E+00 | 626850 | 628127 | 1278 | + |
| Y11_RS02750 | Y11_05881 | | SDR family oxidoreductase | 0,81 | 2,19 | 2,7 | 1,38 | 1,89 | 0,51 | 4,5E-08 | 3,2E-11 | 4,2E-09 | 3,0E-09 | 8,3E-08 | 1,2E-03 | 0,0E+00 | 1,1E-11 | 2,8E-10 | 2,0E-10 | 5,3E-09 | 0,0E+00 | 628190 | 629047 | 858 | - |
| Y11_RS02755 | Y11_05891 | | helix-turn-helix transcriptional regulator | -0,01 | 0,67 | -0,32 | 0,68 | -0,31 | -0,99 | 9,5E-01 | 9,4E-03 | 1,3E-01 | 2,2E-02 | 2,3E-01 | 4,2E-03 | 0,0E+00 | 0,0E+00 | 0,0E+00 | 0,0E+00 | 0,0E+00 | 0,0E+00 | 629317 | 629721 | 405 | + |
| Y11_RS02760 | Y11_05901 | | YciI family protein | 1,14 | 1,36 | 1,63 | 0,22 | 0,49 | 0,27 | 6,8E-06 | 1,8E-07 | 3,0E-08 | 1,3E-01 | 5,3E-03 | 3,3E-02 | 3,6E-07 | 6,7E-09 | 1,2E-09 | 0,0E+00 | 0,0E+00 | 0,0E+00 | 629785 | 630069 | 285 | - |
| Y11_RS02785 | Y11_05981 | hdeB | acid-activated periplasmic chaperone HdeB | 1,06 | 2,46 | 3,45 | 1,4 | 2,39 | 0,99 | 1,4E-03 | 1,0E-04 | 2,0E-06 | 2,2E-03 | 1,0E-05 | 2,6E-02 | 5,1E-05 | 2,2E-06 | 4,2E-08 | 3,6E-05 | 3,4E-07 | 0,0E+00 | 633835 | 634146 | 312 | - |
| Y11_RS02795 | Y11_06001 | | sensor domain-containing diguanylate cyclase | 0,44 | 1,32 | 15 | 0,82 | 15 | 0 | 2,4E-02 | 8,2E-05 | 0,0E+00 | 3,9E-03 | 0,0E+00 | 0,0E+00 | 0,0E+00 | 1,8E-06 | 0,0E+00 | 0,0E+00 | 0,0E+00 | 0,0E+00 | 634930 | 636333 | 1404 | + |
| Y11_RS02810 | Y11_06021 | | serine protein kinase RIO | 0,15 | -1,53 | -1,84 | -1,67 | -1,99 | -0,31 | 2,8E-01 | 1,8E-07 | 2,0E-06 | 8,6E-08 | 1,1E-06 | 3,6E-02 | 0,0E+00 | 6,7E-09 | 4,2E-08 | 2,9E-09 | 4,6E-08 | 0,0E+00 | 638098 | 638946 | 849 | - |
| Y11_RS02815 | Y11_06031 | | ABC transporter ATP-binding protein/permease | 0,51 | -0,39 | -1,24 | -0,9 | -1,75 | -0,84 | 2,2E-04 | 4,6E-03 | 3,1E-04 | 3,4E-06 | 2,0E-05 | 3,4E-03 | 0,0E+00 | 0,0E+00 | 4,0E-06 | 0,0E+00 | 6,0E-07 | 0,0E+00 | 639049 | 640767 | 1719 | - |
| Y11_RS02820 | Y11_06041 | | GNAT family N-acetyltransferase | 0 | -3,32 | -4,17 | -15 | -15 | -0,76 | 0,0E+00 | 2,8E-08 | 4,7E-08 | 0,0E+00 | 0,0E+00 | 6,4E-03 | 0,0E+00 | 1,4E-09 | 1,8E-09 | 0,0E+00 | 0,0E+00 | 0,0E+00 | 640967 | 641386 | 420 | + |
| Y11_RS02825 | Y11_06061 | | amino acid permease | 0,37 | 1,44 | 15 | 1,08 | 0 | 15 | 1,6E-01 | 1,8E-03 | 0,0E+00 | 9,2E-03 | 0,0E+00 | 0,0E+00 | 0,0E+00 | 3,5E-05 | 0,0E+00 | 1,5E-04 | 0,0E+00 | 0,0E+00 | 641654 | 643156 | 1503 | + |
| Y11_RS02845 | Y11_06101 | | DUF2766 domain-containing protein | -0,69 | 0,52 | -0,09 | 1,21 | 0,6 | -0,61 | 2,7E-06 | 2,2E-04 | 3,0E-01 | 5,6E-07 | 8,7E-05 | 2,9E-04 | 0,0E+00 | 0,0E+00 | 0,0E+00 | 1,5E-08 | 0,0E+00 | 0,0E+00 | 644869 | 645108 | 240 | + |
| Y11_RS02850 | Y11_06111 | | membrane protein | 0,19 | 1,54 | 2,54 | 1,35 | 2,35 | 1 | 6,4E-02 | 6,2E-08 | 3,8E-03 | 4,4E-08 | 5,7E-03 | 1,5E-01 | 0,0E+00 | 2,7E-09 | 4,4E-05 | 1,7E-09 | 1,2E-04 | 7,7E-03 | 645201 | 645434 | 234 | - |
| Y11_RS02855 | Y11_06131 | | YebO family protein | 0 | -15 | 0 | 1,64 | 0 | -0,37 | 0,0E+00 | 0,0E+00 | 0,0E+00 | 1,5E-03 | 0,0E+00 | 9,6E-02 | 0,0E+00 | 0,0E+00 | 0,0E+00 | 2,5E-05 | 0,0E+00 | 0,0E+00 | 645973 | 646248 | 276 | + |
| Y11_RS02875 | Y11_06171 | pagP | lipid IV(A) palmitoyltransferase PagP | 15 | -0,89 | -1,1 | -15 | 0 | -0,18 | 0,0E+00 | 3,5E-02 | 2,4E-03 | 0,0E+00 | 0,0E+00 | 6,0E-01 | 0,0E+00 | 0,0E+00 | 2,7E-05 | 0,0E+00 | 0,0E+00 | 0,0E+00 | 648925 | 649539 | 615 | - |
| Y11_RS02885 | | cspE | transcription antiterminator/RNA stability regulator CspE | 0,28 | -0,85 | -0,93 | -1,12 | -1,2 | -0,08 | 2,2E-03 | 3,8E-07 | 2,9E-02 | 2,0E-07 | 8,9E-03 | 8,2E-01 | 0,0E+00 | 0,0E+00 | 0,0E+00 | 6,2E-09 | 1,9E-04 | 0,0E+00 | 650696 | 650905 | 210 | + |
| Y11_RS02890 | Y11_06191 | pgeF | peptidoglycan editing factor PgeF | 0 | 0,27 | 0,23 | 0,27 | 0,23 | -0,04 | 9,8E-01 | 3,1E-02 | 1,6E-01 | 3,6E-02 | 1,7E-01 | 7,8E-01 | 0,0E+00 | 0,0E+00 | 0,0E+00 | 0,0E+00 | 0,0E+00 | 0,0E+00 | 650983 | 651702 | 720 | - |
| Y11_RS02900 | Y11_06221 | rlmA | 23S rRNA (guanine(745)-N(1))-methyltransferase | 0,13 | 0,24 | 0,43 | 0,11 | 0,3 | 0,18 | 2,8E-01 | 8,9E-02 | 6,3E-02 | 3,0E-01 | 1,5E-01 | 3,7E-01 | 0,0E+00 | 0,0E+00 | 0,0E+00 | 0,0E+00 | 0,0E+00 | 0,0E+00 | 653732 | 654592 | 861 | + |
| Y11_RS02915 | Y11_06251 | | TonB-dependent receptor | 1,51 | -0,54 | -1,43 | -2,04 | -2,94 | -0,9 | 5,1E-10 | 8,7E-07 | 2,2E-07 | 2,1E-10 | 1,3E-09 | 1,1E-05 | 1,3E-10 | 0,0E+00 | 6,2E-09 | 2,7E-11 | 2,1E-10 | 0,0E+00 | 657866 | 660145 | 2280 | + |
| Y11_RS02930 | Y11_06281 | | PTS mannose transporter subunit IID | -0,87 | 2,15 | 1,33 | 3,01 | 2,2 | -0,81 | 1,4E-04 | 1,2E-09 | 5,5E-08 | 2,0E-08 | 2,4E-07 | 1,2E-05 | 0,0E+00 | 1,2E-10 | 2,0E-09 | 8,8E-10 | 1,3E-08 | 0,0E+00 | 662010 | 662873 | 864 | - |
| Y11_RS02935 | Y11_06291 | | PTS mannose/fructose/sorbose transporter subunit IIC | -0,53 | 2,33 | 1,76 | 2,87 | 2,29 | -0,57 | 6,0E-02 | 7,0E-06 | 5,2E-05 | 2,2E-07 | 1,1E-06 | 5,4E-03 | 0,0E+00 | 1,7E-07 | 7,5E-07 | 6,6E-09 | 4,6E-08 | 0,0E+00 | 662889 | 663686 | 798 | - |
| Y11_RS02940 | Y11_06301 | manX | PTS mannose transporter subunit IIAB | -0,7 | 1,97 | 1,15 | 2,67 | 1,85 | -0,81 | 3,9E-05 | 1,2E-08 | 1,2E-05 | 5,0E-09 | 7,1E-07 | 2,7E-04 | 0,0E+00 | 6,9E-10 | 1,9E-07 | 2,9E-10 | 3,1E-08 | 0,0E+00 | 663904 | 664869 | 966 | - |
| Y11_RS02945 | Y11_06321 | | TerC family protein | -0,03 | 1,33 | 1,2 | 1,36 | 1,22 | -0,13 | 7,7E-01 | 2,7E-06 | 6,9E-07 | 2,4E-06 | 6,4E-07 | 2,5E-01 | 0,0E+00 | 7,2E-08 | 1,7E-08 | 5,5E-08 | 2,9E-08 | 0,0E+00 | 665639 | 667183 | 1545 | + |
| Y11_RS02950 | Y11_06341 | | L-serine ammonia-lyase | -0,28 | -1,47 | -0,86 | -1,19 | -0,58 | 0,61 | 2,2E-03 | 6,2E-06 | 2,0E-04 | 1,9E-05 | 1,6E-03 | 8,7E-03 | 0,0E+00 | 1,5E-07 | 0,0E+00 | 3,7E-07 | 0,0E+00 | 0,0E+00 | 668286 | 669650 | 1365 | - |
| Y11_RS02960 | Y11_06371 | pabB | aminodeoxychorismate synthase component 1 | -0,88 | -0,07 | -0,31 | 0,82 | 0,57 | -0,24 | 3,8E-06 | 4,5E-01 | 4,9E-03 | 1,1E-05 | 9,8E-05 | 2,2E-02 | 0,0E+00 | 0,0E+00 | 0,0E+00 | 0,0E+00 | 0,0E+00 | 0,0E+00 | 670703 | 672073 | 1371 | - |
| Y11_RS02965 | Y11_06381 | | YoaH family protein | -0,28 | -0,19 | -0,37 | 0,09 | -0,09 | -0,18 | 1,3E-01 | 3,9E-01 | 1,6E-01 | 6,6E-01 | 7,2E-01 | 5,1E-01 | 0,0E+00 | 0,0E+00 | 0,0E+00 | 0,0E+00 | 0,0E+00 | 0,0E+00 | 672256 | 672513 | 258 | + |
| Y11_RS02970 | Y11_06391 | purT | formate-dependent phosphoribosylglycinamide formyltransferase | 0,59 | 2,71 | 3,07 | 2,12 | 2,48 | 0,36 | 6,2E-07 | 1,1E-12 | 1,2E-10 | 4,5E-11 | 1,1E-09 | 1,5E-03 | 0,0E+00 | 1,7E-12 | 2,6E-11 | 1,2E-11 | 1,8E-10 | 0,0E+00 | 672581 | 673762 | 1182 | - |
| Y11_RS02975 | Y11_06401 | dbpA | ATP-dependent RNA helicase DbpA | -0,56 | -0,99 | -1,11 | -0,43 | -0,55 | -0,12 | 8,4E-03 | 2,3E-04 | 4,7E-05 | 3,3E-02 | 6,6E-03 | 4,4E-01 | 0,0E+00 | 0,0E+00 | 6,9E-07 | 0,0E+00 | 0,0E+00 | 0,0E+00 | 674016 | 675398 | 1383 | + |
| Y11_RS02980 | Y11_06411 | | YebG family protein | 0,82 | -3,09 | -3,7 | -3,91 | -4,52 | -0,6 | 1,5E-02 | 4,5E-07 | 6,9E-05 | 1,1E-07 | 1,8E-05 | 2,4E-01 | 0,0E+00 | 1,5E-08 | 9,8E-07 | 3,8E-09 | 5,4E-07 | 0,0E+00 | 675494 | 675814 | 321 | + |
| Y11_RS02985 | Y11_06421 | yqfB | N(4)-acetylcytidine aminohydrolase | -0,1 | -0,58 | -0,03 | -0,48 | 0,07 | 0,55 | 1,0E-01 | 1,3E-04 | 8,4E-01 | 1,6E-04 | 6,0E-01 | 5,5E-03 | 0,0E+00 | 0,0E+00 | 0,0E+00 | 0,0E+00 | 0,0E+00 | 0,0E+00 | 675854 | 676162 | 309 | + |
| Y11_RS02990 | Y11_06431 | | protein YebF | 0,27 | 3,04 | 4,03 | 2,77 | 3,76 | 0,99 | 1,4E-02 | 5,7E-10 | 6,1E-08 | 5,2E-11 | 7,2E-08 | 1,2E-03 | 0,0E+00 | 7,3E-11 | 2,2E-09 | 1,2E-11 | 4,7E-09 | 0,0E+00 | 676268 | 676624 | 357 | + |
| Y11_RS02995 | Y11_06441 | | prolyl oligopeptidase family serine peptidase | 0,12 | -0,74 | -0,71 | -0,87 | -0,83 | 0,04 | 9,1E-02 | 2,0E-07 | 1,3E-05 | 6,0E-07 | 1,0E-05 | 6,3E-01 | 0,0E+00 | 0,0E+00 | 0,0E+00 | 0,0E+00 | 0,0E+00 | 0,0E+00 | 676814 | 678868 | 2055 | + |
| Y11_RS03010 | Y11_06481 | | DNA polymerase III subunit theta | -0,07 | -0,51 | -1,32 | -0,44 | -1,25 | -0,81 | 7,2E-01 | 2,5E-03 | 8,7E-06 | 3,3E-02 | 1,3E-04 | 1,0E-04 | 0,0E+00 | 0,0E+00 | 1,5E-07 | 0,0E+00 | 3,3E-06 | 0,0E+00 | 681058 | 681291 | 234 | - |
| Y11_RS03015 | Y11_06491 | ftnA | non-heme ferritin | -0,27 | 1,46 | 2,04 | 1,73 | 2,31 | 0,58 | 4,4E-02 | 1,1E-07 | 3,3E-06 | 2,9E-07 | 2,3E-06 | 1,2E-02 | 0,0E+00 | 4,6E-09 | 6,3E-08 | 8,4E-09 | 8,7E-08 | 0,0E+00 | 681674 | 682183 | 510 | + |
| Y11_RS03020 | Y11_06511 | yobA | CopC domain-containing protein YobA | 0 | -15 | -15 | -15 | -15 | 0,08 | 0,0E+00 | 0,0E+00 | 0,0E+00 | 0,0E+00 | 0,0E+00 | 8,5E-01 | 0,0E+00 | 0,0E+00 | 0,0E+00 | 0,0E+00 | 0,0E+00 | 0,0E+00 | 682574 | 682960 | 387 | + |
| Y11_RS03030 | Y11_06531 | | YebY family protein | 0,72 | 0,73 | 0,63 | 0,01 | -0,09 | -0,1 | 2,8E-07 | 2,1E-04 | 7,7E-04 | 9,3E-01 | 4,5E-01 | 5,3E-01 | 0,0E+00 | 0,0E+00 | 0,0E+00 | 0,0E+00 | 0,0E+00 | 0,0E+00 | 683943 | 684284 | 342 | + |
| Y11_RS03040 | Y11_06551 | | hypothetical protein | 0,25 | 2,05 | 15 | 0 | 0 | 0 | 3,2E-02 | 1,1E-07 | 0,0E+00 | 0,0E+00 | 0,0E+00 | 0,0E+00 | 0,0E+00 | 4,4E-09 | 0,0E+00 | 0,0E+00 | 0,0E+00 | 0,0E+00 | 685921 | 686736 | 816 | + |
| Y11_RS03085 | Y11_06651 | | class I SAM-dependent methyltransferase | 2,15 | 0,06 | 0,72 | -2,09 | -1,43 | 0,66 | 1,2E-08 | 5,0E-01 | 1,7E-04 | 1,3E-10 | 1,2E-07 | 2,6E-05 | 1,5E-09 | 0,0E+00 | 0,0E+00 | 2,1E-11 | 6,9E-09 | 0,0E+00 | 696178 | 696834 | 657 | + |
| Y11_RS03100 | Y11_06681 | | Cof-type HAD-IIB family hydrolase | 0,81 | 0,64 | 0,88 | -0,17 | 0,08 | 0,24 | 1,5E-06 | 3,0E-06 | 3,4E-05 | 4,7E-03 | 4,7E-01 | 3,4E-02 | 0,0E+00 | 0,0E+00 | 0,0E+00 | 0,0E+00 | 0,0E+00 | 0,0E+00 | 699945 | 700763 | 819 | - |
| Y11_RS03110 | Y11_06701 | | ROK family protein | 1,16 | 1,49 | 2 | 0,33 | 0,85 | 0,52 | 1,3E-04 | 1,3E-06 | 7,8E-07 | 7,1E-02 | 1,7E-03 | 5,4E-03 | 5,4E-06 | 3,7E-08 | 1,9E-08 | 0,0E+00 | 0,0E+00 | 0,0E+00 | 702353 | 703378 | 1026 | - |
| Y11_RS03115 | Y11_06711 | | DUF488 family protein | 0,63 | 1,23 | 2,56 | 0,6 | 1,94 | 1,33 | 3,7E-05 | 2,3E-06 | 2,6E-07 | 6,6E-05 | 1,1E-06 | 3,9E-05 | 0,0E+00 | 6,3E-08 | 7,2E-09 | 0,0E+00 | 4,6E-08 | 3,5E-06 | 703664 | 704017 | 354 | + |
| Y11_RS03130 | Y11_06741 | dcuR | two-component system response regulator DcuR | 0,66 | -1,11 | -1,08 | -1,77 | -1,74 | 0,03 | 3,2E-03 | 2,9E-07 | 1,9E-05 | 3,2E-06 | 1,2E-05 | 8,0E-01 | 0,0E+00 | 9,9E-09 | 3,0E-07 | 7,1E-08 | 4,0E-07 | 0,0E+00 | 708383 | 709102 | 720 | - |
| Y11_RS03140 | Y11_06761 | | DeoR/GlpR family DNA-binding transcription regulator | -0,07 | 0,33 | 0,6 | 0,4 | 0,67 | 0,27 | 5,8E-01 | 1,9E-04 | 1,7E-03 | 1,2E-02 | 4,6E-03 | 7,4E-02 | 0,0E+00 | 0,0E+00 | 0,0E+00 | 0,0E+00 | 0,0E+00 | 0,0E+00 | 711016 | 711795 | 780 | - |
| Y11_RS03160 | Y11_06811 | glsA | glutaminase A | 1,39 | 0,24 | 0,96 | -1,15 | -0,43 | 0,72 | 9,9E-07 | 3,5E-02 | 1,9E-04 | 1,3E-05 | 3,3E-02 | 2,0E-03 | 6,3E-08 | 0,0E+00 | 0,0E+00 | 2,6E-07 | 0,0E+00 | 0,0E+00 | 715547 | 716554 | 1008 | + |
| Y11_RS03165 | Y11_06821 | | MFS transporter | 1,2 | 15 | 15 | 15 | 15 | 0 | 4,9E-05 | 0,0E+00 | 0,0E+00 | 0,0E+00 | 0,0E+00 | 0,0E+00 | 2,2E-06 | 0,0E+00 | 0,0E+00 | 0,0E+00 | 0,0E+00 | 0,0E+00 | 716666 | 718306 | 1641 | + |
| Y11_RS03170 | Y11_06831 | | hypothetical protein | 0,58 | 0,23 | -0,57 | -0,35 | -1,15 | -0,8 | 4,8E-02 | 3,9E-01 | 7,3E-02 | 2,6E-01 | 6,5E-03 | 3,5E-02 | 0,0E+00 | 0,0E+00 | 0,0E+00 | 0,0E+00 | 1,4E-04 | 0,0E+00 | 718423 | 718941 | 519 | - |
| Y11_RS03180 | Y11_06851 | | iron transporter | 15 | 0,15 | -0,31 | -15 | -15 | -0,46 | 0,0E+00 | 5,9E-01 | 4,7E-01 | 0,0E+00 | 0,0E+00 | 2,9E-01 | 0,0E+00 | 0,0E+00 | 0,0E+00 | 0,0E+00 | 0,0E+00 | 0,0E+00 | 721132 | 721659 | 528 | + |
| Y11_RS03200 | Y11_06901 | | ABC transporter ATP-binding protein | 0,07 | -0,39 | -0,19 | -0,46 | -0,26 | 0,2 | 6,1E-01 | 1,4E-02 | 3,1E-01 | 9,4E-03 | 2,0E-01 | 3,0E-01 | 0,0E+00 | 0,0E+00 | 0,0E+00 | 0,0E+00 | 0,0E+00 | 0,0E+00 | 725643 | 726356 | 714 | + |
| Y11_RS03205 | Y11_06911 | | TlpA family protein disulfide reductase | 0 | -15 | -15 | -15 | -15 | -0,25 | 0,0E+00 | 0,0E+00 | 0,0E+00 | 0,0E+00 | 0,0E+00 | 6,6E-01 | 0,0E+00 | 0,0E+00 | 0,0E+00 | 0,0E+00 | 0,0E+00 | 0,0E+00 | 726346 | 726834 | 489 | + |
| Y11_RS03210 | Y11_06921 | | cytochrome c | 0,45 | 0,35 | 0,54 | -0,1 | 0,09 | 0,19 | 5,4E-02 | 1,3E-01 | 9,5E-02 | 4,0E-01 | 7,1E-01 | 4,4E-01 | 0,0E+00 | 0,0E+00 | 0,0E+00 | 0,0E+00 | 0,0E+00 | 0,0E+00 | 726834 | 727148 | 315 | + |
| Y11_RS03215 | Y11_06931 | tehB | SAM-dependent methyltransferase TehB | -0,06 | -1,64 | -1,87 | -1,57 | -1,8 | -0,23 | 3,5E-01 | 1,1E-08 | 7,2E-10 | 7,3E-08 | 8,7E-09 | 2,0E-02 | 0,0E+00 | 6,7E-10 | 8,2E-11 | 2,6E-09 | 8,6E-10 | 0,0E+00 | 727305 | 728174 | 870 | + |
| Y11_RS03220 | Y11_06941 | | LysR family transcriptional regulator | 0,05 | 0 | 0 | 15 | -0,09 | 0 | 8,1E-01 | 0,0E+00 | 0,0E+00 | 0,0E+00 | 6,6E-01 | 0,0E+00 | 0,0E+00 | 0,0E+00 | 0,0E+00 | 0,0E+00 | 0,0E+00 | 0,0E+00 | 728225 | 729157 | 933 | - |
| Y11_RS03225 | Y11_06951 | | hypothetical protein | 0 | -15 | -15 | -0,92 | -0,04 | 0,95 | 0,0E+00 | 0,0E+00 | 0,0E+00 | 4,9E-03 | 8,5E-01 | 1,3E-02 | 0,0E+00 | 0,0E+00 | 0,0E+00 | 0,0E+00 | 0,0E+00 | 0,0E+00 | 729247 | 729873 | 627 | + |
| Y11_RS03235 | Y11_06971 | | M24 family metallopeptidase | 0 | -3,18 | -2,04 | -15 | -15 | 1,17 | 0,0E+00 | 1,1E-06 | 6,2E-05 | 0,0E+00 | 0,0E+00 | 6,6E-03 | 0,0E+00 | 3,2E-08 | 8,8E-07 | 0,0E+00 | 0,0E+00 | 3,9E-04 | 730852 | 732069 | 1218 | + |
| Y11_RS03245 | Y11_07001 | pgaB | poly-beta-1,6-N-acetyl-D-glucosamine N-deacetylase PgaB | -0,18 | -0,04 | 0,6 | 0,14 | 0,78 | 0,64 | 5,8E-01 | 9,1E-01 | 1,2E-01 | 4,6E-01 | 3,4E-03 | 2,4E-02 | 0,0E+00 | 0,0E+00 | 0,0E+00 | 0,0E+00 | 0,0E+00 | 0,0E+00 | 735203 | 737227 | 2025 | + |
| Y11_RS03270 | Y11_07051 | phoH | phosphate starvation-inducible protein PhoH | -1,64 | 1,81 | 2,36 | 3,45 | 4,01 | 0,55 | 2,1E-07 | 2,7E-07 | 3,4E-07 | 1,0E-09 | 4,2E-09 | 8,2E-03 | 1,7E-08 | 9,4E-09 | 9,2E-09 | 8,8E-11 | 4,9E-10 | 0,0E+00 | 740467 | 741255 | 789 | + |
| Y11_RS03280 | Y11_07081 | argS | arginine--tRNA ligase | -0,14 | -0,85 | -0,9 | -0,71 | -0,76 | -0,05 | 1,8E-02 | 2,2E-10 | 5,1E-05 | 4,9E-07 | 2,6E-04 | 6,5E-01 | 0,0E+00 | 0,0E+00 | 0,0E+00 | 0,0E+00 | 0,0E+00 | 0,0E+00 | 742957 | 744687 | 1731 | - |
| Y11_RS03285 | Y11_07091 | | VOC family protein | 0,48 | -1,59 | -0,84 | -2,08 | -1,32 | 0,76 | 1,0E-04 | 8,3E-09 | 4,1E-06 | 1,5E-10 | 3,4E-08 | 1,7E-06 | 0,0E+00 | 5,3E-10 | 0,0E+00 | 2,3E-11 | 2,6E-09 | 0,0E+00 | 745055 | 745615 | 561 | + |
| Y11_RS03290 | Y11_07101 | cutC | copper homeostasis protein CutC | 0,95 | -1,55 | -1,35 | -2,5 | -2,3 | 0,2 | 6,3E-07 | 7,5E-08 | 7,4E-05 | 7,9E-10 | 1,2E-06 | 3,1E-01 | 0,0E+00 | 3,2E-09 | 1,0E-06 | 7,4E-11 | 5,0E-08 | 0,0E+00 | 745857 | 746621 | 765 | + |
| Y11_RS03310 | Y11_07181 | | mechanosensitive ion channel family protein | 0,36 | 0,58 | 0,98 | 0,22 | 0,62 | 0,4 | 8,5E-02 | 1,1E-02 | 5,4E-04 | 1,4E-01 | 1,6E-03 | 9,9E-03 | 0,0E+00 | 0,0E+00 | 0,0E+00 | 0,0E+00 | 0,0E+00 | 0,0E+00 | 748680 | 749720 | 1041 | + |
| Y11_RS03330 | Y11_07241 | | ASCH domain-containing protein | -0,12 | -0,92 | -0,53 | -0,8 | -0,41 | 0,39 | 6,7E-01 | 1,2E-04 | 2,3E-01 | 1,1E-02 | 3,9E-01 | 3,6E-01 | 0,0E+00 | 0,0E+00 | 0,0E+00 | 0,0E+00 | 0,0E+00 | 0,0E+00 | 752617 | 753006 | 390 | + |
| Y11_RS03335 | Y11_07251 | | DUF1480 family protein | 0,9 | 0,4 | 2,2 | -0,5 | 1,3 | 1,8 | 4,0E-05 | 1,3E-01 | 3,0E-06 | 7,1E-02 | 1,3E-04 | 2,3E-04 | 0,0E+00 | 0,0E+00 | 5,8E-08 | 0,0E+00 | 3,4E-06 | 1,7E-05 | 753124 | 753366 | 243 | - |
| Y11_RS03355 | Y11_07291 | | molecular chaperone | 0 | -15 | -15 | -15 | -15 | -0,48 | 0,0E+00 | 0,0E+00 | 0,0E+00 | 0,0E+00 | 0,0E+00 | 4,0E-01 | 0,0E+00 | 0,0E+00 | 0,0E+00 | 0,0E+00 | 0,0E+00 | 0,0E+00 | 758794 | 759543 | 750 | - |
| Y11_RS03385 | Y11_07351 | | PqiB family protein | 0,8 | 0,16 | 0,18 | -0,64 | -0,62 | 0,02 | 4,1E-03 | 4,7E-02 | 2,5E-02 | 1,3E-02 | 1,5E-02 | 7,7E-01 | 0,0E+00 | 0,0E+00 | 0,0E+00 | 0,0E+00 | 0,0E+00 | 0,0E+00 | 764070 | 766700 | 2631 | - |
| Y11_RS03395 | Y11_07371 | | GAF domain-containing protein | 0,03 | -0,31 | -1,1 | -0,34 | -1,13 | -0,79 | 7,4E-01 | 8,9E-04 | 3,7E-03 | 3,7E-03 | 3,6E-03 | 1,9E-02 | 0,0E+00 | 0,0E+00 | 4,3E-05 | 0,0E+00 | 7,9E-05 | 0,0E+00 | 768178 | 768675 | 498 | + |
| Y11_RS03400 | Y11_07381 | proQ | RNA chaperone ProQ | 0,36 | 2,64 | 1,47 | 2,28 | 1,11 | -1,17 | 3,9E-04 | 1,1E-09 | 4,1E-07 | 1,1E-08 | 7,9E-06 | 1,3E-05 | 0,0E+00 | 1,1E-10 | 1,1E-08 | 5,6E-10 | 2,7E-07 | 1,4E-06 | 768771 | 769499 | 729 | + |
| Y11_RS03405 | Y11_07391 | prc | carboxy terminal-processing peptidase | 0,39 | 1,06 | 0,85 | 0,67 | 0,46 | -0,21 | 2,4E-03 | 3,0E-06 | 2,0E-05 | 1,6E-05 | 3,2E-04 | 3,0E-02 | 0,0E+00 | 8,0E-08 | 0,0E+00 | 0,0E+00 | 0,0E+00 | 0,0E+00 | 769519 | 771594 | 2076 | + |
| Y11_RS03410 | Y11_07401 | htpX | protease HtpX | 0,81 | -1,12 | -0,97 | -1,93 | -1,78 | 0,15 | 6,4E-07 | 3,5E-07 | 2,4E-04 | 1,9E-08 | 4,1E-06 | 4,0E-01 | 0,0E+00 | 1,2E-08 | 0,0E+00 | 8,4E-10 | 1,5E-07 | 0,0E+00 | 771889 | 772770 | 882 | + |

| | | | | | | | | | | | | | | | | | | | | | | | | |
|---|---|---|---|---|---|---|---|---|---|---|---|---|---|---|---|---|---|---|---|---|---|---|---|---|
| Y11_RS03420 | Y11_07431 | ogl | oligogalacturonate lyase | -3,26 | -0,34 | 0,33 | 2,92 | 3,59 | 0,67 | 1,8E-11 | 2,5E-03 | 9,0E-06 | 1,1E-09 | 6,0E-12 | 2,3E-05 | 9,7E-12 | 0,0E+00 | 0,0E+00 | 9,4E-11 | 8,8E-12 | 0,0E+00 | 774646 | 775812 | 1167 | + |
| Y11_RS03425 | Y11_07441 | kdgR | DNA-binding transcriptional regulator KdgR | -0,11 | 0,47 | 0,46 | 0,58 | 0,57 | -0,01 | 4,6E-02 | 1,3E-04 | 1,5E-06 | 4,5E-05 | 1,3E-06 | 8,7E-01 | 0,0E+00 | 0,0E+00 | 0,0E+00 | 0,0E+00 | 0,0E+00 | 0,0E+00 | 775999 | 776790 | 792 | + |
| Y11_RS03430 | Y11_07451 | | N-acetylmuramoyl-L-alanine amidase | 0,86 | -0,52 | -0,13 | -1,38 | -0,99 | 0,39 | 8,0E-04 | 4,4E-03 | 3,4E-01 | 2,4E-05 | 1,9E-04 | 9,6E-03 | 0,0E+00 | 0,0E+00 | 0,0E+00 | 4,8E-07 | 0,0E+00 | 0,0E+00 | 777066 | 777917 | 852 | + |
| Y11_RS03465 | Y11_07531 | | ABC transporter substrate-binding protein | 0,08 | -0,19 | 1,04 | -0,26 | 0,96 | 1,23 | 2,5E-01 | 6,1E-03 | 4,0E-06 | 1,5E-03 | 9,0E-06 | 8,3E-07 | 0,0E+00 | 0,0E+00 | 7,4E-08 | 0,0E+00 | 0,0E+00 | 1,6E-07 | 782361 | 783653 | 1293 | - |
| Y11_RS03470 | Y11_07541 | | ABC transporter ATP-binding protein | 0,05 | -1,81 | -0,86 | -1,87 | -0,91 | 0,95 | 6,9E-01 | 3,3E-05 | 5,7E-03 | 2,7E-05 | 4,2E-03 | 1,1E-02 | 0,0E+00 | 7,5E-07 | 0,0E+00 | 5,3E-07 | 0,0E+00 | 0,0E+00 | 783669 | 784796 | 1128 | - |
| Y11_RS03490 | Y11_07591 | kduD | 2-dehydro-3-deoxy-D-gluconate 5-dehydrogenase KduD | -1,06 | -1,4 | -0,29 | -0,34 | 0,77 | 1,11 | 5,3E-07 | 6,6E-08 | 2,5E-02 | 1,6E-03 | 7,2E-05 | 5,1E-06 | 3,7E-08 | 2,9E-09 | 0,0E+00 | 0,0E+00 | 0,0E+00 | 6,6E-07 | 788646 | 789407 | 762 | - |
| Y11_RS03505 | Y11_07621 | hxpB | hexitol phosphatase HxpB | -0,45 | 0,7 | 1,31 | 1,16 | 1,77 | 0,61 | 3,1E-04 | 1,8E-06 | 6,1E-08 | 6,4E-07 | 5,1E-08 | 4,7E-05 | 0,0E+00 | 0,0E+00 | 2,2E-09 | 1,7E-08 | 3,6E-09 | 0,0E+00 | 791036 | 791701 | 666 | - |
| Y11_RS03510 | Y11_07631 | | YniB family protein | -0,28 | 0,08 | 0,09 | 0,36 | 0,37 | 0,01 | 1,6E-01 | 7,0E-01 | 7,1E-01 | 1,8E-01 | 2,0E-01 | 9,8E-01 | 0,0E+00 | 0,0E+00 | 0,0E+00 | 0,0E+00 | 0,0E+00 | 0,0E+00 | 791951 | 792505 | 555 | + |
| Y11_RS03540 | Y11_07691 | | fructosamine kinase family protein | -0,48 | 2,37 | 3,72 | 2,85 | 4,2 | 1,35 | 5,2E-07 | 1,1E-10 | 5,3E-12 | 9,9E-12 | 9,0E-13 | 6,2E-08 | 0,0E+00 | 2,4E-11 | 6,0E-12 | 5,4E-12 | 4,3E-12 | 2,6E-08 | 799327 | 800196 | 870 | - |
| Y11_RS03555 | Y11_07721 | yfeB | iron/manganese ABC transporter ATP-binding protein YfeB | 0,2 | -0,13 | -0,24 | -0,33 | -0,44 | -0,1 | 1,5E-02 | 2,6E-01 | 1,7E-02 | 1,3E-02 | 2,9E-04 | 3,9E-01 | 0,0E+00 | 0,0E+00 | 0,0E+00 | 0,0E+00 | 0,0E+00 | 0,0E+00 | 802584 | 803474 | 891 | - |
| Y11_RS03560 | Y11_07731 | yfeA | iron/manganese ABC transporter substrate-binding protein YfeA | 0,71 | -0,48 | 0,09 | -1,19 | -0,62 | 0,57 | 5,1E-05 | 8,2E-04 | 6,2E-01 | 7,9E-06 | 9,0E-03 | 1,5E-02 | 0,0E+00 | 0,0E+00 | 0,0E+00 | 1,6E-07 | 0,0E+00 | 0,0E+00 | 803471 | 804439 | 969 | - |
| Y11_RS03565 | Y11_07741 | | transglycosylase SLT domain-containing protein | 0,13 | -0,89 | -0,84 | -1,01 | -0,96 | 0,05 | 7,0E-01 | 1,2E-02 | 3,4E-02 | 1,4E-03 | 8,4E-03 | 8,1E-01 | 0,0E+00 | 0,0E+00 | 0,0E+00 | 2,4E-05 | 0,0E+00 | 0,0E+00 | 804651 | 805313 | 663 | + |
| Y11_RS03600 | Y11_07831 | | KTSC domain-containing protein | 0,49 | 1,61 | 1,39 | 1,12 | 0,9 | -0,22 | 2,0E-02 | 1,2E-05 | 1,7E-04 | 1,0E-06 | 3,6E-04 | 1,8E-01 | 0,0E+00 | 2,7E-07 | 2,2E-06 | 2,5E-08 | 0,0E+00 | 0,0E+00 | 809361 | 809573 | 213 | - |
| Y11_RS03605 | Y11_07851 | thrS | threonine--tRNA ligase | 0,26 | 0,49 | 0,27 | 0,22 | 0 | -0,22 | 5,4E-05 | 4,5E-07 | 1,6E-04 | 1,4E-04 | 9,4E-01 | 5,0E-04 | 0,0E+00 | 0,0E+00 | 0,0E+00 | 0,0E+00 | 0,0E+00 | 0,0E+00 | 809897 | 811825 | 1929 | + |
| Y11_RS03615 | Y11_07871 | rpmI | 50S ribosomal protein L35 | 0,54 | -0,03 | 0,11 | -0,56 | -0,43 | 0,13 | 3,5E-02 | 9,2E-01 | 7,0E-01 | 7,2E-02 | 1,8E-01 | 6,9E-01 | 0,0E+00 | 0,0E+00 | 0,0E+00 | 0,0E+00 | 0,0E+00 | 0,0E+00 | 812477 | 812674 | 198 | + |
| Y11_RS03620 | Y11_07881 | rplT | 50S ribosomal protein L20 | 0,35 | -0,9 | -0,91 | -1,25 | -1,26 | -0,02 | 6,9E-04 | 2,6E-06 | 1,7E-03 | 1,4E-07 | 2,0E-04 | 9,4E-01 | 0,0E+00 | 0,0E+00 | 0,0E+00 | 4,4E-09 | 4,9E-06 | 0,0E+00 | 812712 | 813068 | 357 | + |
| Y11_RS03625 | Y11_07891 | pheS | phenylalanine--tRNA ligase subunit alpha | -0,1 | -0,49 | -0,93 | -0,39 | -0,83 | -0,44 | 4,6E-04 | 2,9E-06 | 3,9E-05 | 2,3E-05 | 9,2E-05 | 6,9E-03 | 0,0E+00 | 0,0E+00 | 0,0E+00 | 0,0E+00 | 0,0E+00 | 0,0E+00 | 813533 | 814516 | 984 | + |
| Y11_RS03630 | Y11_07901 | pheT | phenylalanine--tRNA ligase subunit beta | -0,08 | -0,66 | -1,02 | -0,58 | -0,94 | -0,36 | 1,6E-01 | 2,1E-06 | 4,1E-08 | 2,3E-05 | 4,0E-07 | 4,8E-04 | 0,0E+00 | 0,0E+00 | 1,6E-09 | 0,0E+00 | 0,0E+00 | 0,0E+00 | 814531 | 816918 | 2388 | + |
| Y11_RS03635 | Y11_07911 | ihfA | integration host factor subunit alpha | -0,21 | 1,18 | 1,36 | 1,39 | 1,57 | 0,18 | 3,1E-02 | 2,1E-09 | 1,6E-06 | 1,8E-07 | 2,3E-06 | 1,4E-01 | 0,0E+00 | 1,8E-10 | 3,5E-08 | 5,6E-09 | 8,6E-08 | 0,0E+00 | 816923 | 817219 | 297 | + |
| Y11_RS03645 | Y11_07941 | | DUF2502 domain-containing protein | 0 | -15 | -15 | -1,31 | -0,54 | 0,76 | 0,0E+00 | 0,0E+00 | 0,0E+00 | 1,3E-02 | 4,6E-02 | 1,4E-01 | 0,0E+00 | 0,0E+00 | 0,0E+00 | 2,0E-04 | 0,0E+00 | 0,0E+00 | 818599 | 819093 | 495 | + |
| Y11_RS03655 | Y11_07961 | | glutathione peroxidase | 0,43 | 0,68 | 0,64 | 0,25 | 0,2 | -0,05 | 3,1E-07 | 1,9E-07 | 7,7E-05 | 5,4E-04 | 5,0E-02 | 6,3E-01 | 0,0E+00 | 0,0E+00 | 0,0E+00 | 0,0E+00 | 0,0E+00 | 0,0E+00 | 820343 | 820897 | 555 | + |
| Y11_RS03660 | Y11_07971 | btuD | vitamin B12 ABC transporter ATP-binding protein BtuD | 0,47 | 0,61 | 0,51 | 0 | 0 | 0 | 7,8E-04 | 1,9E-03 | 1,3E-02 | 0,0E+00 | 0,0E+00 | 0,0E+00 | 0,0E+00 | 0,0E+00 | 0,0E+00 | 0,0E+00 | 0,0E+00 | 0,0E+00 | 820922 | 821680 | 759 | + |
| Y11_RS03665 | Y11_07981 | arnB | UDP-4-amino-4-deoxy-L-arabinose aminotransferase | 0 | 0,68 | 0,73 | 0,07 | 0,14 | 0,04 | 0,0E+00 | 6,0E-04 | 9,5E-03 | 6,2E-01 | 5,4E-01 | 8,4E-01 | 0,0E+00 | 0,0E+00 | 0,0E+00 | 0,0E+00 | 0,0E+00 | 0,0E+00 | 822027 | 823175 | 1149 | + |
| Y11_RS03670 | Y11_07991 | arnC | undecaprenyl-phosphate 4-deoxy-4-formamido-L-arabinose transferase | 0,14 | -0,55 | -0,53 | -0,69 | -0,68 | 0,02 | 3,2E-02 | 2,6E-04 | 4,3E-04 | 1,1E-04 | 1,6E-04 | 9,0E-01 | 0,0E+00 | 0,0E+00 | 0,0E+00 | 0,0E+00 | 0,0E+00 | 0,0E+00 | 823168 | 824151 | 984 | + |
| Y11_RS03675 | Y11_08001 | arnA | bifunctional UDP-4-amino-4-deoxy-L-arabinose formyltransferase/UDP-glucuronic acid oxidase ArnA | -0,02 | -0,64 | -0,57 | -0,62 | -0,55 | 0,07 | 5,2E-01 | 6,1E-07 | 4,9E-05 | 1,7E-07 | 3,9E-05 | 4,0E-01 | 0,0E+00 | 0,0E+00 | 0,0E+00 | 0,0E+00 | 0,0E+00 | 0,0E+00 | 824148 | 826181 | 2034 | + |
| Y11_RS03680 | Y11_08011 | | 4-deoxy-4-formamido-L-arabinose-phosphoundecaprenol deformylase | 0,28 | 0,07 | -0,09 | -0,21 | -0,38 | -0,17 | 3,3E-04 | 1,6E-01 | 1,3E-01 | 9,4E-03 | 5,3E-04 | 3,8E-02 | 0,0E+00 | 0,0E+00 | 0,0E+00 | 0,0E+00 | 0,0E+00 | 0,0E+00 | 826178 | 827080 | 903 | + |
| Y11_RS03685 | Y11_08021 | arnT | lipid IV(A) 4-amino-4-deoxy-L-arabinosyltransferase | 0,25 | -0,12 | 0,55 | -0,37 | 0,3 | 0,68 | 3,9E-02 | 3,4E-01 | 9,7E-04 | 1,5E-02 | 2,3E-02 | 7,4E-04 | 0,0E+00 | 0,0E+00 | 0,0E+00 | 0,0E+00 | 0,0E+00 | 0,0E+00 | 827080 | 828744 | 1665 | + |
| Y11_RS03700 | Y11_08051 | | NlpC/P60 family protein | 0 | -3,53 | -4,22 | -15 | -15 | -0,68 | 0,0E+00 | 6,5E-10 | 1,8E-07 | 0,0E+00 | 0,0E+00 | 2,2E-02 | 0,0E+00 | 7,9E-11 | 5,4E-09 | 0,0E+00 | 0,0E+00 | 0,0E+00 | 829731 | 830195 | 465 | + |
| Y11_RS03705 | Y11_08061 | | lipoate--protein ligase A | -0,38 | 0,38 | 0,66 | 0,76 | 1,03 | 0,28 | 8,1E-05 | 1,3E-04 | 3,4E-05 | 5,4E-07 | 9,5E-07 | 8,8E-03 | 0,0E+00 | 0,0E+00 | 0,0E+00 | 0,0E+00 | 4,0E-08 | 0,0E+00 | 830442 | 831458 | 1017 | + |
| Y11_RS03710 | Y11_08071 | | YdiU family protein | -0,5 | -1,38 | -1,57 | -0,88 | -1,06 | -0,18 | 1,6E-02 | 3,6E-03 | 3,6E-03 | 2,7E-02 | 2,1E-02 | 7,1E-01 | 0,0E+00 | 7,1E-05 | 4,1E-05 | 0,0E+00 | 4,4E-04 | 0,0E+00 | 831458 | 832957 | 1500 | + |
| Y11_RS03715 | Y11_08081 | | ABC transporter substrate-binding protein | 1,05 | 0,03 | 0,64 | -0,95 | 0 | 0,7 | 4,1E-06 | 8,3E-01 | 1,0E-05 | 1,7E-04 | 0,0E+00 | 4,1E-04 | 2,3E-07 | 0,0E+00 | 0,0E+00 | 0,0E+00 | 0,0E+00 | 0,0E+00 | 833017 | 833850 | 834 | - |
| Y11_RS03720 | Y11_08091 | | hemin-degrading factor | -0,07 | -0,54 | -0,71 | -0,47 | -0,63 | -0,17 | 7,7E-01 | 3,3E-02 | 9,8E-03 | 1,2E-01 | 4,4E-02 | 4,9E-01 | 0,0E+00 | 0,0E+00 | 0,0E+00 | 0,0E+00 | 0,0E+00 | 0,0E+00 | 833843 | 834904 | 1062 | - |
| Y11_RS03735 | Y11_08121 | | 3-deoxy-7-phosphoheptulonate synthase | 0,62 | 0,07 | -0,84 | -0,54 | -1,46 | -0,91 | 4,7E-06 | 3,2E-01 | 1,2E-04 | 5,1E-05 | 2,2E-06 | 9,8E-05 | 0,0E+00 | 0,0E+00 | 0,0E+00 | 0,0E+00 | 8,4E-08 | 0,0E+00 | 837602 | 838651 | 1050 | - |
| Y11_RS03740 | Y11_08131 | | kinase/pyrophosphorylase | -0,01 | 0,32 | 1,26 | 0,33 | 1,27 | 0,94 | 8,2E-01 | 2,9E-03 | 1,4E-08 | 2,3E-03 | 1,4E-08 | 1,1E-06 | 0,0E+00 | 0,0E+00 | 6,8E-10 | 0,0E+00 | 1,2E-09 | 0,0E+00 | 838875 | 839696 | 822 | - |
| Y11_RS03745 | Y11_08141 | ppsA | phosphoenolpyruvate synthase | 0,27 | -1,82 | 0,91 | -2,09 | 0,64 | 2,73 | 3,3E-05 | 1,0E-09 | 1,0E-05 | 5,2E-10 | 1,5E-04 | 5,7E-09 | 0,0E+00 | 1,1E-10 | 0,0E+00 | 5,3E-11 | 0,0E+00 | 4,9E-09 | 839983 | 842364 | 2382 | + |
| Y11_RS03755 | Y11_08161 | | FAD-binding oxidoreductase | -0,72 | 0,17 | -0,31 | 0,89 | 0,41 | -0,48 | 3,1E-05 | 1,1E-02 | 1,6E-04 | 8,0E-06 | 1,4E-03 | 1,4E-05 | 0,0E+00 | 0,0E+00 | 0,0E+00 | 0,0E+00 | 0,0E+00 | 0,0E+00 | 844475 | 847531 | 3057 | + |
| Y11_RS03760 | Y11_08171 | | hotdog fold thioesterase | -0,08 | 0,4 | 0,17 | 0,48 | 0,25 | -0,23 | 4,8E-01 | 1,5E-02 | 1,4E-01 | 5,2E-03 | 3,7E-02 | 1,1E-01 | 0,0E+00 | 0,0E+00 | 0,0E+00 | 0,0E+00 | 0,0E+00 | 0,0E+00 | 847754 | 848176 | 423 | + |
| Y11_RS03770 | Y11_08181 | sufA | Fe-S cluster assembly scaffold SufA | 0,63 | -0,54 | -1,59 | -1,17 | -2,22 | -1,05 | 3,7E-05 | 1,2E-03 | 1,8E-06 | 5,4E-06 | 1,3E-07 | 1,2E-04 | 0,0E+00 | 0,0E+00 | 3,7E-08 | 1,2E-07 | 7,9E-09 | 9,9E-06 | 848925 | 849296 | 372 | + |
| Y11_RS03775 | Y11_08191 | sufB | Fe-S cluster assembly protein SufB | 0,55 | 0,01 | -0,71 | -0,54 | -1,26 | -0,72 | 5,0E-04 | 8,8E-01 | 4,4E-04 | 2,2E-04 | 1,8E-05 | 2,3E-04 | 0,0E+00 | 0,0E+00 | 0,0E+00 | 0,0E+00 | 5,5E-07 | 0,0E+00 | 849311 | 850822 | 1512 | + |
| Y11_RS03780 | Y11_08201 | sufC | Fe-S cluster assembly ATPase SufC | 0,59 | 0,18 | -0,13 | -0,41 | -0,73 | -0,32 | 3,7E-07 | 3,6E-03 | 1,7E-01 | 8,5E-06 | 3,5E-05 | 8,3E-03 | 0,0E+00 | 0,0E+00 | 0,0E+00 | 0,0E+00 | 0,0E+00 | 0,0E+00 | 850879 | 851625 | 747 | + |
| Y11_RS03785 | Y11_08211 | sufD | Fe-S cluster assembly protein SufD | 0,54 | 0,14 | -0,39 | -0,4 | -0,93 | -0,53 | 1,7E-05 | 4,7E-02 | 8,3E-04 | 5,7E-04 | 4,6E-06 | 2,8E-04 | 0,0E+00 | 0,0E+00 | 0,0E+00 | 0,0E+00 | 0,0E+00 | 0,0E+00 | 851600 | 852913 | 1314 | + |
| Y11_RS03790 | Y11_08221 | sufS | cysteine desulfurase SufS | 0,45 | 0,22 | 0,17 | -0,23 | -0,28 | -0,05 | 1,2E-03 | 2,5E-02 | 1,1E-01 | 3,0E-02 | 2,2E-02 | 5,6E-01 | 0,0E+00 | 0,0E+00 | 0,0E+00 | 0,0E+00 | 0,0E+00 | 0,0E+00 | 852910 | 854160 | 1251 | + |
| Y11_RS03795 | Y11_08231 | sufE | cysteine desulfuration protein SufE | 0,29 | 0,02 | -0,49 | -0,27 | -0,78 | -0,51 | 3,1E-02 | 8,6E-01 | 1,3E-02 | 5,6E-02 | 1,2E-03 | 1,3E-02 | 0,0E+00 | 0,0E+00 | 0,0E+00 | 0,0E+00 | 0,0E+00 | 0,0E+00 | 854244 | 854666 | 423 | + |
| Y11_RS03800 | Y11_08241 | | L,D-transpeptidase family protein | 0,15 | -0,68 | -0,86 | -0,83 | -1,01 | -0,18 | 2,5E-02 | 4,7E-07 | 2,5E-05 | 3,3E-07 | 9,8E-06 | 1,1E-01 | 0,0E+00 | 0,0E+00 | 0,0E+00 | 0,0E+00 | 3,2E-07 | 0,0E+00 | 854933 | 856000 | 1068 | + |
| Y11_RS03805 | | | major outer membrane lipoprotein | 0,5 | 1,62 | 1,37 | 1,12 | 0,88 | -0,25 | 1,5E-02 | 4,1E-06 | 1,0E-04 | 1,6E-06 | 4,6E-04 | 1,1E-01 | 0,0E+00 | 1,1E-07 | 1,4E-06 | 3,9E-08 | 0,0E+00 | 0,0E+00 | 856233 | 856469 | 237 | - |
| Y11_RS03810 | Y11_08251 | pykF | pyruvate kinase PykF | 0,32 | 3,02 | 1,52 | 2,7 | 1,2 | -1,5 | 1,7E-04 | 5,9E-12 | 1,6E-09 | 4,7E-12 | 3,4E-09 | 8,1E-10 | 0,0E+00 | 4,8E-12 | 1,4E-10 | 4,1E-12 | 4,2E-10 | 1,4E-09 | 856789 | 858201 | 1413 | - |
| Y11_RS03830 | Y11_08271 | | riboflavin synthase | 1,82 | 2,11 | 0,87 | 0,3 | -0,95 | -1,24 | 3,8E-09 | 8,0E-10 | 6,6E-06 | 7,1E-04 | 1,9E-06 | 1,8E-07 | 7,1E-10 | 9,2E-11 | 0,0E+00 | 0,0E+00 | 0,0E+00 | 5,4E-08 | 861011 | 861664 | 654 | + |
| Y11_RS03835 | Y11_08281 | cfa | cyclopropane fatty acyl phospholipid synthase | 1,52 | 2,79 | 1,81 | 1,28 | 0,29 | -0,98 | 1,2E-08 | 1,2E-10 | 2,1E-08 | 1,4E-08 | 4,3E-03 | 1,2E-06 | 1,5E-09 | 2,5E-11 | 9,3E-10 | 6,7E-10 | 0,0E+00 | 0,0E+00 | 861724 | 862875 | 1152 | - |
| Y11_RS03845 | Y11_08301 | | LysR family transcriptional regulator | -0,03 | -0,28 | -0,21 | -0,25 | -0,19 | 0,07 | 9,0E-01 | 1,5E-01 | 1,5E-01 | 2,6E-01 | 3,1E-01 | 6,5E-01 | 0,0E+00 | 0,0E+00 | 0,0E+00 | 0,0E+00 | 0,0E+00 | 0,0E+00 | 864691 | 865629 | 939 | + |
| Y11_RS03850 | Y11_08311 | purR | HTH-type transcriptional repressor PurR | 0,07 | 0,11 | 0,04 | 0,03 | -0,03 | -0,07 | 2,0E-01 | 2,2E-02 | 6,0E-01 | 4,3E-01 | 6,5E-01 | 3,1E-01 | 0,0E+00 | 0,0E+00 | 0,0E+00 | 0,0E+00 | 0,0E+00 | 0,0E+00 | 865666 | 866691 | 1026 | - |
| Y11_RS03855 | Y11_08321 | sodB | superoxide dismutase [Fe] | 0,06 | 1,73 | 1,96 | 1,67 | 1,91 | 0,23 | 5,3E-01 | 2,8E-07 | 1,4E-07 | 4,8E-08 | 2,9E-08 | 7,7E-02 | 0,0E+00 | 9,9E-09 | 4,4E-09 | 1,8E-09 | 2,3E-09 | 0,0E+00 | 867217 | 867795 | 579 | - |
| Y11_RS03860 | Y11_08331 | | NlpC/P60 family protein | 0 | -15 | -15 | -15 | -15 | 0,76 | 0,0E+00 | 0,0E+00 | 0,0E+00 | 0,0E+00 | 0,0E+00 | 2,7E-03 | 0,0E+00 | 0,0E+00 | 0,0E+00 | 0,0E+00 | 0,0E+00 | 0,0E+00 | 868046 | 868936 | 891 | - |
| Y11_RS03865 | Y11_08341 | | Grx4 family monothiol glutaredoxin | -0,31 | 1,01 | 0,81 | 1,33 | 1,13 | -0,2 | 3,7E-03 | 5,9E-06 | 3,8E-06 | 2,2E-06 | 1,7E-06 | 9,9E-02 | 0,0E+00 | 1,5E-07 | 0,0E+00 | 5,1E-08 | 6,8E-08 | 0,0E+00 | 869370 | 869720 | 351 | + |
| Y11_RS03870 | Y11_08351 | gloA | lactoylglutathione lyase | 0,09 | 0,72 | 0,56 | 0,63 | 0,47 | -0,16 | 5,0E-01 | 2,0E-03 | 5,2E-03 | 4,5E-03 | 1,3E-02 | 3,9E-01 | 0,0E+00 | 0,0E+00 | 0,0E+00 | 0,0E+00 | 0,0E+00 | 0,0E+00 | 869852 | 870259 | 408 | - |
| Y11_RS03875 | Y11_08361 | | alkene reductase | 0,03 | -1,1 | -2,48 | -1,13 | -2,52 | -1,39 | 4,3E-01 | 6,3E-09 | 3,1E-09 | 6,6E-09 | 3,0E-09 | 3,4E-07 | 0,0E+00 | 4,4E-10 | 2,2E-10 | 3,7E-10 | 3,8E-10 | 8,0E-08 | 870441 | 871538 | 1098 | - |
| Y11_RS03880 | Y11_08371 | | TetR/AcrR family transcriptional regulator | -0,14 | -0,44 | -0,38 | -0,3 | -0,24 | 0,06 | 5,3E-01 | 3,7E-02 | 7,7E-02 | 1,1E-01 | 2,2E-01 | 7,0E-01 | 0,0E+00 | 0,0E+00 | 0,0E+00 | 0,0E+00 | 0,0E+00 | 0,0E+00 | 871608 | 872216 | 609 | - |
| Y11_RS03895 | Y11_08401 | | aldo/keto reductase family oxidoreductase | 0,43 | 0,43 | -0,08 | 0 | -0,51 | -0,51 | 3,3E-04 | 5,7E-06 | 1,5E-01 | 9,9E-01 | 1,3E-04 | 3,8E-06 | 0,0E+00 | 0,0E+00 | 0,0E+00 | 0,0E+00 | 0,0E+00 | 0,0E+00 | 874636 | 875532 | 897 | + |
| Y11_RS03905 | Y11_08421 | | HlyD family secretion protein | 1,28 | 2,3 | 1,27 | 1,02 | -0,01 | -1,03 | 2,0E-01 | 3,5E-02 | 2,8E-01 | 8,6E-07 | 9,9E-01 | 1,2E-01 | 6,5E-03 | 6,7E-04 | 3,0E-03 | 2,2E-08 | 0,0E+00 | 6,3E-03 | 877584 | 878441 | 858 | - |
| Y11_RS03915 | Y11_08441 | rovA | virulence master transcriptional regulator RovA | -0,75 | -1,04 | 0,15 | -0,29 | 0,9 | 1,19 | 2,3E-06 | 1,1E-06 | 3,1E-01 | 2,6E-03 | 1,6E-04 | 3,4E-05 | 0,0E+00 | 3,3E-08 | 0,0E+00 | 0,0E+00 | 0,0E+00 | 3,2E-06 | 878865 | 879296 | 432 | + |
| Y11_RS03920 | Y11_08451 | | glycine zipper 2TM domain-containing protein | -0,08 | 1,17 | 0,62 | 1,25 | 0,7 | -0,55 | 7,4E-01 | 1,3E-04 | 1,7E-02 | 8,7E-05 | 9,3E-03 | 4,1E-03 | 0,0E+00 | 2,8E-06 | 0,0E+00 | 1,6E-06 | 0,0E+00 | 0,0E+00 | 879417 | 879884 | 468 | - |
| Y11_RS03925 | Y11_08471 | anmK | anhydro-N-acetylmuramic acid kinase | 0,48 | 0,35 | 0,88 | -0,13 | 0,39 | 0,53 | 1,4E-03 | 7,7E-03 | 4,2E-05 | 3,0E-02 | 4,0E-04 | 3,9E-05 | 0,0E+00 | 0,0E+00 | 0,0E+00 | 0,0E+00 | 0,0E+00 | 0,0E+00 | 880329 | 881441 | 1113 | + |
| Y11_RS03930 | Y11_08481 | | MliC family protein | 0,2 | 0,44 | 1,02 | 0,24 | 0,82 | 0,58 | 2,0E-02 | 1,1E-04 | 3,9E-05 | 8,4E-03 | 2,2E-04 | 1,8E-03 | 0,0E+00 | 0,0E+00 | 5,8E-07 | 0,0E+00 | 0,0E+00 | 0,0E+00 | 881497 | 881814 | 318 | + |
| Y11_RS03935 | Y11_08491 | pdxH | pyridoxamine 5'-phosphate oxidase | 0,02 | 0,05 | -0,02 | 0,04 | -0,04 | -0,07 | 8,6E-01 | 4,6E-01 | 6,2E-01 | 5,3E-01 | 7,5E-01 | 7,2E-01 | 0,0E+00 | 0,0E+00 | 0,0E+00 | 0,0E+00 | 0,0E+00 | 0,0E+00 | 881922 | 882575 | 654 | + |
| Y11_RS03940 | Y11_08501 | tyrS | tyrosine--tRNA ligase | 0,15 | -0,36 | -0,29 | -0,51 | -0,44 | 0,07 | 1,4E-01 | 2,2E-03 | 1,1E-02 | 2,0E-05 | 1,8E-04 | 2,7E-01 | 0,0E+00 | 0,0E+00 | 0,0E+00 | 0,0E+00 | 0,0E+00 | 0,0E+00 | 882691 | 883965 | 1275 | + |
| Y11_RS03945 | Y11_08511 | pdxY | pyridoxal kinase PdxY | -0,32 | 0 | 0,03 | 0,32 | 0,34 | 0,02 | 4,4E-02 | 9,8E-01 | 8,3E-01 | 1,8E-02 | 1,3E-02 | 7,9E-01 | 0,0E+00 | 0,0E+00 | 0,0E+00 | 0,0E+00 | 0,0E+00 | 0,0E+00 | 884040 | 884900 | 861 | + |
| Y11_RS03950 | Y11_08521 | gstA | glutathione transferase GstA | -0,52 | 1,02 | 1,48 | 1,54 | 2,01 | 0,46 | 4,2E-05 | 1,2E-06 | 5,2E-07 | 1,9E-07 | 1,2E-07 | 5,4E-03 | 0,0E+00 | 3,6E-08 | 1,3E-08 | 5,8E-09 | 6,9E-09 | 0,0E+00 | 884952 | 885560 | 609 | - |
| Y11_RS03965 | Y11_08561 | | VOC family protein | 0,82 | -1,41 | 0,86 | -2,23 | 0,05 | 2,28 | 5,1E-05 | 1,6E-07 | 1,1E-04 | 2,7E-09 | 7,1E-01 | 1,8E-08 | 0,0E+00 | 6,2E-09 | 0,0E+00 | 1,8E-10 | 0,0E+00 | 1,1E-08 | 887002 | 887355 | 354 | + |
| Y11_RS03970 | Y11_08571 | dtpA | dipeptide/tripeptide permease DtpA | 0,45 | 15 | 15 | 15 | 15 | 0 | 5,7E-02 | 0,0E+00 | 0,0E+00 | 0,0E+00 | 0,0E+00 | 0,0E+00 | 0,0E+00 | 0,0E+00 | 0,0E+00 | 0,0E+00 | 0,0E+00 | 0,0E+00 | 887742 | 889265 | 1524 | - |

| | | | | | | | | | | | | | | | | | | | | | | | | |
|---|---|---|---|---|---|---|---|---|---|---|---|---|---|---|---|---|---|---|---|---|---|---|---|---|
| Y11_RS03975 | Y11_08591 | | LysR family transcriptional regulator | -0,47 | 15 | 15 | 15 | 15 | 0 | 9,6E-06 | 0,0E+00 | 0,0E+00 | 0,0E+00 | 0,0E+00 | 0,0E+00 | 0,0E+00 | 0,0E+00 | 0,0E+00 | 0,0E+00 | 0,0E+00 | 0,0E+00 | 889804 | 890709 | 906 | + |
| Y11_RS03990 | Y11_08621 | sapF | peptide ABC transporter ATP-binding protein SapF | -0,01 | 15 | 15 | 15 | 15 | 0 | 9,3E-01 | 0,0E+00 | 0,0E+00 | 0,0E+00 | 0,0E+00 | 0,0E+00 | 0,0E+00 | 0,0E+00 | 0,0E+00 | 0,0E+00 | 0,0E+00 | 0,0E+00 | 892342 | 893157 | 816 | - |
| Y11_RS03995 | Y11_08631 | sapD | peptide ABC transporter ATP-binding protein SapD | 0,31 | 15 | 4,23 | 15 | 3,72 | 0 | 1,5E-01 | 0,0E+00 | 2,4E-09 | 0,0E+00 | 1,4E-08 | 0,0E+00 | 0,0E+00 | 0,0E+00 | 1,9E-10 | 0,0E+00 | 1,2E-09 | 0,0E+00 | 893157 | 894149 | 993 | - |
| Y11_RS04010 | Y11_08661 | sapA | peptide ABC transporter substrate-binding protein SapA | -0,14 | 15 | 15 | 15 | 15 | 0 | 9,5E-02 | 0,0E+00 | 0,0E+00 | 0,0E+00 | 0,0E+00 | 0,0E+00 | 0,0E+00 | 0,0E+00 | 0,0E+00 | 0,0E+00 | 0,0E+00 | 0,0E+00 | 895988 | 897661 | 1674 | - |
| Y11_RS04035 | Y11_08701 | pspF | phage shock protein operon transcriptional activator | -0,78 | 0,9 | 1,33 | 1,68 | 2,11 | 0,42 | 5,0E-09 | 4,7E-06 | 1,5E-05 | 3,0E-08 | 4,3E-07 | 3,1E-02 | 0,0E+00 | 0,0E+00 | 2,5E-07 | 1,2E-09 | 2,1E-08 | 0,0E+00 | 901019 | 902029 | 1011 | - |
| Y11_RS04040 | Y11_08711 | pspA | phage shock protein PspA | -0,11 | 1,14 | 1,47 | 1,25 | 1,58 | 0,33 | 6,7E-02 | 2,1E-05 | 7,8E-08 | 6,7E-06 | 1,0E-08 | 4,4E-02 | 0,0E+00 | 4,9E-07 | 2,6E-09 | 1,4E-07 | 9,8E-10 | 0,0E+00 | 902219 | 902884 | 666 | + |
| Y11_RS04045 | Y11_08721 | pspB | envelope stress response membrane protein PspB | 0,02 | 1,39 | 1,85 | 1,36 | 1,82 | 0,46 | 7,5E-01 | 6,2E-09 | 8,2E-07 | 1,4E-08 | 1,0E-06 | 6,9E-03 | 0,0E+00 | 4,4E-10 | 1,9E-08 | 6,8E-10 | 4,3E-08 | 0,0E+00 | 903046 | 903273 | 228 | + |
| Y11_RS04050 | Y11_08731 | pspC | envelope stress response membrane protein PspC | -0,6 | 1,8 | 2,88 | 2,4 | 3,48 | 1,08 | 1,5E-02 | 6,1E-06 | 1,3E-02 | 9,4E-07 | 5,1E-03 | 2,7E-01 | 0,0E+00 | 1,5E-07 | 1,5E-04 | 2,4E-08 | 1,1E-04 | 1,4E-02 | 903273 | 903677 | 405 | + |
| Y11_RS04060 | Y11_08751 | | YcjX family protein | -0,08 | 0,68 | 0,75 | 0,76 | 0,83 | 0,06 | 3,7E-01 | 2,2E-05 | 1,0E-04 | 7,6E-06 | 4,5E-05 | 5,3E-01 | 0,0E+00 | 0,0E+00 | 0,0E+00 | 0,0E+00 | 0,0E+00 | 0,0E+00 | 903943 | 905340 | 1398 | + |
| Y11_RS04065 | Y11_08761 | | YcjF family protein | -0,16 | -0,01 | 0,51 | 0,15 | 0,68 | 0,52 | 1,2E-01 | 9,2E-01 | 2,1E-02 | 7,9E-02 | 5,4E-03 | 1,6E-02 | 0,0E+00 | 0,0E+00 | 0,0E+00 | 0,0E+00 | 0,0E+00 | 0,0E+00 | 905337 | 906401 | 1065 | + |
| Y11_RS04075 | Y11_08781 | tyrR | transcriptional regulator TyrR | -0,45 | 0,25 | 0,65 | 0,71 | 1,1 | 0,39 | 2,7E-04 | 5,0E-03 | 4,0E-06 | 7,3E-06 | 1,1E-07 | 7,4E-05 | 0,0E+00 | 0,0E+00 | 0,0E+00 | 0,0E+00 | 6,5E-09 | 0,0E+00 | 907171 | 908748 | 1578 | + |
| Y11_RS04080 | Y11_08791 | tpx | thiol peroxidase | 0,12 | -0,49 | 0,06 | -0,61 | -0,07 | 0,55 | 2,2E-01 | 3,5E-04 | 6,3E-01 | 2,6E-04 | 6,1E-01 | 1,8E-03 | 0,0E+00 | 0,0E+00 | 0,0E+00 | 0,0E+00 | 0,0E+00 | 0,0E+00 | 908995 | 909498 | 504 | - |
| Y11_RS04085 | Y11_08801 | ycjG | L-Ala-D/L-Glu epimerase | 0,29 | -0,4 | -0,16 | -0,69 | -0,45 | 0,24 | 1,2E-02 | 1,2E-02 | 2,1E-01 | 2,4E-04 | 2,3E-03 | 1,1E-01 | 0,0E+00 | 0,0E+00 | 0,0E+00 | 0,0E+00 | 0,0E+00 | 0,0E+00 | 909651 | 910622 | 972 | + |
| Y11_RS04090 | Y11_08811 | mpaA | murein tripeptide amidase MpaA | 0,02 | -0,44 | -0,73 | -0,46 | -0,75 | -0,3 | 8,7E-01 | 1,3E-03 | 8,2E-04 | 6,7E-04 | 5,9E-04 | 3,6E-02 | 0,0E+00 | 0,0E+00 | 0,0E+00 | 0,0E+00 | 0,0E+00 | 0,0E+00 | 910619 | 911326 | 708 | - |
| Y11_RS04095 | Y11_08821 | | ABC transporter substrate-binding protein | -0,06 | -4,01 | -3,64 | -3,96 | -3,58 | 0,37 | 4,2E-01 | 2,0E-10 | 4,8E-10 | 1,2E-10 | 3,4E-10 | 1,8E-02 | 0,0E+00 | 3,6E-11 | 6,3E-11 | 2,0E-11 | 9,5E-11 | 0,0E+00 | 911612 | 913228 | 1617 | + |
| Y11_RS04100 | Y11_08831 | zntB | zinc transporter ZntB | 0,6 | 2,05 | 2,35 | 1,45 | 1,75 | 0,3 | 4,1E-04 | 3,2E-08 | 2,7E-09 | 1,8E-06 | 1,8E-07 | 1,8E-02 | 0,0E+00 | 1,6E-09 | 2,0E-10 | 4,3E-08 | 1,0E-08 | 0,0E+00 | 913665 | 914648 | 984 | + |
| Y11_RS04105 | Y11_08841 | ttcA | tRNA 2-thiocytidine(32) synthetase TtcA | 0,44 | -0,02 | -1,65 | -0,47 | -2,1 | -1,63 | 3,8E-03 | 8,7E-01 | 4,2E-03 | 2,2E-03 | 8,7E-04 | 4,5E-03 | 0,0E+00 | 0,0E+00 | 4,8E-05 | 0,0E+00 | 2,0E-05 | 2,7E-04 | 914968 | 915909 | 942 | - |
| Y11_RS04115 | Y11_08871 | nifJ | pyruvate:ferredoxin (flavodoxin) oxidoreductase | 0,28 | -0,54 | -1,35 | -0,83 | -1,63 | -0,8 | 7,8E-05 | 1,2E-06 | 6,1E-10 | 1,2E-07 | 3,8E-10 | 1,7E-07 | 0,0E+00 | 0,0E+00 | 7,5E-11 | 0,0E+00 | 9,8E-11 | 0,0E+00 | 917100 | 920633 | 3534 | - |
| Y11_RS04125 | Y11_08901 | | MgtC/SapB family protein | -0,23 | 0,56 | 0,74 | 0,79 | 0,97 | 0,18 | 2,2E-02 | 8,3E-05 | 5,7E-04 | 3,7E-05 | 1,7E-04 | 2,5E-01 | 0,0E+00 | 0,0E+00 | 0,0E+00 | 0,0E+00 | 0,0E+00 | 0,0E+00 | 921489 | 922148 | 660 | - |
| Y11_RS04135 | Y11_08921 | | 2-hydroxyacid dehydrogenase | -0,16 | 0,54 | -0,33 | 0,7 | -0,17 | -0,87 | 3,8E-03 | 3,2E-06 | 2,7E-02 | 1,3E-06 | 2,0E-01 | 1,4E-04 | 0,0E+00 | 0,0E+00 | 0,0E+00 | 0,0E+00 | 0,0E+00 | 0,0E+00 | 922835 | 923827 | 993 | - |
| Y11_RS04140 | Y11_08931 | | YdbH family protein | -0,44 | 0,38 | 0,97 | 0,81 | 1,4 | 0 | 1,9E-01 | 2,1E-01 | 5,2E-03 | 1,0E-02 | 1,6E-04 | 0,0E+00 | 0,0E+00 | 0,0E+00 | 0,0E+00 | 0,0E+00 | 4,2E-06 | 0,0E+00 | 924101 | 926734 | 2634 | + |
| Y11_RS04145 | Y11_08941 | | YnbE family lipoprotein | 0,06 | -0,51 | 1,15 | -0,57 | 1,06 | 1,65 | 8,6E-01 | 1,2E-01 | 4,2E-03 | 1,9E-01 | 2,7E-02 | 1,5E-03 | 0,0E+00 | 0,0E+00 | 4,8E-05 | 0,0E+00 | 5,6E-04 | 9,8E-05 | 926731 | 926925 | 195 | + |
| Y11_RS04150 | Y11_08951 | | YdbL family protein | 0,3 | -2,37 | -2,09 | -2,67 | -2,39 | 0,28 | 1,0E-01 | 3,9E-08 | 1,2E-06 | 1,1E-07 | 1,3E-06 | 9,2E-02 | 0,0E+00 | 1,8E-09 | 2,7E-08 | 3,6E-09 | 5,4E-08 | 0,0E+00 | 926935 | 927306 | 372 | + |
| Y11_RS04155 | Y11_08961 | azoR | FMN-dependent NADH-azoreductase | 0 | -3,28 | -5,63 | -3,28 | -5,63 | -2,35 | 9,9E-01 | 6,7E-08 | 7,4E-11 | 1,0E-07 | 1,6E-10 | 3,8E-06 | 0,0E+00 | 2,9E-09 | 1,9E-11 | 3,4E-09 | 6,2E-11 | 5,2E-07 | 927636 | 928241 | 606 | - |
| Y11_RS04160 | Y11_08971 | hrpA | ATP-dependent RNA helicase HrpA | -0,13 | 0,48 | 0,16 | 0,61 | 0,29 | -0,32 | 1,4E-03 | 1,3E-07 | 2,7E-02 | 1,2E-08 | 1,1E-03 | 5,7E-04 | 0,0E+00 | 0,0E+00 | 0,0E+00 | 0,0E+00 | 0,0E+00 | 0,0E+00 | 928586 | 932473 | 3888 | + |
| Y11_RS04190 | Y11_09041 | rstB | two-component system sensor histidine kinase RstB | 0,32 | -0,56 | -1,14 | -0,88 | -1,47 | -0,58 | 9,4E-02 | 1,9E-04 | 3,2E-06 | 5,4E-04 | 2,3E-05 | 9,7E-05 | 0,0E+00 | 0,0E+00 | 6,1E-08 | 0,0E+00 | 6,8E-07 | 0,0E+00 | 936205 | 937512 | 1308 | - |
| Y11_RS04195 | Y11_09051 | rstA | two-component system response regulator RstA | 0 | -1,13 | -3,18 | -15 | -15 | -2,1 | 0,0E+00 | 4,5E-02 | 2,6E-04 | 0,0E+00 | 0,0E+00 | 9,9E-03 | 0,0E+00 | 8,4E-04 | 3,4E-06 | 0,0E+00 | 0,0E+00 | 5,7E-04 | 937509 | 938237 | 729 | - |
| Y11_RS04200 | Y11_09061 | | hypothetical protein | 1,41 | 1,49 | 1,67 | 0,08 | 0,26 | 0,18 | 4,2E-05 | 5,7E-05 | 2,3E-05 | 6,0E-01 | 1,1E-01 | 3,1E-01 | 2,0E-06 | 1,3E-06 | 3,6E-07 | 0,0E+00 | 0,0E+00 | 0,0E+00 | 938480 | 938896 | 417 | + |
| Y11_RS04210 | Y11_09091 | ydgH | DUF1471 family protein YdgH | 0,44 | -1,63 | -1,22 | -2,07 | -1,66 | 0,41 | 5,1E-04 | 1,0E-08 | 9,2E-06 | 3,0E-10 | 5,7E-07 | 5,4E-03 | 0,0E+00 | 6,3E-10 | 1,6E-07 | 3,5E-11 | 2,6E-08 | 0,0E+00 | 940621 | 941574 | 954 | - |
| Y11_RS04220 | Y11_09101 | pntA | Re/Si-specific NAD(P)(+) transhydrogenase subunit alpha | 0,25 | -0,3 | -0,44 | -0,55 | -0,69 | -0,14 | 4,1E-03 | 4,0E-03 | 1,5E-03 | 4,6E-05 | 5,2E-05 | 1,7E-01 | 0,0E+00 | 0,0E+00 | 0,0E+00 | 0,0E+00 | 0,0E+00 | 0,0E+00 | 942107 | 943633 | 1527 | + |
| Y11_RS04225 | Y11_09111 | pntB | Re/Si-specific NAD(P)(+) transhydrogenase subunit beta | 0,19 | -0,02 | 0,13 | -0,21 | -0,06 | 0,15 | 2,6E-02 | 7,4E-01 | 4,9E-01 | 2,2E-02 | 7,5E-01 | 4,4E-01 | 0,0E+00 | 0,0E+00 | 0,0E+00 | 0,0E+00 | 0,0E+00 | 0,0E+00 | 943754 | 945145 | 1392 | + |
| Y11_RS04230 | Y11_09121 | uspE | universal stress protein UspE | 0,4 | 0,68 | 2,18 | 0,29 | 1,78 | 1,49 | 5,1E-05 | 2,5E-06 | 5,0E-08 | 2,4E-03 | 3,2E-07 | 1,5E-06 | 0,0E+00 | 0,0E+00 | 1,8E-09 | 0,0E+00 | 1,6E-08 | 2,5E-07 | 945511 | 946467 | 957 | - |
| Y11_RS04235 | Y11_09131 | | FNR family transcription factor | 0,2 | -0,04 | 0,28 | -0,24 | 0,08 | 0,32 | 3,2E-02 | 6,9E-01 | 9,3E-03 | 3,4E-02 | 3,4E-01 | 1,1E-02 | 0,0E+00 | 0,0E+00 | 0,0E+00 | 0,0E+00 | 0,0E+00 | 0,0E+00 | 946704 | 947459 | 756 | - |
| Y11_RS04240 | Y11_09141 | ogt | methylated-DNA--[protein]-cysteine S-methyltransferase | -0,81 | -1,14 | -1,13 | -0,32 | -0,31 | 0,01 | 7,1E-03 | 8,2E-05 | 7,4E-04 | 2,3E-01 | 3,1E-01 | 9,6E-01 | 0,0E+00 | 1,8E-06 | 9,0E-06 | 0,0E+00 | 0,0E+00 | 0,0E+00 | 947532 | 948050 | 519 | - |
| Y11_RS04245 | Y11_09151 | | N-formylglutamate amidohydrolase | 0,08 | -0,4 | -0,19 | -0,48 | -0,27 | 0,21 | 5,9E-01 | 8,8E-02 | 3,7E-01 | 3,3E-02 | 1,7E-01 | 4,1E-01 | 0,0E+00 | 0,0E+00 | 0,0E+00 | 0,0E+00 | 0,0E+00 | 0,0E+00 | 948301 | 949089 | 789 | + |
| Y11_RS04255 | Y11_09171 | sbmC | DNA gyrase inhibitor SbmC | 0,08 | 0,7 | 2,55 | 0,62 | 2,47 | 1,85 | 3,6E-01 | 2,1E-03 | 2,9E-09 | 5,8E-03 | 1,2E-08 | 4,3E-06 | 0,0E+00 | 0,0E+00 | 2,1E-10 | 0,0E+00 | 1,1E-09 | 5,8E-07 | 949861 | 950334 | 474 | + |
| Y11_RS04260 | Y11_09181 | | SDR family oxidoreductase | 0 | 1,16 | 2,34 | 1,16 | 2,34 | 1,19 | 9,9E-01 | 7,3E-08 | 4,7E-07 | 2,7E-07 | 5,9E-07 | 7,6E-05 | 0,0E+00 | 3,1E-09 | 1,2E-08 | 7,9E-09 | 2,7E-08 | 6,5E-06 | 950550 | 951290 | 741 | + |
| Y11_RS04270 | Y11_09201 | rbsK | ribokinase | -2,01 | -0,93 | 2,73 | 1,08 | 4,74 | 3,66 | 2,4E-10 | 7,9E-06 | 5,7E-07 | 4,1E-06 | 8,5E-09 | 1,1E-07 | 6,7E-11 | 0,0E+00 | 1,4E-08 | 8,9E-08 | 8,6E-10 | 3,9E-08 | 953179 | 954108 | 930 | + |
| Y11_RS04285 | Y11_09251 | yjjG | pyrimidine 5'-nucleotidase | 0,37 | -2,01 | -2,08 | -2,38 | -2,46 | -0,07 | 6,6E-02 | 5,5E-06 | 1,8E-05 | 1,5E-07 | 1,3E-06 | 7,4E-01 | 0,0E+00 | 1,4E-07 | 2,9E-07 | 4,9E-09 | 5,4E-08 | 0,0E+00 | 958143 | 958817 | 675 | + |
| Y11_RS04290 | Y11_09261 | ilvN | acetolactate synthase small subunit | -0,61 | -3,2 | -1,52 | -2,59 | -0,91 | 1,68 | 2,1E-04 | 1,2E-10 | 5,0E-08 | 2,1E-09 | 7,5E-06 | 1,1E-08 | 0,0E+00 | 2,4E-11 | 1,8E-09 | 1,5E-10 | 0,0E+00 | 7,8E-09 | 958933 | 959238 | 306 | - |
| Y11_RS04345 | Y11_09381 | | inositol monophosphatase | -0,83 | -1,73 | -1,26 | -0,91 | -0,43 | 0,48 | 8,1E-05 | 7,0E-10 | 1,1E-06 | 4,7E-05 | 1,6E-02 | 1,2E-03 | 0,0E+00 | 8,2E-11 | 2,5E-08 | 0,0E+00 | 0,0E+00 | 0,0E+00 | 972982 | 973794 | 813 | - |
| Y11_RS04375 | Y11_09441 | | LysR family transcriptional regulator | 0,65 | 1,96 | 2,71 | 1,31 | 2,05 | 0,73 | 1,6E-03 | 1,3E-04 | 6,8E-08 | 1,4E-03 | 1,7E-07 | 2,9E-02 | 0,0E+00 | 2,7E-06 | 2,4E-09 | 2,3E-05 | 9,8E-09 | 0,0E+00 | 978267 | 979121 | 855 | - |
| Y11_RS04400 | Y11_09521 | | Lrp/AsnC family transcriptional regulator | 0 | 0 | -15 | 0 | -0,27 | -1,6 | 0,0E+00 | 0,0E+00 | 0,0E+00 | 0,0E+00 | 6,4E-01 | 2,4E-02 | 0,0E+00 | 0,0E+00 | 0,0E+00 | 0,0E+00 | 0,0E+00 | 1,3E-03 | 982666 | 983121 | 456 | - |
| Y11_RS04410 | Y11_09541 | | MarR family transcriptional regulator | 0,23 | -0,39 | -0,09 | -0,62 | -0,32 | 0,29 | 4,3E-01 | 1,8E-03 | 5,0E-01 | 5,8E-02 | 3,1E-01 | 5,8E-02 | 0,0E+00 | 0,0E+00 | 0,0E+00 | 0,0E+00 | 0,0E+00 | 0,0E+00 | 984398 | 984919 | 522 | + |
| Y11_RS04425 | Y11_09571 | yghU | glutathione-dependent disulfide-bond oxidoreductase | 0,38 | 1,04 | 1,02 | 0,66 | 0,64 | -0,02 | 1,3E-03 | 2,0E-08 | 1,1E-05 | 7,3E-06 | 5,1E-04 | 8,7E-01 | 0,0E+00 | 1,1E-09 | 1,9E-07 | 0,0E+00 | 0,0E+00 | 0,0E+00 | 987473 | 988324 | 852 | - |
| Y11_RS04465 | Y11_09651 | | hypothetical protein | 0 | -15 | 0 | -15 | 0 | 1,04 | 0,0E+00 | 0,0E+00 | 0,0E+00 | 0,0E+00 | 0,0E+00 | 1,2E-02 | 0,0E+00 | 0,0E+00 | 0,0E+00 | 0,0E+00 | 0,0E+00 | 7,0E-04 | 995476 | 995793 | 318 | - |
| Y11_RS04485 | Y11_09691 | | hypothetical protein | 1,4 | 1,65 | 1,31 | 0,26 | -0,08 | -0,34 | 1,0E-08 | 2,1E-09 | 1,3E-04 | 2,1E-04 | 6,7E-01 | 1,1E-01 | 1,4E-09 | 1,8E-10 | 1,7E-06 | 0,0E+00 | 0,0E+00 | 0,0E+00 | 998316 | 998714 | 399 | + |
| Y11_RS04500 | Y11_09711 | | type II secretion system protein GspD | 0 | 0 | 2,43 | 0 | -15 | 0,03 | 0,0E+00 | 0,0E+00 | 1,2E-03 | 0,0E+00 | 0,0E+00 | 9,6E-01 | 0,0E+00 | 0,0E+00 | 1,4E-05 | 0,0E+00 | 0,0E+00 | 0,0E+00 | 1000090 | 1001352 | 1263 | - |
| Y11_RS04525 | | | HTH domain-containing protein | -0,69 | 0,71 | 1,73 | 1,4 | 2,36 | 0 | 4,2E-02 | 2,4E-04 | 3,1E-07 | 9,1E-04 | 2,6E-05 | 0,0E+00 | 0,0E+00 | 0,0E+00 | 8,5E-09 | 1,5E-05 | 7,5E-07 | 0,0E+00 | 1003330 | 1003680 | 351 | + |
| Y11_RS04545 | Y11_09781 | ydfG | bifunctional NADP-dependent 3-hydroxy acid dehydrogenase/3-hydroxypropionate dehydrogenase YdfG | 0,11 | 0,06 | -0,1 | -0,05 | -0,22 | -0,16 | 2,1E-01 | 5,1E-01 | 2,7E-01 | 3,2E-01 | 4,3E-03 | 2,4E-02 | 0,0E+00 | 0,0E+00 | 0,0E+00 | 0,0E+00 | 0,0E+00 | 0,0E+00 | 1005136 | 1005885 | 750 | + |
| Y11_RS04560 | Y11_09811 | | DUF1283 family protein | 0,08 | -2,21 | -1,63 | -2,28 | -1,7 | 0,58 | 7,2E-01 | 3,9E-07 | 3,9E-04 | 7,3E-07 | 3,6E-04 | 5,1E-02 | 0,0E+00 | 1,3E-08 | 4,9E-06 | 1,9E-08 | 8,7E-06 | 0,0E+00 | 1006643 | 1007014 | 372 | + |
| Y11_RS04565 | Y11_09821 | | DUF1161 domain-containing protein | 1,02 | 0,92 | 2,39 | -0,1 | 1,37 | 1,47 | 1,1E-04 | 1,9E-04 | 1,0E-04 | 5,1E-01 | 3,6E-03 | 2,4E-03 | 4,7E-06 | 0,0E+00 | 1,4E-06 | 0,0E+00 | 7,9E-05 | 1,5E-04 | 1007106 | 1007351 | 246 | + |
| Y11_RS04580 | Y11_09851 | bioD | dethiobiotin synthase | 0,19 | 1,49 | 1,54 | 1,3 | 1,35 | 0,05 | 2,7E-02 | 7,7E-07 | 5,9E-06 | 1,6E-06 | 1,3E-05 | 7,7E-01 | 0,0E+00 | 2,3E-08 | 1,0E-07 | 3,8E-08 | 4,2E-07 | 0,0E+00 | 1009307 | 1009975 | 669 | - |
| Y11_RS04585 | Y11_09861 | mlc | ROK family transcriptional regulator Mlc | -0,09 | 0,62 | 0,55 | 0,71 | 0,64 | -0,07 | 1,0E-01 | 1,4E-06 | 4,7E-05 | 5,3E-06 | 5,7E-05 | 4,1E-01 | 0,0E+00 | 0,0E+00 | 0,0E+00 | 0,0E+00 | 0,0E+00 | 0,0E+00 | 1010198 | 1011415 | 1218 | - |
| Y11_RS04590 | Y11_09871 | | LysR family transcriptional regulator | 0,29 | 1,18 | 1,2 | 0,89 | 0,91 | 0,02 | 5,3E-02 | 6,2E-06 | 1,5E-05 | 3,4E-05 | 8,3E-05 | 8,5E-01 | 0,0E+00 | 1,5E-07 | 2,4E-07 | 0,0E+00 | 0,0E+00 | 0,0E+00 | 1011636 | 1012547 | 912 | - |
| Y11_RS04595 | Y11_09881 | | MFS transporter | -0,26 | 2,55 | 2,32 | 2,73 | 2,57 | -0,16 | 1,2E-01 | 1,8E-06 | 2,1E-05 | 1,2E-06 | 1,1E-05 | 6,1E-01 | 0,0E+00 | 4,9E-08 | 3,4E-07 | 2,9E-08 | 3,7E-07 | 0,0E+00 | 1012723 | 1014015 | 1293 | + |
| Y11_RS04600 | | | SHOCT domain-containing protein | 0,13 | 1,14 | 2,23 | 1,01 | 2,1 | 1,09 | 1,7E-01 | 7,1E-08 | 6,4E-05 | 1,0E-06 | 1,1E-04 | 5,8E-03 | 0,0E+00 | 3,1E-09 | 9,1E-07 | 2,6E-08 | 2,8E-06 | 3,4E-04 | 1014203 | 1014592 | 390 | + |
| Y11_RS04605 | Y11_09911 | blaA | class A beta-lactamase BlaA | 0,45 | -1,37 | -3,01 | -1,83 | -3,46 | -1,63 | 9,4E-05 | 1,2E-06 | 2,2E-12 | 1,5E-07 | 1,4E-12 | 1,6E-07 | 0,0E+00 | 3,4E-08 | 3,9E-12 | 4,7E-09 | 5,4E-12 | 5,0E-08 | 1014817 | 1015701 | 885 | + |
| Y11_RS04610 | Y11_09921 | tus | DNA replication terminus site-binding protein | 0,47 | 0,71 | 1,33 | 0,24 | 0,86 | 0,62 | 3,0E-03 | 7,5E-04 | 1,6E-06 | 8,1E-02 | 1,1E-05 | 7,4E-04 | 0,0E+00 | 0,0E+00 | 3,4E-08 | 0,0E+00 | 0,0E+00 | 0,0E+00 | 1015864 | 1016793 | 930 | + |
| Y11_RS04625 | Y11_09961 | fumC | class II fumarate hydratase | -0,41 | 0,01 | -0,16 | 0,42 | 0,25 | -0,17 | 7,0E-05 | 8,2E-01 | 1,5E-02 | 4,0E-04 | 8,3E-03 | 4,2E-02 | 0,0E+00 | 0,0E+00 | 0,0E+00 | 0,0E+00 | 0,0E+00 | 0,0E+00 | 1019471 | 1020865 | 1395 | - |
| Y11_RS04630 | Y11_09971 | manA | mannose-6-phosphate isomerase | 0,38 | -0,36 | -0,32 | -0,74 | -0,7 | 0,04 | 9,8E-05 | 1,1E-04 | 1,6E-03 | 2,4E-07 | 5,0E-06 | 5,8E-01 | 0,0E+00 | 0,0E+00 | 0,0E+00 | 0,0E+00 | 0,0E+00 | 0,0E+00 | 1021091 | 1022266 | 1176 | + |
| Y11_RS04635 | Y11_09981 | | YdgA family protein | 0,31 | -0,62 | -0,6 | -0,92 | -0,9 | 0,02 | 6,4E-04 | 1,7E-04 | 4,8E-03 | 5,7E-06 | 3,3E-04 | 8,9E-01 | 0,0E+00 | 0,0E+00 | 0,0E+00 | 0,0E+00 | 0,0E+00 | 0,0E+00 | 1022498 | 1024114 | 1617 | + |
| Y11_RS04655 | Y11_10021 | | pyridoxal phosphate-dependent aminotransferase | -0,18 | 1,37 | 1,59 | 1,56 | 1,77 | 0,22 | 2,6E-01 | 2,9E-06 | 4,4E-07 | 3,1E-05 | 8,7E-06 | 1,9E-01 | 0,0E+00 | 7,8E-08 | 1,1E-08 | 5,9E-07 | 2,9E-07 | 0,0E+00 | 1026228 | 1027406 | 1179 | + |
| Y11_RS04660 | Y11_10031 | add | adenosine deaminase | -0,48 | -1,08 | -1,86 | -0,59 | -1,38 | -0,79 | 5,9E-05 | 3,6E-07 | 9,5E-09 | 1,9E-05 | 6,3E-08 | 9,8E-06 | 0,0E+00 | 1,2E-08 | 5,0E-10 | 0,0E+00 | 4,2E-09 | 0,0E+00 | 1027586 | 1028584 | 999 | + |
| Y11_RS04665 | Y11_10041 | | bile acid:sodium symporter | 0,06 | 0,77 | 0,58 | 0,71 | 0,52 | -0,19 | 3,2E-01 | 5,0E-05 | 2,8E-06 | 1,4E-04 | 3,2E-05 | 1,1E-01 | 0,0E+00 | 0,0E+00 | 0,0E+00 | 0,0E+00 | 0,0E+00 | 0,0E+00 | 1028734 | 1029747 | 1014 | - |
| Y11_RS04670 | Y11_10051 | | oxidoreductase | -0,05 | 1,04 | 1,23 | 1,09 | 1,27 | 0,19 | 1,3E-01 | 2,8E-08 | 3,1E-06 | 1,8E-08 | 2,3E-06 | 1,5E-01 | 0,0E+00 | 1,4E-09 | 6,0E-08 | 8,1E-10 | 8,7E-08 | 0,0E+00 | 1029797 | 1030846 | 1050 | - |
| Y11_RS04675 | Y11_10061 | araC | arabinose operon transcriptional regulator AraC | -0,26 | 0,1 | -0,09 | 0,35 | 0,16 | -0,18 | 3,4E-01 | 7,3E-01 | 7,0E-01 | 2,3E-01 | 4,9E-01 | 5,0E-01 | 0,0E+00 | 0,0E+00 | 0,0E+00 | 0,0E+00 | 0,0E+00 | 0,0E+00 | 1030948 | 1031880 | 933 | - |
| Y11_RS04690 | Y11_10101 | | arabinose ABC transporter substrate-binding protein | -1,16 | -2,14 | -2,2 | -0,98 | -1,04 | -0,06 | 1,6E-04 | 4,0E-07 | 2,0E-06 | 5,5E-05 | 2,7E-04 | 6,9E-01 | 6,7E-06 | 1,3E-08 | 4,2E-08 | 0,0E+00 | 6,8E-06 | 0,0E+00 | 1034630 | 1035613 | 984 | - |

| | | | | | | | | | | | | | | | | | | | | | | | | | |
|---|---|---|---|---|---|---|---|---|---|---|---|---|---|---|---|---|---|---|---|---|---|---|---|---|---|
| Y11_RS04695 | Y11_10111 | | ribulokinase | -0,28 | 15 | 15 | 15 | 15 | 0 | 5,8E-02 | 0,0E+00 | 0,0E+00 | 0,0E+00 | 0,0E+00 | 0,0E+00 | 0,0E+00 | 0,0E+00 | 0,0E+00 | 0,0E+00 | 0,0E+00 | 0,0E+00 | 1036192 | 1037907 | 1716 | + |
| Y11_RS04700 | Y11_10121 | araA | L-arabinose isomerase | -0,07 | 0,75 | 0,64 | 0,82 | 0,71 | -0,12 | 4,7E-01 | 1,6E-05 | 2,7E-04 | 3,9E-07 | 2,9E-05 | 1,4E-01 | 0,0E+00 | 0,0E+00 | 0,0E+00 | 0,0E+00 | 0,0E+00 | 0,0E+00 | 1037965 | 1039467 | 1503 | + |
| Y11_RS04730 | Y11_10191 | | formate--tetrahydrofolate ligase | -0,72 | 2,88 | 2,59 | 3,6 | 3,29 | -0,3 | 2,4E-05 | 3,9E-10 | 2,7E-10 | 1,3E-10 | 7,4E-11 | 3,6E-03 | 0,0E+00 | 5,4E-11 | 4,2E-11 | 2,1E-11 | 3,9E-11 | 0,0E+00 | 1045980 | 1047737 | 1758 | + |
| Y11_RS04770 | Y11_10291 | rsxG | electron transport complex subunit RsxG | -0,06 | 15 | 15 | 15 | 15 | 0 | 8,4E-01 | 0,0E+00 | 0,0E+00 | 0,0E+00 | 0,0E+00 | 0,0E+00 | 0,0E+00 | 0,0E+00 | 0,0E+00 | 0,0E+00 | 0,0E+00 | 0,0E+00 | 1054949 | 1055578 | 630 | + |
| Y11_RS04775 | Y11_10301 | | electron transport complex subunit E | -1,57 | 0 | 0 | 15 | 15 | 0 | 4,2E-07 | 0,0E+00 | 0,0E+00 | 0,0E+00 | 0,0E+00 | 0,0E+00 | 3,0E-08 | 0,0E+00 | 0,0E+00 | 0,0E+00 | 0,0E+00 | 0,0E+00 | 1055575 | 1056276 | 702 | + |
| Y11_RS04780 | Y11_10311 | nth | endonuclease III | 0,39 | 15 | 4,92 | 15 | 4,54 | 0 | 5,6E-02 | 0,0E+00 | 9,9E-12 | 0,0E+00 | 4,7E-09 | 0,0E+00 | 0,0E+00 | 0,0E+00 | 7,4E-12 | 0,0E+00 | 5,3E-10 | 0,0E+00 | 1056273 | 1056914 | 642 | + |
| Y11_RS04785 | Y11_10321 | yaxB | pore-forming cytotoxin subunit YaxB | -2,57 | 2,34 | 1,78 | 4,89 | 4,35 | 0 | 1,9E-09 | 1,3E-08 | 1,6E-08 | 4,7E-12 | 3,8E-13 | 0,0E+00 | 4,0E-10 | 7,5E-10 | 7,4E-10 | 4,1E-12 | 2,4E-12 | 0,0E+00 | 1057001 | 1058038 | 1038 | - |
| Y11_RS04790 | Y11_10331 | | alpha-xenorhabdolysin family binary toxin subunit A | -2,49 | 2,11 | 2,38 | 4,6 | 4,87 | 0 | 3,1E-07 | 2,6E-06 | 8,1E-07 | 5,6E-11 | 1,8E-11 | 0,0E+00 | 2,3E-08 | 6,9E-08 | 1,9E-08 | 1,2E-11 | 1,6E-11 | 0,0E+00 | 1058083 | 1059318 | 1236 | - |
| Y11_RS04810 | Y11_10381 | | exoribonuclease II | 0,07 | 15 | 2,6 | 15 | 2,52 | 0 | 6,0E-01 | 0,0E+00 | 7,3E-09 | 0,0E+00 | 9,6E-08 | 0,0E+00 | 0,0E+00 | 0,0E+00 | 4,3E-10 | 0,0E+00 | 6,0E-09 | 0,0E+00 | 1063924 | 1065858 | 1935 | + |
| Y11_RS04815 | Y11_10401 | | carbon starvation protein A | 2,39 | 2,34 | 15 | -0,05 | 15 | 0 | 5,5E-04 | 5,3E-04 | 0,0E+00 | 6,4E-01 | 0,0E+00 | 0,0E+00 | 2,1E-05 | 1,1E-05 | 0,0E+00 | 0,0E+00 | 0,0E+00 | 0,0E+00 | 1066273 | 1068339 | 2067 | + |
| Y11_RS04825 | Y11_10421 | fsa | fructose-6-phosphate aldolase | -0,6 | 4,69 | 4,86 | 5,3 | 5,46 | 0,18 | 3,6E-04 | 1,0E-10 | 1,8E-10 | 1,8E-11 | 3,8E-11 | 1,6E-01 | 0,0E+00 | 2,3E-11 | 3,1E-11 | 6,6E-12 | 2,4E-11 | 0,0E+00 | 1068900 | 1069565 | 666 | - |
| Y11_RS04830 | Y11_10431 | | DeoR/GlpR family DNA-binding transcription regulator | -0,65 | 15 | 15 | 15 | 15 | 0 | 1,6E-02 | 0,0E+00 | 0,0E+00 | 0,0E+00 | 0,0E+00 | 0,0E+00 | 0,0E+00 | 0,0E+00 | 0,0E+00 | 0,0E+00 | 0,0E+00 | 0,0E+00 | 1069761 | 1070507 | 747 | + |
| Y11_RS04835 | Y11_10441 | | DUF2164 domain-containing protein | -0,77 | 15 | 15 | 15 | 15 | 0 | 5,7E-03 | 0,0E+00 | 0,0E+00 | 0,0E+00 | 0,0E+00 | 0,0E+00 | 0,0E+00 | 0,0E+00 | 0,0E+00 | 0,0E+00 | 0,0E+00 | 0,0E+00 | 1070527 | 1070775 | 249 | + |
| Y11_RS04845 | Y11_10461 | osmB | osmotically-inducible lipoprotein OsmB | 0,87 | 15 | 15 | 15 | 15 | 0 | 3,2E-02 | 0,0E+00 | 0,0E+00 | 0,0E+00 | 0,0E+00 | 0,0E+00 | 0,0E+00 | 0,0E+00 | 0,0E+00 | 0,0E+00 | 0,0E+00 | 0,0E+00 | 1071933 | 1072148 | 216 | + |
| Y11_RS04855 | Y11_10491 | yciH | stress response translation initiation inhibitor YciH | 0,17 | -0,29 | -0,41 | -0,46 | -0,58 | -0,12 | 4,3E-01 | 5,1E-02 | 2,2E-02 | 4,5E-02 | 2,3E-02 | 3,7E-01 | 0,0E+00 | 0,0E+00 | 0,0E+00 | 0,0E+00 | 0,0E+00 | 0,0E+00 | 1073900 | 1074226 | 327 | - |
| Y11_RS04860 | Y11_10501 | pyrF | orotidine-5'-phosphate decarboxylase | 0,08 | -0,42 | 0,14 | -0,5 | 0,06 | 0,56 | 1,8E-01 | 1,3E-04 | 2,4E-01 | 3,2E-05 | 5,9E-01 | 1,0E-03 | 0,0E+00 | 0,0E+00 | 0,0E+00 | 0,0E+00 | 0,0E+00 | 0,0E+00 | 1074223 | 1074963 | 741 | - |
| Y11_RS04865 | Y11_10511 | | DUF1471 domain-containing protein | 15 | -7,58 | -2,1 | -15 | -15 | 5,48 | 0,0E+00 | 8,4E-12 | 2,2E-06 | 0,0E+00 | 0,0E+00 | 2,9E-09 | 0,0E+00 | 5,6E-12 | 4,6E-08 | 0,0E+00 | 0,0E+00 | 3,2E-09 | 1075314 | 1075574 | 261 | + |
| Y11_RS04880 | Y11_10551 | | hypothetical protein | 1,24 | -1,11 | -1,75 | -2,35 | -2,99 | -0,64 | 6,0E-04 | 3,4E-05 | 1,5E-05 | 2,1E-06 | 1,4E-06 | 2,4E-03 | 2,3E-05 | 7,8E-07 | 2,3E-07 | 4,8E-08 | 5,5E-08 | 0,0E+00 | 1077757 | 1078044 | 288 | + |
| Y11_RS04885 | Y11_10561 | lapB | lipopolysaccharide assembly protein LapB | -0,16 | 0,63 | 0,63 | 0,79 | 0,79 | 0 | 2,5E-01 | 1,7E-03 | 1,8E-03 | 7,1E-05 | 7,9E-05 | 1,0E+00 | 0,0E+00 | 0,0E+00 | 0,0E+00 | 0,0E+00 | 0,0E+00 | 0,0E+00 | 1078105 | 1079274 | 1170 | - |
| Y11_RS04890 | Y11_10571 | | LapA family protein | 0,07 | 0,65 | 0,61 | 0,58 | 0,54 | -0,05 | 6,2E-01 | 1,2E-03 | 4,3E-02 | 3,5E-03 | 7,0E-02 | 8,6E-01 | 0,0E+00 | 0,0E+00 | 0,0E+00 | 0,0E+00 | 0,0E+00 | 0,0E+00 | 1079281 | 1079592 | 312 | - |
| Y11_RS04900 | Y11_10591 | ribA | GTP cyclohydrolase II | -0,56 | -0,17 | -0,28 | 0,39 | 0,29 | -0,1 | 3,1E-04 | 6,3E-02 | 2,9E-02 | 3,9E-04 | 1,6E-02 | 2,4E-01 | 0,0E+00 | 0,0E+00 | 0,0E+00 | 0,0E+00 | 0,0E+00 | 0,0E+00 | 1081177 | 1081767 | 591 | + |
| Y11_RS04910 | Y11_10601 | acnA | aconitate hydratase AcnA | -0,71 | 1,12 | 1,73 | 1,83 | 2,44 | 0,61 | 1,9E-08 | 6,3E-09 | 2,3E-10 | 3,1E-11 | 3,7E-12 | 1,4E-06 | 0,0E+00 | 4,4E-10 | 3,7E-11 | 8,5E-12 | 7,5E-12 | 0,0E+00 | 1082055 | 1084727 | 2673 | - |
| Y11_RS04920 | Y11_10631 | cysB | HTH-type transcriptional regulator CysB | -0,15 | 0,23 | 0,69 | 0,38 | 0,84 | 0,46 | 7,5E-02 | 4,1E-04 | 6,6E-08 | 1,4E-03 | 5,0E-06 | 1,0E-05 | 0,0E+00 | 0,0E+00 | 0,0E+00 | 0,0E+00 | 0,0E+00 | 0,0E+00 | 1086351 | 1087325 | 975 | - |
| Y11_RS04925 | Y11_10641 | topA | type I DNA topoisomerase | 0,57 | 1,09 | -0,12 | 0,52 | -0,69 | -1,21 | 9,3E-08 | 6,6E-08 | 1,6E-02 | 2,3E-05 | 1,9E-07 | 6,4E-08 | 0,0E+00 | 2,9E-09 | 0,0E+00 | 0,0E+00 | 0,0E+00 | 2,6E-08 | 1087884 | 1090499 | 2616 | - |
| Y11_RS04930 | Y11_10661 | | YciN family protein | -0,14 | 0,49 | 1,09 | 0,63 | 1,22 | 0,6 | 1,6E-01 | 5,7E-04 | 2,0E-03 | 5,1E-05 | 8,8E-04 | 3,7E-02 | 0,0E+00 | 0,0E+00 | 2,3E-05 | 0,0E+00 | 2,0E-05 | 0,0E+00 | 1090940 | 1091191 | 252 | + |
| Y11_RS04935 | Y11_10671 | sohB | protease SohB | 0 | -1,48 | -1,41 | -1,48 | -1,41 | 0,07 | 9,9E-01 | 3,2E-07 | 4,2E-04 | 1,1E-07 | 3,7E-04 | 7,6E-01 | 0,0E+00 | 1,1E-08 | 5,3E-06 | 3,6E-09 | 8,9E-06 | 0,0E+00 | 1091296 | 1092342 | 1047 | - |
| Y11_RS04940 | Y11_10681 | | YciK family oxidoreductase | 0,87 | 0,56 | 0,54 | -0,32 | -0,33 | -0,02 | 4,0E-06 | 7,8E-08 | 1,5E-03 | 3,2E-03 | 3,8E-02 | 8,9E-01 | 0,0E+00 | 0,0E+00 | 0,0E+00 | 0,0E+00 | 0,0E+00 | 0,0E+00 | 1092515 | 1093276 | 762 | + |
| Y11_RS04945 | Y11_10691 | cobO | cob(I)yrinic acid a,c-diamide adenosyltransferase | 0,68 | 0,15 | 0,53 | -0,52 | -0,15 | 0,37 | 9,5E-05 | 9,8E-02 | 2,4E-03 | 7,8E-05 | 2,2E-01 | 6,8E-03 | 0,0E+00 | 0,0E+00 | 0,0E+00 | 0,0E+00 | 0,0E+00 | 0,0E+00 | 1093288 | 1093878 | 591 | + |
| Y11_RS04955 | Y11_10711 | rluB | 23S rRNA pseudouridine(2605) synthase RluB | -0,09 | -0,49 | -1,21 | -0,41 | -1,12 | -0,71 | 5,3E-02 | 5,4E-06 | 4,6E-08 | 3,4E-05 | 1,1E-07 | 6,2E-06 | 0,0E+00 | 0,0E+00 | 1,8E-09 | 0,0E+00 | 6,5E-09 | 0,0E+00 | 1094474 | 1095523 | 1050 | - |
| Y11_RS04960 | Y11_10721 | | threonylcarbamoyl-AMP synthase | 0,05 | 1,2 | 0,75 | 1,15 | 0,7 | -0,45 | 3,9E-01 | 9,6E-08 | 1,1E-03 | 1,8E-07 | 1,8E-03 | 2,1E-02 | 0,0E+00 | 4,0E-09 | 0,0E+00 | 5,7E-09 | 0,0E+00 | 0,0E+00 | 1095764 | 1096384 | 621 | - |
| Y11_RS04965 | Y11_10731 | | PHP domain-containing protein | 0,1 | 0,77 | -0,01 | 0,67 | -0,12 | -0,78 | 8,0E-01 | 1,3E-01 | 9,7E-01 | 3,2E-02 | 5,4E-01 | 2,5E-02 | 0,0E+00 | 0,0E+00 | 0,0E+00 | 0,0E+00 | 0,0E+00 | 0,0E+00 | 1096413 | 1097330 | 918 | - |
| Y11_RS04970 | Y11_10751 | | anthranilate synthase component 1 | 0,02 | -2,79 | -3,29 | -2,81 | -3,31 | -0,5 | 8,7E-01 | 2,5E-08 | 7,0E-09 | 2,6E-09 | 7,4E-10 | 1,9E-03 | 0,0E+00 | 1,3E-09 | 4,1E-10 | 1,8E-10 | 1,4E-10 | 0,0E+00 | 1097888 | 1099450 | 1563 | + |
| Y11_RS04975 | Y11_10761 | | glutamine amidotransferase | 0 | -0,86 | -2,23 | -15 | -15 | -1,43 | 0,0E+00 | 6,2E-02 | 9,2E-07 | 0,0E+00 | 0,0E+00 | 8,6E-03 | 0,0E+00 | 0,0E+00 | 2,1E-08 | 0,0E+00 | 0,0E+00 | 5,0E-04 | 1099465 | 1100055 | 591 | + |
| Y11_RS04980 | Y11_10771 | trpD | anthranilate phosphoribosyltransferase | -0,16 | -1,35 | -0,97 | -1,19 | -0,8 | 0,38 | 6,3E-01 | 2,1E-03 | 1,9E-02 | 1,6E-03 | 2,4E-02 | 1,9E-01 | 0,0E+00 | 4,2E-05 | 0,0E+00 | 2,6E-05 | 0,0E+00 | 0,0E+00 | 1100116 | 1101132 | 1017 | + |
| Y11_RS04985 | Y11_10781 | trpCF | bifunctional indole-3-glycerol-phosphate synthase TrpC/phosphoribosylanthranilate isomerase TrpF | 0,51 | -1,55 | -1,23 | -2,06 | -1,74 | 0,32 | 4,3E-05 | 6,0E-11 | 7,7E-08 | 8,3E-10 | 3,5E-08 | 1,3E-03 | 0,0E+00 | 1,6E-11 | 2,6E-09 | 7,6E-11 | 2,6E-09 | 0,0E+00 | 1101136 | 1102512 | 1377 | + |
| Y11_RS04990 | Y11_10791 | trpB | tryptophan synthase subunit beta | 0,24 | -1,4 | -1,33 | -1,64 | -1,56 | 0,07 | 6,7E-03 | 3,0E-07 | 1,2E-06 | 4,5E-08 | 2,1E-07 | 5,5E-01 | 0,0E+00 | 1,0E-08 | 2,8E-08 | 1,7E-09 | 1,2E-08 | 0,0E+00 | 1102712 | 1103902 | 1191 | + |
| Y11_RS04995 | Y11_10801 | trpA | tryptophan synthase subunit alpha | 0,23 | -0,66 | -0,54 | -0,89 | -0,77 | 0,12 | 5,1E-02 | 9,1E-05 | 2,1E-03 | 4,3E-05 | 4,7E-04 | 3,7E-01 | 0,0E+00 | 0,0E+00 | 0,0E+00 | 0,0E+00 | 0,0E+00 | 0,0E+00 | 1103902 | 1104708 | 807 | + |
| Y11_RS05000 | Y11_10811 | | BON domain-containing protein | 2,05 | 2,95 | 4,08 | 0,9 | 2,03 | 1,12 | 3,9E-08 | 5,1E-10 | 1,5E-11 | 2,4E-05 | 2,8E-08 | 5,6E-07 | 4,0E-09 | 6,7E-11 | 8,0E-12 | 0,0E+00 | 2,2E-09 | 1,2E-07 | 1104965 | 1105279 | 315 | + |
| Y11_RS05005 | Y11_10821 | ompW | outer membrane protein OmpW | 0,92 | -2,55 | -1,52 | -3,47 | -2,44 | 1,03 | 1,8E-03 | 4,2E-06 | 9,5E-04 | 5,5E-07 | 4,5E-05 | 1,4E-02 | 0,0E+00 | 1,1E-07 | 1,1E-05 | 1,5E-08 | 1,3E-06 | 7,7E-04 | 1105405 | 1106040 | 636 | - |
| Y11_RS05015 | Y11_10841 | | envelope biogenesis factor ElyC | -0,34 | -0,86 | -0,39 | -0,52 | -0,06 | 0,47 | 1,9E-01 | 3,1E-04 | 2,5E-01 | 8,1E-02 | 8,9E-01 | 2,0E-01 | 0,0E+00 | 0,0E+00 | 0,0E+00 | 0,0E+00 | 0,0E+00 | 0,0E+00 | 1106795 | 1107562 | 768 | + |
| Y11_RS05030 | Y11_10871 | yciA | acyl-CoA thioester hydrolase YciA | -0,54 | -1,06 | -1,09 | -0,52 | -0,55 | -0,03 | 1,5E-02 | 4,6E-03 | 8,3E-05 | 1,0E-01 | 1,2E-02 | 9,1E-01 | 0,0E+00 | 9,0E-05 | 1,2E-06 | 0,0E+00 | 0,0E+00 | 0,0E+00 | 1110141 | 1110611 | 471 | + |
| Y11_RS05035 | Y11_10881 | tonB | TonB system transport protein TonB | 0,98 | -1,07 | -1,96 | -2,05 | -2,94 | -0,89 | 2,0E-04 | 2,4E-06 | 3,2E-07 | 1,2E-06 | 2,1E-07 | 1,6E-04 | 0,0E+00 | 6,5E-08 | 8,7E-09 | 2,9E-08 | 1,2E-08 | 0,0E+00 | 1110707 | 1111462 | 756 | - |
| Y11_RS05040 | Y11_10891 | | YciI family protein | -0,25 | 0,29 | 0,08 | 0,54 | 0,34 | -0,21 | 4,1E-01 | 3,9E-01 | 8,1E-01 | 5,9E-02 | 2,4E-01 | 5,0E-01 | 0,0E+00 | 0,0E+00 | 0,0E+00 | 0,0E+00 | 0,0E+00 | 0,0E+00 | 1111771 | 1112067 | 297 | + |
| Y11_RS05045 | Y11_10901 | | hypothetical protein | -0,53 | 0,04 | 1,69 | 0,57 | 2,23 | 1,66 | 3,8E-02 | 8,4E-01 | 8,9E-05 | 8,1E-03 | 1,0E-05 | 2,0E-05 | 0,0E+00 | 0,0E+00 | 1,2E-06 | 0,0E+00 | 3,3E-07 | 2,1E-06 | 1112269 | 1112532 | 264 | - |
| Y11_RS05050 | Y11_10911 | | EAL domain-containing protein | 0 | -0,51 | 0,51 | -15 | -15 | 1,07 | 0,0E+00 | 7,6E-02 | 1,5E-01 | 0,0E+00 | 0,0E+00 | 4,6E-03 | 0,0E+00 | 0,0E+00 | 0,0E+00 | 0,0E+00 | 0,0E+00 | 2,8E-04 | 1112684 | 1113406 | 723 | - |
| Y11_RS05055 | Y11_10931 | cls | cardiolipin synthase | 0,03 | -0,03 | 0,39 | -0,06 | 0,36 | 0,42 | 6,7E-01 | 7,0E-01 | 1,8E-02 | 3,7E-01 | 2,0E-02 | 1,2E-02 | 0,0E+00 | 0,0E+00 | 0,0E+00 | 0,0E+00 | 0,0E+00 | 0,0E+00 | 1114472 | 1115932 | 1461 | + |
| Y11_RS05060 | Y11_10941 | | HI1450 family dsDNA-mimic protein | 1,97 | -0,63 | 0,47 | -2,6 | -1,51 | 1,09 | 2,9E-04 | 1,3E-01 | 2,4E-01 | 4,3E-04 | 9,3E-03 | 5,3E-02 | 1,2E-05 | 0,0E+00 | 0,0E+00 | 7,5E-06 | 2,0E-04 | 2,8E-03 | 1116061 | 1116390 | 330 | + |
| Y11_RS05065 | Y11_10951 | oppF | murein tripeptide/oligopeptide ABC transporter ATP binding protein OppF | -0,29 | -0,73 | -0,18 | -0,45 | 0,11 | 0,56 | 1,8E-05 | 2,6E-07 | 6,1E-02 | 9,8E-06 | 2,2E-01 | 2,3E-04 | 0,0E+00 | 0,0E+00 | 0,0E+00 | 0,0E+00 | 0,0E+00 | 0,0E+00 | 1116537 | 1117538 | 1002 | - |
| Y11_RS05070 | Y11_10961 | | ABC transporter ATP-binding protein | -0,35 | -0,82 | -0,41 | -0,47 | -0,06 | 0,41 | 2,2E-05 | 5,2E-09 | 1,7E-03 | 8,7E-06 | 5,4E-01 | 2,1E-03 | 0,0E+00 | 0,0E+00 | 0,0E+00 | 0,0E+00 | 0,0E+00 | 0,0E+00 | 1117535 | 1118536 | 1002 | - |
| Y11_RS05075 | Y11_10971 | oppC | oligopeptide ABC transporter permease OppC | -0,33 | -0,68 | -0,15 | -0,35 | 0,18 | 0,53 | 1,7E-03 | 2,5E-06 | 2,8E-01 | 4,3E-04 | 2,0E-01 | 2,2E-03 | 0,0E+00 | 0,0E+00 | 0,0E+00 | 0,0E+00 | 0,0E+00 | 0,0E+00 | 1118548 | 1119456 | 909 | - |
| Y11_RS05080 | Y11_10981 | oppB | oligopeptide ABC transporter permease OppB | -0,06 | -0,36 | 0,07 | -0,29 | 0,13 | 0,42 | 7,0E-01 | 7,8E-02 | 6,7E-01 | 2,0E-01 | 5,1E-01 | 6,9E-02 | 0,0E+00 | 0,0E+00 | 0,0E+00 | 0,0E+00 | 0,0E+00 | 0,0E+00 | 1119472 | 1120392 | 921 | - |
| Y11_RS05085 | Y11_10991 | oppA | oligopeptide ABC transporter substrate-binding protein OppA | -0,26 | -1,68 | -1,08 | -1,42 | -0,82 | 0,6 | 1,1E-03 | 9,9E-10 | 1,8E-06 | 1,7E-10 | 6,4E-06 | 6,4E-05 | 0,0E+00 | 1,1E-10 | 3,8E-08 | 2,5E-11 | 0,0E+00 | 0,0E+00 | 1120477 | 1122114 | 1638 | - |
| Y11_RS05090 | Y11_11001 | | ABC transporter substrate-binding protein | 0,45 | -0,91 | -0,17 | -1,36 | -0,62 | 0,74 | 2,0E-02 | 1,5E-05 | 2,1E-01 | 2,8E-05 | 7,7E-03 | 4,0E-04 | 0,0E+00 | 0,0E+00 | 0,0E+00 | 5,5E-07 | 0,0E+00 | 0,0E+00 | 1122343 | 1123980 | 1638 | - |
| Y11_RS05100 | Y11_11021 | adhE | bifunctional acetaldehyde-CoA/alcohol dehydrogenase | -0,21 | 1,41 | 0,18 | 1,61 | 0,39 | -1,22 | 1,3E-04 | 2,9E-10 | 1,0E-02 | 1,2E-11 | 4,5E-05 | 1,5E-08 | 0,0E+00 | 4,4E-11 | 0,0E+00 | 5,6E-12 | 0,0E+00 | 9,3E-09 | 1126064 | 1128739 | 2676 | + |
| Y11_RS05105 | Y11_11031 | | thymidine kinase | -1,49 | -0,91 | -1,59 | 0,58 | -0,1 | -0,68 | 6,6E-09 | 3,9E-06 | 2,0E-05 | 4,4E-05 | 5,7E-01 | 6,0E-03 | 1,0E-09 | 0,0E+00 | 3,2E-07 | 0,0E+00 | 0,0E+00 | 0,0E+00 | 1129238 | 1129837 | 600 | - |
| Y11_RS05115 | Y11_11041 | hns | DNA-binding transcriptional regulator H-NS | -0,32 | 0,27 | -0,06 | 0,59 | 0,26 | -0,33 | 3,0E-02 | 2,8E-02 | 6,7E-01 | 3,9E-04 | 8,9E-02 | 2,2E-02 | 0,0E+00 | 0,0E+00 | 0,0E+00 | 0,0E+00 | 0,0E+00 | 0,0E+00 | 1130589 | 1130999 | 411 | + |
| Y11_RS05120 | Y11_11051 | | UDP-glucose/GDP-mannose dehydrogenase family protein | -1,25 | -0,86 | -2,76 | 0,4 | -1,5 | -1,9 | 6,6E-06 | 1,7E-04 | 3,5E-09 | 3,5E-02 | 3,6E-06 | 9,1E-07 | 3,5E-07 | 0,0E+00 | 2,4E-10 | 0,0E+00 | 1,3E-07 | 1,7E-07 | 1131384 | 1132733 | 1350 | - |
| Y11_RS05130 | Y11_11081 | rssA | patatin-like phospholipase RssA | -0,97 | -1,41 | -0,02 | -0,45 | 0,94 | 1,39 | 1,6E-06 | 2,9E-06 | 8,9E-01 | 3,9E-03 | 1,8E-04 | 4,2E-05 | 0,0E+00 | 7,6E-08 | 0,0E+00 | 0,0E+00 | 0,0E+00 | 3,7E-06 | 1134174 | 1135076 | 903 | - |
| Y11_RS05145 | Y11_11131 | purU | formyltetrahydrofolate deformylase | 0,08 | -0,01 | -0,1 | -0,09 | -0,18 | -0,09 | 1,5E-01 | 9,1E-01 | 4,1E-01 | 2,5E-01 | 1,5E-01 | 4,9E-01 | 0,0E+00 | 0,0E+00 | 0,0E+00 | 0,0E+00 | 0,0E+00 | 0,0E+00 | 1136996 | 1137844 | 849 | + |
| Y11_RS05160 | Y11_11161 | xthA | exodeoxyribonuclease III | 0,01 | -0,13 | -0,58 | -0,14 | -0,6 | -0,46 | 8,6E-01 | 1,6E-01 | 3,3E-05 | 2,3E-01 | 3,4E-04 | 1,9E-03 | 0,0E+00 | 0,0E+00 | 0,0E+00 | 0,0E+00 | 0,0E+00 | 0,0E+00 | 1140041 | 1140847 | 807 | + |
| Y11_RS05165 | Y11_11171 | | pyrimidine (deoxy)nucleoside triphosphate diphosphatase | -0,38 | -0,06 | 0,86 | 0,32 | 1,31 | 0,91 | 4,8E-02 | 8,1E-01 | 7,8E-02 | 1,2E-01 | 1,1E-02 | 7,3E-02 | 0,0E+00 | 0,0E+00 | 0,0E+00 | 0,0E+00 | 2,4E-04 | 0,0E+00 | 1140931 | 1141317 | 387 | + |
| Y11_RS05170 | | | DUF1496 domain-containing protein | 0,5 | -0,97 | -1,04 | -1,47 | -1,55 | -0,08 | 1,1E-04 | 1,9E-06 | 5,5E-07 | 1,4E-07 | 5,5E-08 | 4,0E-01 | 0,0E+00 | 0,0E+00 | 1,4E-08 | 4,6E-09 | 3,8E-09 | 0,0E+00 | 1141322 | 1141612 | 291 | - |
| Y11_RS05175 | Y11_11191 | | DNA topoisomerase III | 0,2 | -0,18 | -0,41 | -0,38 | -0,62 | -0,24 | 2,5E-01 | 1,1E-02 | 1,4E-04 | 5,3E-02 | 6,7E-03 | 1,3E-02 | 0,0E+00 | 0,0E+00 | 0,0E+00 | 0,0E+00 | 0,0E+00 | 0,0E+00 | 1141609 | 1143534 | 1926 | - |
| Y11_RS05180 | Y11_11201 | selD | selenide, water dikinase SelD | 0,09 | 0 | 0,6 | -0,1 | 0,51 | 0,6 | 2,3E-01 | 9,7E-01 | 2,7E-05 | 2,3E-01 | 6,3E-05 | 3,8E-05 | 0,0E+00 | 0,0E+00 | 0,0E+00 | 0,0E+00 | 0,0E+00 | 0,0E+00 | 1143581 | 1144627 | 1047 | - |
| Y11_RS05185 | Y11_11211 | | NAD(P)H nitroreductase | -0,59 | 0,38 | -0,2 | 0,97 | 0,39 | -0,58 | 5,5E-06 | 3,7E-04 | 5,4E-02 | 2,0E-07 | 1,7E-03 | 2,1E-04 | 0,0E+00 | 0,0E+00 | 0,0E+00 | 0,0E+00 | 0,0E+00 | 0,0E+00 | 1144989 | 1145540 | 552 | - |
| Y11_RS05190 | Y11_11221 | sppA | signal peptide peptidase SppA | -0,65 | -0,23 | -0,49 | 0,42 | 0,16 | -0,26 | 3,1E-08 | 7,1E-05 | 3,8E-04 | 7,3E-07 | 9,3E-02 | 1,4E-02 | 0,0E+00 | 0,0E+00 | 0,0E+00 | 0,0E+00 | 0,0E+00 | 0,0E+00 | 1145703 | 1147556 | 1854 | + |
| Y11_RS05195 | Y11_11231 | ansA | asparaginase | 0,07 | -1,46 | -1,46 | -1,53 | -1,53 | -0,01 | 4,1E-01 | 3,7E-10 | 1,3E-09 | 4,4E-08 | 6,3E-08 | 8,8E-01 | 0,0E+00 | 5,3E-11 | 1,2E-10 | 1,7E-09 | 4,2E-09 | 0,0E+00 | 1147797 | 1148813 | 1017 | + |
| Y11_RS05215 | Y11_11271 | | DUF1315 family protein | -0,4 | -2,3 | -2,81 | -1,9 | -2,41 | -0,51 | 2,2E-01 | 5,6E-05 | 1,7E-05 | 2,5E-07 | 1,7E-07 | 5,7E-03 | 0,0E+00 | 1,2E-06 | 2,7E-07 | 7,3E-09 | 9,7E-09 | 0,0E+00 | 1151068 | 1151340 | 273 | - |

| | | | | | | | | | | | | | | | | | | | | | | | | | |
|---|---|---|---|---|---|---|---|---|---|---|---|---|---|---|---|---|---|---|---|---|---|---|---|---|---|
| Y11_RS05220 | Y11_11281 | msrB | peptide-methionine (R)-S-oxide reductase MsrB | 0,08 | -1,61 | -1,62 | -1,68 | -1,7 | -0,01 | 2,7E-01 | 4,5E-08 | 4,7E-06 | 3,5E-09 | 2,0E-06 | 9,3E-01 | 0,0E+00 | 2,1E-09 | 8,5E-08 | 2,2E-10 | 7,6E-08 | 0,0E+00 | 1151414 | 1151827 | 414 | - |
| Y11_RS05225 | Y11_11301 | gapA | glyceraldehyde-3-phosphate dehydrogenase | -0,17 | 0,03 | -0,79 | 0,2 | -0,62 | -0,82 | 9,9E-02 | 7,7E-01 | 1,4E-05 | 7,6E-02 | 6,0E-05 | 2,1E-05 | 0,0E+00 | 0,0E+00 | 0,0E+00 | 0,0E+00 | 0,0E+00 | 0,0E+00 | 1152175 | 1153170 | 996 | + |
| Y11_RS05230 | Y11_11311 | | D-hexose-6-phosphate mutarotase | -0,76 | 0,43 | 0,46 | 1,19 | 1,22 | 0,03 | 6,2E-07 | 1,3E-05 | 3,7E-02 | 5,1E-09 | 1,7E-04 | 8,7E-01 | 0,0E+00 | 0,0E+00 | 0,0E+00 | 2,9E-10 | 4,3E-06 | 0,0E+00 | 1153421 | 1154299 | 879 | + |
| Y11_RS05235 | Y11_11321 | | MipA/OmpV family protein | 0,4 | 0,05 | 0,04 | -0,34 | -0,36 | -0,02 | 2,9E-03 | 5,0E-01 | 8,3E-01 | 1,9E-03 | 5,1E-02 | 9,0E-01 | 0,0E+00 | 0,0E+00 | 0,0E+00 | 0,0E+00 | 0,0E+00 | 0,0E+00 | 1154373 | 1155071 | 699 | - |
| Y11_RS05240 | Y11_11331 | yeaG | protein kinase YeaG | 0,02 | 1,36 | 1,9 | 1,35 | 1,89 | 0,54 | 8,1E-01 | 7,6E-09 | 4,3E-09 | 8,1E-10 | 1,4E-09 | 9,7E-06 | 0,0E+00 | 5,0E-10 | 2,8E-10 | 7,5E-11 | 2,1E-10 | 0,0E+00 | 1155565 | 1157499 | 1935 | + |
| Y11_RS05245 | Y11_11341 | | YeaH/YhbH family protein | 0,97 | 2,48 | 2,97 | 1,51 | 2,01 | 0,5 | 2,7E-05 | 5,5E-11 | 6,5E-09 | 8,6E-07 | 1,1E-06 | 2,7E-03 | 0,0E+00 | 1,5E-11 | 3,9E-10 | 2,2E-08 | 4,6E-08 | 0,0E+00 | 1157616 | 1158890 | 1275 | + |
| Y11_RS05265 | Y11_11381 | | pirin family protein | -0,14 | 0,81 | 0,67 | 0,96 | 0,82 | -0,14 | 3,1E-02 | 1,2E-06 | 1,2E-03 | 1,4E-08 | 2,2E-04 | 3,2E-01 | 0,0E+00 | 0,0E+00 | 0,0E+00 | 0,0E+00 | 0,0E+00 | 0,0E+00 | 1161757 | 1162629 | 873 | + |
| Y11_RS05280 | Y11_11411 | dadX | catabolic alanine racemase DadX | 0,21 | 0,2 | 1,86 | -0,01 | 1,65 | 1,66 | 2,6E-04 | 1,8E-04 | 6,1E-08 | 7,1E-01 | 1,8E-07 | 1,5E-07 | 0,0E+00 | 0,0E+00 | 2,2E-09 | 0,0E+00 | 1,0E-08 | 4,9E-08 | 1167132 | 1168223 | 1092 | - |
| Y11_RS05285 | Y11_11421 | | D-amino acid dehydrogenase | 0,69 | 0,46 | 1,71 | -0,23 | 1,02 | 1,25 | 1,3E-06 | 3,8E-05 | 3,9E-09 | 2,8E-03 | 2,1E-07 | 5,3E-08 | 0,0E+00 | 0,0E+00 | 2,6E-10 | 0,0E+00 | 1,1E-08 | 2,2E-08 | 1168407 | 1169711 | 1305 | - |
| Y11_RS05290 | Y11_11441 | | SpoVR family protein | 0,84 | 1,23 | 2,42 | 0,39 | 1,59 | 1,2 | 2,1E-05 | 1,5E-06 | 6,7E-09 | 4,9E-03 | 2,5E-07 | 2,8E-06 | 0,0E+00 | 4,3E-08 | 4,0E-10 | 0,0E+00 | 1,4E-08 | 4,0E-07 | 1170222 | 1171757 | 1536 | + |
| Y11_RS05295 | Y11_11451 | fadR | fatty acid metabolism transcriptional regulator FadR | -0,36 | -0,34 | -0,66 | 0,02 | -0,3 | -0,33 | 3,2E-04 | 3,9E-04 | 2,4E-03 | 5,6E-01 | 7,4E-02 | 5,6E-02 | 0,0E+00 | 0,0E+00 | 0,0E+00 | 0,0E+00 | 0,0E+00 | 0,0E+00 | 1171843 | 1172562 | 720 | - |
| Y11_RS05300 | Y11_11471 | nhaB | sodium/proton antiporter NhaB | -0,51 | -0,36 | 0,36 | 0,15 | 0,87 | 0,72 | 6,4E-02 | 2,7E-01 | 2,5E-01 | 5,3E-01 | 3,7E-03 | 3,5E-02 | 0,0E+00 | 0,0E+00 | 0,0E+00 | 0,0E+00 | 0,0E+00 | 0,0E+00 | 1172884 | 1174458 | 1575 | + |
| Y11_RS05305 | Y11_11491 | dsbB | disulfide bond formation protein DsbB | 0,2 | -0,93 | -0,56 | -1,13 | -0,76 | 0,37 | 2,6E-01 | 2,5E-05 | 2,5E-03 | 1,3E-04 | 2,9E-03 | 2,5E-02 | 0,0E+00 | 0,0E+00 | 0,0E+00 | 2,3E-06 | 0,0E+00 | 0,0E+00 | 1174688 | 1175218 | 531 | + |
| Y11_RS05325 | Y11_11531 | | hypothetical protein | 0,83 | -2,2 | -2,54 | -3,03 | -3,38 | -0,34 | 1,4E-05 | 3,8E-08 | 8,5E-09 | 3,0E-09 | 9,1E-10 | 2,2E-02 | 0,0E+00 | 1,8E-09 | 4,7E-10 | 2,0E-10 | 1,6E-10 | 0,0E+00 | 1177342 | 1177698 | 357 | - |
| Y11_RS05330 | Y11_11541 | | YcgN family cysteine cluster protein | 0 | -15 | 0 | 0,97 | 0 | -0,27 | 0,0E+00 | 0,0E+00 | 0,0E+00 | 4,7E-04 | 0,0E+00 | 3,5E-01 | 0,0E+00 | 0,0E+00 | 0,0E+00 | 0,0E+00 | 0,0E+00 | 0,0E+00 | 1177763 | 1178209 | 447 | - |
| Y11_RS05335 | Y11_11551 | | fumarylacetoacetate hydrolase family protein | -0,37 | 0,09 | 0,72 | 0,47 | 1,1 | 0,63 | 3,5E-03 | 3,6E-01 | 6,7E-05 | 2,0E-03 | 6,3E-06 | 3,7E-04 | 0,0E+00 | 0,0E+00 | 0,0E+00 | 0,0E+00 | 2,1E-07 | 0,0E+00 | 1178324 | 1178980 | 657 | - |
| Y11_RS05340 | Y11_11561 | | lytic murein transglycosylase | -0,13 | -0,64 | -0,45 | -0,52 | -0,33 | 0,19 | 2,5E-01 | 1,6E-04 | 5,8E-04 | 5,0E-03 | 3,1E-02 | 1,6E-01 | 0,0E+00 | 0,0E+00 | 0,0E+00 | 0,0E+00 | 0,0E+00 | 0,0E+00 | 1179061 | 1180131 | 1071 | - |
| Y11_RS05345 | Y11_11571 | | YcgL domain-containing protein | 0,15 | -0,95 | -1,31 | -1,1 | -1,46 | -0,36 | 3,8E-01 | 7,3E-04 | 5,0E-04 | 1,2E-04 | 1,5E-04 | 1,6E-01 | 0,0E+00 | 0,0E+00 | 6,2E-06 | 2,2E-06 | 3,9E-06 | 0,0E+00 | 1180170 | 1180442 | 273 | - |
| Y11_RS05350 | Y11_11581 | | hypothetical protein | 0 | -2,46 | -2,13 | -15 | -15 | 0,33 | 0,0E+00 | 1,8E-07 | 1,1E-05 | 0,0E+00 | 0,0E+00 | 2,0E-01 | 0,0E+00 | 6,7E-09 | 1,9E-07 | 0,0E+00 | 0,0E+00 | 0,0E+00 | 1180720 | 1181082 | 363 | - |
| Y11_RS05355 | Y11_11591 | minC | septum site-determining protein MinC | 0,6 | 0,44 | 0,65 | -0,17 | 0,05 | 0,21 | 5,1E-05 | 3,1E-04 | 7,8E-05 | 4,4E-02 | 6,1E-01 | 3,2E-02 | 0,0E+00 | 0,0E+00 | 0,0E+00 | 0,0E+00 | 0,0E+00 | 0,0E+00 | 1181633 | 1182319 | 687 | + |
| Y11_RS05360 | Y11_11601 | minD | septum site-determining protein MinD | 0,56 | 0,56 | 0,86 | 0 | 0,31 | 0,31 | 6,4E-05 | 1,2E-05 | 6,5E-07 | 9,9E-01 | 5,3E-04 | 1,2E-05 | 0,0E+00 | 0,0E+00 | 0,0E+00 | 0,0E+00 | 0,0E+00 | 0,0E+00 | 1182345 | 1183157 | 813 | + |
| Y11_RS05365 | Y11_11611 | minE | cell division topological specificity factor MinE | 0,51 | 1,11 | 1,6 | 0,61 | 1,09 | 0,48 | 7,2E-04 | 1,4E-05 | 4,8E-06 | 3,2E-04 | 3,9E-05 | 1,3E-02 | 0,0E+00 | 3,2E-07 | 8,8E-08 | 0,0E+00 | 1,1E-06 | 0,0E+00 | 1183161 | 1183430 | 270 | + |
| Y11_RS05370 | Y11_11621 | rnd | ribonuclease D | -0,43 | -0,88 | -1,52 | -0,44 | -1,09 | -0,64 | 5,0E-04 | 5,5E-07 | 8,3E-09 | 1,4E-04 | 1,9E-07 | 7,2E-07 | 0,0E+00 | 0,0E+00 | 4,7E-10 | 0,0E+00 | 1,1E-08 | 0,0E+00 | 1183560 | 1184681 | 1122 | - |
| Y11_RS05375 | Y11_11631 | fadD | long-chain-fatty-acid--CoA ligase FadD | -0,25 | -4,13 | -2,83 | -3,88 | -2,58 | 1,3 | 3,5E-02 | 2,6E-10 | 3,1E-09 | 8,3E-10 | 1,4E-08 | 5,3E-06 | 0,0E+00 | 4,1E-11 | 2,2E-10 | 7,6E-11 | 1,2E-09 | 6,7E-07 | 1184779 | 1186473 | 1695 | - |
| Y11_RS05380 | Y11_11641 | | Slp family lipoprotein | -3,32 | 1,47 | 2,01 | 4,78 | 5,32 | 0,54 | 8,9E-11 | 1,3E-07 | 4,2E-07 | 2,9E-11 | 2,9E-10 | 6,4E-03 | 3,0E-11 | 5,0E-09 | 1,1E-08 | 8,5E-12 | 8,5E-11 | 0,0E+00 | 1186814 | 1187365 | 552 | - |
| Y11_RS05385 | Y11_11651 | tsaB | tRNA (adenosine(37)-N6)-threonylcarbamoyltransferase complex dimerization subunit type 1 TsaB | -0,05 | -1,66 | -1,96 | -1,61 | -1,91 | -0,3 | 7,1E-01 | 4,2E-07 | 6,3E-09 | 2,3E-06 | 1,1E-07 | 1,6E-02 | 0,0E+00 | 1,4E-08 | 3,8E-10 | 5,2E-08 | 6,7E-09 | 0,0E+00 | 1187522 | 1188220 | 699 | - |
| Y11_RS05390 | Y11_11661 | | ATP-dependent DNA helicase | 0,1 | 0 | -0,01 | -0,1 | -0,11 | -0,01 | 5,3E-01 | 1,0E+00 | 9,0E-01 | 6,2E-01 | 4,7E-01 | 9,4E-01 | 0,0E+00 | 0,0E+00 | 0,0E+00 | 0,0E+00 | 0,0E+00 | 0,0E+00 | 1188295 | 1190199 | 1905 | - |
| Y11_RS05395 | Y11_11671 | | RidA family protein | -0,95 | 0,92 | 1,88 | 1,86 | 2,82 | 0,96 | 2,4E-02 | 6,9E-02 | 2,6E-02 | 3,1E-04 | 1,8E-03 | 1,9E-01 | 0,0E+00 | 0,0E+00 | 2,9E-04 | 5,5E-06 | 3,9E-05 | 0,0E+00 | 1190328 | 1190672 | 345 | + |
| Y11_RS05410 | Y11_11711 | | bifunctional 4-hydroxy-2-oxoglutarate aldolase/2-dehydro-3-deoxy-phosphogluconate aldolase | -0,47 | 0,19 | -0,14 | 0,66 | 0,33 | -0,33 | 2,5E-04 | 6,6E-02 | 7,3E-02 | 2,6E-05 | 1,7E-04 | 1,4E-03 | 0,0E+00 | 0,0E+00 | 0,0E+00 | 0,0E+00 | 0,0E+00 | 0,0E+00 | 1192212 | 1192853 | 642 | - |
| Y11_RS05415 | Y11_11721 | zwf | glucose-6-phosphate dehydrogenase | -0,36 | 0,74 | 0,46 | 1,1 | 0,82 | -0,28 | 4,7E-06 | 4,1E-09 | 5,0E-07 | 2,8E-10 | 7,5E-09 | 6,3E-06 | 0,0E+00 | 0,0E+00 | 0,0E+00 | 3,3E-11 | 0,0E+00 | 0,0E+00 | 1193159 | 1194643 | 1485 | - |
| Y11_RS05420 | Y11_11731 | | MurR/RpiR family transcriptional regulator | -0,19 | 0,07 | 0,14 | 0,27 | 0,34 | 0,07 | 1,2E-04 | 5,9E-03 | 1,1E-02 | 8,4E-06 | 9,0E-05 | 1,5E-01 | 0,0E+00 | 0,0E+00 | 0,0E+00 | 0,0E+00 | 0,0E+00 | 0,0E+00 | 1194996 | 1195865 | 870 | + |
| Y11_RS05425 | Y11_11741 | pyk | pyruvate kinase | 0,24 | -0,4 | -0,42 | -0,63 | -0,66 | -0,03 | 1,4E-03 | 2,1E-04 | 1,9E-04 | 4,9E-06 | 5,3E-06 | 7,1E-01 | 0,0E+00 | 0,0E+00 | 0,0E+00 | 0,0E+00 | 0,0E+00 | 0,0E+00 | 1196085 | 1197527 | 1443 | + |
| Y11_RS05435 | Y11_11761 | lpxM | lauroyl-Kdo(2)-lipid IV(A) myristoyltransferase | -0,3 | 0,5 | 0,49 | 0,8 | 0,8 | 0 | 2,4E-04 | 3,1E-08 | 1,4E-05 | 1,0E-07 | 1,9E-06 | 9,3E-01 | 0,0E+00 | 0,0E+00 | 0,0E+00 | 0,0E+00 | 0,0E+00 | 0,0E+00 | 1199356 | 1200321 | 966 | - |
| Y11_RS05445 | Y11_11781 | znuA | zinc ABC transporter substrate-binding protein ZnuA | -0,14 | 1,06 | 0,61 | 1,2 | 0,76 | -0,44 | 1,6E-01 | 2,0E-06 | 1,6E-02 | 1,5E-08 | 4,1E-03 | 4,6E-02 | 0,0E+00 | 5,6E-08 | 0,0E+00 | 6,9E-10 | 0,0E+00 | 0,0E+00 | 1201736 | 1202698 | 963 | - |
| Y11_RS05450 | Y11_11791 | znuC | zinc ABC transporter ATP-binding protein ZnuC | -0,01 | -0,56 | -1,05 | -0,56 | -1,05 | -0,49 | 9,6E-01 | 6,5E-04 | 4,1E-08 | 3,1E-03 | 5,2E-06 | 1,1E-03 | 0,0E+00 | 0,0E+00 | 1,6E-09 | 0,0E+00 | 1,8E-07 | 0,0E+00 | 1202774 | 1203532 | 759 | + |
| Y11_RS05465 | Y11_11821 | ruvB | Holliday junction branch migration DNA helicase RuvB | -0,56 | -0,59 | -0,96 | -0,03 | -0,4 | -0,36 | 3,8E-06 | 1,2E-06 | 1,6E-05 | 4,1E-01 | 4,3E-03 | 6,0E-03 | 0,0E+00 | 0,0E+00 | 0,0E+00 | 0,0E+00 | 0,0E+00 | 0,0E+00 | 1206113 | 1207117 | 1005 | - |
| Y11_RS05470 | Y11_11831 | ruvA | Holliday junction branch migration protein RuvA | -0,3 | -1,16 | -1,31 | -0,86 | -1,01 | -0,15 | 1,6E-02 | 7,4E-07 | 4,0E-04 | 1,9E-05 | 2,3E-03 | 5,1E-01 | 0,0E+00 | 2,2E-08 | 5,0E-06 | 0,0E+00 | 5,1E-05 | 0,0E+00 | 1207153 | 1207770 | 618 | - |
| Y11_RS05475 | Y11_11841 | ruvC | crossover junction endodeoxyribonuclease RuvC | -0,34 | 0,18 | -0,76 | 0,51 | -0,43 | -0,94 | 1,4E-01 | 4,5E-01 | 1,3E-02 | 1,6E-02 | 5,5E-02 | 2,1E-03 | 0,0E+00 | 0,0E+00 | 0,0E+00 | 0,0E+00 | 0,0E+00 | 0,0E+00 | 1207893 | 1208414 | 522 | - |
| Y11_RS05480 | Y11_11851 | | YebC/PmpR family DNA-binding transcriptional regulator | -0,62 | -1,02 | -2,11 | -0,41 | -1,5 | -1,09 | 2,1E-04 | 2,8E-06 | 2,9E-08 | 3,3E-03 | 8,4E-07 | 5,9E-06 | 0,0E+00 | 7,6E-08 | 1,2E-09 | 0,0E+00 | 3,6E-08 | 7,4E-07 | 1208777 | 1209520 | 744 | - |
| Y11_RS05490 | Y11_11871 | aspS | aspartate--tRNA ligase | -0,59 | -0,72 | -0,85 | -0,12 | -0,25 | -0,13 | 4,8E-07 | 2,3E-07 | 6,3E-08 | 4,8E-02 | 1,3E-03 | 4,5E-02 | 0,0E+00 | 0,0E+00 | 0,0E+00 | 0,0E+00 | 0,0E+00 | 0,0E+00 | 1210117 | 1211913 | 1797 | - |
| Y11_RS05500 | Y11_11901 | | hydrolase | -3,1 | 0 | -0,21 | 3,1 | 2,89 | -0,21 | 1,5E-11 | 9,8E-01 | 7,2E-02 | 1,1E-09 | 2,8E-09 | 1,4E-01 | 9,7E-12 | 0,0E+00 | 0,0E+00 | 9,1E-11 | 3,6E-10 | 0,0E+00 | 1212466 | 1213041 | 576 | + |
| Y11_RS05510 | Y11_11921 | | MAPEG family protein | 1,06 | -0,97 | 0,47 | -2,03 | -0,59 | 1,44 | 3,3E-04 | 1,3E-03 | 1,9E-01 | 5,7E-06 | 1,0E-01 | 2,6E-03 | 1,3E-05 | 0,0E+00 | 0,0E+00 | 1,2E-07 | 0,0E+00 | 1,6E-04 | 1214219 | 1214614 | 396 | + |
| Y11_RS05515 | Y11_11931 | cmoA | carboxy-S-adenosyl-L-methionine synthase CmoA | -0,66 | -1,29 | -1,7 | -0,62 | -1,04 | -0,41 | 9,5E-06 | 5,3E-07 | 4,7E-08 | 2,0E-05 | 2,7E-07 | 1,6E-03 | 0,0E+00 | 1,7E-08 | 1,8E-09 | 0,0E+00 | 1,4E-08 | 0,0E+00 | 1214804 | 1215604 | 801 | + |
| Y11_RS05520 | Y11_11941 | cmoB | tRNA 5-methoxyuridine(34)/uridine 5-oxyacetic acid(34) synthase CmoB | -0,83 | -1,16 | -1,54 | -0,33 | -0,71 | -0,38 | 8,5E-07 | 1,5E-06 | 1,9E-08 | 9,4E-03 | 1,5E-05 | 6,5E-03 | 0,0E+00 | 4,3E-08 | 8,6E-10 | 0,0E+00 | 0,0E+00 | 0,0E+00 | 1215601 | 1216572 | 972 | + |
| Y11_RS05530 | Y11_11951 | | hypothetical protein | 0,51 | -0,82 | -0,46 | -1,34 | -0,98 | 0,35 | 6,9E-03 | 1,1E-04 | 2,4E-01 | 1,9E-05 | 3,3E-02 | 3,7E-01 | 0,0E+00 | 0,0E+00 | 0,0E+00 | 3,7E-07 | 0,0E+00 | 0,0E+00 | 1217013 | 1217657 | 645 | - |
| Y11_RS05535 | Y11_11961 | | DeoR family transcriptional regulator | 0,87 | 0,36 | -0,07 | -0,51 | -0,94 | -0,43 | 1,7E-04 | 1,6E-02 | 6,8E-01 | 4,0E-03 | 5,7E-04 | 2,7E-02 | 0,0E+00 | 0,0E+00 | 0,0E+00 | 0,0E+00 | 0,0E+00 | 0,0E+00 | 1217739 | 1218548 | 810 | - |
| Y11_RS05565 | Y11_12021 | | SIS domain-containing protein | -0,65 | -0,63 | 0,14 | 0,02 | 0,79 | 0,77 | 5,9E-04 | 5,4E-03 | 3,1E-01 | 9,2E-01 | 1,6E-04 | 1,7E-03 | 0,0E+00 | 0,0E+00 | 0,0E+00 | 0,0E+00 | 0,0E+00 | 0,0E+00 | 1223076 | 1224230 | 1155 | + |
| Y11_RS05570 | Y11_12031 | kbaY | tagatose-bisphosphate aldolase subunit KbaY | -0,43 | 15 | 15 | 15 | 15 | 0 | 3,5E-02 | 0,0E+00 | 0,0E+00 | 0,0E+00 | 0,0E+00 | 0,0E+00 | 0,0E+00 | 0,0E+00 | 0,0E+00 | 0,0E+00 | 0,0E+00 | 0,0E+00 | 1224243 | 1225121 | 879 | + |
| Y11_RS05575 | Y11_12051 | | peptidoglycan DD-metalloendopeptidase family protein | 2,9 | -0,07 | 0,16 | -2,97 | -2,75 | 0,23 | 8,2E-09 | 5,0E-01 | 5,1E-01 | 3,3E-09 | 3,0E-06 | 3,4E-01 | 1,2E-09 | 0,0E+00 | 0,0E+00 | 2,1E-10 | 1,1E-07 | 0,0E+00 | 1225309 | 1226037 | 729 | + |
| Y11_RS05585 | Y11_12091 | | RpiB/LacA/LacB family sugar-phosphate isomerase | -0,35 | -1,31 | -0,88 | -0,96 | -0,53 | 0,43 | 9,2E-02 | 6,6E-07 | 6,3E-06 | 8,4E-05 | 2,8E-03 | 7,1E-03 | 0,0E+00 | 2,0E-08 | 0,0E+00 | 0,0E+00 | 0,0E+00 | 0,0E+00 | 1227935 | 1228573 | 639 | - |
| Y11_RS05600 | Y11_12121 | zinT | metal-binding protein ZinT | -0,73 | 1,01 | 1,57 | 1,75 | 2,44 | 0 | 2,1E-02 | 4,9E-03 | 2,6E-04 | 4,2E-05 | 3,4E-06 | 0,0E+00 | 0,0E+00 | 9,5E-05 | 3,4E-06 | 8,1E-07 | 1,3E-07 | 0,0E+00 | 1229936 | 1230595 | 660 | - |
| Y11_RS05605 | Y11_12131 | | HutD family protein | -0,17 | -1,25 | 0,94 | -1,09 | 1,1 | 2,19 | 1,5E-02 | 5,6E-08 | 5,5E-04 | 5,9E-08 | 1,6E-04 | 1,3E-06 | 0,0E+00 | 2,5E-09 | 0,0E+00 | 2,2E-09 | 4,1E-06 | 2,1E-07 | 1230782 | 1231354 | 573 | - |
| Y11_RS05610 | Y11_12141 | | formimidoylglutamate deiminase | 0,03 | -0,97 | 0,33 | -0,99 | 0,3 | 1,29 | 7,9E-01 | 5,3E-06 | 1,4E-02 | 1,7E-06 | 1,4E-02 | 4,0E-07 | 0,0E+00 | 0,0E+00 | 0,0E+00 | 0,0E+00 | 0,0E+00 | 8,7E-08 | 1231354 | 1232724 | 1371 | - |
| Y11_RS05615 | Y11_12151 | hutC | histidine utilization repressor | 0,36 | 0,46 | 0,61 | 0,1 | 0,25 | 0,16 | 5,8E-02 | 9,0E-03 | 5,7E-02 | 5,9E-01 | 4,2E-01 | 6,0E-01 | 0,0E+00 | 0,0E+00 | 0,0E+00 | 0,0E+00 | 0,0E+00 | 0,0E+00 | 1232873 | 1233640 | 768 | + |
| Y11_RS05620 | Y11_12161 | hutI | imidazolonepropionase | -0,61 | -0,42 | 1,68 | 0,2 | 2,29 | 2,09 | 6,4E-05 | 2,2E-03 | 1,1E-07 | 4,8E-02 | 4,2E-09 | 2,5E-08 | 0,0E+00 | 0,0E+00 | 3,5E-09 | 0,0E+00 | 4,9E-10 | 1,4E-08 | 1233776 | 1234996 | 1221 | + |
| Y11_RS05625 | Y11_12171 | hutG | N-formylglutamate deformylase | -0,6 | 0,76 | 2,93 | 1,36 | 3,52 | 2,17 | 3,2E-05 | 2,5E-04 | 7,6E-10 | 3,7E-06 | 1,7E-10 | 1,9E-07 | 0,0E+00 | 0,0E+00 | 8,3E-11 | 8,2E-08 | 6,2E-11 | 5,5E-08 | 1234993 | 1235802 | 810 | + |
| Y11_RS05640 | Y11_12201 | | porin | 1,22 | -2,84 | -2,99 | -4,07 | -4,19 | -0,15 | 4,5E-06 | 9,7E-11 | 1,4E-08 | 2,1E-10 | 7,2E-09 | 3,0E-01 | 2,5E-07 | 2,2E-11 | 6,8E-10 | 2,7E-11 | 7,5E-10 | 0,0E+00 | 1237231 | 1238310 | 1080 | - |
| Y11_RS05650 | Y11_12221 | astE | succinylglutamate desuccinylase | 0,21 | 0,21 | 0,52 | 0 | 0,32 | 0,32 | 5,0E-02 | 4,8E-02 | 1,8E-03 | 9,7E-01 | 2,3E-02 | 2,4E-02 | 0,0E+00 | 0,0E+00 | 0,0E+00 | 0,0E+00 | 0,0E+00 | 0,0E+00 | 1240869 | 1241861 | 993 | - |
| Y11_RS05660 | Y11_12251 | astD | succinylglutamate-semialdehyde dehydrogenase | 0,13 | 2,18 | 2,85 | 2,05 | 2,72 | 0,68 | 6,2E-02 | 6,4E-08 | 1,2E-10 | 5,8E-08 | 2,4E-11 | 3,3E-04 | 0,0E+00 | 2,8E-09 | 2,5E-11 | 2,2E-09 | 1,9E-11 | 0,0E+00 | 1242923 | 1244404 | 1482 | - |
| Y11_RS05665 | Y11_12261 | astA | arginine N-succinyltransferase | 0,33 | 2,57 | 3,46 | 2,24 | 3,13 | 0,89 | 4,8E-04 | 2,8E-09 | 5,2E-11 | 1,9E-09 | 9,1E-12 | 8,8E-06 | 0,0E+00 | 2,3E-10 | 1,6E-11 | 1,4E-10 | 1,0E-11 | 0,0E+00 | 1244401 | 1245438 | 1038 | - |
| Y11_RS05670 | Y11_12271 | | aspartate aminotransferase family protein | 0,32 | 2,19 | 3,07 | 1,87 | 2,75 | 0,88 | 8,4E-05 | 5,1E-09 | 1,5E-10 | 1,3E-08 | 2,6E-10 | 2,5E-05 | 0,0E+00 | 3,7E-10 | 2,8E-11 | 6,5E-10 | 7,8E-11 | 0,0E+00 | 1245481 | 1246725 | 1245 | - |
| Y11_RS05675 | Y11_12281 | | PLP-dependent aminotransferase family protein | 0,05 | 0,34 | 0,47 | 0,28 | 0,41 | 0,13 | 5,8E-01 | 2,9E-02 | 1,2E-02 | 6,7E-02 | 2,5E-02 | 4,8E-01 | 0,0E+00 | 0,0E+00 | 0,0E+00 | 0,0E+00 | 0,0E+00 | 0,0E+00 | 1247302 | 1248735 | 1434 | - |
| Y11_RS05680 | Y11_12291 | | cytochrome ubiquinol oxidase subunit I | -0,37 | 1,08 | 15 | 1,45 | 15 | 15 | 8,6E-02 | 1,6E-02 | 0,0E+00 | 2,2E-03 | 0,0E+00 | 0,0E+00 | 0,0E+00 | 3,1E-04 | 0,0E+00 | 3,6E-05 | 0,0E+00 | 0,0E+00 | 1248920 | 1250368 | 1449 | + |
| Y11_RS05690 | Y11_12321 | narX | nitrate/nitrite two-component system sensor histidine kinase NarX | 0,15 | 0,22 | 0,03 | 0,07 | -0,11 | -0,19 | 4,6E-01 | 3,2E-01 | 8,9E-01 | 6,9E-01 | 5,4E-01 | 3,3E-01 | 0,0E+00 | 0,0E+00 | 0,0E+00 | 0,0E+00 | 0,0E+00 | 0,0E+00 | 1251605 | 1253386 | 1782 | + |
| Y11_RS05710 | Y11_12361 | | siderophore-interacting protein | 0,76 | -0,55 | -1,22 | -1,31 | -1,98 | -0,67 | 7,4E-05 | 1,6E-03 | 2,7E-03 | 1,4E-05 | 1,5E-04 | 5,7E-02 | 0,0E+00 | 0,0E+00 | 3,2E-05 | 2,8E-07 | 3,9E-06 | 0,0E+00 | 1257662 | 1258498 | 837 | + |
| Y11_RS05715 | Y11_12371 | chaA | sodium-potassium/proton antiporter ChaA | -0,23 | -1,13 | -0,44 | -0,9 | -0,21 | 0,7 | 2,8E-01 | 5,9E-02 | 2,7E-02 | 1,2E-01 | 2,6E-01 | 2,1E-01 | 0,0E+00 | 1,1E-03 | 0,0E+00 | 0,0E+00 | 0,0E+00 | 0,0E+00 | 1258719 | 1259828 | 1110 | + |
| Y11_RS05720 | Y11_12381 | murJ | murein biosynthesis integral membrane protein MurJ | 0 | 0 | -15 | 0 | -0,33 | -0,43 | 0,0E+00 | 0,0E+00 | 0,0E+00 | 0,0E+00 | 1,8E-03 | 6,7E-03 | 0,0E+00 | 0,0E+00 | 0,0E+00 | 0,0E+00 | 0,0E+00 | 0,0E+00 | 1260148 | 1261683 | 1536 | - |

| | | | | | | | | | | | | | | | | | | | | | | | | | |
|---|---|---|---|---|---|---|---|---|---|---|---|---|---|---|---|---|---|---|---|---|---|---|---|---|---|
| Y11_RS05725 | Y11_12391 | | Gfo/Idh/MocA family oxidoreductase | 0,16 | -0,07 | -0,38 | -0,23 | -0,53 | -0,3 | 2,4E-01 | 4,0E-01 | 2,4E-02 | 8,2E-02 | 1,0E-02 | 4,6E-02 | 0,0E+00 | 0,0E+00 | 0,0E+00 | 0,0E+00 | 0,0E+00 | 0,0E+00 | 1261822 | 1262781 | 960 | - |
| Y11_RS05730 | Y11_12401 | | YceH family protein | 0 | 0 | 0,82 | 0 | -15 | 0,6 | 0,0E+00 | 0,0E+00 | 1,1E-02 | 0,0E+00 | 0,0E+00 | 7,1E-02 | 0,0E+00 | 0,0E+00 | 0,0E+00 | 0,0E+00 | 0,0E+00 | 0,0E+00 | 1262774 | 1263415 | 642 | - |
| Y11_RS05735 | Y11_12411 | rimJ | ribosomal protein S5-alanine N-acetyltransferase | -0,33 | 0,3 | 0,96 | 0,63 | 1,29 | 0,66 | 6,5E-01 | 7,4E-01 | 2,1E-01 | 2,8E-01 | 7,0E-05 | 2,7E-01 | 0,0E+00 | 0,0E+00 | 0,0E+00 | 0,0E+00 | 1,9E-06 | 0,0E+00 | 1263423 | 1264007 | 585 | - |
| Y11_RS05755 | Y11_12441 | | lipoprotein | 0,3 | -0,02 | -0,62 | -0,32 | -0,91 | -0,6 | 6,2E-03 | 8,1E-01 | 2,7E-03 | 4,4E-03 | 1,9E-04 | 3,4E-03 | 0,0E+00 | 0,0E+00 | 0,0E+00 | 0,0E+00 | 0,0E+00 | 0,0E+00 | 1267843 | 1268406 | 564 | + |
| Y11_RS05760 | Y11_12451 | | molecular chaperone | -0,2 | 0,33 | -0,04 | 0,53 | 0,16 | -0,37 | 3,2E-02 | 6,7E-03 | 7,8E-01 | 4,6E-04 | 2,8E-01 | 3,7E-02 | 0,0E+00 | 0,0E+00 | 0,0E+00 | 0,0E+00 | 0,0E+00 | 0,0E+00 | 1268444 | 1269004 | 561 | - |
| Y11_RS05765 | Y11_12461 | | phosphatase | -0,34 | 0,16 | -0,23 | 0,5 | 0,11 | -0,39 | 1,0E-03 | 4,7E-02 | 2,1E-02 | 2,1E-05 | 1,4E-01 | 5,2E-04 | 0,0E+00 | 0,0E+00 | 0,0E+00 | 0,0E+00 | 0,0E+00 | 0,0E+00 | 1269068 | 1269805 | 738 | - |
| Y11_RS05800 | Y11_12511 | kdsA | 3-deoxy-8-phosphooctulonate synthase | 0,26 | -1,34 | -1,29 | -1,6 | -1,55 | 0,05 | 3,0E-03 | 1,8E-09 | 2,5E-06 | 1,4E-08 | 1,4E-06 | 6,6E-01 | 0,0E+00 | 1,7E-10 | 5,0E-08 | 6,7E-10 | 5,5E-08 | 0,0E+00 | 1276241 | 1277095 | 855 | - |
| Y11_RS05805 | Y11_12521 | sirB1 | invasion regulator SirB1 | 0 | 0 | 0,32 | 0 | -0,49 | -0,16 | 0,0E+00 | 0,0E+00 | 4,7E-01 | 0,0E+00 | 2,5E-01 | 7,0E-01 | 0,0E+00 | 0,0E+00 | 0,0E+00 | 0,0E+00 | 0,0E+00 | 0,0E+00 | 1277240 | 1278049 | 810 | - |
| Y11_RS05815 | Y11_12541 | prmC | peptide chain release factor N(5)-glutamine methyltransferase | 0 | -15 | -15 | -15 | -15 | -0,49 | 0,0E+00 | 0,0E+00 | 0,0E+00 | 0,0E+00 | 0,0E+00 | 3,5E-01 | 0,0E+00 | 0,0E+00 | 0,0E+00 | 0,0E+00 | 0,0E+00 | 0,0E+00 | 1278465 | 1279319 | 855 | - |
| Y11_RS05820 | Y11_12551 | prfA | peptide chain release factor 1 | 0,24 | -0,78 | -1,14 | -1,02 | -1,38 | -0,36 | 3,8E-03 | 8,1E-06 | 4,6E-08 | 1,6E-06 | 2,0E-08 | 2,0E-03 | 0,0E+00 | 0,0E+00 | 1,8E-09 | 3,8E-08 | 1,7E-09 | 0,0E+00 | 1279319 | 1280401 | 1083 | - |
| Y11_RS05825 | Y11_12561 | hemA | glutamyl-tRNA reductase | 0,11 | -0,65 | -1,21 | -0,76 | -1,33 | -0,56 | 5,0E-01 | 1,8E-03 | 9,0E-04 | 1,0E-05 | 2,0E-04 | 2,0E-02 | 0,0E+00 | 0,0E+00 | 1,1E-05 | 0,0E+00 | 5,0E-06 | 0,0E+00 | 1280433 | 1281695 | 1263 | - |
| Y11_RS05830 | Y11_12571 | lolB | lipoprotein insertase outer membrane protein LolB | 0,24 | 0,17 | -0,36 | -0,07 | -0,6 | -0,53 | 3,4E-02 | 8,9E-02 | 1,9E-02 | 2,7E-01 | 3,4E-04 | 5,5E-04 | 0,0E+00 | 0,0E+00 | 0,0E+00 | 0,0E+00 | 0,0E+00 | 0,0E+00 | 1282001 | 1282624 | 624 | + |
| Y11_RS05835 | Y11_12581 | ispE | 4-(cytidine 5'-diphospho)-2-C-methyl-D-erythritol kinase | 0,06 | -0,31 | -0,76 | -0,37 | -0,82 | -0,45 | 5,7E-01 | 9,8E-03 | 9,7E-05 | 1,6E-02 | 2,6E-04 | 5,9E-03 | 0,0E+00 | 0,0E+00 | 0,0E+00 | 0,0E+00 | 0,0E+00 | 0,0E+00 | 1282626 | 1283507 | 882 | + |
| Y11_RS05840 | Y11_12591 | prs | ribose-phosphate diphosphokinase | -0,21 | -0,96 | -0,83 | -0,76 | -0,63 | 0,13 | 3,5E-02 | 5,8E-07 | 4,4E-05 | 3,5E-05 | 8,5E-04 | 2,8E-01 | 0,0E+00 | 0,0E+00 | 0,0E+00 | 0,0E+00 | 0,0E+00 | 0,0E+00 | 1283705 | 1284652 | 948 | + |
| Y11_RS05855 | Y11_12621 | pth | aminoacyl-tRNA hydrolase | -0,04 | -0,74 | -1,27 | -0,69 | -1,23 | -0,53 | 5,0E-01 | 7,6E-05 | 2,7E-05 | 2,6E-04 | 5,3E-05 | 1,6E-02 | 0,0E+00 | 0,0E+00 | 4,2E-07 | 0,0E+00 | 1,5E-06 | 0,0E+00 | 1285743 | 1286336 | 594 | + |
| Y11_RS05865 | Y11_12641 | ychF | redox-regulated ATPase YchF | -0,28 | -0,56 | -0,75 | -0,28 | -0,48 | -0,19 | 2,6E-04 | 3,5E-07 | 9,3E-08 | 7,5E-04 | 3,3E-05 | 5,4E-03 | 0,0E+00 | 0,0E+00 | 0,0E+00 | 0,0E+00 | 0,0E+00 | 0,0E+00 | 1287117 | 1288208 | 1092 | + |
| Y11_RS05870 | Y11_12651 | | class C beta-lactamase | 1,03 | -0,03 | -0,97 | -1,05 | -1,99 | -0,94 | 2,9E-03 | 8,1E-01 | 7,0E-04 | 2,4E-03 | 1,1E-04 | 8,0E-04 | 1,0E-04 | 0,0E+00 | 0,0E+00 | 4,0E-05 | 2,9E-06 | 0,0E+00 | 1288355 | 1289521 | 1167 | - |
| Y11_RS05875 | Y11_12661 | ampR | LysR family transcriptional regulator AmpR | 0,46 | -2,15 | -0,65 | -2,6 | -1,11 | 1,49 | 9,1E-02 | 1,4E-02 | 4,2E-01 | 4,1E-03 | 1,7E-01 | 1,7E-01 | 0,0E+00 | 2,6E-04 | 0,0E+00 | 6,8E-05 | 3,5E-03 | 8,5E-03 | 1289656 | 1290540 | 885 | + |
| Y11_RS05880 | Y11_12671 | | DUF533 domain-containing protein | -0,27 | -1,57 | -1,58 | -1,29 | -1,3 | -0,01 | 8,0E-03 | 2,3E-05 | 2,9E-07 | 6,8E-05 | 3,9E-07 | 9,7E-01 | 0,0E+00 | 5,4E-07 | 7,9E-09 | 1,3E-06 | 1,9E-08 | 0,0E+00 | 1290647 | 1291303 | 657 | - |
| Y11_RS05885 | Y11_12691 | | triacylglycerol lipase | 0,56 | 0,02 | 1 | -0,52 | 0,44 | 0,97 | 3,1E-03 | 9,7E-01 | 1,9E-03 | 3,7E-01 | 5,3E-02 | 1,3E-01 | 0,0E+00 | 0,0E+00 | 0,0E+00 | 0,0E+00 | 0,0E+00 | 0,0E+00 | 1291605 | 1292489 | 885 | - |
| Y11_RS05920 | Y11_12731 | | CatB-related O-acetyltransferase | 15 | 2,61 | 15 | 0 | 0 | 0 | 0,0E+00 | 2,2E-04 | 0,0E+00 | 0,0E+00 | 0,0E+00 | 0,0E+00 | 0,0E+00 | 4,7E-06 | 0,0E+00 | 0,0E+00 | 0,0E+00 | 0,0E+00 | 1295268 | 1295912 | 645 | - |
| Y11_RS05925 | Y11_12741 | | helix-turn-helix domain-containing protein | -0,13 | 1,07 | 0,88 | 1,2 | 1,01 | -0,19 | 3,3E-01 | 1,2E-04 | 1,8E-04 | 2,0E-04 | 3,6E-04 | 3,5E-01 | 0,0E+00 | 2,5E-06 | 0,0E+00 | 3,6E-06 | 8,7E-06 | 0,0E+00 | 1296313 | 1296732 | 420 | + |
| Y11_RS05930 | Y11_12751 | | LysR family transcriptional regulator | 0,66 | 0,56 | 0,21 | 0 | 0 | 0 | 4,3E-02 | 1,5E-02 | 2,4E-01 | 0,0E+00 | 0,0E+00 | 0,0E+00 | 0,0E+00 | 0,0E+00 | 0,0E+00 | 0,0E+00 | 0,0E+00 | 0,0E+00 | 1297338 | 1298255 | 918 | - |
| Y11_RS05975 | Y11_12851 | | toll/interleukin-1 receptor domain-containing protein | 0,55 | 0,15 | 0,87 | -0,4 | 0,32 | 0,72 | 3,9E-04 | 2,6E-01 | 7,7E-05 | 2,4E-03 | 4,6E-03 | 2,5E-04 | 0,0E+00 | 0,0E+00 | 0,0E+00 | 0,0E+00 | 0,0E+00 | 0,0E+00 | 1302830 | 1303891 | 1062 | - |
| Y11_RS05980 | Y11_12861 | | BrnA antitoxin family protein | 0,02 | 0,22 | 0,91 | 0,2 | 0,89 | 0,69 | 7,9E-01 | 4,1E-03 | 8,4E-05 | 4,1E-02 | 2,0E-04 | 5,9E-04 | 0,0E+00 | 0,0E+00 | 0,0E+00 | 0,0E+00 | 0,0E+00 | 0,0E+00 | 1303957 | 1304247 | 291 | - |
| Y11_RS05985 | Y11_12871 | | BrnT family toxin | -0,45 | -0,31 | 0,16 | 0,15 | 0,61 | 0,46 | 6,7E-03 | 6,6E-02 | 3,1E-01 | 4,2E-01 | 8,2E-03 | 4,0E-02 | 0,0E+00 | 0,0E+00 | 0,0E+00 | 0,0E+00 | 0,0E+00 | 0,0E+00 | 1304228 | 1304500 | 273 | - |
| Y11_RS06035 | Y11_12961 | | type II toxin-antitoxin system RelB/DinJ family antitoxin | -0,13 | 2,77 | 3,02 | 2,9 | 3,15 | 0,25 | 1,8E-01 | 1,5E-11 | 2,4E-09 | 5,4E-10 | 5,6E-09 | 4,1E-02 | 0,0E+00 | 1,2E-11 | 1,9E-10 | 5,5E-11 | 6,0E-10 | 0,0E+00 | 1309561 | 1309821 | 261 | - |
| Y11_RS06075 | Y11_13051 | | IS256 family transposase | 1,72 | -0,51 | -1,01 | -2,23 | -2,73 | -0,49 | 8,0E-04 | 2,9E-02 | 2,3E-04 | 9,3E-05 | 1,4E-05 | 2,5E-03 | 3,0E-05 | 0,0E+00 | 3,0E-06 | 1,7E-06 | 4,3E-07 | 0,0E+00 | 1314606 | 1315808 | 1203 | + |
| Y11_RS06100 | Y11_13111 | | BRCT domain-containing protein | 0,29 | -0,75 | -0,46 | -1,03 | -0,75 | 0,29 | 5,6E-02 | 1,1E-03 | 9,4E-03 | 2,0E-05 | 5,6E-05 | 4,6E-02 | 0,0E+00 | 0,0E+00 | 0,0E+00 | 4,0E-07 | 0,0E+00 | 0,0E+00 | 1320226 | 1321110 | 885 | - |
| Y11_RS06105 | Y11_13121 | | phage repressor protein CI | -0,43 | 0,2 | 0,28 | 0,63 | 0,71 | 0,08 | 6,4E-04 | 3,9E-02 | 2,6E-02 | 1,1E-04 | 1,8E-04 | 4,9E-01 | 0,0E+00 | 0,0E+00 | 0,0E+00 | 0,0E+00 | 0,0E+00 | 0,0E+00 | 1321151 | 1321726 | 576 | - |
| Y11_RS06315 | Y11_13531 | pgsA | CDP-diacylglycerol--glycerol-3-phosphate 3-phosphatidyltransferase | 0,23 | 0,31 | 15 | 0 | 0 | 0 | 1,4E-01 | 1,8E-01 | 0,0E+00 | 0,0E+00 | 0,0E+00 | 0,0E+00 | 0,0E+00 | 0,0E+00 | 0,0E+00 | 0,0E+00 | 0,0E+00 | 0,0E+00 | 1350055 | 1350603 | 549 | - |
| Y11_RS06320 | Y11_13541 | uvrC | excinuclease ABC subunit UvrC | 0,14 | 0,65 | 0,48 | 0,51 | 0,35 | -0,17 | 2,8E-01 | 8,1E-04 | 2,5E-02 | 6,2E-04 | 5,7E-02 | 3,3E-01 | 0,0E+00 | 0,0E+00 | 0,0E+00 | 0,0E+00 | 0,0E+00 | 0,0E+00 | 1350662 | 1352494 | 1833 | - |
| Y11_RS06325 | Y11_13551 | uvrY | UvrY/SirA/GacA family response regulator transcription factor | -0,92 | 0,81 | 1,15 | 1,73 | 2,06 | 0,34 | 5,5E-07 | 2,5E-06 | 2,3E-07 | 1,4E-09 | 5,5E-10 | 8,1E-04 | 0,0E+00 | 0,0E+00 | 6,5E-09 | 1,1E-10 | 1,2E-10 | 0,0E+00 | 1352487 | 1353143 | 657 | - |
| Y11_RS06330 | Y11_13561 | | DUF2594 family protein | -1,39 | 0,36 | 0,61 | 1,76 | 2 | 0,24 | 9,1E-08 | 1,0E-03 | 2,4E-03 | 2,9E-11 | 2,2E-07 | 8,1E-02 | 8,6E-09 | 0,0E+00 | 0,0E+00 | 8,5E-12 | 1,2E-08 | 0,0E+00 | 1353837 | 1354061 | 225 | + |
| Y11_RS06340 | Y11_13581 | | hypothetical protein | -1,31 | -0,33 | 0,86 | 0,99 | 2,17 | 1,19 | 7,2E-06 | 2,2E-02 | 3,2E-02 | 1,6E-04 | 2,3E-04 | 8,3E-03 | 3,7E-07 | 0,0E+00 | 0,0E+00 | 0,0E+00 | 5,8E-06 | 4,8E-04 | 1354541 | 1354795 | 255 | - |
| Y11_RS06345 | Y11_13591 | amyA | alpha-amylase | -3,32 | 0,23 | 0,13 | 3,55 | 3,45 | -0,1 | 5,2E-11 | 2,6E-01 | 2,5E-01 | 4,0E-08 | 1,6E-10 | 6,2E-01 | 2,2E-11 | 0,0E+00 | 0,0E+00 | 1,6E-09 | 6,2E-11 | 0,0E+00 | 1355047 | 1356546 | 1500 | - |
| Y11_RS06355 | Y11_13611 | | hypothetical protein | -0,98 | 15 | 5,04 | 15 | 5,98 | 0 | 2,4E-04 | 0,0E+00 | 5,2E-10 | 0,0E+00 | 9,6E-11 | 0,0E+00 | 0,0E+00 | 0,0E+00 | 6,6E-11 | 0,0E+00 | 4,5E-11 | 0,0E+00 | 1357102 | 1357599 | 498 | - |
| Y11_RS06360 | Y11_13621 | | class I SAM-dependent methyltransferase | -0,66 | 0,74 | 1,52 | 1,4 | 2,17 | 0,77 | 5,0E-04 | 1,2E-05 | 2,1E-07 | 6,5E-07 | 5,4E-08 | 4,8E-06 | 0,0E+00 | 0,0E+00 | 6,1E-09 | 1,7E-08 | 3,8E-09 | 0,0E+00 | 1358003 | 1358737 | 735 | + |
| Y11_RS06385 | Y11_13651 | cycA | D-serine/D-alanine/glycine transporter | 0,97 | 0,55 | 1,88 | -0,42 | 0,91 | 1,33 | 1,3E-02 | 2,0E-02 | 3,4E-06 | 2,3E-01 | 2,1E-02 | 1,8E-04 | 0,0E+00 | 0,0E+00 | 6,5E-08 | 0,0E+00 | 0,0E+00 | 1,4E-05 | 1360831 | 1362252 | 1422 | + |
| Y11_RS06435 | Y11_13781 | | N-acetylneuraminate epimerase | 0 | 0 | -15 | 0 | -15 | -15 | 0,0E+00 | 0,0E+00 | 0,0E+00 | 0,0E+00 | 0,0E+00 | 0,0E+00 | 0,0E+00 | 0,0E+00 | 0,0E+00 | 0,0E+00 | 0,0E+00 | 0,0E+00 | 1368440 | 1369618 | 1179 | - |
| Y11_RS06440 | Y11_13791 | | helix-turn-helix domain-containing protein | -0,55 | -0,86 | -0,51 | -0,31 | 0,04 | 0,35 | 6,1E-02 | 4,5E-04 | 2,4E-02 | 2,4E-01 | 8,9E-01 | 7,5E-02 | 0,0E+00 | 0,0E+00 | 0,0E+00 | 0,0E+00 | 0,0E+00 | 0,0E+00 | 1369952 | 1370434 | 483 | + |
| Y11_RS06445 | Y11_13801 | wrbA | NAD(P)H:quinone oxidoreductase | 0,23 | 2,39 | 2,99 | 2,16 | 2,76 | 0,59 | 5,5E-02 | 9,7E-08 | 1,2E-09 | 3,1E-08 | 8,1E-12 | 3,0E-04 | 0,0E+00 | 4,0E-09 | 1,2E-10 | 1,3E-09 | 9,7E-12 | 0,0E+00 | 1370604 | 1371203 | 600 | + |
| Y11_RS06455 | Y11_13831 | | iron uptake system protein EfeO | 0,36 | 2,16 | 1,02 | 1,8 | 0,66 | -1,14 | 2,8E-06 | 1,5E-11 | 1,9E-03 | 6,8E-11 | 1,9E-02 | 1,1E-03 | 0,0E+00 | 7,2E-12 | 2,3E-05 | 1,4E-11 | 0,0E+00 | 7,0E-05 | 1372715 | 1373845 | 1131 | - |
| Y11_RS06475 | Y11_13871 | | hypothetical protein | 0,41 | -0,64 | 0,36 | -1,05 | -0,05 | 1 | 3,7E-02 | 2,8E-03 | 3,4E-01 | 3,1E-04 | 8,9E-01 | 2,4E-02 | 0,0E+00 | 0,0E+00 | 0,0E+00 | 5,5E-06 | 0,0E+00 | 1,3E-03 | 1376584 | 1376913 | 330 | + |
| Y11_RS06485 | Y11_13901 | putP | sodium/proline symporter PutP | -0,58 | -1,74 | -1,88 | -1,17 | -1,29 | -0,13 | 2,6E-02 | 9,5E-03 | 3,6E-05 | 4,6E-02 | 7,3E-05 | 8,0E-01 | 0,0E+00 | 1,8E-04 | 5,4E-07 | 7,2E-04 | 2,0E-06 | 0,0E+00 | 1378390 | 1379874 | 1485 | - |
| Y11_RS06490 | Y11_13911 | putA | trifunctional transcriptional regulator/proline dehydrogenase/L-glutamate gamma-semialdehyde dehydrogenase | -0,43 | 0,16 | -1,2 | 0,6 | -0,77 | -1,36 | 1,2E-05 | 9,9E-03 | 3,3E-07 | 8,4E-06 | 1,9E-05 | 2,9E-07 | 0,0E+00 | 0,0E+00 | 9,1E-09 | 0,0E+00 | 0,0E+00 | 7,1E-08 | 1380401 | 1384390 | 3990 | + |
| Y11_RS06500 | Y11_13931 | | MarR family transcriptional regulator | 0,47 | -0,56 | -0,13 | -1,04 | -0,6 | 0,43 | 6,2E-02 | 9,5E-03 | 6,1E-01 | 5,3E-04 | 4,9E-02 | 8,2E-02 | 0,0E+00 | 0,0E+00 | 0,0E+00 | 9,1E-06 | 0,0E+00 | 0,0E+00 | 1384982 | 1385353 | 372 | + |
| Y11_RS06505 | Y11_13941 | yecC | L-cystine ABC transporter ATP-binding protein YecC | -0,57 | -0,09 | 0,23 | 0,48 | 0,8 | 0,32 | 5,5E-04 | 1,9E-01 | 1,8E-02 | 1,8E-03 | 1,0E-04 | 3,4E-03 | 0,0E+00 | 0,0E+00 | 0,0E+00 | 0,0E+00 | 0,0E+00 | 0,0E+00 | 1385425 | 1386180 | 756 | - |
| Y11_RS06510 | Y11_13951 | tcyL | cystine ABC transporter permease | -0,31 | -0,51 | 0,35 | -0,2 | 0,65 | 0,85 | 9,3E-03 | 6,3E-04 | 5,3E-01 | 5,1E-02 | 2,5E-01 | 1,5E-01 | 0,0E+00 | 0,0E+00 | 0,0E+00 | 0,0E+00 | 0,0E+00 | 0,0E+00 | 1386183 | 1386845 | 663 | - |
| Y11_RS06515 | Y11_13961 | tcyJ | cystine ABC transporter substrate-binding protein | -0,36 | 0,12 | 1,55 | 0,47 | 1,91 | 1,44 | 6,4E-04 | 1,1E-01 | 2,0E-06 | 9,6E-06 | 2,5E-07 | 2,1E-06 | 0,0E+00 | 0,0E+00 | 4,1E-08 | 0,0E+00 | 1,3E-08 | 3,3E-07 | 1386845 | 1387645 | 801 | - |
| Y11_RS06520 | Y11_13971 | | D-cysteine desulfhydrase | 0,53 | 0,11 | -0,06 | -0,42 | -0,59 | -0,17 | 3,4E-06 | 4,6E-02 | 3,9E-01 | 3,5E-05 | 2,6E-05 | 3,9E-02 | 0,0E+00 | 0,0E+00 | 0,0E+00 | 0,0E+00 | 0,0E+00 | 0,0E+00 | 1387764 | 1388756 | 993 | - |
| Y11_RS06660 | Y11_14281 | fliN | flagellar motor switch protein FliN | 0 | -2,03 | -3,04 | -2,65 | -3,68 | -1,03 | 0,0E+00 | 3,6E-07 | 7,3E-08 | 6,6E-08 | 2,5E-08 | 1,9E-04 | 0,0E+00 | 1,2E-08 | 2,5E-09 | 2,4E-09 | 2,0E-09 | 1,4E-05 | 1415351 | 1415767 | 417 | + |
| Y11_RS06705 | Y11_14381 | flgH | flagellar basal body L-ring protein FlgH | 0 | 0,09 | 0 | -0,37 | 0 | 0,41 | 0,0E+00 | 8,1E-01 | 0,0E+00 | 2,3E-01 | 0,0E+00 | 2,2E-01 | 0,0E+00 | 0,0E+00 | 0,0E+00 | 0,0E+00 | 0,0E+00 | 0,0E+00 | 1423555 | 1424271 | 717 | - |
| Y11_RS06745 | Y11_14461 | flgM | anti-sigma-28 factor FlgM | -0,56 | -1,01 | -1,56 | -0,44 | -1 | -0,55 | 3,3E-02 | 1,3E-03 | 1,7E-04 | 7,3E-03 | 3,6E-04 | 6,8E-03 | 0,0E+00 | 2,5E-05 | 2,3E-06 | 0,0E+00 | 2,0E-05 | 0,0E+00 | 1430075 | 1430374 | 300 | + |
| Y11_RS06755 | Y11_14481 | inv | autotransporter invasin Inv | 0,53 | 1,28 | 4,99 | 0,75 | 4,47 | 3,71 | 4,0E-06 | 1,6E-09 | 9,3E-11 | 8,0E-08 | 2,2E-10 | 8,2E-10 | 0,0E+00 | 1,5E-10 | 2,3E-11 | 0,0E+00 | 7,2E-11 | 1,4E-09 | 1430869 | 1433376 | 2508 | - |
| Y11_RS06785 | Y11_14551 | cheZ | protein phosphatase CheZ | -0,6 | -2,08 | -2,86 | -1,48 | -2,26 | -0,78 | 8,1E-04 | 9,1E-06 | 6,7E-08 | 1,3E-04 | 6,0E-07 | 1,1E-02 | 0,0E+00 | 2,2E-07 | 2,4E-09 | 2,3E-06 | 2,7E-08 | 0,0E+00 | 1441055 | 1441699 | 645 | - |
| Y11_RS06790 | Y11_14561 | cheY | chemotaxis response regulator CheY | -0,06 | -1,74 | -1,96 | -1,68 | -1,9 | -0,22 | 7,8E-01 | 2,7E-06 | 5,8E-07 | 7,4E-05 | 2,4E-05 | 1,8E-01 | 0,0E+00 | 7,3E-08 | 1,4E-08 | 1,4E-06 | 7,0E-07 | 0,0E+00 | 1441709 | 1442098 | 390 | - |
| Y11_RS06795 | Y11_14571 | | chemotaxis response regulator protein-glutamate methylesterase | 0 | -0,34 | -0,14 | -0,85 | -0,76 | 0,15 | 0,0E+00 | 1,9E-02 | 3,9E-01 | 5,8E-04 | 5,5E-03 | 3,8E-01 | 0,0E+00 | 0,0E+00 | 0,0E+00 | 0,0E+00 | 0,0E+00 | 0,0E+00 | 1442285 | 1443334 | 1050 | - |
| Y11_RS06805 | Y11_14591 | | methyl-accepting chemotaxis protein | 0 | -15 | -15 | -2,19 | -2,48 | -0,31 | 0,0E+00 | 0,0E+00 | 0,0E+00 | 5,0E-06 | 4,5E-06 | 7,0E-02 | 0,0E+00 | 0,0E+00 | 0,0E+00 | 1,1E-07 | 1,6E-07 | 0,0E+00 | 1444223 | 1445860 | 1638 | - |
| Y11_RS06815 | Y11_14621 | cheW | chemotaxis protein CheW | 0 | -2,6 | -3,26 | -2,59 | -3,26 | -0,67 | 9,9E-01 | 6,8E-07 | 5,1E-07 | 3,6E-07 | 3,4E-07 | 2,2E-02 | 0,0E+00 | 2,1E-08 | 1,3E-08 | 1,0E-08 | 1,7E-08 | 0,0E+00 | 1448000 | 1448497 | 498 | - |
| Y11_RS06820 | Y11_14641 | cheA | chemotaxis protein CheA | 0 | -0,75 | -2,55 | -0,94 | -2,71 | -1,8 | 0,0E+00 | 1,2E-03 | 3,7E-08 | 2,9E-04 | 1,2E-08 | 4,8E-06 | 0,0E+00 | 0,0E+00 | 1,5E-09 | 0,0E+00 | 1,2E-09 | 6,2E-07 | 1448678 | 1450726 | 2049 | - |
| Y11_RS06845 | Y11_14701 | uspC | universal stress protein UspC | -0,76 | 1,86 | 2,53 | 2,62 | 3,28 | 0,66 | 1,3E-04 | 4,0E-06 | 1,0E-05 | 8,3E-07 | 2,3E-06 | 6,0E-02 | 0,0E+00 | 1,0E-07 | 1,7E-07 | 2,1E-08 | 8,6E-08 | 0,0E+00 | 1454953 | 1455372 | 420 | + |
| Y11_RS06850 | Y11_14711 | | NADP-dependent oxidoreductase | -0,4 | 0,84 | 1,67 | 1,24 | 2,07 | 0,83 | 4,9E-08 | 4,6E-09 | 6,8E-08 | 4,4E-10 | 1,4E-08 | 2,0E-05 | 0,0E+00 | 0,0E+00 | 2,4E-09 | 4,7E-11 | 1,2E-09 | 0,0E+00 | 1455536 | 1456570 | 1035 | + |
| Y11_RS06855 | Y11_14721 | mgtA | magnesium-translocating P-type ATPase | 0,14 | -0,59 | -0,96 | -0,73 | -1,11 | -0,37 | 1,2E-01 | 4,4E-04 | 7,2E-07 | 1,5E-04 | 6,3E-07 | 6,3E-03 | 0,0E+00 | 0,0E+00 | 0,0E+00 | 0,0E+00 | 2,8E-08 | 0,0E+00 | 1456636 | 1459338 | 2703 | - |
| Y11_RS06885 | Y11_14791 | cspE | transcription antiterminator/RNA stability regulator CspE | 0,12 | -2,34 | -3,8 | -2,47 | -3,92 | -1,46 | 2,5E-01 | 5,0E-06 | 4,6E-05 | 1,6E-06 | 3,2E-05 | 2,1E-02 | 0,0E+00 | 1,3E-07 | 6,6E-07 | 3,9E-08 | 9,2E-07 | 1,1E-03 | 1464239 | 1464451 | 213 | - |
| Y11_RS06895 | Y11_14811 | | beta-galactosidase | 0 | -0,96 | 0 | -1,3 | 0 | 1,2 | 0,0E+00 | 5,5E-07 | 0,0E+00 | 2,5E-06 | 0,0E+00 | 3,8E-05 | 0,0E+00 | 0,0E+00 | 0,0E+00 | 5,6E-08 | 0,0E+00 | 3,5E-06 | 1466146 | 1469298 | 3153 | + |
| Y11_RS06905 | Y11_14831 | | NAD(P)H-quinone oxidoreductase | 0,13 | 1,05 | 1,3 | 0,93 | 1,17 | 0,25 | 5,3E-02 | 2,9E-08 | 8,1E-06 | 3,3E-07 | 2,2E-05 | 9,6E-02 | 0,0E+00 | 1,5E-09 | 1,4E-07 | 0,0E+00 | 6,6E-07 | 0,0E+00 | 1470986 | 1471996 | 1011 | + |
| Y11_RS06915 | Y11_14851 | | Lrp/AsnC family transcriptional regulator | 2,06 | 1,35 | 3,2 | -0,71 | 1,13 | 1,84 | 2,5E-07 | 3,7E-04 | 4,7E-08 | 2,2E-02 | 3,4E-04 | 1,3E-04 | 1,9E-08 | 7,7E-06 | 1,8E-09 | 0,0E+00 | 8,3E-06 | 1,1E-05 | 1473020 | 1473478 | 459 | + |

| | | | | | | | | | | | | | | | | | | | | | | | | |
|---|---|---|---|---|---|---|---|---|---|---|---|---|---|---|---|---|---|---|---|---|---|---|---|---|
| Y11_RS06925 | Y11_14881 | ydfZ | putative selenium delivery protein YdfZ | 1,12 | -3,48 | -3,52 | -4,59 | -4,64 | -0,04 | 1,8E-04 | 7,4E-09 | 1,4E-07 | 1,1E-10 | 7,4E-09 | 7,9E-01 | 7,6E-06 | 5,0E-10 | 4,4E-09 | 1,9E-11 | 7,6E-10 | 0,0E+00 | 1474678 | 1474878 | 201 | + |
| Y11_RS06930 | Y11_14891 | | AppA family phytase/histidine-type acid phosphatase | 1,78 | 0,42 | 0,37 | -1,36 | -1,41 | -0,05 | 7,5E-07 | 2,1E-03 | 2,4E-02 | 7,1E-06 | 2,5E-05 | 7,3E-01 | 4,9E-08 | 0,0E+00 | 0,0E+00 | 1,5E-07 | 7,3E-07 | 0,0E+00 | 1475130 | 1476455 | 1326 | - |
| Y11_RS06935 | Y11_14901 | glgC | glucose-1-phosphate adenylyltransferase | -1,11 | 2,21 | 2,79 | 3,31 | 3,89 | 0,58 | 1,2E-04 | 1,3E-06 | 4,6E-06 | 1,7E-08 | 2,1E-07 | 5,0E-02 | 5,0E-06 | 3,9E-08 | 8,4E-08 | 7,7E-10 | 1,2E-08 | 0,0E+00 | 1476787 | 1478064 | 1278 | + |
| Y11_RS06945 | Y11_14931 | | fimbria/pilus periplasmic chaperone | 0,3 | 1,25 | 1,75 | 0,95 | 1,46 | 0,5 | 1,6E-02 | 4,6E-07 | 3,4E-04 | 1,4E-05 | 1,3E-03 | 1,3E-01 | 0,0E+00 | 1,5E-08 | 4,3E-06 | 0,0E+00 | 2,8E-05 | 0,0E+00 | 1479882 | 1480625 | 744 | - |
| Y11_RS06985 | Y11_15001 | mtfA | DgsA anti-repressor MtfA | -0,11 | 2,03 | -0,31 | 2,14 | -0,2 | -2,34 | 5,8E-01 | 6,2E-06 | 9,7E-02 | 1,3E-06 | 1,6E-01 | 1,3E-07 | 0,0E+00 | 1,5E-07 | 0,0E+00 | 3,1E-08 | 0,0E+00 | 4,4E-08 | 1488361 | 1489164 | 804 | + |
| Y11_RS07050 | Y11_15111 | | MBL fold metallo-hydrolase | 0 | -0,4 | 0 | -0,81 | 0 | 1,06 | 0,0E+00 | 3,0E-02 | 0,0E+00 | 8,7E-04 | 0,0E+00 | 5,5E-04 | 0,0E+00 | 0,0E+00 | 0,0E+00 | 0,0E+00 | 0,0E+00 | 3,9E-05 | 1501300 | 1502352 | 1053 | - |
| Y11_RS07055 | Y11_15121 | | RES family NAD+ phosphorylase | 0,43 | -2,76 | -2,2 | -3,18 | -2,63 | 0,55 | 1,2E-02 | 2,9E-07 | 5,7E-04 | 2,8E-07 | 2,1E-04 | 2,3E-01 | 0,0E+00 | 9,9E-09 | 6,9E-06 | 8,1E-09 | 5,4E-06 | 0,0E+00 | 1502559 | 1504052 | 1494 | - |
| Y11_RS07115 | Y11_15231 | | DUF2058 domain-containing protein | 0,25 | -2,71 | -2,9 | -2,96 | -3,15 | -0,18 | 5,9E-02 | 6,6E-10 | 2,2E-09 | 7,0E-09 | 9,8E-09 | 1,2E-01 | 0,0E+00 | 7,9E-11 | 1,8E-10 | 3,9E-10 | 9,5E-10 | 0,0E+00 | 1517536 | 1518075 | 540 | - |
| Y11_RS07120 | Y11_15241 | | PTS sugar transporter subunit IIA | 0,15 | -1,35 | 1,53 | -1,49 | 1,38 | 2,88 | 1,2E-01 | 2,4E-07 | 6,6E-07 | 6,9E-08 | 1,1E-06 | 3,5E-09 | 0,0E+00 | 8,6E-09 | 1,6E-08 | 2,5E-09 | 4,6E-08 | 3,6E-09 | 1518335 | 1518766 | 432 | - |
| Y11_RS07130 | Y11_15261 | lpxO | lipid A hydroxylase LpxO | 0 | 0,08 | 0,9 | -0,68 | 0,15 | 0,83 | 0,0E+00 | 6,2E-01 | 1,2E-02 | 4,3E-03 | 6,3E-01 | 2,5E-02 | 0,0E+00 | 0,0E+00 | 0,0E+00 | 0,0E+00 | 0,0E+00 | 0,0E+00 | 1520935 | 1521837 | 903 | - |
| Y11_RS07175 | Y11_15341 | citF | citrate lyase subunit alpha | 0 | -15 | -15 | -15 | -15 | -0,44 | 0,0E+00 | 0,0E+00 | 0,0E+00 | 0,0E+00 | 0,0E+00 | 4,9E-03 | 0,0E+00 | 0,0E+00 | 0,0E+00 | 0,0E+00 | 0,0E+00 | 0,0E+00 | 1529213 | 1530730 | 1518 | - |
| Y11_RS07195 | Y11_15381 | dpiB | sensor histidine kinase DpiB | 0,63 | 1,2 | 1,53 | 0,57 | 0,9 | 0,33 | 1,5E-01 | 1,2E-03 | 4,5E-04 | 1,0E-01 | 2,6E-02 | 3,6E-02 | 0,0E+00 | 2,4E-05 | 5,5E-06 | 0,0E+00 | 0,0E+00 | 0,0E+00 | 1533534 | 1535111 | 1578 | + |
| Y11_RS07200 | Y11_15391 | dpiA | two-component response regulator DpiA | 0,45 | -0,47 | 0,27 | -0,92 | -0,17 | 0,74 | 2,9E-03 | 3,5E-02 | 7,6E-02 | 8,6E-04 | 2,1E-01 | 5,4E-03 | 0,0E+00 | 0,0E+00 | 0,0E+00 | 0,0E+00 | 0,0E+00 | 0,0E+00 | 1535133 | 1535819 | 687 | + |
| Y11_RS07220 | Y11_15431 | | ABC transporter substrate-binding protein | 0,04 | -3,52 | -2,36 | -3,56 | -2,4 | 1,16 | 9,0E-01 | 3,3E-07 | 3,5E-06 | 6,2E-07 | 7,1E-06 | 1,2E-05 | 0,0E+00 | 1,1E-08 | 6,7E-08 | 1,6E-08 | 2,4E-07 | 1,4E-06 | 1539751 | 1540782 | 1032 | + |
| Y11_RS07230 | Y11_15451 | fbpC | ferric ABC transporter ATP-binding protein | -0,03 | -1,52 | -1,43 | -1,49 | -1,4 | 0,08 | 8,6E-01 | 2,6E-04 | 5,3E-04 | 1,6E-04 | 3,6E-04 | 7,9E-01 | 0,0E+00 | 5,4E-06 | 6,5E-06 | 2,9E-06 | 8,7E-06 | 0,0E+00 | 1543138 | 1544229 | 1092 | + |
| Y11_RS07240 | Y11_15461 | | oxidoreductase | -0,01 | 0,17 | 0,84 | 0,18 | 0,85 | 0,67 | 8,8E-01 | 2,1E-01 | 2,6E-05 | 2,2E-01 | 6,2E-05 | 1,9E-03 | 0,0E+00 | 0,0E+00 | 0,0E+00 | 0,0E+00 | 0,0E+00 | 0,0E+00 | 1544611 | 1545501 | 891 | - |
| Y11_RS07255 | Y11_15491 | | TetR/AcrR family transcriptional regulator | -0,2 | -0,03 | 0 | 0,16 | 0,2 | 0,04 | 4,2E-02 | 7,0E-01 | 9,8E-01 | 3,7E-03 | 1,6E-01 | 7,9E-01 | 0,0E+00 | 0,0E+00 | 0,0E+00 | 0,0E+00 | 0,0E+00 | 0,0E+00 | 1547702 | 1548286 | 585 | + |
| Y11_RS07270 | Y11_15521 | | DeoR/GlpR family DNA-binding transcription regulator | 0,99 | 2,23 | 3,23 | 1,25 | 2,25 | 1 | 2,9E-04 | 6,9E-06 | 3,3E-06 | 2,6E-04 | 3,6E-05 | 1,2E-02 | 0,0E+00 | 1,7E-07 | 6,4E-08 | 4,6E-06 | 1,0E-06 | 6,8E-04 | 1550761 | 1551522 | 762 | - |
| Y11_RS07280 | Y11_15541 | | hypothetical protein | 15 | -0,81 | -0,24 | -15 | -15 | 0,57 | 0,0E+00 | 4,2E-07 | 2,5E-01 | 0,0E+00 | 0,0E+00 | 2,1E-02 | 0,0E+00 | 0,0E+00 | 0,0E+00 | 0,0E+00 | 0,0E+00 | 0,0E+00 | 1552614 | 1553453 | 840 | - |
| Y11_RS07295 | Y11_15581 | argG | argininosuccinate synthase | -0,25 | 0,31 | 0,26 | 0,56 | 0,5 | -0,05 | 3,4E-05 | 6,7E-05 | 7,0E-03 | 2,3E-06 | 1,5E-04 | 5,2E-01 | 0,0E+00 | 0,0E+00 | 0,0E+00 | 0,0E+00 | 0,0E+00 | 0,0E+00 | 1557120 | 1558484 | 1365 | + |
| Y11_RS07305 | Y11_15601 | | hypothetical protein | 0,45 | -2,5 | -2,19 | -2,96 | -2,64 | 0,32 | 1,1E-01 | 7,4E-06 | 3,0E-04 | 8,7E-08 | 2,8E-05 | 3,3E-01 | 0,0E+00 | 1,8E-07 | 3,9E-06 | 3,0E-09 | 8,2E-07 | 0,0E+00 | 1559095 | 1559922 | 828 | - |
| Y11_RS07370 | Y11_15731 | | fimbria/pilus periplasmic chaperone | -0,05 | 0,16 | 15 | 0,21 | 15 | 0 | 8,3E-01 | 4,9E-01 | 0,0E+00 | 3,1E-01 | 0,0E+00 | 0,0E+00 | 0,0E+00 | 0,0E+00 | 0,0E+00 | 0,0E+00 | 0,0E+00 | 0,0E+00 | 1577737 | 1578483 | 747 | - |
| Y11_RS07420 | Y11_15851 | | DcrB-related protein | 0 | -15 | -15 | -1,98 | -1,45 | 0,54 | 0,0E+00 | 0,0E+00 | 0,0E+00 | 3,8E-06 | 1,1E-04 | 6,8E-02 | 0,0E+00 | 0,0E+00 | 0,0E+00 | 8,4E-08 | 2,8E-06 | 0,0E+00 | 1591595 | 1592080 | 486 | + |
| Y11_RS07440 | Y11_15901 | cobT | nicotinate-nucleotide--dimethylbenzimidazole phosphoribosyltransferase | -0,11 | 0,03 | -0,28 | 0,13 | -0,17 | -0,3 | 4,7E-01 | 8,5E-01 | 1,3E-01 | 3,9E-01 | 3,6E-01 | 1,1E-01 | 0,0E+00 | 0,0E+00 | 0,0E+00 | 0,0E+00 | 0,0E+00 | 0,0E+00 | 1594460 | 1595518 | 1059 | - |
| Y11_RS07455 | Y11_15931 | cobU | bifunctional adenosylcobinamide kinase/adenosylcobinamide-phosphate guanylyltransferase | -0,23 | 1,08 | -0,21 | 1,31 | 0,02 | -1,29 | 4,4E-01 | 1,9E-01 | 6,2E-01 | 1,2E-01 | 9,6E-01 | 1,4E-01 | 0,0E+00 | 3,6E-03 | 0,0E+00 | 1,8E-03 | 0,0E+00 | 7,5E-03 | 1596939 | 1597484 | 546 | - |
| Y11_RS07485 | Y11_16001 | | cobalt-factor II C(20)-methyltransferase | -0,23 | -0,62 | -0,75 | -0,39 | -0,52 | -0,13 | 1,2E-01 | 3,2E-03 | 1,3E-03 | 2,3E-02 | 7,6E-03 | 4,6E-01 | 0,0E+00 | 0,0E+00 | 0,0E+00 | 0,0E+00 | 0,0E+00 | 0,0E+00 | 1601714 | 1602427 | 714 | - |
| Y11_RS07510 | Y11_16051 | | cobalt-precorrin-4 methyltransferase | 0,1 | -0,79 | -0,51 | -0,89 | -0,61 | 0,28 | 4,0E-01 | 5,1E-04 | 3,6E-03 | 2,5E-05 | 6,4E-05 | 4,8E-02 | 0,0E+00 | 0,0E+00 | 0,0E+00 | 0,0E+00 | 0,0E+00 | 0,0E+00 | 1605803 | 1606594 | 792 | - |
| Y11_RS07555 | Y11_16151 | pduA | propanediol utilization microcompartment protein PduA | 0 | -0,38 | 0 | 0,22 | 0 | 0,06 | 0,0E+00 | 2,0E-03 | 0,0E+00 | 8,9E-02 | 0,0E+00 | 8,4E-01 | 0,0E+00 | 0,0E+00 | 0,0E+00 | 0,0E+00 | 0,0E+00 | 0,0E+00 | 1615176 | 1615472 | 297 | + |
| Y11_RS07560 | Y11_16161 | pduB | propanediol utilization microcompartment protein PduB | 0,69 | -0,01 | 0,58 | -0,7 | -0,11 | 0,59 | 1,5E-02 | 9,6E-01 | 1,1E-03 | 1,6E-02 | 6,3E-01 | 1,5E-03 | 0,0E+00 | 0,0E+00 | 0,0E+00 | 0,0E+00 | 0,0E+00 | 0,0E+00 | 1615472 | 1616299 | 828 | + |
| Y11_RS07590 | Y11_16221 | pduJ | propanediol utilization microcompartment protein PduJ | 0,16 | -0,54 | -0,2 | -0,7 | -0,37 | 0,33 | 1,6E-01 | 3,1E-05 | 2,8E-02 | 6,0E-05 | 6,4E-03 | 3,2E-04 | 0,0E+00 | 0,0E+00 | 0,0E+00 | 0,0E+00 | 0,0E+00 | 0,0E+00 | 1621730 | 1622005 | 276 | + |
| Y11_RS07625 | Y11_16291 | | iron-containing alcohol dehydrogenase | 0,11 | -0,36 | -0,22 | -0,47 | -0,33 | 0,14 | 3,5E-01 | 6,8E-02 | 1,6E-01 | 4,2E-02 | 9,0E-02 | 5,3E-01 | 0,0E+00 | 0,0E+00 | 0,0E+00 | 0,0E+00 | 0,0E+00 | 0,0E+00 | 1626480 | 1627595 | 1116 | + |
| Y11_RS07670 | Y11_16381 | | autoinducer 2 ABC transporter substrate-binding protein | -0,34 | -0,55 | 1,41 | -0,21 | 1,75 | 1,96 | 5,4E-03 | 8,3E-05 | 1,2E-06 | 9,7E-02 | 1,2E-06 | 2,9E-07 | 0,0E+00 | 0,0E+00 | 2,6E-08 | 0,0E+00 | 4,9E-08 | 7,1E-08 | 1635035 | 1635970 | 936 | + |
| Y11_RS07675 | Y11_16391 | | sugar ABC transporter ATP-binding protein | -0,39 | -1,85 | 0,46 | -1,46 | 0,85 | 2,31 | 1,2E-02 | 6,5E-07 | 1,4E-03 | 1,7E-05 | 2,1E-04 | 2,2E-07 | 0,0E+00 | 2,0E-08 | 0,0E+00 | 3,4E-07 | 0,0E+00 | 6,0E-08 | 1636207 | 1637688 | 1482 | + |
| Y11_RS07690 | Y11_16421 | | DUF496 family protein | 0,29 | 0,42 | -0,13 | 0,13 | -0,42 | -0,55 | 1,1E-01 | 1,4E-02 | 3,1E-01 | 4,2E-01 | 1,5E-02 | 7,4E-04 | 0,0E+00 | 0,0E+00 | 0,0E+00 | 0,0E+00 | 0,0E+00 | 0,0E+00 | 1639779 | 1640096 | 318 | - |
| Y11_RS07695 | Y11_16431 | | FUSC family protein | 0,42 | 1,21 | 0 | 0,82 | 1,28 | 0,45 | 2,3E-01 | 8,6E-03 | 0,0E+00 | 7,5E-02 | 1,5E-03 | 1,7E-01 | 0,0E+00 | 1,7E-04 | 0,0E+00 | 0,0E+00 | 3,3E-05 | 0,0E+00 | 1640260 | 1641318 | 1059 | - |
| Y11_RS07700 | Y11_16441 | | SDR family oxidoreductase | 0 | 0 | -15 | 0 | -15 | -2,62 | 0,0E+00 | 0,0E+00 | 0,0E+00 | 0,0E+00 | 0,0E+00 | 1,2E-06 | 0,0E+00 | 0,0E+00 | 0,0E+00 | 0,0E+00 | 0,0E+00 | 2,0E-07 | 1641464 | 1642573 | 1110 | - |
| Y11_RS07705 | Y11_16451 | sbcB | exodeoxyribonuclease I | -0,23 | -0,1 | 0,31 | 0,13 | 0,54 | 0,41 | 1,4E-02 | 2,4E-01 | 1,4E-02 | 6,5E-02 | 1,7E-04 | 1,7E-03 | 0,0E+00 | 0,0E+00 | 0,0E+00 | 0,0E+00 | 0,0E+00 | 0,0E+00 | 1642812 | 1644242 | 1431 | + |
| Y11_RS07720 | Y11_16471 | | oxidoreductase | -0,23 | 0,38 | 0,71 | 0,62 | 0,94 | 0,32 | 4,0E-02 | 3,1E-03 | 2,1E-02 | 2,4E-04 | 5,6E-03 | 2,3E-01 | 0,0E+00 | 0,0E+00 | 0,0E+00 | 0,0E+00 | 0,0E+00 | 0,0E+00 | 1646173 | 1647264 | 1092 | - |
| Y11_RS07735 | Y11_16511 | | SDR family oxidoreductase | -1,91 | -1,19 | -0,6 | 0,72 | 1,31 | 0,59 | 6,0E-10 | 2,3E-08 | 6,8E-07 | 9,2E-07 | 1,0E-09 | 4,3E-07 | 1,5E-10 | 1,2E-09 | 0,0E+00 | 0,0E+00 | 1,7E-10 | 0,0E+00 | 1650222 | 1651049 | 828 | - |
| Y11_RS07740 | Y11_16521 | hisG | ATP phosphoribosyltransferase | 1,45 | -1,05 | -1,84 | -2,5 | -3,29 | -0,79 | 1,8E-08 | 3,3E-07 | 3,5E-08 | 8,9E-10 | 6,1E-10 | 4,6E-05 | 2,1E-09 | 1,1E-08 | 1,5E-09 | 8,0E-11 | 1,3E-10 | 0,0E+00 | 1651595 | 1652494 | 900 | + |
| Y11_RS07745 | Y11_16531 | hisD | histidinol dehydrogenase | 1,17 | -1,13 | -1,88 | -2,3 | -3,05 | -0,75 | 9,6E-07 | 1,4E-06 | 1,1E-09 | 2,5E-08 | 4,2E-10 | 2,4E-05 | 6,3E-08 | 3,9E-08 | 1,1E-10 | 1,0E-09 | 1,0E-10 | 0,0E+00 | 1652498 | 1653823 | 1326 | + |
| Y11_RS07750 | Y11_16541 | hisC | histidinol-phosphate transaminase | 1,32 | -0,97 | -1,77 | -2,29 | -3,09 | -0,8 | 2,1E-09 | 2,9E-09 | 3,0E-06 | 7,8E-12 | 4,3E-08 | 8,1E-04 | 4,2E-10 | 0,0E+00 | 5,9E-08 | 5,4E-12 | 3,1E-09 | 0,0E+00 | 1653820 | 1654944 | 1125 | + |
| Y11_RS07755 | Y11_16551 | hisB | bifunctional histidinol-phosphatase/imidazoleglycerol-phosphate dehydratase HisB | 0,93 | -0,83 | -1,15 | -1,75 | -2,08 | -0,33 | 1,9E-07 | 6,5E-05 | 1,2E-05 | 5,6E-07 | 2,8E-07 | 8,0E-02 | 0,0E+00 | 0,0E+00 | 2,0E-07 | 1,5E-08 | 1,4E-08 | 0,0E+00 | 1654941 | 1656008 | 1068 | + |
| Y11_RS07760 | Y11_16561 | hisH | imidazole glycerol phosphate synthase subunit HisH | 0,96 | -0,81 | -1,01 | -1,77 | -1,96 | -0,2 | 2,0E-06 | 3,0E-08 | 8,6E-04 | 3,4E-08 | 1,4E-05 | 3,5E-01 | 0,0E+00 | 0,0E+00 | 1,0E-05 | 1,3E-09 | 4,3E-07 | 0,0E+00 | 1656008 | 1656598 | 591 | + |
| Y11_RS07765 | Y11_16571 | hisA | 1-(5-phosphoribosyl)-5-[(5-phosphoribosylamino)methylideneamino]imidazole-4-carboxamide isomerase | 1,03 | -0,94 | -1,42 | -1,97 | -2,45 | -0,48 | 4,3E-08 | 6,0E-11 | 1,9E-09 | 2,4E-10 | 3,6E-10 | 8,5E-06 | 4,3E-09 | 0,0E+00 | 1,6E-10 | 3,0E-11 | 9,7E-11 | 0,0E+00 | 1656604 | 1657341 | 738 | + |
| Y11_RS07770 | Y11_16581 | hisF | imidazole glycerol phosphate synthase subunit HisF | 0,94 | -0,89 | -1,32 | -1,83 | -2,26 | -0,43 | 2,4E-07 | 1,2E-07 | 2,4E-07 | 1,2E-10 | 2,3E-09 | 3,3E-04 | 0,0E+00 | 0,0E+00 | 6,7E-09 | 2,0E-11 | 3,2E-10 | 0,0E+00 | 1657323 | 1658099 | 777 | + |
| Y11_RS07775 | Y11_16591 | hisIE | bifunctional phosphoribosyl-AMP cyclohydrolase/phosphoribosyl-ATP diphosphatase HisIE | 0,39 | -0,54 | -0,96 | -0,92 | -1,35 | -0,43 | 5,6E-04 | 7,1E-05 | 5,9E-03 | 3,9E-06 | 8,7E-04 | 1,4E-01 | 0,0E+00 | 0,0E+00 | 0,0E+00 | 0,0E+00 | 2,0E-05 | 0,0E+00 | 1658093 | 1658707 | 615 | + |
| Y11_RS07780 | Y11_16601 | gndA | NADP-dependent phosphogluconate dehydrogenase | -0,13 | 1,54 | 2,04 | 1,67 | 2,17 | 0,5 | 1,0E-01 | 1,0E-07 | 2,1E-08 | 4,5E-08 | 1,1E-08 | 1,3E-03 | 0,0E+00 | 4,3E-09 | 9,3E-10 | 1,7E-09 | 1,1E-09 | 0,0E+00 | 1658963 | 1660369 | 1407 | - |
| Y11_RS07790 | Y11_16621 | galU | UTP--glucose-1-phosphate uridylyltransferase GalU | -0,54 | 0,32 | 0 | 0,86 | 0,54 | -0,33 | 5,8E-06 | 8,8E-05 | 9,9E-01 | 6,8E-08 | 6,6E-04 | 9,8E-03 | 0,0E+00 | 0,0E+00 | 0,0E+00 | 0,0E+00 | 0,0E+00 | 0,0E+00 | 1660623 | 1661516 | 894 | - |
| Y11_RS07795 | Y11_16631 | galU | UTP--glucose-1-phosphate uridylyltransferase GalU | -0,64 | 0,58 | -0,24 | 1,22 | 0,4 | -0,82 | 4,9E-06 | 5,0E-06 | 4,2E-02 | 1,7E-08 | 3,6E-03 | 2,8E-05 | 0,0E+00 | 0,0E+00 | 0,0E+00 | 8,0E-10 | 0,0E+00 | 0,0E+00 | 1661582 | 1662472 | 891 | - |
| Y11_RS07825 | Y11_16691 | | hypothetical protein | 0,33 | 0,32 | 0,4 | -0,01 | 0,07 | 0,08 | 6,5E-02 | 3,0E-02 | 2,4E-02 | 9,6E-01 | 7,1E-01 | 6,3E-01 | 0,0E+00 | 0,0E+00 | 0,0E+00 | 0,0E+00 | 0,0E+00 | 0,0E+00 | 1667964 | 1668917 | 954 | - |
| Y11_RS07835 | Y11_16711 | | glycosyltransferase | -0,21 | 0,59 | 0,77 | 0,8 | 0,98 | 0,18 | 8,6E-04 | 3,0E-08 | 2,6E-05 | 3,1E-08 | 6,1E-06 | 7,9E-02 | 0,0E+00 | 0,0E+00 | 0,0E+00 | 0,0E+00 | 0,0E+00 | 0,0E+00 | 1669725 | 1671005 | 1281 | - |
| Y11_RS07840 | Y11_16721 | | SDR family oxidoreductase | 0 | 1,27 | 1,83 | 1,27 | 1,83 | 0,55 | 9,7E-01 | 1,6E-08 | 1,5E-07 | 3,5E-09 | 1,1E-07 | 7,5E-04 | 0,0E+00 | 9,1E-10 | 4,7E-09 | 2,2E-10 | 6,5E-09 | 0,0E+00 | 1671017 | 1671886 | 870 | - |
| Y11_RS07845 | Y11_16731 | rfbC | dTDP-4-dehydrorhamnose 3,5-epimerase | 0,5 | 0,2 | 0,29 | -0,3 | -0,21 | 0,09 | 2,1E-03 | 3,8E-02 | 1,1E-02 | 2,9E-02 | 1,1E-01 | 3,5E-01 | 0,0E+00 | 0,0E+00 | 0,0E+00 | 0,0E+00 | 0,0E+00 | 0,0E+00 | 1671889 | 1672452 | 564 | - |
| Y11_RS07850 | Y11_16741 | rfbE | O-antigen export ABC transporter ATP-binding protein RfbE | -0,42 | 0,34 | 0,41 | 0,76 | 0,83 | 0,08 | 7,3E-03 | 4,2E-04 | 1,7E-02 | 2,2E-04 | 1,3E-03 | 6,0E-01 | 0,0E+00 | 0,0E+00 | 0,0E+00 | 0,0E+00 | 0,0E+00 | 0,0E+00 | 1672672 | 1673391 | 720 | - |
| Y11_RS07855 | Y11_16751 | | ABC transporter permease | -0,13 | 0,3 | 0,64 | 0,43 | 0,77 | 0,34 | 3,6E-01 | 2,2E-01 | 4,7E-02 | 8,0E-02 | 2,0E-02 | 3,2E-01 | 0,0E+00 | 0,0E+00 | 0,0E+00 | 0,0E+00 | 0,0E+00 | 0,0E+00 | 1673394 | 1674173 | 780 | - |
| Y11_RS07860 | Y11_16761 | | rhamnosyltransferase | 0,22 | 0,39 | 1,32 | 0,17 | 1,11 | 0,96 | 9,9E-02 | 1,7E-02 | 2,3E-04 | 3,1E-01 | 1,2E-03 | 3,2E-03 | 0,0E+00 | 0,0E+00 | 3,0E-06 | 0,0E+00 | 2,7E-05 | 0,0E+00 | 1674654 | 1675577 | 924 | - |
| Y11_RS07870 | Y11_16781 | rfbA | glucose-1-phosphate thymidylyltransferase RfbA | 0,41 | 0,38 | 0,64 | -0,03 | 0,23 | 0,26 | 2,2E-05 | 2,0E-04 | 2,3E-04 | 6,5E-01 | 5,4E-02 | 4,4E-02 | 0,0E+00 | 0,0E+00 | 0,0E+00 | 0,0E+00 | 0,0E+00 | 0,0E+00 | 1676811 | 1677680 | 870 | - |
| Y11_RS07880 | Y11_16811 | | TerC family protein | 0,12 | 0,14 | 0,44 | 0,02 | 0,32 | 0,3 | 3,3E-01 | 3,8E-01 | 1,6E-02 | 8,9E-01 | 2,8E-02 | 7,7E-02 | 0,0E+00 | 0,0E+00 | 0,0E+00 | 0,0E+00 | 0,0E+00 | 0,0E+00 | 1680658 | 1682244 | 1587 | + |
| Y11_RS07890 | Y11_16831 | asmA | outer membrane assembly protein AsmA | 0,27 | 0,09 | 0,65 | -0,18 | 0,39 | 0,57 | 7,0E-05 | 7,1E-02 | 1,7E-04 | 6,2E-03 | 5,2E-03 | 6,3E-04 | 0,0E+00 | 0,0E+00 | 0,0E+00 | 0,0E+00 | 0,0E+00 | 0,0E+00 | 1684028 | 1685875 | 1848 | - |
| Y11_RS07895 | Y11_16841 | dcd | dCTP deaminase | 0,51 | -0,15 | 0,22 | -0,66 | -0,28 | 0,38 | 2,0E-03 | 1,6E-01 | 7,7E-02 | 3,9E-05 | 1,8E-02 | 1,3E-03 | 0,0E+00 | 0,0E+00 | 0,0E+00 | 0,0E+00 | 0,0E+00 | 0,0E+00 | 1685940 | 1686521 | 582 | - |
| Y11_RS07900 | Y11_16851 | udk | uridine kinase | 0,51 | -0,1 | 0,17 | -0,62 | -0,35 | 0,27 | 2,8E-03 | 2,8E-01 | 2,0E-01 | 4,5E-04 | 3,3E-02 | 3,6E-02 | 0,0E+00 | 0,0E+00 | 0,0E+00 | 0,0E+00 | 0,0E+00 | 0,0E+00 | 1686607 | 1687248 | 642 | - |
| Y11_RS07905 | Y11_16861 | apbC | iron-sulfur cluster carrier protein ApbC | -0,34 | 0,91 | 0,91 | 1,26 | 1,26 | 0 | 8,3E-04 | 6,0E-09 | 1,0E-05 | 2,9E-08 | 2,2E-06 | 9,9E-01 | 0,0E+00 | 0,0E+00 | 0,0E+00 | 1,2E-09 | 8,4E-08 | 0,0E+00 | 1687604 | 1688716 | 1113 | - |
| Y11_RS07910 | Y11_16881 | metG | methionine--tRNA ligase | -0,02 | 0,11 | 0,33 | 0,13 | 0,35 | 0,22 | 5,2E-01 | 2,7E-04 | 3,5E-02 | 1,6E-03 | 2,9E-02 | 1,3E-01 | 0,0E+00 | 0,0E+00 | 0,0E+00 | 0,0E+00 | 0,0E+00 | 0,0E+00 | 1688996 | 1691023 | 2028 | + |
| Y11_RS07920 | Y11_16901 | | sugar ABC transporter substrate-binding protein | 0,56 | 0,68 | 1,11 | 0,12 | 0,54 | 0,43 | 4,1E-05 | 3,3E-05 | 1,1E-06 | 1,8E-01 | 1,6E-04 | 1,7E-03 | 0,0E+00 | 0,0E+00 | 2,6E-08 | 0,0E+00 | 0,0E+00 | 0,0E+00 | 1692397 | 1693473 | 1077 | + |
| Y11_RS07925 | Y11_16911 | | LVIVD repeat-containing protein | 0,24 | 0,22 | 0,56 | -0,02 | 0,32 | 0,33 | 1,5E-01 | 1,8E-01 | 5,7E-02 | 9,2E-01 | 2,7E-01 | 2,4E-01 | 0,0E+00 | 0,0E+00 | 0,0E+00 | 0,0E+00 | 0,0E+00 | 0,0E+00 | 1693657 | 1694910 | 1254 | + |
| Y11_RS07940 | Y11_16931 | | CidA/LrgA family protein | 0,35 | 0,95 | 0,76 | 0,59 | 0,41 | -0,19 | 7,2E-03 | 2,8E-05 | 8,3E-05 | 1,6E-03 | 9,9E-03 | 2,0E-01 | 0,0E+00 | 0,0E+00 | 0,0E+00 | 0,0E+00 | 0,0E+00 | 0,0E+00 | 1696571 | 1696978 | 408 | + |

| | | | | | | | | | | | | | | | | | | | | | | | | | |
|---|---|---|---|---|---|---|---|---|---|---|---|---|---|---|---|---|---|---|---|---|---|---|---|---|---|
| Y11_RS07945 | Y11_16941 | | CidB/LrgB family autolysis modulator | 0,15 | 0,53 | 0,11 | 0,38 | -0,03 | -0,41 | 6,1E-01 | 1,8E-02 | 5,6E-01 | 1,7E-01 | 9,1E-01 | 3,6E-02 | 0,0E+00 | 0,0E+00 | 0,0E+00 | 0,0E+00 | 0,0E+00 | 0,0E+00 | 1696975 | 1697670 | 696 | + |
| Y11_RS07950 | Y11_16951 | cdd | cytidine deaminase | -0,31 | 1,46 | 1,02 | 1,76 | 1,33 | -0,44 | 7,7E-02 | 3,6E-07 | 2,6E-04 | 3,9E-07 | 7,3E-05 | 1,2E-02 | 0,0E+00 | 1,2E-08 | 3,4E-06 | 1,1E-08 | 2,0E-06 | 0,0E+00 | 1697809 | 1698705 | 897 | + |
| Y11_RS07960 | Y11_16961 | | NAD-dependent malic enzyme | 1,27 | -0,07 | -0,17 | -1,34 | -1,44 | -0,1 | 5,5E-08 | 1,9E-01 | 6,8E-02 | 6,1E-08 | 4,1E-07 | 2,5E-01 | 5,4E-09 | 0,0E+00 | 0,0E+00 | 2,2E-09 | 2,0E-08 | 0,0E+00 | 1699591 | 1701288 | 1698 | + |
| Y11_RS07965 | Y11_16971 | sanA | outer membrane permeability protein SanA | 0,42 | 0,58 | 0,98 | 0,16 | 0,56 | 0,4 | 2,2E-02 | 4,3E-03 | 1,0E-04 | 2,5E-01 | 1,5E-03 | 9,1E-03 | 0,0E+00 | 0,0E+00 | 0,0E+00 | 0,0E+00 | 0,0E+00 | 0,0E+00 | 1701454 | 1702299 | 846 | + |
| Y11_RS08050 | Y11_17151 | mglC | galactose/methyl galactoside ABC transporter permease MglC | -4,09 | -0,02 | -0,01 | 4,07 | 4,09 | 0 | 1,1E-07 | 9,3E-01 | 9,6E-01 | 1,3E-07 | 2,4E-08 | 9,8E-01 | 9,6E-09 | 0,0E+00 | 0,0E+00 | 4,2E-09 | 2,0E-09 | 0,0E+00 | 1720841 | 1721851 | 1011 | - |
| Y11_RS08055 | Y11_17161 | mglA | galactose/methyl galactoside ABC transporter ATP-binding protein MglA | -3,55 | 0,02 | 1,33 | 3,57 | 4,88 | 1,31 | 2,8E-12 | 7,7E-01 | 1,4E-06 | 9,8E-13 | 1,9E-11 | 1,3E-06 | 3,0E-12 | 0,0E+00 | 3,1E-08 | 1,9E-12 | 1,6E-11 | 2,1E-07 | 1721872 | 1723392 | 1521 | - |
| Y11_RS08060 | Y11_17171 | mglB | galactose/glucose ABC transporter substrate-binding protein MglB | -3,1 | 1,08 | 3,03 | 4,18 | 6,13 | 1,95 | 1,9E-08 | 1,9E-05 | 5,8E-08 | 7,0E-11 | 7,8E-11 | 1,9E-07 | 2,1E-09 | 4,5E-07 | 2,1E-09 | 1,4E-11 | 3,9E-11 | 5,5E-08 | 1723585 | 1724577 | 993 | - |
| Y11_RS08065 | Y11_17181 | galS | HTH-type transcriptional regulator GalS | -2,3 | 0,64 | 15 | 2,94 | 15 | 0 | 8,1E-04 | 3,4E-03 | 0,0E+00 | 1,3E-04 | 0,0E+00 | 0,0E+00 | 3,1E-05 | 0,0E+00 | 0,0E+00 | 2,3E-06 | 0,0E+00 | 0,0E+00 | 1724931 | 1725968 | 1038 | - |
| Y11_RS08075 | Y11_17211 | folE | GTP cyclohydrolase I FolE | 0,66 | -0,21 | -0,49 | -0,87 | -1,15 | -0,28 | 2,3E-08 | 1,8E-04 | 6,5E-05 | 4,4E-09 | 1,1E-07 | 2,8E-03 | 0,0E+00 | 0,0E+00 | 0,0E+00 | 0,0E+00 | 6,5E-09 | 0,0E+00 | 1727371 | 1728033 | 663 | - |
| Y11_RS08085 | Y11_17231 | | LysR family transcriptional regulator | 1,04 | 0,59 | 0,75 | -0,45 | -0,29 | 0,16 | 4,9E-02 | 2,5E-01 | 2,4E-01 | 8,7E-02 | 5,0E-01 | 7,3E-01 | 1,6E-03 | 0,0E+00 | 0,0E+00 | 0,0E+00 | 0,0E+00 | 0,0E+00 | 1729515 | 1730384 | 870 | + |
| Y11_RS08090 | Y11_17241 | | S-(hydroxymethyl)glutathione dehydrogenase/class III alcohol dehydrogenase | 0,34 | 0,64 | 0,36 | 0,3 | 0,02 | -0,28 | 7,2E-05 | 2,2E-07 | 1,1E-05 | 6,1E-04 | 7,0E-01 | 4,3E-04 | 0,0E+00 | 0,0E+00 | 0,0E+00 | 0,0E+00 | 0,0E+00 | 0,0E+00 | 1730529 | 1731683 | 1155 | + |
| Y11_RS08095 | Y11_17251 | fghA | S-formylglutathione hydrolase | 0,14 | 0,41 | 0,49 | 0,27 | 0,35 | 0,09 | 4,3E-01 | 2,3E-03 | 6,6E-04 | 1,4E-01 | 6,4E-02 | 3,7E-01 | 0,0E+00 | 0,0E+00 | 0,0E+00 | 0,0E+00 | 0,0E+00 | 0,0E+00 | 1731704 | 1732546 | 843 | + |
| Y11_RS08125 | Y11_17311 | | DUF924 domain-containing protein | -0,25 | 0,26 | 1,08 | 0,51 | 1,33 | 0,82 | 5,8E-04 | 5,4E-02 | 1,8E-04 | 1,8E-03 | 3,7E-05 | 2,9E-03 | 0,0E+00 | 0,0E+00 | 2,4E-06 | 0,0E+00 | 1,0E-06 | 0,0E+00 | 1739092 | 1739631 | 540 | - |
| Y11_RS08130 | Y11_17321 | moeA | molybdopterin molybdotransferase MoeA | 0,31 | 0,26 | 0,17 | -0,06 | -0,14 | -0,09 | 2,4E-03 | 2,1E-02 | 6,5E-02 | 5,1E-01 | 7,7E-02 | 3,6E-01 | 0,0E+00 | 0,0E+00 | 0,0E+00 | 0,0E+00 | 0,0E+00 | 0,0E+00 | 1739832 | 1741067 | 1236 | + |
| Y11_RS08135 | Y11_17331 | moeB | molybdopterin-synthase adenylyltransferase MoeB | 0,24 | 0,2 | 1,14 | -0,04 | 0,9 | 0,94 | 1,4E-02 | 3,8E-03 | 1,2E-06 | 6,5E-01 | 4,2E-05 | 8,8E-06 | 0,0E+00 | 0,0E+00 | 2,6E-08 | 0,0E+00 | 0,0E+00 | 0,0E+00 | 1741104 | 1741859 | 756 | + |
| Y11_RS08145 | Y11_17361 | | ABC-F family ATPase | -0,07 | -1,52 | -2,81 | -1,45 | -2,74 | -1,29 | 5,2E-01 | 3,9E-07 | 1,7E-07 | 4,1E-06 | 4,8E-07 | 1,4E-04 | 0,0E+00 | 1,3E-08 | 5,0E-09 | 8,9E-08 | 2,3E-08 | 1,1E-05 | 1743502 | 1745097 | 1596 | - |
| Y11_RS08165 | Y11_17401 | ompX | outer membrane protein OmpX | -1,21 | 0 | -2,05 | 1,21 | -0,85 | -2,06 | 9,2E-08 | 9,8E-01 | 1,2E-05 | 3,7E-07 | 4,3E-03 | 1,4E-05 | 8,6E-09 | 0,0E+00 | 2,0E-07 | 1,0E-08 | 0,0E+00 | 1,5E-06 | 1748993 | 1749520 | 528 | - |
| Y11_RS08185 | Y11_17431 | | fasciclin domain-containing protein | 4,01 | 2,61 | 3,84 | -1,4 | -0,17 | 1,23 | 2,3E-08 | 8,2E-08 | 8,8E-09 | 7,8E-05 | 4,3E-01 | 6,3E-05 | 2,5E-09 | 3,4E-09 | 4,8E-10 | 1,5E-06 | 0,0E+00 | 5,4E-06 | 1751492 | 1752052 | 561 | + |
| Y11_RS08190 | Y11_17441 | | sigma-70 family RNA polymerase sigma factor | -0,56 | 1,9 | 3,21 | 2,47 | 3,76 | 1,31 | 1,3E-04 | 1,0E-06 | 2,5E-09 | 3,9E-07 | 3,9E-09 | 8,9E-05 | 0,0E+00 | 3,1E-08 | 1,9E-10 | 1,1E-08 | 4,6E-10 | 7,5E-06 | 1752113 | 1752667 | 555 | + |
| Y11_RS08195 | Y11_17451 | | anti-sigma factor | 0,38 | 3,15 | 3,24 | 2,77 | 2,86 | 0,13 | 6,9E-01 | 3,4E-04 | 2,2E-04 | 9,4E-03 | 7,3E-03 | 4,7E-01 | 0,0E+00 | 7,0E-06 | 2,8E-06 | 1,5E-04 | 1,6E-04 | 0,0E+00 | 1752664 | 1753329 | 666 | + |
| Y11_RS08200 | Y11_17461 | tam | trans-aconitate 2-methyltransferase | -0,21 | 1,97 | 2,77 | 2,18 | 2,98 | 0,8 | 2,9E-01 | 6,1E-07 | 6,9E-07 | 7,6E-06 | 2,8E-06 | 7,7E-03 | 0,0E+00 | 1,9E-08 | 1,7E-08 | 1,6E-07 | 1,0E-07 | 0,0E+00 | 1753354 | 1754130 | 777 | + |
| Y11_RS08205 | Y11_17471 | dps | DNA starvation/stationary phase protection protein Dps | -0,69 | 1,54 | 1,94 | 2,24 | 2,63 | 0,4 | 1,5E-04 | 4,1E-08 | 1,5E-07 | 1,5E-08 | 3,6E-08 | 7,1E-03 | 0,0E+00 | 1,9E-09 | 4,6E-09 | 7,0E-10 | 2,7E-09 | 0,0E+00 | 1754445 | 1754948 | 504 | + |
| Y11_RS08215 | Y11_17501 | glnH | glutamine ABC transporter substrate-binding protein GlnH | -0,48 | 0,22 | 0,75 | 0,7 | 1,23 | 0,53 | 1,6E-04 | 3,7E-03 | 8,2E-06 | 1,1E-05 | 6,9E-07 | 1,2E-04 | 0,0E+00 | 0,0E+00 | 0,0E+00 | 0,0E+00 | 3,0E-08 | 0,0E+00 | 1755670 | 1756413 | 744 | + |
| Y11_RS08220 | Y11_17511 | glnP | glutamine ABC transporter permease GlnP | -0,14 | -0,45 | 0,03 | -0,31 | 0,18 | 0,48 | 4,9E-01 | 2,5E-02 | 8,8E-01 | 1,2E-01 | 4,6E-01 | 4,1E-02 | 0,0E+00 | 0,0E+00 | 0,0E+00 | 0,0E+00 | 0,0E+00 | 0,0E+00 | 1756480 | 1757136 | 657 | + |
| Y11_RS08225 | Y11_17521 | glnQ | glutamine ABC transporter ATP-binding protein GlnQ | -0,27 | -0,31 | 0,05 | -0,04 | 0,32 | 0,35 | 2,8E-03 | 1,3E-04 | 5,9E-01 | 5,9E-01 | 1,0E-02 | 2,9E-03 | 0,0E+00 | 0,0E+00 | 0,0E+00 | 0,0E+00 | 0,0E+00 | 0,0E+00 | 1757133 | 1757855 | 723 | + |
| Y11_RS08235 | Y11_17551 | rlmF | 23S rRNA (adenine(1618)-N(6))-methyltransferase RlmF | -0,99 | -0,6 | -0,53 | 0,38 | 0,45 | 0,07 | 9,7E-07 | 5,3E-05 | 8,7E-03 | 1,8E-04 | 1,5E-02 | 6,5E-01 | 0,0E+00 | 0,0E+00 | 0,0E+00 | 0,0E+00 | 0,0E+00 | 0,0E+00 | 1759962 | 1760924 | 963 | - |
| Y11_RS08240 | Y11_17561 | | flavin reductase family protein | 0,48 | 1 | 0,96 | 0,52 | 0,47 | -0,12 | 8,7E-02 | 5,4E-03 | 4,7E-03 | 1,3E-02 | 7,5E-03 | 5,6E-01 | 0,0E+00 | 1,0E-04 | 0,0E+00 | 0,0E+00 | 0,0E+00 | 0,0E+00 | 1761096 | 1761743 | 648 | + |
| Y11_RS08260 | Y11_17611 | ybiB | DNA-binding protein YbiB | 0,02 | 0,63 | 0,47 | 0,61 | 0,44 | -0,16 | 8,7E-01 | 8,5E-03 | 3,6E-03 | 9,0E-03 | 3,4E-03 | 3,4E-01 | 0,0E+00 | 0,0E+00 | 0,0E+00 | 0,0E+00 | 0,0E+00 | 0,0E+00 | 1764349 | 1765329 | 981 | - |
| Y11_RS08265 | Y11_17621 | aqpZ | aquaporin Z | -0,79 | 2,64 | 4,48 | 3,43 | 5,27 | 1,84 | 7,2E-05 | 5,2E-07 | 6,8E-10 | 6,0E-08 | 1,6E-10 | 3,5E-05 | 0,0E+00 | 1,6E-08 | 8,0E-11 | 2,2E-09 | 6,2E-11 | 3,3E-06 | 1765537 | 1766241 | 705 | - |
| Y11_RS08275 | Y11_17651 | ompF2 | porin OmpF2 | 0,19 | -0,96 | -1,92 | -1,15 | -2,12 | -0,96 | 1,4E-01 | 1,5E-04 | 2,8E-05 | 5,5E-05 | 1,6E-05 | 4,3E-03 | 0,0E+00 | 0,0E+00 | 4,4E-07 | 1,0E-06 | 4,9E-07 | 0,0E+00 | 1769217 | 1770332 | 1116 | + |
| Y11_RS08315 | Y11_17731 | | ABC transporter ATP-binding protein | -0,09 | -0,33 | 0,57 | -0,24 | 0,65 | 0,9 | 4,8E-01 | 1,2E-02 | 2,9E-04 | 9,9E-02 | 7,3E-04 | 3,8E-05 | 0,0E+00 | 0,0E+00 | 0,0E+00 | 0,0E+00 | 0,0E+00 | 0,0E+00 | 1776888 | 1777832 | 945 | - |
| Y11_RS08325 | Y11_17751 | | ABC transporter substrate-binding protein | 0,08 | 1,4 | 2,76 | 1,33 | 2,68 | 1,35 | 2,6E-01 | 2,6E-07 | 1,9E-10 | 3,6E-07 | 2,0E-10 | 4,9E-07 | 0,0E+00 | 9,2E-09 | 3,2E-11 | 1,0E-08 | 6,8E-11 | 1,1E-07 | 1779092 | 1780024 | 933 | - |
| Y11_RS08335 | Y11_17771 | | Gfo/Idh/MocA family oxidoreductase | -0,53 | -4,02 | -2,47 | -3,49 | -1,94 | 1,55 | 1,7E-05 | 2,7E-12 | 4,3E-10 | 1,3E-12 | 8,0E-10 | 6,1E-09 | 0,0E+00 | 3,2E-12 | 5,8E-11 | 1,9E-12 | 1,5E-10 | 4,9E-09 | 1781602 | 1782732 | 1131 | - |
| Y11_RS08340 | Y11_17781 | sseA | 3-mercaptopyruvate sulfurtransferase | 0,35 | -0,27 | 1,18 | -0,63 | 0 | 1,46 | 3,5E-01 | 3,2E-01 | 4,2E-04 | 1,2E-01 | 0,0E+00 | 1,5E-04 | 0,0E+00 | 0,0E+00 | 5,3E-06 | 0,0E+00 | 0,0E+00 | 1,1E-05 | 1783123 | 1783977 | 855 | + |
| Y11_RS08370 | Y11_17841 | | DUF1456 family protein | 0,03 | 0,1 | 0,73 | 0,07 | 0,7 | 0,63 | 6,5E-01 | 2,8E-01 | 1,4E-06 | 4,5E-01 | 4,0E-06 | 9,6E-05 | 0,0E+00 | 0,0E+00 | 0,0E+00 | 0,0E+00 | 0,0E+00 | 0,0E+00 | 1790000 | 1790470 | 471 | - |
| Y11_RS08385 | Y11_17871 | phnD | phosphonate ABC transporter substrate-binding protein | 0,87 | 1,2 | 5,59 | 0 | 0 | 0 | 3,1E-02 | 7,0E-03 | 6,3E-07 | 0,0E+00 | 0,0E+00 | 0,0E+00 | 0,0E+00 | 1,4E-04 | 1,5E-08 | 0,0E+00 | 0,0E+00 | 0,0E+00 | 1792558 | 1793487 | 930 | - |
| Y11_RS08420 | Y11_17941 | pbpG | D-alanyl-D-alanine endopeptidase | 0 | -2,07 | -1,69 | -15 | -15 | 0,38 | 0,0E+00 | 2,9E-06 | 2,4E-05 | 0,0E+00 | 0,0E+00 | 1,8E-01 | 0,0E+00 | 7,7E-08 | 3,8E-07 | 0,0E+00 | 0,0E+00 | 0,0E+00 | 1799645 | 1800580 | 936 | - |
| Y11_RS08435 | Y11_17971 | | hypothetical protein | 0,05 | -1,22 | -1,23 | -1,27 | -1,3 | -0,01 | 8,5E-01 | 2,3E-03 | 1,4E-03 | 8,2E-05 | 1,6E-05 | 9,5E-01 | 0,0E+00 | 4,4E-05 | 1,7E-05 | 1,5E-06 | 5,0E-07 | 0,0E+00 | 1802761 | 1803393 | 633 | + |
| Y11_RS08440 | Y11_17981 | dusC | tRNA dihydrouridine(16) synthase DusC | -0,15 | -0,12 | -0,72 | 0,02 | -0,57 | -0,6 | 4,7E-01 | 4,9E-01 | 1,1E-03 | 9,0E-01 | 6,7E-03 | 1,6E-03 | 0,0E+00 | 0,0E+00 | 0,0E+00 | 0,0E+00 | 0,0E+00 | 0,0E+00 | 1803479 | 1804411 | 933 | - |
| Y11_RS08445 | Y11_17991 | rhlE | ATP-dependent RNA helicase RhlE | -0,3 | -3,09 | -4,53 | -2,79 | -4,23 | -1,44 | 9,5E-04 | 9,4E-13 | 4,2E-10 | 5,5E-11 | 1,2E-09 | 3,6E-06 | 0,0E+00 | 1,6E-12 | 5,8E-11 | 1,2E-11 | 1,9E-10 | 4,9E-07 | 1804580 | 1805935 | 1356 | - |
| Y11_RS08450 | Y11_18001 | cecR | transcriptional regulator CecR | 0,34 | 0,62 | 0,45 | 0,28 | 0,12 | -0,16 | 1,2E-02 | 4,8E-04 | 2,4E-02 | 1,5E-02 | 4,6E-01 | 3,2E-01 | 0,0E+00 | 0,0E+00 | 0,0E+00 | 0,0E+00 | 0,0E+00 | 0,0E+00 | 1806338 | 1807048 | 711 | + |
| Y11_RS08455 | Y11_18011 | hlyD | secretion protein HlyD | 0,06 | 0,93 | 1,35 | 0,87 | 1,29 | 0,42 | 4,0E-01 | 6,6E-09 | 4,9E-06 | 7,4E-07 | 1,2E-05 | 9,3E-03 | 0,0E+00 | 0,0E+00 | 8,9E-08 | 0,0E+00 | 3,9E-07 | 0,0E+00 | 1807097 | 1808083 | 987 | + |
| Y11_RS08460 | Y11_18021 | | ATP-binding cassette domain-containing protein | 0,14 | 1,14 | 1,08 | 1 | 0,94 | -0,06 | 3,7E-01 | 2,8E-05 | 8,8E-05 | 6,2E-06 | 3,9E-05 | 5,6E-01 | 0,0E+00 | 6,4E-07 | 1,2E-06 | 1,3E-07 | 0,0E+00 | 0,0E+00 | 1808097 | 1809848 | 1752 | + |
| Y11_RS08465 | Y11_18031 | | ABC transporter permease | 0,11 | 1,91 | 1,99 | 1,8 | 1,88 | 0,08 | 4,9E-01 | 2,2E-04 | 4,8E-07 | 3,0E-04 | 5,4E-07 | 7,8E-01 | 0,0E+00 | 4,6E-06 | 1,2E-08 | 5,3E-06 | 2,5E-08 | 0,0E+00 | 1809845 | 1811035 | 1191 | + |
| Y11_RS08470 | Y11_18041 | | ABC transporter permease | 0,31 | 1,12 | 1,42 | 0,81 | 1,11 | 0,3 | 2,4E-02 | 1,8E-05 | 1,1E-05 | 2,5E-05 | 1,8E-05 | 5,8E-02 | 0,0E+00 | 4,3E-07 | 1,9E-07 | 0,0E+00 | 5,4E-07 | 0,0E+00 | 1811069 | 1812175 | 1107 | + |
| Y11_RS08475 | Y11_18051 | | Bax inhibitor-1/YccA family protein | -0,14 | 0,06 | 0,73 | 0,2 | 0,87 | 0,67 | 4,4E-01 | 6,9E-01 | 6,0E-03 | 2,9E-01 | 4,2E-03 | 1,0E-02 | 0,0E+00 | 0,0E+00 | 0,0E+00 | 0,0E+00 | 0,0E+00 | 0,0E+00 | 1812438 | 1813148 | 711 | - |
| Y11_RS08480 | Y11_18061 | moaE | molybdopterin synthase catalytic subunit MoaE | 0,67 | 1,33 | 0,76 | 0,66 | 0,09 | -0,57 | 1,3E-04 | 2,4E-08 | 1,8E-03 | 1,5E-04 | 6,4E-01 | 8,6E-03 | 0,0E+00 | 1,3E-09 | 0,0E+00 | 0,0E+00 | 0,0E+00 | 0,0E+00 | 1813292 | 1813744 | 453 | - |
| Y11_RS08485 | Y11_18071 | moaD | molybdopterin synthase sulfur carrier subunit | 0,94 | 2,23 | 2,67 | 1,29 | 1,78 | 0,43 | 3,8E-01 | 6,2E-02 | 1,7E-02 | 1,3E-01 | 1,1E-02 | 4,9E-01 | 0,0E+00 | 1,2E-03 | 1,9E-04 | 2,0E-03 | 2,4E-04 | 0,0E+00 | 1813747 | 1813992 | 246 | - |
| Y11_RS08490 | Y11_18081 | moaC | cyclic pyranopterin monophosphate synthase MoaC | 0,7 | 0,93 | 0,75 | 0,23 | 0,04 | -0,18 | 5,0E-04 | 1,1E-04 | 2,2E-04 | 7,8E-02 | 6,5E-01 | 1,2E-01 | 0,0E+00 | 0,0E+00 | 0,0E+00 | 0,0E+00 | 0,0E+00 | 0,0E+00 | 1813989 | 1814468 | 480 | - |
| Y11_RS08495 | Y11_18091 | moaA | GTP 3',8-cyclase MoaA | 0,68 | 1,31 | 1,24 | 0,63 | 0,55 | -0,08 | 2,5E-01 | 5,7E-02 | 7,6E-02 | 3,9E-02 | 9,7E-02 | 8,4E-01 | 0,0E+00 | 1,1E-03 | 8,5E-04 | 0,0E+00 | 0,0E+00 | 0,0E+00 | 1814500 | 1815480 | 981 | - |
| Y11_RS08500 | Y11_18101 | yvcK | uridine diphosphate-N-acetylglucosamine-binding protein YvcK | 0,25 | 0,5 | 0,61 | 0,26 | 0,37 | 0,11 | 3,8E-01 | 4,6E-02 | 3,4E-02 | 3,1E-01 | 2,0E-01 | 6,1E-01 | 0,0E+00 | 0,0E+00 | 0,0E+00 | 0,0E+00 | 0,0E+00 | 0,0E+00 | 1815933 | 1816856 | 924 | + |
| Y11_RS08505 | Y11_18111 | | VF530 family protein | 0,13 | 0,99 | 0,97 | 0,86 | 0,83 | -0,02 | 5,4E-01 | 5,7E-05 | 1,7E-05 | 4,4E-03 | 3,7E-03 | 8,6E-01 | 0,0E+00 | 0,0E+00 | 0,0E+00 | 0,0E+00 | 0,0E+00 | 0,0E+00 | 1816871 | 1817119 | 249 | - |
| Y11_RS08510 | Y11_18121 | uvrB | excinuclease ABC subunit B | 0,42 | 0,73 | 1,04 | 0,32 | 0,62 | 0,3 | 8,0E-05 | 1,0E-06 | 7,7E-07 | 6,7E-04 | 4,7E-05 | 4,4E-03 | 0,0E+00 | 0,0E+00 | 1,8E-08 | 0,0E+00 | 0,0E+00 | 0,0E+00 | 1817188 | 1819200 | 2013 | - |
| Y11_RS08520 | Y11_18151 | bioD | dethiobiotin synthase | 0,01 | -0,38 | -1,08 | -0,4 | -1,1 | -0,7 | 8,5E-01 | 6,8E-03 | 1,2E-05 | 6,4E-03 | 1,3E-05 | 1,1E-03 | 0,0E+00 | 0,0E+00 | 2,0E-07 | 0,0E+00 | 4,2E-07 | 0,0E+00 | 1821172 | 1821885 | 714 | - |
| Y11_RS08545 | Y11_18201 | pgl | 6-phosphogluconolactonase | 0,13 | 1,29 | 1,53 | 1,16 | 1,4 | 0,24 | 3,1E-02 | 2,6E-10 | 2,2E-06 | 1,8E-08 | 6,1E-06 | 1,0E-01 | 0,0E+00 | 4,1E-11 | 4,6E-08 | 8,2E-10 | 2,1E-07 | 0,0E+00 | 1826372 | 1827373 | 1002 | - |
| Y11_RS08550 | Y11_18211 | | pyridoxal phosphatase | 0,49 | 0,65 | -0,22 | 0,17 | -0,71 | -0,88 | 4,3E-05 | 2,3E-05 | 3,9E-02 | 1,3E-02 | 1,1E-05 | 7,0E-06 | 0,0E+00 | 0,0E+00 | 0,0E+00 | 0,0E+00 | 0,0E+00 | 0,0E+00 | 1827787 | 1828608 | 822 | + |
| Y11_RS08555 | Y11_18221 | modC | molybdenum ABC transporter ATP-binding protein ModC | -0,18 | 0,33 | -0,42 | 0,51 | -0,24 | -0,75 | 4,3E-01 | 1,2E-01 | 1,5E-01 | 1,5E-02 | 3,6E-01 | 9,9E-03 | 0,0E+00 | 0,0E+00 | 0,0E+00 | 0,0E+00 | 0,0E+00 | 0,0E+00 | 1828851 | 1829915 | 1065 | - |
| Y11_RS08575 | Y11_18271 | modE | molybdenum-dependent transcriptional regulator | 0,19 | 0,67 | 0,63 | 0,48 | 0,44 | -0,04 | 4,2E-02 | 6,5E-05 | 2,2E-04 | 2,4E-04 | 1,3E-03 | 7,2E-01 | 0,0E+00 | 0,0E+00 | 0,0E+00 | 0,0E+00 | 0,0E+00 | 0,0E+00 | 1831971 | 1832762 | 792 | + |
| Y11_RS08580 | Y11_18291 | modF | molybdate ABC transporter ATP-binding protein ModF | -0,6 | -1,26 | -1,45 | -0,66 | -0,85 | -0,19 | 4,7E-07 | 2,3E-08 | 3,7E-06 | 1,0E-06 | 1,4E-04 | 1,9E-01 | 0,0E+00 | 1,2E-09 | 7,0E-08 | 0,0E+00 | 0,0E+00 | 0,0E+00 | 1832871 | 1834364 | 1494 | + |
| Y11_RS08590 | Y11_18311 | galE | UDP-glucose 4-epimerase GalE | -5,07 | -0,01 | 0,4 | 5,07 | 5,47 | 0,41 | 1,9E-11 | 9,5E-01 | 8,4E-03 | 4,8E-11 | 7,5E-11 | 1,2E-02 | 9,7E-12 | 0,0E+00 | 0,0E+00 | 1,2E-11 | 3,9E-11 | 0,0E+00 | 1835676 | 1836692 | 1017 | + |
| Y11_RS08595 | Y11_18321 | galT | galactose-1-phosphate uridylyltransferase | -4,76 | 0,22 | 0,34 | 4,98 | 5,1 | 0,12 | 9,6E-13 | 2,6E-02 | 1,2E-02 | 3,3E-12 | 2,2E-11 | 3,3E-01 | 1,8E-12 | 0,0E+00 | 0,0E+00 | 3,9E-12 | 2,0E-11 | 0,0E+00 | 1836702 | 1837790 | 1089 | + |
| Y11_RS08600 | Y11_18331 | galK | galactokinase | -5,56 | 0,02 | 0,36 | 5,58 | 5,92 | 0,34 | 2,7E-13 | 8,0E-01 | 5,1E-05 | 1,2E-12 | 5,7E-14 | 6,3E-04 | 1,8E-12 | 0,0E+00 | 0,0E+00 | 1,9E-12 | 5,5E-13 | 0,0E+00 | 1837790 | 1838941 | 1152 | + |
| Y11_RS08605 | Y11_18341 | galM | galactose-1-epimerase | -4,2 | 0,18 | 0,28 | 4,38 | 4,48 | 0,1 | 1,5E-08 | 4,4E-01 | 1,9E-01 | 2,0E-08 | 5,2E-09 | 6,4E-01 | 1,8E-09 | 0,0E+00 | 0,0E+00 | 8,9E-10 | 5,7E-10 | 0,0E+00 | 1838935 | 1839999 | 1065 | + |
| Y11_RS08610 | Y11_18351 | | PsiF family protein | 2,01 | 3,4 | 3,93 | 1,39 | 1,92 | 0,53 | 6,0E-07 | 3,5E-09 | 2,8E-09 | 2,8E-05 | 4,4E-06 | 1,1E-02 | 4,1E-08 | 2,7E-10 | 2,1E-10 | 5,5E-07 | 1,5E-07 | 0,0E+00 | 1840174 | 1840509 | 336 | + |
| Y11_RS08620 | Y11_18381 | | DNA/RNA non-specific endonuclease | -2,85 | -2,99 | -1,2 | -0,14 | 1,64 | 1,78 | 1,2E-10 | 6,5E-10 | 1,5E-04 | 2,0E-01 | 2,1E-05 | 1,6E-05 | 3,6E-11 | 7,9E-11 | 2,0E-06 | 0,0E+00 | 6,3E-07 | 1,7E-06 | 1843890 | 1844741 | 852 | + |
| Y11_RS08625 | Y11_18401 | gpmA | 2,3-diphosphoglycerate-dependent phosphoglycerate mutase | 0,16 | 1,07 | 0,97 | 0,91 | 0,82 | -0,1 | 5,9E-02 | 2,3E-07 | 1,4E-06 | 5,9E-06 | 2,6E-05 | 3,3E-01 | 0,0E+00 | 8,5E-09 | 0,0E+00 | 0,0E+00 | 0,0E+00 | 0,0E+00 | 1844948 | 1845700 | 753 | + |
| Y11_RS08630 | Y11_18411 | aroG | 3-deoxy-7-phosphoheptulonate synthase AroG | 0,36 | 0,55 | 0,08 | 0,19 | -0,27 | -0,46 | 7,8E-04 | 2,3E-05 | 4,2E-01 | 1,5E-02 | 2,2E-02 | 1,3E-03 | 0,0E+00 | 0,0E+00 | 0,0E+00 | 0,0E+00 | 0,0E+00 | 0,0E+00 | 1845942 | 1846994 | 1053 | - |
| Y11_RS08635 | Y11_18421 | zitB | CDF family zinc transporter ZitB | 0,2 | 0,02 | 1,14 | -0,18 | 0,95 | 0 | 2,1E-01 | 8,8E-01 | 7,5E-06 | 3,3E-01 | 4,5E-05 | 0,0E+00 | 0,0E+00 | 0,0E+00 | 1,3E-07 | 0,0E+00 | 0,0E+00 | 0,0E+00 | 1847240 | 1848172 | 933 | + |
| Y11_RS08645 | Y11_18441 | nadA | quinolinate synthase NadA | 0,02 | 0,24 | -0,38 | 0,22 | -0,39 | -0,62 | 9,4E-01 | 3,8E-01 | 9,3E-02 | 3,0E-01 | 8,9E-03 | 1,3E-02 | 0,0E+00 | 0,0E+00 | 0,0E+00 | 0,0E+00 | 0,0E+00 | 0,0E+00 | 1849232 | 1850293 | 1062 | - |

| | | | | | | | | | | | | | | | | | | | | | |
|---|---|---|---|---|---|---|---|---|---|---|---|---|---|---|---|---|---|---|---|---|---|
| Y11_RS08655 | Y11_18451 | cpoB | cell division protein CpoB | 0,65 | 0,57 | 0,53 | -0,08 | -0,12 | -0,05 | 2,3E-06 | 8,8E-05 | 2,9E-02 | 3,4E-01 | 5,5E-01 | 8,3E-01 | 0,0E+00 | 0,0E+00 | 0,0E+00 | 0,0E+00 | 0,0E+00 | 0,0E+00 | 1850978 | 1851787 | 810 | - |
| Y11_RS08660 | Y11_18461 | pal | peptidoglycan-associated lipoprotein Pal | 0,01 | -0,52 | -0,45 | -0,52 | -0,46 | 0,06 | 9,3E-01 | 1,3E-05 | 7,1E-03 | 5,5E-05 | 8,4E-03 | 6,2E-01 | 0,0E+00 | 0,0E+00 | 0,0E+00 | 0,0E+00 | 0,0E+00 | 0,0E+00 | 1851797 | 1852303 | 507 | - |
| Y11_RS08665 | Y11_18471 | tolB | Tol-Pal system beta propeller repeat protein TolB | -0,12 | -0,42 | -0,13 | -0,3 | 0 | 0,3 | 9,6E-03 | 1,9E-06 | 7,8E-02 | 8,9E-07 | 9,6E-01 | 6,7E-04 | 0,0E+00 | 0,0E+00 | 0,0E+00 | 0,0E+00 | 0,0E+00 | 0,0E+00 | 1852352 | 1853644 | 1293 | - |
| Y11_RS08670 | Y11_18481 | tolA | cell envelope integrity protein TolA | 0,35 | 0,18 | 0,37 | -0,17 | 0,02 | 0,18 | 1,4E-02 | 6,5E-02 | 6,1E-02 | 8,3E-02 | 9,1E-01 | 2,6E-01 | 0,0E+00 | 0,0E+00 | 0,0E+00 | 0,0E+00 | 0,0E+00 | 0,0E+00 | 1853764 | 1854933 | 1170 | - |
| Y11_RS08675 | Y11_18491 | tolR | colicin uptake protein TolR | 0,26 | -0,31 | -0,52 | -0,58 | -0,78 | -0,21 | 1,6E-02 | 9,9E-03 | 7,7E-03 | 3,3E-04 | 7,8E-04 | 2,2E-01 | 0,0E+00 | 0,0E+00 | 0,0E+00 | 0,0E+00 | 0,0E+00 | 0,0E+00 | 1855045 | 1855473 | 429 | - |
| Y11_RS08680 | Y11_18501 | tolQ | Tol-Pal system protein TolQ | 0,26 | -0,11 | 0,17 | -0,37 | -0,09 | 0,28 | 7,1E-03 | 9,3E-02 | 8,1E-02 | 7,8E-04 | 3,9E-01 | 1,1E-02 | 0,0E+00 | 0,0E+00 | 0,0E+00 | 0,0E+00 | 0,0E+00 | 0,0E+00 | 1855486 | 1856172 | 687 | - |
| Y11_RS08685 | Y11_18511 | ybgC | tol-pal system-associated acyl-CoA thioesterase | -0,03 | 0,17 | 0,19 | 0,2 | 0,22 | 0,02 | 8,0E-01 | 1,4E-01 | 1,1E-01 | 8,3E-02 | 6,3E-02 | 8,0E-01 | 0,0E+00 | 0,0E+00 | 0,0E+00 | 0,0E+00 | 0,0E+00 | 0,0E+00 | 1856172 | 1856573 | 402 | - |
| Y11_RS08700 | Y11_18541 | cydB | cytochrome d ubiquinol oxidase subunit II | 0,08 | 0,39 | 15 | 0,31 | 15 | 15 | 8,6E-01 | 3,6E-01 | 0,0E+00 | 3,4E-01 | 0,0E+00 | 0,0E+00 | 0,0E+00 | 0,0E+00 | 0,0E+00 | 0,0E+00 | 0,0E+00 | 0,0E+00 | 1857125 | 1858264 | 1140 | - |
| Y11_RS08705 | Y11_18551 | cydA | cytochrome ubiquinol oxidase subunit I | 0,46 | 0,08 | 2,7 | -0,38 | 2,24 | 2,62 | 5,8E-03 | 5,3E-01 | 2,7E-08 | 4,6E-04 | 2,9E-09 | 4,9E-10 | 0,0E+00 | 0,0E+00 | 1,1E-09 | 0,0E+00 | 3,7E-10 | 1,1E-09 | 1858279 | 1859847 | 1569 | - |
| Y11_RS08715 | Y11_18571 | sucD | succinate--CoA ligase subunit alpha | 0,32 | -0,15 | 0,66 | -0,47 | 0,35 | 0,82 | 1,4E-03 | 2,9E-02 | 1,5E-05 | 9,8E-05 | 2,2E-03 | 3,0E-06 | 0,0E+00 | 0,0E+00 | 0,0E+00 | 0,0E+00 | 0,0E+00 | 0,0E+00 | 1862566 | 1863444 | 879 | - |
| Y11_RS08720 | Y11_18581 | sucC | ADP-forming succinate--CoA ligase subunit beta | 0,33 | -0,11 | 0,6 | -0,43 | 0,27 | 0,71 | 8,4E-04 | 5,3E-02 | 3,7E-04 | 6,8E-05 | 3,4E-02 | 9,7E-05 | 0,0E+00 | 0,0E+00 | 0,0E+00 | 0,0E+00 | 0,0E+00 | 0,0E+00 | 1863444 | 1864610 | 1167 | - |
| Y11_RS08725 | Y11_18591 | odhB | 2-oxoglutarate dehydrogenase complex dihydrolipoyllysine-residue succinyltransferase | 0,38 | -0,23 | 0,46 | -0,6 | 0,08 | 0,69 | 2,1E-03 | 3,0E-02 | 3,4E-04 | 1,6E-04 | 3,5E-01 | 3,8E-05 | 0,0E+00 | 0,0E+00 | 0,0E+00 | 0,0E+00 | 0,0E+00 | 0,0E+00 | 1864724 | 1865947 | 1224 | - |
| Y11_RS08730 | Y11_18601 | sucA | 2-oxoglutarate dehydrogenase E1 component | 0,5 | -0,05 | 0,77 | -0,55 | 0,26 | 0,82 | 3,5E-07 | 4,0E-01 | 1,8E-07 | 6,0E-06 | 3,9E-04 | 1,0E-06 | 0,0E+00 | 0,0E+00 | 0,0E+00 | 0,0E+00 | 0,0E+00 | 0,0E+00 | 1865976 | 1868783 | 2808 | - |
| Y11_RS08735 | Y11_18611 | | succinate dehydrogenase iron-sulfur subunit | 0,54 | 0,58 | 1,37 | 0,04 | 0,83 | 0,79 | 7,5E-06 | 1,5E-05 | 1,4E-07 | 5,6E-01 | 7,9E-06 | 1,9E-05 | 0,0E+00 | 0,0E+00 | 4,3E-09 | 0,0E+00 | 0,0E+00 | 0,0E+00 | 1869057 | 1869773 | 717 | - |
| Y11_RS08740 | Y11_18621 | sdhA | succinate dehydrogenase flavoprotein subunit | 0,36 | 0,43 | 1,1 | 0,07 | 0,73 | 0,67 | 1,4E-04 | 5,1E-05 | 2,8E-06 | 3,2E-01 | 7,7E-05 | 1,6E-04 | 0,0E+00 | 0,0E+00 | 5,5E-08 | 0,0E+00 | 0,0E+00 | 0,0E+00 | 1869827 | 1871593 | 1767 | - |
| Y11_RS08745 | Y11_18631 | sdhD | succinate dehydrogenase membrane anchor subunit | 0,66 | 0,88 | 1,33 | 0,23 | 0,67 | 0,45 | 3,2E-03 | 2,8E-04 | 2,2E-04 | 8,7E-02 | 7,9E-03 | 3,8E-02 | 0,0E+00 | 0,0E+00 | 2,9E-06 | 0,0E+00 | 0,0E+00 | 0,0E+00 | 1871594 | 1871941 | 348 | - |
| Y11_RS08750 | Y11_18641 | sdhC | succinate dehydrogenase cytochrome b556 subunit | 0,02 | 0,76 | 15 | 0,75 | 15 | 0 | 9,7E-01 | 3,1E-02 | 0,0E+00 | 8,4E-03 | 0,0E+00 | 0,0E+00 | 0,0E+00 | 0,0E+00 | 0,0E+00 | 0,0E+00 | 0,0E+00 | 0,0E+00 | 1871935 | 1872324 | 390 | - |
| Y11_RS08755 | Y11_18651 | | citrate synthase | 0,17 | -0,9 | -0,34 | -1,07 | -0,51 | 0,56 | 2,8E-02 | 3,1E-06 | 2,1E-04 | 4,5E-06 | 2,0E-04 | 3,0E-04 | 0,0E+00 | 0,0E+00 | 0,0E+00 | 9,9E-08 | 0,0E+00 | 0,0E+00 | 1872973 | 1874253 | 1281 | + |
| Y11_RS08760 | Y11_18661 | pcp | pyroglutamyl-peptidase I | -0,41 | -2,74 | -3,27 | -2,33 | -2,86 | -0,54 | 1,7E-03 | 4,5E-10 | 4,9E-11 | 2,5E-09 | 2,4E-10 | 2,6E-05 | 0,0E+00 | 6,0E-11 | 1,5E-11 | 1,7E-10 | 7,4E-11 | 0,0E+00 | 1874319 | 1874966 | 648 | - |
| Y11_RS08765 | Y11_18671 | | DUF979 domain-containing protein | 0 | -15 | -15 | -15 | -15 | 0,26 | 0,0E+00 | 0,0E+00 | 0,0E+00 | 0,0E+00 | 0,0E+00 | 7,2E-01 | 0,0E+00 | 0,0E+00 | 0,0E+00 | 0,0E+00 | 0,0E+00 | 0,0E+00 | 1874975 | 1875994 | 1020 | - |
| Y11_RS08770 | Y11_18681 | | DUF969 domain-containing protein | -0,22 | -3,92 | -3,84 | -3,7 | -3,62 | 0,08 | 1,7E-01 | 7,1E-09 | 2,2E-04 | 1,4E-10 | 2,8E-04 | 9,0E-01 | 0,0E+00 | 4,9E-10 | 2,9E-06 | 2,1E-11 | 6,9E-06 | 0,0E+00 | 1875991 | 1876722 | 732 | - |
| Y11_RS08775 | Y11_18691 | pxpA | 5-oxoprolinase subunit PxpA | -0,12 | -3,2 | -3,11 | -3,07 | -2,99 | 0,08 | 3,5E-01 | 3,0E-09 | 4,6E-08 | 4,1E-10 | 2,2E-08 | 5,3E-01 | 0,0E+00 | 2,5E-10 | 1,8E-09 | 4,6E-11 | 1,8E-09 | 0,0E+00 | 1876813 | 1877550 | 738 | - |
| Y11_RS08780 | Y11_18701 | | biotin-dependent carboxyltransferase family protein | -0,13 | -0,69 | -1,7 | -0,56 | -1,57 | -1,01 | 1,4E-01 | 1,3E-04 | 4,4E-07 | 1,4E-03 | 2,0E-06 | 1,1E-04 | 0,0E+00 | 0,0E+00 | 1,1E-08 | 0,0E+00 | 7,8E-08 | 8,8E-06 | 1877547 | 1878512 | 966 | - |
| Y11_RS08790 | Y11_18721 | | type 2 GTP cyclohydrolase I | 0,04 | 0,31 | 0,82 | 0,27 | 0,78 | 0,51 | 6,1E-01 | 3,7E-02 | 2,6E-04 | 3,4E-02 | 1,5E-04 | 1,1E-02 | 0,0E+00 | 0,0E+00 | 0,0E+00 | 0,0E+00 | 0,0E+00 | 0,0E+00 | 1879581 | 1880324 | 744 | - |
| Y11_RS08795 | Y11_18731 | | MBL fold metallo-hydrolase | 0,17 | 0,61 | 1,13 | 0,44 | 0,96 | 0,51 | 1,0E-03 | 1,7E-06 | 1,5E-08 | 1,1E-05 | 2,9E-08 | 2,2E-05 | 0,0E+00 | 0,0E+00 | 7,0E-10 | 0,0E+00 | 0,0E+00 | 0,0E+00 | 1880500 | 1881555 | 1056 | - |
| Y11_RS08800 | Y11_18741 | phrB | deoxyribodipyrimidine photo-lyase | 0,63 | 0,34 | 0,56 | -0,29 | -0,07 | 0,22 | 3,2E-05 | 2,7E-04 | 4,2E-08 | 1,1E-02 | 3,9E-01 | 3,3E-03 | 0,0E+00 | 0,0E+00 | 0,0E+00 | 0,0E+00 | 0,0E+00 | 0,0E+00 | 1881571 | 1883016 | 1446 | - |
| Y11_RS08805 | Y11_18751 | | YbgA family protein | 0,19 | 0,05 | 0,7 | -0,14 | 0,51 | 0,65 | 5,3E-01 | 8,7E-01 | 5,4E-02 | 4,9E-01 | 5,1E-02 | 2,1E-02 | 0,0E+00 | 0,0E+00 | 0,0E+00 | 0,0E+00 | 0,0E+00 | 0,0E+00 | 1883020 | 1883496 | 477 | - |
| Y11_RS08810 | Y11_18761 | rtxA | MARTX multifunctional-autoprocessing repeats-in-toxin holotoxin RtxA | 0,24 | 0,59 | 0,3 | 0,35 | 0,05 | -0,29 | 7,4E-01 | 3,7E-01 | 6,9E-01 | 6,0E-01 | 9,4E-01 | 6,6E-01 | 0,0E+00 | 0,0E+00 | 0,0E+00 | 0,0E+00 | 0,0E+00 | 0,0E+00 | 1883643 | 1893380 | 9738 | - |
| Y11_RS08840 | Y11_18811 | kdpB | potassium-transporting ATPase subunit KdpB | -1,89 | 0,37 | 0,72 | 2,26 | 2,61 | 0,36 | 2,7E-07 | 1,6E-02 | 5,0E-04 | 3,1E-07 | 1,4E-07 | 4,9E-02 | 2,1E-08 | 0,0E+00 | 0,0E+00 | 8,7E-09 | 8,3E-09 | 0,0E+00 | 1897185 | 1899251 | 2067 | + |
| Y11_RS08845 | Y11_18821 | kdpC | potassium-transporting ATPase subunit KdpC | -0,84 | 0,94 | 0,7 | 1,79 | 1,53 | 0 | 1,6E-01 | 5,5E-04 | 1,5E-03 | 1,1E-02 | 2,2E-02 | 0,0E+00 | 0,0E+00 | 0,0E+00 | 0,0E+00 | 1,8E-04 | 4,7E-04 | 0,0E+00 | 1899369 | 1899971 | 603 | + |
| Y11_RS08850 | Y11_18831 | kdpD | two-component system sensor histidine kinase KdpD | -0,57 | 0,59 | 0,27 | 1,16 | 0,84 | -0,32 | 1,3E-01 | 3,9E-02 | 2,3E-01 | 9,3E-03 | 3,1E-02 | 1,8E-01 | 0,0E+00 | 0,0E+00 | 0,0E+00 | 1,5E-04 | 0,0E+00 | 0,0E+00 | 1899981 | 1902680 | 2700 | + |
| Y11_RS08855 | Y11_18841 | kdpE | two-component system response regulator KdpE | -0,74 | 1,05 | 0,97 | 1,79 | 1,71 | -0,08 | 1,1E-01 | 3,3E-03 | 6,2E-03 | 6,3E-04 | 9,7E-04 | 3,6E-01 | 0,0E+00 | 6,4E-05 | 0,0E+00 | 1,1E-05 | 2,2E-05 | 0,0E+00 | 1902677 | 1903366 | 690 | + |
| Y11_RS08870 | Y11_18871 | pgm | phosphoglucomutase (alpha-D-glucose-1,6-bisphosphate-dependent) | -0,5 | 0,76 | 1,15 | 1,26 | 1,65 | 0,39 | 3,6E-06 | 5,0E-09 | 5,9E-07 | 8,0E-10 | 5,0E-08 | 1,0E-03 | 0,0E+00 | 0,0E+00 | 1,5E-08 | 7,4E-11 | 3,5E-09 | 0,0E+00 | 1905739 | 1907379 | 1641 | - |
| Y11_RS08875 | Y11_18881 | seqA | replication initiation negative regulator SeqA | -0,17 | 0,35 | 0,03 | 0,52 | 0,2 | -0,32 | 1,1E-01 | 4,5E-03 | 8,1E-01 | 2,1E-03 | 1,9E-01 | 4,3E-02 | 0,0E+00 | 0,0E+00 | 0,0E+00 | 0,0E+00 | 0,0E+00 | 0,0E+00 | 1907472 | 1908011 | 540 | - |
| Y11_RS08880 | Y11_18891 | ybfF | esterase | -0,23 | 0,34 | 0,07 | 0,57 | 0,3 | -0,27 | 2,9E-02 | 9,0E-03 | 5,9E-01 | 1,2E-03 | 6,1E-02 | 1,1E-01 | 0,0E+00 | 0,0E+00 | 0,0E+00 | 0,0E+00 | 0,0E+00 | 0,0E+00 | 1908225 | 1909019 | 795 | + |
| Y11_RS08885 | Y11_18901 | ybfE | LexA regulated protein | 0 | 0 | -15 | 0 | -0,49 | -2,76 | 0,0E+00 | 0,0E+00 | 0,0E+00 | 0,0E+00 | 3,7E-01 | 3,8E-04 | 0,0E+00 | 0,0E+00 | 0,0E+00 | 0,0E+00 | 0,0E+00 | 2,7E-05 | 1909363 | 1909650 | 288 | + |
| Y11_RS08895 | Y11_18921 | fldA | flavodoxin FldA | -0,36 | 0,22 | 0,16 | 0,58 | 0,52 | -0,06 | 9,9E-03 | 1,0E-01 | 1,8E-01 | 8,5E-04 | 9,9E-04 | 6,2E-01 | 0,0E+00 | 0,0E+00 | 0,0E+00 | 0,0E+00 | 0,0E+00 | 0,0E+00 | 1911598 | 1912125 | 528 | + |
| Y11_RS08900 | Y11_18931 | fur | ferric iron uptake transcriptional regulator | -0,18 | 0,43 | 1,37 | 0,61 | 1,55 | 0,94 | 1,8E-01 | 1,0E-02 | 1,3E-04 | 3,6E-05 | 1,7E-05 | 6,3E-04 | 0,0E+00 | 0,0E+00 | 1,8E-06 | 0,0E+00 | 5,3E-07 | 0,0E+00 | 1912494 | 1912940 | 447 | + |
| Y11_RS08905 | Y11_18941 | | carbohydate-binding domain-containing protein | -0,37 | -0,18 | 0,44 | 0,19 | 0,81 | 0,61 | 2,6E-05 | 2,8E-03 | 3,0E-03 | 4,0E-04 | 4,5E-05 | 2,9E-04 | 0,0E+00 | 0,0E+00 | 0,0E+00 | 0,0E+00 | 0,0E+00 | 0,0E+00 | 1913337 | 1916012 | 2676 | - |
| Y11_RS08910 | Y11_18951 | chiQ | ChiQ/YbfN family lipoprotein | 0 | 1,09 | 1,41 | 1,09 | 1,41 | 0,33 | 9,9E-01 | 1,1E-03 | 9,0E-05 | 8,0E-04 | 6,1E-05 | 1,2E-01 | 0,0E+00 | 2,1E-05 | 1,2E-06 | 1,3E-05 | 1,7E-06 | 0,0E+00 | 1916120 | 1916446 | 327 | - |
| Y11_RS08920 | Y11_18971 | glnS | glutamine--tRNA ligase | 0,36 | -1,3 | -1,32 | -1,65 | -1,67 | -0,02 | 6,0E-04 | 3,8E-10 | 2,4E-09 | 3,9E-09 | 7,7E-09 | 6,1E-01 | 0,0E+00 | 5,4E-11 | 1,9E-10 | 2,3E-10 | 7,9E-10 | 0,0E+00 | 1918548 | 1920215 | 1668 | - |
| Y11_RS08925 | Y11_18981 | nagE | N-acetylglucosamine-specific PTS transporter subunit IIBC | -0,21 | 2,45 | 2,77 | 2,67 | 2,98 | 0,32 | 2,9E-03 | 6,8E-12 | 2,7E-10 | 4,7E-11 | 4,1E-10 | 3,7E-03 | 0,0E+00 | 5,1E-12 | 4,2E-11 | 1,2E-11 | 1,0E-10 | 0,0E+00 | 1920408 | 1922441 | 2034 | - |
| Y11_RS08930 | Y11_18991 | nagB | glucosamine-6-phosphate deaminase | -0,35 | 1,79 | 2,35 | 2,15 | 2,7 | 0,55 | 2,0E-02 | 4,8E-09 | 5,5E-07 | 9,0E-08 | 5,9E-07 | 9,1E-03 | 0,0E+00 | 3,7E-10 | 1,4E-08 | 3,1E-09 | 2,7E-08 | 0,0E+00 | 1922794 | 1923594 | 801 | + |
| Y11_RS08935 | Y11_19001 | nagA | N-acetylglucosamine-6-phosphate deacetylase | -0,55 | 1,3 | 2,12 | 1,86 | 2,67 | 0,82 | 2,4E-06 | 1,9E-09 | 1,4E-08 | 3,6E-10 | 2,9E-09 | 2,5E-05 | 0,0E+00 | 1,7E-10 | 6,8E-10 | 4,0E-11 | 3,7E-10 | 0,0E+00 | 1923614 | 1924759 | 1146 | + |
| Y11_RS08940 | Y11_19011 | | ROK family transcriptional regulator | 0,32 | 1,08 | 1,13 | 0,76 | 0,8 | 0,05 | 1,4E-03 | 1,0E-08 | 1,9E-09 | 7,2E-06 | 3,0E-06 | 3,6E-01 | 0,0E+00 | 6,4E-10 | 1,6E-10 | 0,0E+00 | 0,0E+00 | 0,0E+00 | 1924776 | 1925996 | 1221 | + |
| Y11_RS08945 | Y11_19031 | | HAD-IIA family hydrolase | -0,22 | 0,75 | 0,56 | 0,97 | 0,78 | -0,19 | 2,3E-02 | 4,8E-05 | 3,3E-03 | 8,6E-07 | 2,0E-04 | 2,0E-01 | 0,0E+00 | 0,0E+00 | 0,0E+00 | 0,0E+00 | 0,0E+00 | 0,0E+00 | 1926207 | 1926959 | 753 | + |
| Y11_RS08950 | Y11_19041 | asnB | asparagine synthase B | 0 | -2,37 | -2,34 | -2,37 | -2,34 | 0,02 | 9,9E-01 | 3,3E-09 | 1,8E-06 | 5,4E-10 | 1,4E-06 | 9,1E-01 | 0,0E+00 | 2,7E-10 | 3,8E-08 | 5,5E-11 | 5,5E-08 | 0,0E+00 | 1927428 | 1929092 | 1665 | + |
| Y11_RS08985 | Y11_19051 | ubiF | 2-octaprenyl-3-methyl-6-methoxy-1,4-benzoquinol hydroxylase | -0,19 | 0,4 | 0,52 | 0,6 | 0,72 | 0,12 | 2,6E-02 | 9,1E-05 | 2,8E-04 | 4,0E-06 | 2,7E-05 | 1,4E-01 | 0,0E+00 | 0,0E+00 | 0,0E+00 | 0,0E+00 | 0,0E+00 | 0,0E+00 | 1930154 | 1931335 | 1182 | - |
| Y11_RS08990 | Y11_19061 | miaB | tRNA (N6-isopentenyl adenosine(37)-C2)-methylthiotransferase MiaB | 0,74 | 1,4 | 0,96 | 0,66 | 0,22 | -0,44 | 1,2E-07 | 2,2E-09 | 5,6E-07 | 9,9E-08 | 5,7E-03 | 1,3E-04 | 0,0E+00 | 1,9E-10 | 0,0E+00 | 0,0E+00 | 0,0E+00 | 0,0E+00 | 1931578 | 1933002 | 1425 | + |
| Y11_RS08995 | Y11_19071 | | PhoH family protein | 0,21 | 0,54 | 0,41 | 0,33 | 0,2 | -0,13 | 9,3E-03 | 1,3E-09 | 8,1E-04 | 6,6E-04 | 7,8E-02 | 1,3E-01 | 0,0E+00 | 0,0E+00 | 0,0E+00 | 0,0E+00 | 0,0E+00 | 0,0E+00 | 1933300 | 1934373 | 1074 | + |
| Y11_RS09000 | Y11_19081 | ybeY | rRNA maturation RNase YbeY | 0,15 | 0,43 | 0,28 | 0,28 | 0,12 | -0,16 | 5,2E-01 | 9,8E-03 | 4,7E-01 | 2,3E-01 | 7,7E-01 | 6,7E-01 | 0,0E+00 | 0,0E+00 | 0,0E+00 | 0,0E+00 | 0,0E+00 | 0,0E+00 | 1934370 | 1934843 | 474 | + |
| Y11_RS09005 | Y11_19091 | corC | CNNM family magnesium/cobalt transport protein CorC | 0,1 | -0,54 | -0,34 | -0,65 | -0,45 | 0,2 | 1,8E-01 | 9,6E-05 | 2,5E-02 | 1,2E-05 | 5,9E-03 | 1,5E-01 | 0,0E+00 | 0,0E+00 | 0,0E+00 | 0,0E+00 | 0,0E+00 | 0,0E+00 | 1935157 | 1936035 | 879 | + |
| Y11_RS09010 | Y11_19101 | lnt | apolipoprotein N-acyltransferase | -1,2 | -0,15 | -0,57 | 1,05 | 0,63 | -0,42 | 2,4E-06 | 3,2E-01 | 5,2E-04 | 9,1E-05 | 5,4E-04 | 2,0E-02 | 1,4E-07 | 0,0E+00 | 0,0E+00 | 1,7E-06 | 0,0E+00 | 0,0E+00 | 1936043 | 1937620 | 1578 | + |
| Y11_RS09015 | Y11_19111 | | pyridoxamine 5'-phosphate oxidase family protein | -0,49 | 0,72 | -0,8 | 1,21 | -0,31 | -1,52 | 1,8E-03 | 2,6E-05 | 1,0E-03 | 3,5E-06 | 1,1E-01 | 1,2E-05 | 0,0E+00 | 0,0E+00 | 0,0E+00 | 7,7E-08 | 0,0E+00 | 1,4E-06 | 1937629 | 1938213 | 585 | + |
| Y11_RS09020 | Y11_19121 | | amino acid ABC transporter substrate-binding protein | 0,86 | -2,69 | -2,62 | -3,55 | -3,48 | 0,07 | 3,3E-08 | 2,5E-10 | 7,7E-09 | 2,0E-11 | 7,2E-10 | 5,6E-01 | 0,0E+00 | 4,0E-11 | 4,4E-10 | 6,9E-12 | 1,4E-10 | 0,0E+00 | 1938799 | 1939713 | 915 | + |
| Y11_RS09025 | Y11_19131 | | amino acid ABC transporter permease | 3,16 | 15 | 15 | 15 | 15 | 0 | 2,7E-04 | 0,0E+00 | 0,0E+00 | 0,0E+00 | 0,0E+00 | 0,0E+00 | 1,1E-05 | 0,0E+00 | 0,0E+00 | 0,0E+00 | 0,0E+00 | 0,0E+00 | 1940015 | 1940755 | 741 | + |
| Y11_RS09030 | Y11_19141 | gltK | glutamate/aspartate ABC transporter permease GltK | 1,16 | 5,12 | 2,38 | 3,98 | 1,21 | 0 | 1,2E-03 | 2,2E-08 | 1,5E-05 | 6,5E-08 | 8,8E-04 | 0,0E+00 | 4,3E-05 | 1,2E-09 | 2,4E-07 | 2,3E-09 | 2,0E-05 | 0,0E+00 | 1940755 | 1941429 | 675 | + |
| Y11_RS09035 | Y11_19151 | | amino acid ABC transporter ATP-binding protein | 0,67 | 0,72 | 1,65 | 0,05 | 0,98 | 0,93 | 4,8E-05 | 5,9E-05 | 7,4E-09 | 6,8E-01 | 7,7E-06 | 2,0E-05 | 0,0E+00 | 0,0E+00 | 4,3E-10 | 0,0E+00 | 0,0E+00 | 0,0E+00 | 1941429 | 1942154 | 726 | + |
| Y11_RS09040 | Y11_19161 | | zinc ribbon-containing protein | -0,28 | 2,25 | 3,29 | 2,53 | 3,57 | 1,04 | 9,5E-04 | 1,2E-08 | 7,0E-09 | 4,2E-09 | 3,4E-09 | 1,4E-04 | 0,0E+00 | 7,1E-10 | 4,1E-10 | 2,5E-10 | 4,2E-10 | 1,1E-05 | 1942208 | 1942690 | 483 | - |
| Y11_RS09045 | Y11_19181 | leuS | leucine--tRNA ligase | -0,35 | -0,14 | -0,36 | 0,21 | -0,01 | -0,22 | 2,3E-06 | 8,0E-04 | 3,5E-03 | 1,2E-04 | 9,2E-01 | 3,7E-02 | 0,0E+00 | 0,0E+00 | 0,0E+00 | 0,0E+00 | 0,0E+00 | 0,0E+00 | 1943039 | 1945621 | 2583 | + |
| Y11_RS09050 | Y11_19191 | lptE | LPS assembly lipoprotein LptE | -0,36 | 0,29 | 0,04 | 0,65 | 0,4 | -0,25 | 6,9E-05 | 1,1E-02 | 6,4E-01 | 1,1E-04 | 1,8E-03 | 6,0E-02 | 0,0E+00 | 0,0E+00 | 0,0E+00 | 0,0E+00 | 0,0E+00 | 0,0E+00 | 1945636 | 1946259 | 624 | + |
| Y11_RS09055 | Y11_19201 | holA | DNA polymerase III subunit delta | -0,27 | 0,79 | 0,35 | 1,07 | 0,63 | -0,44 | 1,0E-01 | 2,2E-05 | 3,3E-02 | 1,9E-04 | 1,2E-02 | 2,1E-02 | 0,0E+00 | 0,0E+00 | 0,0E+00 | 3,3E-06 | 0,0E+00 | 0,0E+00 | 1946256 | 1947290 | 1035 | + |
| Y11_RS09060 | Y11_19211 | nadD | nicotinate-nucleotide adenylyltransferase | -0,53 | 0,81 | 0,65 | 1,34 | 1,18 | -0,16 | 4,4E-04 | 1,2E-06 | 5,1E-05 | 7,7E-07 | 5,6E-06 | 1,1E-01 | 0,0E+00 | 0,0E+00 | 0,0E+00 | 2,0E-08 | 1,9E-07 | 0,0E+00 | 1947280 | 1947948 | 669 | + |
| Y11_RS09065 | Y11_19221 | rsfS | ribosome silencing factor | 0,02 | -1,16 | -1,73 | -1,18 | -1,75 | -0,56 | 9,0E-01 | 3,2E-06 | 7,3E-05 | 1,9E-04 | 2,3E-04 | 5,1E-02 | 0,0E+00 | 8,5E-08 | 1,0E-06 | 3,3E-06 | 5,6E-06 | 0,0E+00 | 1948430 | 1948747 | 318 | + |
| Y11_RS09070 | Y11_19231 | rlmH | 23S rRNA (pseudouridine(1915)-N(3))-methyltransferase RlmH | -0,34 | -0,09 | -0,53 | 0,25 | -0,18 | -0,43 | 4,5E-03 | 1,3E-01 | 1,5E-04 | 3,3E-02 | 1,5E-01 | 1,3E-03 | 0,0E+00 | 0,0E+00 | 0,0E+00 | 0,0E+00 | 0,0E+00 | 0,0E+00 | 1948751 | 1949221 | 471 | + |
| Y11_RS09075 | Y11_19241 | mrdA | peptidoglycan DD-transpeptidase MrdA | -0,35 | -0,74 | -1,32 | -0,39 | -0,98 | -0,59 | 8,6E-03 | 3,2E-05 | 1,8E-06 | 7,1E-04 | 7,3E-06 | 1,0E-04 | 0,0E+00 | 0,0E+00 | 3,7E-08 | 0,0E+00 | 0,0E+00 | 0,0E+00 | 1949296 | 1951191 | 1896 | + |
| Y11_RS09080 | Y11_19251 | mrdB | peptidoglycan glycosyltransferase MrdB | 1,05 | 1,35 | 1,85 | 0,3 | 0,8 | 0,5 | 1,2E-01 | 4,5E-02 | 8,1E-02 | 7,2E-01 | 4,8E-01 | 6,5E-01 | 4,0E-03 | 8,4E-04 | 9,0E-04 | 0,0E+00 | 0,0E+00 | 0,0E+00 | 1951228 | 1952340 | 1113 | + |
| Y11_RS09085 | Y11_19261 | rlpA | endolytic peptidoglycan transglycosylase RlpA | 0,19 | -0,14 | -0,05 | -0,33 | -0,24 | 0,09 | 1,0E-02 | 9,0E-02 | 6,3E-01 | 1,8E-03 | 4,6E-02 | 4,4E-01 | 0,0E+00 | 0,0E+00 | 0,0E+00 | 0,0E+00 | 0,0E+00 | 0,0E+00 | 1952350 | 1953447 | 1098 | + |
| Y11_RS09090 | Y11_19271 | dacA | D-alanyl-D-alanine carboxypeptidase DacA | -0,04 | -0,54 | -0,67 | -0,49 | -0,63 | -0,14 | 5,2E-01 | 5,2E-06 | 5,0E-05 | 3,2E-05 | 1,2E-04 | 1,4E-01 | 0,0E+00 | 0,0E+00 | 0,0E+00 | 0,0E+00 | 0,0E+00 | 0,0E+00 | 1953661 | 1954872 | 1212 | + |

| | | | | | | | | | | | | | | | | | | | | | | | | | |
|---|---|---|---|---|---|---|---|---|---|---|---|---|---|---|---|---|---|---|---|---|---|---|---|---|---|
| Y11_RS09095 | Y11_19281 | ybeD | DUF493 family protein YbeD | -0,11 | -1,08 | -1,55 | -0,97 | -1,44 | -0,48 | 4,5E-01 | 3,7E-06 | 1,4E-04 | 9,2E-05 | 3,8E-04 | 6,8E-02 | 0,0E+00 | 9,7E-08 | 1,9E-06 | 0,0E+00 | 9,1E-06 | 0,0E+00 | 1955021 | 1955284 | 264 | + |
| Y11_RS09100 | Y11_19291 | lipB | lipoyl(octanoyl) transferase LipB | -0,3 | -1,45 | -1,36 | -1,15 | -1,06 | 0,09 | 1,9E-01 | 5,4E-04 | 9,4E-06 | 6,3E-03 | 1,4E-03 | 7,5E-01 | 0,0E+00 | 1,1E-05 | 1,6E-07 | 1,0E-04 | 3,2E-05 | 0,0E+00 | 1955439 | 1956125 | 687 | + |
| Y11_RS09105 | Y11_19301 | lipA | lipoyl synthase | 0,03 | -1,53 | -2,72 | -1,56 | -2,75 | -1,19 | 7,3E-01 | 1,1E-06 | 5,6E-09 | 1,5E-06 | 9,5E-09 | 2,8E-05 | 0,0E+00 | 3,2E-08 | 3,5E-10 | 3,8E-08 | 9,3E-10 | 2,8E-06 | 1956341 | 1957306 | 966 | + |
| Y11_RS09110 | Y11_19311 | tatA | Sec-independent protein translocase subunit TatA | -1,31 | -0,59 | -2,37 | 0,73 | -1,03 | -1,76 | 5,6E-02 | 8,4E-02 | 1,1E-03 | 2,1E-01 | 1,5E-01 | 2,7E-03 | 1,9E-03 | 0,0E+00 | 1,3E-05 | 0,0E+00 | 3,1E-03 | 1,7E-04 | 1957437 | 1957652 | 216 | - |
| Y11_RS09125 | Y11_19341 | cspE | transcription antiterminator/RNA stability regulator CspE | 0,37 | 0,82 | 1,29 | 0,44 | 0,91 | 0,47 | 2,5E-02 | 8,4E-05 | 8,3E-03 | 1,8E-03 | 3,7E-02 | 2,2E-01 | 0,0E+00 | 0,0E+00 | 9,4E-05 | 0,0E+00 | 0,0E+00 | 0,0E+00 | 1959639 | 1959848 | 210 | - |
| Y11_RS09130 | Y11_19351 | | cold shock domain-containing protein | 0,58 | 0,62 | 0,92 | 0,03 | 0,33 | 0,3 | 3,2E-05 | 2,7E-06 | 6,7E-07 | 6,3E-01 | 2,6E-03 | 1,4E-03 | 0,0E+00 | 0,0E+00 | 0,0E+00 | 0,0E+00 | 0,0E+00 | 0,0E+00 | 1960115 | 1960594 | 480 | + |
| Y11_RS09150 | Y11_19391 | | glycogen synthase | 0,04 | 3,44 | 4,5 | 3,4 | 4,47 | 1,07 | 5,0E-01 | 8,2E-09 | 2,0E-08 | 9,3E-09 | 2,1E-08 | 2,4E-03 | 0,0E+00 | 5,3E-10 | 8,7E-10 | 5,0E-10 | 1,8E-09 | 1,5E-04 | 1963757 | 1963951 | 195 | - |
| Y11_RS09160 | Y11_19411 | | ABC transporter ATP-binding protein/permease | 0,65 | -0,09 | 0,01 | -0,74 | -0,64 | 0,11 | 7,9E-04 | 3,0E-01 | 9,6E-01 | 2,3E-04 | 4,3E-02 | 6,9E-01 | 0,0E+00 | 0,0E+00 | 0,0E+00 | 0,0E+00 | 0,0E+00 | 0,0E+00 | 1964598 | 1966370 | 1773 | - |
| Y11_RS09170 | Y11_19441 | | aromatic amino acid ammonia-lyase | -3,27 | -0,12 | 0,33 | 3,15 | 3,6 | 0,45 | 1,5E-11 | 1,9E-01 | 2,3E-02 | 6,6E-11 | 7,1E-10 | 6,4E-03 | 9,7E-12 | 0,0E+00 | 0,0E+00 | 1,4E-11 | 1,4E-10 | 0,0E+00 | 1968387 | 1969982 | 1596 | - |
| Y11_RS09225 | Y11_19581 | | toprim domain-containing protein | 0,29 | -0,6 | -1,16 | -0,9 | -1,45 | -0,56 | 1,2E-01 | 6,8E-03 | 1,6E-03 | 1,3E-03 | 5,3E-04 | 6,6E-02 | 0,0E+00 | 0,0E+00 | 1,9E-05 | 0,0E+00 | 1,3E-05 | 0,0E+00 | 1976150 | 1978444 | 2295 | + |
| Y11_RS09245 | | | DUF6088 family protein | 0,21 | 1,11 | 2,18 | 0,91 | 1,98 | 1,06 | 3,8E-01 | 1,1E-03 | 1,6E-06 | 3,2E-03 | 2,3E-06 | 2,0E-04 | 0,0E+00 | 2,2E-05 | 3,4E-08 | 0,0E+00 | 8,6E-08 | 1,5E-05 | 1982334 | 1982741 | 408 | + |
| Y11_RS09250 | Y11_19631 | | nucleotidyl transferase AbiEii/AbiGii toxin family protein | 0,06 | -1,04 | 0,18 | -1,1 | 0,12 | 1,22 | 6,4E-01 | 4,3E-04 | 2,4E-01 | 3,4E-04 | 4,4E-01 | 2,8E-04 | 0,0E+00 | 8,8E-06 | 0,0E+00 | 5,9E-06 | 0,0E+00 | 2,0E-05 | 1982722 | 1983813 | 1092 | + |
| Y11_RS09265 | Y11_19651 | folD | bifunctional methylenetetrahydrofolate dehydrogenase/methenyltetrahydrofolate cyclohydrolase FolD | 0,09 | 0,63 | 0,51 | 0,54 | 0,42 | -0,12 | 1,5E-01 | 1,2E-05 | 2,4E-04 | 4,3E-06 | 2,7E-04 | 1,5E-01 | 0,0E+00 | 0,0E+00 | 0,0E+00 | 0,0E+00 | 0,0E+00 | 0,0E+00 | 1985393 | 1986259 | 867 | + |
| Y11_RS09270 | Y11_19661 | ybcJ | ribosome-associated protein YbcJ | -0,19 | -0,14 | -0,66 | 0,05 | -0,47 | -0,52 | 1,6E-01 | 1,8E-01 | 4,8E-04 | 6,7E-01 | 7,2E-03 | 1,3E-03 | 0,0E+00 | 0,0E+00 | 0,0E+00 | 0,0E+00 | 0,0E+00 | 0,0E+00 | 1986274 | 1986486 | 213 | + |
| Y11_RS09275 | Y11_19671 | cysS | cysteine--tRNA ligase | 0,06 | 0,94 | 0,65 | 0,87 | 0,58 | -0,29 | 3,6E-01 | 7,9E-07 | 1,2E-05 | 1,4E-06 | 2,7E-05 | 3,3E-03 | 0,0E+00 | 0,0E+00 | 0,0E+00 | 0,0E+00 | 0,0E+00 | 0,0E+00 | 1986655 | 1988040 | 1386 | - |
| Y11_RS09280 | Y11_19681 | ppiB | peptidylprolyl isomerase B | -0,09 | 0,84 | 0,72 | 0,93 | 0,8 | -0,12 | 3,9E-01 | 3,6E-05 | 2,0E-05 | 2,8E-05 | 1,8E-05 | 2,4E-01 | 0,0E+00 | 0,0E+00 | 0,0E+00 | 0,0E+00 | 0,0E+00 | 0,0E+00 | 1988253 | 1988747 | 495 | + |
| Y11_RS09285 | Y11_19691 | | UDP-2,3-diacylglucosamine diphosphatase | -0,16 | 0,58 | 0,58 | 0,74 | 0,74 | 0 | 4,5E-01 | 4,1E-02 | 2,8E-02 | 3,6E-04 | 1,0E-05 | 1,0E+00 | 0,0E+00 | 0,0E+00 | 0,0E+00 | 0,0E+00 | 0,0E+00 | 0,0E+00 | 1988758 | 1989480 | 723 | + |
| Y11_RS09290 | Y11_19701 | purE | 5-(carboxyamino)imidazole ribonucleotide mutase | 0,24 | 1,5 | 1,58 | 1,26 | 1,34 | 0,08 | 4,8E-02 | 2,1E-07 | 2,2E-06 | 1,2E-07 | 3,5E-06 | 4,9E-01 | 0,0E+00 | 7,6E-09 | 4,5E-08 | 3,8E-09 | 1,3E-07 | 0,0E+00 | 1989656 | 1990180 | 525 | + |
| Y11_RS09295 | Y11_19711 | purK | 5-(carboxyamino)imidazole ribonucleotide synthase | 0,08 | 1,15 | 1,42 | 1,07 | 1,34 | 0,27 | 5,0E-01 | 4,3E-05 | 6,1E-06 | 6,7E-06 | 5,0E-07 | 6,2E-02 | 0,0E+00 | 9,6E-07 | 1,1E-07 | 1,4E-07 | 2,4E-08 | 0,0E+00 | 1990177 | 1991241 | 1065 | + |
| Y11_RS09305 | Y11_19741 | | ABC transporter ATP-binding protein | -0,21 | -3,84 | -2,1 | -3,63 | -1,89 | 1,74 | 3,5E-02 | 6,9E-13 | 9,4E-09 | 2,5E-10 | 2,6E-07 | 1,0E-07 | 0,0E+00 | 1,6E-12 | 5,0E-10 | 3,0E-11 | 1,4E-08 | 3,8E-08 | 1993804 | 1994490 | 687 | - |
| Y11_RS09310 | Y11_19751 | tesA | multifunctional acyl-CoA thioesterase I/protease I/lysophospholipase L1 | 0,51 | -3,51 | -3,03 | -4,02 | -3,54 | 0,49 | 3,0E-04 | 2,2E-11 | 8,2E-10 | 3,2E-11 | 5,5E-10 | 6,4E-04 | 0,0E+00 | 8,7E-12 | 8,6E-11 | 8,8E-12 | 1,2E-10 | 0,0E+00 | 1994461 | 1995096 | 636 | + |
| Y11_RS09315 | Y11_19761 | | SDR family oxidoreductase | 0,64 | -1,49 | -1,32 | -2,12 | -1,96 | 0,17 | 1,8E-05 | 1,6E-09 | 1,1E-05 | 1,0E-09 | 8,0E-07 | 2,5E-01 | 0,0E+00 | 1,5E-10 | 1,8E-07 | 9,0E-11 | 3,4E-08 | 0,0E+00 | 1995142 | 1995918 | 777 | + |
| Y11_RS09320 | Y11_19771 | | co-chaperone YbbN | 0,24 | -0,76 | 0,1 | -1 | -0,14 | 0,86 | 1,6E-03 | 2,8E-09 | 8,3E-01 | 3,2E-08 | 7,7E-01 | 1,1E-01 | 0,0E+00 | 0,0E+00 | 0,0E+00 | 1,3E-09 | 0,0E+00 | 0,0E+00 | 1996115 | 1996984 | 870 | + |
| Y11_RS09325 | Y11_19781 | | SPFH/Band 7/PHB domain protein | 0,93 | -2,88 | -0,78 | -3,81 | -1,71 | 2,1 | 4,4E-08 | 1,2E-10 | 1,5E-04 | 3,0E-11 | 5,8E-07 | 2,2E-07 | 0,0E+00 | 2,4E-11 | 0,0E+00 | 8,5E-12 | 2,7E-08 | 6,0E-08 | 1997181 | 1998095 | 915 | + |
| Y11_RS09330 | Y11_19791 | | NfeD family protein | 0,36 | -1,25 | 0,36 | -1,61 | 0 | 1,61 | 3,2E-01 | 6,3E-02 | 1,6E-01 | 2,6E-02 | 1,0E+00 | 1,8E-02 | 0,0E+00 | 1,2E-03 | 0,0E+00 | 4,2E-04 | 0,0E+00 | 9,8E-04 | 1998098 | 1998547 | 450 | + |
| Y11_RS09335 | Y11_19801 | cueR | Cu(I)-responsive transcriptional regulator | 0,34 | 0,3 | 0,64 | -0,04 | 0,29 | 0,34 | 1,5E-01 | 6,2E-02 | 6,9E-04 | 8,4E-01 | 1,7E-01 | 1,1E-02 | 0,0E+00 | 0,0E+00 | 0,0E+00 | 0,0E+00 | 0,0E+00 | 0,0E+00 | 1998593 | 1999012 | 420 | - |
| Y11_RS09340 | Y11_19811 | copA | copper-exporting P-type ATPase CopA | -0,11 | 1,83 | 2,32 | 1,94 | 2,43 | 0,49 | 8,3E-02 | 2,4E-08 | 2,3E-08 | 1,8E-08 | 1,8E-08 | 5,2E-03 | 0,0E+00 | 1,3E-09 | 9,9E-10 | 8,2E-10 | 1,5E-09 | 0,0E+00 | 1999209 | 2001965 | 2757 | + |
| Y11_RS09350 | Y11_19831 | ybaK | Cys-tRNA(Pro)/Cys-tRNA(Cys) deacylase YbaK | -0,19 | 0,16 | -0,13 | 0,35 | 0,06 | -0,29 | 3,4E-02 | 3,1E-03 | 2,1E-01 | 1,6E-03 | 6,0E-01 | 1,5E-02 | 0,0E+00 | 0,0E+00 | 0,0E+00 | 0,0E+00 | 0,0E+00 | 0,0E+00 | 2003130 | 2003609 | 480 | + |
| Y11_RS09360 | Y11_19841 | ushA | bifunctional UDP-sugar hydrolase/5'-nucleotidase UshA | -0,38 | -0,88 | -0,14 | -0,5 | 0,24 | 0,74 | 3,7E-05 | 1,5E-06 | 2,0E-02 | 7,0E-05 | 7,9E-04 | 5,2E-06 | 0,0E+00 | 0,0E+00 | 0,0E+00 | 0,0E+00 | 0,0E+00 | 0,0E+00 | 2004163 | 2005815 | 1653 | - |
| Y11_RS09370 | Y11_19891 | | Kef family K(+) transporter | -0,46 | 0,69 | 0,89 | 1,15 | 1,35 | 0,2 | 8,9E-03 | 2,5E-03 | 1,4E-03 | 2,9E-05 | 4,1E-05 | 3,2E-01 | 0,0E+00 | 0,0E+00 | 0,0E+00 | 5,6E-07 | 1,1E-06 | 0,0E+00 | 2007752 | 2009443 | 1692 | + |
| Y11_RS09380 | Y11_19911 | galE | UDP-glucose 4-epimerase GalE | -1,24 | 0,35 | 0,03 | 1,59 | 1,28 | -0,31 | 4,1E-06 | 8,2E-03 | 7,8E-01 | 3,3E-07 | 3,4E-06 | 1,3E-02 | 2,3E-07 | 0,0E+00 | 0,0E+00 | 9,2E-09 | 1,2E-07 | 0,0E+00 | 2011029 | 2012039 | 1011 | - |
| Y11_RS09385 | Y11_19921 | | glycosyltransferase | -1,32 | -0,28 | -0,3 | 1,04 | 1,03 | -0,02 | 1,7E-06 | 1,9E-01 | 9,7E-02 | 6,1E-04 | 1,5E-04 | 9,4E-01 | 9,9E-08 | 0,0E+00 | 0,0E+00 | 1,0E-05 | 3,8E-06 | 0,0E+00 | 2012077 | 2013093 | 1017 | - |
| Y11_RS09390 | Y11_19931 | | polysaccharide biosynthesis protein | -0,7 | 0,3 | 0,36 | 1 | 1,05 | 0,05 | 1,3E-04 | 2,1E-02 | 3,5E-02 | 3,7E-07 | 1,5E-05 | 6,6E-01 | 0,0E+00 | 0,0E+00 | 0,0E+00 | 0,0E+00 | 4,8E-07 | 0,0E+00 | 2013148 | 2015064 | 1917 | - |
| Y11_RS09400 | Y11_19951 | | glycosyltransferase | -0,54 | 0,8 | 0,48 | 1,33 | 1,02 | -0,32 | 1,1E-01 | 3,8E-03 | 5,0E-02 | 5,5E-04 | 3,7E-03 | 2,1E-02 | 0,0E+00 | 0,0E+00 | 0,0E+00 | 9,5E-06 | 8,0E-05 | 0,0E+00 | 2016249 | 2017352 | 1104 | - |
| Y11_RS09405 | Y11_19961 | | glycosyltransferase family 4 protein | -0,11 | 0,92 | 1,01 | 1,04 | 1,12 | 0,09 | 5,4E-01 | 7,2E-04 | 9,3E-05 | 4,9E-04 | 7,8E-05 | 5,7E-01 | 0,0E+00 | 0,0E+00 | 1,3E-06 | 8,4E-06 | 2,1E-06 | 0,0E+00 | 2017474 | 2018550 | 1077 | - |
| Y11_RS09410 | Y11_19971 | | glycosyltransferase family 2 protein | -0,07 | 0,34 | 0,35 | 0,41 | 0,41 | 0,01 | 6,1E-01 | 4,1E-02 | 3,9E-02 | 3,5E-03 | 3,5E-03 | 9,6E-01 | 0,0E+00 | 0,0E+00 | 0,0E+00 | 0,0E+00 | 0,0E+00 | 0,0E+00 | 2018559 | 2019437 | 879 | - |
| Y11_RS09415 | Y11_19981 | | glycosyltransferase | -0,61 | 0,11 | -0,05 | 0,72 | 0,56 | -0,16 | 1,2E-04 | 4,5E-01 | 7,9E-01 | 5,3E-04 | 1,9E-02 | 4,8E-01 | 0,0E+00 | 0,0E+00 | 0,0E+00 | 0,0E+00 | 0,0E+00 | 0,0E+00 | 2019439 | 2020395 | 957 | - |
| Y11_RS09425 | Y11_20001 | hemH | ferrochelatase | 0,63 | -0,59 | 1,13 | -1,22 | 0,5 | 1,72 | 2,3E-02 | 3,4E-03 | 2,2E-04 | 6,0E-04 | 7,7E-02 | 9,9E-06 | 0,0E+00 | 0,0E+00 | 2,8E-06 | 1,0E-05 | 0,0E+00 | 1,1E-06 | 2021995 | 2022978 | 984 | - |
| Y11_RS09430 | Y11_20011 | adk | adenylate kinase | -0,26 | 0,51 | 0,22 | 0,77 | 0,49 | -0,28 | 8,9E-04 | 2,0E-06 | 6,8E-04 | 1,8E-06 | 5,4E-05 | 8,9E-04 | 0,0E+00 | 0,0E+00 | 0,0E+00 | 0,0E+00 | 0,0E+00 | 0,0E+00 | 2023094 | 2023738 | 645 | - |
| Y11_RS09435 | Y11_20031 | htpG | molecular chaperone HtpG | 0,08 | 0,7 | 0,42 | 0,63 | 0,34 | -0,29 | 2,8E-01 | 1,4E-07 | 1,8E-06 | 1,1E-05 | 4,8E-04 | 2,4E-05 | 0,0E+00 | 0,0E+00 | 0,0E+00 | 0,0E+00 | 0,0E+00 | 0,0E+00 | 2023964 | 2025832 | 1869 | - |
| Y11_RS09440 | Y11_20041 | recR | recombination mediator RecR | 0,15 | 0,62 | -0,17 | 0,47 | -0,32 | -0,8 | 3,0E-01 | 2,2E-04 | 2,6E-01 | 4,5E-03 | 7,8E-02 | 2,7E-04 | 0,0E+00 | 0,0E+00 | 0,0E+00 | 0,0E+00 | 0,0E+00 | 0,0E+00 | 2026171 | 2026776 | 606 | - |
| Y11_RS09445 | Y11_20051 | | YbaB/EbfC family nucleoid-associated protein | 0,23 | -0,22 | -0,31 | -0,45 | -0,54 | -0,09 | 1,1E-01 | 3,3E-01 | 6,1E-02 | 5,4E-02 | 2,1E-03 | 6,9E-01 | 0,0E+00 | 0,0E+00 | 0,0E+00 | 0,0E+00 | 0,0E+00 | 0,0E+00 | 2026776 | 2027108 | 333 | - |
| Y11_RS09450 | Y11_20061 | dnaX | DNA polymerase III subunit gamma/tau | -0,05 | 0,3 | -0,48 | 0,35 | -0,43 | -0,78 | 3,3E-01 | 1,1E-05 | 1,4E-04 | 1,6E-04 | 8,8E-04 | 4,6E-06 | 0,0E+00 | 0,0E+00 | 0,0E+00 | 0,0E+00 | 0,0E+00 | 0,0E+00 | 2027164 | 2029134 | 1971 | - |
| Y11_RS09455 | Y11_20071 | apt | adenine phosphoribosyltransferase | 0,22 | -0,99 | -1,12 | -1,21 | -1,34 | -0,14 | 1,3E-02 | 1,4E-06 | 3,5E-04 | 2,3E-07 | 1,0E-04 | 4,9E-01 | 0,0E+00 | 0,0E+00 | 4,4E-06 | 6,9E-09 | 2,6E-06 | 0,0E+00 | 2029218 | 2029781 | 564 | - |
| Y11_RS09465 | Y11_20091 | | primosomal replication protein | 0,28 | 0,72 | 0,95 | 0,44 | 0,67 | 0,23 | 7,8E-02 | 3,1E-04 | 7,1E-05 | 4,4E-03 | 5,4E-04 | 4,3E-02 | 0,0E+00 | 0,0E+00 | 0,0E+00 | 0,0E+00 | 0,0E+00 | 0,0E+00 | 2030554 | 2031090 | 537 | + |
| Y11_RS09475 | Y11_20111 | mscK | mechanosensitive channel MscK | -1,55 | 0,73 | 1,41 | 2,28 | 2,96 | 0,68 | 1,9E-11 | 1,3E-07 | 1,4E-08 | 1,7E-11 | 3,9E-11 | 9,7E-06 | 9,7E-12 | 0,0E+00 | 6,8E-10 | 6,5E-12 | 2,4E-11 | 0,0E+00 | 2031625 | 2035074 | 3450 | - |
| Y11_RS09480 | Y11_20121 | | DsrE/DsrF/TusD sulfur relay family protein | 0,29 | 0,92 | 1,44 | 0,63 | 1,15 | 0,52 | 1,1E-01 | 6,2E-06 | 4,8E-04 | 3,1E-03 | 3,6E-03 | 7,2E-02 | 0,0E+00 | 0,0E+00 | 5,9E-06 | 0,0E+00 | 7,9E-05 | 0,0E+00 | 2035265 | 2035630 | 366 | - |
| Y11_RS09490 | Y11_20141 | acrA | efflux RND transporter adaptor subunit AcrA | 0,22 | 0,82 | 0,87 | 0,6 | 0,65 | 0,05 | 2,7E-03 | 2,9E-06 | 6,7E-07 | 1,1E-05 | 1,6E-06 | 4,9E-01 | 0,0E+00 | 0,0E+00 | 0,0E+00 | 0,0E+00 | 0,0E+00 | 0,0E+00 | 2036446 | 2037633 | 1188 | + |
| Y11_RS09495 | Y11_20151 | acrB | efflux RND transporter permease AcrB | 0,14 | 1,2 | 1,08 | 1,06 | 0,94 | -0,11 | 8,6E-03 | 3,5E-08 | 5,3E-08 | 1,4E-07 | 2,5E-07 | 1,6E-01 | 0,0E+00 | 1,7E-09 | 1,9E-09 | 4,6E-09 | 0,0E+00 | 0,0E+00 | 2037650 | 2040802 | 3153 | + |
| Y11_RS09520 | Y11_20231 | | DUF2620 domain-containing protein | -0,63 | 0,8 | 1,05 | 1,44 | 1,68 | 0,25 | 2,6E-04 | 3,2E-05 | 8,7E-04 | 1,3E-07 | 2,8E-05 | 2,4E-01 | 0,0E+00 | 0,0E+00 | 1,1E-05 | 4,4E-09 | 8,2E-07 | 0,0E+00 | 2045968 | 2046333 | 366 | + |
| Y11_RS09540 | Y11_20271 | | metal ABC transporter substrate-binding protein | -1,17 | 0,86 | 0,64 | 2,03 | 1,81 | -0,22 | 7,0E-04 | 2,8E-02 | 4,7E-02 | 2,3E-04 | 1,7E-04 | 5,5E-01 | 2,6E-05 | 0,0E+00 | 0,0E+00 | 4,1E-06 | 4,3E-06 | 0,0E+00 | 2048422 | 2049300 | 879 | - |
| Y11_RS09560 | Y11_20321 | tomB | Hha toxicity modulator TomB | 0 | -15 | 0 | -15 | 0 | 15 | 0,0E+00 | 0,0E+00 | 0,0E+00 | 0,0E+00 | 0,0E+00 | 0,0E+00 | 0,0E+00 | 0,0E+00 | 0,0E+00 | 0,0E+00 | 0,0E+00 | 0,0E+00 | 2052221 | 2052589 | 369 | + |
| Y11_RS09565 | Y11_20331 | ymoA | expression modulating protein YmoA | -0,19 | -2,41 | -3 | -2,22 | -2,81 | -0,59 | 2,0E-01 | 5,0E-09 | 8,8E-09 | 5,7E-07 | 2,4E-07 | 3,9E-03 | 0,0E+00 | 3,7E-10 | 4,8E-10 | 1,5E-08 | 1,3E-08 | 0,0E+00 | 2052636 | 2052839 | 204 | + |
| Y11_RS09575 | Y11_20361 | | YbaY family lipoprotein | -0,34 | 3,64 | 3,49 | 3,98 | 3,83 | -0,15 | 1,0E-02 | 2,8E-08 | 1,2E-07 | 1,2E-08 | 5,0E-08 | 5,5E-01 | 0,0E+00 | 1,4E-09 | 3,7E-09 | 5,9E-10 | 3,5E-09 | 0,0E+00 | 2054033 | 2054563 | 531 | - |
| Y11_RS09580 | Y11_20371 | tesB | acyl-CoA thioesterase II | 0,2 | 1,47 | 1,98 | 1,26 | 1,78 | 0,51 | 1,1E-02 | 1,4E-08 | 3,5E-09 | 2,0E-08 | 4,4E-09 | 8,2E-05 | 0,0E+00 | 8,2E-10 | 2,4E-10 | 8,9E-10 | 5,0E-10 | 0,0E+00 | 2054824 | 2055687 | 864 | + |
| Y11_RS09585 | Y11_20381 | amtB | ammonium transporter AmtB | 1,36 | -0,56 | -0,98 | -1,86 | -2,24 | -0,42 | 3,2E-03 | 9,5E-02 | 3,7E-02 | 1,2E-05 | 1,3E-04 | 2,0E-01 | 1,1E-04 | 0,0E+00 | 0,0E+00 | 2,4E-07 | 3,4E-06 | 0,0E+00 | 2055801 | 2057090 | 1290 | - |
| Y11_RS09590 | Y11_20391 | glnK | P-II family nitrogen regulator | 0,81 | -0,49 | -0,87 | -1,31 | -1,69 | -0,38 | 1,7E-06 | 3,8E-05 | 5,6E-06 | 1,6E-07 | 1,1E-07 | 3,4E-03 | 0,0E+00 | 0,0E+00 | 0,0E+00 | 5,0E-09 | 6,8E-09 | 0,0E+00 | 2057130 | 2057468 | 339 | - |
| Y11_RS09595 | Y11_20401 | | SmdB family multidrug efflux ABC transporter permease/ATP-binding protein | 0,11 | 0,34 | 0,27 | 0,23 | 0,16 | -0,07 | 5,7E-01 | 1,0E-01 | 2,8E-01 | 2,1E-01 | 4,9E-01 | 7,4E-01 | 0,0E+00 | 0,0E+00 | 0,0E+00 | 0,0E+00 | 0,0E+00 | 0,0E+00 | 2057906 | 2059684 | 1779 | - |
| Y11_RS09600 | Y11_20411 | | SmdA family multidrug ABC transporter permease/ATP-binding protein | -0,08 | 0,43 | 0,26 | 0,51 | 0,34 | -0,17 | 3,1E-01 | 1,2E-04 | 1,6E-02 | 1,0E-04 | 6,3E-03 | 8,5E-02 | 0,0E+00 | 0,0E+00 | 0,0E+00 | 0,0E+00 | 0,0E+00 | 0,0E+00 | 2059677 | 2061443 | 1767 | - |
| Y11_RS09605 | Y11_20421 | | Lrp/AsnC family transcriptional regulator | -0,34 | 0,32 | 0,52 | 0,65 | 0,82 | 0,2 | 8,2E-02 | 1,8E-01 | 1,4E-02 | 1,2E-02 | 7,5E-04 | 3,5E-01 | 0,0E+00 | 0,0E+00 | 0,0E+00 | 0,0E+00 | 0,0E+00 | 0,0E+00 | 2061718 | 2062179 | 462 | - |
| Y11_RS09610 | Y11_20431 | | PLP-dependent cysteine synthase family protein | -0,11 | -2 | -2,26 | -1,88 | -2,15 | -0,26 | 5,7E-01 | 1,2E-05 | 1,0E-05 | 1,3E-06 | 2,1E-06 | 2,2E-01 | 0,0E+00 | 2,7E-07 | 1,7E-07 | 3,3E-08 | 8,2E-08 | 0,0E+00 | 2062306 | 2063346 | 1041 | + |
| Y11_RS09625 | Y11_20461 | queC | 7-cyano-7-deazaguanine synthase QueC | 0,21 | 1,93 | 2,42 | 1,72 | 2,21 | 0,49 | 1,1E-01 | 1,6E-07 | 2,2E-07 | 6,9E-08 | 1,8E-07 | 5,3E-03 | 0,0E+00 | 6,1E-09 | 6,4E-09 | 2,5E-09 | 1,0E-08 | 0,0E+00 | 2066214 | 2066912 | 699 | + |
| Y11_RS09640 | Y11_20491 | ppiD | peptidylprolyl isomerase | -0,18 | 0,15 | 0,13 | 0,33 | 0,31 | -0,02 | 8,7E-04 | 7,5E-04 | 6,3E-02 | 3,9E-06 | 8,9E-04 | 7,2E-01 | 0,0E+00 | 0,0E+00 | 0,0E+00 | 0,0E+00 | 0,0E+00 | 0,0E+00 | 2068215 | 2070101 | 1887 | - |
| Y11_RS09645 | Y11_20511 | hupB | DNA-binding protein HU-beta | -0,49 | 0,74 | 0,76 | 1,23 | 1,24 | 0,02 | 1,0E-04 | 1,3E-04 | 4,8E-03 | 3,6E-06 | 2,3E-04 | 9,4E-01 | 0,0E+00 | 0,0E+00 | 0,0E+00 | 8,0E-08 | 5,7E-06 | 0,0E+00 | 2070301 | 2070573 | 273 | - |
| Y11_RS09650 | Y11_20521 | lon | endopeptidase La | 0,15 | 0,64 | 0,48 | 0,49 | 0,33 | -0,16 | 1,4E-02 | 1,3E-08 | 5,0E-05 | 5,3E-06 | 1,7E-03 | 2,8E-02 | 0,0E+00 | 0,0E+00 | 0,0E+00 | 0,0E+00 | 0,0E+00 | 0,0E+00 | 2070791 | 2073145 | 2355 | - |
| Y11_RS09655 | Y11_20541 | clpX | ATP-dependent protease ATP-binding subunit ClpX | -0,39 | -0,43 | -1,01 | -0,04 | -0,63 | -0,59 | 4,0E-05 | 3,2E-04 | 5,1E-07 | 4,9E-01 | 2,7E-06 | 5,2E-05 | 0,0E+00 | 0,0E+00 | 1,3E-08 | 0,0E+00 | 0,0E+00 | 0,0E+00 | 2073340 | 2074611 | 1272 | - |
| Y11_RS09660 | Y11_20551 | clpP | ATP-dependent Clp endopeptidase proteolytic subunit ClpP | -0,76 | 0,72 | 0,76 | 1,49 | 1,52 | 0,04 | 1,6E-05 | 2,1E-05 | 7,6E-06 | 2,3E-07 | 1,2E-07 | 6,6E-01 | 0,0E+00 | 0,0E+00 | 0,0E+00 | 6,8E-09 | 6,9E-09 | 0,0E+00 | 2074817 | 2075440 | 624 | - |
| Y11_RS09665 | Y11_20561 | tig | trigger factor | 0,15 | -0,67 | -1,19 | -0,81 | -1,34 | -0,52 | 2,8E-02 | 2,8E-05 | 1,8E-07 | 1,6E-05 | 2,3E-07 | 8,2E-04 | 0,0E+00 | 0,0E+00 | 5,3E-09 | 0,0E+00 | 1,2E-08 | 0,0E+00 | 2075897 | 2077201 | 1305 | - |

| | | | | | | | | | | | | | | | | | | | | | | | | | |
|---|---|---|---|---|---|---|---|---|---|---|---|---|---|---|---|---|---|---|---|---|---|---|---|---|---|
| Y11_RS09670 | Y11_20571 | bolA | transcriptional regulator BolA | 0,15 | 1,25 | 0,47 | 1,1 | 0,32 | -0,78 | 4,0E-01 | 4,2E-06 | 2,8E-02 | 1,3E-04 | 1,7E-01 | 1,4E-03 | 0,0E+00 | 1,1E-07 | 0,0E+00 | 2,3E-06 | 0,0E+00 | 0,0E+00 | 2077617 | 2077946 | 330 | - |
| Y11_RS09675 | Y11_20581 | | lipoprotein | -0,02 | 0,05 | 0,07 | 0,07 | 0,09 | 0,02 | 7,8E-01 | 5,6E-01 | 5,3E-01 | 3,9E-01 | 4,2E-01 | 8,7E-01 | 0,0E+00 | 0,0E+00 | 0,0E+00 | 0,0E+00 | 0,0E+00 | 0,0E+00 | 2078300 | 2078878 | 579 | + |
| Y11_RS09685 | Y11_20601 | | ion channel protein Tsx | 2,04 | -0,44 | -1,5 | -2,48 | -3,54 | -1,06 | 5,4E-05 | 1,6E-02 | 3,1E-06 | 1,0E-05 | 5,4E-07 | 1,6E-05 | 2,5E-06 | 0,0E+00 | 5,9E-08 | 2,1E-07 | 2,6E-08 | 1,7E-06 | 2080699 | 2081478 | 780 | + |
| Y11_RS09690 | Y11_20611 | cyoA | cytochrome o ubiquinol oxidase subunit II | -0,05 | 1,93 | 1,42 | 1,98 | 1,47 | -0,51 | 6,7E-01 | 6,9E-06 | 5,7E-06 | 1,8E-06 | 2,8E-07 | 1,9E-02 | 0,0E+00 | 1,7E-07 | 1,0E-07 | 4,2E-08 | 1,4E-08 | 0,0E+00 | 2081841 | 2082797 | 957 | + |
| Y11_RS09695 | Y11_20621 | cyoB | cytochrome o ubiquinol oxidase subunit I | 0,24 | 3,02 | 2,64 | 2,78 | 2,4 | -0,38 | 4,2E-02 | 1,1E-07 | 3,7E-05 | 2,4E-07 | 7,8E-05 | 3,1E-01 | 0,0E+00 | 4,3E-09 | 5,5E-07 | 7,1E-09 | 2,1E-06 | 0,0E+00 | 2082802 | 2084793 | 1992 | + |
| Y11_RS09720 | Y11_20671 | | YajQ family cyclic di-GMP-binding protein | 0,01 | -0,3 | 0,04 | -0,31 | 0,03 | 0,34 | 8,0E-01 | 1,8E-04 | 7,3E-01 | 7,4E-04 | 8,2E-01 | 2,0E-02 | 0,0E+00 | 0,0E+00 | 0,0E+00 | 0,0E+00 | 0,0E+00 | 0,0E+00 | 2088240 | 2088731 | 492 | - |
| Y11_RS09725 | Y11_20681 | panE | 2-dehydropantoate 2-reductase | 0 | 2,14 | 2,73 | 2,15 | 2,73 | 0,58 | 9,8E-01 | 6,8E-08 | 1,0E-07 | 3,9E-07 | 3,0E-07 | 1,3E-02 | 0,0E+00 | 3,0E-09 | 3,4E-09 | 1,1E-08 | 1,5E-08 | 0,0E+00 | 2088941 | 2089852 | 912 | + |
| Y11_RS09730 | Y11_20691 | yajL | protein deglycase YajL | 0,53 | 1,44 | 2,4 | 0,91 | 1,86 | 0,95 | 7,0E-05 | 3,7E-09 | 1,0E-06 | 3,3E-06 | 1,0E-05 | 8,9E-04 | 0,0E+00 | 2,9E-10 | 2,4E-08 | 0,0E+00 | 3,4E-07 | 0,0E+00 | 2089815 | 2090405 | 591 | + |
| Y11_RS09735 | Y11_20701 | thiI | tRNA 4-thiouridine(8) synthase ThiI | -0,61 | -0,28 | -0,62 | 0,33 | -0,01 | -0,34 | 2,1E-05 | 4,7E-05 | 1,1E-04 | 1,3E-03 | 9,2E-01 | 4,7E-03 | 0,0E+00 | 0,0E+00 | 0,0E+00 | 0,0E+00 | 0,0E+00 | 0,0E+00 | 2090775 | 2092226 | 1452 | - |
| Y11_RS09740 | Y11_20711 | xseB | exodeoxyribonuclease VII small subunit | 0,26 | -0,04 | 0,02 | -0,3 | -0,25 | 0,05 | 1,9E-03 | 4,9E-01 | 8,2E-01 | 4,8E-04 | 1,0E-02 | 4,5E-01 | 0,0E+00 | 0,0E+00 | 0,0E+00 | 0,0E+00 | 0,0E+00 | 0,0E+00 | 2092506 | 2092760 | 255 | + |
| Y11_RS09745 | Y11_20721 | ispA | (2E,6E)-farnesyl diphosphate synthase | -0,24 | 0,3 | 0,44 | 0,54 | 0,68 | 0,14 | 5,3E-03 | 8,1E-03 | 1,6E-04 | 2,8E-04 | 1,0E-05 | 1,4E-01 | 0,0E+00 | 0,0E+00 | 0,0E+00 | 0,0E+00 | 0,0E+00 | 0,0E+00 | 2092765 | 2093685 | 921 | + |
| Y11_RS09750 | Y11_20731 | dxs | 1-deoxy-D-xylulose-5-phosphate synthase | -0,1 | 0,67 | 0,87 | 0,78 | 0,97 | 0,2 | 8,4E-02 | 8,2E-06 | 5,6E-06 | 7,6E-07 | 1,1E-06 | 5,3E-02 | 0,0E+00 | 0,0E+00 | 0,0E+00 | 0,0E+00 | 0,0E+00 | 0,0E+00 | 2093739 | 2095598 | 1860 | + |
| Y11_RS09765 | Y11_20761 | thiL | thiamine-phosphate kinase | -1,36 | 0,39 | 0,58 | 1,75 | 1,94 | 0,18 | 2,0E-08 | 1,4E-04 | 1,9E-03 | 1,1E-08 | 5,6E-07 | 2,1E-01 | 2,2E-09 | 0,0E+00 | 0,0E+00 | 5,7E-10 | 2,6E-08 | 0,0E+00 | 2097660 | 2098649 | 990 | - |
| Y11_RS09770 | Y11_20771 | nusB | transcription antitermination factor NusB | -0,43 | -0,35 | -0,62 | 0,08 | -0,19 | -0,27 | 6,7E-05 | 1,2E-04 | 5,0E-05 | 3,5E-02 | 2,3E-02 | 2,7E-03 | 0,0E+00 | 0,0E+00 | 0,0E+00 | 0,0E+00 | 0,0E+00 | 0,0E+00 | 2098729 | 2099145 | 417 | - |
| Y11_RS09775 | Y11_20781 | ribE | 6,7-dimethyl-8-ribityllumazine synthase | -0,31 | 0,4 | 0,31 | 0,71 | 0,62 | -0,1 | 1,6E-02 | 5,0E-03 | 2,1E-02 | 1,5E-04 | 4,5E-04 | 4,2E-01 | 0,0E+00 | 0,0E+00 | 0,0E+00 | 0,0E+00 | 0,0E+00 | 0,0E+00 | 2099171 | 2099641 | 471 | - |
| Y11_RS09780 | Y11_20791 | ribD | bifunctional diaminohydroxyphosphoribosylaminopyrimidine deaminase/5-amino-6-(5-phosphoribosylamino)uracil reductase RibD | 0,21 | 0,57 | 0,73 | 0,36 | 0,52 | 0,16 | 1,3E-02 | 4,3E-06 | 1,2E-04 | 1,4E-03 | 2,4E-03 | 1,8E-01 | 0,0E+00 | 0,0E+00 | 0,0E+00 | 0,0E+00 | 0,0E+00 | 0,0E+00 | 2099776 | 2100885 | 1110 | - |
| Y11_RS09785 | Y11_20801 | nrdR | transcriptional regulator NrdR | 0,3 | 1,04 | 0,98 | 0,75 | 0,68 | -0,06 | 1,5E-03 | 2,5E-07 | 1,0E-04 | 3,4E-07 | 8,1E-04 | 6,5E-01 | 0,0E+00 | 8,9E-09 | 0,0E+00 | 0,0E+00 | 0,0E+00 | 0,0E+00 | 2101158 | 2101607 | 450 | - |
| Y11_RS09805 | Y11_20841 | secF | protein translocase subunit SecF | -0,28 | -1,01 | -0,98 | -0,73 | -0,7 | 0,03 | 4,5E-02 | 1,0E-05 | 3,1E-04 | 6,6E-04 | 5,4E-03 | 8,6E-01 | 0,0E+00 | 2,5E-07 | 0,0E+00 | 0,0E+00 | 0,0E+00 | 0,0E+00 | 2106007 | 2106975 | 969 | - |
| Y11_RS09810 | Y11_20851 | secD | protein translocase subunit SecD | -0,46 | -1,17 | -0,91 | -0,71 | -0,46 | 0,26 | 1,2E-06 | 1,9E-09 | 2,1E-08 | 1,3E-08 | 8,6E-07 | 1,4E-04 | 0,0E+00 | 1,7E-10 | 0,0E+00 | 0,0E+00 | 0,0E+00 | 0,0E+00 | 2106986 | 2108833 | 1848 | - |
| Y11_RS09815 | Y11_20861 | yajC | preprotein translocase subunit YajC | -0,2 | -1,03 | -0,68 | -0,83 | -0,48 | 0,35 | 9,6E-02 | 1,0E-05 | 1,8E-02 | 3,0E-04 | 8,5E-02 | 1,9E-01 | 0,0E+00 | 2,5E-07 | 0,0E+00 | 0,0E+00 | 0,0E+00 | 0,0E+00 | 2108861 | 2109193 | 333 | - |
| Y11_RS09820 | Y11_20871 | tgt | tRNA guanosine(34) transglycosylase Tgt | 0,79 | 0,2 | 0,38 | -0,59 | -0,41 | 0,18 | 5,9E-07 | 2,7E-02 | 4,5E-03 | 2,9E-05 | 2,5E-03 | 1,3E-01 | 0,0E+00 | 0,0E+00 | 0,0E+00 | 0,0E+00 | 0,0E+00 | 0,0E+00 | 2109306 | 2110442 | 1137 | - |
| Y11_RS09830 | Y11_20901 | | alpha/beta hydrolase | 0,17 | 3,18 | 3,47 | 3 | 3,29 | 0,29 | 6,3E-02 | 1,2E-09 | 5,6E-09 | 3,9E-09 | 1,3E-08 | 9,8E-02 | 0,0E+00 | 1,2E-10 | 3,5E-10 | 2,3E-10 | 1,2E-09 | 0,0E+00 | 2112318 | 2113007 | 690 | + |
| Y11_RS09835 | Y11_20911 | queA | tRNA preQ1(34) S-adenosylmethionine ribosyltransferase-isomerase QueA | 0,48 | 0,44 | -0,12 | -0,03 | -0,59 | -0,56 | 6,5E-04 | 1,2E-04 | 6,4E-02 | 7,5E-01 | 1,3E-04 | 1,8E-05 | 0,0E+00 | 0,0E+00 | 0,0E+00 | 0,0E+00 | 0,0E+00 | 0,0E+00 | 2113116 | 2114204 | 1089 | - |
| Y11_RS09840 | Y11_20931 | | ACP phosphodiesterase | 0,27 | 1,92 | 2,19 | 1,65 | 1,91 | 0,22 | 2,4E-01 | 3,4E-06 | 2,6E-06 | 4,3E-05 | 2,3E-05 | 2,4E-01 | 0,0E+00 | 9,0E-08 | 5,1E-08 | 8,1E-07 | 6,9E-07 | 0,0E+00 | 2114347 | 2114928 | 582 | + |
| Y11_RS09845 | Y11_20941 | | peroxiredoxin C | 0 | 1,21 | 0,6 | 1,21 | 0,61 | -0,61 | 9,7E-01 | 3,7E-05 | 6,5E-03 | 1,2E-04 | 1,3E-02 | 1,4E-02 | 0,0E+00 | 8,5E-07 | 0,0E+00 | 2,2E-06 | 0,0E+00 | 0,0E+00 | 2115153 | 2115755 | 603 | + |
| Y11_RS09860 | Y11_20971 | | SDR family oxidoreductase | 0,02 | 1,78 | 1,72 | 1,76 | 1,7 | -0,06 | 7,5E-01 | 2,4E-10 | 8,0E-07 | 6,9E-09 | 1,6E-06 | 6,8E-01 | 0,0E+00 | 3,9E-11 | 1,9E-08 | 3,8E-10 | 6,4E-08 | 0,0E+00 | 2118346 | 2119026 | 681 | - |
| Y11_RS09870 | Y11_20991 | proY | proline-specific permease ProY | 0,71 | -1,57 | -0,85 | -2,29 | -1,56 | 0,73 | 1,7E-02 | 1,2E-02 | 5,5E-02 | 1,1E-03 | 1,8E-03 | 2,2E-01 | 0,0E+00 | 2,4E-04 | 0,0E+00 | 1,9E-05 | 4,0E-05 | 0,0E+00 | 2121461 | 2122837 | 1377 | - |
| Y11_RS09880 | Y11_21011 | | phosphate ABC transporter substrate-binding protein | 0,05 | 0,27 | 0,23 | 0,22 | 0,18 | -0,04 | 8,7E-01 | 3,6E-01 | 4,5E-01 | 4,9E-01 | 5,8E-01 | 9,0E-01 | 0,0E+00 | 0,0E+00 | 0,0E+00 | 0,0E+00 | 0,0E+00 | 0,0E+00 | 2124596 | 2125573 | 978 | - |
| Y11_RS09885 | Y11_21021 | phoR | phosphate regulon sensor histidine kinase PhoR | 1,31 | 1,97 | 1,65 | 0,66 | 0,33 | -0,32 | 3,1E-05 | 1,6E-07 | 4,2E-07 | 5,0E-03 | 8,0E-02 | 3,9E-02 | 1,5E-06 | 6,1E-09 | 1,1E-08 | 0,0E+00 | 0,0E+00 | 0,0E+00 | 2125595 | 2126911 | 1317 | - |
| Y11_RS09890 | Y11_21031 | phoB | phosphate response regulator transcription factor PhoB | 0,1 | 0,45 | 0,09 | 0,35 | -0,01 | -0,36 | 4,5E-01 | 2,2E-04 | 3,3E-01 | 3,4E-02 | 9,4E-01 | 6,4E-03 | 0,0E+00 | 0,0E+00 | 0,0E+00 | 0,0E+00 | 0,0E+00 | 0,0E+00 | 2126936 | 2127625 | 690 | - |
| Y11_RS09895 | Y11_21041 | sbcD | exonuclease subunit SbcD | 0,41 | 0,37 | 0,66 | -0,04 | 0,25 | 0,29 | 1,3E-02 | 2,4E-02 | 2,1E-03 | 7,6E-01 | 1,4E-01 | 9,5E-02 | 0,0E+00 | 0,0E+00 | 0,0E+00 | 0,0E+00 | 0,0E+00 | 0,0E+00 | 2127895 | 2129133 | 1239 | + |
| Y11_RS09900 | Y11_21051 | | AAA family ATPase | 0,15 | 0,7 | 1,11 | 0,55 | 0,95 | 0,4 | 2,5E-01 | 3,3E-05 | 4,0E-05 | 2,5E-03 | 4,4E-04 | 1,9E-02 | 0,0E+00 | 0,0E+00 | 5,9E-07 | 0,0E+00 | 0,0E+00 | 0,0E+00 | 2129130 | 2132819 | 3690 | + |
| Y11_RS09905 | Y11_21061 | mak | fructokinase | 0,61 | 1 | 1,41 | 0,38 | 0,8 | 0,41 | 7,1E-06 | 5,5E-08 | 2,4E-06 | 3,9E-04 | 2,3E-04 | 9,8E-03 | 0,0E+00 | 0,0E+00 | 4,9E-08 | 0,0E+00 | 0,0E+00 | 0,0E+00 | 2133011 | 2133925 | 915 | - |
| Y11_RS09910 | Y11_21071 | rdgC | recombination-associated protein RdgC | 0,14 | -0,66 | -0,63 | -0,79 | -0,76 | 0,03 | 8,7E-02 | 2,2E-07 | 3,9E-05 | 6,7E-06 | 6,6E-05 | 7,2E-01 | 0,0E+00 | 0,0E+00 | 0,0E+00 | 0,0E+00 | 0,0E+00 | 0,0E+00 | 2134264 | 2135175 | 912 | + |
| Y11_RS09915 | Y11_21081 | ppnP | pyrimidine/purine nucleoside phosphorylase | -0,15 | 0,69 | 2,61 | 0,84 | 2,77 | 1,93 | 4,1E-01 | 1,9E-04 | 2,4E-08 | 1,0E-03 | 2,9E-07 | 1,1E-07 | 0,0E+00 | 0,0E+00 | 1,0E-09 | 0,0E+00 | 1,5E-08 | 3,8E-08 | 2135247 | 2135534 | 288 | - |
| Y11_RS09920 | Y11_21101 | aroL | shikimate kinase AroL | 0,39 | 0,48 | 0,21 | 0,09 | -0,19 | -0,27 | 1,4E-01 | 9,1E-02 | 5,1E-01 | 5,5E-01 | 4,5E-01 | 3,1E-01 | 0,0E+00 | 0,0E+00 | 0,0E+00 | 0,0E+00 | 0,0E+00 | 0,0E+00 | 2136017 | 2136541 | 525 | - |
| Y11_RS09940 | Y11_21161 | | DUF1471 domain-containing protein | 15 | -4,02 | -2,76 | -15 | -15 | 1,29 | 0,0E+00 | 2,1E-07 | 4,3E-04 | 0,0E+00 | 0,0E+00 | 1,8E-02 | 0,0E+00 | 7,8E-09 | 5,3E-06 | 0,0E+00 | 0,0E+00 | 1,0E-03 | 2140607 | 2140876 | 270 | - |
| Y11_RS09945 | Y11_21171 | | class I SAM-dependent methyltransferase | 0,64 | 1,5 | 1,97 | 0,86 | 1,34 | 0,47 | 4,1E-02 | 4,0E-05 | 1,6E-05 | 2,1E-03 | 3,2E-04 | 5,3E-03 | 0,0E+00 | 9,0E-07 | 2,6E-07 | 0,0E+00 | 7,8E-06 | 0,0E+00 | 2141078 | 2141848 | 771 | - |
| Y11_RS09965 | Y11_21221 | | flagellar brake protein | -0,7 | -1,91 | -1,78 | -1,22 | -1,08 | 0,13 | 6,5E-04 | 2,2E-06 | 4,6E-04 | 4,6E-05 | 8,0E-03 | 7,0E-01 | 0,0E+00 | 6,1E-08 | 5,7E-06 | 8,7E-07 | 1,7E-04 | 0,0E+00 | 2152037 | 2152786 | 750 | - |
| Y11_RS09970 | Y11_21231 | | 6-phospho-alpha-glucosidase | -0,21 | -1,06 | -1,33 | -0,85 | -1,12 | -0,27 | 3,9E-01 | 3,7E-03 | 8,0E-05 | 1,3E-02 | 3,7E-04 | 2,6E-01 | 0,0E+00 | 7,3E-05 | 1,1E-06 | 0,0E+00 | 8,8E-06 | 0,0E+00 | 2152990 | 2154312 | 1323 | - |
| Y11_RS09980 | Y11_21251 | | GntR family transcriptional regulator | 0 | 0,59 | 1,79 | 0,59 | 1,79 | 1,2 | 1,0E+00 | 6,9E-03 | 5,7E-07 | 1,9E-02 | 5,9E-06 | 9,4E-05 | 0,0E+00 | 0,0E+00 | 1,4E-08 | 0,0E+00 | 2,0E-07 | 7,9E-06 | 2156168 | 2156818 | 651 | + |
| Y11_RS09985 | Y11_21261 | proA | glutamate-5-semialdehyde dehydrogenase | 0,28 | -1,69 | -1,33 | -1,97 | -1,61 | 0,36 | 1,3E-04 | 7,3E-09 | 1,5E-08 | 1,5E-09 | 2,1E-09 | 1,7E-03 | 0,0E+00 | 5,0E-10 | 7,2E-10 | 1,2E-10 | 2,9E-10 | 0,0E+00 | 2156888 | 2158141 | 1254 | - |
| Y11_RS09990 | Y11_21271 | proB | glutamate 5-kinase | 0,24 | -2,21 | -2,75 | -2,45 | -2,99 | -0,54 | 3,2E-03 | 1,3E-13 | 9,5E-11 | 1,1E-10 | 4,0E-10 | 2,9E-05 | 0,0E+00 | 9,0E-13 | 2,3E-11 | 1,9E-11 | 1,0E-10 | 0,0E+00 | 2158157 | 2159260 | 1104 | - |
| Y11_RS09995 | Y11_21281 | crl | sigma factor-binding protein Crl | 0,15 | -0,78 | -1,81 | -0,93 | -1,96 | -1,03 | 3,9E-01 | 8,6E-03 | 3,1E-06 | 2,4E-03 | 5,8E-07 | 1,1E-03 | 0,0E+00 | 0,0E+00 | 5,9E-08 | 0,0E+00 | 2,7E-08 | 7,5E-05 | 2159416 | 2159817 | 402 | - |
| Y11_RS10000 | Y11_21291 | frsA | esterase FrsA | 0,64 | 0,87 | 0,47 | 0,23 | -0,17 | -0,4 | 2,4E-04 | 1,3E-06 | 2,6E-04 | 4,1E-02 | 1,2E-01 | 1,9E-04 | 0,0E+00 | 0,0E+00 | 0,0E+00 | 0,0E+00 | 0,0E+00 | 0,0E+00 | 2159877 | 2161124 | 1248 | - |
| Y11_RS10005 | Y11_21301 | gpt | xanthine phosphoribosyltransferase | 0,37 | 0,23 | 0,22 | -0,13 | -0,15 | -0,02 | 5,1E-04 | 6,3E-03 | 1,3E-01 | 6,0E-03 | 2,4E-01 | 9,0E-01 | 0,0E+00 | 0,0E+00 | 0,0E+00 | 0,0E+00 | 0,0E+00 | 0,0E+00 | 2161450 | 2161908 | 459 | - |
| Y11_RS10010 | Y11_21321 | pepD | beta-Ala-His dipeptidase | -0,93 | 1,49 | 1,74 | 2,42 | 2,68 | 0,25 | 6,2E-08 | 1,0E-10 | 5,1E-09 | 5,1E-11 | 5,3E-10 | 6,4E-03 | 0,0E+00 | 2,3E-11 | 3,3E-10 | 1,2E-11 | 1,2E-10 | 0,0E+00 | 2162216 | 2163676 | 1461 | + |
| Y11_RS10015 | Y11_21331 | dinB | DNA polymerase IV | -0,21 | -0,57 | -0,02 | -0,36 | 0,24 | 0,55 | 8,4E-03 | 3,8E-03 | 9,4E-01 | 3,2E-02 | 2,9E-01 | 5,4E-02 | 0,0E+00 | 0,0E+00 | 0,0E+00 | 0,0E+00 | 0,0E+00 | 0,0E+00 | 2163750 | 2164808 | 1059 | - |
| Y11_RS10025 | Y11_21351 | | threo-3-hydroxy-L-aspartate ammonia-lyase | -0,14 | 0,05 | 0,7 | 0,19 | 0,85 | 0,65 | 4,5E-02 | 5,7E-01 | 8,5E-06 | 3,5E-02 | 8,7E-07 | 5,6E-05 | 0,0E+00 | 0,0E+00 | 0,0E+00 | 0,0E+00 | 0,0E+00 | 0,0E+00 | 2165303 | 2166271 | 969 | - |
| Y11_RS10030 | Y11_21361 | | PAS domain-containing protein | -0,17 | -0,4 | -1,36 | -0,23 | -1,19 | -0,96 | 2,0E-01 | 6,8E-02 | 9,7E-06 | 2,5E-01 | 1,8E-05 | 1,1E-03 | 0,0E+00 | 0,0E+00 | 1,7E-07 | 0,0E+00 | 5,4E-07 | 0,0E+00 | 2166274 | 2166906 | 633 | - |
| Y11_RS10035 | Y11_21371 | ddlA | D-alanine--D-alanine ligase | 0,94 | -1,11 | -1,7 | -2,05 | -2,63 | -0,58 | 2,5E-07 | 1,1E-07 | 1,3E-08 | 2,4E-09 | 2,9E-07 | 1,2E-04 | 0,0E+00 | 4,3E-09 | 6,5E-10 | 1,7E-10 | 1,6E-10 | 0,0E+00 | 2167158 | 2168258 | 1101 | - |
| Y11_RS10040 | Y11_21381 | | glycerophosphoryl diester phosphodiesterase | 0,64 | 1,57 | 2,24 | 0,93 | 1,6 | 0,67 | 2,8E-02 | 3,0E-08 | 1,4E-07 | 4,2E-03 | 2,6E-04 | 5,4E-04 | 0,0E+00 | 1,5E-09 | 4,3E-09 | 0,0E+00 | 6,5E-06 | 0,0E+00 | 2168668 | 2169618 | 951 | + |
| Y11_RS10050 | Y11_21401 | | FAD:protein FMN transferase | 0 | 0,3 | 0,6 | 0,29 | 0,6 | 0,31 | 9,7E-01 | 2,5E-02 | 1,3E-04 | 4,2E-02 | 4,4E-04 | 2,6E-02 | 0,0E+00 | 0,0E+00 | 0,0E+00 | 0,0E+00 | 0,0E+00 | 0,0E+00 | 2169998 | 2171020 | 1023 | - |
| Y11_RS10055 | Y11_21411 | nqrF | NADH:ubiquinone reductase (Na(+)-transporting) subunit F | 0,04 | -1,3 | -0,78 | -1,34 | -0,82 | 0,52 | 7,6E-01 | 6,7E-06 | 4,9E-04 | 3,6E-07 | 5,2E-05 | 1,4E-03 | 0,0E+00 | 1,7E-07 | 0,0E+00 | 1,0E-08 | 0,0E+00 | 0,0E+00 | 2171060 | 2172283 | 1224 | - |
| Y11_RS10070 | Y11_21441 | | Na(+)-translocating NADH-quinone reductase subunit C | -0,03 | -0,57 | -0,05 | -0,55 | -0,03 | 0,52 | 7,2E-01 | 1,0E-03 | 6,6E-01 | 6,5E-04 | 8,0E-01 | 4,6E-03 | 0,0E+00 | 0,0E+00 | 0,0E+00 | 0,0E+00 | 0,0E+00 | 0,0E+00 | 2173524 | 2174324 | 801 | - |
| Y11_RS10080 | Y11_21461 | | Na(+)-translocating NADH-quinone reductase subunit A | 0,16 | -0,65 | -0,4 | -0,81 | -0,56 | 0,25 | 3,6E-01 | 9,6E-03 | 5,7E-02 | 1,4E-04 | 5,3E-04 | 1,2E-01 | 0,0E+00 | 0,0E+00 | 0,0E+00 | 0,0E+00 | 0,0E+00 | 0,0E+00 | 2175559 | 2176902 | 1344 | - |
| Y11_RS10090 | Y11_21481 | | class II glutamine amidotransferase | 0,14 | 0,72 | 1,08 | 0,58 | 0,95 | 0,37 | 1,2E-01 | 2,5E-04 | 4,1E-05 | 4,7E-04 | 5,9E-05 | 3,9E-02 | 0,0E+00 | 0,0E+00 | 6,0E-07 | 0,0E+00 | 0,0E+00 | 0,0E+00 | 2178122 | 2178886 | 765 | - |
| Y11_RS10095 | Y11_21491 | lpcA | D-sedoheptulose 7-phosphate isomerase | -0,33 | 0,87 | 0,85 | 1,2 | 1,18 | -0,02 | 1,5E-05 | 6,8E-08 | 1,5E-07 | 2,8E-09 | 6,8E-09 | 7,5E-01 | 0,0E+00 | 0,0E+00 | 0,0E+00 | 1,8E-10 | 7,1E-10 | 0,0E+00 | 2178988 | 2179569 | 582 | - |
| Y11_RS10100 | Y11_21501 | fadE | acyl-CoA dehydrogenase FadE | 0,8 | -1,5 | 0,35 | -2,3 | -0,45 | 1,85 | 3,8E-07 | 1,2E-08 | 1,7E-04 | 5,4E-10 | 3,7E-05 | 2,9E-09 | 0,0E+00 | 7,0E-10 | 0,0E+00 | 5,5E-11 | 0,0E+00 | 3,2E-09 | 2179824 | 2182271 | 2448 | + |
| Y11_RS10130 | Y11_21581 | mtnA | S-methyl-5-thioribose-1-phosphate isomerase | 0,33 | 0,41 | 0,72 | 0,08 | 0,39 | 0,31 | 9,2E-03 | 3,1E-04 | 4,2E-02 | 4,0E-01 | 2,3E-01 | 3,3E-01 | 0,0E+00 | 0,0E+00 | 0,0E+00 | 0,0E+00 | 0,0E+00 | 0,0E+00 | 2195063 | 2196103 | 1041 | + |
| Y11_RS10140 | Y11_21601 | mtnC | acireductone synthase | 0,22 | 1,03 | 1,05 | 0,81 | 0,83 | 0,02 | 2,1E-01 | 4,4E-04 | 5,1E-05 | 1,6E-03 | 1,7E-04 | 9,2E-01 | 0,0E+00 | 9,1E-06 | 7,3E-07 | 0,0E+00 | 0,0E+00 | 0,0E+00 | 2196784 | 2197473 | 690 | - |
| Y11_RS10145 | Y11_21611 | | methylthioribulose 1-phosphate dehydratase | -0,05 | -0,03 | -0,21 | 0,02 | -0,16 | -0,18 | 7,8E-01 | 8,6E-01 | 2,8E-01 | 9,4E-01 | 5,2E-01 | 4,8E-01 | 0,0E+00 | 0,0E+00 | 0,0E+00 | 0,0E+00 | 0,0E+00 | 0,0E+00 | 2197470 | 2198102 | 633 | - |
| Y11_RS10150 | Y11_21621 | | pyridoxal phosphate-dependent aminotransferase | -0,18 | 0,43 | 0,85 | 0,62 | 1,03 | 0,42 | 7,3E-02 | 1,0E-03 | 2,8E-05 | 1,4E-05 | 1,5E-06 | 8,4E-04 | 0,0E+00 | 0,0E+00 | 0,0E+00 | 0,0E+00 | 6,1E-08 | 0,0E+00 | 2198319 | 2199479 | 1161 | + |
| Y11_RS10165 | Y11_21651 | | alanine--glyoxylate aminotransferase family protein | 0,05 | 0,17 | 0,3 | 0,12 | 0,25 | 0,13 | 5,7E-01 | 1,3E-01 | 2,3E-02 | 2,1E-01 | 2,8E-02 | 2,9E-01 | 0,0E+00 | 0,0E+00 | 0,0E+00 | 0,0E+00 | 0,0E+00 | 0,0E+00 | 2201630 | 2202880 | 1251 | - |
| Y11_RS10185 | Y11_21691 | | transporter substrate-binding domain-containing protein | 0,47 | 1,38 | 1,89 | 0,91 | 0 | 0,5 | 8,6E-03 | 4,6E-03 | 2,3E-05 | 3,2E-02 | 0,0E+00 | 2,3E-01 | 0,0E+00 | 8,9E-05 | 3,6E-07 | 0,0E+00 | 0,0E+00 | 0,0E+00 | 2204981 | 2205772 | 792 | - |
| Y11_RS10190 | Y11_21701 | | MurR/RpiR family transcriptional regulator | -0,13 | 0,18 | -0,14 | 0,31 | -0,01 | -0,32 | 1,4E-01 | 8,4E-02 | 4,2E-01 | 3,9E-04 | 9,7E-01 | 6,6E-02 | 0,0E+00 | 0,0E+00 | 0,0E+00 | 0,0E+00 | 0,0E+00 | 0,0E+00 | 2206095 | 2206934 | 840 | - |
| Y11_RS10215 | Y11_21761 | | MFS transporter | -1,97 | 0 | 0 | 15 | 1,82 | 0 | 3,1E-09 | 0,0E+00 | 0,0E+00 | 0,0E+00 | 4,7E-04 | 0,0E+00 | 6,1E-10 | 0,0E+00 | 0,0E+00 | 0,0E+00 | 1,1E-05 | 0,0E+00 | 2211620 | 2212822 | 1203 | + |
| Y11_RS10230 | Y11_21791 | mprA | transcriptional repressor MprA | -0,09 | 0,12 | -0,61 | 0,21 | -0,52 | -0,72 | 3,6E-01 | 2,0E-01 | 9,1E-05 | 1,1E-02 | 2,7E-05 | 1,0E-06 | 0,0E+00 | 0,0E+00 | 0,0E+00 | 0,0E+00 | 0,0E+00 | 0,0E+00 | 2214359 | 2214871 | 513 | + |

| | | | | | | | | | | | | | | | | | | | | | | | | | |
|---|---|---|---|---|---|---|---|---|---|---|---|---|---|---|---|---|---|---|---|---|---|---|---|---|---|
| Y11_RS10240 | Y11_21811 | emrA | multidrug efflux MFS transporter periplasmic adaptor subunit EmrA | 0 | -0,33 | -2,77 | -0,39 | -2,86 | -2,45 | 0,0E+00 | 6,5E-02 | 1,8E-08 | 1,9E-02 | 3,7E-09 | 3,8E-08 | 0,0E+00 | 0,0E+00 | 8,1E-10 | 0,0E+00 | 4,4E-10 | 1,7E-08 | 2216401 | 2217573 | 1173 | + |
| Y11_RS10260 | Y11_21841 | | tRNA/rRNA methyltransferase | 0,24 | 0,94 | 0,75 | 0,7 | 0,5 | -0,2 | 1,2E-03 | 3,1E-07 | 3,8E-04 | 1,8E-06 | 4,0E-03 | 1,7E-01 | 0,0E+00 | 0,0E+00 | 0,0E+00 | 0,0E+00 | 0,0E+00 | 0,0E+00 | 2220753 | 2221859 | 1107 | - |
| Y11_RS10265 | Y11_21851 | trxC | thioredoxin TrxC | 1,29 | 2,52 | 2,02 | 1,23 | 0,73 | -0,51 | 3,4E-07 | 3,3E-09 | 1,7E-07 | 7,9E-07 | 3,0E-04 | 3,7E-03 | 2,5E-08 | 2,6E-10 | 5,1E-09 | 2,0E-08 | 0,0E+00 | 0,0E+00 | 2222080 | 2222520 | 441 | + |
| Y11_RS10275 | Y11_21871 | | bifunctional acetate--CoA ligase family protein/GNAT family N-acetyltransferase | -0,86 | 1,37 | 2,53 | 2,23 | 3,39 | 1,16 | 5,6E-09 | 1,2E-07 | 4,7E-10 | 2,7E-09 | 4,7E-11 | 3,2E-06 | 0,0E+00 | 4,9E-09 | 6,2E-11 | 1,8E-10 | 2,7E-11 | 4,5E-07 | 2223477 | 2226119 | 2643 | + |
| Y11_RS10280 | Y11_21881 | pssA | CDP-diacylglycerol--serine O-phosphatidyltransferase | -0,2 | -0,54 | -0,01 | -0,34 | 0,19 | 0,53 | 2,1E-03 | 1,5E-05 | 9,6E-01 | 6,7E-04 | 4,3E-01 | 5,3E-02 | 0,0E+00 | 0,0E+00 | 0,0E+00 | 0,0E+00 | 0,0E+00 | 0,0E+00 | 2226297 | 2227652 | 1356 | + |
| Y11_RS10310 | Y11_21901 | raiA | ribosome-associated translation inhibitor RaiA | -0,56 | 1,66 | 3,19 | 2,23 | 3,75 | 1,52 | 3,5E-04 | 1,8E-06 | 2,9E-05 | 1,3E-07 | 8,4E-06 | 4,3E-03 | 0,0E+00 | 5,1E-08 | 4,5E-07 | 4,4E-09 | 2,8E-07 | 2,6E-04 | 2234459 | 2234821 | 363 | - |
| Y11_RS10315 | Y11_21911 | bamD | outer membrane protein assembly factor BamD | -0,11 | -0,52 | -0,24 | -0,41 | -0,13 | 0,28 | 9,6E-02 | 4,1E-06 | 9,1E-03 | 1,7E-05 | 9,1E-02 | 1,7E-03 | 0,0E+00 | 0,0E+00 | 0,0E+00 | 0,0E+00 | 0,0E+00 | 0,0E+00 | 2235189 | 2235920 | 732 | - |
| Y11_RS10320 | Y11_21921 | rluD | 23S rRNA pseudouridine(1911/1915/1917) synthase RluD | -0,17 | -0,73 | -1,09 | -0,56 | -0,92 | -0,36 | 4,3E-02 | 5,4E-06 | 1,6E-06 | 4,2E-06 | 1,6E-06 | 1,1E-03 | 0,0E+00 | 0,0E+00 | 3,4E-08 | 0,0E+00 | 0,0E+00 | 0,0E+00 | 2236056 | 2237033 | 978 | + |
| Y11_RS10330 | Y11_21941 | clpB | ATP-dependent chaperone ClpB | -0,14 | 0,44 | 0,29 | 0,58 | 0,43 | -0,15 | 6,7E-02 | 1,1E-04 | 3,0E-03 | 4,7E-05 | 6,1E-04 | 8,1E-02 | 0,0E+00 | 0,0E+00 | 0,0E+00 | 0,0E+00 | 0,0E+00 | 0,0E+00 | 2237895 | 2240468 | 2574 | + |
| Y11_RS10345 | Y11_21981 | pheA | bifunctional chorismate mutase/prephenate dehydratase | 0,27 | 0,35 | 0,58 | 0,08 | 0,31 | 0,23 | 1,3E-03 | 9,5E-06 | 6,2E-05 | 2,0E-01 | 8,5E-03 | 2,1E-02 | 0,0E+00 | 0,0E+00 | 0,0E+00 | 0,0E+00 | 0,0E+00 | 0,0E+00 | 2242153 | 2243310 | 1158 | + |
| Y11_RS10350 | Y11_21991 | tyrA | bifunctional chorismate mutase/prephenate dehydrogenase | 0,34 | 0,4 | 0,62 | 0,05 | 0,28 | 0,23 | 5,7E-03 | 6,9E-03 | 2,0E-03 | 6,3E-01 | 7,1E-02 | 1,6E-01 | 0,0E+00 | 0,0E+00 | 0,0E+00 | 0,0E+00 | 0,0E+00 | 0,0E+00 | 2243590 | 2244711 | 1122 | - |
| Y11_RS10355 | Y11_22001 | | 3-deoxy-7-phosphoheptulonate synthase | -1,01 | 0,29 | 0,46 | 1,3 | 1,47 | 0,17 | 2,7E-08 | 2,9E-05 | 5,7E-03 | 1,3E-09 | 2,5E-06 | 2,1E-01 | 2,9E-09 | 0,0E+00 | 0,0E+00 | 1,1E-10 | 9,4E-08 | 0,0E+00 | 2244714 | 2245793 | 1080 | - |
| Y11_RS10375 | Y11_22041 | rplS | 50S ribosomal protein L19 | 0,25 | -0,56 | -0,93 | -0,81 | -1,18 | -0,37 | 3,9E-03 | 4,6E-05 | 7,8E-07 | 1,2E-05 | 5,6E-07 | 3,3E-03 | 0,0E+00 | 0,0E+00 | 0,0E+00 | 0,0E+00 | 2,6E-08 | 0,0E+00 | 2249643 | 2249999 | 357 | - |
| Y11_RS10380 | Y11_22051 | trmD | tRNA (guanosine(37)-N1)-methyltransferase TrmD | -0,48 | 0 | -0,8 | 0,48 | -0,32 | -0,8 | 3,9E-03 | 9,6E-01 | 1,8E-03 | 1,1E-03 | 1,1E-01 | 1,0E-03 | 0,0E+00 | 0,0E+00 | 0,0E+00 | 0,0E+00 | 0,0E+00 | 0,0E+00 | 2250080 | 2250820 | 741 | - |
| Y11_RS10385 | Y11_22061 | rimM | ribosome maturation factor RimM | -0,35 | -1,8 | -2,75 | -1,46 | -2,4 | -0,95 | 2,3E-03 | 1,1E-07 | 1,4E-09 | 1,7E-06 | 1,6E-08 | 6,5E-05 | 0,0E+00 | 4,4E-09 | 1,3E-10 | 4,0E-08 | 1,4E-09 | 0,0E+00 | 2250859 | 2251407 | 549 | - |
| Y11_RS10390 | Y11_22071 | rpsP | 30S ribosomal protein S16 | -0,07 | 0,03 | -0,44 | 0,09 | -0,37 | -0,47 | 6,3E-01 | 8,1E-01 | 1,7E-01 | 5,4E-01 | 2,7E-01 | 1,6E-01 | 0,0E+00 | 0,0E+00 | 0,0E+00 | 0,0E+00 | 0,0E+00 | 0,0E+00 | 2251429 | 2251677 | 249 | - |
| Y11_RS10395 | Y11_22081 | ffh | signal recognition particle protein | -0,48 | 1,07 | 0,8 | 1,54 | 1,28 | -0,27 | 3,1E-03 | 4,6E-07 | 5,1E-06 | 1,9E-07 | 9,2E-07 | 7,9E-05 | 0,0E+00 | 1,5E-08 | 0,0E+00 | 5,7E-09 | 3,9E-08 | 0,0E+00 | 2251905 | 2253266 | 1362 | - |
| Y11_RS10405 | Y11_22101 | | HlyC/CorC family transporter | 0,19 | 1,02 | 1,25 | 0,83 | 1,06 | 0,23 | 4,5E-01 | 1,7E-03 | 1,7E-03 | 7,3E-03 | 5,3E-03 | 4,0E-01 | 0,0E+00 | 3,5E-05 | 2,0E-05 | 0,0E+00 | 1,1E-04 | 0,0E+00 | 2254296 | 2255573 | 1278 | + |
| Y11_RS10410 | Y11_22111 | luxS | S-ribosylhomocysteine lyase | -0,04 | 0,16 | 0,38 | 0,2 | 0,42 | 0,22 | 6,0E-01 | 3,4E-02 | 5,6E-02 | 1,8E-02 | 4,0E-02 | 2,3E-01 | 0,0E+00 | 0,0E+00 | 0,0E+00 | 0,0E+00 | 0,0E+00 | 0,0E+00 | 2255843 | 2256358 | 516 | - |
| Y11_RS10415 | Y11_22121 | gshA | glutamate--cysteine ligase | -0,56 | -0,27 | -0,21 | 0,29 | 0,35 | 0,06 | 1,6E-06 | 1,4E-04 | 2,3E-02 | 1,7E-05 | 1,3E-03 | 4,2E-01 | 0,0E+00 | 0,0E+00 | 0,0E+00 | 0,0E+00 | 0,0E+00 | 0,0E+00 | 2256525 | 2258123 | 1599 | - |
| Y11_RS10425 | Y11_22141 | yqaB | fructose-1-phosphate/6-phosphogluconate phosphatase | 0,43 | 0,51 | 0,78 | 0,07 | 0,34 | 0,27 | 1,6E-02 | 9,8E-03 | 5,4E-04 | 4,9E-01 | 3,3E-03 | 2,2E-02 | 0,0E+00 | 0,0E+00 | 0,0E+00 | 0,0E+00 | 0,0E+00 | 0,0E+00 | 2258592 | 2259158 | 567 | - |
| Y11_RS10445 | Y11_22151 | csrA | carbon storage regulator CsrA | -0,4 | -0,62 | -0,39 | -0,22 | 0 | 0,22 | 5,6E-02 | 7,3E-03 | 1,5E-01 | 2,1E-01 | 9,9E-01 | 3,7E-01 | 0,0E+00 | 0,0E+00 | 0,0E+00 | 0,0E+00 | 0,0E+00 | 0,0E+00 | 2260082 | 2260267 | 186 | - |
| Y11_RS10450 | Y11_22161 | alaS | alanine--tRNA ligase | 0,59 | -0,29 | -0,28 | -0,88 | -0,87 | 0,01 | 8,6E-08 | 1,6E-05 | 1,2E-02 | 7,5E-09 | 9,4E-06 | 8,7E-01 | 0,0E+00 | 0,0E+00 | 0,0E+00 | 0,0E+00 | 0,0E+00 | 0,0E+00 | 2260521 | 2263148 | 2628 | - |
| Y11_RS10460 | Y11_22181 | recA | recombinase RecA | -0,11 | -0,96 | -0,5 | -0,85 | -0,39 | 0,46 | 4,2E-02 | 9,8E-10 | 2,0E-06 | 2,5E-08 | 4,5E-05 | 1,1E-06 | 0,0E+00 | 0,0E+00 | 0,0E+00 | 0,0E+00 | 0,0E+00 | 0,0E+00 | 2264103 | 2265167 | 1065 | - |
| Y11_RS10465 | Y11_22191 | pncC | nicotinamide-nucleotide amidase | -0,68 | -0,23 | 0,49 | 0,45 | 1,17 | 0,72 | 4,3E-08 | 4,1E-04 | 7,1E-04 | 7,8E-06 | 1,6E-06 | 6,9E-05 | 0,0E+00 | 0,0E+00 | 0,0E+00 | 0,0E+00 | 6,4E-08 | 0,0E+00 | 2265279 | 2265767 | 489 | - |
| Y11_RS10500 | Y11_22271 | | fumarate hydratase | -0,43 | 2,31 | 3,46 | 2,74 | 3,89 | 1,16 | 1,1E-04 | 5,9E-08 | 3,5E-10 | 3,3E-08 | 4,5E-10 | 6,2E-05 | 0,0E+00 | 2,6E-09 | 4,9E-11 | 1,3E-09 | 1,1E-10 | 5,4E-06 | 2274108 | 2275751 | 1644 | + |
| Y11_RS10530 | Y11_22331 | | acetamidase/formamidase family protein | 0 | -0,87 | 0,29 | -15 | -15 | 1,15 | 0,0E+00 | 7,1E-05 | 2,4E-01 | 0,0E+00 | 0,0E+00 | 2,0E-03 | 0,0E+00 | 0,0E+00 | 0,0E+00 | 0,0E+00 | 0,0E+00 | 1,3E-04 | 2280966 | 2281955 | 990 | + |
| Y11_RS10560 | Y11_22391 | | mannose/fructose/sorbose PTS transporter subunit IIB | 0,75 | 0,01 | 0,52 | -0,74 | 0 | 0,54 | 1,0E-01 | 9,8E-01 | 2,3E-01 | 3,5E-03 | 0,0E+00 | 1,6E-02 | 0,0E+00 | 0,0E+00 | 0,0E+00 | 0,0E+00 | 0,0E+00 | 0,0E+00 | 2286983 | 2287474 | 492 | + |
| Y11_RS10580 | Y11_22431 | | ArsR family transcriptional regulator | -0,03 | 0,97 | 1,03 | 1 | 1,06 | 0,06 | 6,3E-01 | 3,0E-05 | 1,5E-03 | 2,8E-05 | 1,3E-03 | 8,0E-01 | 0,0E+00 | 0,0E+00 | 1,8E-05 | 0,0E+00 | 3,0E-05 | 0,0E+00 | 2290634 | 2290981 | 348 | - |
| Y11_RS10585 | Y11_22441 | fusA | elongation factor G | 0,3 | 0,02 | 0,55 | -0,28 | 0,25 | 0,53 | 8,4E-03 | 8,3E-01 | 1,2E-04 | 3,4E-02 | 4,0E-02 | 9,9E-04 | 0,0E+00 | 0,0E+00 | 0,0E+00 | 0,0E+00 | 0,0E+00 | 0,0E+00 | 2291326 | 2293434 | 2109 | + |
| Y11_RS10600 | Y11_22481 | | aminotransferase class I/II-fold pyridoxal phosphate-dependent enzyme | 2,34 | 0,47 | 1,69 | -1,87 | -0,66 | 1,22 | 1,7E-07 | 1,5E-02 | 1,4E-06 | 1,9E-06 | 1,2E-03 | 3,5E-05 | 1,4E-08 | 0,0E+00 | 3,1E-08 | 4,5E-08 | 0,0E+00 | 3,3E-06 | 2295918 | 2297081 | 1164 | + |
| Y11_RS10685 | Y11_22641 | | fimbria/pilus periplasmic chaperone | 0,15 | 0,47 | 0,14 | 0,32 | -0,01 | -0,33 | 5,2E-02 | 3,3E-05 | 2,3E-01 | 2,7E-04 | 9,1E-01 | 9,0E-03 | 0,0E+00 | 0,0E+00 | 0,0E+00 | 0,0E+00 | 0,0E+00 | 0,0E+00 | 2313641 | 2314387 | 747 | - |
| Y11_RS10715 | Y11_22731 | mutS | DNA mismatch repair protein MutS | 0,17 | -0,15 | -0,21 | -0,32 | -0,38 | -0,06 | 7,3E-03 | 2,6E-02 | 8,1E-03 | 1,2E-03 | 6,0E-04 | 4,8E-01 | 0,0E+00 | 0,0E+00 | 0,0E+00 | 0,0E+00 | 0,0E+00 | 0,0E+00 | 2321370 | 2323925 | 2556 | + |
| Y11_RS10720 | Y11_22741 | rpoS | RNA polymerase sigma factor RpoS | -0,55 | 2,67 | 2,83 | 3,22 | 3,38 | 0,16 | 6,5E-06 | 3,0E-09 | 7,4E-10 | 1,2E-09 | 3,9E-10 | 2,2E-01 | 0,0E+00 | 2,4E-10 | 8,3E-11 | 1,0E-10 | 9,9E-11 | 0,0E+00 | 2324013 | 2325011 | 999 | - |
| Y11_RS10725 | Y11_22751 | nlpD | murein hydrolase activator NlpD | -0,63 | 0,64 | 0,3 | 1,27 | 0,93 | -0,34 | 1,4E-05 | 9,6E-06 | 2,7E-02 | 9,2E-08 | 3,7E-05 | 1,6E-02 | 0,0E+00 | 0,0E+00 | 0,0E+00 | 3,1E-09 | 0,0E+00 | 0,0E+00 | 2325066 | 2326031 | 966 | - |
| Y11_RS10730 | Y11_22761 | | protein-L-isoaspartate(D-aspartate) O-methyltransferase | 0,4 | 0,9 | 0,41 | 0,5 | 0,01 | -0,49 | 1,9E-03 | 2,6E-06 | 2,7E-02 | 9,8E-05 | 9,5E-01 | 8,3E-03 | 0,0E+00 | 0,0E+00 | 0,0E+00 | 0,0E+00 | 0,0E+00 | 0,0E+00 | 2326420 | 2327046 | 627 | - |
| Y11_RS10735 | Y11_22771 | surE | 5'/3'-nucleotidase SurE | 1,01 | 1,86 | 1,05 | 0,84 | 0,03 | -0,81 | 2,3E-05 | 3,6E-08 | 4,1E-04 | 2,3E-05 | 8,6E-01 | 1,4E-03 | 1,1E-06 | 1,7E-09 | 5,1E-06 | 0,0E+00 | 0,0E+00 | 0,0E+00 | 2327068 | 2327820 | 753 | - |
| Y11_RS10740 | Y11_22781 | truD | tRNA pseudouridine(13) synthase TruD | 0,31 | -0,59 | -0,84 | -0,9 | -1,14 | -0,24 | 7,1E-02 | 4,7E-06 | 1,8E-04 | 3,0E-04 | 2,9E-04 | 9,6E-02 | 0,0E+00 | 0,0E+00 | 0,0E+00 | 0,0E+00 | 7,2E-06 | 0,0E+00 | 2327798 | 2328847 | 1050 | - |
| Y11_RS10745 | Y11_22791 | ispF | 2-C-methyl-D-erythritol 2,4-cyclodiphosphate synthase | 0,39 | 0,68 | 0,48 | 0,29 | 0,09 | -0,2 | 7,9E-04 | 6,8E-07 | 1,2E-04 | 2,0E-03 | 2,8E-01 | 9,9E-03 | 0,0E+00 | 0,0E+00 | 0,0E+00 | 0,0E+00 | 0,0E+00 | 0,0E+00 | 2328858 | 2329346 | 489 | - |
| Y11_RS10750 | Y11_22801 | ispD | 2-C-methyl-D-erythritol 4-phosphate cytidylyltransferase | 15 | -0,51 | -1,81 | -15 | -15 | -1,3 | 0,0E+00 | 1,5E-01 | 4,6E-04 | 0,0E+00 | 0,0E+00 | 4,5E-03 | 0,0E+00 | 0,0E+00 | 5,7E-06 | 0,0E+00 | 0,0E+00 | 2,7E-04 | 2329361 | 2330005 | 645 | - |
| Y11_RS10755 | Y11_22811 | ftsB | cell division protein FtsB | 0 | -0,27 | -0,6 | -1,44 | -1,8 | -0,32 | 0,0E+00 | 1,3E-01 | 1,7E-02 | 3,2E-05 | 3,6E-05 | 2,2E-01 | 0,0E+00 | 0,0E+00 | 0,0E+00 | 6,1E-07 | 1,0E-06 | 0,0E+00 | 2330123 | 2330446 | 324 | - |
| Y11_RS10765 | Y11_22831 | cysC | adenylyl-sulfate kinase | -0,28 | 0,46 | 1,35 | 0,75 | 1,63 | 0,89 | 5,0E-03 | 5,8E-05 | 2,9E-06 | 1,4E-06 | 6,5E-07 | 3,6E-05 | 0,0E+00 | 0,0E+00 | 5,8E-08 | 0,0E+00 | 2,9E-08 | 0,0E+00 | 2331324 | 2331950 | 627 | - |
| Y11_RS10770 | Y11_22841 | cysN | sulfate adenylyltransferase subunit CysN | 0,35 | -0,77 | 0,13 | -1,12 | -0,22 | 0,9 | 4,3E-03 | 2,0E-05 | 2,6E-01 | 2,4E-07 | 4,5E-02 | 9,3E-06 | 0,0E+00 | 0,0E+00 | 0,0E+00 | 7,0E-09 | 0,0E+00 | 0,0E+00 | 2331976 | 2333475 | 1500 | - |
| Y11_RS10775 | Y11_22851 | cysD | sulfate adenylyltransferase subunit CysD | 0,19 | -0,23 | 0,52 | -0,42 | 0,32 | 0,74 | 6,0E-02 | 4,8E-02 | 6,0E-03 | 2,8E-03 | 5,1E-02 | 9,4E-04 | 0,0E+00 | 0,0E+00 | 0,0E+00 | 0,0E+00 | 0,0E+00 | 0,0E+00 | 2333486 | 2334394 | 909 | - |
| Y11_RS10780 | Y11_22861 | cysG | siroheme synthase CysG | -0,1 | 0,11 | 0,97 | 0,2 | 1,07 | 0,86 | 6,8E-02 | 9,5E-02 | 4,1E-07 | 9,0E-03 | 2,6E-07 | 3,1E-06 | 0,0E+00 | 0,0E+00 | 0,0E+00 | 0,0E+00 | 1,4E-08 | 0,0E+00 | 2334405 | 2335823 | 1419 | - |
| Y11_RS10785 | Y11_22871 | | aminopeptidase | 0,67 | 2,46 | 1,95 | 1,78 | 1,27 | -0,52 | 3,1E-02 | 5,1E-07 | 1,0E-06 | 5,7E-05 | 3,5E-04 | 5,1E-04 | 0,0E+00 | 1,6E-08 | 2,4E-08 | 1,1E-06 | 8,5E-06 | 0,0E+00 | 2336116 | 2337150 | 1035 | + |
| Y11_RS10795 | Y11_22891 | | type II toxin-antitoxin system HicB family antitoxin | -0,09 | -0,65 | -0,23 | -0,56 | -0,14 | 0,42 | 3,4E-01 | 2,6E-06 | 2,4E-02 | 1,6E-04 | 2,2E-01 | 5,5E-04 | 0,0E+00 | 0,0E+00 | 0,0E+00 | 0,0E+00 | 0,0E+00 | 0,0E+00 | 2337665 | 2338072 | 408 | + |
| Y11_RS10800 | Y11_22901 | | phosphoadenylyl-sulfate reductase | 0,22 | 0,76 | 1,6 | 0,53 | 1,38 | 0,84 | 9,4E-04 | 1,0E-07 | 2,9E-08 | 1,4E-06 | 9,2E-08 | 3,8E-06 | 0,0E+00 | 0,0E+00 | 1,2E-09 | 0,0E+00 | 5,8E-09 | 0,0E+00 | 2338185 | 2338919 | 735 | - |
| Y11_RS10805 | Y11_22911 | cysI | assimilatory sulfite reductase (NADPH) hemoprotein subunit | 1,01 | -1,51 | -1,45 | -2,52 | -2,46 | 0,06 | 2,4E-07 | 2,0E-08 | 5,0E-06 | 2,1E-10 | 7,6E-08 | 6,6E-01 | 1,9E-08 | 1,1E-09 | 9,1E-08 | 2,7E-11 | 5,0E-09 | 0,0E+00 | 2338916 | 2340664 | 1749 | - |
| Y11_RS10810 | Y11_22921 | cysJ | NADPH-dependent assimilatory sulfite reductase flavoprotein subunit | 15 | -3,55 | -3,86 | -15 | -15 | -0,31 | 0,0E+00 | 2,1E-08 | 7,6E-10 | 0,0E+00 | 0,0E+00 | 1,3E-01 | 0,0E+00 | 1,1E-09 | 8,3E-11 | 0,0E+00 | 0,0E+00 | 0,0E+00 | 2340664 | 2342463 | 1800 | - |
| Y11_RS10815 | Y11_22931 | queD | 6-carboxytetrahydropterin synthase QueD | -0,73 | -1,45 | -0,65 | -0,71 | 0,08 | 0,79 | 9,9E-04 | 7,3E-06 | 1,3E-03 | 3,7E-03 | 6,6E-01 | 1,5E-03 | 0,0E+00 | 1,8E-07 | 0,0E+00 | 0,0E+00 | 0,0E+00 | 0,0E+00 | 2342831 | 2343193 | 363 | + |
| Y11_RS10835 | Y11_22971 | queE | 7-carboxy-7-deazaguanine synthase QueE | -0,68 | 0,8 | 0,48 | 1,48 | 1,16 | -0,32 | 1,1E-03 | 1,5E-06 | 3,0E-03 | 4,9E-06 | 1,2E-04 | 2,4E-02 | 0,0E+00 | 0,0E+00 | 0,0E+00 | 1,1E-07 | 3,1E-06 | 0,0E+00 | 2345576 | 2346247 | 672 | - |
| Y11_RS10840 | Y11_22981 | | superoxide dismutase family protein | 0,7 | 3,08 | 3,5 | 2,38 | 2,8 | 0,42 | 1,1E-03 | 3,4E-09 | 1,6E-07 | 4,0E-08 | 1,0E-06 | 6,3E-02 | 0,0E+00 | 2,7E-10 | 4,8E-09 | 1,6E-09 | 4,3E-08 | 0,0E+00 | 2346351 | 2346875 | 525 | - |
| Y11_RS10865 | Y11_23031 | serA | phosphoglycerate dehydrogenase | 1,15 | -0,92 | -0,31 | -2,07 | -1,45 | 0,62 | 2,4E-07 | 8,6E-06 | 5,2E-04 | 3,7E-08 | 3,2E-08 | 1,5E-04 | 1,9E-08 | 0,0E+00 | 0,0E+00 | 1,5E-09 | 2,5E-09 | 0,0E+00 | 2349556 | 2350797 | 1242 | - |
| Y11_RS10870 | Y11_23041 | rpiA | ribose-5-phosphate isomerase RpiA | -0,23 | 0,43 | 1,66 | 0,66 | 1,89 | 1,23 | 1,7E-02 | 2,0E-04 | 3,8E-07 | 1,3E-05 | 1,6E-07 | 2,6E-06 | 0,0E+00 | 0,0E+00 | 1,0E-08 | 0,0E+00 | 9,2E-09 | 3,8E-07 | 2351112 | 2351774 | 663 | - |
| Y11_RS10875 | Y11_23051 | | LysR family transcriptional regulator ArgP | 0,01 | 0,44 | 0,79 | 0,43 | 0,78 | 0,35 | 9,3E-01 | 5,2E-04 | 1,6E-05 | 1,1E-03 | 3,3E-05 | 8,8E-03 | 0,0E+00 | 0,0E+00 | 0,0E+00 | 0,0E+00 | 0,0E+00 | 0,0E+00 | 2351994 | 2352887 | 894 | + |
| Y11_RS10880 | Y11_23061 | | oxidative stress defense protein | 0,01 | 0,42 | 1,33 | 0,41 | 1,32 | 0,91 | 8,6E-01 | 9,7E-05 | 5,6E-09 | 4,8E-06 | 3,1E-11 | 6,4E-09 | 0,0E+00 | 0,0E+00 | 3,5E-10 | 0,0E+00 | 2,0E-11 | 0,0E+00 | 2352889 | 2353644 | 756 | - |
| Y11_RS10895 | Y11_23111 | mscS | small-conductance mechanosensitive channel MscS | 0,81 | 0,66 | 0,64 | -0,14 | -0,17 | -0,02 | 4,6E-03 | 1,4E-02 | 1,6E-02 | 5,6E-01 | 5,1E-01 | 9,3E-01 | 0,0E+00 | 0,0E+00 | 0,0E+00 | 0,0E+00 | 0,0E+00 | 0,0E+00 | 2356257 | 2357126 | 870 | - |
| Y11_RS10905 | Y11_23131 | fbaA | class II fructose-bisphosphate aldolase | 0,06 | 1,71 | 1,53 | 1,64 | 1,47 | -0,18 | 2,5E-01 | 7,4E-09 | 2,4E-07 | 1,6E-09 | 1,4E-07 | 1,1E-01 | 0,0E+00 | 5,0E-10 | 6,7E-09 | 1,3E-10 | 8,4E-09 | 0,0E+00 | 2358424 | 2359503 | 1080 | - |
| Y11_RS10910 | Y11_23141 | pgk | phosphoglycerate kinase | 0,03 | 1,08 | 0,33 | 1,06 | 0,3 | -0,75 | 6,3E-01 | 1,9E-08 | 8,7E-04 | 7,9E-08 | 2,7E-03 | 3,3E-06 | 0,0E+00 | 1,0E-09 | 0,0E+00 | 2,8E-09 | 0,0E+00 | 0,0E+00 | 2359647 | 2360810 | 1164 | - |
| Y11_RS10915 | Y11_23151 | epd | erythrose-4-phosphate dehydrogenase | -0,33 | -0,39 | 0,23 | -0,06 | 0,57 | 0,62 | 2,7E-03 | 2,3E-03 | 3,5E-02 | 5,0E-01 | 1,5E-04 | 1,7E-04 | 0,0E+00 | 0,0E+00 | 0,0E+00 | 0,0E+00 | 0,0E+00 | 0,0E+00 | 2360914 | 2361930 | 1017 | - |
| Y11_RS10920 | Y11_23161 | tkt | transketolase | -0,2 | -0,55 | -0,57 | -0,36 | -0,37 | -0,02 | 4,4E-04 | 5,4E-07 | 5,3E-05 | 9,7E-06 | 8,6E-04 | 8,3E-01 | 0,0E+00 | 0,0E+00 | 0,0E+00 | 0,0E+00 | 0,0E+00 | 0,0E+00 | 2362269 | 2364263 | 1995 | - |
| Y11_RS10925 | Y11_23171 | | M48 family metallopeptidase | 0,08 | -1,78 | -0,73 | -1,86 | -0,82 | 1,04 | 1,8E-01 | 6,1E-10 | 1,9E-04 | 6,8E-09 | 1,8E-04 | 2,9E-05 | 0,0E+00 | 7,6E-11 | 0,0E+00 | 3,8E-10 | 0,0E+00 | 2,8E-06 | 2364558 | 2365310 | 753 | + |
| Y11_RS10935 | Y11_23201 | speA | biosynthetic arginine decarboxylase | -1,52 | -1,34 | -1,25 | 0,18 | 0,27 | 0,09 | 4,4E-10 | 2,7E-10 | 1,2E-07 | 6,7E-03 | 1,0E-02 | 2,8E-01 | 1,2E-10 | 4,2E-11 | 3,9E-09 | 0,0E+00 | 0,0E+00 | 0,0E+00 | 2366008 | 2367987 | 1980 | - |
| Y11_RS10940 | Y11_23221 | metK | methionine adenosyltransferase | 0,33 | 1,31 | 1,67 | 0,98 | 1,34 | 0,36 | 1,6E-04 | 1,7E-07 | 1,2E-04 | 6,5E-07 | 4,8E-04 | 1,8E-01 | 0,0E+00 | 6,5E-09 | 1,6E-06 | 0,0E+00 | 1,1E-05 | 0,0E+00 | 2368991 | 2370145 | 1155 | + |
| Y11_RS10955 | Y11_23251 | | metalloregulator ArsR/SmtB family transcription factor | 0,44 | 0,46 | 0,14 | 0,02 | -0,31 | -0,32 | 1,5E-01 | 6,0E-03 | 3,9E-01 | 9,6E-01 | 3,2E-01 | 6,7E-02 | 0,0E+00 | 0,0E+00 | 0,0E+00 | 0,0E+00 | 0,0E+00 | 0,0E+00 | 2371075 | 2371404 | 330 | - |
| Y11_RS10960 | Y11_23261 | | SprT family zinc-dependent metalloprotease | 1 | 0,92 | 1,18 | -0,08 | 0,17 | 0,26 | 1,2E-03 | 5,2E-03 | 2,7E-04 | 6,7E-01 | 1,0E-01 | 1,5E-01 | 4,3E-05 | 0,0E+00 | 3,4E-06 | 0,0E+00 | 0,0E+00 | 0,0E+00 | 2371531 | 2372043 | 513 | + |
| Y11_RS10970 | Y11_23281 | rsmE | 16S rRNA (uracil(1498)-N(3))-methyltransferase | -0,51 | -0,85 | -0,71 | -0,35 | -0,2 | 0,14 | 3,9E-05 | 3,2E-07 | 1,4E-05 | 6,4E-05 | 1,9E-02 | 5,3E-02 | 0,0E+00 | 0,0E+00 | 0,0E+00 | 0,0E+00 | 0,0E+00 | 0,0E+00 | 2372893 | 2373624 | 732 | + |
| Y11_RS10975 | Y11_23291 | gshB | glutathione synthase | -0,27 | 0,26 | 0,62 | 0,54 | 0,89 | 0,36 | 1,0E-03 | 1,3E-03 | 5,0E-06 | 8,4E-06 | 2,6E-07 | 2,4E-04 | 0,0E+00 | 0,0E+00 | 0,0E+00 | 0,0E+00 | 0,0E+00 | 0,0E+00 | 2373650 | 2374603 | 954 | + |

| | | | | | | | | | | | | | | | | | | | | | | | | | |
|---|---|---|---|---|---|---|---|---|---|---|---|---|---|---|---|---|---|---|---|---|---|---|---|---|---|
| Y11_RS10980 | Y11_23301 | | YqgE/AlgH family protein | -0,12 | 3,05 | 3,75 | 3,17 | 3,87 | 0,7 | 1,8E-01 | 3,7E-10 | 7,1E-12 | 1,8E-09 | 1,1E-10 | 7,2E-05 | 0,0E+00 | 5,3E-11 | 6,2E-12 | 1,4E-10 | 4,8E-11 | 0,0E+00 | 2374791 | 2375354 | 564 | + |
| Y11_RS10985 | Y11_23311 | ruvX | Holliday junction resolvase RuvX | 0,31 | -0,31 | -1,06 | -0,62 | -1,37 | -0,75 | 5,3E-02 | 8,2E-03 | 2,5E-05 | 3,3E-03 | 4,6E-05 | 5,8E-04 | 0,0E+00 | 0,0E+00 | 3,9E-07 | 0,0E+00 | 1,3E-06 | 0,0E+00 | 2375354 | 2375776 | 423 | + |
| Y11_RS10990 | Y11_23321 | aguB | N-carbamoylputrescine amidase | 0,2 | 0,76 | 0,94 | 0,56 | 0,74 | 0,18 | 6,2E-02 | 1,1E-05 | 1,1E-06 | 5,6E-04 | 5,7E-05 | 6,5E-02 | 0,0E+00 | 0,0E+00 | 0,0E+00 | 0,0E+00 | 0,0E+00 | 0,0E+00 | 2375958 | 2376842 | 885 | + |
| Y11_RS10995 | Y11_23331 | aguA | agmatine deiminase | 0,1 | 0,87 | 0,62 | 0,77 | 0,53 | -0,25 | 2,2E-01 | 2,7E-06 | 7,2E-05 | 1,1E-05 | 3,3E-04 | 2,6E-02 | 0,0E+00 | 0,0E+00 | 0,0E+00 | 0,0E+00 | 0,0E+00 | 0,0E+00 | 2376846 | 2377970 | 1125 | + |
| Y11_RS11005 | Y11_23351 | | YggS family pyridoxal phosphate-dependent enzyme | -0,16 | 0,79 | 0,84 | 0,95 | 0,99 | 0,04 | 2,1E-01 | 3,6E-06 | 6,6E-04 | 1,6E-05 | 4,2E-04 | 7,7E-01 | 0,0E+00 | 0,0E+00 | 0,0E+00 | 0,0E+00 | 0,0E+00 | 0,0E+00 | 2379191 | 2379886 | 696 | + |
| Y11_RS11010 | Y11_23361 | proC | pyrroline-5-carboxylate reductase | -0,15 | 0,22 | 0,41 | 0,38 | 0,56 | 0,19 | 2,1E-03 | 1,1E-03 | 4,6E-03 | 8,1E-05 | 8,4E-04 | 1,3E-01 | 0,0E+00 | 0,0E+00 | 0,0E+00 | 0,0E+00 | 0,0E+00 | 0,0E+00 | 2379922 | 2380743 | 822 | + |
| Y11_RS11015 | Y11_23371 | | YggT family protein | 0,06 | 0,05 | 1,16 | -0,02 | 1,1 | 1,12 | 8,9E-01 | 9,3E-01 | 6,9E-02 | 9,7E-01 | 3,8E-02 | 7,0E-02 | 0,0E+00 | 0,0E+00 | 7,6E-04 | 0,0E+00 | 7,9E-04 | 3,7E-03 | 2380877 | 2381431 | 555 | + |
| Y11_RS11020 | Y11_23381 | yggU | DUF167 family protein YggU | 0,16 | -0,39 | -0,14 | -0,54 | -0,3 | 0,24 | 3,4E-01 | 9,7E-03 | 4,0E-01 | 1,1E-02 | 1,7E-01 | 2,0E-01 | 0,0E+00 | 0,0E+00 | 0,0E+00 | 0,0E+00 | 0,0E+00 | 0,0E+00 | 2381428 | 2381718 | 291 | + |
| Y11_RS11025 | Y11_23391 | | XTP/dITP diphosphatase | -0,1 | -0,92 | -0,92 | -0,82 | -0,82 | 0 | 1,8E-01 | 3,0E-08 | 1,1E-05 | 6,7E-06 | 1,0E-04 | 9,8E-01 | 0,0E+00 | 0,0E+00 | 0,0E+00 | 0,0E+00 | 0,0E+00 | 0,0E+00 | 2381956 | 2382549 | 594 | + |
| Y11_RS11030 | Y11_23401 | hemW | radical SAM family heme chaperone HemW | 0,03 | -0,38 | -0,1 | -0,41 | -0,13 | 0,28 | 8,5E-01 | 1,3E-02 | 5,5E-01 | 3,1E-02 | 5,1E-01 | 1,2E-01 | 0,0E+00 | 0,0E+00 | 0,0E+00 | 0,0E+00 | 0,0E+00 | 0,0E+00 | 2382542 | 2383672 | 1131 | + |
| Y11_RS11035 | Y11_23411 | | DUF2884 domain-containing protein | 0,18 | 0,1 | 0,3 | -0,08 | 0,12 | 0,2 | 1,1E-01 | 1,4E-01 | 8,1E-04 | 4,3E-01 | 2,1E-01 | 2,4E-03 | 0,0E+00 | 0,0E+00 | 0,0E+00 | 0,0E+00 | 0,0E+00 | 0,0E+00 | 2383757 | 2384497 | 741 | - |
| Y11_RS11040 | Y11_23421 | glsB | glutaminase B | 0,36 | 0,7 | 0,61 | 0,34 | 0,25 | -0,09 | 2,0E-02 | 1,4E-04 | 4,0E-04 | 2,2E-02 | 7,3E-02 | 4,0E-01 | 0,0E+00 | 0,0E+00 | 0,0E+00 | 0,0E+00 | 0,0E+00 | 0,0E+00 | 2384600 | 2385616 | 1017 | - |
| Y11_RS11045 | Y11_23431 | | YggL family protein | -0,18 | -0,93 | -0,36 | -0,75 | -0,18 | 0,57 | 5,7E-01 | 1,3E-03 | 2,7E-01 | 3,7E-02 | 6,5E-01 | 9,3E-02 | 0,0E+00 | 0,0E+00 | 0,0E+00 | 0,0E+00 | 0,0E+00 | 0,0E+00 | 2385656 | 2385982 | 327 | - |
| Y11_RS11050 | Y11_23441 | trmB | tRNA (guanosine(46)-N7)-methyltransferase TrmB | -0,06 | -1,62 | -1,69 | -1,56 | -1,63 | -0,07 | 5,9E-01 | 3,1E-07 | 9,3E-06 | 4,8E-07 | 1,3E-05 | 6,9E-01 | 0,0E+00 | 1,1E-08 | 1,6E-07 | 1,3E-08 | 4,1E-07 | 0,0E+00 | 2385982 | 2386701 | 720 | - |
| Y11_RS11055 | Y11_23451 | mutY | A/G-specific adenine glycosylase | -0,1 | -0,03 | -0,41 | 0,09 | -0,3 | -0,38 | 7,7E-01 | 9,1E-01 | 1,6E-01 | 8,0E-01 | 4,2E-01 | 1,7E-01 | 0,0E+00 | 0,0E+00 | 0,0E+00 | 0,0E+00 | 0,0E+00 | 0,0E+00 | 2387003 | 2388145 | 1143 | + |
| Y11_RS11060 | Y11_23461 | | oxidative damage protection protein | 0,37 | 0,56 | 0,31 | 0,19 | -0,06 | -0,25 | 2,6E-04 | 1,4E-04 | 3,9E-02 | 2,9E-02 | 6,3E-01 | 9,7E-02 | 0,0E+00 | 0,0E+00 | 0,0E+00 | 0,0E+00 | 0,0E+00 | 0,0E+00 | 2388209 | 2388481 | 273 | + |
| Y11_RS11065 | Y11_23471 | mltC | membrane-bound lytic murein transglycosylase MltC | 0,17 | 0,17 | 0,27 | 0 | 0,1 | 0,1 | 1,1E-01 | 1,1E-01 | 2,0E-02 | 9,6E-01 | 1,6E-01 | 2,0E-01 | 0,0E+00 | 0,0E+00 | 0,0E+00 | 0,0E+00 | 0,0E+00 | 0,0E+00 | 2388619 | 2389695 | 1077 | + |
| Y11_RS11075 | Y11_23491 | speF | ornithine decarboxylase SpeF | -0,45 | -0,56 | 0,1 | -0,11 | 0,55 | 0,66 | 3,0E-05 | 9,9E-06 | 6,1E-02 | 5,6E-02 | 4,2E-07 | 3,1E-07 | 0,0E+00 | 0,0E+00 | 0,0E+00 | 0,0E+00 | 0,0E+00 | 0,0E+00 | 2390684 | 2392846 | 2163 | - |
| Y11_RS11120 | Y11_23571 | | linear amide C-N hydrolase | 0,4 | 2,26 | 3,12 | 1,87 | 2,72 | 0,85 | 1,4E-03 | 9,0E-11 | 5,7E-12 | 1,8E-08 | 8,3E-10 | 2,1E-07 | 0,0E+00 | 2,1E-11 | 6,0E-12 | 8,1E-10 | 1,5E-10 | 0,0E+00 | 2397926 | 2398999 | 1074 | + |
| Y11_RS11140 | Y11_23611 | speF | ornithine decarboxylase SpeF | 0,23 | -0,82 | 0,06 | -1,06 | -0,17 | 0,88 | 7,2E-02 | 2,9E-04 | 7,3E-01 | 5,6E-06 | 2,4E-01 | 5,0E-04 | 0,0E+00 | 0,0E+00 | 0,0E+00 | 1,2E-07 | 0,0E+00 | 0,0E+00 | 2402593 | 2404755 | 2163 | - |
| Y11_RS11145 | Y11_23621 | | TolC family protein | 0 | 0,39 | 0,28 | -15 | -15 | -0,09 | 0,0E+00 | 7,4E-02 | 1,3E-01 | 0,0E+00 | 0,0E+00 | 7,2E-01 | 0,0E+00 | 0,0E+00 | 0,0E+00 | 0,0E+00 | 0,0E+00 | 0,0E+00 | 2405501 | 2407021 | 1521 | - |
| Y11_RS11155 | Y11_23641 | mdtN | multidrug transporter subunit MdtN | 1,25 | 1,3 | 0,84 | 0,06 | -0,41 | -0,46 | 1,2E-05 | 1,5E-06 | 2,5E-04 | 6,7E-01 | 3,2E-02 | 8,8E-03 | 6,0E-07 | 4,2E-08 | 0,0E+00 | 0,0E+00 | 0,0E+00 | 0,0E+00 | 2408911 | 2409960 | 1050 | - |
| Y11_RS11165 | Y11_23661 | | amidohydrolase family protein | 0,55 | 2,35 | 3,41 | 1,81 | 2,86 | 1,06 | 5,1E-05 | 1,1E-08 | 2,6E-10 | 9,3E-08 | 1,1E-09 | 1,6E-05 | 0,0E+00 | 6,7E-10 | 4,2E-11 | 3,2E-09 | 1,9E-10 | 1,7E-06 | 2410370 | 2411833 | 1464 | - |
| Y11_RS11175 | Y11_23681 | map | type I methionyl aminopeptidase | 0,08 | 0,45 | 0,58 | 0,38 | 0,5 | 0,13 | 2,3E-01 | 1,3E-03 | 2,2E-04 | 2,5E-03 | 3,2E-04 | 2,9E-01 | 0,0E+00 | 0,0E+00 | 0,0E+00 | 0,0E+00 | 0,0E+00 | 0,0E+00 | 2412485 | 2413270 | 786 | + |
| Y11_RS11180 | Y11_23691 | | ABC transporter ATP-binding protein | 0,51 | 0,44 | -0,12 | -0,07 | -0,63 | -0,56 | 3,3E-05 | 1,2E-04 | 1,9E-01 | 3,1E-01 | 9,1E-05 | 2,3E-04 | 0,0E+00 | 0,0E+00 | 0,0E+00 | 0,0E+00 | 0,0E+00 | 0,0E+00 | 2413334 | 2414092 | 759 | - |
| Y11_RS11195 | Y11_23731 | | siderophore ABC transporter substrate-binding protein | 0,68 | 0,81 | 0,34 | 0,13 | -0,34 | -0,47 | 3,1E-07 | 7,3E-07 | 7,8E-03 | 3,5E-02 | 5,7E-03 | 1,5E-03 | 0,0E+00 | 0,0E+00 | 0,0E+00 | 0,0E+00 | 0,0E+00 | 0,0E+00 | 2416055 | 2417008 | 954 | - |
| Y11_RS11215 | Y11_23771 | | PLP-dependent aminotransferase family protein | 0,19 | 0,14 | 0,4 | -0,04 | 0,21 | 0,26 | 1,7E-01 | 3,3E-01 | 3,7E-03 | 7,7E-01 | 8,4E-02 | 7,4E-02 | 0,0E+00 | 0,0E+00 | 0,0E+00 | 0,0E+00 | 0,0E+00 | 0,0E+00 | 2421209 | 2422699 | 1491 | - |
| Y11_RS11225 | Y11_23801 | | MHS family MFS transporter | 3,42 | 5,92 | 6,09 | 2,5 | 2,66 | 0 | 1,1E-07 | 5,5E-11 | 1,5E-11 | 6,1E-07 | 2,4E-07 | 0,0E+00 | 9,8E-09 | 1,5E-11 | 8,0E-12 | 1,6E-08 | 1,3E-08 | 0,0E+00 | 2423338 | 2424741 | 1404 | + |
| Y11_RS11235 | Y11_23821 | | nucleoside hydrolase | 0,05 | -0,81 | -0,97 | -0,86 | -1,02 | -0,16 | 7,1E-01 | 1,9E-03 | 2,6E-03 | 4,4E-04 | 1,0E-03 | 5,3E-01 | 0,0E+00 | 0,0E+00 | 0,0E+00 | 0,0E+00 | 2,4E-05 | 0,0E+00 | 2426094 | 2427143 | 1050 | + |
| Y11_RS11240 | Y11_23831 | | Na/Pi cotransporter family protein | 0,81 | 0,48 | 0,88 | -0,33 | 0,07 | 0,4 | 1,6E-04 | 2,6E-03 | 3,4E-05 | 2,6E-02 | 5,8E-01 | 5,4E-03 | 0,0E+00 | 0,0E+00 | 0,0E+00 | 0,0E+00 | 0,0E+00 | 0,0E+00 | 2427274 | 2428893 | 1620 | + |
| Y11_RS11245 | Y11_23841 | | isochorismatase family protein | 0 | -0,48 | -0,34 | -0,67 | -0,64 | 0,14 | 0,0E+00 | 3,7E-02 | 1,3E-02 | 8,3E-03 | 2,1E-03 | 4,6E-01 | 0,0E+00 | 0,0E+00 | 0,0E+00 | 0,0E+00 | 0,0E+00 | 0,0E+00 | 2428973 | 2429500 | 528 | - |
| Y11_RS11250 | Y11_23851 | cueP | copper-binding periplasmic metallochaperone CueP | -15 | -15 | -15 | -3,18 | -2,17 | 1 | 0,0E+00 | 0,0E+00 | 0,0E+00 | 2,7E-07 | 3,9E-06 | 3,8E-03 | 0,0E+00 | 0,0E+00 | 0,0E+00 | 7,9E-09 | 1,4E-07 | 2,3E-04 | 2429680 | 2430240 | 561 | - |
| Y11_RS11255 | Y11_23861 | proP | glycine betaine/L-proline transporter ProP | 0,28 | 0,46 | 0,89 | 0,18 | 0,6 | 0,42 | 4,1E-01 | 3,4E-03 | 7,7E-04 | 5,9E-01 | 1,1E-01 | 2,5E-02 | 0,0E+00 | 0,0E+00 | 0,0E+00 | 0,0E+00 | 0,0E+00 | 0,0E+00 | 2430424 | 2431926 | 1503 | - |
| Y11_RS11275 | Y11_23901 | | EAL domain-containing protein | -0,5 | 0,46 | 0,9 | 0,97 | 1,41 | 0,44 | 6,0E-03 | 1,2E-05 | 4,0E-06 | 8,0E-05 | 1,2E-05 | 3,0E-04 | 0,0E+00 | 0,0E+00 | 0,0E+00 | 0,0E+00 | 3,8E-07 | 0,0E+00 | 2437280 | 2439844 | 2565 | + |
| Y11_RS11280 | Y11_23911 | | DUF3313 domain-containing protein | 0,93 | 3,27 | 3,97 | 2,35 | 3,05 | 0,7 | 1,8E-05 | 2,8E-09 | 1,1E-08 | 1,9E-07 | 2,1E-07 | 7,3E-03 | 0,0E+00 | 2,3E-10 | 5,8E-10 | 5,7E-09 | 1,2E-08 | 0,0E+00 | 2439912 | 2440595 | 684 | - |
| Y11_RS11290 | Y11_23921 | thiM | hydroxyethylthiazole kinase | 0,25 | 0,94 | 1,27 | 0,69 | 1,03 | 0,33 | 1,1E-01 | 2,6E-04 | 5,4E-06 | 4,6E-03 | 1,5E-04 | 7,8E-02 | 0,0E+00 | 0,0E+00 | 9,6E-08 | 0,0E+00 | 3,7E-06 | 0,0E+00 | 2440791 | 2441570 | 780 | - |
| Y11_RS11320 | Y11_23981 | hypB | hydrogenase nickel incorporation protein HypB | -0,43 | -1,07 | -1,07 | -0,64 | -0,64 | 0 | 3,9E-02 | 3,2E-04 | 5,0E-06 | 2,4E-02 | 7,0E-03 | 9,9E-01 | 0,0E+00 | 6,6E-06 | 9,1E-08 | 0,0E+00 | 0,0E+00 | 0,0E+00 | 2447709 | 2448764 | 1056 | - |
| Y11_RS11340 | Y11_24021 | hybC | hydrogenase 2 large subunit | 0 | -15 | 0 | -15 | 0 | 0,59 | 0,0E+00 | 0,0E+00 | 0,0E+00 | 0,0E+00 | 0,0E+00 | 1,1E-01 | 0,0E+00 | 0,0E+00 | 0,0E+00 | 0,0E+00 | 0,0E+00 | 0,0E+00 | 2450121 | 2451815 | 1695 | - |
| Y11_RS11360 | Y11_24061 | | glycine zipper 2TM domain-containing protein | 0,17 | 1,81 | 2,58 | 1,64 | 2,41 | 0,77 | 6,7E-01 | 9,8E-04 | 4,3E-05 | 8,7E-04 | 2,4E-05 | 1,2E-02 | 0,0E+00 | 2,0E-05 | 6,3E-07 | 1,5E-05 | 7,0E-07 | 0,0E+00 | 2456030 | 2456497 | 468 | + |
| Y11_RS11405 | Y11_24161 | | flagellin | 0 | -15 | 0 | -15 | 0 | 1,13 | 0,0E+00 | 0,0E+00 | 0,0E+00 | 0,0E+00 | 0,0E+00 | 7,3E-04 | 0,0E+00 | 0,0E+00 | 0,0E+00 | 0,0E+00 | 0,0E+00 | 5,0E-05 | 2462763 | 2464064 | 1302 | - |
| Y11_RS11410 | Y11_24191 | | transcriptional regulator | 0,24 | 0,36 | 0,37 | 0,14 | 0,16 | 0,01 | 4,2E-01 | 2,3E-01 | 2,1E-01 | 4,8E-01 | 3,7E-01 | 9,3E-01 | 0,0E+00 | 0,0E+00 | 0,0E+00 | 0,0E+00 | 0,0E+00 | 0,0E+00 | 2464582 | 2465667 | 1086 | + |
| Y11_RS11490 | Y11_24361 | flgM | flagellar biosynthesis anti-sigma factor FlgM | 0,08 | 1,54 | 1,43 | 1,45 | 1,35 | -0,11 | 6,9E-01 | 1,8E-03 | 2,2E-03 | 3,4E-03 | 4,3E-03 | 8,1E-01 | 0,0E+00 | 3,5E-05 | 2,5E-05 | 5,6E-05 | 9,5E-05 | 0,0E+00 | 2477279 | 2477554 | 276 | + |
| Y11_RS11570 | Y11_24551 | alkB | DNA oxidative demethylase AlkB | 0,2 | 0,79 | 0,89 | 0,59 | 0,69 | 0,09 | 7,2E-01 | 1,3E-03 | 3,6E-03 | 3,3E-01 | 2,7E-01 | 7,3E-01 | 0,0E+00 | 0,0E+00 | 0,0E+00 | 0,0E+00 | 0,0E+00 | 0,0E+00 | 2492879 | 2493544 | 666 | - |
| Y11_RS11645 | Y11_24711 | exbD | TonB system transport protein ExbD | 0,37 | -1,41 | -2,68 | -1,78 | -3,06 | -1,28 | 1,3E-02 | 5,7E-03 | 1,2E-03 | 1,3E-03 | 5,2E-04 | 8,6E-02 | 0,0E+00 | 1,1E-04 | 1,5E-05 | 2,2E-05 | 1,2E-05 | 4,5E-03 | 2505434 | 2505859 | 426 | - |
| Y11_RS11650 | Y11_24721 | exbB | tol-pal system-associated acyl-CoA thioesterase | 1,79 | 0,31 | -0,86 | -1,49 | -2,65 | -1,16 | 8,3E-09 | 1,4E-02 | 6,1E-06 | 3,0E-07 | 7,0E-10 | 3,4E-06 | 1,2E-09 | 0,0E+00 | 0,0E+00 | 8,5E-09 | 1,4E-10 | 4,7E-07 | 2505863 | 2506873 | 1011 | - |
| Y11_RS11655 | Y11_24731 | metC | cystathionine beta-lyase | -0,12 | 0,26 | 0,22 | 0,38 | 0,34 | -0,04 | 4,0E-01 | 9,6E-02 | 1,2E-01 | 1,4E-02 | 1,2E-02 | 7,1E-01 | 0,0E+00 | 0,0E+00 | 0,0E+00 | 0,0E+00 | 0,0E+00 | 0,0E+00 | 2507355 | 2508566 | 1212 | + |
| Y11_RS11665 | Y11_24751 | | AraC family transcriptional regulator | 1,28 | 0,35 | -0,13 | -0,93 | -1,42 | -0,48 | 5,7E-04 | 1,7E-01 | 6,3E-01 | 9,3E-04 | 2,5E-04 | 6,5E-02 | 2,2E-05 | 0,0E+00 | 0,0E+00 | 0,0E+00 | 6,3E-06 | 0,0E+00 | 2509493 | 2510386 | 894 | - |
| Y11_RS11670 | Y11_24761 | yqhD | alcohol dehydrogenase | -0,64 | 1,48 | 1,51 | 2,12 | 2,15 | 0,03 | 4,2E-05 | 8,7E-09 | 1,2E-08 | 1,2E-08 | 1,3E-08 | 7,5E-01 | 0,0E+00 | 5,5E-10 | 6,0E-10 | 6,1E-10 | 1,2E-09 | 0,0E+00 | 2510620 | 2511777 | 1158 | + |
| Y11_RS11675 | Y11_24771 | dkgA | 2,5-didehydrogluconate reductase DkgA | -0,22 | 3,08 | 4,16 | 3,29 | 4,37 | 1,08 | 1,2E-02 | 8,5E-11 | 1,3E-10 | 2,9E-10 | 2,3E-10 | 1,5E-05 | 0,0E+00 | 2,0E-11 | 2,6E-11 | 3,5E-11 | 7,2E-11 | 1,6E-06 | 2511995 | 2512828 | 834 | + |
| Y11_RS11685 | Y11_24791 | ftsP | cell division protein FtsP | -0,01 | -0,58 | -0,2 | -0,57 | -0,19 | 0,38 | 8,8E-01 | 8,2E-06 | 5,0E-03 | 2,6E-06 | 1,9E-03 | 1,3E-05 | 0,0E+00 | 0,0E+00 | 0,0E+00 | 0,0E+00 | 0,0E+00 | 0,0E+00 | 2515317 | 2516744 | 1428 | - |
| Y11_RS11690 | Y11_24801 | | 1-acylglycerol-3-phosphate O-acyltransferase | -0,55 | -0,45 | -0,37 | 0,09 | 0,18 | 0,08 | 1,7E-03 | 1,8E-04 | 6,8E-02 | 3,7E-01 | 3,7E-01 | 6,2E-01 | 0,0E+00 | 0,0E+00 | 0,0E+00 | 0,0E+00 | 0,0E+00 | 0,0E+00 | 2516907 | 2517638 | 732 | - |
| Y11_RS11695 | Y11_24811 | parC | DNA topoisomerase IV subunit A | -0,3 | 0,03 | -0,17 | 0,33 | 0,13 | -0,2 | 4,3E-05 | 5,6E-01 | 8,4E-02 | 3,3E-04 | 1,8E-01 | 6,9E-02 | 0,0E+00 | 0,0E+00 | 0,0E+00 | 0,0E+00 | 0,0E+00 | 0,0E+00 | 2517823 | 2520099 | 2277 | - |
| Y11_RS11700 | Y11_24821 | | NAD(P)H-dependent oxidoreductase | 0,09 | 0,81 | 0,43 | 0,73 | 0,35 | -0,38 | 3,3E-01 | 1,8E-05 | 5,6E-02 | 2,3E-06 | 9,4E-02 | 7,3E-02 | 0,0E+00 | 0,0E+00 | 0,0E+00 | 0,0E+00 | 0,0E+00 | 0,0E+00 | 2520411 | 2521004 | 594 | + |
| Y11_RS11705 | Y11_24831 | | antibiotic biosynthesis monooxygenase | 0,15 | 0,9 | 0,94 | 0,75 | 0,79 | 0,04 | 2,2E-01 | 4,0E-06 | 2,0E-04 | 1,3E-04 | 1,3E-03 | 7,6E-01 | 0,0E+00 | 0,0E+00 | 0,0E+00 | 0,0E+00 | 0,0E+00 | 0,0E+00 | 2521037 | 2521342 | 306 | + |
| Y11_RS11710 | Y11_24841 | | LysR family transcriptional regulator | -0,34 | 2,42 | 1,26 | 2,76 | 1,56 | -1,23 | 5,5E-01 | 2,2E-06 | 1,2E-02 | 1,2E-03 | 4,5E-02 | 2,1E-02 | 0,0E+00 | 6,0E-08 | 1,4E-04 | 1,9E-05 | 9,3E-04 | 1,1E-03 | 2521445 | 2522368 | 924 | - |
| Y11_RS11715 | Y11_24851 | parE | DNA topoisomerase IV subunit B | -0,29 | -0,13 | -0,46 | 0,16 | -0,17 | -0,33 | 1,7E-03 | 8,7E-02 | 1,1E-03 | 3,8E-02 | 1,1E-01 | 8,1E-03 | 0,0E+00 | 0,0E+00 | 0,0E+00 | 0,0E+00 | 0,0E+00 | 0,0E+00 | 2522372 | 2524267 | 1896 | - |
| Y11_RS11720 | Y11_24861 | yqiA | esterase YqiA | -0,23 | 1,76 | 1,76 | 1,99 | 1,98 | -0,01 | 1,8E-02 | 1,6E-07 | 3,9E-07 | 1,7E-07 | 3,5E-07 | 9,6E-01 | 0,0E+00 | 6,2E-09 | 1,0E-08 | 5,3E-09 | 1,7E-08 | 0,0E+00 | 2524382 | 2524963 | 582 | - |
| Y11_RS11725 | Y11_24871 | cpdA | 3',5'-cyclic-AMP phosphodiesterase | 1,49 | 2,66 | 2,09 | 1,17 | 0,63 | 0 | 4,1E-04 | 5,5E-07 | 4,0E-06 | 3,4E-04 | 1,2E-02 | 0,0E+00 | 1,6E-05 | 1,7E-08 | 7,4E-08 | 5,9E-06 | 0,0E+00 | 0,0E+00 | 2524964 | 2525791 | 828 | - |
| Y11_RS11735 | Y11_24891 | nudF | ADP-ribose diphosphatase | 0,45 | 0,05 | 0,26 | -0,4 | -0,18 | 0,21 | 3,0E-04 | 5,5E-01 | 2,7E-02 | 3,2E-05 | 3,5E-02 | 2,8E-02 | 0,0E+00 | 0,0E+00 | 0,0E+00 | 0,0E+00 | 0,0E+00 | 0,0E+00 | 2526308 | 2526940 | 633 | - |
| Y11_RS11740 | Y11_24901 | tolC | outer membrane channel protein TolC | 0,05 | -0,18 | -0,18 | -0,23 | -0,23 | 0 | 5,6E-02 | 3,0E-04 | 2,3E-01 | 4,0E-05 | 1,4E-01 | 1,0E+00 | 0,0E+00 | 0,0E+00 | 0,0E+00 | 0,0E+00 | 0,0E+00 | 0,0E+00 | 2527453 | 2528934 | 1482 | + |
| Y11_RS11745 | Y11_24921 | | DUF1190 family protein | 0,73 | -4,14 | -4,5 | -4,87 | -5,23 | -0,36 | 4,1E-03 | 9,0E-09 | 3,0E-07 | 1,7E-09 | 8,2E-08 | 2,3E-01 | 0,0E+00 | 5,6E-10 | 8,2E-09 | 1,3E-10 | 5,3E-09 | 0,0E+00 | 2529185 | 2529874 | 690 | + |
| Y11_RS11750 | Y11_24931 | | glutathionylspermidine synthase family protein | 0,26 | -1,58 | -1,25 | -1,84 | -1,51 | 0,33 | 8,7E-04 | 1,2E-11 | 4,8E-08 | 1,6E-10 | 3,3E-08 | 6,0E-04 | 0,0E+00 | 6,3E-12 | 1,8E-09 | 2,4E-11 | 2,5E-09 | 0,0E+00 | 2529885 | 2531045 | 1161 | + |
| Y11_RS11755 | Y11_24941 | ygiD | 4,5-DOPA dioxygenase extradiol | 0,29 | 0,61 | 0,7 | 0,32 | 0,41 | 0,09 | 2,0E-01 | 1,2E-02 | 3,7E-02 | 1,8E-01 | 2,1E-01 | 7,6E-01 | 0,0E+00 | 0,0E+00 | 0,0E+00 | 0,0E+00 | 0,0E+00 | 0,0E+00 | 2531218 | 2532000 | 783 | - |
| Y11_RS11760 | Y11_24951 | ribB | 3,4-dihydroxy-2-butanone-4-phosphate synthase | -0,65 | 2,08 | 2,55 | 2,74 | 3,21 | 0,47 | 1,1E-06 | 5,1E-10 | 6,1E-10 | 1,7E-10 | 2,1E-10 | 7,3E-04 | 0,0E+00 | 6,7E-11 | 7,5E-11 | 2,5E-11 | 6,9E-11 | 0,0E+00 | 2532275 | 2532913 | 639 | - |
| Y11_RS11765 | Y11_24961 | | accessory factor UbiK family protein | 0,22 | 1,01 | 0,83 | 0,79 | 0,61 | -0,18 | 3,7E-02 | 1,9E-06 | 7,6E-06 | 2,7E-06 | 1,6E-05 | 1,3E-02 | 0,0E+00 | 5,4E-08 | 0,0E+00 | 0,0E+00 | 0,0E+00 | 0,0E+00 | 2533379 | 2533660 | 282 | + |
| Y11_RS11775 | Y11_24981 | hldE | bifunctional D-glycero-beta-D-manno-heptose-7-phosphate kinase/D-glycero-beta-D-manno-heptose 1-phosphate adenylyltransferase HldE | -0,1 | 0,81 | 1,08 | 0,91 | 1,19 | 0,27 | 4,0E-02 | 3,2E-09 | 2,7E-09 | 6,6E-08 | 2,1E-08 | 3,2E-04 | 0,0E+00 | 0,0E+00 | 2,0E-10 | 0,0E+00 | 1,7E-09 | 0,0E+00 | 2535121 | 2536551 | 1431 | - |
| Y11_RS11780 | Y11_24991 | glnE | bifunctional [glutamate--ammonia ligase]-adenylyl-L-tyrosine phosphorylase/[glutamate--ammonia-ligase] adenylyltransferase | 0,14 | 0,7 | 1,13 | 0,56 | 0,98 | 0,43 | 1,2E-02 | 4,9E-07 | 4,3E-06 | 1,3E-06 | 1,0E-05 | 3,2E-03 | 0,0E+00 | 0,0E+00 | 7,9E-08 | 0,0E+00 | 0,0E+00 | 0,0E+00 | 2536677 | 2539529 | 2853 | - |

| | | | | | | | | | | | | | | | | | | | | | | | | | |
|---|---|---|---|---|---|---|---|---|---|---|---|---|---|---|---|---|---|---|---|---|---|---|---|---|---|
| Y11_RS11785 | Y11_25001 | | inorganic triphosphatase | 0,04 | 0,53 | 0,92 | 0,49 | 0,88 | 0,39 | 4,6E-01 | 3,4E-07 | 2,9E-07 | 5,7E-06 | 1,0E-06 | 1,7E-04 | 0,0E+00 | 0,0E+00 | 0,0E+00 | 0,0E+00 | 0,0E+00 | 0,0E+00 | 2539674 | 2540981 | 1308 | - |
| Y11_RS11790 | Y11_25011 | | SH3 domain-containing protein | 0,53 | 0,03 | -0,32 | -0,5 | -0,86 | -0,36 | 2,6E-05 | 7,1E-01 | 2,7E-03 | 5,7E-04 | 4,2E-06 | 7,4E-03 | 0,0E+00 | 0,0E+00 | 0,0E+00 | 0,0E+00 | 0,0E+00 | 0,0E+00 | 2541307 | 2541927 | 621 | + |
| Y11_RS11795 | Y11_25021 | | multifunctional CCA addition/repair protein | 0,25 | 0,78 | 0,43 | 0,52 | 0,18 | -0,35 | 8,6E-03 | 2,5E-05 | 2,9E-03 | 2,7E-04 | 1,1E-01 | 1,4E-02 | 0,0E+00 | 0,0E+00 | 0,0E+00 | 0,0E+00 | 0,0E+00 | 0,0E+00 | 2541974 | 2543212 | 1239 | + |
| Y11_RS11805 | Y11_25041 | folB | bifunctional dihydroneopterin aldolase/7,8-dihydroneopterin epimerase | 0,29 | -0,12 | -0,14 | -0,41 | -0,42 | -0,02 | 1,5E-01 | 3,6E-01 | 7,1E-01 | 2,9E-02 | 2,8E-01 | 9,6E-01 | 0,0E+00 | 0,0E+00 | 0,0E+00 | 0,0E+00 | 0,0E+00 | 0,0E+00 | 2544553 | 2544912 | 360 | - |
| Y11_RS11810 | Y11_25051 | plsY | glycerol-3-phosphate 1-O-acyltransferase PlsY | 0,03 | 1,06 | 1,89 | 1,03 | 1,86 | 0,83 | 8,4E-01 | 7,7E-05 | 6,4E-05 | 1,0E-04 | 7,4E-05 | 8,0E-03 | 0,0E+00 | 1,7E-06 | 9,0E-07 | 1,9E-06 | 2,0E-06 | 0,0E+00 | 2545020 | 2545673 | 654 | + |
| Y11_RS11815 | Y11_25061 | tsaD | tRNA (adenosine(37)-N6)-threonylcarbamoyltransferase complex transferase subunit TsaD | -0,42 | 0,18 | 0,09 | 0,6 | 0,51 | -0,09 | 4,0E-04 | 1,1E-04 | 3,4E-01 | 2,2E-05 | 1,5E-03 | 3,0E-01 | 0,0E+00 | 0,0E+00 | 0,0E+00 | 0,0E+00 | 0,0E+00 | 0,0E+00 | 2545829 | 2546842 | 1014 | - |
| Y11_RS11820 | Y11_25071 | rpsU | 30S ribosomal protein S21 | 0,42 | -0,27 | -0,51 | -0,69 | -0,93 | -0,24 | 2,8E-04 | 1,0E-02 | 6,4E-02 | 1,4E-05 | 4,1E-03 | 3,5E-01 | 0,0E+00 | 0,0E+00 | 0,0E+00 | 0,0E+00 | 0,0E+00 | 0,0E+00 | 2547216 | 2547431 | 216 | + |
| Y11_RS11825 | Y11_25091 | dnaG | DNA primase | -0,55 | 0,38 | 0,09 | 0,93 | 0,64 | -0,29 | 1,0E-04 | 4,4E-04 | 2,9E-01 | 1,1E-06 | 4,9E-05 | 3,4E-03 | 0,0E+00 | 0,0E+00 | 0,0E+00 | 0,0E+00 | 0,0E+00 | 0,0E+00 | 2547568 | 2549316 | 1749 | + |
| Y11_RS11830 | Y11_25101 | rpoD | RNA polymerase sigma factor RpoD | -0,21 | 0,68 | 0 | 0,89 | 0,21 | -0,68 | 8,4E-02 | 3,0E-05 | 1,0E+00 | 2,9E-06 | 1,8E-01 | 5,4E-04 | 0,0E+00 | 0,0E+00 | 0,0E+00 | 0,0E+00 | 0,0E+00 | 0,0E+00 | 2549474 | 2551312 | 1839 | + |
| Y11_RS11920 | Y11_25291 | | helix-turn-helix domain-containing protein | -1,04 | 0,67 | 0,58 | 1,71 | 1,61 | -0,09 | 3,4E-04 | 1,1E-03 | 2,5E-02 | 2,0E-06 | 6,7E-05 | 6,4E-01 | 1,3E-05 | 0,0E+00 | 0,0E+00 | 4,7E-08 | 1,8E-06 | 0,0E+00 | 2565211 | 2565528 | 318 | - |
| Y11_RS12035 | Y11_25541 | | phage repressor protein | 0,65 | 1,35 | 1,62 | 0,7 | 0,96 | 0,26 | 6,6E-07 | 1,5E-08 | 1,4E-05 | 7,9E-06 | 6,1E-04 | 1,9E-01 | 0,0E+00 | 8,5E-10 | 2,3E-07 | 0,0E+00 | 0,0E+00 | 0,0E+00 | 2581403 | 2581978 | 576 | + |
| Y11_RS12070 | Y11_25621 | murP | PTS N-acetylmuramic acid transporter subunit IIBC | -1,03 | 0,68 | 0,83 | 1,72 | 1,89 | 0,15 | 6,2E-05 | 1,9E-03 | 3,7E-04 | 1,8E-06 | 3,5E-07 | 3,2E-01 | 2,8E-06 | 0,0E+00 | 0,0E+00 | 4,3E-08 | 1,7E-08 | 0,0E+00 | 2590076 | 2591527 | 1452 | + |
| Y11_RS12095 | Y11_25681 | | HdeD family acid-resistance protein | 0,52 | 4,62 | 15 | 4,1 | 15 | 0 | 2,4E-01 | 8,9E-10 | 0,0E+00 | 7,5E-06 | 0,0E+00 | 0,0E+00 | 0,0E+00 | 9,9E-11 | 0,0E+00 | 1,6E-07 | 0,0E+00 | 0,0E+00 | 2596450 | 2597022 | 573 | + |
| Y11_RS12105 | Y11_25701 | | NADPH-dependent 2,4-dienoyl-CoA reductase | 1,48 | -0,99 | 0,91 | -2,5 | -0,59 | 1,89 | 1,3E-04 | 2,0E-03 | 6,6E-04 | 1,9E-05 | 4,1E-02 | 5,4E-05 | 5,4E-06 | 0,0E+00 | 0,0E+00 | 3,7E-07 | 0,0E+00 | 4,7E-06 | 2598223 | 2600244 | 2022 | + |
| Y11_RS12120 | Y11_25731 | rlmG | 23S rRNA (guanine(1835)-N(2))-methyltransferase RlmG | 0,58 | -1,04 | -1,6 | -1,62 | -2,17 | -0,55 | 1,7E-04 | 1,3E-06 | 2,7E-06 | 1,7E-07 | 4,4E-07 | 4,8E-03 | 0,0E+00 | 3,7E-08 | 5,4E-08 | 5,3E-09 | 2,1E-08 | 0,0E+00 | 2602094 | 2603281 | 1188 | - |
| Y11_RS12145 | Y11_25781 | | Gfo/Idh/MocA family oxidoreductase | -0,16 | 0,85 | 2,12 | 1,01 | 2,28 | 1,27 | 1,9E-02 | 1,1E-05 | 1,1E-09 | 2,2E-06 | 3,6E-10 | 7,3E-07 | 0,0E+00 | 0,0E+00 | 1,1E-10 | 5,0E-08 | 9,7E-11 | 1,4E-07 | 2609709 | 2610710 | 1002 | + |
| Y11_RS12160 | Y11_25801 | sstT | serine/threonine transporter SstT | 1,96 | -3,5 | -0,94 | -5,46 | -2,9 | 2,56 | 2,4E-05 | 2,9E-07 | 3,6E-02 | 1,5E-13 | 1,1E-05 | 2,8E-05 | 1,2E-06 | 9,9E-09 | 0,0E+00 | 1,2E-12 | 3,7E-07 | 2,8E-06 | 2612511 | 2613767 | 1257 | + |
| Y11_RS12170 | Y11_25821 | | altronate dehydratase family protein | -1,09 | 0,29 | 0,17 | 1,38 | 1,25 | -0,13 | 3,5E-02 | 4,9E-01 | 7,5E-01 | 5,9E-04 | 1,3E-02 | 7,4E-01 | 1,2E-03 | 0,0E+00 | 0,0E+00 | 1,0E-05 | 2,7E-04 | 0,0E+00 | 2614523 | 2616013 | 1491 | - |
| Y11_RS12175 | Y11_25831 | | tagaturonate reductase | -1,18 | 1,48 | 2 | 2,67 | 3,18 | 0,51 | 1,1E-07 | 1,2E-08 | 5,8E-07 | 2,8E-11 | 1,2E-08 | 5,0E-03 | 9,8E-09 | 7,0E-10 | 1,4E-08 | 8,5E-12 | 1,1E-09 | 0,0E+00 | 2616024 | 2617475 | 1452 | - |
| Y11_RS12180 | Y11_25841 | uxaC | glucuronate isomerase | -0,94 | 1,88 | 3,1 | 2,81 | 4,04 | 1,23 | 6,1E-09 | 5,4E-11 | 2,9E-10 | 7,4E-12 | 4,9E-11 | 6,9E-07 | 0,0E+00 | 1,5E-11 | 4,3E-11 | 5,4E-12 | 2,8E-11 | 1,4E-07 | 2617597 | 2619006 | 1410 | - |
| Y11_RS12190 | Y11_25871 | exuR | transcriptional regulator ExuR | -0,21 | 0,15 | 0,57 | 0,36 | 0,78 | 0,42 | 6,6E-03 | 1,6E-01 | 6,8E-05 | 7,3E-03 | 1,4E-05 | 5,2E-03 | 0,0E+00 | 0,0E+00 | 0,0E+00 | 0,0E+00 | 0,0E+00 | 0,0E+00 | 2621075 | 2621857 | 783 | + |
| Y11_RS12200 | Y11_25891 | | DedA family protein | 0,92 | 1,39 | 1,79 | 0,39 | 0 | 0,29 | 9,6E-02 | 9,0E-03 | 1,3E-03 | 3,8E-01 | 0,0E+00 | 3,2E-01 | 0,0E+00 | 1,7E-04 | 1,5E-05 | 0,0E+00 | 0,0E+00 | 0,0E+00 | 2622998 | 2623690 | 693 | + |
| Y11_RS12210 | Y11_25911 | | DUF1090 domain-containing protein | -0,08 | 1,41 | 1,79 | 1,49 | 1,86 | 0,37 | 3,3E-01 | 7,0E-10 | 4,1E-08 | 6,0E-08 | 1,6E-07 | 4,1E-03 | 0,0E+00 | 8,2E-11 | 1,6E-09 | 2,2E-09 | 9,3E-09 | 0,0E+00 | 2624262 | 2624648 | 387 | + |
| Y11_RS12215 | Y11_25921 | | YqjD family protein | 0,12 | 0,98 | 1,31 | 0,86 | 1,2 | 0,34 | 6,7E-01 | 3,2E-03 | 7,0E-04 | 1,2E-03 | 2,3E-04 | 5,3E-02 | 0,0E+00 | 0,0E+00 | 8,5E-06 | 0,0E+00 | 5,7E-06 | 0,0E+00 | 2624773 | 2625078 | 306 | + |
| Y11_RS12220 | Y11_25931 | | phage holin family protein | 1,08 | 3,93 | 3,05 | 2,85 | 1,95 | -0,9 | 6,5E-02 | 1,9E-11 | 8,2E-08 | 4,8E-04 | 5,9E-03 | 5,6E-04 | 2,2E-03 | 7,7E-12 | 2,7E-09 | 8,3E-06 | 1,3E-04 | 0,0E+00 | 2625081 | 2625476 | 396 | + |
| Y11_RS12230 | Y11_25951 | | DoxX family protein | 0,34 | 1,24 | 1,97 | 0,9 | 1,63 | 0,73 | 5,3E-02 | 1,9E-05 | 2,1E-04 | 1,1E-03 | 1,1E-03 | 5,5E-02 | 0,0E+00 | 4,5E-07 | 2,8E-06 | 0,0E+00 | 2,6E-05 | 0,0E+00 | 2626182 | 2626577 | 396 | + |
| Y11_RS12235 | Y11_25971 | | glutathione S-transferase family protein | -0,81 | 0,58 | 1,08 | 1,39 | 1,89 | 0,5 | 2,4E-05 | 5,8E-05 | 7,4E-07 | 3,1E-07 | 3,4E-08 | 1,7E-04 | 0,0E+00 | 0,0E+00 | 1,8E-08 | 8,7E-09 | 2,6E-09 | 0,0E+00 | 2626748 | 2627734 | 987 | + |
| Y11_RS12240 | Y11_25981 | | LysR family transcriptional regulator | 0,14 | 0,02 | -0,04 | -0,12 | -0,18 | -0,06 | 2,5E-01 | 8,5E-01 | 7,4E-01 | 1,5E-01 | 1,7E-01 | 5,6E-01 | 0,0E+00 | 0,0E+00 | 0,0E+00 | 0,0E+00 | 0,0E+00 | 0,0E+00 | 2627751 | 2628647 | 897 | - |
| Y11_RS12245 | Y11_25991 | | pirin family protein | 0,05 | 0,08 | -0,21 | 0,04 | -0,25 | -0,29 | 6,4E-01 | 3,9E-01 | 1,2E-01 | 7,1E-01 | 7,1E-02 | 4,1E-02 | 0,0E+00 | 0,0E+00 | 0,0E+00 | 0,0E+00 | 0,0E+00 | 0,0E+00 | 2628892 | 2629596 | 705 | + |
| Y11_RS12255 | Y11_26031 | | aldo/keto reductase | -0,19 | 1,7 | 1,91 | 1,89 | 2,1 | 0,21 | 1,0E-02 | 8,0E-10 | 5,3E-08 | 3,4E-09 | 5,3E-08 | 8,0E-02 | 0,0E+00 | 9,2E-11 | 1,9E-09 | 2,1E-10 | 3,7E-09 | 0,0E+00 | 2630759 | 2631754 | 996 | - |
| Y11_RS12270 | Y11_26061 | | fimbrial biogenesis outer membrane usher protein | 1,03 | 2,38 | 2,8 | 1,35 | 1,78 | 0,43 | 3,2E-04 | 6,0E-06 | 1,2E-07 | 1,1E-03 | 4,3E-05 | 1,5E-01 | 1,3E-05 | 1,5E-07 | 3,9E-09 | 1,9E-05 | 1,2E-06 | 0,0E+00 | 2634308 | 2636908 | 2601 | - |
| Y11_RS12275 | Y11_26071 | | fimbria/pilus periplasmic chaperone | 1,17 | 2,88 | 3,18 | 1,71 | 2,03 | 0,3 | 8,8E-07 | 1,4E-07 | 9,7E-10 | 1,0E-05 | 6,8E-08 | 1,3E-01 | 5,8E-08 | 5,6E-09 | 9,9E-11 | 2,2E-07 | 4,5E-09 | 0,0E+00 | 2637007 | 2637726 | 720 | - |
| Y11_RS12290 | | | helix-turn-helix domain-containing protein | -0,19 | 0,41 | 0,06 | 0,59 | 0,25 | -0,34 | 1,2E-01 | 5,8E-03 | 6,9E-01 | 3,5E-04 | 1,2E-01 | 4,5E-02 | 0,0E+00 | 0,0E+00 | 0,0E+00 | 0,0E+00 | 0,0E+00 | 0,0E+00 | 2640513 | 2640746 | 234 | + |
| Y11_RS12295 | | | hypothetical protein | 0,11 | 0,17 | 0,38 | 0,06 | 0,27 | 0,21 | 3,9E-01 | 1,2E-01 | 3,8E-02 | 6,7E-01 | 1,8E-01 | 2,6E-01 | 0,0E+00 | 0,0E+00 | 0,0E+00 | 0,0E+00 | 0,0E+00 | 0,0E+00 | 2640739 | 2641305 | 567 | + |
| Y11_RS12320 | Y11_26151 | rsmI | 16S rRNA (cytidine(1402)-2'-O)-methyltransferase | -0,02 | -0,06 | 0,2 | -0,04 | 0,22 | 0,26 | 7,6E-01 | 3,3E-01 | 5,7E-02 | 4,9E-01 | 3,6E-02 | 1,3E-02 | 0,0E+00 | 0,0E+00 | 0,0E+00 | 0,0E+00 | 0,0E+00 | 0,0E+00 | 2646320 | 2647219 | 900 | - |
| Y11_RS12325 | Y11_26161 | | penicillin-binding protein activator | 0,66 | -0,36 | -0,16 | -1,02 | -0,82 | 0,2 | 4,8E-07 | 2,4E-04 | 3,9E-02 | 7,4E-08 | 1,2E-06 | 2,7E-02 | 0,0E+00 | 0,0E+00 | 0,0E+00 | 2,6E-09 | 0,0E+00 | 0,0E+00 | 2647282 | 2649243 | 1962 | + |
| Y11_RS12335 | Y11_26181 | diaA | DnaA initiator-associating protein DiaA | 0,13 | -0,18 | -0,33 | -0,3 | -0,46 | -0,16 | 2,9E-01 | 2,9E-01 | 9,5E-02 | 1,3E-01 | 4,7E-02 | 5,1E-01 | 0,0E+00 | 0,0E+00 | 0,0E+00 | 0,0E+00 | 0,0E+00 | 0,0E+00 | 2649909 | 2650499 | 591 | + |
| Y11_RS12340 | Y11_26191 | yraP | divisome-associated lipoprotein YraP | -0,08 | 0,29 | 0,36 | 0,38 | 0,44 | 0,06 | 1,9E-01 | 3,1E-03 | 3,8E-03 | 8,9E-05 | 3,7E-04 | 4,7E-01 | 0,0E+00 | 0,0E+00 | 0,0E+00 | 0,0E+00 | 0,0E+00 | 0,0E+00 | 2650510 | 2651085 | 576 | + |
| Y11_RS12350 | Y11_26211 | mtgA | monofunctional biosynthetic peptidoglycan transglycosylase | 0,28 | 1,63 | 1,7 | 1,35 | 1,43 | 0,08 | 2,5E-01 | 8,0E-05 | 9,1E-05 | 5,2E-04 | 5,1E-04 | 7,7E-01 | 0,0E+00 | 1,8E-06 | 1,2E-06 | 8,9E-06 | 1,2E-05 | 0,0E+00 | 2652680 | 2653405 | 726 | - |
| Y11_RS12355 | Y11_26221 | elbB | isoprenoid biosynthesis glyoxalase ElbB | 0,19 | 1,89 | 2,3 | 1,7 | 2,11 | 0,41 | 2,8E-02 | 5,4E-09 | 2,3E-08 | 2,3E-08 | 6,1E-08 | 7,0E-03 | 0,0E+00 | 3,9E-10 | 9,8E-10 | 9,9E-10 | 4,1E-09 | 0,0E+00 | 2653402 | 2654055 | 654 | - |
| Y11_RS12360 | Y11_26231 | arcB | aerobic respiration two-component sensor histidine kinase ArcB | 0,01 | 0,7 | 1,21 | 0,68 | 1,19 | 0,51 | 9,0E-01 | 4,3E-07 | 2,3E-07 | 2,0E-04 | 9,6E-06 | 1,2E-04 | 0,0E+00 | 0,0E+00 | 6,6E-09 | 0,0E+00 | 3,2E-07 | 0,0E+00 | 2654296 | 2656632 | 2337 | - |
| Y11_RS12370 | Y11_26251 | gltB | glutamate synthase large subunit | 0,96 | -0,32 | -0,76 | -1,27 | -1,71 | -0,44 | 6,4E-09 | 3,1E-05 | 8,7E-07 | 2,2E-09 | 2,6E-09 | 8,6E-05 | 0,0E+00 | 0,0E+00 | 0,0E+00 | 1,6E-10 | 3,4E-10 | 0,0E+00 | 2658324 | 2662784 | 4461 | + |
| Y11_RS12375 | Y11_26261 | | glutamate synthase small subunit | 0,77 | -0,29 | -0,72 | -1,06 | -1,49 | -0,43 | 2,2E-06 | 4,5E-03 | 1,7E-04 | 3,0E-06 | 1,8E-06 | 9,7E-03 | 0,0E+00 | 0,0E+00 | 0,0E+00 | 6,8E-08 | 7,1E-08 | 0,0E+00 | 2662794 | 2664212 | 1419 | + |
| Y11_RS12380 | Y11_26271 | | hypothetical protein | -1,84 | 0,21 | -0,11 | 2,06 | 1,73 | -0,32 | 5,3E-08 | 1,5E-02 | 2,0E-01 | 2,7E-09 | 2,1E-08 | 1,2E-04 | 5,3E-09 | 0,0E+00 | 0,0E+00 | 1,8E-10 | 1,7E-09 | 0,0E+00 | 2664666 | 2665151 | 486 | + |
| Y11_RS12390 | Y11_26281 | sspB | ClpXP protease specificity-enhancing factor | 0,81 | -0,33 | -0,76 | -1,14 | -1,57 | -0,43 | 4,7E-05 | 1,5E-01 | 1,1E-02 | 3,0E-04 | 7,4E-05 | 1,6E-01 | 0,0E+00 | 0,0E+00 | 0,0E+00 | 5,3E-06 | 2,0E-06 | 0,0E+00 | 2665393 | 2665908 | 516 | - |
| Y11_RS12395 | Y11_26291 | sspA | stringent starvation protein A | 5,09 | 0,04 | 0,29 | -5,05 | -4,8 | 0,25 | 1,1E-10 | 7,0E-01 | 2,0E-02 | 1,3E-10 | 3,0E-10 | 3,9E-02 | 3,5E-11 | 0,0E+00 | 0,0E+00 | 2,1E-11 | 8,6E-11 | 0,0E+00 | 2665914 | 2666555 | 642 | - |
| Y11_RS12400 | Y11_26311 | rpsI | 30S ribosomal protein S9 | 0,1 | -0,44 | -0,99 | -0,55 | -1,09 | -0,55 | 3,2E-01 | 5,7E-03 | 1,1E-04 | 3,0E-03 | 8,8E-05 | 1,0E-02 | 0,0E+00 | 0,0E+00 | 0,0E+00 | 0,0E+00 | 2,3E-06 | 0,0E+00 | 2666950 | 2667342 | 393 | - |
| Y11_RS12405 | Y11_26321 | rplM | 50S ribosomal protein L13 | 0,28 | -0,49 | -0,91 | -0,77 | -1,19 | -0,42 | 3,1E-03 | 5,5E-06 | 8,1E-04 | 6,3E-06 | 1,9E-04 | 4,6E-02 | 0,0E+00 | 0,0E+00 | 0,0E+00 | 0,0E+00 | 4,8E-06 | 0,0E+00 | 2667357 | 2667785 | 429 | - |
| Y11_RS12410 | Y11_26331 | | cell division protein ZapE | 0,03 | -0,57 | -0,82 | -0,6 | -0,85 | -0,25 | 8,8E-01 | 1,0E-01 | 7,2E-02 | 7,2E-02 | 5,7E-02 | 5,8E-01 | 0,0E+00 | 0,0E+00 | 0,0E+00 | 0,0E+00 | 0,0E+00 | 0,0E+00 | 2668118 | 2669245 | 1128 | - |
| Y11_RS12415 | Y11_26341 | | DUF1043 family protein | 0,06 | 1,73 | 2,06 | 1,67 | 2 | 0,33 | 3,0E-01 | 2,5E-07 | 1,0E-09 | 6,7E-07 | 7,9E-09 | 3,1E-02 | 0,0E+00 | 9,0E-09 | 1,0E-10 | 1,7E-08 | 7,9E-10 | 0,0E+00 | 2669469 | 2669873 | 405 | + |
| Y11_RS12420 | Y11_26351 | degQ | serine endoprotease DegQ | -0,37 | -0,57 | -0,13 | -0,2 | 0,25 | 0,44 | 4,7E-05 | 2,3E-06 | 2,9E-01 | 5,2E-03 | 6,1E-02 | 4,5E-03 | 0,0E+00 | 0,0E+00 | 0,0E+00 | 0,0E+00 | 0,0E+00 | 0,0E+00 | 2670335 | 2671708 | 1374 | + |
| Y11_RS12425 | Y11_26361 | degS | outer membrane-stress sensor serine endopeptidase DegS | -0,57 | 0,18 | 0,42 | 0,75 | 0,99 | 0,24 | 3,9E-05 | 2,9E-02 | 1,4E-03 | 1,7E-06 | 2,2E-06 | 1,8E-02 | 0,0E+00 | 0,0E+00 | 0,0E+00 | 0,0E+00 | 0,0E+00 | 0,0E+00 | 2671797 | 2672885 | 1089 | + |
| Y11_RS12430 | Y11_26371 | murA | UDP-N-acetylglucosamine 1-carboxyvinyltransferase | 0,33 | -0,4 | -0,31 | -0,72 | -0,64 | 0,09 | 9,3E-04 | 9,1E-04 | 6,2E-03 | 7,0E-07 | 6,0E-06 | 2,6E-01 | 0,0E+00 | 0,0E+00 | 0,0E+00 | 0,0E+00 | 0,0E+00 | 0,0E+00 | 2672963 | 2674231 | 1269 | - |
| Y11_RS12435 | Y11_26381 | ibaG | BolA family iron metabolism protein IbaG | 0,29 | -0,37 | -0,31 | -0,66 | -0,61 | 0,06 | 2,4E-01 | 5,3E-02 | 1,6E-01 | 8,3E-03 | 2,7E-02 | 7,3E-01 | 0,0E+00 | 0,0E+00 | 0,0E+00 | 0,0E+00 | 0,0E+00 | 0,0E+00 | 2674353 | 2674607 | 255 | - |
| Y11_RS12440 | Y11_26391 | mlaB | lipid asymmetry maintenance protein MlaB | -0,37 | 0,69 | 0,58 | 1,07 | 0,95 | -0,11 | 5,3E-03 | 1,3E-04 | 2,1E-02 | 6,7E-09 | 7,3E-04 | 5,4E-01 | 0,0E+00 | 0,0E+00 | 0,0E+00 | 3,8E-10 | 0,0E+00 | 0,0E+00 | 2674782 | 2675123 | 342 | - |
| Y11_RS12445 | Y11_26401 | mlaC | phospholipid-binding protein MlaC | -0,6 | 0,59 | 1,18 | 1,19 | 1,78 | 0,59 | 1,7E-04 | 3,4E-05 | 9,9E-07 | 2,7E-07 | 5,6E-08 | 3,7E-05 | 0,0E+00 | 0,0E+00 | 2,3E-08 | 7,9E-09 | 3,9E-09 | 0,0E+00 | 2675126 | 2675752 | 627 | - |
| Y11_RS12450 | Y11_26411 | mlaD | outer membrane lipid asymmetry maintenance protein MlaD | -0,4 | 0,41 | 0,16 | 0,81 | 0,56 | -0,25 | 2,4E-04 | 2,4E-04 | 1,3E-01 | 7,7E-08 | 1,4E-04 | 1,9E-02 | 0,0E+00 | 0,0E+00 | 0,0E+00 | 0,0E+00 | 0,0E+00 | 0,0E+00 | 2675765 | 2676325 | 561 | - |
| Y11_RS12460 | Y11_26431 | mlaF | phospholipid ABC transporter ATP-binding protein MlaF | -0,42 | 0,34 | 0,11 | 0,76 | 0,54 | -0,22 | 7,1E-03 | 2,6E-02 | 4,1E-01 | 1,3E-04 | 1,7E-03 | 9,9E-02 | 0,0E+00 | 0,0E+00 | 0,0E+00 | 0,0E+00 | 0,0E+00 | 0,0E+00 | 2677127 | 2677930 | 804 | - |
| Y11_RS12470 | Y11_26451 | kdsD | arabinose-5-phosphate isomerase KdsD | 0,4 | 0,78 | 1,18 | 0,38 | 0,78 | 0,4 | 2,2E-03 | 3,0E-06 | 6,3E-07 | 1,1E-03 | 2,6E-05 | 3,5E-04 | 0,0E+00 | 0,0E+00 | 1,5E-08 | 0,0E+00 | 0,0E+00 | 0,0E+00 | 2679379 | 2680365 | 987 | + |
| Y11_RS12475 | Y11_26461 | kdsC | 3-deoxy-manno-octulosonate-8-phosphatase KdsC | 0,44 | 1,04 | 2,01 | 0,6 | 1,56 | 0,96 | 3,1E-03 | 7,4E-05 | 6,3E-05 | 6,2E-03 | 4,8E-04 | 1,1E-02 | 0,0E+00 | 1,6E-06 | 8,9E-07 | 0,0E+00 | 1,1E-05 | 0,0E+00 | 2680379 | 2680942 | 564 | + |
| Y11_RS12480 | Y11_26471 | lptC | LPS export ABC transporter periplasmic protein LptC | -0,06 | 0,67 | 1,25 | 0,74 | 1,31 | 0,58 | 7,6E-01 | 1,1E-02 | 1,9E-03 | 4,5E-03 | 1,1E-03 | 6,2E-02 | 0,0E+00 | 0,0E+00 | 2,2E-05 | 0,0E+00 | 2,4E-05 | 0,0E+00 | 2680939 | 2681502 | 564 | + |
| Y11_RS12485 | Y11_26481 | lptA | lipopolysaccharide ABC transporter substrate-binding protein LptA | -0,1 | 0,38 | 0,53 | 0,48 | 0,63 | 0,15 | 2,1E-01 | 1,2E-04 | 4,5E-05 | 2,0E-04 | 6,8E-05 | 6,0E-02 | 0,0E+00 | 0,0E+00 | 0,0E+00 | 0,0E+00 | 0,0E+00 | 0,0E+00 | 2681486 | 2682031 | 546 | + |
| Y11_RS12490 | Y11_26491 | lptB | LPS export ABC transporter ATP-binding protein | 0,01 | 0,46 | 0,23 | 0,45 | 0,23 | -0,23 | 9,1E-01 | 5,0E-05 | 5,0E-03 | 6,0E-05 | 6,1E-03 | 7,9E-03 | 0,0E+00 | 0,0E+00 | 0,0E+00 | 0,0E+00 | 0,0E+00 | 0,0E+00 | 2682038 | 2682763 | 726 | + |
| Y11_RS12495 | Y11_26501 | rpoN | RNA polymerase factor sigma-54 | 0,37 | 0,41 | -0,4 | 0,04 | -0,77 | -0,81 | 5,4E-04 | 3,9E-04 | 1,2E-02 | 6,0E-01 | 2,0E-04 | 1,6E-04 | 0,0E+00 | 0,0E+00 | 0,0E+00 | 0,0E+00 | 0,0E+00 | 0,0E+00 | 2682967 | 2684400 | 1434 | + |
| Y11_RS12500 | Y11_26511 | hpf | ribosome hibernation promoting factor | 0,63 | 1,92 | 3,07 | 1,29 | 2,44 | 1,15 | 1,8E-02 | 4,7E-06 | 6,1E-07 | 8,4E-05 | 3,5E-06 | 2,4E-04 | 0,0E+00 | 1,2E-07 | 1,5E-08 | 1,6E-06 | 1,3E-07 | 1,8E-05 | 2684424 | 2684711 | 288 | + |
| Y11_RS12505 | Y11_26521 | ptsN | PTS IIA-like nitrogen regulatory protein PtsN | -0,04 | -0,03 | -0,28 | 0,01 | -0,24 | -0,26 | 4,9E-01 | 7,4E-01 | 1,3E-01 | 8,2E-01 | 1,8E-01 | 1,7E-01 | 0,0E+00 | 0,0E+00 | 0,0E+00 | 0,0E+00 | 0,0E+00 | 0,0E+00 | 2684882 | 2685364 | 483 | + |
| Y11_RS12510 | Y11_26531 | rapZ | RNase adapter RapZ | -1,2 | 0,38 | 0,51 | 1,58 | 1,71 | 0,13 | 1,3E-06 | 2,0E-03 | 6,2E-04 | 5,3E-07 | 4,8E-07 | 2,7E-01 | 8,0E-08 | 0,0E+00 | 0,0E+00 | 1,4E-08 | 2,3E-08 | 0,0E+00 | 2685443 | 2686294 | 852 | + |
| Y11_RS12515 | Y11_26541 | npr | PTS phosphocarrier protein NPr | -0,51 | 0,21 | 0,01 | 0,72 | 0,51 | -0,2 | 3,4E-03 | 4,2E-01 | 9,6E-01 | 1,4E-02 | 1,0E-02 | 4,6E-01 | 0,0E+00 | 0,0E+00 | 0,0E+00 | 0,0E+00 | 0,0E+00 | 0,0E+00 | 2686295 | 2686567 | 273 | + |
| Y11_RS12520 | Y11_26561 | pyrB | aspartate carbamoyltransferase | 1,09 | 3,82 | 4,34 | 2,73 | 3,25 | 0,52 | 1,0E-06 | 3,1E-09 | 4,1E-11 | 3,5E-08 | 2,3E-10 | 5,8E-03 | 6,3E-08 | 2,5E-10 | 1,4E-11 | 1,4E-09 | 7,2E-11 | 0,0E+00 | 2687689 | 2688624 | 936 | + |

| | | | | | | | | | | | | | | | | | | | | | | | | | |
|---|---|---|---|---|---|---|---|---|---|---|---|---|---|---|---|---|---|---|---|---|---|---|---|---|---|
| Y11_RS12525 | Y11_26571 | pyrI | aspartate carbamoyltransferase regulatory subunit | 0,97 | 2,09 | 2,25 | 1,13 | 1,29 | 0,16 | 1,9E-03 | 2,7E-06 | 2,8E-06 | 1,3E-04 | 9,9E-05 | 2,9E-01 | 0,0E+00 | 7,2E-08 | 5,5E-08 | 2,3E-06 | 2,6E-06 | 0,0E+00 | 2688636 | 2689100 | 465 | + |
| Y11_RS12530 | Y11_26581 | ridA | 2-iminobutanoate/2-iminopropanoate deaminase | 0,08 | 0,97 | 1,19 | 0,89 | 1,11 | 0,23 | 5,0E-01 | 2,8E-07 | 1,3E-05 | 4,6E-05 | 1,0E-04 | 1,1E-01 | 0,0E+00 | 0,0E+00 | 2,2E-07 | 0,0E+00 | 2,7E-06 | 0,0E+00 | 2689594 | 2689980 | 387 | + |
| Y11_RS12555 | Y11_26641 | treC | alpha,alpha-phosphotrehalase | 0,37 | 0,85 | 1,34 | 0,49 | 0,97 | 0,48 | 4,5E-06 | 2,1E-06 | 3,9E-11 | 1,2E-04 | 3,3E-10 | 1,0E-04 | 0,0E+00 | 0,0E+00 | 1,4E-11 | 0,0E+00 | 0,0E+00 | 0,0E+00 | 2694525 | 2696186 | 1662 | - |
| Y11_RS12560 | Y11_26651 | treB | PTS trehalose transporter subunit IIBC | 0,35 | 1,13 | 2,35 | 0,78 | 2 | 1,21 | 5,5E-03 | 8,9E-06 | 3,9E-06 | 1,3E-04 | 1,3E-05 | 6,0E-04 | 0,0E+00 | 2,2E-07 | 7,3E-08 | 0,0E+00 | 4,2E-07 | 4,2E-05 | 2696252 | 2697667 | 1416 | - |
| Y11_RS12565 | Y11_26661 | treR | trehalose operon repressor TreR | -0,81 | 0,98 | 0,62 | 1,79 | 1,43 | -0,36 | 3,6E-06 | 7,2E-08 | 2,6E-03 | 8,6E-09 | 1,3E-05 | 3,8E-02 | 0,0E+00 | 0,0E+00 | 0,0E+00 | 4,7E-10 | 4,3E-07 | 0,0E+00 | 2697884 | 2698831 | 948 | - |
| Y11_RS12570 | Y11_26671 | mgtA | magnesium-translocating P-type ATPase | 0,18 | -0,02 | 0,05 | -0,2 | -0,13 | 0,07 | 5,7E-01 | 9,3E-01 | 8,2E-01 | 4,9E-01 | 6,6E-01 | 7,1E-01 | 0,0E+00 | 0,0E+00 | 0,0E+00 | 0,0E+00 | 0,0E+00 | 0,0E+00 | 2699603 | 2702323 | 2721 | + |
| Y11_RS12575 | | | type II toxin-antitoxin system RelE/ParE family toxin | 0 | -0,9 | 0 | -0,87 | 0 | 0,17 | 0,0E+00 | 2,3E-03 | 0,0E+00 | 2,7E-03 | 0,0E+00 | 6,0E-01 | 0,0E+00 | 0,0E+00 | 0,0E+00 | 0,0E+00 | 0,0E+00 | 0,0E+00 | 2702403 | 2702681 | 279 | + |
| Y11_RS12580 | Y11_26691 | | HigA family addiction module antitoxin | 0,28 | -0,57 | -0,07 | -0,85 | -0,35 | 0,49 | 1,4E-01 | 1,0E-03 | 3,7E-01 | 2,0E-03 | 6,9E-02 | 2,0E-03 | 0,0E+00 | 0,0E+00 | 0,0E+00 | 0,0E+00 | 0,0E+00 | 0,0E+00 | 2702681 | 2702986 | 306 | + |
| Y11_RS12590 | Y11_26711 | | lactonase family protein | 1,48 | -1,95 | -1,79 | -3,44 | -3,27 | 0,17 | 3,8E-06 | 1,4E-06 | 1,7E-06 | 1,4E-09 | 8,3E-10 | 2,3E-01 | 2,2E-07 | 3,9E-08 | 3,7E-08 | 1,1E-10 | 1,5E-10 | 0,0E+00 | 2705059 | 2706300 | 1242 | - |
| Y11_RS12600 | Y11_26741 | | DgaE family pyridoxal phosphate-dependent ammonia lyase | 0,08 | 0,21 | -0,54 | 0,13 | -0,61 | -0,74 | 7,2E-01 | 5,7E-01 | 5,8E-02 | 6,6E-01 | 7,9E-04 | 3,8E-02 | 0,0E+00 | 0,0E+00 | 0,0E+00 | 0,0E+00 | 0,0E+00 | 0,0E+00 | 2707174 | 2708286 | 1113 | - |
| Y11_RS12605 | Y11_26751 | | amidohydrolase/deacetylase family metallohydrolase | 0,32 | 0,16 | 0,73 | -0,16 | 0,4 | 0,57 | 7,0E-04 | 5,9E-02 | 8,3E-06 | 2,4E-02 | 1,4E-04 | 4,6E-05 | 0,0E+00 | 0,0E+00 | 0,0E+00 | 0,0E+00 | 0,0E+00 | 0,0E+00 | 2708270 | 2709439 | 1170 | - |
| Y11_RS12625 | Y11_26791 | | glycine-rich SFCGS family protein | 0,9 | 0,15 | 1,23 | -0,75 | 0,33 | 1,08 | 1,5E-06 | 3,8E-02 | 3,4E-05 | 8,7E-06 | 6,6E-02 | 9,5E-05 | 0,0E+00 | 0,0E+00 | 5,1E-07 | 0,0E+00 | 0,0E+00 | 7,9E-06 | 2711280 | 2711645 | 366 | - |
| Y11_RS12635 | Y11_26821 | rnk | nucleoside diphosphate kinase regulator | -0,09 | -0,18 | 1,95 | -0,1 | 2,04 | 2,13 | 3,3E-01 | 8,7E-02 | 2,8E-04 | 3,7E-01 | 2,2E-04 | 1,7E-04 | 0,0E+00 | 0,0E+00 | 3,6E-06 | 0,0E+00 | 5,5E-06 | 1,3E-05 | 2712545 | 2712955 | 411 | + |
| Y11_RS12640 | Y11_26831 | cybC | cytochrome b562 | -1,26 | 0,76 | 1,34 | 2,02 | 2,6 | 0,58 | 1,6E-04 | 5,4E-05 | 3,6E-05 | 2,2E-06 | 1,9E-06 | 3,1E-03 | 6,6E-06 | 0,0E+00 | 5,4E-07 | 5,0E-08 | 7,3E-08 | 0,0E+00 | 2713003 | 2713389 | 387 | - |
| Y11_RS12650 | Y11_26851 | pmbA | metalloprotease PmbA | -0,09 | 0,97 | 0,83 | 1,06 | 0,92 | -0,14 | 8,4E-02 | 2,1E-08 | 4,1E-05 | 2,3E-08 | 2,3E-05 | 2,2E-01 | 0,0E+00 | 0,0E+00 | 0,0E+00 | 9,7E-10 | 0,0E+00 | 0,0E+00 | 2715109 | 2716449 | 1341 | - |
| Y11_RS12655 | Y11_26861 | | ribosome-associated protein | 0,48 | 0,83 | 1,03 | 0,35 | 0,55 | 0,2 | 3,8E-04 | 4,5E-05 | 4,5E-05 | 5,4E-03 | 1,8E-03 | 1,8E-01 | 0,0E+00 | 0,0E+00 | 6,5E-07 | 0,0E+00 | 0,0E+00 | 0,0E+00 | 2716618 | 2717163 | 546 | + |
| Y11_RS12660 | Y11_26871 | | barstar family protein | 0,21 | 1,57 | 2,55 | 1,35 | 2,34 | 0,99 | 4,4E-01 | 2,5E-04 | 6,4E-06 | 8,6E-05 | 1,1E-06 | 3,7E-04 | 0,0E+00 | 5,3E-06 | 1,1E-07 | 1,6E-06 | 4,5E-08 | 0,0E+00 | 2717195 | 2717461 | 267 | - |
| Y11_RS12665 | Y11_26881 | | ribonuclease Ba | 0,54 | 0,17 | 0,52 | -0,37 | -0,01 | 0,36 | 3,6E-04 | 8,0E-02 | 2,7E-02 | 5,5E-03 | 9,5E-01 | 1,1E-01 | 0,0E+00 | 0,0E+00 | 0,0E+00 | 0,0E+00 | 0,0E+00 | 0,0E+00 | 2717466 | 2717939 | 474 | - |
| Y11_RS12670 | Y11_26891 | | NAD-dependent succinate-semialdehyde dehydrogenase | 0,41 | 2,78 | 3,05 | 2,37 | 2,64 | 0,27 | 4,2E-05 | 9,8E-10 | 6,7E-08 | 2,5E-09 | 1,9E-07 | 1,6E-01 | 0,0E+00 | 1,1E-10 | 2,4E-09 | 1,7E-10 | 1,1E-08 | 0,0E+00 | 2718037 | 2719530 | 1494 | - |
| Y11_RS12690 | Y11_26931 | aaeR | HTH-type transcriptional activator AaeR | -0,07 | 0,77 | 1,53 | 0,84 | 1,6 | 0,76 | 6,8E-01 | 1,0E-04 | 1,5E-06 | 5,4E-04 | 8,9E-06 | 1,3E-04 | 0,0E+00 | 0,0E+00 | 3,3E-08 | 0,0E+00 | 2,9E-07 | 0,0E+00 | 2723081 | 2723992 | 912 | + |
| Y11_RS12745 | Y11_27051 | | helix-turn-helix domain-containing protein | 0 | -0,5 | 0 | -0,42 | 0 | -0,69 | 0,0E+00 | 5,6E-02 | 0,0E+00 | 1,5E-01 | 0,0E+00 | 4,3E-02 | 0,0E+00 | 0,0E+00 | 0,0E+00 | 0,0E+00 | 0,0E+00 | 0,0E+00 | 2739968 | 2740246 | 279 | + |
| Y11_RS12750 | Y11_27061 | | type II toxin-antitoxin system HipA family toxin | 0,12 | 1,3 | 1,66 | 1,18 | 1,53 | 0,36 | 3,4E-01 | 6,9E-06 | 5,6E-07 | 7,8E-05 | 7,7E-06 | 5,2E-02 | 0,0E+00 | 1,7E-07 | 1,4E-08 | 1,5E-06 | 2,6E-07 | 0,0E+00 | 2740243 | 2741529 | 1287 | + |
| Y11_RS12755 | Y11_27071 | tldD | metalloprotease TldD | 0,09 | -0,19 | 0,12 | -0,28 | 0,04 | 0,31 | 1,5E-02 | 2,0E-03 | 1,9E-01 | 9,9E-05 | 6,8E-01 | 7,8E-03 | 0,0E+00 | 0,0E+00 | 0,0E+00 | 0,0E+00 | 0,0E+00 | 0,0E+00 | 2741583 | 2743028 | 1446 | - |
| Y11_RS12760 | Y11_27081 | | carbon-nitrogen hydrolase family protein | 0,08 | -0,38 | 0,43 | -0,46 | 0,35 | 0,81 | 1,5E-01 | 4,7E-05 | 1,6E-05 | 1,2E-05 | 6,8E-05 | 8,4E-08 | 0,0E+00 | 0,0E+00 | 0,0E+00 | 0,0E+00 | 0,0E+00 | 0,0E+00 | 2743040 | 2743900 | 861 | - |
| Y11_RS12765 | Y11_27091 | yhdP | AsmA2 domain-containing protein YhdP | -0,04 | 0,07 | -0,28 | 0,12 | -0,23 | -0,35 | 8,8E-01 | 7,3E-01 | 1,8E-01 | 6,9E-01 | 4,0E-01 | 1,3E-01 | 0,0E+00 | 0,0E+00 | 0,0E+00 | 0,0E+00 | 0,0E+00 | 0,0E+00 | 2743897 | 2747772 | 3876 | - |
| Y11_RS12770 | Y11_27101 | rng | ribonuclease G | -0,28 | -0,03 | 0,1 | 0,25 | 0,38 | 0,13 | 1,8E-04 | 6,6E-01 | 1,3E-01 | 1,3E-03 | 1,2E-04 | 9,2E-02 | 0,0E+00 | 0,0E+00 | 0,0E+00 | 0,0E+00 | 0,0E+00 | 0,0E+00 | 2747974 | 2749443 | 1470 | - |
| Y11_RS12775 | Y11_27111 | | Maf-like protein | -0,11 | -0,06 | 0,09 | 0,04 | 0,2 | 0,16 | 4,4E-01 | 5,9E-01 | 4,0E-01 | 7,4E-01 | 1,4E-01 | 1,6E-01 | 0,0E+00 | 0,0E+00 | 0,0E+00 | 0,0E+00 | 0,0E+00 | 0,0E+00 | 2749433 | 2750026 | 594 | - |
| Y11_RS12785 | Y11_27131 | mreC | rod shape-determining protein MreC | -0,09 | -0,44 | -0,54 | -0,35 | -0,45 | -0,1 | 5,8E-01 | 1,0E-02 | 2,6E-02 | 1,6E-02 | 4,4E-02 | 5,7E-01 | 0,0E+00 | 0,0E+00 | 0,0E+00 | 0,0E+00 | 0,0E+00 | 0,0E+00 | 2750525 | 2751511 | 987 | - |
| Y11_RS12790 | Y11_27141 | mreB | rod shape-determining protein MreB | -0,37 | -0,44 | -0,84 | -0,07 | -0,47 | -0,39 | 1,8E-04 | 1,0E-04 | 5,3E-07 | 3,7E-01 | 2,2E-04 | 1,0E-03 | 0,0E+00 | 0,0E+00 | 0,0E+00 | 0,0E+00 | 0,0E+00 | 0,0E+00 | 2751611 | 2752654 | 1044 | - |
| Y11_RS12795 | Y11_27151 | csrD | RNase E specificity factor CsrD | -0,06 | 0,55 | 1,18 | 0,61 | 1,24 | 0,63 | 8,4E-01 | 4,8E-02 | 2,3E-03 | 3,5E-02 | 1,9E-03 | 1,6E-02 | 0,0E+00 | 0,0E+00 | 2,7E-05 | 0,0E+00 | 4,2E-05 | 0,0E+00 | 2752989 | 2754908 | 1920 | - |
| Y11_RS12800 | Y11_27161 | | oxidoreductase | 0,36 | 0,05 | 0,26 | -0,31 | -0,1 | 0,21 | 2,0E-04 | 4,4E-01 | 6,7E-03 | 8,2E-04 | 1,9E-01 | 2,3E-02 | 0,0E+00 | 0,0E+00 | 0,0E+00 | 0,0E+00 | 0,0E+00 | 0,0E+00 | 2755198 | 2756175 | 978 | + |
| Y11_RS12805 | Y11_27171 | msrP | protein-methionine-sulfoxide reductase catalytic subunit MsrP | 0,71 | 1,27 | 1,17 | 0,56 | 0,46 | -0,1 | 2,7E-07 | 1,8E-08 | 1,4E-08 | 1,6E-06 | 1,5E-06 | 7,5E-02 | 0,0E+00 | 9,9E-10 | 6,7E-10 | 0,0E+00 | 0,0E+00 | 0,0E+00 | 2756329 | 2757363 | 1035 | + |
| Y11_RS12810 | Y11_27181 | msrQ | protein-methionine-sulfoxide reductase heme-binding subunit MsrQ | -0,36 | 0,24 | 0,03 | 0,6 | 0,38 | -0,21 | 1,2E-01 | 3,0E-01 | 9,6E-01 | 2,9E-02 | 4,7E-01 | 6,9E-01 | 0,0E+00 | 0,0E+00 | 0,0E+00 | 0,0E+00 | 0,0E+00 | 0,0E+00 | 2757363 | 2757962 | 600 | + |
| Y11_RS12815 | Y11_27191 | aroQ | type II 3-dehydroquinate dehydratase | -0,16 | 0,84 | 1,16 | 1 | 1,32 | 0,32 | 3,3E-01 | 2,7E-04 | 1,0E-04 | 1,4E-06 | 4,2E-06 | 1,3E-02 | 0,0E+00 | 0,0E+00 | 1,4E-06 | 0,0E+00 | 1,5E-07 | 0,0E+00 | 2758171 | 2758623 | 453 | + |
| Y11_RS12820 | Y11_27201 | accB | acetyl-CoA carboxylase biotin carboxyl carrier protein | 0,24 | -0,65 | -0,79 | -0,88 | -1,02 | -0,14 | 7,1E-02 | 2,3E-04 | 1,5E-03 | 2,5E-05 | 2,8E-04 | 4,1E-01 | 0,0E+00 | 0,0E+00 | 0,0E+00 | 0,0E+00 | 7,0E-06 | 0,0E+00 | 2758656 | 2759120 | 465 | + |
| Y11_RS12825 | Y11_27211 | accC | acetyl-CoA carboxylase biotin carboxylase subunit | 0,12 | -0,52 | -0,68 | -0,63 | -0,79 | -0,16 | 5,4E-02 | 4,9E-05 | 5,3E-07 | 2,9E-05 | 9,1E-07 | 6,0E-02 | 0,0E+00 | 0,0E+00 | 0,0E+00 | 0,0E+00 | 0,0E+00 | 0,0E+00 | 2759132 | 2760481 | 1350 | + |
| Y11_RS12840 | Y11_27241 | prmA | 50S ribosomal protein L11 methyltransferase | 0,05 | 0,86 | 0,96 | 0,82 | 0,91 | 0,1 | 4,5E-01 | 4,3E-07 | 2,3E-07 | 7,8E-08 | 4,5E-08 | 8,0E-02 | 0,0E+00 | 0,0E+00 | 0,0E+00 | 0,0E+00 | 0,0E+00 | 0,0E+00 | 2762398 | 2763279 | 882 | + |
| Y11_RS12845 | Y11_27251 | dusB | tRNA dihydrouridine synthase DusB | 0,01 | -1,31 | -1,62 | -1,32 | -1,63 | -0,31 | 9,3E-01 | 1,7E-06 | 8,4E-05 | 3,7E-06 | 9,8E-05 | 2,1E-01 | 0,0E+00 | 4,7E-08 | 1,2E-06 | 8,2E-08 | 2,6E-06 | 0,0E+00 | 2763722 | 2764687 | 966 | + |
| Y11_RS12850 | Y11_27261 | fis | DNA-binding transcriptional regulator Fis | -0,31 | -1,7 | -2,59 | -1,38 | -2,28 | -0,89 | 4,2E-02 | 2,2E-04 | 5,8E-07 | 8,0E-04 | 1,4E-06 | 1,6E-02 | 0,0E+00 | 4,7E-06 | 1,4E-08 | 1,3E-05 | 5,5E-08 | 0,0E+00 | 2764708 | 2765004 | 297 | + |
| Y11_RS12860 | Y11_27281 | | glycine zipper 2TM domain-containing protein | -0,02 | 2,35 | 3,15 | 2,37 | 3,17 | 0,8 | 8,8E-01 | 2,1E-07 | 7,5E-09 | 1,2E-07 | 3,6E-09 | 4,9E-04 | 0,0E+00 | 7,7E-09 | 4,3E-10 | 3,9E-09 | 4,3E-10 | 0,0E+00 | 2766878 | 2767348 | 471 | + |
| Y11_RS12865 | Y11_27291 | cspA | RNA chaperone/antiterminator CspA | 0 | 0,61 | 0,64 | -15 | -15 | -0,01 | 0,0E+00 | 1,1E-02 | 9,9E-03 | 0,0E+00 | 0,0E+00 | 9,6E-01 | 0,0E+00 | 0,0E+00 | 0,0E+00 | 0,0E+00 | 0,0E+00 | 0,0E+00 | 2767730 | 2767942 | 213 | + |
| Y11_RS12890 | Y11_27361 | | PhoPQ-regulated protein | 1,5 | 15 | 15 | 0 | 0 | 0 | 2,0E-05 | 0,0E+00 | 0,0E+00 | 0,0E+00 | 0,0E+00 | 0,0E+00 | 9,9E-07 | 0,0E+00 | 0,0E+00 | 0,0E+00 | 0,0E+00 | 0,0E+00 | 2773271 | 2774809 | 1539 | - |
| Y11_RS12925 | Y11_27431 | | LysR family transcriptional regulator | 0,05 | 1,71 | 1,84 | 1,66 | 1,79 | 0,13 | 7,3E-01 | 2,2E-07 | 7,1E-06 | 3,5E-06 | 2,5E-05 | 4,9E-01 | 0,0E+00 | 8,0E-09 | 1,2E-07 | 7,8E-08 | 7,3E-07 | 0,0E+00 | 2780844 | 2781770 | 927 | + |
| Y11_RS12930 | Y11_27441 | | Na+/H+ antiporter | -0,63 | 0,36 | -0,22 | 0,99 | 0,42 | -0,57 | 1,9E-04 | 1,9E-02 | 1,7E-01 | 9,4E-06 | 1,1E-02 | 4,3E-03 | 0,0E+00 | 0,0E+00 | 0,0E+00 | 0,0E+00 | 0,0E+00 | 0,0E+00 | 2781777 | 2783426 | 1650 | - |
| Y11_RS12935 | Y11_27451 | | NCS2 family permease | 1,45 | 1,66 | 1,56 | 0,21 | 0,11 | -0,1 | 3,1E-06 | 2,9E-08 | 4,2E-06 | 9,3E-02 | 5,2E-01 | 4,5E-01 | 1,8E-07 | 1,5E-09 | 7,7E-08 | 0,0E+00 | 0,0E+00 | 0,0E+00 | 2783641 | 2785008 | 1368 | - |
| Y11_RS12940 | Y11_27461 | | glutathione S-transferase | 0,35 | 2,99 | 3,26 | 2,64 | 2,91 | 0,27 | 1,6E-02 | 4,2E-11 | 9,4E-09 | 1,1E-08 | 1,0E-07 | 8,0E-02 | 0,0E+00 | 1,3E-11 | 5,0E-10 | 5,8E-10 | 6,5E-09 | 0,0E+00 | 2785369 | 2786040 | 672 | - |
| Y11_RS12945 | Y11_27471 | | helix-turn-helix transcriptional regulator | -0,14 | 0,18 | 0,02 | 0,32 | 0,16 | -0,16 | 2,9E-01 | 2,4E-01 | 9,1E-01 | 2,4E-03 | 2,2E-01 | 2,8E-01 | 0,0E+00 | 0,0E+00 | 0,0E+00 | 0,0E+00 | 0,0E+00 | 0,0E+00 | 2786335 | 2786634 | 300 | + |
| Y11_RS12950 | Y11_27481 | | type II toxin-antitoxin system HipA family toxin | -0,16 | 0,62 | 1,07 | 0,78 | 1,23 | 0,45 | 1,8E-01 | 2,2E-04 | 3,7E-05 | 2,9E-05 | 1,0E-05 | 4,9E-03 | 0,0E+00 | 0,0E+00 | 5,5E-07 | 0,0E+00 | 3,4E-07 | 0,0E+00 | 2786636 | 2787964 | 1329 | + |
| Y11_RS12955 | Y11_27491 | ssb1 | single-stranded DNA-binding protein SSB1 | -0,1 | 0,49 | 1,14 | 0,59 | 1,24 | 0,65 | 4,5E-02 | 2,0E-06 | 3,5E-09 | 3,2E-06 | 1,1E-08 | 1,5E-06 | 0,0E+00 | 0,0E+00 | 2,4E-10 | 0,0E+00 | 1,1E-09 | 0,0E+00 | 2788053 | 2788601 | 549 | - |
| Y11_RS12960 | Y11_27501 | uvrA | excinuclease ABC subunit UvrA | -0,67 | -0,25 | 0,18 | 0,42 | 0,85 | 0,43 | 2,1E-06 | 8,7E-04 | 2,7E-02 | 2,9E-05 | 1,8E-06 | 1,4E-04 | 0,0E+00 | 0,0E+00 | 0,0E+00 | 0,0E+00 | 0,0E+00 | 0,0E+00 | 2789130 | 2791961 | 2832 | + |
| Y11_RS12970 | Y11_27521 | | aspartate/tyrosine/aromatic aminotransferase | 0,29 | 0,43 | 1,06 | 0,14 | 0,77 | 0,63 | 7,8E-04 | 7,4E-06 | 4,9E-08 | 6,4E-02 | 6,5E-06 | 1,1E-05 | 0,0E+00 | 0,0E+00 | 1,8E-09 | 0,0E+00 | 0,0E+00 | 0,0E+00 | 2792691 | 2793884 | 1194 | - |
| Y11_RS12975 | Y11_27531 | alr | alanine racemase | 0,11 | -0,25 | -0,14 | -0,36 | -0,25 | 0,11 | 1,8E-01 | 1,2E-02 | 3,7E-01 | 6,4E-03 | 1,5E-01 | 5,0E-01 | 0,0E+00 | 0,0E+00 | 0,0E+00 | 0,0E+00 | 0,0E+00 | 0,0E+00 | 2793942 | 2795021 | 1080 | - |
| Y11_RS12980 | Y11_27541 | dnaB | replicative DNA helicase | 0,04 | -0,04 | -0,16 | -0,07 | -0,2 | -0,13 | 6,5E-01 | 5,6E-01 | 5,2E-02 | 2,8E-01 | 2,4E-02 | 6,7E-02 | 0,0E+00 | 0,0E+00 | 0,0E+00 | 0,0E+00 | 0,0E+00 | 0,0E+00 | 2795051 | 2796457 | 1407 | - |
| Y11_RS12985 | Y11_27551 | | quinone oxidoreductase | 0,07 | -0,74 | -0,66 | -0,81 | -0,73 | 0,08 | 4,5E-01 | 6,7E-05 | 4,2E-04 | 3,0E-06 | 4,6E-05 | 4,6E-01 | 0,0E+00 | 0,0E+00 | 0,0E+00 | 0,0E+00 | 0,0E+00 | 0,0E+00 | 2796630 | 2797613 | 984 | + |
| Y11_RS13000 | Y11_27581 | dusA | tRNA dihydrouridine(20/20a) synthase DusA | -0,76 | -0,97 | -1,45 | -0,21 | -0,7 | -0,49 | 1,4E-07 | 1,2E-08 | 8,2E-08 | 4,5E-03 | 4,2E-05 | 4,6E-04 | 0,0E+00 | 0,0E+00 | 2,8E-09 | 0,0E+00 | 0,0E+00 | 0,0E+00 | 2798904 | 2799950 | 1047 | - |
| Y11_RS13010 | Y11_27601 | zur | zinc uptake transcriptional repressor Zur | 0,39 | -0,06 | 0,48 | -0,45 | 0,09 | 0,54 | 1,7E-04 | 3,6E-01 | 1,1E-02 | 7,9E-05 | 5,4E-01 | 6,2E-03 | 0,0E+00 | 0,0E+00 | 0,0E+00 | 0,0E+00 | 0,0E+00 | 0,0E+00 | 2802525 | 2803034 | 510 | + |
| Y11_RS13015 | Y11_27621 | | CsbD family protein | 1,01 | 0,4 | 2,34 | -0,61 | 1,33 | 1,93 | 2,4E-03 | 5,5E-02 | 1,0E-03 | 5,1E-02 | 3,0E-02 | 3,9E-03 | 8,4E-05 | 0,0E+00 | 1,2E-05 | 0,0E+00 | 6,2E-04 | 2,4E-04 | 2803319 | 2803528 | 210 | - |
| Y11_RS13020 | Y11_27631 | lexA | transcriptional repressor LexA | 0,18 | 0,72 | 0,98 | 0,54 | 0,79 | 0,26 | 2,8E-02 | 9,4E-07 | 4,1E-06 | 1,7E-05 | 2,5E-05 | 1,3E-02 | 0,0E+00 | 0,0E+00 | 0,0E+00 | 0,0E+00 | 0,0E+00 | 0,0E+00 | 2803837 | 2804445 | 609 | - |
| Y11_RS13030 | Y11_27651 | plsB | glycerol-3-phosphate 1-O-acyltransferase PlsB | 0,09 | 0,65 | 0,93 | 0,57 | 0,85 | 0,28 | 6,3E-02 | 1,0E-05 | 2,2E-08 | 5,1E-05 | 1,9E-07 | 5,7E-03 | 0,0E+00 | 0,0E+00 | 0,0E+00 | 0,0E+00 | 0,0E+00 | 0,0E+00 | 2805067 | 2807535 | 2469 | + |
| Y11_RS13040 | Y11_27671 | ubiC | chorismate lyase | 0,38 | -1,11 | -0,83 | -1,5 | -1,21 | 0,28 | 7,8E-02 | 4,4E-06 | 8,9E-05 | 7,7E-05 | 4,1E-04 | 6,8E-02 | 0,0E+00 | 1,1E-07 | 0,0E+00 | 1,4E-06 | 9,7E-06 | 0,0E+00 | 2808484 | 2808990 | 507 | - |
| Y11_RS13050 | Y11_27691 | malM | maltose operon protein MalM | 0,75 | -1,46 | -1,36 | -2,2 | -2,1 | 0,1 | 6,8E-03 | 2,9E-06 | 4,9E-05 | 3,3E-06 | 1,5E-05 | 5,5E-01 | 0,0E+00 | 7,7E-08 | 7,1E-07 | 7,4E-08 | 4,7E-07 | 0,0E+00 | 2812184 | 2813101 | 918 | - |
| Y11_RS13055 | Y11_27701 | | maltoporin | 1,81 | -1,23 | -1,29 | -3,04 | -3,11 | -0,06 | 4,6E-07 | 1,3E-05 | 1,2E-05 | 7,1E-09 | 8,6E-09 | 6,5E-01 | 3,3E-08 | 3,2E-07 | 2,0E-07 | 4,0E-10 | 8,6E-10 | 0,0E+00 | 2813232 | 2814530 | 1299 | - |
| Y11_RS13060 | Y11_27711 | malK | maltose/maltodextrin ABC transporter ATP-binding protein MalK | 0,06 | -1,56 | -1,44 | -1,63 | -1,51 | 0,12 | 5,2E-01 | 4,0E-08 | 1,4E-07 | 6,3E-08 | 1,9E-07 | 1,4E-01 | 0,0E+00 | 1,9E-09 | 4,3E-09 | 2,3E-09 | 1,1E-08 | 0,0E+00 | 2814602 | 2815717 | 1116 | - |
| Y11_RS13065 | Y11_27721 | malE | maltose/maltodextrin ABC transporter substrate-binding protein MalE | 1,24 | 0,65 | 1,26 | -0,59 | 0,03 | 0,61 | 3,5E-09 | 2,1E-05 | 4,9E-06 | 6,4E-05 | 8,4E-01 | 1,8E-03 | 6,7E-10 | 0,0E+00 | 8,8E-08 | 0,0E+00 | 0,0E+00 | 0,0E+00 | 2816362 | 2817573 | 1212 | + |
| Y11_RS13070 | Y11_27731 | malF | maltose ABC transporter permease MalF | 0,42 | -1,25 | -1,57 | -1,68 | -2,02 | -0,32 | 7,4E-03 | 4,4E-04 | 5,8E-05 | 2,5E-05 | 4,6E-06 | 2,5E-01 | 0,0E+00 | 9,1E-06 | 8,3E-07 | 4,9E-07 | 1,6E-07 | 0,0E+00 | 2817929 | 2819506 | 1578 | + |
| Y11_RS13085 | Y11_27761 | pgi | glucose-6-phosphate isomerase | 0,11 | 0,79 | 0,4 | 0,68 | 0,28 | -0,4 | 5,8E-03 | 2,1E-11 | 1,8E-04 | 1,1E-08 | 2,5E-03 | 1,7E-04 | 0,0E+00 | 0,0E+00 | 0,0E+00 | 0,0E+00 | 0,0E+00 | 0,0E+00 | 2821426 | 2823072 | 1647 | - |
| Y11_RS13090 | Y11_27771 | lysC | lysine-sensitive aspartokinase 3 | 0,02 | -0,89 | -1,27 | -0,92 | -1,29 | -0,37 | 6,4E-01 | 4,7E-08 | 3,2E-07 | 7,2E-08 | 3,4E-07 | 1,9E-03 | 0,0E+00 | 0,0E+00 | 8,7E-09 | 0,0E+00 | 1,7E-08 | 0,0E+00 | 2823385 | 2824800 | 1416 | + |
| Y11_RS13110 | Y11_27811 | metH | methionine synthase | -0,62 | 2,08 | 1,8 | 2,7 | 2,41 | -0,28 | 2,4E-07 | 9,5E-12 | 1,9E-08 | 3,0E-13 | 1,5E-09 | 7,0E-03 | 0,0E+00 | 6,0E-12 | 8,6E-10 | 1,2E-12 | 2,3E-10 | 0,0E+00 | 2832849 | 2836544 | 3696 | - |
| Y11_RS13115 | Y11_27821 | iclR | glyoxylate bypass operon transcriptional repressor IclR | 0,71 | 0,19 | 0,87 | -0,52 | 0,16 | 0,68 | 5,9E-05 | 1,2E-01 | 1,7E-05 | 1,6E-03 | 1,5E-01 | 3,3E-04 | 0,0E+00 | 0,0E+00 | 0,0E+00 | 0,0E+00 | 0,0E+00 | 0,0E+00 | 2836749 | 2837588 | 840 | + |

| | | | | | | | | | | | | | | | | | | | | | | | | | |
|---|---|---|---|---|---|---|---|---|---|---|---|---|---|---|---|---|---|---|---|---|---|---|---|---|---|
| Y11_RS13125 | Y11_27841 | aceA | isocitrate lyase | 1,29 | -0,69 | 1,56 | -1,98 | 0,27 | 2,25 | 4,9E-06 | 1,6E-02 | 3,5E-05 | 1,5E-05 | 1,5E-01 | 2,4E-05 | 2,7E-07 | 0,0E+00 | 5,3E-07 | 3,0E-07 | 0,0E+00 | 2,5E-06 | 2839686 | 2840993 | 1308 | - |
| Y11_RS13130 | Y11_27851 | aceB | malate synthase A | 0 | -15 | 0 | -15 | 0 | 3,63 | 0,0E+00 | 0,0E+00 | 0,0E+00 | 0,0E+00 | 0,0E+00 | 1,3E-05 | 0,0E+00 | 0,0E+00 | 0,0E+00 | 0,0E+00 | 0,0E+00 | 1,5E-06 | 2841166 | 2842764 | 1599 | - |
| Y11_RS13170 | Y11_27891 | murI | glutamate racemase | -0,62 | 0,33 | 0,59 | 0,95 | 1,21 | 0,26 | 3,1E-05 | 1,9E-03 | 1,8E-03 | 9,5E-09 | 5,0E-06 | 4,8E-02 | 0,0E+00 | 0,0E+00 | 0,0E+00 | 0,0E+00 | 1,7E-07 | 0,0E+00 | 2851021 | 2851884 | 864 | - |
| Y11_RS13175 | Y11_27901 | btuB | TonB-dependent vitamin B12 receptor BtuB | 1 | 0,57 | -0,46 | -0,43 | -1,46 | -1,03 | 4,4E-07 | 6,1E-05 | 1,2E-02 | 7,7E-04 | 8,2E-06 | 1,2E-04 | 0,0E+00 | 0,0E+00 | 0,0E+00 | 0,0E+00 | 2,7E-07 | 9,6E-06 | 2851829 | 2853727 | 1899 | - |
| Y11_RS13180 | Y11_27911 | trmA | tRNA (uridine(54)-C5)-methyltransferase TrmA | 0,73 | -0,99 | -0,6 | -1,72 | -1,33 | 0,39 | 2,0E-06 | 8,3E-08 | 7,3E-05 | 1,3E-08 | 6,8E-07 | 2,9E-03 | 0,0E+00 | 0,0E+00 | 0,0E+00 | 6,2E-10 | 3,0E-08 | 0,0E+00 | 2854153 | 2855256 | 1104 | + |
| Y11_RS13190 | Y11_27931 | fabR | HTH-type transcriptional repressor FabR | -1,04 | 0,4 | 0,67 | 1,45 | 1,72 | 0,27 | 1,5E-07 | 6,4E-04 | 6,1E-04 | 2,0E-07 | 1,2E-06 | 8,9E-02 | 1,2E-08 | 0,0E+00 | 0,0E+00 | 6,2E-09 | 4,9E-08 | 0,0E+00 | 2855985 | 2856617 | 633 | - |
| Y11_RS13195 | Y11_27941 | sthA | Si-specific NAD(P)(+) transhydrogenase | -0,23 | -0,07 | 0,23 | 0,16 | 0,47 | 0,3 | 1,6E-04 | 3,3E-02 | 4,1E-03 | 1,3E-03 | 6,8E-05 | 7,8E-04 | 0,0E+00 | 0,0E+00 | 0,0E+00 | 0,0E+00 | 0,0E+00 | 0,0E+00 | 2856835 | 2858235 | 1401 | + |
| Y11_RS13200 | Y11_27951 | oxyR | DNA-binding transcriptional regulator OxyR | 0,12 | 0,24 | -0,11 | 0,12 | -0,23 | -0,35 | 1,1E-02 | 3,1E-04 | 2,4E-01 | 3,2E-02 | 3,1E-02 | 4,2E-03 | 0,0E+00 | 0,0E+00 | 0,0E+00 | 0,0E+00 | 0,0E+00 | 0,0E+00 | 2858218 | 2859150 | 933 | - |
| Y11_RS13205 | Y11_27961 | | glutathione peroxidase | 0 | 0 | -15 | 0 | -15 | -1,89 | 0,0E+00 | 0,0E+00 | 0,0E+00 | 0,0E+00 | 0,0E+00 | 4,5E-06 | 0,0E+00 | 0,0E+00 | 0,0E+00 | 0,0E+00 | 0,0E+00 | 5,9E-07 | 2859298 | 2860029 | 732 | + |
| Y11_RS13210 | Y11_27971 | | dihydrolipoyl dehydrogenase | -0,44 | -1,33 | -1,09 | -0,89 | -0,65 | 0,24 | 3,7E-05 | 8,7E-11 | 3,7E-06 | 1,0E-07 | 2,6E-04 | 3,6E-02 | 0,0E+00 | 2,0E-11 | 7,0E-08 | 0,0E+00 | 0,0E+00 | 0,0E+00 | 2860322 | 2861770 | 1449 | + |
| Y11_RS13255 | Y11_28061 | argH | argininosuccinate lyase | 0,05 | -1,04 | -1,06 | -1,1 | -1,12 | -0,02 | 5,6E-01 | 2,7E-05 | 2,3E-05 | 1,0E-05 | 8,7E-06 | 9,0E-01 | 0,0E+00 | 6,2E-07 | 3,6E-07 | 2,1E-07 | 2,9E-07 | 0,0E+00 | 2872431 | 2873804 | 1374 | - |
| Y11_RS13260 | Y11_28081 | argB | acetylglutamate kinase | 0,06 | -1,13 | -1,24 | -1,2 | -1,31 | -0,11 | 2,7E-01 | 7,9E-07 | 4,2E-07 | 2,0E-07 | 1,1E-07 | 2,9E-01 | 0,0E+00 | 2,4E-08 | 1,1E-08 | 6,0E-09 | 6,8E-09 | 0,0E+00 | 2873976 | 2874752 | 777 | - |
| Y11_RS13265 | Y11_28091 | argC | N-acetyl-gamma-glutamyl-phosphate reductase | -0,03 | -0,42 | -1,05 | -0,38 | -1,02 | -0,63 | 8,6E-01 | 5,5E-02 | 8,8E-04 | 1,2E-03 | 1,4E-05 | 9,5E-04 | 0,0E+00 | 0,0E+00 | 1,1E-05 | 0,0E+00 | 4,4E-07 | 0,0E+00 | 2874828 | 2875832 | 1005 | - |
| Y11_RS13270 | Y11_28101 | argE | acetylornithine deacetylase | 0,1 | -0,27 | -0,64 | -0,37 | -0,74 | -0,37 | 3,7E-02 | 1,1E-03 | 5,3E-04 | 1,1E-04 | 2,0E-04 | 1,5E-02 | 0,0E+00 | 0,0E+00 | 0,0E+00 | 0,0E+00 | 0,0E+00 | 0,0E+00 | 2876080 | 2877258 | 1179 | + |
| Y11_RS13275 | Y11_28121 | ppc | phosphoenolpyruvate carboxylase | 0,33 | -0,17 | -1,08 | -0,51 | -1,42 | -0,91 | 6,6E-06 | 1,7E-03 | 1,2E-05 | 3,1E-06 | 1,8E-06 | 5,2E-05 | 0,0E+00 | 0,0E+00 | 1,9E-07 | 0,0E+00 | 7,0E-08 | 0,0E+00 | 2877482 | 2880121 | 2640 | + |
| Y11_RS13280 | Y11_28131 | metF | methylenetetrahydrofolate reductase | 1,1 | 1,98 | 15 | 0,88 | 15 | 15 | 1,1E-04 | 5,9E-09 | 0,0E+00 | 4,9E-04 | 0,0E+00 | 0,0E+00 | 4,8E-06 | 4,2E-10 | 0,0E+00 | 0,0E+00 | 0,0E+00 | 0,0E+00 | 2880966 | 2881850 | 885 | - |
| Y11_RS13285 | Y11_28141 | | bifunctional aspartate kinase/homoserine dehydrogenase II | -0,21 | 0,5 | 0,46 | 0,71 | 0,67 | -0,04 | 2,0E-03 | 4,8E-07 | 1,0E-06 | 7,1E-07 | 1,1E-06 | 3,8E-01 | 0,0E+00 | 0,0E+00 | 0,0E+00 | 0,0E+00 | 0,0E+00 | 0,0E+00 | 2882092 | 2884527 | 2436 | - |
| Y11_RS13290 | Y11_28151 | metB | cystathionine gamma-synthase | -0,05 | 0,29 | 0,3 | 0,34 | 0,36 | 0,02 | 5,5E-01 | 2,1E-02 | 9,0E-03 | 9,8E-04 | 7,8E-05 | 8,2E-01 | 0,0E+00 | 0,0E+00 | 0,0E+00 | 0,0E+00 | 0,0E+00 | 0,0E+00 | 2884530 | 2885690 | 1161 | - |
| Y11_RS13295 | Y11_28161 | metJ | met regulon transcriptional regulator MetJ | 0,21 | 1,08 | 1,12 | 0,87 | 0,91 | 0,04 | 1,9E-02 | 2,9E-07 | 1,2E-05 | 1,0E-06 | 4,7E-05 | 7,1E-01 | 0,0E+00 | 1,0E-08 | 2,0E-07 | 0,0E+00 | 0,0E+00 | 0,0E+00 | 2886066 | 2886383 | 318 | + |
| Y11_RS13300 | Y11_28171 | rpmE | 50S ribosomal protein L31 | 0,4 | -0,75 | -1,18 | -1,15 | -1,58 | -0,43 | 5,9E-04 | 6,0E-06 | 3,6E-05 | 2,0E-07 | 4,0E-06 | 1,7E-02 | 0,0E+00 | 0,0E+00 | 5,4E-07 | 6,0E-09 | 1,4E-07 | 0,0E+00 | 2886476 | 2886691 | 216 | - |
| Y11_RS13305 | Y11_28181 | priA | primosomal protein N' | -0,46 | 0,94 | 1,06 | 1,39 | 1,52 | 0,12 | 4,0E-02 | 3,1E-04 | 3,5E-05 | 4,7E-05 | 8,9E-06 | 3,0E-01 | 0,0E+00 | 0,0E+00 | 5,3E-07 | 8,9E-07 | 2,9E-07 | 0,0E+00 | 2886939 | 2889137 | 2199 | + |
| Y11_RS13310 | Y11_28191 | cytR | DNA-binding transcriptional regulator CytR | -0,35 | 0,29 | 0,85 | 0,64 | 1,2 | 0,56 | 1,1E-03 | 4,3E-03 | 1,2E-04 | 1,2E-06 | 4,4E-06 | 1,0E-03 | 0,0E+00 | 0,0E+00 | 0,0E+00 | 0,0E+00 | 1,6E-07 | 0,0E+00 | 2889379 | 2890407 | 1029 | + |
| Y11_RS13315 | Y11_28201 | ftsN | cell division protein FtsN | -0,15 | 0,18 | -0,31 | 0,33 | -0,16 | -0,49 | 7,0E-02 | 8,8E-02 | 7,7E-02 | 6,0E-03 | 3,3E-01 | 1,6E-02 | 0,0E+00 | 0,0E+00 | 0,0E+00 | 0,0E+00 | 0,0E+00 | 0,0E+00 | 2890473 | 2891342 | 870 | + |
| Y11_RS13320 | Y11_28211 | hslV | ATP-dependent protease subunit HslV | -0,05 | 0,89 | 1,19 | 0,94 | 1,24 | 0,3 | 5,5E-01 | 3,0E-06 | 1,6E-06 | 3,3E-06 | 1,7E-06 | 1,2E-02 | 0,0E+00 | 0,0E+00 | 3,5E-08 | 0,0E+00 | 6,5E-08 | 0,0E+00 | 2891442 | 2891966 | 525 | + |
| Y11_RS13325 | Y11_28221 | hslU | HslU--HslV peptidase ATPase subunit | 0,01 | 0,99 | 0,92 | 0,98 | 0,91 | -0,07 | 9,2E-01 | 8,1E-08 | 5,7E-08 | 1,5E-06 | 1,7E-06 | 2,7E-01 | 0,0E+00 | 0,0E+00 | 0,0E+00 | 0,0E+00 | 0,0E+00 | 0,0E+00 | 2892003 | 2893334 | 1332 | + |
| Y11_RS13335 | Y11_28241 | rraA | ribonuclease E activity regulator RraA | -0,48 | 0,76 | 1,65 | 1,24 | 2,13 | 0,89 | 2,4E-03 | 2,6E-05 | 6,0E-10 | 1,2E-05 | 4,3E-08 | 6,2E-06 | 0,0E+00 | 0,0E+00 | 7,5E-11 | 2,5E-07 | 3,1E-09 | 0,0E+00 | 2894548 | 2895033 | 486 | + |
| Y11_RS13340 | Y11_28251 | zapB | cell division protein ZapB | -0,86 | 2,13 | 1,75 | 2,98 | 2,6 | -0,38 | 7,3E-09 | 5,4E-08 | 1,1E-05 | 5,2E-09 | 5,8E-07 | 1,1E-01 | 0,0E+00 | 2,4E-09 | 1,8E-07 | 3,0E-10 | 2,7E-08 | 0,0E+00 | 2895170 | 2895409 | 240 | - |
| Y11_RS13350 | Y11_28261 | | aquaporin family protein | -1,23 | 0,43 | -0,22 | 1,65 | 1,01 | -0,65 | 3,3E-04 | 1,0E-01 | 4,8E-01 | 5,0E-06 | 3,1E-03 | 3,9E-02 | 1,3E-05 | 0,0E+00 | 0,0E+00 | 1,1E-07 | 6,8E-05 | 0,0E+00 | 2896004 | 2896852 | 849 | + |
| Y11_RS13355 | Y11_28271 | glpK | glycerol kinase GlpK | -0,52 | 0,96 | 1,16 | 1,48 | 1,68 | 0,2 | 1,3E-07 | 4,0E-08 | 1,8E-08 | 6,7E-10 | 5,6E-10 | 1,2E-02 | 0,0E+00 | 0,0E+00 | 8,3E-10 | 6,5E-11 | 1,2E-10 | 0,0E+00 | 2896929 | 2898452 | 1524 | + |
| Y11_RS13360 | Y11_28281 | glpX | class II fructose-bisphosphatase | -0,55 | 3,5 | 2,93 | 4,05 | 3,47 | -0,58 | 1,1E-05 | 1,9E-11 | 3,3E-08 | 9,4E-13 | 6,2E-09 | 3,6E-03 | 0,0E+00 | 7,7E-12 | 1,4E-09 | 1,9E-12 | 6,5E-10 | 0,0E+00 | 2898634 | 2899644 | 1011 | + |
| Y11_RS13385 | Y11_28331 | fpr | ferredoxin--NADP(+) reductase | 0,33 | 1,02 | 1,57 | 0,69 | 1,24 | 0,55 | 8,8E-05 | 3,1E-08 | 3,1E-10 | 1,2E-06 | 4,9E-09 | 5,0E-06 | 0,0E+00 | 1,5E-09 | 4,4E-11 | 0,0E+00 | 5,4E-10 | 0,0E+00 | 2909455 | 2910201 | 747 | + |
| Y11_RS13395 | Y11_28351 | | DUF1454 family protein | 0,36 | 0,42 | 0,8 | 0,07 | 0,44 | 0,37 | 1,2E-02 | 6,2E-04 | 1,9E-04 | 6,2E-01 | 2,5E-02 | 2,6E-02 | 0,0E+00 | 0,0E+00 | 0,0E+00 | 0,0E+00 | 0,0E+00 | 0,0E+00 | 2910856 | 2911461 | 606 | + |
| Y11_RS13400 | Y11_28361 | tpiA | triose-phosphate isomerase | -0,19 | 1,52 | 1,44 | 1,71 | 1,63 | -0,08 | 2,3E-02 | 2,6E-08 | 1,5E-07 | 3,9E-09 | 2,8E-08 | 3,7E-01 | 0,0E+00 | 1,3E-09 | 4,4E-09 | 2,3E-10 | 2,2E-09 | 0,0E+00 | 2911590 | 2912357 | 768 | + |
| Y11_RS13410 | Y11_28381 | | sulfate ABC transporter substrate-binding protein | -0,5 | 2,88 | 4,28 | 3,38 | 4,78 | 1,4 | 1,4E-03 | 1,6E-08 | 6,4E-11 | 3,5E-09 | 1,6E-11 | 2,1E-06 | 0,0E+00 | 8,9E-10 | 1,8E-11 | 2,2E-10 | 1,6E-11 | 3,3E-07 | 2913460 | 2914449 | 990 | - |
| Y11_RS13415 | Y11_28391 | pfkA | 6-phosphofructokinase | -0,17 | 1,82 | 0,71 | 1,99 | 0,88 | -1,11 | 1,4E-02 | 3,5E-10 | 4,9E-04 | 1,4E-09 | 1,7E-04 | 2,8E-05 | 0,0E+00 | 5,2E-11 | 0,0E+00 | 1,1E-10 | 0,0E+00 | 2,8E-06 | 2914660 | 2915643 | 984 | - |
| Y11_RS13470 | Y11_28521 | | type II toxin-antitoxin system HipA family toxin | 0,26 | 0,36 | 0,57 | 0,1 | 0,31 | 0,21 | 2,0E-03 | 3,5E-03 | 1,4E-04 | 2,5E-01 | 3,6E-03 | 7,3E-02 | 0,0E+00 | 0,0E+00 | 0,0E+00 | 0,0E+00 | 0,0E+00 | 0,0E+00 | 2924673 | 2926016 | 1344 | - |
| Y11_RS13475 | Y11_28531 | | helix-turn-helix domain-containing protein | 0,02 | 0,77 | 0,43 | 0,75 | 0,41 | -0,34 | 9,1E-01 | 2,1E-03 | 3,2E-01 | 1,1E-02 | 3,8E-01 | 4,4E-01 | 0,0E+00 | 0,0E+00 | 0,0E+00 | 0,0E+00 | 0,0E+00 | 0,0E+00 | 2926013 | 2926276 | 264 | - |
| Y11_RS13545 | Y11_28691 | | helix-turn-helix domain-containing protein | 0,05 | -0,1 | 0,19 | -0,15 | 0,14 | 0,29 | 8,3E-01 | 2,1E-01 | 5,4E-01 | 5,3E-01 | 7,1E-01 | 3,7E-01 | 0,0E+00 | 0,0E+00 | 0,0E+00 | 0,0E+00 | 0,0E+00 | 0,0E+00 | 2939389 | 2939664 | 276 | + |
| Y11_RS13555 | Y11_28711 | cpxP | cell-envelope stress modulator CpxP | -1,22 | -0,05 | 1,01 | 1,17 | 2,23 | 1,06 | 1,9E-06 | 5,7E-01 | 1,6E-04 | 1,1E-05 | 1,2E-06 | 2,1E-04 | 1,1E-07 | 0,0E+00 | 2,1E-06 | 2,2E-07 | 5,0E-08 | 1,6E-05 | 2940913 | 2941410 | 498 | - |
| Y11_RS13560 | Y11_28721 | cpxR | envelope stress response regulator transcription factor CpxR | -0,13 | -1,07 | -0,46 | -0,94 | -0,33 | 0,61 | 9,3E-02 | 3,6E-07 | 5,3E-03 | 4,0E-06 | 3,6E-02 | 1,7E-03 | 0,0E+00 | 1,2E-08 | 0,0E+00 | 0,0E+00 | 0,0E+00 | 0,0E+00 | 2941598 | 2942296 | 699 | + |
| Y11_RS13565 | Y11_28731 | cpxA | envelope stress sensor histidine kinase CpxA | -0,25 | -2,01 | -2,1 | -1,76 | -1,85 | -0,09 | 3,6E-02 | 2,0E-08 | 5,5E-08 | 9,9E-10 | 1,4E-08 | 2,3E-01 | 0,0E+00 | 1,1E-09 | 2,0E-09 | 8,8E-11 | 1,2E-09 | 0,0E+00 | 2942293 | 2943669 | 1377 | + |
| Y11_RS13570 | Y11_28741 | ada | bifunctional DNA-binding transcriptional regulator/O6-methylguanine-DNA methyltransferase Ada | 0,17 | 1,65 | 1,63 | 1,48 | 1,46 | -0,02 | 1,9E-01 | 1,7E-05 | 1,7E-04 | 7,4E-05 | 5,1E-04 | 9,4E-01 | 0,0E+00 | 4,0E-07 | 2,3E-06 | 1,4E-06 | 1,2E-05 | 0,0E+00 | 2943739 | 2944815 | 1077 | - |
| Y11_RS13575 | Y11_28751 | trmL | tRNA (uridine(34)/cytosine(34)/5-carboxymethylaminomethyluridine(34)-2'-O)-methyltransferase TrmL | 0,14 | -0,47 | -0,57 | -0,61 | -0,71 | -0,1 | 5,5E-01 | 2,8E-02 | 3,0E-02 | 4,7E-03 | 7,8E-03 | 5,4E-01 | 0,0E+00 | 0,0E+00 | 0,0E+00 | 0,0E+00 | 0,0E+00 | 0,0E+00 | 2945009 | 2945512 | 504 | + |
| Y11_RS13580 | Y11_28761 | cysE | serine O-acetyltransferase | -0,15 | 0,85 | 0,66 | 1 | 0,8 | -0,2 | 5,8E-02 | 6,0E-06 | 2,6E-03 | 6,8E-07 | 6,1E-04 | 2,4E-01 | 0,0E+00 | 0,0E+00 | 0,0E+00 | 0,0E+00 | 0,0E+00 | 0,0E+00 | 2945920 | 2946741 | 822 | - |
| Y11_RS13585 | Y11_28771 | gpsA | NAD(P)H-dependent glycerol-3-phosphate dehydrogenase | -0,02 | 0,62 | 0,29 | 0,64 | 0,31 | -0,33 | 7,2E-01 | 3,7E-05 | 2,7E-02 | 3,9E-05 | 2,2E-02 | 2,3E-02 | 0,0E+00 | 0,0E+00 | 0,0E+00 | 0,0E+00 | 0,0E+00 | 0,0E+00 | 2946867 | 2947886 | 1020 | - |
| Y11_RS13590 | Y11_28781 | secB | protein-export chaperone SecB | -0,03 | 0,2 | -0,34 | 0,24 | -0,31 | -0,55 | 6,2E-01 | 4,0E-03 | 1,4E-01 | 7,0E-04 | 1,7E-01 | 2,9E-02 | 0,0E+00 | 0,0E+00 | 0,0E+00 | 0,0E+00 | 0,0E+00 | 0,0E+00 | 2947886 | 2948356 | 471 | - |
| Y11_RS13595 | Y11_28791 | grxC | glutaredoxin 3 | 0,14 | 1,07 | 1,54 | 0,93 | 1,4 | 0,47 | 2,5E-01 | 3,0E-04 | 4,8E-04 | 3,7E-04 | 6,7E-04 | 1,5E-01 | 0,0E+00 | 6,3E-06 | 5,9E-06 | 0,0E+00 | 1,6E-05 | 0,0E+00 | 2948448 | 2948696 | 249 | - |
| Y11_RS13600 | Y11_28801 | | rhodanese-like domain-containing protein | -0,08 | 0,29 | 0,41 | 0,38 | 0,49 | 0,12 | 5,0E-01 | 1,0E-01 | 1,6E-02 | 1,9E-02 | 7,5E-04 | 4,5E-01 | 0,0E+00 | 0,0E+00 | 0,0E+00 | 0,0E+00 | 0,0E+00 | 0,0E+00 | 2948746 | 2949180 | 435 | - |
| Y11_RS13605 | Y11_28811 | gpmM | 2,3-bisphosphoglycerate-independent phosphoglycerate mutase | -0,56 | -0,74 | -1,29 | -0,18 | -0,73 | -0,55 | 1,3E-05 | 2,2E-07 | 1,2E-07 | 1,1E-02 | 1,7E-05 | 5,7E-05 | 0,0E+00 | 0,0E+00 | 3,7E-09 | 0,0E+00 | 0,0E+00 | 0,0E+00 | 2949422 | 2950969 | 1548 | + |
| Y11_RS13610 | Y11_28821 | envC | murein hydrolase activator EnvC | 0,03 | 0,99 | 1 | 0,95 | 0,96 | 0,01 | 7,0E-01 | 6,9E-06 | 7,6E-07 | 1,2E-05 | 1,5E-06 | 8,9E-01 | 0,0E+00 | 0,0E+00 | 0,0E+00 | 0,0E+00 | 0,0E+00 | 0,0E+00 | 2950979 | 2952346 | 1368 | + |
| Y11_RS13615 | Y11_28831 | | divergent polysaccharide deacetylase family protein | 0,19 | 0,39 | 0,04 | 0,2 | -0,15 | -0,35 | 1,7E-01 | 2,0E-02 | 8,1E-01 | 1,7E-01 | 4,0E-01 | 7,8E-02 | 0,0E+00 | 0,0E+00 | 0,0E+00 | 0,0E+00 | 0,0E+00 | 0,0E+00 | 2952370 | 2953374 | 1005 | + |
| Y11_RS13625 | Y11_28851 | | MASE1 domain-containing protein | 0 | -0,47 | 0 | -0,5 | 0 | 0,84 | 0,0E+00 | 2,0E-01 | 0,0E+00 | 1,7E-01 | 0,0E+00 | 5,2E-02 | 0,0E+00 | 0,0E+00 | 0,0E+00 | 0,0E+00 | 0,0E+00 | 0,0E+00 | 2955115 | 2956389 | 1275 | - |
| Y11_RS13630 | Y11_28871 | tdh | L-threonine 3-dehydrogenase | 0 | 0,56 | 1,36 | 0,55 | 1,36 | 0,81 | 9,7E-01 | 4,8E-03 | 2,0E-07 | 4,2E-03 | 9,7E-08 | 3,5E-04 | 0,0E+00 | 0,0E+00 | 5,8E-09 | 0,0E+00 | 6,1E-09 | 0,0E+00 | 2956714 | 2957739 | 1026 | - |
| Y11_RS13635 | Y11_28881 | kbl | glycine C-acetyltransferase | -0,18 | 0,24 | 1,75 | 0,42 | 1,93 | 1,5 | 3,5E-02 | 1,9E-02 | 7,0E-10 | 1,0E-03 | 3,8E-10 | 2,0E-08 | 0,0E+00 | 0,0E+00 | 8,1E-11 | 0,0E+00 | 9,8E-11 | 1,2E-08 | 2957764 | 2958975 | 1212 | - |
| Y11_RS13640 | Y11_28891 | rfaD | ADP-glyceromanno-heptose 6-epimerase | -0,07 | 0,31 | 0,25 | 0,38 | 0,32 | -0,06 | 1,9E-01 | 1,9E-03 | 7,3E-02 | 5,3E-04 | 2,9E-02 | 6,4E-01 | 0,0E+00 | 0,0E+00 | 0,0E+00 | 0,0E+00 | 0,0E+00 | 0,0E+00 | 2959219 | 2960151 | 933 | + |
| Y11_RS13645 | Y11_28901 | rfaF | ADP-heptose--LPS heptosyltransferase RfaF | -0,24 | 0,13 | 0,21 | 0,37 | 0,45 | 0,08 | 9,9E-03 | 1,4E-01 | 6,7E-02 | 2,2E-04 | 7,1E-04 | 3,9E-01 | 0,0E+00 | 0,0E+00 | 0,0E+00 | 0,0E+00 | 0,0E+00 | 0,0E+00 | 2960182 | 2961246 | 1065 | + |
| Y11_RS13650 | Y11_28911 | rfaC | lipopolysaccharide heptosyltransferase RfaC | -0,12 | 0,48 | 0,37 | 0,6 | 0,49 | -0,11 | 2,3E-01 | 1,7E-03 | 2,6E-02 | 1,7E-04 | 4,6E-03 | 4,3E-01 | 0,0E+00 | 0,0E+00 | 0,0E+00 | 0,0E+00 | 0,0E+00 | 0,0E+00 | 2961246 | 2962211 | 966 | + |
| Y11_RS13655 | Y11_28921 | waaA | lipid IV(A) 3-deoxy-D-manno-octulosonic acid transferase | 0,12 | 0,47 | 1,15 | 0,35 | 1,03 | 0,68 | 2,4E-01 | 1,3E-03 | 9,7E-06 | 1,9E-03 | 7,3E-06 | 1,7E-04 | 0,0E+00 | 0,0E+00 | 1,7E-07 | 0,0E+00 | 2,5E-07 | 0,0E+00 | 2962759 | 2964033 | 1275 | + |
| Y11_RS13660 | Y11_28931 | | glycosyltransferase family 2 protein | -0,07 | 0,3 | 1,01 | 0,36 | 1,07 | 0,71 | 2,5E-01 | 6,7E-03 | 1,4E-04 | 7,5E-04 | 6,5E-05 | 1,8E-03 | 0,0E+00 | 0,0E+00 | 1,9E-06 | 0,0E+00 | 1,8E-06 | 0,0E+00 | 2964034 | 2964816 | 783 | + |
| Y11_RS13665 | Y11_28941 | coaD | pantetheine-phosphate adenylyltransferase | 0,32 | 0,86 | 1,58 | 0,54 | 1,26 | 0,72 | 2,1E-03 | 4,1E-05 | 1,5E-06 | 2,3E-03 | 1,8E-05 | 1,9E-03 | 0,0E+00 | 0,0E+00 | 3,2E-08 | 0,0E+00 | 5,4E-07 | 0,0E+00 | 2964813 | 2965292 | 480 | + |
| Y11_RS13670 | Y11_28951 | mutM | bifunctional DNA-formamidopyrimidine glycosylase/DNA-(apurinic or apyrimidinic site) lyase | -0,25 | -0,37 | -0,58 | -0,13 | -0,33 | -0,2 | 4,2E-02 | 1,1E-02 | 2,4E-02 | 3,3E-01 | 1,6E-01 | 3,8E-01 | 0,0E+00 | 0,0E+00 | 0,0E+00 | 0,0E+00 | 0,0E+00 | 0,0E+00 | 2965590 | 2966399 | 810 | - |
| Y11_RS13675 | Y11_28961 | rpmG | 50S ribosomal protein L33 | 0,22 | -0,66 | -1,18 | -0,88 | -1,4 | -0,52 | 2,1E-01 | 1,1E-03 | 5,6E-02 | 5,8E-05 | 2,9E-02 | 3,4E-01 | 0,0E+00 | 0,0E+00 | 6,3E-04 | 0,0E+00 | 5,9E-04 | 0,0E+00 | 2966486 | 2966653 | 168 | - |
| Y11_RS13680 | Y11_28971 | rpmB | 50S ribosomal protein L28 | 0,42 | 0,07 | -0,16 | -0,35 | -0,58 | -0,23 | 7,0E-03 | 5,9E-01 | 9,6E-02 | 6,7E-02 | 2,5E-03 | 1,5E-01 | 0,0E+00 | 0,0E+00 | 0,0E+00 | 0,0E+00 | 0,0E+00 | 0,0E+00 | 2966665 | 2966901 | 237 | - |
| Y11_RS13690 | Y11_28991 | coaBC | bifunctional phosphopantothenoylcysteine decarboxylase/phosphopantothenate--cysteine ligase CoaBC | -0,73 | -0,52 | -0,28 | 0,2 | 0,44 | 0,24 | 2,7E-06 | 2,2E-08 | 1,4E-02 | 1,0E-02 | 3,1E-03 | 2,7E-02 | 0,0E+00 | 0,0E+00 | 0,0E+00 | 0,0E+00 | 0,0E+00 | 0,0E+00 | 2968025 | 2969257 | 1233 | + |
| Y11_RS13695 | Y11_29001 | dut | dUTP diphosphatase | -0,62 | -0,2 | -0,58 | 0,42 | 0,04 | -0,38 | 1,8E-03 | 3,6E-01 | 6,8E-04 | 1,2E-01 | 8,0E-01 | 1,3E-01 | 0,0E+00 | 0,0E+00 | 0,0E+00 | 0,0E+00 | 0,0E+00 | 0,0E+00 | 2969235 | 2969693 | 459 | + |

| | | | | | | | | | | | | | | | | | | | | | | | | |
|---|---|---|---|---|---|---|---|---|---|---|---|---|---|---|---|---|---|---|---|---|---|---|---|---|
| Y11_RS13700 | Y11_29011 | slmA | nucleoid occlusion factor SlmA | -0,48 | 0,39 | 0,33 | 0,87 | 0,81 | -0,06 | 6,7E-05 | 1,2E-04 | 1,5E-03 | 1,4E-06 | 7,6E-06 | 4,1E-01 | 0,0E+00 | 0,0E+00 | 0,0E+00 | 0,0E+00 | 0,0E+00 | 0,0E+00 | 2969815 | 2970411 | 597 | + |
| Y11_RS13710 | Y11_29051 | pyrE | orotate phosphoribosyltransferase | 0,24 | 1,72 | 2,97 | 1,48 | 2,72 | 1,25 | 2,6E-03 | 3,1E-09 | 1,3E-10 | 2,6E-08 | 5,1E-10 | 3,6E-07 | 0,0E+00 | 2,5E-10 | 2,6E-11 | 1,1E-09 | 1,2E-10 | 8,3E-08 | 2972107 | 2972748 | 642 | - |
| Y11_RS13715 | Y11_29061 | rph | ribonuclease PH | -0,29 | -1,06 | -0,72 | -0,77 | -0,44 | 0,34 | 1,2E-05 | 1,2E-07 | 2,7E-05 | 1,2E-06 | 8,2E-04 | 9,2E-03 | 0,0E+00 | 4,6E-09 | 0,0E+00 | 0,0E+00 | 0,0E+00 | 0,0E+00 | 2972902 | 2973618 | 717 | - |
| Y11_RS13720 | Y11_29071 | | YicC family protein | -0,17 | 0 | -0,22 | 0,18 | -0,04 | -0,22 | 3,0E-03 | 9,3E-01 | 6,3E-03 | 3,3E-03 | 5,2E-01 | 6,4E-03 | 0,0E+00 | 0,0E+00 | 0,0E+00 | 0,0E+00 | 0,0E+00 | 0,0E+00 | 2973745 | 2974608 | 864 | + |
| Y11_RS13745 | Y11_29121 | | DUF3574 domain-containing protein | 0,69 | 0,98 | 1,14 | 0,29 | 0,45 | 0,16 | 1,6E-03 | 1,3E-05 | 2,4E-05 | 5,8E-02 | 1,8E-02 | 1,8E-01 | 0,0E+00 | 0,0E+00 | 3,7E-07 | 0,0E+00 | 0,0E+00 | 0,0E+00 | 2980815 | 2981279 | 465 | - |
| Y11_RS13760 | Y11_29151 | gmk | guanylate kinase | -0,07 | -0,07 | -0,32 | 0 | -0,25 | -0,25 | 2,2E-01 | 4,2E-01 | 3,3E-02 | 9,8E-01 | 7,1E-02 | 1,0E-01 | 0,0E+00 | 0,0E+00 | 0,0E+00 | 0,0E+00 | 0,0E+00 | 0,0E+00 | 2984201 | 2984824 | 624 | + |
| Y11_RS13765 | Y11_29161 | rpoZ | DNA-directed RNA polymerase subunit omega | 0,02 | 0,1 | -0,03 | 0,08 | -0,04 | -0,12 | 6,9E-01 | 1,7E-01 | 8,2E-01 | 1,9E-01 | 7,1E-01 | 3,6E-01 | 0,0E+00 | 0,0E+00 | 0,0E+00 | 0,0E+00 | 0,0E+00 | 0,0E+00 | 2984879 | 2985154 | 276 | + |
| Y11_RS13770 | Y11_29171 | spoT | bifunctional GTP diphosphokinase/guanosine-3',5'-bis pyrophosphate 3'-pyrophosphohydrolase | -0,23 | 0,11 | -0,11 | 0,34 | 0,13 | -0,22 | 4,0E-05 | 1,4E-01 | 3,9E-02 | 9,2E-04 | 1,7E-02 | 2,0E-02 | 0,0E+00 | 0,0E+00 | 0,0E+00 | 0,0E+00 | 0,0E+00 | 0,0E+00 | 2985173 | 2987275 | 2103 | + |
| Y11_RS13775 | Y11_29181 | trmH | tRNA (guanosine(18)-2'-O)-methyltransferase TrmH | -0,1 | 0,11 | -0,36 | 0,21 | -0,25 | -0,47 | 6,2E-01 | 4,8E-01 | 8,3E-02 | 4,1E-01 | 3,7E-01 | 7,8E-02 | 0,0E+00 | 0,0E+00 | 0,0E+00 | 0,0E+00 | 0,0E+00 | 0,0E+00 | 2987281 | 2987973 | 693 | + |
| Y11_RS13780 | Y11_29191 | recG | ATP-dependent DNA helicase RecG | 0,31 | -0,49 | -0,03 | -0,8 | -0,34 | 0,46 | 8,7E-02 | 1,6E-03 | 8,0E-01 | 4,6E-04 | 6,1E-02 | 2,0E-03 | 0,0E+00 | 0,0E+00 | 0,0E+00 | 0,0E+00 | 0,0E+00 | 0,0E+00 | 2987974 | 2990055 | 2082 | + |
| Y11_RS13790 | Y11_29221 | gltS | sodium/glutamate symporter | 1,61 | -0,5 | -0,69 | -2,1 | -2,3 | -0,2 | 4,2E-05 | 2,8E-02 | 2,8E-02 | 1,9E-08 | 3,4E-06 | 3,2E-01 | 2,0E-06 | 0,0E+00 | 0,0E+00 | 8,4E-10 | 1,2E-07 | 0,0E+00 | 2991673 | 2992887 | 1215 | - |
| Y11_RS13800 | Y11_29241 | | hypothetical protein | 0,59 | -0,22 | -0,04 | -0,8 | -0,63 | 0,17 | 6,3E-05 | 2,1E-02 | 7,8E-01 | 2,6E-06 | 3,1E-03 | 2,9E-01 | 0,0E+00 | 0,0E+00 | 0,0E+00 | 0,0E+00 | 0,0E+00 | 0,0E+00 | 2994657 | 2996366 | 1710 | + |
| Y11_RS13825 | Y11_29291 | fabY | fatty acid biosynthesis protein FabY | -0,21 | -0,33 | -0,76 | -0,12 | -0,55 | -0,43 | 8,1E-02 | 2,8E-03 | 7,7E-04 | 3,4E-01 | 1,1E-02 | 2,1E-02 | 0,0E+00 | 0,0E+00 | 0,0E+00 | 0,0E+00 | 0,0E+00 | 0,0E+00 | 2999183 | 3000106 | 924 | - |
| Y11_RS13830 | Y11_29301 | dtd | D-aminoacyl-tRNA deacylase | -0,03 | 0,49 | 0,52 | 0,51 | 0,55 | 0,04 | 9,0E-01 | 5,4E-03 | 1,1E-02 | 4,8E-02 | 5,0E-02 | 8,4E-01 | 0,0E+00 | 0,0E+00 | 0,0E+00 | 0,0E+00 | 0,0E+00 | 0,0E+00 | 3000363 | 3000800 | 438 | - |
| Y11_RS13840 | Y11_29321 | yihX | glucose-1-phosphatase | -0,03 | 0,51 | 1,3 | 0,54 | 1,32 | 0,79 | 6,9E-01 | 6,3E-11 | 7,9E-08 | 3,2E-05 | 6,7E-07 | 3,8E-06 | 0,0E+00 | 0,0E+00 | 2,7E-09 | 0,0E+00 | 3,0E-08 | 0,0E+00 | 3001790 | 3002377 | 588 | - |
| Y11_RS13845 | Y11_29331 | typA | ribosome-dependent GTPase TypA | -0,28 | -0,16 | -0,84 | 0,12 | -0,56 | -0,68 | 6,0E-03 | 6,2E-02 | 8,6E-07 | 2,0E-01 | 9,8E-05 | 1,8E-05 | 0,0E+00 | 0,0E+00 | 0,0E+00 | 0,0E+00 | 0,0E+00 | 0,0E+00 | 3002672 | 3004495 | 1824 | - |
| Y11_RS13850 | Y11_29341 | glnA | glutamate--ammonia ligase | 0,03 | -0,34 | -0,46 | -0,37 | -0,5 | -0,13 | 4,0E-01 | 5,9E-03 | 7,4E-04 | 5,4E-03 | 8,0E-04 | 3,4E-01 | 0,0E+00 | 0,0E+00 | 0,0E+00 | 0,0E+00 | 0,0E+00 | 0,0E+00 | 3004997 | 3006406 | 1410 | + |
| Y11_RS13855 | Y11_29361 | glnL | nitrogen regulation protein NR(II) | 0,36 | -0,26 | -0,86 | -0,63 | -1,22 | -0,6 | 4,4E-03 | 7,7E-02 | 1,1E-05 | 7,5E-04 | 1,7E-07 | 8,7E-04 | 0,0E+00 | 0,0E+00 | 0,0E+00 | 0,0E+00 | 9,5E-09 | 0,0E+00 | 3006774 | 3007823 | 1050 | + |
| Y11_RS13860 | Y11_29371 | glnG | nitrogen regulation protein NR(I) | 0,14 | 0,03 | -0,63 | -0,11 | -0,76 | -0,66 | 1,7E-01 | 6,4E-01 | 2,3E-04 | 2,8E-01 | 1,9E-04 | 1,6E-04 | 0,0E+00 | 0,0E+00 | 0,0E+00 | 0,0E+00 | 0,0E+00 | 0,0E+00 | 3007831 | 3009243 | 1413 | + |
| Y11_RS13865 | Y11_29381 | hemN | oxygen-independent coproporphyrinogen III oxidase | 0,36 | 0,73 | 0,55 | 0,36 | 0,19 | -0,18 | 1,1E-04 | 2,3E-06 | 3,9E-04 | 6,5E-04 | 9,5E-02 | 1,2E-01 | 0,0E+00 | 0,0E+00 | 0,0E+00 | 0,0E+00 | 0,0E+00 | 0,0E+00 | 3009392 | 3010765 | 1374 | - |
| Y11_RS13870 | Y11_29391 | yihI | Der GTPase-activating protein YihI | 0,47 | 0,66 | 0,7 | 0,19 | 0,23 | 0,04 | 2,9E-03 | 3,3E-04 | 2,9E-02 | 1,8E-01 | 4,2E-01 | 8,9E-01 | 0,0E+00 | 0,0E+00 | 0,0E+00 | 0,0E+00 | 0,0E+00 | 0,0E+00 | 3010948 | 3011514 | 567 | - |
| Y11_RS13875 | Y11_29401 | yihA | ribosome biogenesis GTP-binding protein YihA/YsxC | -0,52 | 0,38 | 0,37 | 0,91 | 0,9 | -0,01 | 5,1E-07 | 7,7E-06 | 1,7E-03 | 3,3E-08 | 5,2E-06 | 9,1E-01 | 0,0E+00 | 0,0E+00 | 0,0E+00 | 0,0E+00 | 0,0E+00 | 0,0E+00 | 3012277 | 3012930 | 654 | + |
| Y11_RS13880 | Y11_29411 | polA | DNA polymerase I | -0,03 | -0,28 | -0,17 | -0,25 | -0,13 | 0,12 | 3,8E-01 | 2,9E-04 | 4,4E-03 | 9,1E-04 | 1,9E-02 | 5,8E-02 | 0,0E+00 | 0,0E+00 | 0,0E+00 | 0,0E+00 | 0,0E+00 | 0,0E+00 | 3013364 | 3016162 | 2799 | - |
| Y11_RS13885 | Y11_29421 | dsbA | thiol:disulfide interchange protein DsbA | -0,4 | -1,11 | -0,54 | -0,71 | -0,13 | 0,57 | 1,5E-04 | 7,9E-07 | 1,8E-04 | 4,3E-06 | 7,7E-02 | 1,5E-04 | 0,0E+00 | 2,4E-08 | 0,0E+00 | 0,0E+00 | 0,0E+00 | 0,0E+00 | 3016753 | 3017376 | 624 | - |
| Y11_RS13890 | Y11_29431 | | serine/threonine protein kinase | -0,25 | -0,99 | -0,53 | -0,74 | -0,28 | 0,46 | 3,0E-03 | 7,0E-06 | 2,8E-04 | 1,1E-04 | 1,9E-02 | 5,7E-03 | 0,0E+00 | 0,0E+00 | 0,0E+00 | 0,0E+00 | 0,0E+00 | 0,0E+00 | 3017404 | 3018390 | 987 | - |
| Y11_RS13895 | Y11_29441 | | YihD family protein | -0,23 | -1,11 | -1,04 | -0,88 | -0,81 | 0,07 | 2,6E-03 | 6,9E-07 | 4,1E-06 | 3,5E-06 | 2,3E-05 | 5,6E-01 | 0,0E+00 | 2,1E-08 | 7,6E-08 | 0,0E+00 | 0,0E+00 | 0,0E+00 | 3018482 | 3018751 | 270 | - |
| Y11_RS13900 | Y11_29451 | mobA | molybdenum cofactor guanylyltransferase MobA | -0,27 | -0,65 | 0,5 | -0,38 | 0,77 | 1,15 | 3,6E-02 | 1,0E-03 | 1,9E-03 | 1,3E-02 | 4,8E-05 | 1,2E-05 | 0,0E+00 | 0,0E+00 | 0,0E+00 | 0,0E+00 | 0,0E+00 | 1,4E-06 | 3018905 | 3019495 | 591 | + |
| Y11_RS13905 | Y11_29461 | mobB | molybdopterin-guanine dinucleotide biosynthesis protein MobB | 0 | -1,42 | -0,51 | -1,42 | -0,51 | 0,91 | 9,5E-01 | 1,5E-06 | 5,0E-04 | 1,7E-06 | 5,9E-04 | 1,2E-04 | 0,0E+00 | 4,2E-08 | 0,0E+00 | 4,0E-08 | 0,0E+00 | 0,0E+00 | 3019492 | 3020025 | 534 | + |
| Y11_RS13940 | Y11_29491 | | FadR family transcriptional regulator | 0,08 | 0,74 | 0,86 | 0,66 | 0,78 | 0,12 | 5,6E-01 | 1,1E-07 | 3,6E-04 | 1,3E-03 | 4,0E-03 | 4,5E-01 | 0,0E+00 | 0,0E+00 | 0,0E+00 | 0,0E+00 | 0,0E+00 | 0,0E+00 | 3027522 | 3028211 | 690 | + |
| Y11_RS13950 | Y11_29521 | rbsR | ribose operon transcriptional repressor RbsR | -0,22 | -0,02 | -0,24 | 0,2 | -0,02 | -0,23 | 3,3E-01 | 9,1E-01 | 2,4E-01 | 2,5E-01 | 9,1E-01 | 1,4E-01 | 0,0E+00 | 0,0E+00 | 0,0E+00 | 0,0E+00 | 0,0E+00 | 0,0E+00 | 3029709 | 3030710 | 1002 | - |
| Y11_RS13955 | Y11_29531 | rbsK | ribokinase | 0,35 | -0,19 | 0,38 | -0,53 | 0,03 | 0,56 | 2,9E-03 | 1,2E-01 | 5,0E-03 | 8,6E-04 | 7,7E-01 | 1,3E-03 | 0,0E+00 | 0,0E+00 | 0,0E+00 | 0,0E+00 | 0,0E+00 | 0,0E+00 | 3030713 | 3031639 | 927 | - |
| Y11_RS13960 | Y11_29541 | rbsB | ribose ABC transporter substrate-binding protein RbsB | 0,41 | 1,3 | 1,95 | 0,89 | 1,55 | 0,65 | 8,0E-04 | 5,3E-08 | 1,7E-07 | 1,2E-07 | 6,1E-07 | 2,1E-04 | 0,0E+00 | 2,4E-09 | 5,2E-09 | 0,0E+00 | 2,7E-08 | 0,0E+00 | 3031730 | 3032620 | 891 | - |
| Y11_RS13970 | Y11_29561 | rbsA | ribose ABC transporter ATP-binding protein RbsA | 0,38 | 1,31 | 1,61 | 0,93 | 1,23 | 0,3 | 2,5E-02 | 2,4E-05 | 2,5E-07 | 8,9E-04 | 2,5E-05 | 8,7E-02 | 0,0E+00 | 5,5E-07 | 6,9E-09 | 0,0E+00 | 7,4E-07 | 0,0E+00 | 3033739 | 3035241 | 1503 | - |
| Y11_RS13975 | Y11_29571 | rbsD | D-ribose pyranase | 0,14 | 0,6 | 1,46 | 0,46 | 1,32 | 0,87 | 4,9E-01 | 6,7E-02 | 2,4E-02 | 1,4E-01 | 3,6E-02 | 1,6E-01 | 0,0E+00 | 0,0E+00 | 2,8E-04 | 0,0E+00 | 7,5E-04 | 0,0E+00 | 3035249 | 3035668 | 420 | - |
| Y11_RS13980 | Y11_29581 | kup | low affinity potassium transporter Kup | -0,11 | -1,13 | -0,82 | -1,02 | -0,72 | 0,31 | 3,9E-01 | 8,1E-09 | 1,8E-05 | 3,0E-05 | 1,1E-03 | 1,1E-02 | 0,0E+00 | 5,2E-10 | 0,0E+00 | 5,9E-07 | 0,0E+00 | 0,0E+00 | 3036005 | 3037876 | 1872 | - |
| Y11_RS13985 | Y11_29591 | ravA | ATPase RavA | 0,54 | 0,48 | 1,18 | -0,06 | 0,64 | 0,7 | 1,0E-04 | 1,1E-03 | 2,7E-04 | 4,7E-01 | 8,2E-03 | 6,7E-03 | 0,0E+00 | 0,0E+00 | 3,5E-06 | 0,0E+00 | 0,0E+00 | 0,0E+00 | 3038164 | 3039672 | 1509 | + |
| Y11_RS13995 | Y11_29611 | | DUF1345 domain-containing protein | 1,1 | 0,43 | 1,7 | -0,67 | 0,55 | 1,27 | 1,2E-03 | 1,9E-01 | 1,4E-04 | 1,3E-02 | 1,9E-03 | 6,4E-04 | 4,6E-05 | 0,0E+00 | 1,9E-06 | 0,0E+00 | 0,0E+00 | 4,5E-05 | 3041152 | 3041823 | 672 | - |
| Y11_RS14000 | Y11_29621 | asnA | aspartate--ammonia ligase | -1,21 | -0,8 | -2,27 | 0,42 | -1,06 | -1,47 | 9,4E-06 | 7,9E-05 | 1,1E-07 | 4,3E-03 | 3,0E-05 | 1,0E-06 | 4,8E-07 | 0,0E+00 | 3,5E-09 | 0,0E+00 | 8,8E-07 | 1,9E-07 | 3042056 | 3043048 | 993 | - |
| Y11_RS14005 | Y11_29631 | asnC | transcriptional regulator AsnC | 0,11 | -0,04 | 0,22 | -0,16 | 0,11 | 0,26 | 4,7E-01 | 6,9E-01 | 7,3E-02 | 2,8E-01 | 4,6E-01 | 1,6E-02 | 0,0E+00 | 0,0E+00 | 0,0E+00 | 0,0E+00 | 0,0E+00 | 0,0E+00 | 3043233 | 3043694 | 462 | + |
| Y11_RS14010 | Y11_29641 | mioC | FMN-binding protein MioC | 0,19 | 1,03 | 1,51 | 0,84 | 1,31 | 0,47 | 1,2E-01 | 5,4E-05 | 7,3E-06 | 6,8E-04 | 5,5E-05 | 3,4E-02 | 0,0E+00 | 1,2E-06 | 1,3E-07 | 0,0E+00 | 1,5E-06 | 0,0E+00 | 3043786 | 3044226 | 441 | + |
| Y11_RS14015 | Y11_29661 | mnmG | tRNA uridine-5-carboxymethylaminomethyl(34) synthesis enzyme MnmG | 0,68 | -1,17 | -1,09 | -1,47 | -1,39 | 0,08 | 7,8E-03 | 6,7E-08 | 2,7E-07 | 1,6E-10 | 2,1E-09 | 1,6E-01 | 0,0E+00 | 2,9E-09 | 7,6E-09 | 2,4E-11 | 2,9E-10 | 0,0E+00 | 3044607 | 3046496 | 1890 | + |
| Y11_RS14020 | Y11_29671 | rsmG | 16S rRNA (guanine(527)-N(7))-methyltransferase RsmG | 0,28 | 0,16 | 0,43 | -0,12 | 0,15 | 0,27 | 3,2E-04 | 6,8E-03 | 7,5E-04 | 4,8E-02 | 1,2E-01 | 1,3E-02 | 0,0E+00 | 0,0E+00 | 0,0E+00 | 0,0E+00 | 0,0E+00 | 0,0E+00 | 3046645 | 3047265 | 621 | + |
| Y11_RS14030 | Y11_29691 | atpB | F0F1 ATP synthase subunit A | 0,38 | -0,12 | 0,8 | -0,5 | 0,42 | 0,92 | 6,1E-03 | 3,1E-01 | 7,8E-05 | 2,3E-03 | 6,2E-03 | 5,4E-05 | 0,0E+00 | 0,0E+00 | 0,0E+00 | 0,0E+00 | 0,0E+00 | 0,0E+00 | 3048371 | 3049195 | 825 | + |
| Y11_RS14035 | Y11_29701 | atpE | F0F1 ATP synthase subunit C | 0,37 | -0,43 | 0,72 | -0,8 | 0,35 | 1,14 | 5,8E-02 | 8,8E-03 | 2,3E-01 | 1,4E-04 | 5,4E-01 | 6,5E-02 | 0,0E+00 | 0,0E+00 | 0,0E+00 | 0,0E+00 | 0,0E+00 | 3,4E-03 | 3049245 | 3049484 | 240 | + |
| Y11_RS14040 | Y11_29711 | atpF | F0F1 ATP synthase subunit B | 0,36 | -0,5 | 0,08 | -0,85 | -0,27 | 0,58 | 4,0E-03 | 3,5E-04 | 5,8E-01 | 5,6E-06 | 8,4E-02 | 2,9E-03 | 0,0E+00 | 0,0E+00 | 0,0E+00 | 0,0E+00 | 0,0E+00 | 0,0E+00 | 3049545 | 3050015 | 471 | + |
| Y11_RS14045 | Y11_29721 | atpH | F0F1 ATP synthase subunit delta | 0,29 | -0,71 | -0,3 | -1 | -0,58 | 0,42 | 5,2E-04 | 6,0E-10 | 9,3E-05 | 3,7E-08 | 1,1E-05 | 4,3E-06 | 0,0E+00 | 0,0E+00 | 0,0E+00 | 1,4E-09 | 0,0E+00 | 0,0E+00 | 3050030 | 3050563 | 534 | + |
| Y11_RS14050 | Y11_29731 | atpA | F0F1 ATP synthase subunit alpha | 0,27 | -0,73 | -0,12 | -1 | -0,38 | 0,62 | 1,9E-03 | 4,0E-06 | 3,2E-02 | 1,4E-06 | 2,7E-04 | 2,0E-05 | 0,0E+00 | 0,0E+00 | 0,0E+00 | 3,4E-08 | 0,0E+00 | 0,0E+00 | 3050578 | 3052119 | 1542 | + |
| Y11_RS14055 | Y11_29741 | atpG | F0F1 ATP synthase subunit gamma | 0,3 | -0,72 | 0 | -1,02 | -0,3 | 0,72 | 2,3E-05 | 1,2E-08 | 9,8E-01 | 2,2E-09 | 3,5E-04 | 3,8E-07 | 0,0E+00 | 0,0E+00 | 0,0E+00 | 1,6E-10 | 0,0E+00 | 0,0E+00 | 3052181 | 3053044 | 864 | + |
| Y11_RS14060 | Y11_29751 | atpD | F0F1 ATP synthase subunit beta | 0,23 | -0,75 | -0,06 | -0,98 | -0,29 | 0,69 | 2,2E-02 | 2,5E-04 | 3,8E-01 | 6,6E-05 | 8,1E-03 | 4,8E-04 | 0,0E+00 | 0,0E+00 | 0,0E+00 | 0,0E+00 | 0,0E+00 | 0,0E+00 | 3053201 | 3054583 | 1383 | + |
| Y11_RS14065 | Y11_29761 | atpC | F0F1 ATP synthase subunit epsilon | 0,2 | -0,28 | 0,35 | -0,48 | 0,14 | 0,63 | 1,9E-02 | 8,2E-03 | 4,0E-02 | 7,7E-05 | 3,1E-01 | 2,0E-03 | 0,0E+00 | 0,0E+00 | 0,0E+00 | 0,0E+00 | 0,0E+00 | 0,0E+00 | 3054605 | 3055027 | 423 | + |
| Y11_RS14070 | Y11_29771 | glmU | bifunctional UDP-N-acetylglucosamine diphosphorylase/glucosamine-1-phosphate N-acetyltransferase GlmU | 0,94 | -0,76 | -0,64 | -1,7 | -1,57 | 0,13 | 3,0E-08 | 8,8E-07 | 2,1E-06 | 1,7E-09 | 1,9E-09 | 7,9E-02 | 0,0E+00 | 0,0E+00 | 0,0E+00 | 1,3E-10 | 2,8E-10 | 0,0E+00 | 3055541 | 3056911 | 1371 | + |
| Y11_RS14075 | Y11_29781 | glmS | glutamine--fructose-6-phosphate transaminase (isomerizing) | 0,83 | -1,03 | -0,86 | -1,86 | -1,69 | 0,17 | 3,7E-09 | 1,4E-06 | 7,2E-06 | 1,7E-08 | 4,6E-08 | 1,7E-01 | 0,0E+00 | 4,1E-08 | 0,0E+00 | 7,8E-10 | 3,3E-09 | 0,0E+00 | 3057259 | 3059088 | 1830 | + |
| Y11_RS14080 | Y11_29791 | pstS | phosphate ABC transporter substrate-binding protein PstS | 0,61 | 0,44 | 0,3 | -0,17 | -0,31 | -0,14 | 1,5E-06 | 9,0E-08 | 2,0E-02 | 7,6E-03 | 2,2E-02 | 1,9E-01 | 0,0E+00 | 0,0E+00 | 0,0E+00 | 0,0E+00 | 0,0E+00 | 0,0E+00 | 3059726 | 3060766 | 1041 | + |
| Y11_RS14090 | Y11_29811 | pstA | phosphate ABC transporter permease PstA | -0,65 | -0,41 | -0,76 | 0,18 | -0,14 | -0,33 | 5,0E-02 | 3,6E-02 | 1,5E-03 | 6,0E-01 | 6,8E-01 | 1,8E-01 | 0,0E+00 | 0,0E+00 | 0,0E+00 | 0,0E+00 | 0,0E+00 | 0,0E+00 | 3061907 | 3062794 | 888 | + |
| Y11_RS14095 | Y11_29821 | pstB | phosphate ABC transporter ATP-binding protein PstB | -0,97 | -0,31 | -0,76 | 0,66 | 0,2 | -0,45 | 9,4E-09 | 3,8E-05 | 1,6E-07 | 1,1E-08 | 5,8E-04 | 1,2E-06 | 0,0E+00 | 0,0E+00 | 0,0E+00 | 0,0E+00 | 0,0E+00 | 0,0E+00 | 3062895 | 3063671 | 777 | + |
| Y11_RS14100 | Y11_29831 | phoU | phosphate signaling complex protein PhoU | -0,63 | 0,05 | -0,31 | 0,68 | 0,31 | -0,36 | 2,3E-04 | 6,1E-01 | 7,8E-03 | 7,9E-05 | 7,1E-03 | 2,0E-03 | 0,0E+00 | 0,0E+00 | 0,0E+00 | 0,0E+00 | 0,0E+00 | 0,0E+00 | 3063736 | 3064458 | 723 | + |
| Y11_RS14110 | Y11_29861 | | ABC transporter substrate-binding protein | 15 | -7,54 | -5,58 | -15 | -15 | 1,96 | 0,0E+00 | 1,2E-11 | 1,5E-10 | 0,0E+00 | 0,0E+00 | 1,3E-06 | 0,0E+00 | 6,3E-12 | 2,8E-11 | 0,0E+00 | 0,0E+00 | 2,1E-07 | 3065688 | 3066542 | 855 | + |
| Y11_RS14115 | Y11_29871 | | amino acid ABC transporter permease | -0,16 | -1,23 | 2,86 | -1,07 | 3,02 | 4,08 | 6,8E-01 | 5,0E-04 | 1,8E-06 | 1,7E-02 | 3,5E-05 | 7,6E-08 | 0,0E+00 | 1,0E-05 | 3,7E-08 | 2,7E-04 | 1,0E-06 | 2,9E-08 | 3066771 | 3067508 | 738 | + |
| Y11_RS14125 | Y11_29891 | yieH | 6-phosphogluconate phosphatase | 0,78 | 1,17 | 1,41 | 0,39 | 0,63 | 0,24 | 1,8E-05 | 4,1E-07 | 4,2E-05 | 2,8E-03 | 8,3E-03 | 2,1E-01 | 0,0E+00 | 1,3E-08 | 6,2E-07 | 0,0E+00 | 0,0E+00 | 0,0E+00 | 3068262 | 3068930 | 669 | - |
| Y11_RS14130 | Y11_29901 | | NCS2 family permease | 0,9 | 1,01 | 0,49 | 0,11 | -0,41 | -0,52 | 2,5E-02 | 6,4E-03 | 1,6E-01 | 6,6E-01 | 2,0E-01 | 5,7E-02 | 0,0E+00 | 1,2E-04 | 0,0E+00 | 0,0E+00 | 0,0E+00 | 0,0E+00 | 3069200 | 3070528 | 1329 | + |
| Y11_RS14135 | Y11_29911 | | heme-binding protein | 0,63 | 0,58 | 0,85 | -0,06 | 0,22 | 0,28 | 4,1E-03 | 2,1E-03 | 1,7E-03 | 7,5E-01 | 3,3E-01 | 1,8E-01 | 0,0E+00 | 0,0E+00 | 0,0E+00 | 0,0E+00 | 0,0E+00 | 0,0E+00 | 3070614 | 3071105 | 492 | - |
| Y11_RS14140 | Y11_29921 | | response regulator | -0,1 | 0,22 | 0,34 | 0,33 | 0,45 | 0,12 | 7,1E-01 | 5,7E-01 | 3,3E-01 | 3,9E-01 | 1,9E-01 | 7,8E-01 | 0,0E+00 | 0,0E+00 | 0,0E+00 | 0,0E+00 | 0,0E+00 | 0,0E+00 | 3071345 | 3071962 | 618 | - |
| Y11_RS14145 | Y11_29931 | | ATP-binding protein | 0,08 | -0,12 | -0,19 | -0,2 | -0,27 | -0,07 | 6,4E-01 | 5,5E-01 | 2,0E-01 | 3,1E-01 | 5,7E-02 | 6,9E-01 | 0,0E+00 | 0,0E+00 | 0,0E+00 | 0,0E+00 | 0,0E+00 | 0,0E+00 | 3071937 | 3073286 | 1350 | - |
| Y11_RS14150 | Y11_29941 | | NAD(P)H-dependent oxidoreductase | 0,44 | 1,54 | 1,45 | 1,1 | 1,01 | -0,1 | 3,3E-04 | 5,8E-10 | 3,4E-07 | 3,1E-07 | 1,6E-05 | 3,4E-01 | 0,0E+00 | 7,3E-11 | 9,2E-09 | 8,8E-09 | 5,1E-07 | 0,0E+00 | 3073394 | 3073966 | 573 | - |
| Y11_RS14175 | Y11_29991 | fabV | enoyl-ACP reductase FabV | 0,34 | 0,45 | 0,14 | 0,1 | -0,2 | -0,31 | 9,0E-05 | 4,8E-06 | 3,5E-01 | 3,4E-02 | 1,9E-01 | 5,9E-02 | 0,0E+00 | 0,0E+00 | 0,0E+00 | 0,0E+00 | 0,0E+00 | 0,0E+00 | 3078998 | 3080197 | 1200 | - |
| Y11_RS14180 | Y11_30001 | mnmE | tRNA uridine-5-carboxymethylaminomethyl(34) synthesis GTPase MnmE | -0,1 | -1,17 | -1,39 | -1,07 | -1,29 | -0,22 | 1,2E-01 | 1,1E-08 | 2,1E-08 | 1,0E-08 | 2,5E-08 | 4,0E-03 | 0,0E+00 | 6,5E-10 | 9,3E-10 | 5,3E-10 | 2,0E-09 | 0,0E+00 | 3080306 | 3081670 | 1365 | - |
| Y11_RS14185 | Y11_30011 | yidC | membrane protein insertase YidC | 0,18 | -0,38 | -0,44 | -0,57 | -0,63 | -0,06 | 3,0E-03 | 3,2E-04 | 7,6E-04 | 3,4E-05 | 9,9E-05 | 5,5E-01 | 0,0E+00 | 0,0E+00 | 0,0E+00 | 0,0E+00 | 0,0E+00 | 0,0E+00 | 3081838 | 3083484 | 1647 | - |

| | | | | | | | | | | | | | | | | | | | | | | | | | |
|---|---|---|---|---|---|---|---|---|---|---|---|---|---|---|---|---|---|---|---|---|---|---|---|---|---|
| Y11_RS14190 | | rnpA | ribonuclease P protein component | 0,25 | -0,56 | -0,73 | -0,81 | -0,97 | -0,17 | 3,1E-01 | 2,0E-02 | 1,7E-03 | 1,1E-02 | 2,0E-03 | 3,7E-01 | 0,0E+00 | 0,0E+00 | 0,0E+00 | 0,0E+00 | 0,0E+00 | 0,0E+00 | 3083708 | 3084067 | 360 | - |
| Y11_RS14195 | Y11_30051 | rpmH | 50S ribosomal protein L34 | 0,55 | -1,23 | -0,98 | -1,78 | -1,54 | 0,25 | 2,1E-02 | 8,6E-06 | 1,1E-02 | 3,9E-05 | 2,3E-03 | 4,6E-01 | 0,0E+00 | 2,1E-07 | 0,0E+00 | 7,5E-07 | 5,1E-05 | 0,0E+00 | 3084086 | 3084226 | 141 | - |
| Y11_RS14200 | Y11_30071 | dnaA | chromosomal replication initiator protein DnaA | -0,22 | -0,33 | -0,48 | -0,11 | -0,25 | -0,15 | 2,0E-03 | 2,4E-04 | 7,6E-05 | 4,3E-02 | 2,3E-03 | 3,8E-02 | 0,0E+00 | 0,0E+00 | 0,0E+00 | 0,0E+00 | 0,0E+00 | 0,0E+00 | 3084931 | 3086319 | 1389 | + |
| Y11_RS14205 | Y11_30081 | dnaN | DNA polymerase III subunit beta | -0,07 | 0,1 | -0,06 | 0,17 | 0,01 | -0,16 | 3,2E-01 | 2,5E-01 | 4,9E-01 | 6,0E-02 | 9,2E-01 | 1,3E-01 | 0,0E+00 | 0,0E+00 | 0,0E+00 | 0,0E+00 | 0,0E+00 | 0,0E+00 | 3086324 | 3087424 | 1101 | + |
| Y11_RS14215 | Y11_30101 | gyrB | DNA topoisomerase (ATP-hydrolyzing) subunit B | 0,03 | -0,51 | -0,58 | -0,54 | -0,61 | -0,07 | 1,7E-01 | 1,0E-06 | 6,7E-05 | 6,4E-07 | 4,5E-05 | 4,0E-01 | 0,0E+00 | 0,0E+00 | 0,0E+00 | 0,0E+00 | 0,0E+00 | 0,0E+00 | 3088702 | 3091116 | 2415 | + |
| Y11_RS14220 | Y11_30111 | yidA | sugar-phosphatase | 0 | 1,13 | 0,27 | 1,13 | 0,27 | -0,86 | 1,0E+00 | 3,7E-05 | 7,4E-02 | 6,3E-06 | 2,7E-02 | 9,2E-05 | 0,0E+00 | 8,3E-07 | 0,0E+00 | 1,3E-07 | 0,0E+00 | 0,0E+00 | 3091287 | 3092096 | 810 | + |
| Y11_RS14225 | Y11_30121 | | ornithine cyclodeaminase family protein | 0,33 | 1,15 | 1,64 | 0,82 | 1,32 | 0,49 | 3,9E-02 | 3,6E-08 | 1,5E-08 | 2,2E-04 | 1,2E-05 | 8,1E-05 | 0,0E+00 | 1,7E-09 | 7,1E-10 | 0,0E+00 | 3,8E-07 | 0,0E+00 | 3092278 | 3093216 | 939 | - |
| Y11_RS14245 | Y11_30171 | ibpA | heat shock chaperone IbpA | -0,69 | 0,76 | 0,8 | 1,45 | 1,49 | 0,04 | 1,8E-05 | 6,9E-06 | 9,9E-05 | 1,3E-09 | 3,1E-07 | 6,9E-01 | 0,0E+00 | 0,0E+00 | 0,0E+00 | 1,1E-10 | 1,6E-08 | 0,0E+00 | 3096340 | 3096753 | 414 | + |
| Y11_RS14250 | Y11_30181 | ibpB | heat shock chaperone IbpB | -0,66 | 0,99 | 1,01 | 1,65 | 1,67 | 0,02 | 7,7E-04 | 5,6E-07 | 3,8E-06 | 6,6E-07 | 1,4E-06 | 8,3E-01 | 0,0E+00 | 0,0E+00 | 7,1E-08 | 1,7E-08 | 5,6E-08 | 0,0E+00 | 3096878 | 3097351 | 474 | + |
| Y11_RS14270 | Y11_30211 | | putative transporter | -2,19 | 0,65 | 15 | 2,81 | 15 | 0 | 6,3E-05 | 5,4E-02 | 0,0E+00 | 1,1E-07 | 0,0E+00 | 0,0E+00 | 2,9E-06 | 0,0E+00 | 0,0E+00 | 3,7E-09 | 0,0E+00 | 0,0E+00 | 3099412 | 3101073 | 1662 | + |
| Y11_RS14275 | Y11_30221 | | valine--pyruvate transaminase | 0,65 | -0,06 | -0,26 | -0,7 | -0,91 | -0,2 | 2,8E-05 | 5,0E-01 | 1,2E-01 | 1,0E-05 | 2,2E-04 | 2,0E-01 | 0,0E+00 | 0,0E+00 | 0,0E+00 | 0,0E+00 | 0,0E+00 | 0,0E+00 | 3101154 | 3102407 | 1254 | - |
| Y11_RS14285 | Y11_30241 | ghrB | glyoxylate/hydroxypyruvate reductase GhrB | -1,61 | 1,52 | 0,96 | 3,13 | 2,57 | -0,56 | 8,2E-09 | 9,9E-10 | 5,1E-06 | 5,4E-11 | 6,5E-09 | 2,9E-04 | 1,2E-09 | 1,1E-10 | 0,0E+00 | 1,2E-11 | 6,9E-10 | 0,0E+00 | 3104759 | 3105739 | 981 | - |
| Y11_RS14290 | Y11_30251 | | MFS transporter | -2,97 | -1,16 | -1,54 | 1,81 | 1,43 | -0,38 | 3,3E-04 | 2,2E-02 | 2,0E-03 | 8,1E-03 | 1,5E-02 | 3,3E-01 | 1,3E-05 | 4,2E-04 | 2,3E-05 | 1,3E-04 | 3,2E-04 | 0,0E+00 | 3106043 | 3107485 | 1443 | - |
| Y11_RS14295 | Y11_30261 | | transcriptional regulator | 1,46 | 1,38 | 1,23 | -0,12 | -0,26 | -0,18 | 1,4E-02 | 2,1E-02 | 3,8E-02 | 6,1E-01 | 3,1E-01 | 5,6E-01 | 4,7E-04 | 4,0E-04 | 4,2E-04 | 0,0E+00 | 0,0E+00 | 0,0E+00 | 3107671 | 3108315 | 645 | + |
| Y11_RS14320 | Y11_30311 | | DNA-3-methyladenine glycosylase I | 0,16 | -0,57 | 0,05 | -0,73 | -0,11 | 0,62 | 3,4E-01 | 4,6E-03 | 7,8E-01 | 1,6E-04 | 5,0E-01 | 2,4E-03 | 0,0E+00 | 0,0E+00 | 0,0E+00 | 0,0E+00 | 0,0E+00 | 0,0E+00 | 3115607 | 3116179 | 573 | - |
| Y11_RS14325 | Y11_30321 | glyQ | glycine--tRNA ligase subunit alpha | -0,23 | -0,92 | -0,87 | -0,69 | -0,63 | 0,05 | 2,7E-02 | 1,8E-07 | 3,0E-03 | 8,3E-05 | 2,0E-02 | 8,0E-01 | 0,0E+00 | 0,0E+00 | 0,0E+00 | 0,0E+00 | 0,0E+00 | 0,0E+00 | 3116435 | 3117349 | 915 | + |
| Y11_RS14330 | Y11_30331 | glyS | glycine--tRNA ligase subunit beta | -0,16 | -0,47 | -0,46 | -0,31 | -0,3 | 0,01 | 2,1E-04 | 7,9E-07 | 6,7E-04 | 5,3E-05 | 9,0E-03 | 8,9E-01 | 0,0E+00 | 0,0E+00 | 0,0E+00 | 0,0E+00 | 0,0E+00 | 0,0E+00 | 3117359 | 3119428 | 2070 | + |
| Y11_RS14335 | Y11_30341 | | DUF3053 domain-containing protein | -0,13 | -0,4 | -0,36 | -0,28 | -0,23 | 0,05 | 1,0E-01 | 5,6E-04 | 1,1E-04 | 1,6E-02 | 1,5E-02 | 5,8E-01 | 0,0E+00 | 0,0E+00 | 0,0E+00 | 0,0E+00 | 0,0E+00 | 0,0E+00 | 3119551 | 3120270 | 720 | + |
| Y11_RS14340 | Y11_30351 | | PTS mannitol transporter subunit IICBA | 1,48 | 3,34 | 3,36 | 1,86 | 1,88 | 0,02 | 5,2E-08 | 4,1E-11 | 3,7E-08 | 1,3E-08 | 4,1E-06 | 9,0E-01 | 5,2E-09 | 1,3E-11 | 1,5E-09 | 6,4E-10 | 1,5E-07 | 0,0E+00 | 3121453 | 3123375 | 1923 | + |
| Y11_RS14345 | Y11_30361 | | mannitol-1-phosphate 5-dehydrogenase | 0,77 | 2,52 | 2,55 | 1,74 | 1,78 | 0,04 | 1,2E-06 | 3,9E-12 | 1,3E-08 | 2,2E-09 | 4,1E-07 | 7,4E-01 | 0,0E+00 | 3,9E-12 | 6,5E-10 | 1,6E-10 | 2,0E-08 | 0,0E+00 | 3123640 | 3124803 | 1164 | + |
| Y11_RS14350 | Y11_30371 | | MltR family transcriptional regulator | 0,11 | -0,53 | -0,35 | -0,65 | -0,46 | 0,19 | 3,8E-01 | 1,1E-02 | 9,8E-02 | 7,1E-03 | 5,2E-02 | 4,3E-01 | 0,0E+00 | 0,0E+00 | 0,0E+00 | 0,0E+00 | 0,0E+00 | 0,0E+00 | 3125035 | 3125589 | 555 | + |
| Y11_RS14365 | Y11_30401 | | YibL family ribosome-associated protein | -0,11 | 0,85 | 0,66 | 0,96 | 0,77 | -0,19 | 4,8E-01 | 3,7E-04 | 6,6E-04 | 2,3E-04 | 3,8E-04 | 1,6E-01 | 0,0E+00 | 0,0E+00 | 0,0E+00 | 0,0E+00 | 0,0E+00 | 0,0E+00 | 3128039 | 3128395 | 357 | + |
| Y11_RS14370 | Y11_30411 | vapB | type II toxin-antitoxin system VapB family antitoxin | 0,29 | 0,91 | 1,31 | 0,63 | 1,03 | 0,4 | 3,6E-01 | 3,6E-02 | 1,8E-03 | 1,1E-01 | 5,1E-03 | 2,8E-01 | 0,0E+00 | 0,0E+00 | 2,1E-05 | 0,0E+00 | 1,1E-04 | 0,0E+00 | 3128527 | 3128754 | 228 | + |
| Y11_RS14375 | Y11_30421 | | type II toxin-antitoxin system VapC family toxin | 0,7 | 1,08 | 1,49 | 0,38 | 0,77 | 0,41 | 1,1E-01 | 2,0E-03 | 2,5E-04 | 2,9E-01 | 5,0E-02 | 5,3E-03 | 0,0E+00 | 4,0E-05 | 3,3E-06 | 0,0E+00 | 0,0E+00 | 0,0E+00 | 3128751 | 3129170 | 420 | + |
| Y11_RS14380 | Y11_30431 | yiiM | 6-N-hydroxylaminopurine resistance protein | 1,34 | 2,36 | 2,59 | 1,02 | 1,25 | 0,23 | 5,0E-03 | 9,5E-06 | 3,5E-05 | 2,2E-02 | 1,6E-02 | 5,0E-01 | 1,8E-04 | 2,3E-07 | 5,2E-07 | 3,5E-04 | 3,4E-04 | 0,0E+00 | 3129324 | 3129995 | 672 | - |
| Y11_RS14385 | Y11_30441 | | methyl-accepting chemotaxis protein | 0,11 | 0,54 | 0,97 | 0,42 | 0,86 | 0,44 | 6,5E-01 | 1,1E-01 | 1,2E-02 | 7,0E-02 | 3,6E-03 | 1,5E-01 | 0,0E+00 | 0,0E+00 | 0,0E+00 | 0,0E+00 | 0,0E+00 | 0,0E+00 | 3130074 | 3132023 | 1950 | - |
| Y11_RS14390 | Y11_30461 | sodA | superoxide dismutase [Mn] | 0,33 | 1,09 | 1,13 | 0,75 | 0,79 | 0,04 | 5,3E-03 | 8,2E-07 | 1,4E-06 | 3,8E-05 | 4,7E-05 | 6,8E-01 | 0,0E+00 | 2,4E-08 | 3,1E-08 | 0,0E+00 | 0,0E+00 | 0,0E+00 | 3132347 | 3132970 | 624 | - |
| Y11_RS14395 | | fdhD | formate dehydrogenase accessory sulfurtransferase FdhD | 0,11 | 0,24 | 0,56 | 0,12 | 0,45 | 0,32 | 4,4E-01 | 1,2E-01 | 9,9E-02 | 4,8E-01 | 2,0E-01 | 3,4E-01 | 0,0E+00 | 0,0E+00 | 0,0E+00 | 0,0E+00 | 0,0E+00 | 0,0E+00 | 3133097 | 3133951 | 855 | - |
| Y11_RS14405 | Y11_30471 | fdnG | formate dehydrogenase-N subunit alpha | 0,36 | 1,89 | 2,33 | 1,53 | 1,97 | 0,44 | 5,0E-05 | 2,1E-09 | 1,5E-09 | 2,7E-08 | 1,1E-08 | 1,8E-03 | 0,0E+00 | 1,8E-10 | 1,3E-10 | 1,1E-09 | 1,0E-09 | 0,0E+00 | 3134135 | 3137182 | 3048 | + |
| Y11_RS14410 | Y11_30481 | fdxH | formate dehydrogenase subunit beta | 0,68 | 1,65 | 1,94 | 0,96 | 1,26 | 0,3 | 2,6E-03 | 8,1E-07 | 3,7E-07 | 2,3E-04 | 4,8E-05 | 4,2E-02 | 0,0E+00 | 2,4E-08 | 1,0E-08 | 0,0E+00 | 1,3E-06 | 0,0E+00 | 3137195 | 3138166 | 972 | + |
| Y11_RS14415 | Y11_30491 | fdoI | formate dehydrogenase cytochrome b556 subunit | 0,45 | 1,95 | 3,03 | 1,5 | 2,58 | 1,08 | 2,2E-01 | 1,3E-05 | 2,2E-05 | 1,4E-03 | 2,7E-04 | 9,7E-03 | 0,0E+00 | 3,0E-07 | 3,4E-07 | 2,3E-05 | 6,6E-06 | 5,6E-04 | 3138163 | 3138813 | 651 | + |
| Y11_RS14420 | Y11_30501 | fdhE | formate dehydrogenase accessory protein FdhE | 0,02 | 0,51 | 0,52 | 0,49 | 0,5 | 0,01 | 8,0E-01 | 2,8E-05 | 5,3E-03 | 2,3E-05 | 6,1E-03 | 9,3E-01 | 0,0E+00 | 0,0E+00 | 0,0E+00 | 0,0E+00 | 0,0E+00 | 0,0E+00 | 3138813 | 3139742 | 930 | + |
| Y11_RS14425 | Y11_30511 | selA | L-seryl-tRNA(Sec) selenium transferase | -0,17 | 0,25 | 0,01 | 0,41 | 0,17 | -0,24 | 2,7E-01 | 4,1E-02 | 9,7E-01 | 5,9E-03 | 2,6E-01 | 5,0E-02 | 0,0E+00 | 0,0E+00 | 0,0E+00 | 0,0E+00 | 0,0E+00 | 0,0E+00 | 3139774 | 3141162 | 1389 | + |
| Y11_RS14430 | Y11_30521 | selB | selenocysteine-specific translation elongation factor | -0,31 | -0,76 | -0,32 | -0,45 | 0 | 0,44 | 2,9E-02 | 9,4E-05 | 1,8E-02 | 1,4E-02 | 9,7E-01 | 1,1E-02 | 0,0E+00 | 0,0E+00 | 0,0E+00 | 0,0E+00 | 0,0E+00 | 0,0E+00 | 3141159 | 3143051 | 1893 | + |
| Y11_RS14435 | Y11_30531 | | MFS transporter | -0,02 | -0,69 | -1,41 | -0,69 | -1,41 | -0,71 | 9,2E-01 | 5,3E-03 | 7,0E-04 | 1,1E-02 | 1,1E-03 | 1,7E-02 | 0,0E+00 | 0,0E+00 | 8,5E-06 | 0,0E+00 | 2,4E-05 | 0,0E+00 | 3143117 | 3144346 | 1230 | - |
| Y11_RS14460 | Y11_30581 | xylB | xylulokinase | 0,26 | 0,21 | 0,43 | -0,05 | 0,17 | 0,22 | 1,8E-02 | 2,3E-02 | 6,8E-02 | 6,3E-01 | 4,7E-01 | 3,3E-01 | 0,0E+00 | 0,0E+00 | 0,0E+00 | 0,0E+00 | 0,0E+00 | 0,0E+00 | 3148788 | 3150248 | 1461 | - |
| Y11_RS14470 | Y11_30611 | xylF | D-xylose ABC transporter substrate-binding protein | 0,47 | 1,96 | 3,29 | 1,5 | 2,83 | 1,33 | 5,9E-04 | 6,6E-07 | 1,6E-11 | 5,9E-06 | 9,2E-11 | 6,9E-06 | 0,0E+00 | 2,0E-08 | 8,0E-12 | 1,2E-07 | 4,4E-11 | 8,3E-07 | 3152142 | 3153137 | 996 | + |
| Y11_RS14520 | Y11_30731 | | amino acid permease | 0,36 | -1,46 | 0,56 | -1,83 | 0,19 | 2,02 | 1,3E-01 | 4,3E-04 | 3,9E-02 | 5,6E-05 | 3,7E-01 | 3,4E-05 | 0,0E+00 | 8,9E-06 | 0,0E+00 | 1,1E-06 | 0,0E+00 | 3,2E-06 | 3165507 | 3166904 | 1398 | - |
| Y11_RS14525 | Y11_30741 | hutH | histidine ammonia-lyase | 0,55 | -1,27 | 0,48 | -1,82 | -0,07 | 1,75 | 4,9E-07 | 3,8E-09 | 2,1E-04 | 4,4E-10 | 3,8E-01 | 2,6E-08 | 0,0E+00 | 2,9E-10 | 0,0E+00 | 4,7E-11 | 0,0E+00 | 1,4E-08 | 3167014 | 3168549 | 1536 | - |
| Y11_RS14530 | Y11_30751 | hutU | urocanate hydratase | 0,41 | -0,71 | 0,89 | -1,13 | 0,48 | 1,6 | 1,0E-03 | 1,9E-05 | 4,2E-06 | 1,2E-08 | 1,4E-05 | 5,0E-10 | 0,0E+00 | 0,0E+00 | 0,0E+00 | 6,2E-10 | 0,0E+00 | 1,1E-09 | 3168546 | 3170237 | 1692 | - |
| Y11_RS14550 | Y11_30811 | | glycoside hydrolase family 1 protein | 0 | -15 | 0 | -15 | 0 | 15 | 0,0E+00 | 0,0E+00 | 0,0E+00 | 0,0E+00 | 0,0E+00 | 0,0E+00 | 0,0E+00 | 0,0E+00 | 0,0E+00 | 0,0E+00 | 0,0E+00 | 0,0E+00 | 3176117 | 3177517 | 1401 | - |
| Y11_RS14570 | Y11_30861 | | PTS sugar transporter subunit IIB | 0 | -15 | -15 | -15 | -15 | 1,68 | 0,0E+00 | 0,0E+00 | 0,0E+00 | 0,0E+00 | 0,0E+00 | 2,7E-02 | 0,0E+00 | 0,0E+00 | 0,0E+00 | 0,0E+00 | 0,0E+00 | 1,4E-03 | 3182822 | 3183136 | 315 | + |
| Y11_RS14590 | Y11_30901 | | LacI family transcriptional regulator | -0,43 | 1,29 | 1,04 | 1,67 | 1,48 | -0,21 | 8,8E-02 | 1,0E-04 | 2,6E-03 | 1,8E-05 | 3,0E-04 | 3,5E-01 | 0,0E+00 | 2,2E-06 | 3,0E-05 | 3,7E-07 | 7,2E-06 | 0,0E+00 | 3185889 | 3186851 | 963 | + |
| Y11_RS14615 | Y11_30971 | | alpha/beta hydrolase | 0,33 | 0,59 | 1,02 | 0,26 | 0,69 | 0,43 | 1,0E-05 | 3,2E-08 | 1,4E-06 | 2,6E-05 | 2,8E-05 | 6,2E-04 | 0,0E+00 | 0,0E+00 | 3,1E-08 | 0,0E+00 | 0,0E+00 | 0,0E+00 | 3191707 | 3193170 | 1464 | + |
| Y11_RS14655 | Y11_31041 | uhpB | signal transduction histidine-protein kinase/phosphatase UhpB | -1,16 | 1,12 | 4 | 2,28 | 5,09 | 2,85 | 6,6E-05 | 3,9E-03 | 2,7E-04 | 1,5E-05 | 5,1E-05 | 2,9E-03 | 3,0E-06 | 7,7E-05 | 3,4E-06 | 3,0E-07 | 1,4E-06 | 1,8E-04 | 3199905 | 3201437 | 1533 | - |
| Y11_RS14660 | Y11_31051 | uhpA | transcriptional regulator UhpA | -1,3 | 0,25 | 1,28 | 1,55 | 2,57 | 1,02 | 9,6E-05 | 6,5E-02 | 5,2E-05 | 3,7E-05 | 2,6E-06 | 3,5E-04 | 4,3E-06 | 0,0E+00 | 7,4E-07 | 7,0E-07 | 9,7E-08 | 2,6E-05 | 3201434 | 3202024 | 591 | - |
| Y11_RS14670 | Y11_31071 | | organic hydroperoxide resistance protein | 0,79 | 3,54 | 4,44 | 2,75 | 3,65 | 0,9 | 1,1E-04 | 1,8E-09 | 9,0E-09 | 8,5E-08 | 1,0E-07 | 2,6E-03 | 0,0E+00 | 1,7E-10 | 4,9E-10 | 2,9E-09 | 6,4E-09 | 0,0E+00 | 3204087 | 3204512 | 426 | - |
| Y11_RS14680 | Y11_31101 | | ABC transporter substrate-binding protein | -0,56 | -1,29 | -1,09 | -0,73 | -0,53 | 0,2 | 4,7E-07 | 3,4E-08 | 1,2E-08 | 2,7E-06 | 3,2E-06 | 1,8E-02 | 0,0E+00 | 1,6E-09 | 6,0E-10 | 0,0E+00 | 0,0E+00 | 0,0E+00 | 3206812 | 3208419 | 1608 | + |
| Y11_RS14695 | Y11_31131 | dppD | dipeptide ABC transporter ATP-binding protein | -0,42 | -0,43 | -0,51 | -0,01 | -0,09 | -0,07 | 8,5E-06 | 2,3E-04 | 1,2E-03 | 8,3E-01 | 4,2E-01 | 5,5E-01 | 0,0E+00 | 0,0E+00 | 0,0E+00 | 0,0E+00 | 0,0E+00 | 0,0E+00 | 3210650 | 3211630 | 981 | + |
| Y11_RS14700 | Y11_31141 | | ABC transporter ATP-binding protein | -0,56 | -0,01 | -0,69 | 0,55 | -0,13 | -0,68 | 3,9E-05 | 9,0E-01 | 3,8E-04 | 3,1E-04 | 2,6E-01 | 8,9E-04 | 0,0E+00 | 0,0E+00 | 0,0E+00 | 0,0E+00 | 0,0E+00 | 0,0E+00 | 3211645 | 3212649 | 1005 | + |
| Y11_RS14705 | Y11_31151 | bcsG | cellulose biosynthesis protein BcsG | 0,53 | 1,14 | 1,73 | 0,61 | 1,2 | 0,59 | 6,0E-03 | 4,2E-05 | 2,1E-04 | 2,4E-03 | 2,2E-03 | 6,0E-02 | 0,0E+00 | 9,4E-07 | 2,8E-06 | 0,0E+00 | 4,8E-05 | 0,0E+00 | 3212992 | 3214644 | 1653 | - |
| Y11_RS14715 | Y11_31171 | bcsE | cellulose biosynthesis protein BcsE | -0,86 | 0 | 0 | 15 | 0,07 | 0 | 3,3E-03 | 0,0E+00 | 0,0E+00 | 0,0E+00 | 6,9E-01 | 0,0E+00 | 0,0E+00 | 0,0E+00 | 0,0E+00 | 0,0E+00 | 0,0E+00 | 0,0E+00 | 3214832 | 3216391 | 1560 | - |
| Y11_RS14735 | Y11_31211 | bcsB | cellulose biosynthesis cyclic di-GMP-binding regulatory protein BcsB | 0,75 | 0,74 | 0,4 | -0,01 | -0,35 | -0,34 | 1,6E-06 | 7,3E-06 | 4,2E-04 | 8,7E-01 | 6,8E-04 | 2,3E-03 | 0,0E+00 | 0,0E+00 | 0,0E+00 | 0,0E+00 | 0,0E+00 | 0,0E+00 | 3220156 | 3222459 | 2304 | + |
| Y11_RS14755 | Y11_31251 | | EAL domain-containing protein | 0,01 | 0,66 | 0,32 | 0,65 | 0,31 | -0,34 | 9,3E-01 | 1,0E-06 | 6,9E-03 | 4,0E-04 | 5,0E-02 | 2,4E-03 | 0,0E+00 | 0,0E+00 | 0,0E+00 | 0,0E+00 | 0,0E+00 | 0,0E+00 | 3227311 | 3228840 | 1530 | + |
| Y11_RS14765 | Y11_31271 | | pectate lyase | 0 | -4,06 | -3,34 | -3,77 | -3,1 | 0,7 | 0,0E+00 | 2,6E-11 | 3,3E-09 | 9,0E-10 | 9,0E-09 | 3,3E-04 | 0,0E+00 | 3,6E-11 | 1,2E-10 | 8,1E-11 | 9,0E-10 | 0,0E+00 | 3230221 | 3231939 | 1719 | + |
| Y11_RS14770 | Y11_31281 | | discoidin domain-containing protein | 0,45 | -0,83 | 1,17 | -1,28 | 0,73 | 2,01 | 3,6E-04 | 1,8E-05 | 4,9E-02 | 2,0E-07 | 1,9E-01 | 4,2E-03 | 0,0E+00 | 0,0E+00 | 5,5E-04 | 6,0E-09 | 0,0E+00 | 2,6E-04 | 3232064 | 3232543 | 480 | + |
| Y11_RS14775 | Y11_31291 | | dicarboxylate/amino acid:cation symporter | -0,01 | -1,32 | 1,56 | -1,31 | 1,57 | 2,88 | 9,7E-01 | 3,1E-04 | 4,8E-04 | 1,1E-03 | 9,9E-04 | 1,1E-06 | 0,0E+00 | 6,5E-06 | 5,9E-06 | 1,8E-05 | 2,3E-05 | 1,9E-07 | 3232921 | 3234213 | 1293 | + |
| Y11_RS14780 | Y11_31301 | | insulinase family protein | -1,02 | 0,39 | 1,14 | 1,41 | 2,16 | 0,75 | 1,3E-08 | 2,6E-04 | 2,1E-09 | 1,4E-08 | 6,2E-12 | 1,3E-06 | 1,6E-09 | 0,0E+00 | 1,7E-10 | 6,7E-10 | 8,8E-12 | 0,0E+00 | 3234628 | 3236118 | 1491 | + |
| Y11_RS14785 | Y11_31311 | | sugar kinase | 0 | 0,07 | 0,85 | 0,07 | 0,85 | 0,78 | 9,8E-01 | 3,9E-01 | 3,6E-05 | 4,4E-01 | 5,2E-05 | 2,7E-05 | 0,0E+00 | 0,0E+00 | 0,0E+00 | 0,0E+00 | 0,0E+00 | 0,0E+00 | 3236278 | 3237222 | 945 | - |
| Y11_RS14790 | Y11_31321 | | 4-oxalocrotonate tautomerase family protein | -0,68 | 1,36 | 1,6 | 2,05 | 2,29 | 0,24 | 1,9E-06 | 1,8E-07 | 4,6E-05 | 3,7E-09 | 3,0E-06 | 2,8E-01 | 0,0E+00 | 6,8E-09 | 6,6E-07 | 2,3E-10 | 1,1E-07 | 0,0E+00 | 3237367 | 3237567 | 201 | - |
| Y11_RS14800 | Y11_31351 | | AsmA family protein | 0,75 | 0,42 | 0,09 | -0,33 | -0,66 | -0,33 | 6,2E-05 | 5,8E-03 | 5,4E-01 | 1,6E-02 | 1,4E-03 | 5,7E-02 | 0,0E+00 | 0,0E+00 | 0,0E+00 | 0,0E+00 | 0,0E+00 | 0,0E+00 | 3238696 | 3240837 | 2142 | + |
| Y11_RS14805 | Y11_31361 | | CDP-diacylglycerol diphosphatase | 0,01 | 0,03 | 0,86 | 0,02 | 0,86 | 0,83 | 9,7E-01 | 6,7E-01 | 3,0E-05 | 8,7E-01 | 8,3E-04 | 2,0E-05 | 0,0E+00 | 0,0E+00 | 0,0E+00 | 0,0E+00 | 0,0E+00 | 0,0E+00 | 3240842 | 3241606 | 765 | - |
| Y11_RS14810 | Y11_31371 | yhjD | inner membrane protein YhjD | -1,09 | 1,29 | 1,8 | 2,38 | 2,88 | 0,5 | 1,8E-05 | 3,4E-06 | 3,9E-06 | 4,3E-09 | 4,3E-08 | 7,2E-03 | 8,9E-07 | 8,9E-08 | 7,2E-08 | 2,6E-10 | 3,1E-09 | 0,0E+00 | 3241630 | 3242619 | 990 | - |
| Y11_RS14820 | Y11_31391 | gorA | glutathione-disulfide reductase | -0,43 | 0,45 | 0,17 | 0,88 | 0,6 | -0,28 | 2,0E-07 | 1,4E-07 | 1,1E-02 | 2,5E-09 | 3,3E-09 | 7,3E-04 | 0,0E+00 | 0,0E+00 | 0,0E+00 | 0,0E+00 | 0,0E+00 | 0,0E+00 | 3246637 | 3247989 | 1353 | - |
| Y11_RS14825 | Y11_31401 | | 23S rRNA (adenine(2030)-N(6))-methyltransferase RlmJ | -0,43 | 1,02 | 1,19 | 1,45 | 1,63 | 0,17 | 6,6E-05 | 2,7E-08 | 6,2E-06 | 1,8E-09 | 6,0E-07 | 1,6E-01 | 0,0E+00 | 1,4E-09 | 1,1E-07 | 1,4E-10 | 2,7E-08 | 0,0E+00 | 3248233 | 3249075 | 843 | - |
| Y11_RS14830 | Y11_31411 | prlC | oligopeptidase A | 0,12 | 0,71 | 0,68 | 0,59 | 0,56 | -0,03 | 6,4E-04 | 8,9E-10 | 6,1E-08 | 2,7E-08 | 6,3E-07 | 5,2E-01 | 0,0E+00 | 0,0E+00 | 0,0E+00 | 0,0E+00 | 0,0E+00 | 0,0E+00 | 3249412 | 3251454 | 2043 | + |
| Y11_RS14835 | Y11_31421 | rsmJ | 16S rRNA (guanine(1516)-N(2))-methyltransferase RsmJ | -0,68 | -0,75 | -2,01 | -0,06 | -1,32 | -1,26 | 6,1E-02 | 2,5E-02 | 1,8E-05 | 8,5E-01 | 1,8E-03 | 7,9E-04 | 0,0E+00 | 0,0E+00 | 2,9E-07 | 0,0E+00 | 4,0E-05 | 5,4E-05 | 3251458 | 3252216 | 759 | + |
| Y11_RS14845 | Y11_31441 | gdhA | NADP-specific glutamate dehydrogenase | 0,41 | -0,55 | -0,95 | -0,96 | -1,36 | -0,4 | 2,7E-05 | 4,0E-07 | 2,2E-09 | 3,8E-08 | 1,6E-09 | 2,0E-06 | 0,0E+00 | 0,0E+00 | 0,0E+00 | 0,0E+00 | 2,4E-10 | 0,0E+00 | 3253871 | 3255214 | 1344 | - |
| Y11_RS14850 | Y11_31451 | uspA | universal stress protein UspA | -0,76 | 2,54 | 3,05 | 3,3 | 3,81 | 0,51 | 6,0E-06 | 2,5E-10 | 9,4E-09 | 1,3E-10 | 2,5E-09 | 4,7E-03 | 0,0E+00 | 4,1E-11 | 5,0E-10 | 2,1E-11 | 3,4E-10 | 0,0E+00 | 3255478 | 3255924 | 447 | - |

| | | | | | | | | | | | | | | | | | | | | | | | | | |
|---|---|---|---|---|---|---|---|---|---|---|---|---|---|---|---|---|---|---|---|---|---|---|---|---|---|
| Y11_RS14860 | Y11_31471 | pitA | inorganic phosphate transporter PitA | -0,31 | 15 | 0,61 | 15 | 1,02 | 0 | 2,9E-01 | 0,0E+00 | 1,9E-02 | 0,0E+00 | 7,0E-03 | 0,0E+00 | 0,0E+00 | 0,0E+00 | 0,0E+00 | 0,0E+00 | 1,5E-04 | 0,0E+00 | 3257243 | 3258739 | 1497 | - |
| Y11_RS14865 | Y11_31481 | | NAD(P)/FAD-dependent oxidoreductase | 0,41 | -0,14 | -0,36 | -0,55 | -0,77 | -0,22 | 4,0E-03 | 3,8E-01 | 2,6E-03 | 1,0E-02 | 1,3E-04 | 1,8E-01 | 0,0E+00 | 0,0E+00 | 0,0E+00 | 0,0E+00 | 0,0E+00 | 0,0E+00 | 3259051 | 3260250 | 1200 | + |
| Y11_RS14885 | Y11_31521 | | methylaspartate ammonia-lyase | 0,31 | 0,1 | 0,63 | -0,21 | 0,33 | 0,54 | 2,2E-02 | 3,7E-01 | 3,0E-04 | 1,1E-01 | 2,5E-02 | 1,6E-03 | 0,0E+00 | 0,0E+00 | 0,0E+00 | 0,0E+00 | 0,0E+00 | 0,0E+00 | 3263513 | 3264757 | 1245 | - |
| Y11_RS14895 | Y11_31541 | | glutamate mutase L | 0,87 | 2,21 | 2,37 | 1,29 | 1,5 | 0,17 | 6,4E-02 | 4,7E-06 | 4,5E-06 | 7,8E-03 | 4,0E-03 | 2,4E-01 | 0,0E+00 | 1,2E-07 | 8,2E-08 | 1,3E-04 | 8,8E-05 | 0,0E+00 | 3266217 | 3267608 | 1392 | - |
| Y11_RS14950 | Y11_31661 | | CoA-acylating methylmalonate-semialdehyde dehydrogenase | -0,92 | -4,74 | -4,93 | -3,83 | -4,01 | -0,18 | 1,8E-07 | 2,5E-11 | 1,7E-13 | 5,7E-11 | 1,4E-14 | 5,7E-02 | 0,0E+00 | 9,5E-12 | 9,2E-13 | 1,2E-11 | 2,7E-13 | 0,0E+00 | 3279259 | 3280764 | 1506 | + |
| Y11_RS14955 | Y11_31671 | iolB | 5-deoxy-glucuronate isomerase | -0,51 | -2,76 | -2,03 | -2,26 | -1,52 | 0,73 | 5,0E-05 | 9,2E-10 | 6,3E-06 | 4,4E-10 | 3,8E-05 | 5,6E-03 | 0,0E+00 | 1,0E-10 | 1,1E-07 | 4,7E-11 | 1,1E-06 | 0,0E+00 | 3280820 | 3281644 | 825 | + |
| Y11_RS14960 | Y11_31681 | | MurR/RpiR family transcriptional regulator | -0,2 | 1,81 | 2,63 | 2,01 | 2,83 | 0,82 | 5,8E-03 | 1,2E-07 | 1,1E-10 | 2,2E-08 | 2,5E-12 | 3,3E-05 | 0,0E+00 | 4,7E-09 | 2,5E-11 | 9,4E-10 | 6,2E-12 | 0,0E+00 | 3281739 | 3282572 | 834 | - |
| Y11_RS14965 | Y11_31691 | iolC | 5-dehydro-2-deoxygluconokinase | -0,44 | -4,26 | -3,77 | -3,82 | -3,33 | 0,48 | 5,9E-06 | 7,4E-12 | 1,6E-11 | 8,2E-12 | 1,9E-11 | 5,2E-04 | 0,0E+00 | 5,1E-12 | 8,0E-12 | 5,4E-12 | 1,6E-11 | 0,0E+00 | 3282857 | 3284776 | 1920 | - |
| Y11_RS14970 | Y11_31701 | iolD | 3D-(3,5/4)-trihydroxycyclohexane-1,2-dione acylhydrolase (decyclizing) | -0,44 | -3,16 | -1,36 | -2,72 | -0,92 | 1,8 | 7,3E-05 | 1,1E-11 | 2,3E-09 | 1,9E-11 | 2,0E-08 | 5,0E-11 | 0,0E+00 | 6,2E-12 | 1,8E-10 | 6,6E-12 | 0,0E+00 | 3,8E-10 | 3285247 | 3287187 | 1941 | + |
| Y11_RS14975 | Y11_31711 | | Gfo/Idh/MocA family oxidoreductase | -0,23 | -2,82 | -0,79 | -2,59 | -0,56 | 2,03 | 8,2E-04 | 4,0E-12 | 2,3E-06 | 7,5E-11 | 8,1E-05 | 4,7E-09 | 0,0E+00 | 3,9E-12 | 0,0E+00 | 1,5E-11 | 0,0E+00 | 4,2E-09 | 3287267 | 3288280 | 1014 | + |
| Y11_RS14980 | Y11_31721 | iolE | myo-inosose-2 dehydratase | -0,2 | -2,23 | -0,16 | -2,03 | 0,04 | 2,07 | 1,0E-02 | 1,1E-10 | 3,1E-01 | 2,0E-09 | 8,0E-01 | 7,7E-07 | 0,0E+00 | 2,4E-11 | 0,0E+00 | 1,5E-10 | 0,0E+00 | 1,5E-07 | 3288414 | 3289310 | 897 | + |
| Y11_RS14985 | Y11_31731 | | GNAT family N-acetyltransferase | 0,54 | 0,25 | -0,25 | -0,29 | -0,79 | -0,51 | 1,4E-02 | 1,8E-01 | 2,5E-01 | 2,3E-02 | 9,4E-04 | 1,1E-02 | 0,0E+00 | 0,0E+00 | 0,0E+00 | 0,0E+00 | 0,0E+00 | 0,0E+00 | 3289378 | 3289878 | 501 | - |
| Y11_RS14990 | Y11_31741 | | pirin family protein | 0,04 | 0,43 | 0,23 | 0,4 | 0,19 | -0,21 | 6,4E-01 | 4,1E-04 | 4,5E-02 | 4,1E-03 | 1,4E-01 | 1,2E-01 | 0,0E+00 | 0,0E+00 | 0,0E+00 | 0,0E+00 | 0,0E+00 | 0,0E+00 | 3290265 | 3290960 | 696 | + |
| Y11_RS14995 | Y11_31751 | gntR | gluconate operon transcriptional repressor GntR | -0,45 | -0,16 | -0,14 | 0,29 | 0,31 | 0,02 | 1,5E-01 | 5,7E-01 | 7,1E-01 | 1,6E-01 | 3,1E-01 | 9,3E-01 | 0,0E+00 | 0,0E+00 | 0,0E+00 | 0,0E+00 | 0,0E+00 | 0,0E+00 | 3291215 | 3292210 | 996 | + |
| Y11_RS15000 | Y11_31761 | gntT | gluconate transporter | -0,79 | 0,41 | -0,12 | 1,21 | 0,68 | -0,53 | 5,2E-02 | 2,0E-02 | 6,1E-01 | 6,1E-03 | 1,1E-01 | 2,4E-02 | 0,0E+00 | 0,0E+00 | 0,0E+00 | 9,9E-05 | 0,0E+00 | 0,0E+00 | 3292235 | 3293551 | 1317 | - |
| Y11_RS15005 | Y11_31771 | | gluconokinase | -0,1 | 0,63 | 0,2 | 0,73 | 0,3 | -0,43 | 5,2E-01 | 1,5E-05 | 3,7E-01 | 1,2E-03 | 2,5E-01 | 6,6E-02 | 0,0E+00 | 0,0E+00 | 0,0E+00 | 0,0E+00 | 0,0E+00 | 0,0E+00 | 3293773 | 3294285 | 513 | + |
| Y11_RS15015 | Y11_31791 | asd | aspartate-semialdehyde dehydrogenase | 0,91 | 0,27 | 0,28 | -0,64 | -0,63 | 0,02 | 2,9E-05 | 2,0E-02 | 2,1E-02 | 4,7E-04 | 7,3E-04 | 8,9E-01 | 0,0E+00 | 0,0E+00 | 0,0E+00 | 0,0E+00 | 0,0E+00 | 0,0E+00 | 3295222 | 3296325 | 1104 | + |
| Y11_RS15020 | Y11_31801 | alsS | acetolactate synthase AlsS | 0,81 | 1,3 | 1,92 | 0,49 | 1,12 | 0,63 | 4,2E-01 | 2,0E-01 | 6,7E-02 | 3,1E-01 | 2,5E-02 | 9,4E-02 | 0,0E+00 | 3,7E-03 | 7,4E-04 | 0,0E+00 | 5,3E-04 | 0,0E+00 | 3296425 | 3298104 | 1680 | - |
| Y11_RS15040 | Y11_31841 | glgB | 1,4-alpha-glucan branching protein GlgB | -0,19 | 0,68 | 1,22 | 0,87 | 1,4 | 0,53 | 2,0E-01 | 3,2E-04 | 1,9E-04 | 5,6E-04 | 2,0E-04 | 3,2E-02 | 0,0E+00 | 0,0E+00 | 2,5E-06 | 0,0E+00 | 4,9E-06 | 0,0E+00 | 3302122 | 3304305 | 2184 | + |
| Y11_RS15045 | Y11_31851 | glgX | glycogen debranching protein GlgX | 0,35 | 1,34 | 1,46 | 0,99 | 1,1 | 0,12 | 4,1E-02 | 2,0E-07 | 3,3E-04 | 2,0E-04 | 3,9E-03 | 6,5E-01 | 0,0E+00 | 7,3E-09 | 4,2E-06 | 0,0E+00 | 8,6E-05 | 0,0E+00 | 3304307 | 3306295 | 1989 | + |
| Y11_RS15050 | Y11_31861 | glgC | glucose-1-phosphate adenylyltransferase | 0,01 | 1,88 | 2,39 | 1,87 | 2,38 | 0,52 | 9,4E-01 | 9,4E-10 | 3,5E-09 | 1,4E-07 | 7,0E-08 | 7,1E-04 | 0,0E+00 | 1,0E-10 | 2,4E-10 | 4,6E-09 | 4,6E-09 | 0,0E+00 | 3306305 | 3307591 | 1287 | + |
| Y11_RS15055 | Y11_31871 | glgA | glycogen synthase GlgA | 0,14 | 1,6 | 2,23 | 1,46 | 2,08 | 0,62 | 3,0E-02 | 6,2E-08 | 3,2E-11 | 2,7E-07 | 4,9E-10 | 1,1E-04 | 0,0E+00 | 2,8E-09 | 1,3E-11 | 7,9E-09 | 1,1E-10 | 0,0E+00 | 3307775 | 3309205 | 1431 | + |
| Y11_RS15060 | Y11_31881 | glgP | glycogen phosphorylase | 0,22 | 1,93 | 2,72 | 1,71 | 2,5 | 0,79 | 3,1E-03 | 6,0E-10 | 7,5E-10 | 4,6E-09 | 2,5E-09 | 2,4E-05 | 0,0E+00 | 7,4E-11 | 8,3E-11 | 2,7E-10 | 3,4E-10 | 0,0E+00 | 3309252 | 3311699 | 2448 | + |
| Y11_RS15065 | Y11_31891 | glpD | glycerol-3-phosphate dehydrogenase | 0,75 | 0,19 | -1,33 | -0,56 | -2,08 | -1,52 | 2,5E-07 | 1,0E-02 | 1,2E-07 | 6,1E-07 | 9,7E-10 | 2,7E-08 | 0,0E+00 | 0,0E+00 | 3,7E-09 | 0,0E+00 | 1,7E-10 | 1,4E-08 | 3311923 | 3313437 | 1515 | - |
| Y11_RS15075 | Y11_31911 | | DeoR/GlpR family transcriptional regulator | -0,39 | 0,33 | 0,63 | 0,72 | 1,02 | 0,3 | 9,9E-04 | 2,0E-05 | 5,8E-06 | 1,0E-05 | 2,7E-06 | 6,9E-04 | 0,0E+00 | 0,0E+00 | 0,0E+00 | 0,0E+00 | 1,0E-07 | 0,0E+00 | 3314275 | 3315033 | 759 | - |
| Y11_RS15095 | Y11_31951 | malT | HTH-type transcriptional regulator MalT | -0,32 | -1,02 | -0,98 | -0,7 | -0,69 | 0,04 | 3,9E-02 | 5,5E-04 | 2,2E-04 | 6,1E-03 | 1,2E-03 | 8,6E-01 | 0,0E+00 | 1,1E-05 | 0,0E+00 | 0,0E+00 | 0,0E+00 | 0,0E+00 | 3317846 | 3320557 | 2712 | - |
| Y11_RS15100 | Y11_31961 | malP | maltodextrin phosphorylase | 0,68 | -0,37 | -1,48 | -1,05 | -2,16 | -1,11 | 1,3E-04 | 1,9E-03 | 1,0E-05 | 8,7E-06 | 5,2E-07 | 1,0E-04 | 0,0E+00 | 0,0E+00 | 1,7E-07 | 1,8E-07 | 2,5E-08 | 8,4E-06 | 3320876 | 3323281 | 2406 | + |
| Y11_RS15105 | Y11_31971 | malQ | 4-alpha-glucanotransferase | 0,49 | 0,33 | -0,33 | -0,17 | -0,83 | -0,66 | 1,8E-02 | 1,2E-02 | 1,8E-02 | 3,6E-01 | 2,0E-03 | 8,3E-04 | 0,0E+00 | 0,0E+00 | 0,0E+00 | 0,0E+00 | 0,0E+00 | 0,0E+00 | 3323294 | 3325390 | 2097 | + |
| Y11_RS15115 | Y11_31991 | nfuA | Fe-S biogenesis protein NfuA | 0,21 | 0 | 0,08 | -0,22 | -0,13 | 0,08 | 5,2E-03 | 9,5E-01 | 5,6E-01 | 2,2E-03 | 3,5E-01 | 5,3E-01 | 0,0E+00 | 0,0E+00 | 0,0E+00 | 0,0E+00 | 0,0E+00 | 0,0E+00 | 3327152 | 3327727 | 576 | - |
| Y11_RS15125 | Y11_32021 | bioH | pimeloyl-ACP methyl ester esterase BioH | 0,03 | 0,6 | 0,26 | 0,56 | 0,23 | -0,33 | 7,4E-01 | 9,8E-05 | 2,6E-01 | 1,3E-03 | 3,5E-01 | 1,8E-01 | 0,0E+00 | 0,0E+00 | 0,0E+00 | 0,0E+00 | 0,0E+00 | 0,0E+00 | 3328708 | 3329484 | 777 | + |
| Y11_RS15130 | Y11_32031 | | DUF1471 domain-containing protein | 3,63 | 4,37 | 5,47 | 0,74 | 1,84 | 1,09 | 7,1E-10 | 2,2E-10 | 5,6E-11 | 4,9E-04 | 9,2E-07 | 5,4E-05 | 1,7E-10 | 3,8E-11 | 1,7E-11 | 0,0E+00 | 3,9E-08 | 4,7E-06 | 3329599 | 3329862 | 264 | + |
| Y11_RS15135 | Y11_32041 | | ferrous iron transporter C | 0 | 0,73 | 0 | -0,69 | 0 | 1,18 | 0,0E+00 | 1,3E-03 | 0,0E+00 | 2,6E-03 | 0,0E+00 | 1,1E-03 | 0,0E+00 | 0,0E+00 | 0,0E+00 | 0,0E+00 | 0,0E+00 | 7,5E-05 | 3329900 | 3330148 | 249 | - |
| Y11_RS15140 | Y11_32051 | feoB | Fe(2+) transporter permease subunit FeoB | -0,76 | -1,25 | -0,59 | -0,49 | 0,17 | 0,66 | 4,5E-07 | 8,7E-08 | 7,4E-03 | 6,7E-05 | 3,3E-01 | 4,7E-03 | 0,0E+00 | 3,6E-09 | 0,0E+00 | 0,0E+00 | 0,0E+00 | 0,0E+00 | 3330162 | 3332477 | 2316 | - |
| Y11_RS15150 | Y11_32081 | | RNA-binding transcriptional accessory protein | 0,13 | -0,41 | -1,2 | -0,54 | -1,33 | -0,79 | 7,5E-03 | 5,0E-06 | 3,0E-09 | 8,1E-07 | 1,7E-09 | 1,4E-07 | 0,0E+00 | 0,0E+00 | 2,2E-10 | 0,0E+00 | 2,5E-10 | 0,0E+00 | 3333555 | 3335939 | 2385 | - |
| Y11_RS15175 | Y11_32141 | greB | transcription elongation factor GreB | -0,53 | -0,74 | -0,69 | -0,21 | -0,16 | 0,06 | 1,5E-05 | 8,2E-07 | 1,9E-04 | 1,4E-02 | 2,1E-01 | 6,1E-01 | 0,0E+00 | 0,0E+00 | 0,0E+00 | 0,0E+00 | 0,0E+00 | 0,0E+00 | 3340501 | 3340998 | 498 | + |
| Y11_RS15180 | Y11_32151 | ompR | two-component system response regulator OmpR | -1,92 | 0,83 | 0,58 | 2,75 | 2,5 | -0,25 | 6,2E-09 | 5,9E-07 | 1,4E-02 | 1,9E-10 | 1,0E-06 | 2,1E-01 | 1,0E-09 | 0,0E+00 | 0,0E+00 | 2,7E-11 | 4,3E-08 | 0,0E+00 | 3341119 | 3341838 | 720 | + |
| Y11_RS15185 | Y11_32161 | envZ | two-component system sensor histidine kinase EnvZ | -1,26 | 0,69 | 0 | 1,95 | 1,26 | -0,69 | 5,3E-07 | 7,9E-07 | 9,7E-01 | 1,5E-08 | 1,3E-06 | 8,2E-06 | 3,7E-08 | 0,0E+00 | 0,0E+00 | 7,2E-10 | 5,4E-08 | 0,0E+00 | 3341835 | 3343187 | 1353 | + |
| Y11_RS15190 | Y11_32191 | pckA | phosphoenolpyruvate carboxykinase (ATP) | -0,73 | 0,51 | 2,4 | 1,23 | 3,13 | 1,9 | 1,7E-05 | 1,3E-04 | 2,4E-09 | 6,7E-07 | 9,0E-10 | 3,1E-08 | 0,0E+00 | 0,0E+00 | 1,9E-10 | 1,7E-08 | 1,6E-10 | 1,5E-08 | 3344017 | 3345636 | 1620 | - |
| Y11_RS15195 | Y11_32201 | hslO | Hsp33 family molecular chaperone HslO | 0,23 | -0,99 | -0,75 | -1,21 | -0,97 | 0,24 | 2,4E-02 | 1,5E-05 | 1,9E-04 | 4,5E-07 | 6,4E-06 | 7,3E-02 | 0,0E+00 | 0,0E+00 | 0,0E+00 | 1,2E-08 | 0,0E+00 | 0,0E+00 | 3345836 | 3346750 | 915 | - |
| Y11_RS15205 | Y11_32221 | yrfG | GMP/IMP nucleotidase | -0,16 | 0,05 | -0,4 | 0,2 | -0,24 | -0,44 | 3,8E-01 | 7,8E-01 | 6,1E-02 | 2,6E-02 | 7,0E-02 | 2,6E-03 | 0,0E+00 | 0,0E+00 | 0,0E+00 | 0,0E+00 | 0,0E+00 | 0,0E+00 | 3347224 | 3347901 | 678 | - |
| Y11_RS15215 | Y11_32261 | nudE | ADP compounds hydrolase NudE | -0,17 | -0,81 | -0,64 | -0,63 | -0,47 | 0,16 | 2,8E-02 | 3,4E-06 | 1,8E-04 | 7,1E-05 | 2,6E-03 | 1,8E-01 | 0,0E+00 | 0,0E+00 | 0,0E+00 | 0,0E+00 | 0,0E+00 | 0,0E+00 | 3350663 | 3351220 | 558 | + |
| Y11_RS15220 | Y11_32271 | mrcA | peptidoglycan glycosyltransferase/peptidoglycan DD-transpeptidase MrcA | -0,27 | -0,32 | -0,52 | -0,04 | -0,24 | -0,2 | 2,7E-04 | 1,8E-04 | 2,2E-03 | 3,2E-01 | 6,7E-02 | 1,3E-01 | 0,0E+00 | 0,0E+00 | 0,0E+00 | 0,0E+00 | 0,0E+00 | 0,0E+00 | 3351290 | 3353845 | 2556 | - |
| Y11_RS15250 | Y11_32331 | aroK | shikimate kinase AroK | -0,17 | -1,59 | -1,76 | -1,42 | -1,6 | -0,18 | 1,2E-02 | 4,6E-10 | 1,2E-08 | 3,1E-10 | 1,8E-08 | 3,1E-02 | 0,0E+00 | 6,2E-11 | 6,2E-10 | 3,5E-11 | 1,5E-09 | 0,0E+00 | 3358017 | 3358538 | 522 | + |
| Y11_RS15255 | Y11_32341 | aroB | 3-dehydroquinate synthase | -0,25 | -1,43 | -1,01 | -1,18 | -0,76 | 0,42 | 7,8E-03 | 8,2E-09 | 6,0E-07 | 4,3E-07 | 2,8E-05 | 8,0E-04 | 0,0E+00 | 5,3E-10 | 1,5E-08 | 1,2E-08 | 0,0E+00 | 0,0E+00 | 3358595 | 3359683 | 1089 | + |
| Y11_RS15260 | Y11_32351 | | SPOR domain-containing protein | 0,03 | -1,76 | -1,1 | -1,79 | -1,13 | 0,66 | 7,9E-01 | 2,0E-07 | 9,7E-06 | 3,0E-07 | 1,3E-05 | 6,4E-04 | 0,0E+00 | 7,4E-09 | 1,7E-07 | 8,4E-09 | 4,1E-07 | 0,0E+00 | 3359825 | 3360823 | 999 | + |
| Y11_RS15265 | Y11_32361 | dam | adenine-specific DNA-methyltransferase | -0,18 | -0,86 | -0,68 | -0,67 | -0,5 | 0,18 | 7,2E-02 | 5,4E-04 | 2,0E-02 | 1,7E-03 | 5,9E-02 | 5,0E-01 | 0,0E+00 | 0,0E+00 | 0,0E+00 | 0,0E+00 | 0,0E+00 | 0,0E+00 | 3360907 | 3361719 | 813 | + |
| Y11_RS15270 | Y11_32371 | rpe | ribulose-phosphate 3-epimerase | 0,04 | -0,51 | -0,16 | -0,55 | -0,21 | 0,34 | 6,3E-01 | 6,4E-04 | 8,2E-02 | 3,1E-04 | 3,1E-02 | 4,1E-03 | 0,0E+00 | 0,0E+00 | 0,0E+00 | 0,0E+00 | 0,0E+00 | 0,0E+00 | 3361824 | 3362501 | 678 | + |
| Y11_RS15275 | Y11_32381 | | phosphoglycolate phosphatase | -0,04 | -0,3 | -0,29 | -0,26 | -0,25 | 0,01 | 6,9E-01 | 2,0E-02 | 1,2E-01 | 5,6E-02 | 1,9E-01 | 9,6E-01 | 0,0E+00 | 0,0E+00 | 0,0E+00 | 0,0E+00 | 0,0E+00 | 0,0E+00 | 3362494 | 3363192 | 699 | + |
| Y11_RS15280 | Y11_32391 | trpS | tryptophan--tRNA ligase | 0,19 | -0,63 | -0,48 | -0,82 | -0,67 | 0,15 | 5,8E-03 | 3,6E-06 | 1,7E-04 | 6,3E-09 | 2,4E-06 | 4,4E-02 | 0,0E+00 | 0,0E+00 | 0,0E+00 | 0,0E+00 | 0,0E+00 | 0,0E+00 | 3363194 | 3364234 | 1041 | + |
| Y11_RS15290 | Y11_32411 | nirC | nitrite transporter NirC | 0 | -15 | -15 | 0,67 | 0 | -0,7 | 0,0E+00 | 0,0E+00 | 0,0E+00 | 6,4E-02 | 9,9E-01 | 2,0E-01 | 0,0E+00 | 0,0E+00 | 0,0E+00 | 0,0E+00 | 0,0E+00 | 0,0E+00 | 3365872 | 3366678 | 807 | - |
| Y11_RS15295 | Y11_32421 | nirD | nitrite reductase small subunit NirD | 0,45 | 0,14 | -0,82 | -0,3 | -1,26 | -0,95 | 5,1E-02 | 4,9E-01 | 1,0E-01 | 1,1E-01 | 2,1E-02 | 5,9E-02 | 0,0E+00 | 0,0E+00 | 0,0E+00 | 0,0E+00 | 4,3E-04 | 0,0E+00 | 3366770 | 3367096 | 327 | - |
| Y11_RS15305 | Y11_32451 | | cytosine deaminase | 0,23 | 1,57 | 1,83 | 1,33 | 1,6 | 0,27 | 3,2E-01 | 5,4E-06 | 3,7E-06 | 3,3E-04 | 1,3E-04 | 1,4E-01 | 0,0E+00 | 1,4E-07 | 7,0E-08 | 5,7E-06 | 3,4E-06 | 0,0E+00 | 3369920 | 3371215 | 1296 | + |
| Y11_RS15315 | Y11_32471 | | LacI family transcriptional regulator | 0,41 | 1,3 | 1,22 | 0,89 | 0,81 | -0,08 | 1,3E-02 | 5,7E-06 | 5,8E-06 | 1,9E-05 | 1,9E-05 | 3,5E-01 | 0,0E+00 | 1,4E-07 | 1,0E-07 | 0,0E+00 | 0,0E+00 | 0,0E+00 | 3372762 | 3373742 | 981 | - |
| Y11_RS15320 | Y11_32481 | | 6-phospho-beta-glucosidase | -2,06 | 0,36 | 1,86 | 2,42 | 3,92 | 1,5 | 5,2E-09 | 4,2E-03 | 1,2E-06 | 3,7E-09 | 3,5E-09 | 8,2E-06 | 9,1E-10 | 0,0E+00 | 2,7E-08 | 2,3E-10 | 4,3E-10 | 9,7E-07 | 3373824 | 3375137 | 1314 | - |
| Y11_RS15325 | Y11_32501 | ppiA | peptidylprolyl isomerase A | -0,01 | -0,92 | 0,26 | -0,91 | 0,27 | 1,18 | 9,1E-01 | 5,9E-06 | 9,0E-02 | 2,5E-05 | 1,0E-01 | 4,2E-05 | 0,0E+00 | 0,0E+00 | 0,0E+00 | 0,0E+00 | 0,0E+00 | 3,7E-06 | 3375490 | 3376059 | 570 | + |
| Y11_RS15330 | Y11_32511 | | DUF853 family protein | -0,55 | -0,15 | 1,34 | 0,4 | 1,88 | 1,48 | 1,5E-04 | 6,6E-02 | 1,8E-06 | 2,1E-03 | 3,4E-07 | 1,3E-06 | 0,0E+00 | 0,0E+00 | 3,8E-08 | 0,0E+00 | 1,7E-08 | 2,2E-07 | 3376193 | 3377704 | 1512 | + |
| Y11_RS15335 | Y11_32521 | | aminodeoxychorismate synthase component II | -0,6 | -0,74 | -0,36 | -0,13 | 0,24 | 0,38 | 3,4E-04 | 3,3E-05 | 3,0E-02 | 1,2E-01 | 9,5E-02 | 1,3E-02 | 0,0E+00 | 0,0E+00 | 0,0E+00 | 0,0E+00 | 0,0E+00 | 0,0E+00 | 3377725 | 3378300 | 576 | + |
| Y11_RS15345 | Y11_32551 | | YccS/YhfK family putative transporter | 0,18 | 0,68 | 1,1 | 0,5 | 0,91 | 0,41 | 1,8E-01 | 6,2E-04 | 4,0E-05 | 2,2E-03 | 7,9E-05 | 1,3E-02 | 0,0E+00 | 0,0E+00 | 5,9E-07 | 0,0E+00 | 0,0E+00 | 0,0E+00 | 3379948 | 3382068 | 2121 | - |
| Y11_RS15350 | Y11_32561 | crp | cAMP-activated global transcriptional regulator CRP | 0,23 | 0,64 | 1 | 0,41 | 0,77 | 0,36 | 5,1E-03 | 4,5E-07 | 1,4E-05 | 3,0E-04 | 1,9E-04 | 1,2E-02 | 0,0E+00 | 0,0E+00 | 2,3E-07 | 0,0E+00 | 0,0E+00 | 0,0E+00 | 3382346 | 3382978 | 633 | - |
| Y11_RS15355 | Y11_32571 | | OsmC family protein | 0,16 | -0,08 | 0,13 | -0,24 | -0,03 | 0,21 | 1,4E-02 | 2,6E-01 | 4,7E-01 | 1,1E-02 | 8,6E-01 | 2,8E-01 | 0,0E+00 | 0,0E+00 | 0,0E+00 | 0,0E+00 | 0,0E+00 | 0,0E+00 | 3383313 | 3383720 | 408 | + |
| Y11_RS15360 | Y11_32581 | | phosphoribulokinase | 0,47 | -0,09 | 0,17 | -0,55 | -0,29 | 0,26 | 1,3E-04 | 1,8E-01 | 2,7E-01 | 3,1E-05 | 8,4E-02 | 1,1E-01 | 0,0E+00 | 0,0E+00 | 0,0E+00 | 0,0E+00 | 0,0E+00 | 0,0E+00 | 3383863 | 3384732 | 870 | - |
| Y11_RS15365 | Y11_32591 | | YheU family protein | 0 | -15 | -15 | -15 | -15 | 0,55 | 0,0E+00 | 0,0E+00 | 0,0E+00 | 0,0E+00 | 0,0E+00 | 2,8E-01 | 0,0E+00 | 0,0E+00 | 0,0E+00 | 0,0E+00 | 0,0E+00 | 0,0E+00 | 3384874 | 3385110 | 237 | - |
| Y11_RS15370 | Y11_32601 | | hydrolase | 0 | 0,87 | 0,51 | 0,2 | -0,13 | -0,33 | 0,0E+00 | 4,7E-03 | 3,6E-02 | 4,0E-01 | 5,4E-01 | 3,0E-01 | 0,0E+00 | 0,0E+00 | 0,0E+00 | 0,0E+00 | 0,0E+00 | 0,0E+00 | 3385107 | 3386087 | 981 | - |
| Y11_RS15380 | Y11_32621 | | PhzF family phenazine biosynthesis protein | 0 | 0,45 | 0,73 | -15 | -15 | 0,33 | 0,0E+00 | 5,5E-02 | 3,2E-02 | 0,0E+00 | 0,0E+00 | 3,6E-01 | 0,0E+00 | 0,0E+00 | 0,0E+00 | 0,0E+00 | 0,0E+00 | 0,0E+00 | 3386812 | 3387594 | 783 | + |
| Y11_RS15385 | Y11_32641 | tauA | taurine ABC transporter substrate-binding protein | 1,1 | -2,4 | 1,28 | -3,5 | 0,17 | 3,67 | 1,9E-04 | 6,2E-07 | 5,7E-04 | 9,8E-11 | 3,6E-01 | 3,2E-08 | 7,8E-06 | 1,9E-08 | 7,0E-06 | 1,9E-11 | 0,0E+00 | 1,5E-08 | 3387781 | 3388809 | 1029 | + |
| Y11_RS15390 | Y11_32651 | tauB | taurine ABC transporter ATP-binding subunit | -1,63 | -1,52 | 0,13 | 0,1 | 1,74 | 1,63 | 4,3E-03 | 4,3E-03 | 7,2E-01 | 7,1E-01 | 4,4E-05 | 9,1E-06 | 1,5E-04 | 8,5E-05 | 0,0E+00 | 0,0E+00 | 1,2E-06 | 1,1E-06 | 3388818 | 3389585 | 768 | + |
| Y11_RS15400 | Y11_32671 | tauD | taurine dioxygenase | -1,67 | -0,82 | 1,68 | 0,85 | 3,35 | 2,51 | 5,6E-06 | 1,8E-03 | 1,3E-04 | 2,4E-04 | 2,8E-07 | 4,5E-06 | 3,0E-07 | 0,0E+00 | 1,7E-06 | 0,0E+00 | 1,4E-08 | 5,9E-07 | 3390469 | 3391317 | 849 | + |
| Y11_RS15405 | Y11_32681 | | glycosyltransferase family 9 protein | -0,55 | -0,3 | 1,01 | 0,26 | 1,56 | 1,31 | 2,7E-06 | 5,4E-04 | 3,8E-06 | 1,2E-03 | 1,3E-07 | 6,7E-07 | 0,0E+00 | 0,0E+00 | 7,1E-08 | 0,0E+00 | 7,4E-09 | 1,4E-07 | 3391352 | 3392323 | 972 | - |
| Y11_RS15415 | Y11_32701 | | ABC transporter ATP-binding protein | 0,09 | 0,05 | -0,04 | -0,03 | -0,13 | -0,09 | 8,7E-02 | 4,4E-01 | 7,0E-01 | 6,0E-01 | 2,7E-01 | 4,5E-01 | 0,0E+00 | 0,0E+00 | 0,0E+00 | 0,0E+00 | 0,0E+00 | 0,0E+00 | 3393446 | 3395362 | 1917 | - |

| | | | | | | | | | | | | | | | | | | | | | | | | | |
|---|---|---|---|---|---|---|---|---|---|---|---|---|---|---|---|---|---|---|---|---|---|---|---|---|---|
| Y11_RS15425 | Y11_32721 | kefB | glutathione-regulated potassium-efflux system protein KefB | -0,1 | -0,63 | -0,23 | -0,53 | -0,14 | 0,4 | 5,3E-01 | 3,5E-03 | 2,3E-01 | 4,2E-03 | 4,3E-01 | 4,9E-02 | 0,0E+00 | 0,0E+00 | 0,0E+00 | 0,0E+00 | 0,0E+00 | 0,0E+00 | 3396277 | 3398085 | 1809 | + |
| Y11_RS15430 | Y11_32731 | | YheV family putative metal-binding protein | 1,82 | 0,73 | 0,48 | -1,09 | -1,34 | -0,24 | 4,0E-03 | 4,7E-02 | 1,8E-01 | 1,6E-02 | 7,6E-03 | 2,0E-01 | 1,4E-04 | 0,0E+00 | 0,0E+00 | 2,5E-04 | 1,6E-04 | 0,0E+00 | 3398100 | 3398321 | 222 | + |
| Y11_RS15435 | Y11_32741 | slyD | peptidylprolyl isomerase | -0,39 | -0,67 | -0,84 | -0,28 | -0,45 | -0,17 | 1,1E-04 | 1,1E-05 | 4,1E-06 | 5,4E-03 | 5,2E-04 | 1,0E-01 | 0,0E+00 | 0,0E+00 | 0,0E+00 | 0,0E+00 | 0,0E+00 | 0,0E+00 | 3398416 | 3399000 | 585 | + |
| Y11_RS15440 | Y11_32751 | | SlyX family protein | 0,08 | -0,19 | 0,46 | -0,27 | 0,38 | 0,65 | 8,3E-01 | 6,0E-01 | 2,2E-01 | 4,7E-01 | 3,3E-01 | 8,7E-02 | 0,0E+00 | 0,0E+00 | 0,0E+00 | 0,0E+00 | 0,0E+00 | 0,0E+00 | 3399066 | 3399284 | 219 | - |
| Y11_RS15445 | Y11_32761 | fkpA | FKBP-type peptidyl-prolyl cis-trans isomerase | -0,54 | 0,16 | 0,41 | 0,69 | 0,95 | 0,26 | 9,3E-05 | 8,8E-04 | 8,8E-02 | 1,2E-05 | 2,8E-03 | 2,6E-01 | 0,0E+00 | 0,0E+00 | 0,0E+00 | 0,0E+00 | 0,0E+00 | 0,0E+00 | 3399569 | 3400393 | 825 | + |
| Y11_RS15450 | Y11_32771 | | transcriptional regulator | -0,18 | -0,34 | -0,93 | -0,15 | -0,74 | -0,59 | 5,6E-03 | 1,5E-04 | 9,1E-06 | 2,9E-02 | 6,3E-05 | 3,4E-04 | 0,0E+00 | 0,0E+00 | 0,0E+00 | 0,0E+00 | 0,0E+00 | 0,0E+00 | 3400659 | 3401381 | 723 | + |
| Y11_RS15455 | Y11_32781 | tusD | sulfurtransferase complex subunit TusD | -0,58 | -1,09 | -1,64 | -0,52 | -1,07 | -0,55 | 2,1E-02 | 4,9E-04 | 2,1E-05 | 2,8E-02 | 4,2E-04 | 1,5E-02 | 0,0E+00 | 1,0E-05 | 3,3E-07 | 0,0E+00 | 1,0E-05 | 0,0E+00 | 3401381 | 3401791 | 411 | + |
| Y11_RS15465 | Y11_32801 | tusB | sulfurtransferase complex subunit TusB | 0,19 | -0,45 | -0,72 | -0,64 | -0,9 | -0,27 | 1,6E-01 | 1,2E-03 | 1,0E-03 | 5,9E-04 | 4,8E-04 | 9,1E-02 | 0,0E+00 | 0,0E+00 | 0,0E+00 | 0,0E+00 | 0,0E+00 | 0,0E+00 | 3402176 | 3402463 | 288 | + |
| Y11_RS15470 | Y11_32811 | rpsL | 30S ribosomal protein S12 | 0,39 | -0,25 | -0,71 | -0,65 | -1,1 | -0,45 | 7,3E-03 | 3,2E-02 | 7,2E-05 | 4,7E-04 | 9,5E-06 | 1,8E-03 | 0,0E+00 | 0,0E+00 | 0,0E+00 | 0,0E+00 | 3,1E-07 | 0,0E+00 | 3402601 | 3402975 | 375 | + |
| Y11_RS15475 | Y11_32821 | rpsG | 30S ribosomal protein S7 | 0,16 | -0,22 | -0,56 | -0,37 | -0,71 | -0,34 | 4,6E-03 | 1,0E-02 | 1,1E-04 | 1,8E-04 | 9,4E-06 | 5,1E-03 | 0,0E+00 | 0,0E+00 | 0,0E+00 | 0,0E+00 | 0,0E+00 | 0,0E+00 | 3403072 | 3403542 | 471 | + |
| Y11_RS15480 | Y11_32831 | fusA | elongation factor G | -0,37 | -0,82 | -1,15 | -0,46 | -0,78 | -0,33 | 3,5E-04 | 1,4E-04 | 9,1E-08 | 6,3E-03 | 3,7E-06 | 3,2E-02 | 0,0E+00 | 0,0E+00 | 3,0E-09 | 0,0E+00 | 0,0E+00 | 0,0E+00 | 3403636 | 3405744 | 2109 | + |
| Y11_RS15485 | Y11_32841 | tuf | elongation factor Tu | 0,07 | -0,4 | -1,04 | -0,47 | -1,1 | -0,64 | 7,1E-01 | 1,4E-01 | 7,2E-05 | 9,9E-02 | 9,9E-05 | 2,3E-02 | 0,0E+00 | 0,0E+00 | 1,0E-06 | 0,0E+00 | 2,6E-06 | 0,0E+00 | 3405816 | 3407000 | 1185 | + |
| Y11_RS15495 | Y11_32861 | bfr | bacterioferritin | 0,68 | 3,87 | 4,66 | 3,19 | 3,98 | 0,79 | 5,2E-05 | 2,7E-10 | 1,1E-09 | 2,2E-10 | 1,8E-09 | 5,9E-04 | 0,0E+00 | 4,2E-11 | 1,1E-10 | 2,8E-11 | 2,7E-10 | 0,0E+00 | 3407417 | 3407890 | 474 | + |
| Y11_RS15500 | Y11_32871 | rpsJ | 30S ribosomal protein S10 | 0,12 | -0,28 | -0,74 | -0,4 | -0,87 | -0,46 | 2,4E-01 | 3,4E-02 | 2,7E-03 | 1,6E-02 | 1,8E-03 | 4,7E-02 | 0,0E+00 | 0,0E+00 | 0,0E+00 | 0,0E+00 | 0,0E+00 | 0,0E+00 | 3408271 | 3408582 | 312 | + |
| Y11_RS15505 | Y11_32881 | rplC | 50S ribosomal protein L3 | 0,37 | -0,92 | -1,13 | -1,29 | -1,5 | -0,21 | 1,3E-04 | 3,2E-10 | 1,7E-03 | 7,5E-09 | 3,1E-04 | 4,2E-01 | 0,0E+00 | 0,0E+00 | 2,0E-05 | 4,1E-10 | 7,6E-06 | 0,0E+00 | 3408615 | 3409244 | 630 | + |
| Y11_RS15510 | Y11_32891 | rplD | 50S ribosomal protein L4 | 0,2 | -0,76 | -1,07 | -0,96 | -1,26 | -0,3 | 9,0E-03 | 1,2E-04 | 2,4E-06 | 4,1E-05 | 1,5E-06 | 5,8E-02 | 0,0E+00 | 0,0E+00 | 4,8E-08 | 0,0E+00 | 6,0E-08 | 0,0E+00 | 3409255 | 3409860 | 606 | + |
| Y11_RS15515 | Y11_32901 | rplW | 50S ribosomal protein L23 | 0,03 | -0,89 | -1,29 | -0,92 | -1,33 | -0,41 | 7,5E-01 | 2,2E-05 | 2,0E-05 | 7,6E-05 | 3,8E-05 | 3,6E-02 | 0,0E+00 | 0,0E+00 | 3,2E-07 | 0,0E+00 | 1,1E-06 | 0,0E+00 | 3409857 | 3410159 | 303 | + |
| Y11_RS15520 | Y11_32911 | rplB | 50S ribosomal protein L2 | 0,41 | -0,58 | -1 | -0,99 | -1,4 | -0,42 | 2,9E-05 | 1,2E-05 | 5,3E-05 | 2,2E-07 | 4,3E-06 | 1,4E-02 | 0,0E+00 | 0,0E+00 | 0,0E+00 | 0,0E+00 | 1,5E-07 | 0,0E+00 | 3410176 | 3411000 | 825 | + |
| Y11_RS15525 | | rpsS | 30S ribosomal protein S19 | 0,16 | 0,06 | -0,45 | -0,1 | -0,61 | -0,51 | 2,1E-01 | 4,7E-01 | 2,0E-03 | 4,7E-01 | 2,6E-03 | 2,1E-03 | 0,0E+00 | 0,0E+00 | 0,0E+00 | 0,0E+00 | 0,0E+00 | 0,0E+00 | 3411015 | 3411293 | 279 | + |
| Y11_RS15530 | Y11_32931 | rplV | 50S ribosomal protein L22 | 0,29 | -0,42 | -0,77 | -0,7 | -1,05 | -0,35 | 6,1E-03 | 2,7E-05 | 1,2E-04 | 1,1E-05 | 2,7E-05 | 1,2E-02 | 0,0E+00 | 0,0E+00 | 0,0E+00 | 0,0E+00 | 7,9E-07 | 0,0E+00 | 3411308 | 3411640 | 333 | + |
| Y11_RS15535 | Y11_32941 | rpsC | 30S ribosomal protein S3 | 0,15 | -0,39 | -0,83 | -0,54 | -0,98 | -0,44 | 1,7E-02 | 1,1E-04 | 1,7E-05 | 1,2E-05 | 5,1E-06 | 1,7E-03 | 0,0E+00 | 0,0E+00 | 0,0E+00 | 0,0E+00 | 0,0E+00 | 0,0E+00 | 3411658 | 3412356 | 699 | + |
| Y11_RS15540 | Y11_32951 | rplP | 50S ribosomal protein L16 | 0,31 | -0,37 | -0,76 | -0,69 | -1,07 | -0,38 | 3,2E-03 | 2,8E-03 | 9,5E-06 | 1,3E-04 | 2,3E-06 | 5,2E-03 | 0,0E+00 | 0,0E+00 | 0,0E+00 | 0,0E+00 | 8,7E-08 | 0,0E+00 | 3412369 | 3412779 | 411 | + |
| Y11_RS15545 | Y11_32961 | rpmC | 50S ribosomal protein L29 | 0,32 | -0,63 | -1,14 | -0,95 | -1,45 | -0,51 | 6,2E-02 | 1,9E-03 | 6,4E-04 | 3,1E-08 | 2,0E-05 | 1,3E-02 | 0,0E+00 | 0,0E+00 | 7,8E-06 | 0,0E+00 | 6,1E-07 | 0,0E+00 | 3412779 | 3412970 | 192 | + |
| Y11_RS15550 | | rpsQ | 30S ribosomal protein S17 | -0,01 | 0,29 | -0,14 | 0,29 | -0,14 | -0,43 | 9,4E-01 | 1,6E-02 | 5,8E-01 | 3,1E-02 | 6,0E-01 | 1,3E-01 | 0,0E+00 | 0,0E+00 | 0,0E+00 | 0,0E+00 | 0,0E+00 | 0,0E+00 | 3412970 | 3413224 | 255 | + |
| Y11_RS15555 | Y11_32971 | rplN | 50S ribosomal protein L14 | 0,27 | -0,76 | -1,36 | -1,03 | -1,62 | -0,6 | 1,0E-02 | 9,5E-06 | 3,8E-06 | 4,1E-07 | 7,1E-07 | 8,5E-04 | 0,0E+00 | 0,0E+00 | 7,2E-08 | 1,1E-08 | 3,1E-08 | 0,0E+00 | 3413390 | 3413761 | 372 | + |
| Y11_RS15560 | Y11_32981 | rplX | 50S ribosomal protein L24 | 0,31 | -0,69 | -0,82 | -1 | -1,13 | -0,13 | 2,1E-02 | 1,5E-03 | 1,8E-03 | 3,5E-05 | 1,0E-04 | 5,1E-01 | 0,0E+00 | 0,0E+00 | 0,0E+00 | 0,0E+00 | 2,7E-06 | 0,0E+00 | 3413772 | 3414086 | 315 | + |
| Y11_RS15565 | Y11_32991 | rplE | 50S ribosomal protein L5 | 0,31 | -0,69 | -1,15 | -1 | -1,46 | -0,47 | 3,0E-03 | 3,1E-04 | 3,4E-05 | 1,2E-05 | 3,8E-06 | 1,8E-02 | 0,0E+00 | 0,0E+00 | 5,1E-07 | 0,0E+00 | 1,4E-07 | 0,0E+00 | 3414101 | 3414640 | 540 | + |
| Y11_RS15570 | Y11_33001 | rpsN | 30S ribosomal protein S14 | 0,34 | -0,11 | -0,24 | -0,46 | -0,58 | -0,13 | 1,2E-03 | 5,8E-02 | 8,4E-02 | 5,4E-05 | 1,6E-03 | 3,0E-01 | 0,0E+00 | 0,0E+00 | 0,0E+00 | 0,0E+00 | 0,0E+00 | 0,0E+00 | 3414654 | 3414959 | 306 | + |
| Y11_RS15575 | Y11_33011 | rpsH | 30S ribosomal protein S8 | 0,14 | -0,02 | -0,45 | -0,16 | -0,59 | -0,43 | 2,3E-01 | 8,1E-01 | 1,6E-02 | 1,8E-01 | 5,3E-03 | 2,2E-02 | 0,0E+00 | 0,0E+00 | 0,0E+00 | 0,0E+00 | 0,0E+00 | 0,0E+00 | 3414993 | 3415385 | 393 | + |
| Y11_RS15580 | Y11_33021 | rplF | 50S ribosomal protein L6 | 0,17 | -0,52 | -0,97 | -0,69 | -1,14 | -0,45 | 8,1E-02 | 1,2E-03 | 4,4E-04 | 7,0E-04 | 2,7E-04 | 5,0E-02 | 0,0E+00 | 0,0E+00 | 0,0E+00 | 0,0E+00 | 6,6E-06 | 0,0E+00 | 3415400 | 3415933 | 534 | + |
| Y11_RS15585 | Y11_33031 | rplR | 50S ribosomal protein L18 | 0,37 | -1,18 | -1,7 | -1,55 | -2,07 | -0,52 | 1,2E-04 | 2,4E-05 | 2,4E-07 | 2,6E-06 | 3,7E-08 | 1,2E-02 | 0,0E+00 | 5,6E-07 | 6,7E-09 | 5,9E-08 | 2,7E-09 | 0,0E+00 | 3415943 | 3416296 | 354 | + |
| Y11_RS15590 | Y11_33041 | rpsE | 30S ribosomal protein S5 | 0,03 | -0,52 | -1,3 | -0,55 | -1,33 | -0,78 | 8,1E-01 | 4,6E-03 | 7,8E-05 | 3,9E-03 | 7,5E-05 | 3,2E-03 | 0,0E+00 | 0,0E+00 | 1,1E-06 | 0,0E+00 | 2,0E-06 | 0,0E+00 | 3416311 | 3416814 | 504 | + |
| Y11_RS15595 | Y11_33051 | rpmD | 50S ribosomal protein L30 | 0,19 | -1,03 | -1,69 | -1,22 | -1,88 | -0,66 | 2,9E-02 | 1,8E-05 | 2,6E-06 | 5,0E-06 | 1,1E-06 | 4,3E-03 | 0,0E+00 | 4,3E-07 | 5,1E-08 | 1,1E-07 | 4,6E-08 | 0,0E+00 | 3416817 | 3416996 | 180 | + |
| Y11_RS15600 | Y11_33061 | rplO | 50S ribosomal protein L15 | 0,28 | -0,9 | -1,41 | -1,18 | -1,7 | -0,51 | 2,4E-04 | 2,4E-07 | 3,0E-05 | 8,2E-08 | 8,9E-06 | 1,8E-02 | 0,0E+00 | 0,0E+00 | 4,6E-07 | 2,9E-09 | 2,9E-07 | 0,0E+00 | 3417000 | 3417434 | 435 | + |
| Y11_RS15605 | Y11_33071 | secY | preprotein translocase subunit SecY | -0,68 | -0,8 | -0,76 | -0,11 | -0,08 | 0,03 | 1,1E-05 | 4,5E-05 | 2,0E-05 | 2,2E-01 | 2,7E-01 | 7,5E-01 | 0,0E+00 | 0,0E+00 | 0,0E+00 | 0,0E+00 | 0,0E+00 | 0,0E+00 | 3417442 | 3418773 | 1332 | + |
| Y11_RS15615 | Y11_33091 | rpsM | 30S ribosomal protein S13 | 0,03 | -0,36 | -0,76 | -0,38 | -0,78 | -0,4 | 7,0E-01 | 7,7E-03 | 4,9E-04 | 1,1E-02 | 6,7E-04 | 4,3E-02 | 0,0E+00 | 0,0E+00 | 0,0E+00 | 0,0E+00 | 0,0E+00 | 0,0E+00 | 3419072 | 3419428 | 357 | + |
| Y11_RS15620 | Y11_33101 | rpsK | 30S ribosomal protein S11 | 0,33 | -0,26 | -0,8 | -0,58 | -1,13 | -0,54 | 4,2E-03 | 8,4E-03 | 7,2E-03 | 6,5E-05 | 1,0E-03 | 4,0E-02 | 0,0E+00 | 0,0E+00 | 0,0E+00 | 0,0E+00 | 2,4E-05 | 0,0E+00 | 3419445 | 3419834 | 390 | + |
| Y11_RS15625 | Y11_33111 | rpsD | 30S ribosomal protein S4 | 0,18 | -0,36 | -0,89 | -0,54 | -1,07 | -0,53 | 1,5E-02 | 5,2E-04 | 5,8E-07 | 1,2E-04 | 6,4E-07 | 2,0E-04 | 0,0E+00 | 0,0E+00 | 0,0E+00 | 0,0E+00 | 2,9E-08 | 0,0E+00 | 3419865 | 3420485 | 621 | + |
| Y11_RS15630 | Y11_33121 | rpoA | DNA-directed RNA polymerase subunit alpha | -0,3 | -0,35 | -0,55 | -0,05 | -0,25 | -0,2 | 2,3E-04 | 2,2E-04 | 1,9E-04 | 3,1E-01 | 1,2E-02 | 3,9E-02 | 0,0E+00 | 0,0E+00 | 0,0E+00 | 0,0E+00 | 0,0E+00 | 0,0E+00 | 3420511 | 3421503 | 993 | + |
| Y11_RS15635 | Y11_33131 | rplQ | 50S ribosomal protein L17 | 0,2 | -0,92 | -1,47 | -1,12 | -1,67 | -0,55 | 1,3E-02 | 7,1E-06 | 3,9E-08 | 2,2E-06 | 2,3E-08 | 5,8E-04 | 0,0E+00 | 0,0E+00 | 1,6E-09 | 5,0E-08 | 1,9E-09 | 0,0E+00 | 3421544 | 3421933 | 390 | + |
| Y11_RS15660 | Y11_33191 | mscL | large-conductance mechanosensitive channel protein MscL | 0,57 | 2,44 | 3,33 | 1,87 | 2,76 | 0,89 | 1,3E-03 | 6,6E-08 | 6,9E-08 | 6,8E-08 | 1,1E-07 | 6,5E-04 | 0,0E+00 | 2,9E-09 | 2,4E-09 | 2,4E-09 | 6,9E-09 | 0,0E+00 | 3424838 | 3425248 | 411 | - |
| Y11_RS15665 | Y11_33201 | trkA | Trk system potassium transporter TrkA | 0,36 | 0,77 | 0,81 | 0,42 | 0,45 | 0,04 | 2,0E-02 | 6,1E-06 | 1,6E-04 | 3,6E-03 | 1,2E-02 | 7,1E-01 | 0,0E+00 | 0,0E+00 | 0,0E+00 | 0,0E+00 | 0,0E+00 | 0,0E+00 | 3425368 | 3426744 | 1377 | - |
| Y11_RS15670 | Y11_33211 | rsmB | 16S rRNA (cytosine(967)-C(5))-methyltransferase RsmB | 0,18 | 0,09 | -0,27 | -0,09 | -0,46 | -0,37 | 9,0E-03 | 2,0E-01 | 4,6E-04 | 1,5E-01 | 1,4E-06 | 1,2E-04 | 0,0E+00 | 0,0E+00 | 0,0E+00 | 0,0E+00 | 0,0E+00 | 0,0E+00 | 3426796 | 3428085 | 1290 | - |
| Y11_RS15675 | Y11_33221 | fmt | methionyl-tRNA formyltransferase | 0,19 | 0,94 | 0,7 | 0,75 | 0,5 | -0,25 | 4,5E-05 | 7,9E-10 | 4,0E-06 | 1,9E-08 | 6,0E-05 | 6,2E-03 | 0,0E+00 | 0,0E+00 | 0,0E+00 | 0,0E+00 | 0,0E+00 | 0,0E+00 | 3428447 | 3429394 | 948 | - |
| Y11_RS15680 | Y11_33231 | def | peptide deformylase | 0,6 | 0,35 | 0,08 | -0,26 | -0,52 | -0,26 | 3,5E-05 | 3,3E-03 | 4,8E-01 | 2,7E-02 | 2,7E-03 | 7,5E-02 | 0,0E+00 | 0,0E+00 | 0,0E+00 | 0,0E+00 | 0,0E+00 | 0,0E+00 | 3429419 | 3429931 | 513 | - |
| Y11_RS15695 | Y11_33261 | | topoisomerase DNA-binding C4 zinc finger domain-containing protein | 0,13 | 0,78 | 0,56 | 0,65 | 0,42 | -0,23 | 4,0E-01 | 4,3E-04 | 5,5E-02 | 3,0E-04 | 1,1E-01 | 3,4E-01 | 0,0E+00 | 0,0E+00 | 0,0E+00 | 0,0E+00 | 0,0E+00 | 0,0E+00 | 3431680 | 3432216 | 537 | + |
| Y11_RS15700 | Y11_33271 | tsaC | L-threonylcarbamoyladenylate synthase type 1 TsaC | 0,2 | -0,36 | -0,91 | -0,56 | -1,11 | -0,55 | 1,9E-02 | 2,8E-03 | 3,7E-03 | 2,0E-04 | 1,2E-03 | 4,5E-02 | 0,0E+00 | 0,0E+00 | 0,0E+00 | 0,0E+00 | 2,6E-05 | 0,0E+00 | 3432224 | 3432796 | 573 | + |
| Y11_RS15705 | Y11_33281 | aroE | shikimate dehydrogenase | 0,12 | 0,32 | -0,24 | 0,2 | -0,37 | -0,57 | 2,6E-01 | 4,8E-02 | 6,7E-02 | 1,3E-01 | 3,6E-03 | 2,4E-03 | 0,0E+00 | 0,0E+00 | 0,0E+00 | 0,0E+00 | 0,0E+00 | 0,0E+00 | 3432805 | 3433626 | 822 | + |
| Y11_RS15710 | Y11_33291 | | gamma carbonic anhydrase family protein | 0,75 | 0,43 | 0,3 | -0,31 | -0,43 | -0,13 | 3,6E-03 | 3,7E-02 | 5,1E-01 | 1,5E-02 | 3,3E-01 | 7,7E-01 | 0,0E+00 | 0,0E+00 | 0,0E+00 | 0,0E+00 | 0,0E+00 | 0,0E+00 | 3433686 | 3434228 | 543 | - |
| Y11_RS15750 | Y11_33331 | | ABC transporter substrate-binding protein | -0,31 | 3,48 | 3,85 | 3,79 | 4,15 | 0,37 | 7,8E-03 | 1,3E-11 | 6,8E-10 | 1,2E-10 | 1,1E-09 | 1,4E-02 | 0,0E+00 | 6,5E-12 | 8,0E-11 | 2,0E-11 | 1,8E-10 | 0,0E+00 | 3441186 | 3442142 | 957 | + |
| Y11_RS15755 | Y11_33341 | | sugar ABC transporter ATP-binding protein | -0,38 | 1,71 | 1,99 | 2,09 | 2,37 | 0,28 | 3,0E-03 | 1,3E-09 | 2,6E-09 | 1,6E-08 | 1,3E-08 | 5,8E-03 | 0,0E+00 | 1,3E-10 | 1,9E-10 | 7,5E-10 | 1,2E-09 | 0,0E+00 | 3442227 | 3443717 | 1491 | + |
| Y11_RS15770 | Y11_33371 | hdfR | HTH-type transcriptional regulator HdfR | 0,88 | 1,63 | 1,95 | 0,76 | 1,08 | 0,32 | 1,4E-03 | 5,0E-07 | 6,4E-07 | 4,5E-03 | 8,9E-04 | 6,9E-02 | 0,0E+00 | 1,6E-08 | 1,6E-08 | 0,0E+00 | 2,1E-05 | 0,0E+00 | 3445727 | 3446608 | 882 | - |
| Y11_RS15775 | Y11_33381 | | DUF413 domain-containing protein | 0,49 | 0,02 | 0,59 | -0,47 | 0,1 | 0,57 | 9,7E-05 | 7,8E-01 | 2,4E-05 | 1,4E-04 | 1,9E-01 | 3,3E-05 | 0,0E+00 | 0,0E+00 | 0,0E+00 | 0,0E+00 | 0,0E+00 | 0,0E+00 | 3446728 | 3447066 | 339 | + |
| Y11_RS15795 | Y11_33431 | | branched-chain amino acid transaminase | -0,31 | 1,08 | 0,88 | 1,39 | 1,19 | -0,2 | 1,6E-02 | 8,9E-07 | 1,3E-05 | 1,0E-07 | 1,2E-06 | 2,0E-02 | 0,0E+00 | 2,6E-08 | 0,0E+00 | 3,4E-09 | 4,8E-08 | 0,0E+00 | 3451189 | 3452115 | 927 | + |
| Y11_RS15800 | Y11_33451 | ilvD | dihydroxy-acid dehydratase | -0,98 | 0,29 | 0,41 | 1,28 | 1,39 | 0,12 | 8,9E-09 | 9,0E-05 | 6,1E-06 | 2,3E-10 | 7,6E-11 | 6,1E-03 | 0,0E+00 | 0,0E+00 | 0,0E+00 | 2,9E-11 | 3,9E-11 | 0,0E+00 | 3452195 | 3454045 | 1851 | + |
| Y11_RS15805 | Y11_33461 | ilvA | threonine ammonia-lyase, biosynthetic | -0,93 | 0,51 | 0,48 | 1,44 | 1,41 | -0,03 | 1,8E-07 | 5,9E-05 | 7,9E-05 | 7,0E-08 | 7,2E-08 | 7,1E-01 | 0,0E+00 | 0,0E+00 | 0,0E+00 | 2,5E-09 | 4,7E-09 | 0,0E+00 | 3454051 | 3455595 | 1545 | + |
| Y11_RS15820 | Y11_33491 | ilvY | HTH-type transcriptional activator IlvY | 0,26 | 0,86 | 0,97 | 0,6 | 0,71 | 0,11 | 2,7E-01 | 1,1E-02 | 1,3E-03 | 1,6E-02 | 1,5E-04 | 5,4E-01 | 0,0E+00 | 0,0E+00 | 0,0E+00 | 0,0E+00 | 0,0E+00 | 0,0E+00 | 3459191 | 3460087 | 897 | - |
| Y11_RS15825 | Y11_33501 | ilvC | ketol-acid reductoisomerase | 0,22 | 1,03 | 0,54 | 0,81 | 0,32 | -0,49 | 2,3E-02 | 3,5E-07 | 7,2E-05 | 4,0E-05 | 1,5E-02 | 8,5E-04 | 0,0E+00 | 1,2E-08 | 0,0E+00 | 0,0E+00 | 0,0E+00 | 0,0E+00 | 3460340 | 3461818 | 1479 | + |
| Y11_RS15830 | Y11_33511 | | S-type pyocin domain-containing protein | 0,54 | 0,26 | 0,23 | -0,28 | -0,31 | -0,04 | 1,7E-02 | 1,7E-01 | 2,3E-01 | 7,3E-02 | 4,7E-02 | 7,8E-01 | 0,0E+00 | 0,0E+00 | 0,0E+00 | 0,0E+00 | 0,0E+00 | 0,0E+00 | 3462190 | 3464904 | 2715 | + |
| Y11_RS15835 | Y11_33521 | | bacteriocin immunity protein | -0,35 | 0,5 | 0,41 | 0,86 | 0,77 | -0,09 | 3,3E-02 | 4,3E-03 | 6,1E-02 | 8,0E-05 | 3,0E-03 | 6,2E-01 | 0,0E+00 | 0,0E+00 | 0,0E+00 | 0,0E+00 | 0,0E+00 | 0,0E+00 | 3464906 | 3465160 | 255 | + |
| Y11_RS15850 | Y11_33551 | | molecular chaperone | 0,33 | 2,23 | 2,98 | 1,9 | 2,65 | 0,75 | 2,5E-03 | 8,7E-08 | 1,9E-07 | 4,3E-07 | 5,6E-07 | 6,9E-03 | 0,0E+00 | 3,6E-09 | 5,6E-09 | 1,2E-08 | 2,6E-08 | 0,0E+00 | 3466519 | 3467277 | 759 | - |
| Y11_RS15865 | Y11_33581 | | molecular chaperone | 0,58 | 3,66 | 5,66 | 3,08 | 5,07 | 2 | 1,1E-06 | 7,6E-09 | 7,9E-11 | 2,9E-08 | 1,8E-10 | 5,8E-06 | 0,0E+00 | 5,0E-10 | 2,0E-11 | 1,2E-09 | 6,3E-11 | 7,3E-07 | 3471356 | 3472084 | 729 | - |
| Y11_RS15875 | Y11_33601 | | peptidyl-prolyl cis-trans isomerase | -0,36 | 0,91 | 0,91 | 1,28 | 1,28 | 0 | 3,0E-03 | 2,5E-04 | 1,8E-03 | 6,5E-05 | 3,6E-04 | 1,0E+00 | 0,0E+00 | 0,0E+00 | 0,0E+00 | 1,2E-06 | 8,7E-06 | 0,0E+00 | 3473364 | 3473645 | 282 | - |
| Y11_RS15880 | Y11_33611 | rep | DNA helicase Rep | -0,42 | -0,37 | -0,1 | 0,05 | 0,32 | 0,27 | 5,3E-06 | 2,6E-05 | 1,3E-01 | 3,1E-01 | 1,2E-03 | 4,2E-03 | 0,0E+00 | 0,0E+00 | 0,0E+00 | 0,0E+00 | 0,0E+00 | 0,0E+00 | 3473859 | 3475883 | 2025 | + |
| Y11_RS15890 | Y11_33631 | ppx | exopolyphosphatase | 0,23 | 2,92 | 15 | 2,63 | 15 | 0 | 4,8E-02 | 5,5E-09 | 0,0E+00 | 1,8E-09 | 0,0E+00 | 0,0E+00 | 0,0E+00 | 3,9E-10 | 0,0E+00 | 1,4E-10 | 0,0E+00 | 0,0E+00 | 3477922 | 3479415 | 1494 | - |
| Y11_RS15895 | Y11_33641 | rhlB | ATP-dependent RNA helicase RhlB | 0,29 | 0,9 | 0,37 | 0,6 | 0,08 | -0,53 | 5,3E-03 | 3,5E-06 | 9,2E-03 | 1,2E-06 | 4,1E-01 | 3,5E-04 | 0,0E+00 | 0,0E+00 | 0,0E+00 | 0,0E+00 | 0,0E+00 | 0,0E+00 | 3479422 | 3480708 | 1287 | - |
| Y11_RS15900 | Y11_33651 | trxA | thioredoxin TrxA | 0,41 | 0,99 | 1,44 | 0,58 | 1,03 | 0,46 | 6,7E-04 | 7,4E-06 | 7,1E-05 | 3,9E-04 | 7,1E-04 | 5,5E-02 | 0,0E+00 | 0,0E+00 | 1,0E-06 | 0,0E+00 | 1,7E-05 | 0,0E+00 | 3480827 | 3481153 | 327 | + |
| Y11_RS15905 | Y11_33661 | rho | transcription termination factor Rho | 0,15 | 0,28 | -0,07 | 0,13 | -0,21 | -0,35 | 2,5E-04 | 1,6E-05 | 5,6E-01 | 1,4E-03 | 8,8E-02 | 1,4E-02 | 0,0E+00 | 0,0E+00 | 0,0E+00 | 0,0E+00 | 0,0E+00 | 0,0E+00 | 3481639 | 3482898 | 1260 | + |

| | | | | | | | | | | | | | | | | | | | | | | | | | |
|---|---|---|---|---|---|---|---|---|---|---|---|---|---|---|---|---|---|---|---|---|---|---|---|---|---|
| Y11_RS15910 | Y11_33671 | wecA | UDP-N-acetylglucosamine--undecaprenyl-phosphate N-acetylglucosaminephosphotransferase | 1,94 | 2,74 | 3,45 | 0,81 | 1,52 | 0 | 1,0E-02 | 1,6E-03 | 1,9E-04 | 4,7E-02 | 2,2E-04 | 0,0E+00 | 3,5E-04 | 3,2E-05 | 2,5E-06 | 0,0E+00 | 5,6E-06 | 0,0E+00 | 3483569 | 3484666 | 1098 | + |
| Y11_RS15915 | Y11_33681 | wzzE | ECA polysaccharide chain length modulation protein | -0,14 | 0,92 | 1,09 | 1,06 | 1,23 | 0,17 | 2,8E-03 | 3,6E-09 | 1,2E-05 | 1,3E-08 | 6,3E-06 | 1,9E-01 | 0,0E+00 | 0,0E+00 | 1,9E-07 | 6,5E-10 | 2,1E-07 | 0,0E+00 | 3484687 | 3485763 | 1077 | + |
| Y11_RS15920 | Y11_33691 | wecB | UDP-N-acetylglucosamine 2-epimerase (non-hydrolyzing) | -0,01 | 0,18 | 0,37 | 0,19 | 0,38 | 0,2 | 7,8E-01 | 6,7E-05 | 1,9E-03 | 6,6E-05 | 1,7E-03 | 4,4E-02 | 0,0E+00 | 0,0E+00 | 0,0E+00 | 0,0E+00 | 0,0E+00 | 0,0E+00 | 3485843 | 3486973 | 1131 | + |
| Y11_RS15925 | Y11_33701 | wecC | UDP-N-acetyl-D-mannosamine dehydrogenase | 0,32 | 1,69 | 1,4 | 1,37 | 1,09 | -0,29 | 6,7E-04 | 1,8E-09 | 1,5E-09 | 4,8E-09 | 4,3E-09 | 5,5E-05 | 0,0E+00 | 1,6E-10 | 1,4E-10 | 2,8E-10 | 4,9E-10 | 0,0E+00 | 3486970 | 3488232 | 1263 | + |
| Y11_RS15930 | Y11_33711 | rffG | dTDP-glucose 4,6-dehydratase | -0,03 | 1,01 | 1,41 | 1,03 | 1,44 | 0,41 | 4,7E-01 | 1,2E-07 | 3,3E-09 | 1,5E-07 | 4,9E-09 | 4,0E-04 | 0,0E+00 | 4,9E-09 | 2,3E-10 | 4,9E-09 | 5,4E-10 | 0,0E+00 | 3488223 | 3489296 | 1074 | + |
| Y11_RS15935 | Y11_33721 | rfbA | glucose-1-phosphate thymidylyltransferase RfbA | 0,14 | 0,75 | 1,4 | 0,61 | 1,26 | 0,65 | 7,4E-03 | 5,9E-07 | 5,7E-09 | 4,5E-07 | 2,2E-09 | 2,7E-06 | 0,0E+00 | 0,0E+00 | 3,5E-10 | 0,0E+00 | 3,1E-10 | 0,0E+00 | 3489556 | 3490437 | 882 | + |
| Y11_RS15940 | Y11_33731 | rffC | dTDP-4-amino-4,6-dideoxy-D-galactose acyltransferase | -0,06 | 0,04 | 0,09 | 0,11 | 0,15 | 0,04 | 5,7E-01 | 6,9E-01 | 2,4E-01 | 4,5E-01 | 1,9E-01 | 7,0E-01 | 0,0E+00 | 0,0E+00 | 0,0E+00 | 0,0E+00 | 0,0E+00 | 0,0E+00 | 3490415 | 3491152 | 738 | + |
| Y11_RS15945 | Y11_33741 | rffA | dTDP-4-amino-4,6-dideoxygalactose transaminase | -0,07 | 0,55 | 0,58 | 0,62 | 0,65 | 0,02 | 5,6E-01 | 8,0E-04 | 4,4E-03 | 5,5E-05 | 1,1E-03 | 8,5E-01 | 0,0E+00 | 0,0E+00 | 0,0E+00 | 0,0E+00 | 0,0E+00 | 0,0E+00 | 3491154 | 3492284 | 1131 | + |
| Y11_RS15955 | Y11_33761 | | TDP-N-acetylfucosamine:lipid II N-acetylfucosaminyltransferase | 0,13 | -0,01 | -0,16 | -0,14 | -0,29 | -0,14 | 1,1E-01 | 8,3E-01 | 6,0E-02 | 4,1E-03 | 4,0E-04 | 4,3E-03 | 0,0E+00 | 0,0E+00 | 0,0E+00 | 0,0E+00 | 0,0E+00 | 0,0E+00 | 3493565 | 3494650 | 1086 | + |
| Y11_RS15965 | Y11_33781 | rffM | lipopolysaccharide N-acetylmannosaminouronosyltransferase | 0,22 | 0,25 | 0,17 | 0,03 | -0,04 | -0,08 | 1,3E-02 | 1,4E-02 | 1,2E-02 | 7,1E-01 | 4,5E-01 | 3,0E-01 | 0,0E+00 | 0,0E+00 | 0,0E+00 | 0,0E+00 | 0,0E+00 | 0,0E+00 | 3496032 | 3496790 | 759 | + |
| Y11_RS16005 | Y11_33831 | hemY | protoheme IX biogenesis protein HemY | 0,13 | 1,29 | 1,56 | 1,17 | 1,44 | 0,27 | 4,1E-02 | 1,3E-09 | 3,1E-09 | 1,1E-08 | 1,3E-08 | 9,6E-04 | 0,0E+00 | 1,3E-10 | 2,2E-10 | 5,6E-10 | 1,2E-09 | 0,0E+00 | 3501545 | 3502744 | 1200 | - |
| Y11_RS16010 | Y11_33841 | hemX | uroporphyrinogen-III C-methyltransferase | 0,51 | 1,09 | 1,57 | 0,57 | 1,06 | 0,48 | 2,8E-06 | 2,2E-10 | 4,0E-07 | 8,3E-07 | 1,1E-05 | 1,7E-03 | 0,0E+00 | 3,8E-11 | 1,1E-08 | 0,0E+00 | 3,7E-07 | 0,0E+00 | 3502747 | 3503871 | 1125 | - |
| Y11_RS16015 | Y11_33851 | hemD | uroporphyrinogen-III synthase | 0,05 | -0,56 | -0,57 | -0,61 | -0,62 | -0,01 | 6,9E-01 | 1,9E-05 | 5,7E-05 | 1,2E-03 | 1,5E-03 | 8,9E-01 | 0,0E+00 | 0,0E+00 | 0,0E+00 | 0,0E+00 | 0,0E+00 | 0,0E+00 | 3503904 | 3504650 | 747 | - |
| Y11_RS16020 | Y11_33861 | hemC | hydroxymethylbilane synthase | 0,13 | 0,28 | 0,14 | 0,14 | 0 | -0,14 | 8,0E-02 | 3,6E-04 | 1,5E-01 | 6,4E-02 | 9,9E-01 | 1,3E-01 | 0,0E+00 | 0,0E+00 | 0,0E+00 | 0,0E+00 | 0,0E+00 | 0,0E+00 | 3504647 | 3505588 | 942 | - |
| Y11_RS16030 | Y11_33891 | cyaY | iron donor protein CyaY | 0,2 | 1,05 | 1,83 | 0,85 | 1,63 | 0,78 | 1,9E-01 | 4,2E-05 | 4,6E-05 | 2,2E-04 | 1,1E-04 | 9,3E-03 | 0,0E+00 | 9,4E-07 | 6,7E-07 | 0,0E+00 | 2,9E-06 | 0,0E+00 | 3508680 | 3509000 | 321 | - |
| Y11_RS16035 | Y11_33901 | | lipoprotein | 1,29 | 1,15 | 1,85 | -0,13 | 0,56 | 0,69 | 2,4E-05 | 1,2E-04 | 8,4E-04 | 5,4E-01 | 1,8E-01 | 1,1E-01 | 1,1E-06 | 2,6E-06 | 1,0E-05 | 0,0E+00 | 0,0E+00 | 0,0E+00 | 3509114 | 3509317 | 204 | + |
| Y11_RS16040 | Y11_33911 | dapF | diaminopimelate epimerase | 0,58 | 0,61 | 0,75 | 0,03 | 0,18 | 0,15 | 3,1E-03 | 2,0E-03 | 2,7E-04 | 8,6E-01 | 2,3E-01 | 3,0E-01 | 0,0E+00 | 0,0E+00 | 0,0E+00 | 0,0E+00 | 0,0E+00 | 0,0E+00 | 3509406 | 3510230 | 825 | + |
| Y11_RS16045 | Y11_33921 | | DUF484 domain-containing protein | 1,61 | 15 | 15 | 15 | 15 | 0 | 1,4E-03 | 0,0E+00 | 0,0E+00 | 0,0E+00 | 0,0E+00 | 0,0E+00 | 5,0E-05 | 0,0E+00 | 0,0E+00 | 0,0E+00 | 0,0E+00 | 0,0E+00 | 3510370 | 3511074 | 705 | + |
| Y11_RS16055 | Y11_33941 | yigB | 5-amino-6-(5-phospho-D-ribitylamino)uracil phosphatase YigB | 1,14 | 3,34 | 3,31 | 2,2 | 2,29 | 0 | 1,8E-05 | 9,7E-10 | 1,3E-08 | 2,9E-07 | 2,5E-07 | 0,0E+00 | 8,9E-07 | 1,1E-10 | 6,4E-10 | 8,4E-09 | 1,4E-08 | 0,0E+00 | 3511982 | 3512698 | 717 | + |
| Y11_RS16060 | Y11_33951 | uvrD | DNA helicase II | 0,33 | -0,1 | -0,28 | -0,43 | -0,61 | -0,18 | 3,2E-04 | 1,2E-01 | 6,6E-03 | 8,8E-07 | 9,9E-06 | 1,8E-02 | 0,0E+00 | 0,0E+00 | 0,0E+00 | 0,0E+00 | 0,0E+00 | 0,0E+00 | 3512798 | 3514960 | 2163 | + |
| Y11_RS16065 | Y11_33961 | | TetR/AcrR family transcriptional regulator | 2,3 | 2,82 | 3,45 | 0,52 | 1,15 | 0,63 | 1,1E-10 | 1,3E-07 | 9,7E-08 | 1,4E-02 | 3,7E-04 | 3,6E-02 | 3,6E-11 | 5,2E-09 | 3,2E-09 | 0,0E+00 | 8,9E-06 | 0,0E+00 | 3515876 | 3516535 | 660 | - |
| Y11_RS16070 | Y11_33971 | | Tm-1-like ATP-binding domain-containing protein | 5,59 | 6,13 | 6,33 | 0,55 | 0,75 | 0,2 | 2,8E-12 | 1,0E-11 | 7,8E-10 | 4,1E-04 | 3,8E-03 | 3,4E-01 | 3,0E-12 | 6,2E-12 | 8,4E-11 | 0,0E+00 | 0,0E+00 | 0,0E+00 | 3516755 | 3517987 | 1233 | + |
| Y11_RS16075 | Y11_33981 | | phosphoenolpyruvate hydrolase family protein | 5,47 | 5,44 | 5,61 | -0,03 | 0,14 | 0,17 | 1,6E-09 | 8,1E-10 | 2,0E-09 | 7,4E-01 | 3,2E-01 | 1,5E-01 | 3,6E-10 | 3,9E-11 | 1,6E-10 | 0,0E+00 | 0,0E+00 | 0,0E+00 | 3518014 | 3518844 | 831 | + |
| Y11_RS16080 | Y11_33991 | corA | magnesium/cobalt transporter CorA | -1,11 | -2,44 | -2,06 | -1,34 | -0,96 | 0,38 | 1,3E-07 | 1,0E-10 | 3,5E-06 | 2,4E-08 | 8,5E-04 | 7,1E-02 | 1,1E-08 | 2,3E-11 | 6,6E-08 | 1,0E-09 | 0,0E+00 | 0,0E+00 | 3519347 | 3520297 | 951 | + |
| Y11_RS16090 | Y11_34011 | | thioesterase family protein | 0,93 | 1,49 | 2,48 | 0,57 | 1,56 | 0,99 | 5,8E-06 | 4,8E-05 | 1,4E-06 | 1,5E-02 | 3,6E-05 | 4,5E-03 | 0,0E+00 | 1,1E-06 | 3,0E-08 | 0,0E+00 | 1,0E-06 | 0,0E+00 | 3521435 | 3521905 | 471 | - |
| Y11_RS16095 | Y11_34021 | pldA | phospholipase A | 1,02 | 2,82 | 2,91 | 1,8 | 1,89 | 0,09 | 2,3E-07 | 2,9E-10 | 1,4E-11 | 1,8E-08 | 1,3E-09 | 2,5E-01 | 1,8E-08 | 4,4E-11 | 8,0E-12 | 8,2E-10 | 2,1E-10 | 0,0E+00 | 3522086 | 3522964 | 879 | + |
| Y11_RS16100 | Y11_34031 | recQ | ATP-dependent DNA helicase RecQ | 0,16 | 1,46 | 0,62 | 1,3 | 0,46 | -0,84 | 7,4E-02 | 5,8E-08 | 2,2E-04 | 1,8E-06 | 5,5E-03 | 1,2E-04 | 0,0E+00 | 2,6E-09 | 0,0E+00 | 4,3E-08 | 0,0E+00 | 0,0E+00 | 3523041 | 3524891 | 1851 | + |
| Y11_RS16105 | Y11_34041 | rhtC | threonine export protein RhtC | -0,43 | 0,77 | 1,86 | 1,2 | 0 | 1,09 | 1,5E-01 | 7,3E-02 | 2,5E-04 | 1,0E-02 | 0,0E+00 | 2,1E-02 | 0,0E+00 | 0,0E+00 | 3,3E-06 | 1,6E-04 | 0,0E+00 | 1,1E-03 | 3524954 | 3525574 | 621 | + |
| Y11_RS16115 | Y11_34061 | pldB | lysophospholipase L2 | 0,24 | 1,06 | 1,17 | 0,82 | 0,93 | 0,11 | 3,5E-02 | 1,6E-08 | 2,2E-08 | 4,9E-05 | 2,5E-05 | 1,5E-01 | 0,0E+00 | 9,1E-10 | 9,7E-10 | 0,0E+00 | 0,0E+00 | 0,0E+00 | 3526433 | 3527488 | 1056 | + |
| Y11_RS16120 | Y11_34071 | yigL | sugar/pyridoxal phosphate phosphatase YigL | 0,03 | 0,47 | 0,43 | 0,45 | 0,4 | -0,04 | 7,2E-01 | 1,3E-03 | 2,8E-03 | 2,1E-04 | 6,2E-04 | 6,6E-01 | 0,0E+00 | 0,0E+00 | 0,0E+00 | 0,0E+00 | 0,0E+00 | 0,0E+00 | 3527500 | 3528309 | 810 | + |
| Y11_RS16130 | Y11_34091 | glpQ | glycerophosphodiester phosphodiesterase | 0,64 | 0,72 | 0,91 | 0,08 | 0,27 | 0,19 | 6,6E-06 | 1,9E-06 | 9,8E-06 | 2,5E-01 | 2,4E-02 | 8,5E-02 | 0,0E+00 | 0,0E+00 | 0,0E+00 | 0,0E+00 | 0,0E+00 | 0,0E+00 | 3530026 | 3531132 | 1107 | - |
| Y11_RS16135 | Y11_34101 | glpT | glycerol-3-phosphate transporter | 1,45 | 0,73 | 0,5 | -0,72 | -0,96 | -0,23 | 1,0E-04 | 8,3E-03 | 4,3E-02 | 2,0E-03 | 2,9E-04 | 1,9E-01 | 4,5E-06 | 0,0E+00 | 0,0E+00 | 0,0E+00 | 0,0E+00 | 0,0E+00 | 3531286 | 3532641 | 1356 | - |
| Y11_RS16140 | Y11_34121 | glpA | anaerobic glycerol-3-phosphate dehydrogenase subunit A | 0,34 | 15 | -1,17 | 15 | -1,5 | -15 | 1,1E-01 | 0,0E+00 | 2,1E-03 | 0,0E+00 | 2,4E-04 | 0,0E+00 | 0,0E+00 | 0,0E+00 | 2,4E-05 | 0,0E+00 | 5,9E-06 | 0,0E+00 | 3533038 | 3534693 | 1656 | + |
| Y11_RS16155 | Y11_34151 | | DcrB family lipoprotein | 0,77 | 1,41 | 1,55 | 0,64 | 0,78 | 0,13 | 3,1E-05 | 2,8E-06 | 4,1E-06 | 1,5E-04 | 1,8E-04 | 3,8E-01 | 0,0E+00 | 7,4E-08 | 7,5E-08 | 0,0E+00 | 0,0E+00 | 0,0E+00 | 3537375 | 3537899 | 525 | - |
| Y11_RS16165 | Y11_34171 | tusA | sulfurtransferase TusA | 1,04 | -1,72 | -2,16 | -2,76 | -3,2 | -0,43 | 9,0E-04 | 4,2E-05 | 1,4E-03 | 5,8E-06 | 1,5E-04 | 3,9E-01 | 3,4E-05 | 9,4E-07 | 1,6E-05 | 1,2E-07 | 3,8E-06 | 0,0E+00 | 3538887 | 3539141 | 255 | + |
| Y11_RS16180 | Y11_34201 | | DUF1820 family protein | 0,15 | 0,47 | 1,34 | 0,32 | 1,18 | 0,87 | 7,3E-02 | 1,5E-05 | 4,2E-04 | 3,0E-03 | 1,1E-03 | 5,8E-03 | 0,0E+00 | 0,0E+00 | 5,3E-06 | 0,0E+00 | 2,5E-05 | 0,0E+00 | 3542692 | 3543012 | 321 | + |
| Y11_RS16185 | Y11_34211 | | DUF2500 domain-containing protein | 0,8 | 0,41 | 1,4 | -0,39 | 0,6 | 0,99 | 1,7E-04 | 7,1E-03 | 3,4E-07 | 1,7E-02 | 6,3E-04 | 1,2E-05 | 0,0E+00 | 0,0E+00 | 9,1E-09 | 0,0E+00 | 0,0E+00 | 0,0E+00 | 3543051 | 3543434 | 384 | + |
| Y11_RS16195 | Y11_34231 | rsmD | 16S rRNA (guanine(966)-N(2))-methyltransferase | 0,29 | 1,17 | 1,03 | 0,87 | 0,74 | -0,13 | 4,6E-02 | 1,0E-04 | 6,9E-05 | 4,1E-04 | 2,9E-04 | 4,4E-01 | 0,0E+00 | 2,2E-06 | 9,7E-07 | 0,0E+00 | 0,0E+00 | 0,0E+00 | 3543737 | 3544420 | 684 | - |
| Y11_RS16200 | Y11_34251 | ftsY | signal recognition particle-docking protein FtsY | 0,05 | 0,42 | 0,62 | 0,37 | 0,57 | 0,2 | 2,9E-01 | 1,1E-05 | 1,5E-03 | 9,1E-05 | 2,9E-03 | 1,7E-01 | 0,0E+00 | 0,0E+00 | 0,0E+00 | 0,0E+00 | 0,0E+00 | 0,0E+00 | 3544674 | 3546161 | 1488 | + |
| Y11_RS16205 | Y11_34261 | ftsE | cell division ATP-binding protein FtsE | 0,03 | 0,7 | 0,74 | 0,67 | 0,71 | 0,04 | 7,7E-01 | 1,7E-04 | 1,7E-04 | 1,8E-04 | 1,8E-04 | 7,6E-01 | 0,0E+00 | 0,0E+00 | 0,0E+00 | 0,0E+00 | 0,0E+00 | 0,0E+00 | 3546167 | 3546835 | 669 | + |
| Y11_RS16210 | Y11_34271 | ftsX | permease-like cell division protein FtsX | 0,3 | 0,66 | 0,91 | 0,36 | 0,61 | 0,25 | 3,6E-02 | 1,0E-03 | 5,5E-04 | 9,8E-03 | 3,1E-03 | 1,5E-01 | 0,0E+00 | 0,0E+00 | 0,0E+00 | 0,0E+00 | 0,0E+00 | 0,0E+00 | 3546825 | 3547784 | 960 | + |
| Y11_RS16230 | Y11_34311 | panM | aspartate 1-decarboxylase autocleavage activator PanM | -0,25 | 0,32 | 0,17 | 0,57 | 0,42 | -0,15 | 8,7E-03 | 8,8E-02 | 6,9E-02 | 1,3E-02 | 4,1E-03 | 4,3E-01 | 0,0E+00 | 0,0E+00 | 0,0E+00 | 0,0E+00 | 0,0E+00 | 0,0E+00 | 3551565 | 3551957 | 393 | - |
| Y11_RS16235 | Y11_34321 | | branched-chain amino acid ABC transporter substrate-binding protein | 0,28 | 3,3 | 3,2 | 3,02 | 2,92 | -0,1 | 6,0E-03 | 7,3E-08 | 3,1E-09 | 2,4E-07 | 1,9E-08 | 6,3E-01 | 0,0E+00 | 3,1E-09 | 2,2E-10 | 7,0E-09 | 1,6E-09 | 0,0E+00 | 3552379 | 3553494 | 1116 | + |
| Y11_RS16250 | Y11_34351 | livG | high-affinity branched-chain amino acid ABC transporter ATP-binding protein LivG | -0,4 | 1,08 | 0,75 | 1,49 | 1,15 | -0,33 | 1,4E-02 | 1,1E-04 | 2,7E-04 | 1,7E-05 | 2,4E-05 | 7,0E-02 | 0,0E+00 | 2,3E-06 | 0,0E+00 | 3,3E-07 | 7,2E-07 | 0,0E+00 | 3555859 | 3556626 | 768 | + |
| Y11_RS16255 | Y11_34361 | livF | high-affinity branched-chain amino acid ABC transporter ATP-binding protein LivF | -0,11 | 0,72 | 1,21 | 0,83 | 1,32 | 0,49 | 5,1E-01 | 6,3E-07 | 3,0E-06 | 8,3E-04 | 9,6E-05 | 1,7E-03 | 0,0E+00 | 0,0E+00 | 5,9E-08 | 0,0E+00 | 2,6E-06 | 0,0E+00 | 3556671 | 3557372 | 702 | + |
| Y11_RS16275 | Y11_34401 | | fimbria/pilus periplasmic chaperone | 0,86 | 1,88 | 2,93 | 1,01 | 2,06 | 1,05 | 2,3E-02 | 4,3E-04 | 1,6E-05 | 4,9E-03 | 4,1E-05 | 5,6E-03 | 0,0E+00 | 8,8E-06 | 2,6E-07 | 8,0E-05 | 1,1E-06 | 3,3E-04 | 3559739 | 3560422 | 684 | - |
| Y11_RS16310 | Y11_34491 | ugpQ | glycerophosphodiester phosphodiesterase | 0,38 | -1,96 | -2,23 | -2,35 | -2,61 | -0,26 | 1,0E-01 | 2,8E-07 | 2,0E-06 | 1,9E-06 | 3,8E-06 | 1,5E-01 | 0,0E+00 | 9,7E-09 | 4,1E-08 | 4,5E-08 | 1,4E-07 | 0,0E+00 | 3568388 | 3569128 | 741 | + |
| Y11_RS16330 | Y11_34521 | metE | 5-methyltetrahydropteroyltriglutamate--homocysteine S-methyltransferase | 2,95 | 2,79 | 2,41 | -0,19 | -0,55 | 0 | 3,4E-07 | 1,7E-07 | 5,4E-08 | 3,4E-01 | 6,8E-03 | 0,0E+00 | 2,5E-08 | 6,3E-09 | 2,0E-09 | 0,0E+00 | 0,0E+00 | 0,0E+00 | 3572221 | 3574497 | 2277 | + |
| Y11_RS16335 | Y11_34531 | | dienelactone hydrolase family protein | -0,09 | 3,52 | 4,34 | 3,61 | 4,43 | 0,82 | 1,8E-01 | 1,2E-11 | 2,2E-12 | 7,2E-11 | 1,4E-11 | 8,2E-06 | 0,0E+00 | 6,4E-12 | 3,9E-12 | 1,4E-11 | 1,4E-11 | 0,0E+00 | 3574563 | 3575417 | 855 | - |
| Y11_RS16340 | Y11_34541 | udp | uridine phosphorylase | 0,17 | 1,56 | 1,44 | 1,38 | 1,27 | -0,12 | 1,0E-01 | 4,9E-08 | 3,4E-07 | 4,9E-08 | 4,9E-07 | 1,5E-01 | 0,0E+00 | 2,2E-09 | 9,2E-09 | 1,9E-09 | 2,3E-08 | 0,0E+00 | 3576058 | 3576819 | 762 | + |
| Y11_RS16350 | Y11_34591 | | tyrosine-protein phosphatase | 0,52 | 0,69 | 0,77 | 0,17 | 0,25 | 0,08 | 5,5E-02 | 1,4E-02 | 2,8E-02 | 3,8E-01 | 3,7E-01 | 7,5E-01 | 0,0E+00 | 0,0E+00 | 0,0E+00 | 0,0E+00 | 0,0E+00 | 0,0E+00 | 3578535 | 3579332 | 798 | + |
| Y11_RS16360 | Y11_34611 | | DedA family protein | 0,53 | 1,14 | 0,73 | 0,62 | 0,2 | -0,42 | 3,1E-03 | 6,4E-05 | 4,1E-02 | 3,3E-03 | 5,2E-01 | 2,2E-01 | 0,0E+00 | 1,4E-06 | 0,0E+00 | 0,0E+00 | 0,0E+00 | 0,0E+00 | 3581616 | 3582146 | 531 | + |
| Y11_RS16365 | Y11_34621 | rmuC | DNA recombination protein RmuC | 0,63 | -0,16 | -0,23 | -0,79 | -0,86 | -0,08 | 4,2E-06 | 5,1E-03 | 7,8E-02 | 1,3E-06 | 1,1E-04 | 5,3E-01 | 0,0E+00 | 0,0E+00 | 0,0E+00 | 0,0E+00 | 0,0E+00 | 0,0E+00 | 3582347 | 3583873 | 1527 | + |
| Y11_RS16370 | Y11_34631 | ubiE | bifunctional demethylmenaquinone methyltransferase/2-methoxy-6-polyprenyl-1,4-benzoquinol methylase UbiE | -0,02 | 0,63 | 0,35 | 0,64 | 0,37 | -0,28 | 7,4E-01 | 2,5E-06 | 9,5E-02 | 1,1E-07 | 7,7E-02 | 1,7E-01 | 0,0E+00 | 0,0E+00 | 0,0E+00 | 0,0E+00 | 0,0E+00 | 0,0E+00 | 3584007 | 3584762 | 756 | + |
| Y11_RS16375 | Y11_34641 | | SCP2 domain-containing protein | 0,04 | 0,59 | 0,61 | 0,55 | 0,57 | 0,01 | 6,2E-01 | 2,7E-05 | 2,6E-03 | 1,2E-04 | 4,9E-03 | 9,2E-01 | 0,0E+00 | 0,0E+00 | 0,0E+00 | 0,0E+00 | 0,0E+00 | 0,0E+00 | 3584776 | 3585429 | 654 | + |
| Y11_RS16380 | Y11_34651 | ubiB | ubiquinone biosynthesis regulatory protein kinase UbiB | 0,23 | 0,98 | 1,3 | 0,74 | 1,07 | 0,33 | 7,7E-03 | 2,9E-08 | 1,2E-08 | 1,4E-06 | 2,0E-07 | 9,2E-05 | 0,0E+00 | 0,0E+00 | 6,1E-10 | 0,0E+00 | 1,1E-08 | 0,0E+00 | 3585426 | 3587057 | 1632 | + |
| Y11_RS16385 | Y11_34661 | tatA | Sec-independent protein translocase subunit TatA | 0,09 | 1,19 | 1,85 | 1,1 | 1,75 | 0,65 | 2,3E-01 | 1,2E-05 | 3,8E-05 | 2,0E-05 | 5,3E-05 | 3,0E-02 | 0,0E+00 | 2,8E-07 | 5,6E-07 | 3,9E-07 | 1,5E-06 | 0,0E+00 | 3587258 | 3587524 | 267 | + |
| Y11_RS16390 | Y11_34671 | tatB | Sec-independent protein translocase protein TatB | 0,09 | 1,24 | 1,49 | 1,15 | 1,39 | 0,24 | 8,3E-02 | 1,1E-08 | 1,3E-06 | 3,6E-08 | 2,4E-06 | 7,5E-02 | 0,0E+00 | 6,7E-10 | 2,8E-08 | 1,4E-09 | 8,9E-08 | 0,0E+00 | 3587528 | 3588184 | 657 | + |
| Y11_RS16395 | Y11_34681 | tatC | Sec-independent protein translocase subunit TatC | 0,15 | 1,33 | 1,65 | 1,18 | 1,5 | 0,33 | 4,1E-01 | 3,9E-05 | 9,8E-05 | 1,3E-04 | 2,3E-04 | 2,0E-01 | 0,0E+00 | 8,7E-07 | 1,3E-06 | 2,3E-06 | 5,6E-06 | 0,0E+00 | 3588187 | 3588966 | 780 | + |
| Y11_RS16405 | Y11_34701 | hemB | porphobilinogen synthase | 0,05 | -0,14 | 0,18 | -0,19 | 0,13 | 0,32 | 2,8E-01 | 5,6E-02 | 1,7E-01 | 1,9E-02 | 3,1E-01 | 3,9E-02 | 0,0E+00 | 0,0E+00 | 0,0E+00 | 0,0E+00 | 0,0E+00 | 0,0E+00 | 3589819 | 3590841 | 1023 | + |
| Y11_RS16410 | Y11_34711 | rfaH | transcription/translation regulatory transformer protein RfaH | 0,5 | 2,21 | 1,53 | 1,71 | 1,02 | -0,68 | 1,8E-01 | 4,7E-05 | 2,6E-04 | 1,5E-03 | 1,6E-02 | 4,0E-02 | 0,0E+00 | 1,0E-06 | 3,4E-06 | 2,5E-05 | 3,5E-04 | 0,0E+00 | 3590930 | 3591418 | 489 | - |
| Y11_RS16415 | Y11_34721 | ubiD | 4-hydroxy-3-polyprenylbenzoate decarboxylase | 0 | 0,21 | 0,47 | 0,2 | 0,47 | 0,27 | 9,8E-01 | 6,5E-02 | 1,2E-03 | 2,9E-02 | 2,3E-04 | 5,9E-03 | 0,0E+00 | 0,0E+00 | 0,0E+00 | 0,0E+00 | 0,0E+00 | 0,0E+00 | 3591683 | 3593179 | 1497 | + |
| Y11_RS16420 | Y11_34731 | fre | NAD(P)H-flavin reductase | 0,07 | 0,64 | 0,75 | 0,57 | 0,68 | 0,11 | 4,1E-01 | 1,3E-06 | 4,1E-05 | 4,2E-05 | 2,0E-04 | 2,5E-01 | 0,0E+00 | 0,0E+00 | 0,0E+00 | 0,0E+00 | 0,0E+00 | 0,0E+00 | 3593234 | 3593935 | 702 | + |
| Y11_RS16425 | Y11_34741 | fadA | acetyl-CoA C-acyltransferase FadA | 0,34 | 1,18 | 3,37 | 0,84 | 3,03 | 2,19 | 1,4E-02 | 4,2E-07 | 1,1E-08 | 8,2E-05 | 7,7E-08 | 3,8E-07 | 0,0E+00 | 1,4E-08 | 5,5E-10 | 0,0E+00 | 5,0E-09 | 8,5E-08 | 3594035 | 3595198 | 1164 | - |
| Y11_RS16430 | Y11_34751 | fadB | fatty acid oxidation complex subunit alpha FadB | 0,56 | 1,17 | 3,43 | 0,61 | 2,87 | 2,26 | 3,2E-05 | 3,2E-08 | 1,1E-10 | 1,1E-05 | 7,4E-10 | 2,3E-09 | 0,0E+00 | 1,6E-09 | 2,5E-11 | 0,0E+00 | 1,4E-10 | 3,0E-09 | 3595210 | 3597399 | 2190 | - |
| Y11_RS16435 | Y11_34761 | pepQ | Xaa-Pro dipeptidase | -0,49 | 1,04 | 1,97 | 1,53 | 2,46 | 0,94 | 3,2E-05 | 2,1E-07 | 8,8E-10 | 2,4E-09 | 3,0E-11 | 1,6E-07 | 0,0E+00 | 7,6E-09 | 9,1E-11 | 1,7E-10 | 2,0E-11 | 0,0E+00 | 3597727 | 3599058 | 1332 | + |
| Y11_RS16450 | Y11_34791 | hemG | menaquinone-dependent protoporphyrinogen IX dehydrogenase | 0,23 | -0,44 | 0,45 | -0,66 | 0,22 | 0,88 | 5,5E-02 | 6,6E-03 | 4,7E-03 | 2,4E-05 | 1,3E-02 | 1,4E-05 | 0,0E+00 | 0,0E+00 | 0,0E+00 | 0,0E+00 | 0,0E+00 | 0,0E+00 | 3601182 | 3601715 | 534 | + |

| | | | | | | | | | | | | | | | | | | | | | | | | | |
|---|---|---|---|---|---|---|---|---|---|---|---|---|---|---|---|---|---|---|---|---|---|---|---|---|---|
| Y11_RS16485 | Y11_34821 | murB | UDP-N-acetylmuramate dehydrogenase | -0,36 | -1,11 | -1,23 | -0,75 | -0,87 | -0,12 | 2,6E-05 | 8,3E-10 | 4,0E-07 | 3,6E-08 | 6,4E-06 | 1,7E-01 | 0,0E+00 | 9,5E-11 | 1,0E-08 | 0,0E+00 | 0,0E+00 | 0,0E+00 | 3609230 | 3610267 | 1038 | + |
| Y11_RS16490 | Y11_34831 | birA | bifunctional biotin--[acetyl-CoA-carboxylase] ligase/biotin operon repressor BirA | -0,53 | 0,37 | 0,57 | 0,9 | 1,11 | 0,21 | 7,0E-04 | 2,3E-05 | 1,8E-04 | 1,0E-05 | 1,6E-05 | 3,1E-02 | 0,0E+00 | 0,0E+00 | 0,0E+00 | 0,0E+00 | 4,9E-07 | 0,0E+00 | 3610264 | 3611223 | 960 | + |
| Y11_RS16495 | Y11_34841 | coaA | type I pantothenate kinase | 0,5 | 0,52 | 1,13 | 0,02 | 0,63 | 0,61 | 1,3E-03 | 1,1E-04 | 1,1E-05 | 8,1E-01 | 1,0E-03 | 3,5E-04 | 0,0E+00 | 0,0E+00 | 1,9E-07 | 0,0E+00 | 0,0E+00 | 0,0E+00 | 3611263 | 3612213 | 951 | - |
| Y11_RS16525 | Y11_34871 | tuf | elongation factor Tu | 0,11 | -0,23 | -0,07 | -0,34 | -0,19 | 0,16 | 1,1E-01 | 5,3E-03 | 6,3E-01 | 2,3E-03 | 2,6E-01 | 3,3E-01 | 0,0E+00 | 0,0E+00 | 0,0E+00 | 0,0E+00 | 0,0E+00 | 0,0E+00 | 3614139 | 3615323 | 1185 | + |
| Y11_RS16530 | Y11_34881 | secE | preprotein translocase subunit SecE | -0,71 | -0,96 | -0,03 | -0,25 | 0,68 | 0,93 | 1,7E-03 | 3,6E-03 | 9,7E-01 | 3,5E-01 | 3,1E-01 | 1,9E-01 | 0,0E+00 | 0,0E+00 | 0,0E+00 | 0,0E+00 | 0,0E+00 | 0,0E+00 | 3615563 | 3615946 | 384 | + |
| Y11_RS16535 | Y11_34891 | nusG | transcription termination/antitermination protein NusG | 0,04 | 0,03 | 0,28 | -0,01 | 0,23 | 0,24 | 6,7E-01 | 6,3E-01 | 1,2E-01 | 8,9E-01 | 2,0E-01 | 1,4E-01 | 0,0E+00 | 0,0E+00 | 0,0E+00 | 0,0E+00 | 0,0E+00 | 0,0E+00 | 3615948 | 3616493 | 546 | + |
| Y11_RS16540 | Y11_34901 | rplK | 50S ribosomal protein L11 | 0,14 | -0,77 | -1,51 | -0,9 | -1,65 | -0,74 | 2,2E-01 | 4,0E-04 | 1,3E-06 | 8,2E-05 | 3,4E-07 | 5,9E-04 | 0,0E+00 | 0,0E+00 | 2,8E-08 | 0,0E+00 | 1,7E-08 | 0,0E+00 | 3616687 | 3617115 | 429 | + |
| Y11_RS16545 | Y11_34911 | rplA | 50S ribosomal protein L1 | 0,22 | -0,71 | -1,05 | -0,94 | -1,27 | -0,34 | 1,4E-02 | 1,2E-03 | 8,5E-07 | 2,1E-04 | 1,8E-07 | 5,4E-02 | 0,0E+00 | 0,0E+00 | 2,0E-08 | 0,0E+00 | 1,0E-08 | 0,0E+00 | 3617119 | 3617823 | 705 | + |
| Y11_RS16550 | Y11_34921 | rplJ | 50S ribosomal protein L10 | 0,12 | -0,4 | -0,93 | -0,51 | -1,04 | -0,53 | 1,7E-01 | 2,1E-03 | 3,2E-04 | 2,2E-03 | 3,0E-04 | 1,8E-02 | 0,0E+00 | 0,0E+00 | 0,0E+00 | 0,0E+00 | 7,2E-06 | 0,0E+00 | 3618196 | 3618699 | 504 | + |
| Y11_RS16555 | Y11_34931 | rplL | 50S ribosomal protein L7/L12 | -0,1 | -0,32 | -0,75 | -0,22 | -0,65 | -0,43 | 4,4E-01 | 4,2E-02 | 7,4E-04 | 2,1E-01 | 5,0E-03 | 4,1E-02 | 0,0E+00 | 0,0E+00 | 0,0E+00 | 0,0E+00 | 0,0E+00 | 0,0E+00 | 3618768 | 3619136 | 369 | + |
| Y11_RS16560 | Y11_34941 | rpoB | DNA-directed RNA polymerase subunit beta | -0,15 | -0,28 | -0,84 | -0,13 | -0,69 | -0,56 | 8,6E-03 | 6,5E-04 | 2,7E-07 | 1,1E-02 | 1,6E-07 | 3,1E-06 | 0,0E+00 | 0,0E+00 | 0,0E+00 | 0,0E+00 | 0,0E+00 | 0,0E+00 | 3619480 | 3623508 | 4029 | + |
| Y11_RS16565 | Y11_34951 | rpoC | DNA-directed RNA polymerase subunit beta' | -0,1 | -0,06 | -0,41 | 0,05 | -0,3 | -0,35 | 3,7E-02 | 2,4E-01 | 4,6E-08 | 4,6E-01 | 1,3E-04 | 8,9E-05 | 0,0E+00 | 0,0E+00 | 0,0E+00 | 0,0E+00 | 0,0E+00 | 0,0E+00 | 3623645 | 3627865 | 4221 | + |
| Y11_RS16605 | Y11_35031 | | Rsd/AlgQ family anti-sigma factor | 2,05 | 1,9 | 1,86 | -0,16 | -0,2 | -0,04 | 1,4E-07 | 1,9E-07 | 1,7E-06 | 2,5E-01 | 2,5E-01 | 8,0E-01 | 1,1E-08 | 7,1E-09 | 3,6E-08 | 0,0E+00 | 0,0E+00 | 0,0E+00 | 3635463 | 3635966 | 504 | - |
| Y11_RS16610 | Y11_35041 | nudC | NAD(+) diphosphatase | 0,12 | 1,07 | 0,63 | 0,95 | 0,51 | -0,44 | 8,7E-02 | 5,0E-07 | 5,9E-04 | 3,9E-06 | 3,2E-03 | 9,4E-03 | 0,0E+00 | 1,6E-08 | 0,0E+00 | 0,0E+00 | 0,0E+00 | 0,0E+00 | 3636058 | 3636843 | 786 | + |
| Y11_RS16615 | Y11_35051 | hemE | uroporphyrinogen decarboxylase | -0,17 | 1,04 | 1,28 | 1,21 | 1,46 | 0,25 | 6,8E-02 | 7,3E-07 | 2,2E-07 | 1,3E-07 | 4,9E-08 | 6,9E-03 | 0,0E+00 | 2,2E-08 | 6,2E-09 | 4,2E-09 | 3,5E-09 | 0,0E+00 | 3636924 | 3637991 | 1068 | + |
| Y11_RS16620 | Y11_35061 | nfi | deoxyribonuclease V | 0 | 0,87 | 0,08 | 0,87 | 0,07 | -0,79 | 9,9E-01 | 5,8E-04 | 7,7E-01 | 9,8E-07 | 7,3E-01 | 4,0E-03 | 0,0E+00 | 0,0E+00 | 0,0E+00 | 0,0E+00 | 0,0E+00 | 0,0E+00 | 3638015 | 3638719 | 705 | + |
| Y11_RS16625 | Y11_35071 | | YjaG family protein | 0,13 | 0,33 | 0,16 | 0,19 | 0,03 | -0,16 | 1,9E-01 | 8,0E-03 | 1,1E-01 | 1,3E-01 | 7,9E-01 | 2,0E-01 | 0,0E+00 | 0,0E+00 | 0,0E+00 | 0,0E+00 | 0,0E+00 | 0,0E+00 | 3638765 | 3639355 | 591 | + |
| Y11_RS16630 | Y11_35081 | hupA | DNA-binding protein HU-alpha | 0,22 | 2,03 | 2,45 | 1,81 | 2,23 | 0,42 | 2,3E-02 | 1,1E-08 | 5,3E-04 | 1,7E-07 | 1,0E-03 | 3,7E-01 | 0,0E+00 | 6,5E-10 | 6,5E-06 | 5,3E-09 | 2,4E-05 | 0,0E+00 | 3639544 | 3639819 | 276 | + |
| Y11_RS16635 | Y11_35091 | | DUF1481 domain-containing protein | 0,58 | 2,57 | 2,35 | 2 | 1,77 | -0,23 | 4,6E-02 | 1,9E-07 | 2,5E-10 | 1,1E-04 | 9,1E-05 | 1,8E-01 | 0,0E+00 | 7,0E-09 | 4,1E-11 | 2,0E-06 | 2,4E-06 | 0,0E+00 | 3639872 | 3640525 | 654 | + |
| Y11_RS16640 | Y11_35101 | purD | phosphoribosylamine--glycine ligase | 1,11 | 2,91 | 3,28 | 1,79 | 2,17 | 0,38 | 8,7E-09 | 2,0E-12 | 1,1E-11 | 1,8E-10 | 4,3E-10 | 1,8E-04 | 1,2E-09 | 2,5E-12 | 7,7E-12 | 2,6E-11 | 1,0E-10 | 0,0E+00 | 3640603 | 3641898 | 1296 | - |
| Y11_RS16645 | Y11_35111 | purH | bifunctional phosphoribosylaminoimidazolecarboxamide formyltransferase/IMP cyclohydrolase | 1,29 | 2,59 | 2,8 | 1,29 | 1,51 | 0,22 | 6,2E-09 | 1,6E-11 | 2,8E-10 | 3,9E-09 | 3,7E-08 | 1,8E-02 | 1,0E-09 | 7,2E-12 | 4,2E-11 | 2,3E-10 | 2,7E-09 | 0,0E+00 | 3641924 | 3643513 | 1590 | - |
| Y11_RS16685 | Y11_35131 | actP | cation/acetate symporter ActP | -0,51 | 15 | 15 | 15 | 15 | 0 | 3,3E-01 | 0,0E+00 | 0,0E+00 | 0,0E+00 | 0,0E+00 | 0,0E+00 | 0,0E+00 | 0,0E+00 | 0,0E+00 | 0,0E+00 | 0,0E+00 | 0,0E+00 | 3650311 | 3651966 | 1656 | - |
| Y11_RS16690 | Y11_35141 | | DUF485 domain-containing protein | 0,22 | 2,05 | 1,99 | 1,83 | 1,78 | -0,05 | 5,9E-01 | 2,9E-04 | 3,1E-04 | 5,7E-04 | 6,2E-04 | 8,6E-01 | 0,0E+00 | 6,1E-06 | 3,9E-06 | 9,7E-06 | 1,4E-05 | 0,0E+00 | 3651963 | 3652274 | 312 | - |
| Y11_RS16695 | Y11_35151 | acs | acetate--CoA ligase | -0,29 | 3,98 | 15 | 4,27 | 15 | 15 | 7,6E-04 | 4,4E-10 | 0,0E+00 | 1,2E-10 | 0,0E+00 | 0,0E+00 | 0,0E+00 | 5,9E-11 | 0,0E+00 | 2,0E-11 | 0,0E+00 | 0,0E+00 | 3652328 | 3654286 | 1959 | - |
| Y11_RS16700 | Y11_35161 | gltP | glutamate/aspartate:proton symporter GltP | -0,43 | -0,8 | -1,06 | -0,37 | -0,64 | -0,26 | 4,6E-02 | 7,0E-04 | 1,8E-03 | 3,4E-02 | 2,4E-02 | 2,4E-01 | 0,0E+00 | 0,0E+00 | 2,1E-05 | 0,0E+00 | 0,0E+00 | 0,0E+00 | 3655205 | 3656518 | 1314 | + |
| Y11_RS16905 | | yefM | YoeB-YefM toxin-antitoxin system antitoxin YefM | 0,02 | 0,15 | 0,01 | 0,13 | -0,01 | -0,14 | 9,1E-01 | 1,7E-01 | 9,1E-01 | 4,6E-01 | 9,7E-01 | 2,8E-01 | 0,0E+00 | 0,0E+00 | 0,0E+00 | 0,0E+00 | 0,0E+00 | 0,0E+00 | 3692037 | 3692288 | 252 | + |
| Y11_RS16915 | Y11_35631 | | DUF3772 domain-containing protein | 0,22 | -0,08 | 0,26 | -0,3 | 0,04 | 0,34 | 5,6E-01 | 5,1E-01 | 4,4E-01 | 4,3E-01 | 9,3E-01 | 3,2E-01 | 0,0E+00 | 0,0E+00 | 0,0E+00 | 0,0E+00 | 0,0E+00 | 0,0E+00 | 3692563 | 3694899 | 2337 | - |
| Y11_RS16920 | Y11_35641 | | TonB-dependent siderophore receptor | 0,49 | -2,35 | -3,5 | -2,84 | -3,99 | -1,15 | 8,1E-06 | 1,5E-11 | 1,2E-09 | 1,4E-11 | 4,9E-10 | 7,5E-06 | 0,0E+00 | 7,2E-12 | 1,1E-10 | 6,1E-12 | 1,1E-10 | 9,0E-07 | 3695318 | 3697450 | 2133 | + |
| Y11_RS16925 | Y11_35661 | | PLP-dependent aminotransferase family protein | -0,14 | -0,31 | 0,35 | -0,17 | 0,49 | 0,66 | 2,6E-01 | 8,3E-02 | 2,6E-02 | 2,7E-01 | 2,0E-03 | 2,4E-03 | 0,0E+00 | 0,0E+00 | 0,0E+00 | 0,0E+00 | 0,0E+00 | 0,0E+00 | 3697930 | 3699381 | 1452 | + |
| Y11_RS16940 | Y11_35691 | | hemin ABC transporter substrate-binding protein | 0,88 | -1,2 | -2,82 | -2,08 | -3,7 | -1,62 | 7,8E-05 | 8,4E-06 | 8,8E-09 | 7,4E-08 | 5,9E-10 | 3,9E-07 | 0,0E+00 | 2,0E-07 | 4,8E-10 | 2,6E-09 | 1,2E-10 | 8,7E-08 | 3701340 | 3702179 | 840 | - |
| Y11_RS16945 | Y11_35701 | | hemin-degrading factor | 0,29 | -2,71 | -4,85 | -2,99 | -5,13 | -2,14 | 4,2E-03 | 3,5E-07 | 1,4E-10 | 2,1E-07 | 1,7E-10 | 6,6E-06 | 0,0E+00 | 1,2E-08 | 2,7E-11 | 6,4E-09 | 6,2E-11 | 8,1E-07 | 3702176 | 3703213 | 1038 | - |
| Y11_RS16950 | Y11_35711 | | TonB-dependent hemoglobin/transferrin/lactoferrin family receptor | 1,85 | -1,42 | -2,95 | -3,27 | -4,79 | -1,52 | 1,0E-07 | 1,7E-06 | 1,9E-07 | 1,9E-11 | 1,1E-09 | 1,2E-05 | 9,4E-09 | 4,8E-08 | 5,7E-09 | 6,6E-12 | 1,8E-10 | 1,4E-06 | 3703330 | 3705393 | 2064 | - |
| Y11_RS16965 | Y11_35741 | hutX | heme utilization cystosolic carrier protein HutX | 0 | -0,18 | -0,74 | -0,12 | -0,77 | -0,6 | 0,0E+00 | 1,3E-01 | 3,8E-03 | 3,3E-01 | 3,3E-03 | 5,5E-03 | 0,0E+00 | 0,0E+00 | 0,0E+00 | 0,0E+00 | 0,0E+00 | 0,0E+00 | 3706490 | 3707035 | 546 | - |
| Y11_RS16975 | Y11_35761 | tdcB | bifunctional threonine ammonia-lyase/L-serine ammonia-lyase TdcB | -1,94 | 0,97 | 1,61 | 2,9 | 3,55 | 0,65 | 6,4E-06 | 7,0E-04 | 3,2E-05 | 5,0E-09 | 3,6E-09 | 7,4E-04 | 3,4E-07 | 0,0E+00 | 4,9E-07 | 2,9E-10 | 4,3E-10 | 0,0E+00 | 3708983 | 3709975 | 993 | + |
| Y11_RS16980 | Y11_35771 | tdcC | threonine/serine transporter TdcC | -2,05 | 1 | 1,18 | 3,05 | 3,28 | 0,28 | 1,4E-05 | 1,3E-03 | 1,1E-03 | 4,9E-10 | 1,1E-09 | 2,1E-02 | 7,0E-07 | 2,6E-05 | 1,3E-05 | 5,2E-11 | 1,8E-10 | 0,0E+00 | 3710000 | 3711328 | 1329 | + |
| Y11_RS17000 | Y11_35801 | | transcriptional regulator | -0,04 | 0,93 | 1,69 | 0,97 | 1,73 | 0,75 | 6,9E-01 | 5,0E-06 | 1,1E-06 | 1,9E-06 | 7,1E-07 | 1,7E-04 | 0,0E+00 | 0,0E+00 | 2,6E-08 | 0,0E+00 | 3,1E-08 | 0,0E+00 | 3715273 | 3715848 | 576 | - |
| Y11_RS17015 | Y11_35831 | | protein-disulfide reductase DsbD | -0,89 | 0,41 | 0,44 | 1,3 | 1,32 | 0,03 | 1,5E-04 | 7,8E-03 | 8,4E-03 | 1,8E-05 | 2,3E-05 | 8,5E-01 | 0,0E+00 | 0,0E+00 | 0,0E+00 | 3,6E-07 | 6,8E-07 | 0,0E+00 | 3717399 | 3719162 | 1764 | - |
| Y11_RS17020 | Y11_35841 | cutA | divalent cation tolerance protein CutA | -0,53 | 0,47 | 0,78 | 1 | 1,31 | 0,31 | 1,0E-01 | 8,3E-02 | 2,9E-02 | 4,0E-04 | 6,9E-04 | 1,3E-01 | 0,0E+00 | 0,0E+00 | 0,0E+00 | 6,9E-06 | 1,6E-05 | 0,0E+00 | 3719138 | 3719476 | 339 | - |
| Y11_RS17025 | Y11_35851 | | anaerobic C4-dicarboxylate transporter | -1,44 | 2,31 | 2,03 | 3,74 | 3,47 | 0 | 1,0E-02 | 4,2E-04 | 7,6E-04 | 1,6E-07 | 1,1E-07 | 0,0E+00 | 3,6E-04 | 8,6E-06 | 9,2E-06 | 5,1E-09 | 6,7E-09 | 0,0E+00 | 3719633 | 3720934 | 1302 | - |
| Y11_RS17030 | Y11_35861 | aspA | aspartate ammonia-lyase | -1,29 | 1,27 | 3,24 | 2,57 | 4,54 | 1,97 | 9,6E-09 | 6,0E-09 | 4,1E-12 | 3,4E-10 | 3,9E-12 | 2,1E-09 | 1,3E-09 | 4,3E-10 | 5,4E-12 | 3,9E-11 | 7,5E-12 | 3,0E-09 | 3721054 | 3722490 | 1437 | - |
| Y11_RS17035 | Y11_35871 | fxsA | membrane protein FxsA | -0,3 | 0,81 | 0,93 | 1,11 | 1,23 | 0,12 | 4,1E-01 | 1,5E-02 | 5,8E-03 | 5,7E-03 | 2,6E-03 | 5,2E-01 | 0,0E+00 | 0,0E+00 | 0,0E+00 | 9,2E-05 | 5,7E-05 | 0,0E+00 | 3722932 | 3723459 | 528 | + |
| Y11_RS17040 | Y11_35881 | | co-chaperone GroES | -0,19 | 0,49 | 1,03 | 0,68 | 1,23 | 0,55 | 6,5E-03 | 1,5E-05 | 4,3E-06 | 2,4E-07 | 7,7E-07 | 2,9E-04 | 0,0E+00 | 0,0E+00 | 7,8E-08 | 0,0E+00 | 3,3E-08 | 0,0E+00 | 3723713 | 3724006 | 294 | + |
| Y11_RS17045 | Y11_35891 | groL | chaperonin GroEL | -0,22 | 0,05 | 0,14 | 0,26 | 0,35 | 0,09 | 2,0E-01 | 8,1E-01 | 3,8E-01 | 2,0E-01 | 4,0E-02 | 6,3E-01 | 0,0E+00 | 0,0E+00 | 0,0E+00 | 0,0E+00 | 0,0E+00 | 0,0E+00 | 3724063 | 3725715 | 1653 | + |
| Y11_RS17055 | Y11_35901 | | DUF4156 domain-containing protein | -0,16 | 0,42 | 0,46 | 0,57 | 0,61 | 0,04 | 3,7E-01 | 3,1E-02 | 2,1E-02 | 7,8E-03 | 5,4E-03 | 8,1E-01 | 0,0E+00 | 0,0E+00 | 0,0E+00 | 0,0E+00 | 0,0E+00 | 0,0E+00 | 3726100 | 3726438 | 339 | + |
| Y11_RS17060 | Y11_35911 | epmB | EF-P beta-lysylation protein EpmB | 0 | -0,49 | -0,88 | -0,5 | -0,89 | -0,39 | 9,7E-01 | 2,6E-04 | 1,5E-05 | 4,5E-03 | 2,0E-04 | 8,3E-03 | 0,0E+00 | 0,0E+00 | 0,0E+00 | 0,0E+00 | 0,0E+00 | 0,0E+00 | 3726495 | 3727526 | 1032 | - |
| Y11_RS17065 | Y11_35921 | efp | elongation factor P | -0,71 | -0,11 | -0,26 | 0,6 | 0,45 | -0,16 | 7,2E-05 | 3,9E-01 | 3,8E-01 | 3,5E-03 | 1,7E-01 | 6,2E-01 | 0,0E+00 | 0,0E+00 | 0,0E+00 | 0,0E+00 | 0,0E+00 | 0,0E+00 | 3727562 | 3728128 | 567 | + |
| Y11_RS17070 | Y11_35931 | | entericidin A/B family lipoprotein | 0,47 | 3,7 | 4,18 | 3,23 | 3,71 | 0,48 | 2,4E-01 | 1,3E-06 | 5,2E-06 | 2,8E-06 | 1,1E-05 | 1,6E-01 | 0,0E+00 | 3,9E-08 | 9,3E-08 | 6,2E-08 | 3,5E-07 | 0,0E+00 | 3728314 | 3728445 | 132 | + |
| Y11_RS17080 | Y11_35951 | blc | outer membrane lipoprotein Blc | 0,37 | 0,57 | 0,19 | 0,2 | -0,19 | -0,38 | 5,7E-05 | 2,1E-06 | 1,0E-01 | 8,0E-05 | 8,2E-02 | 3,4E-03 | 0,0E+00 | 0,0E+00 | 0,0E+00 | 0,0E+00 | 0,0E+00 | 0,0E+00 | 3728940 | 3729479 | 540 | - |
| Y11_RS17095 | Y11_35981 | | succinate dehydrogenase/fumarate reductase iron-sulfur subunit | -0,82 | 0,63 | 1,46 | 1,45 | 2,28 | 0,83 | 2,1E-03 | 1,3E-03 | 3,4E-06 | 2,4E-05 | 7,9E-07 | 4,6E-05 | 0,0E+00 | 0,0E+00 | 6,5E-08 | 4,7E-07 | 3,4E-08 | 0,0E+00 | 3730411 | 3731145 | 735 | - |
| Y11_RS17100 | Y11_35991 | frdA | fumarate reductase (quinol) flavoprotein subunit | -0,88 | 0,37 | 1,3 | 1,25 | 2,17 | 0,93 | 8,9E-08 | 4,1E-04 | 1,9E-06 | 4,4E-08 | 3,2E-08 | 3,8E-05 | 0,0E+00 | 0,0E+00 | 3,9E-08 | 1,7E-09 | 2,5E-09 | 0,0E+00 | 3731138 | 3732961 | 1824 | - |
| Y11_RS17105 | Y11_36001 | epmA | elongation factor P--(R)-beta-lysine ligase | 0,04 | -0,74 | -0,48 | -0,78 | -0,52 | 0,26 | 5,0E-01 | 1,2E-05 | 4,1E-04 | 1,5E-05 | 3,8E-04 | 3,6E-02 | 0,0E+00 | 0,0E+00 | 0,0E+00 | 0,0E+00 | 0,0E+00 | 0,0E+00 | 3733539 | 3734516 | 978 | + |
| Y11_RS17110 | Y11_36011 | mscM | miniconductance mechanosensitive channel MscM | -0,24 | 2,59 | 2,75 | 2,83 | 2,99 | 0,16 | 1,2E-02 | 7,6E-11 | 7,2E-08 | 1,3E-10 | 4,7E-08 | 2,9E-01 | 0,0E+00 | 1,9E-11 | 2,5E-09 | 2,1E-11 | 3,3E-09 | 0,0E+00 | 3734658 | 3738017 | 3360 | - |
| Y11_RS17115 | Y11_36021 | asd | archaetidylserine decarboxylase | 0,03 | 0,88 | 0,62 | 0,85 | 0,58 | -0,26 | 6,9E-01 | 8,6E-07 | 2,1E-04 | 7,3E-07 | 2,6E-04 | 1,1E-02 | 0,0E+00 | 0,0E+00 | 0,0E+00 | 0,0E+00 | 0,0E+00 | 0,0E+00 | 3738058 | 3738939 | 882 | - |
| Y11_RS17120 | Y11_36031 | rsgA | small ribosomal subunit biogenesis GTPase RsgA | -0,11 | 0,73 | 0,87 | 0,84 | 0,99 | 0,14 | 4,8E-02 | 1,3E-04 | 9,2E-05 | 3,5E-05 | 3,1E-05 | 3,7E-01 | 0,0E+00 | 0,0E+00 | 0,0E+00 | 0,0E+00 | 0,0E+00 | 0,0E+00 | 3739048 | 3740100 | 1053 | - |
| Y11_RS17125 | Y11_36041 | orn | oligoribonuclease | -0,34 | 0,79 | 1,29 | 1,13 | 1,63 | 0,5 | 4,3E-03 | 1,4E-06 | 1,5E-08 | 2,5E-06 | 1,1E-07 | 1,0E-04 | 0,0E+00 | 0,0E+00 | 7,2E-10 | 5,7E-08 | 6,7E-09 | 0,0E+00 | 3740278 | 3740823 | 546 | + |
| Y11_RS17140 | Y11_36051 | queG | tRNA epoxyqueuosine(34) reductase QueG | 0 | 0 | -1,12 | 0 | -0,47 | -0,75 | 0,0E+00 | 0,0E+00 | 1,3E-02 | 0,0E+00 | 1,7E-01 | 4,4E-02 | 0,0E+00 | 0,0E+00 | 1,5E-04 | 0,0E+00 | 0,0E+00 | 0,0E+00 | 3741688 | 3742842 | 1155 | - |
| Y11_RS17145 | Y11_36061 | nnr | bifunctional ADP-dependent NAD(P)H-hydrate dehydratase/NAD(P)H-hydrate epimerase | 0,08 | -0,76 | -0,01 | -0,83 | -0,09 | 0,74 | 6,8E-02 | 1,8E-07 | 8,1E-01 | 3,6E-07 | 2,0E-01 | 5,4E-06 | 0,0E+00 | 0,0E+00 | 0,0E+00 | 0,0E+00 | 0,0E+00 | 0,0E+00 | 3742841 | 3744355 | 1515 | + |
| Y11_RS17150 | Y11_36071 | tsaE | tRNA (adenosine(37)-N6)-threonylcarbamoyltransferase complex ATPase subunit type 1 TsaE | -0,31 | -0,45 | -0,24 | -0,14 | 0,07 | 0,21 | 3,7E-03 | 2,7E-03 | 3,8E-01 | 2,2E-01 | 8,1E-01 | 4,7E-01 | 0,0E+00 | 0,0E+00 | 0,0E+00 | 0,0E+00 | 0,0E+00 | 0,0E+00 | 3744366 | 3744836 | 471 | + |
| Y11_RS17155 | Y11_36081 | amiB | N-acetylmuramoyl-L-alanine amidase AmiB | 0,43 | -1,72 | -2,1 | -2,16 | -2,53 | -0,37 | 8,4E-04 | 1,9E-10 | 5,8E-07 | 3,3E-09 | 2,7E-07 | 3,3E-02 | 0,0E+00 | 3,5E-11 | 1,4E-08 | 2,1E-10 | 1,4E-08 | 0,0E+00 | 3744916 | 3746781 | 1866 | + |
| Y11_RS17160 | Y11_36091 | mutL | DNA mismatch repair endonuclease MutL | 0,21 | 0,07 | 0,46 | -0,14 | 0,24 | 0,38 | 2,0E-03 | 2,7E-01 | 4,0E-03 | 7,3E-02 | 7,4E-02 | 1,5E-02 | 0,0E+00 | 0,0E+00 | 0,0E+00 | 0,0E+00 | 0,0E+00 | 0,0E+00 | 3746797 | 3748704 | 1908 | + |
| Y11_RS17170 | Y11_36121 | hfq | RNA chaperone Hfq | 0,03 | 0,25 | 0,47 | 0,22 | 0,44 | 0,23 | 7,9E-01 | 8,5E-02 | 8,5E-02 | 2,9E-02 | 8,1E-02 | 3,5E-01 | 0,0E+00 | 0,0E+00 | 0,0E+00 | 0,0E+00 | 0,0E+00 | 0,0E+00 | 3749754 | 3750059 | 306 | + |
| Y11_RS17175 | Y11_36131 | hflX | GTPase HflX | 0,21 | 0,34 | -0,21 | 0,13 | -0,41 | -0,55 | 4,5E-02 | 3,5E-03 | 5,9E-02 | 3,9E-02 | 3,8E-04 | 3,4E-05 | 0,0E+00 | 0,0E+00 | 0,0E+00 | 0,0E+00 | 0,0E+00 | 0,0E+00 | 3750158 | 3751459 | 1302 | + |
| Y11_RS17180 | Y11_36141 | hflK | FtsH protease activity modulator HflK | 0,12 | -1,35 | -1,32 | -1,47 | -1,44 | 0,03 | 3,4E-02 | 3,5E-08 | 1,2E-06 | 3,3E-09 | 3,4E-07 | 7,9E-01 | 0,0E+00 | 1,7E-09 | 2,7E-08 | 2,1E-10 | 1,7E-08 | 0,0E+00 | 3751557 | 3752840 | 1284 | + |
| Y11_RS17185 | Y11_36151 | hflC | protease modulator HflC | 0,13 | -1,06 | -0,93 | -1,19 | -1,06 | 0,13 | 2,1E-02 | 4,7E-07 | 5,9E-05 | 3,3E-07 | 2,7E-05 | 3,6E-01 | 0,0E+00 | 1,5E-08 | 0,0E+00 | 9,1E-09 | 7,9E-07 | 0,0E+00 | 3752844 | 3753848 | 1005 | + |
| Y11_RS17195 | Y11_36171 | | adenylosuccinate synthase | -0,04 | -0,97 | -1 | -0,93 | -0,96 | -0,03 | 5,4E-01 | 1,5E-08 | 1,3E-07 | 2,1E-07 | 7,0E-07 | 5,4E-01 | 0,0E+00 | 0,0E+00 | 3,9E-09 | 0,0E+00 | 0,0E+00 | 0,0E+00 | 3754248 | 3755546 | 1299 | + |
| Y11_RS17200 | Y11_36181 | nsrR | nitric oxide-sensing transcriptional repressor NsrR | 1,91 | 5,54 | 5,02 | 3,63 | 3,11 | -0,53 | 8,9E-02 | 1,0E-03 | 8,2E-06 | 3,4E-02 | 1,6E-02 | 6,5E-01 | 2,9E-03 | 2,1E-05 | 1,4E-07 | 5,3E-04 | 3,3E-04 | 0,0E+00 | 3755972 | 3756397 | 426 | + |
| Y11_RS17205 | Y11_36191 | rnr | ribonuclease R | 0,16 | 1,87 | 2 | 1,71 | 1,84 | 0,14 | 4,1E-04 | 4,4E-12 | 1,5E-10 | 1,2E-11 | 3,2E-10 | 2,8E-02 | 0,0E+00 | 4,0E-12 | 2,8E-11 | 5,6E-12 | 9,0E-11 | 0,0E+00 | 3756451 | 3758997 | 2547 | + |

| | | | | | | | | | | | | | | | | | | | | | | | | | |
|---|---|---|---|---|---|---|---|---|---|---|---|---|---|---|---|---|---|---|---|---|---|---|---|---|---|
| Y11_RS17210 | Y11_36201 | rlmB | 23S rRNA (guanosine(2251)-2'-O)-methyltransferase RlmB | 0,09 | 0,13 | -0,4 | 0,04 | -0,48 | -0,52 | 1,9E-01 | 5,2E-02 | 1,7E-02 | 6,0E-01 | 9,6E-03 | 6,0E-03 | 0,0E+00 | 0,0E+00 | 0,0E+00 | 0,0E+00 | 0,0E+00 | 0,0E+00 | 3759195 | 3759935 | 741 | + |
| Y11_RS17225 | Y11_36241 | bsmA | biofilm peroxide resistance protein BsmA | 0,65 | 3,15 | 3,91 | 2,5 | 3,33 | 0 | 7,3E-03 | 3,8E-05 | 1,7E-08 | 3,0E-04 | 2,3E-07 | 0,0E+00 | 0,0E+00 | 8,7E-07 | 8,0E-10 | 5,4E-06 | 1,2E-08 | 0,0E+00 | 3761929 | 3762237 | 309 | - |
| Y11_RS17230 | Y11_36251 | yjfP | esterase | -0,55 | 1,28 | 1,71 | 1,83 | 2,26 | 0,44 | 4,7E-03 | 2,1E-05 | 2,9E-06 | 3,9E-08 | 1,3E-08 | 2,6E-03 | 0,0E+00 | 4,8E-07 | 5,7E-08 | 1,5E-09 | 1,2E-09 | 0,0E+00 | 3762423 | 3763172 | 750 | + |
| Y11_RS17240 | Y11_36271 | rpsF | 30S ribosomal protein S6 | 0,06 | -0,16 | -0,78 | -0,22 | -0,85 | -0,62 | 4,3E-01 | 4,9E-02 | 6,0E-04 | 1,0E-02 | 3,5E-04 | 2,2E-03 | 0,0E+00 | 0,0E+00 | 0,0E+00 | 0,0E+00 | 0,0E+00 | 0,0E+00 | 3764103 | 3764495 | 393 | + |
| Y11_RS17250 | Y11_36291 | rpsR | 30S ribosomal protein S18 | 0,09 | -0,44 | -1,13 | -0,53 | -1,22 | -0,69 | 1,7E-01 | 3,4E-03 | 1,4E-04 | 1,3E-03 | 8,6E-05 | 6,8E-03 | 0,0E+00 | 0,0E+00 | 1,9E-06 | 0,0E+00 | 2,3E-06 | 0,0E+00 | 3764826 | 3765053 | 228 | + |
| Y11_RS17255 | Y11_36301 | rplI | 50S ribosomal protein L9 | 0,25 | -0,43 | -0,97 | -0,67 | -1,22 | -0,54 | 6,6E-03 | 9,1E-04 | 4,4E-05 | 1,1E-04 | 1,4E-05 | 4,6E-03 | 0,0E+00 | 0,0E+00 | 0,0E+00 | 0,0E+00 | 4,4E-07 | 0,0E+00 | 3765093 | 3765545 | 453 | + |
| Y11_RS17265 | Y11_36321 | | OapA family protein | 0,55 | 1,6 | 1,9 | 1,04 | 1,35 | 0,31 | 9,8E-04 | 2,4E-07 | 1,4E-08 | 1,7E-05 | 8,9E-07 | 9,2E-03 | 0,0E+00 | 8,5E-09 | 7,0E-10 | 3,4E-07 | 3,8E-08 | 0,0E+00 | 3766215 | 3766976 | 762 | - |
| Y11_RS17270 | Y11_36331 | | peptidylprolyl isomerase | -0,45 | 0,66 | 0,7 | 1,1 | 1,15 | 0,04 | 5,8E-05 | 7,2E-06 | 4,3E-02 | 6,7E-08 | 4,3E-03 | 8,8E-01 | 0,0E+00 | 0,0E+00 | 0,0E+00 | 2,4E-09 | 9,4E-05 | 0,0E+00 | 3767263 | 3767883 | 621 | + |
| Y11_RS17275 | Y11_36341 | ytfE | iron-sulfur cluster repair protein YtfE | 0 | -15 | -15 | -0,62 | 0,13 | 0,75 | 0,0E+00 | 0,0E+00 | 0,0E+00 | 2,1E-01 | 6,0E-01 | 1,8E-01 | 0,0E+00 | 0,0E+00 | 0,0E+00 | 0,0E+00 | 0,0E+00 | 0,0E+00 | 3767968 | 3768633 | 666 | - |
| Y11_RS17280 | Y11_36351 | | bifunctional 2',3'-cyclic-nucleotide 2'-phosphodiesterase/3'-nucleotidase | -0,29 | -0,08 | 1,39 | 0,22 | 1,69 | 1,47 | 1,3E-02 | 4,3E-01 | 7,9E-07 | 7,5E-02 | 4,1E-07 | 1,2E-06 | 0,0E+00 | 0,0E+00 | 1,9E-08 | 0,0E+00 | 2,0E-08 | 2,1E-07 | 3768761 | 3770734 | 1974 | - |
| Y11_RS17285 | Y11_36361 | cysQ | 3'(2'),5'-bisphosphate nucleotidase CysQ | 0,56 | 2,48 | 2,47 | 1,92 | 1,91 | -0,01 | 1,9E-06 | 1,4E-09 | 1,6E-10 | 1,1E-08 | 1,2E-09 | 9,5E-01 | 0,0E+00 | 1,4E-10 | 2,9E-11 | 5,5E-10 | 2,0E-10 | 0,0E+00 | 3771213 | 3771953 | 741 | + |
| Y11_RS17290 | Y11_36371 | | YtfJ family protein | 0,39 | 3,63 | 3,86 | 3,25 | 3,48 | 0,23 | 1,1E-02 | 5,9E-10 | 1,9E-08 | 4,3E-10 | 2,8E-08 | 1,8E-01 | 0,0E+00 | 7,4E-11 | 8,4E-10 | 4,7E-11 | 2,2E-09 | 0,0E+00 | 3771956 | 3772519 | 564 | - |
| Y11_RS17295 | Y11_36381 | | DUF1107 domain-containing protein | 0 | -2,68 | 0 | -2,39 | 0 | 15 | 0,0E+00 | 6,7E-05 | 0,0E+00 | 1,5E-04 | 0,0E+00 | 0,0E+00 | 0,0E+00 | 1,5E-06 | 0,0E+00 | 2,6E-06 | 0,0E+00 | 0,0E+00 | 3772843 | 3773046 | 204 | + |
| Y11_RS17300 | Y11_36391 | | hemolysin family protein | -0,12 | 0,51 | 0,22 | 0,63 | 0,34 | -0,28 | 3,3E-01 | 2,5E-03 | 1,5E-01 | 2,3E-05 | 1,2E-02 | 2,8E-02 | 0,0E+00 | 0,0E+00 | 0,0E+00 | 0,0E+00 | 0,0E+00 | 0,0E+00 | 3773125 | 3774456 | 1332 | - |
| Y11_RS17305 | Y11_36401 | msrA | peptide-methionine (S)-S-oxide reductase MsrA | 0,35 | -0,48 | -0,76 | -0,83 | -1,12 | -0,29 | 1,6E-01 | 1,1E-01 | 2,0E-01 | 1,9E-02 | 7,9E-02 | 6,2E-01 | 0,0E+00 | 0,0E+00 | 0,0E+00 | 0,0E+00 | 1,6E-03 | 0,0E+00 | 3774623 | 3775261 | 639 | - |
| Y11_RS17310 | Y11_36421 | | autotransporter assembly complex protein TamA | 0,26 | 0,23 | 0,24 | -0,03 | -0,02 | 0,01 | 8,4E-03 | 7,1E-03 | 7,7E-02 | 6,6E-01 | 8,6E-01 | 9,6E-01 | 0,0E+00 | 0,0E+00 | 0,0E+00 | 0,0E+00 | 0,0E+00 | 0,0E+00 | 3775735 | 3777471 | 1737 | + |
| Y11_RS17315 | Y11_36431 | | translocation/assembly module TamB | 0,3 | 0,41 | 0,93 | 0,11 | 0,63 | 0,52 | 5,4E-05 | 4,3E-06 | 2,1E-07 | 3,3E-02 | 7,2E-06 | 2,9E-05 | 0,0E+00 | 0,0E+00 | 0,0E+00 | 0,0E+00 | 0,0E+00 | 0,0E+00 | 3777540 | 3781352 | 3813 | + |
| Y11_RS17320 | Y11_36441 | | gamma-glutamylcyclotransferase | 0,03 | 0,07 | -0,05 | 0,04 | -0,08 | -0,12 | 8,3E-01 | 6,4E-01 | 7,5E-01 | 7,5E-01 | 5,5E-01 | 4,0E-01 | 0,0E+00 | 0,0E+00 | 0,0E+00 | 0,0E+00 | 0,0E+00 | 0,0E+00 | 3781355 | 3781717 | 363 | + |
| Y11_RS17325 | Y11_36451 | ppa | inorganic diphosphatase | -0,31 | 0,41 | 0,77 | 0,71 | 1,08 | 0,37 | 7,3E-05 | 7,7E-04 | 5,6E-02 | 1,8E-05 | 1,4E-02 | 3,3E-01 | 0,0E+00 | 0,0E+00 | 0,0E+00 | 0,0E+00 | 3,0E-04 | 0,0E+00 | 3781989 | 3782516 | 528 | - |
| Y11_RS17335 | Y11_36481 | fbp | class 1 fructose-bisphosphatase | 0,11 | 1,61 | 1,29 | 1,5 | 1,18 | -0,32 | 1,1E-02 | 5,9E-11 | 7,9E-09 | 4,1E-10 | 3,2E-08 | 5,4E-04 | 0,0E+00 | 1,6E-11 | 4,5E-10 | 4,6E-11 | 2,5E-09 | 0,0E+00 | 3784583 | 3785596 | 1014 | - |
| Y11_RS17340 | Y11_36491 | mpl | UDP-N-acetylmuramate:L-alanyl-gamma-D-glutamyl-meso-diaminopimelate ligase | 0,23 | 0,21 | 0 | -0,03 | -0,23 | -0,21 | 3,0E-03 | 1,0E-02 | 1,0E+00 | 7,0E-01 | 1,5E-02 | 3,3E-02 | 0,0E+00 | 0,0E+00 | 0,0E+00 | 0,0E+00 | 0,0E+00 | 0,0E+00 | 3785766 | 3787151 | 1386 | + |
| Y11_RS17350 | Y11_36511 | argR | transcriptional regulator ArgR | -0,38 | 0,12 | 0,58 | 0,5 | 0,96 | 0,46 | 1,6E-03 | 1,4E-01 | 4,5E-03 | 1,3E-03 | 3,9E-04 | 2,3E-02 | 0,0E+00 | 0,0E+00 | 0,0E+00 | 0,0E+00 | 0,0E+00 | 0,0E+00 | 3788081 | 3788551 | 471 | - |
| Y11_RS17355 | Y11_36521 | mdh | malate dehydrogenase | -0,36 | 0,48 | 0,73 | 0,85 | 1,09 | 0,25 | 1,9E-03 | 5,9E-04 | 2,7E-05 | 7,5E-05 | 1,0E-05 | 6,8E-02 | 0,0E+00 | 0,0E+00 | 0,0E+00 | 0,0E+00 | 3,3E-07 | 0,0E+00 | 3789015 | 3789950 | 936 | + |
| Y11_RS17370 | Y11_36541 | ispB | octaprenyl diphosphate synthase | 0,53 | 0,42 | 0,43 | -0,11 | -0,1 | 0,01 | 2,3E-03 | 6,3E-02 | 7,2E-03 | 5,6E-01 | 2,6E-01 | 9,7E-01 | 0,0E+00 | 0,0E+00 | 0,0E+00 | 0,0E+00 | 0,0E+00 | 0,0E+00 | 3790889 | 3791860 | 972 | - |
| Y11_RS17375 | Y11_36551 | rplU | 50S ribosomal protein L21 | 0,24 | -0,54 | -1,06 | -0,77 | -1,29 | -0,52 | 9,2E-04 | 2,8E-03 | 1,0E-03 | 3,2E-04 | 2,8E-04 | 6,3E-02 | 0,0E+00 | 0,0E+00 | 1,2E-05 | 0,0E+00 | 7,0E-06 | 0,0E+00 | 3792120 | 3792431 | 312 | + |
| Y11_RS17380 | Y11_36561 | rpmA | 50S ribosomal protein L27 | 0,22 | -0,36 | -0,71 | -0,58 | -0,93 | -0,36 | 5,7E-02 | 5,5E-02 | 2,8E-03 | 4,4E-03 | 3,4E-04 | 1,1E-01 | 0,0E+00 | 0,0E+00 | 0,0E+00 | 0,0E+00 | 0,0E+00 | 0,0E+00 | 3792451 | 3792708 | 258 | + |
| Y11_RS17390 | Y11_36581 | cgtA | Obg family GTPase CgtA | 0,04 | 0,04 | -0,82 | 0 | -0,86 | -0,86 | 2,7E-01 | 4,5E-01 | 8,6E-07 | 9,6E-01 | 1,5E-06 | 5,0E-06 | 0,0E+00 | 0,0E+00 | 0,0E+00 | 0,0E+00 | 0,0E+00 | 0,0E+00 | 3793786 | 3794958 | 1173 | + |
| Y11_RS17395 | Y11_36591 | pmrB | two-component system sensor histidine kinase PmrB | -0,05 | -0,05 | 0,35 | 0 | 0,4 | 0,4 | 7,0E-01 | 7,0E-01 | 4,7E-02 | 9,8E-01 | 2,7E-03 | 2,3E-03 | 0,0E+00 | 0,0E+00 | 0,0E+00 | 0,0E+00 | 0,0E+00 | 0,0E+00 | 3795062 | 3796123 | 1062 | - |
| Y11_RS17400 | Y11_36601 | pmrA | two-component system response regulator PmrA | -0,19 | 0,17 | -0,16 | 0,36 | 0,03 | -0,33 | 1,2E-01 | 3,9E-01 | 3,4E-01 | 4,7E-02 | 7,9E-01 | 1,2E-01 | 0,0E+00 | 0,0E+00 | 0,0E+00 | 0,0E+00 | 0,0E+00 | 0,0E+00 | 3796123 | 3796782 | 660 | - |
| Y11_RS17405 | Y11_36611 | dacB | serine-type D-Ala-D-Ala carboxypeptidase | 0,25 | -0,49 | -0,94 | -0,74 | -1,18 | -0,44 | 9,1E-03 | 1,1E-03 | 2,1E-03 | 9,7E-05 | 5,1E-04 | 8,2E-02 | 0,0E+00 | 0,0E+00 | 0,0E+00 | 0,0E+00 | 1,2E-05 | 0,0E+00 | 3796788 | 3798236 | 1449 | - |
| Y11_RS17410 | Y11_36621 | greA | transcription elongation factor GreA | -0,12 | -1,44 | -1,87 | -1,32 | -1,76 | -0,44 | 2,9E-01 | 3,8E-07 | 8,4E-08 | 1,7E-06 | 3,0E-07 | 3,2E-03 | 0,0E+00 | 1,2E-08 | 2,8E-09 | 4,0E-08 | 1,5E-08 | 0,0E+00 | 3798495 | 3798971 | 477 | + |
| Y11_RS17415 | Y11_36631 | yhbY | ribosome assembly RNA-binding protein YhbY | -0,15 | 0,47 | 0,38 | 0,62 | 0,53 | -0,09 | 6,9E-01 | 6,8E-02 | 1,3E-01 | 8,4E-02 | 1,3E-01 | 5,8E-01 | 0,0E+00 | 0,0E+00 | 0,0E+00 | 0,0E+00 | 0,0E+00 | 0,0E+00 | 3799098 | 3799391 | 294 | - |
| Y11_RS17420 | Y11_36641 | rlmE | 23S rRNA (uridine(2552)-2'-O)-methyltransferase RlmE | 0,36 | 15 | 2,4 | 15 | 2,09 | 0 | 2,5E-02 | 0,0E+00 | 1,6E-08 | 0,0E+00 | 7,4E-07 | 0,0E+00 | 0,0E+00 | 0,0E+00 | 7,3E-10 | 0,0E+00 | 3,2E-08 | 0,0E+00 | 3799538 | 3800167 | 630 | + |
| Y11_RS17425 | Y11_36651 | ftsH | ATP-dependent zinc metalloprotease FtsH | 0,01 | 0,91 | 0,47 | 0,9 | 0,46 | -0,44 | 7,4E-01 | 1,7E-07 | 2,1E-06 | 4,2E-07 | 1,0E-05 | 7,5E-05 | 0,0E+00 | 0,0E+00 | 0,0E+00 | 0,0E+00 | 0,0E+00 | 0,0E+00 | 3800224 | 3802158 | 1935 | + |
| Y11_RS17430 | Y11_36661 | folP | dihydropteroate synthase | 0,34 | 0,24 | 0 | -0,1 | -0,35 | -0,24 | 1,9E-03 | 3,8E-03 | 9,9E-01 | 3,1E-01 | 9,0E-03 | 2,6E-02 | 0,0E+00 | 0,0E+00 | 0,0E+00 | 0,0E+00 | 0,0E+00 | 0,0E+00 | 3802474 | 3803307 | 834 | + |
| Y11_RS17435 | Y11_36671 | glmM | phosphoglucosamine mutase | -0,01 | 0,71 | 0,97 | 0,71 | 0,97 | 0,26 | 9,2E-01 | 6,4E-06 | 2,3E-07 | 2,2E-07 | 1,0E-09 | 6,6E-04 | 0,0E+00 | 0,0E+00 | 0,0E+00 | 0,0E+00 | 0,0E+00 | 0,0E+00 | 3803317 | 3804657 | 1341 | + |
| Y11_RS17440 | Y11_36681 | secG | preprotein translocase subunit SecG | -0,7 | -1,23 | -1,33 | -0,53 | -0,63 | -0,09 | 4,8E-04 | 1,1E-06 | 1,9E-05 | 1,1E-03 | 3,9E-03 | 5,0E-01 | 0,0E+00 | 3,1E-08 | 3,0E-07 | 0,0E+00 | 0,0E+00 | 0,0E+00 | 3804881 | 3805216 | 336 | + |
| Y11_RS17455 | Y11_36691 | rimP | ribosome maturation factor RimP | -0,12 | -0,05 | 0,03 | 0,06 | 0,15 | 0,09 | 4,8E-01 | 7,3E-01 | 8,2E-01 | 7,4E-01 | 4,1E-01 | 6,1E-01 | 0,0E+00 | 0,0E+00 | 0,0E+00 | 0,0E+00 | 0,0E+00 | 0,0E+00 | 3805747 | 3806199 | 453 | + |
| Y11_RS17460 | Y11_36701 | nusA | transcription termination factor NusA | -0,33 | -0,64 | -1,24 | -0,31 | -0,91 | -0,6 | 5,6E-05 | 1,3E-06 | 2,0E-07 | 4,0E-04 | 2,8E-06 | 9,1E-05 | 0,0E+00 | 0,0E+00 | 5,9E-09 | 0,0E+00 | 0,0E+00 | 0,0E+00 | 3806221 | 3807708 | 1488 | + |
| Y11_RS17465 | Y11_36711 | infB | translation initiation factor IF-2 | -0,15 | -0,73 | -1,42 | -0,58 | -1,27 | -0,69 | 3,5E-03 | 3,9E-07 | 2,2E-11 | 8,0E-06 | 1,4E-09 | 1,1E-06 | 0,0E+00 | 0,0E+00 | 9,9E-12 | 0,0E+00 | 2,1E-10 | 0,0E+00 | 3807733 | 3810411 | 2679 | + |
| Y11_RS17470 | Y11_36721 | rbfA | 30S ribosome-binding factor RbfA | -0,35 | -0,85 | -1,73 | -0,51 | -1,38 | -0,88 | 1,6E-05 | 1,3E-05 | 2,0E-08 | 3,4E-04 | 6,0E-08 | 4,5E-05 | 0,0E+00 | 0,0E+00 | 8,7E-10 | 0,0E+00 | 4,1E-09 | 0,0E+00 | 3810478 | 3810888 | 411 | + |
| Y11_RS17475 | Y11_36731 | truB | tRNA pseudouridine(55) synthase TruB | -0,03 | -0,78 | -2,06 | -0,75 | -2,03 | -1,27 | 6,8E-01 | 2,6E-06 | 1,3E-09 | 1,0E-05 | 4,7E-09 | 1,4E-07 | 0,0E+00 | 0,0E+00 | 1,2E-10 | 0,0E+00 | 5,3E-10 | 4,5E-08 | 3810888 | 3811877 | 990 | + |
| Y11_RS17480 | Y11_36741 | rpsO | 30S ribosomal protein S15 | 0,02 | 0,26 | -0,25 | 0,24 | -0,27 | -0,51 | 8,3E-01 | 1,8E-03 | 2,8E-02 | 2,9E-02 | 5,1E-02 | 1,0E-03 | 0,0E+00 | 0,0E+00 | 0,0E+00 | 0,0E+00 | 0,0E+00 | 0,0E+00 | 3811999 | 3812268 | 270 | + |
| Y11_RS17485 | Y11_36751 | pnp | polyribonucleotide nucleotidyltransferase | -0,06 | -0,1 | -0,78 | -0,03 | -0,71 | -0,68 | 3,6E-02 | 2,8E-02 | 8,9E-08 | 3,2E-01 | 8,8E-08 | 5,1E-07 | 0,0E+00 | 0,0E+00 | 0,0E+00 | 0,0E+00 | 0,0E+00 | 0,0E+00 | 3812531 | 3814651 | 2121 | + |
| Y11_RS17490 | Y11_36761 | nlpI | lipoprotein NlpI | -0,06 | 1,12 | 1,27 | 1,18 | 1,33 | 0,14 | 6,2E-01 | 6,5E-05 | 2,5E-06 | 1,7E-04 | 1,9E-05 | 4,3E-01 | 0,0E+00 | 1,4E-06 | 5,0E-08 | 3,0E-06 | 5,7E-07 | 0,0E+00 | 3814774 | 3815658 | 885 | + |
| Y11_RS17495 | Y11_36771 | | DEAD/DEAH family ATP-dependent RNA helicase | 0,36 | -0,91 | -2,88 | -1,28 | -3,24 | -1,96 | 3,4E-04 | 1,4E-06 | 1,1E-08 | 1,6E-08 | 2,4E-09 | 1,9E-07 | 0,0E+00 | 0,0E+00 | 5,7E-10 | 7,4E-10 | 3,3E-10 | 5,5E-08 | 3815844 | 3817817 | 1974 | + |
| Y11_RS17500 | Y11_36781 | | hypothetical protein | 0,02 | -4,46 | -4,53 | -4,47 | -4,55 | -0,07 | 8,3E-01 | 1,6E-12 | 2,8E-08 | 1,0E-11 | 3,3E-08 | 7,4E-01 | 0,0E+00 | 2,2E-12 | 1,2E-09 | 5,4E-12 | 2,5E-09 | 0,0E+00 | 3818177 | 3818449 | 273 | + |
| Y11_RS17505 | Y11_36791 | | LLM class flavin-dependent oxidoreductase | -0,39 | 1,99 | 1,96 | 2,38 | 2,35 | -0,03 | 2,0E-05 | 7,7E-09 | 3,1E-07 | 1,2E-09 | 6,2E-08 | 8,4E-01 | 0,0E+00 | 5,0E-10 | 8,5E-09 | 9,8E-11 | 4,2E-09 | 0,0E+00 | 3818867 | 3819889 | 1023 | - |
| Y11_RS17540 | Y11_36861 | | N-acetyltransferase | -1,23 | 0,18 | -0,3 | 1,41 | 0,93 | -0,48 | 1,3E-04 | 3,4E-01 | 4,0E-01 | 9,4E-09 | 1,2E-02 | 1,4E-01 | 5,4E-06 | 0,0E+00 | 0,0E+00 | 5,0E-10 | 0,0E+00 | 0,0E+00 | 3827962 | 3828465 | 504 | + |
| Y11_RS17555 | Y11_36891 | speF | ornithine decarboxylase SpeF | -0,39 | 0,06 | 0,39 | 0,46 | 0,78 | 0,32 | 7,2E-03 | 6,0E-01 | 1,2E-01 | 9,4E-04 | 6,0E-03 | 1,7E-01 | 0,0E+00 | 0,0E+00 | 0,0E+00 | 0,0E+00 | 0,0E+00 | 0,0E+00 | 3830292 | 3832457 | 2166 | - |
| Y11_RS17560 | Y11_36911 | | heparinase II/III family protein | 0,24 | -2,23 | 0,21 | -2,46 | -0,01 | 2,44 | 2,2E-01 | 2,7E-07 | 8,5E-02 | 2,0E-06 | 9,4E-01 | 1,8E-07 | 0,0E+00 | 9,4E-09 | 0,0E+00 | 4,6E-08 | 0,0E+00 | 5,4E-08 | 3833602 | 3835626 | 2025 | - |
| Y11_RS17570 | Y11_36931 | | sugar ABC transporter substrate-binding protein | 0,41 | 0,36 | 3,76 | -0,05 | 3,35 | 3,4 | 5,1E-03 | 3,4E-02 | 8,1E-10 | 6,3E-01 | 2,6E-12 | 9,2E-10 | 0,0E+00 | 0,0E+00 | 8,6E-11 | 0,0E+00 | 6,2E-12 | 1,4E-09 | 3837447 | 3838760 | 1314 | + |
| Y11_RS17595 | Y11_36981 | | YhbP family protein | -15 | 0 | 0 | 15 | 15 | 0 | 0,0E+00 | 0,0E+00 | 0,0E+00 | 0,0E+00 | 0,0E+00 | 0,0E+00 | 0,0E+00 | 0,0E+00 | 0,0E+00 | 0,0E+00 | 0,0E+00 | 0,0E+00 | 3842891 | 3843343 | 453 | + |
| Y11_RS17600 | Y11_36991 | | type II toxin-antitoxin system Phd/YefM family antitoxin | 0,03 | -0,03 | -0,14 | -0,06 | -0,17 | -0,11 | 8,4E-01 | 7,5E-01 | 4,1E-01 | 6,8E-01 | 4,1E-01 | 4,6E-01 | 0,0E+00 | 0,0E+00 | 0,0E+00 | 0,0E+00 | 0,0E+00 | 0,0E+00 | 3843551 | 3843802 | 252 | + |
| Y11_RS17605 | Y11_37001 | | type II toxin-antitoxin system RelE/ParE family toxin | -0,29 | 0,49 | 0,08 | 0,79 | 0,37 | -0,41 | 2,5E-02 | 1,3E-01 | 5,5E-01 | 3,0E-02 | 2,9E-02 | 2,1E-01 | 0,0E+00 | 0,0E+00 | 0,0E+00 | 0,0E+00 | 0,0E+00 | 0,0E+00 | 3843799 | 3844086 | 288 | + |
| Y11_RS17610 | Y11_37011 | | NAD(P)H-binding protein | -0,14 | 1,12 | 1,54 | 1,26 | 1,69 | 0,42 | 1,9E-01 | 1,4E-06 | 1,7E-06 | 8,1E-08 | 3,8E-07 | 3,6E-03 | 0,0E+00 | 4,0E-08 | 3,7E-08 | 2,8E-09 | 1,9E-08 | 0,0E+00 | 3844162 | 3844791 | 630 | + |
| Y11_RS17680 | Y11_37171 | nrdD | anaerobic ribonucleoside-triphosphate reductase | 0,06 | -0,48 | 0,11 | -0,54 | 0,05 | 0,59 | 8,4E-01 | 3,6E-02 | 6,7E-01 | 1,1E-01 | 8,9E-01 | 4,3E-02 | 0,0E+00 | 0,0E+00 | 0,0E+00 | 0,0E+00 | 0,0E+00 | 0,0E+00 | 3856725 | 3858863 | 2139 | - |
| Y11_RS17690 | Y11_37201 | | ABC transporter ATP-binding protein | -0,17 | -0,85 | -0,73 | -0,68 | -0,56 | 0,12 | 1,9E-02 | 2,6E-06 | 1,9E-05 | 1,4E-06 | 2,5E-05 | 1,6E-01 | 0,0E+00 | 0,0E+00 | 0,0E+00 | 0,0E+00 | 0,0E+00 | 0,0E+00 | 3859875 | 3860591 | 717 | - |
| Y11_RS17695 | Y11_37211 | | ABC transporter ATP-binding protein | -0,25 | -0,74 | -0,49 | -0,49 | -0,24 | 0,25 | 2,3E-03 | 2,4E-05 | 2,3E-02 | 6,8E-05 | 1,8E-01 | 2,0E-01 | 0,0E+00 | 0,0E+00 | 0,0E+00 | 0,0E+00 | 0,0E+00 | 0,0E+00 | 3860578 | 3861456 | 879 | - |
| Y11_RS17700 | Y11_37221 | | ABC transporter permease | -0,33 | -0,71 | -0,3 | -0,38 | 0,03 | 0,41 | 2,0E-02 | 6,3E-04 | 2,7E-01 | 9,2E-03 | 9,1E-01 | 1,4E-01 | 0,0E+00 | 0,0E+00 | 0,0E+00 | 0,0E+00 | 0,0E+00 | 0,0E+00 | 3861449 | 3862288 | 840 | - |
| Y11_RS17705 | Y11_37231 | | ABC transporter permease | -0,04 | -1,16 | -0,76 | -1,12 | -0,72 | 0,4 | 8,3E-01 | 8,5E-06 | 5,5E-02 | 2,2E-04 | 8,1E-02 | 2,9E-01 | 0,0E+00 | 2,1E-07 | 0,0E+00 | 3,8E-06 | 0,0E+00 | 0,0E+00 | 3862290 | 3863339 | 1050 | - |
| Y11_RS17710 | Y11_37241 | | ABC transporter substrate-binding protein | 0,3 | -0,97 | -5,25 | -1,28 | -5,56 | -4,28 | 5,3E-03 | 2,8E-05 | 4,3E-14 | 1,0E-05 | 1,8E-12 | 2,7E-10 | 0,0E+00 | 0,0E+00 | 4,6E-13 | 2,1E-07 | 5,6E-12 | 1,0E-09 | 3863502 | 3865070 | 1569 | - |
| Y11_RS17715 | Y11_37261 | | YhcH/YjgK/YiaL family protein | -1,28 | -1,17 | -0,35 | 0,1 | 0,93 | 0,83 | 3,0E-05 | 5,9E-06 | 1,0E-02 | 5,6E-01 | 4,9E-04 | 2,2E-04 | 1,4E-06 | 1,5E-07 | 0,0E+00 | 0,0E+00 | 0,0E+00 | 0,0E+00 | 3865623 | 3866081 | 459 | + |
| Y11_RS17720 | Y11_37271 | argF | ornithine carbamoyltransferase | 0 | -0,61 | 0 | -1,01 | 0 | 1,73 | 0,0E+00 | 4,0E-02 | 0,0E+00 | 3,6E-03 | 0,0E+00 | 8,3E-05 | 0,0E+00 | 0,0E+00 | 0,0E+00 | 5,9E-05 | 0,0E+00 | 7,1E-06 | 3866234 | 3867241 | 1008 | - |
| Y11_RS17725 | Y11_37281 | rraB | ribonuclease E inhibitor RraB | 0,78 | -0,06 | 1,02 | -0,84 | 0,24 | 1,08 | 2,3E-04 | 2,0E-01 | 7,2E-02 | 1,3E-04 | 6,5E-01 | 5,9E-02 | 0,0E+00 | 0,0E+00 | 8,0E-04 | 0,0E+00 | 0,0E+00 | 3,1E-03 | 3867412 | 3867843 | 432 | + |
| Y11_RS17735 | Y11_37311 | | GNAT family N-acetyltransferase | 0,31 | -3,5 | -4,76 | -3,81 | -5,07 | -1,26 | 7,2E-02 | 1,1E-07 | 1,5E-08 | 7,8E-09 | 1,5E-09 | 3,6E-04 | 0,0E+00 | 4,4E-09 | 7,1E-10 | 4,3E-10 | 2,3E-10 | 2,6E-05 | 3869649 | 3870152 | 504 | - |
| Y11_RS17740 | Y11_37321 | | valine--tRNA ligase | -0,04 | -0,11 | -0,18 | -0,07 | -0,14 | -0,07 | 2,6E-01 | 9,3E-03 | 2,8E-02 | 7,3E-02 | 7,9E-02 | 3,5E-01 | 0,0E+00 | 0,0E+00 | 0,0E+00 | 0,0E+00 | 0,0E+00 | 0,0E+00 | 3870485 | 3873382 | 2898 | - |
| Y11_RS17745 | Y11_37331 | | DNA polymerase III subunit chi | 0,15 | 0,46 | 0,69 | 0,31 | 0,55 | 0,23 | 1,7E-01 | 6,3E-04 | 8,0E-02 | 2,1E-02 | 1,6E-01 | 5,2E-01 | 0,0E+00 | 0,0E+00 | 0,0E+00 | 0,0E+00 | 0,0E+00 | 0,0E+00 | 3873396 | 3873845 | 450 | - |
| Y11_RS17750 | Y11_37341 | pepA | leucyl aminopeptidase | -0,05 | 0,4 | 0,47 | 0,45 | 0,52 | 0,06 | 3,8E-01 | 6,7E-07 | 4,6E-07 | 1,7E-05 | 8,1E-06 | 3,6E-02 | 0,0E+00 | 0,0E+00 | 0,0E+00 | 0,0E+00 | 0,0E+00 | 0,0E+00 | 3874095 | 3875606 | 1512 | - |

| | | | | | | | | | | | | | | | | | | | | | | | | | |
|---|---|---|---|---|---|---|---|---|---|---|---|---|---|---|---|---|---|---|---|---|---|---|---|---|---|
| Y11_RS17755 | Y11_37351 | lptF | LPS export ABC transporter permease LptF | 0,03 | 0,42 | 0,74 | 0,39 | 0,7 | 0,31 | 5,7E-01 | 1,2E-04 | 4,0E-05 | 3,4E-04 | 7,0E-05 | 1,3E-02 | 0,0E+00 | 0,0E+00 | 0,0E+00 | 0,0E+00 | 0,0E+00 | 0,0E+00 | 3875886 | 3876980 | 1095 | + |
| Y11_RS17760 | Y11_37361 | lptG | LPS export ABC transporter permease LptG | 0,11 | 0,2 | 0,43 | 0,09 | 0,32 | 0,23 | 4,8E-01 | 1,2E-01 | 6,3E-03 | 5,2E-01 | 4,8E-02 | 5,2E-02 | 0,0E+00 | 0,0E+00 | 0,0E+00 | 0,0E+00 | 0,0E+00 | 0,0E+00 | 3876980 | 3878050 | 1071 | + |
| Y11_RS17775 | Y11_37391 | | ribbon-helix-helix protein, CopG family | -0,09 | 0 | 0,38 | 0,09 | 0,47 | 0,38 | 5,8E-01 | 1,0E+00 | 4,7E-02 | 6,1E-01 | 6,3E-03 | 6,3E-02 | 0,0E+00 | 0,0E+00 | 0,0E+00 | 0,0E+00 | 0,0E+00 | 0,0E+00 | 3879286 | 3879534 | 249 | + |
| Y11_RS17780 | Y11_37401 | | type II toxin-antitoxin system RelE/ParE family toxin | -0,37 | 0,02 | -0,13 | 0,39 | 0,24 | -0,16 | 1,5E-04 | 7,3E-01 | 2,7E-01 | 1,3E-04 | 5,3E-02 | 2,0E-01 | 0,0E+00 | 0,0E+00 | 0,0E+00 | 0,0E+00 | 0,0E+00 | 0,0E+00 | 3879538 | 3879813 | 276 | + |
| Y11_RS17785 | Y11_37411 | | helix-turn-helix domain-containing protein | -0,36 | -0,03 | -0,17 | 0,33 | 0,19 | -0,14 | 2,2E-04 | 6,3E-01 | 2,3E-01 | 1,7E-04 | 1,7E-01 | 3,1E-01 | 0,0E+00 | 0,0E+00 | 0,0E+00 | 0,0E+00 | 0,0E+00 | 0,0E+00 | 3879932 | 3880720 | 789 | + |
| Y11_RS17810 | Y11_37461 | | GNAT family N-acetyltransferase | -0,16 | 1,43 | 2,87 | 1,59 | 3,04 | 1,44 | 9,2E-02 | 9,2E-07 | 2,0E-06 | 3,6E-08 | 9,0E-07 | 2,7E-04 | 0,0E+00 | 2,7E-08 | 4,1E-08 | 1,4E-09 | 3,8E-08 | 2,0E-05 | 3885398 | 3885808 | 411 | + |
| Y11_RS17815 | Y11_37471 | | PAS and helix-turn-helix domain-containing protein | 0,07 | 0,17 | 15 | 0,1 | 15 | 15 | 7,2E-01 | 2,7E-01 | 0,0E+00 | 5,4E-01 | 0,0E+00 | 0,0E+00 | 0,0E+00 | 0,0E+00 | 0,0E+00 | 0,0E+00 | 0,0E+00 | 0,0E+00 | 3885834 | 3886523 | 690 | + |
| Y11_RS17845 | Y11_37551 | | DUF2501 domain-containing protein | 0 | -15 | -15 | -15 | -15 | -1,24 | 0,0E+00 | 0,0E+00 | 0,0E+00 | 0,0E+00 | 0,0E+00 | 9,0E-05 | 0,0E+00 | 0,0E+00 | 0,0E+00 | 0,0E+00 | 0,0E+00 | 7,5E-06 | 3892742 | 3893224 | 483 | - |
| Y11_RS17855 | Y11_37571 | lsrG | (4S)-4-hydroxy-5-phosphonooxypentane-2,3-dione isomerase | -0,3 | 1,2 | 1,94 | 1,49 | 2,23 | 0,74 | 1,5E-02 | 9,3E-05 | 4,2E-07 | 2,7E-05 | 2,5E-07 | 5,4E-03 | 0,0E+00 | 2,0E-06 | 1,1E-08 | 5,2E-07 | 1,3E-08 | 0,0E+00 | 3893687 | 3893977 | 291 | - |
| Y11_RS17860 | Y11_37581 | lsrF | 3-hydroxy-5-phosphonooxypentane-2,4-dione thiolase | -0,51 | 2,05 | 2,68 | 2,56 | 3,18 | 0,62 | 3,4E-03 | 4,5E-08 | 1,9E-08 | 1,3E-08 | 7,1E-09 | 4,6E-04 | 0,0E+00 | 2,1E-09 | 8,6E-10 | 6,5E-10 | 7,4E-10 | 0,0E+00 | 3894039 | 3894914 | 876 | - |
| Y11_RS17865 | Y11_37591 | lsrB | autoinducer 2 ABC transporter substrate-binding protein LsrB | 0,03 | 2,81 | 3,84 | 2,78 | 3,81 | 1,03 | 4,8E-01 | 3,8E-09 | 9,2E-09 | 3,6E-09 | 9,2E-09 | 5,2E-04 | 0,0E+00 | 2,9E-10 | 5,0E-10 | 2,2E-10 | 9,1E-10 | 3,7E-05 | 3894970 | 3896004 | 1035 | - |
| Y11_RS17875 | Y11_37611 | lsrC | autoinducer 2 ABC transporter permease LsrC | -0,91 | 0,08 | 15 | 0,98 | 15 | 0 | 1,1E-02 | 7,0E-01 | 0,0E+00 | 2,9E-03 | 0,0E+00 | 0,0E+00 | 0,0E+00 | 0,0E+00 | 0,0E+00 | 0,0E+00 | 0,0E+00 | 0,0E+00 | 3897053 | 3898150 | 1098 | - |
| Y11_RS17880 | Y11_37621 | lsrA | autoinducer 2 ABC transporter ATP-binding protein LsrA | -0,45 | 15 | 15 | 15 | 15 | 0 | 8,4E-02 | 0,0E+00 | 0,0E+00 | 0,0E+00 | 0,0E+00 | 0,0E+00 | 0,0E+00 | 0,0E+00 | 0,0E+00 | 0,0E+00 | 0,0E+00 | 0,0E+00 | 3898144 | 3899709 | 1566 | - |
| Y11_RS17885 | Y11_37631 | lsrR | transcriptional regulator LsrR | 0,15 | 1,25 | 1,43 | 1,1 | 1,28 | 0,18 | 1,6E-01 | 6,7E-07 | 2,1E-05 | 5,7E-07 | 3,4E-05 | 2,6E-01 | 0,0E+00 | 2,1E-08 | 3,3E-07 | 1,5E-08 | 9,9E-07 | 0,0E+00 | 3899927 | 3900922 | 996 | + |
| Y11_RS17890 | Y11_37641 | lsrK | autoinducer-2 kinase | -0,15 | 0,81 | 1,3 | 0,96 | 1,45 | 0,49 | 5,4E-01 | 5,8E-03 | 1,4E-04 | 2,2E-03 | 6,3E-05 | 1,6E-02 | 0,0E+00 | 0,0E+00 | 1,8E-06 | 0,0E+00 | 1,7E-06 | 0,0E+00 | 3901263 | 3902843 | 1581 | + |
| Y11_RS17895 | Y11_37651 | rfaQ | putative lipopolysaccharide heptosyltransferase III | -0,12 | 0,28 | 1,08 | 0,4 | 1,2 | 0,8 | 1,6E-01 | 7,9E-03 | 3,9E-05 | 4,4E-04 | 1,3E-05 | 2,4E-04 | 0,0E+00 | 0,0E+00 | 5,7E-07 | 0,0E+00 | 4,2E-07 | 0,0E+00 | 3903215 | 3904372 | 1158 | + |
| Y11_RS17900 | Y11_37661 | waaL-ps | O-antigen polysaccharide ligase WaaL-ps | 0,17 | -0,13 | 0,57 | -0,3 | 0,4 | 0,7 | 1,6E-01 | 1,5E-01 | 4,1E-03 | 2,9E-02 | 4,0E-02 | 1,3E-03 | 0,0E+00 | 0,0E+00 | 0,0E+00 | 0,0E+00 | 0,0E+00 | 0,0E+00 | 3904438 | 3905682 | 1245 | + |
| Y11_RS17905 | Y11_37671 | | amino acid permease | 15 | 2,22 | 15 | 0 | 0 | 0 | 0,0E+00 | 1,1E-05 | 0,0E+00 | 0,0E+00 | 0,0E+00 | 0,0E+00 | 0,0E+00 | 2,6E-07 | 0,0E+00 | 0,0E+00 | 0,0E+00 | 0,0E+00 | 3906939 | 3908447 | 1509 | + |
| Y11_RS17910 | Y11_37681 | arcA | arginine deiminase | 3,41 | 2,08 | 2,79 | -1,32 | -0,62 | 0,71 | 9,3E-12 | 1,9E-09 | 2,0E-09 | 1,2E-08 | 9,1E-05 | 6,5E-05 | 7,8E-12 | 1,7E-10 | 1,6E-10 | 6,1E-10 | 0,0E+00 | 0,0E+00 | 3908463 | 3909746 | 1284 | + |
| Y11_RS17915 | Y11_37691 | | efflux RND transporter periplasmic adaptor subunit | 1,89 | 0,54 | 0,45 | -1,35 | -1,44 | -0,09 | 8,9E-10 | 1,4E-07 | 4,5E-04 | 1,1E-08 | 2,8E-07 | 2,8E-01 | 2,1E-10 | 0,0E+00 | 0,0E+00 | 5,8E-10 | 1,4E-08 | 0,0E+00 | 3910106 | 3911188 | 1083 | + |
| Y11_RS17920 | Y11_37701 | | efflux RND transporter permease subunit | 1,88 | 0,91 | 0,59 | -0,97 | -1,29 | -0,33 | 5,4E-09 | 4,7E-06 | 1,4E-02 | 4,6E-06 | 1,3E-04 | 1,3E-01 | 9,3E-10 | 0,0E+00 | 0,0E+00 | 0,0E+00 | 3,4E-06 | 0,0E+00 | 3911188 | 3914274 | 3087 | + |
| Y11_RS17925 | Y11_37711 | | glycerol dehydrogenase | -0,9 | -0,19 | -0,03 | 0,71 | 0,87 | 0,16 | 1,1E-06 | 2,5E-01 | 8,2E-01 | 2,1E-03 | 2,7E-04 | 4,4E-01 | 0,0E+00 | 0,0E+00 | 0,0E+00 | 0,0E+00 | 0,0E+00 | 0,0E+00 | 3914584 | 3915678 | 1095 | + |
| Y11_RS17930 | Y11_37721 | dhaK | dihydroxyacetone kinase subunit DhaK | -1,14 | 0,61 | 1,21 | 1,74 | 2,35 | 0,61 | 1,0E-06 | 8,3E-05 | 2,9E-05 | 4,5E-07 | 6,0E-07 | 6,2E-03 | 6,3E-08 | 0,0E+00 | 4,4E-07 | 1,2E-08 | 2,7E-08 | 0,0E+00 | 3915773 | 3916840 | 1068 | + |
| Y11_RS17935 | Y11_37731 | dhaL | dihydroxyacetone kinase ADP-binding subunit DhaL | -0,9 | 0,81 | 1,35 | 1,71 | 2,25 | 0,54 | 6,8E-05 | 4,1E-06 | 7,9E-07 | 6,7E-07 | 2,1E-07 | 7,5E-04 | 0,0E+00 | 0,0E+00 | 1,9E-08 | 1,7E-08 | 1,2E-08 | 0,0E+00 | 3916858 | 3917490 | 633 | + |
| Y11_RS17940 | Y11_37741 | dhaM | dihydroxyacetone kinase subunit DhaM | -0,65 | 1,25 | 0,66 | 1,91 | 1,31 | -0,6 | 3,8E-02 | 8,6E-03 | 8,0E-02 | 3,6E-04 | 1,7E-03 | 1,5E-01 | 0,0E+00 | 1,7E-04 | 0,0E+00 | 6,3E-06 | 3,8E-05 | 0,0E+00 | 3917524 | 3918957 | 1434 | + |
| Y11_RS17985 | Y11_37831 | | LacI family DNA-binding transcriptional regulator | -0,13 | 0,4 | 0,74 | 0,53 | 0,87 | 0,34 | 2,5E-01 | 7,9E-03 | 3,9E-03 | 5,0E-04 | 1,0E-03 | 9,5E-02 | 0,0E+00 | 0,0E+00 | 0,0E+00 | 0,0E+00 | 0,0E+00 | 0,0E+00 | 3930658 | 3931692 | 1035 | - |
| Y11_RS17990 | Y11_37841 | | sucrose-6-phosphate hydrolase | -0,21 | -0,65 | -1,7 | -0,43 | -1,49 | -1,05 | 2,6E-01 | 9,4E-03 | 6,9E-06 | 6,7E-02 | 3,8E-05 | 7,3E-04 | 0,0E+00 | 0,0E+00 | 1,2E-07 | 0,0E+00 | 1,1E-06 | 5,0E-05 | 3931739 | 3933148 | 1410 | - |
| Y11_RS17995 | Y11_37851 | | sucrose-specific PTS transporter subunit IIBC | -0,27 | -0,43 | -0,95 | -0,16 | -0,68 | -0,51 | 9,5E-05 | 2,9E-05 | 7,4E-03 | 2,4E-02 | 3,5E-02 | 9,4E-02 | 0,0E+00 | 0,0E+00 | 0,0E+00 | 0,0E+00 | 0,0E+00 | 0,0E+00 | 3933148 | 3934518 | 1371 | - |
| Y11_RS18000 | Y11_37861 | | carbohydrate porin | -0,17 | -0,16 | -1,09 | 0 | -0,93 | -0,93 | 4,6E-02 | 2,6E-01 | 4,7E-04 | 9,8E-01 | 1,2E-03 | 2,9E-03 | 0,0E+00 | 0,0E+00 | 5,8E-06 | 0,0E+00 | 0,0E+00 | 0,0E+00 | 3934586 | 3936124 | 1539 | - |
| Y11_RS18005 | Y11_37871 | | aminoimidazole riboside kinase | 0 | 0,5 | -3,38 | 0 | -3,88 | -3,88 | 0,0E+00 | 2,0E-02 | 2,0E-06 | 9,9E-01 | 3,5E-07 | 6,6E-07 | 0,0E+00 | 0,0E+00 | 4,1E-08 | 0,0E+00 | 1,7E-08 | 1,4E-07 | 3936423 | 3937358 | 936 | - |
| Y11_RS18010 | Y11_37881 | | ABC transporter substrate-binding protein | 0,95 | 0,97 | -1,47 | 0,03 | -2,43 | -2,44 | 3,6E-03 | 3,4E-03 | 3,2E-02 | 8,7E-01 | 2,1E-03 | 2,1E-03 | 0,0E+00 | 0,0E+00 | 3,6E-04 | 0,0E+00 | 4,8E-05 | 1,3E-04 | 3937562 | 3938707 | 1146 | + |
| Y11_RS18015 | Y11_37891 | | TonB-dependent siderophore receptor | 2 | 0,26 | -0,74 | -1,73 | -2,75 | -1,01 | 1,0E-06 | 4,5E-03 | 2,2E-06 | 3,2E-06 | 7,7E-08 | 2,7E-07 | 6,3E-08 | 0,0E+00 | 0,0E+00 | 7,2E-08 | 5,0E-09 | 6,9E-08 | 3939193 | 3941349 | 2157 | + |
| Y11_RS18050 | Y11_37961 | rsmC | 16S rRNA (guanine(1207)-N(2))-methyltransferase RsmC | 0,2 | 0,46 | 0,75 | 0,26 | 0,55 | 0,28 | 1,7E-02 | 3,6E-05 | 2,7E-06 | 4,1E-03 | 6,9E-05 | 2,1E-03 | 0,0E+00 | 0,0E+00 | 0,0E+00 | 0,0E+00 | 0,0E+00 | 0,0E+00 | 3946285 | 3947328 | 1044 | - |
| Y11_RS18055 | Y11_37971 | | DNA polymerase III subunit psi | -0,15 | -0,15 | -0,35 | -0,01 | -0,2 | -0,19 | 3,2E-01 | 2,6E-01 | 1,1E-01 | 9,5E-01 | 3,1E-01 | 3,0E-01 | 0,0E+00 | 0,0E+00 | 0,0E+00 | 0,0E+00 | 0,0E+00 | 0,0E+00 | 3947441 | 3947878 | 438 | + |
| Y11_RS18060 | Y11_37981 | rimI | ribosomal protein S18-alanine N-acetyltransferase | 0,92 | 1,39 | 2,56 | 0,47 | 1,64 | 1,16 | 8,4E-03 | 1,3E-03 | 9,7E-05 | 9,2E-02 | 1,0E-03 | 9,8E-03 | 0,0E+00 | 2,6E-05 | 1,3E-06 | 0,0E+00 | 2,4E-05 | 5,7E-04 | 3947823 | 3948266 | 444 | + |
| Y11_RS18065 | Y11_37991 | prfC | peptide chain release factor 3 | 0,32 | -0,01 | 0,06 | -0,34 | -0,27 | 0,07 | 4,6E-03 | 8,8E-01 | 5,2E-01 | 5,8E-04 | 1,6E-03 | 3,0E-01 | 0,0E+00 | 0,0E+00 | 0,0E+00 | 0,0E+00 | 0,0E+00 | 0,0E+00 | 3948517 | 3950106 | 1590 | + |
| Y11_RS18070 | Y11_38001 | osmY | molecular chaperone OsmY | 0,66 | 1,9 | 2,88 | 1,24 | 2,22 | 0,98 | 2,3E-05 | 4,1E-08 | 3,3E-08 | 2,0E-06 | 3,3E-07 | 2,4E-04 | 0,0E+00 | 1,9E-09 | 1,4E-09 | 4,6E-08 | 1,7E-08 | 0,0E+00 | 3950418 | 3951032 | 615 | + |
| Y11_RS18075 | Y11_38021 | | patatin family protein | 0,15 | 0,47 | 0,19 | 0,32 | 0,04 | -0,28 | 5,4E-01 | 1,7E-02 | 1,3E-01 | 2,4E-01 | 8,6E-01 | 8,2E-02 | 0,0E+00 | 0,0E+00 | 0,0E+00 | 0,0E+00 | 0,0E+00 | 0,0E+00 | 3951476 | 3952729 | 1254 | + |
| Y11_RS18090 | Y11_38051 | deoC | deoxyribose-phosphate aldolase | 0,1 | 1,72 | 1,03 | 1,62 | 0,93 | -0,69 | 1,2E-01 | 1,6E-09 | 3,2E-04 | 3,1E-09 | 6,4E-04 | 3,8E-03 | 0,0E+00 | 1,5E-10 | 4,1E-06 | 2,0E-10 | 0,0E+00 | 0,0E+00 | 3955396 | 3956193 | 798 | + |
| Y11_RS18095 | Y11_38061 | deoA | thymidine phosphorylase | 0,08 | 1,35 | 0,3 | 1,27 | 0,22 | -1,05 | 1,3E-01 | 1,4E-09 | 2,4E-02 | 2,0E-10 | 6,9E-02 | 6,0E-06 | 0,0E+00 | 1,4E-10 | 0,0E+00 | 2,7E-11 | 0,0E+00 | 7,4E-07 | 3956296 | 3957618 | 1323 | + |
| Y11_RS18100 | Y11_38071 | deoB | phosphopentomutase | 0,24 | 1,27 | 0,46 | 1,04 | 0,22 | -0,82 | 2,8E-04 | 4,9E-08 | 9,8E-03 | 6,7E-08 | 1,3E-01 | 4,1E-04 | 0,0E+00 | 2,2E-09 | 0,0E+00 | 2,4E-09 | 0,0E+00 | 0,0E+00 | 3957672 | 3958895 | 1224 | + |
| Y11_RS18105 | Y11_38081 | deoD | purine-nucleoside phosphorylase | 0,06 | 1,49 | 0,46 | 1,43 | 0,41 | -1,02 | 5,9E-01 | 1,6E-07 | 1,9E-02 | 2,5E-07 | 3,4E-02 | 1,6E-04 | 0,0E+00 | 6,1E-09 | 0,0E+00 | 7,3E-09 | 0,0E+00 | 1,3E-05 | 3959115 | 3959834 | 720 | + |
| Y11_RS18110 | Y11_38091 | | YtjB family periplasmic protein | -0,44 | 3,49 | 15 | 0 | 0 | 0 | 6,7E-01 | 9,6E-03 | 0,0E+00 | 0,0E+00 | 0,0E+00 | 0,0E+00 | 0,0E+00 | 1,8E-04 | 0,0E+00 | 0,0E+00 | 0,0E+00 | 0,0E+00 | 3959889 | 3960566 | 678 | - |
| Y11_RS18115 | Y11_38101 | serB | phosphoserine phosphatase | -15 | -15 | -15 | -0,99 | -2,09 | -1,09 | 0,0E+00 | 0,0E+00 | 0,0E+00 | 1,0E-02 | 1,2E-04 | 1,1E-02 | 0,0E+00 | 0,0E+00 | 0,0E+00 | 0,0E+00 | 3,1E-06 | 6,3E-04 | 3960802 | 3961779 | 978 | + |
| Y11_RS18120 | Y11_38111 | radA | DNA repair protein RadA | -0,36 | -0,99 | -1,22 | -0,63 | -0,87 | -0,24 | 3,4E-03 | 5,1E-08 | 3,0E-06 | 7,5E-05 | 1,3E-04 | 5,4E-02 | 0,0E+00 | 0,0E+00 | 5,9E-08 | 0,0E+00 | 0,0E+00 | 0,0E+00 | 3961952 | 3963334 | 1383 | + |
| Y11_RS18135 | Y11_38131 | nadR | multifunctional transcriptional regulator/nicotinamide-nucleotide adenylyltransferase/ribosylnicotinamide kinase NadR | -0,13 | 0,22 | 0,25 | 0,36 | 0,38 | 0,02 | 7,3E-02 | 3,9E-02 | 8,2E-02 | 4,4E-03 | 1,5E-02 | 8,8E-01 | 0,0E+00 | 0,0E+00 | 0,0E+00 | 0,0E+00 | 0,0E+00 | 0,0E+00 | 3964654 | 3965937 | 1284 | + |
| Y11_RS18140 | Y11_38141 | | zinc-binding alcohol dehydrogenase family protein | 0,84 | 2,55 | 2,48 | 1,71 | 1,64 | -0,07 | 2,0E-04 | 7,0E-08 | 6,9E-07 | 3,7E-06 | 2,6E-05 | 7,3E-01 | 0,0E+00 | 3,0E-09 | 1,7E-08 | 8,1E-08 | 7,7E-07 | 0,0E+00 | 3965934 | 3966935 | 1002 | - |
| Y11_RS18155 | Y11_38171 | ettA | energy-dependent translational throttle protein EttA | -0,17 | -0,7 | -1,07 | -0,53 | -0,91 | -0,38 | 5,2E-04 | 6,6E-09 | 5,6E-07 | 7,8E-09 | 1,7E-06 | 7,6E-04 | 0,0E+00 | 0,0E+00 | 1,4E-08 | 0,0E+00 | 0,0E+00 | 0,0E+00 | 3969261 | 3970928 | 1668 | - |
| Y11_RS18160 | Y11_38181 | | OmpA family protein | 0,58 | 0,95 | 0,96 | 0,37 | 0,37 | 0 | 1,3E-04 | 8,9E-05 | 1,8E-04 | 4,0E-02 | 5,3E-02 | 9,9E-01 | 0,0E+00 | 0,0E+00 | 0,0E+00 | 0,0E+00 | 0,0E+00 | 0,0E+00 | 3971287 | 3971796 | 510 | - |
| Y11_RS18165 | Y11_38191 | | diguanylate cyclase | 0,16 | 1,25 | 15 | 0 | 0 | 0 | 3,1E-01 | 4,7E-05 | 0,0E+00 | 0,0E+00 | 0,0E+00 | 0,0E+00 | 0,0E+00 | 1,0E-06 | 0,0E+00 | 0,0E+00 | 0,0E+00 | 0,0E+00 | 3971802 | 3973163 | 1362 | - |
| Y11_RS18175 | Y11_38211 | | cation-transporting P-type ATPase | -0,41 | 0,84 | 1,17 | 1,25 | 1,58 | 0,33 | 7,1E-04 | 5,8E-06 | 2,0E-07 | 2,4E-07 | 1,6E-08 | 1,9E-03 | 0,0E+00 | 0,0E+00 | 5,9E-09 | 7,1E-09 | 1,4E-09 | 0,0E+00 | 3973997 | 3976708 | 2712 | - |
| Y11_RS18180 | Y11_38221 | sltY | murein transglycosylase | 1,6 | -2,11 | -1,85 | -3,71 | -3,45 | 0,26 | 8,3E-06 | 6,2E-07 | 1,2E-05 | 8,6E-12 | 6,9E-09 | 7,2E-02 | 4,3E-07 | 1,9E-08 | 1,9E-07 | 5,4E-12 | 7,2E-10 | 0,0E+00 | 3977256 | 3979175 | 1920 | + |
| Y11_RS18185 | Y11_38231 | trpR | trp operon repressor | -0,08 | -0,44 | -0,34 | -0,36 | -0,27 | 0,1 | 4,9E-01 | 8,9E-04 | 1,5E-02 | 1,7E-03 | 3,6E-02 | 3,0E-01 | 0,0E+00 | 0,0E+00 | 0,0E+00 | 0,0E+00 | 0,0E+00 | 0,0E+00 | 3979449 | 3979781 | 333 | + |
| Y11_RS18190 | Y11_38241 | yjjX | inosine/xanthosine triphosphatase | -0,21 | -0,26 | -0,57 | -0,05 | -0,36 | -0,31 | 1,6E-01 | 8,3E-02 | 3,1E-04 | 7,7E-01 | 3,1E-02 | 4,8E-02 | 0,0E+00 | 0,0E+00 | 0,0E+00 | 0,0E+00 | 0,0E+00 | 0,0E+00 | 3979790 | 3980317 | 528 | - |
| Y11_RS18195 | Y11_38251 | gpmB | 2,3-diphosphoglycerate-dependent phosphoglycerate mutase GpmB | -0,11 | -0,29 | 0,25 | -0,18 | 0,37 | 0,55 | 9,5E-02 | 9,2E-03 | 1,3E-02 | 5,3E-02 | 9,4E-04 | 4,2E-04 | 0,0E+00 | 0,0E+00 | 0,0E+00 | 0,0E+00 | 0,0E+00 | 0,0E+00 | 3980374 | 3981021 | 648 | + |
| Y11_RS18200 | Y11_38261 | robA | MDR efflux pump AcrAB transcriptional activator RobA | -0,86 | 0,76 | 1,2 | 0,13 | 2,06 | 0,44 | 3,7E-06 | 4,9E-06 | 1,2E-05 | 5,3E-01 | 7,1E-08 | 3,1E-03 | 0,0E+00 | 0,0E+00 | 1,9E-07 | 0,0E+00 | 4,7E-09 | 0,0E+00 | 3981018 | 3981884 | 867 | - |
| Y11_RS18205 | Y11_38271 | creA | protein CreA | -0,26 | 1,13 | 1,35 | 1,39 | 1,61 | 0,22 | 1,9E-03 | 1,0E-08 | 1,0E-07 | 2,1E-08 | 7,5E-08 | 2,7E-02 | 0,0E+00 | 6,4E-10 | 3,3E-09 | 9,0E-10 | 4,9E-09 | 0,0E+00 | 3982109 | 3982576 | 468 | + |
| Y11_RS18210 | Y11_38281 | arcA | two-component system response regulator ArcA | 0,65 | 0,37 | 0,17 | -0,28 | -0,48 | -0,2 | 3,0E-07 | 8,3E-05 | 1,3E-02 | 5,7E-04 | 2,3E-05 | 1,3E-02 | 0,0E+00 | 0,0E+00 | 0,0E+00 | 0,0E+00 | 0,0E+00 | 0,0E+00 | 3982658 | 3983374 | 717 | - |
| Y11_RS18220 | Y11_38311 | thrA | bifunctional aspartate kinase/homoserine dehydrogenase I | -1,15 | -3,42 | -2,81 | -2,27 | -1,67 | 0,61 | 5,0E-09 | 6,7E-13 | 8,5E-09 | 9,2E-13 | 3,8E-07 | 5,9E-04 | 9,0E-10 | 1,6E-12 | 4,7E-10 | 1,9E-12 | 1,9E-08 | 0,0E+00 | 3984539 | 3986998 | 2460 | + |
| Y11_RS18225 | Y11_38321 | thrB | homoserine kinase | -0,74 | -2,9 | -2,12 | -2,17 | -1,39 | 0,78 | 3,2E-08 | 5,9E-11 | 7,3E-05 | 7,0E-10 | 1,3E-03 | 2,8E-02 | 0,0E+00 | 1,6E-11 | 1,0E-06 | 6,7E-11 | 2,8E-05 | 0,0E+00 | 3987001 | 3987930 | 930 | + |
| Y11_RS18230 | Y11_38331 | thrC | threonine synthase | -0,61 | -1,21 | -1,11 | -0,6 | -0,49 | 0,11 | 4,8E-05 | 3,9E-07 | 1,6E-04 | 1,1E-06 | 1,2E-02 | 5,1E-01 | 0,0E+00 | 1,3E-08 | 2,1E-06 | 0,0E+00 | 0,0E+00 | 0,0E+00 | 3987934 | 3989223 | 1290 | + |
| Y11_RS18240 | Y11_38351 | yaaA | peroxide stress protein YaaA | -1,29 | -1,22 | -1,3 | 0,07 | -0,01 | -0,08 | 6,5E-09 | 9,5E-08 | 2,2E-07 | 3,6E-01 | 9,3E-01 | 4,4E-01 | 1,0E-09 | 3,9E-09 | 6,4E-09 | 0,0E+00 | 0,0E+00 | 0,0E+00 | 3990837 | 3991613 | 777 | - |
| Y11_RS18245 | Y11_38361 | tal | transaldolase | 0,32 | 0,5 | 0,28 | 0,18 | -0,03 | -0,22 | 8,9E-04 | 1,1E-04 | 9,8E-05 | 6,2E-02 | 5,8E-01 | 1,4E-02 | 0,0E+00 | 0,0E+00 | 0,0E+00 | 0,0E+00 | 0,0E+00 | 0,0E+00 | 3992024 | 3992977 | 954 | + |
| Y11_RS18250 | Y11_38371 | mog | molybdopterin adenylyltransferase | -1,19 | 0,65 | -0,07 | 1,84 | 1,12 | -0,72 | 2,8E-04 | 5,0E-05 | 5,3E-01 | 1,1E-05 | 5,6E-04 | 1,0E-04 | 1,1E-05 | 0,0E+00 | 0,0E+00 | 2,3E-07 | 1,3E-05 | 0,0E+00 | 3993135 | 3993719 | 585 | + |
| Y11_RS18265 | Y11_38401 | satP | acetate uptake transporter | 0,08 | -4,17 | -4,93 | -4,25 | -5 | -0,76 | 8,1E-01 | 3,6E-09 | 9,8E-10 | 1,4E-06 | 4,1E-07 | 5,3E-03 | 0,0E+00 | 2,8E-10 | 1,0E-10 | 3,5E-08 | 2,0E-08 | 0,0E+00 | 3998000 | 3998593 | 594 | - |
| Y11_RS18270 | Y11_38411 | dnaK | molecular chaperone DnaK | -0,03 | 0,55 | 0,43 | 0,57 | 0,46 | -0,12 | 6,3E-01 | 1,2E-07 | 3,3E-03 | 2,9E-06 | 3,4E-03 | 3,0E-01 | 0,0E+00 | 0,0E+00 | 0,0E+00 | 0,0E+00 | 0,0E+00 | 0,0E+00 | 3998966 | 4000873 | 1908 | + |
| Y11_RS18275 | Y11_38421 | dnaJ | molecular chaperone DnaJ | -0,23 | -0,74 | -1,3 | -0,51 | -1,07 | -0,57 | 8,0E-03 | 3,0E-07 | 2,4E-06 | 1,4E-05 | 1,3E-05 | 6,2E-04 | 0,0E+00 | 0,0E+00 | 4,9E-08 | 0,0E+00 | 4,2E-07 | 0,0E+00 | 4000985 | 4002115 | 1131 | + |
| Y11_RS18280 | Y11_38431 | nhaA | Na+/H+ antiporter NhaA | 0,45 | -0,07 | 0,12 | -0,52 | -0,33 | 0,19 | 3,3E-01 | 8,4E-01 | 7,9E-01 | 2,7E-01 | 5,4E-01 | 6,8E-01 | 0,0E+00 | 0,0E+00 | 0,0E+00 | 0,0E+00 | 0,0E+00 | 0,0E+00 | 4002347 | 4003525 | 1179 | + |

| | | | | | | | | | | | | | | | | | | | | | | | | | |
|---|---|---|---|---|---|---|---|---|---|---|---|---|---|---|---|---|---|---|---|---|---|---|---|---|---|
| Y11_RS18285 | Y11_38441 | nhaR | transcriptional activator NhaR | -0,26 | -0,8 | -0,6 | -0,53 | -0,34 | 0,19 | 5,8E-03 | 1,7E-05 | 5,1E-05 | 4,3E-05 | 1,8E-04 | 2,8E-02 | 0,0E+00 | 0,0E+00 | 0,0E+00 | 0,0E+00 | 0,0E+00 | 0,0E+00 | 4003676 | 4004575 | 900 | + |
| Y11_RS18290 | Y11_38451 | rpsT | 30S ribosomal protein S20 | 0,39 | -0,27 | -0,7 | -0,66 | -1,08 | -0,43 | 3,0E-03 | 1,5E-02 | 6,4E-03 | 2,9E-05 | 4,0E-04 | 5,0E-02 | 0,0E+00 | 0,0E+00 | 0,0E+00 | 0,0E+00 | 9,5E-06 | 0,0E+00 | 4004719 | 4004982 | 264 | - |
| Y11_RS18300 | Y11_38461 | ribF | bifunctional riboflavin kinase/FAD synthetase | -0,1 | -0,6 | -0,24 | -0,5 | -0,14 | 0,36 | 2,5E-01 | 6,3E-06 | 8,1E-03 | 3,1E-04 | 1,7E-01 | 7,1E-04 | 0,0E+00 | 0,0E+00 | 0,0E+00 | 0,0E+00 | 0,0E+00 | 0,0E+00 | 4005396 | 4006334 | 939 | + |
| Y11_RS18305 | Y11_38471 | ileS | isoleucine--tRNA ligase | -0,02 | -0,51 | -0,58 | -0,49 | -0,56 | -0,07 | 5,4E-01 | 7,2E-07 | 4,8E-06 | 1,8E-06 | 8,6E-06 | 2,6E-01 | 0,0E+00 | 0,0E+00 | 0,0E+00 | 0,0E+00 | 0,0E+00 | 0,0E+00 | 4006366 | 4009182 | 2817 | + |
| Y11_RS18315 | Y11_38491 | fkpB | FKBP-type peptidyl-prolyl cis-trans isomerase | 0 | -0,78 | -1,52 | -15 | -15 | -0,74 | 0,0E+00 | 1,2E-03 | 7,1E-05 | 0,0E+00 | 0,0E+00 | 1,4E-02 | 0,0E+00 | 0,0E+00 | 1,0E-06 | 0,0E+00 | 0,0E+00 | 0,0E+00 | 4009693 | 4010193 | 501 | + |
| Y11_RS18320 | Y11_38501 | ispH | 4-hydroxy-3-methylbut-2-enyl diphosphate reductase | 0,92 | -1,63 | -2,25 | -2,55 | -3,17 | -0,62 | 2,2E-06 | 3,8E-09 | 8,1E-09 | 2,0E-10 | 7,1E-10 | 6,8E-05 | 0,0E+00 | 2,9E-10 | 4,6E-10 | 2,7E-11 | 1,4E-10 | 0,0E+00 | 4010174 | 4011127 | 954 | + |
| Y11_RS18325 | Y11_38511 | dapB | 4-hydroxy-tetrahydrodipicolinate reductase | 0,08 | -0,79 | -0,58 | -0,87 | -0,66 | 0,21 | 4,5E-01 | 5,9E-05 | 1,3E-03 | 7,2E-06 | 2,4E-04 | 8,4E-02 | 0,0E+00 | 0,0E+00 | 0,0E+00 | 0,0E+00 | 0,0E+00 | 0,0E+00 | 4011646 | 4012467 | 822 | + |
| Y11_RS18330 | Y11_38521 | carA | glutamine-hydrolyzing carbamoyl-phosphate synthase small subunit | 0,28 | 2,68 | 3,78 | 2,4 | 3,5 | 1,1 | 7,5E-04 | 3,4E-10 | 1,1E-09 | 2,1E-09 | 3,0E-09 | 3,8E-05 | 0,0E+00 | 5,1E-11 | 1,1E-10 | 1,5E-10 | 3,7E-10 | 3,5E-06 | 4012938 | 4014101 | 1164 | + |
| Y11_RS18335 | Y11_38531 | carB | carbamoyl-phosphate synthase large subunit | 0,39 | 2,31 | 3,08 | 1,92 | 2,69 | 0,77 | 2,4E-05 | 2,6E-08 | 8,3E-12 | 1,8E-07 | 1,4E-10 | 2,0E-04 | 0,0E+00 | 1,3E-09 | 6,7E-12 | 5,4E-09 | 5,9E-11 | 0,0E+00 | 4014118 | 4017351 | 3234 | + |
| Y11_RS18355 | Y11_38571 | folA | type 3 dihydrofolate reductase | 0,31 | 0,75 | 0,53 | 0,44 | 0,22 | -0,22 | 2,5E-03 | 1,5E-04 | 1,4E-04 | 4,2E-03 | 2,2E-02 | 9,5E-02 | 0,0E+00 | 0,0E+00 | 0,0E+00 | 0,0E+00 | 0,0E+00 | 0,0E+00 | 4019864 | 4020346 | 483 | + |
| Y11_RS18360 | Y11_38581 | apaH | bis(5'-nucleosyl)-tetraphosphatase (symmetrical) ApaH | 0,54 | 1,09 | 0,94 | 0,56 | 0,4 | -0,16 | 1,9E-05 | 3,8E-09 | 1,6E-04 | 5,2E-05 | 3,0E-02 | 3,2E-01 | 0,0E+00 | 2,9E-10 | 0,0E+00 | 0,0E+00 | 0,0E+00 | 0,0E+00 | 4020540 | 4021418 | 879 | - |
| Y11_RS18365 | Y11_38591 | apaG | Co2+/Mg2+ efflux protein ApaG | 0,16 | 0,77 | 1,37 | 0,61 | 1,21 | 0,6 | 1,2E-01 | 3,3E-05 | 9,7E-06 | 7,5E-05 | 1,7E-05 | 1,8E-03 | 0,0E+00 | 0,0E+00 | 1,7E-07 | 0,0E+00 | 5,2E-07 | 0,0E+00 | 4021433 | 4021810 | 378 | - |
| Y11_RS18370 | Y11_38601 | rsmA | 16S rRNA (adenine(1518)-N(6)/adenine(1519)-N(6))-dimethyltransferase RsmA | -0,28 | -0,5 | -0,85 | -0,22 | -0,57 | -0,35 | 3,4E-02 | 1,7E-03 | 4,7E-05 | 3,8E-02 | 2,0E-04 | 3,7E-03 | 0,0E+00 | 0,0E+00 | 0,0E+00 | 0,0E+00 | 0,0E+00 | 0,0E+00 | 4021824 | 4022642 | 819 | - |
| Y11_RS18375 | Y11_38611 | pdxA | 4-hydroxythreonine-4-phosphate dehydrogenase PdxA | 0,02 | 0,06 | 0,07 | 0,04 | 0,05 | 0,01 | 8,1E-01 | 3,8E-01 | 2,7E-01 | 6,0E-01 | 4,9E-01 | 9,0E-01 | 0,0E+00 | 0,0E+00 | 0,0E+00 | 0,0E+00 | 0,0E+00 | 0,0E+00 | 4022635 | 4023630 | 996 | - |
| Y11_RS18380 | Y11_38621 | surA | peptidylprolyl isomerase SurA | -0,01 | -0,08 | 0,23 | -0,07 | 0,24 | 0,31 | 6,9E-01 | 1,3E-01 | 2,5E-02 | 1,7E-01 | 1,9E-02 | 9,8E-03 | 0,0E+00 | 0,0E+00 | 0,0E+00 | 0,0E+00 | 0,0E+00 | 0,0E+00 | 4023614 | 4024918 | 1305 | - |
| Y11_RS18385 | Y11_38631 | lptD | LPS assembly protein LptD | 0 | -0,37 | -0,49 | -0,37 | -0,5 | -0,12 | 9,4E-01 | 9,5E-06 | 1,0E-03 | 8,9E-05 | 1,4E-03 | 2,4E-01 | 0,0E+00 | 0,0E+00 | 0,0E+00 | 0,0E+00 | 0,0E+00 | 0,0E+00 | 4024986 | 4027352 | 2367 | - |
| Y11_RS18390 | Y11_38641 | djlA | co-chaperone DjlA | 0,31 | -0,61 | -0,34 | -0,92 | -0,65 | 0,27 | 1,2E-01 | 1,0E-03 | 4,1E-02 | 1,9E-04 | 3,3E-03 | 8,8E-03 | 0,0E+00 | 0,0E+00 | 0,0E+00 | 0,0E+00 | 0,0E+00 | 0,0E+00 | 4027537 | 4028364 | 828 | + |
| Y11_RS18395 | Y11_38651 | rluA | bifunctional tRNA pseudouridine(32) synthase/23S rRNA pseudouridine(746) synthase RluA | 0,24 | -1,87 | -3,06 | -2,11 | -3,3 | -1,19 | 1,5E-01 | 2,1E-10 | 1,1E-06 | 4,2E-07 | 1,9E-06 | 8,7E-04 | 0,0E+00 | 3,7E-11 | 2,5E-08 | 1,2E-08 | 7,2E-08 | 5,9E-05 | 4028450 | 4029070 | 621 | - |
| Y11_RS18405 | Y11_38671 | rapA | RNA polymerase-associated protein RapA | 0,05 | -0,41 | -0,95 | -0,46 | -0,99 | -0,54 | 1,4E-01 | 1,8E-06 | 6,6E-07 | 1,6E-06 | 5,6E-07 | 6,8E-05 | 0,0E+00 | 0,0E+00 | 0,0E+00 | 0,0E+00 | 0,0E+00 | 0,0E+00 | 4033608 | 4036514 | 2907 | - |
| Y11_RS18410 | Y11_38681 | | DNA polymerase II | 0,2 | -0,24 | -0,37 | -0,44 | -0,56 | -0,12 | 1,5E-01 | 1,1E-01 | 1,2E-01 | 2,3E-03 | 1,7E-02 | 5,5E-01 | 0,0E+00 | 0,0E+00 | 0,0E+00 | 0,0E+00 | 0,0E+00 | 0,0E+00 | 4036783 | 4039155 | 2373 | - |
| Y11_RS18420 | Y11_38701 | thiQ | thiamine ABC transporter ATP-binding protein ThiQ | -0,36 | -0,05 | -0,08 | 0,31 | 0,28 | -0,03 | 4,5E-02 | 7,0E-01 | 6,2E-01 | 8,0E-02 | 1,5E-01 | 8,6E-01 | 0,0E+00 | 0,0E+00 | 0,0E+00 | 0,0E+00 | 0,0E+00 | 0,0E+00 | 4040247 | 4040948 | 702 | - |
| Y11_RS18430 | Y11_38721 | thiB | thiamine ABC transporter substrate binding subunit | -0,9 | 0,62 | 0,79 | 1,53 | 1,69 | 0,16 | 6,9E-07 | 2,0E-04 | 2,5E-04 | 6,4E-07 | 1,6E-06 | 3,2E-01 | 0,0E+00 | 0,0E+00 | 0,0E+00 | 1,7E-08 | 6,3E-08 | 0,0E+00 | 4042518 | 4043507 | 990 | - |
| Y11_RS18445 | Y11_38751 | sgrR | HTH-type transcriptional regulator SgrR | -0,33 | -0,04 | 0,66 | 0,28 | 0,99 | 0,69 | 1,7E-01 | 8,9E-01 | 1,3E-02 | 2,7E-01 | 1,6E-05 | 1,7E-02 | 0,0E+00 | 0,0E+00 | 0,0E+00 | 0,0E+00 | 0,0E+00 | 0,0E+00 | 4045263 | 4046924 | 1662 | - |
| Y11_RS18480 | Y11_38831 | leuD | 3-isopropylmalate dehydratase small subunit | 0 | -1,38 | -3,75 | -15 | -15 | -2,37 | 0,0E+00 | 3,0E-06 | 4,6E-10 | 0,0E+00 | 0,0E+00 | 3,3E-08 | 0,0E+00 | 7,8E-08 | 6,1E-11 | 0,0E+00 | 0,0E+00 | 1,5E-08 | 4055443 | 4056063 | 621 | - |
| Y11_RS18485 | Y11_38841 | leuC | 3-isopropylmalate dehydratase large subunit | -0,03 | -1,53 | -3,18 | -1,51 | -3,16 | -1,65 | 7,6E-01 | 6,1E-06 | 1,7E-10 | 6,7E-06 | 1,7E-10 | 3,3E-06 | 0,0E+00 | 1,5E-07 | 3,0E-11 | 1,4E-07 | 6,2E-11 | 4,7E-07 | 4056078 | 4057490 | 1413 | - |
| Y11_RS18490 | Y11_38851 | leuB | 3-isopropylmalate dehydrogenase | 0,11 | -2,79 | -3,87 | -2,9 | -3,98 | -1,08 | 1,7E-01 | 4,2E-10 | 3,9E-11 | 2,6E-10 | 2,7E-11 | 1,1E-06 | 0,0E+00 | 5,8E-11 | 1,4E-11 | 3,2E-11 | 2,0E-11 | 1,9E-07 | 4057492 | 4058583 | 1092 | - |
| Y11_RS18495 | Y11_38861 | leuA | 2-isopropylmalate synthase | 0,1 | -2,14 | -3,38 | -2,24 | -3,48 | -1,24 | 2,9E-01 | 1,4E-09 | 2,0E-11 | 2,0E-08 | 4,6E-10 | 3,2E-07 | 0,0E+00 | 1,4E-10 | 9,4E-12 | 8,8E-10 | 1,1E-10 | 7,7E-08 | 4058586 | 4060148 | 1563 | - |
| Y11_RS18505 | Y11_38891 | | long-chain fatty acid--CoA ligase | -0,63 | -2,72 | -0,46 | -2,1 | 0,17 | 2,27 | 1,5E-03 | 3,6E-06 | 4,1E-02 | 1,2E-05 | 3,2E-01 | 2,1E-05 | 0,0E+00 | 9,5E-08 | 0,0E+00 | 2,5E-07 | 0,0E+00 | 2,2E-06 | 4062608 | 4064407 | 1800 | + |
| Y11_RS18510 | Y11_38911 | ilvI | acetolactate synthase 3 large subunit | 0,43 | -0,79 | -0,21 | -1,22 | -0,64 | 0,58 | 9,9E-06 | 5,7E-07 | 6,2E-04 | 2,8E-08 | 3,8E-07 | 5,3E-06 | 0,0E+00 | 0,0E+00 | 0,0E+00 | 1,1E-09 | 0,0E+00 | 0,0E+00 | 4064856 | 4066583 | 1728 | + |
| Y11_RS18515 | Y11_38921 | ilvN | acetolactate synthase small subunit | 0,32 | -1,16 | -0,63 | -1,48 | -0,95 | 0,53 | 1,8E-03 | 1,2E-07 | 8,1E-05 | 3,6E-08 | 6,1E-06 | 3,0E-04 | 0,0E+00 | 4,6E-09 | 0,0E+00 | 1,4E-09 | 0,0E+00 | 0,0E+00 | 4066586 | 4067080 | 495 | + |
| Y11_RS18520 | Y11_38931 | cra | catabolite repressor/activator | 0,39 | -0,22 | -0,27 | -0,61 | -0,67 | -0,06 | 1,4E-03 | 5,0E-02 | 8,9E-03 | 6,6E-04 | 1,8E-04 | 6,2E-01 | 0,0E+00 | 0,0E+00 | 0,0E+00 | 0,0E+00 | 0,0E+00 | 0,0E+00 | 4067426 | 4068430 | 1005 | + |
| Y11_RS18535 | Y11_38961 | mraZ | division/cell wall cluster transcriptional repressor MraZ | -0,11 | 0,99 | 1,35 | 1,1 | 1,45 | 0,36 | 2,7E-01 | 2,2E-06 | 1,0E-07 | 8,6E-06 | 6,9E-07 | 6,3E-03 | 0,0E+00 | 0,0E+00 | 3,3E-09 | 1,8E-07 | 3,0E-08 | 0,0E+00 | 4070038 | 4070496 | 459 | + |
| Y11_RS18540 | Y11_38971 | rsmH | 16S rRNA (cytosine(1402)-N(4))-methyltransferase RsmH | 0 | -1,22 | -2,85 | -1,22 | -2,85 | -1,63 | 9,8E-01 | 2,3E-05 | 1,4E-10 | 4,8E-05 | 2,0E-09 | 4,3E-06 | 0,0E+00 | 5,2E-07 | 2,7E-11 | 9,2E-07 | 2,8E-10 | 5,8E-07 | 4070499 | 4071461 | 963 | + |
| Y11_RS18545 | Y11_38981 | ftsL | cell division protein FtsL | -0,05 | -0,45 | -1,27 | -0,4 | -1,22 | -0,82 | 8,4E-01 | 2,4E-02 | 7,2E-03 | 9,9E-02 | 1,3E-02 | 4,3E-02 | 0,0E+00 | 0,0E+00 | 8,2E-05 | 0,0E+00 | 2,7E-04 | 0,0E+00 | 4071458 | 4071778 | 321 | + |
| Y11_RS18550 | Y11_38991 | | peptidoglycan glycosyltransferase FtsI | 0,23 | 0,53 | 0,04 | 0,3 | -0,19 | -0,49 | 6,4E-03 | 4,4E-05 | 7,4E-01 | 9,2E-06 | 6,4E-02 | 7,0E-04 | 0,0E+00 | 0,0E+00 | 0,0E+00 | 0,0E+00 | 0,0E+00 | 0,0E+00 | 4071839 | 4073602 | 1764 | + |
| Y11_RS18555 | Y11_39001 | murE | UDP-N-acetylmuramoyl-L-alanyl-D-glutamate--2,6-diaminopimelate ligase | -0,23 | 0,41 | 0,23 | 0,64 | 0,46 | -0,18 | 5,4E-04 | 9,0E-05 | 5,4E-03 | 4,2E-06 | 7,8E-05 | 4,1E-02 | 0,0E+00 | 0,0E+00 | 0,0E+00 | 0,0E+00 | 0,0E+00 | 0,0E+00 | 4073589 | 4075079 | 1491 | + |
| Y11_RS18560 | Y11_39011 | murF | UDP-N-acetylmuramoyl-tripeptide--D-alanyl-D-alanine ligase | -0,2 | 0,65 | 0,65 | 0,85 | 0,84 | 0 | 4,3E-02 | 6,7E-06 | 1,3E-03 | 5,4E-06 | 3,5E-04 | 1,0E+00 | 0,0E+00 | 0,0E+00 | 0,0E+00 | 0,0E+00 | 0,0E+00 | 0,0E+00 | 4075072 | 4076436 | 1365 | + |
| Y11_RS18565 | Y11_39021 | mraY | phospho-N-acetylmuramoyl-pentapeptide-transferase | 0,88 | 0,76 | 2,65 | -0,12 | 1,77 | 1,89 | 1,2E-01 | 2,8E-03 | 1,9E-03 | 8,2E-01 | 4,8E-02 | 1,3E-02 | 0,0E+00 | 0,0E+00 | 2,2E-05 | 0,0E+00 | 1,0E-03 | 7,4E-04 | 4076430 | 4077512 | 1083 | + |
| Y11_RS18570 | Y11_39031 | murD | UDP-N-acetylmuramoyl-L-alanine--D-glutamate ligase | 0,09 | -0,35 | -1,21 | -0,45 | -1,31 | -0,86 | 1,2E-01 | 2,6E-05 | 6,7E-06 | 3,7E-05 | 5,3E-06 | 8,3E-05 | 0,0E+00 | 0,0E+00 | 1,2E-07 | 0,0E+00 | 1,9E-07 | 0,0E+00 | 4077515 | 4078849 | 1335 | + |
| Y11_RS18580 | Y11_39051 | murG | undecaprenyldiphospho-muramoylpentapeptide beta-N-acetylglucosaminyltransferase | -0,13 | -0,28 | -0,4 | -0,14 | -0,26 | -0,12 | 2,7E-02 | 1,2E-02 | 8,9E-04 | 8,4E-02 | 2,6E-03 | 2,3E-01 | 0,0E+00 | 0,0E+00 | 0,0E+00 | 0,0E+00 | 0,0E+00 | 0,0E+00 | 4080048 | 4081115 | 1068 | + |
| Y11_RS18585 | Y11_39061 | murC | UDP-N-acetylmuramate--L-alanine ligase | -0,29 | -0,5 | -0,79 | -0,21 | -0,5 | -0,29 | 9,1E-03 | 3,0E-05 | 1,7E-04 | 2,4E-02 | 4,6E-03 | 3,3E-02 | 0,0E+00 | 0,0E+00 | 0,0E+00 | 0,0E+00 | 0,0E+00 | 0,0E+00 | 4081189 | 4082664 | 1476 | + |
| Y11_RS18590 | Y11_39071 | | D-alanine--D-alanine ligase | -0,18 | -0,19 | -0,32 | -0,01 | -0,14 | -0,12 | 1,5E-02 | 1,3E-02 | 6,2E-04 | 8,7E-01 | 9,4E-02 | 1,3E-01 | 0,0E+00 | 0,0E+00 | 0,0E+00 | 0,0E+00 | 0,0E+00 | 0,0E+00 | 4082657 | 4083577 | 921 | + |
| Y11_RS18595 | Y11_39081 | ftsQ | cell division protein FtsQ | -0,02 | -0,31 | -0,78 | -0,29 | -0,76 | -0,47 | 8,8E-01 | 4,0E-02 | 7,5E-04 | 7,2E-02 | 1,4E-03 | 1,1E-02 | 0,0E+00 | 0,0E+00 | 0,0E+00 | 0,0E+00 | 0,0E+00 | 0,0E+00 | 4083579 | 4084436 | 858 | + |
| Y11_RS18600 | Y11_39091 | ftsA | cell division protein FtsA | -0,45 | -0,04 | 0,09 | 0,41 | 0,54 | 0,13 | 4,1E-05 | 6,2E-01 | 2,0E-01 | 2,3E-04 | 1,6E-05 | 1,2E-01 | 0,0E+00 | 0,0E+00 | 0,0E+00 | 0,0E+00 | 0,0E+00 | 0,0E+00 | 4084433 | 4085689 | 1257 | + |
| Y11_RS18605 | Y11_39101 | ftsZ | cell division protein FtsZ | -0,38 | -0,91 | -1,14 | -0,53 | -0,76 | -0,23 | 3,1E-05 | 4,2E-08 | 2,6E-06 | 6,5E-07 | 4,0E-05 | 4,3E-02 | 0,0E+00 | 0,0E+00 | 5,1E-08 | 0,0E+00 | 0,0E+00 | 0,0E+00 | 4085762 | 4086913 | 1152 | + |
| Y11_RS18610 | Y11_39111 | lpxC | UDP-3-O-acyl-N-acetylglucosamine deacetylase | -0,54 | 1,4 | 1,82 | 1,94 | 2,37 | 0,43 | 2,0E-06 | 3,4E-08 | 7,8E-07 | 1,9E-09 | 9,6E-08 | 1,7E-02 | 0,0E+00 | 1,6E-09 | 1,9E-08 | 1,4E-10 | 6,0E-09 | 0,0E+00 | 4087014 | 4087934 | 921 | + |
| Y11_RS18615 | Y11_39121 | | DUF721 domain-containing protein | -0,02 | 1,19 | 1,04 | 1,22 | 1,06 | -0,16 | 8,9E-01 | 1,0E-04 | 2,6E-03 | 1,1E-04 | 2,5E-03 | 5,5E-01 | 0,0E+00 | 2,2E-06 | 3,0E-05 | 2,0E-06 | 5,5E-05 | 0,0E+00 | 4088070 | 4088594 | 525 | - |
| Y11_RS18625 | Y11_39141 | secA | preprotein translocase subunit SecA | 0,11 | 0,7 | 0,85 | 0,59 | 0,74 | 0,15 | 2,3E-02 | 1,2E-07 | 4,0E-06 | 9,8E-07 | 1,4E-05 | 9,5E-02 | 0,0E+00 | 0,0E+00 | 0,0E+00 | 0,0E+00 | 0,0E+00 | 0,0E+00 | 4089234 | 4091948 | 2715 | + |
| Y11_RS18630 | Y11_39151 | mutT | 8-oxo-dGTP diphosphatase MutT | 0 | -0,42 | -1,42 | -0,94 | -1,96 | -1,01 | 0,0E+00 | 4,2E-02 | 7,0E-06 | 5,4E-03 | 2,8E-05 | 1,8E-03 | 0,0E+00 | 0,0E+00 | 1,2E-07 | 0,0E+00 | 8,2E-07 | 1,1E-04 | 4092082 | 4092468 | 387 | + |
| Y11_RS18635 | Y11_39161 | yacG | DNA gyrase inhibitor YacG | -0,05 | 0,57 | 0,47 | 0,63 | 0,53 | -0,1 | 7,8E-01 | 2,3E-02 | 4,2E-02 | 2,9E-03 | 4,7E-03 | 5,3E-01 | 0,0E+00 | 0,0E+00 | 0,0E+00 | 0,0E+00 | 0,0E+00 | 0,0E+00 | 4092539 | 4092745 | 207 | - |
| Y11_RS18640 | Y11_39171 | zapD | cell division protein ZapD | 0,39 | 0,06 | -0,56 | -0,33 | -0,95 | -0,62 | 4,6E-04 | 7,0E-01 | 4,9E-03 | 4,7E-02 | 1,1E-04 | 1,2E-02 | 0,0E+00 | 0,0E+00 | 0,0E+00 | 0,0E+00 | 0,0E+00 | 0,0E+00 | 4092966 | 4093718 | 753 | - |
| Y11_RS18645 | Y11_39181 | coaE | dephospho-CoA kinase | -0,2 | 0,21 | 0,47 | 0,41 | 0,67 | 0,25 | 3,2E-03 | 1,5E-02 | 2,9E-02 | 7,6E-04 | 5,7E-03 | 2,1E-01 | 0,0E+00 | 0,0E+00 | 0,0E+00 | 0,0E+00 | 0,0E+00 | 0,0E+00 | 4093715 | 4094335 | 621 | - |
| Y11_RS18650 | Y11_39191 | | GMP reductase | 1,33 | 2,25 | 2,72 | 0,92 | 1,39 | 0,47 | 1,7E-08 | 1,9E-11 | 1,1E-10 | 9,9E-08 | 4,6E-08 | 4,1E-05 | 1,9E-09 | 7,7E-12 | 2,5E-11 | 0,0E+00 | 3,3E-09 | 0,0E+00 | 4094629 | 4095672 | 1044 | + |
| Y11_RS18670 | Y11_39231 | nadC | carboxylating nicotinate-nucleotide diphosphorylase | 0,47 | 0,58 | 0,78 | 0,11 | 0,31 | 0,2 | 4,2E-05 | 1,1E-06 | 2,2E-05 | 1,8E-01 | 2,0E-02 | 7,6E-02 | 0,0E+00 | 0,0E+00 | 0,0E+00 | 0,0E+00 | 0,0E+00 | 0,0E+00 | 4099114 | 4100022 | 909 | - |
| Y11_RS18675 | Y11_39241 | ampD | 1,6-anhydro-N-acetylmuramyl-L-alanine amidase AmpD | 0 | -0,06 | 0,19 | -0,15 | 0,05 | 0,24 | 0,0E+00 | 7,3E-01 | 4,1E-01 | 5,1E-01 | 8,6E-01 | 3,6E-01 | 0,0E+00 | 0,0E+00 | 0,0E+00 | 0,0E+00 | 0,0E+00 | 0,0E+00 | 4100141 | 4100716 | 576 | + |
| Y11_RS18680 | Y11_39251 | ampE | beta-lactamase regulator AmpE | -0,49 | 0,82 | 2,58 | 1,31 | 3,06 | 1,75 | 7,4E-04 | 3,2E-04 | 1,1E-08 | 5,0E-06 | 5,8E-10 | 1,2E-06 | 0,0E+00 | 0,0E+00 | 5,5E-10 | 1,1E-07 | 1,2E-10 | 2,1E-07 | 4100860 | 4101714 | 855 | + |
| Y11_RS18685 | Y11_39261 | | amino acid permease | -0,52 | 1,86 | 1,99 | 2,37 | 2,5 | 0,13 | 2,0E-02 | 1,0E-06 | 2,9E-05 | 5,3E-06 | 2,6E-05 | 6,5E-01 | 0,0E+00 | 3,1E-08 | 4,4E-07 | 1,1E-07 | 7,6E-07 | 0,0E+00 | 4101872 | 4103236 | 1365 | - |
| Y11_RS18690 | Y11_39291 | pdhR | pyruvate dehydrogenase complex transcriptional repressor PdhR | -0,03 | -0,56 | -1,08 | -0,53 | -1,05 | -0,52 | 8,0E-01 | 5,3E-04 | 1,3E-06 | 4,0E-03 | 2,4E-05 | 7,1E-04 | 0,0E+00 | 0,0E+00 | 2,8E-08 | 0,0E+00 | 7,1E-07 | 0,0E+00 | 4103937 | 4104701 | 765 | + |
| Y11_RS18695 | Y11_39301 | aceE | pyruvate dehydrogenase (acetyl-transferring), homodimeric type | -0,1 | -0,92 | -1,67 | -0,82 | -1,58 | -0,75 | 3,0E-02 | 7,1E-08 | 6,6E-09 | 1,9E-08 | 3,6E-09 | 3,4E-06 | 0,0E+00 | 0,0E+00 | 3,9E-10 | 0,0E+00 | 4,4E-10 | 0,0E+00 | 4104878 | 4107541 | 2664 | + |
| Y11_RS18700 | Y11_39311 | aceF | pyruvate dehydrogenase complex dihydrolipoyllysine-residue acetyltransferase | -0,07 | -0,51 | -1,13 | -0,44 | -1,06 | -0,62 | 2,3E-01 | 2,5E-04 | 2,8E-07 | 1,0E-03 | 9,3E-07 | 2,7E-04 | 0,0E+00 | 0,0E+00 | 7,8E-09 | 0,0E+00 | 3,9E-08 | 0,0E+00 | 4107555 | 4109417 | 1863 | + |
| Y11_RS18705 | Y11_39321 | lpdA | dihydrolipoyl dehydrogenase | -0,11 | -0,19 | -0,32 | -0,08 | -0,21 | -0,13 | 2,2E-01 | 5,5E-02 | 2,3E-02 | 4,9E-01 | 1,6E-01 | 3,5E-01 | 0,0E+00 | 0,0E+00 | 0,0E+00 | 0,0E+00 | 0,0E+00 | 0,0E+00 | 4109905 | 4111329 | 1425 | + |
| Y11_RS18715 | Y11_39341 | | AraC family transcriptional regulator | 0,17 | 0,34 | 0,27 | 0,17 | 0,09 | -0,07 | 4,0E-01 | 1,4E-01 | 4,4E-01 | 4,5E-01 | 7,8E-01 | 8,3E-01 | 0,0E+00 | 0,0E+00 | 0,0E+00 | 0,0E+00 | 0,0E+00 | 0,0E+00 | 4112965 | 4113972 | 1008 | - |
| Y11_RS18725 | Y11_39361 | acnB | bifunctional aconitate hydratase 2/2-methylisocitrate dehydratase | 0,2 | 0,07 | 0,88 | -0,13 | 0,68 | 0,81 | 5,2E-04 | 1,1E-01 | 1,4E-06 | 7,8E-04 | 4,3E-06 | 1,5E-06 | 0,0E+00 | 0,0E+00 | 0,0E+00 | 0,0E+00 | 0,0E+00 | 0,0E+00 | 4117065 | 4119662 | 2598 | + |
| Y11_RS18730 | Y11_39371 | yacL | protein YacL | -0,04 | 0,71 | 0,6 | 0,75 | 0,64 | -0,1 | 6,3E-01 | 4,7E-04 | 2,4E-03 | 1,6E-04 | 9,6E-04 | 5,2E-01 | 0,0E+00 | 0,0E+00 | 0,0E+00 | 0,0E+00 | 0,0E+00 | 0,0E+00 | 4119864 | 4120226 | 363 | + |
| Y11_RS18745 | Y11_39411 | speD | adenosylmethionine decarboxylase | -0,29 | 0,54 | 0,26 | 0,83 | 0,54 | -0,29 | 2,3E-03 | 1,5E-05 | 1,6E-02 | 1,6E-07 | 1,1E-04 | 4,1E-03 | 0,0E+00 | 0,0E+00 | 0,0E+00 | 0,0E+00 | 0,0E+00 | 0,0E+00 | 4121984 | 4122778 | 795 | - |
| Y11_RS18750 | Y11_39421 | speE | polyamine aminopropyltransferase | -0,14 | 0,41 | 0,48 | 0,55 | 0,62 | 0,08 | 1,3E-01 | 1,5E-03 | 1,2E-04 | 8,9E-05 | 3,7E-06 | 2,5E-01 | 0,0E+00 | 0,0E+00 | 0,0E+00 | 0,0E+00 | 0,0E+00 | 0,0E+00 | 4122803 | 4123666 | 864 | - |
| Y11_RS18760 | Y11_39441 | cueO | multicopper oxidase CueO | 0,19 | 1,62 | 2,29 | 1,42 | 2,1 | 0,68 | 3,6E-02 | 1,1E-07 | 2,8E-10 | 2,1E-07 | 2,6E-10 | 1,7E-05 | 0,0E+00 | 4,4E-09 | 4,2E-11 | 6,4E-09 | 7,8E-11 | 0,0E+00 | 4124359 | 4125951 | 1593 | + |
| Y11_RS18765 | Y11_39451 | hpt | hypoxanthine phosphoribosyltransferase | -0,33 | -0,31 | -0,51 | 0,02 | -0,18 | -0,19 | 3,6E-05 | 5,7E-04 | 2,1E-02 | 8,1E-01 | 3,5E-01 | 3,2E-01 | 0,0E+00 | 0,0E+00 | 0,0E+00 | 0,0E+00 | 0,0E+00 | 0,0E+00 | 4126067 | 4126603 | 537 | + |
| Y11_RS18770 | Y11_39461 | can | carbonate dehydratase | 0,4 | 0,72 | 0,32 | 0,31 | -0,08 | -0,4 | 5,7E-04 | 2,5E-05 | 4,0E-02 | 9,7E-03 | 5,5E-01 | 2,4E-02 | 0,0E+00 | 0,0E+00 | 0,0E+00 | 0,0E+00 | 0,0E+00 | 0,0E+00 | 4126692 | 4127354 | 663 | - |

| | | | | | | | | | | | | | | | | | | | | | | | | |
|---|---|---|---|---|---|---|---|---|---|---|---|---|---|---|---|---|---|---|---|---|---|---|---|---|
| Y11_RS18775 | Y11_39471 | | ABC transporter ATP-binding protein | -0,23 | -0,47 | -1,11 | -0,23 | -0,88 | -0,64 | 7,9E-03 | 2,4E-05 | 3,4E-07 | 4,7E-03 | 3,5E-06 | 1,2E-05 | 0,0E+00 | 0,0E+00 | 9,2E-09 | 0,0E+00 | 0,0E+00 | 0,0E+00 | 4127586 | 4128512 | 927 | + |
| Y11_RS18790 | Y11_39511 | panD | aspartate 1-decarboxylase | 1,06 | -0,47 | -1,12 | -1,53 | -2,18 | -0,66 | 2,9E-06 | 4,7E-03 | 5,4E-06 | 3,1E-06 | 8,7E-08 | 1,8E-03 | 1,7E-07 | 0,0E+00 | 9,7E-08 | 6,9E-08 | 5,6E-09 | 0,0E+00 | 4131097 | 4131477 | 381 | - |
| Y11_RS18795 | Y11_39521 | panC | pantoate--beta-alanine ligase | -0,36 | -0,89 | -0,72 | -0,53 | -0,36 | 0,17 | 6,8E-03 | 3,2E-06 | 9,4E-05 | 1,3E-03 | 2,1E-02 | 1,7E-01 | 0,0E+00 | 0,0E+00 | 0,0E+00 | 0,0E+00 | 0,0E+00 | 0,0E+00 | 4131635 | 4132489 | 855 | - |
| Y11_RS18800 | Y11_39531 | panB | 3-methyl-2-oxobutanoate hydroxymethyltransferase | -0,48 | -1,09 | -1,25 | -0,61 | -0,77 | -0,16 | 2,3E-04 | 2,6E-06 | 1,5E-05 | 1,5E-04 | 4,2E-04 | 2,9E-01 | 0,0E+00 | 6,9E-08 | 2,4E-07 | 0,0E+00 | 0,0E+00 | 0,0E+00 | 4132739 | 4133536 | 798 | - |
| Y11_RS18805 | Y11_39541 | folK | 2-amino-4-hydroxy-6-hydroxymethyldihydropteridine diphosphokinase | -0,52 | -0,21 | -0,09 | 0,31 | 0,43 | 0,11 | 2,9E-05 | 4,3E-04 | 4,3E-01 | 9,4E-04 | 8,0E-03 | 3,3E-01 | 0,0E+00 | 0,0E+00 | 0,0E+00 | 0,0E+00 | 0,0E+00 | 0,0E+00 | 4133616 | 4134095 | 480 | - |
| Y11_RS18810 | Y11_39551 | pcnB | polynucleotide adenylyltransferase PcnB | -0,08 | 4,28 | 4,13 | 4,37 | 4,21 | 0 | 4,1E-01 | 8,2E-13 | 6,3E-10 | 7,1E-11 | 2,3E-09 | 0,0E+00 | 0,0E+00 | 1,6E-12 | 7,6E-11 | 1,4E-11 | 3,2E-10 | 0,0E+00 | 4134104 | 4135426 | 1323 | - |
| Y11_RS18815 | Y11_39561 | gluQRS | tRNA glutamyl-Q(34) synthetase GluQRS | -0,58 | 15 | 15 | 15 | 15 | 0 | 2,0E-02 | 0,0E+00 | 0,0E+00 | 0,0E+00 | 0,0E+00 | 0,0E+00 | 0,0E+00 | 0,0E+00 | 0,0E+00 | 0,0E+00 | 0,0E+00 | 0,0E+00 | 4135708 | 4136637 | 930 | - |
| Y11_RS18820 | Y11_39571 | dksA | RNA polymerase-binding protein DksA | -0,03 | 4,47 | 4,42 | 4,51 | 4,46 | -0,05 | 8,4E-01 | 2,8E-09 | 4,3E-10 | 1,2E-08 | 3,9E-09 | 7,7E-01 | 0,0E+00 | 2,3E-10 | 5,8E-11 | 6,1E-10 | 4,6E-10 | 0,0E+00 | 4136779 | 4137234 | 456 | - |
| Y11_RS18825 | Y11_39581 | sfsA | DNA/RNA nuclease SfsA | 0,53 | 0,42 | 0,08 | -0,11 | -0,45 | -0,34 | 2,6E-03 | 2,9E-02 | 4,3E-01 | 5,5E-01 | 1,0E-02 | 7,6E-02 | 0,0E+00 | 0,0E+00 | 0,0E+00 | 0,0E+00 | 0,0E+00 | 0,0E+00 | 4137414 | 4138157 | 744 | - |
| Y11_RS18830 | Y11_39591 | thpR | RNA 2',3'-cyclic phosphodiesterase | 0,6 | 0,75 | 0,19 | 0,15 | -0,42 | -0,56 | 1,8E-02 | 6,6E-03 | 4,4E-01 | 2,7E-01 | 3,0E-02 | 8,0E-03 | 0,0E+00 | 0,0E+00 | 0,0E+00 | 0,0E+00 | 0,0E+00 | 0,0E+00 | 4138169 | 4138726 | 558 | - |
| Y11_RS18835 | Y11_39601 | hrpB | ATP-dependent helicase HrpB | -0,47 | 0,07 | 0,03 | 0,55 | 0,51 | -0,04 | 3,7E-04 | 3,6E-01 | 8,3E-01 | 6,3E-05 | 9,6E-03 | 8,0E-01 | 0,0E+00 | 0,0E+00 | 0,0E+00 | 0,0E+00 | 0,0E+00 | 0,0E+00 | 4138770 | 4141289 | 2520 | + |
| Y11_RS18840 | Y11_39611 | mrcB | bifunctional glycosyl transferase/transpeptidase | 0,13 | -0,89 | -0,72 | -1,01 | -0,85 | 0,17 | 1,9E-02 | 2,2E-08 | 8,0E-07 | 1,4E-09 | 9,1E-08 | 6,4E-03 | 0,0E+00 | 0,0E+00 | 0,0E+00 | 1,1E-10 | 0,0E+00 | 0,0E+00 | 4141593 | 4144097 | 2505 | + |
| Y11_RS18845 | Y11_39621 | fhuC | Fe3+-hydroxamate ABC transporter ATP-binding protein FhuC | 0,96 | 0,33 | -0,74 | -0,63 | -1,71 | -1,07 | 2,5E-02 | 4,2E-01 | 2,3E-01 | 5,9E-02 | 9,5E-03 | 7,8E-02 | 0,0E+00 | 0,0E+00 | 0,0E+00 | 0,0E+00 | 2,0E-04 | 4,1E-03 | 4144354 | 4145148 | 795 | + |
| Y11_RS18850 | Y11_39631 | fhuD | Fe(3+)-hydroxamate ABC transporter substrate-binding protein FhuD | 0 | 0 | -0,42 | 0 | -0,6 | -0,2 | 0,0E+00 | 0,0E+00 | 3,4E-02 | 0,0E+00 | 2,2E-03 | 2,1E-01 | 0,0E+00 | 0,0E+00 | 0,0E+00 | 0,0E+00 | 0,0E+00 | 0,0E+00 | 4145148 | 4146077 | 930 | + |
| Y11_RS18865 | Y11_39661 | hemL | glutamate-1-semialdehyde 2,1-aminomutase | 0,48 | 0,56 | 1,21 | 0,08 | 0,73 | 0,65 | 7,9E-05 | 2,7E-04 | 7,1E-06 | 3,9E-01 | 2,3E-04 | 1,2E-03 | 0,0E+00 | 0,0E+00 | 1,2E-07 | 0,0E+00 | 0,0E+00 | 0,0E+00 | 4150153 | 4151433 | 1281 | - |
| Y11_RS18870 | Y11_39681 | clcA | H(+)/Cl(-) exchange transporter ClcA | 0,6 | 0,86 | -0,01 | 0,26 | -0,61 | -0,87 | 8,1E-03 | 2,8E-05 | 9,1E-01 | 2,1E-01 | 1,8E-02 | 4,5E-04 | 0,0E+00 | 0,0E+00 | 0,0E+00 | 0,0E+00 | 0,0E+00 | 0,0E+00 | 4151700 | 4153121 | 1422 | + |
| Y11_RS18875 | Y11_39691 | erpA | iron-sulfur cluster insertion protein ErpA | 0,64 | -0,18 | 0,01 | -0,82 | -0,64 | 0,19 | 2,5E-03 | 1,8E-01 | 9,7E-01 | 2,1E-04 | 4,4E-03 | 2,2E-01 | 0,0E+00 | 0,0E+00 | 0,0E+00 | 0,0E+00 | 0,0E+00 | 0,0E+00 | 4153251 | 4153595 | 345 | + |
| Y11_RS18885 | Y11_39721 | btuF | vitamin B12 ABC transporter substrate-binding protein BtuF | -0,18 | 1,34 | 0,52 | 1,52 | 0,7 | -0,82 | 3,1E-01 | 3,5E-04 | 1,3E-02 | 5,3E-04 | 1,4E-02 | 1,6E-02 | 0,0E+00 | 7,3E-06 | 0,0E+00 | 9,2E-06 | 0,0E+00 | 0,0E+00 | 4155062 | 4155925 | 864 | - |
| Y11_RS18890 | Y11_39731 | mtnN | 5'-methylthioadenosine/S-adenosylhomocysteine nucleosidase | 0,34 | 1,64 | 1,53 | 1,3 | 1,19 | -0,11 | 2,1E-04 | 1,6E-09 | 1,1E-07 | 2,8E-08 | 1,2E-06 | 2,5E-01 | 0,0E+00 | 1,5E-10 | 3,6E-09 | 1,2E-09 | 4,9E-08 | 0,0E+00 | 4155925 | 4156626 | 702 | - |
| Y11_RS18895 | Y11_39741 | dgt | dGTPase | 0,33 | -0,37 | 0,43 | -0,7 | 0,11 | 0,81 | 3,0E-02 | 5,5E-03 | 3,5E-02 | 5,8E-04 | 5,9E-01 | 1,6E-03 | 0,0E+00 | 0,0E+00 | 0,0E+00 | 0,0E+00 | 0,0E+00 | 0,0E+00 | 4156835 | 4158352 | 1518 | + |
| Y11_RS18900 | Y11_39751 | degP | serine endoprotease DegP | -2,99 | -0,05 | 0,89 | 2,94 | 3,88 | 0,94 | 1,9E-11 | 3,1E-01 | 1,0E-05 | 4,1E-11 | 2,4E-10 | 8,8E-06 | 9,7E-12 | 0,0E+00 | 0,0E+00 | 1,1E-11 | 7,4E-11 | 0,0E+00 | 4158651 | 4160087 | 1437 | + |
| Y11_RS18905 | Y11_39771 | barA | two-component sensor histidine kinase BarA | 0,19 | -0,05 | -0,04 | -0,24 | -0,23 | 0,01 | 2,4E-01 | 7,1E-01 | 7,9E-01 | 1,1E-01 | 1,5E-01 | 9,2E-01 | 0,0E+00 | 0,0E+00 | 0,0E+00 | 0,0E+00 | 0,0E+00 | 0,0E+00 | 4160563 | 4163106 | 2544 | - |
| Y11_RS18915 | Y11_39791 | relA | GTP diphosphokinase | 0,01 | -0,02 | 0,12 | -0,03 | 0,1 | 0,13 | 7,9E-01 | 7,5E-01 | 7,2E-02 | 5,2E-01 | 9,7E-02 | 2,9E-02 | 0,0E+00 | 0,0E+00 | 0,0E+00 | 0,0E+00 | 0,0E+00 | 0,0E+00 | 4164504 | 4166738 | 2235 | + |
| Y11_RS18920 | Y11_39801 | mazG | nucleoside triphosphate pyrophosphohydrolase | -0,24 | -0,62 | -1,24 | -0,38 | -1 | -0,62 | 1,0E-01 | 1,9E-03 | 2,7E-03 | 2,0E-02 | 8,3E-03 | 6,4E-02 | 0,0E+00 | 0,0E+00 | 3,1E-05 | 0,0E+00 | 1,8E-04 | 0,0E+00 | 4167026 | 4167862 | 837 | + |
| Y11_RS18925 | Y11_39811 | pyrG | CTP synthase (glutamine hydrolyzing) | -0,03 | -0,38 | -0,41 | -0,35 | -0,38 | -0,03 | 6,1E-01 | 2,6E-04 | 1,3E-03 | 7,0E-04 | 2,5E-03 | 7,9E-01 | 0,0E+00 | 0,0E+00 | 0,0E+00 | 0,0E+00 | 0,0E+00 | 0,0E+00 | 4168074 | 4169711 | 1638 | + |
| Y11_RS18930 | Y11_39821 | eno | phosphopyruvate hydratase | -0,06 | -0,08 | -1,3 | -0,03 | -1,24 | -1,21 | 3,9E-01 | 4,5E-01 | 1,7E-06 | 8,1E-01 | 2,9E-06 | 2,1E-05 | 0,0E+00 | 0,0E+00 | 3,5E-08 | 0,0E+00 | 1,1E-07 | 2,2E-06 | 4169791 | 4171086 | 1296 | + |
| Y11_RS18940 | Y11_39841 | zapA | cell division protein ZapA | -0,14 | 0,31 | 0,4 | 0,45 | 0,54 | 0,1 | 9,9E-02 | 2,0E-03 | 3,0E-02 | 4,8E-05 | 6,1E-03 | 5,2E-01 | 0,0E+00 | 0,0E+00 | 0,0E+00 | 0,0E+00 | 0,0E+00 | 0,0E+00 | 4172326 | 4172655 | 330 | - |
| Y11_RS18945 | Y11_39851 | | YecA family protein | 0,39 | 0,04 | 0,49 | -0,35 | 0,1 | 0,46 | 2,9E-02 | 8,4E-01 | 7,9E-02 | 1,1E-01 | 7,0E-01 | 1,4E-01 | 0,0E+00 | 0,0E+00 | 0,0E+00 | 0,0E+00 | 0,0E+00 | 0,0E+00 | 4173111 | 4173689 | 579 | + |
| Y11_RS18950 | Y11_39861 | pepP | Xaa-Pro aminopeptidase | 0,22 | 1,04 | 1,56 | 0,82 | 1,34 | 0,52 | 7,2E-04 | 3,3E-07 | 6,0E-13 | 5,8E-06 | 1,3E-09 | 6,8E-05 | 0,0E+00 | 1,1E-08 | 1,6E-12 | 0,0E+00 | 2,0E-10 | 0,0E+00 | 4173744 | 4175057 | 1314 | + |
| Y11_RS18955 | Y11_39871 | ubiH | 2-octaprenyl-6-methoxyphenyl hydroxylase | 0,07 | 0,85 | 1,09 | 0,78 | 1,02 | 0,24 | 3,9E-01 | 2,6E-06 | 8,8E-08 | 2,4E-06 | 4,8E-08 | 1,8E-03 | 0,0E+00 | 0,0E+00 | 2,9E-09 | 0,0E+00 | 3,4E-09 | 0,0E+00 | 4175061 | 4176230 | 1170 | + |
| Y11_RS18960 | Y11_39881 | ubiI | FAD-dependent 2-octaprenylphenol hydroxylase | -0,01 | 0,57 | 0,89 | 0,58 | 0,9 | 0,32 | 8,7E-01 | 4,2E-05 | 3,4E-06 | 2,7E-05 | 2,4E-06 | 6,5E-03 | 0,0E+00 | 0,0E+00 | 0,0E+00 | 0,0E+00 | 0,0E+00 | 0,0E+00 | 4176318 | 4177532 | 1215 | + |
| Y11_RS18965 | Y11_39901 | gcvT | glycine cleavage system aminomethyltransferase GcvT | -1,43 | 2,95 | 3,51 | 4,38 | 4,94 | 0,56 | 9,8E-07 | 1,5E-10 | 6,2E-12 | 2,1E-10 | 4,2E-11 | 3,4E-05 | 6,3E-08 | 2,9E-11 | 6,0E-12 | 2,7E-11 | 2,5E-11 | 0,0E+00 | 4177924 | 4179021 | 1098 | + |
| Y11_RS18970 | Y11_39911 | gcvH | glycine cleavage system protein GcvH | -1,36 | 2,6 | 3,99 | 3,97 | 5,35 | 1,39 | 4,7E-06 | 3,3E-08 | 2,5E-06 | 2,0E-09 | 2,8E-07 | 3,7E-03 | 2,5E-07 | 1,6E-09 | 4,9E-08 | 1,5E-10 | 1,4E-08 | 2,3E-04 | 4179088 | 4179474 | 387 | + |
| Y11_RS18975 | Y11_39921 | gcvP | aminomethyl-transferring glycine dehydrogenase | -1,27 | 2,34 | 2,89 | 3,6 | 4,16 | 0,55 | 1,0E-07 | 1,3E-10 | 6,5E-11 | 9,5E-12 | 6,9E-12 | 7,5E-06 | 9,4E-09 | 2,6E-11 | 1,8E-11 | 5,4E-12 | 8,8E-12 | 0,0E+00 | 4179574 | 4182462 | 2889 | + |
| Y11_RS18985 | Y11_39931 | | lipocalin-like domain-containing protein | -0,31 | 0,59 | 0,75 | 0,9 | 1,05 | 0,16 | 9,1E-02 | 1,9E-03 | 3,9E-04 | 9,5E-05 | 2,8E-05 | 8,9E-02 | 0,0E+00 | 0,0E+00 | 0,0E+00 | 0,0E+00 | 8,0E-07 | 0,0E+00 | 4182863 | 4183300 | 438 | + |
| Y11_RS18990 | Y11_39941 | | SDR family oxidoreductase | -0,17 | 0,79 | 1,25 | 0,96 | 1,42 | 0,46 | 9,0E-02 | 1,5E-06 | 2,6E-08 | 7,5E-06 | 4,4E-07 | 2,4E-04 | 0,0E+00 | 0,0E+00 | 2,2E-09 | 0,0E+00 | 2,1E-08 | 0,0E+00 | 4183316 | 4184065 | 750 | + |
| Y11_RS19015 | Y11_39991 | ygfZ | tRNA-modifying protein YgfZ | 0,94 | 0,49 | 0,72 | -0,45 | -0,21 | 0,23 | 4,0E-07 | 2,8E-05 | 2,8E-04 | 3,1E-05 | 1,0E-01 | 7,4E-02 | 0,0E+00 | 0,0E+00 | 0,0E+00 | 0,0E+00 | 0,0E+00 | 0,0E+00 | 4187220 | 4188212 | 993 | - |
| Y11_RS19020 | Y11_40001 | sdhE | FAD assembly factor SdhE | 0,55 | 0,66 | 1,48 | 0,11 | 0,93 | 0,82 | 3,4E-02 | 1,6E-02 | 1,8E-04 | 5,7E-01 | 1,4E-03 | 3,1E-03 | 0,0E+00 | 0,0E+00 | 2,4E-06 | 0,0E+00 | 0,0E+00 | 0,0E+00 | 4188479 | 4188745 | 267 | + |
| Y11_RS19045 | Y11_40061 | fldB | flavodoxin FldB | -0,47 | -0,1 | 0,25 | 0,37 | 0,72 | 0,35 | 4,0E-02 | 6,6E-01 | 2,5E-01 | 7,6E-02 | 1,8E-03 | 1,1E-01 | 0,0E+00 | 0,0E+00 | 0,0E+00 | 0,0E+00 | 0,0E+00 | 0,0E+00 | 4192854 | 4193372 | 519 | - |
| Y11_RS19050 | Y11_40071 | xerD | site-specific tyrosine recombinase XerD | -0,09 | 0,44 | 0,71 | 0,53 | 0,8 | 0,27 | 4,2E-01 | 3,8E-03 | 8,1E-05 | 2,6E-04 | 2,8E-06 | 6,9E-03 | 0,0E+00 | 0,0E+00 | 0,0E+00 | 0,0E+00 | 0,0E+00 | 0,0E+00 | 4193479 | 4194378 | 900 | + |
| Y11_RS19055 | Y11_40081 | dsbC | bifunctional protein-disulfide isomerase/oxidoreductase DsbC | 0,15 | -0,02 | -0,32 | -0,17 | -0,47 | -0,3 | 1,7E-01 | 8,0E-01 | 3,6E-03 | 9,2E-02 | 7,4E-04 | 2,0E-03 | 0,0E+00 | 0,0E+00 | 0,0E+00 | 0,0E+00 | 0,0E+00 | 0,0E+00 | 4194427 | 4195143 | 717 | + |
| Y11_RS19060 | Y11_40091 | recJ | single-stranded-DNA-specific exonuclease RecJ | 0,12 | -0,44 | -0,3 | -0,56 | -0,42 | 0,14 | 6,3E-02 | 1,4E-05 | 3,1E-05 | 1,9E-05 | 5,6E-05 | 1,5E-02 | 0,0E+00 | 0,0E+00 | 0,0E+00 | 0,0E+00 | 0,0E+00 | 0,0E+00 | 4195150 | 4196883 | 1734 | + |
| Y11_RS19065 | Y11_40101 | prfB | peptide chain release factor 2 | -0,24 | -0,87 | -1,15 | -0,63 | -0,91 | -0,28 | 1,1E-02 | 1,3E-05 | 1,4E-06 | 2,9E-05 | 1,6E-06 | 1,7E-02 | 0,0E+00 | 0,0E+00 | 3,1E-08 | 0,0E+00 | 0,0E+00 | 0,0E+00 | 4197405 | 4198427 | 1023 | + |
| Y11_RS19070 | Y11_40111 | lysS | lysine--tRNA ligase | -0,21 | -0,81 | -0,66 | -0,61 | -0,45 | 0,16 | 4,9E-04 | 2,2E-08 | 1,5E-03 | 3,1E-07 | 1,2E-02 | 3,0E-01 | 0,0E+00 | 0,0E+00 | 0,0E+00 | 0,0E+00 | 0,0E+00 | 0,0E+00 | 4198437 | 4199954 | 1518 | + |
| Y11_RS19085 | Y11_40141 | | helix-turn-helix domain-containing protein | 0,89 | 1,36 | 1,7 | 0,47 | 0,81 | 0,35 | 5,7E-07 | 6,5E-06 | 1,5E-07 | 6,4E-03 | 3,0E-05 | 5,0E-02 | 0,0E+00 | 1,6E-07 | 4,6E-09 | 0,0E+00 | 0,0E+00 | 0,0E+00 | 4201337 | 4201699 | 363 | + |
| Y11_RS19095 | Y11_40161 | | type II toxin-antitoxin system MqsA family antitoxin | 1,01 | 3,12 | 2,77 | 2,12 | 1,76 | -0,36 | 1,3E-02 | 1,7E-06 | 5,2E-06 | 4,5E-04 | 1,6E-03 | 3,0E-01 | 4,4E-04 | 4,9E-08 | 9,4E-08 | 7,8E-06 | 3,5E-05 | 0,0E+00 | 4202152 | 4202550 | 399 | + |
| Y11_RS19115 | Y11_40211 | | SgcJ/EcaC family oxidoreductase | 0,43 | 0,29 | 1,3 | -0,14 | 0,87 | 1,01 | 1,3E-01 | 1,9E-01 | 2,4E-04 | 5,5E-01 | 5,4E-03 | 3,8E-04 | 0,0E+00 | 0,0E+00 | 3,2E-06 | 0,0E+00 | 0,0E+00 | 2,7E-05 | 4205434 | 4205829 | 396 | + |
| Y11_RS19135 | Y11_40251 | cbpA | curved DNA-binding protein | 4,58 | 2,5 | 3,23 | -2,08 | -1,34 | 0,73 | 1,7E-12 | 2,2E-10 | 7,3E-11 | 2,9E-10 | 3,2E-08 | 3,6E-06 | 2,5E-12 | 3,8E-11 | 1,9E-11 | 3,5E-11 | 2,5E-09 | 0,0E+00 | 4208815 | 4209768 | 954 | + |
| Y11_RS19140 | Y11_40261 | cbpM | chaperone modulator CbpM | 15 | 3,15 | 4,52 | -15 | -15 | 1,35 | 0,0E+00 | 9,8E-08 | 1,2E-07 | 0,0E+00 | 0,0E+00 | 1,4E-03 | 0,0E+00 | 4,0E-09 | 3,7E-09 | 0,0E+00 | 0,0E+00 | 9,4E-05 | 4209775 | 4210080 | 306 | + |
| Y11_RS19150 | Y11_40281 | | helix-turn-helix transcriptional regulator | 15 | 0,34 | 0,61 | -15 | -15 | 0,27 | 0,0E+00 | 1,5E-01 | 2,5E-01 | 0,0E+00 | 0,0E+00 | 6,1E-01 | 0,0E+00 | 0,0E+00 | 0,0E+00 | 0,0E+00 | 0,0E+00 | 0,0E+00 | 4211000 | 4211383 | 384 | + |
| Y11_RS19155 | Y11_40291 | | carbonic anhydrase | 4,96 | 3,78 | 4,53 | -1,18 | -0,43 | 0,76 | 8,4E-12 | 7,9E-11 | 1,3E-10 | 4,1E-06 | 9,6E-03 | 3,5E-04 | 7,8E-12 | 1,9E-11 | 2,7E-11 | 8,9E-08 | 0,0E+00 | 0,0E+00 | 4211533 | 4212288 | 756 | + |
| Y11_RS19255 | Y11_40521 | | MBL fold metallo-hydrolase | 0 | 0 | -15 | 0 | -15 | -15 | 0,0E+00 | 0,0E+00 | 0,0E+00 | 0,0E+00 | 0,0E+00 | 0,0E+00 | 0,0E+00 | 0,0E+00 | 0,0E+00 | 0,0E+00 | 0,0E+00 | 0,0E+00 | 4227737 | 4228624 | 888 | - |
| Y11_RS19270 | Y11_40551 | | LysR family transcriptional regulator | 1,75 | 1,01 | 1,44 | -0,74 | -0,31 | 0,43 | 9,4E-08 | 2,3E-05 | 8,0E-05 | 1,8E-04 | 1,5E-01 | 6,8E-02 | 8,7E-09 | 5,2E-07 | 1,1E-06 | 0,0E+00 | 0,0E+00 | 0,0E+00 | 4231122 | 4232075 | 954 | - |
| Y11_RS19275 | Y11_40561 | lysA | diaminopimelate decarboxylase | 15 | -1,02 | -0,42 | -15 | -15 | 0,6 | 0,0E+00 | 2,6E-03 | 5,3E-02 | 0,0E+00 | 0,0E+00 | 6,5E-02 | 0,0E+00 | 5,1E-05 | 0,0E+00 | 0,0E+00 | 0,0E+00 | 0,0E+00 | 4232242 | 4233504 | 1263 | + |
| Y11_RS19280 | Y11_40571 | galR | HTH-type transcriptional regulator GalR | 15 | -0,59 | 0,22 | -15 | -15 | 0,81 | 0,0E+00 | 6,1E-02 | 3,4E-01 | 0,0E+00 | 0,0E+00 | 1,3E-02 | 0,0E+00 | 0,0E+00 | 0,0E+00 | 0,0E+00 | 0,0E+00 | 0,0E+00 | 4233523 | 4234557 | 1035 | - |
| Y11_RS19295 | Y11_40601 | aas | bifunctional acyl-ACP--phospholipid O-acyltransferase/long-chain-fatty-acid--ACP ligase | -0,7 | -0,81 | -0,32 | -0,11 | 0,38 | 0,49 | 4,5E-06 | 3,3E-07 | 1,8E-03 | 4,0E-02 | 3,4E-04 | 1,4E-05 | 0,0E+00 | 0,0E+00 | 0,0E+00 | 0,0E+00 | 0,0E+00 | 0,0E+00 | 4239188 | 4241344 | 2157 | + |
| Y11_RS19305 | Y11_40621 | | NADP(H)-dependent aldo-keto reductase | 0,5 | -0,33 | 0,62 | -0,84 | 0,12 | 0,96 | 4,7E-06 | 1,7E-04 | 4,0E-04 | 2,6E-08 | 2,6E-01 | 1,6E-05 | 0,0E+00 | 0,0E+00 | 0,0E+00 | 0,0E+00 | 0,0E+00 | 0,0E+00 | 4242615 | 4243655 | 1041 | - |
| Y11_RS19315 | Y11_40641 | mutH | DNA mismatch repair endonuclease MutH | -0,56 | 1,54 | 2,92 | 2,1 | 3,48 | 1,38 | 7,2E-03 | 2,4E-03 | 2,3E-06 | 3,9E-04 | 8,7E-07 | 9,4E-03 | 0,0E+00 | 4,6E-05 | 4,7E-08 | 6,8E-06 | 3,7E-08 | 5,5E-04 | 4244619 | 4245305 | 687 | - |
| Y11_RS19320 | Y11_40661 | rppH | RNA pyrophosphohydrolase | 0,56 | 0,66 | 0,88 | 0,1 | 0,32 | 0,22 | 5,3E-02 | 9,3E-03 | 4,3E-03 | 6,1E-01 | 1,8E-01 | 1,8E-01 | 0,0E+00 | 0,0E+00 | 0,0E+00 | 0,0E+00 | 0,0E+00 | 0,0E+00 | 4246003 | 4246530 | 528 | + |
| Y11_RS19325 | Y11_40671 | ptsP | phosphoenolpyruvate--protein phosphotransferase | 0,07 | 0,32 | 0,47 | 0,25 | 0,4 | 0,15 | 3,2E-01 | 9,5E-04 | 4,6E-04 | 6,7E-07 | 1,0E-04 | 2,7E-02 | 0,0E+00 | 0,0E+00 | 0,0E+00 | 0,0E+00 | 0,0E+00 | 0,0E+00 | 4246543 | 4248789 | 2247 | + |
| Y11_RS19330 | Y11_40681 | lgt | prolipoprotein diacylglyceryl transferase | -2,32 | -1,69 | -0,89 | 0,63 | 1,43 | 0,8 | 1,2E-09 | 9,1E-09 | 4,1E-03 | 7,1E-05 | 2,4E-04 | 8,0E-03 | 2,7E-10 | 5,7E-10 | 0,0E+00 | 0,0E+00 | 6,0E-06 | 0,0E+00 | 4248917 | 4249789 | 873 | + |
| Y11_RS19335 | Y11_40691 | thyA | thymidylate synthase | -2,34 | -1,2 | -0,28 | 1,14 | 2,06 | 0,92 | 4,5E-11 | 1,6E-07 | 5,1E-03 | 2,8E-07 | 2,9E-09 | 7,0E-06 | 2,1E-11 | 6,1E-09 | 0,0E+00 | 8,2E-09 | 3,7E-10 | 0,0E+00 | 4249796 | 4250590 | 795 | + |
| Y11_RS19340 | Y11_40701 | | prepilin peptidase-dependent protein | 0,2 | -0,74 | -0,83 | -0,94 | -1,03 | -0,09 | 2,4E-01 | 8,5E-04 | 1,3E-02 | 3,6E-04 | 5,2E-03 | 7,3E-01 | 0,0E+00 | 0,0E+00 | 0,0E+00 | 0,0E+00 | 1,1E-04 | 0,0E+00 | 4250822 | 4251358 | 537 | + |
| Y11_RS19360 | Y11_40741 | recC | exodeoxyribonuclease V subunit gamma | 0 | -0,25 | 0,19 | -0,26 | 0,18 | 0,44 | 9,6E-01 | 8,9E-03 | 1,1E-01 | 1,6E-03 | 8,1E-02 | 2,6E-03 | 0,0E+00 | 0,0E+00 | 0,0E+00 | 0,0E+00 | 0,0E+00 | 0,0E+00 | 4252962 | 4256333 | 3372 | + |
| Y11_RS19365 | Y11_40751 | ptrA | pitrilysin | -0,47 | -0,41 | 0,8 | 0,06 | 1,27 | 1,21 | 1,1E-07 | 1,9E-06 | 3,4E-08 | 1,1E-01 | 1,0E-09 | 3,8E-09 | 0,0E+00 | 0,0E+00 | 0,0E+00 | 0,0E+00 | 1,7E-10 | 3,6E-09 | 4256433 | 4259324 | 2892 | + |
| Y11_RS19370 | Y11_40761 | recB | exodeoxyribonuclease V subunit beta | 0 | -0,01 | 0,46 | -0,01 | 0,46 | 0,47 | 9,5E-01 | 8,8E-01 | 2,1E-06 | 8,6E-01 | 3,3E-05 | 1,1E-05 | 0,0E+00 | 0,0E+00 | 0,0E+00 | 0,0E+00 | 0,0E+00 | 0,0E+00 | 4259321 | 4262950 | 3630 | + |
| Y11_RS19375 | Y11_40771 | recD | exodeoxyribonuclease V subunit alpha | -0,64 | 0,18 | 0,9 | 0,82 | 1,54 | 0,71 | 1,6E-05 | 7,9E-02 | 5,3E-05 | 1,8E-05 | 9,4E-07 | 5,7E-04 | 0,0E+00 | 0,0E+00 | 0,0E+00 | 0,0E+00 | 4,0E-08 | 0,0E+00 | 4262947 | 4264848 | 1902 | + |
| Y11_RS19385 | Y11_40791 | argA | amino-acid N-acetyltransferase | 0,58 | -0,88 | -0,79 | -1,46 | -1,37 | 0,09 | 4,0E-03 | 5,0E-03 | 2,3E-03 | 1,1E-04 | 1,9E-05 | 7,1E-01 | 0,0E+00 | 0,0E+00 | 0,0E+00 | 2,0E-06 | 5,7E-07 | 0,0E+00 | 4266435 | 4267760 | 1326 | - |
| Y11_RS19390 | Y11_40801 | amiC | N-acetylmuramoyl-L-alanine amidase AmiC | -0,01 | 1,04 | 0,77 | 1,04 | 0,77 | -0,27 | 9,6E-01 | 2,4E-06 | 9,0E-05 | 9,2E-05 | 1,1E-03 | 5,1E-02 | 0,0E+00 | 6,5E-08 | 0,0E+00 | 1,7E-06 | 0,0E+00 | 0,0E+00 | 4267995 | 4269245 | 1251 | + |

| | | | | | | | | | | | | | | | | | | | | | | | | | |
|---|---|---|---|---|---|---|---|---|---|---|---|---|---|---|---|---|---|---|---|---|---|---|---|---|---|
| Y11_RS19400 | Y11_40811 | mltA | murein transglycosylase A | 0,16 | -2,72 | -3,64 | -2,97 | -3,79 | -0,91 | 3,9E-01 | 6,8E-09 | 9,5E-10 | 5,3E-09 | 1,1E-08 | 4,6E-07 | 0,0E+00 | 4,7E-10 | 9,8E-11 | 3,0E-10 | 1,1E-09 | 0,0E+00 | 4269922 | 4271094 | 1173 | + |
| Y11_RS19405 | Y11_40821 | tcdA | tRNA cyclic N6-threonylcarbamoyladenosine(37) synthase TcdA | -0,16 | -0,9 | -1,19 | -0,74 | -1,03 | -0,29 | 6,2E-02 | 2,9E-05 | 1,3E-03 | 8,7E-05 | 3,0E-03 | 3,0E-01 | 0,0E+00 | 0,0E+00 | 1,6E-05 | 0,0E+00 | 6,6E-05 | 0,0E+00 | 4271209 | 4272018 | 810 | + |
| Y11_RS19410 | Y11_40831 | csdE | cysteine desulfurase sulfur acceptor subunit CsdE | 0,07 | 0,38 | 0,32 | 0,3 | 0,24 | -0,06 | 2,2E-01 | 4,1E-05 | 5,9E-04 | 1,1E-03 | 8,4E-03 | 3,7E-01 | 0,0E+00 | 0,0E+00 | 0,0E+00 | 0,0E+00 | 0,0E+00 | 0,0E+00 | 4272022 | 4272462 | 441 | - |
| Y11_RS19415 | Y11_40841 | csdA | cysteine desulfurase CsdA | -0,1 | 0,31 | 0,21 | 0,42 | 0,32 | -0,1 | 4,0E-01 | 4,9E-03 | 6,6E-02 | 3,8E-03 | 2,8E-02 | 2,6E-01 | 0,0E+00 | 0,0E+00 | 0,0E+00 | 0,0E+00 | 0,0E+00 | 0,0E+00 | 4272481 | 4273686 | 1206 | - |
| Y11_RS19420 | Y11_40851 | | YgdI/YgdR family lipoprotein | 1,08 | 2,33 | 2,64 | 1,24 | 1,55 | 0,31 | 3,4E-06 | 5,8E-08 | 4,8E-05 | 3,7E-05 | 2,1E-03 | 4,1E-01 | 1,9E-07 | 2,6E-09 | 7,0E-07 | 7,0E-07 | 4,6E-05 | 0,0E+00 | 4273869 | 4274090 | 222 | + |
| Y11_RS19425 | Y11_40861 | | transcriptional regulator GcvA | 0,21 | 0,23 | 0,32 | 0,02 | 0,11 | 0,09 | 1,6E-01 | 1,1E-01 | 5,7E-03 | 8,9E-01 | 4,6E-01 | 5,5E-01 | 0,0E+00 | 0,0E+00 | 0,0E+00 | 0,0E+00 | 0,0E+00 | 0,0E+00 | 4274571 | 4275488 | 918 | + |
| Y11_RS19435 | Y11_40881 | rlmM | 23S rRNA (cytidine(2498)-2'-O)-methyltransferase RlmM | 0,36 | 0,04 | -0,15 | -0,32 | -0,5 | -0,18 | 6,7E-03 | 6,9E-01 | 1,5E-01 | 1,0E-02 | 6,1E-04 | 7,1E-02 | 0,0E+00 | 0,0E+00 | 0,0E+00 | 0,0E+00 | 0,0E+00 | 0,0E+00 | 4275916 | 4277022 | 1107 | + |
| Y11_RS19440 | Y11_40891 | | IS110 family transposase | -0,98 | 1,33 | 1,04 | 2,32 | 2,02 | -0,3 | 2,9E-03 | 1,4E-05 | 6,0E-06 | 1,5E-05 | 2,0E-05 | 5,5E-02 | 0,0E+00 | 1,1E-07 | 2,9E-07 | 6,1E-07 | 0,0E+00 | 0,0E+00 | 4277426 | 4278445 | 1020 | - |
| Y11_RS19455 | Y11_40921 | ppnN | nucleotide 5'-monophosphate nucleosidase PpnN | -2,28 | -1,93 | -1,43 | 0,34 | 0,84 | 0,5 | 3,2E-08 | 8,8E-09 | 9,2E-09 | 2,5E-02 | 6,5E-05 | 2,1E-04 | 3,3E-09 | 5,5E-10 | 5,0E-10 | 0,0E+00 | 0,0E+00 | 0,0E+00 | 4280846 | 4282210 | 1365 | - |
| Y11_RS19460 | Y11_40931 | queF | NADPH-dependent 7-cyano-7-deazaguanine reductase QueF | -15 | -15 | -15 | 0,65 | 0,77 | 0,12 | 0,0E+00 | 0,0E+00 | 0,0E+00 | 4,6E-03 | 8,3E-03 | 5,2E-01 | 0,0E+00 | 0,0E+00 | 0,0E+00 | 0,0E+00 | 0,0E+00 | 0,0E+00 | 4282424 | 4283269 | 846 | - |
| Y11_RS19465 | Y11_40941 | syd | SecY-interacting protein | -15 | -15 | -15 | -0,4 | -0,12 | 0,28 | 0,0E+00 | 0,0E+00 | 0,0E+00 | 1,0E-01 | 5,4E-01 | 7,4E-02 | 0,0E+00 | 0,0E+00 | 0,0E+00 | 0,0E+00 | 0,0E+00 | 0,0E+00 | 4283401 | 4283952 | 552 | + |
| Y11_RS19480 | Y11_40971 | | flavodoxin | -0,27 | -1,04 | -0,79 | -0,77 | -0,52 | 0,25 | 7,2E-03 | 1,8E-06 | 1,8E-04 | 1,0E-05 | 2,2E-03 | 7,2E-02 | 0,0E+00 | 5,0E-08 | 0,0E+00 | 0,0E+00 | 0,0E+00 | 0,0E+00 | 4285760 | 4286209 | 450 | + |
| Y11_RS19485 | Y11_40981 | | DUF3461 family protein | -0,8 | 0,79 | 1,33 | 1,59 | 2,13 | 0,54 | 2,7E-05 | 1,3E-07 | 9,8E-05 | 8,6E-08 | 5,4E-06 | 1,9E-02 | 0,0E+00 | 0,0E+00 | 1,3E-06 | 2,9E-09 | 1,9E-07 | 0,0E+00 | 4286345 | 4286734 | 390 | - |
| Y11_RS19490 | Y11_40991 | dapD | 2,3,4,5-tetrahydropyridine-2,6-dicarboxylate N-succinyltransferase | 0,32 | -1,25 | -1,57 | -1,57 | -1,89 | -0,32 | 2,8E-04 | 8,1E-09 | 1,6E-06 | 8,3E-09 | 5,6E-07 | 4,2E-02 | 0,0E+00 | 5,2E-10 | 3,5E-08 | 4,5E-10 | 2,6E-08 | 0,0E+00 | 4286855 | 4287679 | 825 | - |
| Y11_RS19495 | Y11_41001 | glnD | bifunctional uridylyltransferase/uridylyl-removing protein GlnD | 0,49 | -0,18 | -0,03 | -0,67 | -0,52 | 0,15 | 2,7E-04 | 1,2E-01 | 6,7E-01 | 5,7E-05 | 4,8E-06 | 1,2E-01 | 0,0E+00 | 0,0E+00 | 0,0E+00 | 0,0E+00 | 0,0E+00 | 0,0E+00 | 4287796 | 4290474 | 2679 | - |
| Y11_RS19500 | Y11_41011 | map | type I methionyl aminopeptidase | 0,39 | 0 | 0 | -0,39 | -0,39 | 0 | 1,2E-03 | 1,0E+00 | 9,9E-01 | 8,0E-05 | 1,1E-02 | 9,9E-01 | 0,0E+00 | 0,0E+00 | 0,0E+00 | 0,0E+00 | 0,0E+00 | 0,0E+00 | 4290537 | 4291328 | 792 | - |
| Y11_RS19505 | Y11_41021 | rpsB | 30S ribosomal protein S2 | 0,25 | -0,18 | -0,55 | -0,42 | -0,79 | -0,37 | 4,1E-03 | 2,3E-02 | 6,6E-04 | 8,6E-04 | 1,2E-04 | 1,2E-02 | 0,0E+00 | 0,0E+00 | 0,0E+00 | 0,0E+00 | 0,0E+00 | 0,0E+00 | 4291858 | 4292583 | 726 | + |
| Y11_RS19510 | Y11_41031 | tsf | translation elongation factor Ts | 0,05 | 0,14 | -0,03 | 0,09 | -0,08 | -0,17 | 5,1E-01 | 7,6E-02 | 7,7E-01 | 3,3E-01 | 4,5E-01 | 1,1E-01 | 0,0E+00 | 0,0E+00 | 0,0E+00 | 0,0E+00 | 0,0E+00 | 0,0E+00 | 4292712 | 4293569 | 858 | + |
| Y11_RS19515 | Y11_41041 | pyrH | UMP kinase | -0,07 | -0,13 | 0,3 | -0,06 | 0,37 | 0,43 | 2,4E-01 | 1,4E-01 | 1,1E-02 | 4,4E-01 | 2,6E-03 | 2,9E-03 | 0,0E+00 | 0,0E+00 | 0,0E+00 | 0,0E+00 | 0,0E+00 | 0,0E+00 | 4293837 | 4294583 | 747 | + |
| Y11_RS19520 | Y11_41051 | frr | ribosome recycling factor | 0,11 | 0,02 | -0,1 | -0,08 | -0,21 | -0,13 | 2,7E-01 | 6,9E-01 | 2,8E-01 | 3,1E-01 | 8,6E-02 | 1,5E-01 | 0,0E+00 | 0,0E+00 | 0,0E+00 | 0,0E+00 | 0,0E+00 | 0,0E+00 | 4294716 | 4295273 | 558 | + |
| Y11_RS19525 | Y11_41061 | ispC | 1-deoxy-D-xylulose-5-phosphate reductoisomerase | 0,27 | 0,21 | 0,79 | -0,06 | 0,52 | 0,58 | 1,7E-02 | 6,2E-03 | 5,4E-06 | 5,5E-01 | 1,4E-03 | 1,1E-04 | 0,0E+00 | 0,0E+00 | 0,0E+00 | 0,0E+00 | 0,0E+00 | 0,0E+00 | 4295583 | 4296779 | 1197 | + |
| Y11_RS19530 | Y11_41071 | ispU | (2E,6E)-farnesyl-diphosphate-specific ditrans,polycis-undecaprenyl-diphosphate synthase | -0,2 | -0,83 | -0,6 | -0,63 | -0,4 | 0,23 | 6,4E-02 | 3,5E-06 | 5,0E-04 | 1,7E-05 | 4,9E-03 | 3,0E-02 | 0,0E+00 | 0,0E+00 | 0,0E+00 | 0,0E+00 | 0,0E+00 | 0,0E+00 | 4297005 | 4297763 | 759 | + |
| Y11_RS19535 | Y11_41081 | cdsA | phosphatidate cytidylyltransferase | 0,5 | 0,95 | 15 | 0,44 | 15 | 0 | 1,7E-01 | 9,6E-05 | 0,0E+00 | 2,0E-01 | 0,0E+00 | 0,0E+00 | 0,0E+00 | 0,0E+00 | 0,0E+00 | 0,0E+00 | 0,0E+00 | 0,0E+00 | 4297773 | 4298621 | 849 | + |
| Y11_RS19540 | Y11_41091 | rseP | sigma E protease regulator RseP | 0,26 | -0,79 | -0,5 | -1,05 | -0,76 | 0,29 | 1,3E-02 | 2,3E-07 | 1,8E-06 | 3,2E-06 | 2,5E-05 | 1,1E-03 | 0,0E+00 | 0,0E+00 | 0,0E+00 | 7,1E-08 | 0,0E+00 | 0,0E+00 | 4298650 | 4300005 | 1356 | + |
| Y11_RS19545 | Y11_41101 | bamA | outer membrane protein assembly factor BamA | -0,33 | -0,71 | -0,66 | -0,38 | -0,33 | 0,05 | 2,0E-06 | 1,3E-10 | 2,4E-05 | 1,6E-06 | 3,3E-03 | 5,3E-01 | 0,0E+00 | 0,0E+00 | 0,0E+00 | 0,0E+00 | 0,0E+00 | 0,0E+00 | 4300042 | 4302426 | 2385 | + |
| Y11_RS19550 | Y11_41111 | skp | molecular chaperone Skp | -0,14 | 0,36 | 0,49 | 0,5 | 0,64 | 0,13 | 3,6E-02 | 3,5E-05 | 5,2E-05 | 3,1E-05 | 3,0E-05 | 7,7E-02 | 0,0E+00 | 0,0E+00 | 0,0E+00 | 0,0E+00 | 0,0E+00 | 0,0E+00 | 4302585 | 4303082 | 498 | + |
| Y11_RS19555 | Y11_41121 | lpxD | UDP-3-O-(3-hydroxymyristoyl)glucosamine N-acyltransferase | -0,18 | 0 | -0,08 | 0,17 | 0,09 | -0,08 | 1,3E-01 | 9,8E-01 | 6,3E-01 | 2,3E-01 | 6,1E-01 | 6,8E-01 | 0,0E+00 | 0,0E+00 | 0,0E+00 | 0,0E+00 | 0,0E+00 | 0,0E+00 | 4303086 | 4304108 | 1023 | + |
| Y11_RS19560 | Y11_41131 | fabZ | 3-hydroxyacyl-ACP dehydratase FabZ | -0,33 | 0,67 | 0,19 | 1 | 0,52 | -0,47 | 3,0E-04 | 1,8E-06 | 1,1E-01 | 3,7E-07 | 1,8E-03 | 3,2E-03 | 0,0E+00 | 0,0E+00 | 0,0E+00 | 0,0E+00 | 0,0E+00 | 0,0E+00 | 4304271 | 4304807 | 537 | + |
| Y11_RS19565 | Y11_41141 | lpxA | acyl-ACP--UDP-N-acetylglucosamine O-acyltransferase | -0,32 | 0,75 | 0,59 | 1,07 | 0,91 | -0,16 | 5,2E-02 | 1,4E-05 | 9,2E-05 | 8,7E-05 | 2,8E-04 | 1,3E-01 | 0,0E+00 | 0,0E+00 | 0,0E+00 | 1,6E-06 | 0,0E+00 | 0,0E+00 | 4304811 | 4305599 | 789 | + |
| Y11_RS19570 | Y11_41151 | lpxB | lipid-A-disaccharide synthase | -0,13 | 0,25 | 0,02 | 0,38 | 0,14 | -0,23 | 1,7E-01 | 7,2E-03 | 8,5E-01 | 3,0E-04 | 1,1E-01 | 7,5E-03 | 0,0E+00 | 0,0E+00 | 0,0E+00 | 0,0E+00 | 0,0E+00 | 0,0E+00 | 4305603 | 4306787 | 1185 | + |
| Y11_RS19580 | Y11_41171 | dnaE | DNA polymerase III subunit alpha | -0,03 | 0,11 | -0,73 | 0,15 | -0,7 | -0,85 | 6,3E-01 | 1,1E-02 | 1,1E-05 | 6,7E-02 | 9,1E-05 | 4,0E-06 | 0,0E+00 | 0,0E+00 | 0,0E+00 | 0,0E+00 | 0,0E+00 | 0,0E+00 | 4307491 | 4310973 | 3483 | + |
| Y11_RS19585 | Y11_41181 | accA | acetyl-CoA carboxylase carboxyl transferase subunit alpha | 0,23 | 0,12 | -0,11 | -0,11 | -0,35 | -0,24 | 9,5E-03 | 1,3E-01 | 2,1E-01 | 2,4E-01 | 6,6E-03 | 4,2E-02 | 0,0E+00 | 0,0E+00 | 0,0E+00 | 0,0E+00 | 0,0E+00 | 0,0E+00 | 4310986 | 4311945 | 960 | + |
| Y11_RS19590 | Y11_41191 | | lysine decarboxylase LdcC | -0,08 | 0,39 | 0 | 0,47 | 1,18 | 0,67 | 8,0E-01 | 6,6E-02 | 0,0E+00 | 1,9E-01 | 4,1E-03 | 5,3E-03 | 0,0E+00 | 0,0E+00 | 0,0E+00 | 0,0E+00 | 8,9E-05 | 0,0E+00 | 4312250 | 4314391 | 2142 | + |
| Y11_RS19595 | Y11_41201 | | VOC family protein | 0,23 | 0,62 | 0,6 | 0,39 | 0,37 | -0,02 | 1,2E-01 | 1,4E-03 | 6,4E-02 | 4,7E-02 | 2,5E-01 | 9,4E-01 | 0,0E+00 | 0,0E+00 | 0,0E+00 | 0,0E+00 | 0,0E+00 | 0,0E+00 | 4314405 | 4314803 | 399 | + |
| Y11_RS19600 | Y11_41211 | tilS | tRNA lysidine(34) synthetase TilS | 0,1 | -0,14 | 0,24 | -0,24 | 0,14 | 0,38 | 3,2E-01 | 2,9E-01 | 9,4E-02 | 6,4E-02 | 2,7E-01 | 2,6E-02 | 0,0E+00 | 0,0E+00 | 0,0E+00 | 0,0E+00 | 0,0E+00 | 0,0E+00 | 4314805 | 4316190 | 1386 | + |
| Y11_RS19610 | Y11_41231 | rof | Rho-binding antiterminator | -0,27 | 0,85 | 0,86 | 1,12 | 1,13 | 0,01 | 3,5E-02 | 1,8E-04 | 2,5E-04 | 2,3E-05 | 3,5E-05 | 9,4E-01 | 0,0E+00 | 0,0E+00 | 0,0E+00 | 4,6E-07 | 9,9E-07 | 0,0E+00 | 4316613 | 4316873 | 261 | - |
| Y11_RS19615 | Y11_41241 | | YaeP family protein | -0,5 | 0,16 | 0,22 | 0,65 | 0,72 | 0,07 | 4,0E-04 | 1,8E-01 | 3,5E-01 | 3,8E-05 | 1,0E-02 | 7,8E-01 | 0,0E+00 | 0,0E+00 | 0,0E+00 | 0,0E+00 | 0,0E+00 | 0,0E+00 | 4316860 | 4317060 | 201 | - |
| Y11_RS19620 | Y11_41251 | | YaeQ family protein | 0,01 | 0,49 | 0,44 | 0,48 | 0,43 | -0,05 | 9,2E-01 | 2,3E-03 | 1,2E-02 | 1,6E-03 | 1,1E-02 | 6,5E-01 | 0,0E+00 | 0,0E+00 | 0,0E+00 | 0,0E+00 | 0,0E+00 | 0,0E+00 | 4317298 | 4317846 | 549 | + |
| Y11_RS19630 | Y11_41271 | nlpE | envelope stress response activation lipoprotein NlpE | 0,13 | 1,54 | 1,66 | 1,41 | 1,53 | 0,12 | 3,8E-01 | 2,1E-06 | 4,8E-07 | 2,6E-06 | 5,2E-07 | 2,4E-01 | 0,0E+00 | 5,7E-08 | 1,2E-08 | 5,9E-08 | 2,4E-08 | 0,0E+00 | 4318338 | 4319021 | 684 | + |
| Y11_RS19635 | Y11_41281 | proS | proline--tRNA ligase | 0,05 | -0,16 | -0,58 | -0,21 | -0,64 | -0,42 | 3,5E-01 | 2,7E-02 | 3,5E-04 | 7,0E-03 | 2,0E-04 | 2,9E-03 | 0,0E+00 | 0,0E+00 | 0,0E+00 | 0,0E+00 | 0,0E+00 | 0,0E+00 | 4319095 | 4320813 | 1719 | - |
| Y11_RS19645 | | rcsF | Rcs stress response system protein RcsF | 0,67 | -0,57 | -0,75 | -1,24 | -1,42 | -0,18 | 1,3E-04 | 4,0E-04 | 1,1E-03 | 9,1E-07 | 1,1E-05 | 2,5E-01 | 0,0E+00 | 0,0E+00 | 0,0E+00 | 2,3E-08 | 3,6E-07 | 0,0E+00 | 4321746 | 4322153 | 408 | - |
| Y11_RS19650 | Y11_41321 | | MetQ/NlpA family lipoprotein | 0,06 | 1,59 | 1,24 | 1,53 | 1,18 | -0,35 | 2,4E-01 | 1,2E-08 | 3,3E-08 | 2,3E-08 | 7,1E-08 | 2,5E-03 | 0,0E+00 | 7,2E-10 | 1,4E-09 | 9,8E-10 | 4,7E-09 | 0,0E+00 | 4322301 | 4323116 | 816 | - |
| Y11_RS19655 | Y11_41331 | | methionine ABC transporter permease MetI | 0,21 | 1,04 | 0,64 | 0,83 | 0,43 | -0,4 | 4,0E-01 | 2,5E-02 | 3,7E-01 | 3,6E-02 | 5,2E-01 | 5,9E-01 | 0,0E+00 | 4,7E-04 | 0,0E+00 | 0,0E+00 | 0,0E+00 | 0,0E+00 | 4323311 | 4323964 | 654 | - |
| Y11_RS19660 | Y11_41341 | metN | methionine ABC transporter ATP-binding protein MetN | -0,02 | 0,79 | 0,17 | 0,82 | 0,2 | -0,62 | 6,7E-01 | 1,7E-07 | 4,2E-02 | 4,9E-07 | 3,6E-02 | 3,9E-05 | 0,0E+00 | 0,0E+00 | 0,0E+00 | 0,0E+00 | 0,0E+00 | 0,0E+00 | 4323957 | 4324988 | 1032 | - |
| Y11_RS19665 | Y11_41351 | gmhB | D-glycero-beta-D-manno-heptose 1,7-bisphosphate 7-phosphatase | -0,46 | 0,63 | 0,73 | 1,08 | 1,18 | 0,1 | 1,9E-03 | 3,7E-06 | 8,0E-04 | 2,2E-06 | 6,5E-05 | 4,7E-01 | 0,0E+00 | 0,0E+00 | 0,0E+00 | 5,1E-08 | 1,8E-06 | 0,0E+00 | 4325175 | 4325741 | 567 | + |
| Y11_RS19695 | Y11_41361 | | IS256 family transposase | 0,81 | 0,13 | -0,09 | -0,68 | -0,9 | -0,22 | 1,2E-03 | 2,1E-01 | 5,7E-01 | 2,5E-03 | 1,5E-03 | 1,4E-01 | 0,0E+00 | 0,0E+00 | 0,0E+00 | 0,0E+00 | 0,0E+00 | 0,0E+00 | 4332303 | 4333505 | 1203 | + |
| Y11_RS19705 | Y11_41381 | dkgB | 2,5-didehydrogluconate reductase DkgB | 0,19 | -0,21 | 0,13 | -0,41 | -0,06 | 0,34 | 1,8E-01 | 6,6E-02 | 1,6E-01 | 2,0E-02 | 6,4E-01 | 7,4E-03 | 0,0E+00 | 0,0E+00 | 0,0E+00 | 0,0E+00 | 0,0E+00 | 0,0E+00 | 4334632 | 4335435 | 804 | + |
| Y11_RS19710 | Y11_41391 | | LysR family transcriptional regulator | -0,06 | 0,25 | 0,25 | 0,31 | 0,3 | 0 | 6,6E-01 | 1,5E-01 | 5,0E-02 | 1,3E-01 | 6,2E-02 | 9,8E-01 | 0,0E+00 | 0,0E+00 | 0,0E+00 | 0,0E+00 | 0,0E+00 | 0,0E+00 | 4335480 | 4336382 | 903 | - |
| Y11_RS19720 | Y11_41431 | | endonuclease/exonuclease/phosphatase family protein | 0,8 | 1,21 | 1,77 | 0,41 | 0,97 | 0,57 | 1,3E-04 | 4,3E-07 | 3,8E-05 | 4,7E-03 | 2,7E-03 | 2,7E-02 | 0,0E+00 | 1,4E-08 | 5,6E-07 | 0,0E+00 | 0,0E+00 | 0,0E+00 | 4337949 | 4338710 | 762 | + |
| Y11_RS19735 | Y11_41471 | gloB | hydroxyacylglutathione hydrolase | 0,17 | 1,28 | 1,2 | 1,1 | 1,03 | -0,07 | 8,4E-02 | 2,2E-07 | 1,4E-06 | 5,8E-06 | 2,2E-05 | 5,2E-01 | 0,0E+00 | 8,1E-09 | 3,1E-08 | 1,2E-07 | 6,7E-07 | 0,0E+00 | 4341749 | 4342504 | 756 | - |
| Y11_RS19740 | Y11_41481 | | class I SAM-dependent methyltransferase | 0,96 | 0,71 | 1,02 | -0,25 | 0,06 | 0,31 | 9,0E-04 | 1,5E-03 | 5,6E-04 | 1,7E-01 | 7,7E-01 | 9,3E-02 | 0,0E+00 | 0,0E+00 | 6,9E-06 | 0,0E+00 | 0,0E+00 | 0,0E+00 | 4342551 | 4343270 | 720 | + |
| Y11_RS19745 | Y11_41491 | rnhA | ribonuclease HI | 0,77 | 0,85 | 1,39 | 0,08 | 0,62 | 0,54 | 2,1E-02 | 5,8E-04 | 3,2E-05 | 7,7E-01 | 4,4E-02 | 5,3E-03 | 0,0E+00 | 0,0E+00 | 4,9E-07 | 0,0E+00 | 0,0E+00 | 0,0E+00 | 4343286 | 4343750 | 465 | - |
| Y11_RS19750 | Y11_41501 | dnaQ | DNA polymerase III subunit epsilon | 0,26 | 0,18 | -0,48 | -0,08 | -0,74 | -0,65 | 8,1E-02 | 1,0E-01 | 9,9E-03 | 6,0E-01 | 4,2E-03 | 4,0E-03 | 0,0E+00 | 0,0E+00 | 0,0E+00 | 0,0E+00 | 0,0E+00 | 0,0E+00 | 4343821 | 4344603 | 783 | + |
| Y11_RS19760 | Y11_41511 | proX | glycine betaine/L-proline ABC transporter substrate-binding protein ProX | 1,7 | 1,57 | 2,22 | -0,14 | 0,51 | 0,65 | 1,6E-07 | 4,2E-08 | 2,3E-09 | 1,5E-01 | 2,9E-04 | 2,2E-06 | 1,3E-08 | 1,9E-09 | 1,8E-10 | 0,0E+00 | 0,0E+00 | 0,0E+00 | 4345034 | 4346038 | 1005 | - |
| Y11_RS19770 | Y11_41531 | proV | glycine betaine/L-proline ABC transporter ATP-binding protein ProV | 2,48 | 2,16 | 2,31 | -0,33 | 0 | 0,19 | 1,2E-04 | 3,1E-04 | 3,4E-04 | 6,8E-03 | 0,0E+00 | 3,0E-01 | 5,1E-06 | 6,4E-06 | 3,8E-06 | 0,0E+00 | 0,0E+00 | 0,0E+00 | 4347367 | 4348566 | 1200 | - |
| Y11_RS19775 | Y11_41541 | nrdF | class 1b ribonucleoside-diphosphate reductase subunit beta | -0,1 | -0,38 | 0,48 | -0,28 | 0,58 | 0,86 | 5,6E-01 | 1,6E-02 | 6,2E-02 | 6,7E-02 | 3,3E-02 | 2,4E-03 | 0,0E+00 | 0,0E+00 | 0,0E+00 | 0,0E+00 | 0,0E+00 | 0,0E+00 | 4349062 | 4350033 | 972 | - |
| Y11_RS19780 | Y11_41551 | nrdE | class 1b ribonucleoside-diphosphate reductase subunit alpha | 0,35 | -0,71 | -0,29 | -1,06 | -0,64 | 0,42 | 2,5E-02 | 3,0E-04 | 9,2E-02 | 3,6E-05 | 3,9E-03 | 2,3E-02 | 0,0E+00 | 0,0E+00 | 0,0E+00 | 6,9E-07 | 0,0E+00 | 0,0E+00 | 4350203 | 4352353 | 2151 | - |
| Y11_RS19795 | Y11_41591 | | carboxymuconolactone decarboxylase family protein | 0 | -15 | -15 | -2,43 | -2,08 | 0,36 | 0,0E+00 | 0,0E+00 | 0,0E+00 | 7,0E-05 | 3,1E-04 | 4,2E-01 | 0,0E+00 | 0,0E+00 | 0,0E+00 | 1,3E-06 | 7,5E-06 | 0,0E+00 | 4353272 | 4353703 | 432 | - |
| Y11_RS19800 | Y11_41601 | | PLP-dependent aminotransferase family protein | -0,35 | 0,95 | 1,38 | 1,3 | 1,72 | 0,42 | 6,0E-02 | 9,1E-04 | 3,0E-04 | 9,5E-05 | 5,5E-05 | 1,2E-01 | 0,0E+00 | 0,0E+00 | 3,8E-06 | 1,7E-06 | 1,5E-06 | 0,0E+00 | 4353800 | 4355254 | 1455 | + |
| Y11_RS19805 | Y11_41611 | | YqjD family protein | 0,7 | 2,26 | 2,11 | 1,56 | 1,41 | -0,15 | 4,6E-01 | 1,1E-02 | 2,7E-02 | 4,2E-02 | 9,4E-02 | 7,5E-01 | 0,0E+00 | 2,1E-04 | 3,0E-04 | 6,6E-04 | 1,9E-03 | 0,0E+00 | 4355402 | 4355710 | 309 | - |
| Y11_RS19835 | Y11_41661 | | DUF2002 family protein | -0,03 | -1,82 | -1,84 | -1,79 | -1,83 | -0,02 | 8,4E-01 | 6,4E-05 | 3,4E-04 | 3,0E-05 | 2,2E-04 | 9,5E-01 | 0,0E+00 | 1,4E-06 | 4,3E-06 | 5,8E-07 | 5,5E-06 | 0,0E+00 | 4359763 | 4360098 | 336 | + |
| Y11_RS19845 | Y11_41681 | | hypothetical protein | 0,14 | 0,11 | 0,67 | -0,02 | 0,54 | 0,56 | 1,2E-01 | 4,2E-01 | 7,6E-04 | 8,6E-01 | 1,1E-03 | 6,5E-03 | 0,0E+00 | 0,0E+00 | 0,0E+00 | 0,0E+00 | 0,0E+00 | 0,0E+00 | 4360689 | 4361027 | 339 | - |
| Y11_RS19870 | Y11_41731 | | hypothetical protein | 0,99 | -0,12 | 0,2 | -1,11 | -0,79 | 0,31 | 1,4E-02 | 6,8E-01 | 3,8E-01 | 1,8E-02 | 4,5E-02 | 3,2E-01 | 0,0E+00 | 0,0E+00 | 0,0E+00 | 2,9E-04 | 0,0E+00 | 0,0E+00 | 4363007 | 4363315 | 309 | + |
| Y11_RS19875 | Y11_41741 | | hypothetical protein | 0,76 | -2,85 | -1,83 | -3,6 | -2,59 | 1,02 | 2,6E-03 | 4,0E-09 | 1,8E-04 | 1,8E-08 | 2,9E-05 | 5,5E-03 | 0,0E+00 | 3,0E-10 | 2,4E-06 | 8,3E-10 | 8,5E-07 | 3,3E-04 | 4363430 | 4364239 | 810 | + |
| Y11_RS19880 | Y11_41751 | | ABC transporter substrate-binding protein | 0,77 | 0,21 | 1,35 | -0,56 | 0,58 | 1,14 | 3,5E-07 | 9,5E-04 | 3,7E-04 | 6,2E-06 | 3,7E-02 | 1,1E-03 | 0,0E+00 | 0,0E+00 | 4,6E-06 | 0,0E+00 | 0,0E+00 | 7,3E-05 | 4364842 | 4366425 | 1584 | + |
| Y11_RS19885 | Y11_41761 | | ABC transporter permease | 0,49 | -0,41 | 0,23 | -0,83 | 0 | 0,65 | 1,2E-01 | 6,8E-02 | 1,6E-01 | 1,9E-02 | 0,0E+00 | 1,7E-03 | 0,0E+00 | 0,0E+00 | 0,0E+00 | 0,0E+00 | 0,0E+00 | 0,0E+00 | 4366425 | 4367393 | 969 | + |
| Y11_RS19895 | Y11_41781 | | ATP-binding cassette domain-containing protein | 0,28 | 0,99 | 15 | 0,73 | 15 | 0 | 3,1E-01 | 6,5E-03 | 0,0E+00 | 5,5E-02 | 0,0E+00 | 0,0E+00 | 0,0E+00 | 0,0E+00 | 0,0E+00 | 0,0E+00 | 0,0E+00 | 0,0E+00 | 4368196 | 4369002 | 807 | + |
| Y11_RS19910 | Y11_41821 | | urease subunit gamma | 0,43 | 2,88 | 2,96 | 2,46 | 2,54 | 0,08 | 3,3E-03 | 3,3E-10 | 5,5E-08 | 9,6E-09 | 3,5E-07 | 6,1E-01 | 0,0E+00 | 4,9E-11 | 2,0E-09 | 5,1E-10 | 1,8E-08 | 0,0E+00 | 4370802 | 4371104 | 303 | + |
| Y11_RS19915 | Y11_41831 | | urease subunit beta | 0,36 | 3,16 | 3,41 | 2,8 | 3,05 | 0,25 | 5,4E-04 | 4,8E-11 | 5,8E-09 | 2,1E-12 | 6,8E-09 | 8,0E-02 | 0,0E+00 | 1,4E-11 | 3,6E-10 | 2,8E-12 | 7,1E-10 | 0,0E+00 | 4371158 | 4371652 | 495 | + |

| | | | | | | | | | | | | | | | | | | | | | | | | | |
|---|---|---|---|---|---|---|---|---|---|---|---|---|---|---|---|---|---|---|---|---|---|---|---|---|---|
| Y11_RS19920 | Y11_41841 | | urease subunit alpha | 0,37 | 2,79 | 2,94 | 2,42 | 2,57 | 0,15 | 2,4E-04 | 3,3E-11 | 1,9E-08 | 4,8E-12 | 3,7E-08 | 2,6E-01 | 0,0E+00 | 1,1E-11 | 8,6E-10 | 4,1E-12 | 2,7E-09 | 0,0E+00 | 4371817 | 4373535 | 1719 | + |
| Y11_RS19925 | Y11_41851 | ureE | urease accessory protein UreE | 0,19 | 2,22 | 1,98 | 2,03 | 1,8 | -0,24 | 5,1E-02 | 3,4E-09 | 2,3E-06 | 1,9E-10 | 2,7E-06 | 1,7E-01 | 0,0E+00 | 2,7E-10 | 4,7E-08 | 2,7E-11 | 1,0E-07 | 0,0E+00 | 4373709 | 4374389 | 681 | + |
| Y11_RS19930 | Y11_41861 | | urease accessory protein UreF | 0,05 | 2,32 | 1,67 | 2,27 | 1,62 | -0,65 | 5,5E-01 | 5,8E-09 | 8,2E-06 | 1,4E-08 | 1,3E-05 | 6,0E-03 | 0,0E+00 | 4,1E-10 | 1,4E-07 | 6,7E-10 | 4,1E-07 | 0,0E+00 | 4374411 | 4375097 | 687 | + |
| Y11_RS19935 | Y11_41871 | ureG | urease accessory protein UreG | 0,2 | 2,49 | 2,34 | 2,29 | 2,14 | -0,14 | 7,3E-02 | 3,5E-09 | 1,6E-07 | 4,5E-10 | 1,2E-07 | 2,6E-01 | 0,0E+00 | 2,8E-10 | 4,7E-09 | 4,7E-11 | 7,2E-09 | 0,0E+00 | 4375215 | 4375880 | 666 | + |
| Y11_RS19940 | Y11_41881 | | urease accessory protein UreD | 0,03 | 2,43 | 1,78 | 2,4 | 1,75 | -0,65 | 7,4E-01 | 9,9E-09 | 2,6E-06 | 7,6E-11 | 6,9E-07 | 1,1E-03 | 0,0E+00 | 6,1E-10 | 5,1E-08 | 1,5E-11 | 3,1E-08 | 0,0E+00 | 4375877 | 4376860 | 984 | + |
| Y11_RS19950 | Y11_41901 | | HoxN/HupN/NixA family nickel/cobalt transporter | -0,32 | 15 | 15 | 15 | 15 | 0 | 5,5E-01 | 0,0E+00 | 0,0E+00 | 0,0E+00 | 0,0E+00 | 0,0E+00 | 0,0E+00 | 0,0E+00 | 0,0E+00 | 0,0E+00 | 0,0E+00 | 0,0E+00 | 4377989 | 4379047 | 1059 | + |
| Y11_RS19955 | Y11_41911 | hdeB | acid-activated periplasmic chaperone HdeB | 0,19 | 1,88 | 3,27 | 1,69 | 3,09 | 1,4 | 7,4E-02 | 4,3E-08 | 9,2E-08 | 2,8E-08 | 1,1E-07 | 4,8E-05 | 0,0E+00 | 2,0E-09 | 3,0E-09 | 1,2E-09 | 6,6E-09 | 4,2E-06 | 4379113 | 4379445 | 333 | - |
| Y11_RS19960 | Y11_41941 | kch | voltage-gated potassium channel protein | 0,58 | 0,99 | 1,46 | 0,41 | 0,88 | 0,48 | 5,2E-04 | 2,3E-06 | 3,4E-04 | 4,1E-03 | 8,1E-03 | 8,7E-02 | 0,0E+00 | 0,0E+00 | 4,3E-06 | 0,0E+00 | 0,0E+00 | 0,0E+00 | 4379789 | 4380964 | 1176 | + |
| Y11_RS19965 | Y11_41951 | crcB | fluoride efflux transporter CrcB | 0,16 | 1,73 | 1,73 | 1,57 | 1,58 | 0 | 6,2E-01 | 1,1E-05 | 8,9E-07 | 2,3E-03 | 1,4E-03 | 9,9E-01 | 0,0E+00 | 2,7E-07 | 2,1E-08 | 3,7E-05 | 3,3E-05 | 0,0E+00 | 4381047 | 4381451 | 405 | - |
| Y11_RS19975 | Y11_41971 | mgrA | L-glyceraldehyde 3-phosphate reductase | 0,36 | 0,22 | 1,33 | -0,13 | 0,98 | 1,11 | 1,8E-05 | 4,1E-04 | 6,9E-08 | 8,3E-03 | 7,2E-07 | 2,6E-07 | 0,0E+00 | 0,0E+00 | 2,4E-09 | 0,0E+00 | 0,0E+00 | 6,9E-08 | 4382090 | 4383124 | 1035 | + |
| Y11_RS19995 | Y11_42011 | | PTS sugar transporter subunit IIB | -0,18 | 2,52 | 2,87 | 2,7 | 3,05 | 0,36 | 4,9E-02 | 1,9E-06 | 2,2E-08 | 1,6E-06 | 3,0E-08 | 1,8E-01 | 0,0E+00 | 5,4E-08 | 9,7E-10 | 3,8E-08 | 2,4E-09 | 0,0E+00 | 4386546 | 4386824 | 279 | + |
| Y11_RS20000 | Y11_42021 | chbC | PTS N,N'-diacetylchitobiose transporter subunit IIC | -0,15 | 1,38 | 1,6 | 1,53 | 1,74 | 0,21 | 4,2E-01 | 1,9E-04 | 1,3E-04 | 8,6E-05 | 6,6E-05 | 4,4E-01 | 0,0E+00 | 4,1E-06 | 1,8E-06 | 1,6E-06 | 1,8E-06 | 0,0E+00 | 4386835 | 4388196 | 1362 | + |
| Y11_RS20005 | Y11_42031 | | glycoside hydrolase family 1 protein | 0 | 1,61 | 2,75 | 1,61 | 2,75 | 1,14 | 9,9E-01 | 1,3E-09 | 3,3E-09 | 3,5E-09 | 4,4E-09 | 1,6E-06 | 0,0E+00 | 1,3E-10 | 2,3E-10 | 2,2E-10 | 5,0E-10 | 2,6E-07 | 4388235 | 4389620 | 1386 | + |
| Y11_RS20010 | Y11_42041 | chbA | PTS N,N'-diacetylchitobiose transporter subunit IIA | 0,45 | 1,23 | 0,84 | 0,78 | 0,39 | -0,39 | 2,0E-03 | 8,2E-07 | 2,0E-04 | 6,3E-05 | 2,3E-02 | 2,0E-02 | 0,0E+00 | 2,4E-08 | 0,0E+00 | 0,0E+00 | 0,0E+00 | 0,0E+00 | 4389649 | 4389996 | 348 | + |
| Y11_RS20020 | Y11_42061 | chbG | chitin disaccharide deacetylase | 0,72 | 1,07 | 1,45 | 0,35 | 0,73 | 0,37 | 7,7E-06 | 1,7E-06 | 5,8E-07 | 2,4E-03 | 6,7E-05 | 8,5E-03 | 0,0E+00 | 4,7E-08 | 1,4E-08 | 0,0E+00 | 0,0E+00 | 0,0E+00 | 4390927 | 4391688 | 762 | + |
| Y11_RS20045 | Y11_42101 | smpB | SsrA-binding protein SmpB | 0,33 | 0,04 | 0,47 | -0,28 | 0,14 | 0,43 | 1,3E-02 | 7,9E-01 | 2,9E-02 | 9,1E-02 | 4,1E-01 | 7,3E-02 | 0,0E+00 | 0,0E+00 | 0,0E+00 | 0,0E+00 | 0,0E+00 | 0,0E+00 | 4396877 | 4397359 | 483 | - |
| Y11_RS20050 | Y11_42111 | | type II toxin-antitoxin system RatA family toxin | -0,2 | -0,34 | -2,08 | -0,14 | -1,88 | -1,73 | 1,7E-01 | 1,2E-02 | 1,3E-05 | 2,2E-01 | 2,4E-05 | 2,7E-05 | 0,0E+00 | 0,0E+00 | 2,1E-07 | 0,0E+00 | 7,2E-07 | 2,7E-06 | 4397520 | 4397954 | 435 | + |
| Y11_RS20060 | Y11_42131 | bamE | outer membrane protein assembly factor BamE | 0,19 | -0,39 | 0,26 | -0,58 | 0,07 | 0,64 | 1,9E-01 | 1,9E-02 | 1,0E-01 | 9,5E-05 | 4,7E-01 | 1,3E-04 | 0,0E+00 | 0,0E+00 | 0,0E+00 | 0,0E+00 | 0,0E+00 | 0,0E+00 | 4398383 | 4398724 | 342 | - |
| Y11_RS20065 | Y11_42141 | recN | DNA repair protein RecN | 15 | -2,08 | -0,71 | -15 | -15 | 1,36 | 0,0E+00 | 5,3E-05 | 2,7E-04 | 0,0E+00 | 0,0E+00 | 9,2E-04 | 0,0E+00 | 1,2E-06 | 0,0E+00 | 0,0E+00 | 0,0E+00 | 6,2E-05 | 4398838 | 4400499 | 1662 | - |
| Y11_RS20070 | Y11_42151 | nadK | NAD(+) kinase | 0,23 | 0,67 | 1,15 | 0,44 | 0,92 | 0,48 | 5,3E-02 | 3,8E-04 | 3,2E-05 | 8,5E-03 | 2,6E-04 | 1,5E-02 | 0,0E+00 | 0,0E+00 | 4,9E-07 | 0,0E+00 | 0,0E+00 | 0,0E+00 | 4400586 | 4401467 | 882 | - |
| Y11_RS20075 | Y11_42171 | grpE | nucleotide exchange factor GrpE | 0,25 | -0,27 | -0,3 | -0,52 | -0,55 | -0,03 | 1,9E-03 | 1,9E-04 | 4,7E-03 | 7,8E-06 | 1,6E-04 | 7,3E-01 | 0,0E+00 | 0,0E+00 | 0,0E+00 | 0,0E+00 | 0,0E+00 | 0,0E+00 | 4401591 | 4402169 | 579 | + |
| Y11_RS20080 | Y11_42181 | ung | uracil-DNA glycosylase | 0,46 | 1,06 | 1,64 | 0,6 | 1,18 | 0,58 | 1,5E-04 | 1,7E-05 | 7,0E-08 | 7,1E-04 | 5,5E-07 | 1,8E-03 | 0,0E+00 | 4,1E-07 | 2,4E-09 | 0,0E+00 | 2,6E-08 | 0,0E+00 | 4402238 | 4402924 | 687 | - |
| Y11_RS20085 | Y11_42191 | grcA | autonomous glycyl radical cofactor GrcA | 0,81 | -0,27 | -0,73 | -1,09 | -1,54 | -0,45 | 1,6E-06 | 7,6E-03 | 8,4E-07 | 3,1E-06 | 4,3E-08 | 8,6E-04 | 0,0E+00 | 0,0E+00 | 0,0E+00 | 7,1E-08 | 3,1E-09 | 0,0E+00 | 4403276 | 4403659 | 384 | + |
| Y11_RS20120 | Y11_42261 | srmB | ATP-dependent RNA helicase SrmB | -0,48 | -0,19 | -0,31 | 0,29 | 0,16 | -0,13 | 3,9E-07 | 1,2E-03 | 6,7E-06 | 4,2E-05 | 3,6E-04 | 6,7E-03 | 0,0E+00 | 0,0E+00 | 0,0E+00 | 0,0E+00 | 0,0E+00 | 0,0E+00 | 4408598 | 4409923 | 1326 | - |
| Y11_RS20125 | Y11_42271 | | tRNA1(Val) (adenine(37)-N6)-methyltransferase | -0,27 | -1,09 | -1,34 | -0,81 | -1,07 | -0,25 | 5,6E-02 | 7,1E-07 | 1,3E-05 | 3,7E-05 | 1,3E-04 | 7,5E-02 | 0,0E+00 | 2,2E-08 | 2,2E-07 | 0,0E+00 | 3,3E-06 | 0,0E+00 | 4410056 | 4410802 | 747 | + |
| Y11_RS20130 | Y11_42281 | nadB | L-aspartate oxidase | 0,39 | -2,85 | -2,62 | -3,24 | -3,01 | 0,23 | 2,3E-05 | 4,9E-13 | 4,2E-08 | 3,0E-13 | 1,5E-08 | 1,2E-01 | 0,0E+00 | 1,6E-12 | 1,7E-09 | 1,2E-12 | 1,3E-09 | 0,0E+00 | 4410898 | 4412499 | 1602 | - |
| Y11_RS20135 | | rpoE | RNA polymerase sigma factor RpoE | -0,51 | 0,21 | 0,29 | 0,72 | 0,8 | 0,08 | 8,5E-04 | 1,1E-01 | 2,4E-01 | 7,6E-05 | 7,2E-03 | 7,2E-01 | 0,0E+00 | 0,0E+00 | 0,0E+00 | 0,0E+00 | 0,0E+00 | 0,0E+00 | 4412730 | 4413305 | 576 | + |
| Y11_RS20140 | Y11_42301 | rseA | anti-sigma-E factor RseA | -0,79 | -0,16 | -0,45 | 0,63 | 0,34 | -0,28 | 6,7E-05 | 2,0E-01 | 2,2E-02 | 8,8E-04 | 6,5E-02 | 1,3E-01 | 0,0E+00 | 0,0E+00 | 0,0E+00 | 0,0E+00 | 0,0E+00 | 0,0E+00 | 4413330 | 4413986 | 657 | + |
| Y11_RS20145 | Y11_42311 | rseB | sigma-E factor regulatory protein RseB | -0,55 | 1,02 | 1,47 | 1,57 | 2,02 | 0,45 | 3,3E-05 | 2,7E-07 | 9,9E-07 | 1,0E-11 | 2,3E-08 | 1,2E-03 | 0,0E+00 | 9,4E-09 | 2,3E-08 | 5,4E-12 | 1,9E-09 | 0,0E+00 | 4413986 | 4414942 | 957 | + |
| Y11_RS20160 | Y11_42341 | lepA | translation elongation factor 4 | -0,19 | 0,23 | -0,1 | 0,43 | 0,1 | -0,33 | 2,4E-03 | 2,2E-04 | 5,5E-01 | 4,4E-05 | 5,6E-01 | 7,1E-02 | 0,0E+00 | 0,0E+00 | 0,0E+00 | 0,0E+00 | 0,0E+00 | 0,0E+00 | 4416623 | 4418422 | 1800 | + |
| Y11_RS20165 | Y11_42351 | lepB | signal peptidase I | -0,84 | 0,77 | 0,55 | 1,61 | 1,39 | -0,22 | 1,4E-07 | 9,6E-09 | 3,4E-06 | 1,8E-09 | 2,5E-08 | 3,5E-03 | 0,0E+00 | 0,0E+00 | 0,0E+00 | 1,4E-10 | 2,0E-09 | 0,0E+00 | 4418432 | 4419430 | 999 | + |
| Y11_RS20170 | Y11_42361 | rnc | ribonuclease III | -0,37 | 0,06 | -0,29 | 0,43 | 0,08 | -0,35 | 1,2E-03 | 3,3E-01 | 3,7E-02 | 6,6E-04 | 5,4E-01 | 1,9E-02 | 0,0E+00 | 0,0E+00 | 0,0E+00 | 0,0E+00 | 0,0E+00 | 0,0E+00 | 4419676 | 4420356 | 681 | + |
| Y11_RS20175 | Y11_42371 | era | GTPase Era | -0,21 | -0,19 | -0,51 | 0,02 | -0,3 | -0,32 | 6,2E-03 | 2,6E-02 | 8,3E-04 | 8,3E-01 | 2,6E-02 | 2,5E-02 | 0,0E+00 | 0,0E+00 | 0,0E+00 | 0,0E+00 | 0,0E+00 | 0,0E+00 | 4420353 | 4421264 | 912 | + |
| Y11_RS20185 | Y11_42391 | pdxJ | pyridoxine 5'-phosphate synthase | -0,37 | 0,54 | 0,95 | 0,91 | 1,32 | 0,41 | 5,5E-03 | 4,8E-05 | 3,8E-07 | 1,4E-05 | 6,2E-07 | 1,2E-04 | 0,0E+00 | 0,0E+00 | 0,0E+00 | 0,0E+00 | 2,8E-08 | 0,0E+00 | 4422168 | 4422899 | 732 | + |
| Y11_RS20190 | Y11_42401 | acpS | holo-ACP synthase | 0,12 | 1,01 | 0,91 | 0,9 | 0,8 | -0,1 | 7,4E-01 | 9,5E-03 | 5,4E-03 | 6,1E-02 | 6,6E-02 | 7,7E-01 | 0,0E+00 | 1,8E-04 | 0,0E+00 | 0,0E+00 | 0,0E+00 | 0,0E+00 | 4422899 | 4423279 | 381 | + |
| Y11_RS20200 | | | YfhL family 4Fe-4S dicluster ferredoxin | 0 | -15 | 0 | -15 | 0 | 0,96 | 0,0E+00 | 0,0E+00 | 0,0E+00 | 0,0E+00 | 0,0E+00 | 3,2E-02 | 0,0E+00 | 0,0E+00 | 0,0E+00 | 0,0E+00 | 0,0E+00 | 0,0E+00 | 4425312 | 4425572 | 261 | - |
| Y11_RS20205 | Y11_42421 | | MurR/RpiR family transcriptional regulator | -0,65 | 0,96 | 1,1 | 1,6 | 1,75 | 0,14 | 8,1E-08 | 5,4E-07 | 3,8E-05 | 4,4E-09 | 1,0E-06 | 3,4E-01 | 0,0E+00 | 0,0E+00 | 5,6E-07 | 2,6E-10 | 4,2E-08 | 0,0E+00 | 4425634 | 4426476 | 843 | - |
| Y11_RS20210 | Y11_42431 | murQ | N-acetylmuramic acid 6-phosphate etherase | 0,45 | -0,27 | -0,52 | -0,73 | -0,97 | -0,25 | 4,2E-05 | 3,2E-03 | 9,6E-04 | 4,0E-06 | 1,3E-05 | 5,0E-02 | 0,0E+00 | 0,0E+00 | 0,0E+00 | 0,0E+00 | 0,0E+00 | 0,0E+00 | 4426833 | 4427720 | 888 | + |
| Y11_RS20215 | Y11_42441 | yfhb | phosphatidylglycerophosphatase C | 0,03 | -0,05 | 0,01 | -0,07 | -0,02 | 0,05 | 8,9E-01 | 7,9E-01 | 9,7E-01 | 6,5E-01 | 9,0E-01 | 7,1E-01 | 0,0E+00 | 0,0E+00 | 0,0E+00 | 0,0E+00 | 0,0E+00 | 0,0E+00 | 4427828 | 4428511 | 684 | + |
| Y11_RS20220 | Y11_42451 | | DUF2157 domain-containing protein | 0 | -0,34 | 1,99 | -0,97 | 1,44 | 2,33 | 0,0E+00 | 7,2E-01 | 4,3E-02 | 3,1E-01 | 1,2E-01 | 9,0E-02 | 0,0E+00 | 0,0E+00 | 4,8E-04 | 0,0E+00 | 2,5E-03 | 4,6E-03 | 4428588 | 4429637 | 1050 | - |
| Y11_RS20225 | Y11_42461 | tadA | tRNA adenosine(34) deaminase TadA | 0,09 | -0,25 | -0,28 | -0,34 | -0,37 | -0,03 | 2,8E-01 | 3,4E-03 | 7,0E-02 | 5,7E-03 | 4,0E-02 | 8,4E-01 | 0,0E+00 | 0,0E+00 | 0,0E+00 | 0,0E+00 | 0,0E+00 | 0,0E+00 | 4429771 | 4430361 | 591 | + |
| Y11_RS20230 | Y11_42471 | mltF | membrane-bound lytic murein transglycosylase MltF | 0 | 0 | -0,04 | 0 | -0,8 | -0,04 | 0,0E+00 | 0,0E+00 | 8,1E-01 | 0,0E+00 | 1,2E-03 | 8,8E-01 | 0,0E+00 | 0,0E+00 | 0,0E+00 | 0,0E+00 | 0,0E+00 | 0,0E+00 | 4430525 | 4431985 | 1461 | - |
| Y11_RS20235 | Y11_42491 | purL | phosphoribosylformylglycinamidine synthase | 0,97 | 1,56 | 2,34 | 0,59 | 1,37 | 0,78 | 1,4E-07 | 2,5E-08 | 9,8E-11 | 8,1E-06 | 1,4E-10 | 8,4E-07 | 0,0E+00 | 1,3E-09 | 2,3E-11 | 0,0E+00 | 5,9E-11 | 0,0E+00 | 4432300 | 4436190 | 3891 | + |
| Y11_RS20240 | Y11_42501 | | hypothetical protein | 1,07 | 15 | 15 | 0 | 0 | 0 | 4,4E-04 | 0,0E+00 | 0,0E+00 | 0,0E+00 | 0,0E+00 | 0,0E+00 | 1,7E-05 | 0,0E+00 | 0,0E+00 | 0,0E+00 | 0,0E+00 | 0,0E+00 | 4436279 | 4436707 | 429 | - |
| Y11_RS20245 | Y11_42511 | | HAMP domain-containing histidine kinase | 0,36 | 0,54 | 0,83 | 0,18 | 0,47 | 0,28 | 2,9E-03 | 5,7E-04 | 8,1E-05 | 9,7E-02 | 3,0E-03 | 4,7E-02 | 0,0E+00 | 0,0E+00 | 0,0E+00 | 0,0E+00 | 0,0E+00 | 0,0E+00 | 4437848 | 4439284 | 1437 | + |
| Y11_RS20250 | Y11_42521 | qseG | two-component system QseEF-associated lipoprotein QseG | 0,42 | 0,18 | 0,09 | -0,24 | -0,33 | -0,09 | 1,1E-04 | 2,0E-01 | 4,3E-01 | 8,0E-02 | 1,2E-02 | 5,9E-01 | 0,0E+00 | 0,0E+00 | 0,0E+00 | 0,0E+00 | 0,0E+00 | 0,0E+00 | 4439288 | 4440340 | 1053 | + |
| Y11_RS20255 | Y11_42531 | glrR | two-component system response regulator GlrR | 0,27 | 0,76 | 1,05 | 0,49 | 0,78 | 0,29 | 1,5E-04 | 1,7E-06 | 3,3E-05 | 2,7E-05 | 2,4E-04 | 6,2E-02 | 0,0E+00 | 0,0E+00 | 5,0E-07 | 0,0E+00 | 0,0E+00 | 0,0E+00 | 4440303 | 4441640 | 1338 | + |
| Y11_RS20260 | Y11_42541 | | NAD+ synthase | 0,06 | -0,33 | 0,07 | -0,39 | 0,01 | 0,4 | 1,3E-01 | 7,6E-05 | 5,2E-01 | 5,3E-05 | 9,5E-01 | 6,7E-03 | 0,0E+00 | 0,0E+00 | 0,0E+00 | 0,0E+00 | 0,0E+00 | 0,0E+00 | 4441685 | 4443307 | 1623 | + |
| Y11_RS20265 | Y11_42551 | glnB | nitrogen regulatory protein P-II | -0,35 | 0,67 | 0,93 | 1,02 | 1,28 | 0,27 | 1,6E-02 | 1,6E-04 | 1,0E-04 | 1,7E-06 | 4,5E-06 | 3,3E-02 | 0,0E+00 | 0,0E+00 | 0,0E+00 | 4,0E-08 | 1,6E-07 | 0,0E+00 | 4443322 | 4443660 | 339 | + |
| Y11_RS20275 | Y11_42571 | | DUF2620 domain-containing protein | -0,12 | -0,9 | -0,5 | -0,78 | -0,37 | 0,4 | 2,5E-01 | 2,7E-05 | 1,2E-01 | 1,9E-05 | 2,2E-01 | 1,9E-01 | 0,0E+00 | 0,0E+00 | 0,0E+00 | 0,0E+00 | 0,0E+00 | 0,0E+00 | 4445166 | 4445519 | 354 | - |
| Y11_RS20280 | Y11_42581 | | phosphotriesterase-related protein | -0,21 | -1,61 | -0,66 | -1,4 | -0,44 | 0,96 | 1,1E-01 | 2,6E-07 | 2,5E-05 | 3,1E-06 | 2,4E-03 | 2,2E-06 | 0,0E+00 | 9,4E-09 | 0,0E+00 | 6,9E-08 | 0,0E+00 | 0,0E+00 | 4445522 | 4446409 | 888 | - |
| Y11_RS20295 | Y11_42611 | | GntR family transcriptional regulator | 0,28 | -0,26 | -0,26 | -0,54 | -0,54 | -0,01 | 9,1E-02 | 9,8E-02 | 9,3E-02 | 6,9E-03 | 6,6E-03 | 9,7E-01 | 0,0E+00 | 0,0E+00 | 0,0E+00 | 0,0E+00 | 0,0E+00 | 0,0E+00 | 4448216 | 4449115 | 900 | - |
| Y11_RS20300 | Y11_42621 | hmpA | NO-inducible flavohemoprotein | -0,26 | -0,59 | -0,54 | -0,33 | -0,28 | 0,05 | 5,7E-02 | 8,8E-04 | 3,7E-05 | 6,4E-02 | 5,2E-02 | 6,6E-01 | 0,0E+00 | 0,0E+00 | 0,0E+00 | 0,0E+00 | 0,0E+00 | 0,0E+00 | 4449379 | 4450569 | 1191 | - |
| Y11_RS20305 | Y11_42631 | | serine hydroxymethyltransferase | -0,16 | 0,87 | 1,4 | 1,04 | 1,56 | 0,52 | 1,4E-02 | 6,3E-08 | 9,9E-09 | 2,1E-08 | 4,9E-09 | 1,3E-05 | 0,0E+00 | 0,0E+00 | 5,2E-10 | 9,2E-10 | 5,4E-10 | 0,0E+00 | 4450957 | 4452210 | 1254 | + |
| Y11_RS20320 | Y11_42661 | | DUF1007 family protein | 0,16 | -1,78 | -1,06 | -1,94 | -1,21 | 0,73 | 6,1E-02 | 5,0E-09 | 4,3E-05 | 2,3E-09 | 1,5E-05 | 5,1E-04 | 0,0E+00 | 3,7E-10 | 6,3E-07 | 1,6E-10 | 4,8E-07 | 0,0E+00 | 4454958 | 4455632 | 675 | + |
| Y11_RS20325 | Y11_42671 | | nickel/cobalt transporter | 1,12 | 2,34 | 2,33 | 1,3 | 1,24 | -0,04 | 2,3E-01 | 1,7E-02 | 1,8E-02 | 7,5E-03 | 1,3E-02 | 8,2E-01 | 7,4E-03 | 3,1E-04 | 2,1E-04 | 1,2E-04 | 2,8E-04 | 0,0E+00 | 4455623 | 4456639 | 1017 | + |
| Y11_RS20330 | Y11_42681 | suhB | inositol-1-monophosphatase | -0,13 | -0,24 | -0,97 | -0,11 | -0,84 | -0,72 | 5,6E-02 | 9,1E-03 | 2,5E-05 | 6,7E-02 | 3,5E-05 | 1,6E-04 | 0,0E+00 | 0,0E+00 | 0,0E+00 | 0,0E+00 | 0,0E+00 | 0,0E+00 | 4456847 | 4457653 | 807 | - |
| Y11_RS20335 | Y11_42691 | trmJ | tRNA (cytosine(32)/uridine(32)-2'-O)-methyltransferase TrmJ | -0,05 | -0,17 | -0,37 | -0,12 | -0,32 | -0,2 | 6,1E-01 | 4,1E-03 | 1,0E-02 | 2,2E-01 | 4,6E-02 | 1,2E-01 | 0,0E+00 | 0,0E+00 | 0,0E+00 | 0,0E+00 | 0,0E+00 | 0,0E+00 | 4457897 | 4458670 | 774 | + |
| Y11_RS20340 | Y11_42701 | iscR | Fe-S cluster assembly transcriptional regulator IscR | -0,43 | -0,32 | 1,17 | 0,1 | 1,59 | 1,49 | 9,6E-05 | 6,0E-04 | 5,2E-08 | 2,5E-02 | 1,3E-10 | 2,0E-10 | 0,0E+00 | 0,0E+00 | 1,9E-09 | 0,0E+00 | 5,7E-11 | 1,0E-09 | 4458827 | 4459321 | 495 | + |
| Y11_RS20345 | Y11_42711 | | IscS subfamily cysteine desulfurase | 0 | -0,01 | -0,4 | -0,01 | -0,39 | -0,38 | 9,5E-01 | 8,4E-01 | 4,8E-05 | 9,1E-01 | 6,4E-04 | 8,0E-04 | 0,0E+00 | 0,0E+00 | 0,0E+00 | 0,0E+00 | 0,0E+00 | 0,0E+00 | 4459390 | 4460604 | 1215 | + |
| Y11_RS20350 | Y11_42721 | iscU | Fe-S cluster assembly scaffold IscU | 0,07 | -0,47 | -0,94 | -0,54 | -1,01 | -0,46 | 3,2E-01 | 4,4E-05 | 5,7E-05 | 9,6E-06 | 3,1E-05 | 4,2E-03 | 0,0E+00 | 0,0E+00 | 0,0E+00 | 0,0E+00 | 8,8E-07 | 0,0E+00 | 4460635 | 4461021 | 387 | + |
| Y11_RS20355 | Y11_42731 | iscA | iron-sulfur cluster assembly protein IscA | 0,03 | -1,95 | -1,71 | -1,98 | -1,74 | 0,25 | 8,3E-01 | 2,9E-08 | 9,2E-05 | 1,9E-07 | 1,1E-04 | 3,1E-01 | 0,0E+00 | 1,5E-09 | 1,3E-06 | 5,7E-09 | 3,0E-06 | 0,0E+00 | 4461125 | 4461448 | 324 | + |
| Y11_RS20360 | Y11_42741 | hscB | co-chaperone HscB | -0,09 | -0,36 | -0,18 | -0,26 | -0,08 | 0,18 | 3,3E-01 | 4,4E-03 | 2,2E-02 | 5,2E-02 | 4,1E-01 | 9,1E-02 | 0,0E+00 | 0,0E+00 | 0,0E+00 | 0,0E+00 | 0,0E+00 | 0,0E+00 | 4461556 | 4462083 | 528 | + |
| Y11_RS20365 | Y11_42751 | hscA | Fe-S protein assembly chaperone HscA | -0,21 | -0,57 | -0,62 | -0,35 | -0,4 | -0,05 | 3,6E-03 | 6,6E-07 | 2,6E-07 | 6,9E-05 | 2,2E-05 | 1,5E-01 | 0,0E+00 | 0,0E+00 | 0,0E+00 | 0,0E+00 | 0,0E+00 | 0,0E+00 | 4462266 | 4464116 | 1851 | + |
| Y11_RS20370 | Y11_42761 | fdx | ISC system 2Fe-2S type ferredoxin | -0,3 | -0,33 | -0,17 | -0,03 | 0,13 | 0,16 | 2,6E-02 | 1,9E-03 | 1,5E-01 | 7,4E-01 | 3,1E-01 | 9,7E-02 | 0,0E+00 | 0,0E+00 | 0,0E+00 | 0,0E+00 | 0,0E+00 | 0,0E+00 | 4464119 | 4464454 | 336 | + |
| Y11_RS20375 | Y11_42771 | iscX | Fe-S cluster assembly protein IscX | -0,47 | -1,25 | -1,01 | -0,78 | -0,54 | 0,24 | 1,8E-03 | 7,3E-08 | 4,9E-08 | 8,6E-05 | 5,6E-04 | 4,4E-03 | 0,0E+00 | 3,1E-09 | 1,8E-09 | 0,0E+00 | 0,0E+00 | 0,0E+00 | 4464475 | 4464675 | 201 | + |
| Y11_RS20380 | Y11_42781 | pepB | aminopeptidase PepB | -0,24 | 0,9 | 0,77 | 1,14 | 1,01 | -0,12 | 9,7E-04 | 3,3E-08 | 1,2E-04 | 2,7E-09 | 1,6E-05 | 2,9E-01 | 0,0E+00 | 0,0E+00 | 0,0E+00 | 1,8E-10 | 4,9E-07 | 0,0E+00 | 4464829 | 4466124 | 1296 | + |
| Y11_RS20385 | Y11_42791 | sseB | enhanced serine sensitivity protein SseB | -0,23 | -0,67 | -1,14 | -0,43 | -0,91 | -0,47 | 1,6E-01 | 4,9E-04 | 5,7E-06 | 2,3E-02 | 2,6E-04 | 3,0E-03 | 0,0E+00 | 0,0E+00 | 1,0E-07 | 0,0E+00 | 0,0E+00 | 0,0E+00 | 4466281 | 4467168 | 888 | + |
| Y11_RS20390 | Y11_42801 | | autotransporter outer membrane beta-barrel domain-containing protein | 0 | 0,64 | 0,47 | 0,25 | 0,12 | -0,11 | 0,0E+00 | 8,3E-02 | 2,4E-01 | 2,9E-01 | 7,0E-01 | 7,6E-01 | 0,0E+00 | 0,0E+00 | 0,0E+00 | 0,0E+00 | 0,0E+00 | 0,0E+00 | 4467248 | 4469773 | 2526 | - |

| | | | | | | | | | | | | | | | | | | | | | | | | | |
|---|---|---|---|---|---|---|---|---|---|---|---|---|---|---|---|---|---|---|---|---|---|---|---|---|---|
| Y11_RS20400 | Y11_42841 | | alpha-2-macroglobulin family protein | 0,57 | 0,44 | 0,93 | -0,13 | 0,35 | 0,48 | 5,9E-05 | 5,3E-05 | 1,6E-04 | 1,9E-01 | 5,5E-02 | 1,2E-02 | 0,0E+00 | 0,0E+00 | 0,0E+00 | 0,0E+00 | 0,0E+00 | 0,0E+00 | 4471909 | 4476945 | 5037 | + |
| Y11_RS20415 | Y11_42871 | ndk | nucleoside-diphosphate kinase | -0,03 | -0,53 | -0,43 | -0,51 | -0,4 | 0,11 | 7,5E-01 | 1,9E-03 | 3,6E-03 | 5,1E-03 | 1,1E-02 | 4,8E-01 | 0,0E+00 | 0,0E+00 | 0,0E+00 | 0,0E+00 | 0,0E+00 | 0,0E+00 | 4482579 | 4483007 | 429 | + |
| Y11_RS20420 | Y11_42881 | | bifunctional tRNA (adenosine(37)-C2)-methyltransferase TrmG/ribosomal RNA large subunit methyltransferase RlmN | 0,11 | -1,27 | -2,92 | -1,38 | -3,03 | -1,65 | 1,8E-01 | 2,3E-06 | 5,2E-10 | 7,9E-07 | 1,8E-10 | 3,4E-07 | 0,0E+00 | 6,4E-08 | 6,6E-11 | 2,0E-08 | 6,3E-11 | 8,0E-08 | 4483275 | 4484441 | 1167 | + |
| Y11_RS20425 | Y11_42891 | pilW | type IV pilus biogenesis/stability protein PilW | -0,28 | -3,96 | -3,21 | -3,68 | -2,93 | 0,75 | 2,2E-01 | 2,6E-09 | 3,9E-07 | 2,0E-08 | 1,6E-06 | 2,8E-03 | 0,0E+00 | 2,2E-10 | 1,0E-08 | 9,0E-10 | 6,1E-08 | 0,0E+00 | 4484635 | 4485384 | 750 | + |
| Y11_RS20430 | Y11_42901 | rodZ | cytoskeleton protein RodZ | -0,19 | 0,24 | -0,28 | 0,43 | -0,09 | -0,52 | 3,2E-03 | 7,9E-04 | 3,2E-03 | 2,2E-05 | 2,3E-01 | 7,6E-05 | 0,0E+00 | 0,0E+00 | 0,0E+00 | 0,0E+00 | 0,0E+00 | 0,0E+00 | 4485374 | 4486390 | 1017 | + |
| Y11_RS20435 | Y11_42911 | ispG | flavodoxin-dependent (E)-4-hydroxy-3-methylbut-2-enyl-diphosphate synthase | 0,32 | -0,04 | -0,61 | -0,36 | -0,93 | -0,57 | 3,8E-04 | 3,9E-01 | 2,9E-06 | 1,6E-05 | 5,7E-08 | 2,6E-07 | 0,0E+00 | 0,0E+00 | 0,0E+00 | 0,0E+00 | 0,0E+00 | 0,0E+00 | 4486511 | 4487635 | 1125 | + |
| Y11_RS20440 | Y11_42931 | hisS | histidine--tRNA ligase | -0,28 | 0,05 | -0,25 | 0,33 | 0,03 | -0,3 | 7,2E-06 | 6,1E-02 | 7,4E-02 | 4,4E-07 | 8,1E-01 | 3,6E-02 | 0,0E+00 | 0,0E+00 | 0,0E+00 | 0,0E+00 | 0,0E+00 | 0,0E+00 | 4487852 | 4489126 | 1275 | + |
| Y11_RS20445 | Y11_42941 | | YfgM family protein | -0,3 | 0,05 | -0,07 | 0,35 | 0,23 | -0,12 | 2,3E-04 | 5,7E-01 | 3,5E-01 | 1,9E-03 | 1,2E-02 | 2,5E-01 | 0,0E+00 | 0,0E+00 | 0,0E+00 | 0,0E+00 | 0,0E+00 | 0,0E+00 | 4489140 | 4489760 | 621 | + |
| Y11_RS20450 | Y11_42951 | bamB | outer membrane protein assembly factor BamB | -0,12 | 0,5 | 0,5 | 0,61 | 0,62 | 0 | 9,7E-02 | 9,9E-06 | 9,8E-04 | 1,6E-06 | 2,4E-04 | 9,7E-01 | 0,0E+00 | 0,0E+00 | 0,0E+00 | 0,0E+00 | 0,0E+00 | 0,0E+00 | 4489772 | 4490953 | 1182 | + |
| Y11_RS20460 | Y11_42971 | der | ribosome biogenesis GTPase Der | -0,7 | -0,01 | -0,85 | 0,69 | -0,15 | -0,84 | 1,9E-06 | 9,1E-01 | 6,8E-07 | 4,6E-05 | 5,8E-02 | 1,3E-05 | 0,0E+00 | 0,0E+00 | 0,0E+00 | 0,0E+00 | 0,0E+00 | 0,0E+00 | 4491153 | 4492637 | 1485 | + |
| Y11_RS20470 | Y11_43001 | | zinc ribbon domain-containing protein | -0,21 | 0,79 | 0,5 | 1 | 0,71 | -0,29 | 5,3E-01 | 2,4E-02 | 1,2E-01 | 1,4E-03 | 1,2E-02 | 1,1E-01 | 0,0E+00 | 0,0E+00 | 0,0E+00 | 0,0E+00 | 0,0E+00 | 0,0E+00 | 4493908 | 4494132 | 225 | + |
| Y11_RS20475 | Y11_43011 | xseA | exodeoxyribonuclease VII large subunit | 0,32 | 0,19 | 0,1 | -0,12 | -0,22 | -0,09 | 1,7E-03 | 4,2E-02 | 5,5E-01 | 1,9E-01 | 2,0E-01 | 5,8E-01 | 0,0E+00 | 0,0E+00 | 0,0E+00 | 0,0E+00 | 0,0E+00 | 0,0E+00 | 4494176 | 4495552 | 1377 | - |
| Y11_RS20480 | Y11_43021 | guaB | IMP dehydrogenase | -0,18 | 0,79 | 0,61 | 0,97 | 0,79 | -0,18 | 1,4E-02 | 2,2E-08 | 1,6E-03 | 1,1E-07 | 4,3E-04 | 2,1E-01 | 0,0E+00 | 0,0E+00 | 0,0E+00 | 0,0E+00 | 0,0E+00 | 0,0E+00 | 4495721 | 4497184 | 1464 | + |
| Y11_RS20485 | Y11_43041 | guaA | glutamine-hydrolyzing GMP synthase | -0,39 | 0,41 | 0,65 | 0,81 | 1,05 | 0,24 | 1,9E-04 | 6,2E-04 | 1,5E-04 | 8,6E-06 | 6,5E-06 | 6,3E-02 | 0,0E+00 | 0,0E+00 | 0,0E+00 | 0,0E+00 | 2,2E-07 | 0,0E+00 | 4497371 | 4498948 | 1578 | + |
| Y11_RS20520 | Y11_43141 | thiD | bifunctional hydroxymethylpyrimidine kinase/phosphomethylpyrimidine kinase | -0,36 | 1 | 1,24 | 1,36 | 1,6 | 0,25 | 1,9E-02 | 1,0E-05 | 4,7E-06 | 3,5E-06 | 1,9E-06 | 5,9E-02 | 0,0E+00 | 0,0E+00 | 8,5E-08 | 7,7E-08 | 7,3E-08 | 0,0E+00 | 4507439 | 4508239 | 801 | + |
| Y11_RS20530 | Y11_43161 | | inhibitor of vertebrate lysozyme family protein | 1,09 | 0,46 | 0,6 | -0,63 | -0,49 | 0,14 | 1,1E-03 | 1,5E-04 | 4,1E-03 | 2,0E-02 | 9,3E-02 | 3,7E-01 | 4,2E-05 | 0,0E+00 | 0,0E+00 | 0,0E+00 | 0,0E+00 | 0,0E+00 | 4509431 | 4509976 | 546 | + |
| Y11_RS20535 | Y11_43171 | gutQ | arabinose-5-phosphate isomerase GutQ | -0,2 | 0,39 | 0,91 | 0,58 | 1,11 | 0,52 | 4,1E-02 | 3,1E-03 | 1,8E-05 | 1,2E-03 | 2,2E-05 | 4,5E-03 | 0,0E+00 | 0,0E+00 | 0,0E+00 | 0,0E+00 | 6,7E-07 | 0,0E+00 | 4510056 | 4511021 | 966 | - |
| Y11_RS20540 | Y11_43181 | | DNA-binding transcriptional repressor | -0,16 | 0,25 | 0,79 | 0,41 | 0,95 | 0,54 | 1,9E-01 | 3,0E-02 | 4,4E-05 | 6,8E-03 | 3,9E-05 | 5,6E-04 | 0,0E+00 | 0,0E+00 | 0,0E+00 | 0,0E+00 | 0,0E+00 | 0,0E+00 | 4511014 | 4511784 | 771 | - |
| Y11_RS20550 | Y11_43201 | srlD | sorbitol-6-phosphate dehydrogenase | 0,26 | 2,74 | 3,37 | 2,48 | 3,11 | 0,63 | 3,3E-01 | 2,8E-08 | 7,5E-09 | 1,4E-05 | 2,8E-06 | 5,9E-03 | 0,0E+00 | 1,4E-09 | 4,3E-10 | 2,8E-07 | 1,0E-07 | 0,0E+00 | 4512591 | 4513370 | 780 | - |
| Y11_RS20560 | Y11_43221 | | PTS glucitol/sorbitol transporter subunit IIB | -0,27 | 2,83 | 3,8 | 3,1 | 4,07 | 0,97 | 4,7E-03 | 1,1E-08 | 3,8E-08 | 1,7E-09 | 1,4E-08 | 1,4E-03 | 0,0E+00 | 6,8E-10 | 1,6E-09 | 1,3E-10 | 1,3E-09 | 0,0E+00 | 4513945 | 4514970 | 1026 | - |
| Y11_RS20570 | Y11_43241 | yegS | lipid kinase YegS | 0,21 | 1,07 | 1,79 | 0,86 | 1,58 | 0,72 | 3,8E-01 | 8,8E-05 | 2,6E-06 | 2,7E-03 | 5,7E-05 | 2,5E-04 | 0,0E+00 | 1,9E-06 | 5,1E-08 | 0,0E+00 | 1,6E-06 | 0,0E+00 | 4515933 | 4516823 | 891 | - |
| Y11_RS20575 | Y11_43251 | yegQ | tRNA 5-hydroxyuridine modification protein YegQ | 0,39 | -1,23 | -1,85 | -1,62 | -2,24 | -0,62 | 1,1E-04 | 7,9E-09 | 3,6E-08 | 3,3E-10 | 6,1E-09 | 7,7E-05 | 0,0E+00 | 5,1E-10 | 1,5E-09 | 3,8E-11 | 6,5E-10 | 0,0E+00 | 4517306 | 4518694 | 1389 | - |
| Y11_RS20580 | Y11_43261 | | YegP family protein | 0,57 | 3,38 | 5,61 | 2,81 | 5,05 | 2,24 | 1,5E-04 | 1,3E-08 | 2,7E-13 | 1,6E-07 | 3,1E-11 | 5,8E-07 | 0,0E+00 | 7,7E-10 | 9,5E-13 | 5,0E-09 | 2,0E-11 | 1,2E-07 | 4518933 | 4519271 | 339 | - |
| Y11_RS20585 | Y11_43271 | baeR | two-component system response regulator BaeR | 0,07 | 0,03 | 0,3 | -0,03 | 0,24 | 0,27 | 7,5E-01 | 7,8E-01 | 1,6E-01 | 8,5E-01 | 3,3E-01 | 1,5E-01 | 0,0E+00 | 0,0E+00 | 0,0E+00 | 0,0E+00 | 0,0E+00 | 0,0E+00 | 4519350 | 4520069 | 720 | - |
| Y11_RS20590 | Y11_43281 | baeS | two-component system sensor histidine kinase BaeS | -0,24 | 0,13 | 0,75 | 0,37 | 0,99 | 0,62 | 1,5E-02 | 7,0E-02 | 5,7E-04 | 1,6E-04 | 7,2E-05 | 1,2E-03 | 0,0E+00 | 0,0E+00 | 0,0E+00 | 0,0E+00 | 0,0E+00 | 0,0E+00 | 4520078 | 4521454 | 1377 | - |
| Y11_RS20600 | Y11_43311 | mdtC | multidrug efflux RND transporter permease subunit MdtC | -0,16 | -0,51 | -0,37 | -0,34 | -0,21 | 0,14 | 3,0E-01 | 1,1E-03 | 1,5E-01 | 7,4E-02 | 4,6E-01 | 5,9E-01 | 0,0E+00 | 0,0E+00 | 0,0E+00 | 0,0E+00 | 0,0E+00 | 0,0E+00 | 4523024 | 4526098 | 3075 | - |
| Y11_RS20605 | Y11_43321 | | MdtB/MuxB family multidrug efflux RND transporter permease subunit | -0,06 | 0,11 | 0,28 | 0,17 | 0,34 | 0,17 | 5,9E-01 | 3,5E-01 | 9,8E-02 | 1,9E-01 | 6,1E-02 | 3,0E-01 | 0,0E+00 | 0,0E+00 | 0,0E+00 | 0,0E+00 | 0,0E+00 | 0,0E+00 | 4526095 | 4529238 | 3144 | - |
| Y11_RS20610 | Y11_43331 | | MdtA/MuxA family multidrug efflux RND transporter periplasmic adaptor subunit | -0,53 | -0,64 | -0,04 | -0,12 | 0,49 | 0,61 | 3,2E-04 | 4,7E-05 | 8,5E-01 | 1,5E-01 | 3,3E-02 | 1,2E-02 | 0,0E+00 | 0,0E+00 | 0,0E+00 | 0,0E+00 | 0,0E+00 | 0,0E+00 | 4529238 | 4530572 | 1335 | - |
| Y11_RS20620 | Y11_43341 | yegD | molecular chaperone | 0 | -2,72 | -4,33 | -2,85 | -4,46 | -1,61 | 0,0E+00 | 5,6E-10 | 3,4E-10 | 9,1E-09 | 1,5E-09 | 9,2E-07 | 0,0E+00 | 7,2E-11 | 4,8E-11 | 4,9E-10 | 2,3E-10 | 1,7E-07 | 4531584 | 4532936 | 1353 | - |
| Y11_RS20630 | Y11_43361 | pstB | phosphate ABC transporter ATP-binding protein PstB | 0,01 | -0,57 | -0,17 | -0,58 | -0,18 | 0,4 | 9,5E-01 | 1,6E-02 | 4,5E-01 | 1,0E-02 | 4,0E-01 | 1,4E-01 | 0,0E+00 | 0,0E+00 | 0,0E+00 | 0,0E+00 | 0,0E+00 | 0,0E+00 | 4533811 | 4534623 | 813 | - |
| Y11_RS20640 | Y11_43381 | | ABC transporter permease subunit | 0 | -0,95 | -0,54 | -0,03 | 0,33 | 0,41 | 0,0E+00 | 8,8E-03 | 2,1E-02 | 9,1E-01 | 1,1E-01 | 2,4E-01 | 0,0E+00 | 0,0E+00 | 0,0E+00 | 0,0E+00 | 0,0E+00 | 0,0E+00 | 4536316 | 4538502 | 2187 | - |
| Y11_RS20645 | Y11_43391 | ppk1 | polyphosphate kinase 1 | 0,71 | 0,16 | 1,27 | -0,56 | 0,56 | 1,11 | 5,2E-08 | 7,2E-04 | 3,8E-07 | 1,1E-08 | 1,5E-04 | 6,6E-07 | 0,0E+00 | 0,0E+00 | 1,0E-08 | 0,0E+00 | 0,0E+00 | 1,4E-07 | 4538748 | 4540817 | 2070 | + |
| Y11_RS20650 | Y11_43401 | ppx | exopolyphosphatase | 0,38 | -0,29 | -0,03 | -0,68 | -0,42 | 0,26 | 1,0E-05 | 7,1E-05 | 2,6E-01 | 1,2E-06 | 1,2E-05 | 3,1E-04 | 0,0E+00 | 0,0E+00 | 0,0E+00 | 0,0E+00 | 0,0E+00 | 0,0E+00 | 4540822 | 4542384 | 1563 | + |
| Y11_RS20655 | Y11_43411 | | DUF2633 family protein | 0,18 | 0,09 | 0,79 | -0,09 | 0,6 | 0,69 | 2,5E-01 | 6,4E-01 | 1,4E-03 | 7,2E-01 | 2,1E-02 | 2,3E-02 | 0,0E+00 | 0,0E+00 | 0,0E+00 | 0,0E+00 | 0,0E+00 | 0,0E+00 | 4542447 | 4542638 | 192 | + |
| Y11_RS20660 | Y11_43421 | mgtE | magnesium transporter | 0,19 | -1,31 | -0,65 | -1,5 | -0,84 | 0,66 | 3,0E-02 | 4,0E-07 | 8,1E-06 | 2,2E-07 | 2,7E-06 | 7,6E-05 | 0,0E+00 | 1,3E-08 | 0,0E+00 | 6,7E-09 | 0,0E+00 | 0,0E+00 | 4542686 | 4544161 | 1476 | + |
| Y11_RS20670 | Y11_43441 | speG | spermidine N1-acetyltransferase | 0,93 | 1,34 | 1,64 | 0,4 | 0,71 | 0,31 | 1,3E-03 | 2,0E-05 | 3,7E-06 | 6,6E-02 | 5,2E-03 | 5,9E-02 | 0,0E+00 | 4,7E-07 | 7,0E-08 | 0,0E+00 | 0,0E+00 | 0,0E+00 | 4548674 | 4549219 | 546 | - |
| Y11_RS20675 | Y11_43451 | purN | phosphoribosylglycinamide formyltransferase | 1 | 2,14 | 2,89 | 1,14 | 1,88 | 0,72 | 1,6E-03 | 1,4E-06 | 1,8E-07 | 9,4E-04 | 3,6E-05 | 4,6E-03 | 5,8E-05 | 3,9E-08 | 5,3E-09 | 1,6E-05 | 1,0E-06 | 0,0E+00 | 4549355 | 4549993 | 639 | - |
| Y11_RS20680 | Y11_43461 | purM | phosphoribosylformylglycinamidine cyclo-ligase | 0,64 | 2,25 | 3,17 | 1,62 | 2,54 | 0,92 | 4,1E-07 | 5,5E-09 | 1,4E-08 | 7,4E-08 | 8,2E-08 | 4,0E-04 | 0,0E+00 | 4,0E-10 | 6,9E-10 | 2,6E-09 | 5,3E-09 | 0,0E+00 | 4550034 | 4551077 | 1044 | - |
| Y11_RS20685 | Y11_43471 | upp | uracil phosphoribosyltransferase | 0,44 | 0,72 | 1,11 | 0,28 | 0,67 | 0,39 | 1,7E-03 | 1,7E-04 | 2,1E-05 | 3,5E-02 | 7,1E-04 | 2,1E-02 | 0,0E+00 | 0,0E+00 | 3,3E-07 | 0,0E+00 | 0,0E+00 | 0,0E+00 | 4551311 | 4551937 | 627 | + |
| Y11_RS20690 | Y11_43481 | uraA | uracil permease | 0,82 | 1,02 | 1,58 | 0,2 | 0,75 | 0,55 | 2,5E-01 | 1,7E-01 | 4,4E-02 | 3,8E-01 | 3,6E-03 | 3,8E-02 | 0,0E+00 | 3,1E-03 | 4,9E-04 | 0,0E+00 | 0,0E+00 | 0,0E+00 | 4552018 | 4553307 | 1290 | + |
| Y11_RS20730 | Y11_p0061 | traT | complement resistance protein TraT | 4,72 | 15 | 15 | 0 | 0 | 0 | 1,2E-08 | 0,0E+00 | 0,0E+00 | 0,0E+00 | 0,0E+00 | 0,0E+00 | 1,5E-09 | 0,0E+00 | 0,0E+00 | 0,0E+00 | 0,0E+00 | 0,0E+00 | 8791 | 9528 | 738 | - |
| Y11_RS20740 | Y11_p0081 | yopK | type III secretion system effector YopK | 15 | 1,65 | 15 | 0 | 0 | 0 | 0,0E+00 | 5,3E-04 | 0,0E+00 | 0,0E+00 | 0,0E+00 | 0,0E+00 | 0,0E+00 | 1,1E-05 | 0,0E+00 | 0,0E+00 | 0,0E+00 | 0,0E+00 | 10010 | 10558 | 549 | - |
| Y11_RS20770 | Y11_p0171 | lcrQ | type III secretion system exported negative regulator LcrQ/YscM1 | 15 | 15 | 15 | 0 | 0 | 0 | 0,0E+00 | 0,0E+00 | 0,0E+00 | 0,0E+00 | 0,0E+00 | 0,0E+00 | 0,0E+00 | 0,0E+00 | 0,0E+00 | 0,0E+00 | 0,0E+00 | 0,0E+00 | 14297 | 14647 | 351 | + |
| Y11_RS20790 | Y11_p0221 | yadA | trimeric autotransporter adhesin YadA | 6,34 | 6,44 | 6,55 | 0,09 | 0 | 0,11 | 1,4E-10 | 6,7E-11 | 3,4E-11 | 5,5E-01 | 0,0E+00 | 3,8E-01 | 4,1E-11 | 1,7E-11 | 1,3E-11 | 0,0E+00 | 0,0E+00 | 0,0E+00 | 16908 | 18320 | 1413 | + |
| Y11_RS20865 | Y11_p0411 | sopA | plasmid-partitioning protein SopA | 15 | 5,24 | 4,16 | 0 | 0 | 0 | 0,0E+00 | 4,0E-11 | 2,3E-09 | 0,0E+00 | 0,0E+00 | 0,0E+00 | 0,0E+00 | 1,3E-11 | 1,8E-10 | 0,0E+00 | 0,0E+00 | 0,0E+00 | 28080 | 29246 | 1167 | + |
| Y11_RS20870 | Y11_p0421 | | ParB/RepB/Spo0J family plasmid partition protein | 15 | 4,51 | 15 | 0 | 0 | 0 | 0,0E+00 | 2,2E-07 | 0,0E+00 | 0,0E+00 | 0,0E+00 | 0,0E+00 | 0,0E+00 | 8,0E-09 | 0,0E+00 | 0,0E+00 | 0,0E+00 | 0,0E+00 | 29243 | 30208 | 966 | + |
| Y11_RS20885 | Y11_p0471 | yopE | type III secretion system effector GTPase activator YopE | 4,33 | 4,44 | 5,04 | 0 | 0 | 0 | 6,8E-13 | 1,0E-13 | 6,0E-11 | 0,0E+00 | 0,0E+00 | 0,0E+00 | 1,8E-12 | 9,0E-13 | 1,8E-11 | 0,0E+00 | 0,0E+00 | 0,0E+00 | 31765 | 32424 | 660 | - |
| Y11_RS20890 | Y11_p0481 | sycE | type III secretion system chaperone SycE/YerA | 15 | 4,4 | 15 | 0 | 0 | 0 | 0,0E+00 | 1,6E-07 | 0,0E+00 | 0,0E+00 | 0,0E+00 | 0,0E+00 | 0,0E+00 | 6,1E-09 | 0,0E+00 | 0,0E+00 | 0,0E+00 | 0,0E+00 | 32617 | 33009 | 393 | + |
| Y11_RS20905 | Y11_p0531 | sycH | type III secretion system chaperone SycH | 15 | 6,02 | 15 | 0 | 0 | 0 | 0,0E+00 | 4,6E-12 | 0,0E+00 | 0,0E+00 | 0,0E+00 | 0,0E+00 | 0,0E+00 | 4,0E-12 | 0,0E+00 | 0,0E+00 | 0,0E+00 | 0,0E+00 | 34622 | 35047 | 426 | + |
| Y11_RS20910 | Y11_p0541 | yopH | T3SS effector protein-tyrosine-phosphatase YopH | 2,77 | 3,05 | 2,78 | 0,29 | 0,01 | -0,27 | 1,0E-08 | 3,0E-08 | 2,8E-09 | 9,8E-02 | 8,9E-01 | 9,0E-02 | 1,4E-09 | 1,5E-09 | 2,1E-10 | 0,0E+00 | 0,0E+00 | 0,0E+00 | 35275 | 36681 | 1407 | - |
| Y11_RS20915 | Y11_p0551 | lcrQ | type III secretion system exported negative regulator LcrQ/YscM1 | 3,97 | 15 | 15 | 15 | 15 | 0 | 7,0E-08 | 0,0E+00 | 0,0E+00 | 0,0E+00 | 0,0E+00 | 0,0E+00 | 6,8E-09 | 0,0E+00 | 0,0E+00 | 0,0E+00 | 0,0E+00 | 0,0E+00 | 37366 | 37713 | 348 | - |
| Y11_RS20945 | Y11_p0611 | yscG | type III secretion system chaperone YscG | 15 | 15 | 15 | 0 | 0 | 0 | 0,0E+00 | 0,0E+00 | 0,0E+00 | 0,0E+00 | 0,0E+00 | 0,0E+00 | 0,0E+00 | 0,0E+00 | 0,0E+00 | 0,0E+00 | 0,0E+00 | 0,0E+00 | 40761 | 41108 | 348 | - |
| Y11_RS20955 | Y11_p0631 | | EscE/YscE/SsaE family type III secretion system needle protein co-chaperone | 15 | 15 | 15 | 0 | 0 | 0 | 0,0E+00 | 0,0E+00 | 0,0E+00 | 0,0E+00 | 0,0E+00 | 0,0E+00 | 0,0E+00 | 0,0E+00 | 0,0E+00 | 0,0E+00 | 0,0E+00 | 0,0E+00 | 41370 | 41573 | 204 | - |
| Y11_RS20965 | Y11_p0661 | yscC | SctC family type III secretion system outer membrane ring subunit YscC | 3,36 | 15 | 15 | 0 | 0 | 0 | 5,1E-07 | 0,0E+00 | 0,0E+00 | 0,0E+00 | 0,0E+00 | 0,0E+00 | 3,6E-08 | 0,0E+00 | 0,0E+00 | 0,0E+00 | 0,0E+00 | 0,0E+00 | 42823 | 44646 | 1824 | - |
| Y11_RS20970 | Y11_p0671 | yscB | type III secretion system chaperone YscB | 15 | 15 | 3,53 | 0 | 0 | 0 | 0,0E+00 | 0,0E+00 | 5,9E-09 | 0,0E+00 | 0,0E+00 | 0,0E+00 | 0,0E+00 | 0,0E+00 | 3,6E-10 | 0,0E+00 | 0,0E+00 | 0,0E+00 | 44652 | 45065 | 414 | - |
| Y11_RS20980 | Y11_p0691 | virF | virulence regulon transcriptional activator VirF | 15 | 3,73 | 3,69 | 0 | 0 | 0 | 0,0E+00 | 3,4E-07 | 2,7E-11 | 0,0E+00 | 0,0E+00 | 0,0E+00 | 0,0E+00 | 1,1E-08 | 1,2E-11 | 0,0E+00 | 0,0E+00 | 0,0E+00 | 45468 | 46283 | 816 | - |
| Y11_RS20985 | Y11_p0701 | yscW | type III secretion system pilotin YscW | 15 | 5,06 | 15 | 0 | 0 | 0 | 0,0E+00 | 1,2E-08 | 0,0E+00 | 0,0E+00 | 0,0E+00 | 0,0E+00 | 0,0E+00 | 6,9E-10 | 0,0E+00 | 0,0E+00 | 0,0E+00 | 0,0E+00 | 46407 | 46802 | 396 | - |
| Y11_RS21010 | Y11_p0751 | yscQ | SctQ family type III secretion system cytoplasmic ring protein YscQ | 5,58 | 5,58 | 5,91 | -0,01 | 0,34 | 0 | 2,7E-09 | 2,3E-09 | 1,3E-09 | 8,8E-01 | 1,1E-04 | 0,0E+00 | 5,3E-10 | 1,9E-10 | 1,2E-10 | 0,0E+00 | 0,0E+00 | 0,0E+00 | 50142 | 51065 | 924 | - |
| Y11_RS21015 | Y11_p0761 | | type III secretion system needle length determinant | 15 | 15 | 15 | 0 | 0 | 0 | 0,0E+00 | 0,0E+00 | 0,0E+00 | 0,0E+00 | 0,0E+00 | 0,0E+00 | 0,0E+00 | 0,0E+00 | 0,0E+00 | 0,0E+00 | 0,0E+00 | 0,0E+00 | 51062 | 52609 | 1548 | - |
| Y11_RS21025 | Y11_p0781 | yscN | SctN family type III secretion system ATPase YscN | 15 | 15 | 15 | 0 | 0 | 0 | 0,0E+00 | 0,0E+00 | 0,0E+00 | 0,0E+00 | 0,0E+00 | 0,0E+00 | 0,0E+00 | 0,0E+00 | 0,0E+00 | 0,0E+00 | 0,0E+00 | 0,0E+00 | 53070 | 54389 | 1320 | - |
| Y11_RS21030 | Y11_p0791 | yopN | SctW family type III secretion system gatekeeper subunit YopN | 3,89 | 3,64 | 3,97 | -0,23 | 0,08 | 0,34 | 1,1E-08 | 7,5E-09 | 4,6E-08 | 2,2E-02 | 6,5E-01 | 3,8E-02 | 1,5E-09 | 5,0E-10 | 1,8E-09 | 0,0E+00 | 0,0E+00 | 0,0E+00 | 54587 | 55468 | 882 | + |
| Y11_RS21040 | Y11_p0811 | sycN | type III secretion chaperone SycN | 15 | 15 | 15 | 0 | 0 | 0 | 0,0E+00 | 0,0E+00 | 0,0E+00 | 0,0E+00 | 0,0E+00 | 0,0E+00 | 0,0E+00 | 0,0E+00 | 0,0E+00 | 0,0E+00 | 0,0E+00 | 0,0E+00 | 55714 | 56085 | 372 | + |
| Y11_RS21050 | Y11_p0831 | yscY | type III secretion system chaperone YscY | 15 | 15 | 15 | 0 | 0 | 0 | 0,0E+00 | 0,0E+00 | 0,0E+00 | 0,0E+00 | 0,0E+00 | 0,0E+00 | 0,0E+00 | 0,0E+00 | 0,0E+00 | 0,0E+00 | 0,0E+00 | 0,0E+00 | 56447 | 56791 | 345 | + |
| Y11_RS21055 | Y11_p0841 | lcrD | SctV family type III secretion system export apparatus subunit LcrD | 15 | 15 | -0,5 | 0 | 0 | 0 | 0,0E+00 | 0,0E+00 | 3,1E-04 | 0,0E+00 | 0,0E+00 | 0,0E+00 | 0,0E+00 | 0,0E+00 | 0,0E+00 | 0,0E+00 | 0,0E+00 | 0,0E+00 | 56778 | 58892 | 2115 | + |
| Y11_RS21065 | Y11_p0861 | lcrG | type III secretion system chaperone LcrG | 15 | 15 | 15 | 0 | 0 | 0 | 0,0E+00 | 0,0E+00 | 0,0E+00 | 0,0E+00 | 0,0E+00 | 0,0E+00 | 0,0E+00 | 0,0E+00 | 0,0E+00 | 0,0E+00 | 0,0E+00 | 0,0E+00 | 59371 | 59658 | 288 | + |
| Y11_RS21070 | Y11_p0871 | lcrV | type III secretion system needle tip protein LcrV | 15 | 3,95 | 4,01 | 0 | 0 | 0 | 0,0E+00 | 2,5E-09 | 5,6E-09 | 0,0E+00 | 0,0E+00 | 0,0E+00 | 0,0E+00 | 2,1E-10 | 3,5E-10 | 0,0E+00 | 0,0E+00 | 0,0E+00 | 59660 | 60634 | 975 | + |
| Y11_RS21075 | | lcrH | type III secretion system chaperone SycD/LcrH | 15 | 4,69 | 15 | 0 | 0 | 0 | 0,0E+00 | 2,0E-09 | 0,0E+00 | 0,0E+00 | 0,0E+00 | 0,0E+00 | 0,0E+00 | 1,8E-10 | 0,0E+00 | 0,0E+00 | 0,0E+00 | 0,0E+00 | 60637 | 61143 | 507 | + |
| Y11_RS21085 | Y11_p0901 | yopD | type III secretion system translocon subunit YopD | 15 | 3,08 | 15 | 0 | 0 | 0 | 0,0E+00 | 1,1E-07 | 0,0E+00 | 0,0E+00 | 0,0E+00 | 0,0E+00 | 0,0E+00 | 4,5E-09 | 0,0E+00 | 0,0E+00 | 0,0E+00 | 0,0E+00 | 62345 | 63265 | 921 | + |

| | | | | | | | | | | | | | | | | | | | | | | | | | |
|---|---|---|---|---|---|---|---|---|---|---|---|---|---|---|---|---|---|---|---|---|---|---|---|---|---|
| Y11_RS21100 | Y11_p0931 | yopM | type III secretion system effector YopM | 15 | 3,98 | 15 | 0 | 0 | 0 | 0,0E+00 | 1,5E-09 | 0,0E+00 | 0,0E+00 | 0,0E+00 | 0,0E+00 | 0,0E+00 | 1,4E-10 | 0,0E+00 | 0,0E+00 | 0,0E+00 | 0,0E+00 | 64040 | 65143 | 1104 | - |
| Y11_RS21380 | Y11_04571 | rmf | ribosome modulation factor | 3,41 | 1,83 | 3,99 | -1,58 | 0,58 | 2,13 | 4,6E-06 | 4,2E-04 | 4,6E-07 | 4,2E-04 | 3,0E-02 | 2,4E-05 | 2,5E-07 | 8,7E-06 | 1,2E-08 | 7,3E-06 | 0,0E+00 | 2,5E-06 | 488942 | 489109 | 168 | + |
| Y11_RS21435 | | | ricin-type beta-trefoil lectin domain protein | 2,98 | 1,12 | 1,46 | -1,86 | -1,52 | 0,34 | 4,0E-07 | 2,6E-06 | 2,6E-05 | 1,4E-05 | 2,4E-04 | 8,0E-02 | 2,9E-08 | 7,0E-08 | 4,0E-07 | 2,7E-07 | 5,9E-06 | 0,0E+00 | 696972 | 697493 | 522 | + |
| Y11_RS21460 | | infC | translation initiation factor IF-3 | 0,11 | 0,4 | -0,12 | 0,29 | -0,24 | -0,53 | 5,7E-01 | 3,9E-02 | 4,6E-01 | 9,0E-02 | 1,4E-01 | 1,1E-03 | 0,0E+00 | 0,0E+00 | 0,0E+00 | 0,0E+00 | 0,0E+00 | 0,0E+00 | 811829 | 812380 | 552 | + |
| Y11_RS23215 | | | hypothetical protein | -0,43 | 0,78 | 1,19 | 1,2 | 1,62 | 0,41 | 4,1E-05 | 3,5E-06 | 1,6E-05 | 5,0E-08 | 1,2E-06 | 1,4E-02 | 0,0E+00 | 0,0E+00 | 2,5E-07 | 1,9E-09 | 5,0E-08 | 0,0E+00 | 2490754 | 2491149 | 396 | - |
| Y11_RS23630 | Y11_07231 | | YnfU family zinc-binding protein | 1,64 | 2,25 | 9,22 | 0,61 | 7,42 | 6,97 | 3,8E-01 | 2,4E-01 | 7,3E-04 | 7,4E-01 | 3,1E-03 | 4,3E-03 | 1,2E-02 | 4,4E-03 | 8,9E-06 | 0,0E+00 | 6,8E-05 | 2,6E-04 | 752152 | 752319 | 168 | + |

**Table S7.** Published *Ye* transcriptomes comparisons to which the proteomes comparisons were correlated

| Number | Comparison | Reference |
|---|---|---|
| 1 | Ye.1 vs Ye.2 | This sudy |
| 2 | Ye.1 vs Ye.3 | This sudy |
| 3 | Ye.1 vs Ye.4 | This sudy |
| 4 | Ye.2 vs Ye.3 | This sudy |
| 5 | Ye.2 vs Ye.4 | This sudy |
| 6 | Ye.3 vs Ye.4 | This sudy |
| 7 | 37C 2014 vs 22C 2014 | Leskinen K et al., Microbiology 2015 (PMID: 25416689) |
| 8 | Mutant hfq 22C 2015 vs WT 22C 2015 | Leskinen K et al., Mol Microbiol. 2017 (Pubmed:28010054) |
| 9 | 37C 2015 vs 22C 2015 | Leskinen K et al., Microbiology 2015 (PMID: 25767108) |
| 10 | Mutant rfaH 22C 2015 vs WT 22C 2015 | Leskinen K et al., Microbiology 2015 (PMID: 25767108) |
| 11 | Mutant ybeY 37C 2014 vs Mutant ybeY 22C 2014 | Leskinen K et al., Microbiology 2015 (PMID: 25416689) |
| 12 | Mutant ybeY 37C 2014 vs WT 37C 2014 | Leskinen K et al., Microbiology 2015 (PMID: 25416689) |
| 13 | Mutant hfq 37C 2015 vs Mutant hfq 22C 2015 | Leskinen K et al., Mol Microbiol. 2017 (Pubmed:28010054) |
| 14 | Mutant hfq 37C 2015 vs WT 37C 2015 | Leskinen K et al., Mol Microbiol. 2017 (Pubmed:28010054) |
| 15 | Mutant rfaH 37C 2015 vs WT 37C 2015 | Leskinen K et al., Microbiology 2015 (PMID: 25767108) |
| 16 | Mutant rfaH 37C 2015 vs Mutant rfaH 22C 2015 | Leskinen K et al., Microbiology 2015 (PMID: 25767108) |
| 17 | Mutant OAntigen 22C 2015 vs WT 22C 2015 | Leskinen K et al., Microbiology 2015 (PMID: 25767108) |
| 18 | Mutant ybeY 22C 2014 vs WT 22C 2014 | Leskinen K et al., Microbiology 2015 (PMID: 25416689) |
| 19 | 26C 120min TYE vs 26C 30min TYE | Bent et al., Infect Immun. 2015 (PMID: 25895974) |
| 20 | Mutant yenR OverExpr ytxR 26.37C 5h vs Mutant yenR 26.37C 5h | Axler-DiPerte GL et al., J Bacteriol 2009 (PMID: 19011024) |
| 21 | Mutant rcsB 26C M63 Log vs WT 26C M63 Log | Meng et al., Curr Genet. 2020 (PMID: 32488337) |
| 22 | 26C 60min TYE vs 26C 30min TYE | Bent et al., Infect Immun. 2015 (PMID: 25895974) |
| 23 | 37C 120min RPMI vs 26C 120min TYE | Bent et al., Infect Immun. 2015 (PMID: 25895974) |
| 24 | 37C 120min RPMI vs 37C 30min RPMI | Bent et al., Infect Immun. 2015 (PMID: 25895974) |
| 25 | 37C 60min RPMI vs 37C 30min RPMI | Bent et al., Infect Immun. 2015 (PMID: 25895974) |
| 26 | 37C 60min RPMI vs 26C 60min TYE | Bent et al., Infect Immun. 2015 (PMID: 25895974) |
| 27 | 26C 240min TYE vs 26C 30min TYE | Bent et al., Infect Immun. 2015 (PMID: 25895974) |
| 28 | 37C 30min P388D1 RPMI vs 37C 30min RPMI | Bent et al., Infect Immun. 2015 (PMID: 25895974) |
| 29 | 37C 240min RPMI vs 26C 240min TYE | Bent et al., Infect Immun. 2015 (PMID: 25895974) |
| 30 | 37C 240min RPMI vs 37C 30min RPMI | Bent et al., Infect Immun. 2015 (PMID: 25895974) |
| 31 | 37C 60min P388D1 RPMI vs 37C 30min P388D1 RPMI | Bent et al., Infect Immun. 2015 (PMID: 25895974) |
| 32 | 37C 60min P388D1 RPMI vs 37C 60min RPMI | Bent et al., Infect Immun. 2015 (PMID: 25895974) |
| 33 | Mutant yenR ytxR OverExpr ytxR 26C vs Mutant yenR ytxR 26C | Axler-DiPerte GL et al., J Bacteriol 2009 (PMID: 19011024) |
| 34 | 8081 25C Log LB vs Y1 25C Log LB | Schmuhl et al., mSystems 2019 (PMID: 31020044) |
| 35 | 8081 25C Log LB vs 8081 25C Stat LB | Schmuhl et al., mSystems 2019 (PMID: 31020044) |
| 36 | 37C 240min P388D1 RPMI vs 37C 30min P388D1 RPMI | Bent et al., Infect Immun. 2015 (PMID: 25895974) |
| 37 | 37C 240min P388D1 RPMI vs 37C 240min RPMI | Bent et al., Infect Immun. 2015 (PMID: 25895974) |
| 38 | 37C 120min Intracellular P388D1 RPMI vs 37C 120min RPMI | Bent et al., Infect Immun. 2015 (PMID: 25895974) |
| 39 | 37C 120min Intracellular P388D1 RPMI vs 37C 120min P388D1 RPMI | Bent et al., Infect Immun. 2015 (PMID: 25895974) |
| 40 | Mutant yenR rovA 26C 8h vs Mutant yenR 26C 8h | Cathelyn JS et al., Mol Microbiol 2007 (PMID: 17784909) |
| 41 | 37C 30min RPMI vs 26C 30min TYE | Bent et al., Infect Immun. 2015 (PMID: 25895974) |
| 42 | 8081 25C Stat LB vs Y1 25C Stat LB | Schmuhl et al., mSystems 2019 (PMID: 31020044) |
| 43 | 37C 120min P388D1 RPMI vs 37C 120min RPMI | Bent et al., Infect Immun. 2015 (PMID: 25895974) |
| 44 | 37C 120min P388D1 RPMI vs 37C 30min P388D1 RPMI | Bent et al., Infect Immun. 2015 (PMID: 25895974) |
| 45 | 8081 37C Stat LB vs Y1 37C Stat LB | Schmuhl et al., mSystems 2019 (PMID: 31020044) |
| 46 | 8081 37C Stat LB vs 8081 25C Stat LB | Schmuhl et al., mSystems 2019 (PMID: 31020044) |
| 47 | 8081 37C Log LB vs 8081 37C Stat LB | Schmuhl et al., mSystems 2019 (PMID: 31020044) |
| 48 | 8081 37C Log LB vs Y1 37C Log LB | Schmuhl et al., mSystems 2019 (PMID: 31020044) |
| 49 | 8081 37C Log LB vs 8081 25C Log LB | Schmuhl et al., mSystems 2019 (PMID: 31020044) |
| 50 | relA spoT vs WT | Huang et al., Int J Mol Sci. 2023 (PMID: 37108773) |
| 51 | dksA vs WT | Huang et al., Int J Mol Sci. 2023 (PMID: 37108773) |
| 52 | dksA relA spoT vs dksA | Huang et al., Int J Mol Sci. 2023 (PMID: 37108773) |
| 53 | dksA relA spoT vs WT | Huang et al., Int J Mol Sci. 2023 (PMID: 37108773) |
| 54 | dksA vs relA spoT | Huang et al., Int J Mol Sci. 2023 (PMID: 37108773) |
| 55 | dksA relA spoT vs relA spoT | Huang et al., Int J Mol Sci. 2023 (PMID: 37108773) |
| 56 | Y1 25C Stat LB vs 8081 25C Stat LB | Schmuhl et al., mSystems 2019 (PMID: 31020044) |
| 57 | Y1 37C Log LB vs Y1 37C Stat LB | Schmuhl et al., mSystems 2019 (PMID: 31020044) |
| 58 | Y1 37C Log LB vs Y1 25C Log LB | Schmuhl et al., mSystems 2019 (PMID: 31020044) |
| 59 | Y1 37C Log LB vs 8081 37C Log LB | Schmuhl et al., mSystems 2019 (PMID: 31020044) |
| 60 | Y1 37C Stat LB vs Y1 25C Stat LB | Schmuhl et al., mSystems 2019 (PMID: 31020044) |
| 61 | Y1 37C Stat LB vs 8081 37C Stat LB | Schmuhl et al., mSystems 2019 (PMID: 31020044) |
| 62 | Y1 25C Log LB vs Y1 25C Stat LB | Schmuhl et al., mSystems 2019 (PMID: 31020044) |
| 63 | Y1 25C Log LB vs 8081 25C Log LB | Schmuhl et al., mSystems 2019 (PMID: 31020044) |

**Table S8.** Correlation between published transcriptome comparisons and proteome comparisons of Ye.1 to Ye.4. Comparisons corresponding to row and column numbers are detailled in Table S7

| row | column | cor | p |
|---|---|---|---|
| 1 | 2 | 0.092 | 4.8e-05 |
| 1 | 3 | 0.1 | 6.4e-06 |
| 1 | 4 | -0.5 | 8,00E-123 |
| 1 | 5 | -0.44 | 2,00E-92 |
| 1 | 10 | 0.052 | 0.022 |
| 1 | 14 | 0.048 | 0.033 |
| 1 | 15 | 0.09 | 7.2e-05 |
| 1 | 19 | -0.049 | 0.031 |
| 1 | 21 | 0.014 | 0.55 |
| 1 | 22 | 0.00035 | 0.99 |
| 1 | 27 | -0.027 | 0.24 |
| 1 | 34 | -0.025 | 0.27 |
| 1 | 38 | 0.027 | 0.24 |
| 1 | 39 | -0.0061 | 0.79 |
| 1 | 40 | -0.0038 | 0.87 |
| 1 | 41 | 0.043 | 0.056 |
| 1 | 42 | 0.01 | 0.65 |
| 1 | 45 | 0.014 | 0.55 |
| 1 | 46 | 0.019 | 0.4 |
| 1 | 48 | 0.0014 | 0.95 |
| 1 | 49 | 0.048 | 0.034 |
| 1 | 52 | -0.14 | 9.9e-10 |
| 1 | 60 | 0.028 | 0.22 |
| 2 | 3 | 0.85 | 0 |
| 2 | 34 | -0.046 | 0.044 |
| 2 | 38 | 0.058 | 0.011 |
| 2 | 39 | -0.024 | 0.3 |
| 2 | 40 | 0.0097 | 0.67 |
| 2 | 42 | 0.098 | 1.6e-05 |
| 2 | 48 | -0.095 | 2.9e-05 |
| 2 | 60 | -0.023 | 0.32 |
| 3 | 34 | -0.015 | 0.5 |
| 3 | 38 | 0.15 | 6.2e-11 |
| 3 | 39 | 0.0047 | 0.84 |
| 3 | 40 | 0.044 | 0.054 |
| 3 | 42 | 0.12 | 4.2e-08 |
| 3 | 48 | -0.09 | 7.3e-05 |
| 3 | 60 | -0.013 | 0.58 |
| 4 | 2 | 0.79 | 0 |
| 4 | 3 | 0.67 | 2,00E-253 |
| 4 | 5 | 0.89 | 0 |
| 4 | 34 | -0.029 | 0.21 |
| 4 | 38 | 0.048 | 0.035 |
| 4 | 39 | -0.011 | 0.61 |
| 4 | 40 | 0.014 | 0.54 |
| 4 | 42 | 0.074 | 0.0011 |
| 4 | 48 | -0.081 | 0.00039 |
| 4 | 60 | -0.026 | 0.25 |
| 5 | 2 | 0.7 | 4.9e-281 |
| 5 | 3 | 0.83 | 0 |
| 5 | 34 | -0.0067 | 0.77 |
| 5 | 38 | 0.13 | 2.4e-08 |
| 5 | 39 | 0.013 | 0.56 |
| 5 | 40 | 0.045 | 0.046 |
| 5 | 42 | 0.1 | 5.9e-06 |
| 5 | 48 | -0.083 | 0.00026 |
| 5 | 60 | -0.023 | 0.32 |
| 6 | 1 | 0.055 | 0.016 |
| 6 | 2 | 0.02 | 0.39 |
| 6 | 3 | 0.54 | 9.5e-148 |
| 6 | 4 | 0.018 | 0.42 |
| 6 | 5 | 0.47 | 1.3e-105 |
| 6 | 10 | 0.11 | 6.9e-07 |
| 6 | 14 | 0.17 | 8.9e-15 |
| 6 | 15 | 0.1 | 7,00E-06 |
| 6 | 19 | 0.1 | 1.1e-05 |
| 6 | 21 | 0.16 | 5.7e-13 |
| 6 | 22 | 0.084 | 0.00021 |
| 6 | 23 | 0.13 | 6.5e-09 |
| 6 | 26 | 0.14 | 1.8e-10 |
| 6 | 27 | 0.17 | 2.3e-13 |
| 6 | 29 | 0.3 | 4.1e-41 |
| 6 | 30 | 0.27 | 1.1e-32 |
| 6 | 34 | 0.049 | 0.03 |
| 6 | 38 | 0.2 | 3.9e-18 |
| 6 | 39 | 0.048 | 0.036 |
| 6 | 40 | 0.071 | 0.0019 |
| 6 | 41 | 0.19 | 3.5e-17 |
| 6 | 42 | 0.084 | 0.00021 |
| 6 | 45 | 0.056 | 0.013 |
| 6 | 46 | -0.012 | 0.61 |
| 6 | 48 | -0.016 | 0.47 |
| 6 | 49 | 0.16 | 2.8e-13 |
| 6 | 52 | -0.028 | 0.22 |
| 6 | 54 | 0.11 | 6.4e-07 |
| 6 | 55 | 0.12 | 1.5e-07 |
| 6 | 60 | 0.015 | 0.51 |
| 7 | 1 | -0.0089 | 0.69 |
| 7 | 2 | 0.16 | 1.3e-12 |
| 7 | 3 | 0.19 | 8.3e-17 |
| 7 | 4 | 0.15 | 3.9e-11 |
| 7 | 5 | 0.18 | 5.8e-15 |
| 7 | 6 | 0.1 | 9.5e-06 |
| 7 | 9 | 1 | 0 |
| 7 | 10 | 0.55 | 1.5e-153 |
| 7 | 11 | 0.41 | 2,00E-77 |
| 7 | 13 | 0.57 | 3.9e-170 |
| 7 | 14 | -0.053 | 0.019 |
| 7 | 15 | 0.15 | 9.4e-11 |
| 7 | 19 | 0.02 | 0.38 |
| 7 | 21 | -0.03 | 0.19 |
| 7 | 22 | -0.043 | 0.059 |

| row | column | cor | p |
|---|---|---|---|
| 15 | 40 | -0.015 | 0.51 |
| 15 | 41 | 0.049 | 0.032 |
| 15 | 42 | 2.2e-05 | 1 |
| 15 | 45 | 0.068 | 0.0026 |
| 15 | 46 | 0.12 | 2.3e-07 |
| 15 | 48 | -0.069 | 0.0023 |
| 15 | 49 | 0.17 | 1.5e-14 |
| 15 | 52 | -0.0038 | 0.87 |
| 15 | 60 | 0.07 | 0.002 |
| 16 | 1 | 0.0068 | 0.76 |
| 16 | 2 | 0.038 | 0.095 |
| 16 | 3 | 0.074 | 0.0011 |
| 16 | 4 | 0.045 | 0.049 |
| 16 | 5 | 0.077 | 0.00071 |
| 16 | 6 | 0.081 | 0.00036 |
| 16 | 7 | 0.7 | 1.8e-290 |
| 16 | 9 | 0.7 | 1.2e-290 |
| 16 | 10 | 0.083 | 0.00027 |
| 16 | 11 | 0.15 | 7.4e-12 |
| 16 | 13 | 0.48 | 7.8e-114 |
| 16 | 14 | 0.029 | 0.21 |
| 16 | 15 | 0.35 | 3.7e-57 |
| 16 | 19 | 0.025 | 0.27 |
| 16 | 21 | -0.14 | 9.7e-10 |
| 16 | 22 | -0.027 | 0.23 |
| 16 | 23 | 0.16 | 1.9e-12 |
| 16 | 24 | 0.21 | 2.8e-20 |
| 16 | 25 | 0.12 | 8.6e-08 |
| 16 | 26 | 0.1 | 4.5e-06 |
| 16 | 27 | 0.033 | 0.14 |
| 16 | 28 | 0.024 | 0.29 |
| 16 | 29 | 0.18 | 8.3e-16 |
| 16 | 30 | 0.2 | 1.2e-19 |
| 16 | 32 | -0.0011 | 0.96 |
| 16 | 34 | -0.066 | 0.0036 |
| 16 | 38 | -0.026 | 0.26 |
| 16 | 39 | -0.016 | 0.48 |
| 16 | 40 | -0.098 | 1.5e-05 |
| 16 | 41 | -0.013 | 0.55 |
| 16 | 42 | -0.062 | 0.006 |
| 16 | 43 | -0.011 | 0.62 |
| 16 | 45 | -0.092 | 4.6e-05 |
| 16 | 46 | -0.06 | 0.0083 |
| 16 | 48 | -0.06 | 0.0082 |
| 16 | 49 | 0.29 | 1.7e-39 |
| 16 | 52 | -0.019 | 0.4 |
| 16 | 54 | -0.063 | 0.0055 |
| 16 | 55 | -0.087 | 0.00012 |
| 16 | 58 | 0.27 | 3.7e-34 |
| 16 | 60 | -0.047 | 0.039 |
| 17 | 1 | 0.034 | 0.13 |
| 17 | 2 | -0.036 | 0.11 |
| 17 | 3 | -0.13 | 2.6e-08 |
| 17 | 4 | -0.065 | 0.0042 |
| 17 | 5 | -0.14 | 4.5e-10 |
| 17 | 6 | -0.19 | 2.3e-17 |
| 17 | 7 | 0.09 | 7.7e-05 |
| 17 | 9 | 0.09 | 7.4e-05 |
| 17 | 10 | 0.12 | 1.1e-07 |
| 17 | 11 | -0.2 | 3,00E-18 |
| 17 | 12 | 0.25 | 5.4e-29 |
| 17 | 13 | -0.22 | 2.7e-22 |
| 17 | 14 | -0.0042 | 0.85 |
| 17 | 15 | 0.34 | 1.3e-54 |
| 17 | 16 | 0.26 | 1.4e-30 |
| 17 | 18 | 0.46 | 1.4e-100 |
| 17 | 19 | -0.076 | 0.00086 |
| 17 | 20 | 0.041 | 0.071 |
| 17 | 21 | -0.25 | 2.5e-28 |
| 17 | 22 | -0.019 | 0.41 |
| 17 | 23 | -0.047 | 0.038 |
| 17 | 24 | -0.027 | 0.24 |
| 17 | 25 | -0.066 | 0.0037 |
| 17 | 26 | -0.14 | 3.5e-10 |
| 17 | 27 | -0.23 | 1.1e-24 |
| 17 | 28 | -0.079 | 0.00049 |
| 17 | 29 | -0.037 | 0.11 |
| 17 | 30 | -0.18 | 5.8e-16 |
| 17 | 31 | -0.071 | 0.0018 |
| 17 | 32 | -0.12 | 1.7e-07 |
| 17 | 33 | 0.087 | 0.00013 |
| 17 | 34 | -0.096 | 2.3e-05 |
| 17 | 35 | 0.25 | 2.5e-29 |
| 17 | 36 | 0.042 | 0.065 |
| 17 | 37 | 0.16 | 3,00E-13 |
| 17 | 38 | -0.18 | 4.4e-16 |
| 17 | 39 | -0.079 | 0.00048 |
| 17 | 40 | -0.034 | 0.13 |
| 17 | 41 | -0.095 | 3.1e-05 |
| 17 | 42 | -0.052 | 0.022 |
| 17 | 43 | -0.12 | 1,00E-07 |
| 17 | 44 | -0.0071 | 0.76 |
| 17 | 45 | -0.01 | 0.65 |
| 17 | 46 | 0.049 | 0.032 |
| 17 | 47 | 0.19 | 1.7e-17 |
| 17 | 48 | -0.029 | 0.19 |
| 17 | 49 | -0.068 | 0.0026 |
| 17 | 50 | 0.24 | 2.3e-26 |
| 17 | 51 | 0.17 | 9.8e-14 |
| 17 | 52 | 0.042 | 0.067 |
| 17 | 53 | 0.17 | 1.6e-13 |
| 17 | 54 | -0.15 | 4.7e-11 |

| row | column | cor | p |
|---|---|---|---|
| 28 | 11 | 0.098 | 1.5e-05 |
| 28 | 13 | 0.16 | 1.3e-12 |
| 28 | 14 | 0.055 | 0.015 |
| 28 | 15 | 0.016 | 0.48 |
| 28 | 19 | 0.043 | 0.059 |
| 28 | 21 | 0.074 | 0.0011 |
| 28 | 22 | 0.046 | 0.042 |
| 28 | 23 | 0.036 | 0.11 |
| 28 | 26 | -0.019 | 0.41 |
| 28 | 27 | 0.1 | 9.6e-06 |
| 28 | 29 | 0.2 | 8.3e-19 |
| 28 | 30 | 0.31 | 1.1e-44 |
| 28 | 32 | 0.41 | 7.9e-79 |
| 28 | 34 | 0.079 | 0.00047 |
| 28 | 38 | 0.19 | 5.3e-17 |
| 28 | 39 | -0.043 | 0.061 |
| 28 | 40 | 0.0033 | 0.89 |
| 28 | 41 | -0.044 | 0.054 |
| 28 | 42 | 0.1 | 3.8e-06 |
| 28 | 45 | 0.041 | 0.068 |
| 28 | 46 | -0.046 | 0.044 |
| 28 | 48 | 0.016 | 0.48 |
| 28 | 49 | -0.012 | 0.61 |
| 28 | 52 | 0.026 | 0.25 |
| 28 | 54 | 0.0056 | 0.81 |
| 28 | 55 | 0.022 | 0.33 |
| 28 | 58 | 0.11 | 1.2e-06 |
| 28 | 60 | 0.027 | 0.23 |
| 29 | 1 | 0.05 | 0.028 |
| 29 | 2 | 0.098 | 1.6e-05 |
| 29 | 3 | 0.25 | 2.3e-28 |
| 29 | 4 | 0.066 | 0.0035 |
| 29 | 5 | 0.2 | 1.9e-18 |
| 29 | 10 | 0.2 | 2.6e-19 |
| 29 | 14 | 0.19 | 6.5e-18 |
| 29 | 15 | 0.18 | 1.7e-15 |
| 29 | 19 | -0.047 | 0.04 |
| 29 | 21 | 0.043 | 0.058 |
| 29 | 22 | 0.056 | 0.013 |
| 29 | 23 | 0.29 | 4.6e-40 |
| 29 | 26 | 0.16 | 3.8e-12 |
| 29 | 27 | -0.16 | 3.4e-12 |
| 29 | 30 | 0.51 | 9.1e-131 |
| 29 | 34 | -0.0027 | 0.91 |
| 29 | 38 | 0.32 | 1.1e-46 |
| 29 | 39 | -0.024 | 0.29 |
| 29 | 40 | -0.0092 | 0.69 |
| 29 | 41 | 0.22 | 8.8e-23 |
| 29 | 42 | 0.11 | 1.5e-06 |
| 29 | 45 | 0.078 | 0.00057 |
| 29 | 46 | 0.012 | 0.61 |
| 29 | 48 | -0.01 | 0.66 |
| 29 | 49 | 0.24 | 1.5e-27 |
| 29 | 52 | -0.063 | 0.0055 |
| 29 | 54 | 0.12 | 8.7e-08 |
| 29 | 55 | 0.11 | 1.9e-06 |
| 29 | 60 | 0.061 | 0.0077 |
| 30 | 1 | -0.025 | 0.27 |
| 30 | 2 | 0.12 | 2.7e-07 |
| 30 | 3 | 0.24 | 6.5e-27 |
| 30 | 4 | 0.12 | 5.3e-08 |
| 30 | 5 | 0.24 | 3.8e-26 |
| 30 | 10 | 0.24 | 3.8e-27 |
| 30 | 14 | 0.14 | 1.4e-09 |
| 30 | 15 | 0.16 | 2.3e-12 |
| 30 | 19 | 0.19 | 5.8e-17 |
| 30 | 21 | 0.18 | 9.2e-16 |
| 30 | 22 | 0.089 | 9.2e-05 |
| 30 | 23 | 0.13 | 1.6e-08 |
| 30 | 26 | 0.036 | 0.12 |
| 30 | 27 | 0.39 | 2,00E-72 |
| 30 | 34 | 0.058 | 0.011 |
| 30 | 38 | 0.18 | 4.2e-16 |
| 30 | 39 | 0.023 | 0.31 |
| 30 | 40 | 0.12 | 2.3e-07 |
| 30 | 41 | -0.16 | 3,00E-12 |
| 30 | 42 | 0.12 | 4.4e-08 |
| 30 | 45 | 0.0097 | 0.67 |
| 30 | 46 | -0.1 | 6,00E-06 |
| 30 | 48 | -0.0047 | 0.84 |
| 30 | 49 | 0.22 | 3.1e-22 |
| 30 | 52 | 0.016 | 0.49 |
| 30 | 54 | 0.017 | 0.46 |
| 30 | 55 | 0.03 | 0.19 |
| 30 | 60 | 0.011 | 0.61 |
| 31 | 1 | 0.0096 | 0.67 |
| 31 | 2 | 0.07 | 0.002 |
| 31 | 3 | 0.043 | 0.059 |
| 31 | 4 | 0.059 | 0.0089 |
| 31 | 5 | 0.035 | 0.12 |
| 31 | 6 | -0.02 | 0.38 |
| 31 | 7 | 0.054 | 0.017 |
| 31 | 9 | 0.054 | 0.017 |
| 31 | 10 | 0.056 | 0.013 |
| 31 | 11 | 0.1 | 4.9e-06 |
| 31 | 13 | 0.11 | 2.4e-06 |
| 31 | 14 | 0.08 | 4,00E-04 |
| 31 | 15 | 0.034 | 0.13 |
| 31 | 16 | 0.032 | 0.15 |
| 31 | 19 | 0.048 | 0.033 |
| 31 | 21 | 0.056 | 0.013 |
| 31 | 22 | -0.015 | 0.5 |

| row | column | cor | p |
|---|---|---|---|
| 43 | 6 | 0.16 | 6.1e-13 |
| 43 | 10 | 0.087 | 0.00013 |
| 43 | 11 | 0.15 | 1.5e-11 |
| 43 | 13 | 0.31 | 9.3e-44 |
| 43 | 14 | 0.19 | 4.9e-17 |
| 43 | 15 | 0.078 | 0.00055 |
| 43 | 19 | 0.035 | 0.12 |
| 43 | 21 | 0.17 | 5.7e-14 |
| 43 | 22 | 0.077 | 0.00065 |
| 43 | 23 | -0.099 | 1.4e-05 |
| 43 | 26 | 0.093 | 4,00E-05 |
| 43 | 27 | 0.095 | 2.8e-05 |
| 43 | 28 | 0.25 | 1.7e-28 |
| 43 | 29 | 0.38 | 9.1e-69 |
| 43 | 30 | 0.17 | 1.5e-13 |
| 43 | 32 | 0.34 | 3.3e-55 |
| 43 | 34 | -0.041 | 0.071 |
| 43 | 38 | 0.54 | 1.5e-147 |
| 43 | 39 | -0.39 | 4.2e-70 |
| 43 | 40 | -0.029 | 0.21 |
| 43 | 41 | 0.3 | 8.7e-41 |
| 43 | 42 | 0.13 | 2.9e-08 |
| 43 | 45 | 0.086 | 0.00014 |
| 43 | 46 | 0.016 | 0.47 |
| 43 | 48 | -0.08 | 0.00044 |
| 43 | 49 | 0.17 | 3.1e-14 |
| 43 | 52 | -0.025 | 0.27 |
| 43 | 54 | 0.098 | 1.5e-05 |
| 43 | 55 | 0.1 | 5.4e-06 |
| 43 | 58 | 0.22 | 1.8e-22 |
| 43 | 60 | 0.068 | 0.0026 |
| 44 | 1 | -0.0079 | 0.73 |
| 44 | 2 | 0.053 | 0.019 |
| 44 | 3 | 0.012 | 0.61 |
| 44 | 4 | 0.05 | 0.027 |
| 44 | 5 | 0.013 | 0.56 |
| 44 | 6 | -0.05 | 0.029 |
| 44 | 7 | 0.11 | 3.4e-06 |
| 44 | 9 | 0.11 | 3.4e-06 |
| 44 | 10 | 0.097 | 2,00E-05 |
| 44 | 11 | 0.12 | 2,00E-07 |
| 44 | 13 | 0.11 | 6.4e-07 |
| 44 | 14 | 0.032 | 0.16 |
| 44 | 15 | 0.097 | 1.9e-05 |
| 44 | 16 | 0.099 | 1.4e-05 |
| 44 | 19 | 0.1 | 1.1e-05 |
| 44 | 21 | 0.022 | 0.34 |
| 44 | 22 | 0.067 | 0.0034 |
| 44 | 23 | 0.097 | 1.7e-05 |
| 44 | 24 | 0.25 | 4.6e-28 |
| 44 | 25 | 0.18 | 6.4e-15 |
| 44 | 26 | 0.04 | 0.08 |
| 44 | 27 | 0.1 | 1,00E-05 |
| 44 | 28 | -0.77 | 0 |
| 44 | 29 | 0.0039 | 0.86 |
| 44 | 30 | 0.13 | 1.1e-08 |
| 44 | 32 | -0.22 | 6.7e-22 |
| 44 | 34 | -0.082 | 3,00E-04 |
| 44 | 38 | -0.012 | 0.58 |
| 44 | 39 | -0.12 | 3.9e-08 |
| 44 | 40 | 0.028 | 0.22 |
| 44 | 41 | -0.017 | 0.45 |
| 44 | 42 | -0.032 | 0.16 |
| 44 | 43 | 0.11 | 2.2e-06 |
| 44 | 45 | 0.01 | 0.66 |
| 44 | 46 | 0.047 | 0.038 |
| 44 | 48 | -0.027 | 0.24 |
| 44 | 49 | 0.18 | 4.9e-15 |
| 44 | 52 | -0.048 | 0.035 |
| 44 | 54 | -0.00029 | 0.99 |
| 44 | 55 | -0.029 | 0.2 |
| 44 | 58 | 0.078 | 0.00057 |
| 44 | 60 | -0.00057 | 0.98 |
| 45 | 2 | -0.021 | 0.35 |
| 45 | 3 | 0.011 | 0.63 |
| 45 | 4 | -0.033 | 0.15 |
| 45 | 5 | -0.003 | 0.89 |
| 45 | 19 | -0.017 | 0.45 |
| 45 | 21 | 0.019 | 0.4 |
| 45 | 22 | 0.0064 | 0.78 |
| 45 | 27 | -0.049 | 0.033 |
| 45 | 34 | 0.41 | 1.7e-78 |
| 45 | 38 | 0.11 | 1.8e-06 |
| 45 | 39 | 0.031 | 0.18 |
| 45 | 40 | 0.092 | 5.1e-05 |
| 45 | 41 | 0.012 | 0.6 |
| 45 | 42 | 0.47 | 2.8e-105 |
| 45 | 46 | 0.49 | 8.7e-120 |
| 45 | 48 | 0.52 | 2.7e-133 |
| 45 | 60 | -0.19 | 5.8e-17 |
| 46 | 2 | -0.13 | 1.8e-08 |
| 46 | 3 | -0.11 | 7,00E-07 |
| 46 | 4 | -0.12 | 2.1e-07 |
| 46 | 5 | -0.11 | 8.3e-07 |
| 46 | 19 | -0.083 | 0.00026 |
| 46 | 21 | -0.052 | 0.023 |
| 46 | 22 | -0.03 | 0.18 |
| 46 | 27 | -0.16 | 5.7e-12 |
| 46 | 34 | 0.039 | 0.086 |
| 46 | 38 | 0.14 | 8.2e-10 |
| 46 | 39 | 0.13 | 3.1e-09 |
| 46 | 40 | 0.0056 | 0.8 |

| row | column | cor | p |
|---|---|---|---|
| 54 | 40 | 0.026 | 0.25 |
| 54 | 41 | 0.21 | 1.1e-20 |
| 54 | 42 | 0.044 | 0.051 |
| 54 | 45 | 0.0048 | 0.83 |
| 54 | 46 | -0.052 | 0.023 |
| 54 | 48 | -0.051 | 0.024 |
| 54 | 49 | 0.13 | 1.9e-08 |
| 54 | 52 | -0.56 | 2.3e-159 |
| 54 | 55 | 0.87 | 0 |
| 54 | 60 | -0.03 | 0.19 |
| 55 | 1 | -0.0084 | 0.71 |
| 55 | 2 | -0.093 | 4.3e-05 |
| 55 | 3 | -0.016 | 0.47 |
| 55 | 4 | -0.074 | 0.0011 |
| 55 | 5 | -0.0067 | 0.77 |
| 55 | 10 | 0.038 | 0.096 |
| 55 | 14 | 0.12 | 3.9e-07 |
| 55 | 15 | 0.048 | 0.035 |
| 55 | 19 | 0.013 | 0.56 |
| 55 | 21 | 0.15 | 6.1e-11 |
| 55 | 22 | 0.0028 | 0.9 |
| 55 | 23 | 0.17 | 2,00E-14 |
| 55 | 26 | 0.24 | 8.6e-28 |
| 55 | 27 | 0.15 | 7.2e-11 |
| 55 | 34 | -0.038 | 0.095 |
| 55 | 38 | 0.0073 | 0.75 |
| 55 | 39 | -0.098 | 1.5e-05 |
| 55 | 40 | 0.072 | 0.0015 |
| 55 | 41 | 0.23 | 1.2e-25 |
| 55 | 42 | 0.052 | 0.023 |
| 55 | 45 | 0.031 | 0.18 |
| 55 | 46 | -0.02 | 0.38 |
| 55 | 48 | -0.048 | 0.034 |
| 55 | 49 | 0.12 | 7.8e-08 |
| 55 | 52 | -0.072 | 0.0015 |
| 55 | 60 | -0.0082 | 0.72 |
| 56 | 1 | -0.024 | 0.29 |
| 56 | 2 | -0.05 | 0.027 |
| 56 | 3 | -0.08 | 0.00041 |
| 56 | 4 | -0.024 | 0.29 |
| 56 | 5 | -0.055 | 0.015 |
| 56 | 6 | -0.078 | 0.00063 |
| 56 | 7 | 0.004 | 0.86 |
| 56 | 9 | 0.004 | 0.86 |
| 56 | 10 | -0.062 | 0.0067 |
| 56 | 11 | -0.0034 | 0.88 |
| 56 | 13 | -0.047 | 0.037 |
| 56 | 14 | -0.048 | 0.035 |
| 56 | 15 | -0.033 | 0.14 |
| 56 | 16 | 0.04 | 0.075 |
| 56 | 19 | -0.092 | 5.4e-05 |
| 56 | 21 | -0.078 | 0.00058 |
| 56 | 22 | -0.065 | 0.0041 |
| 56 | 23 | -0.01 | 0.65 |
| 56 | 24 | 0.001 | 0.96 |
| 56 | 25 | 0.054 | 0.018 |
| 56 | 26 | 0.0029 | 0.9 |
| 56 | 27 | -0.12 | 3.6e-08 |
| 56 | 28 | -0.069 | 0.0023 |
| 56 | 29 | -0.076 | 0.00084 |
| 56 | 30 | -0.11 | 5.3e-07 |
| 56 | 31 | -0.027 | 0.23 |
| 56 | 32 | -0.12 | 1.8e-07 |
| 56 | 34 | -0.044 | 0.051 |
| 56 | 36 | 0.078 | 0.00062 |
| 56 | 38 | 0.026 | 0.25 |
| 56 | 39 | 0.1 | 7.3e-06 |
| 56 | 40 | -0.15 | 6.4e-11 |
| 56 | 41 | -0.086 | 0.00015 |
| 56 | 42 | -0.59 | 1.4e-181 |
| 56 | 43 | -0.071 | 0.0018 |
| 56 | 44 | 0.022 | 0.34 |
| 56 | 45 | -0.095 | 3.1e-05 |
| 56 | 46 | 0.4 | 7.3e-74 |
| 56 | 48 | -0.057 | 0.012 |
| 56 | 49 | -0.034 | 0.13 |
| 56 | 52 | -0.0012 | 0.96 |
| 56 | 54 | -0.055 | 0.016 |
| 56 | 55 | -0.066 | 0.0035 |
| 56 | 58 | -0.028 | 0.22 |
| 56 | 59 | 0.55 | 2.7e-152 |
| 56 | 60 | -0.094 | 3.2e-05 |
| 56 | 63 | 0.53 | 3.5e-138 |
| 57 | 1 | -0.056 | 0.013 |
| 57 | 2 | -0.16 | 1.9e-12 |
| 57 | 3 | -0.24 | 1.1e-26 |
| 57 | 4 | -0.11 | 7,00E-07 |
| 57 | 5 | -0.19 | 4.3e-17 |
| 57 | 6 | -0.21 | 3.2e-20 |
| 57 | 7 | -0.16 | 3.5e-13 |
| 57 | 9 | -0.16 | 3.5e-13 |
| 57 | 10 | -0.15 | 3.4e-11 |
| 57 | 11 | -0.32 | 3.3e-48 |
| 57 | 12 | 0.054 | 0.016 |
| 57 | 13 | -0.36 | 1.3e-61 |
| 57 | 14 | -0.14 | 8.8e-10 |
| 57 | 15 | 0.017 | 0.46 |
| 57 | 16 | -0.025 | 0.27 |
| 57 | 18 | 0.17 | 5.3e-14 |
| 57 | 19 | 0.011 | 0.63 |
| 57 | 20 | -0.011 | 0.62 |
| 57 | 21 | -0.25 | 4.6e-28 |

| | | | |
|---|---|---|---|
| 7 | 23 | 0.23 | 3.6e-25 |
| 7 | 24 | 0.25 | 7.2e-29 |
| 7 | 25 | 0.17 | 9.4e-15 |
| 7 | 26 | 0.19 | 2.5e-17 |
| 7 | 27 | 0.11 | 1.7e-06 |
| 7 | 28 | 0.057 | 0.012 |
| 7 | 29 | 0.21 | 1.3e-21 |
| 7 | 30 | 0.28 | 5.2e-37 |
| 7 | 32 | 0.025 | 0.27 |
| 7 | 34 | 0.067 | 0.0034 |
| 7 | 38 | 0.0096 | 0.67 |
| 7 | 39 | -0.0018 | 0.94 |
| 7 | 40 | -0.048 | 0.033 |
| 7 | 41 | 0.012 | 0.61 |
| 7 | 42 | 0.0089 | 0.69 |
| 7 | 43 | 0.013 | 0.56 |
| 7 | 45 | -0.026 | 0.25 |
| 7 | 46 | -0.03 | 0.19 |
| 7 | 48 | -0.029 | 0.2 |
| 7 | 49 | 0.26 | 6.1e-31 |
| 7 | 52 | -0.049 | 0.033 |
| 7 | 54 | -0.038 | 0.092 |
| 7 | 55 | -0.075 | 0.00093 |
| 7 | 58 | 0.42 | 5.4e-84 |
| 7 | 60 | 0.0028 | 0.9 |
| 8 | 1 | -0.044 | 0.053 |
| 8 | 2 | -0.18 | 1.1e-15 |
| 8 | 3 | -0.24 | 1.5e-27 |
| 8 | 4 | -0.14 | 1.5e-09 |
| 8 | 5 | -0.2 | 1.9e-19 |
| 8 | 6 | -0.18 | 1.2e-15 |
| 8 | 7 | 0.071 | 0.0017 |
| 8 | 9 | 0.071 | 0.0017 |
| 8 | 10 | 0.16 | 2.3e-12 |
| 8 | 11 | -0.093 | 3.8e-05 |
| 8 | 12 | 0.21 | 1.5e-20 |
| 8 | 13 | -0.44 | 2.2e-93 |
| 8 | 14 | 0.34 | 3.5e-53 |
| 8 | 15 | 0.1 | 8.2e-06 |
| 8 | 16 | 0.0015 | 0.95 |
| 8 | 17 | 0.37 | 6.9e-64 |
| 8 | 18 | 0.33 | 9.7e-51 |
| 8 | 19 | -0.05 | 0.028 |
| 8 | 20 | 0.03 | 0.19 |
| 8 | 21 | -0.24 | 2.6e-26 |
| 8 | 22 | -0.047 | 0.039 |
| 8 | 23 | -0.087 | 0.00012 |
| 8 | 24 | -0.018 | 0.43 |
| 8 | 25 | -2,00E-04 | 0.99 |
| 8 | 26 | -0.077 | 0.00065 |
| 8 | 27 | -0.12 | 7.3e-08 |
| 8 | 28 | -0.12 | 3.5e-07 |
| 8 | 29 | -0.17 | 1.1e-14 |
| 8 | 30 | -0.18 | 6.4e-15 |
| 8 | 31 | -0.019 | 0.4 |
| 8 | 32 | -0.13 | 9.1e-09 |
| 8 | 33 | 0.14 | 1.4e-09 |
| 8 | 34 | 0.084 | 0.00021 |
| 8 | 35 | 0.35 | 1.6e-56 |
| 8 | 36 | 0.05 | 0.029 |
| 8 | 37 | 0.11 | 9,00E-07 |
| 8 | 38 | -0.29 | 1.1e-39 |
| 8 | 39 | -0.092 | 4.9e-05 |
| 8 | 40 | 0.039 | 0.084 |
| 8 | 41 | -0.11 | 3.4e-06 |
| 8 | 42 | -0.054 | 0.017 |
| 8 | 43 | -0.23 | 1.9e-24 |
| 8 | 44 | -0.029 | 0.21 |
| 8 | 45 | 0.069 | 0.0024 |
| 8 | 46 | 0.13 | 1.1e-08 |
| 8 | 47 | 0.22 | 6.7e-22 |
| 8 | 48 | 0.14 | 2.1e-09 |
| 8 | 49 | -0.088 | 0.00011 |
| 8 | 50 | 0.25 | 1,00E-28 |
| 8 | 51 | 0.26 | 2.7e-30 |
| 8 | 52 | 0.046 | 0.045 |
| 8 | 53 | 0.24 | 9.4e-28 |
| 8 | 54 | -0.087 | 0.00013 |
| 8 | 55 | -0.077 | 0.00071 |
| 8 | 56 | 0.022 | 0.34 |
| 8 | 57 | 0.22 | 3.6e-22 |
| 8 | 58 | -0.15 | 1.2e-11 |
| 8 | 59 | -0.17 | 2.6e-13 |
| 8 | 60 | 0.02 | 0.38 |
| 8 | 61 | -0.096 | 2.2e-05 |
| 8 | 62 | 0.28 | 1.1e-35 |
| 8 | 63 | -0.11 | 3.2e-06 |
| 9 | 1 | -0.009 | 0.69 |
| 9 | 2 | 0.16 | 1.3e-12 |
| 9 | 3 | 0.19 | 8.1e-17 |
| 9 | 4 | 0.15 | 3.9e-11 |
| 9 | 5 | 0.18 | 5.6e-15 |
| 9 | 6 | 0.1 | 9.2e-06 |
| 9 | 10 | 0.55 | 1.4e-153 |
| 9 | 11 | 0.41 | 1.8e-77 |
| 9 | 13 | 0.57 | 3.2e-170 |
| 9 | 14 | -0.053 | 0.019 |
| 9 | 15 | 0.15 | 8.7e-11 |
| 9 | 19 | 0.02 | 0.38 |
| 9 | 21 | -0.03 | 0.19 |
| 9 | 22 | -0.043 | 0.059 |
| 9 | 23 | 0.23 | 3.5e-25 |
| 9 | 24 | 0.25 | 7.1e-29 |
| 9 | 25 | 0.17 | 9.4e-15 |
| 9 | 26 | 0.19 | 2.5e-17 |
| 17 | 55 | -0.15 | 9.9e-12 |
| 17 | 56 | 0.017 | 0.47 |
| 17 | 57 | 0.23 | 6.3e-24 |
| 17 | 58 | -0.19 | 2,00E-17 |
| 17 | 59 | -0.011 | 0.63 |
| 17 | 60 | 0.015 | 0.5 |
| 17 | 61 | -0.022 | 0.34 |
| 17 | 62 | 0.3 | 1.4e-41 |
| 17 | 63 | 0.064 | 0.0051 |
| 18 | 1 | 0.044 | 0.054 |
| 18 | 2 | -0.16 | 5.7e-12 |
| 18 | 3 | -0.29 | 3.5e-39 |
| 18 | 4 | -0.17 | 7.1e-14 |
| 18 | 5 | -0.3 | 7.7e-41 |
| 18 | 6 | -0.32 | 2.5e-46 |
| 18 | 7 | 0.15 | 2.4e-11 |
| 18 | 9 | 0.15 | 2.4e-11 |
| 18 | 10 | 0.15 | 4.5e-11 |
| 18 | 11 | -0.29 | 9.3e-39 |
| 18 | 13 | -0.21 | 5.2e-20 |
| 18 | 14 | -0.094 | 3.6e-05 |
| 18 | 15 | 0.26 | 4,00E-30 |
| 18 | 16 | 0.22 | 1.1e-22 |
| 18 | 19 | -0.088 | 0.00011 |
| 18 | 21 | -0.34 | 3.1e-53 |
| 18 | 22 | -0.039 | 0.086 |
| 18 | 23 | -0.14 | 5.6e-10 |
| 18 | 24 | 0.059 | 0.0097 |
| 18 | 25 | 0.061 | 0.007 |
| 18 | 26 | -0.2 | 1.9e-19 |
| 18 | 27 | -0.32 | 1.3e-46 |
| 18 | 28 | -0.043 | 0.06 |
| 18 | 29 | -0.1 | 4.7e-06 |
| 18 | 30 | -0.19 | 2.1e-17 |
| 18 | 31 | -0.063 | 0.0059 |
| 18 | 32 | -0.14 | 1.9e-09 |
| 18 | 34 | -0.024 | 0.28 |
| 18 | 36 | 0.07 | 0.0021 |
| 18 | 38 | -0.13 | 7.6e-09 |
| 18 | 39 | -0.016 | 0.48 |
| 18 | 40 | -0.038 | 0.093 |
| 18 | 41 | -0.24 | 7.9e-28 |
| 18 | 42 | -0.054 | 0.018 |
| 18 | 43 | -0.13 | 2.7e-08 |
| 18 | 44 | 0.0074 | 0.75 |
| 18 | 45 | 0.02 | 0.37 |
| 18 | 46 | 0.12 | 1.7e-07 |
| 18 | 48 | 0.072 | 0.0015 |
| 18 | 49 | -0.17 | 1.5e-13 |
| 18 | 52 | -0.0065 | 0.78 |
| 18 | 54 | -0.25 | 1,00E-28 |
| 18 | 55 | -0.3 | 1.9e-42 |
| 18 | 56 | 0.043 | 0.059 |
| 18 | 58 | -0.32 | 1,00E-46 |
| 18 | 59 | -0.083 | 0.00025 |
| 18 | 60 | 0.074 | 0.0012 |
| 18 | 61 | -0.031 | 0.17 |
| 18 | 63 | 0.022 | 0.32 |
| 19 | 2 | 0.043 | 0.057 |
| 19 | 3 | 0.085 | 0.00019 |
| 19 | 4 | 0.077 | 0.00071 |
| 19 | 5 | 0.12 | 3.3e-07 |
| 19 | 21 | 0.14 | 4.6e-10 |
| 19 | 22 | 0.75 | 0 |
| 19 | 27 | 0.67 | 4,00E-256 |
| 19 | 34 | -0.0026 | 0.91 |
| 19 | 38 | 0.0092 | 0.68 |
| 19 | 39 | -0.025 | 0.27 |
| 19 | 40 | 0.089 | 8.6e-05 |
| 19 | 41 | 0.53 | 1,00E-137 |
| 19 | 42 | 0.08 | 4,00E-04 |
| 19 | 48 | -0.064 | 0.0049 |
| 19 | 60 | 0.031 | 0.17 |
| 20 | 1 | 0.0059 | 0.8 |
| 20 | 2 | -0.15 | 6.3e-11 |
| 20 | 3 | -0.17 | 1,00E-14 |
| 20 | 4 | -0.14 | 1.3e-09 |
| 20 | 5 | -0.17 | 1.1e-13 |
| 20 | 6 | -0.097 | 1.8e-05 |
| 20 | 7 | -0.12 | 1.6e-07 |
| 20 | 9 | -0.12 | 1.6e-07 |
| 20 | 10 | -0.14 | 1.5e-09 |
| 20 | 11 | 0.0057 | 0.8 |
| 20 | 12 | 0.18 | 3.2e-16 |
| 20 | 13 | -0.17 | 3.2e-14 |
| 20 | 14 | -0.1 | 1.1e-05 |
| 20 | 15 | -0.096 | 2.3e-05 |
| 20 | 16 | -0.073 | 0.0014 |
| 20 | 18 | 0.092 | 5,00E-05 |
| 20 | 19 | -0.09 | 8,00E-05 |
| 20 | 21 | -0.18 | 3.5e-15 |
| 20 | 22 | -0.00066 | 0.98 |
| 20 | 23 | -0.084 | 0.00021 |
| 20 | 24 | -0.017 | 0.45 |
| 20 | 25 | 0.063 | 0.0053 |
| 20 | 26 | -0.12 | 2.2e-07 |
| 20 | 27 | -0.2 | 1.2e-19 |
| 20 | 28 | 0.021 | 0.35 |
| 20 | 29 | -0.022 | 0.34 |
| 20 | 30 | -0.093 | 4.4e-05 |
| 20 | 31 | -0.029 | 0.21 |
| 20 | 32 | -0.038 | 0.098 |
| 20 | 33 | 0.52 | 1.6e-134 |
| 20 | 34 | 0.086 | 0.00014 |
| 20 | 36 | -0.0024 | 0.92 |
| 31 | 23 | -0.0051 | 0.82 |
| 31 | 24 | 0.11 | 4.1e-07 |
| 31 | 25 | 0.17 | 1.1e-13 |
| 31 | 26 | 0.076 | 0.00085 |
| 31 | 27 | 0.13 | 3.2e-08 |
| 31 | 28 | -0.51 | 2.4e-127 |
| 31 | 29 | -0.017 | 0.47 |
| 31 | 30 | 0.16 | 3.7e-13 |
| 31 | 32 | 0.46 | 2.1e-103 |
| 31 | 34 | -0.0095 | 0.68 |
| 31 | 36 | 0.6 | 4,00E-192 |
| 31 | 38 | 0.03 | 0.19 |
| 31 | 39 | 0.0017 | 0.94 |
| 31 | 40 | 0.029 | 0.2 |
| 31 | 41 | -0.054 | 0.018 |
| 31 | 42 | 0.015 | 0.5 |
| 31 | 43 | 0.018 | 0.43 |
| 31 | 44 | 0.62 | 1.4e-202 |
| 31 | 45 | 0.0094 | 0.68 |
| 31 | 46 | 0.011 | 0.62 |
| 31 | 48 | -0.00056 | 0.98 |
| 31 | 49 | 0.1 | 5.1e-06 |
| 31 | 52 | -0.002 | 0.93 |
| 31 | 54 | -0.017 | 0.45 |
| 31 | 55 | -0.022 | 0.33 |
| 31 | 58 | 0.09 | 7.9e-05 |
| 31 | 60 | 0.029 | 0.2 |
| 32 | 1 | 0.037 | 0.11 |
| 32 | 2 | 0.071 | 0.0016 |
| 32 | 3 | 0.13 | 1.2e-08 |
| 32 | 4 | 0.048 | 0.034 |
| 32 | 5 | 0.1 | 1,00E-05 |
| 32 | 6 | 0.13 | 4.9e-09 |
| 32 | 10 | 0.04 | 0.077 |
| 32 | 11 | 0.13 | 2.8e-08 |
| 32 | 13 | 0.21 | 1.9e-20 |
| 32 | 14 | 0.14 | 9.9e-10 |
| 32 | 15 | 0.014 | 0.54 |
| 32 | 19 | 0.042 | 0.067 |
| 32 | 21 | 0.16 | 6.5e-13 |
| 32 | 22 | 0.013 | 0.57 |
| 32 | 23 | 0.00093 | 0.97 |
| 32 | 26 | -0.064 | 0.0047 |
| 32 | 27 | 0.19 | 2.2e-16 |
| 32 | 29 | 0.2 | 2.4e-18 |
| 32 | 30 | 0.28 | 1.1e-35 |
| 32 | 34 | 0.063 | 0.0052 |
| 32 | 38 | 0.23 | 5.1e-25 |
| 32 | 39 | -0.1 | 1.1e-05 |
| 32 | 40 | 0.045 | 0.049 |
| 32 | 41 | 0.088 | 0.00011 |
| 32 | 42 | 0.13 | 1.6e-08 |
| 32 | 45 | 0.041 | 0.069 |
| 32 | 46 | -0.047 | 0.038 |
| 32 | 48 | 0.00079 | 0.97 |
| 32 | 49 | 0.047 | 0.04 |
| 32 | 52 | 0.05 | 0.027 |
| 32 | 54 | -0.0091 | 0.69 |
| 32 | 55 | 0.019 | 0.4 |
| 32 | 58 | 0.16 | 4.8e-12 |
| 32 | 60 | 0.073 | 0.0014 |
| 33 | 1 | 0.0036 | 0.87 |
| 33 | 2 | -0.23 | 5.7e-25 |
| 33 | 3 | -0.21 | 4.2e-20 |
| 33 | 4 | -0.18 | 8.3e-15 |
| 33 | 5 | -0.18 | 8,00E-15 |
| 33 | 6 | -0.023 | 0.32 |
| 33 | 7 | 0.1 | 4.7e-06 |
| 33 | 9 | 0.1 | 4.7e-06 |
| 33 | 10 | -0.0036 | 0.88 |
| 33 | 11 | 0.13 | 3.8e-09 |
| 33 | 12 | 0.14 | 1.5e-09 |
| 33 | 13 | -0.068 | 0.0026 |
| 33 | 14 | -0.052 | 0.022 |
| 33 | 15 | 0.0038 | 0.87 |
| 33 | 16 | 0.13 | 2.8e-08 |
| 33 | 18 | 0.12 | 2.8e-07 |
| 33 | 19 | -0.054 | 0.017 |
| 33 | 21 | -0.22 | 7.9e-23 |
| 33 | 22 | -0.00056 | 0.98 |
| 33 | 23 | 0.017 | 0.46 |
| 33 | 24 | 0.0066 | 0.77 |
| 33 | 25 | 0.083 | 0.00026 |
| 33 | 26 | 0.013 | 0.56 |
| 33 | 27 | -0.13 | 3.4e-09 |
| 33 | 28 | -0.0078 | 0.73 |
| 33 | 29 | -0.0016 | 0.94 |
| 33 | 30 | -0.097 | 1.8e-05 |
| 33 | 31 | -0.006 | 0.79 |
| 33 | 32 | -0.058 | 0.01 |
| 33 | 34 | 0.056 | 0.013 |
| 33 | 36 | 0.048 | 0.036 |
| 33 | 37 | 0.18 | 8.6e-16 |
| 33 | 38 | 0.051 | 0.026 |
| 33 | 39 | 0.042 | 0.065 |
| 33 | 40 | -0.15 | 1,00E-11 |
| 33 | 41 | -0.043 | 0.058 |
| 33 | 42 | -0.099 | 1.2e-05 |
| 33 | 43 | 0.017 | 0.47 |
| 33 | 44 | 0.02 | 0.39 |
| 33 | 45 | 0.058 | 0.011 |
| 33 | 46 | 0.15 | 1.3e-11 |
| 33 | 48 | 0.076 | 0.00077 |
| 33 | 49 | -0.0078 | 0.73 |
| 33 | 52 | -0.013 | 0.56 |
| 46 | 41 | -0.05 | 0.029 |
| 46 | 42 | -0.36 | 5,00E-59 |
| 46 | 48 | 0.065 | 0.0043 |
| 46 | 60 | 0.26 | 1.1e-31 |
| 47 | 1 | -0.06 | 0.0084 |
| 47 | 2 | -0.18 | 7.1e-16 |
| 47 | 3 | -0.28 | 1.4e-35 |
| 47 | 4 | -0.12 | 6.6e-08 |
| 47 | 5 | -0.22 | 4.8e-22 |
| 47 | 6 | -0.24 | 5.5e-27 |
| 47 | 7 | -0.14 | 2.6e-10 |
| 47 | 9 | -0.14 | 2.5e-10 |
| 47 | 10 | -0.25 | 5.9e-29 |
| 47 | 11 | -0.36 | 1.1e-59 |
| 47 | 12 | 0.018 | 0.44 |
| 47 | 13 | -0.4 | 3.8e-76 |
| 47 | 14 | -0.21 | 1.4e-21 |
| 47 | 15 | -0.087 | 0.00014 |
| 47 | 16 | 0.02 | 0.37 |
| 47 | 18 | 0.17 | 9.7e-15 |
| 47 | 19 | -0.0092 | 0.68 |
| 47 | 20 | 0.015 | 0.51 |
| 47 | 21 | -0.25 | 2.8e-29 |
| 47 | 22 | -0.023 | 0.32 |
| 47 | 23 | -0.19 | 4.9e-17 |
| 47 | 24 | 0.058 | 0.011 |
| 47 | 25 | 0.019 | 0.41 |
| 47 | 26 | -0.21 | 2.4e-20 |
| 47 | 27 | -0.12 | 5.4e-08 |
| 47 | 28 | -0.17 | 5,00E-14 |
| 47 | 29 | -0.32 | 2,00E-47 |
| 47 | 30 | -0.21 | 2.2e-21 |
| 47 | 31 | -0.078 | 0.00056 |
| 47 | 32 | -0.26 | 2,00E-30 |
| 47 | 33 | 0.0039 | 0.86 |
| 47 | 34 | -0.029 | 0.2 |
| 47 | 36 | 0.071 | 0.0019 |
| 47 | 37 | 0.1 | 4.5e-06 |
| 47 | 38 | -0.49 | 9,00E-116 |
| 47 | 39 | -0.18 | 3.3e-16 |
| 47 | 40 | -0.058 | 0.011 |
| 47 | 41 | -0.21 | 7.7e-20 |
| 47 | 42 | -0.026 | 0.25 |
| 47 | 43 | -0.35 | 2.6e-55 |
| 47 | 44 | -0.0026 | 0.91 |
| 47 | 45 | -0.37 | 2.8e-62 |
| 47 | 46 | -0.48 | 1.2e-110 |
| 47 | 48 | 0.14 | 1,00E-09 |
| 47 | 49 | -0.03 | 0.19 |
| 47 | 52 | 0.013 | 0.56 |
| 47 | 54 | -0.07 | 0.002 |
| 47 | 55 | -0.077 | 0.00075 |
| 47 | 56 | -0.088 | 1,00E-04 |
| 47 | 57 | 0.68 | 3.3e-258 |
| 47 | 58 | -0.3 | 2.6e-40 |
| 47 | 59 | -0.24 | 4.6e-27 |
| 47 | 60 | -0.21 | 4.5e-21 |
| 47 | 61 | 0.23 | 2.2e-25 |
| 47 | 62 | 0.64 | 1.2e-224 |
| 47 | 63 | -0.062 | 0.0062 |
| 48 | 38 | -0.095 | 2.9e-05 |
| 48 | 39 | -0.023 | 0.31 |
| 48 | 40 | 0.051 | 0.025 |
| 48 | 60 | 0.023 | 0.32 |
| 49 | 2 | 0.076 | 0.00086 |
| 49 | 3 | 0.14 | 1.5e-10 |
| 49 | 4 | 0.077 | 0.00069 |
| 49 | 5 | 0.13 | 6.1e-09 |
| 49 | 19 | 0.035 | 0.12 |
| 49 | 21 | 0.085 | 0.00019 |
| 49 | 22 | 0.014 | 0.54 |
| 49 | 27 | 0.16 | 3.7e-13 |
| 49 | 34 | -0.3 | 1.2e-41 |
| 49 | 38 | 0.044 | 0.052 |
| 49 | 39 | -0.13 | 3.2e-08 |
| 49 | 40 | 0.018 | 0.43 |
| 49 | 41 | 0.18 | 6.2e-16 |
| 49 | 42 | 0.0022 | 0.92 |
| 49 | 45 | 0.031 | 0.17 |
| 49 | 46 | 0.037 | 0.11 |
| 49 | 48 | 0.038 | 0.094 |
| 49 | 52 | -0.052 | 0.021 |
| 49 | 60 | 0.0036 | 0.88 |
| 50 | 1 | -0.072 | 0.0014 |
| 50 | 2 | -0.18 | 2.4e-16 |
| 50 | 3 | -0.31 | 5.6e-43 |
| 50 | 4 | -0.12 | 3.8e-08 |
| 50 | 5 | -0.24 | 1,00E-27 |
| 50 | 6 | -0.29 | 3.8e-40 |
| 50 | 7 | -0.15 | 1.6e-11 |
| 50 | 9 | -0.15 | 1.6e-11 |
| 50 | 10 | -0.24 | 3.9e-27 |
| 50 | 11 | -0.19 | 8.8e-18 |
| 50 | 12 | 0.4 | 1.7e-76 |
| 50 | 13 | -0.44 | 9.5e-93 |
| 50 | 14 | -0.23 | 8.3e-24 |
| 50 | 15 | -0.13 | 3.4e-09 |
| 50 | 16 | -0.034 | 0.13 |
| 50 | 18 | 0.43 | 2.9e-89 |
| 50 | 19 | -0.13 | 1.2e-08 |
| 50 | 20 | 0.23 | 1.1e-23 |
| 50 | 21 | -0.33 | 5.4e-49 |
| 50 | 22 | -0.056 | 0.014 |
| 50 | 23 | -0.25 | 2.2e-28 |
| 50 | 24 | 0.057 | 0.012 |
| 57 | 22 | -0.018 | 0.42 |
| 57 | 23 | -0.14 | 9.2e-10 |
| 57 | 24 | 0.034 | 0.14 |
| 57 | 25 | 0.028 | 0.22 |
| 57 | 26 | -0.12 | 1.8e-07 |
| 57 | 27 | -0.1 | 4.8e-06 |
| 57 | 28 | -0.17 | 5,00E-14 |
| 57 | 29 | -0.28 | 1.7e-36 |
| 57 | 30 | -0.22 | 1.9e-23 |
| 57 | 31 | -0.071 | 0.0019 |
| 57 | 32 | -0.26 | 3.9e-30 |
| 57 | 33 | 0.0083 | 0.72 |
| 57 | 34 | -0.14 | 7.4e-10 |
| 57 | 36 | 0.086 | 0.00015 |
| 57 | 37 | 0.14 | 3.2e-10 |
| 57 | 38 | -0.38 | 1.9e-67 |
| 57 | 39 | -0.16 | 6.9e-12 |
| 57 | 40 | -0.014 | 0.55 |
| 57 | 41 | -0.13 | 4.3e-09 |
| 57 | 42 | 0.068 | 0.003 |
| 57 | 43 | -0.26 | 1.8e-31 |
| 57 | 44 | 0.034 | 0.13 |
| 57 | 45 | 0.22 | 1.1e-22 |
| 57 | 46 | -0.088 | 9.8e-05 |
| 57 | 48 | 0.0083 | 0.72 |
| 57 | 49 | -0.032 | 0.16 |
| 57 | 52 | 0.041 | 0.072 |
| 57 | 54 | -0.039 | 0.088 |
| 57 | 55 | -0.022 | 0.33 |
| 57 | 56 | -0.15 | 1.8e-11 |
| 57 | 58 | -0.29 | 2.3e-38 |
| 57 | 59 | -0.08 | 0.00043 |
| 57 | 60 | -0.46 | 1,00E-100 |
| 57 | 61 | -0.33 | 2.4e-49 |
| 57 | 62 | 0.81 | 0 |
| 57 | 63 | 0.084 | 0.00021 |
| 58 | 1 | -0.00014 | 1 |
| 58 | 2 | 0.15 | 2.3e-11 |
| 58 | 3 | 0.26 | 5.8e-32 |
| 58 | 4 | 0.16 | 4.6e-12 |
| 58 | 5 | 0.25 | 1.3e-29 |
| 58 | 6 | 0.27 | 4.3e-34 |
| 58 | 10 | 0.27 | 7,00E-34 |
| 58 | 14 | 0.24 | 1,00E-26 |
| 58 | 15 | 0.08 | 4,00E-04 |
| 58 | 19 | 0.14 | 7.9e-10 |
| 58 | 21 | 0.23 | 1.2e-25 |
| 58 | 22 | 0.079 | 5,00E-04 |
| 58 | 23 | 0.25 | 6.2e-30 |
| 58 | 26 | 0.29 | 6.7e-40 |
| 58 | 27 | 0.35 | 1.1e-56 |
| 58 | 29 | 0.27 | 2.3e-33 |
| 58 | 30 | 0.35 | 3.8e-57 |
| 58 | 34 | 0.18 | 2.8e-15 |
| 58 | 38 | 0.27 | 2.2e-34 |
| 58 | 39 | 0.078 | 0.00063 |
| 58 | 40 | 0.065 | 0.0043 |
| 58 | 41 | 0.26 | 1.3e-31 |
| 58 | 42 | 0.056 | 0.014 |
| 58 | 45 | -0.07 | 0.0021 |
| 58 | 46 | 0.0052 | 0.82 |
| 58 | 48 | -0.17 | 5.7e-14 |
| 58 | 49 | 0.42 | 9.8e-86 |
| 58 | 52 | 0.052 | 0.021 |
| 58 | 54 | 0.079 | 0.00051 |
| 58 | 55 | 0.13 | 2.5e-08 |
| 58 | 60 | 0.23 | 1.4e-24 |
| 59 | 1 | -0.016 | 0.48 |
| 59 | 2 | 0.16 | 2.7e-12 |
| 59 | 3 | 0.15 | 5.2e-11 |
| 59 | 4 | 0.15 | 6.5e-11 |
| 59 | 5 | 0.15 | 1.3e-10 |
| 59 | 6 | 0.027 | 0.24 |
| 59 | 7 | 0.055 | 0.016 |
| 59 | 9 | 0.055 | 0.016 |
| 59 | 10 | 0.041 | 0.069 |
| 59 | 11 | 0.0068 | 0.77 |
| 59 | 13 | 0.16 | 1.1e-12 |
| 59 | 14 | 0.0076 | 0.74 |
| 59 | 15 | 0.024 | 0.29 |
| 59 | 16 | 0.04 | 0.081 |
| 59 | 19 | 0.048 | 0.034 |
| 59 | 21 | 0.076 | 0.00077 |
| 59 | 22 | 0.017 | 0.46 |
| 59 | 23 | 0.075 | 0.00098 |
| 59 | 24 | -0.038 | 0.095 |
| 59 | 25 | 0.0025 | 0.91 |
| 59 | 26 | 0.15 | 1.3e-10 |
| 59 | 27 | 0.1 | 3.8e-06 |
| 59 | 28 | 0.019 | 0.4 |
| 59 | 29 | 0.05 | 0.029 |
| 59 | 30 | 0.025 | 0.27 |
| 59 | 31 | -0.013 | 0.57 |
| 59 | 32 | 0.0029 | 0.9 |
| 59 | 34 | -0.14 | 1.5e-10 |
| 59 | 36 | -0.016 | 0.48 |
| 59 | 38 | 0.17 | 1.7e-14 |
| 59 | 39 | 0.07 | 0.0021 |
| 59 | 40 | -0.078 | 0.00057 |
| 59 | 41 | 0.14 | 2.1e-09 |
| 59 | 42 | -0.065 | 0.0043 |
| 59 | 43 | 0.12 | 1.4e-07 |
| 59 | 44 | 0.025 | 0.28 |
| 59 | 45 | -0.079 | 0.00053 |
| 59 | 46 | -0.0085 | 0.71 |

| | | | |
|---|---|---|---|
| 9 | 27 | 0.11 | 1.7e-06 |
| 9 | 28 | 0.057 | 0.012 |
| 9 | 29 | 0.21 | 1.2e-21 |
| 9 | 30 | 0.28 | 4.9e-37 |
| 9 | 32 | 0.025 | 0.27 |
| 9 | 34 | 0.067 | 0.0034 |
| 9 | 38 | 0.0097 | 0.67 |
| 9 | 39 | -0.0017 | 0.94 |
| 9 | 40 | -0.048 | 0.033 |
| 9 | 41 | 0.012 | 0.61 |
| 9 | 42 | 0.009 | 0.69 |
| 9 | 43 | 0.013 | 0.55 |
| 9 | 45 | -0.026 | 0.25 |
| 9 | 46 | -0.03 | 0.19 |
| 9 | 48 | -0.029 | 0.2 |
| 9 | 49 | 0.26 | 5.9e-31 |
| 9 | 52 | -0.048 | 0.033 |
| 9 | 54 | -0.038 | 0.092 |
| 9 | 55 | -0.075 | 0.00093 |
| 9 | 58 | 0.42 | 4.9e-84 |
| 9 | 60 | 0.0028 | 0.9 |
| 10 | 2 | 0.16 | 7.3e-13 |
| 10 | 3 | 0.19 | 1.1e-17 |
| 10 | 4 | 0.11 | 9.6e-07 |
| 10 | 5 | 0.14 | 1.5e-10 |
| 10 | 14 | 0.17 | 2.8e-14 |
| 10 | 15 | 0.59 | 2.1e-184 |
| 10 | 19 | 0.038 | 0.099 |
| 10 | 21 | 0.056 | 0.014 |
| 10 | 22 | 0.00096 | 0.97 |
| 10 | 27 | 0.13 | 2.1e-08 |
| 10 | 34 | 0.049 | 0.03 |
| 10 | 38 | 0.093 | 4.4e-05 |
| 10 | 39 | 0.014 | 0.54 |
| 10 | 40 | 0.031 | 0.17 |
| 10 | 41 | 0.064 | 0.0049 |
| 10 | 42 | 0.072 | 0.0015 |
| 10 | 45 | 0.11 | 5.1e-07 |
| 10 | 46 | 0.12 | 3.2e-07 |
| 10 | 48 | -0.027 | 0.23 |
| 10 | 49 | 0.14 | 1.5e-09 |
| 10 | 52 | -0.039 | 0.086 |
| 10 | 60 | 0.1 | 4.3e-06 |
| 11 | 1 | 0.0033 | 0.89 |
| 11 | 2 | 0.07 | 0.0021 |
| 11 | 3 | 0.17 | 1.6e-13 |
| 11 | 4 | 0.063 | 0.0058 |
| 11 | 5 | 0.15 | 2,00E-11 |
| 11 | 6 | 0.22 | 6.3e-22 |
| 11 | 10 | 0.39 | 5.7e-73 |
| 11 | 13 | 0.43 | 2.6e-86 |
| 11 | 14 | 0.13 | 1.9e-08 |
| 11 | 15 | 0.11 | 6.9e-07 |
| 11 | 19 | 0.059 | 0.0091 |
| 11 | 21 | 0.12 | 2,00E-07 |
| 11 | 22 | 0.041 | 0.074 |
| 11 | 23 | 0.24 | 2.4e-26 |
| 11 | 26 | 0.18 | 4.6e-15 |
| 11 | 27 | 0.17 | 5.1e-14 |
| 11 | 29 | 0.26 | 3.9e-32 |
| 11 | 30 | 0.32 | 9.3e-49 |
| 11 | 34 | 0.22 | 1.1e-22 |
| 11 | 38 | 0.28 | 2.3e-37 |
| 11 | 39 | 0.16 | 3.3e-12 |
| 11 | 40 | 0.078 | 0.00063 |
| 11 | 41 | 0.087 | 0.00012 |
| 11 | 42 | 0.032 | 0.16 |
| 11 | 45 | 0.11 | 2.6e-06 |
| 11 | 46 | 0.15 | 8.7e-12 |
| 11 | 48 | 0.039 | 0.086 |
| 11 | 49 | 0.19 | 1.7e-16 |
| 11 | 52 | 0.037 | 0.1 |
| 11 | 54 | -0.00022 | 0.99 |
| 11 | 55 | 0.022 | 0.34 |
| 11 | 58 | 0.5 | 6.5e-125 |
| 11 | 60 | 0.13 | 1.8e-08 |
| 12 | 1 | 0.052 | 0.023 |
| 12 | 2 | -0.22 | 1.9e-23 |
| 12 | 3 | -0.31 | 1.3e-43 |
| 12 | 4 | -0.23 | 2.1e-25 |
| 12 | 5 | -0.32 | 6.3e-46 |
| 12 | 6 | -0.23 | 1.5e-24 |
| 12 | 7 | -0.3 | 3.1e-40 |
| 12 | 9 | -0.3 | 3.1e-40 |
| 12 | 10 | 0.029 | 0.2 |
| 12 | 11 | 0.14 | 2.2e-10 |
| 12 | 13 | -0.32 | 2.6e-47 |
| 12 | 14 | 0.039 | 0.083 |
| 12 | 15 | 0.23 | 2.2e-24 |
| 12 | 16 | -0.19 | 3,00E-17 |
| 12 | 18 | 0.67 | 5.4e-257 |
| 12 | 19 | -0.058 | 0.01 |
| 12 | 21 | -0.23 | 2.3e-24 |
| 12 | 22 | 0.023 | 0.3 |
| 12 | 23 | -0.14 | 7,00E-10 |
| 12 | 24 | 0.023 | 0.31 |
| 12 | 25 | 0.04 | 0.081 |
| 12 | 26 | -0.22 | 1,00E-21 |
| 12 | 27 | -0.27 | 3,00E-34 |
| 12 | 28 | -0.012 | 0.59 |
| 12 | 29 | -0.07 | 0.0021 |
| 12 | 30 | -0.16 | 6.1e-13 |
| 12 | 31 | -0.027 | 0.23 |
| 12 | 32 | -0.061 | 0.0069 |
| 12 | 34 | 0.089 | 8.3e-05 |

| | | | |
|---|---|---|---|
| 20 | 37 | 0.15 | 8,00E-11 |
| 20 | 38 | 0.084 | 0.00023 |
| 20 | 39 | 0.084 | 2,00E-04 |
| 20 | 40 | -0.1 | 4,00E-06 |
| 20 | 41 | -0.14 | 5.5e-10 |
| 20 | 42 | -0.048 | 0.033 |
| 20 | 43 | 0.0077 | 0.74 |
| 20 | 44 | -0.03 | 0.19 |
| 20 | 45 | 0.051 | 0.025 |
| 20 | 46 | 0.085 | 0.00018 |
| 20 | 48 | 0.1 | 7.2e-06 |
| 20 | 49 | -0.14 | 1.5e-09 |
| 20 | 52 | 0.071 | 0.0017 |
| 20 | 54 | -0.069 | 0.0026 |
| 20 | 55 | -0.04 | 0.081 |
| 20 | 56 | 0.071 | 0.0017 |
| 20 | 58 | -0.14 | 2.1e-10 |
| 20 | 59 | -0.096 | 2.4e-05 |
| 20 | 60 | -0.012 | 0.6 |
| 20 | 61 | -0.016 | 0.49 |
| 20 | 63 | -0.081 | 0.00039 |
| 21 | 2 | 0.14 | 4.3e-10 |
| 21 | 3 | 0.2 | 1.1e-18 |
| 21 | 4 | 0.12 | 9.6e-08 |
| 21 | 5 | 0.18 | 4.1e-15 |
| 21 | 34 | 0.062 | 0.0065 |
| 21 | 38 | 0.25 | 1.8e-29 |
| 21 | 39 | 0.11 | 2.5e-06 |
| 21 | 40 | 0.12 | 1.3e-07 |
| 21 | 42 | 0.12 | 8.6e-08 |
| 21 | 48 | -0.029 | 0.2 |
| 21 | 60 | 0.073 | 0.0012 |
| 22 | 2 | 0.016 | 0.48 |
| 22 | 3 | 0.053 | 0.019 |
| 22 | 4 | 0.01 | 0.65 |
| 22 | 5 | 0.046 | 0.042 |
| 22 | 21 | 0.075 | 0.00091 |
| 22 | 27 | 0.51 | 2.9e-130 |
| 22 | 34 | 0.016 | 0.49 |
| 22 | 38 | 0.045 | 0.047 |
| 22 | 39 | -0.018 | 0.44 |
| 22 | 40 | 0.039 | 0.085 |
| 22 | 41 | 0.54 | 8,00E-148 |
| 22 | 42 | 0.07 | 0.0021 |
| 22 | 48 | -0.021 | 0.35 |
| 22 | 60 | 0.067 | 0.0032 |
| 23 | 1 | 0.083 | 0.00026 |
| 23 | 2 | 0.066 | 0.0038 |
| 23 | 3 | 0.12 | 1.1e-07 |
| 23 | 4 | 0.011 | 0.63 |
| 23 | 5 | 0.061 | 0.0072 |
| 23 | 10 | 0.19 | 6.7e-17 |
| 23 | 14 | 0.12 | 8.5e-08 |
| 23 | 15 | 0.11 | 2.8e-06 |
| 23 | 19 | -0.31 | 1.3e-44 |
| 23 | 21 | 0.043 | 0.058 |
| 23 | 22 | -0.097 | 2.1e-05 |
| 23 | 26 | 0.47 | 4.7e-107 |
| 23 | 27 | 0.11 | 1.3e-06 |
| 23 | 34 | -0.056 | 0.014 |
| 23 | 38 | -0.039 | 0.088 |
| 23 | 39 | 0.038 | 0.096 |
| 23 | 40 | -0.02 | 0.37 |
| 23 | 41 | 0.25 | 5.7e-28 |
| 23 | 42 | 0.012 | 0.61 |
| 23 | 45 | 0.036 | 0.11 |
| 23 | 46 | 0.02 | 0.37 |
| 23 | 48 | -0.045 | 0.048 |
| 23 | 49 | 0.27 | 8.9e-35 |
| 23 | 52 | -0.082 | 0.00031 |
| 23 | 60 | -0.015 | 0.52 |
| 24 | 1 | -0.02 | 0.37 |
| 24 | 2 | -0.0061 | 0.79 |
| 24 | 3 | -0.00099 | 0.97 |
| 24 | 4 | 0.0071 | 0.76 |
| 24 | 5 | 0.012 | 0.61 |
| 24 | 6 | 0.012 | 0.6 |
| 24 | 10 | 0.16 | 1.7e-12 |
| 24 | 11 | 0.21 | 8,00E-20 |
| 24 | 13 | 0.14 | 3.3e-10 |
| 24 | 14 | -0.052 | 0.023 |
| 24 | 15 | 0.11 | 3,00E-06 |
| 24 | 19 | 0.17 | 1.7e-14 |
| 24 | 21 | -0.009 | 0.69 |
| 24 | 22 | 0.094 | 3.6e-05 |
| 24 | 23 | 0.33 | 2.3e-49 |
| 24 | 25 | 0.68 | 1.7e-264 |
| 24 | 26 | -0.032 | 0.16 |
| 24 | 27 | 0.21 | 7.3e-20 |
| 24 | 28 | 0.13 | 4.4e-09 |
| 24 | 29 | -0.021 | 0.36 |
| 24 | 30 | 0.54 | 2.6e-149 |
| 24 | 32 | -0.073 | 0.0013 |
| 24 | 34 | 0.032 | 0.16 |
| 24 | 38 | -0.22 | 5.3e-23 |
| 24 | 39 | 0.14 | 2.4e-09 |
| 24 | 40 | 0.077 | 0.00068 |
| 24 | 41 | -0.39 | 1.7e-70 |
| 24 | 42 | -0.008 | 0.73 |
| 24 | 43 | -0.4 | 3.7e-76 |
| 24 | 45 | -0.0016 | 0.94 |
| 24 | 46 | -0.015 | 0.52 |
| 24 | 48 | 0.058 | 0.01 |
| 24 | 49 | 0.099 | 1.3e-05 |
| 24 | 52 | -0.013 | 0.58 |

| | | | |
|---|---|---|---|
| 33 | 54 | 0.056 | 0.014 |
| 33 | 55 | 0.059 | 0.0091 |
| 33 | 56 | 0.12 | 1.6e-07 |
| 33 | 58 | -0.032 | 0.16 |
| 33 | 59 | -0.058 | 0.011 |
| 33 | 60 | -3,00E-04 | 0.99 |
| 33 | 61 | -0.035 | 0.12 |
| 33 | 63 | -0.036 | 0.11 |
| 34 | 38 | 0.05 | 0.028 |
| 34 | 39 | 0.098 | 1.7e-05 |
| 34 | 40 | 0.073 | 0.0014 |
| 34 | 48 | 0.78 | 0 |
| 34 | 60 | 0.15 | 6.6e-11 |
| 35 | 1 | -0.061 | 0.0074 |
| 35 | 2 | -0.31 | 2.8e-44 |
| 35 | 3 | -0.42 | 4.3e-82 |
| 35 | 4 | -0.24 | 8.4e-28 |
| 35 | 5 | -0.35 | 1.2e-57 |
| 35 | 6 | -0.31 | 2.7e-43 |
| 35 | 7 | -0.26 | 4.3e-32 |
| 35 | 9 | -0.26 | 4.1e-32 |
| 35 | 10 | -0.2 | 1,00E-18 |
| 35 | 11 | -0.29 | 5.9e-39 |
| 35 | 12 | 0.31 | 1.7e-45 |
| 35 | 13 | -0.59 | 6,00E-183 |
| 35 | 14 | -0.23 | 3.5e-25 |
| 35 | 15 | -0.055 | 0.016 |
| 35 | 16 | -0.14 | 1.8e-10 |
| 35 | 18 | 0.33 | 9,00E-51 |
| 35 | 19 | -0.091 | 6.2e-05 |
| 35 | 20 | 0.14 | 1.1e-09 |
| 35 | 21 | -0.32 | 1.2e-46 |
| 35 | 22 | -0.052 | 0.021 |
| 35 | 23 | -0.27 | 2.3e-34 |
| 35 | 24 | 0.0045 | 0.84 |
| 35 | 25 | 0.0081 | 0.72 |
| 35 | 26 | -0.31 | 9.1e-44 |
| 35 | 27 | -0.31 | 1.1e-44 |
| 35 | 28 | -0.2 | 2.4e-18 |
| 35 | 29 | -0.39 | 5.8e-73 |
| 35 | 30 | -0.38 | 8.5e-66 |
| 35 | 31 | -0.11 | 2.5e-06 |
| 35 | 32 | -0.3 | 1.3e-42 |
| 35 | 33 | 0.13 | 4.6e-09 |
| 35 | 34 | 0.12 | 7.4e-08 |
| 35 | 36 | 0.081 | 0.00036 |
| 35 | 37 | 0.29 | 1.7e-38 |
| 35 | 38 | -0.37 | 2.1e-64 |
| 35 | 39 | -0.018 | 0.44 |
| 35 | 40 | -0.058 | 0.011 |
| 35 | 41 | -0.31 | 2.7e-44 |
| 35 | 42 | -0.32 | 3.9e-47 |
| 35 | 43 | -0.39 | 8.5e-70 |
| 35 | 44 | -0.033 | 0.15 |
| 35 | 45 | 0.043 | 0.06 |
| 35 | 46 | 0.35 | 1.2e-56 |
| 35 | 47 | 0.58 | 1.7e-175 |
| 35 | 48 | 0.17 | 3.2e-14 |
| 35 | 49 | -0.39 | 1.5e-71 |
| 35 | 52 | 0.091 | 5.7e-05 |
| 35 | 54 | -0.16 | 1.4e-12 |
| 35 | 55 | -0.14 | 1.1e-09 |
| 35 | 56 | 0.25 | 5.3e-30 |
| 35 | 57 | 0.59 | 1,00E-181 |
| 35 | 58 | -0.45 | 1.4e-95 |
| 35 | 59 | -0.21 | 3,00E-21 |
| 35 | 60 | 0.01 | 0.66 |
| 35 | 61 | -0.095 | 2.8e-05 |
| 35 | 62 | 0.74 | 0 |
| 35 | 63 | -0.15 | 2.6e-11 |
| 36 | 1 | 0.012 | 0.6 |
| 36 | 2 | 0.011 | 0.63 |
| 36 | 3 | -0.046 | 0.044 |
| 36 | 4 | 0.0021 | 0.93 |
| 36 | 5 | -0.048 | 0.035 |
| 36 | 6 | -0.094 | 3.5e-05 |
| 36 | 7 | 0.082 | 0.00033 |
| 36 | 9 | 0.082 | 0.00033 |
| 36 | 10 | 0.057 | 0.013 |
| 36 | 11 | 0.071 | 0.0017 |
| 36 | 13 | 0.028 | 0.22 |
| 36 | 14 | 0.014 | 0.54 |
| 36 | 15 | 0.086 | 0.00016 |
| 36 | 16 | 0.1 | 5.8e-06 |
| 36 | 19 | 0.037 | 0.1 |
| 36 | 21 | -0.052 | 0.023 |
| 36 | 22 | 0.049 | 0.031 |
| 36 | 23 | 0.069 | 0.0025 |
| 36 | 24 | 0.3 | 1.6e-40 |
| 36 | 25 | 0.22 | 6.6e-23 |
| 36 | 26 | -0.051 | 0.024 |
| 36 | 27 | -0.0017 | 0.94 |
| 36 | 28 | -0.8 | 0 |
| 36 | 29 | -0.073 | 0.0012 |
| 36 | 30 | 0.066 | 0.0036 |
| 36 | 32 | -0.28 | 9.3e-36 |
| 36 | 34 | -0.091 | 6.5e-05 |
| 36 | 38 | -0.12 | 3.9e-07 |
| 36 | 39 | 0.049 | 0.03 |
| 36 | 40 | 0.023 | 0.32 |
| 36 | 41 | -0.14 | 1.8e-10 |
| 36 | 42 | -0.1 | 4.8e-06 |
| 36 | 43 | -0.18 | 2.8e-16 |
| 36 | 44 | 0.88 | 0 |
| 36 | 45 | -0.0044 | 0.85 |

| | | | |
|---|---|---|---|
| 50 | 25 | 0.021 | 0.34 |
| 50 | 26 | -0.36 | 6.6e-59 |
| 50 | 27 | -0.35 | 2.3e-56 |
| 50 | 28 | -0.11 | 1.5e-06 |
| 50 | 29 | -0.27 | 5.3e-33 |
| 50 | 30 | -0.25 | 1.1e-29 |
| 50 | 31 | -0.088 | 0.00011 |
| 50 | 32 | -0.21 | 2.6e-20 |
| 50 | 33 | 0.071 | 0.0017 |
| 50 | 34 | 0.095 | 2.6e-05 |
| 50 | 35 | 0.55 | 1.7e-155 |
| 50 | 36 | 0.047 | 0.037 |
| 50 | 37 | 0.22 | 8.1e-22 |
| 50 | 38 | -0.2 | 1.2e-18 |
| 50 | 39 | 0.082 | 0.00033 |
| 50 | 40 | -0.044 | 0.055 |
| 50 | 41 | -0.37 | 3.7e-62 |
| 50 | 42 | -0.12 | 1.4e-07 |
| 50 | 43 | -0.29 | 3.3e-39 |
| 50 | 44 | -0.033 | 0.14 |
| 50 | 45 | 0.026 | 0.26 |
| 50 | 46 | 0.14 | 1.4e-09 |
| 50 | 47 | 0.34 | 1.5e-52 |
| 50 | 48 | 0.17 | 1.7e-13 |
| 50 | 49 | -0.3 | 1.3e-40 |
| 50 | 51 | 0.59 | 1.5e-183 |
| 50 | 52 | 0.45 | 1.3e-97 |
| 50 | 53 | 0.74 | 0 |
| 50 | 54 | -0.72 | 6.0e-312 |
| 50 | 55 | -0.6 | 4.6e-188 |
| 50 | 56 | 0.1 | 9,00E-06 |
| 50 | 57 | 0.29 | 6.9e-39 |
| 50 | 58 | -0.39 | 3.5e-71 |
| 50 | 59 | -0.19 | 1,00E-16 |
| 50 | 60 | -0.00087 | 0.97 |
| 50 | 61 | -0.041 | 0.069 |
| 50 | 62 | 0.44 | 1.4e-90 |
| 50 | 63 | -0.11 | 1.9e-06 |
| 51 | 1 | -0.032 | 0.16 |
| 51 | 2 | -0.34 | 1.4e-53 |
| 51 | 3 | -0.43 | 9.7e-89 |
| 51 | 4 | -0.29 | 1.2e-38 |
| 51 | 5 | -0.38 | 1.4e-68 |
| 51 | 6 | -0.29 | 3.7e-39 |
| 51 | 7 | -0.26 | 5.4e-32 |
| 51 | 9 | -0.26 | 5.1e-32 |
| 51 | 10 | -0.29 | 4.4e-38 |
| 51 | 11 | -0.28 | 1.3e-35 |
| 51 | 12 | 0.32 | 8.3e-48 |
| 51 | 13 | -0.52 | 3.2e-135 |
| 51 | 14 | -0.23 | 1.8e-24 |
| 51 | 15 | -0.14 | 2.4e-10 |
| 51 | 16 | -0.12 | 5.7e-08 |
| 51 | 18 | 0.33 | 1.3e-50 |
| 51 | 19 | -0.19 | 2.2e-17 |
| 51 | 20 | 0.24 | 2,00E-27 |
| 51 | 21 | -0.34 | 1.1e-52 |
| 51 | 22 | -0.086 | 0.00017 |
| 51 | 23 | -0.14 | 6.6e-10 |
| 51 | 24 | -0.0089 | 0.7 |
| 51 | 25 | 0.025 | 0.27 |
| 51 | 26 | -0.22 | 6.1e-23 |
| 51 | 27 | -0.38 | 1.9e-68 |
| 51 | 28 | -0.15 | 3.6e-11 |
| 51 | 29 | -0.24 | 3.8e-27 |
| 51 | 30 | -0.34 | 1.4e-54 |
| 51 | 31 | -0.15 | 1,00E-10 |
| 51 | 32 | -0.31 | 6.4e-44 |
| 51 | 33 | 0.17 | 1.4e-13 |
| 51 | 34 | 0.053 | 0.019 |
| 51 | 35 | 0.61 | 1.3e-194 |
| 51 | 36 | 0.046 | 0.043 |
| 51 | 37 | 0.27 | 1.3e-34 |
| 51 | 38 | -0.28 | 3.1e-37 |
| 51 | 39 | -0.00044 | 0.98 |
| 51 | 40 | -0.032 | 0.16 |
| 51 | 41 | -0.28 | 5.1e-36 |
| 51 | 42 | -0.12 | 1.4e-07 |
| 51 | 43 | -0.3 | 1.7e-42 |
| 51 | 44 | -0.048 | 0.035 |
| 51 | 45 | 0.042 | 0.062 |
| 51 | 46 | 0.14 | 1.8e-09 |
| 51 | 47 | 0.4 | 1.2e-75 |
| 51 | 48 | 0.18 | 2.3e-15 |
| 51 | 49 | -0.28 | 2.3e-35 |
| 51 | 52 | -0.0044 | 0.85 |
| 51 | 53 | 0.86 | 0 |
| 51 | 54 | 0.13 | 1,00E-08 |
| 51 | 55 | 0.15 | 1.2e-11 |
| 51 | 56 | 0.081 | 0.00037 |
| 51 | 57 | 0.37 | 3.4e-64 |
| 51 | 58 | -0.47 | 3.3e-105 |
| 51 | 59 | -0.22 | 2.5e-22 |
| 51 | 60 | -0.036 | 0.11 |
| 51 | 61 | -0.075 | 0.00094 |
| 51 | 62 | 0.52 | 1.4e-137 |
| 51 | 63 | -0.085 | 0.00017 |
| 52 | 2 | -0.026 | 0.26 |
| 52 | 3 | -0.037 | 0.1 |
| 52 | 4 | 0.067 | 0.0033 |
| 52 | 5 | 0.045 | 0.049 |
| 52 | 19 | 0.033 | 0.15 |
| 52 | 21 | 0.024 | 0.29 |
| 52 | 22 | 0.014 | 0.55 |
| 52 | 27 | 0.044 | 0.052 |

| | | | |
|---|---|---|---|
| 59 | 48 | -0.36 | 9.4e-60 |
| 59 | 49 | -0.068 | 0.0029 |
| 59 | 52 | -0.025 | 0.26 |
| 59 | 54 | 0.043 | 0.06 |
| 59 | 55 | 0.036 | 0.11 |
| 59 | 58 | 0.22 | 9.3e-23 |
| 59 | 60 | 0.019 | 0.41 |
| 59 | 63 | 0.76 | 0 |
| 60 | 38 | 0.13 | 1.9e-08 |
| 60 | 39 | 0.071 | 0.0017 |
| 60 | 40 | 0.051 | 0.025 |
| 61 | 1 | -0.02 | 0.39 |
| 61 | 2 | 0.067 | 0.0032 |
| 61 | 3 | 0.035 | 0.12 |
| 61 | 4 | 0.077 | 0.00071 |
| 61 | 5 | 0.048 | 0.033 |
| 61 | 6 | -0.042 | 0.065 |
| 61 | 7 | 0.034 | 0.14 |
| 61 | 9 | 0.033 | 0.14 |
| 61 | 10 | -0.092 | 4.6e-05 |
| 61 | 11 | -0.075 | 0.001 |
| 61 | 13 | 0.023 | 0.3 |
| 61 | 14 | -0.097 | 2.1e-05 |
| 61 | 15 | -0.09 | 7.3e-05 |
| 61 | 16 | 0.062 | 0.0064 |
| 61 | 19 | 0.013 | 0.58 |
| 61 | 21 | 0.026 | 0.25 |
| 61 | 22 | 0.0059 | 0.79 |
| 61 | 23 | -0.034 | 0.13 |
| 61 | 24 | -0.01 | 0.66 |
| 61 | 25 | -0.0053 | 0.82 |
| 61 | 26 | -0.011 | 0.61 |
| 61 | 27 | 0.03 | 0.19 |
| 61 | 28 | 0.0058 | 0.8 |
| 61 | 29 | -0.027 | 0.24 |
| 61 | 30 | 0.0052 | 0.82 |
| 61 | 31 | -0.029 | 0.21 |
| 61 | 32 | -0.02 | 0.37 |
| 61 | 34 | 0.0076 | 0.74 |
| 61 | 36 | -0.034 | 0.13 |
| 61 | 38 | -0.025 | 0.27 |
| 61 | 39 | 0.009 | 0.69 |
| 61 | 40 | -0.11 | 1.6e-06 |
| 61 | 41 | 0.00063 | 0.98 |
| 61 | 42 | -0.13 | 8.7e-09 |
| 61 | 43 | -0.031 | 0.17 |
| 61 | 44 | -0.035 | 0.12 |
| 61 | 45 | -0.65 | 3.6e-232 |
| 61 | 46 | -0.42 | 2.5e-83 |
| 61 | 48 | -0.1 | 4.4e-06 |
| 61 | 49 | -0.065 | 0.0045 |
| 61 | 52 | -0.04 | 0.076 |
| 61 | 54 | -0.014 | 0.55 |
| 61 | 55 | -0.041 | 0.074 |
| 61 | 56 | 0.46 | 6,00E-104 |
| 61 | 58 | 0.11 | 2.8e-06 |
| 61 | 59 | 0.51 | 5.8e-130 |
| 61 | 60 | 0.25 | 5.2e-29 |
| 61 | 63 | 0.38 | 6.3e-68 |
| 62 | 1 | -0.037 | 0.1 |
| 62 | 2 | -0.22 | 1.7e-23 |
| 62 | 3 | -0.34 | 3.7e-54 |
| 62 | 4 | -0.19 | 2,00E-16 |
| 62 | 5 | -0.3 | 1.8e-40 |
| 62 | 6 | -0.3 | 8.8e-42 |
| 62 | 7 | -0.33 | 6.6e-51 |
| 62 | 9 | -0.33 | 6.2e-51 |
| 62 | 10 | -0.2 | 6.1e-19 |
| 62 | 11 | -0.45 | 3,00E-96 |
| 62 | 12 | 0.25 | 6.5e-29 |
| 62 | 13 | -0.55 | 2.4e-150 |
| 62 | 14 | -0.18 | 5.6e-15 |
| 62 | 15 | 0.018 | 0.43 |
| 62 | 16 | -0.16 | 3.1e-13 |
| 62 | 18 | 0.33 | 2.4e-51 |
| 62 | 19 | -0.033 | 0.14 |
| 62 | 20 | 0.045 | 0.048 |
| 62 | 21 | -0.29 | 7.8e-39 |
| 62 | 22 | -0.015 | 0.5 |
| 62 | 23 | -0.25 | 6,00E-28 |
| 62 | 24 | -0.024 | 0.29 |
| 62 | 25 | -0.024 | 0.3 |
| 62 | 26 | -0.26 | 1.1e-30 |
| 62 | 27 | -0.25 | 2.7e-28 |
| 62 | 28 | -0.19 | 2.7e-17 |
| 62 | 29 | -0.34 | 6.5e-55 |
| 62 | 30 | -0.35 | 1.2e-57 |
| 62 | 31 | -0.089 | 9.6e-05 |
| 62 | 32 | -0.27 | 1.6e-32 |
| 62 | 33 | 0.021 | 0.35 |
| 62 | 34 | -0.13 | 1.7e-08 |
| 62 | 36 | 0.072 | 0.0015 |
| 62 | 37 | 0.26 | 2.4e-30 |
| 62 | 38 | -0.4 | 4.9e-76 |
| 62 | 39 | -0.14 | 7.4e-10 |
| 62 | 40 | -0.013 | 0.55 |
| 62 | 41 | -0.21 | 1.3e-21 |
| 62 | 42 | 0.1 | 7.4e-06 |
| 62 | 43 | -0.3 | 1.8e-41 |
| 62 | 44 | -0.0023 | 0.92 |
| 62 | 45 | 0.13 | 2.8e-09 |
| 62 | 46 | 0.054 | 0.018 |
| 62 | 48 | 0.093 | 4.6e-05 |
| 62 | 49 | -0.21 | 9.5e-21 |
| 62 | 52 | 0.04 | 0.077 |

| | | | |
|---|---|---|---|
| 12 | 36 | 0.061 | 0.0074 |
| 12 | 38 | 0.073 | 0.0014 |
| 12 | 39 | 0.1 | 7.7e-06 |
| 12 | 40 | 0.056 | 0.015 |
| 12 | 41 | -0.19 | 5.2e-17 |
| 12 | 42 | -0.036 | 0.11 |
| 12 | 43 | -0.023 | 0.32 |
| 12 | 44 | 0.015 | 0.52 |
| 12 | 45 | 0.12 | 1.6e-07 |
| 12 | 46 | 0.25 | 5.9e-30 |
| 12 | 48 | 0.12 | 5.9e-08 |
| 12 | 49 | -0.22 | 2.5e-23 |
| 12 | 52 | 0.057 | 0.012 |
| 12 | 54 | -0.22 | 1.2e-22 |
| 12 | 55 | -0.23 | 9.8e-25 |
| 12 | 56 | 0.037 | 0.1 |
| 12 | 58 | -0.26 | 2.7e-31 |
| 12 | 59 | -0.12 | 1.6e-07 |
| 12 | 60 | 0.17 | 2.2e-13 |
| 12 | 61 | -0.11 | 9.2e-07 |
| 12 | 63 | -0.089 | 8.8e-05 |
| 13 | 1 | 0.06 | 0.0083 |
| 13 | 2 | 0.25 | 2.4e-28 |
| 13 | 3 | 0.37 | 4.2e-64 |
| 13 | 4 | 0.2 | 1.1e-19 |
| 13 | 5 | 0.32 | 3,00E-47 |
| 13 | 6 | 0.32 | 1.8e-47 |
| 13 | 10 | 0.37 | 6,00E-64 |
| 13 | 14 | 0.42 | 2.3e-84 |
| 13 | 15 | 0.26 | 1,00E-30 |
| 13 | 19 | 0.088 | 1,00E-04 |
| 13 | 21 | 0.22 | 1.6e-22 |
| 13 | 22 | 0.03 | 0.19 |
| 13 | 23 | 0.3 | 2.1e-42 |
| 13 | 26 | 0.31 | 6.8e-45 |
| 13 | 27 | 0.26 | 9.1e-31 |
| 13 | 29 | 0.41 | 2.2e-77 |
| 13 | 30 | 0.41 | 5.1e-80 |
| 13 | 34 | -0.019 | 0.4 |
| 13 | 38 | 0.3 | 1.3e-40 |
| 13 | 39 | 0.015 | 0.5 |
| 13 | 40 | -0.0027 | 0.9 |
| 13 | 41 | 0.22 | 4.4e-22 |
| 13 | 42 | 0.11 | 3.6e-06 |
| 13 | 45 | 0.03 | 0.19 |
| 13 | 46 | -0.028 | 0.22 |
| 13 | 48 | -0.085 | 0.00017 |
| 13 | 49 | 0.46 | 1.9e-103 |
| 13 | 52 | -0.045 | 0.046 |
| 13 | 54 | 0.095 | 2.9e-05 |
| 13 | 55 | 0.087 | 0.00013 |
| 13 | 58 | 0.56 | 2,00E-159 |
| 13 | 60 | 0.06 | 0.0078 |
| 14 | 2 | 0.014 | 0.54 |
| 14 | 3 | 0.099 | 1.2e-05 |
| 14 | 4 | 0.0086 | 0.7 |
| 14 | 5 | 0.079 | 0.00047 |
| 14 | 19 | 0.055 | 0.016 |
| 14 | 21 | 0.095 | 2.9e-05 |
| 14 | 22 | 0.035 | 0.13 |
| 14 | 27 | 0.14 | 1,00E-09 |
| 14 | 34 | -0.0033 | 0.88 |
| 14 | 38 | 0.11 | 1.1e-06 |
| 14 | 39 | -0.072 | 0.0016 |
| 14 | 40 | 0.083 | 0.00025 |
| 14 | 41 | 0.19 | 5.1e-17 |
| 14 | 42 | 0.084 | 0.00022 |
| 14 | 45 | 0.14 | 1.5e-09 |
| 14 | 46 | 0.12 | 6.8e-08 |
| 14 | 48 | 0.045 | 0.047 |
| 14 | 49 | 0.32 | 2.1e-48 |
| 14 | 52 | 0.029 | 0.2 |
| 14 | 60 | 0.1 | 3.9e-06 |
| 15 | 2 | 0.023 | 0.31 |
| 15 | 3 | 0.069 | 0.0023 |
| 15 | 4 | -0.018 | 0.42 |
| 15 | 5 | 0.027 | 0.23 |
| 15 | 14 | 0.34 | 3.1e-53 |
| 15 | 19 | 0.051 | 0.024 |
| 15 | 21 | -0.062 | 0.0062 |
| 15 | 22 | 0.03 | 0.19 |
| 15 | 27 | 0.048 | 0.036 |
| 15 | 34 | -0.12 | 1.3e-07 |
| 15 | 38 | 0.073 | 0.0013 |
| 15 | 39 | 4,00E-04 | 0.99 |

| | | | |
|---|---|---|---|
| 24 | 54 | -0.078 | 6,00E-04 |
| 24 | 55 | -0.1 | 7.9e-06 |
| 24 | 58 | 0.088 | 1,00E-04 |
| 24 | 60 | -0.033 | 0.15 |
| 25 | 1 | -0.026 | 0.26 |
| 25 | 2 | -0.028 | 0.22 |
| 25 | 3 | -0.024 | 0.28 |
| 25 | 4 | -0.0034 | 0.88 |
| 25 | 5 | -0.0044 | 0.85 |
| 25 | 6 | -0.00093 | 0.97 |
| 25 | 10 | 0.13 | 2.9e-08 |
| 25 | 11 | 0.15 | 5.9e-11 |
| 25 | 13 | 0.11 | 1.1e-06 |
| 25 | 14 | -0.0081 | 0.72 |
| 25 | 15 | 0.061 | 0.0068 |
| 25 | 19 | 0.078 | 0.00063 |
| 25 | 21 | -0.069 | 0.0025 |
| 25 | 22 | -0.019 | 0.4 |
| 25 | 23 | 0.092 | 4.9e-05 |
| 25 | 26 | 0.3 | 8.6e-43 |
| 25 | 27 | 0.099 | 1.3e-05 |
| 25 | 28 | 0.1 | 8.1e-06 |
| 25 | 29 | -0.067 | 0.003 |
| 25 | 30 | 0.41 | 2.8e-80 |
| 25 | 32 | -0.2 | 5.1e-19 |
| 25 | 34 | 0.006 | 0.79 |
| 25 | 38 | -0.079 | 0.00047 |
| 25 | 39 | 0.12 | 2.4e-07 |
| 25 | 40 | -0.024 | 0.3 |
| 25 | 41 | -0.4 | 3.5e-77 |
| 25 | 42 | -0.029 | 0.2 |
| 25 | 43 | -0.22 | 1.6e-23 |
| 25 | 45 | 0.014 | 0.53 |
| 25 | 46 | 0.032 | 0.17 |
| 25 | 48 | 0.032 | 0.16 |
| 25 | 49 | 0.091 | 5.6e-05 |
| 25 | 52 | -0.054 | 0.017 |
| 25 | 54 | -0.0049 | 0.83 |
| 25 | 55 | -0.038 | 0.09 |
| 25 | 58 | 0.076 | 0.00086 |
| 25 | 60 | -0.032 | 0.16 |
| 26 | 1 | 0.031 | 0.17 |
| 26 | 2 | 0.086 | 0.00014 |
| 26 | 3 | 0.15 | 9.6e-11 |
| 26 | 4 | 0.077 | 0.00072 |
| 26 | 5 | 0.13 | 1.6e-08 |
| 26 | 10 | 0.17 | 3.9e-14 |
| 26 | 14 | 0.18 | 4.9e-16 |
| 26 | 15 | 0.074 | 0.001 |
| 26 | 19 | -0.077 | 0.00064 |
| 26 | 21 | 0.082 | 0.00029 |
| 26 | 22 | -0.37 | 1.6e-64 |
| 26 | 27 | 0.2 | 8.8e-19 |
| 26 | 34 | -0.098 | 1.4e-05 |
| 26 | 38 | 0.096 | 2.4e-05 |
| 26 | 39 | -0.0044 | 0.85 |
| 26 | 40 | -0.055 | 0.016 |
| 26 | 41 | 0.33 | 4.2e-51 |
| 26 | 42 | 0.013 | 0.58 |
| 26 | 45 | 0.021 | 0.36 |
| 26 | 46 | -0.0018 | 0.94 |
| 26 | 48 | -0.12 | 1.3e-07 |
| 26 | 49 | 0.27 | 5.6e-34 |
| 26 | 52 | -0.089 | 8.9e-05 |
| 26 | 60 | -0.041 | 0.074 |
| 27 | 2 | 0.13 | 1,00E-08 |
| 27 | 3 | 0.19 | 1.9e-16 |
| 27 | 4 | 0.14 | 2.4e-09 |
| 27 | 5 | 0.19 | 7.5e-18 |
| 27 | 21 | 0.31 | 4.1e-43 |
| 27 | 34 | -0.0083 | 0.72 |
| 27 | 38 | 0.059 | 0.01 |
| 27 | 39 | -0.043 | 0.059 |
| 27 | 40 | 0.12 | 4.9e-08 |
| 27 | 41 | 0.54 | 8.3e-148 |
| 27 | 42 | 0.11 | 2.2e-06 |
| 27 | 48 | -0.12 | 5.9e-08 |
| 27 | 60 | -0.0015 | 0.95 |
| 28 | 1 | 0.015 | 0.51 |
| 28 | 2 | -0.007 | 0.76 |
| 28 | 3 | 0.077 | 7,00E-04 |
| 28 | 4 | -0.0084 | 0.71 |
| 28 | 5 | 0.065 | 0.0041 |
| 28 | 6 | 0.15 | 2.4e-11 |
| 28 | 10 | 0.053 | 0.02 |

| | | | |
|---|---|---|---|
| 36 | 46 | 0.098 | 1.6e-05 |
| 36 | 48 | 0.0019 | 0.93 |
| 36 | 49 | 0.17 | 2.1e-14 |
| 36 | 52 | -0.044 | 0.054 |
| 36 | 54 | -0.019 | 0.4 |
| 36 | 55 | -0.049 | 0.031 |
| 36 | 58 | 0.019 | 0.41 |
| 36 | 60 | 5,00E-04 | 0.98 |
| 37 | 1 | 0.077 | 0.00072 |
| 37 | 2 | -0.14 | 8.8e-10 |
| 37 | 3 | -0.25 | 7.9e-29 |
| 37 | 4 | -0.17 | 2.6e-13 |
| 37 | 5 | -0.27 | 1.9e-33 |
| 37 | 6 | -0.24 | 7.9e-27 |
| 37 | 7 | -0.14 | 2.3e-10 |
| 37 | 9 | -0.14 | 2.2e-10 |
| 37 | 10 | -0.14 | 2.5e-09 |
| 37 | 11 | -0.14 | 3.6e-10 |
| 37 | 12 | 0.28 | 1.9e-35 |
| 37 | 13 | -0.22 | 1.2e-23 |
| 37 | 14 | -0.07 | 0.002 |
| 37 | 15 | -0.044 | 0.054 |
| 37 | 16 | -0.063 | 0.0058 |
| 37 | 18 | 0.28 | 4.7e-35 |
| 37 | 19 | -0.1 | 4.1e-06 |
| 37 | 21 | -0.18 | 3.7e-15 |
| 37 | 22 | 0.04 | 0.082 |
| 37 | 23 | -0.00045 | 0.98 |
| 37 | 24 | -0.018 | 0.44 |
| 37 | 25 | -0.026 | 0.26 |
| 37 | 26 | -0.16 | 5.7e-12 |
| 37 | 27 | -0.33 | 9.3e-52 |
| 37 | 28 | -0.12 | 2.2e-07 |
| 37 | 29 | -0.43 | 1.7e-88 |
| 37 | 30 | -0.65 | 5.1e-236 |
| 37 | 31 | -0.05 | 0.028 |
| 37 | 32 | -0.17 | 1.2e-13 |
| 37 | 34 | -0.092 | 4.6e-05 |
| 37 | 36 | 0.25 | 5.9e-29 |
| 37 | 38 | -0.12 | 2.2e-07 |
| 37 | 39 | -0.016 | 0.49 |
| 37 | 40 | -0.11 | 1.2e-06 |
| 37 | 41 | -0.086 | 0.00015 |
| 37 | 42 | -0.16 | 5.9e-12 |
| 37 | 43 | -0.12 | 3.7e-07 |
| 37 | 44 | 0.048 | 0.036 |
| 37 | 45 | 0.046 | 0.041 |
| 37 | 46 | 0.22 | 6.7e-22 |
| 37 | 48 | 0.032 | 0.16 |
| 37 | 49 | -0.027 | 0.23 |
| 37 | 52 | -0.054 | 0.018 |
| 37 | 54 | -0.03 | 0.18 |
| 37 | 55 | -0.069 | 0.0024 |
| 37 | 56 | 0.16 | 1.5e-12 |
| 37 | 58 | -0.25 | 5.1e-28 |
| 37 | 59 | -0.021 | 0.36 |
| 37 | 60 | 0.035 | 0.12 |
| 37 | 61 | -0.049 | 0.032 |
| 37 | 63 | 0.11 | 1.9e-06 |
| 38 | 39 | 0.56 | 2.6e-163 |
| 40 | 38 | 0.078 | 0.00064 |
| 40 | 39 | 0.11 | 1,00E-06 |
| 41 | 2 | 0.11 | 1.7e-06 |
| 41 | 3 | 0.19 | 1.7e-16 |
| 41 | 4 | 0.079 | 0.00049 |
| 41 | 5 | 0.15 | 2.3e-11 |
| 41 | 21 | 0.18 | 2.2e-15 |
| 41 | 34 | -0.077 | 0.00068 |
| 41 | 38 | 0.17 | 7.7e-14 |
| 41 | 39 | -0.1 | 5.2e-06 |
| 41 | 40 | 0.0017 | 0.94 |
| 41 | 42 | 0.086 | 0.00014 |
| 41 | 48 | -0.14 | 6,00E-10 |
| 41 | 60 | 0.04 | 0.081 |
| 42 | 34 | 0.52 | 7.1e-133 |
| 42 | 38 | 0.056 | 0.014 |
| 42 | 39 | -0.064 | 0.0048 |
| 42 | 40 | 0.12 | 7,00E-08 |
| 42 | 48 | 0.54 | 3.7e-144 |
| 42 | 60 | 0.12 | 1.3e-07 |
| 43 | 1 | 0.036 | 0.11 |
| 43 | 2 | 0.089 | 8.2e-05 |
| 43 | 3 | 0.16 | 3.7e-12 |
| 43 | 4 | 0.065 | 0.004 |
| 43 | 5 | 0.12 | 3.9e-08 |

| | | | |
|---|---|---|---|
| 52 | 34 | 0.08 | 0.00043 |
| 52 | 38 | 0.011 | 0.62 |
| 52 | 39 | 0.038 | 0.095 |
| 52 | 40 | 0.068 | 0.0028 |
| 52 | 41 | -0.029 | 0.2 |
| 52 | 42 | -0.0028 | 0.9 |
| 52 | 45 | 0.041 | 0.069 |
| 52 | 46 | 0.07 | 0.0019 |
| 52 | 48 | 0.022 | 0.33 |
| 52 | 60 | 0.047 | 0.039 |
| 53 | 1 | -0.097 | 1.9e-05 |
| 53 | 2 | -0.31 | 1.2e-43 |
| 53 | 3 | -0.39 | 2.2e-72 |
| 53 | 4 | -0.22 | 4.6e-22 |
| 53 | 5 | -0.31 | 4.1e-44 |
| 53 | 6 | -0.27 | 8.2e-33 |
| 53 | 7 | -0.25 | 1.7e-29 |
| 53 | 9 | -0.25 | 1.7e-29 |
| 53 | 10 | -0.27 | 2.7e-33 |
| 53 | 11 | -0.22 | 5,00E-23 |
| 53 | 12 | 0.31 | 1.4e-43 |
| 53 | 13 | -0.47 | 4.2e-109 |
| 53 | 14 | -0.18 | 3.5e-16 |
| 53 | 15 | -0.13 | 2.6e-08 |
| 53 | 16 | -0.12 | 3,00E-07 |
| 53 | 18 | 0.28 | 5.1e-37 |
| 53 | 19 | -0.15 | 4.3e-11 |
| 53 | 20 | 0.25 | 3.6e-28 |
| 53 | 21 | -0.28 | 3.5e-36 |
| 53 | 22 | -0.067 | 0.0031 |
| 53 | 23 | -0.16 | 6.8e-13 |
| 53 | 24 | -0.014 | 0.54 |
| 53 | 25 | -0.0056 | 0.8 |
| 53 | 26 | -0.24 | 4.8e-26 |
| 53 | 27 | -0.31 | 4.1e-44 |
| 53 | 28 | -0.12 | 2.7e-07 |
| 53 | 29 | -0.24 | 5.1e-27 |
| 53 | 30 | -0.29 | 1.2e-38 |
| 53 | 31 | -0.13 | 1.7e-08 |
| 53 | 32 | -0.24 | 3.8e-27 |
| 53 | 33 | 0.14 | 1.1e-09 |
| 53 | 34 | 0.087 | 0.00014 |
| 53 | 35 | 0.57 | 5.5e-168 |
| 53 | 36 | 0.018 | 0.43 |
| 53 | 37 | 0.21 | 1,00E-20 |
| 53 | 38 | -0.24 | 7.6e-27 |
| 53 | 39 | 0.019 | 0.41 |
| 53 | 40 | 0.0062 | 0.78 |
| 53 | 41 | -0.26 | 1.8e-30 |
| 53 | 42 | -0.1 | 3.9e-06 |
| 53 | 43 | -0.28 | 5.1e-35 |
| 53 | 44 | -0.066 | 0.0039 |
| 53 | 45 | 0.057 | 0.011 |
| 53 | 46 | 0.15 | 1.2e-11 |
| 53 | 47 | 0.35 | 3.1e-58 |
| 53 | 48 | 0.17 | 1.9e-13 |
| 53 | 49 | -0.27 | 9.7e-33 |
| 53 | 52 | 0.5 | 8.7e-123 |
| 53 | 54 | -0.17 | 7.3e-14 |
| 53 | 55 | 0.097 | 2.1e-05 |
| 53 | 56 | 0.069 | 0.0022 |
| 53 | 57 | 0.34 | 3.2e-54 |
| 53 | 58 | -0.38 | 1,00E-66 |
| 53 | 59 | -0.2 | 2.9e-19 |
| 53 | 60 | -0.0079 | 0.73 |
| 53 | 61 | -0.085 | 0.00017 |
| 53 | 62 | 0.47 | 1.6e-109 |
| 53 | 63 | -0.12 | 2.5e-07 |
| 54 | 1 | 0.062 | 0.0063 |
| 54 | 2 | -0.064 | 0.0046 |
| 54 | 3 | 0.0048 | 0.83 |
| 54 | 4 | -0.095 | 2.8e-05 |
| 54 | 5 | -0.028 | 0.22 |
| 54 | 10 | 0.051 | 0.025 |
| 54 | 14 | 0.081 | 0.00034 |
| 54 | 15 | 0.042 | 0.067 |
| 54 | 19 | -0.0054 | 0.81 |
| 54 | 21 | 0.11 | 9.7e-07 |
| 54 | 22 | -0.0045 | 0.84 |
| 54 | 23 | 0.18 | 2.8e-16 |
| 54 | 26 | 0.25 | 1.7e-28 |
| 54 | 27 | 0.1 | 9.4e-06 |
| 54 | 34 | -0.071 | 0.0017 |
| 54 | 38 | 0.00046 | 0.98 |
| 54 | 39 | -0.1 | 9.1e-06 |

| | | | |
|---|---|---|---|
| 62 | 54 | -0.086 | 0.00016 |
| 62 | 55 | -0.079 | 0.00051 |
| 62 | 56 | -0.18 | 2.2e-15 |
| 62 | 58 | -0.58 | 4.3e-171 |
| 62 | 59 | -0.16 | 2,00E-12 |
| 62 | 60 | 0.0055 | 0.81 |
| 62 | 61 | -0.22 | 6.4e-22 |
| 62 | 63 | 0.084 | 0.00021 |
| 63 | 1 | 0.01 | 0.65 |
| 63 | 2 | 0.098 | 1.4e-05 |
| 63 | 3 | 0.06 | 0.0078 |
| 63 | 4 | 0.089 | 9.5e-05 |
| 63 | 5 | 0.057 | 0.012 |
| 63 | 6 | -0.048 | 0.036 |
| 63 | 7 | -0.054 | 0.017 |
| 63 | 9 | -0.054 | 0.017 |
| 63 | 10 | -0.057 | 0.012 |
| 63 | 11 | -0.2 | 1.2e-19 |
| 63 | 13 | 0.078 | 0.00059 |
| 63 | 14 | 0.054 | 0.018 |
| 63 | 15 | 0.067 | 0.0031 |
| 63 | 16 | 0.047 | 0.037 |
| 63 | 19 | -0.0068 | 0.76 |
| 63 | 21 | -0.027 | 0.23 |
| 63 | 22 | -0.023 | 0.31 |
| 63 | 23 | 0.057 | 0.013 |
| 63 | 24 | -0.028 | 0.21 |
| 63 | 25 | 0.025 | 0.28 |
| 63 | 26 | 0.12 | 3.7e-07 |
| 63 | 27 | -0.012 | 0.59 |
| 63 | 28 | -0.057 | 0.012 |
| 63 | 29 | 0.018 | 0.42 |
| 63 | 30 | -0.053 | 0.02 |
| 63 | 31 | 0.02 | 0.39 |
| 63 | 32 | -0.045 | 0.046 |
| 63 | 34 | -0.41 | 1.1e-77 |
| 63 | 36 | 0.08 | 0.00044 |
| 63 | 38 | 0.0063 | 0.78 |
| 63 | 39 | -0.073 | 0.0012 |
| 63 | 40 | -0.1 | 4.7e-06 |
| 63 | 41 | 0.066 | 0.0037 |
| 63 | 42 | -0.086 | 0.00015 |
| 63 | 43 | 0.078 | 0.00055 |
| 63 | 44 | 0.085 | 0.00017 |
| 63 | 45 | 0.001 | 0.96 |
| 63 | 46 | 0.016 | 0.47 |
| 63 | 48 | -0.19 | 1.4e-16 |
| 63 | 49 | 0.27 | 8.4e-33 |
| 63 | 52 | -0.086 | 0.00016 |
| 63 | 54 | 0.06 | 0.0086 |
| 63 | 55 | 0.02 | 0.37 |
| 63 | 58 | -0.17 | 1.8e-13 |
| 63 | 60 | -0.12 | 4.5e-08 |

**Table S9.** Log2(foldchange) of the proteome pairwise comparisons and Δ*dksA* , Δ*relA* Δ*spoT* and Δ*dksA* Δ*relA* Δ*spoT* transcriptomes. Proteins were mapped on the Y11 reference genome. P-value and adjusted p-value can be found in Table S6. 0 corresponds to proteins not detected in any conditions of the comparison. +15 and -15 corresponds to proteins detected only in the first or second condition of the comparison, respectively.

| Locus | Old Locus | Gene | Product | Log2(foldchange) Ye.1 vs Ye.2 | Ye.1 vs Ye.3 | Ye.1 vs Ye.4 | Ye.2 vs Ye.3 | Ye.2 vs Ye.4 | Ye.3 vs Ye.4 | ΔrelA ΔspoT vs WT | ΔdksA vs WT | ΔdksA ΔrelA ΔspoT vs WT |
|---|---|---|---|---|---|---|---|---|---|---|---|---|
| **Ribosomal and ribosome-associated proteins** | | | | | | | | | | | | |
| Y11_RS00715 | Y11_01471 | *prmB* | 50S ribosomal protein L3 N(5)-glutamine methyltransferase | 0,22 | 0,33 | 0,80 | 0,12 | 0,58 | 0,47 | 0,00 | -0,80 | -0,86 |
| Y11_RS01390 | Y11_02981 | *rplY* | 50S ribosomal protein L25 | 0,29 | -0,47 | -0,65 | -0,77 | -0,94 | -0,18 | 0,91 | 1,02 | 0,88 |
| Y11_RS01925 | Y11_04141 | *ycaO* | 30S ribosomal protein S12 methylthiotransferase accessory protein YcaO | -0,15 | -2,56 | -2,45 | -2,41 | -2,29 | 0,12 | 2,10 | 1,65 | 1,74 |
| Y11_RS01955 | Y11_04211 | *rpsA* | 30S ribosomal protein S1 | 0,10 | -0,15 | -0,55 | -0,25 | -0,65 | -0,40 | 1,56 | 0,97 | 0,89 |
| Y11_RS02435 | Y11_05241 | *rpmF* | 50S ribosomal protein L32 | 0,10 | -0,09 | 0,06 | -0,20 | -0,04 | 0,16 | 1,43 | 0,74 | 0,86 |
| Y11_RS03615 | Y11_07871 | *rpmI* | 50S ribosomal protein L35 | 0,54 | -0,03 | 0,11 | -0,56 | -0,43 | 0,13 | 0,94 | 0,64 | 0,25 |
| Y11_RS03620 | Y11_07881 | *rplT* | 50S ribosomal protein L20 | 0,35 | -0,90 | -0,91 | -1,25 | -1,26 | -0,02 | 1,19 | 0,99 | 0,52 |
| Y11_RS05735 | Y11_12411 | *rimJ* | ribosomal protein S5-alanine N-acetyltransferase | -0,33 | 0,30 | 0,96 | 0,63 | 1,29 | 0,66 | -0,21 | -0,73 | -0,44 |
| Y11_RS08145 | Y11_17361 | | ABC-F family ATPase | -0,07 | -1,52 | -2,81 | -1,45 | -2,74 | -1,29 | 2,02 | 0,76 | 0,81 |
| Y11_RS09065 | Y11_19221 | *rsfS* | ribosome silencing factor | 0,02 | -1,16 | -1,73 | -1,18 | -1,75 | -0,56 | 1,59 | 0,67 | 1,21 |
| Y11_RS09270 | Y11_19661 | *ybcJ* | ribosome-associated protein YbcJ | -0,19 | -0,14 | -0,66 | 0,05 | -0,47 | -0,52 | 2,01 | 0,52 | 1,45 |
| Y11_RS09530 | Y11_20251 | | type B 50S ribosomal protein L31 | 0,00 | 0,00 | 0,00 | 0,00 | 0,00 | 0,00 | 0,76 | 0,08 | 0,17 |
| Y11_RS09535 | Y11_20261 | *ykgO* | type B 50S ribosomal protein L36 | 0,00 | 0,00 | 0,00 | 0,00 | 0,00 | 0,00 | -0,90 | -0,67 | -1,85 |
| Y11_RS10310 | Y11_21901 | *raiA* | ribosome-associated translation inhibitor RaiA | -0,56 | 1,66 | 3,19 | 2,23 | 3,75 | 1,52 | -1,92 | -2,13 | -1,88 |
| Y11_RS10375 | Y11_22041 | *rplS* | 50S ribosomal protein L19 | 0,25 | -0,56 | -0,93 | -0,81 | -1,18 | -0,37 | 1,70 | 0,80 | 0,59 |
| Y11_RS10385 | Y11_22061 | *rimM* | ribosome maturation factor RimM | -0,35 | -1,80 | -2,75 | -1,46 | -2,40 | -0,95 | 2,40 | 1,18 | 1,47 |
| Y11_RS10390 | Y11_22071 | *rpsP* | 30S ribosomal protein S16 | -0,07 | 0,03 | -0,44 | 0,09 | -0,37 | -0,47 | 2,14 | 1,10 | 1,26 |
| Y11_RS11820 | Y11_25071 | *rpsU* | 30S ribosomal protein S21 | 0,42 | -0,27 | -0,51 | -0,69 | -0,93 | -0,24 | 1,37 | 1,20 | 1,07 |
| Y11_RS12400 | Y11_26311 | *rpsI* | 30S ribosomal protein S9 | 0,10 | -0,44 | -0,99 | -0,55 | -1,09 | -0,55 | 1,25 | 1,40 | 1,00 |
| Y11_RS12405 | Y11_26321 | *rplM* | 50S ribosomal protein L13 | 0,28 | -0,49 | -0,91 | -0,77 | -1,19 | -0,42 | 1,84 | 1,67 | 1,64 |
| Y11_RS12500 | Y11_26511 | *hpf* | ribosome hibernation promoting factor | 0,63 | 1,92 | 3,07 | 1,29 | 2,44 | 1,15 | -1,97 | -1,08 | -1,13 |
| Y11_RS12655 | Y11_26861 | | ribosome-associated protein | 0,48 | 0,83 | 1,03 | 0,35 | 0,55 | 0,20 | 0,63 | -0,36 | -0,30 |
| Y11_RS12840 | Y11_27241 | *prmA* | 50S ribosomal protein L11 methyltransferase | 0,05 | 0,86 | 0,96 | 0,82 | 0,91 | 0,10 | 0,82 | 0,69 | 1,27 |
| Y11_RS13300 | Y11_28171 | *rpmE* | 50S ribosomal protein L31 | 0,40 | -0,75 | -1,18 | -1,15 | -1,58 | -0,43 | 1,96 | 0,76 | 1,18 |
| Y11_RS13675 | Y11_28961 | *rpmG* | 50S ribosomal protein L33 | 0,22 | -0,66 | -1,18 | -0,88 | -1,40 | -0,52 | 1,75 | 1,25 | 0,96 |
| Y11_RS13680 | Y11_28971 | *rpmB* | 50S ribosomal protein L28 | 0,42 | 0,07 | -0,16 | -0,35 | -0,58 | -0,23 | 2,33 | 1,52 | 1,56 |
| Y11_RS13845 | Y11_29331 | *typA* | ribosome-dependent GTPase TypA | -0,28 | -0,16 | -0,84 | 0,12 | -0,56 | -0,68 | 2,36 | 0,67 | 0,94 |
| Y11_RS13875 | Y11_29401 | *yihA* | ribosome biogenesis GTP-binding protein YihA/YsxC | -0,52 | 0,38 | 0,37 | 0,91 | 0,90 | -0,01 | 0,49 | 0,06 | -0,10 |
| Y11_RS14195 | Y11_30051 | *rpmH* | 50S ribosomal protein L34 | 0,55 | -1,23 | -0,98 | -1,78 | -1,54 | 0,25 | 2,60 | 0,79 | 1,16 |
| Y11_RS14365 | Y11_30401 | | YibL family ribosome-associated protein | -0,11 | 0,85 | 0,66 | 0,96 | 0,77 | -0,19 | 0,12 | -0,25 | -0,50 |
| Y11_RS15200 | Y11_32211 | *hslR* | ribosome-associated heat shock protein Hsp15 | 0,00 | 0,00 | 0,00 | 0,00 | 0,00 | 0,00 | -0,46 | 0,82 | 1,52 |
| Y11_RS15470 | Y11_32811 | *rpsL* | 30S ribosomal protein S12 | 0,39 | -0,25 | -0,71 | -0,65 | -1,10 | -0,45 | 2,37 | 1,38 | 1,50 |
| Y11_RS15475 | Y11_32821 | *rpsG* | 30S ribosomal protein S7 | 0,16 | -0,22 | -0,56 | -0,37 | -0,71 | -0,34 | 2,48 | 1,40 | 1,66 |
| Y11_RS15500 | Y11_32871 | *rpsJ* | 30S ribosomal protein S10 | 0,12 | -0,28 | -0,74 | -0,40 | -0,87 | -0,46 | 2,88 | 1,66 | 2,07 |
| Y11_RS15505 | Y11_32881 | *rplC* | 50S ribosomal protein L3 | 0,37 | -0,92 | -1,13 | -1,29 | -1,50 | -0,21 | 2,65 | 1,53 | 1,96 |
| Y11_RS15510 | Y11_32891 | *rplD* | 50S ribosomal protein L4 | 0,20 | -0,76 | -1,07 | -0,96 | -1,26 | -0,30 | 2,13 | 1,29 | 1,34 |
| Y11_RS15515 | Y11_32901 | *rplW* | 50S ribosomal protein L23 | 0,03 | -0,89 | -1,29 | -0,92 | -1,33 | -0,41 | 2,42 | 1,31 | 1,58 |
| Y11_RS15520 | Y11_32911 | *rplB* | 50S ribosomal protein L2 | 0,41 | -0,58 | -1,00 | -0,99 | -1,40 | -0,42 | 1,62 | 0,98 | 0,77 |
| Y11_RS15525 | | *rpsS* | 30S ribosomal protein S19 | 0,16 | 0,06 | -0,45 | -0,10 | -0,61 | -0,51 | 1,34 | 1,11 | 0,50 |
| Y11_RS15530 | Y11_32931 | *rplV* | 50S ribosomal protein L22 | 0,29 | -0,42 | -0,77 | -0,70 | -1,05 | -0,35 | 1,69 | 1,03 | 0,82 |
| Y11_RS15535 | Y11_32941 | *rpsC* | 30S ribosomal protein S3 | 0,15 | -0,39 | -0,83 | -0,54 | -0,98 | -0,44 | 1,64 | 0,99 | 0,74 |
| Y11_RS15540 | Y11_32951 | *rplP* | 50S ribosomal protein L16 | 0,31 | -0,37 | -0,76 | -0,69 | -1,07 | -0,38 | 1,60 | 0,88 | 0,72 |
| Y11_RS15545 | Y11_32961 | *rpmC* | 50S ribosomal protein L29 | 0,32 | -0,63 | -1,14 | -0,95 | -1,45 | -0,51 | 1,85 | 1,03 | 0,99 |
| Y11_RS15550 | | *rpsQ* | 30S ribosomal protein S17 | -0,01 | 0,29 | -0,14 | 0,29 | -0,14 | -0,43 | 1,11 | 0,61 | 0,11 |
| Y11_RS15555 | Y11_32971 | *rplN* | 50S ribosomal protein L14 | 0,27 | -0,76 | -1,36 | -1,03 | -1,62 | -0,60 | 2,13 | 1,33 | 1,49 |
| Y11_RS15560 | Y11_32981 | *rplX* | 50S ribosomal protein L24 | 0,31 | -0,69 | -0,82 | -1,00 | -1,13 | -0,13 | 2,13 | 1,33 | 1,28 |
| Y11_RS15565 | Y11_32991 | *rplE* | 50S ribosomal protein L5 | 0,31 | -0,69 | -1,15 | -1,00 | -1,46 | -0,47 | 1,82 | 1,13 | 0,85 |
| Y11_RS15570 | Y11_33001 | *rpsN* | 30S ribosomal protein S14 | 0,34 | -0,11 | -0,24 | -0,46 | -0,58 | -0,13 | 2,13 | 1,34 | 1,18 |
| Y11_RS15575 | Y11_33011 | *rpsH* | 30S ribosomal protein S8 | 0,14 | -0,02 | -0,45 | -0,16 | -0,59 | -0,43 | 2,43 | 1,23 | 1,42 |
| Y11_RS15580 | Y11_33021 | *rplF* | 50S ribosomal protein L6 | 0,17 | -0,52 | -0,97 | -0,69 | -1,14 | -0,45 | 1,91 | 1,22 | 0,98 |
| Y11_RS15585 | Y11_33031 | *rplR* | 50S ribosomal protein L18 | 0,37 | -1,18 | -1,70 | -1,55 | -2,07 | -0,52 | 2,20 | 1,28 | 1,29 |
| Y11_RS15590 | Y11_33041 | *rpsE* | 30S ribosomal protein S5 | 0,03 | -0,52 | -1,30 | -0,55 | -1,33 | -0,78 | 1,73 | 1,23 | 0,84 |
| Y11_RS15595 | Y11_33051 | *rpmD* | 50S ribosomal protein L30 | 0,19 | -1,03 | -1,69 | -1,22 | -1,88 | -0,66 | 1,67 | 1,19 | 0,73 |
| Y11_RS15600 | Y11_33061 | *rplO* | 50S ribosomal protein L15 | 0,28 | -0,90 | -1,41 | -1,18 | -1,70 | -0,51 | 1,32 | 1,11 | 0,36 |
| Y11_RS15610 | Y11_33081 | *rpmJ* | 50S ribosomal protein L36 | 0,00 | 0,00 | 0,00 | 0,00 | 0,00 | 0,00 | 1,80 | 1,43 | 1,42 |
| Y11_RS15615 | Y11_33091 | *rpsM* | 30S ribosomal protein S13 | 0,03 | -0,36 | -0,76 | -0,38 | -0,78 | -0,40 | 1,64 | 1,46 | 1,25 |
| Y11_RS15620 | Y11_33101 | *rpsK* | 30S ribosomal protein S11 | 0,33 | -0,26 | -0,80 | -0,58 | -1,13 | -0,54 | 1,72 | 1,48 | 1,26 |
| Y11_RS15625 | Y11_33111 | *rpsD* | 30S ribosomal protein S4 | 0,18 | -0,36 | -0,89 | -0,54 | -1,07 | -0,53 | 2,71 | 1,59 | 2,24 |
| Y11_RS15635 | Y11_33131 | *rplQ* | 50S ribosomal protein L17 | 0,20 | -0,92 | -1,47 | -1,12 | -1,67 | -0,55 | 1,80 | 1,69 | 1,47 |
| Y11_RS15650 | Y11_33161 | | alternative ribosome-rescue factor A | 0,00 | 0,00 | 0,00 | 0,00 | 0,00 | 0,00 | 1,84 | 0,70 | 0,53 |
| Y11_RS16540 | Y11_34901 | *rplK* | 50S ribosomal protein L11 | 0,14 | -0,77 | -1,51 | -0,90 | -1,65 | -0,74 | 2,83 | 1,37 | 1,58 |
| Y11_RS16545 | Y11_34911 | *rplA* | 50S ribosomal protein L1 | 0,22 | -0,71 | -1,05 | -0,94 | -1,27 | -0,34 | 2,32 | 1,25 | 1,05 |
| Y11_RS16550 | Y11_34921 | *rplJ* | 50S ribosomal protein L10 | 0,12 | -0,40 | -0,93 | -0,51 | -1,04 | -0,53 | 1,93 | 1,10 | 0,86 |
| Y11_RS16555 | Y11_34931 | *rplL* | 50S ribosomal protein L7/L12 | -0,10 | -0,32 | -0,75 | -0,22 | -0,65 | -0,43 | 1,45 | 0,86 | 0,38 |
| Y11_RS17120 | Y11_36031 | *rsgA* | small ribosomal subunit biogenesis GTPase RsgA | -0,11 | 0,73 | 0,87 | 0,84 | 0,99 | 0,14 | 0,62 | -0,38 | -0,38 |
| Y11_RS17240 | Y11_36271 | *rpsF* | 30S ribosomal protein S6 | 0,06 | -0,16 | -0,78 | -0,22 | -0,85 | -0,62 | 1,52 | 1,39 | 1,22 |
| Y11_RS17250 | Y11_36291 | *rpsR* | 30S ribosomal protein S18 | 0,09 | -0,44 | -1,13 | -0,53 | -1,22 | -0,69 | 1,70 | 1,12 | 1,21 |
| Y11_RS17255 | Y11_36301 | *rplI* | 50S ribosomal protein L9 | 0,25 | -0,43 | -0,97 | -0,67 | -1,22 | -0,54 | 0,79 | 0,94 | 0,40 |
| Y11_RS17375 | Y11_36551 | *rplU* | 50S ribosomal protein L21 | 0,24 | -0,54 | -1,06 | -0,77 | -1,29 | -0,52 | 1,84 | 1,29 | 1,34 |
| Y11_RS17380 | Y11_36561 | *rpmA* | 50S ribosomal protein L27 | 0,22 | -0,36 | -0,71 | -0,58 | -0,93 | -0,36 | 1,34 | 1,14 | 0,78 |
| Y11_RS17415 | Y11_36631 | *yhbY* | ribosome assembly RNA-binding protein YhbY | -0,15 | 0,47 | 0,38 | 0,62 | 0,53 | -0,09 | 1,09 | -0,02 | -0,33 |
| Y11_RS17455 | Y11_36691 | *rimP* | ribosome maturation factor RimP | -0,12 | -0,05 | 0,03 | 0,06 | 0,15 | 0,09 | 2,00 | 1,00 | 1,09 |
| Y11_RS17470 | Y11_36721 | *rbfA* | 30S ribosome-binding factor RbfA | -0,35 | -0,85 | -1,73 | -0,51 | -1,38 | -0,88 | 1,64 | 0,59 | 0,60 |
| Y11_RS17480 | Y11_36741 | *rpsO* | 30S ribosomal protein S15 | 0,02 | 0,26 | -0,25 | 0,24 | -0,27 | -0,51 | 2,38 | 1,25 | 1,21 |
| Y11_RS18060 | Y11_37981 | *rimI* | ribosomal protein S18-alanine N-acetyltransferase | 0,92 | 1,39 | 2,56 | 0,47 | 1,64 | 1,16 | 1,14 | -0,23 | 0,20 |
| Y11_RS18290 | Y11_38451 | *rpsT* | 30S ribosomal protein S20 | 0,39 | -0,27 | -0,70 | -0,66 | -1,08 | -0,43 | 2,49 | 0,99 | 1,36 |
| Y11_RS19505 | Y11_41021 | *rpsB* | 30S ribosomal protein S2 | 0,25 | -0,18 | -0,55 | -0,42 | -0,79 | -0,37 | 2,19 | 1,18 | 1,27 |
| Y11_RS19520 | Y11_41051 | *frr* | ribosome recycling factor | 0,11 | 0,02 | -0,10 | -0,08 | -0,21 | -0,13 | 1,37 | 1,03 | 1,53 |
| Y11_RS20460 | Y11_42971 | *der* | ribosome biogenesis GTPase Der | -0,70 | -0,01 | -0,85 | 0,69 | -0,15 | -0,84 | 1,70 | 0,79 | 1,35 |
| Y11_RS21380 | Y11_04571 | *rmf* | ribosome modulation factor | 3,41 | 1,83 | 3,99 | -1,58 | 0,58 | 2,13 | -5,61 | -2,78 | -4,02 |
| **tRNA-associated proteins** | | | | | | | | | | | | |
| Y11_RS00390 | Y11_00831 | *gltX* | glutamate--tRNA ligase | 0,27 | 0,52 | 0,84 | 0,26 | 0,58 | 0,32 | 1,466 | 0,89 | 1,095 |
| Y11_RS00745 | Y11_01531 | *mnmC* | bifunctional tRNA (5-methylaminomethyl-2-thiouridine)(34)-methyltransferase MnmD/FAD-dependent 5-carboxymethylaminomethyl-2-thiouridine(34) oxidoreductase MnmC | 0,24 | -1 | -1,04 | -1,24 | -1,28 | -0,04 | 0,495 | 1,017 | 0,771 |
| Y11_RS00860 | Y11_01781 | *truA* | tRNA pseudouridine(38-40) synthase TruA | -0,3 | -0,23 | -0,47 | 0,07 | -0,18 | -0,24 | 0,963 | 0,571 | 0,671 |
| Y11_RS01840 | Y11_03951 | *aat* | leucyl/phenylalanyl-tRNA--protein transferase | 0,97 | 1,58 | 2,06 | 0,61 | 1,09 | 0,48 | 1,593 | 1,006 | 1,328 |
| Y11_RS01880 | Y11_04031 | *serS* | serine--tRNA ligase | 0,18 | -0,55 | -0,87 | -0,74 | -1,06 | -0,32 | 0,845 | 0,299 | 0,732 |
| Y11_RS02015 | Y11_04351 | *cmoM* | tRNA uridine 5-oxyacetic acid(34) methyltransferase CmoM | 0,38 | -0,97 | -0,81 | -1,34 | -1,19 | 0,16 | 1,318 | 0,19 | 0,557 |
| Y11_RS02070 | Y11_04451 | *asnS* | asparagine--tRNA ligase | 0,86 | 0,13 | 0,08 | -0,72 | -0,77 | -0,05 | 1,365 | 0,542 | 1,171 |
| Y11_RS02635 | Y11_05651 | *mnmA* | tRNA 2-thiouridine(34) synthase MnmA | -0,31 | -0,78 | -0,81 | -0,48 | -0,5 | -0,03 | 1,652 | 1,055 | 1,273 |
| Y11_RS03280 | Y11_07081 | *argS* | arginine--tRNA ligase | -0,14 | -0,85 | -0,9 | -0,71 | -0,76 | -0,05 | 1,707 | 0,883 | 1,291 |
| Y11_RS03605 | Y11_07851 | *thrS* | threonine--tRNA ligase | 0,26 | 0,49 | 0,27 | 0,22 | 0 | -0,22 | 0,866 | 0,51 | 0,386 |
| Y11_RS03625 | Y11_07891 | *pheS* | phenylalanine--tRNA ligase subunit alpha | -0,1 | -0,49 | -0,93 | -0,39 | -0,83 | -0,44 | 0,855 | 0,744 | 1,143 |
| Y11_RS03630 | Y11_07901 | *pheT* | phenylalanine--tRNA ligase subunit beta | -0,08 | -0,66 | -1,02 | -0,58 | -0,94 | -0,36 | 0,583 | 0,352 | 0,665 |

| | | | | | | | | | | | | |
|---|---|---|---|---|---|---|---|---|---|---|---|---|
| Y11_RS03940 | Y11_08501 | *tyrS* | tyrosine--tRNA ligase | 0,15 | -0,36 | -0,29 | -0,51 | -0,44 | 0,07 | 1,527 | 0,387 | 1,215 |
| Y11_RS04105 | Y11_08841 | *ttcA* | tRNA 2-thiocytidine(32) synthetase TtcA | 0,44 | -0,02 | -1,65 | -0,47 | -2,1 | -1,63 | 1,552 | -0,31 | 0,16 |
| Y11_RS05385 | Y11_11651 | *tsaB* | tRNA (adenosine(37)-N6)-threonylcarbamoyltransferase complex dimerization subunit type 1 TsaB | -0,05 | -1,66 | -1,96 | -1,61 | -1,91 | -0,3 | 1,399 | 1,76 | 1,687 |
| Y11_RS05490 | Y11_11871 | *aspS* | aspartate--tRNA ligase | -0,59 | -0,72 | -0,85 | -0,12 | -0,25 | -0,13 | 1,039 | 0,437 | 0,632 |
| Y11_RS05520 | Y11_11941 | *cmoB* | tRNA 5-methoxyuridine(34)/uridine 5-oxyacetic acid(34) synthase CmoB | -0,83 | -1,16 | -1,54 | -0,33 | -0,71 | -0,38 | 2,276 | 1,329 | 1,79 |
| Y11_RS05825 | Y11_12561 | *hemA* | glutamyl-tRNA reductase | 0,11 | -0,65 | -1,21 | -0,76 | -1,33 | -0,56 | 2,071 | 0,305 | 0,986 |
| Y11_RS05855 | Y11_12621 | *pth* | aminoacyl-tRNA hydrolase | -0,04 | -0,74 | -1,27 | -0,69 | -1,23 | -0,53 | 1,695 | 1,136 | 1,26 |
| Y11_RS07910 | Y11_16881 | *metG* | methionine--tRNA ligase | -0,02 | 0,11 | 0,33 | 0,13 | 0,35 | 0,22 | 0,574 | -0,24 | -0,01 |
| Y11_RS08440 | Y11_17981 | *dusC* | tRNA dihydrouridine(16) synthase DusC | -0,15 | -0,12 | -0,72 | 0,02 | -0,57 | -0,6 | 0,988 | 0,789 | 1,147 |
| Y11_RS08920 | Y11_18971 | *glnS* | glutamine--tRNA ligase | 0,36 | -1,3 | -1,32 | -1,65 | -1,67 | -0,02 | 1,58 | 0,743 | 1,058 |
| Y11_RS08990 | Y11_19061 | *miaB* | tRNA (N6-isopentenyl adenosine(37)-C2)-methylthiotransferase MiaB | 0,74 | 1,4 | 0,96 | 0,66 | 0,22 | -0,44 | 0,892 | -0,6 | -0,34 |
| Y11_RS09045 | Y11_19181 | *leuS* | leucine--tRNA ligase | -0,35 | -0,14 | -0,36 | 0,21 | -0,01 | -0,22 | 1,43 | 0,258 | 0,399 |
| Y11_RS09275 | Y11_19671 | *cysS* | cysteine--tRNA ligase | 0,06 | 0,94 | 0,65 | 0,87 | 0,58 | -0,29 | 0,852 | -0,34 | -0,07 |
| Y11_RS09350 | Y11_19831 | *ybaK* | Cys-tRNA(Pro)/Cys-tRNA(Cys) deacylase YbaK | -0,19 | 0,16 | -0,13 | 0,35 | 0,06 | -0,29 | 0,194 | 0,493 | 0,614 |
| Y11_RS09735 | Y11_20701 | *thiI* | tRNA 4-thiouridine(8) synthase ThiI | -0,61 | -0,28 | -0,62 | 0,33 | -0,01 | -0,34 | 1,707 | 1,002 | 1,147 |
| Y11_RS09820 | Y11_20871 | *tgt* | tRNA guanosine(34) transglycosylase Tgt | 0,79 | 0,2 | 0,38 | -0,59 | -0,41 | 0,18 | 1,842 | 1,346 | 1,411 |
| Y11_RS09835 | Y11_20911 | *queA* | tRNA preQ1(34) S-adenosylmethionine ribosyltransferase-isomerase QueA | 0,48 | 0,44 | -0,12 | -0,03 | -0,59 | -0,56 | 1,898 | 1,346 | 1,495 |
| Y11_RS10260 | Y11_21841 | | tRNA/rRNA methyltransferase | 0,24 | 0,94 | 0,75 | 0,7 | 0,5 | -0,2 | 0,862 | 2,587 | 2,528 |
| Y11_RS10380 | Y11_22051 | *trmD* | tRNA (guanosine(37)-N1)-methyltransferase TrmD | -0,48 | 0 | -0,8 | 0,48 | -0,32 | -0,8 | 2,635 | 1,262 | 1,687 |
| Y11_RS10450 | Y11_22161 | *alaS* | alanine--tRNA ligase | 0,59 | -0,29 | -0,28 | -0,88 | -0,87 | 0,01 | 1,296 | 0,91 | 0,915 |
| Y11_RS10740 | Y11_22781 | *truD* | tRNA pseudouridine(13) synthase TruD | 0,31 | -0,59 | -0,84 | -0,9 | -1,14 | -0,24 | 0,475 | 0,299 | 0,508 |
| Y11_RS11050 | Y11_23441 | *trmB* | tRNA (guanosine(46)-N7)-methyltransferase TrmB | -0,06 | -1,62 | -1,69 | -1,56 | -1,63 | -0,07 | 0,669 | 1,274 | 1,259 |
| Y11_RS11815 | Y11_25061 | *tsaD* | tRNA (adenosine(37)-N6)-threonylcarbamoyltransferase complex transferase subunit TsaD | -0,42 | 0,18 | 0,09 | 0,6 | 0,51 | -0,09 | 1,486 | 0,336 | 0,411 |
| Y11_RS12845 | Y11_27251 | *dusB* | tRNA dihydrouridine synthase DusB | 0,01 | -1,31 | -1,62 | -1,32 | -1,63 | -0,31 | 2,405 | 2,397 | 2,839 |
| Y11_RS13000 | Y11_27581 | *dusA* | tRNA dihydrouridine(20/20a) synthase DusA | -0,76 | -0,97 | -1,45 | -0,21 | -0,7 | -0,49 | 2,044 | -0,09 | 0,303 |
| Y11_RS13180 | Y11_27911 | *trmA* | tRNA (uridine(54)-C5)-methyltransferase TrmA | 0,73 | -0,99 | -0,6 | -1,72 | -1,33 | 0,39 | 1,399 | 2,103 | 1,988 |
| Y11_RS13575 | Y11_28751 | *trmL* | tRNA (uridine(34)/cytosine(34)/5-carboxymethylaminomethyluridine(34)-2'-O)-methyltransferase TrmL | 0,14 | -0,47 | -0,57 | -0,61 | -0,71 | -0,1 | 2,009 | 2,176 | 2,535 |
| Y11_RS13775 | Y11_29181 | *trmH* | tRNA (guanosine(18)-2'-O)-methyltransferase TrmH | -0,1 | 0,11 | -0,36 | 0,21 | -0,25 | -0,47 | 3,065 | 0,229 | 2,14 |
| Y11_RS13830 | Y11_29301 | *dtd* | D-aminoacyl-tRNA deacylase | -0,03 | 0,49 | 0,52 | 0,51 | 0,55 | 0,04 | -0,99 | -0,32 | -0,43 |
| Y11_RS14015 | Y11_29661 | *mnmG* | tRNA uridine-5-carboxymethylaminomethyl(34) synthesis enzyme MnmG | 0,68 | -1,17 | -1,09 | -1,47 | -1,39 | 0,08 | 2,064 | 1,435 | 1,503 |
| Y11_RS14180 | Y11_30001 | *mnmE* | tRNA uridine-5-carboxymethylaminomethyl(34) synthesis GTPase MnmE | -0,1 | -1,17 | -1,39 | -1,07 | -1,29 | -0,22 | 1,455 | 1,069 | 1,671 |
| Y11_RS14325 | Y11_30321 | *glyQ* | glycine--tRNA ligase subunit alpha | -0,23 | -0,92 | -0,87 | -0,69 | -0,63 | 0,05 | 1,59 | 0,512 | 1,005 |
| Y11_RS14330 | Y11_30331 | *glyS* | glycine--tRNA ligase subunit beta | -0,16 | -0,47 | -0,46 | -0,31 | -0,3 | 0,01 | 0,938 | 0,553 | 0,504 |
| Y11_RS14425 | Y11_30511 | *selA* | L-seryl-tRNA(Sec) selenium transferase | -0,17 | 0,25 | 0,01 | 0,41 | 0,17 | -0,24 | 0,181 | -0,29 | -0,04 |
| Y11_RS15280 | Y11_32391 | *trpS* | tryptophan--tRNA ligase | 0,19 | -0,63 | -0,48 | -0,82 | -0,67 | 0,15 | 0,435 | 0,472 | 0,631 |
| Y11_RS15675 | Y11_33221 | *fmt* | methionyl-tRNA formyltransferase | 0,19 | 0,94 | 0,7 | 0,75 | 0,5 | -0,25 | -0,23 | -0,87 | -0,93 |
| Y11_RS17140 | Y11_36051 | *queG* | tRNA epoxyqueuosine(34) reductase QueG | 0 | 0 | -1,12 | 0 | -0,47 | -0,75 | 1,282 | 0,099 | 0,328 |
| Y11_RS17150 | Y11_36071 | *tsaE* | tRNA (adenosine(37)-N6)-threonylcarbamoyltransferase complex ATPase subunit type 1 TsaE | -0,31 | -0,45 | -0,24 | -0,14 | 0,07 | 0,21 | 0,426 | -0,16 | 0,128 |
| Y11_RS17475 | Y11_36731 | *truB* | tRNA pseudouridine(55) synthase TruB | -0,03 | -0,78 | -2,06 | -0,75 | -2,03 | -1,27 | 1,835 | 0,846 | 0,686 |
| Y11_RS17740 | Y11_37321 | | valine--tRNA ligase | -0,04 | -0,11 | -0,18 | -0,07 | -0,14 | -0,07 | 1,321 | 0,363 | 0,582 |
| Y11_RS18305 | Y11_38471 | *ileS* | isoleucine--tRNA ligase | -0,02 | -0,51 | -0,58 | -0,49 | -0,56 | -0,07 | 1,301 | 0,073 | 0,33 |
| Y11_RS18395 | Y11_38651 | *rluA* | bifunctional tRNA pseudouridine(32) synthase/23S rRNA pseudouridine(746) synthase RluA | 0,24 | -1,87 | -3,06 | -2,11 | -3,3 | -1,19 | 2,088 | 1,291 | 1,661 |
| Y11_RS18815 | Y11_39561 | *gluQRS* | tRNA glutamyl-Q(34) synthetase GluQRS | -0,58 | 15 | 15 | 15 | 15 | 0 | -0,37 | -0,11 | -0,28 |
| Y11_RS19015 | Y11_39991 | *ygfZ* | tRNA-modifying protein YgfZ | 0,94 | 0,49 | 0,72 | -0,45 | -0,21 | 0,23 | -0,3 | -0,12 | -0,39 |
| Y11_RS19070 | Y11_40111 | *lysS* | lysine--tRNA ligase | -0,21 | -0,81 | -0,66 | -0,61 | -0,45 | 0,16 | 1,557 | 0,308 | 0,397 |
| Y11_RS19405 | Y11_40821 | *tcdA* | tRNA cyclic N6-threonylcarbamoyladenosine(37) synthase TcdA | -0,16 | -0,9 | -1,19 | -0,74 | -1,03 | -0,29 | 1,854 | 0,965 | 1,828 |
| Y11_RS19475 | Y11_40961 | *truC* | tRNA pseudouridine(65) synthase TruC | 0 | 0 | 0 | 0 | 0 | 0 | 2,147 | 1,218 | 2,031 |
| Y11_RS19600 | Y11_41211 | *tilS* | tRNA lysidine(34) synthetase TilS | 0,1 | -0,14 | 0,24 | -0,24 | 0,14 | 0,38 | -0,56 | -0,05 | -0,19 |
| Y11_RS19625 | Y11_41261 | *arfB* | aminoacyl-tRNA hydrolase | 0 | 0 | 0 | 0 | 0 | 0 | -0,22 | -0,49 | -0,34 |
| Y11_RS19635 | Y11_41281 | *proS* | proline--tRNA ligase | 0,05 | -0,16 | -0,58 | -0,21 | -0,64 | -0,42 | 1,2 | 0,926 | 0,997 |
| Y11_RS19640 | Y11_41301 | *tsaA* | tRNA (N6-threonylcarbamoyladenosine(37)-N6)-methyltransferase TrmO | 0 | 0 | 0 | 0 | 0 | 0 | 1,098 | 0,783 | 1,337 |
| Y11_RS20125 | Y11_42271 | | tRNA1(Val) (adenine(37)-N6)-methyltransferase | -0,27 | -1,09 | -1,34 | -0,81 | -1,07 | -0,25 | 1,52 | 1,483 | 1,791 |
| Y11_RS20225 | Y11_42461 | *tadA* | tRNA adenosine(34) deaminase TadA | 0,09 | -0,25 | -0,28 | -0,34 | -0,37 | -0,03 | 0,92 | 1,689 | 1,797 |
| Y11_RS20335 | Y11_42691 | *trmJ* | tRNA (cytosine(32)/uridine(32)-2'-O)-methyltransferase TrmJ | -0,05 | -0,17 | -0,37 | -0,12 | -0,32 | -0,2 | 1,778 | 0,385 | 0,861 |
| Y11_RS20420 | Y11_42881 | *rlmN* | bifunctional tRNA (adenosine(37)-C2)-methyltransferase TrmG/ribosomal RNA large subunit methyltransferase RlmN | 0,11 | -1,27 | -2,92 | -1,38 | -3,03 | -1,65 | 1,572 | 0,456 | 0,722 |
| Y11_RS20440 | Y11_42931 | *hisS* | histidine--tRNA ligase | -0,28 | 0,05 | -0,25 | 0,33 | 0,03 | -0,3 | -0,01 | -0,1 | -0,06 |
| Y11_RS20575 | Y11_43251 | *yegQ* | tRNA 5-hydroxyuridine modification protein YegQ | 0,39 | -1,23 | -1,85 | -1,62 | -2,24 | -0,62 | 1,079 | 1,469 | 1,632 |
| **other protein which expression pattern correlates with *dksA* inactivation** | | | | | | | | | | | | |
| Y11_RS00165 | Y11_00341 | | response regulator | 0,28 | 1,29 | 1,91 | 1,01 | 1,63 | 0,63 | 1,209 | -1,36 | -0,6 |
| Y11_RS00435 | Y11_00891 | | alpha-keto acid decarboxylase family protein | -0,19 | 2,35 | 2,2 | 2,54 | 2,39 | -0,14 | -1,89 | -1,18 | -1,82 |
| Y11_RS00795 | Y11_01641 | *agp* | bifunctional glucose-1-phosphatase/inositol phosphatase | -0,24 | 2,95 | 4,48 | 3,18 | 4,72 | 1,53 | -2,47 | -1,83 | -1,21 |
| Y11_RS00820 | Y11_01701 | | DsbA family protein | 0,03 | -3,35 | -2,48 | -3,37 | -2,52 | 0,87 | 0,967 | 1,019 | 0,682 |
| Y11_RS00885 | Y11_01841 | *cvpA* | colicin V production protein | 0,01 | -1,25 | -1,58 | -1,26 | -1,58 | -0,32 | 1,353 | 1,534 | 1,49 |
| Y11_RS00900 | Y11_01871 | *hisJ* | histidine ABC transporter substrate-binding protein HisJ | -0,39 | 1,02 | 3,72 | 1,41 | 4,11 | 2,7 | -1,54 | -3,19 | -2,14 |
| Y11_RS00930 | Y11_01931 | | TIGR01777 family oxidoreductase | -0,03 | 0,94 | 2,14 | 0,97 | 2,17 | 1,2 | -1,43 | -1,68 | -1,31 |
| Y11_RS01140 | Y11_02411 | *elaB* | stress response protein ElaB | 0,58 | 4,97 | 4,77 | 4,39 | 4,19 | -0,2 | -3,18 | -2,39 | -3,44 |
| Y11_RS01145 | Y11_02421 | *menF* | isochorismate synthase MenF | -0,1 | -1,56 | -2,08 | -1,46 | -1,97 | -0,52 | 1,584 | 1,528 | 1,575 |
| Y11_RS01485 | Y11_03161 | *yeiP* | elongation factor P-like protein YeiP | -0,26 | -2,75 | -3,17 | -2,49 | -2,91 | -0,42 | 1,017 | 3,023 | 2,848 |
| Y11_RS01595 | Y11_03421 | | ligand-gated channel protein | 1,21 | -1,56 | -1,96 | -2,77 | -3,17 | -0,4 | -0,9 | 1,704 | 1,714 |
| Y11_RS01725 | Y11_03711 | *rlmC* | 23S rRNA (uracil(747)-C(5))-methyltransferase RlmC | -0,04 | -0,82 | -1,7 | -0,78 | -1,65 | -0,87 | 2,186 | 1,41 | 2,167 |
| Y11_RS01985 | Y11_04271 | | cold-shock protein | 0,02 | -4,68 | -5,3 | -4,7 | -5,31 | -0,62 | 7,567 | 1,288 | 1,067 |
| Y11_RS02095 | Y11_04511 | | YcbX family protein | 0,54 | -2,03 | -2,06 | -2,57 | -2,6 | -0,03 | 0,156 | 1,441 | 1,235 |
| Y11_RS02145 | Y11_04621 | *matP* | macrodomain Ter protein MatP | 0,11 | 0,92 | 1,08 | 0,81 | 0,97 | 0,17 | -0,89 | -1,16 | -1,44 |
| Y11_RS02275 | Y11_04891 | | acyl-homoserine-lactone synthase | -0,36 | -2,42 | -2,17 | -2,06 | -1,79 | 0,26 | 0,669 | 0,822 | 0,835 |
| Y11_RS02465 | Y11_05301 | *fabF* | beta-ketoacyl-ACP synthase II | -0,12 | -1,02 | -1,2 | -0,9 | -1,09 | -0,19 | 1,736 | 1,448 | 1,859 |
| Y11_RS02625 | Y11_05631 | *purB* | adenylosuccinate lyase | -0,28 | -1,13 | -1,17 | -0,85 | -0,89 | -0,04 | 1,02 | 1,185 | 1,244 |
| Y11_RS02630 | Y11_05641 | *hflD* | high frequency lysogenization protein HflD | -0,37 | -1,17 | -0,74 | -0,8 | -0,38 | 0,43 | 1,876 | 1,072 | 1,492 |
| Y11_RS02810 | Y11_06021 | | serine protein kinase RIO | 0,15 | -1,53 | -1,84 | -1,67 | -1,99 | -0,31 | -0,06 | 1,164 | 0,978 |
| Y11_RS02815 | Y11_06031 | | ABC transporter ATP-binding protein/permease | 0,51 | -0,39 | -1,24 | -0,9 | -1,75 | -0,84 | 1,124 | 1,065 | 0,823 |
| Y11_RS02825 | Y11_06061 | | amino acid permease | 0,37 | 1,44 | 15 | 1,08 | 0 | 15 | -3 | -1,08 | -1,65 |
| Y11_RS02915 | Y11_06251 | | TonB-dependent receptor | 1,51 | -0,54 | -1,43 | -2,04 | -2,94 | -0,9 | -0,06 | 2,246 | 1,793 |
| Y11_RS03015 | Y11_06491 | *ftnA* | non-heme ferritin | -0,27 | 1,46 | 2,04 | 1,73 | 2,31 | 0,58 | 0,608 | -1,93 | -0,97 |
| Y11_RS03540 | Y11_07691 | | fructosamine kinase family protein | -0,48 | 2,37 | 3,72 | 2,85 | 4,2 | 1,35 | -2 | -1,71 | -1,97 |
| Y11_RS04155 | Y11_08961 | *azoR* | FMN-dependent NADH-azoreductase | 0 | -3,28 | -5,63 | -3,28 | -5,63 | -2,35 | -0,17 | 1,233 | 1,24 |
| Y11_RS04260 | Y11_09181 | | SDR family oxidoreductase | 0 | 1,16 | 2,34 | 1,16 | 2,34 | 1,19 | -1,09 | -1,1 | -0,87 |
| Y11_RS04290 | Y11_09261 | *ilvN* | acetolactate synthase small subunit | -0,61 | -3,2 | -1,52 | -2,59 | -0,91 | 1,68 | 0,489 | 1,13 | 1,001 |
| Y11_RS04600 | | | SHOCT domain-containing protein | 0,13 | 1,14 | 2,23 | 1,01 | 2,1 | 1,09 | -1,75 | -1,26 | -1,56 |
| Y11_RS04790 | Y11_10331 | | alpha-xenorhabdolysin family binary toxin subunit A | -2,49 | 2,11 | 2,38 | 4,6 | 4,87 | 0 | -1,66 | -1,98 | -0,61 |
| Y11_RS04865 | Y11_10511 | | DUF1471 domain-containing protein | 15 | -7,58 | -2,1 | -15 | -15 | 5,48 | 1,261 | 1,999 | 2,303 |
| Y11_RS04910 | Y11_10601 | *acnA* | aconitate hydratase AcnA | -0,71 | 1,12 | 1,73 | 1,83 | 2,44 | 0,61 | -0,85 | -1,91 | -1,63 |
| Y11_RS05000 | Y11_10811 | | BON domain-containing protein | 2,05 | 2,95 | 4,08 | 0,9 | 2,03 | 1,12 | -0,85 | -1,08 | -1,29 |
| Y11_RS05035 | Y11_10881 | *tonB* | TonB system transport protein TonB | 0,98 | -1,07 | -1,96 | -2,05 | -2,94 | -0,89 | 1,023 | 1,148 | 1,334 |
| Y11_RS05195 | Y11_11231 | *ansA* | asparaginase | 0,07 | -1,46 | -1,46 | -1,53 | -1,53 | -0,01 | 1,204 | 1,079 | 1,323 |
| Y11_RS05240 | Y11_11331 | *yeaG* | protein kinase YeaG | 0,02 | 1,36 | 1,9 | 1,35 | 1,89 | 0,54 | -2,93 | -1,84 | -2,14 |
| Y11_RS05245 | Y11_11341 | | YeaH/YhbH family protein | 0,97 | 2,48 | 2,97 | 1,51 | 2,01 | 0,5 | -2,87 | -1,46 | -1,8 |
| Y11_RS05345 | Y11_11571 | | YcgL domain-containing protein | 0,15 | -0,95 | -1,31 | -1,1 | -1,46 | -0,36 | 1,058 | 1,036 | 1,462 |
| Y11_RS05350 | Y11_11581 | | hypothetical protein | 0 | -2,46 | -2,13 | -15 | -15 | 0,33 | -0,03 | 1,735 | 1,762 |
| Y11_RS05625 | Y11_12171 | *hutG* | N-formylglutamate deformylase | -0,6 | 0,76 | 2,93 | 1,36 | 3,52 | 2,17 | -4,09 | -2,9 | -2,89 |
| Y11_RS05640 | Y11_12201 | | porin | 1,22 | -2,84 | -2,99 | -4,07 | -4,19 | -0,15 | -4,16 | 1,055 | 0,565 |
| Y11_RS05660 | Y11_12251 | *astD* | succinylglutamate-semialdehyde dehydrogenase | 0,13 | 2,18 | 2,85 | 2,05 | 2,72 | 0,68 | -4,06 | -3,29 | -3,31 |
| Y11_RS05665 | Y11_12261 | *astA* | arginine N-succinyltransferase | 0,33 | 2,57 | 3,46 | 2,24 | 3,13 | 0,89 | -4,2 | -3,91 | -4,03 |
| Y11_RS05670 | Y11_12271 | | aspartate aminotransferase family protein | 0,32 | 2,19 | 3,07 | 1,87 | 2,75 | 0,88 | -4,56 | -4,01 | -4,64 |
| Y11_RS05680 | Y11_12291 | | cytochrome ubiquinol oxidase subunit I | -0,37 | 1,08 | 15 | 1,45 | 15 | 15 | -1,55 | -1,14 | -1,86 |
| Y11_RS05710 | Y11_12361 | | siderophore-interacting protein | 0,76 | -0,55 | -1,22 | -1,31 | -1,98 | -0,67 | 0,008 | 1,935 | 1,748 |
| Y11_RS06445 | Y11_13801 | *wrbA* | NAD(P)H:quinone oxidoreductase | 0,23 | 2,39 | 2,99 | 2,16 | 2,76 | 0,59 | -3,17 | -1,25 | -1,62 |

| | | | | | | | | | | | | |
|---|---|---|---|---|---|---|---|---|---|---|---|---|
| Y11_RS06845 | Y11_14701 | *uspC* | universal stress protein UspC | -0,76 | 1,86 | 2,53 | 2,62 | 3,28 | 0,66 | -0,77 | -1,15 | -1,01 |
| Y11_RS06850 | Y11_14711 | | NADP-dependent oxidoreductase | -0,4 | 0,84 | 1,67 | 1,24 | 2,07 | 0,83 | -1,25 | -1,09 | -1,27 |
| Y11_RS06935 | Y11_14901 | *glgC* | glucose-1-phosphate adenylyltransferase | -1,11 | 2,21 | 2,79 | 3,31 | 3,89 | 0,58 | -5,45 | -3,38 | -4,21 |
| Y11_RS08060 | Y11_17171 | *mglB* | galactose/glucose ABC transporter substrate-binding protein MglB | -3,1 | 1,08 | 3,03 | 4,18 | 6,13 | 1,95 | 0,558 | -1,68 | 0,372 |
| Y11_RS08190 | Y11_17441 | | sigma-70 family RNA polymerase sigma factor | -0,56 | 1,9 | 3,21 | 2,47 | 3,76 | 1,31 | -1,36 | -1,08 | -1,07 |
| Y11_RS08200 | Y11_17461 | *tam* | trans-aconitate 2-methyltransferase | -0,21 | 1,97 | 2,77 | 2,18 | 2,98 | 0,8 | -3,41 | -1,17 | -1,91 |
| Y11_RS08265 | Y11_17621 | *aqpZ* | aquaporin Z | -0,79 | 2,64 | 4,48 | 3,43 | 5,27 | 1,84 | -1,26 | -0,91 | -1,73 |
| Y11_RS08275 | Y11_17651 | *ompF2* | porin OmpF2 | 0,19 | -0,96 | -1,92 | -1,15 | -2,12 | -0,96 | 1,181 | 1,901 | 1,894 |
| Y11_RS08445 | Y11_17991 | *rhlE* | ATP-dependent RNA helicase RhlE | -0,3 | -3,09 | -4,53 | -2,79 | -4,23 | -1,44 | 4,037 | 1,415 | 1,066 |
| Y11_RS08465 | Y11_18031 | | ABC transporter permease | 0,11 | 1,91 | 1,99 | 1,8 | 1,88 | 0,08 | -1,38 | -1,07 | -1,03 |
| Y11_RS08610 | Y11_18351 | | PsiF family protein | 2,01 | 3,4 | 3,93 | 1,39 | 1,92 | 0,53 | -3,3 | -2,05 | -2,88 |
| Y11_RS09025 | Y11_19131 | | amino acid ABC transporter permease | 3,16 | 15 | 15 | 15 | 15 | 0 | -2,18 | -1,81 | -1,4 |
| Y11_RS09030 | Y11_19141 | *gltK* | glutamate/aspartate ABC transporter permease GltK | 1,16 | 5,12 | 2,38 | 3,98 | 1,21 | 0 | -1,19 | -1,54 | -1,25 |
| Y11_RS09040 | Y11_19161 | | zinc ribbon-containing protein | -0,28 | 2,25 | 3,29 | 2,53 | 3,57 | 1,04 | -2,72 | -2,5 | -2,96 |
| Y11_RS09455 | Y11_20071 | *apt* | adenine phosphoribosyltransferase | 0,22 | -0,99 | -1,12 | -1,21 | -1,34 | -0,14 | 1,628 | 1,165 | 1,511 |
| Y11_RS09575 | Y11_20361 | | YbaY family lipoprotein | -0,34 | 3,64 | 3,49 | 3,98 | 3,83 | -0,15 | -0,49 | -1,39 | -1,25 |
| Y11_RS09725 | Y11_20681 | *panE* | 2-dehydropantoate 2-reductase | 0 | 2,14 | 2,73 | 2,15 | 2,73 | 0,58 | -0,18 | -1,05 | -1,12 |
| Y11_RS09730 | Y11_20691 | *yajL* | protein deglycase YajL | 0,53 | 1,44 | 2,4 | 0,91 | 1,86 | 0,95 | -0,71 | -1,01 | -1,3 |
| Y11_RS09985 | Y11_21261 | *proA* | glutamate-5-semialdehyde dehydrogenase | 0,28 | -1,69 | -1,33 | -1,97 | -1,61 | 0,36 | -0,6 | 1,167 | 1,003 |
| Y11_RS09990 | Y11_21271 | *proB* | glutamate 5-kinase | 0,24 | -2,21 | -2,75 | -2,45 | -2,99 | -0,54 | -0,04 | 1,401 | 1,386 |
| Y11_RS10275 | Y11_21871 | | bifunctional acetate--CoA ligase family protein/GNAT family N-acetyltransferase | -0,86 | 1,37 | 2,53 | 2,23 | 3,39 | 1,16 | -1,18 | -1,44 | -0,95 |
| Y11_RS10500 | Y11_22271 | | fumarate hydratase | -0,43 | 2,31 | 3,46 | 2,74 | 3,89 | 1,16 | 0,074 | -1,17 | 0,004 |
| Y11_RS10720 | Y11_22741 | *rpoS* | RNA polymerase sigma factor RpoS | -0,55 | 2,67 | 2,83 | 3,22 | 3,38 | 0,16 | -2,16 | -2,11 | -2,53 |
| Y11_RS10840 | Y11_22981 | *sodC* | superoxide dismutase family protein | 0,7 | 3,08 | 3,5 | 2,38 | 2,8 | 0,42 | -1,67 | -1,15 | -1,44 |
| Y11_RS11280 | Y11_23911 | | DUF3313 domain-containing protein | 0,93 | 3,27 | 3,97 | 2,35 | 3,05 | 0,7 | -3,96 | -1,51 | -1,82 |
| Y11_RS11645 | Y11_24711 | *exbD* | TonB system transport protein ExbD | 0,37 | -1,41 | -2,68 | -1,78 | -3,06 | -1,28 | 0,775 | 1,312 | 0,986 |
| Y11_RS11650 | Y11_24721 | *exbB* | tol-pal system-associated acyl-CoA thioesterase | 1,79 | 0,31 | -0,86 | -1,49 | -2,65 | -1,16 | 1,333 | 1,325 | 1,325 |
| Y11_RS11675 | Y11_24771 | *dkgA* | 2,5-didehydrogluconate reductase DkgA | -0,22 | 3,08 | 4,16 | 3,29 | 4,37 | 1,08 | -1,67 | -1,12 | -1,49 |
| Y11_RS11760 | Y11_24951 | *ribB* | 3,4-dihydroxy-2-butanone-4-phosphate synthase | -0,65 | 2,08 | 2,55 | 2,74 | 3,21 | 0,47 | -1 | -1,34 | -1,3 |
| Y11_RS12120 | Y11_25731 | *rlmG* | 23S rRNA (guanine(1835)-N(2))-methyltransferase RlmG | 0,58 | -1,04 | -1,6 | -1,62 | -2,17 | -0,55 | 0,583 | 3,006 | 2,838 |
| Y11_RS12145 | Y11_25781 | | Gfo/Idh/MocA family oxidoreductase | -0,16 | 0,85 | 2,12 | 1,01 | 2,28 | 1,27 | -0,98 | -1,05 | -0,77 |
| Y11_RS12175 | Y11_25831 | | tagaturonate reductase | -1,18 | 1,48 | 2 | 2,67 | 3,18 | 0,51 | -1 | -1,4 | -1,09 |
| Y11_RS12180 | Y11_25841 | *uxaC* | glucuronate isomerase | -0,94 | 1,88 | 3,1 | 2,81 | 4,04 | 1,23 | -0,81 | -1,56 | -1,12 |
| Y11_RS12215 | Y11_25921 | | YqjD family protein | 0,12 | 0,98 | 1,31 | 0,86 | 1,2 | 0,34 | -1,25 | -0,85 | -1,28 |
| Y11_RS12355 | Y11_26221 | *elbB* | isoprenoid biosynthesis glyoxalase ElbB | 0,19 | 1,89 | 2,3 | 1,7 | 2,11 | 0,41 | -1,11 | -1,08 | -1,02 |
| Y11_RS12360 | Y11_26231 | *arcB* | aerobic respiration two-component sensor histidine kinase ArcB | 0,01 | 0,7 | 1,21 | 0,68 | 1,19 | 0,51 | -0,71 | -0,92 | -0,71 |
| Y11_RS12640 | Y11_26831 | *cybC* | cytochrome b562 | -1,26 | 0,76 | 1,34 | 2,02 | 2,6 | 0,58 | -1,48 | -1,53 | -1,26 |
| Y11_RS12670 | Y11_26891 | | NAD-dependent succinate-semialdehyde dehydrogenase | 0,41 | 2,78 | 3,05 | 2,37 | 2,64 | 0,27 | -1,69 | -1,01 | -1,4 |
| Y11_RS12850 | Y11_27261 | *fis* | DNA-binding transcriptional regulator Fis | -0,31 | -1,7 | -2,59 | -1,38 | -2,28 | -0,89 | 2,735 | 2,361 | 3,512 |
| Y11_RS12860 | Y11_27281 | | glycine zipper 2TM domain-containing protein | -0,02 | 2,35 | 3,15 | 2,37 | 3,17 | 0,8 | -2,84 | -1,67 | -2,1 |
| Y11_RS13060 | Y11_27711 | *malK* | maltose/maltodextrin ABC transporter ATP-binding protein MalK | 0,06 | -1,56 | -1,44 | -1,63 | -1,51 | 0,12 | -0,13 | 1,218 | 0,951 |
| Y11_RS13335 | Y11_28241 | *rraA* | ribonuclease E activity regulator RraA | -0,48 | 0,76 | 1,65 | 1,24 | 2,13 | 0,89 | 0,356 | -1,05 | -1,1 |
| Y11_RS13355 | Y11_28271 | *glpK* | glycerol kinase GlpK | -0,52 | 0,96 | 1,16 | 1,48 | 1,68 | 0,2 | -1,09 | -2,16 | -1,85 |
| Y11_RS13770 | Y11_29171 | *spoT* | bifunctional GTP diphosphokinase/guanosine-3',5'-bis pyrophosphate 3'-pyrophosphohydrolase | -0,23 | 0,11 | -0,11 | 0,34 | 0,13 | -0,22 | -13 | 0,123 | -12,5 |
| Y11_RS13980 | Y11_29581 | *kup* | low affinity potassium transporter Kup | -0,11 | -1,13 | -0,82 | -1,02 | -0,72 | 0,31 | 0,406 | 1,072 | 0,865 |
| Y11_RS14150 | Y11_29941 | | NAD(P)H-dependent oxidoreductase | 0,44 | 1,54 | 1,45 | 1,1 | 1,01 | -0,1 | -2,23 | -1,14 | -1,17 |
| Y11_RS14340 | Y11_30351 | | PTS mannitol transporter subunit IICBA | 1,48 | 3,34 | 3,36 | 1,86 | 1,88 | 0,02 | -0,88 | -4,37 | -4,19 |
| Y11_RS14345 | Y11_30361 | | mannitol-1-phosphate 5-dehydrogenase | 0,77 | 2,52 | 2,55 | 1,74 | 1,78 | 0,04 | -0,99 | -1,97 | -2,12 |
| Y11_RS14370 | Y11_30411 | *vapB* | type II toxin-antitoxin system VapB family antitoxin | 0,29 | 0,91 | 1,31 | 0,63 | 1,03 | 0,4 | -0,27 | -1,01 | -1,01 |
| Y11_RS14470 | Y11_30611 | *xylF* | D-xylose ABC transporter substrate-binding protein | 0,47 | 1,96 | 3,29 | 1,5 | 2,83 | 1,33 | -3,26 | -1,03 | -1,01 |
| Y11_RS14570 | Y11_30861 | | PTS sugar transporter subunit IIB | 0 | -15 | -15 | -15 | -15 | 1,68 | -0,63 | 1,266 | 1,109 |
| Y11_RS14590 | Y11_30901 | | LacI family transcriptional regulator | -0,43 | 1,29 | 1,04 | 1,67 | 1,48 | -0,21 | 0,361 | -1,03 | -0,89 |
| Y11_RS14670 | Y11_31071 | | organic hydroperoxide resistance protein | 0,79 | 3,54 | 4,44 | 2,75 | 3,65 | 0,9 | -2,01 | -1,46 | -1,48 |
| Y11_RS14850 | Y11_31451 | *uspA* | universal stress protein UspA | -0,76 | 2,54 | 3,05 | 3,3 | 3,81 | 0,51 | -0,68 | -1,1 | -0,66 |
| Y11_RS14960 | Y11_31681 | | MurR/RpiR family transcriptional regulator | -0,2 | 1,81 | 2,63 | 2,01 | 2,83 | 0,82 | -0,04 | -1,45 | -1,31 |
| Y11_RS15040 | Y11_31841 | *glgB* | 1,4-alpha-glucan branching protein GlgB | -0,19 | 0,68 | 1,22 | 0,87 | 1,4 | 0,53 | -3,38 | -2,61 | -2,95 |
| Y11_RS15045 | Y11_31851 | *glgX* | glycogen debranching protein GlgX | 0,35 | 1,34 | 1,46 | 0,99 | 1,1 | 0,12 | -3,52 | -2,32 | -2,76 |
| Y11_RS15050 | Y11_31861 | *glgC* | glucose-1-phosphate adenylyltransferase | 0,01 | 1,88 | 2,39 | 1,87 | 2,38 | 0,52 | -3,73 | -2,2 | -2,84 |
| Y11_RS15055 | Y11_31871 | *glgA* | glycogen synthase GlgA | 0,14 | 1,6 | 2,23 | 1,46 | 2,08 | 0,62 | -3,38 | -1,8 | -2,1 |
| Y11_RS15060 | Y11_31881 | *glgP* | glycogen phosphorylase | 0,22 | 1,93 | 2,72 | 1,71 | 2,5 | 0,79 | -3,59 | -1,79 | -2,17 |
| Y11_RS15190 | Y11_32191 | *pckA* | phosphoenolpyruvate carboxykinase (ATP) | -0,73 | 0,51 | 2,4 | 1,23 | 3,13 | 1,9 | -1,73 | -1,08 | -0,54 |
| Y11_RS15250 | Y11_32331 | *aroK* | shikimate kinase AroK | -0,17 | -1,59 | -1,76 | -1,42 | -1,6 | -0,18 | 1,427 | 1,339 | 1,515 |
| Y11_RS15255 | Y11_32341 | *aroB* | 3-dehydroquinate synthase | -0,25 | -1,43 | -1,01 | -1,18 | -0,76 | 0,42 | 1,443 | 1,241 | 1,507 |
| Y11_RS15495 | Y11_32861 | *bfr* | bacterioferritin | 0,68 | 3,87 | 4,66 | 3,19 | 3,98 | 0,79 | -0,63 | -0,9 | -0,72 |
| Y11_RS15750 | Y11_33331 | | ABC transporter substrate-binding protein | -0,31 | 3,48 | 3,85 | 3,79 | 4,15 | 0,37 | -2,45 | -1,68 | -1,06 |
| Y11_RS15755 | Y11_33341 | | sugar ABC transporter ATP-binding protein | -0,38 | 1,71 | 1,99 | 2,09 | 2,37 | 0,28 | -2,63 | -1,51 | -1,13 |
| Y11_RS15865 | Y11_33581 | | molecular chaperone | 0,58 | 3,66 | 5,66 | 3,08 | 5,07 | 2 | -1,61 | -1,69 | -1,77 |
| Y11_RS15890 | Y11_33631 | *ppx* | exopolyphosphatase | 0,23 | 2,92 | 15 | 2,63 | 15 | 0 | 0,647 | 0,023 | -0,31 |
| Y11_RS16005 | Y11_33831 | *hemY* | protoheme IX biogenesis protein HemY | 0,13 | 1,29 | 1,56 | 1,17 | 1,44 | 0,27 | -0,82 | -0,56 | -0,94 |
| Y11_RS16010 | Y11_33841 | *hemX* | uroporphyrinogen-III C-methyltransferase | 0,51 | 1,09 | 1,57 | 0,57 | 1,06 | 0,48 | -0,34 | -0,7 | -0,59 |
| Y11_RS16080 | Y11_33991 | *corA* | magnesium/cobalt transporter CorA | -1,11 | -2,44 | -2,06 | -1,34 | -0,96 | 0,38 | 2,1 | 2,496 | 2,827 |
| Y11_RS16165 | Y11_34171 | *tusA* | sulfurtransferase TusA | 1,04 | -1,72 | -2,16 | -2,76 | -3,2 | -0,43 | 1,311 | 1,893 | 1,845 |
| Y11_RS16335 | Y11_34531 | | dienelactone hydrolase family protein | -0,09 | 3,52 | 4,34 | 3,61 | 4,43 | 0,82 | -3,36 | -2,34 | -3,21 |
| Y11_RS16425 | Y11_34741 | *fadA* | acetyl-CoA C-acyltransferase FadA | 0,34 | 1,18 | 3,37 | 0,84 | 3,03 | 2,19 | -2,26 | -3,15 | -2,87 |
| Y11_RS16430 | Y11_34751 | *fadB* | fatty acid oxidation complex subunit alpha FadB | 0,56 | 1,17 | 3,43 | 0,61 | 2,87 | 2,26 | -1,97 | -2,89 | -2,14 |
| Y11_RS16685 | Y11_35131 | *actP* | cation/acetate symporter ActP | -0,51 | 15 | 15 | 15 | 15 | 0 | -3,01 | -3,71 | -3,66 |
| Y11_RS16690 | Y11_35141 | | DUF485 domain-containing protein | 0,22 | 2,05 | 1,99 | 1,83 | 1,78 | -0,05 | -3,67 | -4,38 | -3,96 |
| Y11_RS16695 | Y11_35151 | *acs* | acetate--CoA ligase | -0,29 | 3,98 | 15 | 4,27 | 15 | 15 | -2,74 | -4,29 | -4,08 |
| Y11_RS16920 | Y11_35641 | | TonB-dependent siderophore receptor | 0,49 | -2,35 | -3,5 | -2,84 | -3,99 | -1,15 | -0,06 | 4,271 | 3,926 |
| Y11_RS16925 | Y11_35661 | | PLP-dependent aminotransferase family protein | -0,14 | -0,31 | 0,35 | -0,17 | 0,49 | 0,66 | -0,89 | -1,03 | -0,88 |
| Y11_RS16940 | Y11_35691 | *hmuT* | hemin ABC transporter substrate-binding protein | 0,88 | -1,2 | -2,82 | -2,08 | -3,7 | -1,62 | -2,02 | 1,696 | 0,995 |
| Y11_RS16945 | Y11_35701 | *hmuS* | hemin-degrading factor | 0,29 | -2,71 | -4,85 | -2,99 | -5,13 | -2,14 | -1,87 | 1,768 | 1,275 |
| Y11_RS16950 | Y11_35711 | *hmuR* | TonB-dependent hemoglobin/transferrin/lactoferrin family receptor | 1,85 | -1,42 | -2,95 | -3,27 | -4,79 | -1,52 | -0,81 | 1,776 | 1,67 |
| Y11_RS17025 | Y11_35851 | | anaerobic C4-dicarboxylate transporter | -1,44 | 2,31 | 2,03 | 3,74 | 3,47 | 0 | -0,79 | -0,96 | -0,69 |
| Y11_RS17030 | Y11_35861 | *aspA* | aspartate ammonia-lyase | -1,29 | 1,27 | 3,24 | 2,57 | 4,54 | 1,97 | -0,52 | -1,28 | -0,92 |
| Y11_RS17070 | Y11_35931 | | entericidin A/B family lipoprotein | 0,47 | 3,7 | 4,18 | 3,23 | 3,71 | 0,48 | -1,45 | -1,56 | -2,41 |
| Y11_RS17095 | Y11_35981 | | succinate dehydrogenase/fumarate reductase iron-sulfur subunit | -0,82 | 0,63 | 1,46 | 1,45 | 2,28 | 0,83 | -0,97 | -2,08 | -1,45 |
| Y11_RS17100 | Y11_35991 | *frdA* | fumarate reductase (quinol) flavoprotein subunit | -0,88 | 0,37 | 1,3 | 1,25 | 2,17 | 0,93 | -0,39 | -1,85 | -1,12 |
| Y11_RS17495 | Y11_36771 | | DEAD/DEAH family ATP-dependent RNA helicase | 0,36 | -0,91 | -2,88 | -1,28 | -3,24 | -1,96 | 4,249 | 1,387 | 1,652 |
| Y11_RS17855 | Y11_37571 | *lsrG* | (4S)-4-hydroxy-5-phosphonooxypentane-2,3-dione isomerase | -0,3 | 1,2 | 1,94 | 1,49 | 2,23 | 0,74 | -2,61 | -0,12 | -0,45 |
| Y11_RS17860 | Y11_37581 | *lsrF* | 3-hydroxy-5-phosphonooxypentane-2,4-dione thiolase | -0,51 | 2,05 | 2,68 | 2,56 | 3,18 | 0,62 | -2,52 | -0,57 | -0,64 |
| Y11_RS17865 | Y11_37591 | *lsrB* | autoinducer 2 ABC transporter substrate-binding protein LsrB | 0,03 | 2,81 | 3,84 | 2,78 | 3,81 | 1,03 | -2,92 | -1,69 | -1,6 |
| Y11_RS17880 | Y11_37621 | *lsrA* | autoinducer 2 ABC transporter ATP-binding protein LsrA | -0,45 | 15 | 15 | 15 | 15 | 0 | -2 | -1,33 | -0,71 |
| Y11_RS17885 | Y11_37631 | *lsrR* | transcriptional regulator LsrR | 0,15 | 1,25 | 1,43 | 1,1 | 1,28 | 0,18 | 0,203 | -0,53 | 0,046 |
| Y11_RS17890 | Y11_37641 | *lsrK* | autoinducer-2 kinase | -0,15 | 0,81 | 1,3 | 0,96 | 1,45 | 0,49 | -1,57 | -0,35 | -0,88 |
| Y11_RS18015 | Y11_37891 | *fecA* | Iron(III) dicitrate transport protein FecA | 2 | 0,26 | -0,74 | -1,73 | -2,75 | -1,01 | 1,772 | 2,645 | 3,056 |
| Y11_RS18070 | Y11_38001 | *osmY* | molecular chaperone OsmY | 0,66 | 1,9 | 2,88 | 1,24 | 2,22 | 0,98 | -1,79 | -1,28 | -0,9 |
| Y11_RS18120 | Y11_38111 | *radA* | DNA repair protein RadA | -0,36 | -0,99 | -1,22 | -0,63 | -0,87 | -0,24 | 0,095 | 1,303 | 0,87 |
| Y11_RS18265 | Y11_38401 | *satP* | acetate uptake transporter | 0,08 | -4,17 | -4,93 | -4,25 | -5 | -0,76 | 2,941 | 3,876 | 4,647 |
| Y11_RS18915 | Y11_39791 | *relA* | GTP diphosphokinase | 0,01 | -0,02 | 0,12 | -0,03 | 0,1 | 0,13 | -8,3 | -0,32 | -9,22 |
| Y11_RS18965 | Y11_39901 | *gcvT* | glycine cleavage system aminomethyltransferase GcvT | -1,43 | 2,95 | 3,51 | 4,38 | 4,94 | 0,56 | -0,87 | -1,05 | -0,62 |
| Y11_RS19420 | Y11_40851 | | YgdI/YgdR family lipoprotein | 1,08 | 2,33 | 2,64 | 1,24 | 1,55 | 0,31 | -0,85 | -1,63 | -1,23 |
| Y11_RS19485 | Y11_40981 | | DUF3461 family protein | -0,8 | 0,79 | 1,33 | 1,59 | 2,13 | 0,54 | -2,49 | -1,06 | -1,59 |
| Y11_RS19805 | Y11_41611 | | YqjD family protein | 0,7 | 2,26 | 2,11 | 1,56 | 1,41 | -0,15 | -1,48 | -1,39 | -1,37 |
| Y11_RS19805 | Y11_41611 | | YqjD family protein | 0,7 | 2,26 | 2,11 | 1,56 | 1,41 | -0,15 | -1,48 | -1,39 | -1,37 |

| | | | | | | | | | | | | |
|---|---|---|---|---|---|---|---|---|---|---|---|---|
| Y11_RS19910 | Y11_41821 | *ureA* | urease subunit gamma | 0,43 | 2,88 | 2,96 | 2,46 | 2,54 | 0,08 | -4,14 | -3,14 | -3,33 |
| Y11_RS19915 | Y11_41831 | *ureB* | urease subunit beta | 0,36 | 3,16 | 3,41 | 2,8 | 3,05 | 0,25 | -3,76 | -3,01 | -3,07 |
| Y11_RS19920 | Y11_41841 | *ureC* | urease subunit alpha | 0,37 | 2,79 | 2,94 | 2,42 | 2,57 | 0,15 | -4,45 | -2,79 | -3,16 |
| Y11_RS19925 | Y11_41851 | *ureE* | urease accessory protein UreE | 0,19 | 2,22 | 1,98 | 2,03 | 1,8 | -0,24 | -4,42 | -2,87 | -3,22 |
| Y11_RS19930 | Y11_41861 | *ureF* | urease accessory protein UreF | 0,05 | 2,32 | 1,67 | 2,27 | 1,62 | -0,65 | -4,73 | -2,84 | -3,34 |
| Y11_RS19935 | Y11_41871 | *ureG* | urease accessory protein UreG | 0,2 | 2,49 | 2,34 | 2,29 | 2,14 | -0,14 | -3,7 | -2,31 | -2,79 |
| Y11_RS19940 | Y11_41881 | *ureD* | urease accessory protein UreD | 0,03 | 2,43 | 1,78 | 2,4 | 1,75 | -0,65 | -3,54 | -2,19 | -2,49 |
| Y11_RS19950 | Y11_41901 | | HoxN/HupN/NixA family nickel/cobalt transporter | -0,32 | 15 | 15 | 15 | 15 | 0 | -3,62 | -1,96 | -2,26 |
| Y11_RS19955 | Y11_41911 | *hdeB* | acid-activated periplasmic chaperone HdeB | 0,19 | 1,88 | 3,27 | 1,69 | 3,09 | 1,4 | -2,67 | -2,46 | -2,68 |
| Y11_RS20320 | Y11_42661 | | DUF1007 family protein | 0,16 | -1,78 | -1,06 | -1,94 | -1,21 | 0,73 | 0,751 | 1,035 | 1,475 |
| Y11_RS20560 | Y11_43221 | | PTS glucitol/sorbitol transporter subunit IIB | -0,27 | 2,83 | 3,8 | 3,1 | 4,07 | 0,97 | -1,69 | -0,85 | 0,021 |
| Y11_RS20580 | Y11_43261 | | YegP family protein | 0,57 | 3,38 | 5,61 | 2,81 | 5,05 | 2,24 | -3,05 | -1,79 | -2,58 |
| Y11_RS20620 | Y11_43341 | *yegD* | molecular chaperone | 0 | -2,72 | -4,33 | -2,85 | -4,46 | -1,61 | 2,749 | 5,343 | 5,95 |
| Y11_RS20650 | Y11_43401 | *ppx* | exopolyphosphatase | 0,38 | -0,29 | -0,03 | -0,68 | -0,42 | 0,26 | -1,27 | 0,185 | 0,133 |
